# Mating strategy predicts gene presence/absence patterns in a genus of simultaneously hermaphroditic flatworms

**DOI:** 10.1101/2022.04.25.489193

**Authors:** R. Axel W. Wiberg, Gudrun Viktorin, Lukas Schärer

## Abstract

Gene repertoire turnover is a characteristic of genome evolution. However, we lack well-replicated analyses of presence/absence patterns associated with different selection contexts. Here, we study ∼100 transcriptome assemblies across *Macrostomum*, a genus of simultaneously hermaphroditic flatworms exhibiting multiple convergent shifts in mating strategy and associated reproductive morphologies. Many species mate reciprocally, with partners donating and receiving sperm at the same time. Other species convergently evolved to mate by hypodermic injection of sperm into the partner. We find that for orthologous transcripts annotated as expressed in the body region containing the testes, sequences from hypodermically inseminating species diverge more rapidly from the model species, *M. lignano*, and have a lower probability of being observed in other species. For other annotation categories, simpler models with a constant rate of similarity decay with increasing genetic distance from *M. lignano* match the observed patterns well. Thus, faster rates of sequence evolution for hypodermically inseminating species in testis-region genes result in higher rates of homology detection failure, yielding a signal of rapid evolution in sequence presence/absence patterns. Our results highlight the utility of considering appropriate null models for unobserved genes, as well as associating patterns of gene presence/absence with replicated evolutionary events in a phylogenetic context.

## Introduction

Gene repertoire turnover characterises the evolution of genomes throughout the tree of life (Albalat and Cañestro 2016; Fernández and Gabaldón 2020). A common observation at many taxonomic scales is that some lineages have no detectable homologs of genes present in other, related lineages (Zhaxabayeva et al. 2008; Vakirlis et al. 2020; Weisman et al. 2020). Often, the inference is that such patterns are the result of adaptive gene gain and/or loss. For example, the study of the origins of novel genes has received a lot of attention (e.g. Neme et al. 2017), and gene loss is often adaptive (Albalat and Cañestro 2016). However, a well-understood, but less exotic, explanation is that “missing” genes have simply evolved, at a constant rate, to the point of being unrecognisable by sequence-similarity approaches to homology detection, so-called “homology detection failure” (Zhaxabayeva et al. 2008; Vakirlis et al. 2020; Weisman et al. 2020). As many as 20-55% of so-called “lineage-specific” genes may be due to homology detection failure in outgroups (e.g. Zhaxabayeva et al. 2008; Vakirlis et al. 2020; Weisman et al. 2020). For genes with overall rapid rates of sequence evolution (e.g. reproduction-related genes, see below) homology detection failure may be more common and failure to consider it as a hypothesis will lead to errors of inference. Recent efforts have aimed to develop methods to test whether an observation of gene absence is expected on the basis of sequence similarity at different evolutionary distances from a focal species, when assuming a constant rate of sequence evolution (Weisman et al. 2020). Deviations from this “null” model can then be due to a variety of factors, including changes in the rate of evolution due to changes in selective pressures on existing genes or *bona fide* gene turnover (gain/loss; Weisman et al. 2020).

### Gene presence/absence and the evolution of reproduction-related genes

Reproduction-related genes are typically found to be evolving at higher rates compared to the genomic background (Swanson and Vacquier 2002; Wilburn and Swanson 2016). A commonly proposed explanation for these observations is diversifying sexual selection (Swanson and Vacquier 2002; Dapper and Wade 2016; 2020; for reviews see also e.g., Vacquier and Swanson, 2011; Wilburn and Swanson, 2016). However, an alternative explanation is that relaxed selection is the driving force behind many of these patterns (Dapper and Wade 2016; 2020). Indeed, there are several reasons why reproduction-related genes might be expected to be under relaxed selection. First, in separate-sexed organisms, reproduction-related genes are often sex-biased or even sex-limited in expression (Dapper and Wade 2020). In the extreme case of sex-limited expression, the efficiency of selection is half that of a gene with unbiased expression, because it is only exposed to selection when residing in one sex (Dapper and Wade 2020). However, it is unclear to what extent sex-biases in expression could operate under other modes of reproduction (e.g. simultaneous hermaphroditism, where individuals are both male and female at the same time; Wiberg et al. 2021). Second, also in separate-sexed organisms, reproduction-related genes are often sexually antagonistic, where the fitness effects differ, and are sometimes in opposition, in the two sexes. In this case the net selection is the average of that experienced in either sex. All else being equal, sexually antagonistic loci therefore necessarily experience lower net directional selection (with a similar effect to relaxed selection) compared to the rest of the genome. Finally, regardless of the mode of reproduction (i.e. hermaphroditism or gonochorism), genes involved in sperm competition or cryptic female choice experience weaker selection because the genetic variation exposed to selection within females is typically only a fraction of that present in the population at large, especially in mating systems with low female remating rates (Dapper and Wade 2016; 2020).

Be it due to relaxed selection or positive selection, the faster rates of evolution of reproduction-related genes would also predict higher rates of homology detection failure for such genes. A few studies have found evidence that, in addition to rapid rates of sequence evolution, turnover of gene repertoires is a characteristic feature of reproduction-related genes (Zhang et al 2004; Cutter & Ward 2005, Ellegren and Parsch 2007; Hahn et al. 2007; Ahmed-Braimah et al. 2017). In particular genes expressed in male reproductive tissues (e.g. the testes, the prostate, or accessory glands) often show high rates of turnover (Zhang et al 2004; Cutter & Ward 2005, Ellegren and Parsch 2007; Hahn et al. 2007; Ahmed-Braimah et al. 2017). In contrast, some studies, for example a recent study of 11 passerine species, found no evidence for higher turnover rates for male-biased reproduction-related genes (Rowe et al. 2020). However, compared to studies of rates of molecular sequence evolution, relatively little work has focused on patterns of presence/absence of reproduction-related genes in different lineages, especially in systems where a link can be drawn to consistent differences in the sexual selection context.

Although there is considerable taxonomic bias in the above studies, they suggest that, in general, reproduction-related genes may not only be rapidly evolving in sequence but also show higher rates of missing genes than other categories of genes. However, this pattern is not strictly universal, raising the question of why such patterns are observed in some systems but not others. Such a pattern is not easily explained as a simple consequence of the higher rates of sequence evolution due to relaxed selection at reproduction-related genes. Since sexual selection is thought to be an important force affecting the evolution of reproduction-related genes, an obvious explanation is that differences in the sexual selection context between systems might account for the observed differences in patterns of gain and loss. This is especially true since changes in copy number may be an important component in the evolution sexual dimorphism; redundant copies of genes are more free to evolve sex-specific expression patterns and functions, partially resolving intra-locus sexual conflict (Ellegren and Parsch 2007). For example, sex-biased expression, allowing sex-specific phenotypes, seems to be more common at recently duplicated genes (Wyman et al. 2011). Moreover, turnover in sex-biased gene expression is associated with differences in the sexual selection context in birds (Harrison et al. 2015). Thus, both a reduced efficiency of selection at reproduction-related genes as well as changes in the intensity of selection at particular loci due to changing sexual selection contexts, might result in the loss of genes from a species’ repertoire. To date, few studies have associated such changes in gene repertoires with changes in the sexual selection context (but see Thomas et al. 2012; Fierst et al. 2015; Yin et al. 2018, reviewed in Cutter et al. 2019, for changes in repertoires associated with transitions to self-fertilisation, which likely also involves a change in the sexual selection context).

A further technical complication is that it is often difficult to distinguish actual absence of a gene from artefacts arising from homology detection failure (e.g. Weisman et al. 2020) or poor gene assembly quality. Poor assembly quality may also in some cases mis-label paralogs as orthologs, as ortholog identification typically relies at least in part on the detection of similar sequences. If true orthologs are not successfully assembled, then (likely more diverged) paralogs may be the most similar sequences detected. Additionally, in the case of transcriptome data, variation in expression levels may also create apparently missing genes if the transcriptome is not sampled to high enough coverage or when expression of some genes is temporally or spatially restricted. Some of this variation in expression, at least across species, may also be a signal of adaptive differences (e.g. Harrison et al. 2015). It is clear that more work is needed to understand whether and how patterns of rapid evolution of reproduction-related genes translates into patterns of gene presence/absence. Additionally, there is a need for data from more taxonomic groups, ideally combined with replicated contrasts of different sexual selection contexts.

### The genus Macrostomum

Free-living flatworms of the genus *Macrostomum* have become excellent models for studying mating behaviour and sexual selection, including sexual conflict (Schärer et al. 2004; Vizoso et al. 2010; Ramm et al. 2015; Giannakara et al. 2016; Giannakara and Ramm 2017; Marie-Orleach et al. 2017; Patlar et al. 2020; Singh et al. 2020a, 2020b). In particular *M. lignano* has been the focus of numerous experiments and has substantial genetic resources (Ladurner et al. 2005; Wudarski et al. 2020). For example, expression studies in *M. lignano* have provided information about body regions containing important reproductive tissues that can be used to annotate reproduction-related transcripts and genes (Arbore et al. 2015; Brand et al. 2020). *Macrostomum* worms are simultaneously hermaphroditic, producing sperm and eggs at the same time. Hermaphroditism is expected to lead to conflicts between individuals over the sex roles they assume during copulation, with the role of sperm donor thought to be preferred over the role of sperm recipient in many scenarios (Charnov 1979; Michiels 1998; Schärer et al. 2015). In *Macrostomum* such conflicts likely explain much of the diversity of reproductive morphologies and behaviours that are observed among species (Schärer et al. 2011; Brand et al. 2022a).

One of the main aspects of this diversity is variation in the mating strategies. Many species of *Macrostomum* engage in reciprocal copulation, where both partners donate and receive sperm at the same time (Schärer et al. 2004; Vizoso et al. 2010). Other species inseminate hypodermically by injecting sperm into the body of their partner using a “needle-like” stylet (Schärer et al. 2011; Ramm et al. 2015). These contrasting strategies have been shown to be strongly associated with other axes of variation in reproductive morphology and behaviour (Schärer et al. 2011; Brand et al. 2022a). In particular, reciprocally copulating species have sperm with stiff lateral bristles that likely serve as an anchoring mechanism to prevent postcopulatory sperm removal (Vizoso et al. 2010; Schärer et al. 2011). Meanwhile hypodermically inseminating species have simpler sperm that lack such bristles. A recent phylogenomic study has shown that hypodermic insemination has evolved at least nine times independently within the genus (Brand et al. 2022a; Brand et al. 2022b). A handful of species also show intermediate sperm morphologies, with short or reduced bristles, possibly representing a transitional state between the reciprocal and hypodermic mating strategies (Schärer et al. 2011; Brand et al. 2022a). The state of the sperm bristle (present, reduced, or absent) is therefore a good indicator of the mating strategy (Schärer et al. 2011; Brand et al. 2022b). Thus, contrasting mating strategies, themselves the result of conflicts over sex roles, represent different sexual selection contexts resulting in repeated evolution of different reproductive morphologies.

We have previously shown that reproduction-related genes evolve rapidly in *Macrostomum* (Brand et al. 2020; Wiberg et al. 2021), and that species with absent sperm bristles (taken as a proxy for hypodermic mating) show faster genome-wide rates of sequence evolution than species with bristles (Wiberg et al. 2021). However, an open question is whether the bristle state also predicts whether orthologs of reproduction-related genes are observed, or not, across species. Transcriptome assemblies show substantial variation in gene presence/absence patterns, although there is a strong association with genetic distance from *M. lignano*, the species from which annotations are derived, there is a conspicuous lack of representative sequences within a clade of exclusively hypodermically mating species (figure S1). A strong effect of bristle state would indicate that different sexual selection contexts alter the nature of selection acting on particular genes, sets of genes, or the genome as a whole. Such effects might inform us about candidate genes involved in, for example, sperm bristle development in the case of genes expressed in the testes. If shifts to a hypodermic mating strategy impose strong selection to lose the bristles, then the genes involved might degrade and be lost rapidly, making them difficult to detect in studies of the rate of sequence evolution (e.g. Brand et al. 2020 and Wiberg et al. 2021). Alternatively, if genes underlying bristle development were also important in other contexts, the loss or reduction of bristles may involve changes in gene expression patterns rather than complete loss. Addressing this possibility will require comparative gene expression datasets from different tissues, which are currently unavailable.

If there were no overall effect of bristle state then the reason for unobserved transcripts in other species is more difficult to infer and could be linked to one of multiple, non-mutually exclusive factors. For example, the lack of an effect could indicate that there is no overall turnover of gene repertoires due to sexual selection. Technical effects due to the use of transcriptome data, such as variation in the quality of assemblies or low expression of certain genes/transcripts, will lead to noise in presence/absence data (see above).

Here we use a set of previously established orthogroups (OGs – a collection of orthologous sequences across multiple species; Brand et al. 2022a; Wiberg et al. 2021) that have been annotated on the basis of reproduction-related transcripts from the model species *M. lignano* (Arbore et al. 2015; Brand et al. 2020). We then test for a signal of decay in similarity scores with increasing phylogenetic distance from the model species *M. lignano* and evaluate the probability of detection given the observed rates of decay (Weisman et al. 2020). Next, we apply phylogenetically controlled logistic regression to test the hypothesis that different sexual selection contexts predict the probability of observing a representative sequence in an OG. We also extend the annotation of these OGs using RNA-Seq data, novel to *M. lignano*, comparing gene expression between adults and hatchlings in the model species *M. lignano*. Elevated expression in adults presents an additional source of evidence for a reproduction-related function of a gene because only adults are reproductively active.

## Methods

### Orthogroups and pairwise similarity scores

Data have previously been generated in *de novo* transcriptome assemblies from 98 species of *Macrostomum* (Brand et al. 2022a). Orthogroups (OG) were produced from this data using the PDC pipeline of Yang & Smith (2014), resulting in OGs with a single representative sequence from each species (though they may not represent strict 1-to-1 orthologs, as from *de novo* transcriptome data it is difficult to tell the difference between allelic variants, isoforms, and duplications). These OGs were processed and filtered to produce codon alignments in a previous study on molecular evolution in *Macrostomum* (Wiberg et al. 2021). We obtained the amino acid alignments corresponding to the codon alignments used in Wiberg et al. (2021). These alignments represent orthogroups (OGs) and are broadly annotated into reproduction-related categories, namely by their expression in the testis-, ovary-, and tail-region of the worm (Arbore et al. 2015; Brand et al. 2020; Wiberg et al. 2021; table S1 – sheet 2). There is also a randomly chosen set of OGs that are annotated as ubiquitously expressed with no obvious reproductive function that serves as a comparison to the reproduction-related OGs. A random subset, rather than all ubiquitously expressed transcripts, was chosen also to mitigate the inflated power of any statistical analyses that assume statistical independence between OGs. Annotations are based on data from a previous study (Arbore et al. 2015), which were used to re-annotate a more recent version of the *M. lignano* genome-guided transcriptome assembly (Brand et al. 2020). Briefly, worms were cut at different levels along the body axis to produce fragments containing the head only; the head and the body region including the testes; the head and the body regions containing the testes and ovaries; the whole worm. RNA-Seq data from pools of these body fragments were mapped to the *M. lignano* transcriptome assembly (Grudniewska et al. 2018) and transcripts were annotated on the basis of expression differences between fragments (Brand et al. 2020). Note that body fragments contained other tissues than the “target” tissues (e.g. the region containing the testes also contains part of the gut).

The amino acid alignments from Wiberg et al. (2021) were trimmed using ZORRO (v. 1.0; Wu et al. 2012) scores, removing columns with a score ≤ 3 (note that some columns will still contain gaps in some representative sequences). For each trimmed alignment, we then removed gap characters (-) and created protein blast databases (blast v. 2.9.0; Camacho et al. 2009). We also refined the annotation of these OGs using expression data from adults and hatchlings of *M. lignano* to identify transcripts that are more highly expressed in adults (see supplementary text T1 in the supporting information for more details as well as figure S2, table S2, and table S3).

### Evaluating rates of homology detection failure

For all OGs we collected similarity scores (bit-scores) for all hits between the focal *M. lignano* sequence and all other species present in the OG using a BLASTP (v. 2.9.0; Camacho et al. 2009) search of all sequences in an OG against a database created from the same OG always treating the *M. lignano* sequence as the “target”. We then applied an approach to model the decay in the bit-score with increasing genetic distance from *M. lignano* (table S1 – sheet 3). Weisman et al. (2020) defined an exponential decay model based on a simple model of sequence evolution:

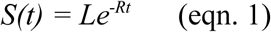

Where *S(t)* is the bit-score at a genetic distance *t* from the focal species (see table S1 – sheet 4 for the genetic distances of all species). *L* is the bit-score of the gene of the focal species with itself (and thus related to the protein’s length), and *R* is the rate of decay in the bit-score. The expected variance at genetic distance *t* is given by:

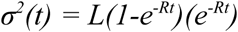

Because linear and non-linear regression fitting approaches minimise slightly different sums of squares, and non-linear regression approaches require specified starting values for estimated parameters, we took a double-fitting approach to estimate the *L* and *R* parameters. First we fit a linear model to log()-transformed bit-scores, and used the *L* and *R* parameter estimates as starting values to fit a non-linear least squared (nls) regression, with untransformed bit-scores, using the “nls()” R function. To fit these models, we only used species with bristles present, and only OGs with data from ≥ 5 species (table S1 – sheet 4). This approach allowed us to evaluate whether the species with reduced or absent bristles deviated from the patterns observed in the species with bristles present. We also created a weighting variable in an attempt to account for variation in assembly quality (*weight = 1 – P*_*miss*_; where *P*_*miss*_ is the proportion of BUSCO (v. 5.0.0; Simão et al. 2015; metazoa_odb10) genes that were reported missing from a transcriptome; see table S1 – sheet 3). We also performed the analysis excluding species with high rates (>50%) of missing BUSCO genes. Despite a substantial reduction in sample sizes, particularly for species with reduced or absent bristles, our results were qualitatively unchanged (see the supplementary text T2). Any OGs that produced *R* values > 0 (i.e. an estimated increase in similarity score with increasing distance from *M. lignano;* N = 7/999, 0.7%) were treated as spurious, likely due to noisy input data, and were not considered further.

We next computed residuals for all species, including those that were not used to estimate the best-fit line, by subtracting the empirical bit-score from that predicted by the parameter estimates and the non-linear equation given above (eqn. 1). Because none of the species used in the model fitting had genetic distances to *M. lignano >* 0.7, the accuracy of predictions could be compromised when extrapolating to larger genetic distances. We therefore only used residuals computed for species with genetic distances to *M. lignano* < 0.7. This effectively excludes all species in a clade of exclusively hypodermically inseminating species (called the “hypodermic mating clade” in Brand et al. 2022a; yellow box in figure 1A). This had the added benefit that there was no longer a detectable systematic difference between the species with different bristle states in the genetic distance to *M. lignano* (figure S3A and B). Moreover, there was no systematic difference in assembly “completeness”, as measured by BUSCO, between species with different bristle states (figure S3C and D). We summarised the residuals for each bristle state within each OG as the median value across representative species (figure 1B). We present a validation of the above approach, namely to fit models using one group of species and to evaluate their fit to another group of species not included in the original model, in the supplementary materials (see the supplementary text T3).

**Figure 1.**
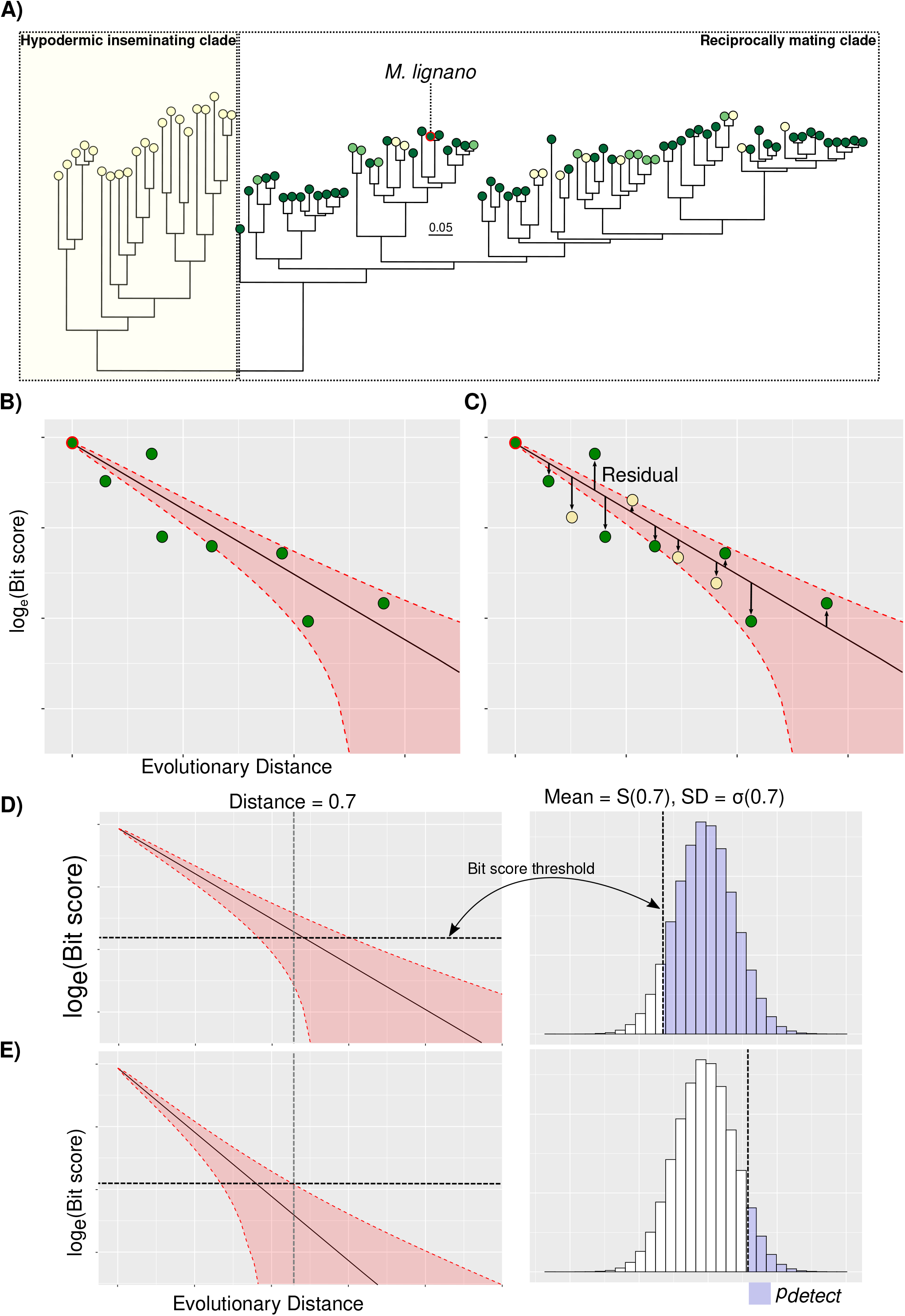
General approach employed. **A)** A depiction of the phylogenetic distribution of species used in this study. The hypodermic mating clade, which is excluded from analyses (see Methods), is highlighted in the yellow box, and the focal species (*M. lignano*) is outlined in red. Dark green, light green, and yellow circles at branch tips denote species with present, reduced, and absent sperm bristles, respectively. **B)** and **C)** are diagrams of the 2-step analysis approach taken for each orthogroup (see Methods): **B)** species with bristles are used to fit a line describing the decay in bit scores, a measure of pairwise sequence similarity from a BLAST alignment, as a function of evolutionary distance from the focal species (*M. lignano*). **C)** Residuals from other species (here only species with absent bristles are shown) to the line of best fit are then computed and summarised. Negative and positive residuals mean less similar and more similar sequences than expected, respectively. Shaded areas in **B)** and **C)** represent the estimated variance in expected bit scores, from which the probability of detection (*p*_*detect*_ ; see Methods) can be calculated. **D)** and **E)** are representations of the approach to compute *p*_*detect*_ from the estimated variance in expected bit scores at a given evolutionary distance. A bit-score threshold is defined, and the proportion of a normal distribution with mean S(0.7) and standard deviation σ(0.7), given by the fitted model, that lies above this threshold is defined as *p*_*detect*_. **D)** shows a case with a high *p*_*detect*_, where most of the predicted distribution of bit-scores lies above the threshold bit-score value, while **E)**, in contrast, shows a case with a low *p*_*detect*_

Given this model, fit using all available data for species with bristles present, and the parameter estimates of *L* and *R* for each OG, it is then possible to compute the probability of detecting a homolog at some genetic distance *t*, by setting an e-value threshold, and assuming the sequence length. We set an e-value threshold of 0.001 (a commonly used value), and assume that the length of a hypothetical sequence is the same as that of the relevant *M. lignano* sequence. The database size was set to 853,740,050 amino acids, the size of the initial database in Brand et al. (2022a) used for homology detection. We then computed the probability of detection (*p*_*detect*_) as the proportion of a normal distribution with mean *S(0*.*7)* and standard deviation *σ(0*.*7)* that lies above the given bit-score threshold (Weisman et al. 2020; figure 1C). We identified OGs with a high probability of detection (*p*_*detect*_ > 0.90; table S1 – sheet 4). If the fit of data from species with absent or reduced bristles are a good fit to the model, then these *p*_*detect*_ values should also be representative for these species. Thus, for OGs with *p*_*detect*_ > 0.90, but for which representative sequences are unobserved for some species, alternative explanations can then be explored (see also next section).

The approach by Weisman et al. (2020) has some limitations. First, because bit-scores are dependent on the length of the sequences being compared, and sequence lengths varied both within OG alignments, due to alignment gaps even after trimming, as well as across OGs, due to variation in sequence length across genes, this results in substantial variation in the bit-scores (figure S4A). This is likely to be a source of increased variation in bit-scores more generally, especially in non-model systems with fewer and less well characterised genetic resources. Second, as in any comparative analysis, species may have similar bit-scores, and genetic distances to the focal species, because they are themselves closely related (i.e. phylogenetic non-independence; Felsenstein 1985; Uyeda et al. 2017). Thus, modelling approaches that explicitly account for phylogeny would be preferable. We therefore also modelled the rate of decay in the bit-scores with increasing phylogenetic distance from *M. lignano* in a way that tried to address these limitations. However, because these additional analyses produced very similar results to those above, we only present the relevant methods and results in the supplementary information (supplementary text T4; table S1 – sheet 5).

### Gene presence/absence

Our above approach to compute residuals from the fitted decay models does not capture cases of complete absence of a sequence from an orthogroup, because in such cases there are no empirical bit-scores to compare against predicted values. We therefore took a complementary approach to analyse transcript presence/absence data. We fit phylogenetically-controlled logistic regression models to presence/absence data for those OGs with a high probability of detecting homologs among species with bristles present (*p*_*detect*_ > 0.9; see above). These models test whether the bristle state or reproduction-related annotations had an effect on the probability of observing a representative sequence. Because the decay rate analyses establish that simple models of similarity decay fit the data well also for species with reduced and absent bristles, any significant effect of bristle state on the probability can be attributable to other processes. To incorporate the phylogenetic relationships as a random effect, we fit these logistic regression models with the “MCMCglmm” package (v. 2.32; Hadfield 2010), using a probit link function (family = “categorical”). To incorporate the phylogenetic random effect, we transformed the phylogenetic tree (H-IQ-TREE from Brand et al. 2022a; see also figure 1A) to be ultrametric with a root depth of 1 using the penalised marginal likelihood approach (Sanderson 2002) in TreePL (Smith and O’Meara 2012), as in Brand et al. (2022b). As above, we excluded data from species with a genetic distance from *M. lignano* > 0.7. For a single OG, model fitting was not possible because all species with genetic distances to *M. lignano* < 0.7 had a representative sequence. We summarised the effects of bristle state by collecting, for each OG, the posterior means (on the probit scale) for each bristle state (table S1 – sheet 6). We then tested for an overall difference in the posterior means between bristle states within each annotation category, and between annotations within bristle states, using Kruskal-Wallis rank-sum tests and Dwass-Steele-Critchlow-Fligner all-pairs tests for *post-hoc* contrasts. We also identified a set of candidate OGs by comparing the 95% confidence intervals of the posterior means between bristle states (table S1 – sheet 6). If the entire interval for species with absent bristles was below that for species with bristles present, indicating a much lower probability of observing a representative sequence, we called the posterior means significantly different.

### Statistical software and packages

In addition to the above-named packages, several other R packages were used in the analysis pipeline of these data; “ddpcr” (v. 1.15; Attali 2020), “plyr” (v. 1.8.6; Wickham 2011), “ggplot2” (v. 3.3.3; Wickham 2016), “here” (v. 1.0.1; Müller 2020), as well as the packages for handling phylogenetic trees; “ape” (v. 5.5; Paradis & Schliep 2019), “ggtree” (v. 3.2.1; Yu et al. 2017; Yu et al. 2018; Yu 2020), “ggimage” (v. 0.3.0; Yu 2021), “TDbook” (v. 0.0.5; Yu et al. 2022) and “phytools” (v. 0.7-70; Revell 2012). Bioinformatic analyses involved cutadapt (v. 3.0; Martin 2011), salmon (v. 1.4.0; Patro et al. 2017), DESeq2 (v. 1.24.0; Love et al. 2014) as well as the *M. lignano* transcriptome assembly (Wudarski et al. 2017; Grudniewska et al. 2018; see supplementary text T1).

## Results and Discussion

Reproduction-related genes have been shown to evolve rapidly in the genus *Macrostomum*, with rates of sequence evolution being associated with the state of the sperm bristles (Wiberg et al. 2021), a diagnostic feature of divergent mating systems in these flatworms (Brand et al. 2022b). Meanwhile, patterns of presence/absence hint at an association with bristle states (figure S1), a pattern also seen for reproduction-related genes in other taxa, and often attributed to sexually-selected turnover of gene repertoires (Zhang et al 2004; Cutter & Ward 2005, Ellegren and Parsch 2007; Hahn et al. 2007; Ahmed-Braimah et al. 2017; Cutter 2019). However, homology detection failure may also produce patterns of presence/absence, and such artefacts may be even more likely among rapidly evolving genes. We apply a recently proposed approach (Weisman et al. 2020) to evaluate whether patterns of presence/absence are related to bristle states beyond what is expected from simple models of sequence similarity decay as a function of genetic distance (homology detection failure). To reject this hypothesis, the probability of observing a representative sequence predicted by the sperm bristle states must be lower than expected from differences between bristle states in the rate of sequence similarity decay.

We were able to estimate decay rates for 959 OGs (figure S5 A-D; table S1 – sheet 4), and for 796 of these OGs (83%) a homologous sequence had an estimated probability (*p*_*detect*_) > 0.9 of being detected at a genetic distance of 0.7 from *M. lignano*. The percentage of genes with *p*_*detect*_ > 0.9 varied among the reproduction-related OGs (Testis-region 81.7%; Ovary-region 83.6%; Tail-region 78.2%; Ubiquitously expressed 88.0%). As expected, OGs annotated as ubiquitously-expressed had the highest percentage. The ubiquitously-expressed OGs are expected to be more conserved and thus probably have slower rates of evolution on average, making representative sequences easier to detect at greater genetic distances.

There were significant differences in the median residuals across bristle states for OGs with the testis-region annotations (Kruskal-Wallis rank-sum test, χ^2^ = 53.51, p < 0.001), with the pattern being driven by more negative residuals in species with reduced and absent bristles (figure 2A). This suggests that the simple model of a constant rate of similarity decay estimated from species with bristles does not adequately explain similarity scores for species with either reduced or absent bristles. This is in line with previous observations that genes with testis-region annotations show faster rates of sequence evolution in species with absent bristles compared to species with bristles (Wiberg et al. 2021). There were also significant effects of bristle state on the probability of observing representative sequences within OGs with testis-region annotations (Kruskal-Wallis rank-sum test; χ^2^ = 14.26, d.f. = 2, p < 0.001). This effect was driven by differences between species with bristles and those with either reduced or absent bristles (figure 2B). Thus, for OGs with the testis-region annotations, results indicated rapid rates of sequence evolution as well as lower probabilities of observing a sequence in species with absent or reduced bristles. Although it remains possible that there is true gene loss among testis-region genes driving the lower probability of observing a sequence, rather than homology detection failure, the combination of low probabilities of observing a sequence and the faster rates of sequence evolution mean that it is not possible to tell these hypotheses apart. Indeed, these hypotheses are also not mutually exclusive and either processes could be at work for any particular OG. The most parsimonious explanation, effectively the null hypothesis, is that the more rapid rates of sequence evolution in species with reduced and absent bristles lead to higher rates of homology detection failure.

**Figure 2.**
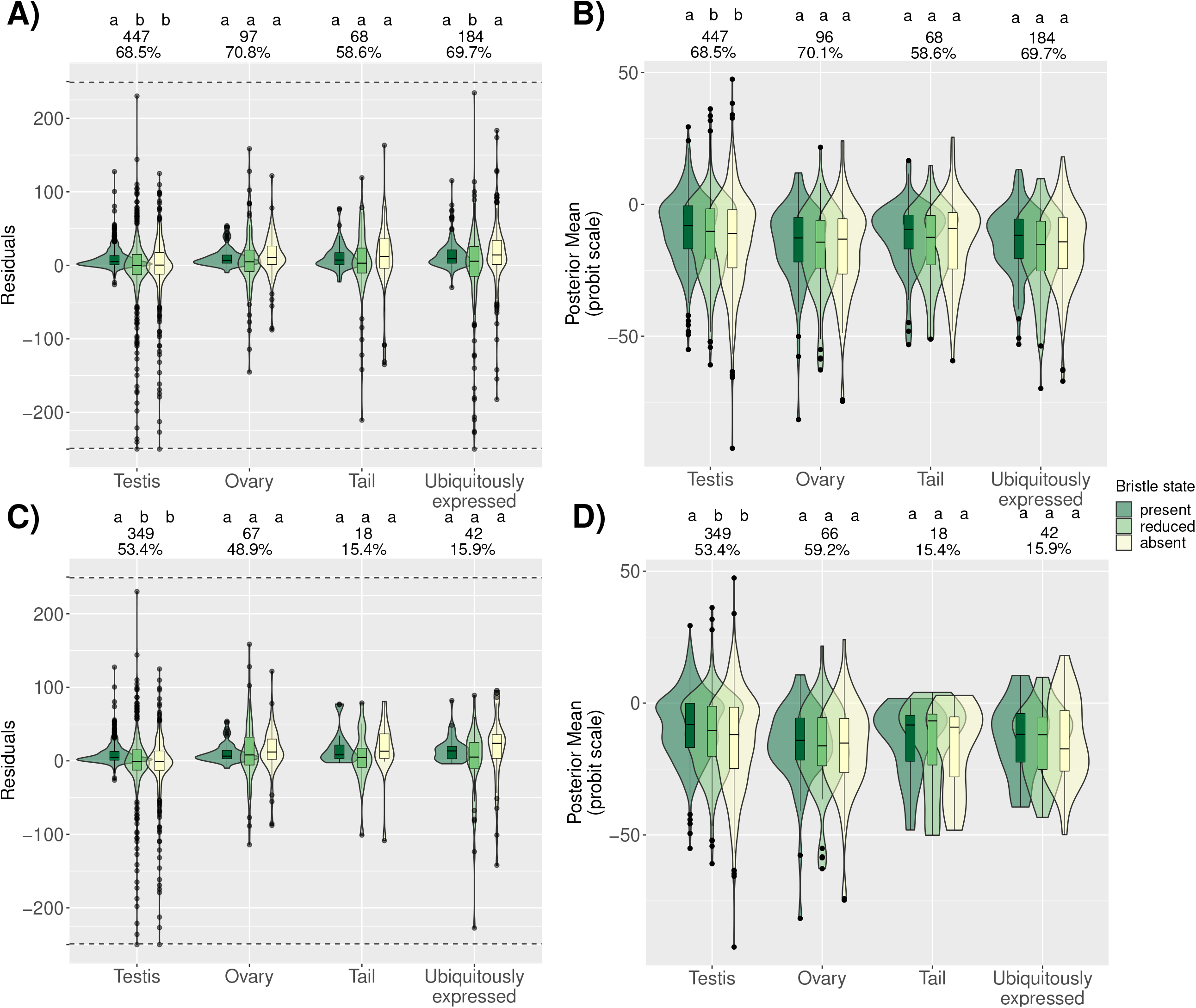
Distributions across orthogroups (OGs) with different functional annotations for species with different bristle states. **A)** Distributions across OGs with different functional annotations of the median residuals for species with different bristle states, when fitting similarity decay models for the species with bristles present. Negative and positive residuals mean less similar and more similar sequences than expected, respectively. **B)** Distributions across OGs with different annotations of the posterior mean estimate of the probability (on the probit scale) of observing a gene for species with different bristle states. **C)** and **D)** are the same as in in **A)** and **B)**, respectively, but with data subset to OGs annotated as having higher expression in adults compared to hatchlings. In all panels, letters above the data for each annotation category give the results of Dwass-Steele-Critchlow-Fligner all-pairs *post-hoc* tests, with different letters denoting significant (p < 0.05) differences between bristle states. The inset numbers above each group of box- and violin-plots give the number and percent of all OGs in each set (see also table S1 – sheet 4 and sheet 6). In **A)** and **C)** to make the figure clearer, OGs with values < −250 (N = 3) and > 250 (N = 0) have been plotted at *y =* −250 and *y =* 250 respectively, and separated from the remaining points by dashed horizontal lines.

These findings potentially have implications for interpreting previous findings in other taxa. Several studies have found male-biased genes, in particular those expressed in testes and the male germline, to show both high rates of sequence evolution and gene turnover. For example, in both *Caenorhabditis briggsae* (Cutter & Ward 2005) and *Drosophila melanogaster* (Zhang et al. 2004) there is evidence of high turnover for genes expressed in the testes and the male germline. Additionally, in a comparative study of *Drosophila*, Hahn et al. (2007) showed evidence for expansions in gene families associated with spermatogenesis in the lineage to *D. melanogaster*. Another comparative study of *Drosophila*, this time in the *D. virilis* group, found lineage-specific transcripts only for genes expressed in testes of *D. americana* and the ejaculatory bulb of *D. lummei* (Ahmed-Braimah et al. 2017). In the light of our findings, such results might be due to detection failure in some lineages stemming from more rapid sequence evolution of genes expressed in testes as compared to other tissues. Our results suggest that this hypothesis should be considered and tested before explanations involving adaptive gain/loss are invoked.

In other taxa, different groups of reproduction-related genes also show interesting patterns of presence/absence. For example, for lineage-specific gene families unique to the *Drosophila* sub-genus, many of the genes unique to *D. melanogaster* were found to be expressed in the accessory glands (Hahn et al. 2007; Clark et al. 2007; Haerty et al. 2007), which are also known to evolve particularly rapidly in *Drosophila* (Haerty et al. 2007). More recently, it was shown that several members of the CheB family of gustatory receptors, which show sexually dimorphic expression and are involved in modulating mating and aggression behaviours in *D. melanogaster*, show high rates of gene turnover (Torres-Oliva et al. 2016). Our results suggest that homology detection failure might provide an explanation more generally and gene turnover rates should therefore be placed in the context of sequence evolution for a full picture.

Among OGs with ubiquitously-expressed annotations there was a significant effect of bristle state (χ^2^ = 12.95, p < 0.001), also in line with previous findings (Wiberg et al. 2021). However, this effect was driven by slightly more negative residuals in species with reduced bristles compared to either species with present bristles or species with absent bristles (figure 2A), which is difficult to explain. This might be a spurious effect of the relatively low sample sizes for species with reduced bristles. This predicts that there should be no overall difference among bristle states in the probability of observing a sequence for OGs annotated as ubiquitously-expressed. For OGs annotated as ubiquitously-expressed there was a marginally non-significant effect of bristle state on the probability of observing a representative sequence (χ^2^ = 5.26, d.f. = 2, p = 0.07).

In contrast, we did not detect an effect of bristle state for ovary- or tail-region OGs (χ^2^ = 4.53, d.f. = 2, p = 0.1 and χ^2^ = 2.78, d.f. = 2, p = 0.25, respectively; figure 2A). For these annotation groups, the results therefore differ from previous analyses, where an effect of bristle state was observed for all reproduction-related annotation categories (Wiberg et al. 2021). However, given that the model of Weisman et al. (2020) employed here is a fairly simplistic way of describing sequence evolution, these contrasting results are not so surprising. Codon models, as applied in Wiberg et al. (2021), are likely more sensitive to variation in rates of sequence evolution, and we therefore consider these results largely consistent with those of previous work (Brand et al. 2020; Wiberg et al. 2021). On the other hand, the absence of a significant difference in the residuals between species with present bristles and those with absent bristles suggests that the probability of a sequence being detectable should also not depend on the bristle state for these annotation categories. As expected from these results on bit-score decay, there were no detectable effects of bristle state for ovary- or tail-region OGs on the probability of observing a representative sequence (χ^2^ = 3.0, d.f. = 2, p = 0.22 and χ^2^ = 4.37, d.f. = 2, p = 0.11, respectively; figure 2B).

When OGs were subset to those annotated as having higher expression in adults than in hatchlings, the results were largely similar. However, effects for OGs annotated as ubiquitously-expressed, although similar in magnitude, were no longer statistically significant for differences in median residuals (χ^2^ = 5.11, d.f. = 2, p = 0.078), nor for probabilities of observing a sequence (χ^2^ = 0.65, d.f. = 2, p = 0.72; figure 2C and D). Only OGs with the testis-region annotation had significant differences between bristle states in median residuals (χ^2^ = 55.86, d.f. = 2, p < 0.001; figure 2C) and in the probability of observing a sequence (χ^2^ = 13.98, d.f. = 2 p < 0.001; figure 2D). Effects for ovary- and tail-region OGs remained non-significant both for differences in median residuals (χ^2^ = 1.1, d.f. = 2, p = 0.58 and χ^2^ = 2.44, d.f. = 2, p = 0.29, respectively) and for the probability in observing a representative sequence (χ^2^ = 0.98, d.f. = 2, p = 0.61 and χ^2^ = 1.29, d.f. = 2, p = 0.53, respectively). These results therefore suggest that the strongest and most consistent response in both analyses is for testis-region annotated OGs.

Some important caveats should be borne in mind. There is substantial variation in assembly quality and completeness across species and this probably explains some of the variation in unobserved sequences. For any given species we cannot exclude the possibility that a transcript is unobserved due to it not being assembled, even though it occurs in the species. Moreover, lower assembly quality due to unassembled genes may result in the mis-labelling of paralogs as orthologs. Ortholog identification typically involves at least an initial search for similar sequences (homologs). If orthologs are unassembled then the most similar sequence will sometimes be a paralog. Unassembled genes might result for related technical reasons, with low-expressed genes being less likely to be sampled by an RNA-Seq strategy. Moreover, gene expression levels may vary across species. Many of the analysed worms were caught in the field and are thus not standardised in condition or environment, which might also result in gene expression variation. In addition, turnover of expression across species, for example, due to selection for different expression patterns, might also results in variation. However, sources of bias from systematic differences between species with different bristle states should produce artefactual results throughout the genome (i.e. in all annotation categories). Together with our substantial efforts to test our method by cross-validation and account for sources of bias and variation in our analyses, our finding of an effect largely isolated to testis-region annotated OGs provides an additional validation of our approach.

Despite the above caveats, a number of OGs (N = 15) can be highlighted as potentially interesting candidates for follow-up studies (table S1 – sheet 7) on the basis of non-overlapping confidence intervals between bristle states (see Methods) from logistic regression analysis in the presence/absence data (table S1 – sheet 6). Of these, 4 OGs are annotated as expressed in the testis-region, all with significantly higher expression in adults compared to hatchlings, making them strong candidates for genes affecting reproduction-related functions. In particular, the transcript Mlig016310.g1, which annotates OG0001273_1_MIortho10 as being expressed in the testis region, has been confirmed, by *in situ* hybridisation (ISH) in *M. lignano*, to be limited in expression to the testes (Arbore et al. 2015; called RNA815_7008 therein). Moreover, RNAi knockdown of this transcript in *M. lignano* produces an aberrant sperm bristle phenotype (Arbore et al. 2015), in which the usually backwards pointing bristles are no longer stably anchored in that position, pointing to a possible role in the formation of the bristle complex (Willems et al. 2004). While a previous study did identify orthologs of this transcript both in a species with bristles (*M. spirale*) and two species with absent bristles (*M. pusillum*, and *M. hystrix*), all showing expression localised to the testes (Brand et al. 2020), no species with absent bristles are represented within this OG in this study. This discrepancy can probably be attributed to differences in the orthology detection approach. Brand et al. (2020) relied exclusively on the sequence similarity-based OrthoFinder pipeline (Emms and Kelly 2015), while we here use orthologs generated from the PDC pipeline of Yang & Smith (2014), which incorporates phylogenetic information from gene trees of initial homolog groups. Thus, previously identified orthogroups may have contained spuriously identified paralogs with similar tissue expression patterns across species that are correctly identified as paralogs in more recent analyses including more species and tree-based methods (Yang & Smith 2014; Wiberg et al. 2021; Brand et al. 2022a). Further validation of these findings will require a more thorough, manual investigation of ortholog and paralog relationships of genes with homology to Mlig016310.g1, validation of their absence from genome sequences, as well as confirmation of expression patterns in species with different bristle states, all of which are well outside the scope of this study. An additional 7, 3, and one candidate OGs are annotated as ubiquitously-expressed, or specific to the ovary- and tail-regions, respectively. No further ISH data are currently available for the transcripts in these OGs. As above, these OGs will require more thorough validation, such as more targeted identification of homologs, orthologs, and paralogs, as well as ISH experiments.

## Conclusions

We explored patterns of missing genes in ∼100 transcriptome assemblies from diverse *Macrostomum* species with contrasting mating strategies. We also expand annotations for *M. lignano* and identify a more stringent set of reproduction-related transcripts with higher expression in adults compared to hatchlings. Our results are consistent with previous findings of rapid sequence evolution at reproduction-related genes, at least for OGs annotated as testis-region specific, and in species with reduced and/or absent sperm bristles, a proxy for the hypodermic mating strategy. Our novel findings also show that species with absent bristles have a lower probability of observing a sequence within OGs annotated as testis-region specific. We suggest that the most parsimonious explanation is that the faster rate of evolution seen also for these OGs results in a higher rate of homology detection failure for such genes, resulting in a signal of rapid evolution of reproduction-related genes also in sequence presence/absence data. Our study highlights at least one exciting candidate OG for further study where representative transcripts have already been associated with phenotypic effects on sperm bristle phenotypes in *M. lignano*. Thus, our results highlight the utility of considering an appropriate null expectation for missing or unobserved genes as well as associating patterns of gene presence/absence with replicated evolutionary events in a phylogenetic context.

## Data accessibility

Transcriptome assemblies and annotated orthogroups forming the basis of these analyses are available from https://doi.org/10.5281/zenodo.4543289 and https://doi.org/10.5281/zenodo.4972188. Raw-sequencing files for the adult-hatchling contrast of *M. lignano* has been deposited with the European Nucleotide Archive (ENA) at EMBL-EBI under accession number PRJEB51403 (https://www.ebi.ac.uk/ena/browser/view/PRJEB51403). Data and analysis scripts used in this study are available from https://doi.org/10.5281/zenodo.6984747.

## Acknowledgements

This work was supported by grants from the Swiss National Science Foundation (grant numbers 31003A_162543 and 310030_184916 to LS). We also thank Lukas Zimmermann for IT support. We thank Elena Finkler for help with the culturing of worms. We would also like to thank Michael Matschiner and Jeremias Brand for valuable discussions on this work. The authors are also grateful to Peter Ladurner for sending specimens of *M. lignano* to produced material for RNA extractions. We are grateful to Christian Beisel, Ina Nissen, Elodie Burcklen Philippe Demougin, and the D-BSSE ETH Zürich Genomics Facility Basel for support in library preparation and sequencing.

## References

Albalat R. & Cañestro, C. (2016). Evolution by gene loss. Nat. Rev. Genet. 17: 379–391.

Arbore, R., Sekii, K., Beisel, C., Ladurner, P., Berezikov, E. & Schärer, L. (2015). Positional RNA-Seq identifies candidate genes for phenotypic engineering of sexual traits. Front. Zool. 12: 14.

Attali, D. (2020). ddpcr: Analysis and Visualization of Droplet Digital PCR in R and on the Web. R package version 1.15. https://CRAN.R-project.org/package=ddpcr

Brand, J.N., Wiberg, R.A.W., Pjeta, R., Bertemes, P., Beisel, C., Ladurner, P. et al. (2020). RNA-Seq of three free-living flatworm species suggests rapid evolution of reproduction-related genes. BMC Genom. 21: 462.

Brand, J.N., Viktorin, G., Wiberg, R.A.W., Beisel, C. & Schärer, L. (2022a). Large-scale phylogenomics of the genus Macrostomum (Platyhelminthes) reveals cryptic diversity and novel sexual traits. Mol. Phylogenet. Evol. 166: 107296.

Brand, J.N., Harmon, L.J. & Schärer, L. (2022b). Frequent origins of traumatic insemination involve convergent shifts in sperm and genital morphology. Evol. Lett. 6: 63–82.

Camacho, C., Coulouris, G., Avagyan, V., Ma, N., Papadopoulos, J., Bealer, K. et al. (2009). BLAST+: architecture and applications. BMC Bioinform. 10: 421.

Charnov, E.L. (1979). Simultaneous hermaphroditism and sexual selection. Proc. Natl. Acad. Sci. U.S.A. 76: 2480–2484.

Cutter, A.D. (2019). Reproductive transitions in plants and animals: selfing syndrome, sexual selection and speciation. New Phytol. 224: 1080–1094.

Cutter, A.D. & Ward, S. (2005). Sexual and temporal dynamics of molecular evolution in C. elegans development. Mol. Biol. Evol. 22: 178–188.

Dapper, A. & Wade, M.J. (2016). The evolution of sperm competition genes: The effect of mating system on levels of genetic variation within and between species. Evolution. 70: 502–511.

Dapper, A. & Wade, M.J. (2020). Relaxed selection and the evolution of reproductive genes. Trends Genet. 36: 640–649.

Drosophila 12 Genomes Consortium (2007). Evolution of genes and genomes on the Drosophila phylogeny. Nature. 450: 203–218.

Emms, D. & Kelly, S. (2015). OrthoFinder: solving fundamental biases in whole genome comparisons dramatically improves orthogroup inference accuracy. Genome Biol. 16: 157.

Fernández R. & Gabaldón, T. (2020). Gene gain and loss across the metazoan tree of life. Nat. Ecol. Evol. 4: 524–533.

Fierst, J.L., Willis, J.H., Thomas, C.G., Wang, W., Reynolds, R.M., Ahearne, T.E., Cutter, A.D. & Phillips, P.C. (2015). Reproductive mode and the evolution of genome size and structure in Caenorhabditis nematodes. PLoS. Genet. 11: e1005323.

Giannakara, A., Schärer, L. & Ramm, S.A. (2016). Sperm competition-induced plasticity in the speed of spermatogenesis. BMC Evol. Biol. 16: 60.

Giannakara, A., & Ramm, S.A. (2017). Self-fertilization, sex allocation and spermatogenesis kinetics in the hypodermically inseminating Macrostomum pusillum. J. Exp. Biol. 220: 1568–1577.

Grudniewska, M., Mouton, S., Grelling, M., Wolters, A.H.G., Kuipers, J., Giepmans, B.N.G. et al. (2018). A novel flatworm-specific gene implicated in reproduction in Macrostomum lignano. Sci. Rep. 8: 3192.

Hahn, M.W., Han, M.V. & Han, S-G. (2007). Gene family evolution across 12 Drosophila genomes. PLoS Genet. 3: e197.

Hadfield, J.D. (2010). MCMC Methods for Multi-Response Generalized Linear Mixed Models: The MCMCglmm R Package. J. Stat. Softw. 33: 1–22.

Haerty, W., Jagadeeshan, S., Kulathinal, R.J., Wong, A., Ravi Ram, K., Sirot, L.K. et al. (2007). Evolution in the fast lane: Rapidly evolving sex-related genes in Drosophila. Genetics. 177: 1321–1335.

Ladurner, P., Schärer, L., Salvenmoser, W., Rieger. R.M., (2005). A new model organism among the lower Bilateria and the use of digital microscopy in taxonomy of meiobenthic Platyhelminthes: Macrostomum lignano, n. sp. (Rhabditophora, Macrostomorpha). J. Zoolog. Syst. Evol. 43: 114–126.

Love, M.I., Huber, W. & Anders, S. (2014). Moderated estimation of fold change and dispersion for RNA-seq data with DESeq2. Genome Biol. 15: 550.

Marie-Orleach, L., Vogt-Burri, N., Mouginot, P., Schlatter, A., Vizoso, D.B., Bailey, N.W. et al. (2017). Indirect genetic effects and sexual conflicts: Partner genotype influences multiple morphological and behavioral reproductive traits in a flatworm. Evolution. 71: 1232–1245.

Martin, M. (2011). Cutadapt removes adapter sequences from high-throughput sequencing reads. EMBnet.Journal. 17: 10–12.

Michiels, N. (1998). Mating conflicts and sperm competition in simultaneous hermaphrodites. Pp 219–254 in T. R. Birkhead and A. P. Møller, eds. Sperm competition and sexual selection. Academic Press. London.

Müller, K. (2020). here: A Simpler Way to Find Your Files. R package version 1.0.1. https://CRAN.R-project.org/package=here

Neme, R., Amador, C., Yildirim, B., McConnell, E. & Tautz, D. (2017). Random sequences are an abundant source of bioactive RNAs or peptides. Nat. Ecol. Evol. 1: 0127.

Patlar, B., Weber, M., Temizyürek, T. & Ramm, S.A. (2020). Seminal fluid-mediated manipulation of post-mating behavior in a simultaneous hermaphrodite. Curr. Biol. 30: 143–149.

Paradis, E. & Schliep, K. (2019). ape 5.0: an environment for modern phylogenetics and evolutionary analyses in R. Bioinformatics. 35: 526–528.

Patro, R., Duggal, G., Love, M.I., Irizzary, R.A. & Kingsford, C. (2017). Salmon provides fast and bias-aware quantification of transcript expression. Nat. Methods. 14: 417–419.

Ramm, S. A., Schlatter, A., Poirer, M. & Schärer, L. (2015). Hypodermic self-insemination as a reproductive assurance strategy. Proc. Royal Soc. B. 282: 20150660.

Revell, L. J. (2012). phytools: An R package for phylogenetic comparative biology (and other things). Methods Ecol. Evol. 3: 217–223.

Rowe, M., Whittington, E., Borziak, K., Ravinet, M., Eroukhmanoff, F., Sætre, G.P. & Dorus, S. (2020). Molecular diversification of the seminal fluid proteome in a recently diverged passerine species pair. Mol. Biol. Evol. 37: 488–506.

Schärer, L., Joss, G. & Sandner, P. (2004). Mating behaviour of the marine turbellarian Macrostomum sp.: these worms suck. Mar. Biol. 145: 373–380.

Schärer, L., Littlewood, D.T.J., Waeschenbach, A., Yoshida, W. & Vizoso, D.B. (2011). Mating behavior and the evolution of sperm design. Proc. Natl. Acad. Sci. U.S.A. 108: 1490–1495.

Schärer, L., Janicke, T. & Ramm, S.A. (2015). Sexual conflict in hermaphrodites. Cold Spring Harb. Perspect. Biol. 7: a017673.

Sanderson, M.J. (2002). Estimating absolute rates of molecular evolution and divergence times: A penalized likelihood approach. Mol. Biol. Evol. 19: 101–109.

Simão, F.A., Waterhouse, R.M., Ioannidis, P., Kriventseva, E.V. & Zbodnov, E.M. (2015). BUSCO: assessing genome assembly and annotation completeness with single-copy orthologs. Bioinformatics. 31: 3210–3212.

Singh, P., Ballmer, D.N., Laubscher, M. & Schärer, L. (2020a). Successful mating and hybridisation in two closely related flatworm species despite significant differences in reproductive morphology and behaviour. Sci. Rep. 10: 12830.

Singh, P., Vellnow, N. & Schärer, L. (2020b). Variation in sex allocation plasticity in three closely related flatworm species. Ecol. Evol. 10: 26–37.

Smith, S.A. & O’Meara, B.C. (2012). treePL: Divergence time estimation using penalized likelihood for large phylogenies. Bioinformatics. 28: 2689–2690.

Swanson, W.J. & Vacquier, V.D. (2002). The rapid evolution of reproductive proteins. Nat. Rev. Genet. 3: 137–144.

Thomas, C.G., Li, R., Smith, H.E., Woodruff, G.C., Oliver, B. & Haag, E.S. (2012). Simplification and desexualization of gene expression in self-fertile nematodes. Curr. Biol. 22: 2167–2172.

Vacquier, V.D. & Swanson, W.J. (2011). Selection in the rapid evolution of gamete recognition proteins in marine invertebrates. Cold Spring Harb. Perspect. Biol. 3: e002931.

Vizoso, D. B., Rieger, G. & Schärer, L. (2010). Goings-on inside a worm: functional hypotheses derived from sexual conflict thinking. Biol. J. Linn. Soc. 99: 370–383.

Weisman, C.M., Murray, A.W. & Eddy, S.R. (2020). Many, but not all, lineage-specific genes can be explained by homology detection failure. PLoS Biol. 18: e3000862.

Wiberg, R.A.W., Brand, J.N. & Schärer, L. (2021). Faster rates of molecular sequence evolution in reproduction-related genes and in species with hypodermic sperm morphologies. Mol. Biol. Evol. 38: 5685–5703.

Wickham, H. (2011). The split-apply-combine strategy for data analysis. J. Stat. Softw. 40: 1–29.

Wickham, H. (2016). ggplot2: Elegant Graphics for Data Analysis. Springer-Verlag. New York.

Wilburn, D.B. & Swanson, W.J. (2016). From molecules to mating: Rapid evolution and biochemical studies of reproductive proteins. J. Proteom. 135: 12–25.

Willems, M., Leroux, F., Claeys, M., Boone, M., Mouton, S., Artois, T. & Borgonie, G. (2008). Ontogeny of the complex sperm in the macrostomid flatworm Macrostomum lignano (Macrostomorpha, Rhabditophora). J. Morphol. 270: 162–174.

Wu, M., Chatterji, S. & Eisen, J.A. (2012). Accounting for alignment uncertainty in phylogenomics. PLoS ONE. 7:e30288.

Wudarski, J., Simanov, D., Ustyantsev, K., De Mulder, K., Grelling, M., Grudniewska, M. et al. (2017). Efficient transgenesis and annotated genome sequence of the regenerative flatworm model Macrostomum lignano. Nat. Comm. 8: 2120

Wudarski, J. Egger, B., Ramm, S.A., Schärer, L., Ladurner, P., Zadesenets, K.S. et al. (2020). The free-living flatworm Macrostomum lignano. EvoDevo. 11: 5.

Yang, Y. & Smith, S.A. (2014). Orthology inference in non-model organisms using transcriptomes and low-coverage genomes: improving accuracy and matrix occupancy for phylogenomics. Mol. Biol. Evol. 31: 3081–3092.

Yin, D., Schwarz, E.M., Thomas, C.G., Felde, R.L., Korf, I.F., Cutter, A.D., Schartner, C.M., Ralston, E.J., Meyer, B.J., Haag, E.S. (2018). Rapid genome shrinkage in a self-fertile nematode reveals sperm competition proteins. Science. 359: 55–61.

Yu G. Smith, D. Zhu, H. Guan, Y. & Tsan-Yuk Lam, T. (2017). ggtree: an R package for visualization and annotation of phylogenetic trees with their covariates and other associated data. Methods Ecol. Evol. 8: 28–36

Yu G. Tsan-Yuk Lam, T. Zhu, H. & Guan. Y. (2018). Two methods for mapping and visualizing associated data on phylogeny using ggtree. Mol. Biol. Evol. 35: 3041–3043

Yu G. (2020). Using ggtree to visualize data on tree-like structures. Curr. Protoc. Bioinform. 69: e96.

Yu G. (2021). ggimage: Use Image in ‘ggplot2’. https://CRAN.R-project.org/package=ggimage

Yu G. Xu, S. & Li, L. (2022). TDbook: Companion Package for the Book “Data Integration, Manipulation and Visualization of Phylogenetic Trees”by Guangchuang Yu (2022, ISBN:9781032233574). https://CRAN.R-project.org/package=TDbook

Zhang, Z., Hambuch, T.M. & Parsch, J. (2004). Molecular evolution of sex-biased genes in Drosophila. Mol. Biol. Evol. 21: 2130–2139.

