## supplementary materials for "Mating strategy predicts gene presence/absence patterns in a genus of simultaneously hermaphroditic flatworms"

### Supplementary text T1: Adult *versus* hatchling gene expression contrast in *M. lignano*

#### *Methods*

8 samples of pooled worms were created; 4 adult pools (~3-4 weeks post-hatching, N = 60 individuals per pool) and 4 hatchling pools (~2-5 days post-hatching, N = 230 individuals per pool). Animals were starved for 24 hours, after which RNA was extracted using NucleoSpin™ RNA XS columns (Macherey-Nagel) for each pool separately. Illumina cDNA libraries were prepared from 100ng total RNA, using the High Throughput TruSeq Stranded mRNA Library Prep Kit (Cat# RS-122-2103, Illumina, San Diego, CA, USA) and the TruSeq RNA CD Index Plate (Cat# 20019792, Illumina). Following 15 cycles of PCR amplification, the libraries were multiplexed, and sequenced on an SP flowcell to produce paired-end 101bp reads (SP 200), on one lane of the NovaSeq6000 platform at the Genomics Facility Basel (D-BSSE, ETH Zurich). Raw reads are deposited in the European Nucleotide Archive (ENA) at EMBL-EBI under accession number PRJEB51403 (https://www.ebi.ac.uk/ena/browser/view/PRJEB51403; see also table S2 for individual sample information).

We trimmed remaining TruSeq adapter sequences using cutadapt (v. 3.0; Martin 2011) and then quantified expression of all transcripts in the *M. lignano* transcriptome (Mlig_RNA_3_7_DV1.v3.coregenes; Grudniewska et al. 2018) using the quasi-mapping approach of Salmon (v. 1.4.0; Patro et al. 2017). Salmon discarded N = 2,134 transcripts as exact duplicates of other transcripts. We performed minimal filtering to exclude transcripts with a read count of 0 in all samples (N = 8,859), as well as all transcripts with read counts < 10 across all samples (N = 3,676). The final set of transcripts (N = 75,346) represents 83.5% of all transcripts. Finally, we tested for differential expression between adults and hatchlings with the DESeq2 R package (v. 1.24.0; Love et al. 2014). We called transcripts as differentially expressed if they had an FDR corrected p-value < 0.05. We then identified all transcripts that were differentially expressed *and* had higher expression in adults compared to hatchlings ("Up-in-A") or higher expression in hatchlings compared to adults ("Up-in-H"). Orthogroups (OGs; see main text) that contained *M. lignano* transcripts with higher expression in adults were then also annotated as Up-in-A.

#### *Results & Discussion*

Of the final set of 75,346 *M. lignano* transcripts, 29,947 were differentially expressed between adults and hatchlings (figure S2; table S2 – sheet 3). Of these, 17,728 are called as Up-in-A, and 12,219 are called as Up-in-H, representing 23.5% and 16.2% of the final set, respectively. As in previous studies of other *Macrostomum* species, there were more transcripts annotated as Up-in-A than Up-in-H (Brand et al. 2020). Additionally, the large group of transcripts with much higher expression in adults seen in the region below the diagonal in figure S2 is also observed in other species, indicating the presence of a conserved set of highly adult-specific or adult-biased transcripts (Brand et al. 2020). Note that while the observed differences may reflect differences in expression within cells, they may also arise due to differences in the relative amounts of different tissue types (i.e. hatchlings and adults have different shapes, and the former have no gonads yet). The overlap between OGs with annotation categories is also noteworthy. As expected, most (74.2%) *M. lignano* transcripts with testis-region annotations also had higher expression in adults compared to hatchlings. In contrast, a relatively small percentage (19.4%) of transcripts annotated as ubiquitously expressed, had higher expression in adults. Overall, transcripts annotated as expressed in body regions containing reproductive tissues (reproduction-related genes) had higher percentages (42.9-74.2%) of transcripts with higher expression in adults (table S3). Such a pattern is only expected if transcripts annotated on the basis of their expression in body regions containing reproductive tissues are indeed from predominantly reproduction-related genes. Meanwhile, ubiquitously-expressed transcripts should not be particularly enriched for transcripts with reproductive functions. These patterns are similar also when considering the overlap in annotation categories for OGs annotated on the basis of *M. lignano* transcripts (table S3). Overall, we conclude that the intersection of transcripts expressed in body regions containing reproductive tissues and also expressed at higher levels in adults compared to hatchlings represents a set of more stringently annotated reproduction-related transcripts. In the main text, we always present results on the full set of reproduction-related transcripts as well as the more stringently annotated set.

**Table S3.** The number of transcripts and orthogroups (OGs) with different annotations, based on their positional expression in the testis-, ovary-, or tail-region of *M. lignano* (as well as those that are ubiquitously expressed). Also shown are the number and percentage (%; of the total in each positional category) of transcripts/OGs with additional annotation based on whether they have higher expression in adults compared to hatchlings in *M. lignano* (Up-in-A).

| | | **Testis-region** | **Ovary-region** | **Tail-region** | **Ubiquitously expressed** |
|---|---|---|---|---|---|
| | **Up-in-A** | 2911 | 355 | 585 | 9170 |
| **Transcripts** | **(%)** | (74.2) | (42.9) | (69.9) | (19.4) |
| | **Total** | 3924 | 828 | 837 | 47371 |
| | **Up-in-A** | 499 | 94 | 42 | 59 |
| **OGs** | **(%)** | (76.4) | (68.6) | (36.9) | (22.3) |
| | **Total** | 653 | 137 | 117 | 264 |

**Supplementary text T2: Similarity score decay models excluding species with high rates (>50%) of missing BUSCO genes.**

There was variation in the BUSCO completeness scores across the transcriptome assemblies we used, with some assemblies having high rates of missing BUSCO genes when using the metazoa_odb10 database (Simão et al. 2015). The expectations for BUSCO score completeness using the metazoa_odb dataset are ~80% for flatworm assemblies. This level of missing genes is observed even for the best, genome-guided, transcriptome assemblies available (including for *M. lignano* in Wudarski et al. 2017 and Grudniewska et al. 2018). This may be due to the poor representation of Lophotrochozoa in the metazoan_odb10 dataset. Although our analysis accounts for the variation in transcriptome assembly quality by applying lower weights to data from assemblies with high rates of missing genes, we here evaluate whether our results are robust to the complete exclusion of low-quality assemblies. We do this by sub-setting our data to exclude assemblies with >50% missing BUSCO genes. We reason that this is a good threshold because a) remaining species will have more BUSCO genes present than absent, and b) it is only a 30 percentage point difference from the "expected" maximum of ~80% for these species. We then fit decay models and compute residuals for all remaining data as described in the main text.

*Results & Discussion*

We find that the results remain largely the same as in the main analysis (figure S6). Although the number of species included in the analysis is substantially reduced, the number of orthogroups (OGs) remains very similar. This is because the species with low BUSCO scores are not contributing to much to this analysis in the first place. There remains a significant effect of bristle state for OGs with the testis-region annotation (Kruskal-Wallis $\chi^2$ =35.11, d.f. = 2, $p < 0.001$). For OGs annotated as ubiquitously-expressed there was also a significant effect (Kruskal-Wallis $\chi^2$ = 8.12, d.f. = 2, p = 0.02). However, the pairwise contrast between species with present bristles and species with reduced bristles was no longer significant, probably as a consequence of the increased variance observed in species with reduced bristles (figure S6). There were no significant effects for OGs with the ovary- or tail-region annotations (Kruskal-Wallis $\chi^2$ =2, d.f. = 2, p = 0.37 and Kruskal-Wallis $\chi^2$ = 3.73, d.f. = 2, p = 0.16, respectively). We therefore conclude that our results and conclusions are robust to the exclusion of low-quality transcriptome assemblies with high rates of missing BUSCO genes.

**Supplementary text T3: Validation of our approach to fit models using one group of species and to evaluate their fit in another group species not included in the original model fits.**

*Methods*

In order to validate our methodological approach, we perform a type of cross-validation test. We first subset our data to only species with bristles present. We then identify all orthogroups (OGs) with representative sequences and bit-scores from at least $n = 10$ species. For each OG, we then split the species (rounding down for the 'fitting' set, if $n$ is uneven) into a 'fitting' ($n_f$) set and a 'testing' ($n_t$) set (by sampling from the total set of species without replacement). We always include *M. lignano* in the 'fitting' set to ensure that our validation is comparable to the real data-set where it is also always included. We then test for an overall difference in evolutionary distance from *M. lignano* between the 'fitting' species and the 'testing' species. If the difference is significant (at a threshold of $p < 0.1$), we take a new random sample of species. Finally, we use the 'fitting' set to fit a model of decay in bit-scores with evolutionary distance exactly as described in the main text. We then compute residuals from the fitted line for all species in the 'fitting' and the 'testing' sets, as in the main text, and compare residuals within each annotation category. If our method did not systematically biased toward producing larger residuals in the 'testing' sets, we should not observe a statistically significant difference in residuals between the fitting and the testing sets.

*Results & Discussion*

Sub-setting species and OGs to include only species with bristles and only OGs with $n \geq 10$ representative species resulted in final sample sizes of N = 590 for OGs annotated as testis-region ( N = 332), ovary-region ( N = 72), tail-region ( N = 50), and ubiquitously-expressed ( N = 136). Kruskal-Wallis rank-sum tests showed no significant differences in residuals between 'fitting' and 'testing' species sets for any annotation category (figure S7). We thus conclude that our approach of evaluating the fit of a set of species to a model fit using another set of species does not produce a systematic bias in average residuals.

**Supplementary text T4: Similarity score decay models using normalised bit-scores and phylogenetically controlled regression.**

*Methods*

We here attempted to overcome some limitations of the approach described by Weisman et al. (2020; see the main text and figure 4A). In order to remove the effect of variation in sequence length on bit-scores we adopted a normalisation procedure similar to Emms and Kelly (2015). We placed all bit-scores into bins on the basis of $L_{qh}$ where,

$$L_{qh} = L_q * L_h$$

here $L_q$ is the length of the query sequence and $L_h$ is the length of the hit sequence. As in Emms and Kelly (2015), each bin was constructed to contain ~1000 hits. We then used the top 5% of bit-scores from each bin (figure S4B) to fit a regression model,

$$log_e(B_{qh}) = a\ log_e(L_{qh}) + b$$

where $B_{qh}$ is the bit-score between sequences *q* and *h.* We then normalise all bit-scores according to,

$$B'_{qh} = B_{qh}/e^b L^a_{qh}$$

such that $B'_{qh}$ is the bit-score divided by the best bit-score that could be expected from hits between sequences of the same length class. The resulting normalised scores did not show a dependence on length (figure S4C). Note that it is still possible to fit a decay function to these data, but the explicit model of Weisman et al. (2020), including the expected variance, may no longer apply. We therefore rely on probabilities of detection from the initial model fitting, and instead here only compare the patterns of residuals across bristles states and annotation groups.

To account for similarity of normalised bit-scores among closely related species, we aimed to control for phylogeny. With the ultrametric tree from the main text we used the R package "MCMCglmm" (v. 2.32; Hadfield 2010) to fit phylogenetic linear mixed models with $log_e(B'_{qh})$ as the dependent variable, and distance from *M. lignano* as an independent linear predictor, and included the covariance matrix defined by the phylogeny as a random effect. In order to not include genetic distance from *M. lignano* in the model twice (once as a fixed effect, and once in the phylogenetic covariance matrix) we removed the data for *M. lignano* from all analyses. As above, to fit these models, we only used species with bristles present, and only OGs with data from ≥ 5 species. As before, OGs that produced $R$ values > 0 (N = 58/915, 6.3%) were treated as spurious and not considered further. We then computed residuals for all species, by subtracting the empirical normalised bit-score from that predicted by the linear equation

$$log_e(B'_{qh}) = c + bx$$

where $c$ and $b$ are, respectively, the posterior estimates of the intercept and the slope from the phylogenetic linear mixed model. Again, we only used residuals computed for species with genetic distances to *M. lignano* < 0.7. We summarised residuals as the median across all species of a bristle state within an OG and tested for overall differences between bristle states, across OGs, using Kruskal-Wallis rank-sum tests and Dwass-Steele-Critchlow-Fligner all-pairs tests for *post-hoc* testing of individual contrasts.

*Results & Discussion*

Results using bit-scores normalised by sequence lengths and accounting for phylogeny in estimating the decay rates were qualitatively similar to those presented in the main text (figure S8A; table S1 – sheet 5). For testis-region transcripts there was a significant effect of bristle state ($\chi^2$ = 45.53, p < 0.001). Species with absent bristles had the most negative residuals overall, and species with reduced bristles had more negative residuals than species with bristles, but more positive than species with absent bristles (figure S8A). For OGs annotated as ubiquitously expressed, there was also a significant effect of bristle state ($\chi^2$ = 8.66, p = 0.01), driven by differences between species with reduced bristles and species with absent bristles (figure S8A). Effects of bristle states in ovary- ($\chi^2$ = 0.94, p = 0.62) and tail- ($\chi^2$ = 0.19, p = 0.91) region annotated OGs remain non-significant. These results hold also when restricting this analysis to OGs annotated as having higher expression in adults except that there is no longer a significant effect of bristle state for OGs annotated as ubiquitously expressed (figure S8B).

**Supplementary figures and legends**

**Figure S1. (see the separate .pdf files)** Gene presence/absence patterns for OGs with **A)** testis-region, **B)** ovary-region, and **C)** tail-region annotations, as well as OGs annotated as **D)** ubiquitously expressed. In panels A-D, the columns represent different OGs, and the rows represent transcriptome assemblies from different species. Each tile is coloured in red if a representative sequence is present in the given species for the given OG, if a representative is not present the tile is instead coloured by the bristle state of the species. Each panel also shows the phylogenetic tree with points at the tips coloured by the bristle states (green – present; pale green – reduced; yellow – absent). Tip labels give a short version of the species names for plotting convenience (see table S1 – sheet 3 for full species names), the tip labels also give the number and proportion (in brackets) of OGs with a representative sequence for that species, i.e. the number and proportion of tiles that are red. In **E)** the percentage of OGs with a representative sequence from each species is plotted as function of genetic distance from *M. lignano*.

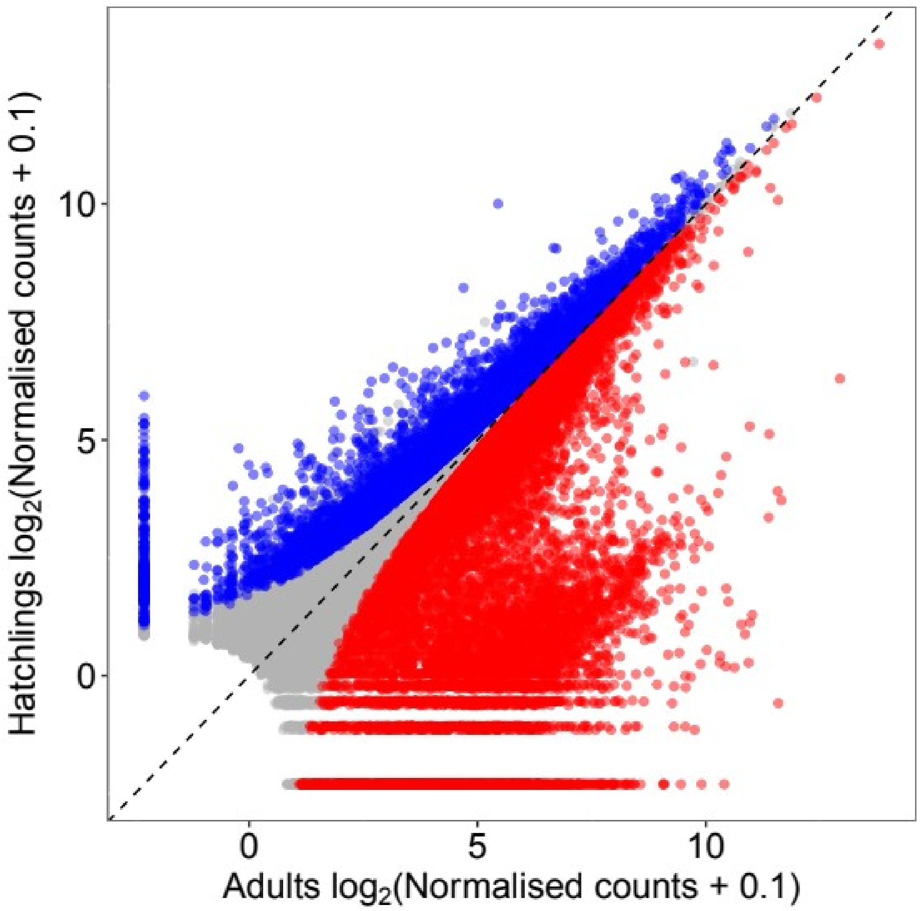


**Figure S2** Comparison of mean $\log_2()$ normalised counts for transcripts across replicates of adult and hatchlings pools. Red and blue points, below and above the diagonal (dashed, black line), are those transcripts that have significantly different expression (FDR p-value < 0.05) between adults and hatchlings. Grey points are transcripts that are not differentially expressed.

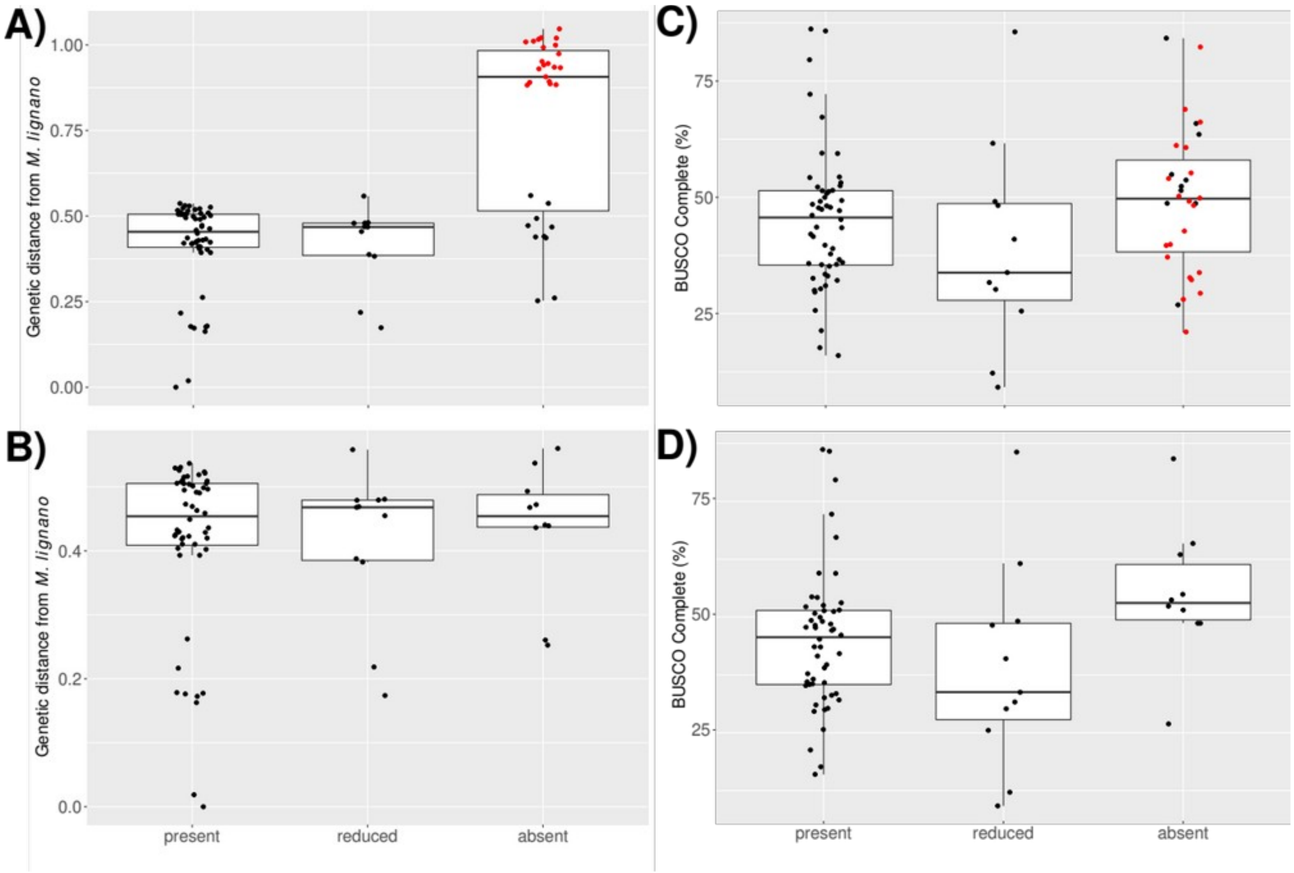


**Figure S3.** The distributions of the genetic distance to *M. lignano* (A and B) and the BUSCO completeness scores (C and D) for species with different bristle states, both including and excluding the species with genetic distances from *M. lignano* > 0.7 (i.e. the "hypodermic mating clade", red points; yellow box in Figure 1A). **A)** There is a significant effect of bristle state on genetic distance when all species are included (ANOVA $F_{2,\ 91} = 43.48$, $p < 0.001$), but not **B)** when the species in the "hypodermic mating clade" are removed ($F_{2,\ 70} = 0.12$, $p = 0.89$). **C)** There is no significant effect of bristle state on BUSCO completeness scores when all species are included ($F_{2,\ 91} = 1.87$, $p = 0.16$), but **D)** when the species in the "hypodermic mating clade" are removed there is a marginally non-significant effect, with species with absent bristles tending to have on average higher completeness than species with reduced or present bristles ($F_{2,\ 70} = 2.68$, $p = 0.07$). Note that statistical tests are indicative only as individual points are not phylogenetically independent.

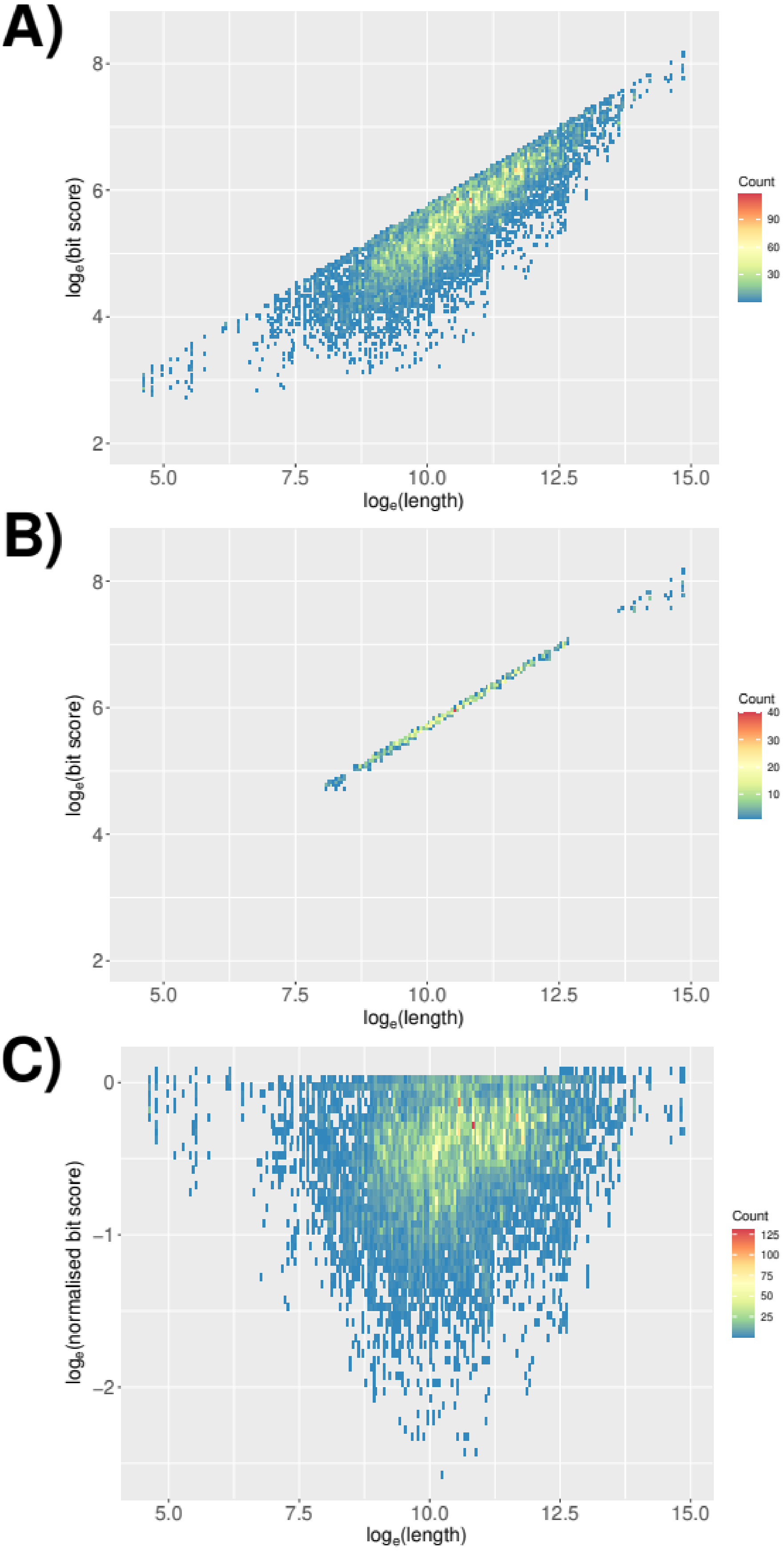


**Figure S4. A)** The relationship between bit-scores and the length of sequences before normalisation. **B)** The same data as in **A)** but only showing the top 5% of data points in each length bin (see Methods). **C)** The relationship between bit-scores and the length of sequences following the normalisation procedure (see

Methods). In all panels, each tile represents a bin of size 0.05 along each axis, the colour represents the number of data points within each bin.

**Figure S5 A-D. (see separate .pdf files)** Individual similarity score decay plots for orthogroups (OGs) annotated as expressed in **A)** the testis-region, **B)** the ovary-region, **C)** the tail-region, or **D)** annotated as ubiquitously expressed. Each panel shows the $log_e$(bit-score) as a function of increasing evolutionary distance from *M. lignano.* Inset text in each panel gives the $R^2$ value from a linear regression, as well as the exponential decay function with estimated parameter values. If the decay function is given in red coloured font, the *R* parameter (see Methods) was positive and the OG was not considered further. Points are shown also for all species that lack a representative sequence in each OG at the bottom of each panel (split by bristle state), separated from the rest of the points by a grey dashed line. The horizontal black dashed line denotes the estimated bit-score threshold value to achieve an e-value of 0.001 in a BLAST search of the full database (see Methods). The red dashed line gives the line of best-fit, and the red area around the dashed line gives the area corresponding to ±3 standard deviations around the best-fit line.

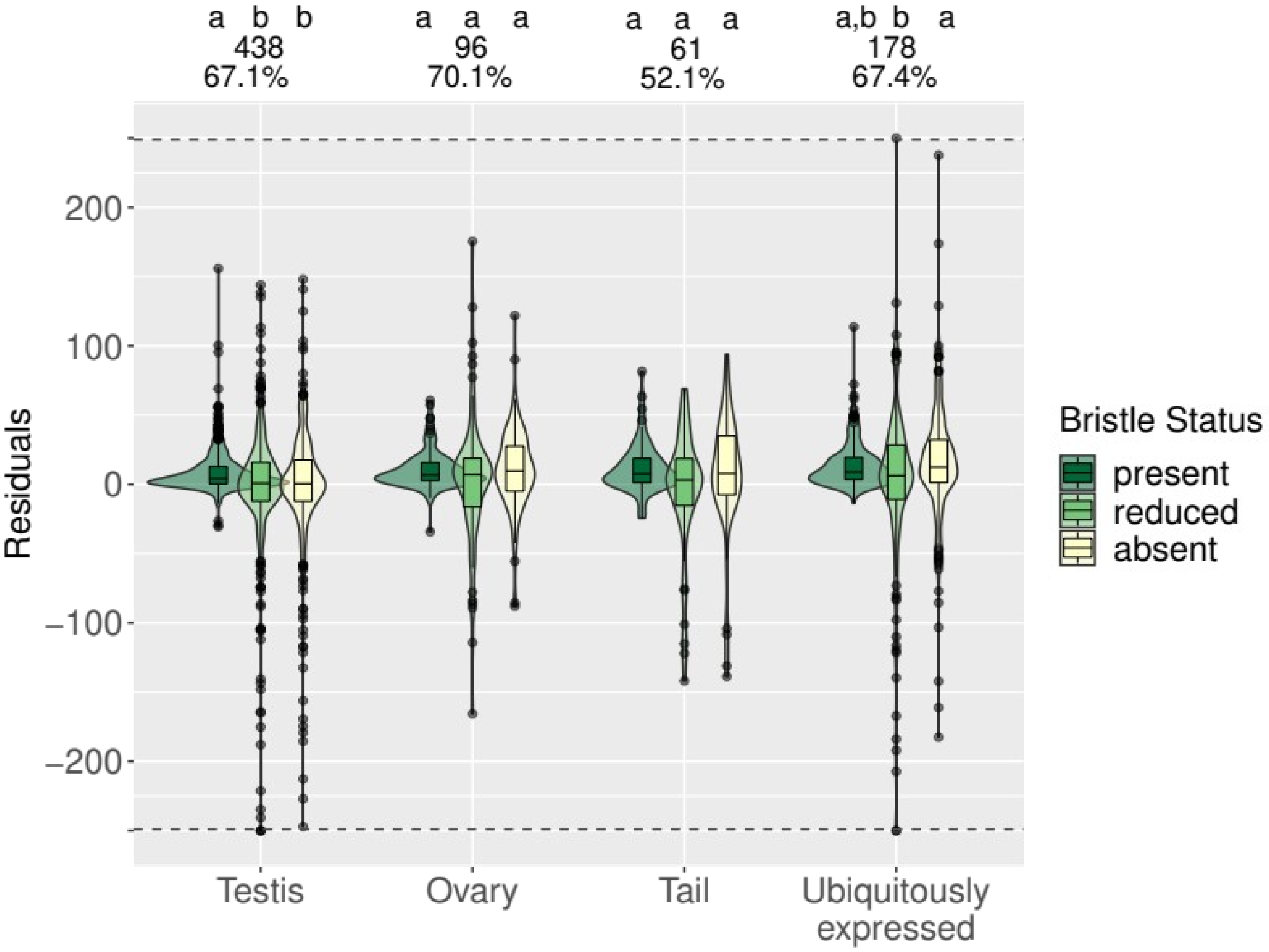
a b b
438
67.1%
a a a
96
70.1%
a a a
61
52.1%
a,b b a
178
67.4%
Residuals
200
100
0
−100
−200
Testis
Ovary
Tail
Ubiquitously expressed
Bristle Status
present
reduced
absent

**Figure S6.**

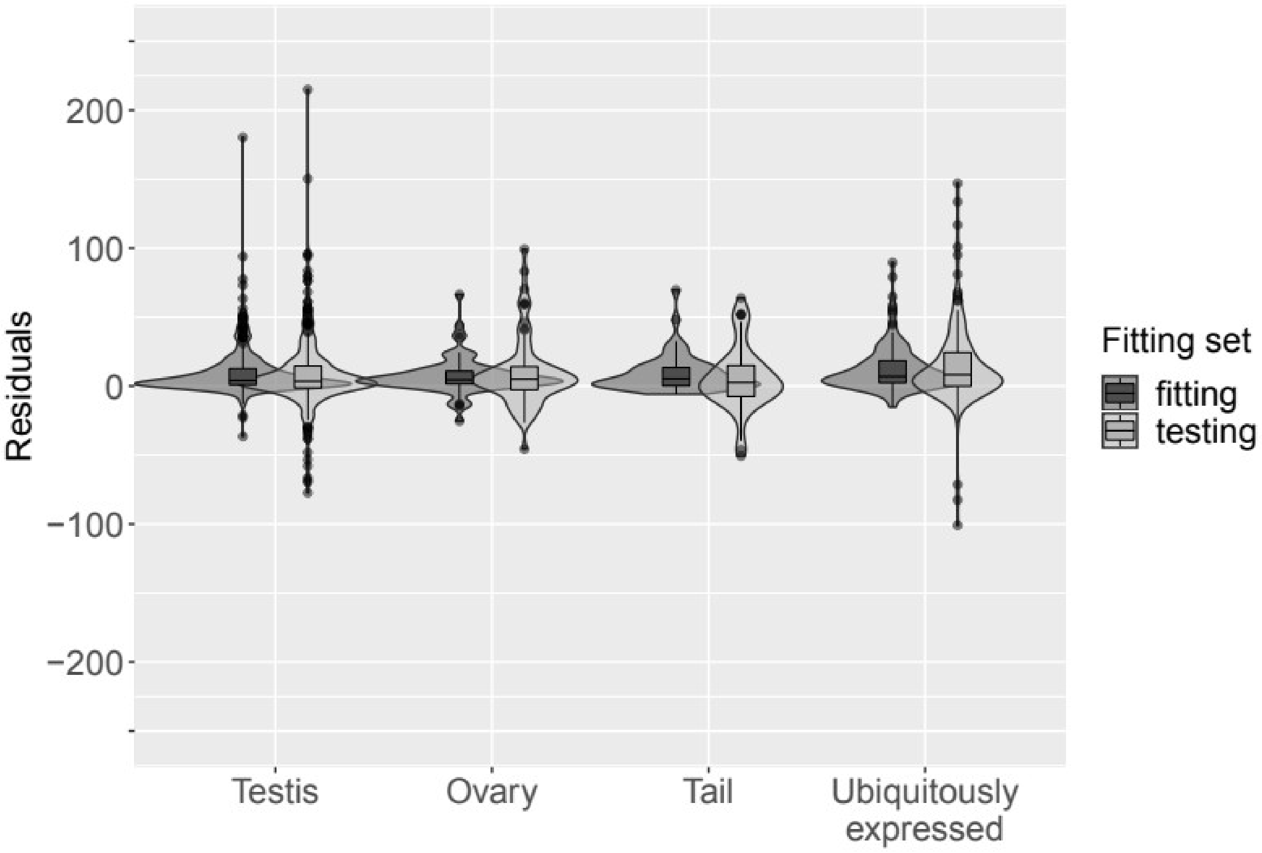


**Figure S7.** Distributions across orthogroups with different annotations of the median residuals for species within the ‘fitting’ and ‘testing’ sets. See supplementary text T2 for the methods and results.

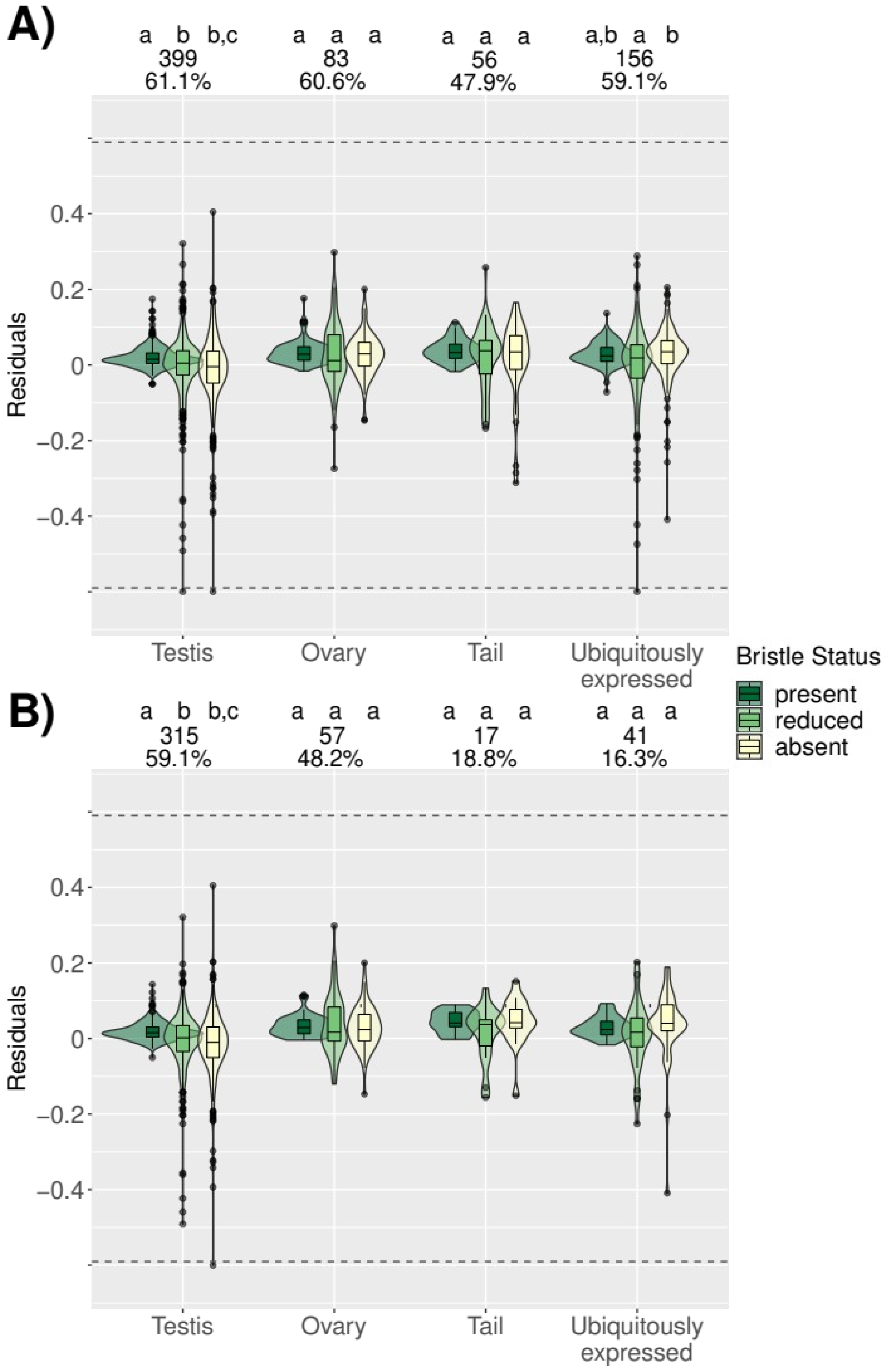


**Figure S8.** Distributions across orthogroups with different annotations of the median residuals for species with different bristle states, when the data were analysed using normalised bit-scores and phylogenetically controlled regression (see main text for a simpler analysis). Results are shown for

**A)** All OGs and **B)** OGs annotated also as having higher expression in adults compared to hatchlings. Large points indicate the mean of the distribution. Letters above the data for each annotation category give the results of Dwass-Steele-Critchlow-Fligner all-pairs *post-hoc* tests, with different letters denoting significant ($p < 0.05$) differences between bristle states. The inset numbers above each group of boxplots give the number and percent of all OGs in each annotation group for which decay models were fit (see also table S3). To make the figure clearer, OGs with values < -0.6 ($N = 3$) and > 0.6 ($N = 0$) have been plotted at $y = -0.6$ and $y = 0.6$ respectively, and separated from the remaining points by dashed horizontal lines.

**References**

Brand, J.N., Wiberg, R.A.W., Pjeta, R., Bertemes, P., Beisel, C., Ladurner, P. & Schärer, L. (2020). RNA-Seq of three free-living flatworm species suggests rapid evolution of reproduction-related genes. *BMC Genom.* 21: 462.

Brand, J.N., Viktorin, G., Wiberg, R.A.W., Beisel, C. & Schärer, L. (2022a). Large-scale phylogenomics of the genus *Macrostomum* (Platyhelminthes) reveals cryptic diversity and novel sexual traits. *Mol. Phylogenet. Evol.* 166: 107296.

Brand, J.N., Harmon, L.J. & L. Schärer, L. (2022b). Frequent origins of traumatic insemination involve convergent shifts in sperm and genital morphology. *Evol. Lett.* 6: 63-82.

Emms, D. & Kelly, S. (2015). OrthoFinder: solving fundamental biases in whole genome comparisons dramatically improves orthogroup inference accuracy. *Genome Biol.* 16: 157.

Grudniewska, M., Mouton, S., Grelling, M., Wolters, A.H.G., Kuipers, J., Giepmans, B.N.G. *et al.* (2018). A novel flatworm-specific gene implicated in reproduction in *Macrostomum lignano*. *Sci. Rep.* 8: 3192.

Hadfield, J.D. (2010). MCMC Methods for Multi-Response Generalized Linear Mixed Models: The MCMCglmm R Package. *J. Stat. Softw.* 33: 1-22.

Love, M.I., Huber, W. & Anders, S. (2014). Moderated estimation of fold change and dispersion for RNA-seq data with DESeq2. *Genome Biol.* 15: 550.

Martin, M. 2011. Cutadapt removes adapter sequences from high-throughput sequencing reads. *EMBnet.Journal.* 17: 10-12.

Patro, R., Duggal, G., Love, M.I., Irizzary, R.A. & Kingsford, C. (2017). Salmon provides fast and bias-aware quantification of transcript expression. *Nat. Methods.* 14: 417-419.

Sanderson, M. J. 2002. Estimating absolute rates of molecular evolution and divergence times: A penalized likelihood approach. *Mol. Biol. Evol.* 19: 101–109.

Simão, F.A., Waterhouse, R.M., Ioannidis, P., Kriventseva, E.V. & Zbodnov, E.M. (2015). BUSCO: assessing genome assembly and annotation completeness with single-copy orthologs. *Bioinformatics*. 31: 3210-3212.

Smith, S.A., & O'Meara, B.C. (2012). treePL: Divergence time estimation using penalized likelihood for large phylogenies. *Bioinformatics.* 28: 2689–2690.

Weisman, C.M., Murray., A.W. & Eddy, S.R. (2020). Many, but not all, lineage-specific genes can be explained by homology detection failure. *PLoS Biol.* 18: e3000862.
