## Supplementary figures and images for "Mating strategy predicts gene presence/absence patterns in a genus of simultaneously hermaphroditic flatworms"

### figure S1A

A) Testis region

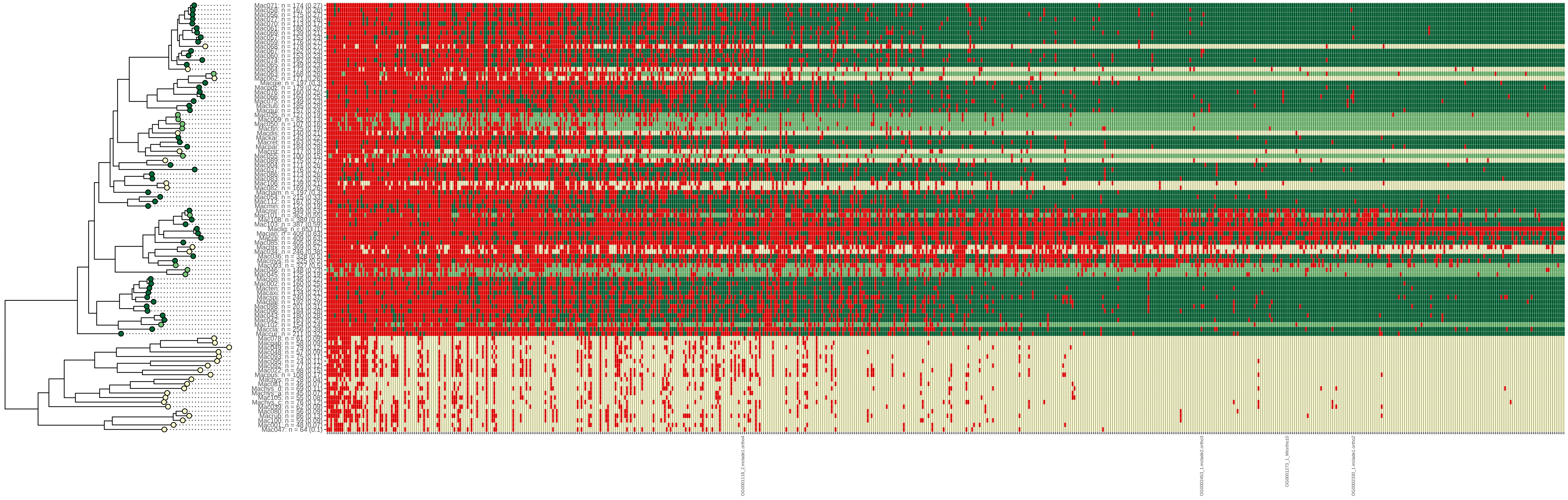

### figure S1B

B) Ovary region

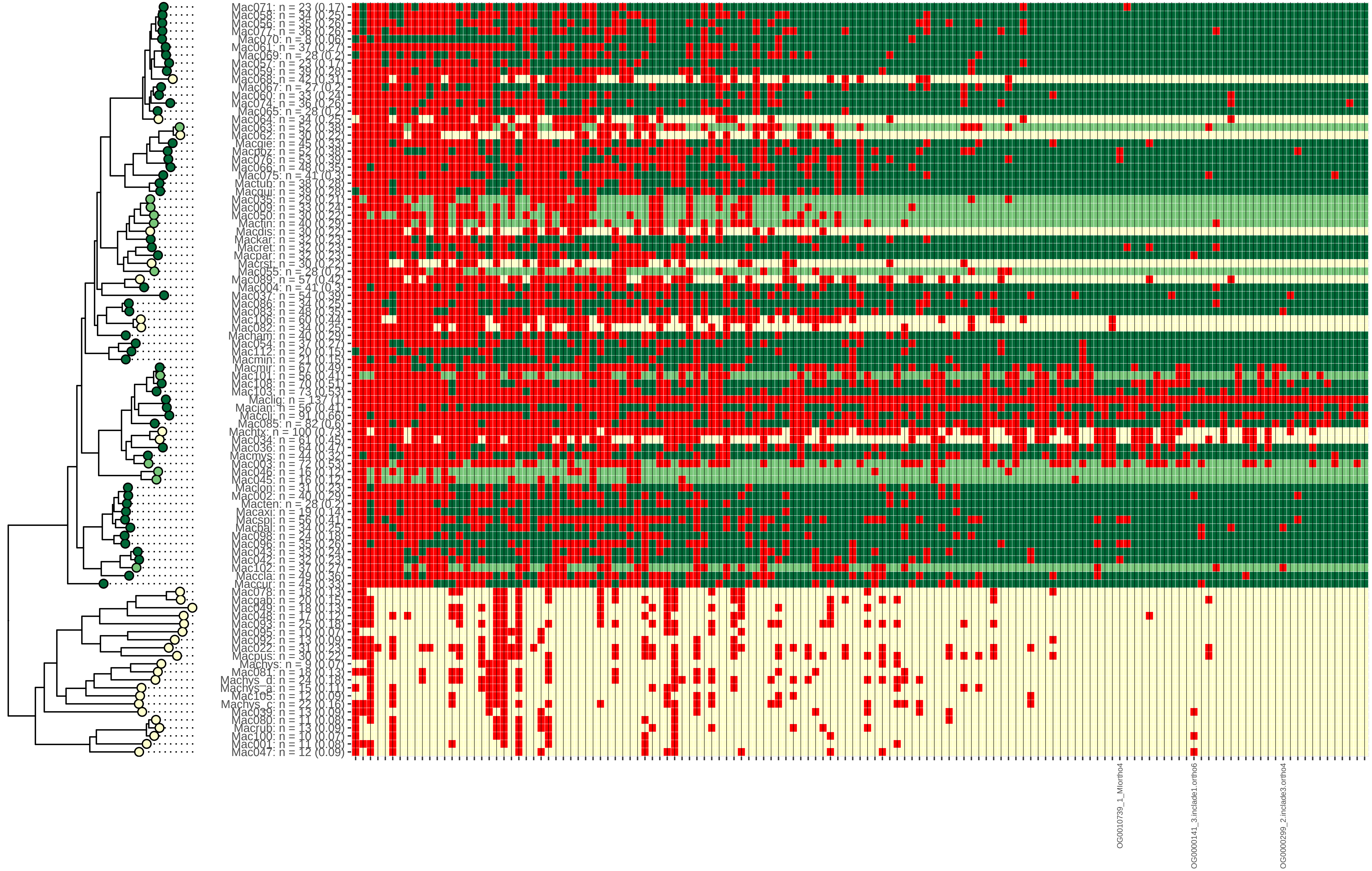

### figure S1C

C) Tail region

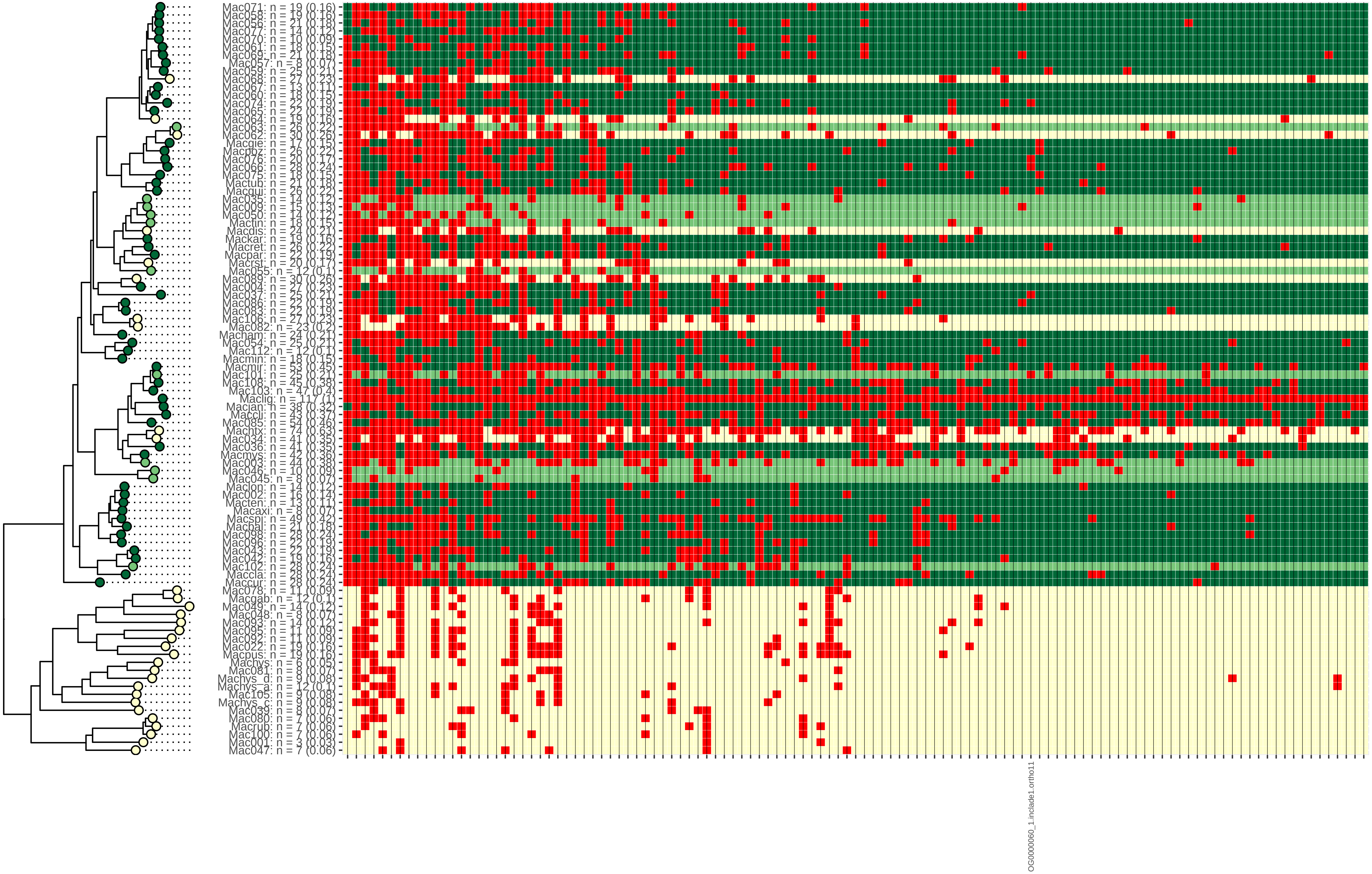

### figure S1D

D) Ubiquitously expressed

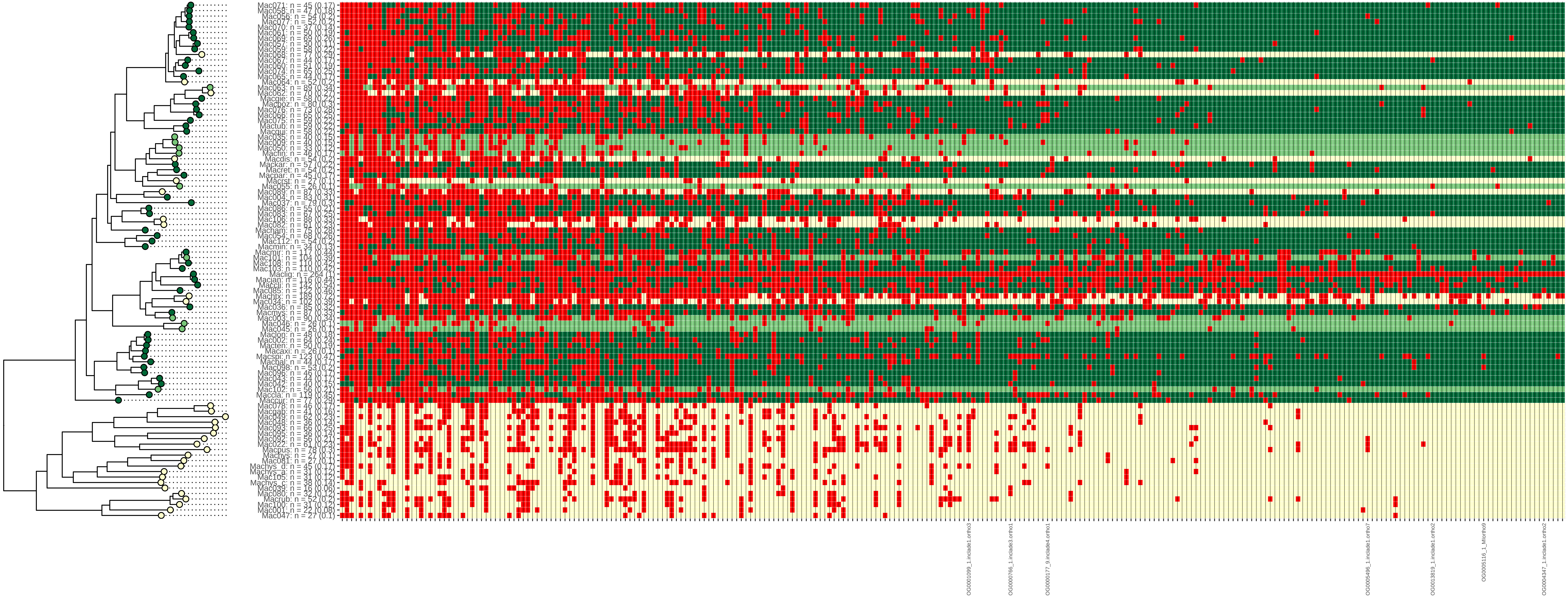

### figure S1E

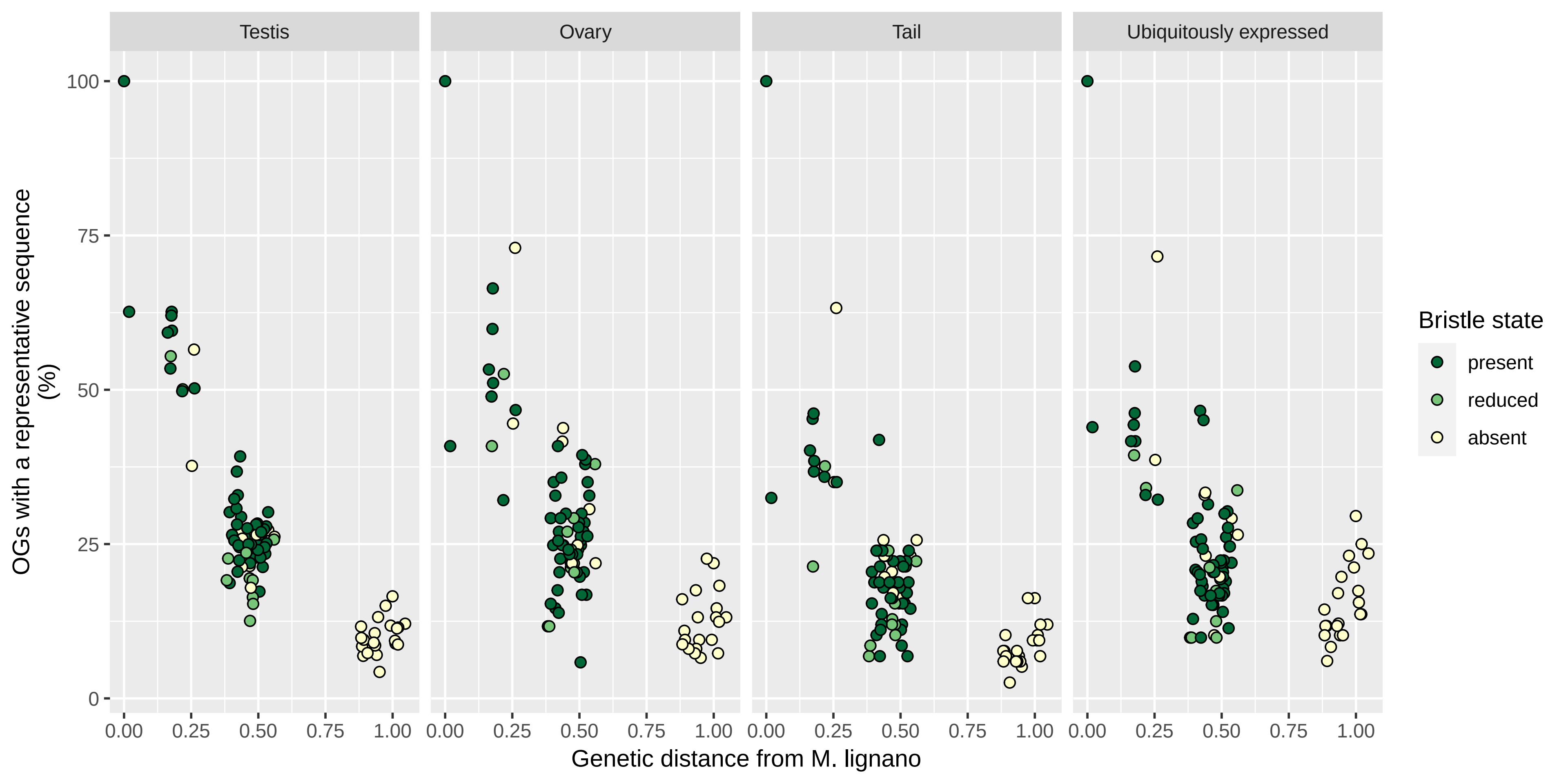
