## Supplementary material for "Mating strategy predicts gene presence/absence patterns in a genus of simultaneously hermaphroditic flatworms": figure S5D

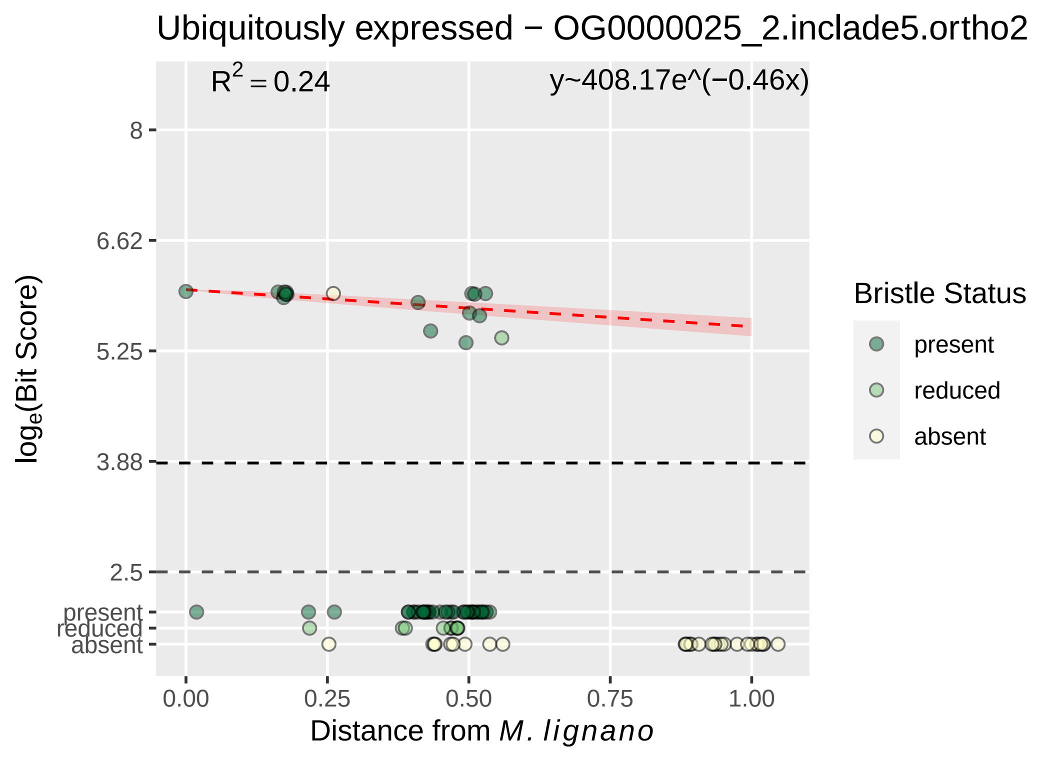

Ubiquitously expressed − OG0000025_2.inclade5.ortho2
$R^2 = 0.24$
y~408.17e^(−0.46x)
8
6.62
5.25
3.88
2.5
present
reduced
absent
$log_e$(Bit Score)
0.00
0.25
0.50
0.75
1.00
Distance from *M. lignano*
Bristle Status
present
reduced
absent

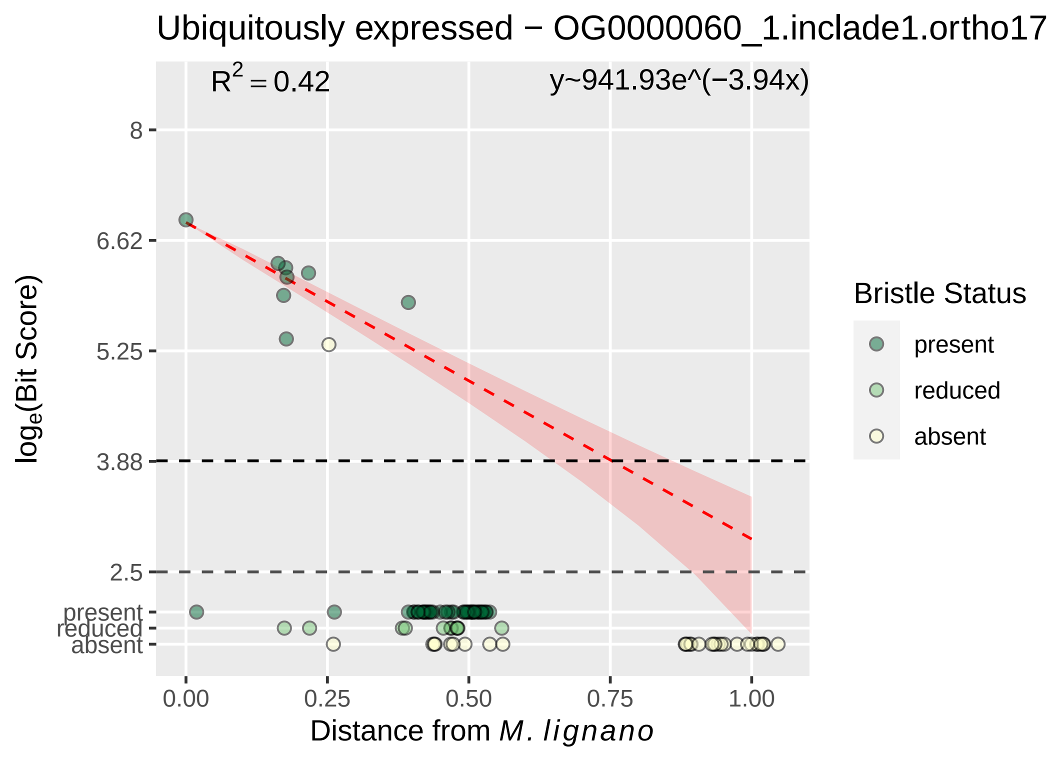

Ubiquitously expressed − OG0000060_1.inclade1.ortho17
R^2 = 0.42
y~941.93e^(−3.94x)
8
6.62
5.25
3.88
2.5
present
reduced
absent
0.00
0.25
0.50
0.75
1.00
log_e(Bit Score)
Distance from M. lignano
Bristle Status
present
reduced
absent

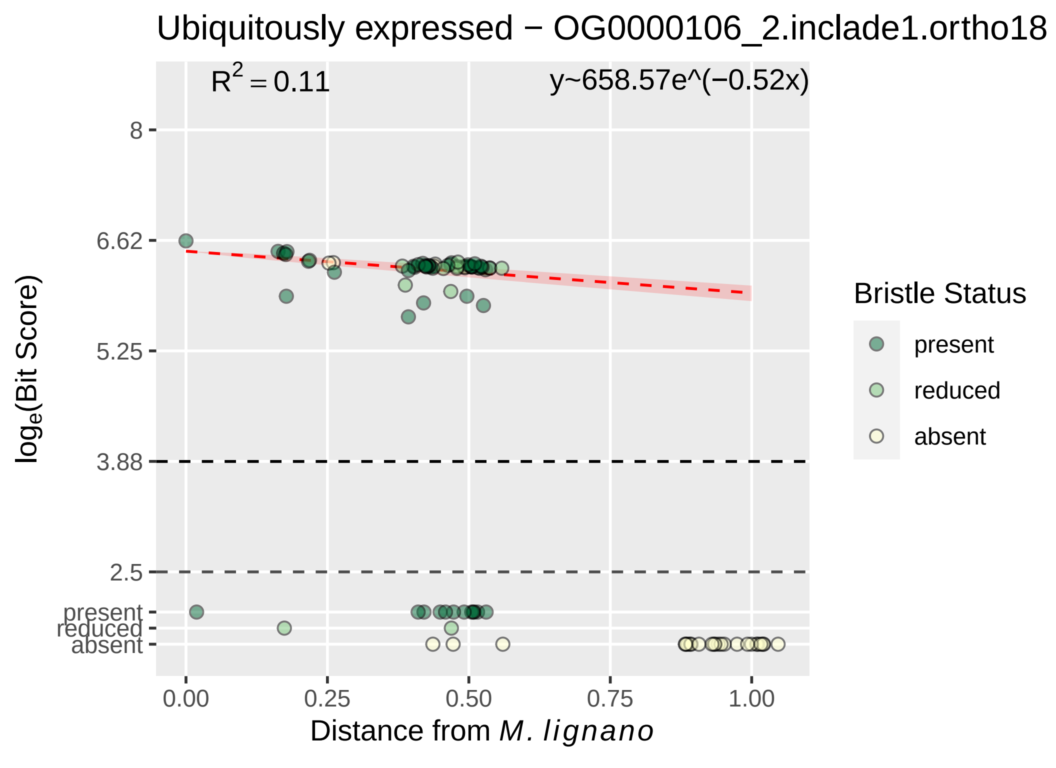
Ubiquitously expressed − OG0000106_2.inclade1.ortho18
$R^2 = 0.11$
y~658.57e^(−0.52x)
8
6.62
5.25
3.88
2.5
present
reduced
absent
$log_e$(Bit Score)
0.00
0.25
0.50
0.75
1.00
Distance from *M. lignano*
Bristle Status
present
reduced
absent

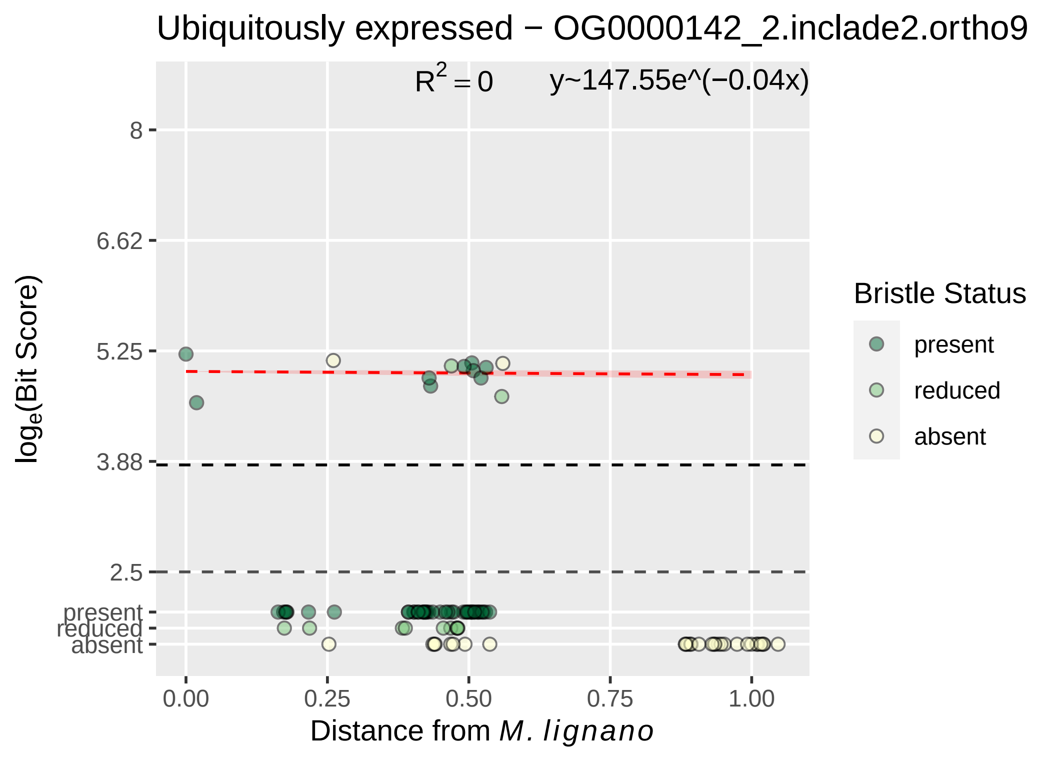

Ubiquitously expressed − OG0000142_2.inclade2.ortho9
R^2 = 0
y~147.55e^(−0.04x)
8
6.62
5.25
3.88
2.5
present
reduced
absent
0.00
0.25
0.50
0.75
1.00
Distance from M. lignano
loge(Bit Score)
Bristle Status
present
reduced
absent

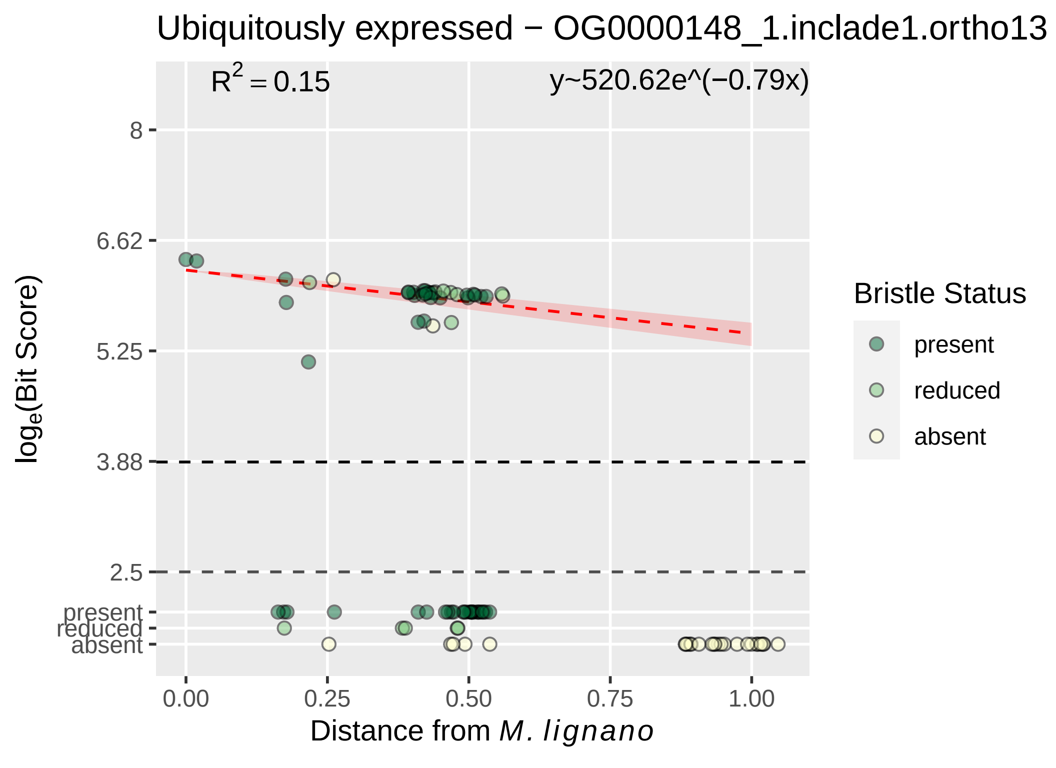

Ubiquitously expressed − OG0000148_1.inclade1.ortho13
$R^2 = 0.15$
y~520.62e^(−0.79x)
8
6.62
5.25
3.88
2.5
present
reduced
absent
0.00
0.25
0.50
0.75
1.00
Distance from M. lignano
loge(Bit Score)
Bristle Status
present
reduced
absent

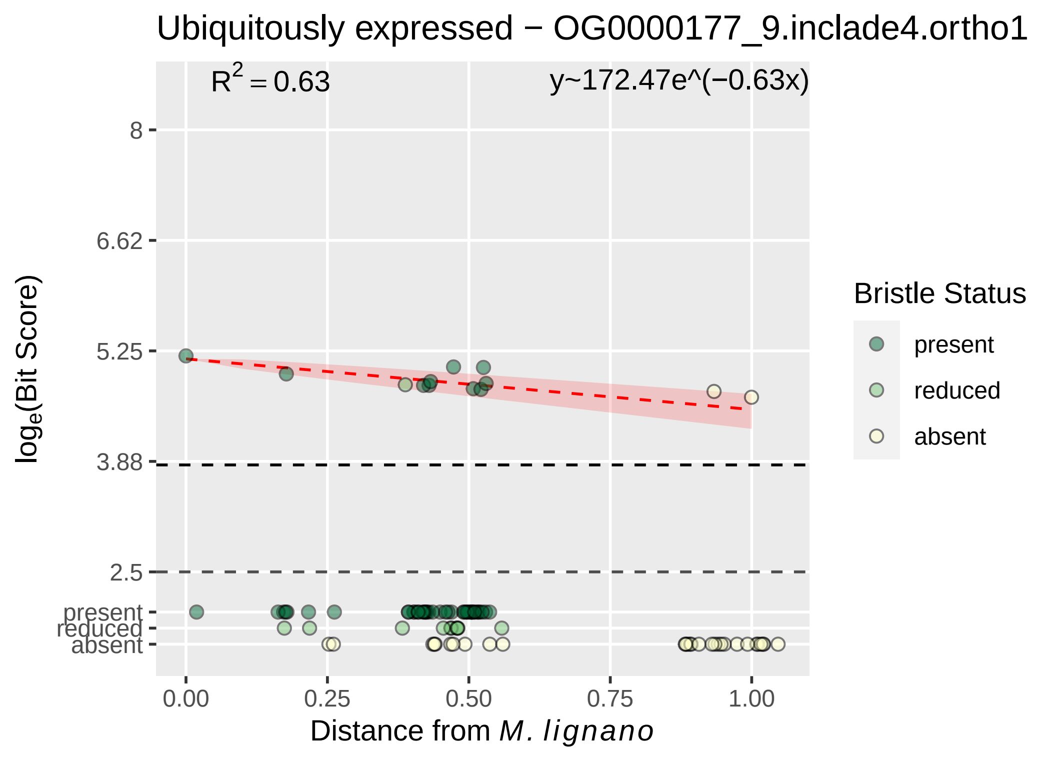
Ubiquitously expressed − OG0000177_9.inclade4.ortho1
R^2 = 0.63
y~172.47e^(−0.63x)
8
6.62
5.25
3.88
2.5
present
reduced
absent
0.00
0.25
0.50
0.75
1.00
log_e(Bit Score)
Distance from M. lignano
Bristle Status
present
reduced
absent

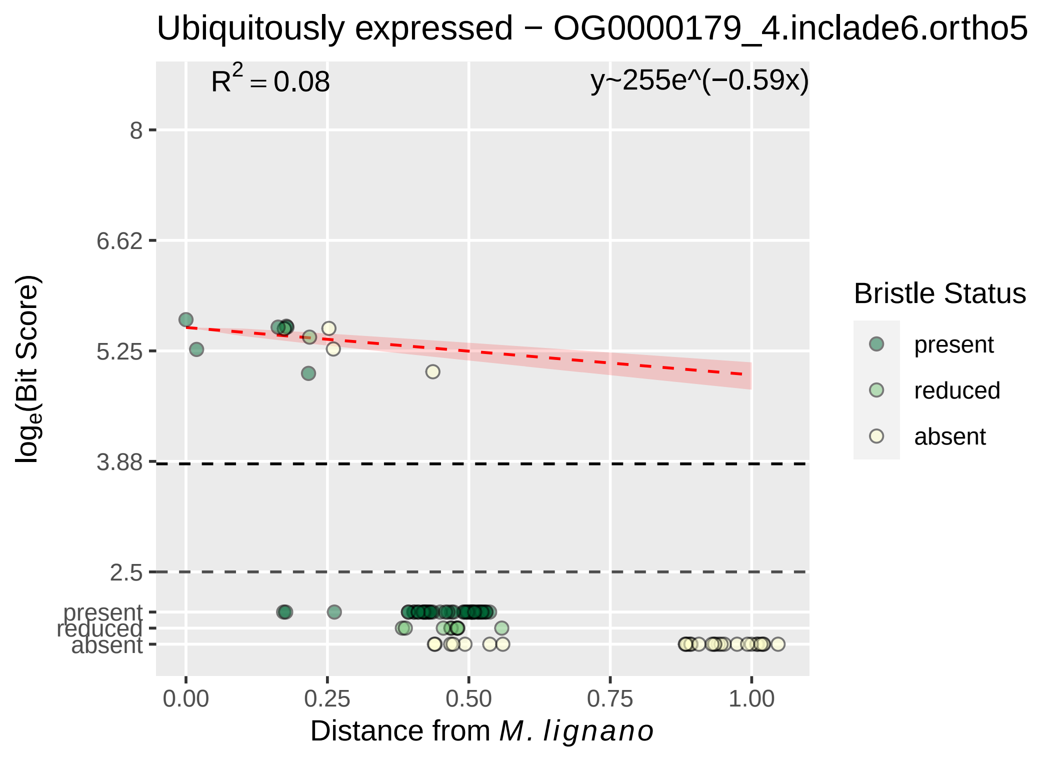
Ubiquitously expressed − OG0000179_4.inclade6.ortho5
R2 = 0.08
y~255e^(−0.59x)
8
6.62
5.25
3.88
2.5
present
reduced
absent
log e(Bit Score)
0.00
0.25
0.50
0.75
1.00
Distance from M. lignano
Bristle Status
present
reduced
absent

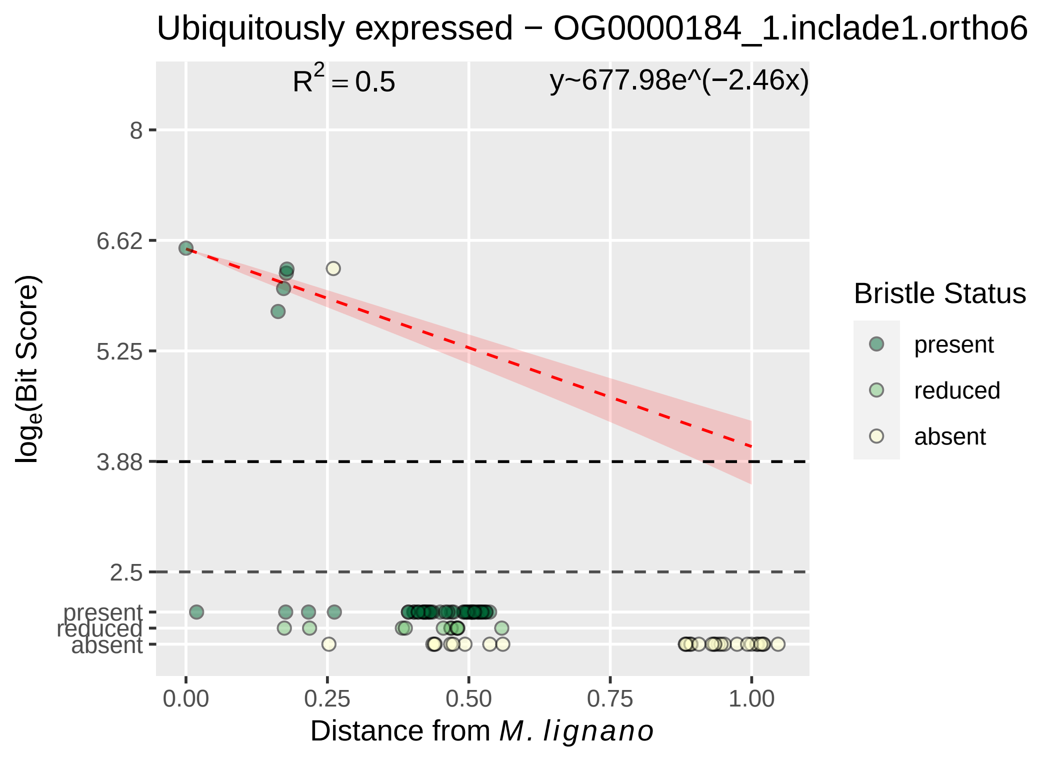
Ubiquitously expressed − OG0000184_1.inclade1.ortho6
R2 = 0.5
y~677.98e^(−2.46x)
8
6.62
5.25
3.88
2.5
present
reduced
absent
0.00
0.25
0.50
0.75
1.00
Distance from M. lignano
loge(Bit Score)
Bristle Status
present
reduced
absent

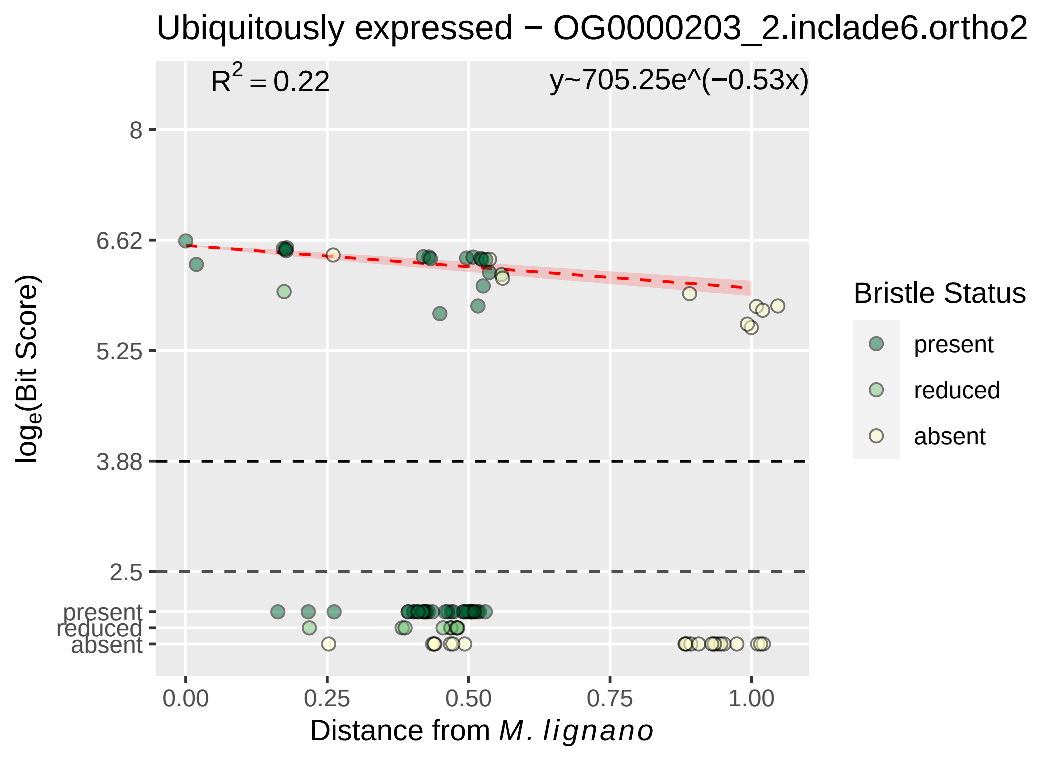
Ubiquitously expressed − OG0000203_2.inclade6.ortho2
R^2 = 0.22
y~705.25e^(−0.53x)
8
6.62
5.25
3.88
2.5
present
reduced
absent
log_e(Bit Score)
0.00
0.25
0.50
0.75
1.00
Distance from M. lignano
Bristle Status
present
reduced
absent

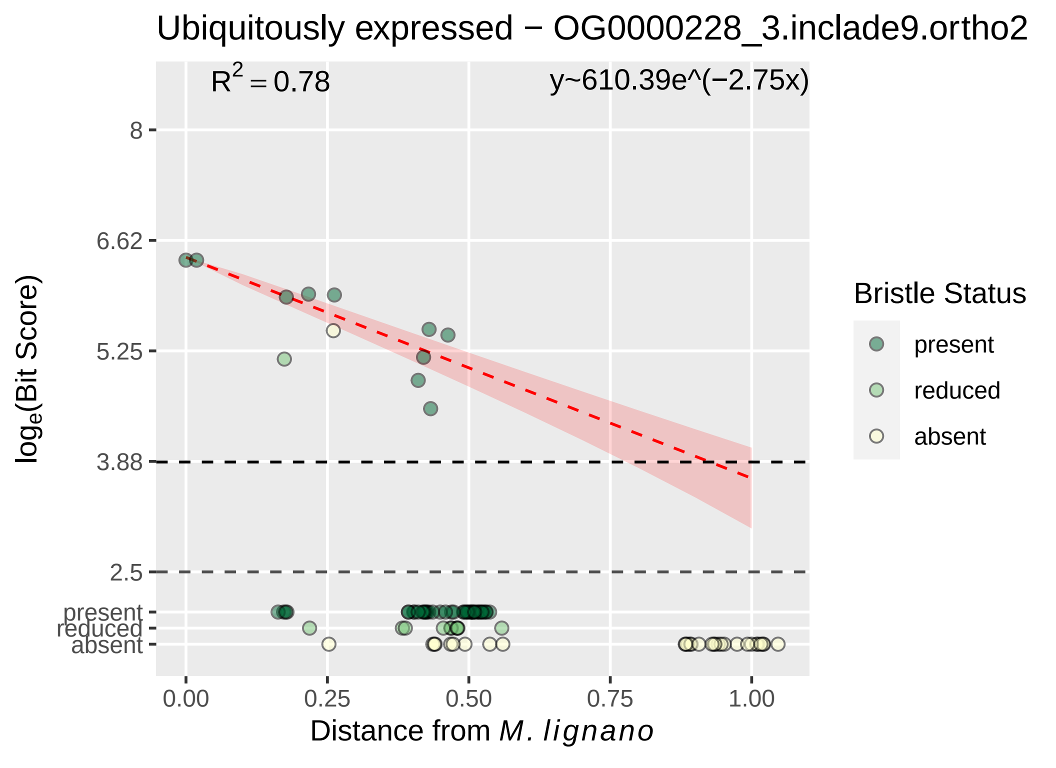

Ubiquitously expressed − OG0000228_3.inclade9.ortho2
R^2 = 0.78
y~610.39e^(−2.75x)
8
6.62
5.25
3.88
2.5
present
reduced
absent
0.00
0.25
0.50
0.75
1.00
Distance from M. lignano
loge(Bit Score)
Bristle Status
present
reduced
absent

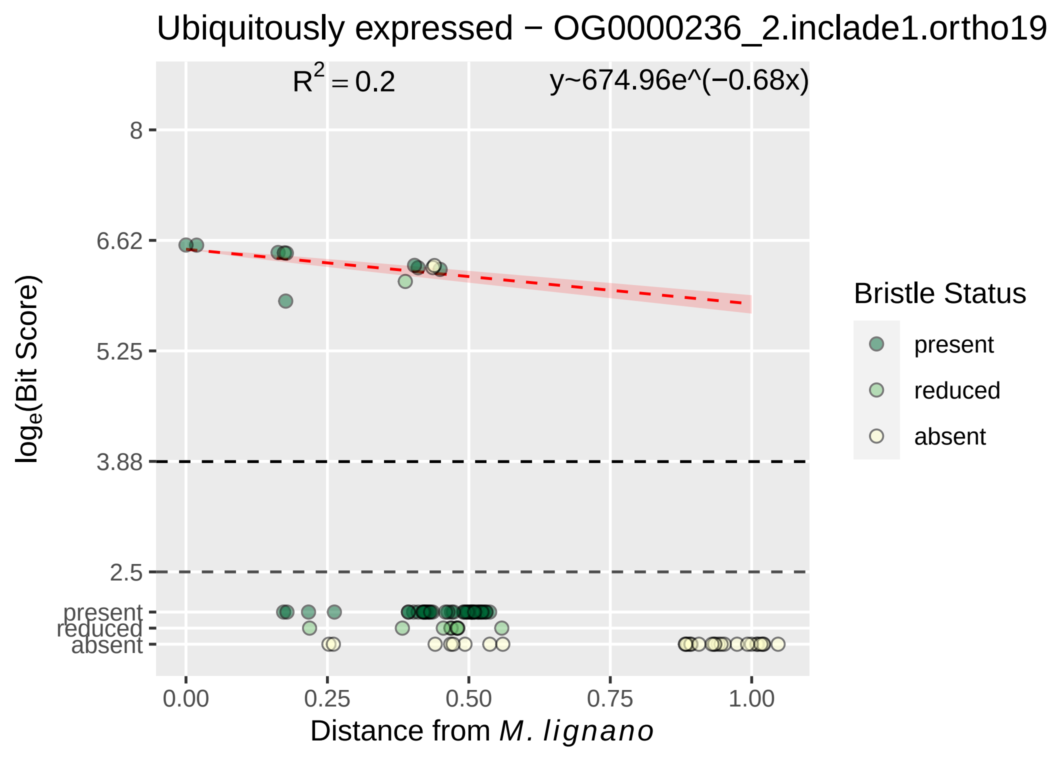

Ubiquitously expressed − OG0000236_2.inclade1.ortho19
R2 = 0.2
y~674.96e^(−0.68x)
8
6.62
5.25
3.88
2.5
present
reduced
absent
0.00
0.25
0.50
0.75
1.00
loge(Bit Score)
Distance from M. lignano
Bristle Status
present
reduced
absent

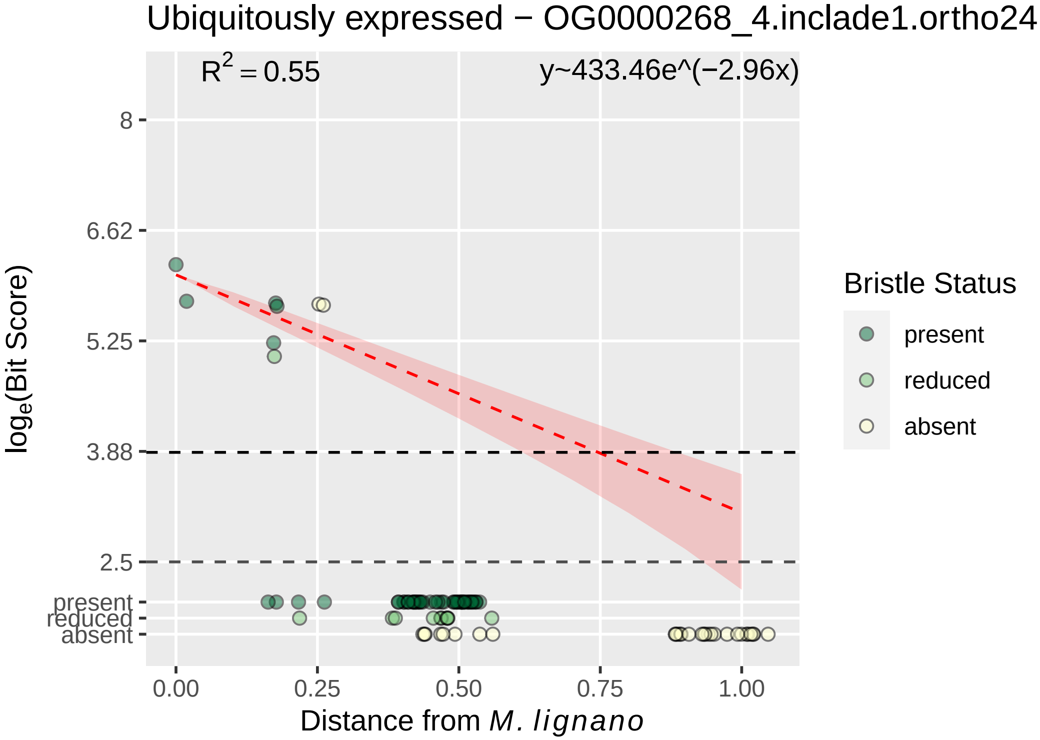
Ubiquitously expressed − OG0000268_4.inclade1.ortho24
$R^2 = 0.55$
y~433.46e^(−2.96x)
8
6.62
5.25
3.88
2.5
present
reduced
absent
0.00
0.25
0.50
0.75
1.00
Distance from *M. lignano*
$\log_e$(Bit Score)
Bristle Status
present
reduced
absent

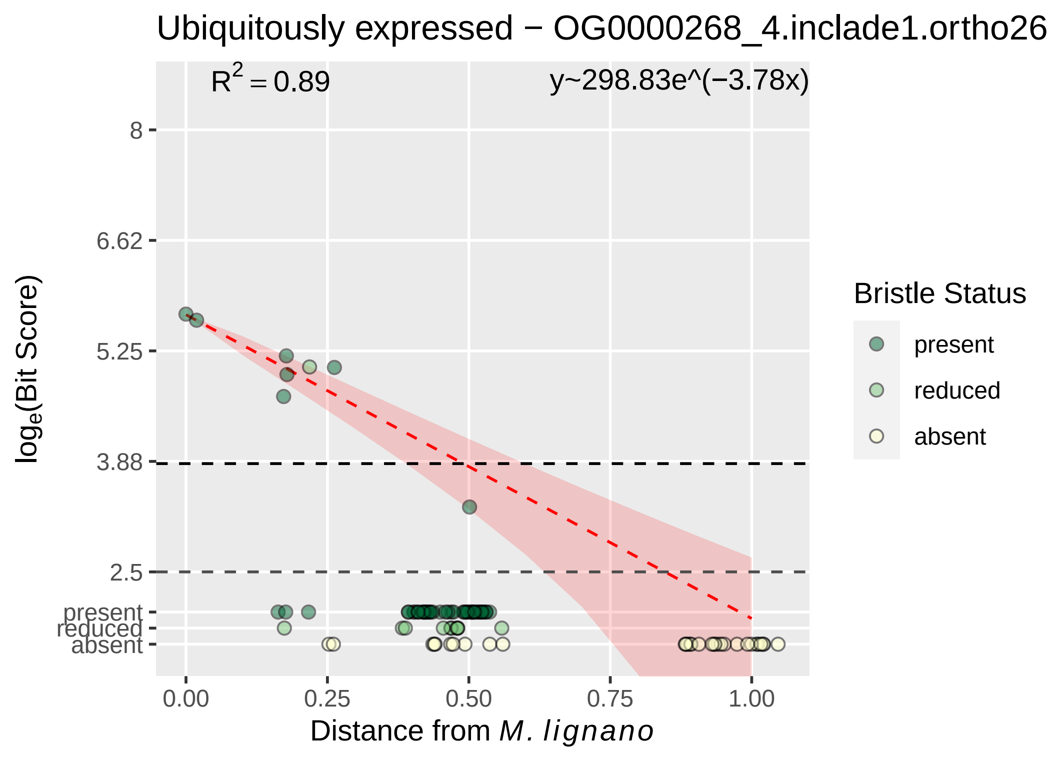

Ubiquitously expressed − OG0000268_4.inclade1.ortho26
R^2 = 0.89
y~298.83e^(−3.78x)
log_e(Bit Score)
8
6.62
5.25
3.88
2.5
present
reduced
absent
0.00
0.25
0.50
0.75
1.00
Distance from M. lignano
Bristle Status
present
reduced
absent

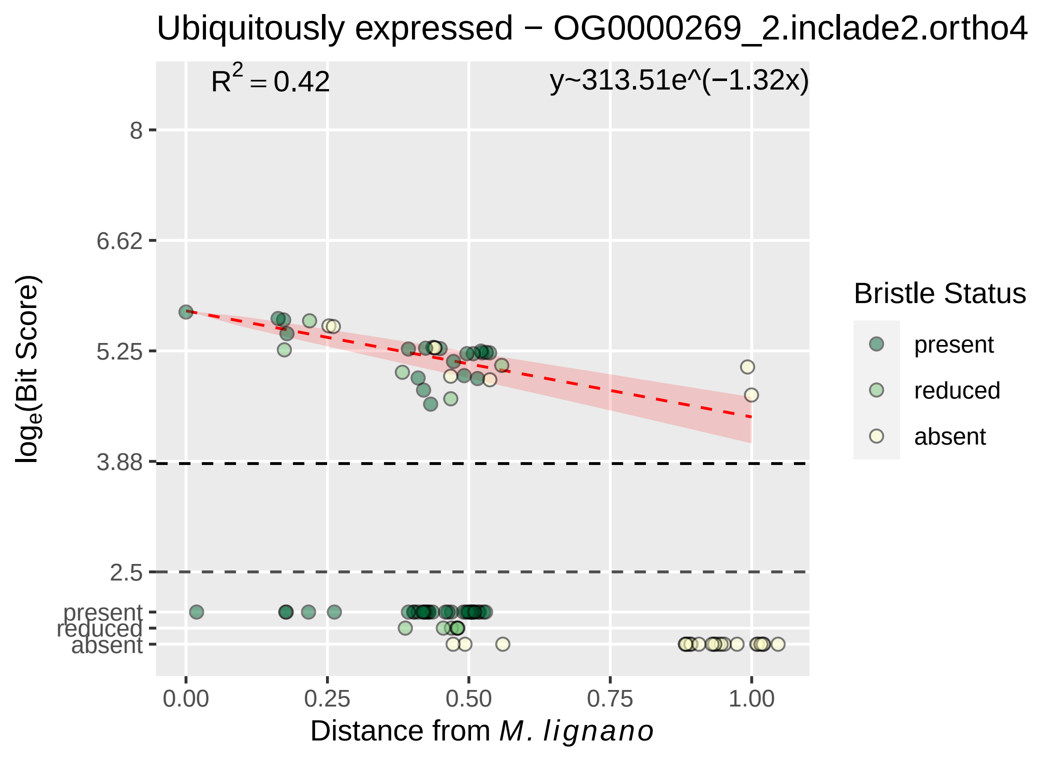

Ubiquitously expressed − OG0000269_2.inclade2.ortho4
R^2 = 0.42
y~313.51e^(−1.32x)
8
6.62
5.25
3.88
2.5
present
reduced
absent
log_e(Bit Score)
0.00
0.25
0.50
0.75
1.00
Distance from M. lignano
Bristle Status
present
reduced
absent

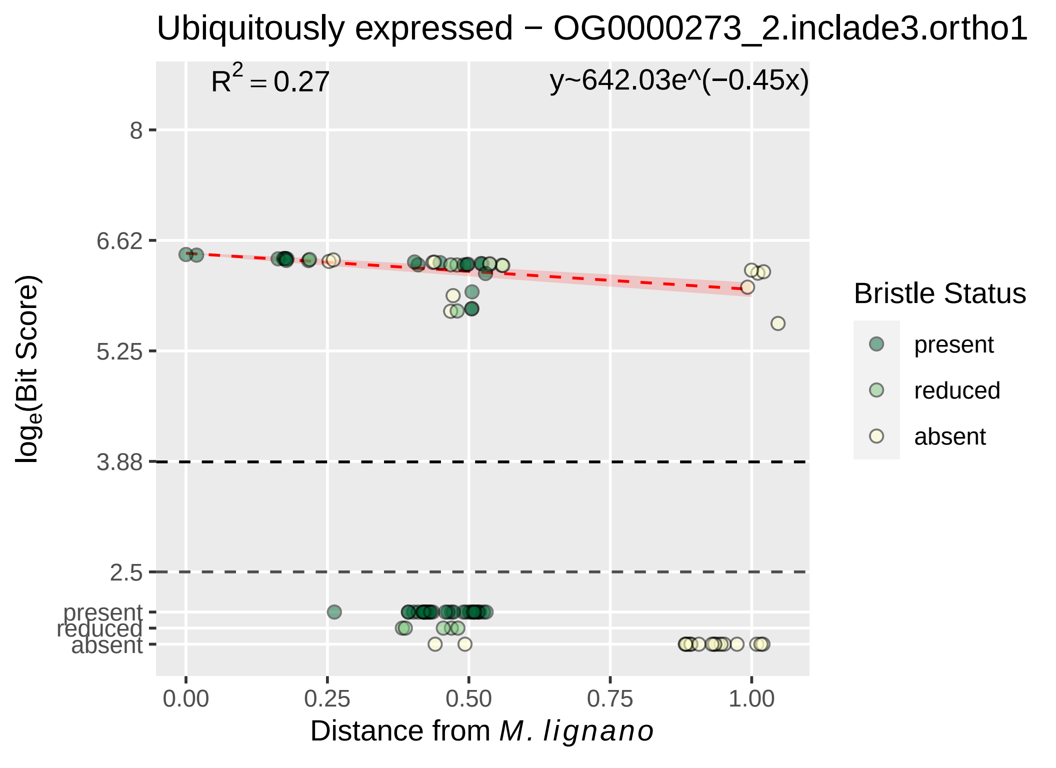

Ubiquitously expressed − OG0000273_2.inclade3.ortho1
R2 = 0.27
y~642.03e^(−0.45x)
8
6.62
5.25
3.88
2.5
present
reduced
absent
0.00
0.25
0.50
0.75
1.00
loge(Bit Score)
Distance from M. lignano
Bristle Status
present
reduced
absent

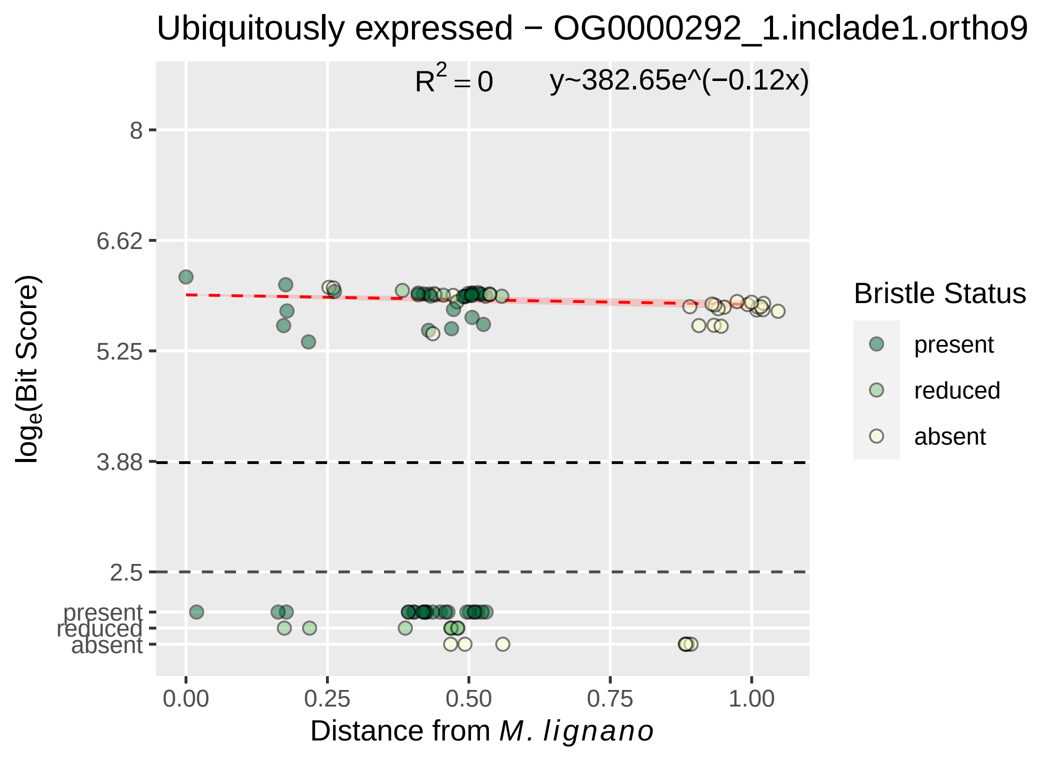

Ubiquitously expressed − OG0000292_1.inclade1.ortho9
R² = 0
y~382.65e^(−0.12x)
loge(Bit Score)
8
6.62
5.25
3.88
2.5
present
reduced
absent
0.00
0.25
0.50
0.75
1.00
Distance from M. lignano
Bristle Status
present
reduced
absent

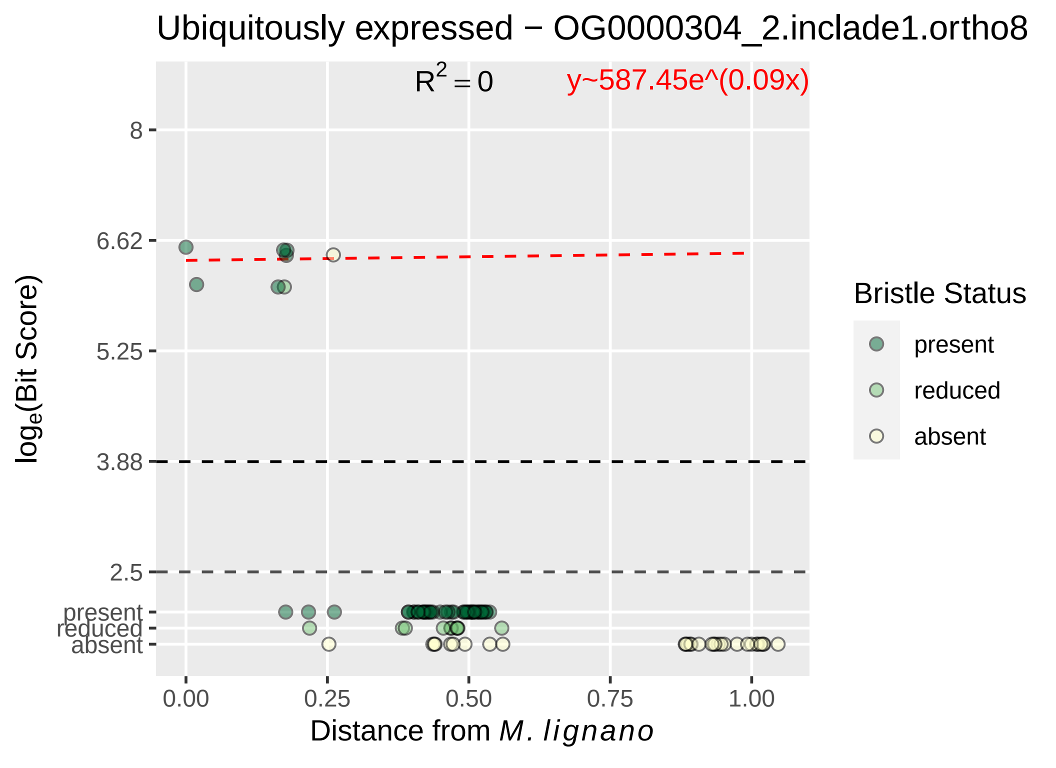
Ubiquitously expressed − OG0000304_2.inclade1.ortho8
R^2 = 0
y~587.45e^(0.09x)
8
6.62
5.25
3.88
2.5
present
reduced
absent
log_e(Bit Score)
0.00
0.25
0.50
0.75
1.00
Distance from M. lignano
Bristle Status
present
reduced
absent

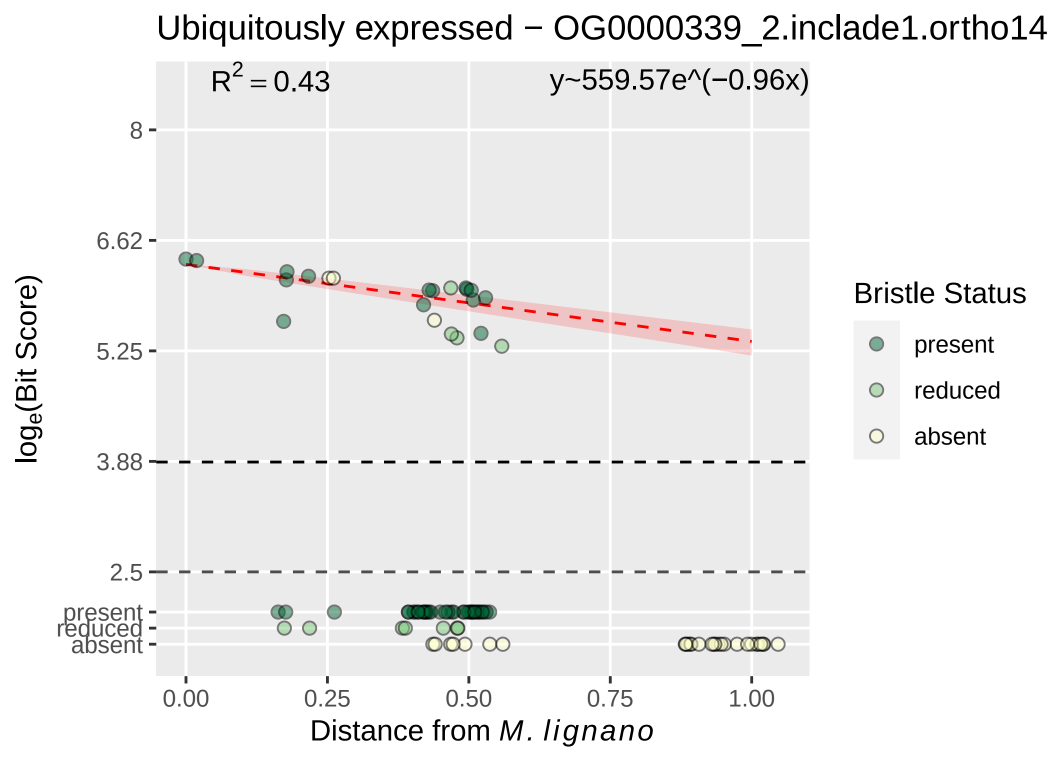

Ubiquitously expressed − OG0000339_2.inclade1.ortho14
$R^2 = 0.43$
y~559.57e^(−0.96x)
8
6.62
5.25
3.88
2.5
present
reduced
absent
$log_e$(Bit Score)
0.00
0.25
0.50
0.75
1.00
Distance from *M. lignano*
Bristle Status
present
reduced
absent

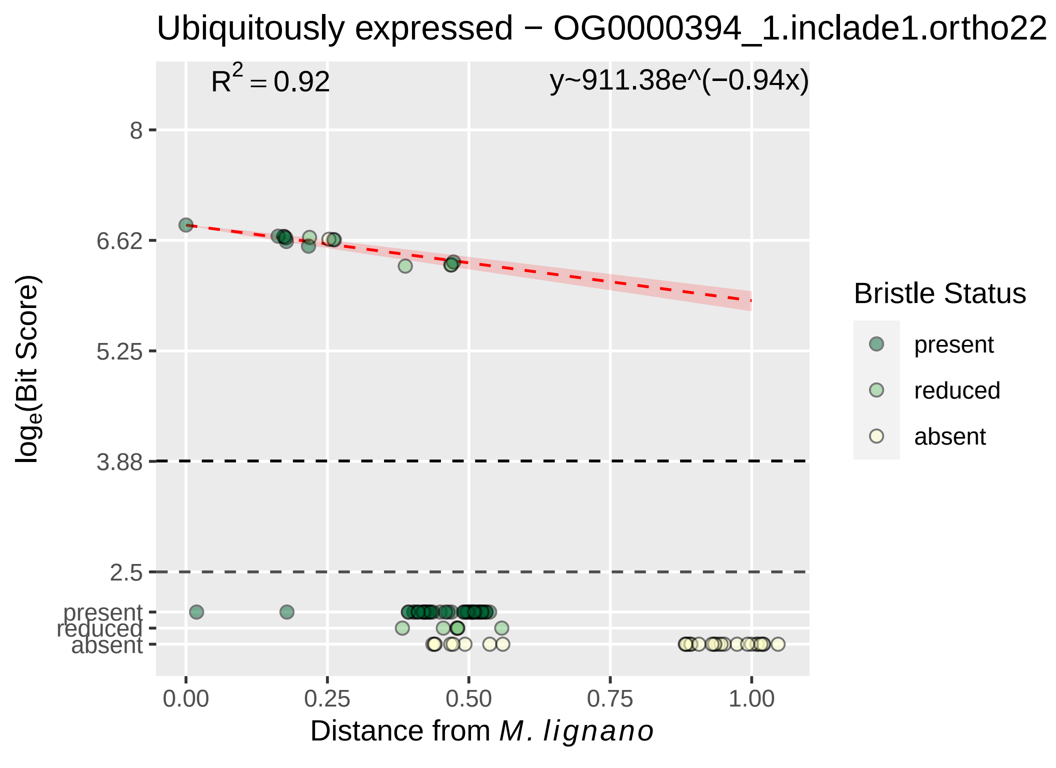

Ubiquitously expressed − OG0000394_1.inclade1.ortho22
R2 = 0.92
y~911.38e^(−0.94x)
8
6.62
5.25
3.88
2.5
present
reduced
absent
loge(Bit Score)
0.00
0.25
0.50
0.75
1.00
Distance from M. lignano
Bristle Status
present
reduced
absent

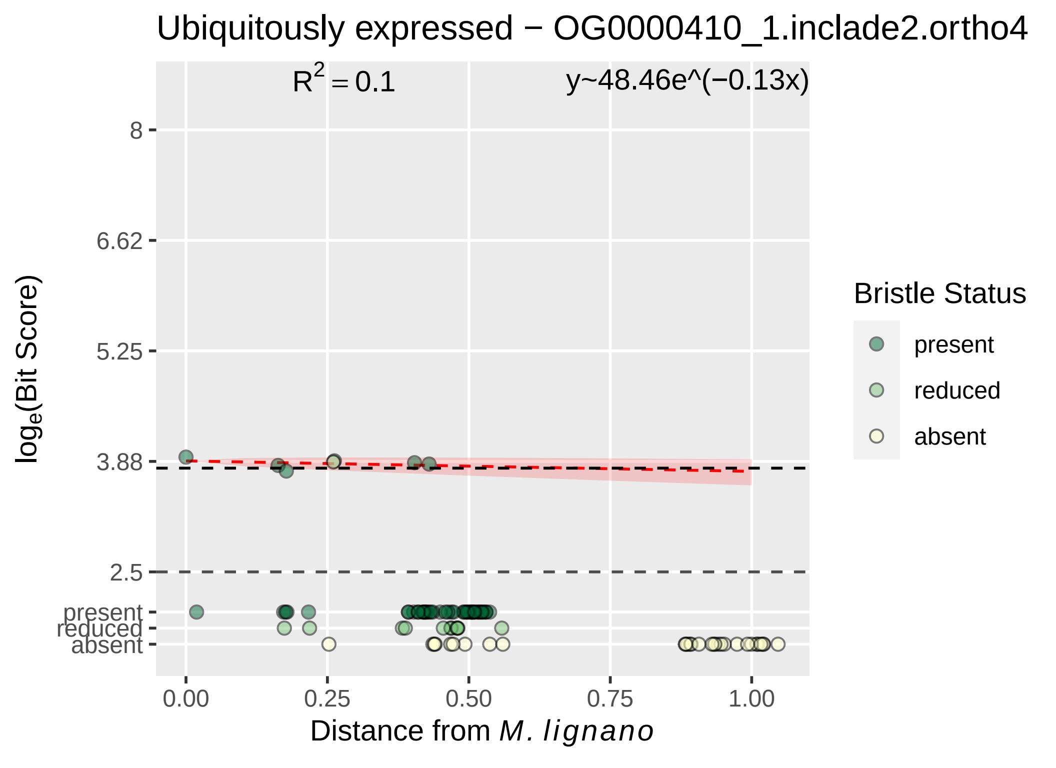

Ubiquitously expressed − OG0000410_1.inclade2.ortho4
R² = 0.1
y~48.46e^(−0.13x)
log_e(Bit Score)
8
6.62
5.25
3.88
2.5
present
reduced
absent
0.00
0.25
0.50
0.75
1.00
Distance from M. lignano
Bristle Status
present
reduced
absent

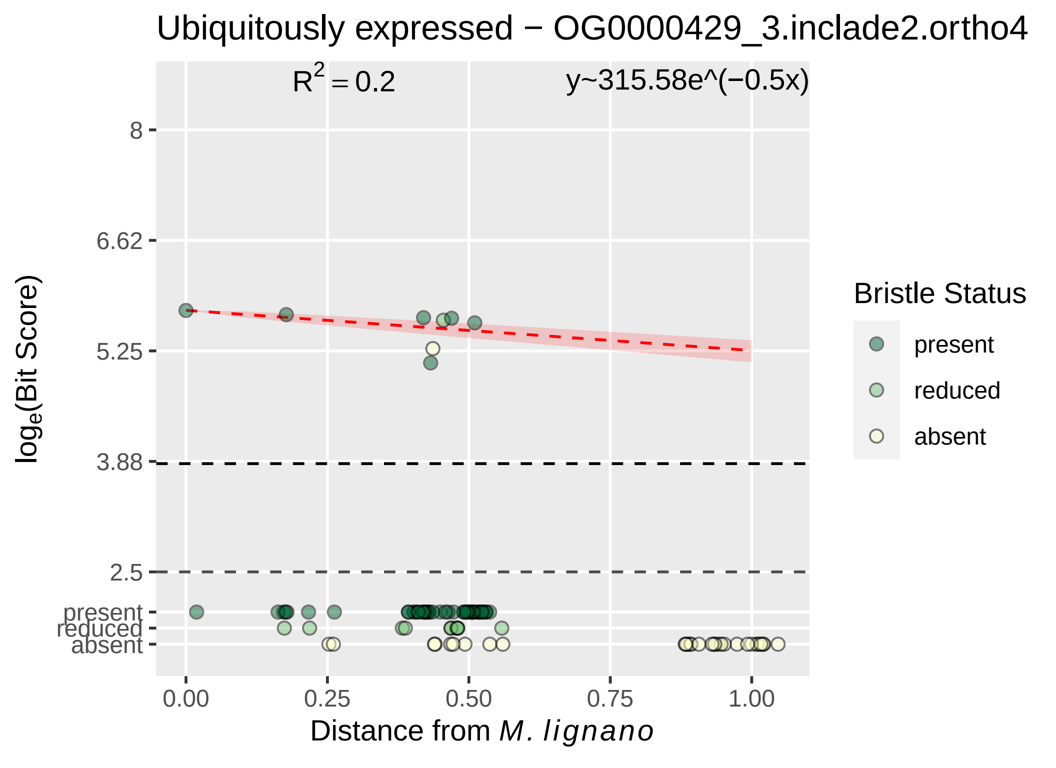

Ubiquitously expressed − OG0000429_3.inclade2.ortho4
R^2 = 0.2
y~315.58e^(−0.5x)
8
6.62
5.25
3.88
2.5
present
reduced
absent
0.00
0.25
0.50
0.75
1.00
log_e(Bit Score)
Distance from M. lignano
Bristle Status
present
reduced
absent

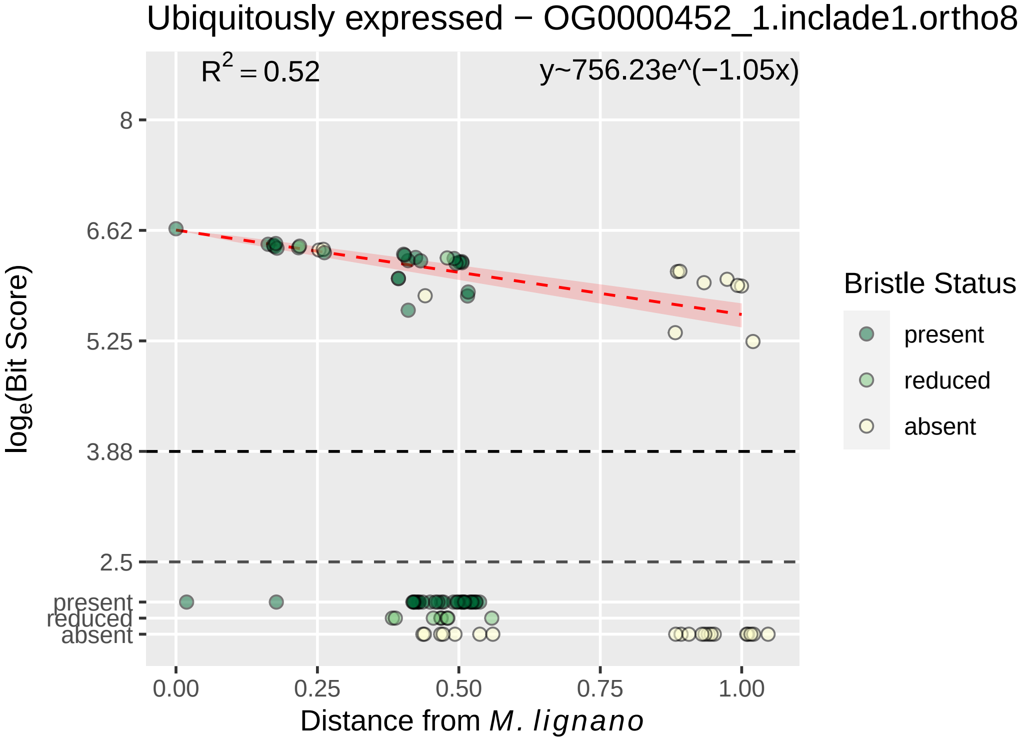

Ubiquitously expressed − OG0000452_1.inclade1.ortho8
R² = 0.52
y~756.23e^(−1.05x)
8
6.62
5.25
3.88
2.5
present
reduced
absent
0.00
0.25
0.50
0.75
1.00
loge(Bit Score)
Distance from M. lignano
Bristle Status
present
reduced
absent

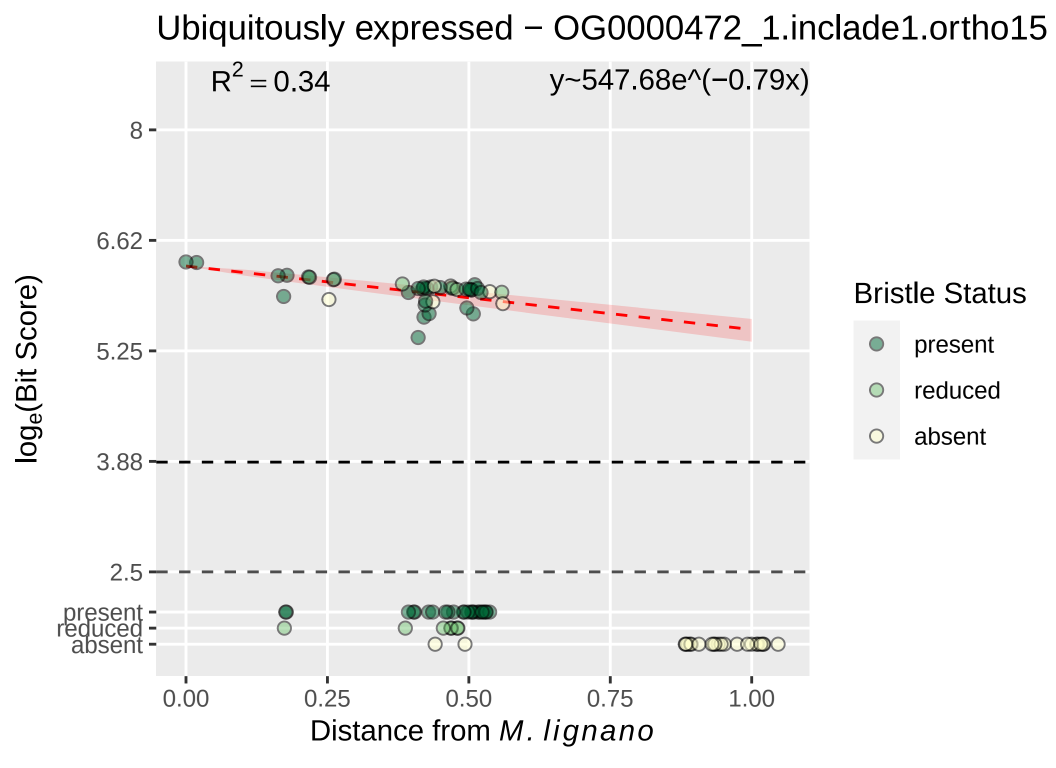

Ubiquitously expressed − OG0000472_1.inclade1.ortho15
R² = 0.34
y~547.68e^(−0.79x)
8
6.62
5.25
3.88
2.5
present
reduced
absent
loge(Bit Score)
0.00
0.25
0.50
0.75
1.00
Distance from M. lignano
Bristle Status
present
reduced
absent

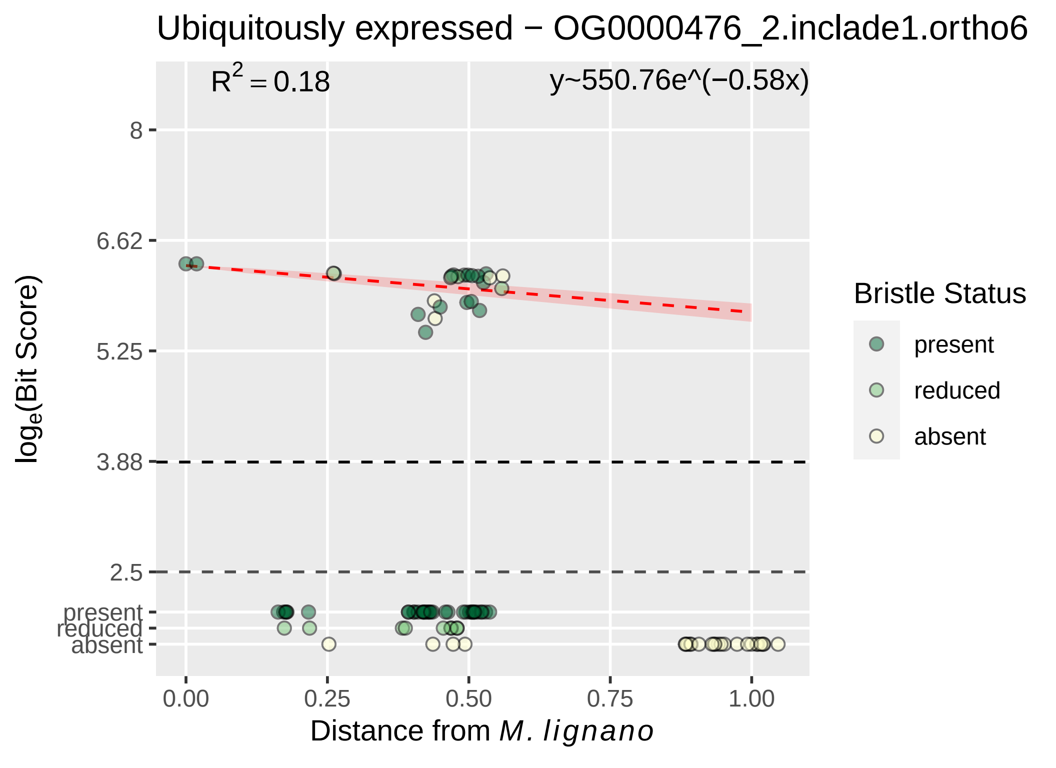

Ubiquitously expressed − OG0000476_2.inclade1.ortho6
$R^2 = 0.18$
y~550.76e^(−0.58x)
8
6.62
5.25
3.88
2.5
present
reduced
absent
0.00
0.25
0.50
0.75
1.00
Distance from M. lignano
loge(Bit Score)
Bristle Status
present
reduced
absent

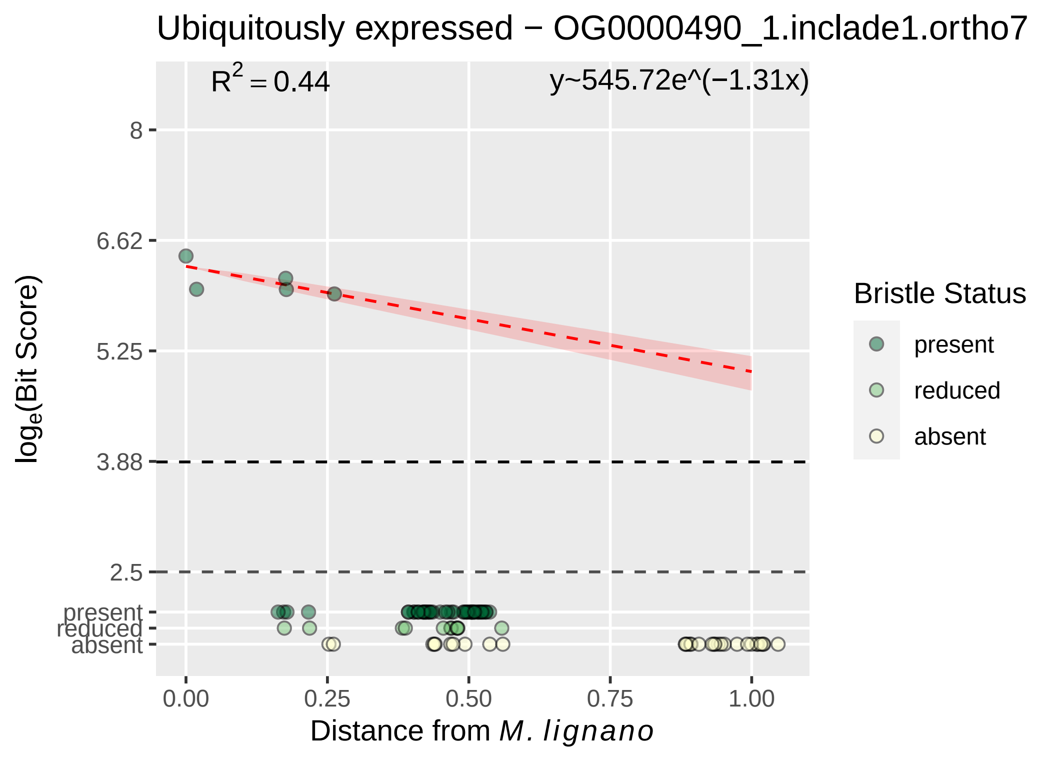

Ubiquitously expressed − OG0000490_1.inclade1.ortho7
R^2 = 0.44
y~545.72e^(−1.31x)
8
6.62
5.25
3.88
2.5
present
reduced
absent
0.00
0.25
0.50
0.75
1.00
Distance from M. lignano
log_e(Bit Score)
Bristle Status
present
reduced
absent

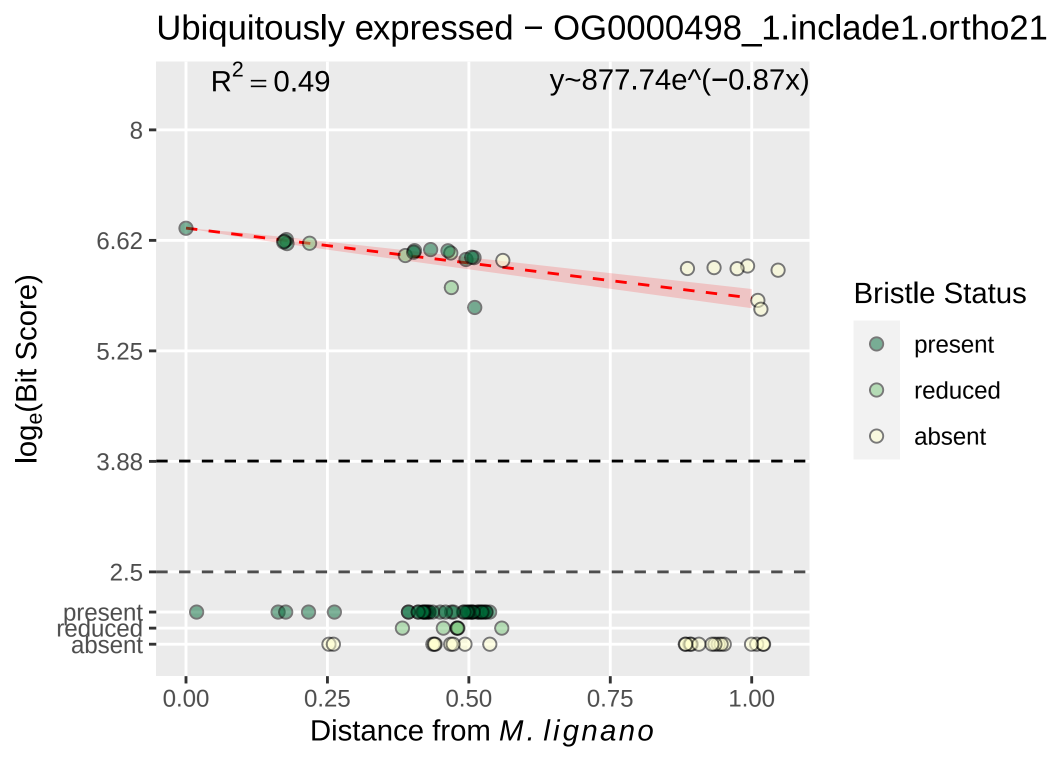

Ubiquitously expressed − OG0000498_1.inclade1.ortho21
R2 = 0.49
y~877.74e^(−0.87x)
8
6.62
5.25
3.88
2.5
present
reduced
absent
0.00
0.25
0.50
0.75
1.00
Distance from M. lignano
loge(Bit Score)
Bristle Status
present
reduced
absent

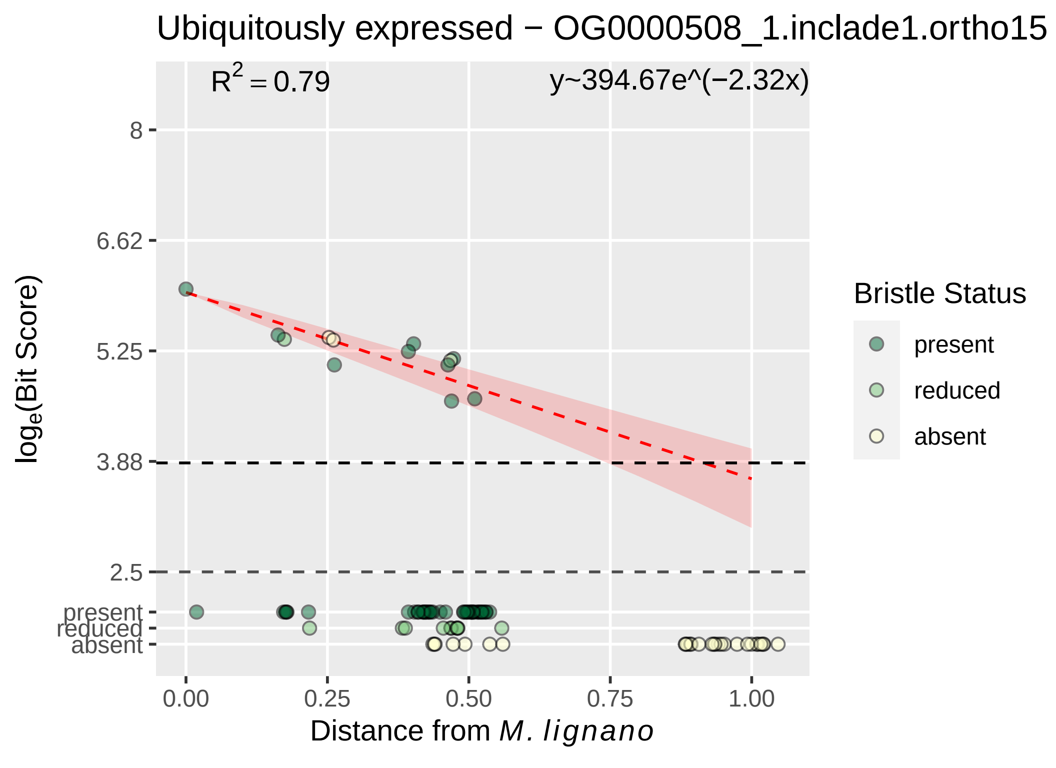

Ubiquitously expressed − OG0000508_1.inclade1.ortho15
R² = 0.79
y~394.67e^(−2.32x)
8
6.62
5.25
3.88
2.5
present
reduced
absent
log_e(Bit Score)
0.00
0.25
0.50
0.75
1.00
Distance from M. lignano
Bristle Status
present
reduced
absent

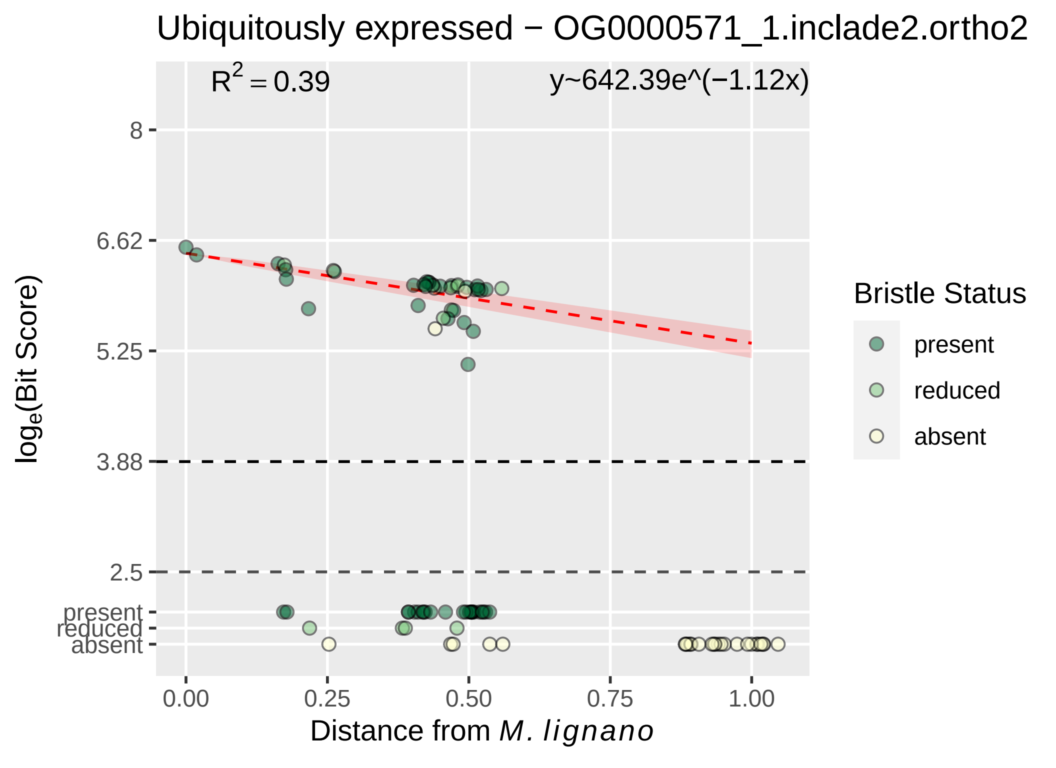

Ubiquitously expressed − OG0000571_1.inclade2.ortho2
R2 = 0.39
y~642.39e^(−1.12x)
8
6.62
5.25
3.88
2.5
present
reduced
absent
loge(Bit Score)
0.00
0.25
0.50
0.75
1.00
Distance from M. lignano
Bristle Status
present
reduced
absent

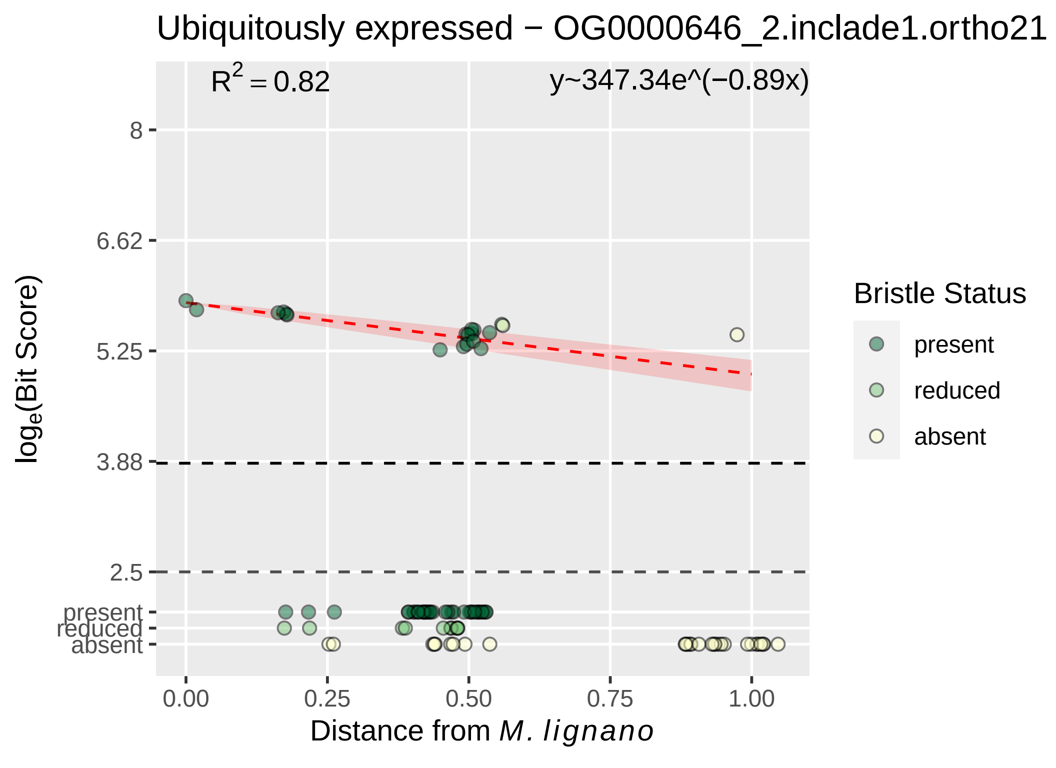
Ubiquitously expressed − OG0000646_2.inclade1.ortho21
R^2 = 0.82
y~347.34e^(−0.89x)
8
6.62
5.25
3.88
2.5
present
reduced
absent
0.00
0.25
0.50
0.75
1.00
Distance from M. lignano
log_e(Bit Score)
Bristle Status
present
reduced
absent

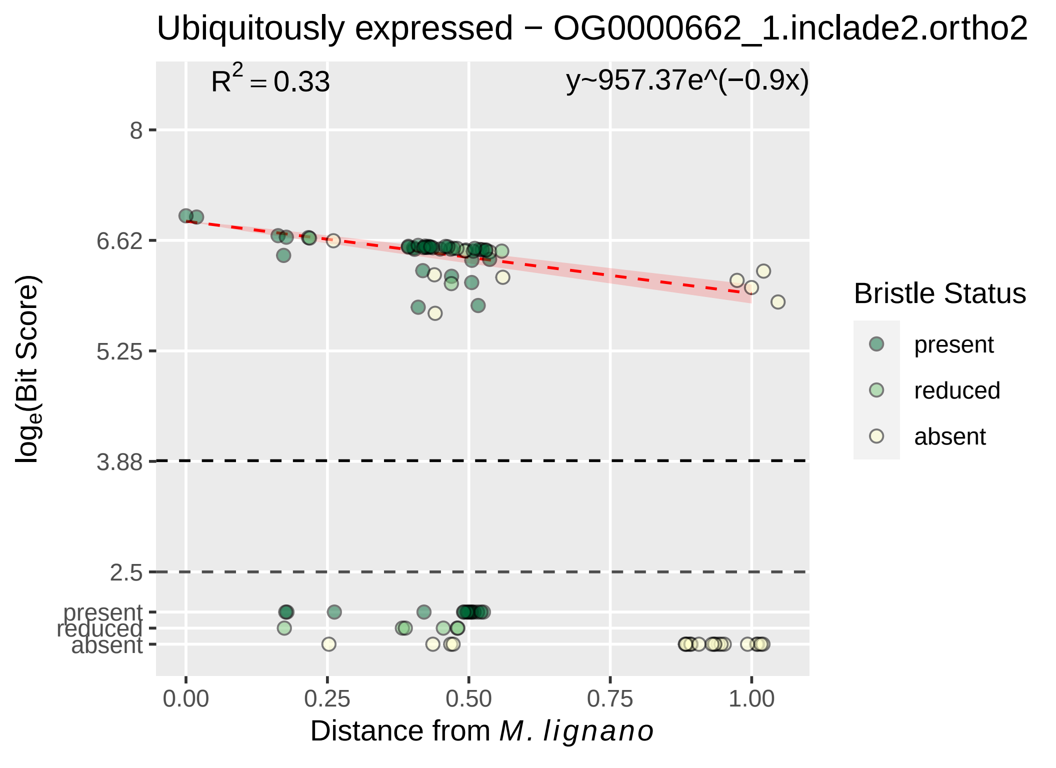

Ubiquitously expressed − OG0000662_1.inclade2.ortho2
R^2 = 0.33
y~957.37e^(−0.9x)
8
6.62
5.25
3.88
2.5
present
reduced
absent
log_e(Bit Score)
0.00
0.25
0.50
0.75
1.00
Distance from M. lignano
Bristle Status
present
reduced
absent

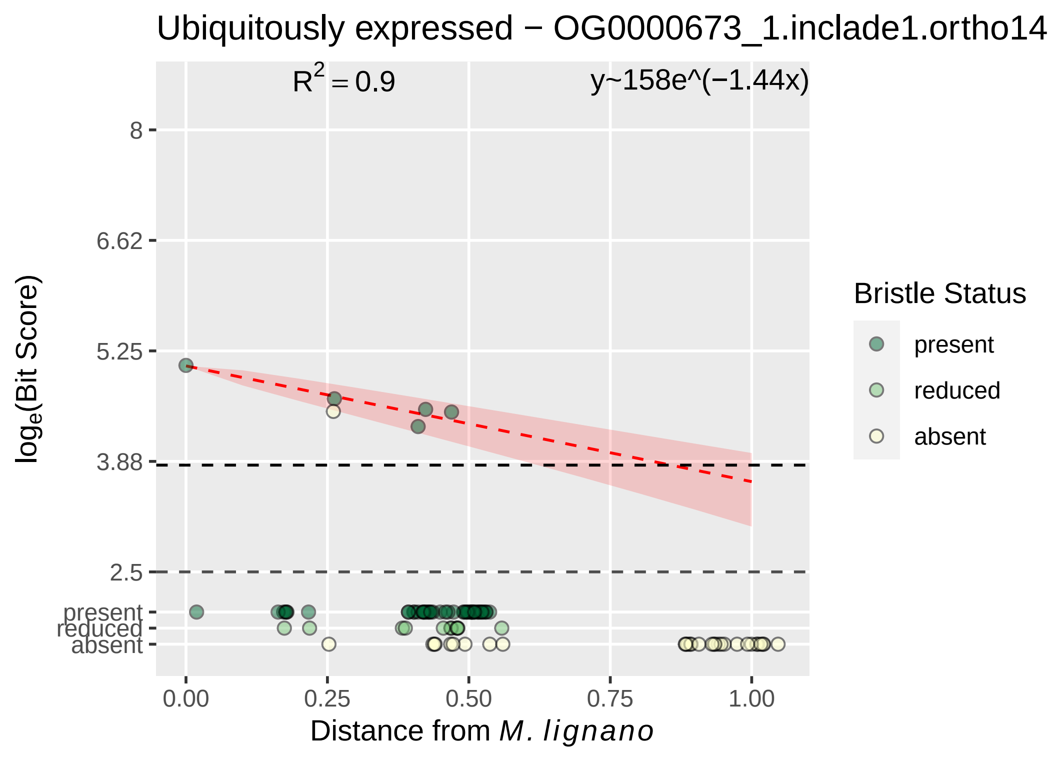

Ubiquitously expressed − OG0000673_1.inclade1.ortho14
R^2 = 0.9
y~158e^(−1.44x)
8
6.62
5.25
3.88
2.5
present
reduced
absent
0.00
0.25
0.50
0.75
1.00
log_e(Bit Score)
Distance from M. lignano
Bristle Status
present
reduced
absent

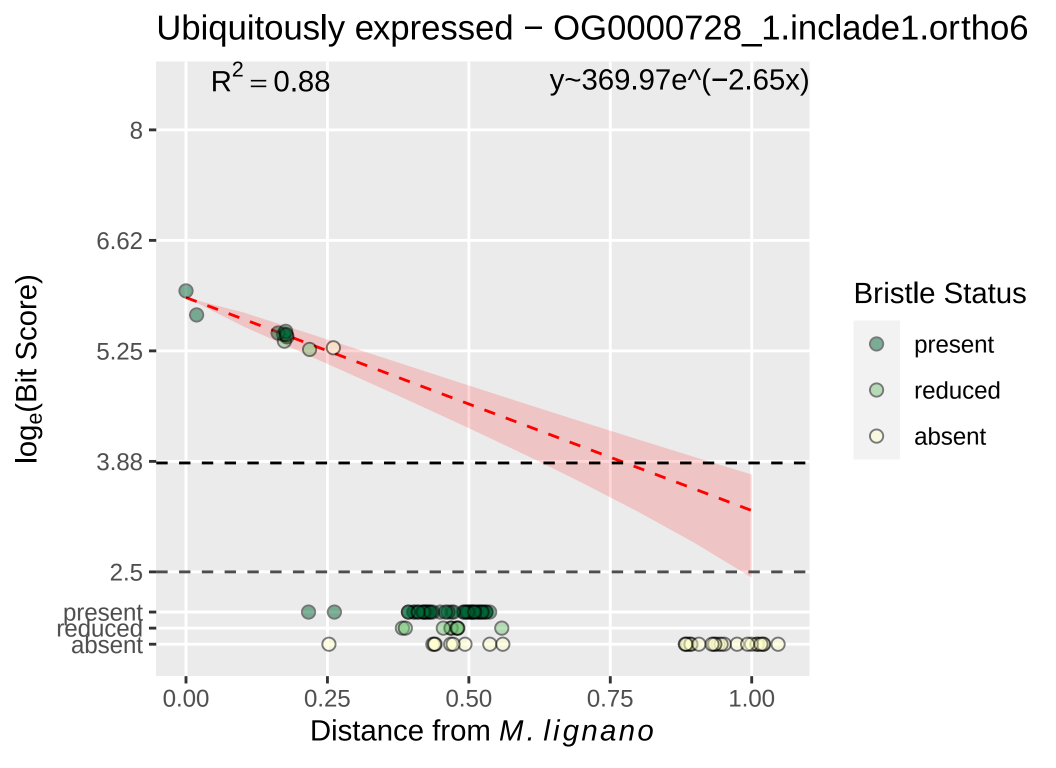

Ubiquitously expressed − OG0000728_1.inclade1.ortho6
R^2 = 0.88
y~369.97e^(−2.65x)
8
6.62
5.25
3.88
2.5
present
reduced
absent
log_e(Bit Score)
0.00
0.25
0.50
0.75
1.00
Distance from M. lignano
Bristle Status
present
reduced
absent

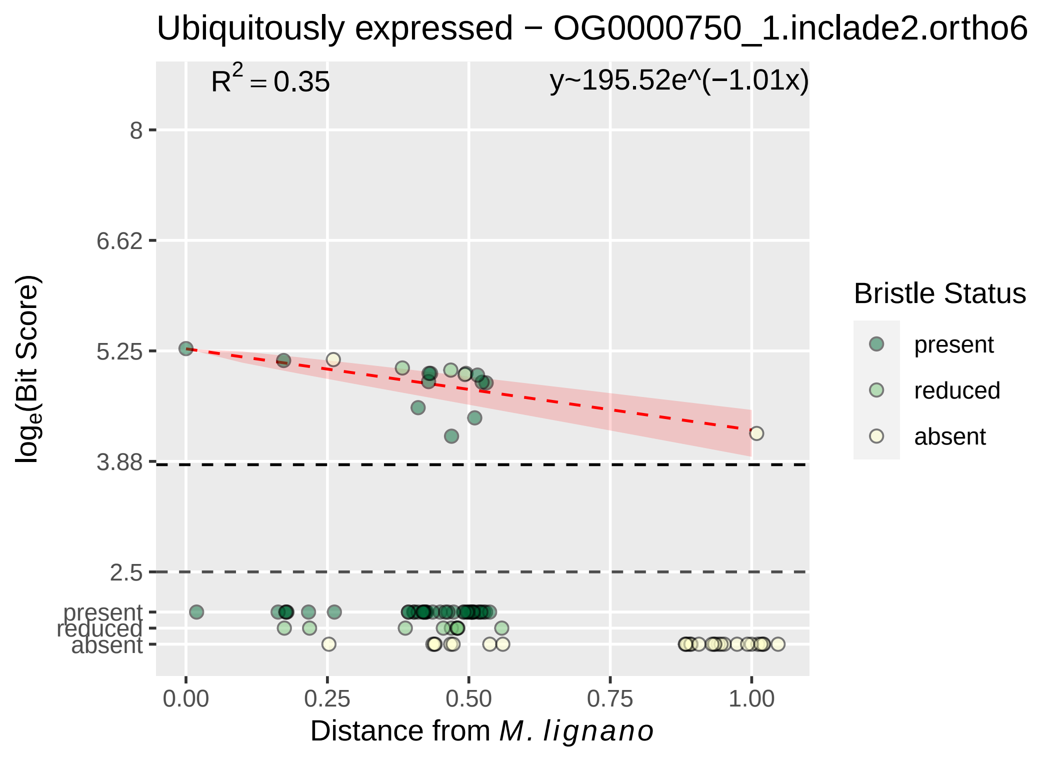

Ubiquitously expressed − OG0000750_1.inclade2.ortho6
R^2 = 0.35
y~195.52e^(−1.01x)
8
6.62
5.25
3.88
2.5
present
reduced
absent
0.00
0.25
0.50
0.75
1.00
log_e(Bit Score)
Distance from M. lignano
Bristle Status
present
reduced
absent

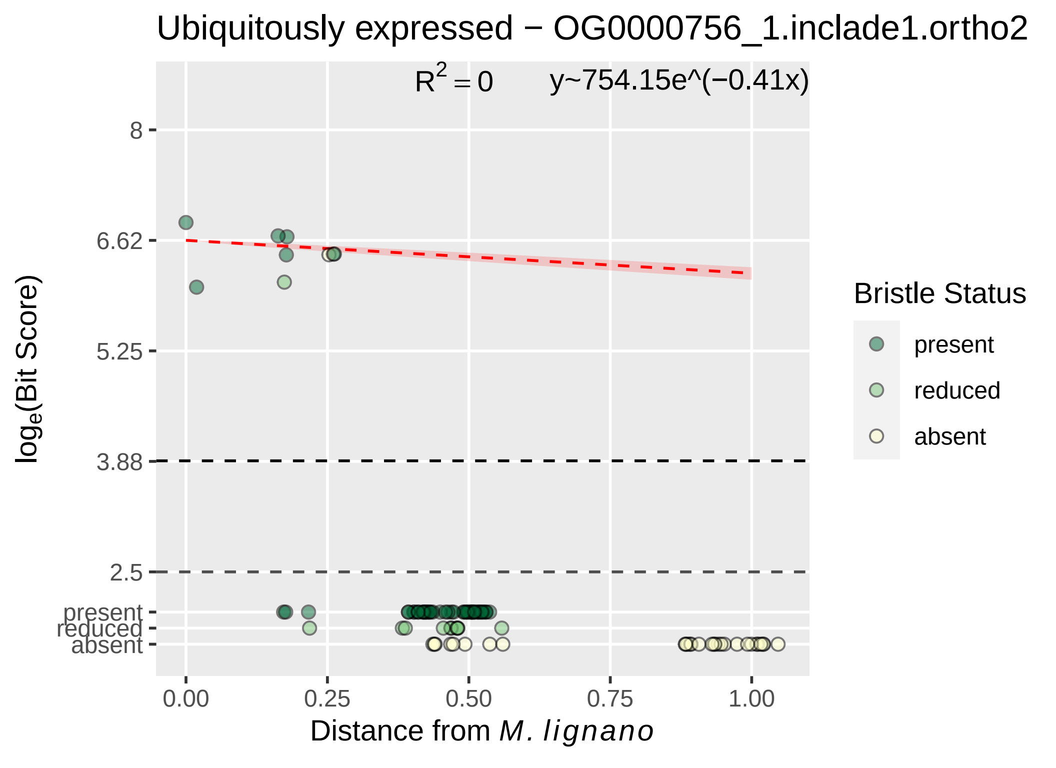

Ubiquitously expressed − OG0000756_1.inclade1.ortho2
R^2 = 0
y~754.15e^(−0.41x)
8
6.62
5.25
3.88
2.5
present
reduced
absent
log_e(Bit Score)
0.00
0.25
0.50
0.75
1.00
Distance from M. lignano
Bristle Status
present
reduced
absent

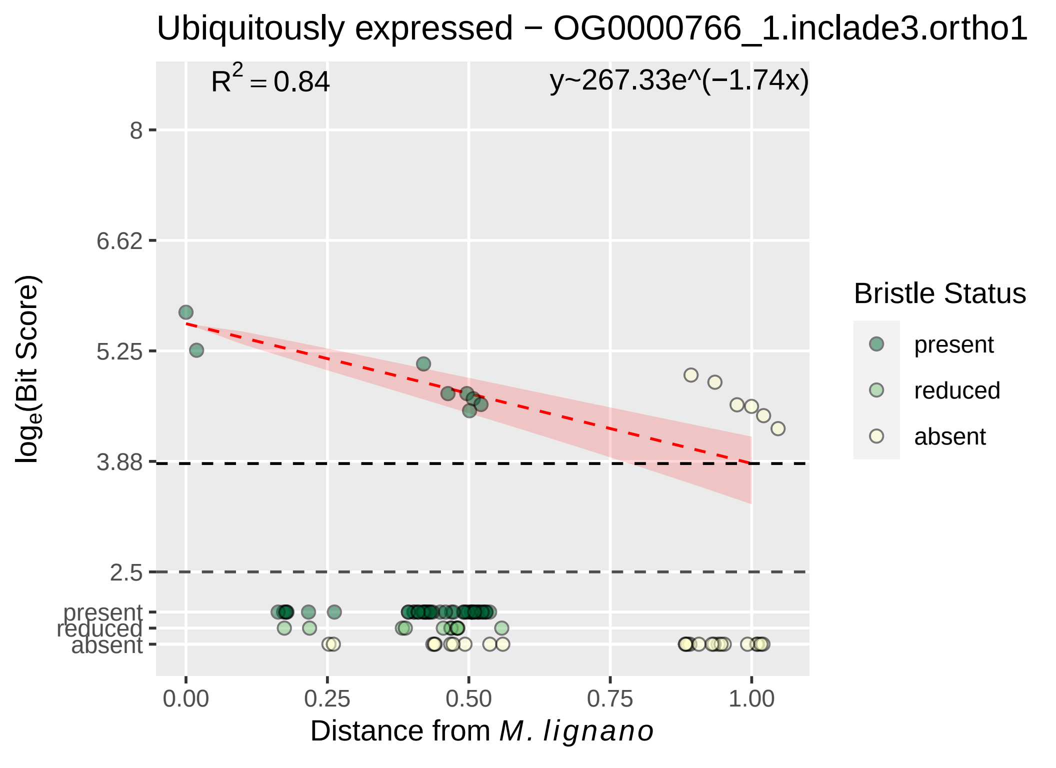

Ubiquitously expressed − OG0000766_1.inclade3.ortho1
$R^2 = 0.84$
y~267.33e^(−1.74x)
8
6.62
5.25
3.88
2.5
present
reduced
absent
0.00
0.25
0.50
0.75
1.00
Distance from M. lignano
$\log_e$(Bit Score)
Bristle Status
present
reduced
absent

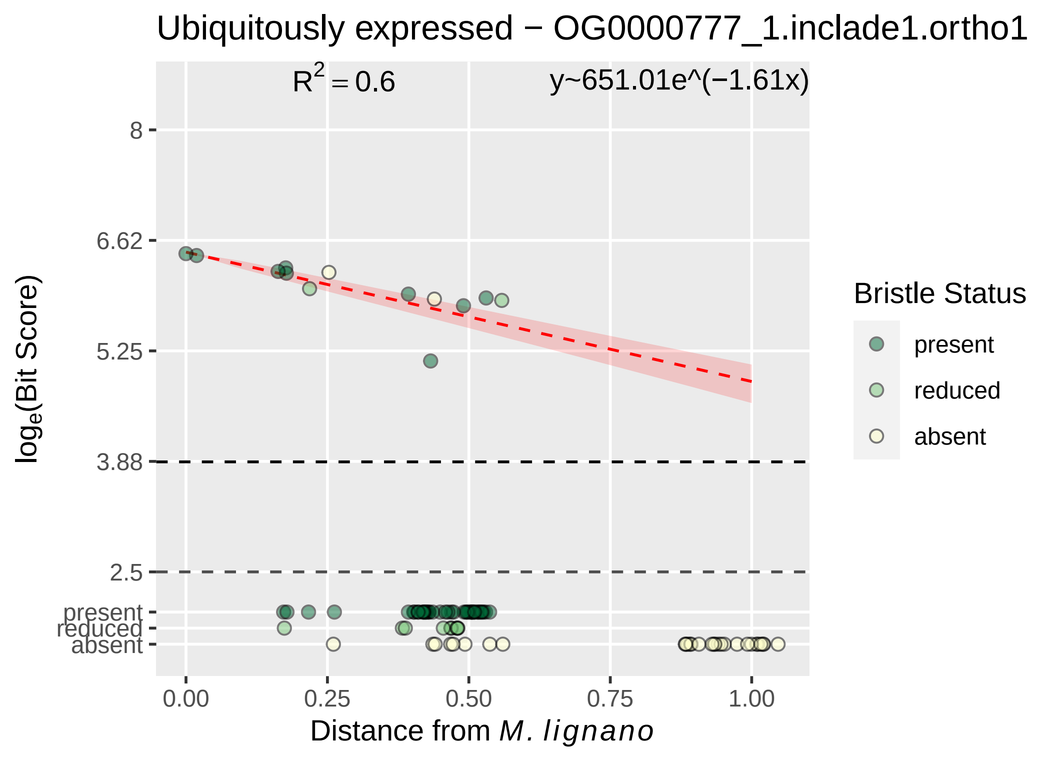

Ubiquitously expressed − OG0000777_1.inclade1.ortho1
R^2 = 0.6
y~651.01e^(−1.61x)
8
6.62
5.25
3.88
2.5
present
reduced
absent
log_e(Bit Score)
0.00
0.25
0.50
0.75
1.00
Distance from M. lignano
Bristle Status
present
reduced
absent

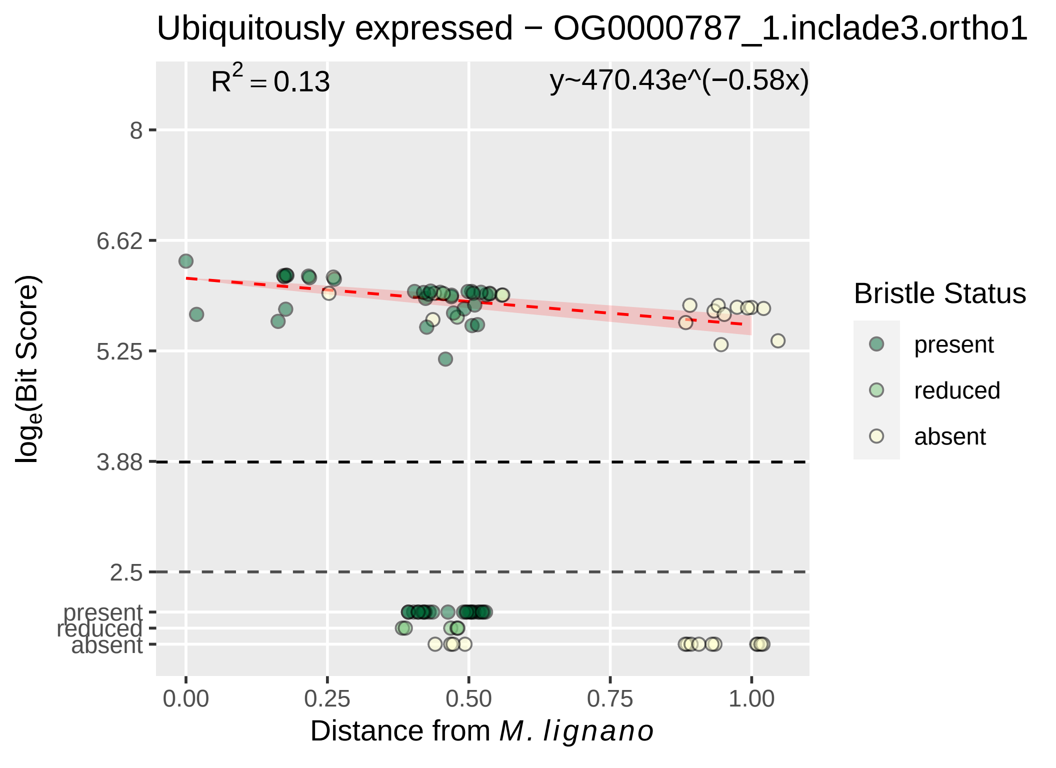

Ubiquitously expressed − OG0000787_1.inclade3.ortho1
R² = 0.13
y~470.43e^(−0.58x)
8
6.62
5.25
3.88
2.5
present
reduced
absent
loge(Bit Score)
0.00
0.25
0.50
0.75
1.00
Distance from M. lignano
Bristle Status
present
reduced
absent

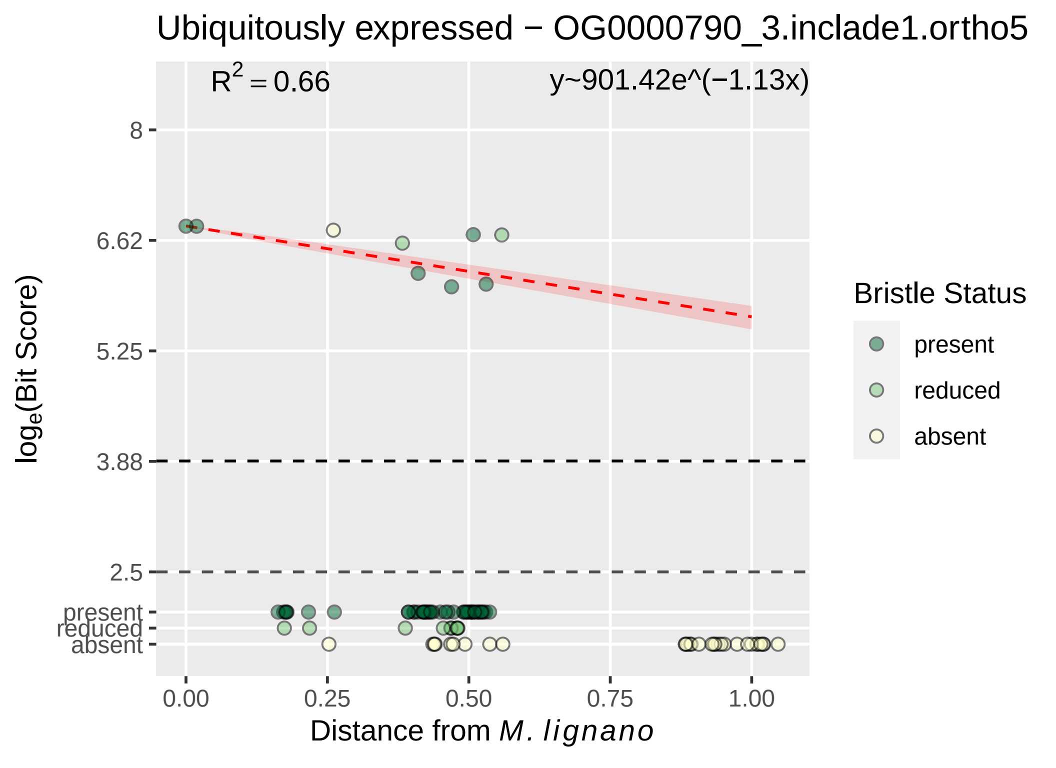

Ubiquitously expressed − OG0000790_3.inclade1.ortho5
$R^2 = 0.66$
y~901.42e^(−1.13x)
8
6.62
5.25
3.88
2.5
present
reduced
absent
$\log_e$(Bit Score)
0.00
0.25
0.50
0.75
1.00
Distance from *M. lignano*
Bristle Status
present
reduced
absent

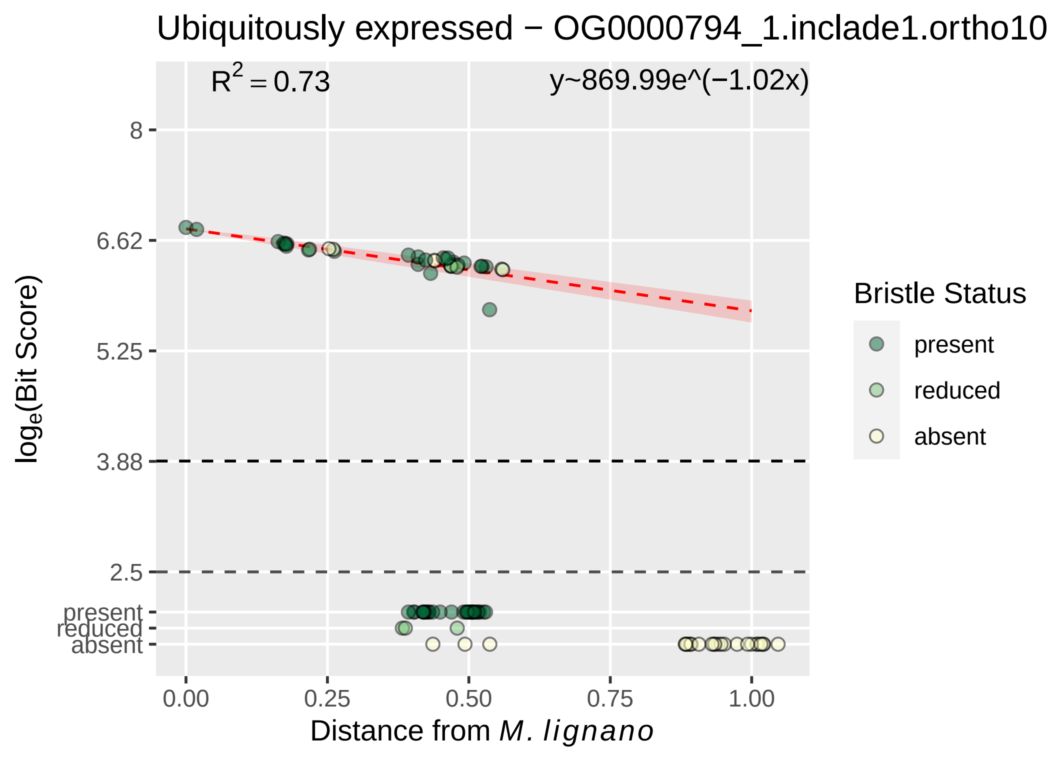
Ubiquitously expressed − OG0000794_1.inclade1.ortho10
R^2 = 0.73
y~869.99e^(−1.02x)
loge(Bit Score)
8
6.62
5.25
3.88
2.5
present
reduced
absent
0.00
0.25
0.50
0.75
1.00
Distance from M. lignano
Bristle Status
present
reduced
absent

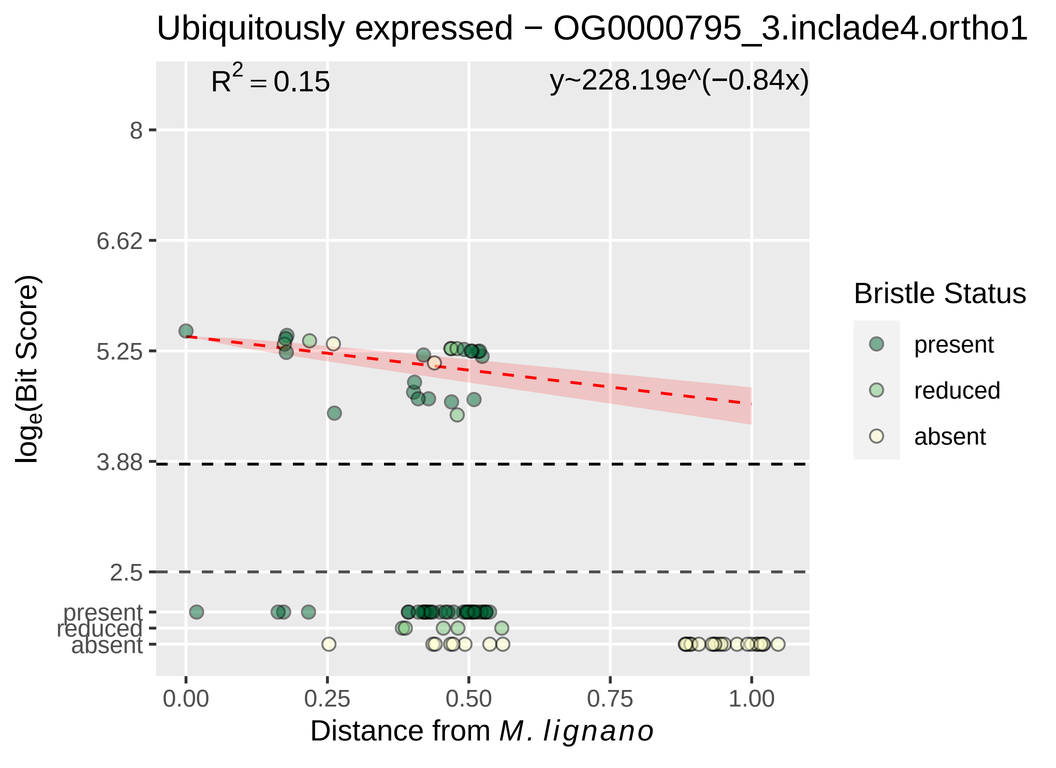

Ubiquitously expressed − OG0000795_3.inclade4.ortho1
R^2 = 0.15
y~228.19e^(−0.84x)
8
6.62
5.25
3.88
2.5
present
reduced
absent
0.00
0.25
0.50
0.75
1.00
log_e(Bit Score)
Distance from M. lignano
Bristle Status
present
reduced
absent

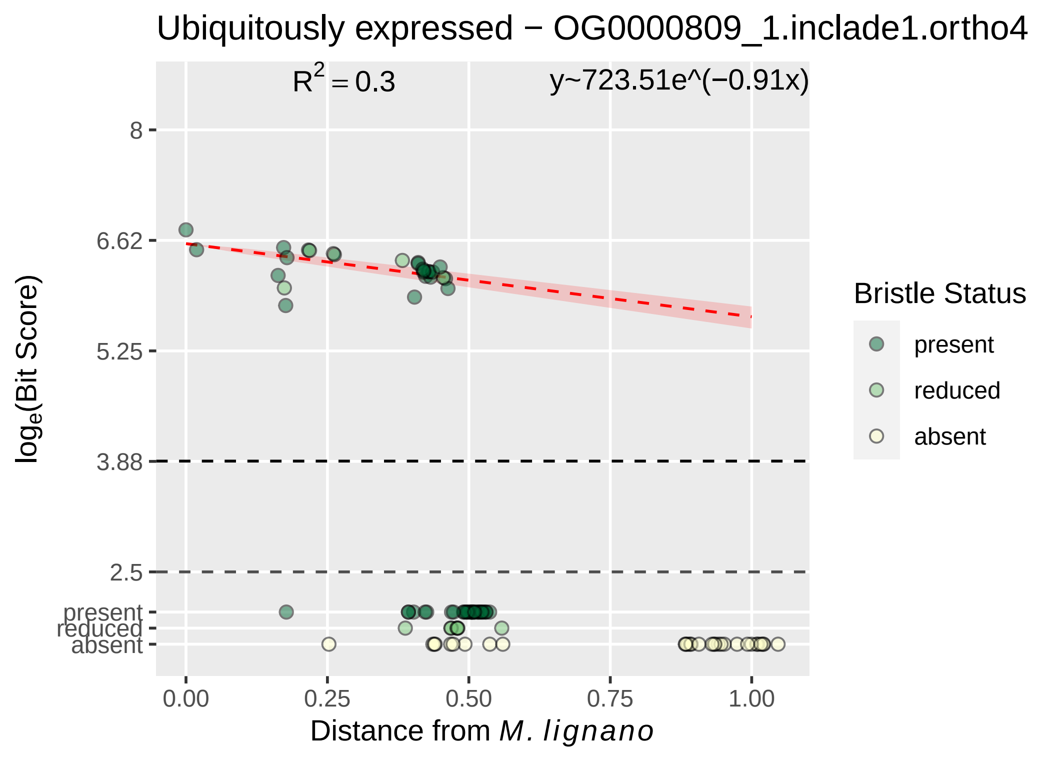

Ubiquitously expressed − OG0000809_1.inclade1.ortho4
R2 = 0.3
y~723.51e^(−0.91x)
8
6.62
5.25
3.88
2.5
present
reduced
absent
loge(Bit Score)
0.00
0.25
0.50
0.75
1.00
Distance from M. lignano
Bristle Status
present
reduced
absent

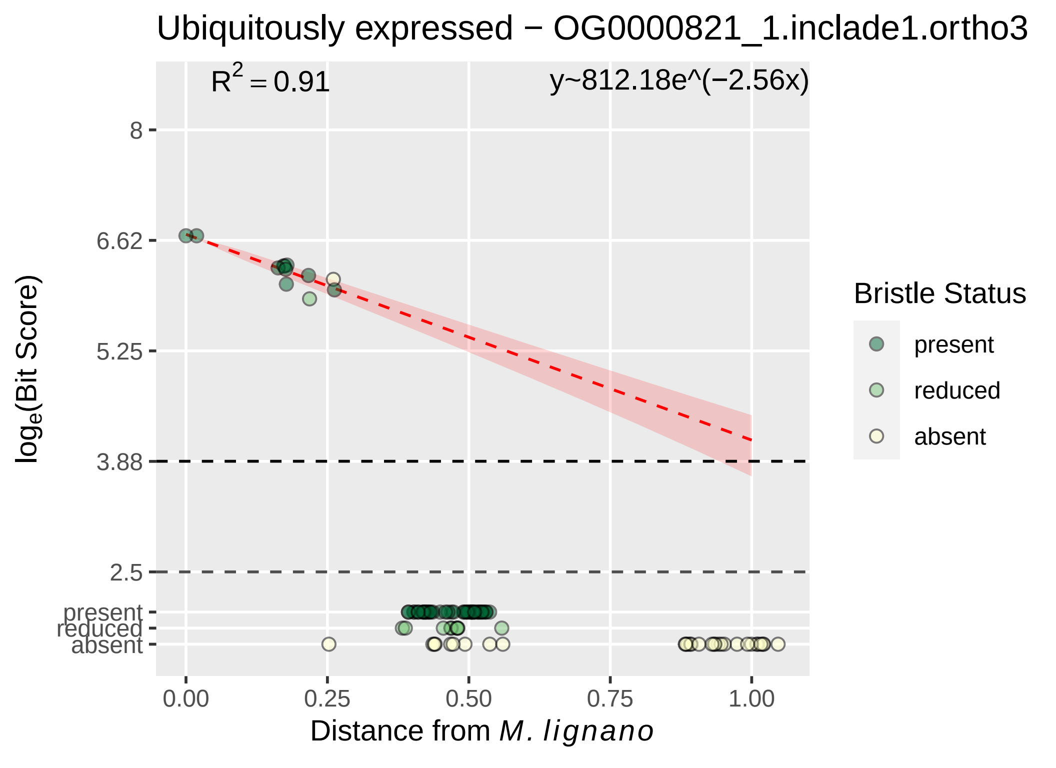

Ubiquitously expressed − OG0000821_1.inclade1.ortho3
R^2 = 0.91
y~812.18e^(−2.56x)
8
6.62
5.25
3.88
2.5
present
reduced
absent
log_e(Bit Score)
0.00
0.25
0.50
0.75
1.00
Distance from M. lignano
Bristle Status
present
reduced
absent

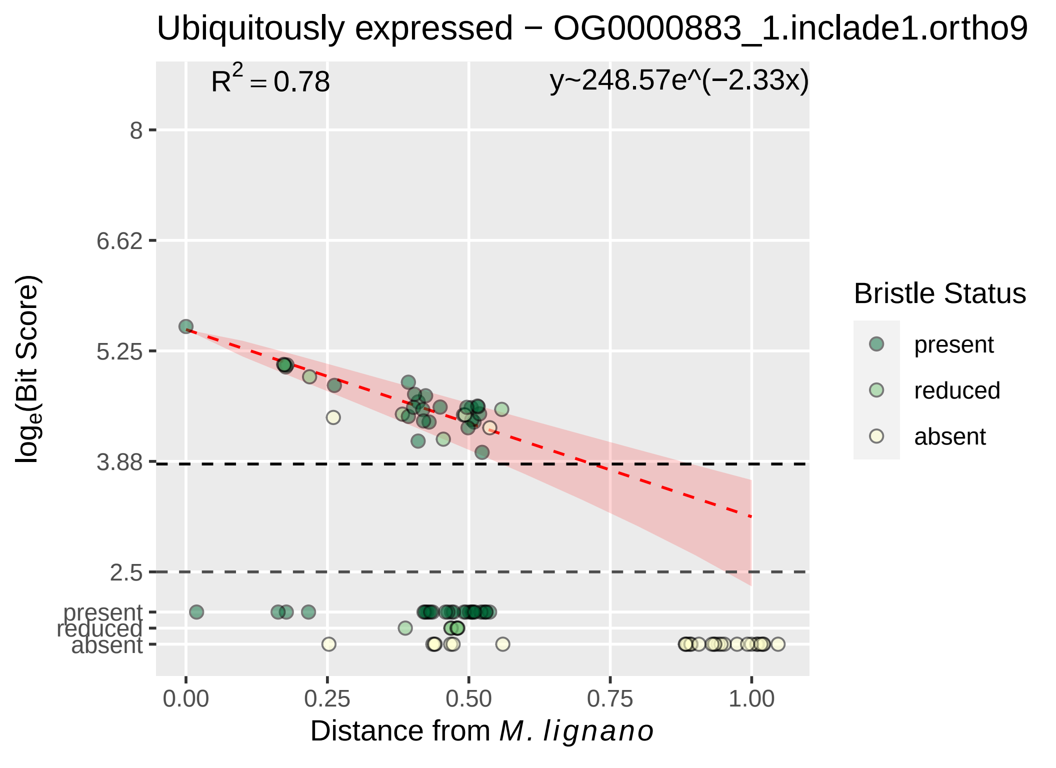
Ubiquitously expressed − OG0000883_1.inclade1.ortho9
R^2 = 0.78
y~248.57e^(−2.33x)
8
6.62
5.25
3.88
2.5
present
reduced
absent
log_e(Bit Score)
0.00
0.25
0.50
0.75
1.00
Distance from M. lignano
Bristle Status
present
reduced
absent

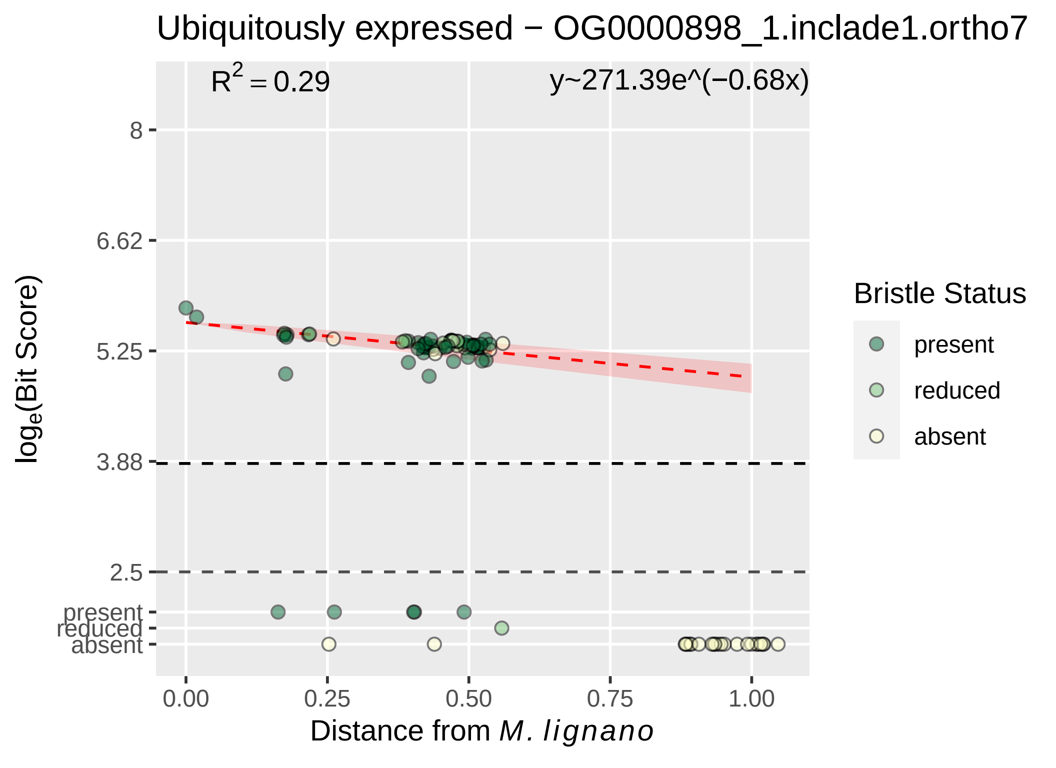

Ubiquitously expressed − OG0000898_1.inclade1.ortho7
$R^2 = 0.29$
y~271.39e^(−0.68x)
8
6.62
5.25
3.88
2.5
present
reduced
absent
$\log_e$(Bit Score)
0.00
0.25
0.50
0.75
1.00
Distance from *M. lignano*
Bristle Status
present
reduced
absent

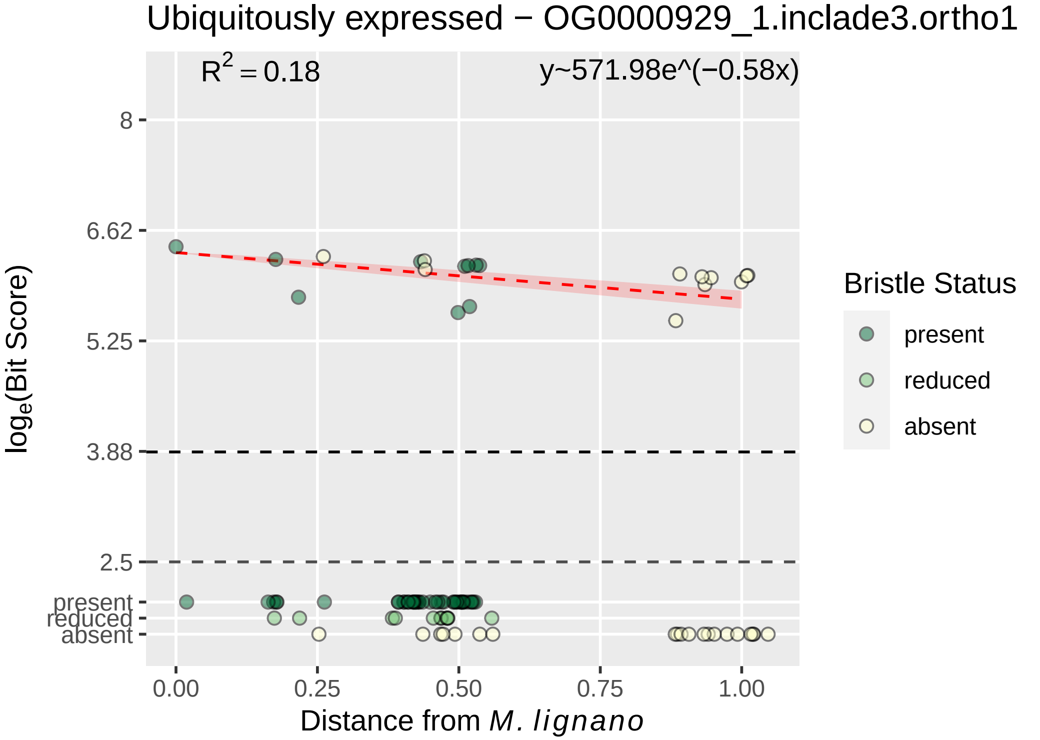

Ubiquitously expressed − OG0000929_1.inclade3.ortho1
$R^2 = 0.18$
y~571.98e^(−0.58x)
8
6.62
5.25
3.88
2.5
present
reduced
absent
$\log_e$(Bit Score)
0.00
0.25
0.50
0.75
1.00
Distance from *M. lignano*
Bristle Status
present
reduced
absent

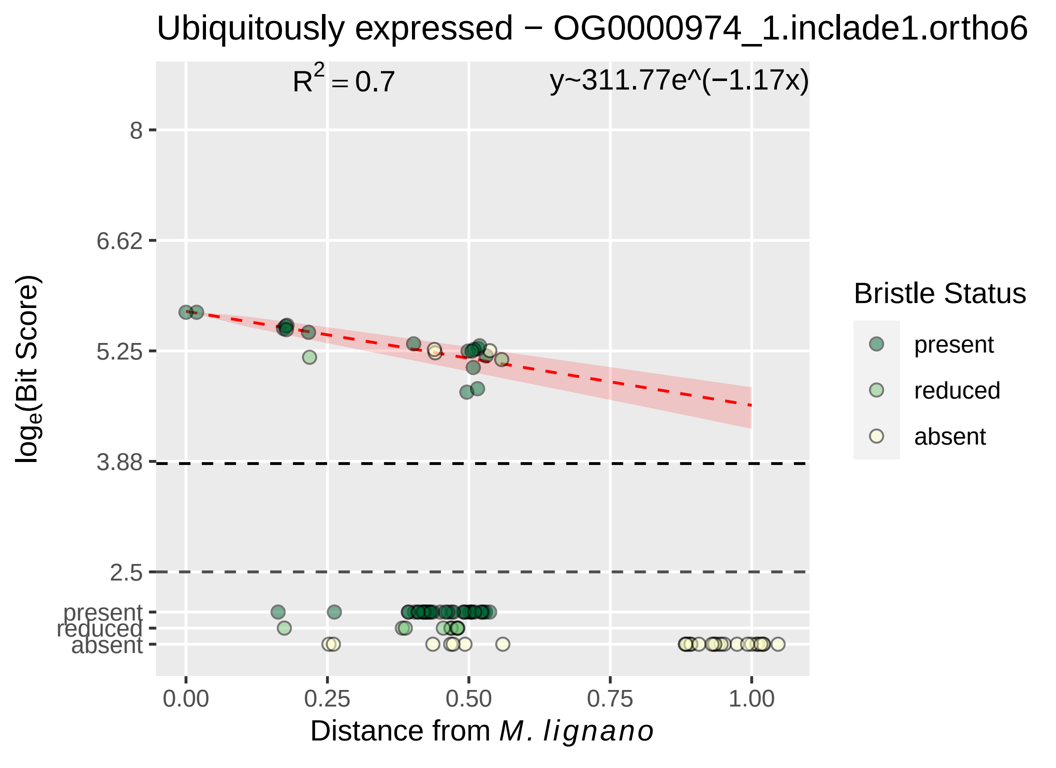

Ubiquitously expressed − OG0000974_1.inclade1.ortho6
$R^2 = 0.7$
y~311.77e^(−1.17x)
8
6.62
5.25
3.88
2.5
present
reduced
absent
$\log_e$(Bit Score)
0.00
0.25
0.50
0.75
1.00
Distance from *M. lignano*
Bristle Status
present
reduced
absent

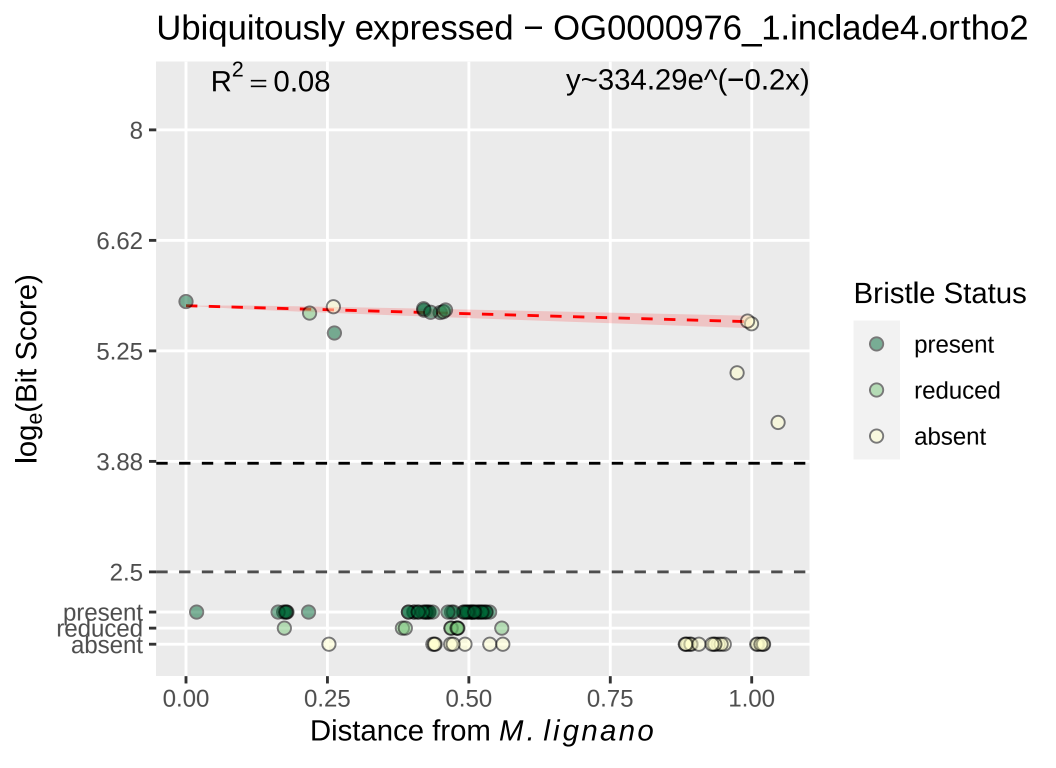

Ubiquitously expressed − OG0000976_1.inclade4.ortho2
R² = 0.08
y~334.29e^(−0.2x)
8
6.62
5.25
3.88
2.5
present
reduced
absent
0.00
0.25
0.50
0.75
1.00
loge(Bit Score)
Distance from M. lignano
Bristle Status
present
reduced
absent

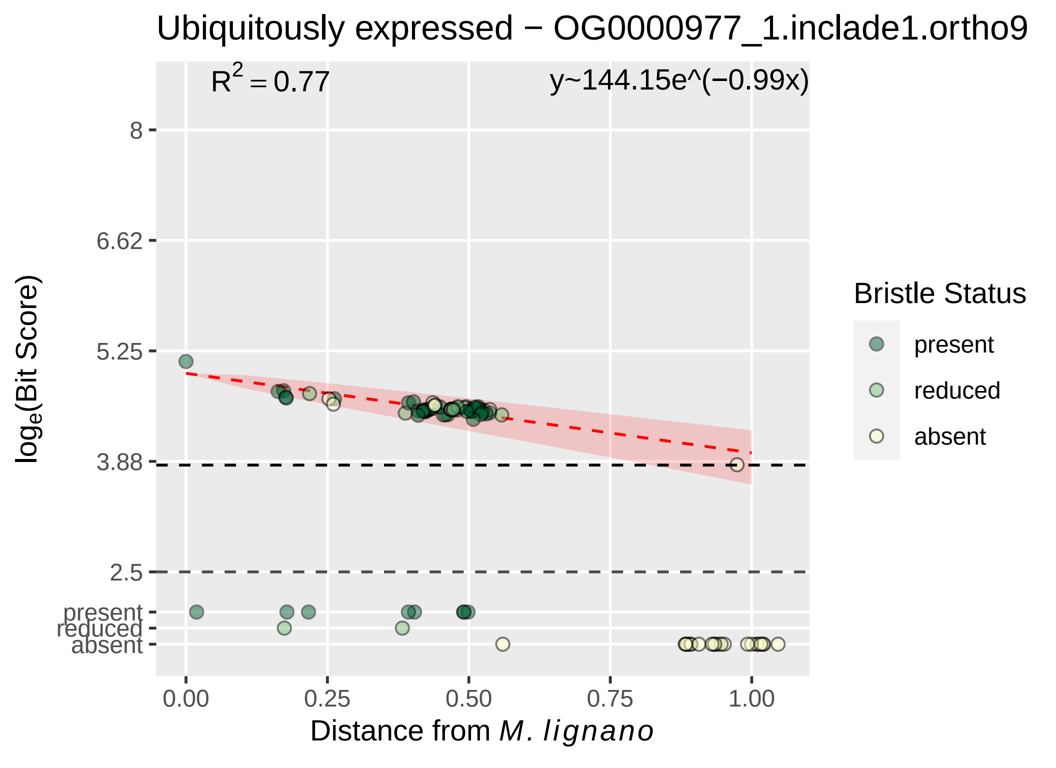

Ubiquitously expressed − OG0000977_1.inclade1.ortho9
R2 = 0.77
y~144.15e^(−0.99x)
8
6.62
5.25
3.88
2.5
present
reduced
absent
0.00
0.25
0.50
0.75
1.00
Distance from M. lignano
loge(Bit Score)
Bristle Status
present
reduced
absent

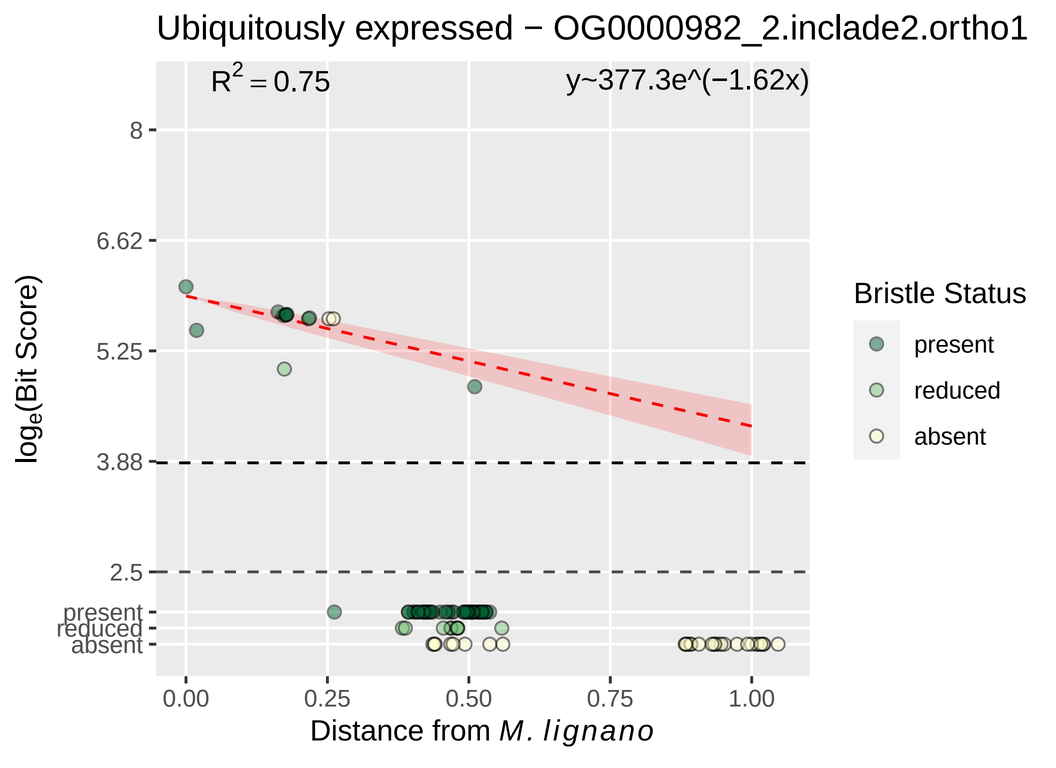

Ubiquitously expressed − OG0000982_2.inclade2.ortho1
$R^2 = 0.75$
y~377.3e^(−1.62x)
8
6.62
5.25
3.88
2.5
present
reduced
absent
0.00
0.25
0.50
0.75
1.00
Distance from M. lignano
log_e(Bit Score)
Bristle Status
present
reduced
absent

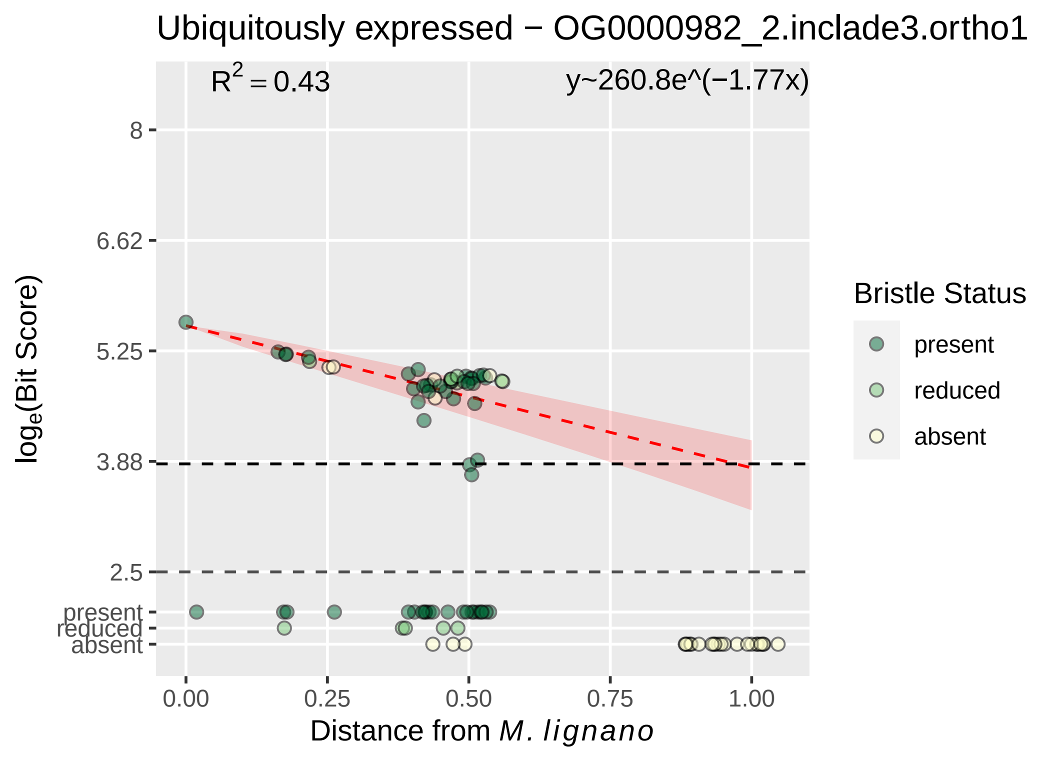

Ubiquitously expressed − OG0000982_2.inclade3.ortho1
R^2 = 0.43
y~260.8e^(−1.77x)
8
6.62
5.25
3.88
2.5
present
reduced
absent
log_e(Bit Score)
0.00
0.25
0.50
0.75
1.00
Distance from M. lignano
Bristle Status
present
reduced
absent

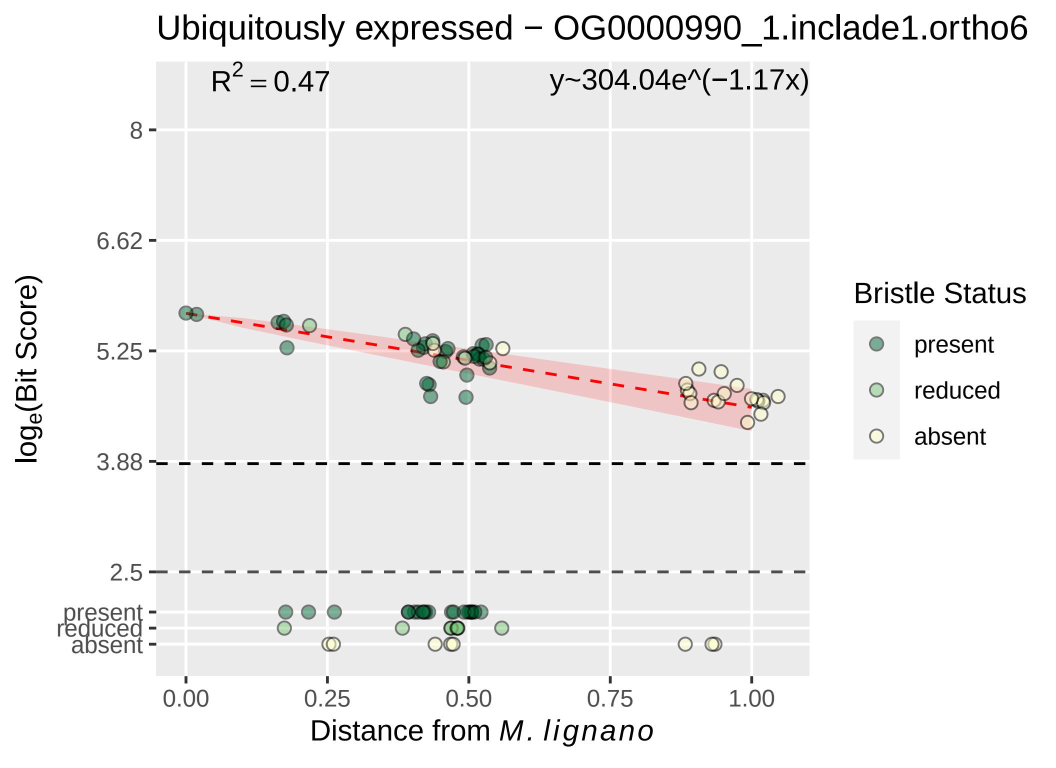
Ubiquitously expressed − OG0000990_1.inclade1.ortho6
R^2 = 0.47
y~304.04e^(−1.17x)
8
6.62
5.25
3.88
2.5
present
reduced
absent
log_e(Bit Score)
0.00
0.25
0.50
0.75
1.00
Distance from M. lignano
Bristle Status
present
reduced
absent

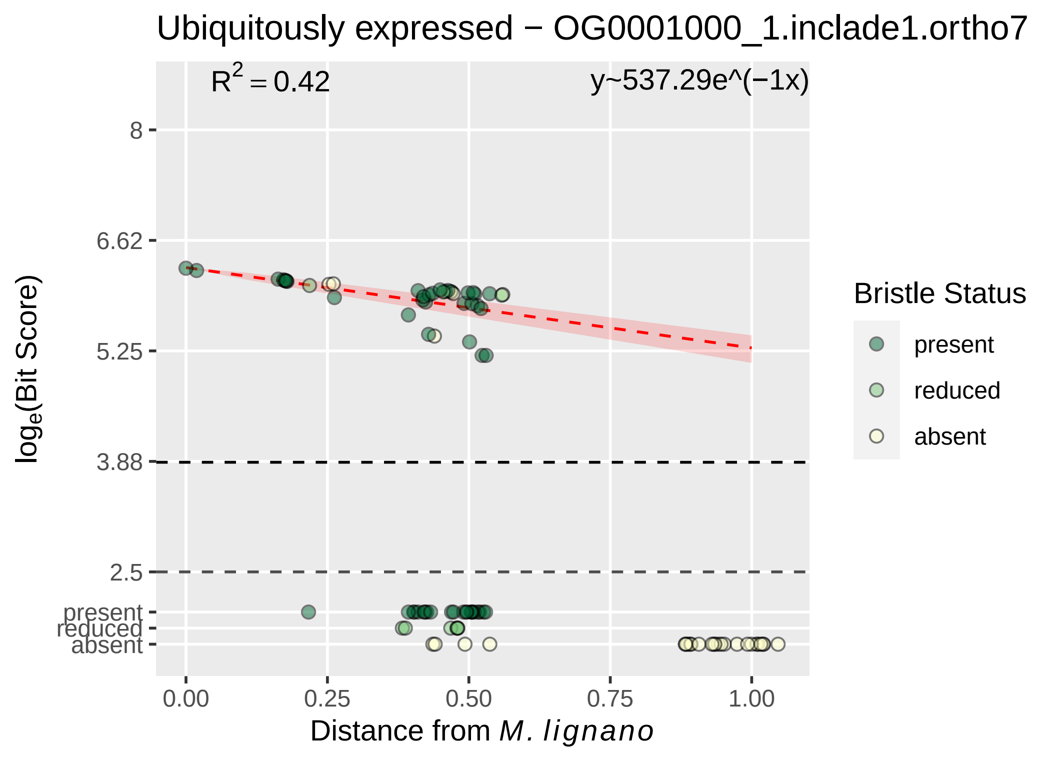

Ubiquitously expressed − OG0001000_1.inclade1.ortho7
R^2 = 0.42
y~537.29e^(−1x)
8
6.62
5.25
3.88
2.5
present
reduced
absent
log_e(Bit Score)
0.00
0.25
0.50
0.75
1.00
Distance from M. lignano
Bristle Status
present
reduced
absent

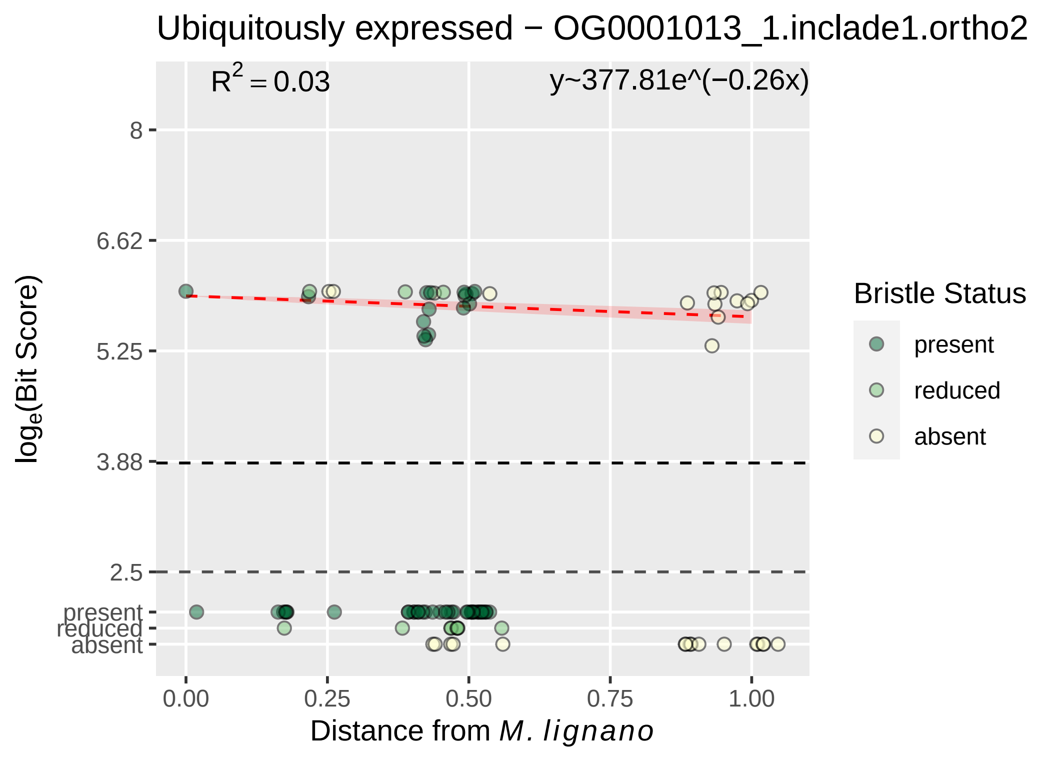

Ubiquitously expressed − OG0001013_1.inclade1.ortho2
R2 = 0.03
y~377.81e^(−0.26x)
8
6.62
5.25
3.88
2.5
present
reduced
absent
0.00
0.25
0.50
0.75
1.00
Distance from M. lignano
loge(Bit Score)
Bristle Status
present
reduced
absent

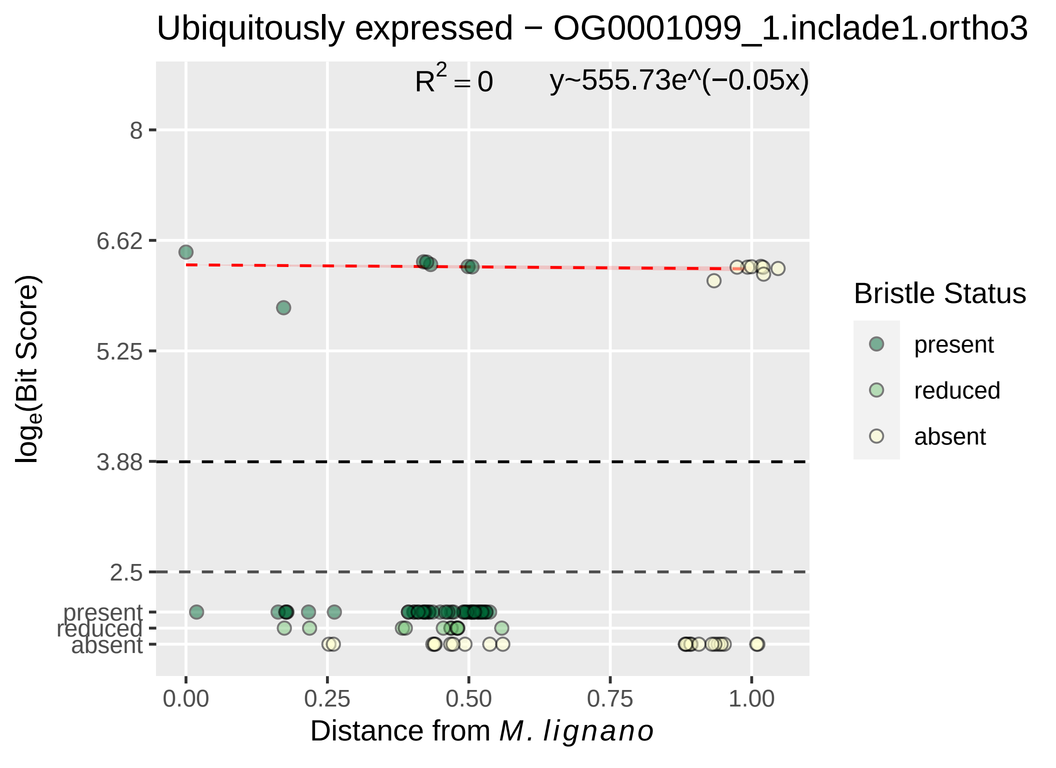

Ubiquitously expressed − OG0001099_1.inclade1.ortho3
R^2 = 0
y~555.73e^(−0.05x)
log_e(Bit Score)
8
6.62
5.25
3.88
2.5
present
reduced
absent
0.00
0.25
0.50
0.75
1.00
Distance from M. lignano
Bristle Status
present
reduced
absent

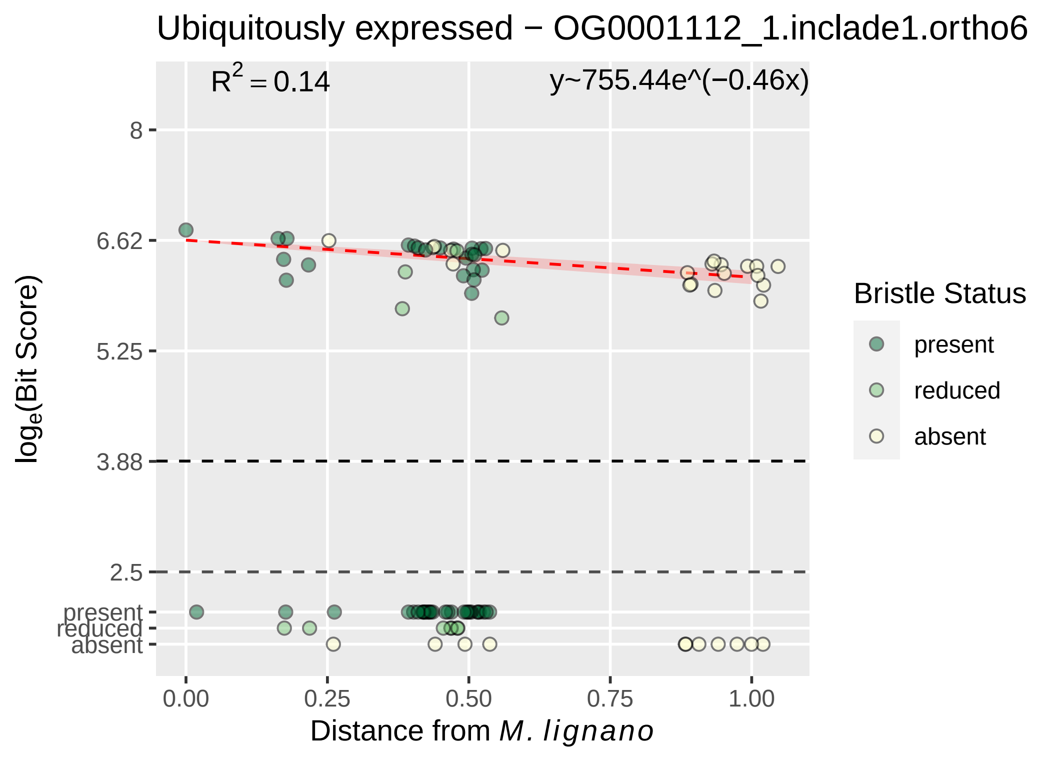

Ubiquitously expressed − OG0001112_1.inclade1.ortho6
R2 = 0.14
y~755.44e^(−0.46x)
8
6.62
5.25
3.88
2.5
present
reduced
absent
loge(Bit Score)
0.00
0.25
0.50
0.75
1.00
Distance from M. lignano
Bristle Status
present
reduced
absent

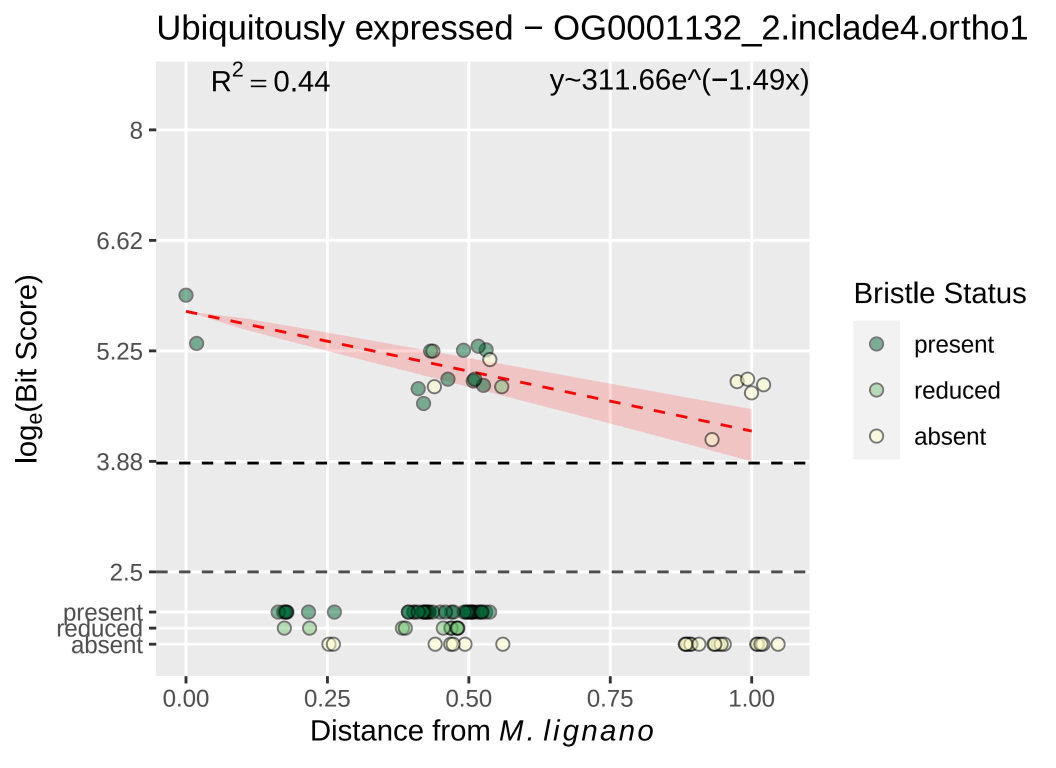

Ubiquitously expressed − OG0001132_2.inclade4.ortho1
R2 = 0.44
y~311.66e^(−1.49x)
loge(Bit Score)
8
6.62
5.25
3.88
2.5
present
reduced
absent
0.00
0.25
0.50
0.75
1.00
Distance from M. lignano
Bristle Status
present
reduced
absent

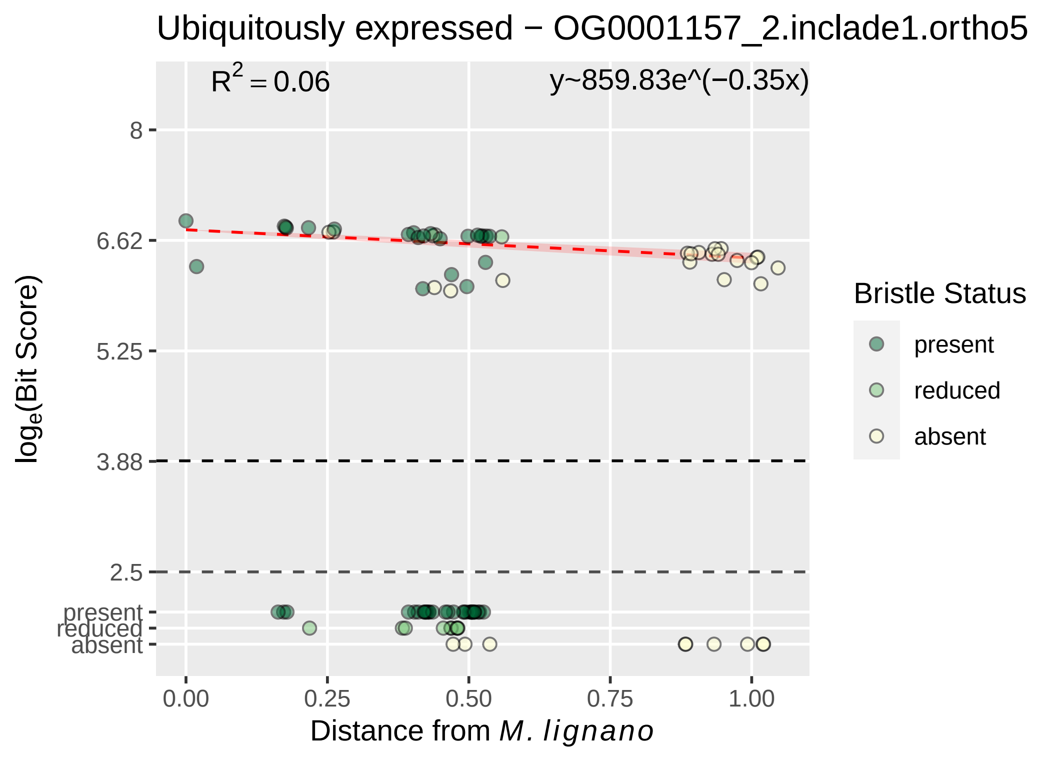
Ubiquitously expressed − OG0001157_2.inclade1.ortho5
R^2 = 0.06
y~859.83e^(−0.35x)
8
6.62
5.25
3.88
2.5
present
reduced
absent
0.00
0.25
0.50
0.75
1.00
log_e(Bit Score)
Distance from M. lignano
Bristle Status
present
reduced
absent

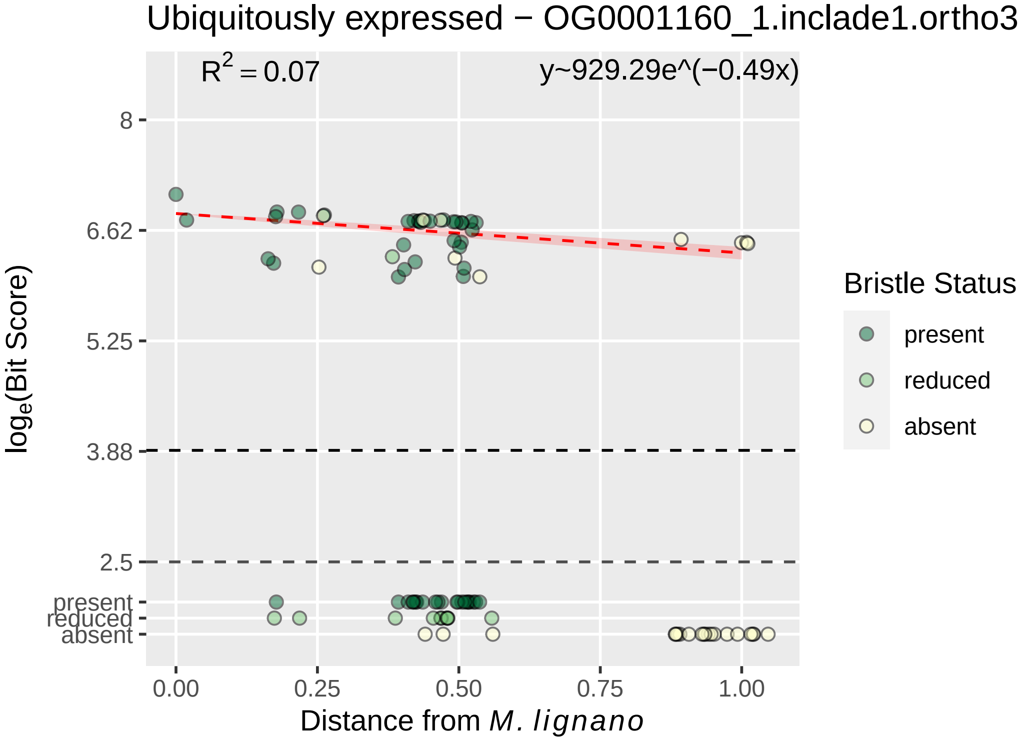

Ubiquitously expressed − OG0001160_1.inclade1.ortho3
$R^2 = 0.07$
y~929.29e^(−0.49x)
8
6.62
5.25
3.88
2.5
present
reduced
absent
0.00
0.25
0.50
0.75
1.00
log$_e$(Bit Score)
Distance from *M. lignano*
Bristle Status
present
reduced
absent

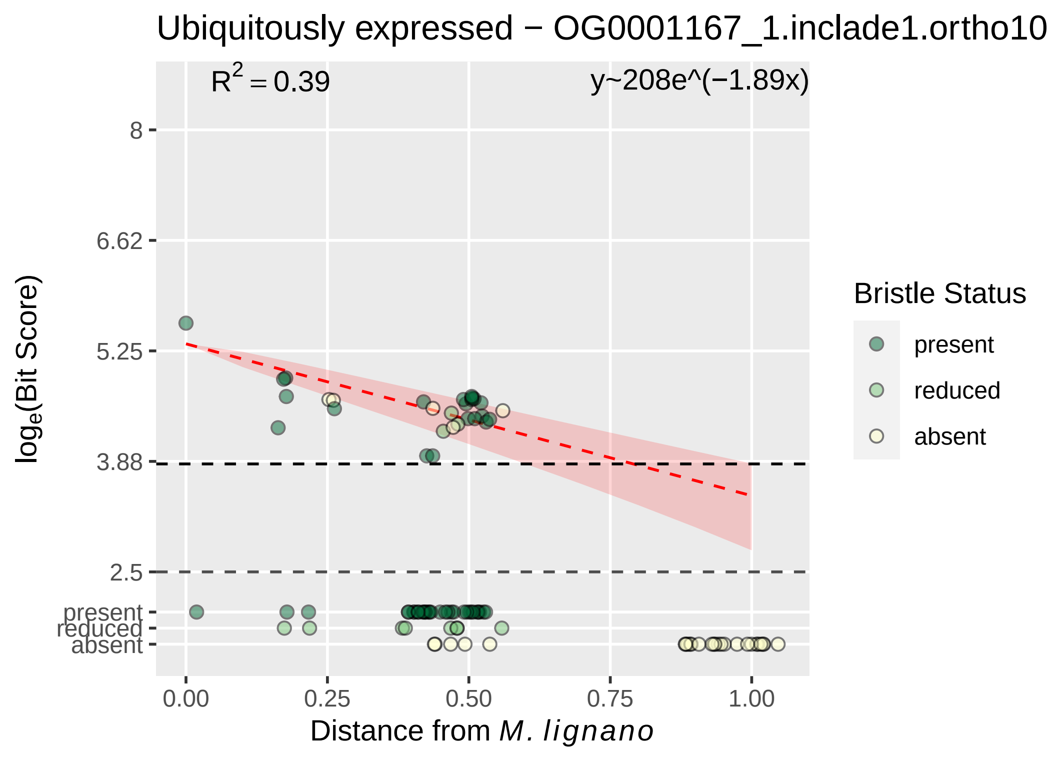

Ubiquitously expressed − OG0001167_1.inclade1.ortho10
R^2 = 0.39
y~208e^(−1.89x)
8
6.62
5.25
3.88
2.5
present
reduced
absent
log_e(Bit Score)
0.00
0.25
0.50
0.75
1.00
Distance from M. lignano
Bristle Status
present
reduced
absent

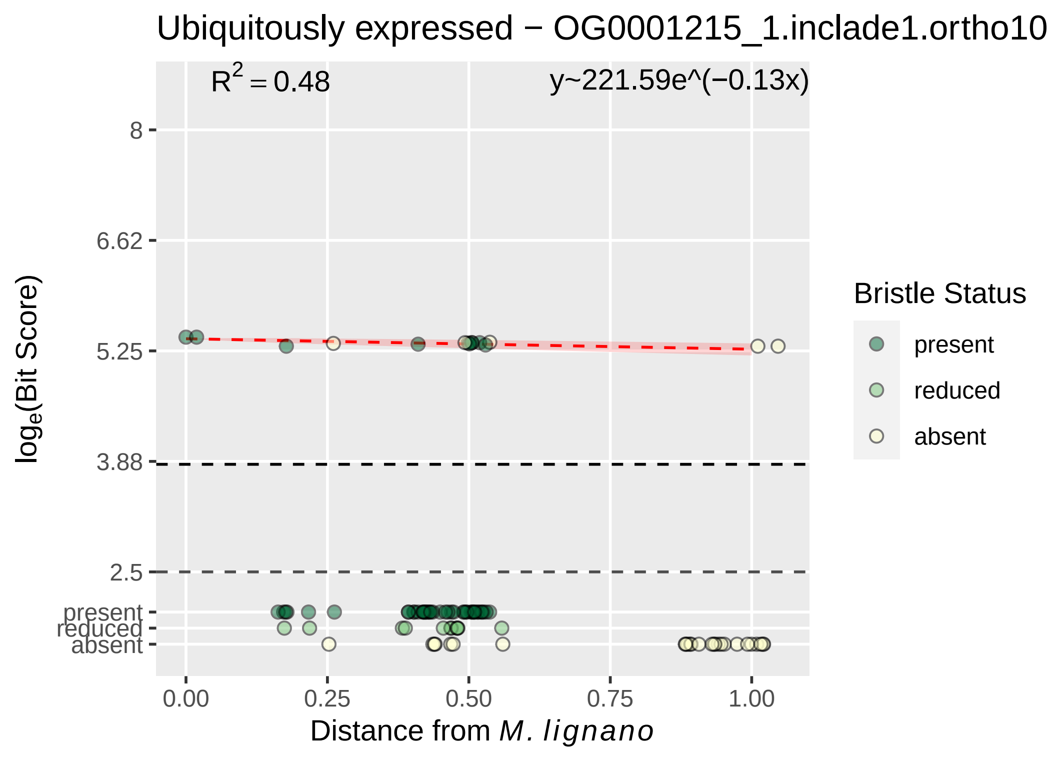
Ubiquitously expressed − OG0001215_1.inclade1.ortho10
$R^2 = 0.48$
y~221.59e^(−0.13x)
8
6.62
5.25
3.88
2.5
present
reduced
absent
$\log_e$(Bit Score)
0.00
0.25
0.50
0.75
1.00
Distance from *M. lignano*
Bristle Status
present
reduced
absent

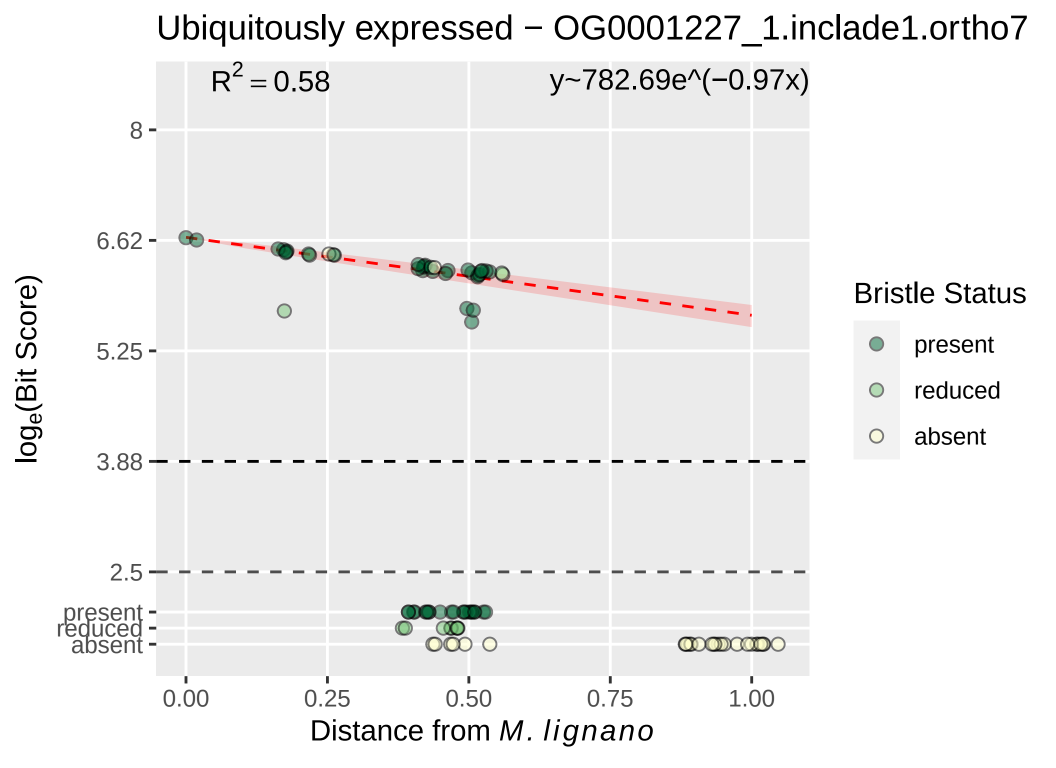

Ubiquitously expressed − OG0001227_1.inclade1.ortho7
R^2 = 0.58
y~782.69e^(−0.97x)
log_e(Bit Score)
8
6.62
5.25
3.88
2.5
present
reduced
absent
0.00
0.25
0.50
0.75
1.00
Distance from M. lignano
Bristle Status
present
reduced
absent

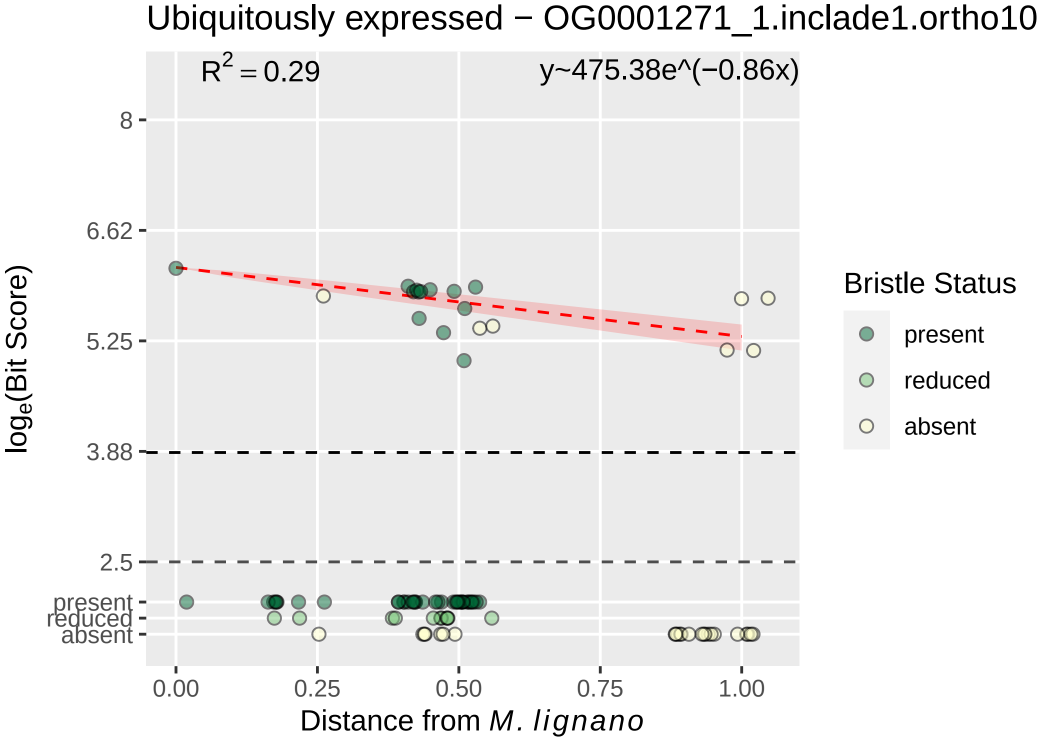

Ubiquitously expressed − OG0001271_1.inclade1.ortho10
$R^2 = 0.29$
y~475.38e^(−0.86x)
8
6.62
5.25
3.88
2.5
present
reduced
absent
$\log_e$(Bit Score)
0.00
0.25
0.50
0.75
1.00
Distance from *M. lignano*
Bristle Status
present
reduced
absent

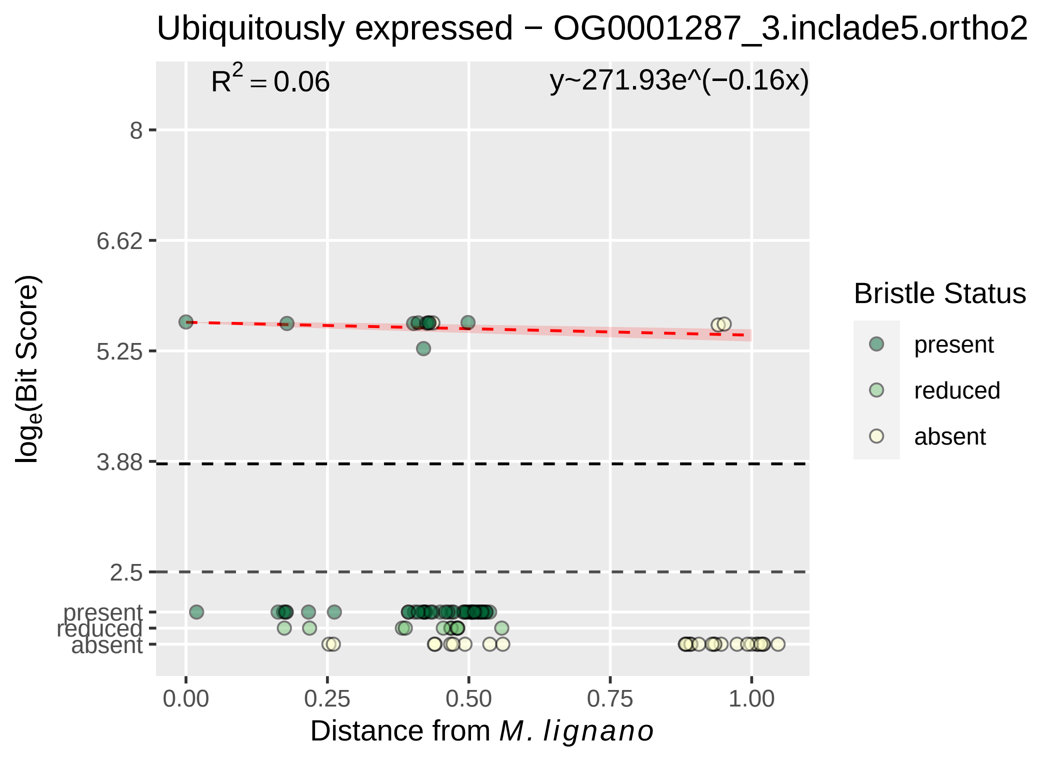

Ubiquitously expressed − OG0001287_3.inclade5.ortho2
$R^2 = 0.06$
y~271.93e^(−0.16x)
8
6.62
5.25
3.88
2.5
present
reduced
absent
$log_e$(Bit Score)
0.00
0.25
0.50
0.75
1.00
Distance from M. lignano
Bristle Status
present
reduced
absent

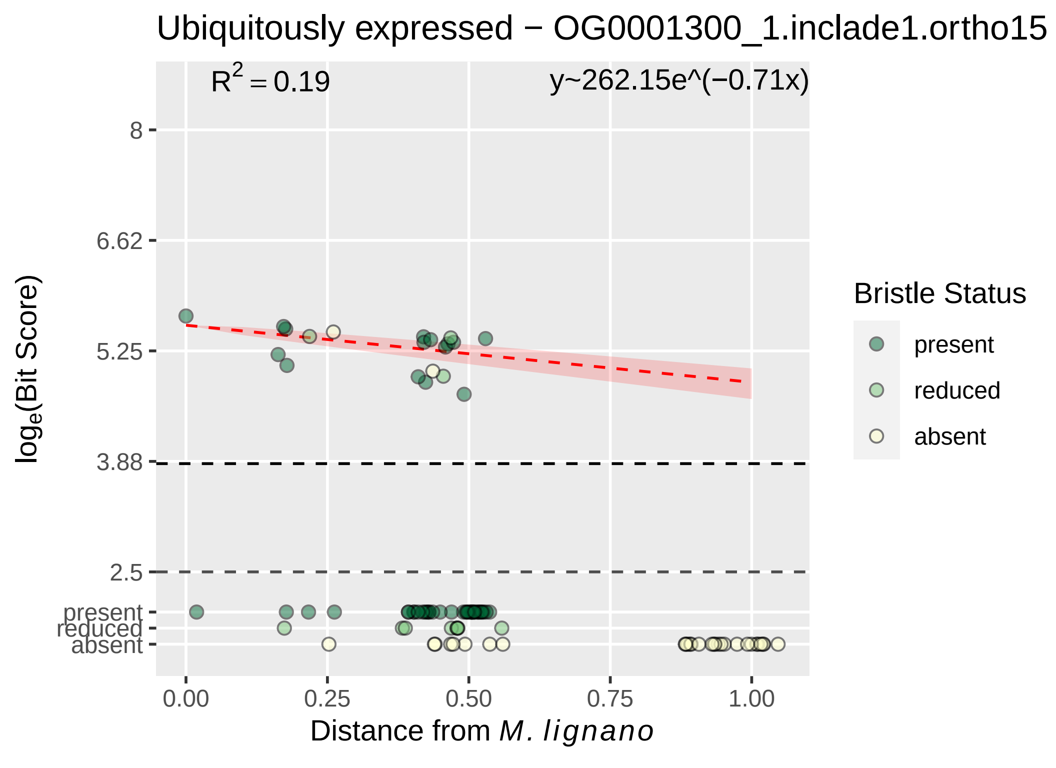

Ubiquitously expressed − OG0001300_1.inclade1.ortho15
R^2 = 0.19
y~262.15e^(−0.71x)
8
6.62
5.25
3.88
2.5
present
reduced
absent
log_e(Bit Score)
0.00
0.25
0.50
0.75
1.00
Distance from M. lignano
Bristle Status
present
reduced
absent

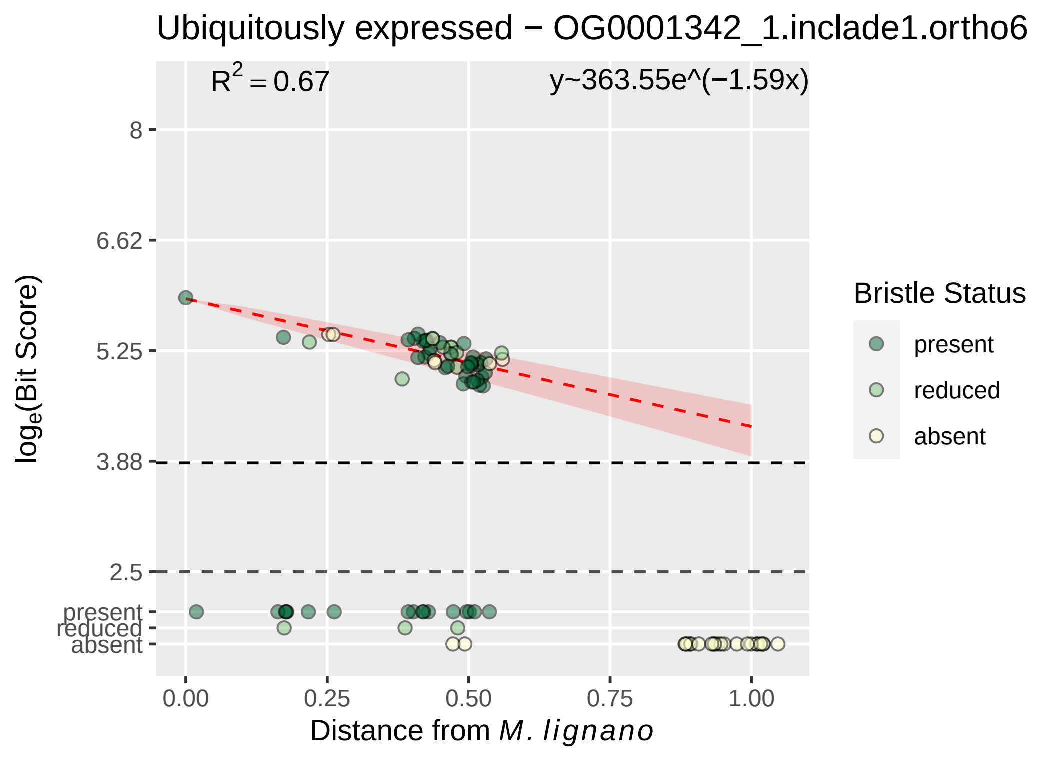

Ubiquitously expressed − OG0001342_1.inclade1.ortho6
R^2 = 0.67
y~363.55e^(−1.59x)
8
6.62
5.25
3.88
2.5
present
reduced
absent
log_e(Bit Score)
0.00
0.25
0.50
0.75
1.00
Distance from M. lignano
Bristle Status
present
reduced
absent

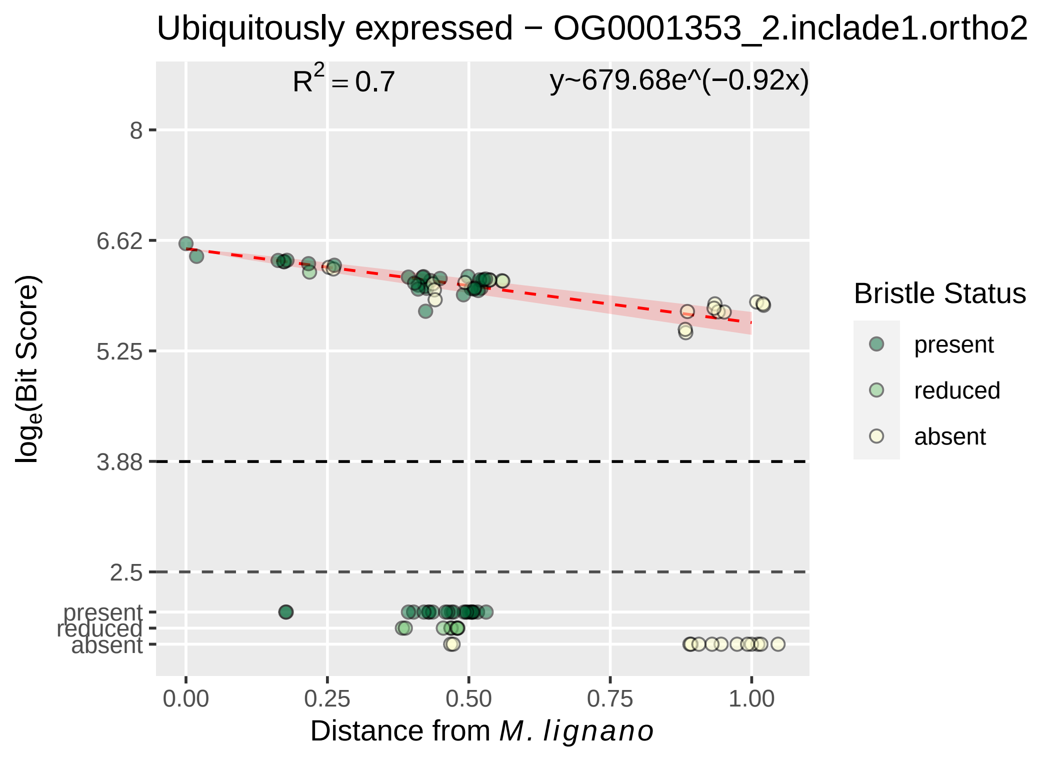

Ubiquitously expressed − OG0001353_2.inclade1.ortho2
R^2 = 0.7
y~679.68e^(−0.92x)
8
6.62
5.25
3.88
2.5
present
reduced
absent
0.00
0.25
0.50
0.75
1.00
loge(Bit Score)
Distance from M. lignano
Bristle Status
present
reduced
absent

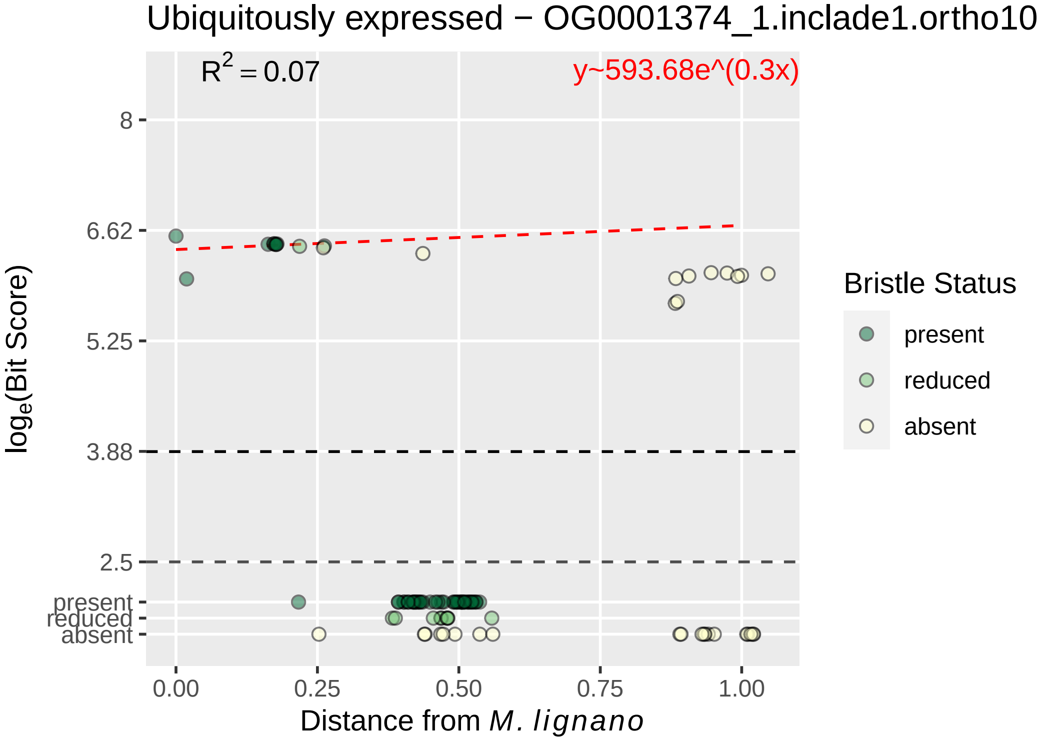
Ubiquitously expressed − OG0001374_1.inclade1.ortho10
$R^2 = 0.07$
y~593.68e^(0.3x)
8
6.62
5.25
3.88
2.5
present
reduced
absent
0.00
0.25
0.50
0.75
1.00
log$_e$(Bit Score)
Distance from *M. lignano*
Bristle Status
present
reduced
absent

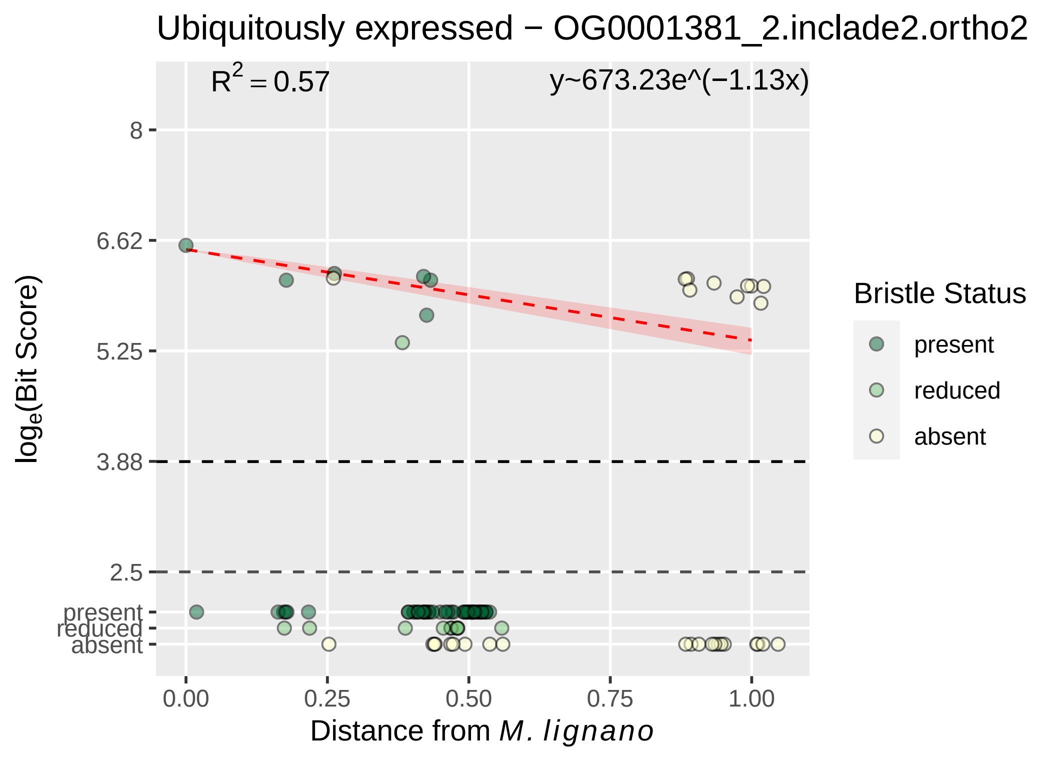

Ubiquitously expressed − OG0001381_2.inclade2.ortho2
$R^2 = 0.57$
y~673.23e^(−1.13x)
8
6.62
5.25
3.88
2.5
present
reduced
absent
0.00
0.25
0.50
0.75
1.00
$\log_e$(Bit Score)
Distance from *M. lignano*
Bristle Status
present
reduced
absent

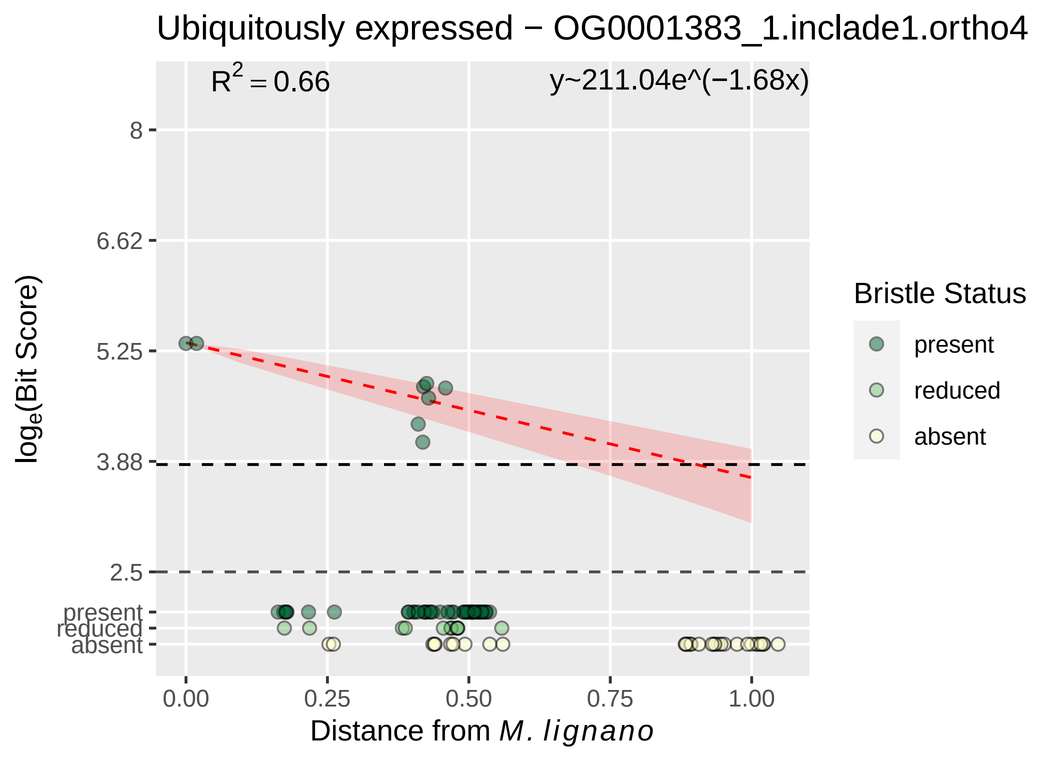

Ubiquitously expressed − OG0001383_1.inclade1.ortho4
R^2 = 0.66
y~211.04e^(−1.68x)
8
6.62
5.25
3.88
2.5
present
reduced
absent
log_e(Bit Score)
0.00
0.25
0.50
0.75
1.00
Distance from M. lignano
Bristle Status
present
reduced
absent

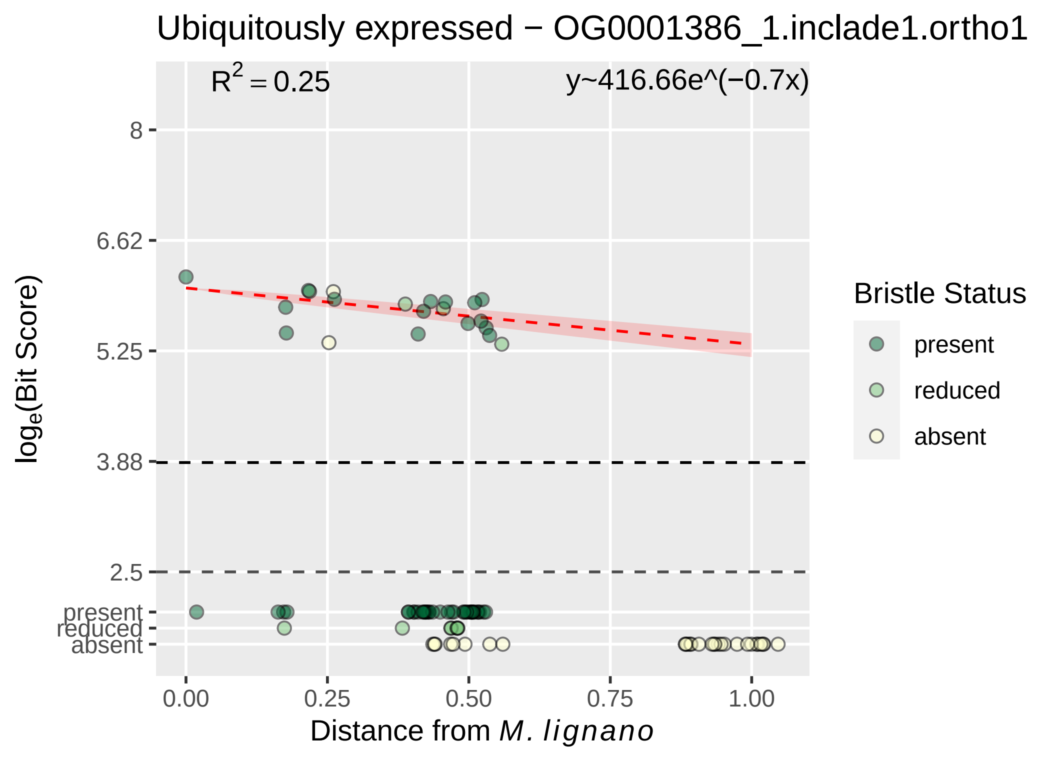

Ubiquitously expressed − OG0001386_1.inclade1.ortho1
$R^2 = 0.25$
y~416.66e^(−0.7x)
8
6.62
5.25
3.88
2.5
present
reduced
absent
$\log_e$(Bit Score)
0.00
0.25
0.50
0.75
1.00
Distance from *M. lignano*
Bristle Status
present
reduced
absent

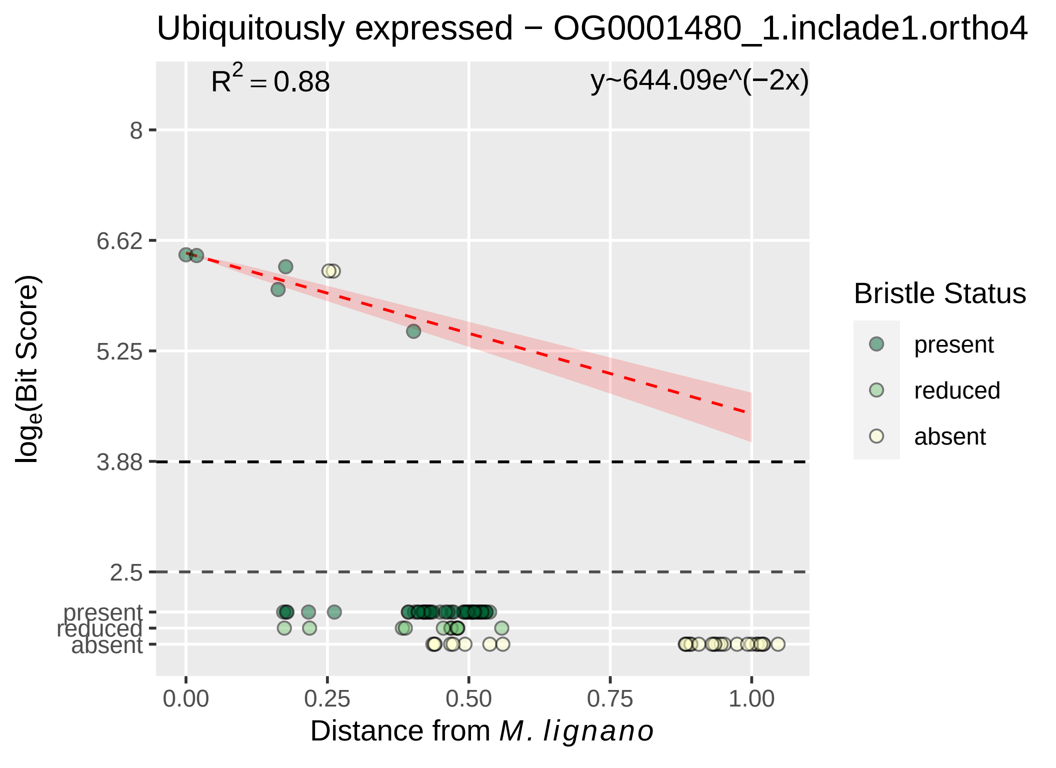

Ubiquitously expressed − OG0001480_1.inclade1.ortho4
R^2 = 0.88
y~644.09e^(−2x)
8
6.62
5.25
3.88
2.5
present
reduced
absent
log_e(Bit Score)
0.00
0.25
0.50
0.75
1.00
Distance from M. lignano
Bristle Status
present
reduced
absent

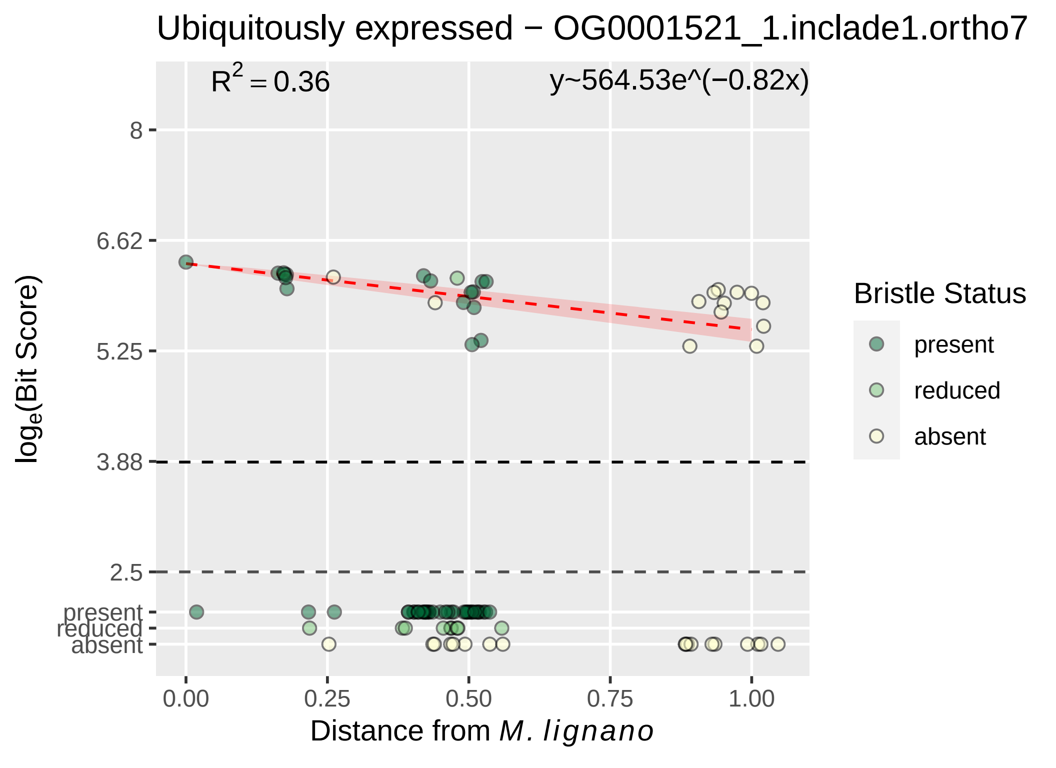
Ubiquitously expressed − OG0001521_1.inclade1.ortho7
$R^2 = 0.36$
y~564.53e^(−0.82x)
8
6.62
5.25
3.88
2.5
present
reduced
absent
0.00
0.25
0.50
0.75
1.00
$log_e$(Bit Score)
Distance from *M. lignano*
Bristle Status
present
reduced
absent

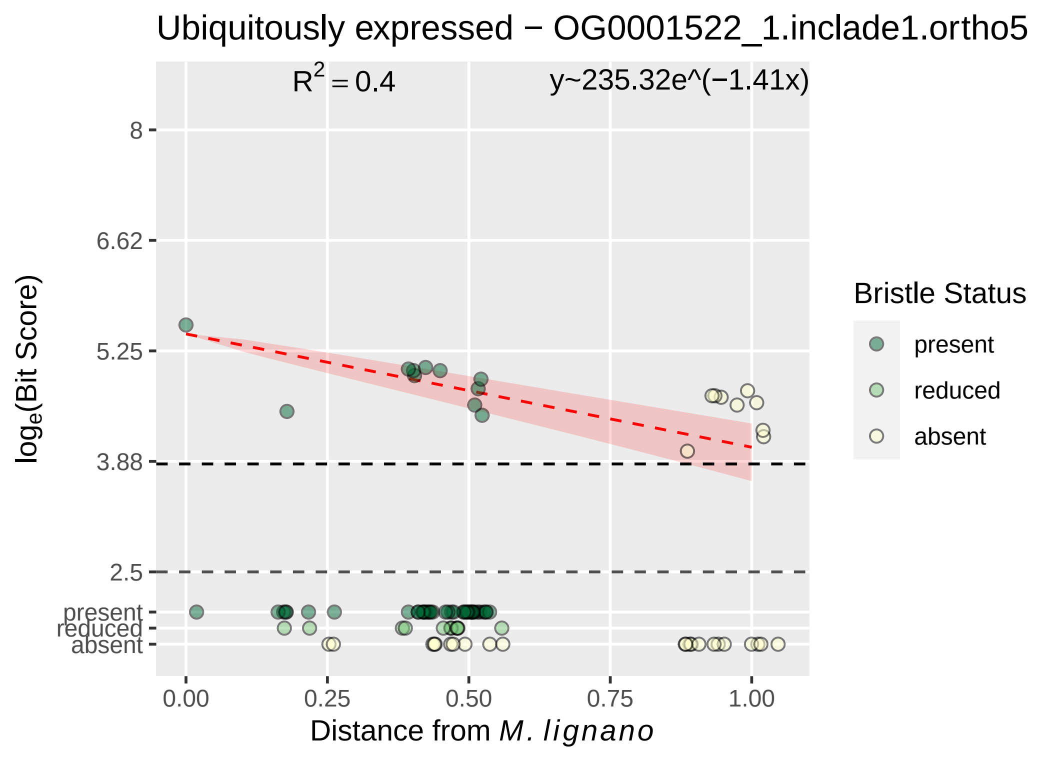

Ubiquitously expressed − OG0001522_1.inclade1.ortho5
R2 = 0.4
y~235.32e^(−1.41x)
8
6.62
5.25
3.88
2.5
present
reduced
absent
0.00
0.25
0.50
0.75
1.00
Distance from M. lignano
loge(Bit Score)
Bristle Status
present
reduced
absent

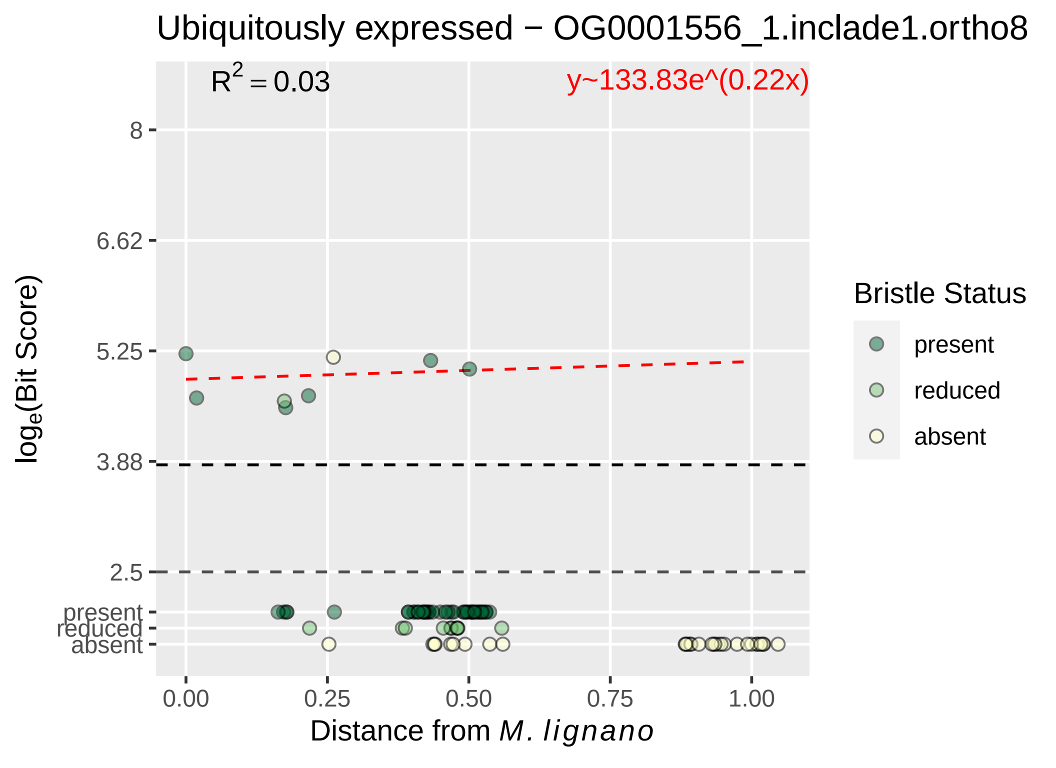
Ubiquitously expressed − OG0001556_1.inclade1.ortho8
R^2 = 0.03
y~133.83e^(0.22x)
8
6.62
5.25
3.88
2.5
present
reduced
absent
0.00
0.25
0.50
0.75
1.00
Distance from M. lignano
loge(Bit Score)
Bristle Status
present
reduced
absent

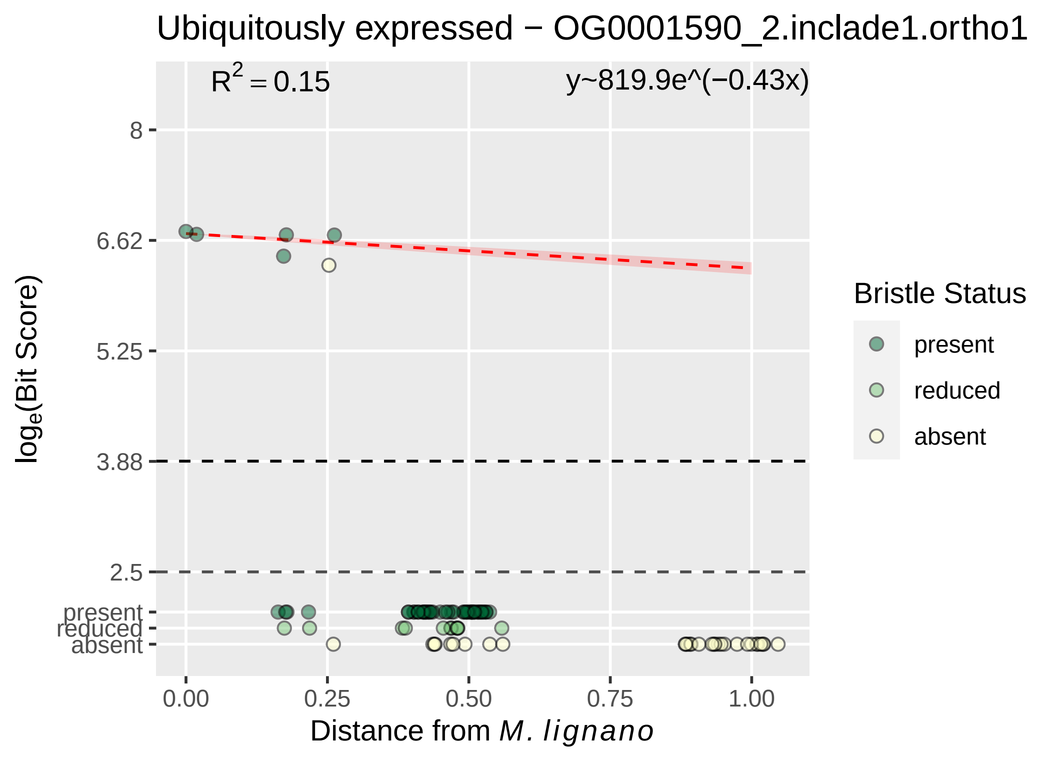

Ubiquitously expressed − OG0001590_2.inclade1.ortho1
R^2 = 0.15
y~819.9e^(−0.43x)
8
6.62
5.25
3.88
2.5
present
reduced
absent
0.00
0.25
0.50
0.75
1.00
log_e(Bit Score)
Distance from M. lignano
Bristle Status
present
reduced
absent

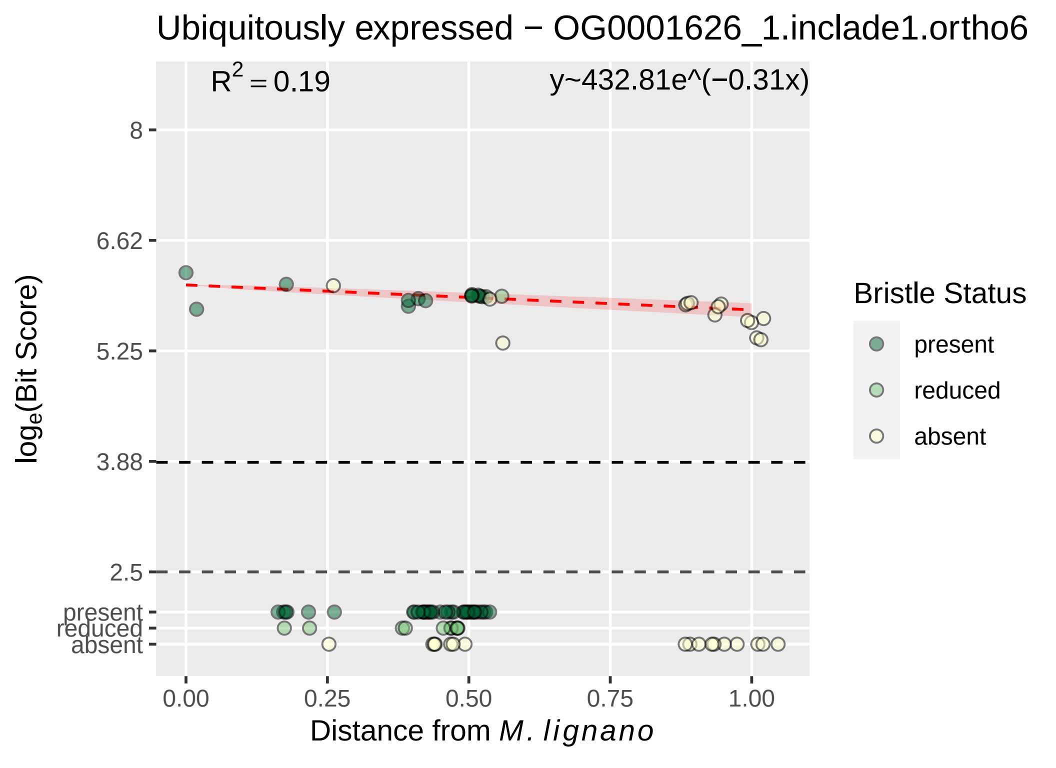

Ubiquitously expressed − OG0001626_1.inclade1.ortho6
$R^2 = 0.19$
y~432.81e^(−0.31x)
8
6.62
5.25
3.88
2.5
present
reduced
absent
0.00
0.25
0.50
0.75
1.00
Distance from M. lignano
log$_e$(Bit Score)
Bristle Status
present
reduced
absent

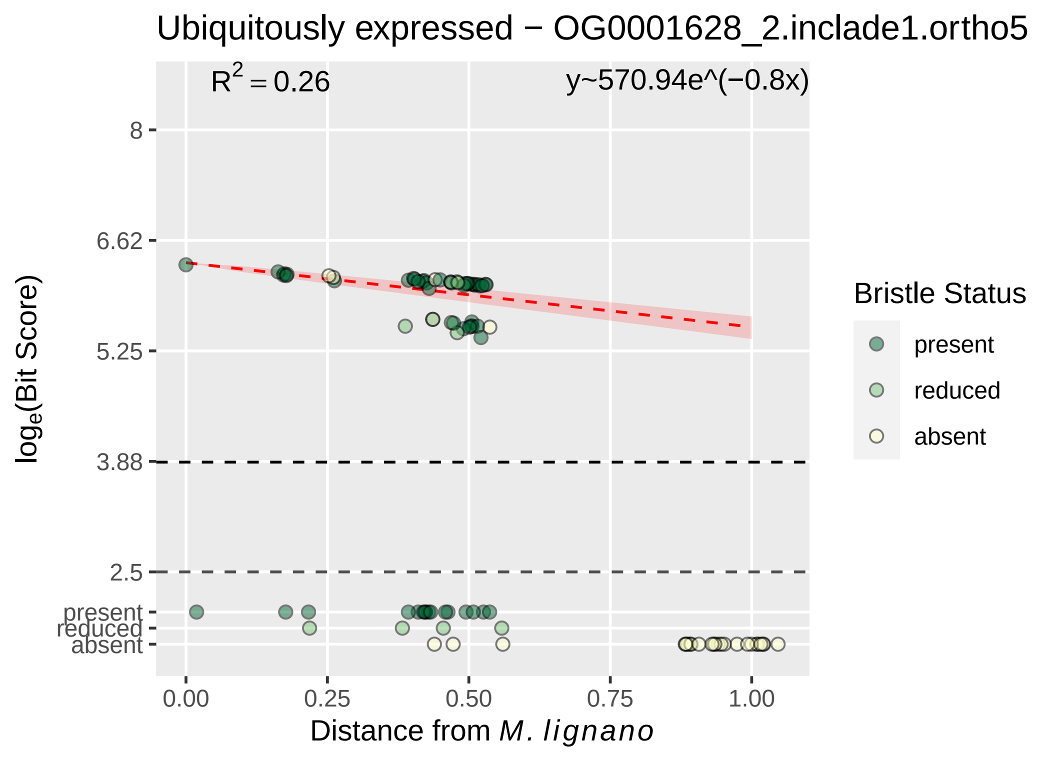

Ubiquitously expressed − OG0001628_2.inclade1.ortho5
R^2 = 0.26
y~570.94e^(−0.8x)
8
6.62
5.25
3.88
2.5
present
reduced
absent
0.00
0.25
0.50
0.75
1.00
Distance from M. lignano
log_e(Bit Score)
Bristle Status
present
reduced
absent

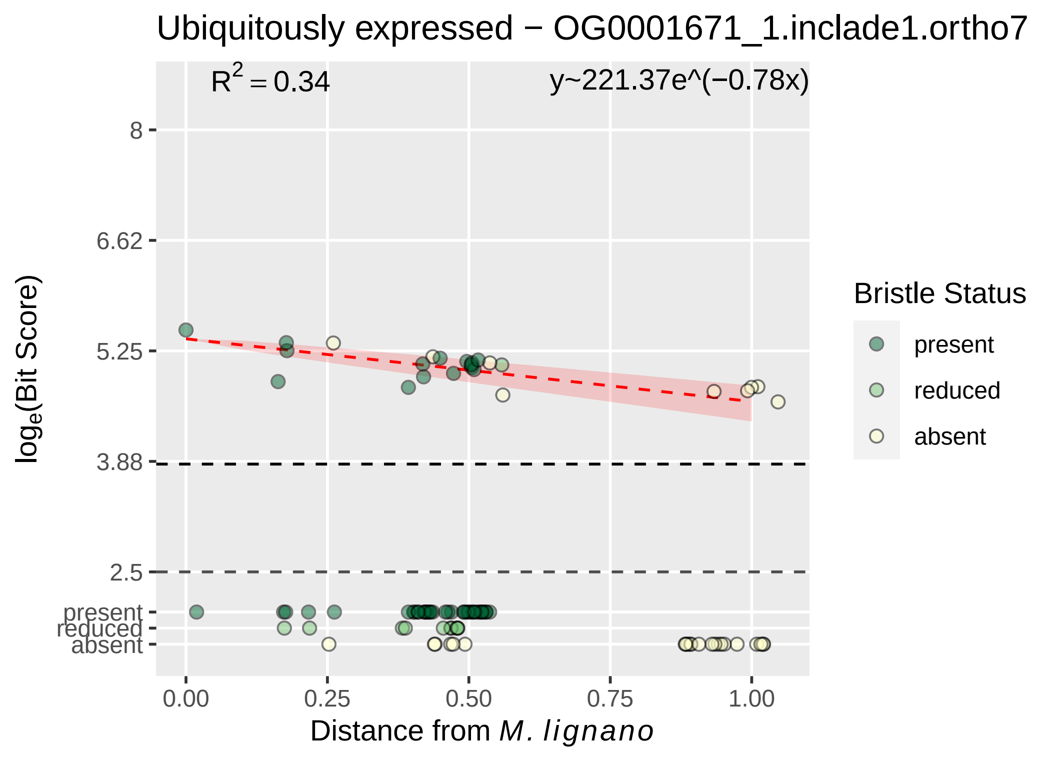

Ubiquitously expressed − OG0001671_1.inclade1.ortho7
R² = 0.34
y~221.37e^(−0.78x)
8
6.62
5.25
3.88
2.5
present
reduced
absent
0.00
0.25
0.50
0.75
1.00
loge(Bit Score)
Distance from M. lignano
Bristle Status
present
reduced
absent

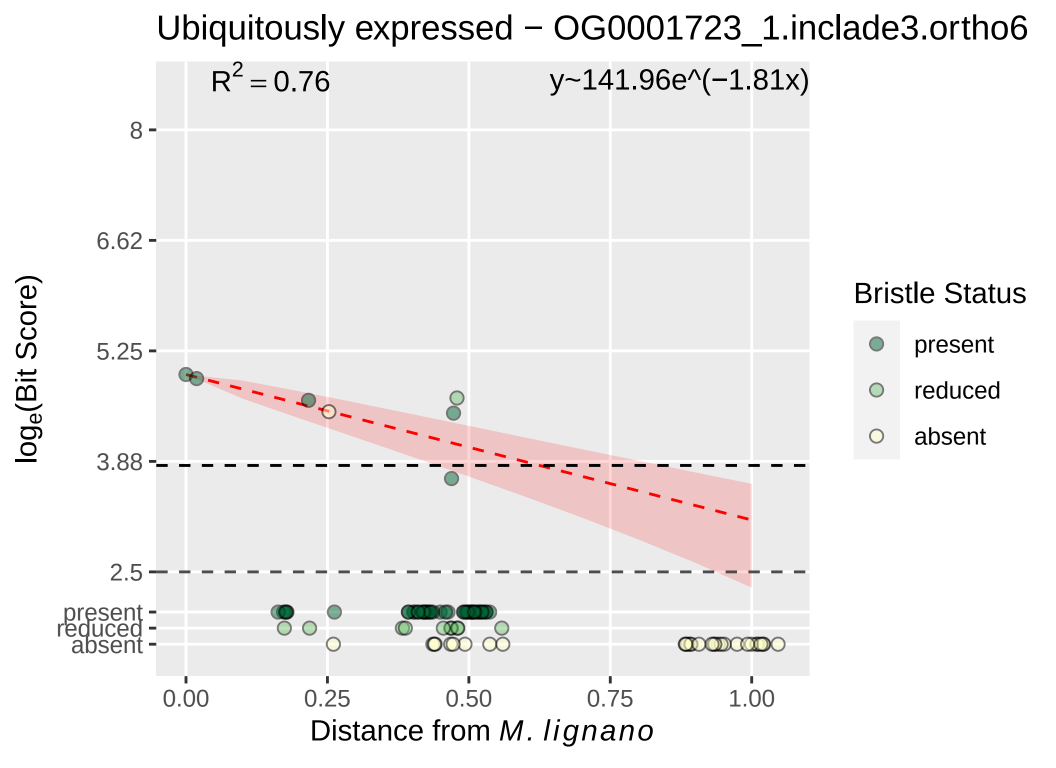

Ubiquitously expressed − OG0001723_1.inclade3.ortho6
R^2 = 0.76
y~141.96e^(−1.81x)
8
6.62
5.25
3.88
2.5
present
reduced
absent
log_e(Bit Score)
0.00
0.25
0.50
0.75
1.00
Distance from M. lignano
Bristle Status
present
reduced
absent

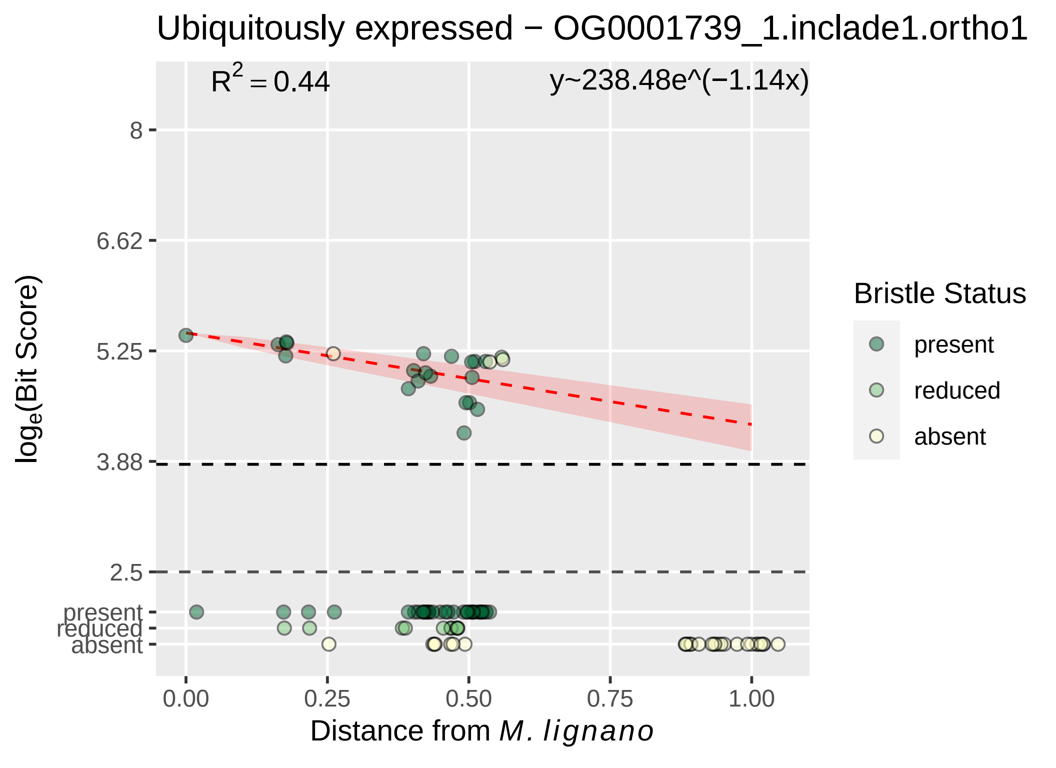

Ubiquitously expressed − OG0001739_1.inclade1.ortho1
R2 = 0.44
y~238.48e^(−1.14x)
8
6.62
5.25
3.88
2.5
present
reduced
absent
log(Bit Score)
0.00
0.25
0.50
0.75
1.00
Distance from M. lignano
Bristle Status
present
reduced
absent

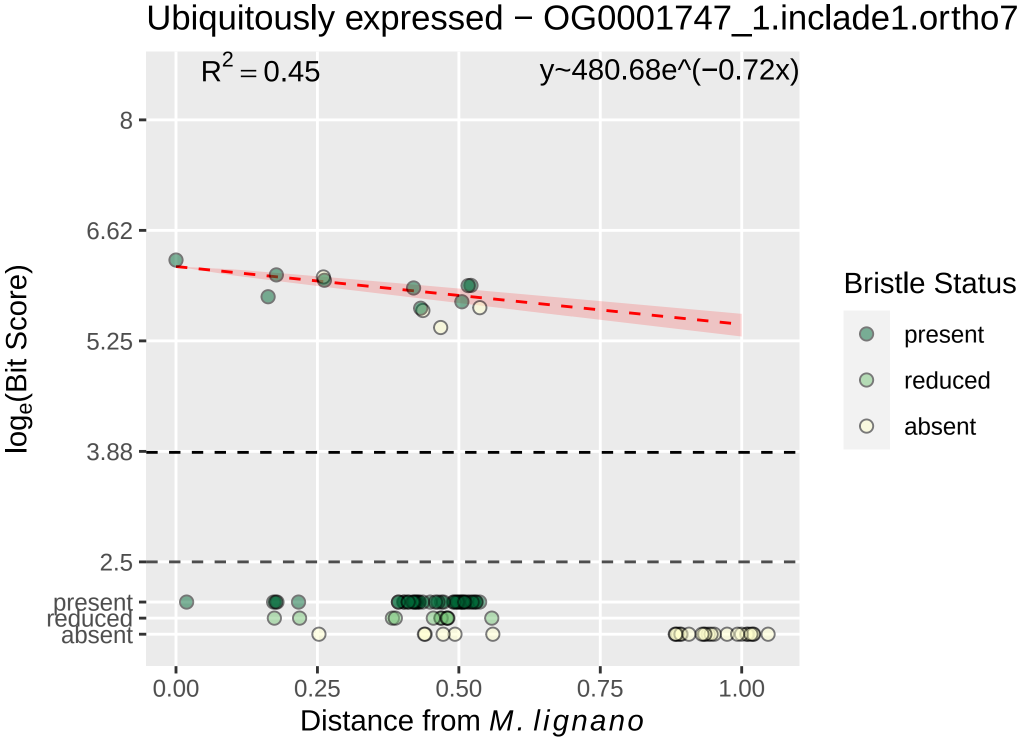

Ubiquitously expressed − OG0001747_1.inclade1.ortho7
$R^2 = 0.45$
y~480.68e^(−0.72x)
8
6.62
5.25
3.88
2.5
present
reduced
absent
$\log_e$(Bit Score)
0.00
0.25
0.50
0.75
1.00
Distance from *M. lignano*
Bristle Status
present
reduced
absent

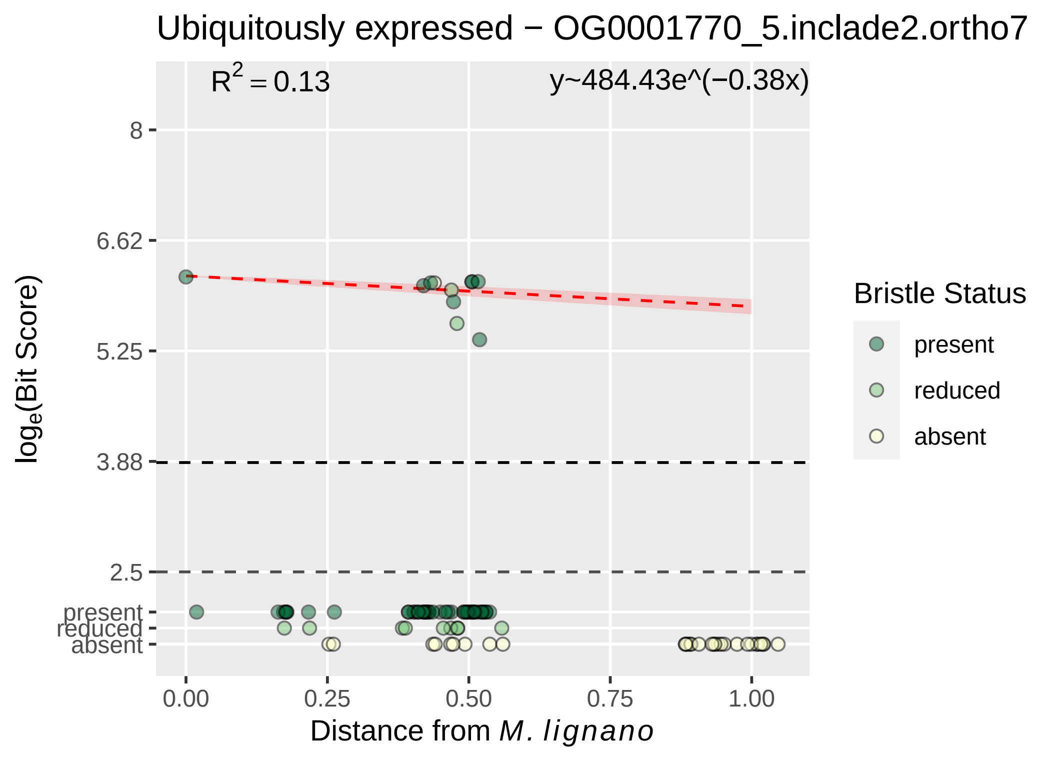
Ubiquitously expressed − OG0001770_5.inclade2.ortho7
R^2 = 0.13
y~484.43e^(−0.38x)
8
6.62
5.25
3.88
2.5
present
reduced
absent
0.00
0.25
0.50
0.75
1.00
Distance from M. lignano
log_e(Bit Score)
Bristle Status
present
reduced
absent

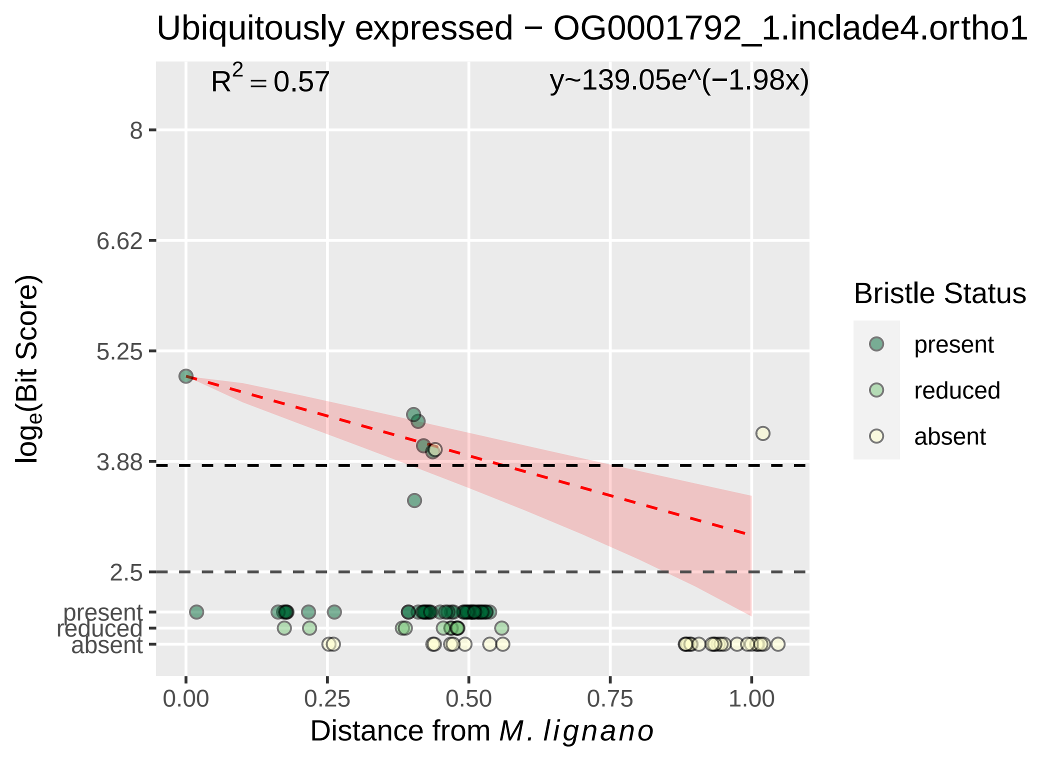

Ubiquitously expressed − OG0001792_1.inclade4.ortho1
$R^2 = 0.57$
y~139.05e^(−1.98x)
8
6.62
5.25
3.88
2.5
present
reduced
absent
$\log_e$(Bit Score)
0.00
0.25
0.50
0.75
1.00
Distance from *M. lignano*
Bristle Status
present
reduced
absent

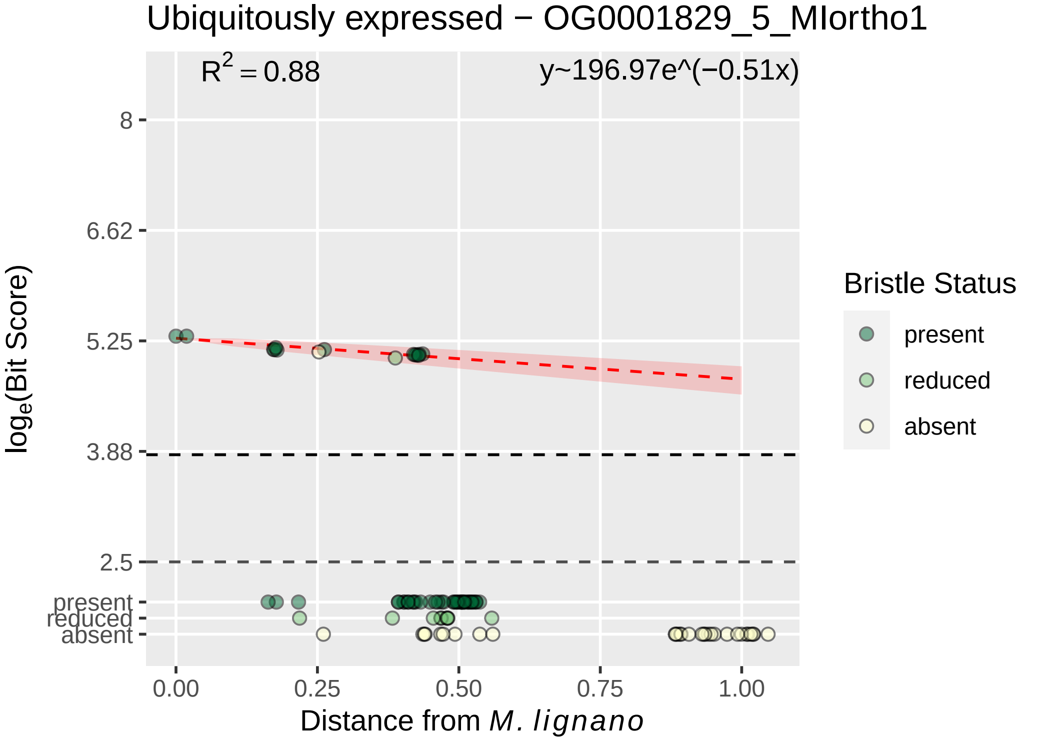

Ubiquitously expressed − OG0001829_5_MIortho1
$R^2 = 0.88$
y~196.97e^(−0.51x)
8
6.62
5.25
3.88
2.5
present
reduced
absent
$\log_e$(Bit Score)
0.00
0.25
0.50
0.75
1.00
Distance from *M. lignano*
Bristle Status
present
reduced
absent

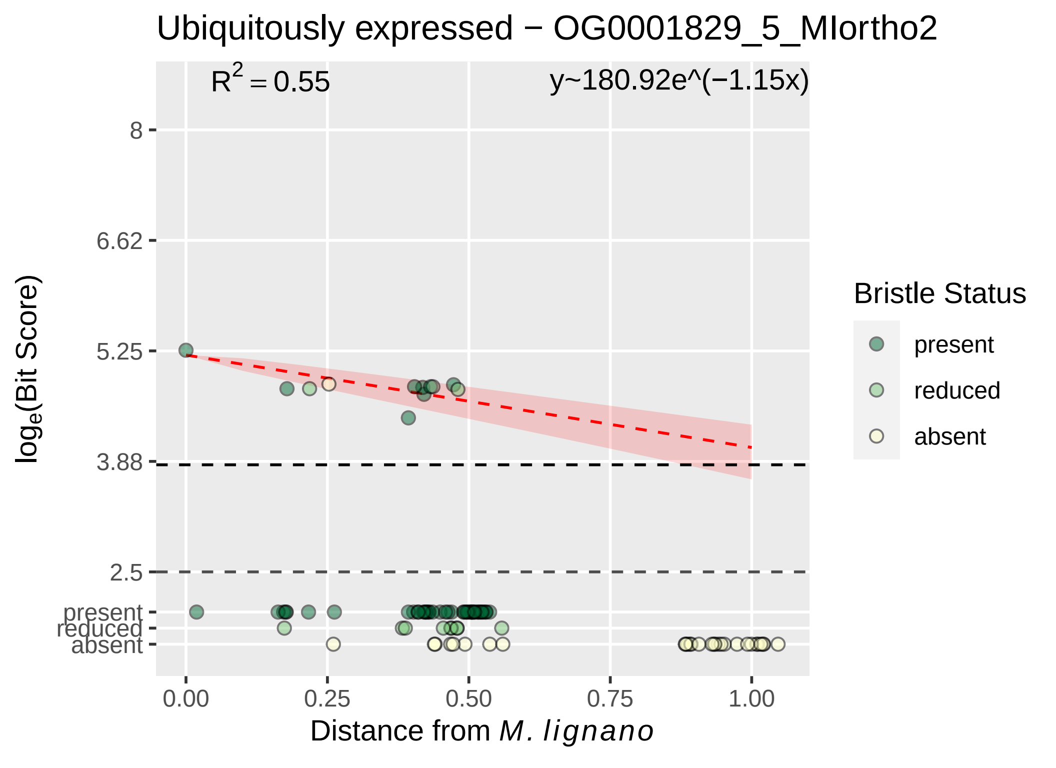

Ubiquitously expressed − OG0001829_5_MIortho2
R2 = 0.55
y~180.92e^(−1.15x)
8
6.62
5.25
3.88
2.5
present
reduced
absent
loge(Bit Score)
0.00
0.25
0.50
0.75
1.00
Distance from M. lignano
Bristle Status
present
reduced
absent

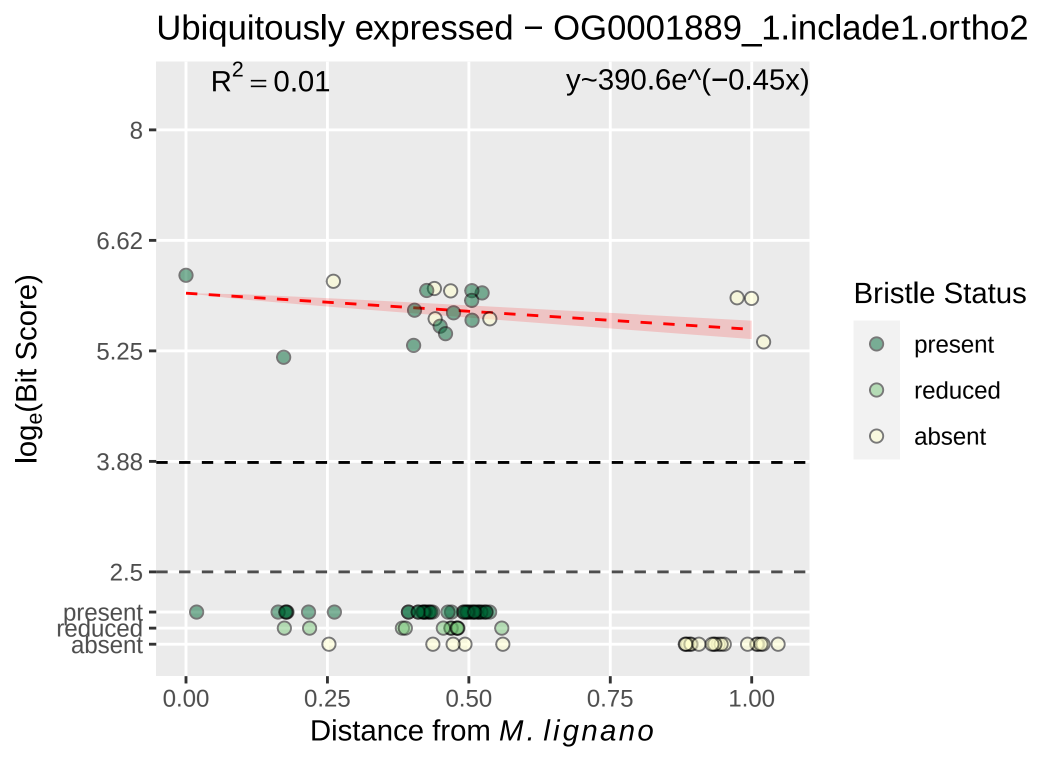

Ubiquitously expressed − OG0001889_1.inclade1.ortho2
$R^2 = 0.01$
y~390.6e^(−0.45x)
8
6.62
5.25
3.88
2.5
present
reduced
absent
$\log_e$(Bit Score)
0.00
0.25
0.50
0.75
1.00
Distance from *M. lignano*
Bristle Status
present
reduced
absent

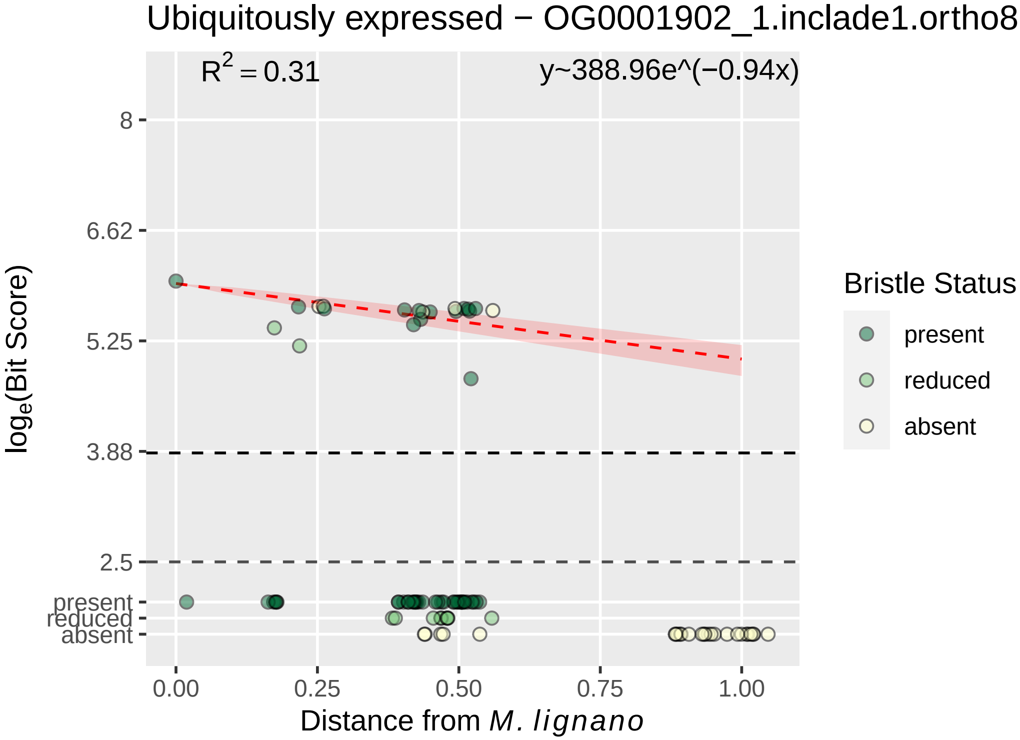
Ubiquitously expressed − OG0001902_1.inclade1.ortho8
R2 = 0.31
y~388.96e^(−0.94x)
8
6.62
5.25
3.88
2.5
present
reduced
absent
loge(Bit Score)
0.00
0.25
0.50
0.75
1.00
Distance from M. lignano
Bristle Status
present
reduced
absent

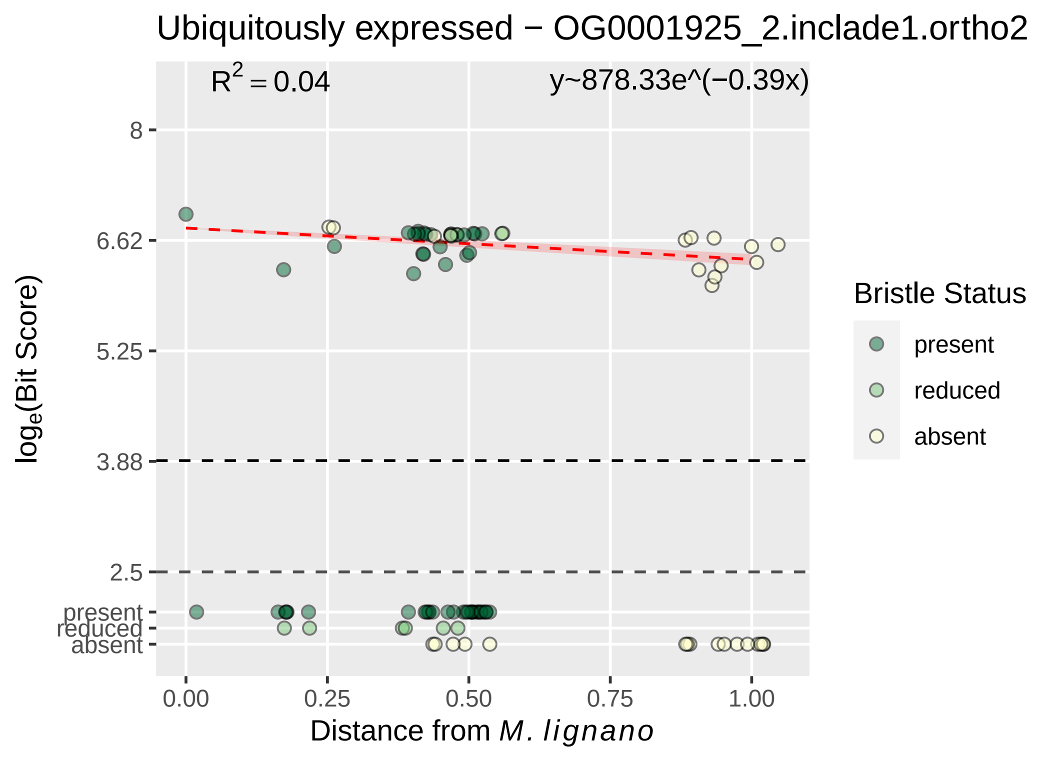

Ubiquitously expressed − OG0001925_2.inclade1.ortho2
$R^2 = 0.04$
y~878.33e^(−0.39x)
8
6.62
5.25
3.88
2.5
present
reduced
absent
log$_e$(Bit Score)
0.00
0.25
0.50
0.75
1.00
Distance from *M. lignano*
Bristle Status
present
reduced
absent

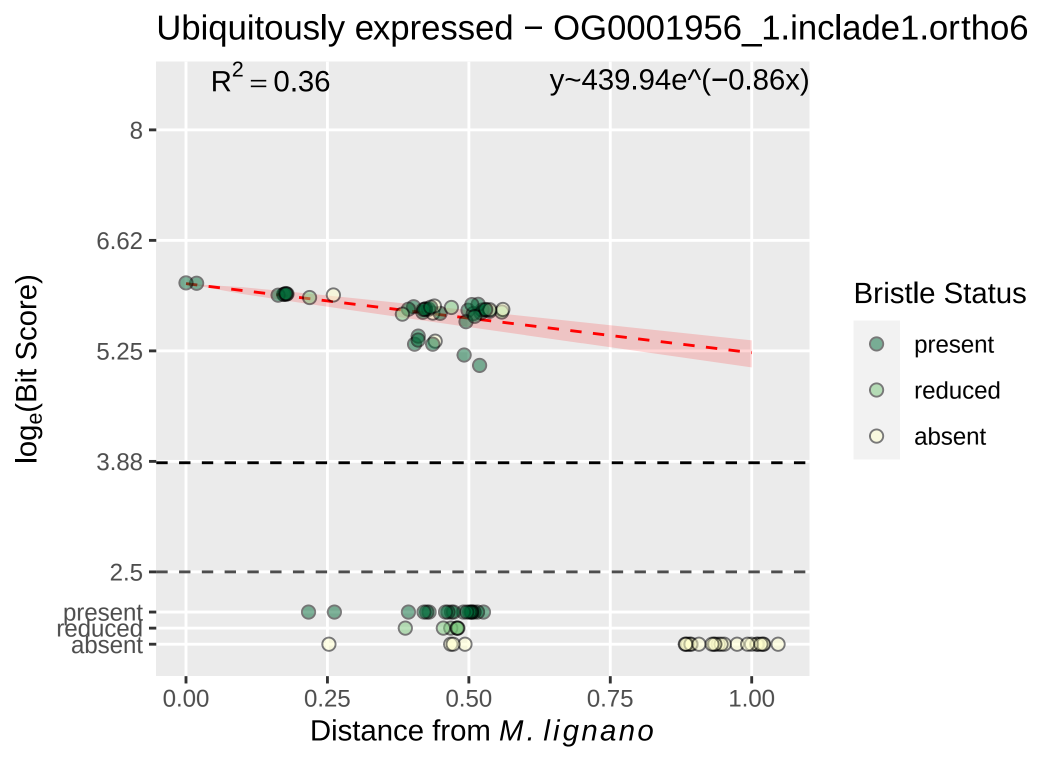

Ubiquitously expressed − OG0001956_1.inclade1.ortho6
$R^2 = 0.36$
y~439.94e^(−0.86x)
8
6.62
5.25
3.88
2.5
present
reduced
absent
$\log_e$(Bit Score)
0.00
0.25
0.50
0.75
1.00
Distance from *M. lignano*
Bristle Status
present
reduced
absent

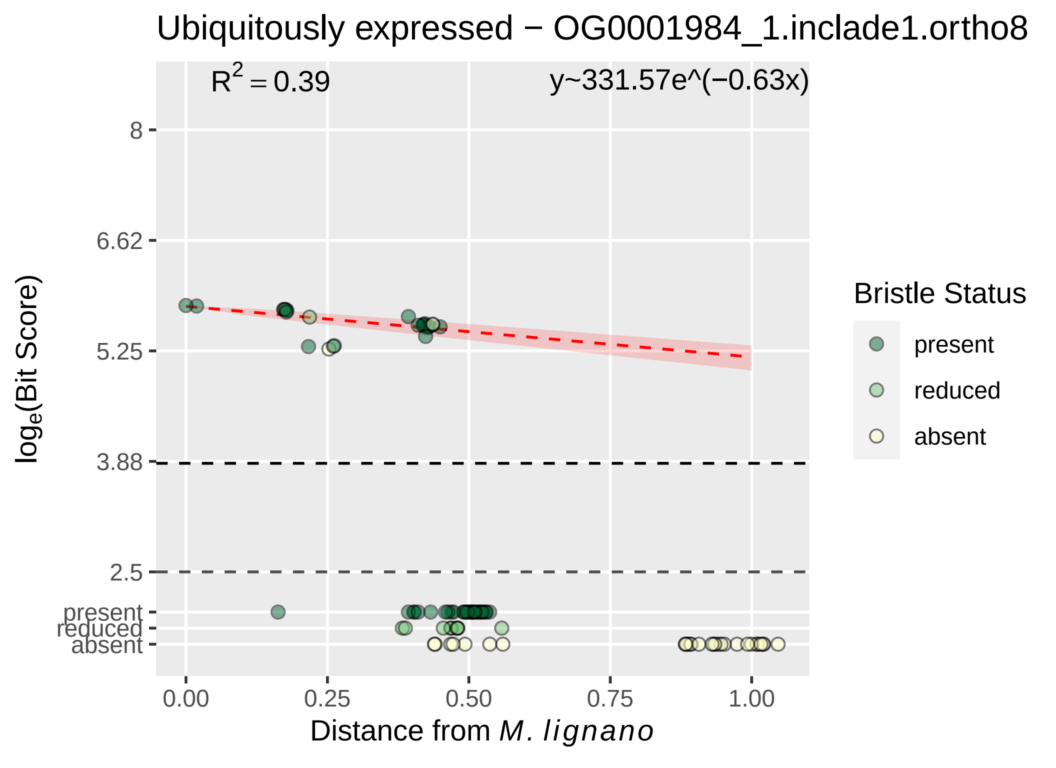

Ubiquitously expressed − OG0001984_1.inclade1.ortho8
R² = 0.39
y~331.57e^(−0.63x)
log_e(Bit Score)
8
6.62
5.25
3.88
2.5
present
reduced
absent
0.00
0.25
0.50
0.75
1.00
Distance from M. lignano
Bristle Status
present
reduced
absent

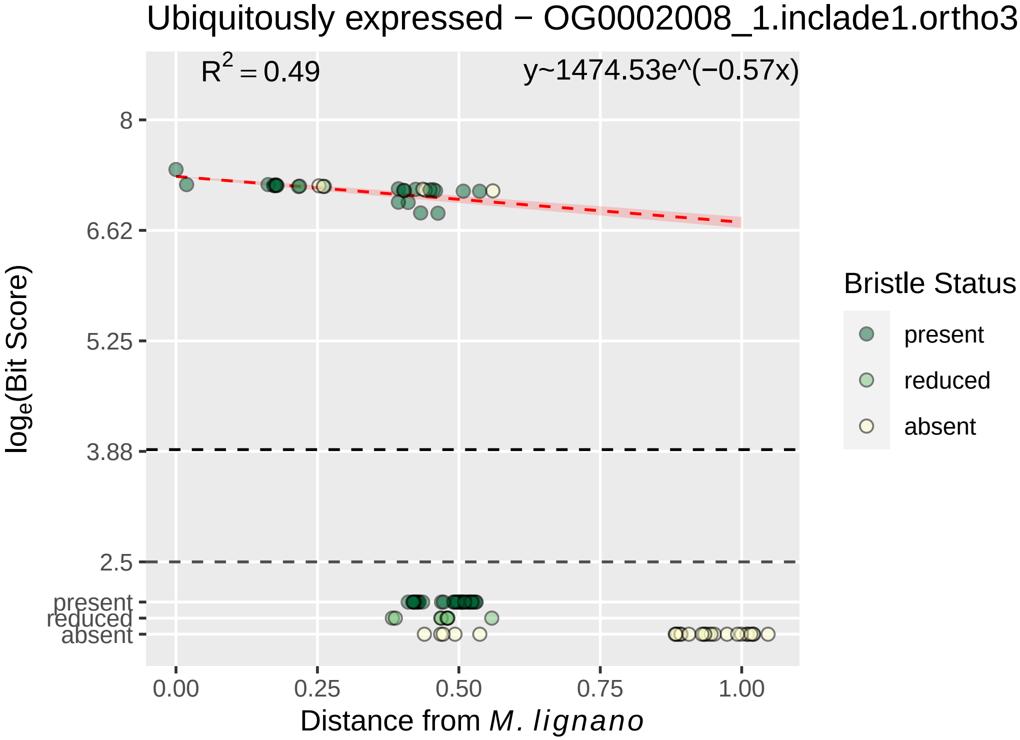
Ubiquitously expressed − OG0002008_1.inclade1.ortho3
R2 = 0.49
y~1474.53e^(−0.57x)
loge(Bit Score)
8
6.62
5.25
3.88
2.5
present
reduced
absent
0.00
0.25
0.50
0.75
1.00
Distance from M. lignano
Bristle Status
present
reduced
absent

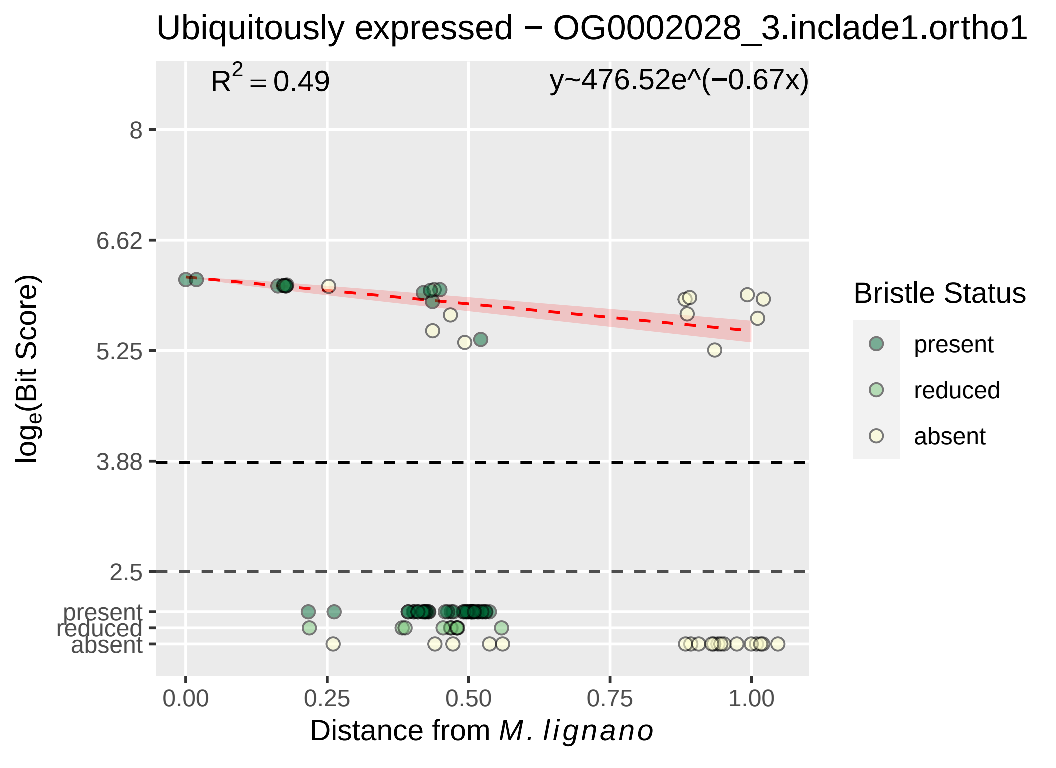

Ubiquitously expressed − OG0002028_3.inclade1.ortho1
R^2 = 0.49
y~476.52e^(−0.67x)
8
6.62
5.25
3.88
2.5
present
reduced
absent
log_e(Bit Score)
0.00
0.25
0.50
0.75
1.00
Distance from M. lignano
Bristle Status
present
reduced
absent

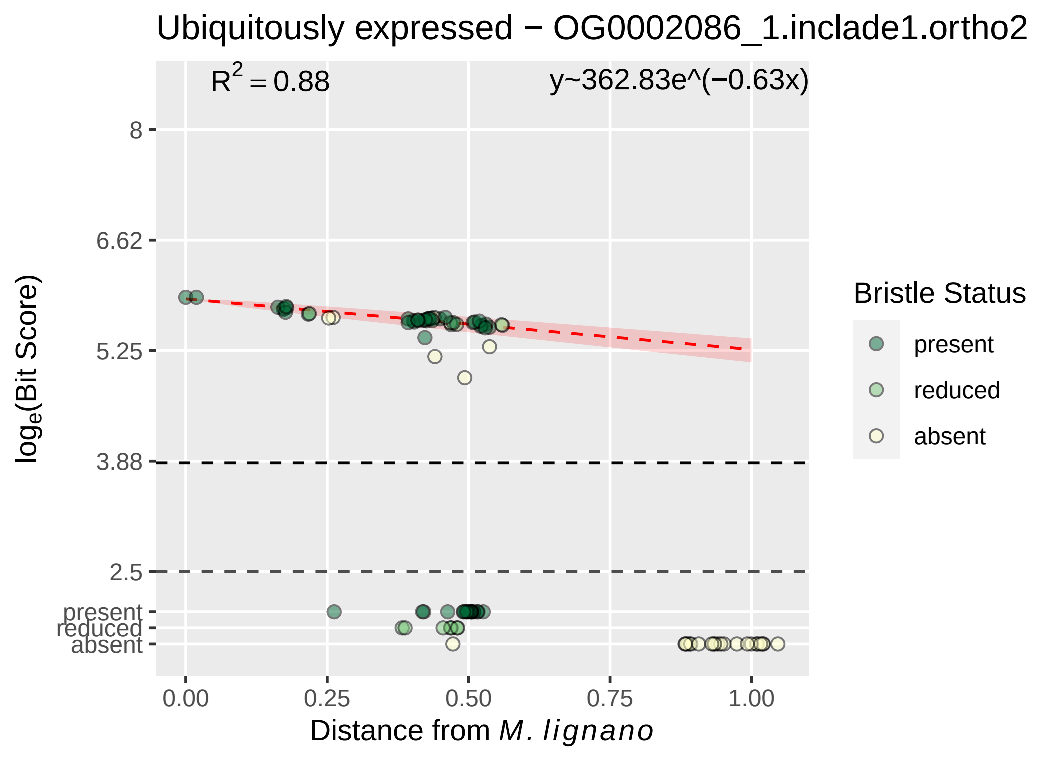

Ubiquitously expressed − OG0002086_1.inclade1.ortho2
$R^2 = 0.88$
y~362.83e^(−0.63x)
8
6.62
5.25
3.88
2.5
present
reduced
absent
$log_e$(Bit Score)
0.00
0.25
0.50
0.75
1.00
Distance from *M. lignano*
Bristle Status
present
reduced
absent

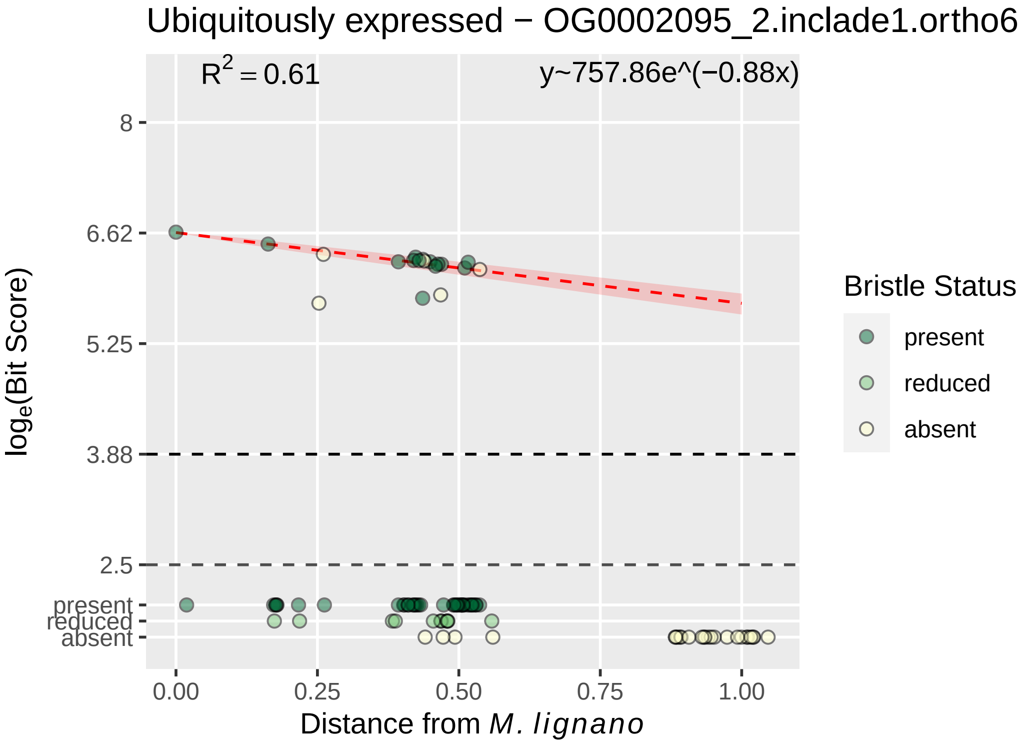

Ubiquitously expressed − OG0002095_2.inclade1.ortho6
$R^2 = 0.61$
y~757.86e^(−0.88x)
8
6.62
5.25
3.88
2.5
present
reduced
absent
$\log_e$(Bit Score)
0.00
0.25
0.50
0.75
1.00
Distance from *M. lignano*
Bristle Status
present
reduced
absent

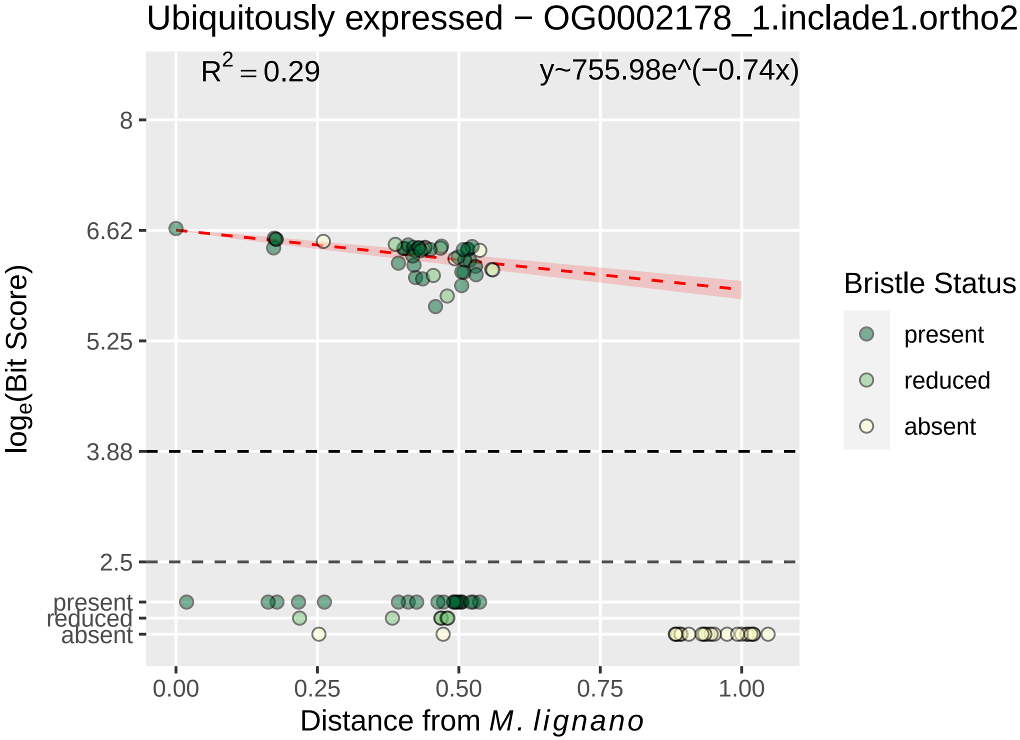

Ubiquitously expressed − OG0002178_1.inclade1.ortho2
$R^2 = 0.29$
y~755.98e^(−0.74x)
8
6.62
5.25
3.88
2.5
present
reduced
absent
0.00
0.25
0.50
0.75
1.00
Distance from M. lignano
$log_e$(Bit Score)
Bristle Status
present
reduced
absent

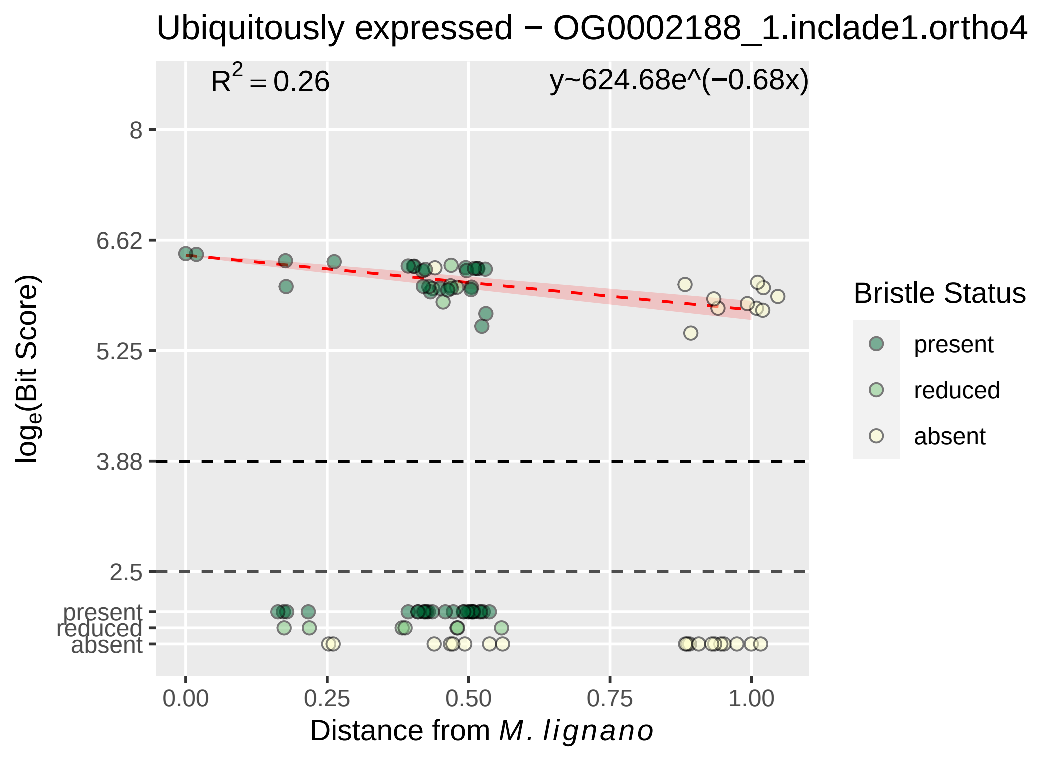

Ubiquitously expressed − OG0002188_1.inclade1.ortho4
R² = 0.26
y~624.68e^(−0.68x)
8
6.62
5.25
3.88
2.5
present
reduced
absent
0.00
0.25
0.50
0.75
1.00
Distance from M. lignano
log_e(Bit Score)
Bristle Status
present
reduced
absent

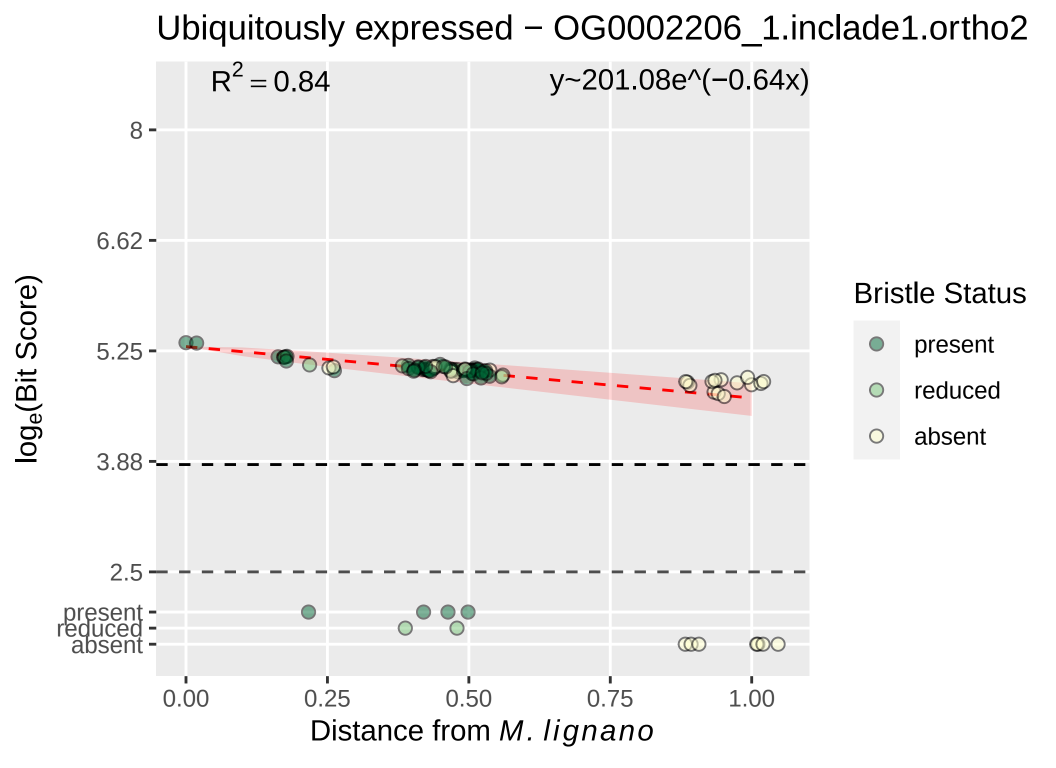

Ubiquitously expressed − OG0002206_1.inclade1.ortho2
R2 = 0.84
y~201.08e^(−0.64x)
8
6.62
5.25
3.88
2.5
present
reduced
absent
loge(Bit Score)
0.00
0.25
0.50
0.75
1.00
Distance from M. lignano
Bristle Status
present
reduced
absent

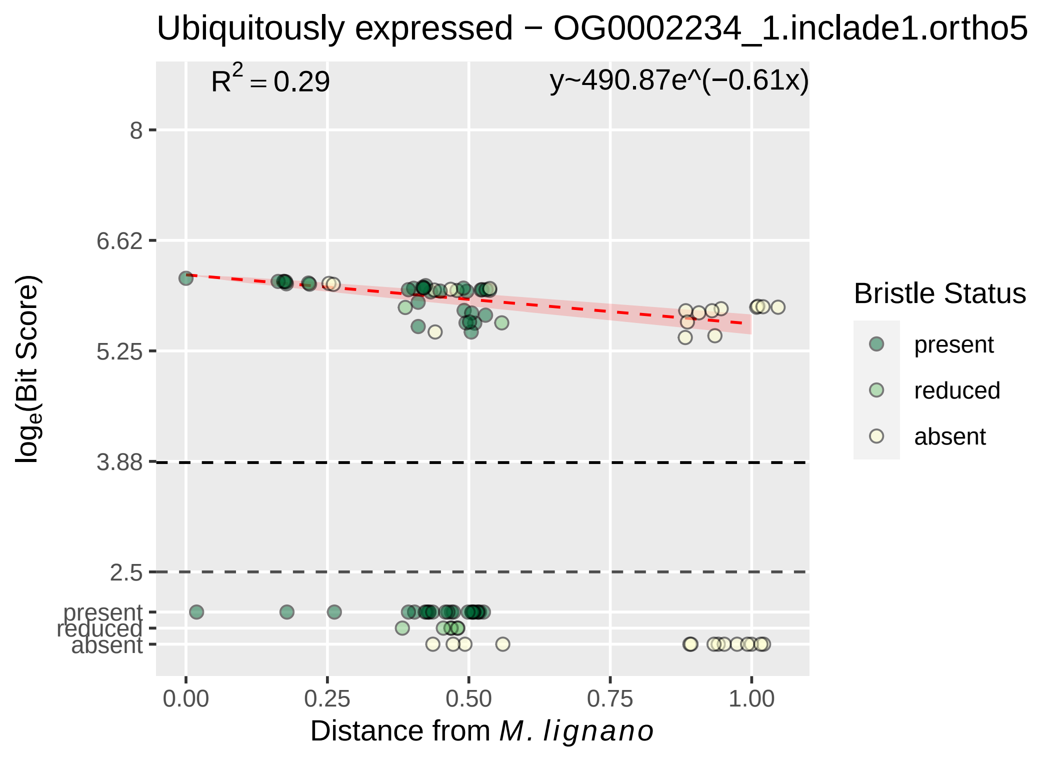

Ubiquitously expressed − OG0002234_1.inclade1.ortho5
R² = 0.29
y~490.87e^(−0.61x)
8
6.62
5.25
3.88
2.5
present
reduced
absent
log_e(Bit Score)
0.00
0.25
0.50
0.75
1.00
Distance from M. lignano
Bristle Status
present
reduced
absent

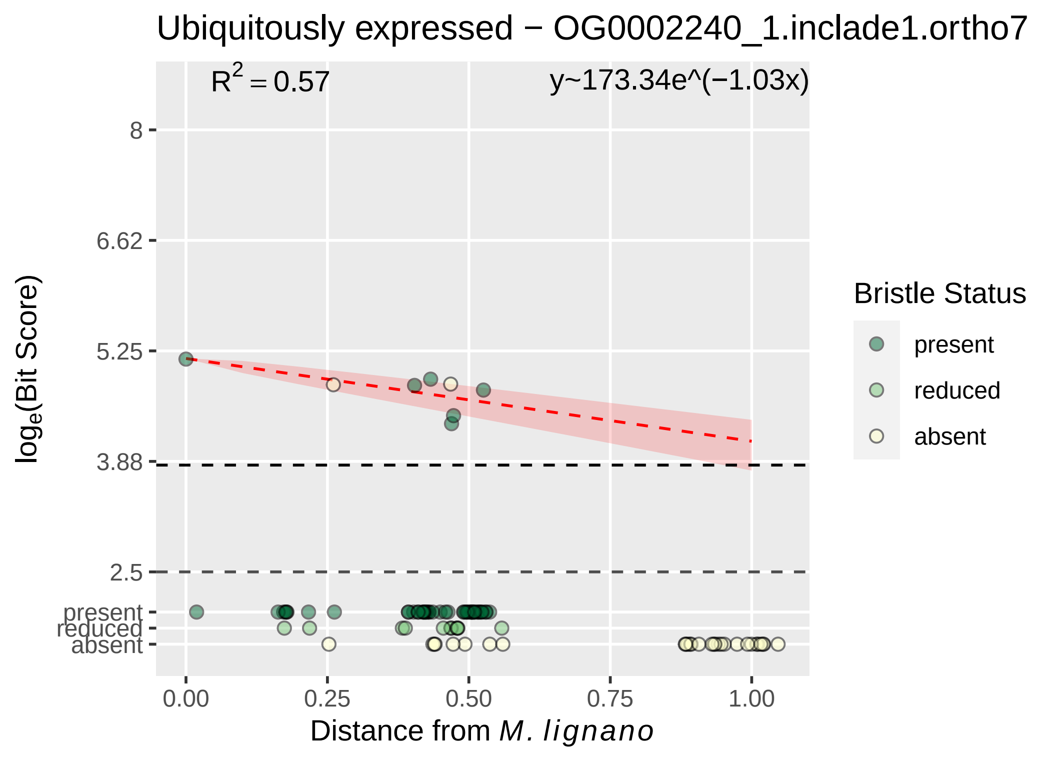

Ubiquitously expressed − OG0002240_1.inclade1.ortho7
$R^2 = 0.57$
y~173.34e^(−1.03x)
8
6.62
5.25
3.88
2.5
present
reduced
absent
0.00
0.25
0.50
0.75
1.00
log$_e$(Bit Score)
Distance from *M. lignano*
Bristle Status
present
reduced
absent

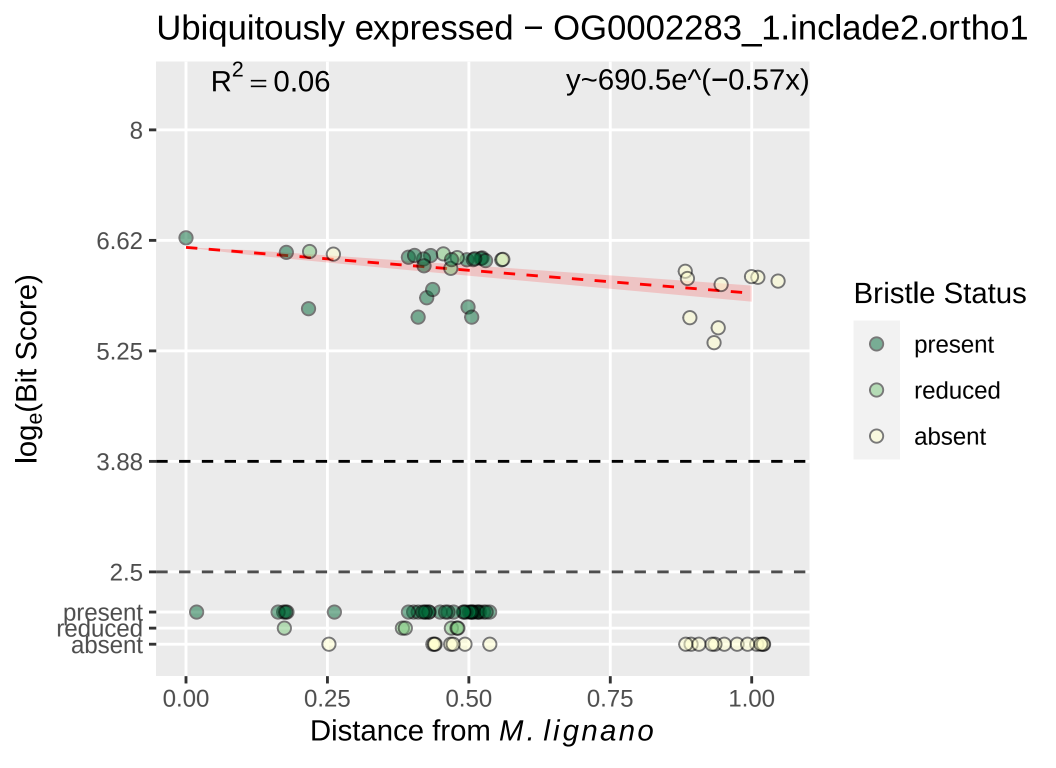

Ubiquitously expressed − OG0002283_1.inclade2.ortho1
R2 = 0.06
y~690.5e^(−0.57x)
8
6.62
5.25
3.88
2.5
present
reduced
absent
loge(Bit Score)
0.00
0.25
0.50
0.75
1.00
Distance from M. lignano
Bristle Status
present
reduced
absent

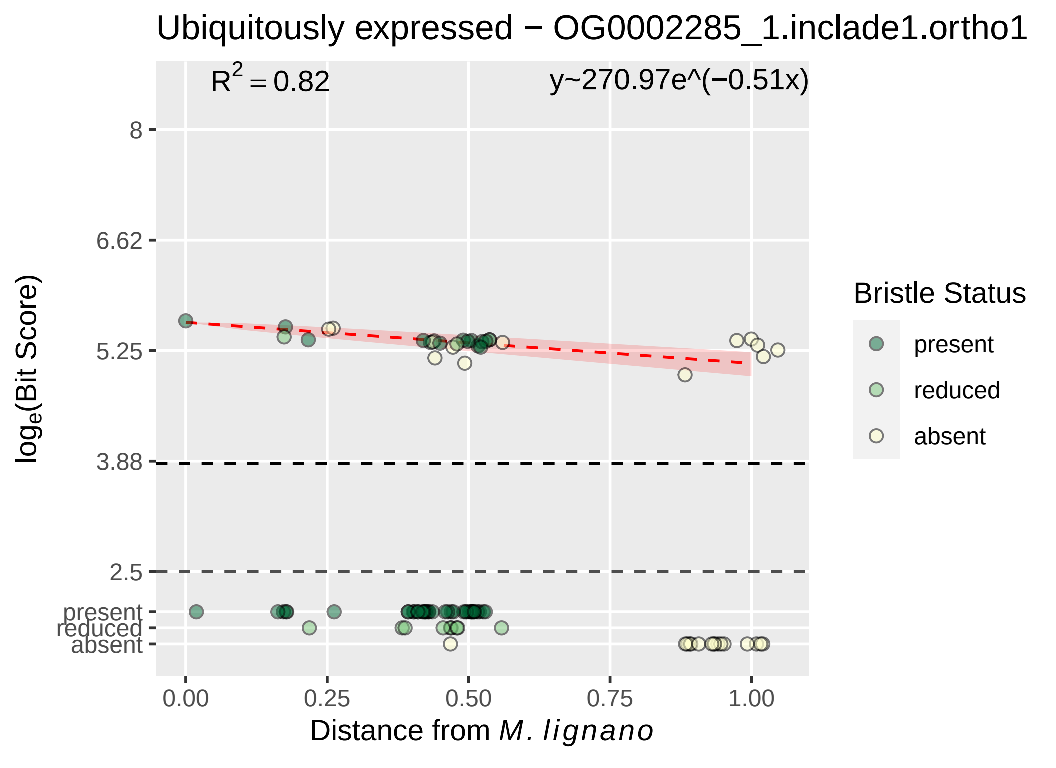
Ubiquitously expressed − OG0002285_1.inclade1.ortho1
$R^2 = 0.82$
y~270.97e^(−0.51x)
8
6.62
5.25
3.88
2.5
present
reduced
absent
0.00
0.25
0.50
0.75
1.00
$log_e$(Bit Score)
Distance from M. lignano
Bristle Status
present
reduced
absent

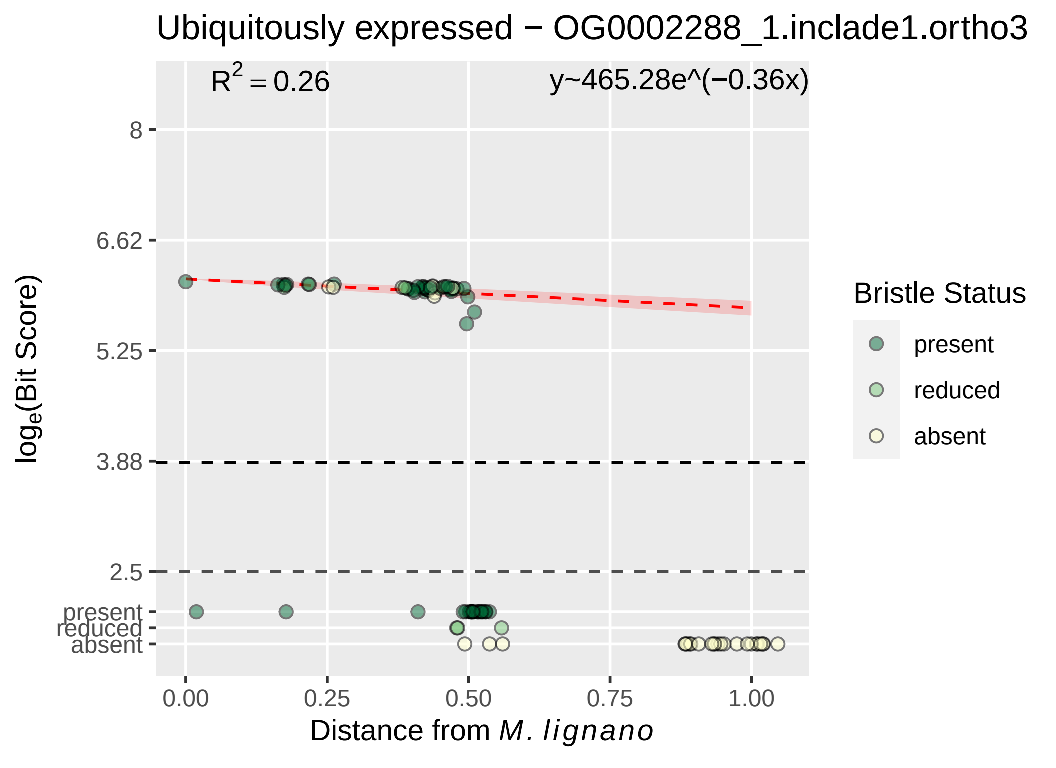
Ubiquitously expressed − OG0002288_1.inclade1.ortho3
R² = 0.26
y~465.28e^(−0.36x)
8
6.62
5.25
3.88
2.5
present
reduced
absent
log_e(Bit Score)
0.00
0.25
0.50
0.75
1.00
Distance from M. lignano
Bristle Status
present
reduced
absent

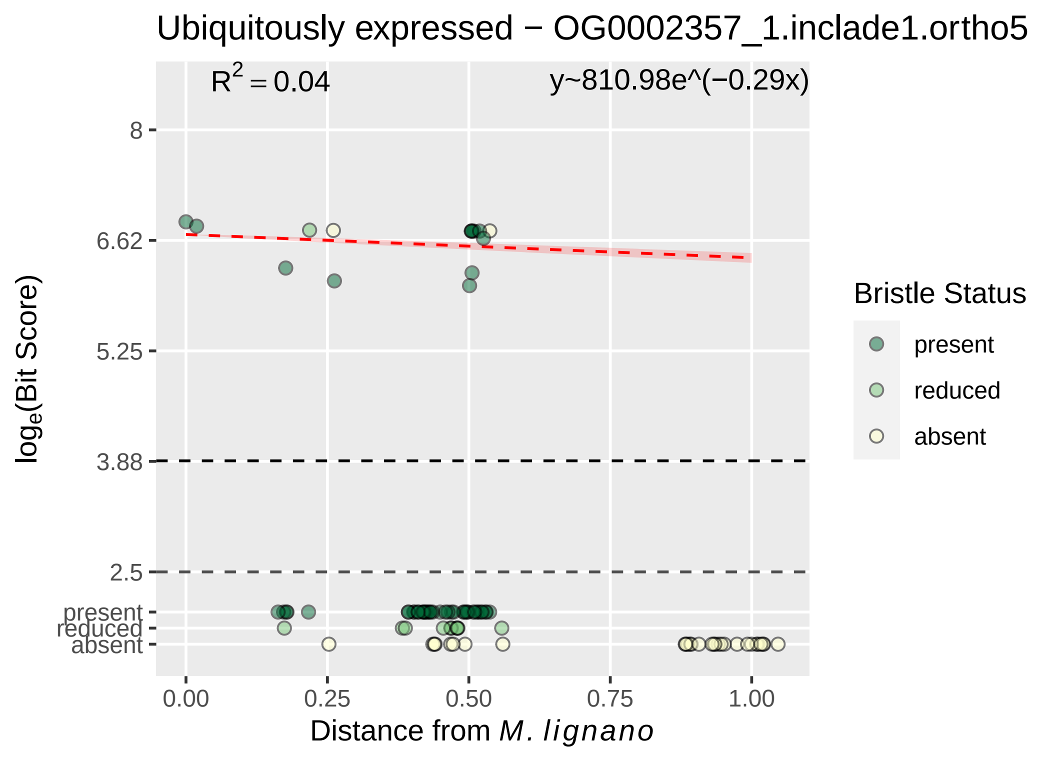

Ubiquitously expressed − OG0002357_1.inclade1.ortho5
R^2 = 0.04
y~810.98e^(−0.29x)
8
6.62
5.25
3.88
2.5
present
reduced
absent
log_e(Bit Score)
0.00
0.25
0.50
0.75
1.00
Distance from M. lignano
Bristle Status
present
reduced
absent

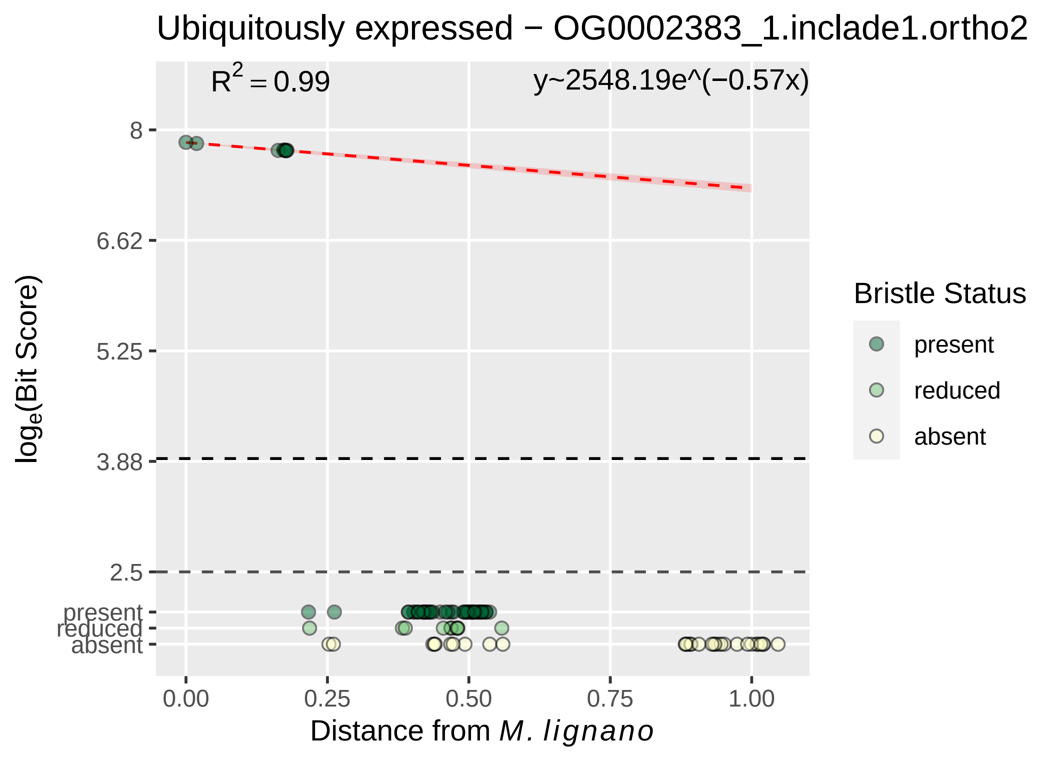

Ubiquitously expressed − OG0002383_1.inclade1.ortho2
R² = 0.99
y~2548.19e^(−0.57x)
log_e(Bit Score)
8
6.62
5.25
3.88
2.5
present
reduced
absent
0.00
0.25
0.50
0.75
1.00
Distance from M. lignano
Bristle Status
present
reduced
absent

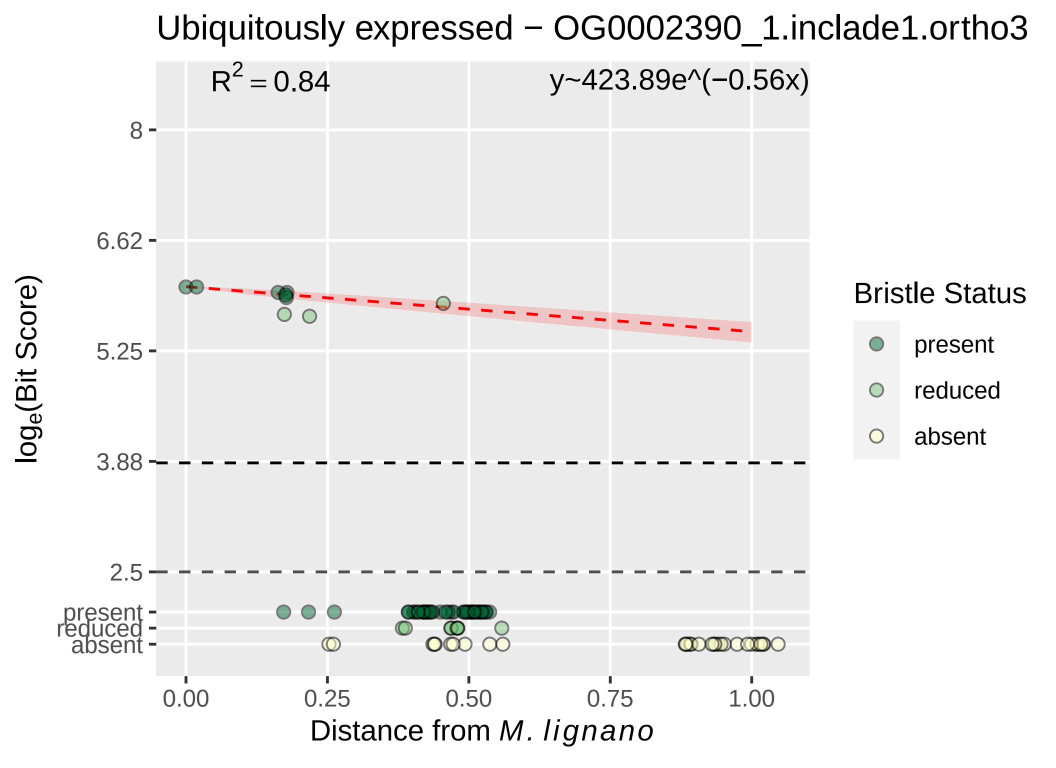

Ubiquitously expressed − OG0002390_1.inclade1.ortho3
$R^2 = 0.84$
y~423.89e^(−0.56x)
8
6.62
5.25
3.88
2.5
present
reduced
absent
0.00
0.25
0.50
0.75
1.00
$\log_e$(Bit Score)
Distance from *M. lignano*
Bristle Status
present
reduced
absent

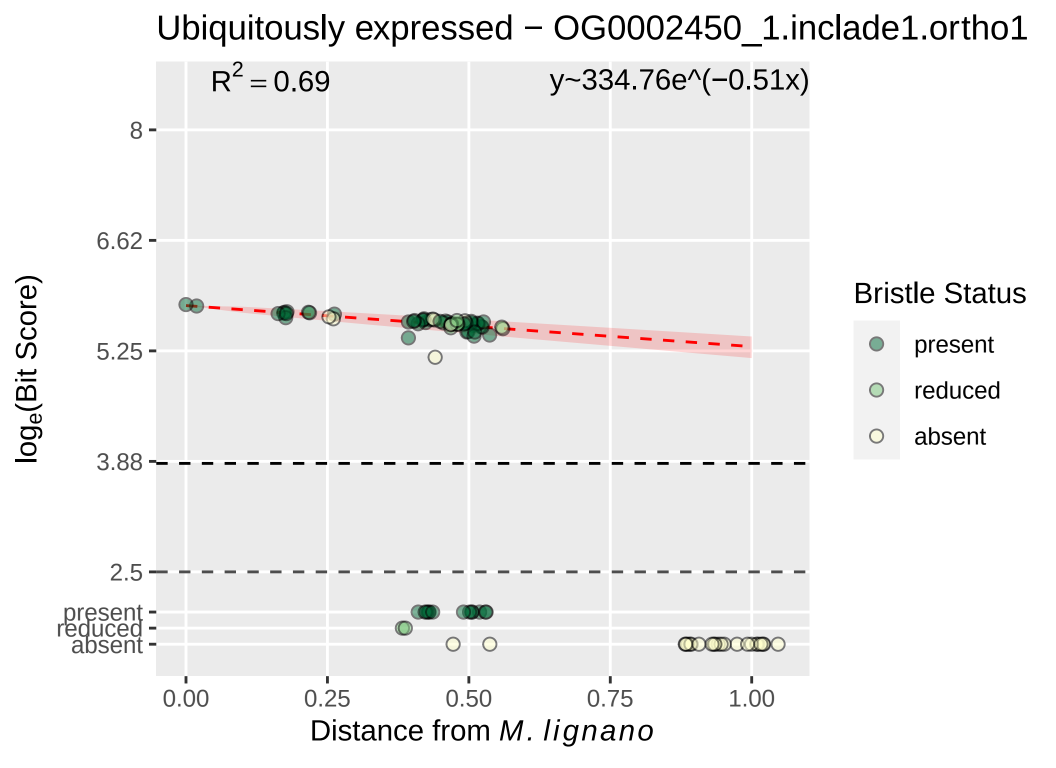

Ubiquitously expressed − OG0002450_1.inclade1.ortho1
R2 = 0.69
y~334.76e^(−0.51x)
8
6.62
5.25
3.88
2.5
present
reduced
absent
loge(Bit Score)
0.00
0.25
0.50
0.75
1.00
Distance from M. lignano
Bristle Status
present
reduced
absent

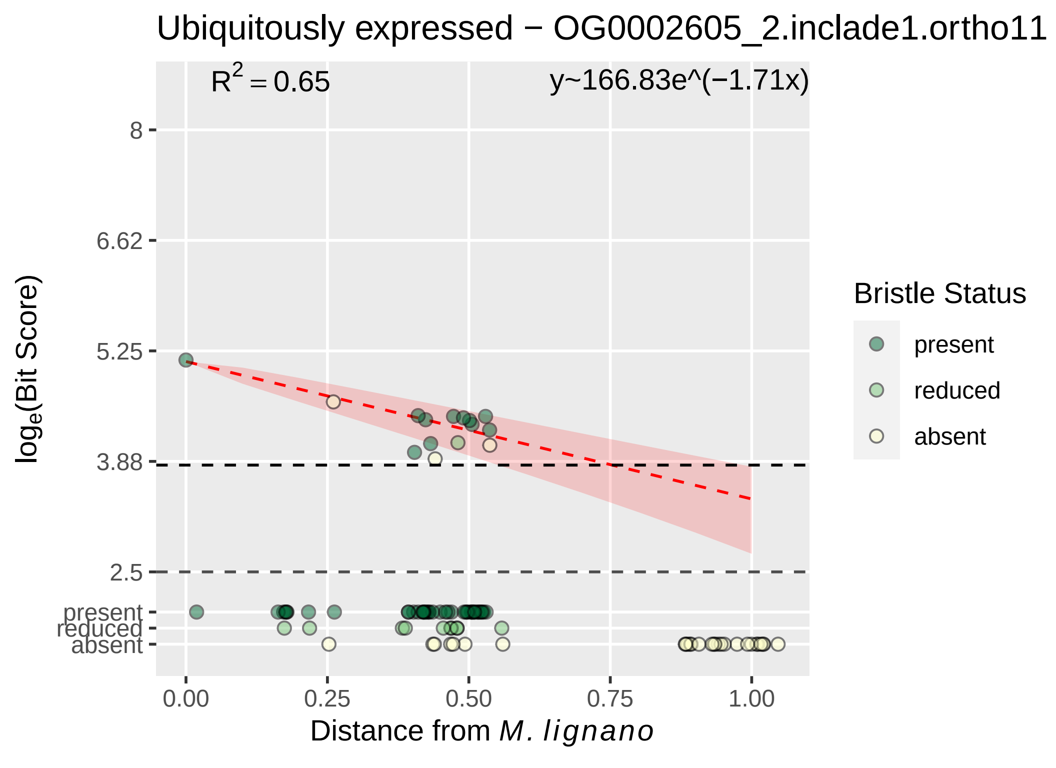

Ubiquitously expressed − OG0002605_2.inclade1.ortho11
$R^2 = 0.65$
y~166.83e^(−1.71x)
8
6.62
5.25
3.88
2.5
present
reduced
absent
0.00
0.25
0.50
0.75
1.00
Distance from M. lignano
log$_e$(Bit Score)
Bristle Status
present
reduced
absent

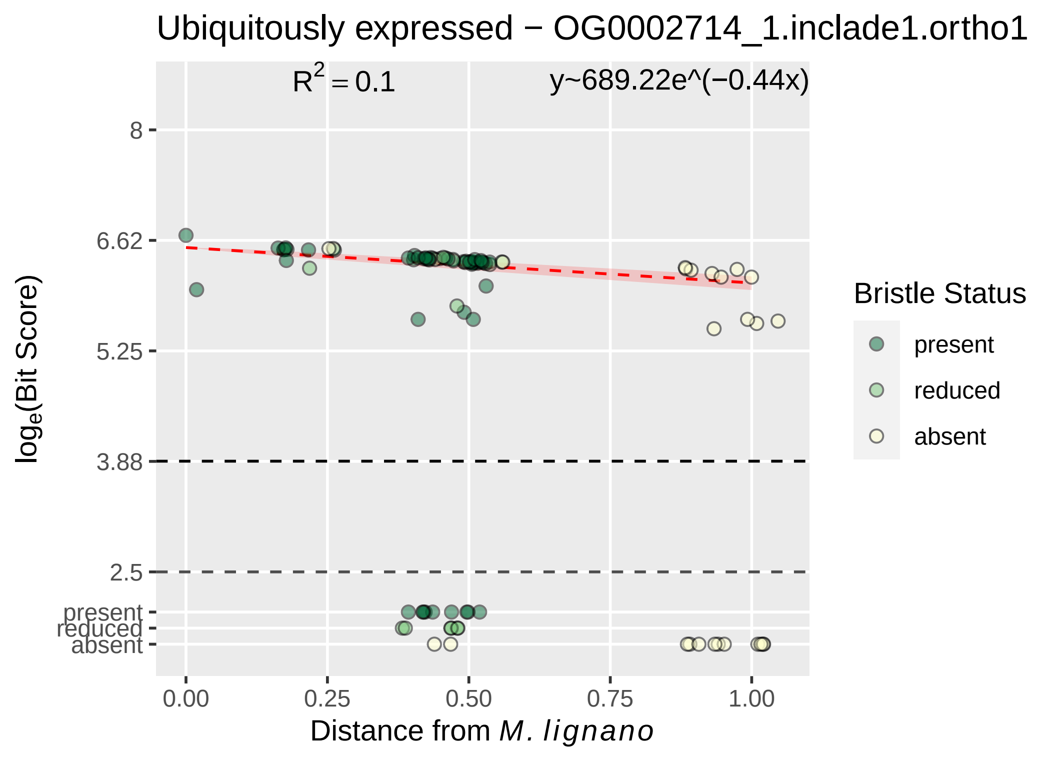

Ubiquitously expressed − OG0002714_1.inclade1.ortho1
R2 = 0.1
y~689.22e^(−0.44x)
8
6.62
5.25
3.88
2.5
present
reduced
absent
loge(Bit Score)
0.00
0.25
0.50
0.75
1.00
Distance from M. lignano
Bristle Status
present
reduced
absent

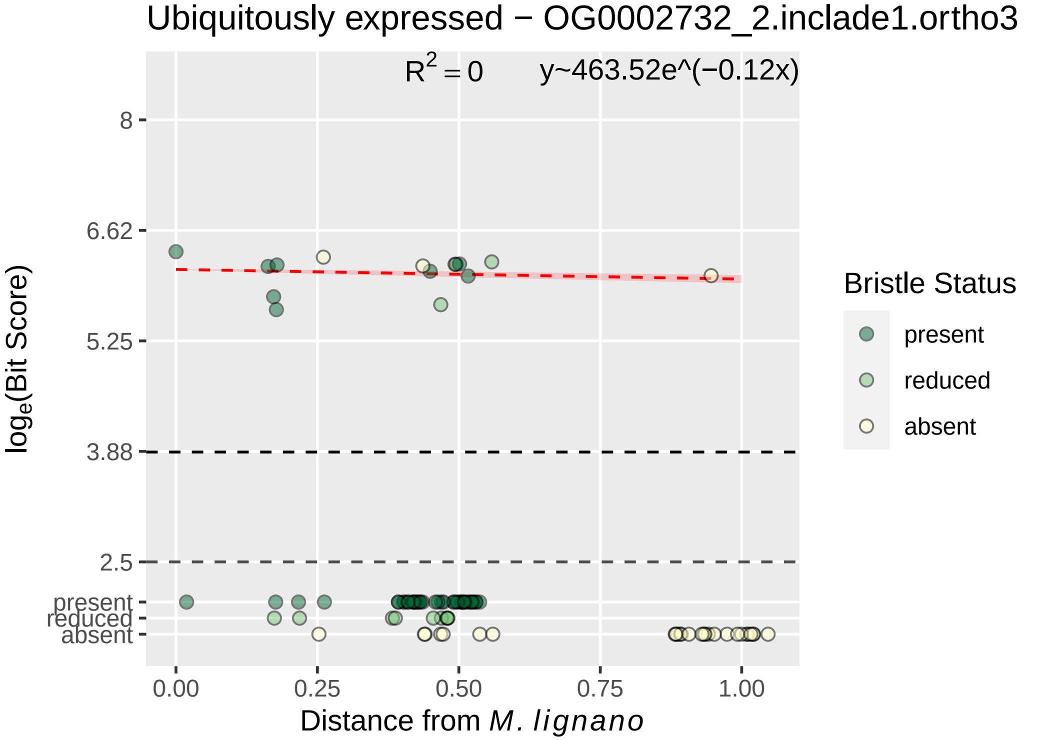
Ubiquitously expressed − OG0002732_2.inclade1.ortho3
R^2 = 0
y~463.52e^(−0.12x)
8
6.62
5.25
3.88
2.5
present
reduced
absent
0.00
0.25
0.50
0.75
1.00
log_e(Bit Score)
Distance from M. lignano
Bristle Status
present
reduced
absent

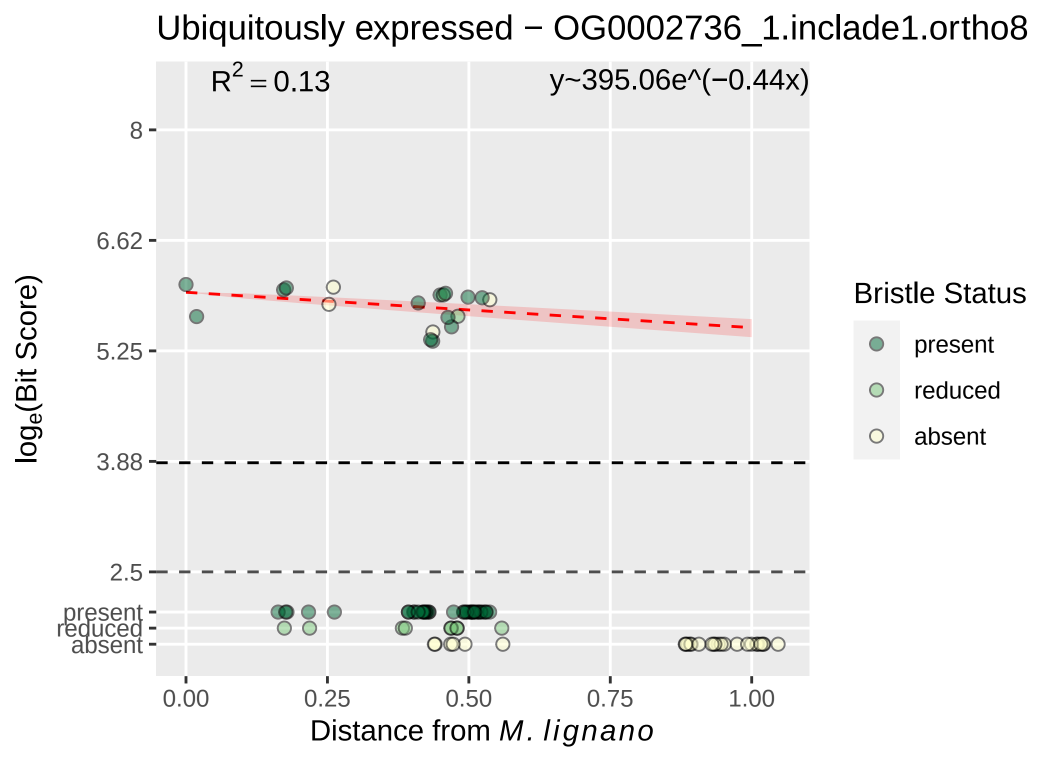

Ubiquitously expressed − OG0002736_1.inclade1.ortho8
R² = 0.13
y~395.06e^(−0.44x)
8
6.62
5.25
3.88
2.5
present
reduced
absent
0.00
0.25
0.50
0.75
1.00
Distance from M. lignano
loge(Bit Score)
Bristle Status
present
reduced
absent

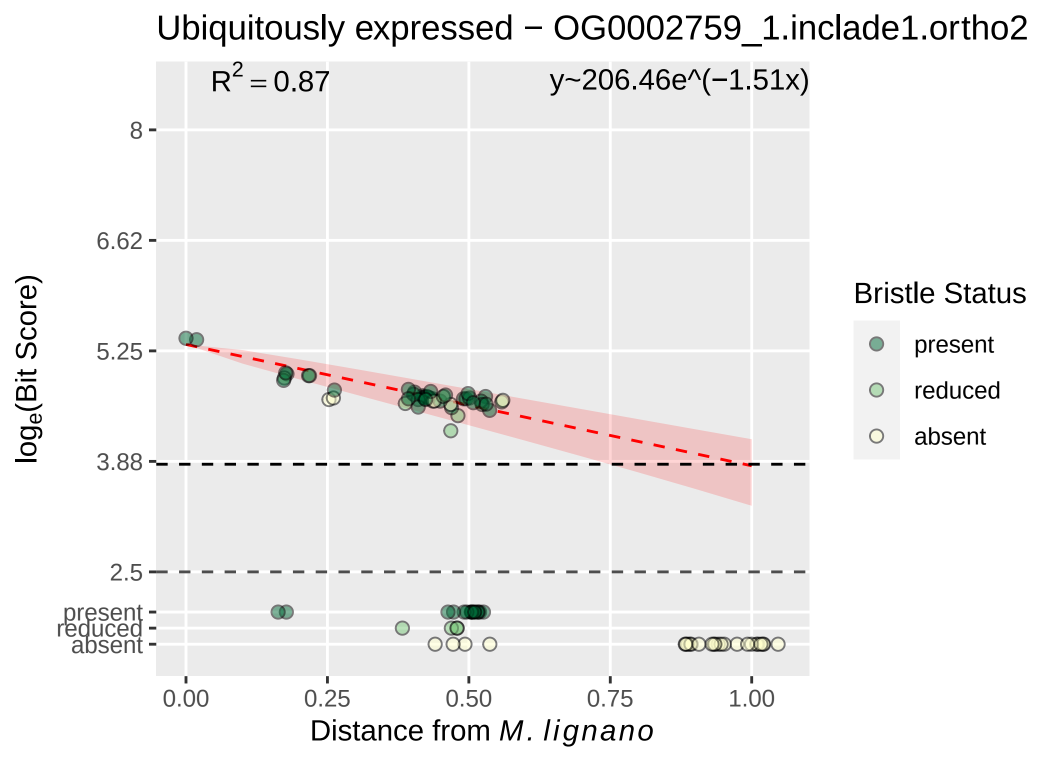
Ubiquitously expressed − OG0002759_1.inclade1.ortho2
R^2 = 0.87
y~206.46e^(−1.51x)
8
6.62
5.25
3.88
2.5
present
reduced
absent
log_e(Bit Score)
0.00
0.25
0.50
0.75
1.00
Distance from M. lignano
Bristle Status
present
reduced
absent

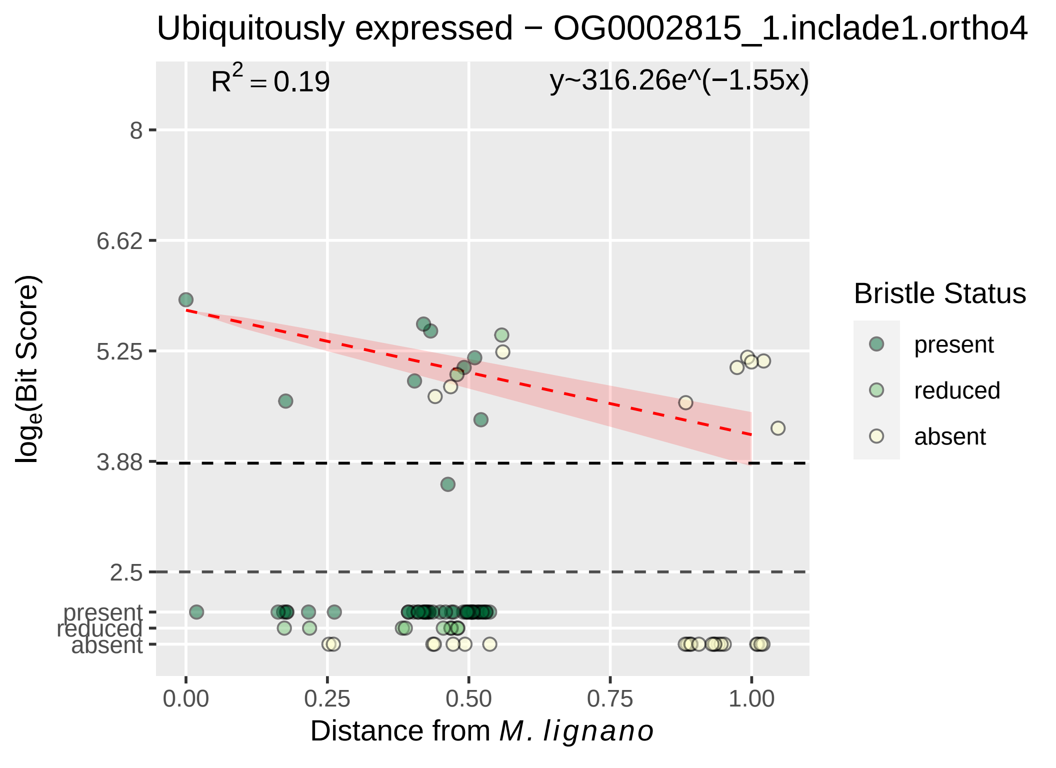
Ubiquitously expressed − OG0002815_1.inclade1.ortho4
R2 = 0.19
y~316.26e^(−1.55x)
8
6.62
5.25
3.88
2.5
present
reduced
absent
0.00
0.25
0.50
0.75
1.00
Distance from M. lignano
loge(Bit Score)
Bristle Status
present
reduced
absent

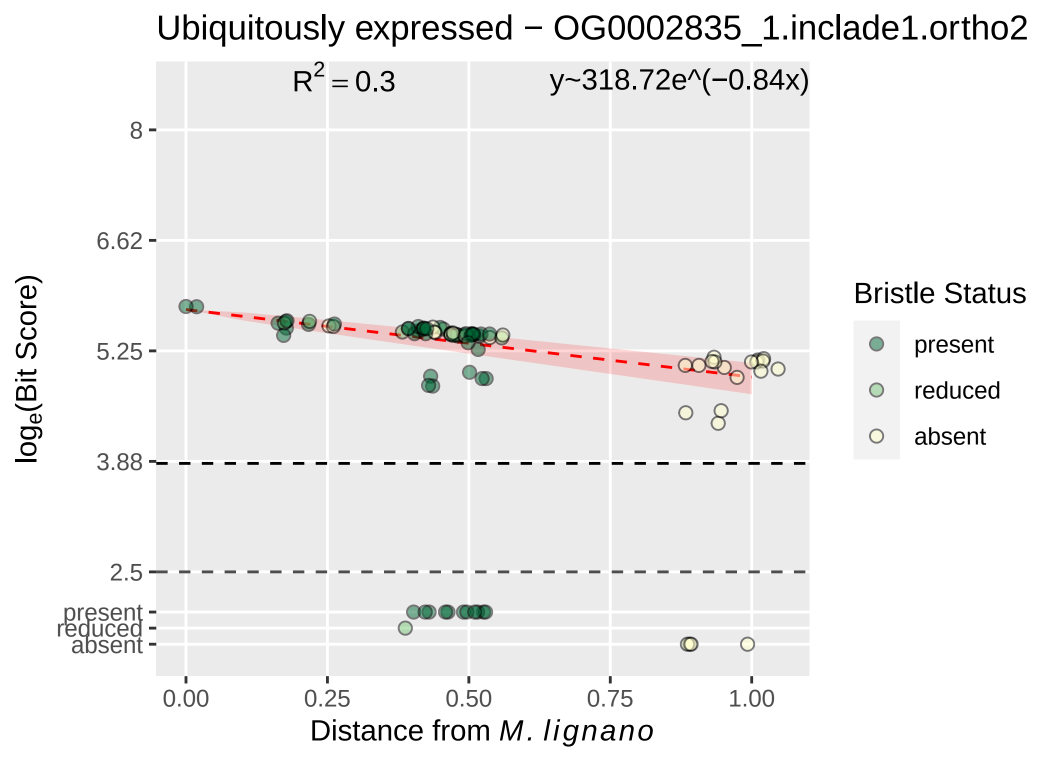
Ubiquitously expressed − OG0002835_1.inclade1.ortho2
R2 = 0.3
y~318.72e^(−0.84x)
8
6.62
5.25
3.88
2.5
present
reduced
absent
loge(Bit Score)
0.00
0.25
0.50
0.75
1.00
Distance from M. lignano
Bristle Status
present
reduced
absent

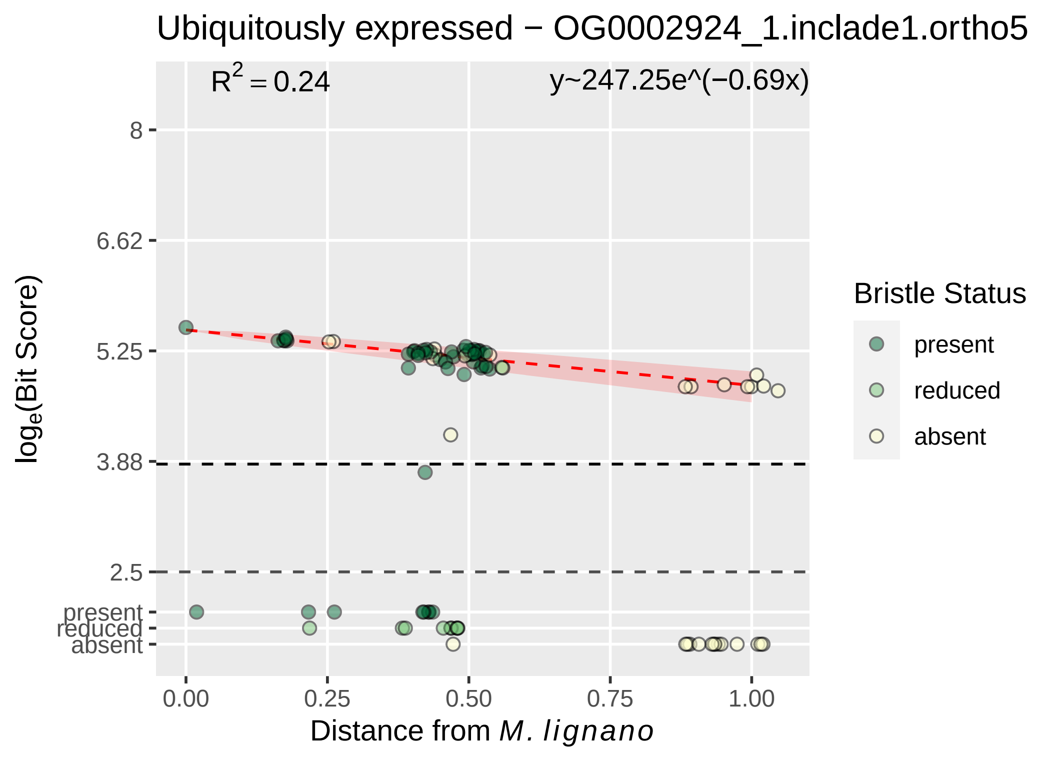

Ubiquitously expressed − OG0002924_1.inclade1.ortho5
R^2 = 0.24
y~247.25e^(−0.69x)
8
6.62
5.25
3.88
2.5
present
reduced
absent
log_e(Bit Score)
0.00
0.25
0.50
0.75
1.00
Distance from M. lignano
Bristle Status
present
reduced
absent

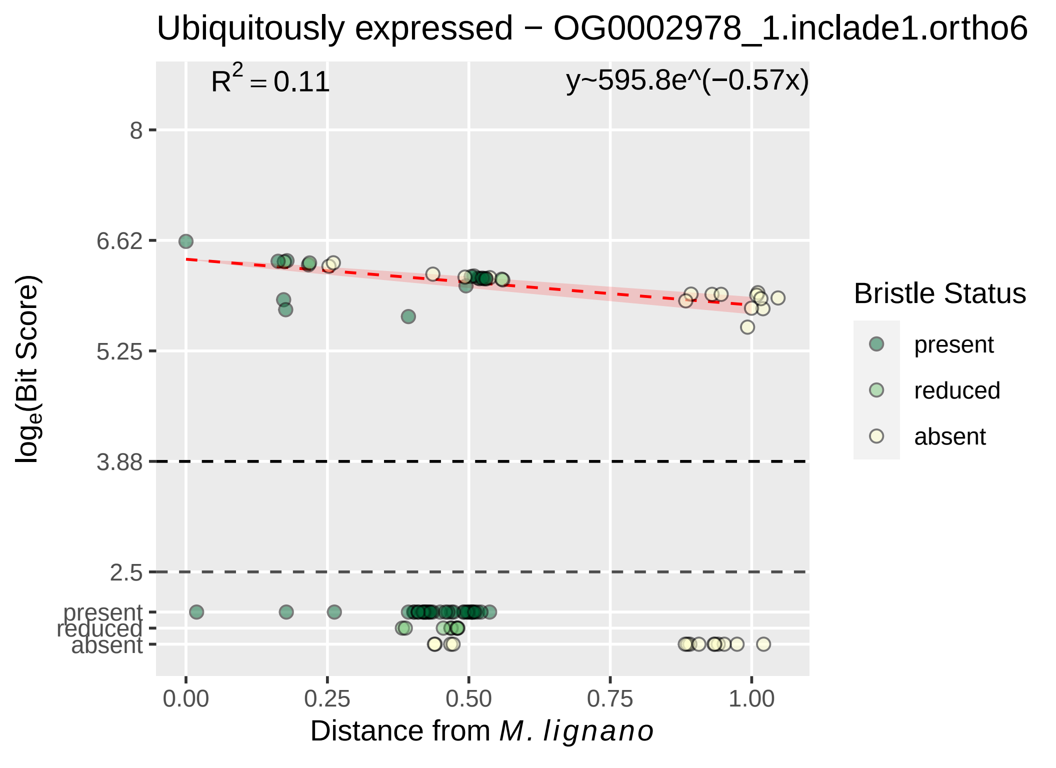

Ubiquitously expressed − OG0002978_1.inclade1.ortho6
R² = 0.11
y~595.8e^(−0.57x)
8
6.62
5.25
3.88
2.5
present
reduced
absent
log_e(Bit Score)
0.00
0.25
0.50
0.75
1.00
Distance from M. lignano
Bristle Status
present
reduced
absent

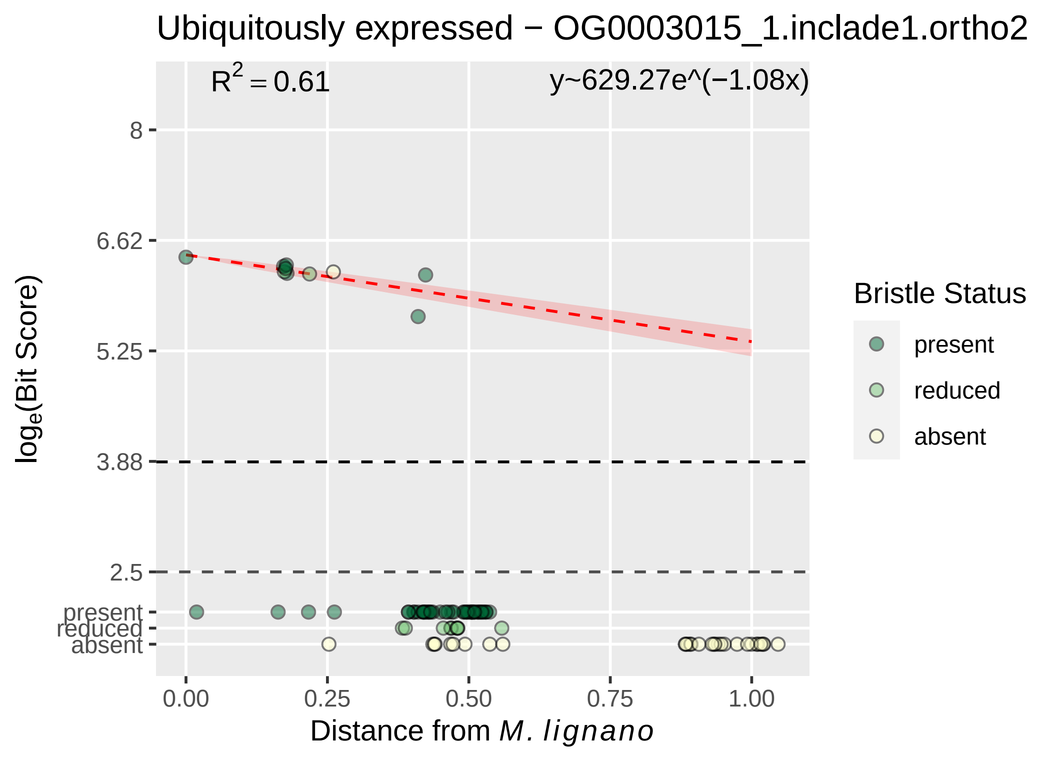

Ubiquitously expressed − OG0003015_1.inclade1.ortho2
R^2 = 0.61
y~629.27e^(−1.08x)
8
6.62
5.25
3.88
2.5
present
reduced
absent
log_e(Bit Score)
0.00
0.25
0.50
0.75
1.00
Distance from M. lignano
Bristle Status
present
reduced
absent

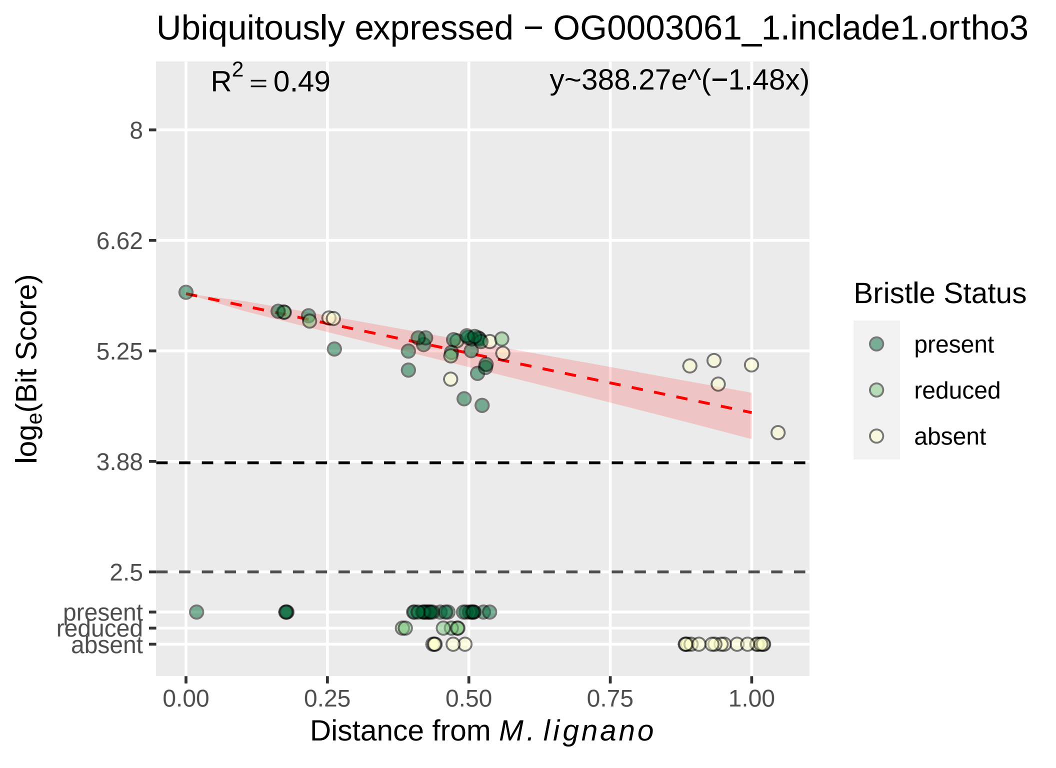

Ubiquitously expressed − OG0003061_1.inclade1.ortho3
R2 = 0.49
y~388.27e^(−1.48x)
8
6.62
5.25
3.88
2.5
present
reduced
absent
log_e(Bit Score)
0.00
0.25
0.50
0.75
1.00
Distance from M. lignano
Bristle Status
present
reduced
absent

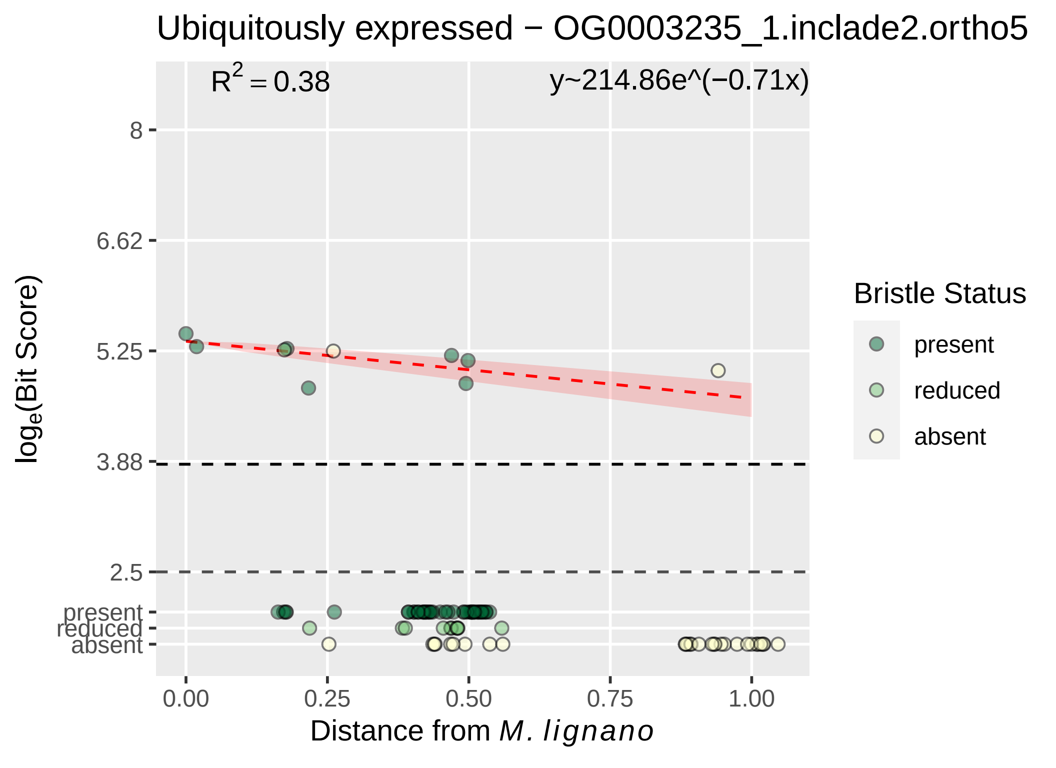

Ubiquitously expressed − OG0003235_1.inclade2.ortho5
R2 = 0.38
y~214.86e^(−0.71x)
8
6.62
5.25
3.88
2.5
present
reduced
absent
0.00
0.25
0.50
0.75
1.00
Distance from M. lignano
loge(Bit Score)
Bristle Status
present
reduced
absent

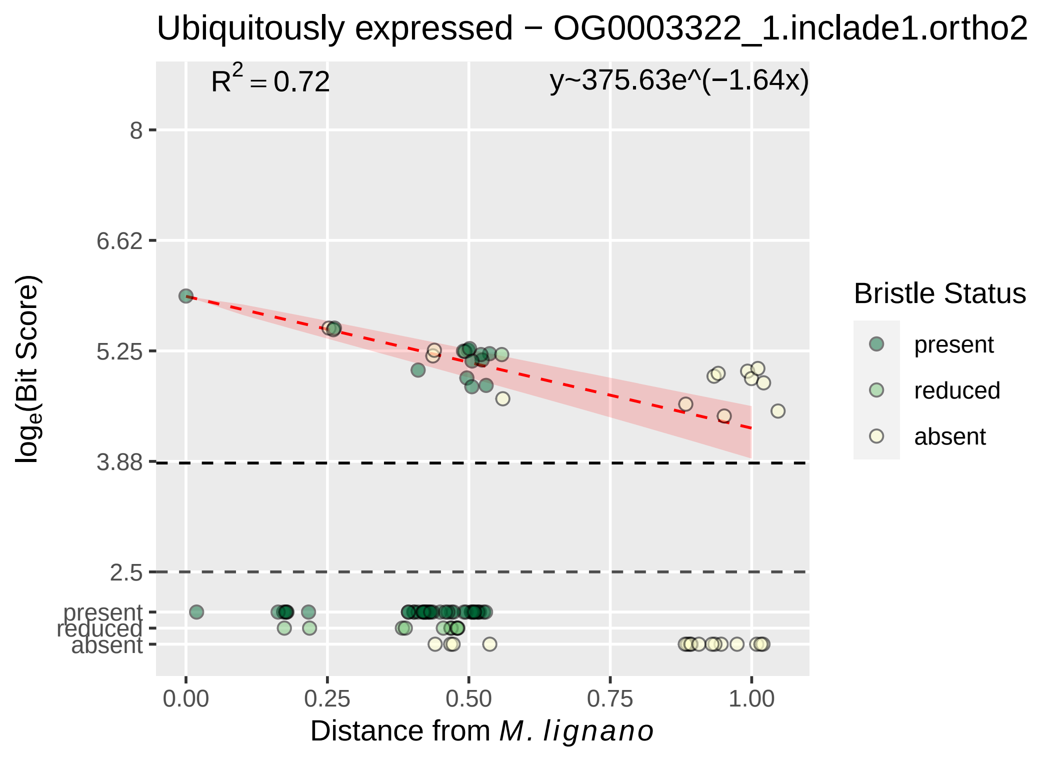

Ubiquitously expressed − OG0003322_1.inclade1.ortho2
R² = 0.72
y~375.63e^(−1.64x)
8
6.62
5.25
3.88
2.5
present
reduced
absent
log_e(Bit Score)
0.00
0.25
0.50
0.75
1.00
Distance from M. lignano
Bristle Status
present
reduced
absent

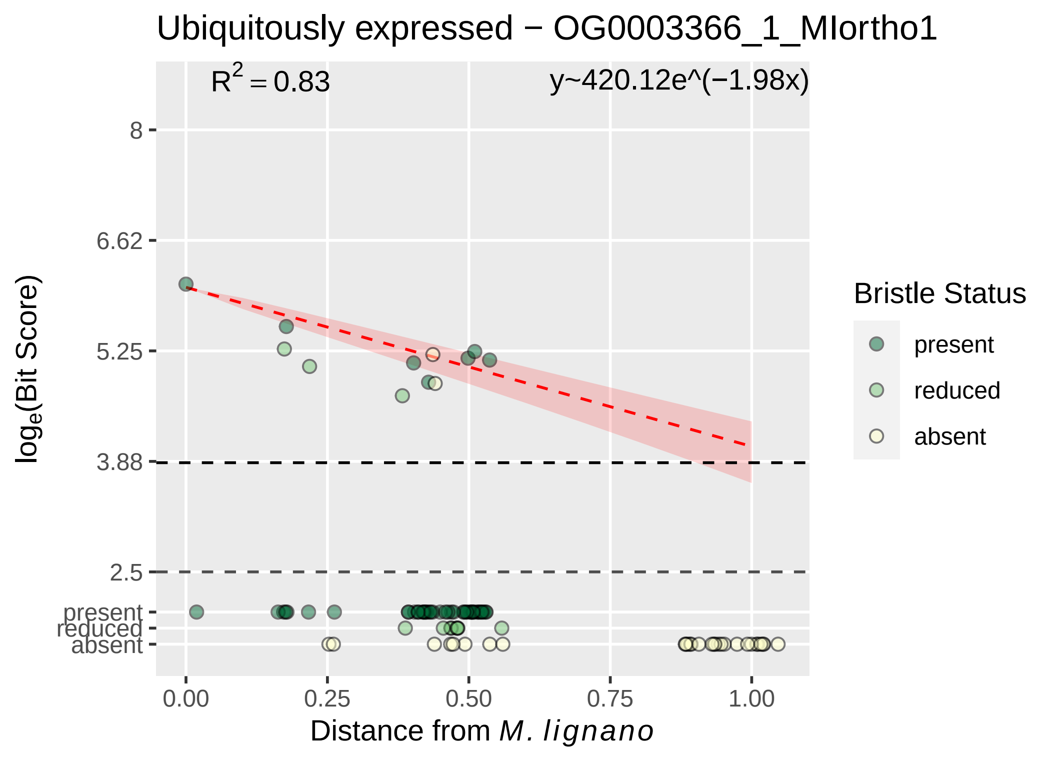

Ubiquitously expressed − OG0003366_1_MIortho1
R² = 0.83
y~420.12e^(−1.98x)
8
6.62
5.25
3.88
2.5
present
reduced
absent
0.00
0.25
0.50
0.75
1.00
Distance from M. lignano
log_e(Bit Score)
Bristle Status
present
reduced
absent

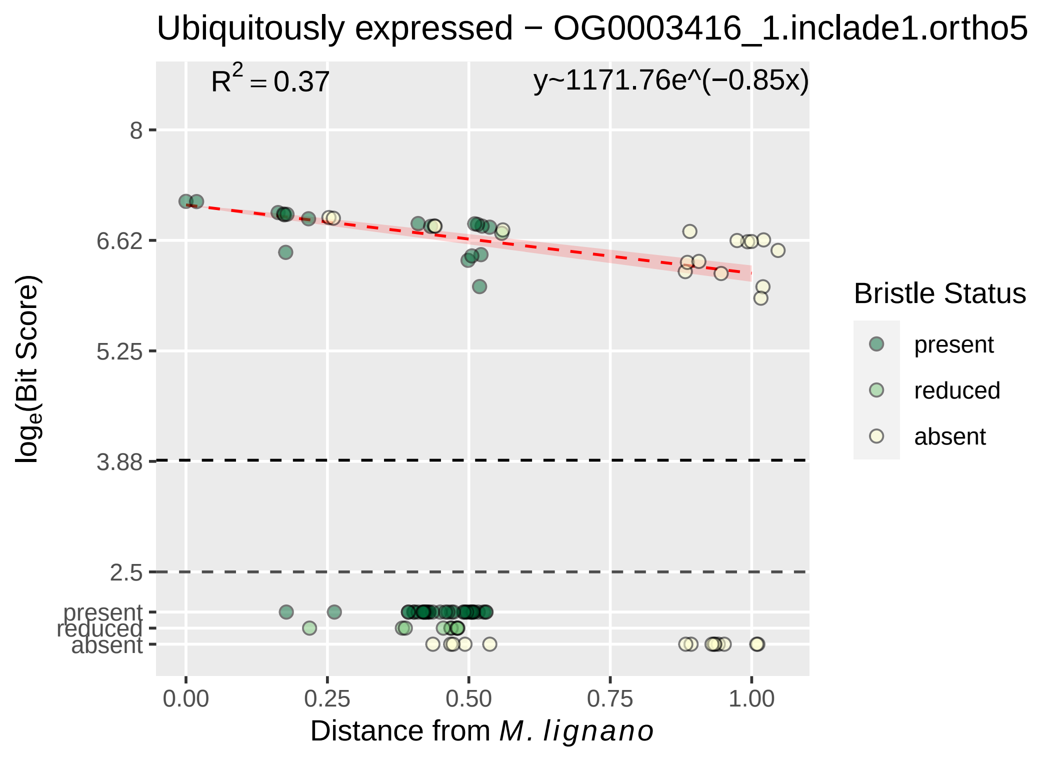

Ubiquitously expressed − OG0003416_1.inclade1.ortho5
R^2 = 0.37
y~1171.76e^(−0.85x)
8
6.62
5.25
3.88
2.5
present
reduced
absent
log_e(Bit Score)
0.00
0.25
0.50
0.75
1.00
Distance from M. lignano
Bristle Status
present
reduced
absent

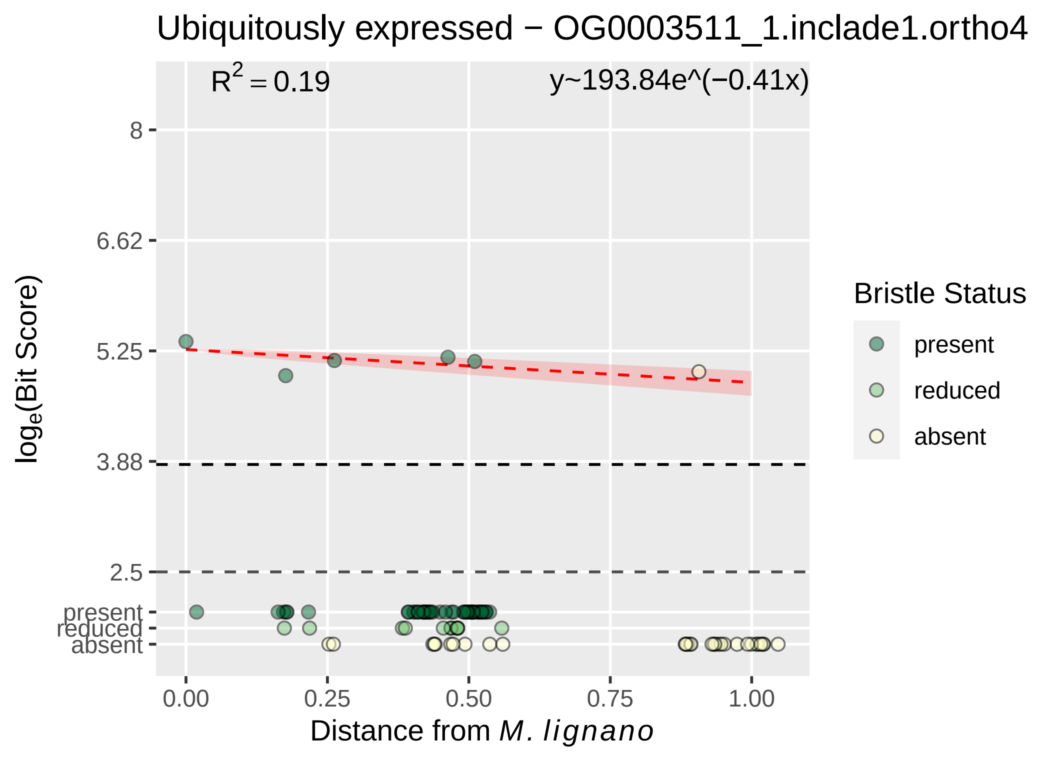

Ubiquitously expressed − OG0003511_1.inclade1.ortho4
R2 = 0.19
y~193.84e^(−0.41x)
8
6.62
5.25
3.88
2.5
present
reduced
absent
loge(Bit Score)
0.00
0.25
0.50
0.75
1.00
Distance from M. lignano
Bristle Status
present
reduced
absent

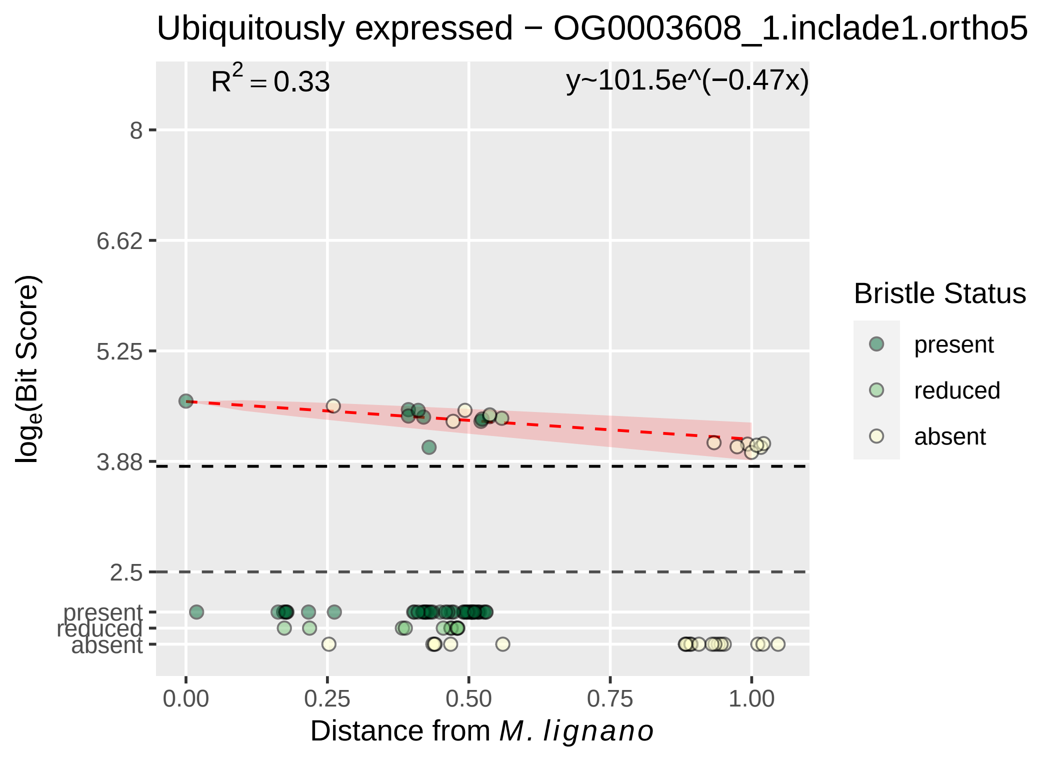

Ubiquitously expressed − OG0003608_1.inclade1.ortho5
R² = 0.33
y~101.5e^(−0.47x)
8
6.62
5.25
3.88
2.5
present
reduced
absent
log_e(Bit Score)
0.00
0.25
0.50
0.75
1.00
Distance from M. lignano
Bristle Status
present
reduced
absent

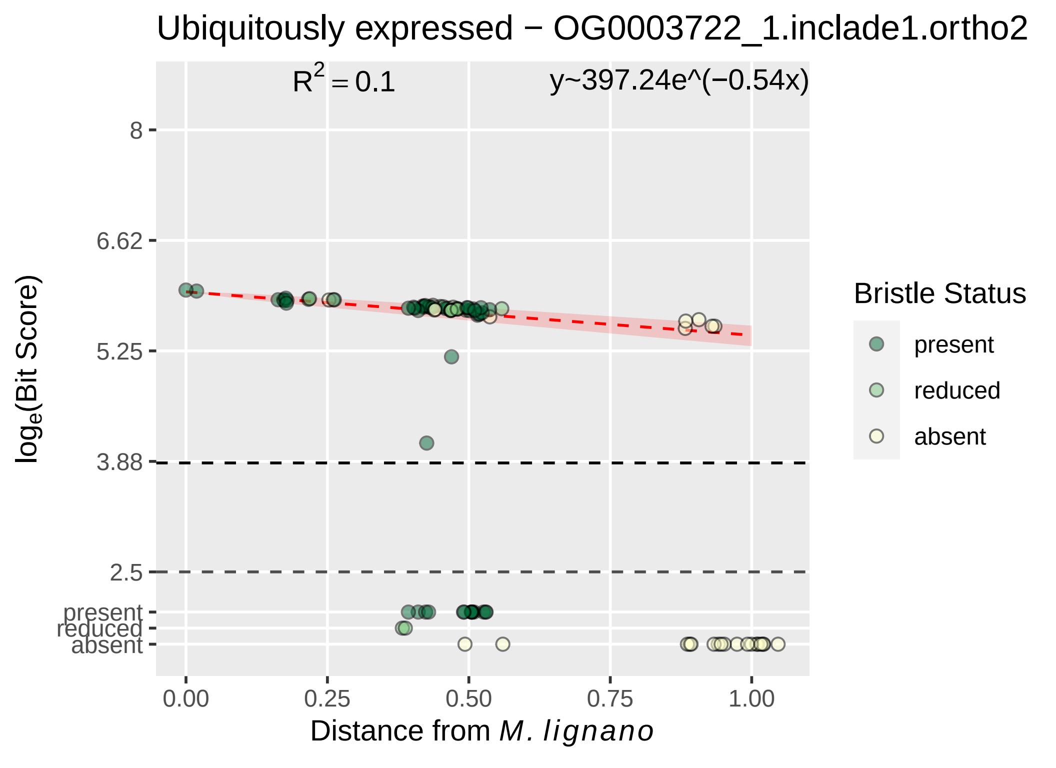

Ubiquitously expressed − OG0003722_1.inclade1.ortho2
R^2 = 0.1
y~397.24e^(−0.54x)
8
6.62
5.25
3.88
2.5
present
reduced
absent
0.00
0.25
0.50
0.75
1.00
log_e(Bit Score)
Distance from M. lignano
Bristle Status
present
reduced
absent

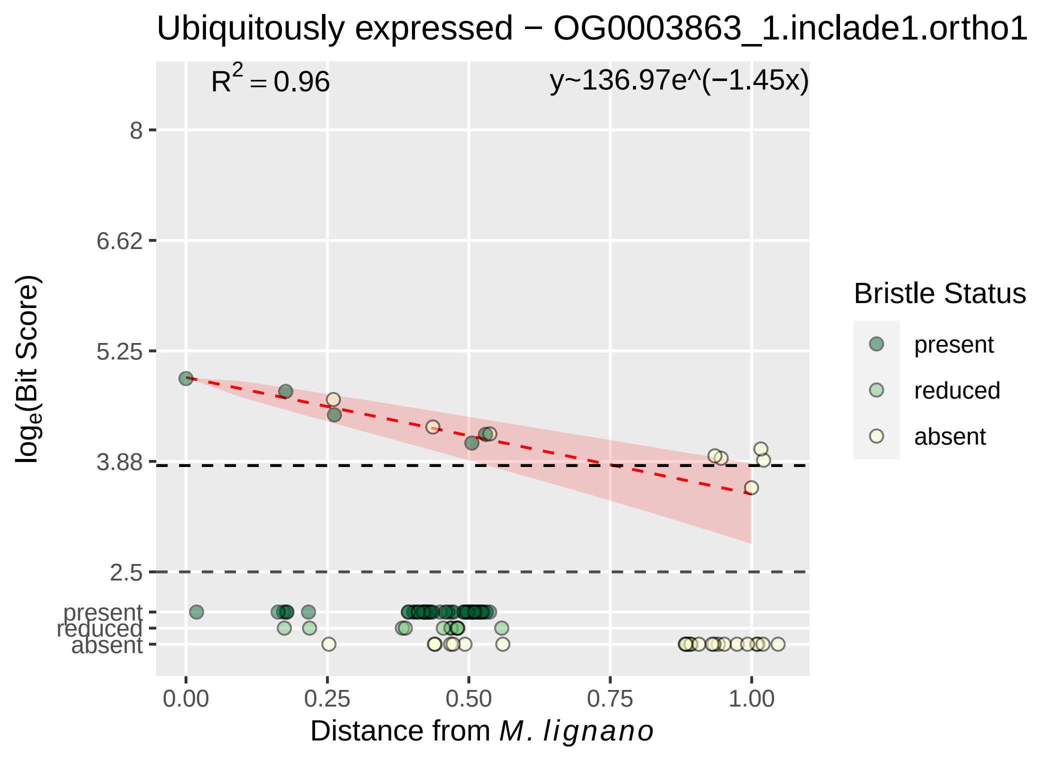

Ubiquitously expressed − OG0003863_1.inclade1.ortho1
$R^2 = 0.96$
y~136.97e^(−1.45x)
8
6.62
5.25
3.88
2.5
present
reduced
absent
log_e(Bit Score)
0.00
0.25
0.50
0.75
1.00
Distance from M. lignano
Bristle Status
present
reduced
absent

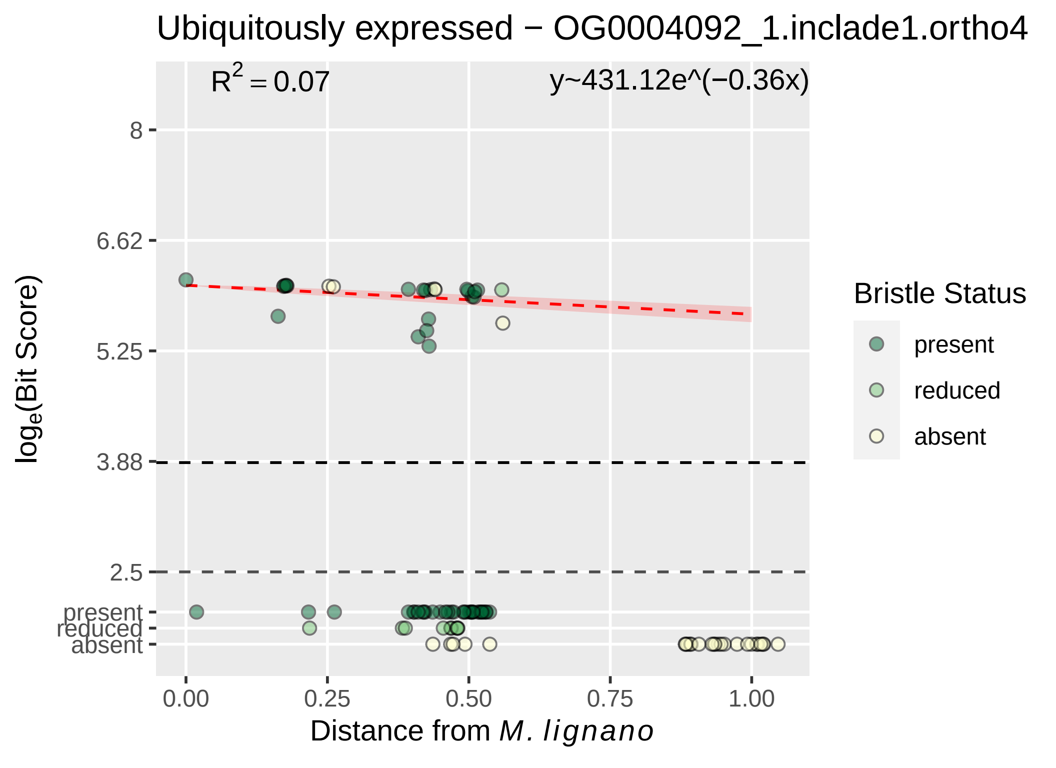

Ubiquitously expressed − OG0004092_1.inclade1.ortho4
R^2 = 0.07
y~431.12e^(−0.36x)
8
6.62
5.25
3.88
2.5
present
reduced
absent
log_e(Bit Score)
0.00
0.25
0.50
0.75
1.00
Distance from M. lignano
Bristle Status
present
reduced
absent

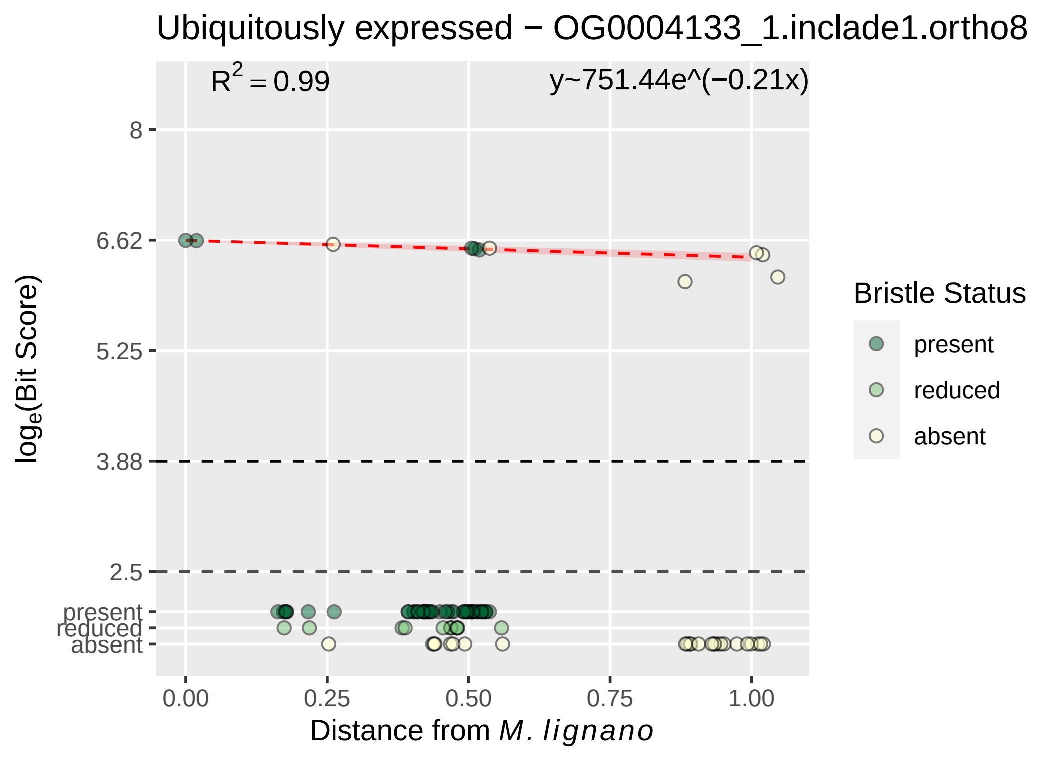

Ubiquitously expressed − OG0004133_1.inclade1.ortho8
R^2 = 0.99
y~751.44e^(−0.21x)
8
6.62
5.25
3.88
2.5
present
reduced
absent
log_e(Bit Score)
0.00
0.25
0.50
0.75
1.00
Distance from M. lignano
Bristle Status
present
reduced
absent

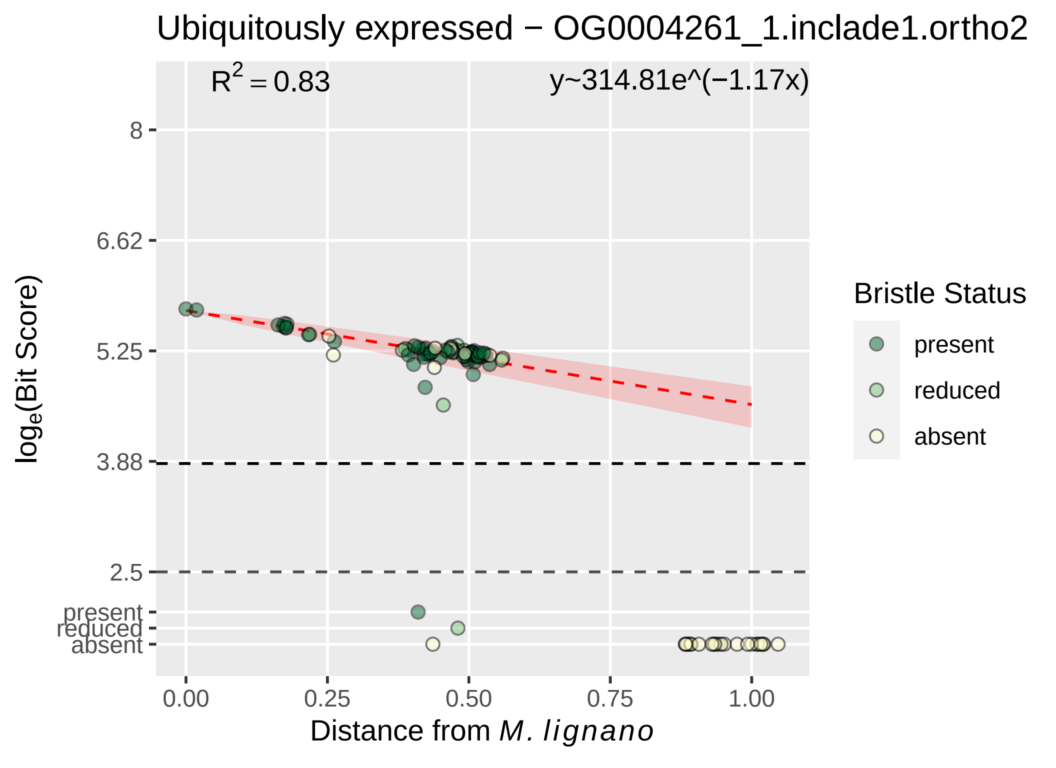

Ubiquitously expressed − OG0004261_1.inclade1.ortho2
R2 = 0.83
y~314.81e^(−1.17x)
8
6.62
5.25
3.88
2.5
present
reduced
absent
0.00
0.25
0.50
0.75
1.00
Distance from M. lignano
loge(Bit Score)
Bristle Status
present
reduced
absent

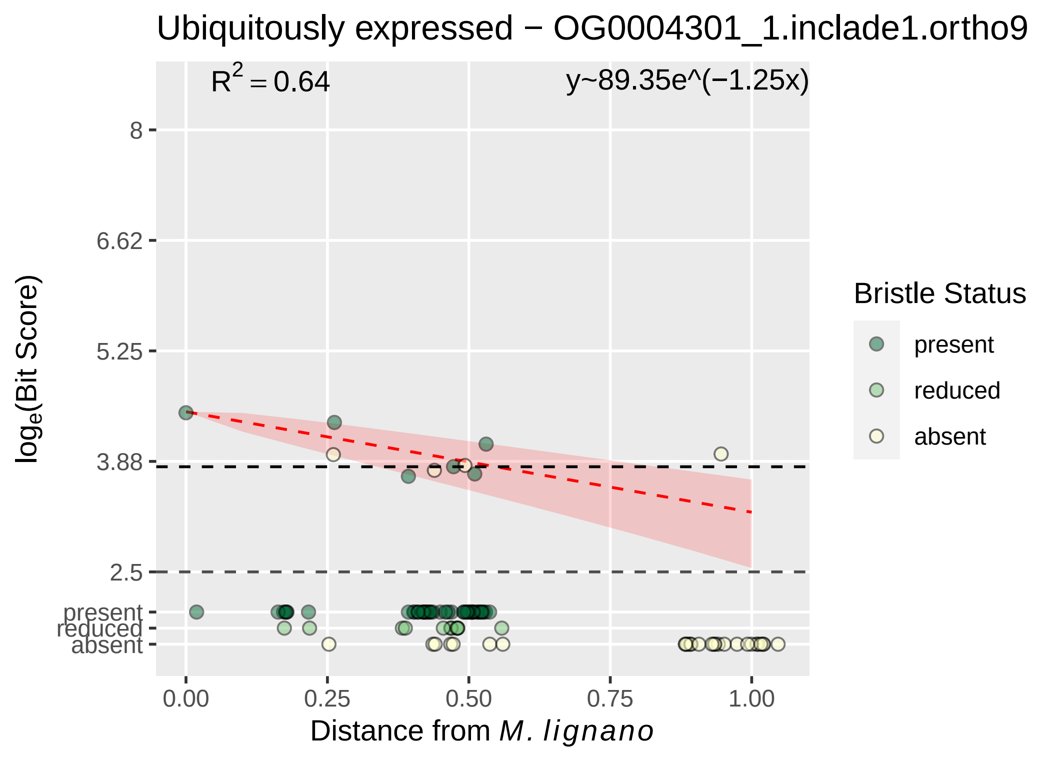

Ubiquitously expressed − OG0004301_1.inclade1.ortho9
$R^2 = 0.64$
y~89.35e^(−1.25x)
8
6.62
5.25
3.88
2.5
present
reduced
absent
0.00
0.25
0.50
0.75
1.00
log$_e$(Bit Score)
Distance from M. lignano
Bristle Status
present
reduced
absent

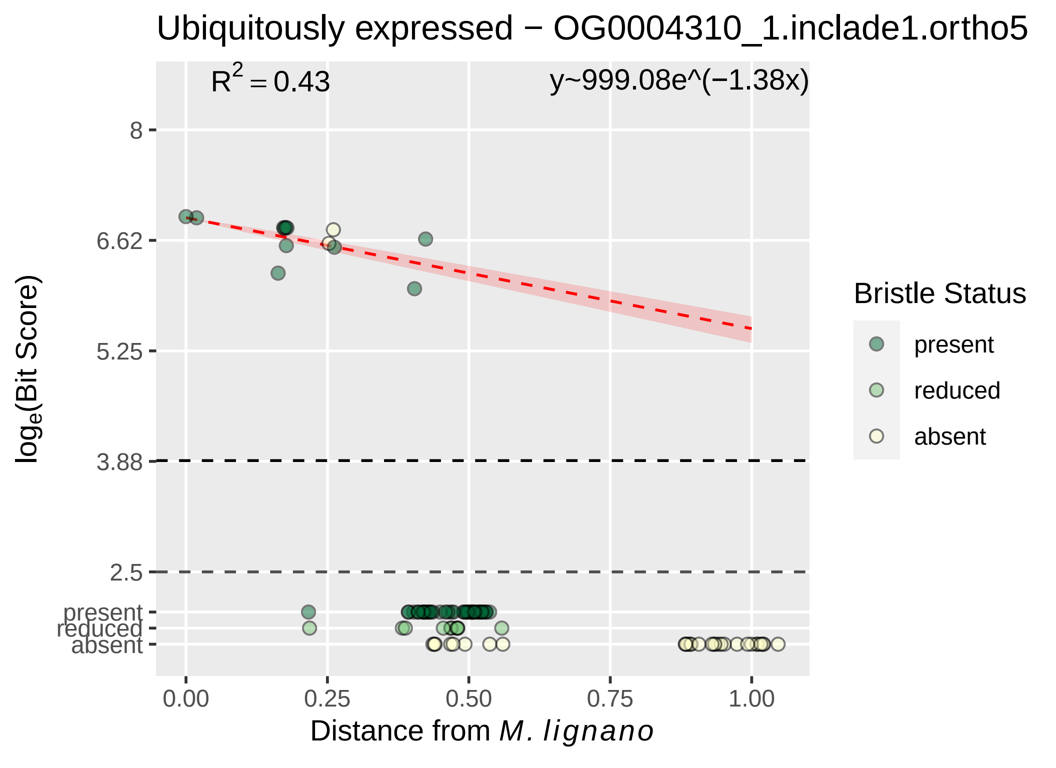

Ubiquitously expressed − OG0004310_1.inclade1.ortho5
R^2 = 0.43
y~999.08e^(−1.38x)
8
6.62
5.25
3.88
2.5
present
reduced
absent
0.00
0.25
0.50
0.75
1.00
Distance from M. lignano
log_e(Bit Score)
Bristle Status
present
reduced
absent

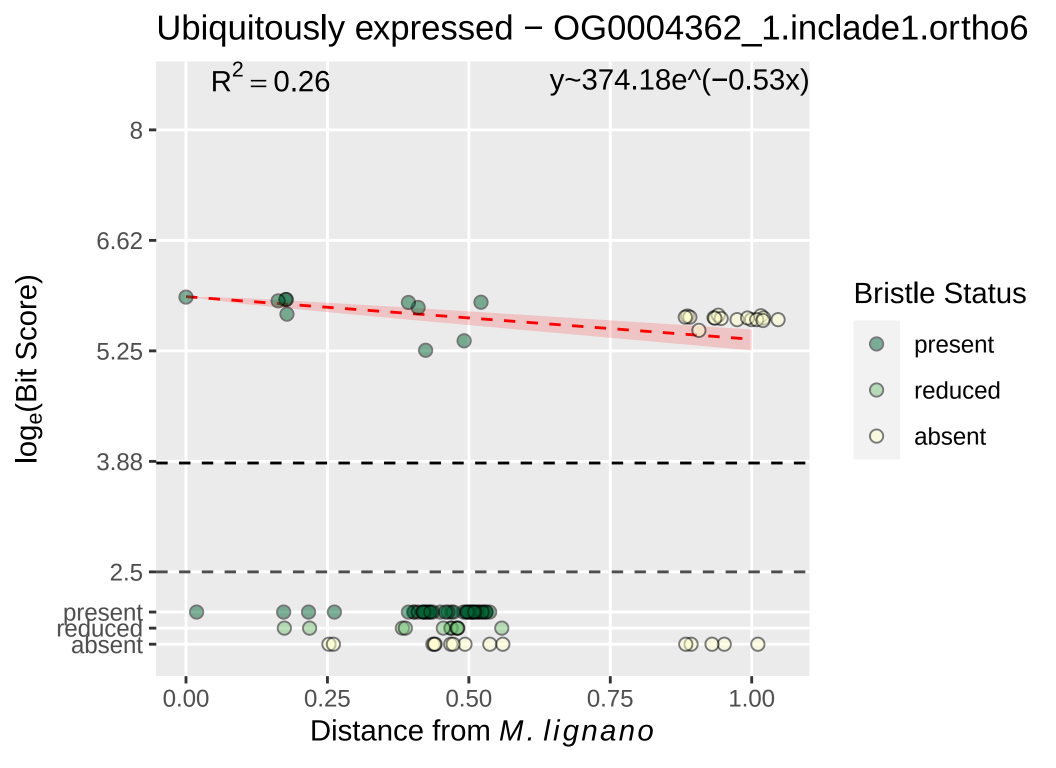

Ubiquitously expressed − OG0004362_1.inclade1.ortho6
$R^2 = 0.26$
y~374.18e^(−0.53x)
8
6.62
5.25
3.88
2.5
present
reduced
absent
$\log_e$(Bit Score)
0.00
0.25
0.50
0.75
1.00
Distance from *M. lignano*
Bristle Status
present
reduced
absent

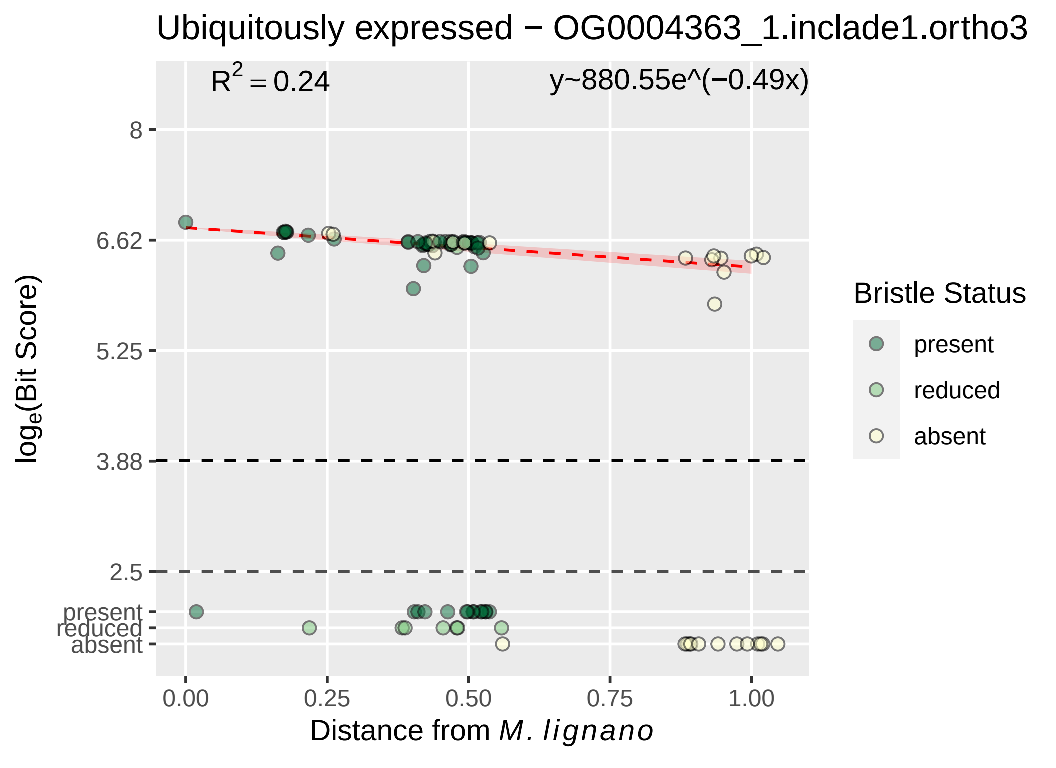

Ubiquitously expressed − OG0004363_1.inclade1.ortho3
$R^2 = 0.24$
y~880.55e^(−0.49x)
8
6.62
5.25
3.88
2.5
present
reduced
absent
0.00
0.25
0.50
0.75
1.00
Distance from M. lignano
$\log_e$(Bit Score)
Bristle Status
present
reduced
absent

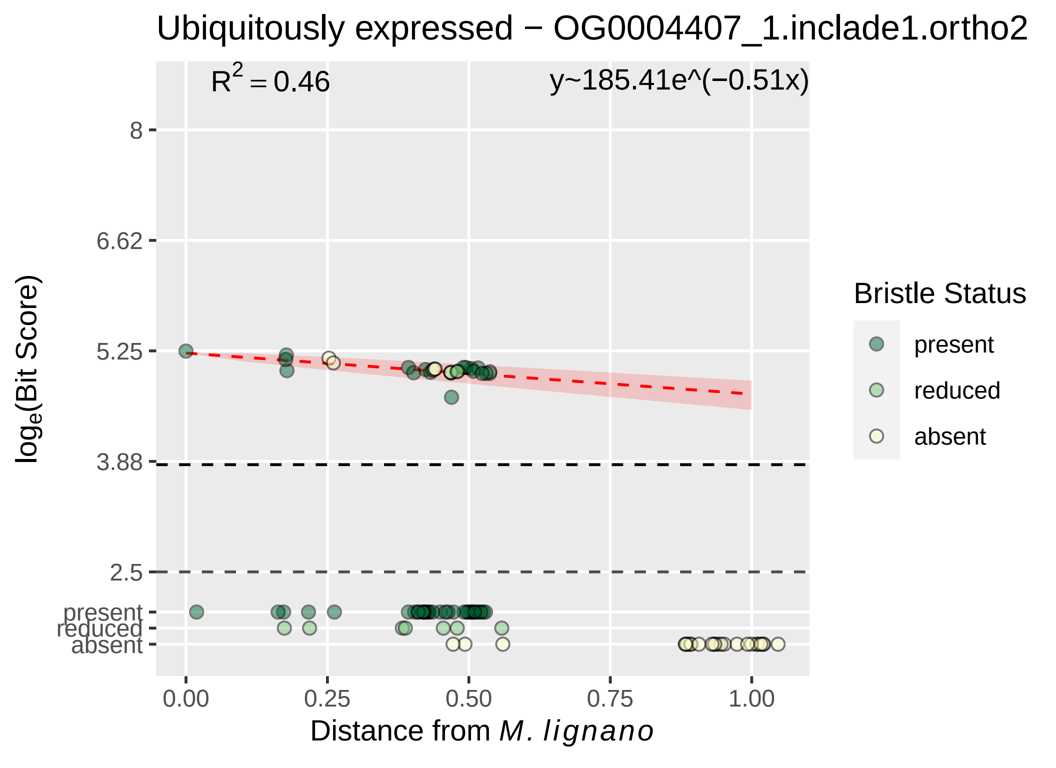
Ubiquitously expressed − OG0004407_1.inclade1.ortho2
$R^2 = 0.46$
y~185.41e^(−0.51x)
8
6.62
5.25
3.88
2.5
present
reduced
absent
$\log_e$(Bit Score)
0.00
0.25
0.50
0.75
1.00
Distance from *M. lignano*
Bristle Status
present
reduced
absent

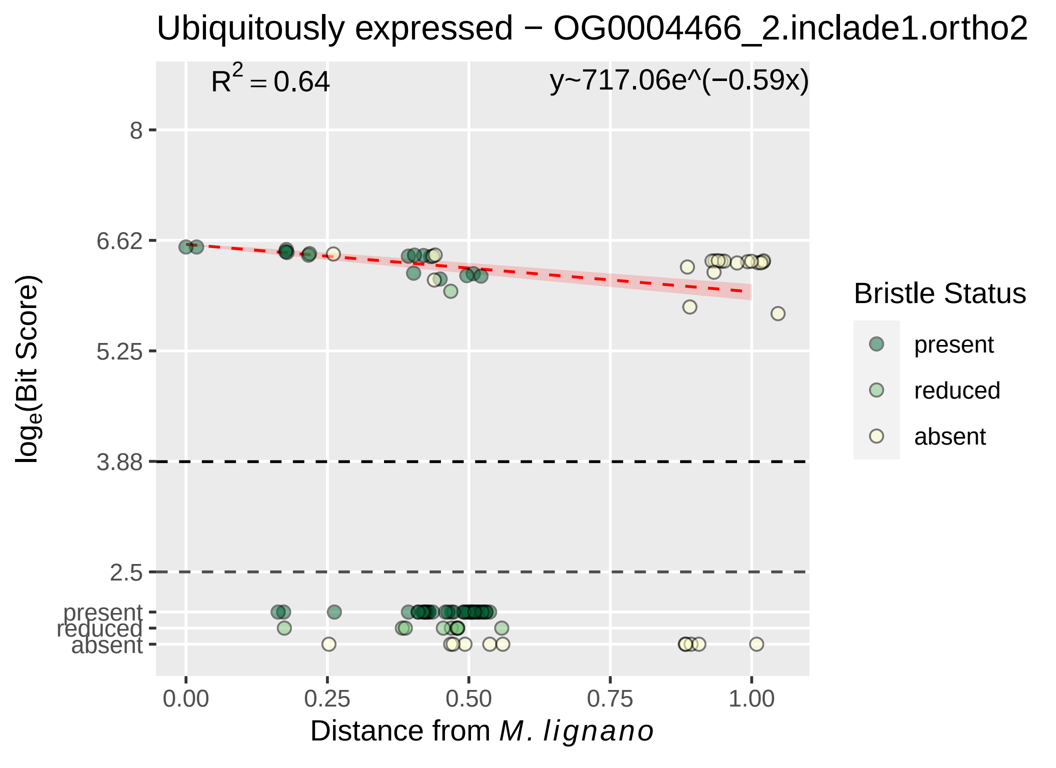
Ubiquitously expressed − OG0004466_2.inclade1.ortho2
$R^2 = 0.64$
y~717.06e^(−0.59x)
8
6.62
5.25
3.88
2.5
present
reduced
absent
0.00
0.25
0.50
0.75
1.00
Distance from M. lignano
log$_e$(Bit Score)
Bristle Status
present
reduced
absent

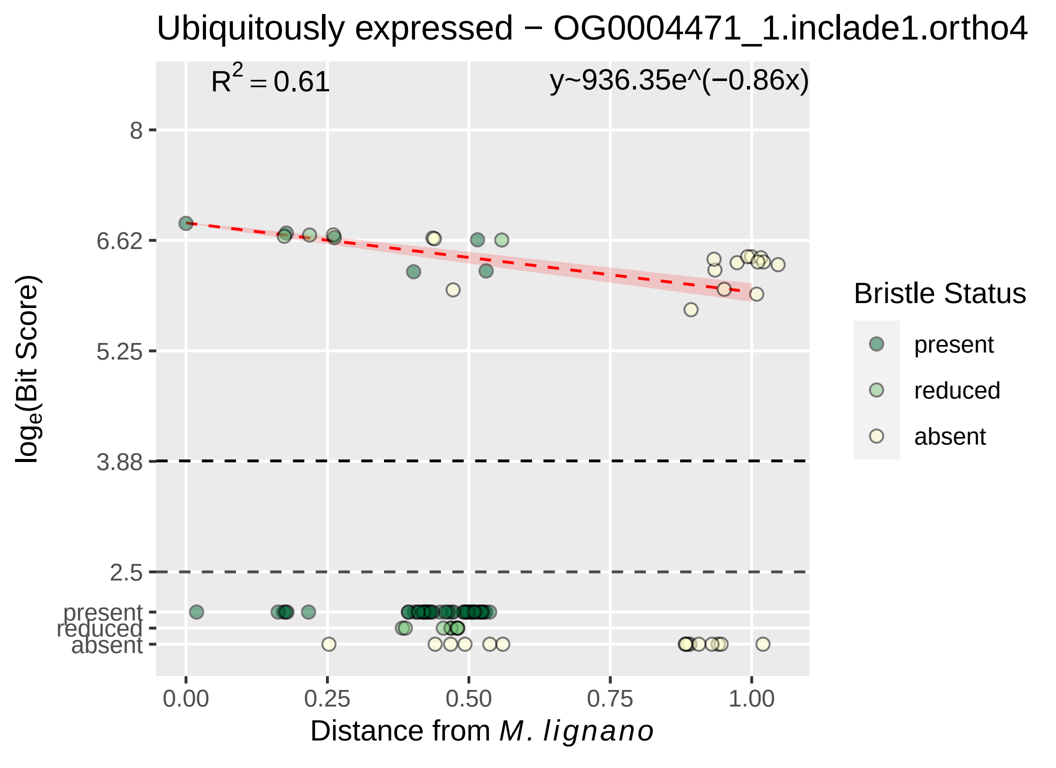
Ubiquitously expressed − OG0004471_1.inclade1.ortho4
R² = 0.61
y~936.35e^(−0.86x)
log_e(Bit Score)
8
6.62
5.25
3.88
2.5
present
reduced
absent
0.00
0.25
0.50
0.75
1.00
Distance from M. lignano
Bristle Status
present
reduced
absent

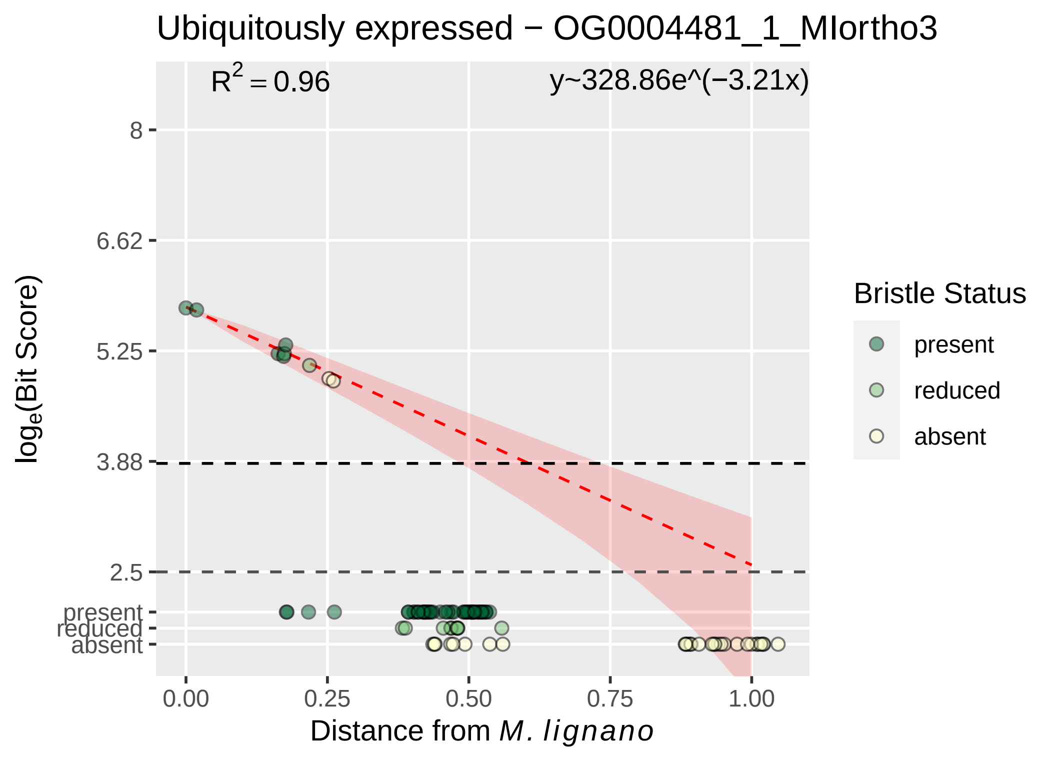
Ubiquitously expressed − OG0004481_1_MIortho3
R² = 0.96
y~328.86e^(−3.21x)
8
6.62
5.25
3.88
2.5
present
reduced
absent
log_e(Bit Score)
0.00
0.25
0.50
0.75
1.00
Distance from M. lignano
Bristle Status
present
reduced
absent

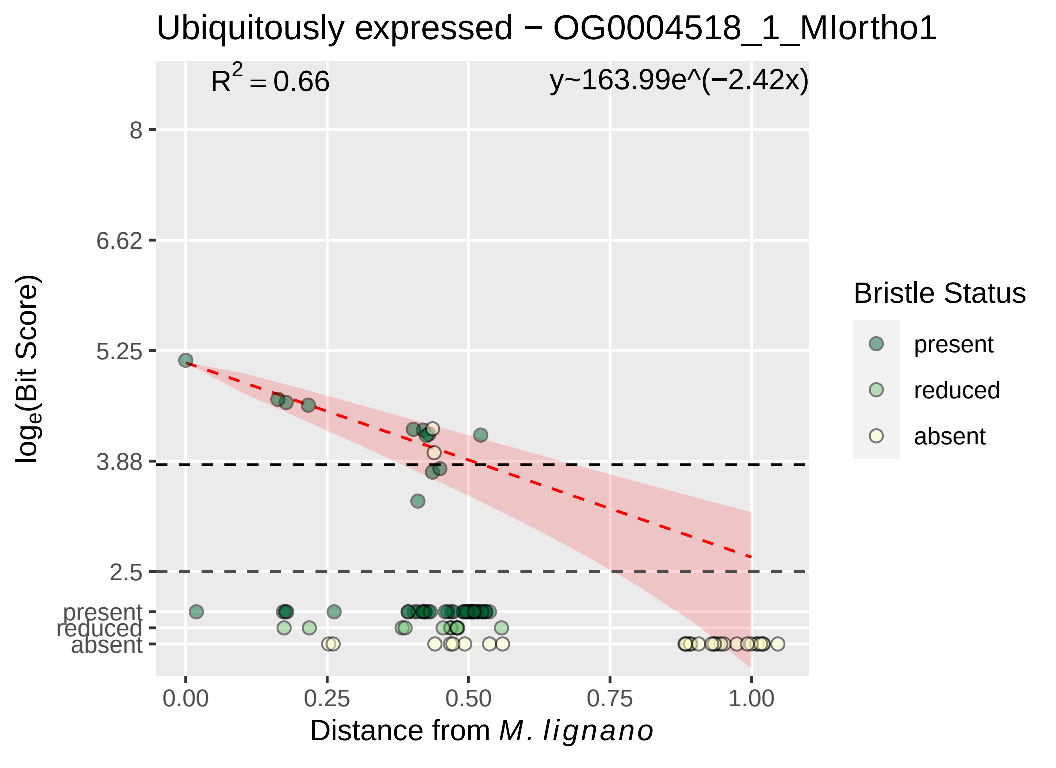
Ubiquitously expressed − OG0004518_1_MIortho1
R^2 = 0.66
y~163.99e^(−2.42x)
8
6.62
5.25
3.88
2.5
present
reduced
absent
0.00
0.25
0.50
0.75
1.00
loge(Bit Score)
Distance from M. lignano
Bristle Status
present
reduced
absent

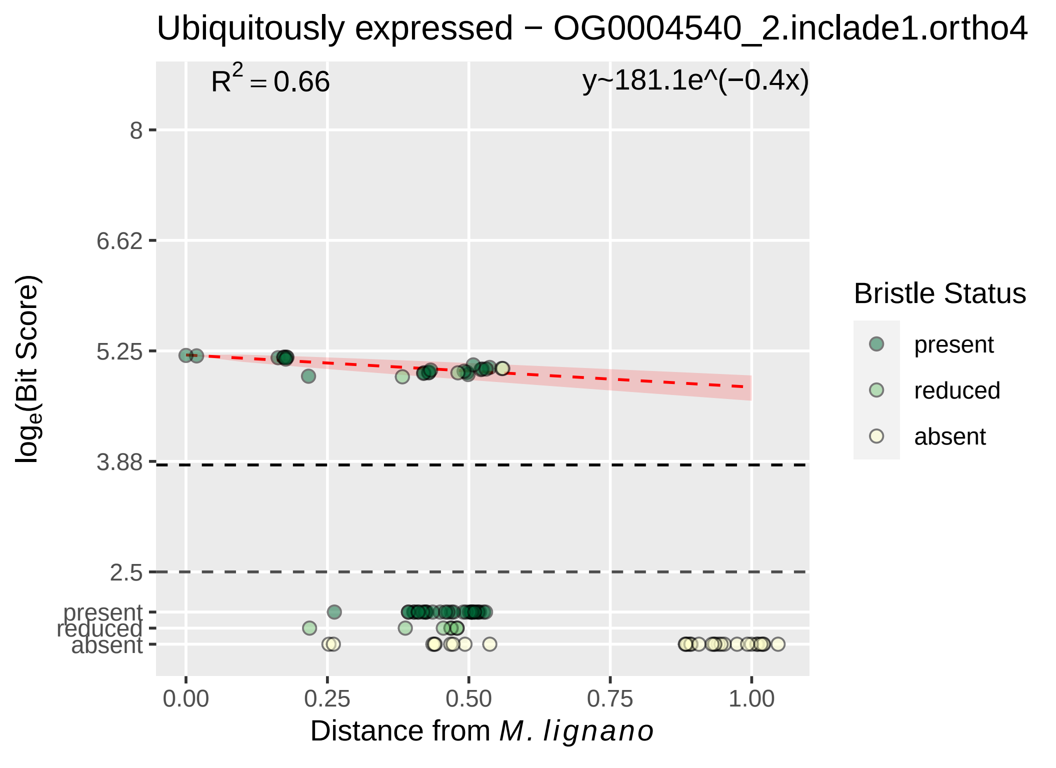

Ubiquitously expressed − OG0004540_2.inclade1.ortho4
R2 = 0.66
y~181.1e^(−0.4x)
loge(Bit Score)
8
6.62
5.25
3.88
2.5
present
reduced
absent
0.00
0.25
0.50
0.75
1.00
Distance from M. lignano
Bristle Status
present
reduced
absent

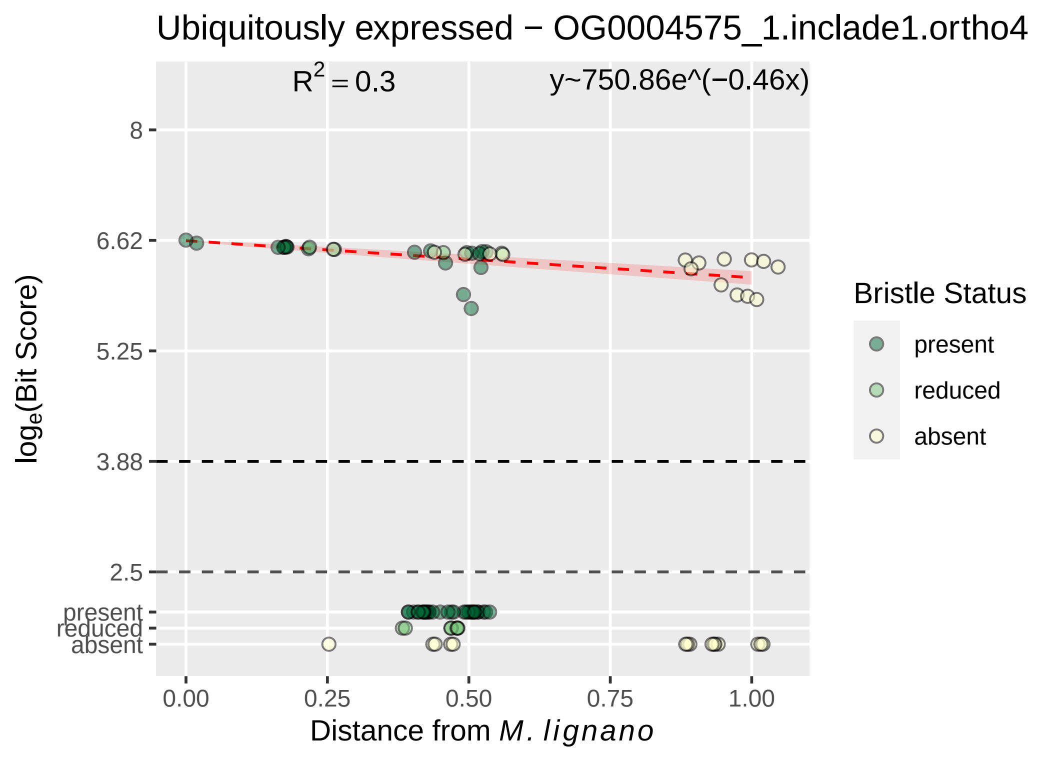

Ubiquitously expressed − OG0004575_1.inclade1.ortho4
R^2 = 0.3
y~750.86e^(−0.46x)
8
6.62
5.25
3.88
2.5
present
reduced
absent
0.00
0.25
0.50
0.75
1.00
Distance from M. lignano
loge(Bit Score)
Bristle Status
present
reduced
absent

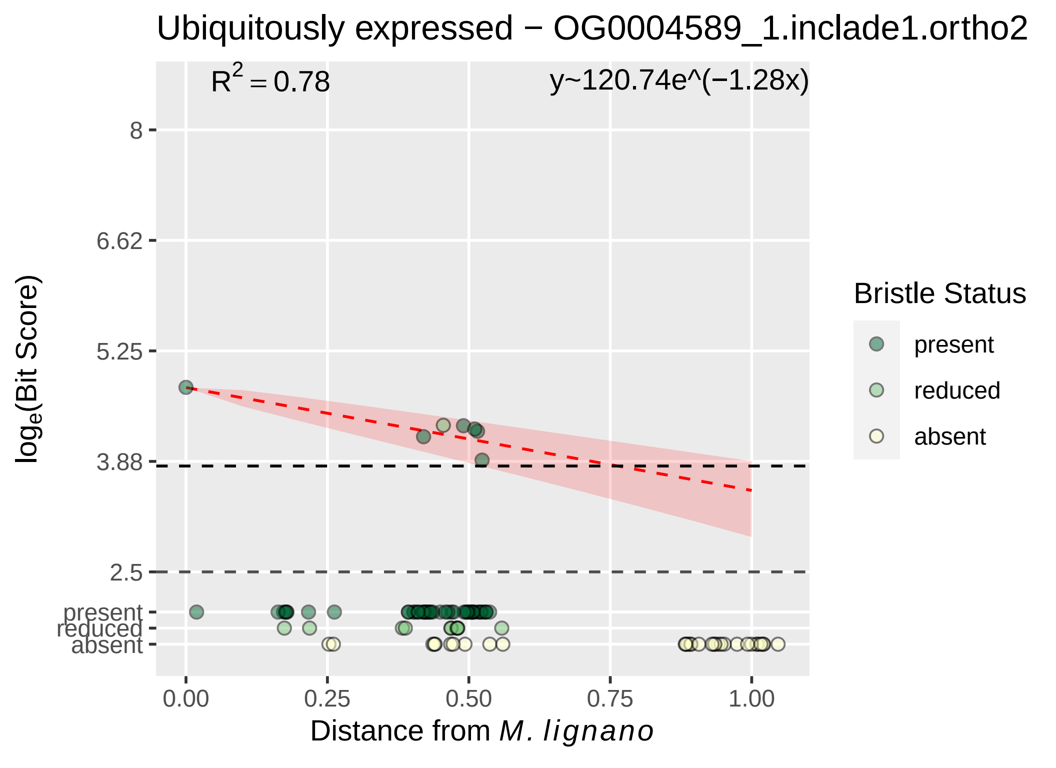

Ubiquitously expressed − OG0004589_1.inclade1.ortho2
$R^2 = 0.78$
y~120.74e^(−1.28x)
8
6.62
5.25
3.88
2.5
present
reduced
absent
$\log_e$(Bit Score)
0.00
0.25
0.50
0.75
1.00
Distance from *M. lignano*
Bristle Status
present
reduced
absent

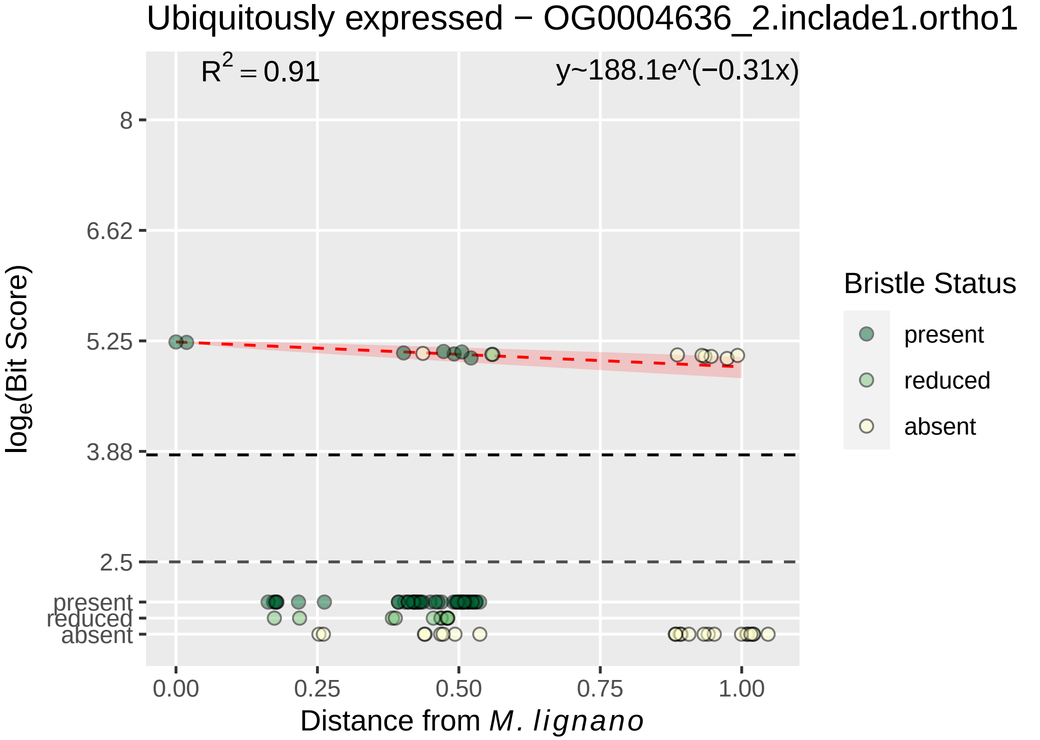

Ubiquitously expressed − OG0004636_2.inclade1.ortho1
$R^2 = 0.91$
y~188.1e^(−0.31x)
8
6.62
5.25
3.88
2.5
present
reduced
absent
$\log_e$(Bit Score)
0.00
0.25
0.50
0.75
1.00
Distance from *M. lignano*
Bristle Status
present
reduced
absent

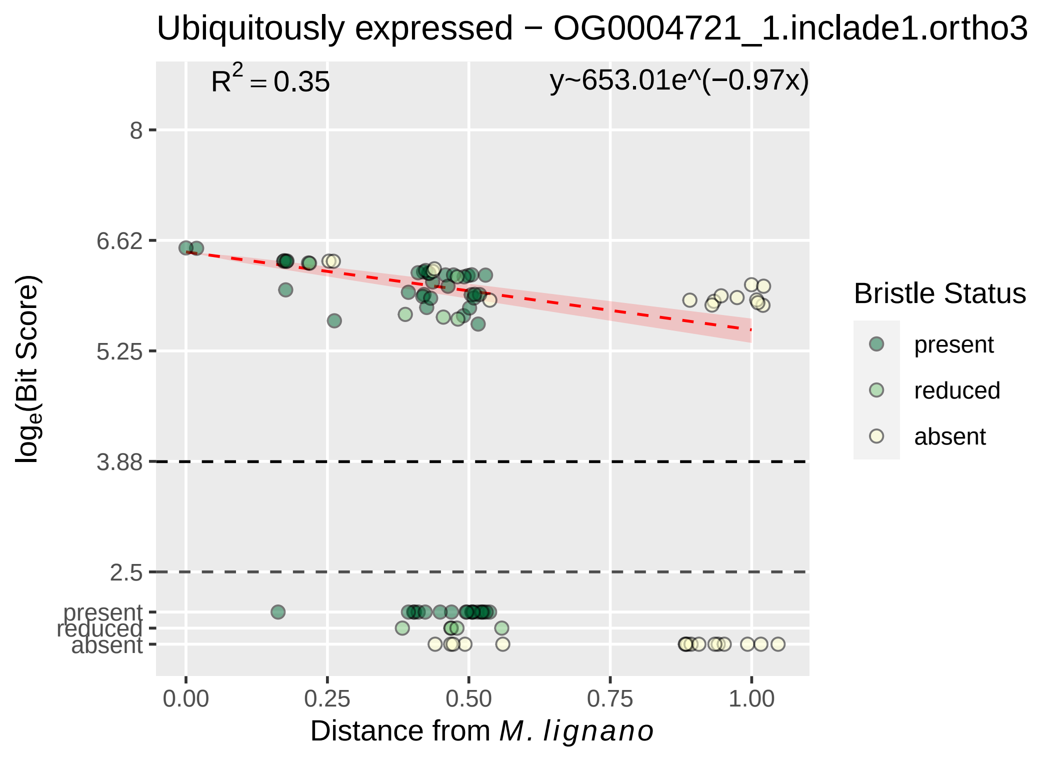

Ubiquitously expressed − OG0004721_1.inclade1.ortho3
R2 = 0.35
y~653.01e^(−0.97x)
8
6.62
5.25
3.88
2.5
present
reduced
absent
loge(Bit Score)
0.00
0.25
0.50
0.75
1.00
Distance from M. lignano
Bristle Status
present
reduced
absent

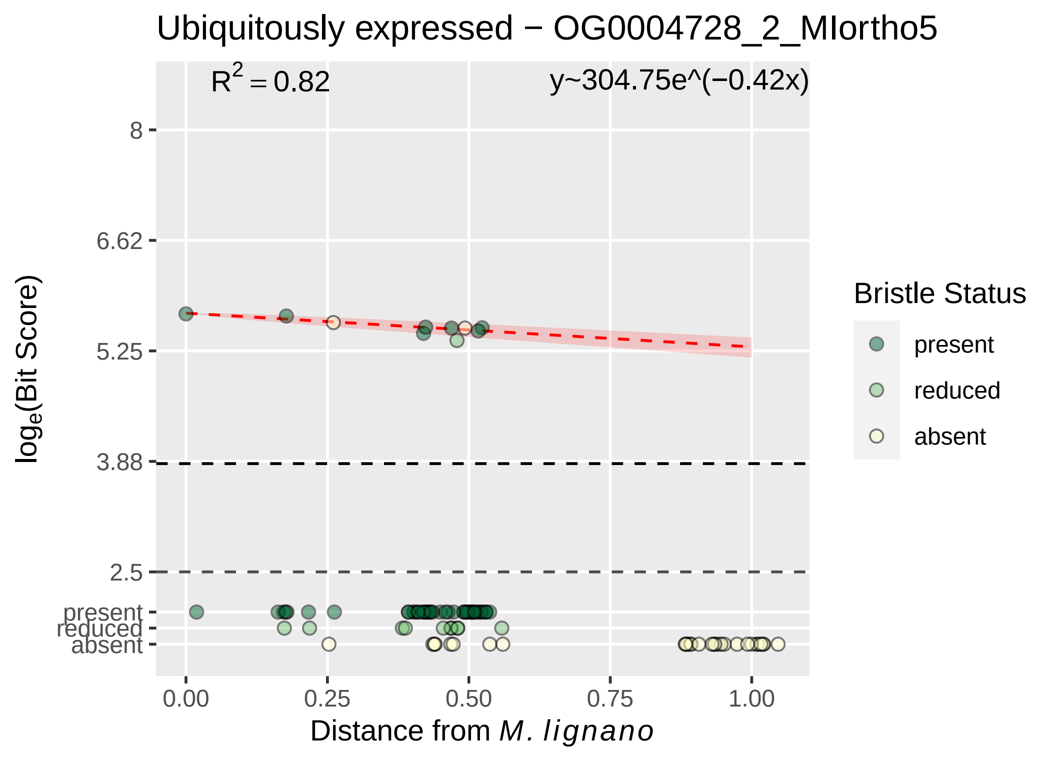

Ubiquitously expressed − OG0004728_2_MIortho5
R^2 = 0.82
y~304.75e^(−0.42x)
8
6.62
5.25
3.88
2.5
present
reduced
absent
log_e(Bit Score)
0.00
0.25
0.50
0.75
1.00
Distance from M. lignano
Bristle Status
present
reduced
absent

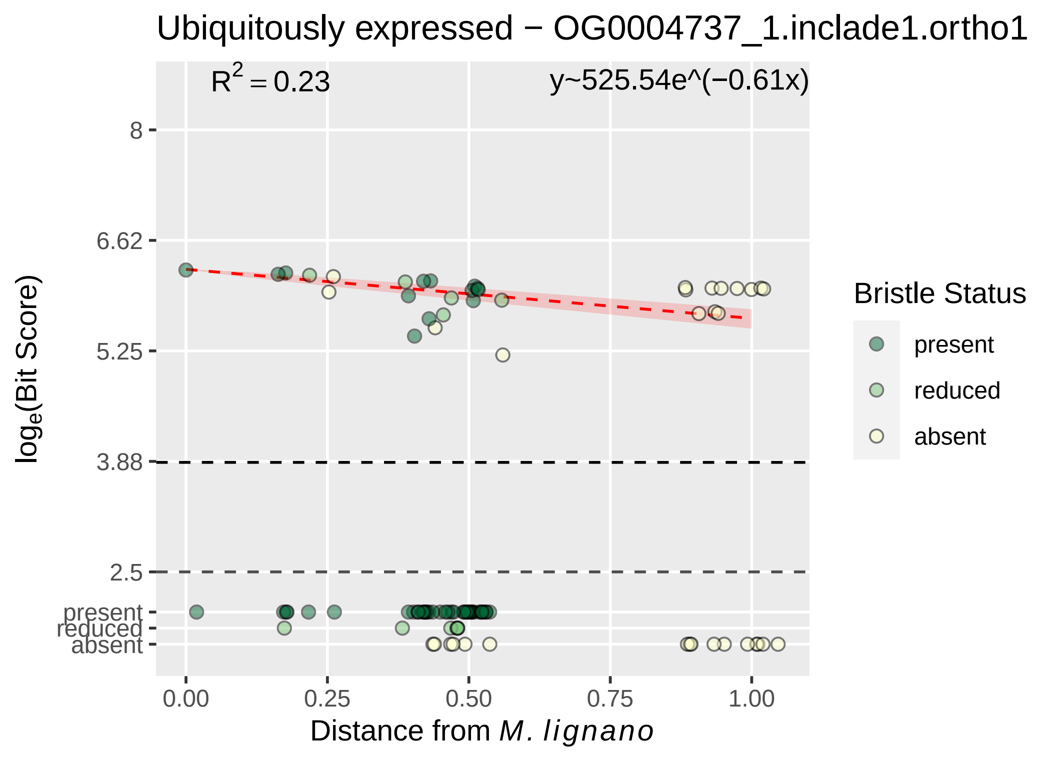

Ubiquitously expressed − OG0004737_1.inclade1.ortho1
R2 = 0.23
y~525.54e^(−0.61x)
8
6.62
5.25
3.88
2.5
present
reduced
absent
loge(Bit Score)
0.00
0.25
0.50
0.75
1.00
Distance from M. lignano
Bristle Status
present
reduced
absent

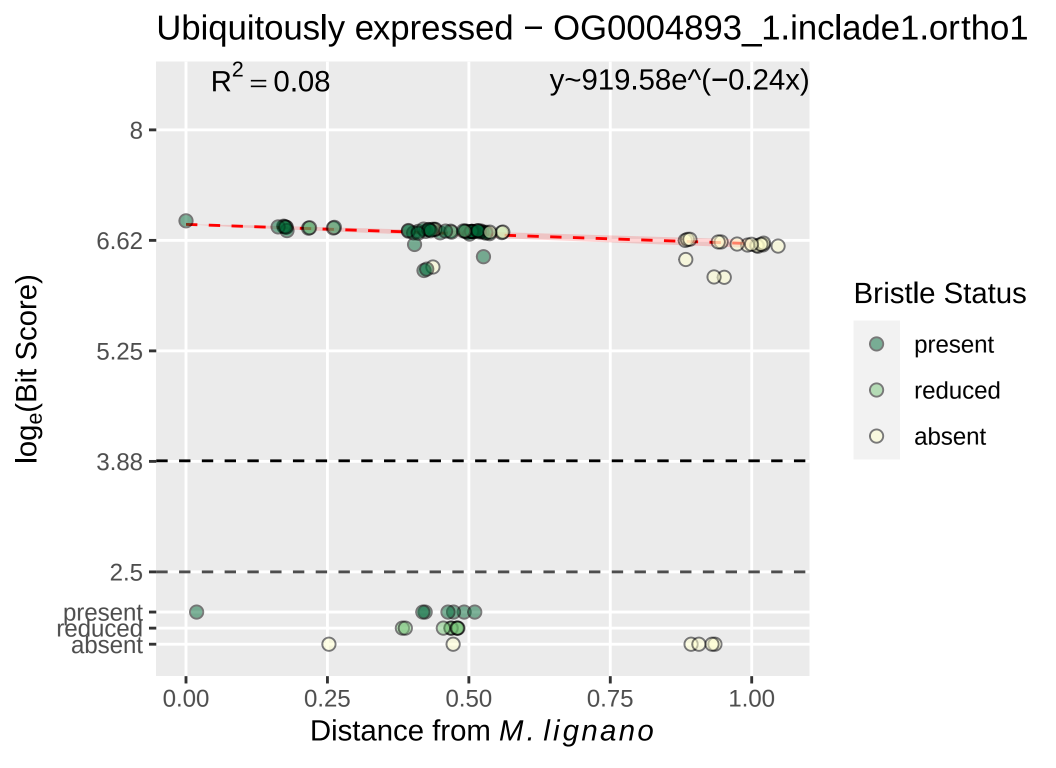

Ubiquitously expressed − OG0004893_1.inclade1.ortho1
$R^2 = 0.08$
y~919.58e^(−0.24x)
8
6.62
5.25
3.88
2.5
present
reduced
absent
0.00
0.25
0.50
0.75
1.00
Distance from M. lignano
log$_e$(Bit Score)
Bristle Status
present
reduced
absent

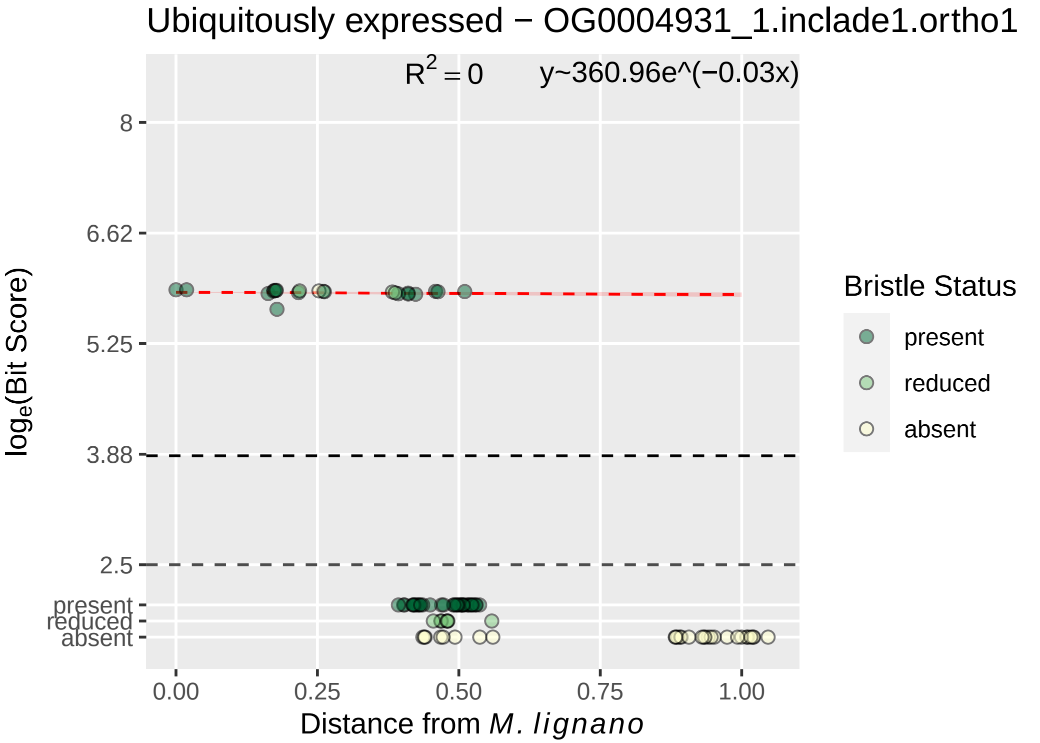

Ubiquitously expressed − OG0004931_1.inclade1.ortho1
R^2 = 0
y~360.96e^(−0.03x)
8
6.62
5.25
3.88
2.5
present
reduced
absent
0.00
0.25
0.50
0.75
1.00
loge(Bit Score)
Distance from M. lignano
Bristle Status
present
reduced
absent

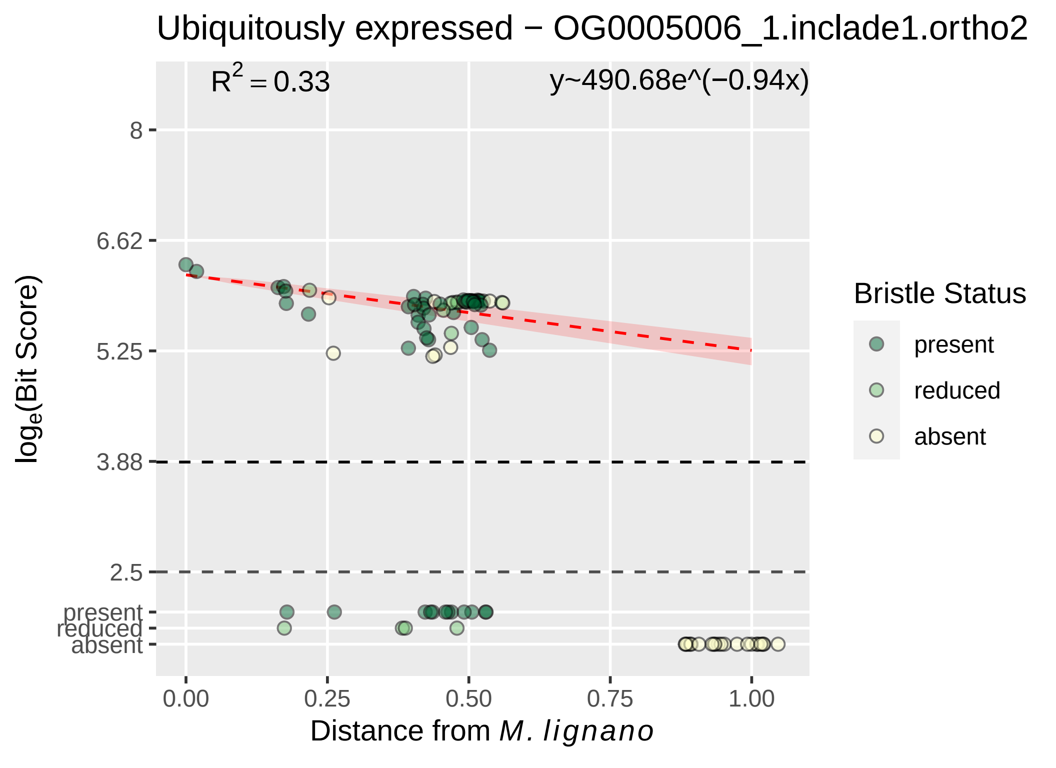
Ubiquitously expressed − OG0005006_1.inclade1.ortho2
R2 = 0.33
y~490.68e^(−0.94x)
8
6.62
5.25
3.88
2.5
present
reduced
absent
0.00
0.25
0.50
0.75
1.00
loge(Bit Score)
Distance from M. lignano
Bristle Status
present
reduced
absent

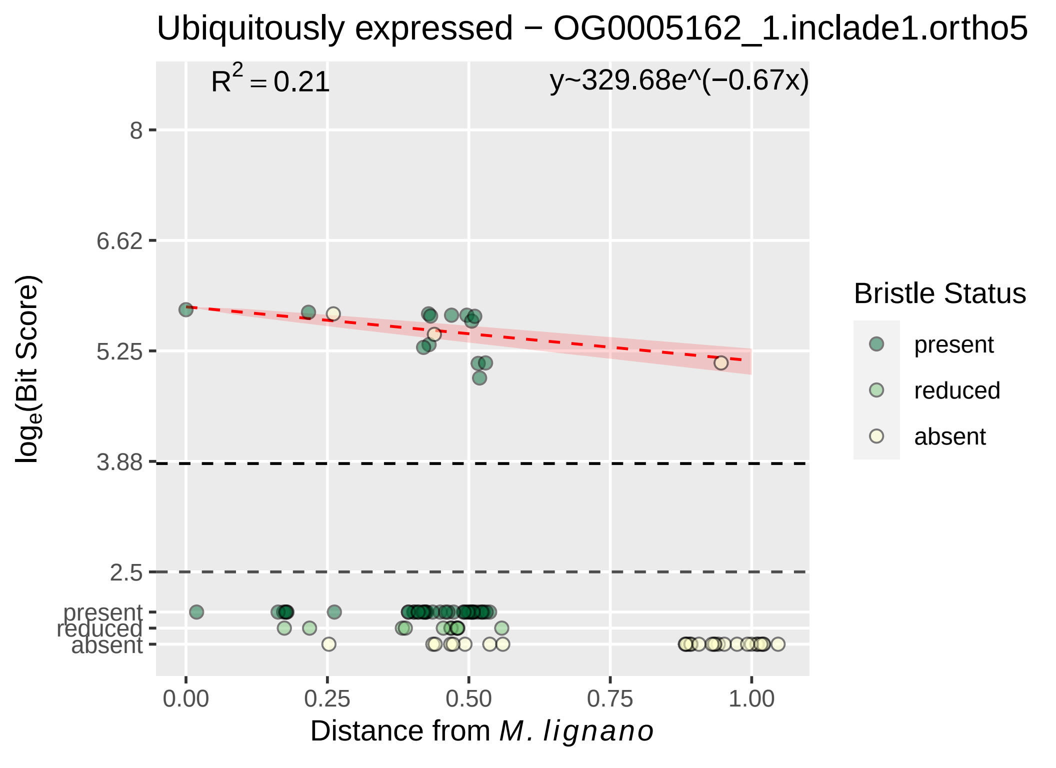

Ubiquitously expressed − OG0005162_1.inclade1.ortho5
R^2 = 0.21
y~329.68e^(−0.67x)
8
6.62
5.25
3.88
2.5
present
reduced
absent
0.00
0.25
0.50
0.75
1.00
log_e(Bit Score)
Distance from M. lignano
Bristle Status
present
reduced
absent

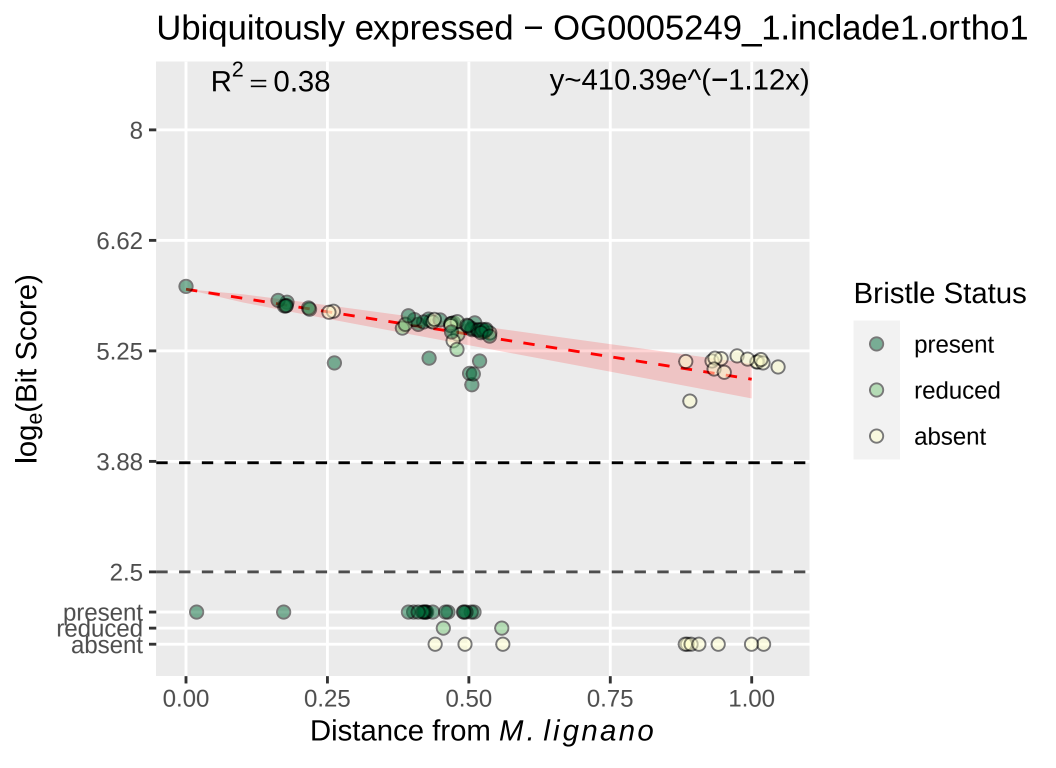

Ubiquitously expressed − OG0005249_1.inclade1.ortho1
R² = 0.38
y~410.39e^(−1.12x)
8
6.62
5.25
3.88
2.5
present
reduced
absent
log_e(Bit Score)
0.00
0.25
0.50
0.75
1.00
Distance from M. lignano
Bristle Status
present
reduced
absent

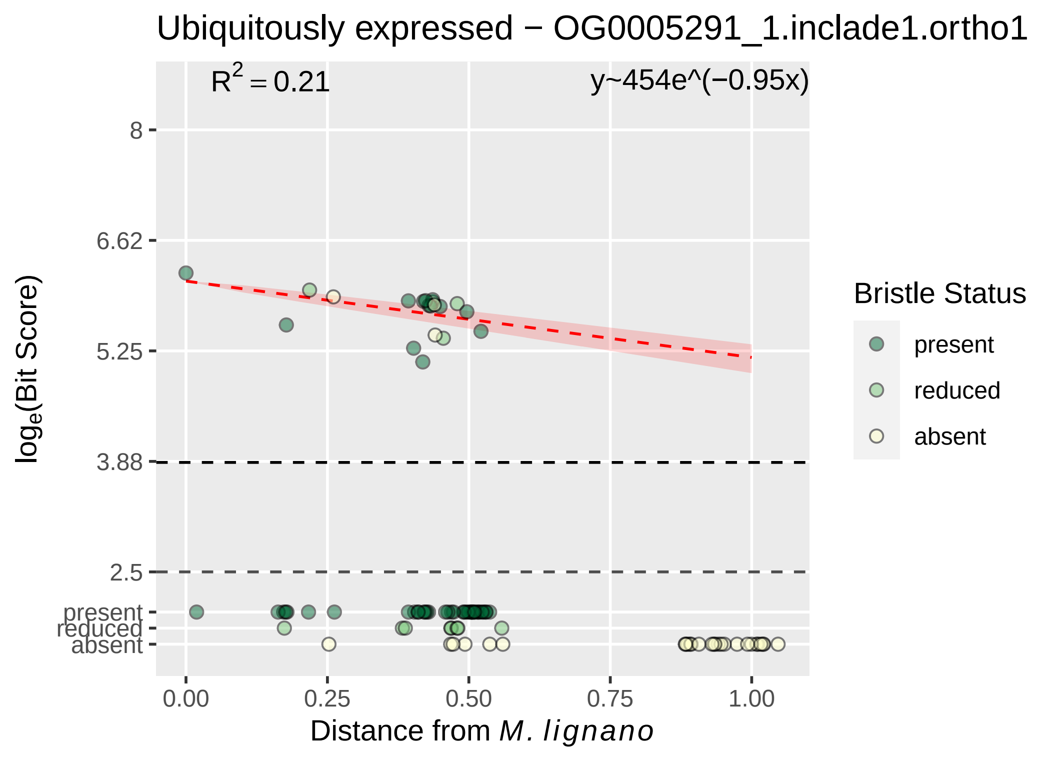

Ubiquitously expressed − OG0005291_1.inclade1.ortho1
R2 = 0.21
y~454e^(−0.95x)
8
6.62
5.25
3.88
2.5
present
reduced
absent
loge(Bit Score)
0.00
0.25
0.50
0.75
1.00
Distance from M. lignano
Bristle Status
present
reduced
absent

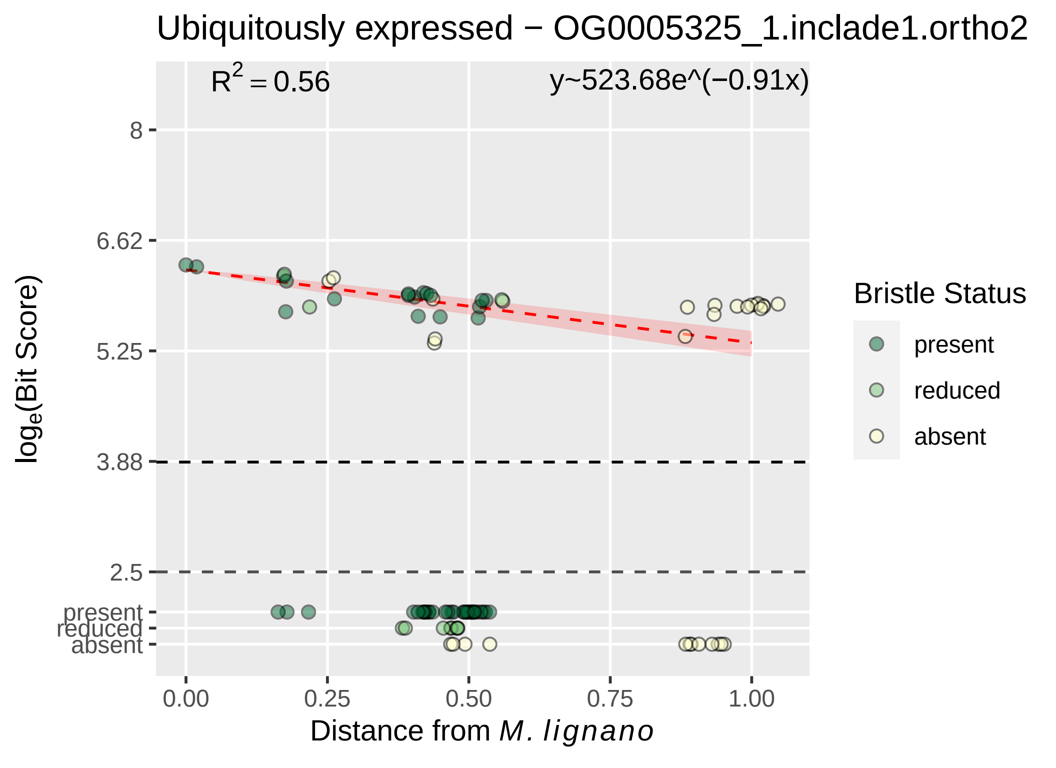

Ubiquitously expressed − OG0005325_1.inclade1.ortho2
$R^2 = 0.56$
y~523.68e^(−0.91x)
8
6.62
5.25
3.88
2.5
present
reduced
absent
0.00
0.25
0.50
0.75
1.00
Distance from M. lignano
$\log_e$(Bit Score)
Bristle Status
present
reduced
absent

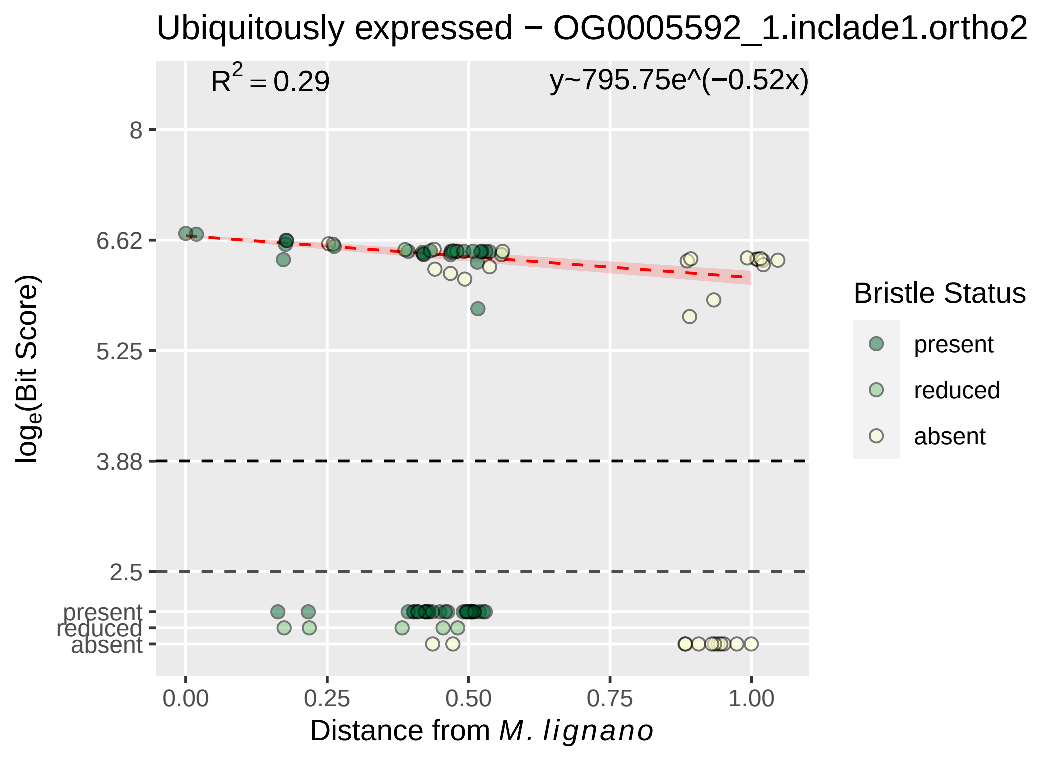

Ubiquitously expressed − OG0005592_1.inclade1.ortho2
R² = 0.29
y~795.75e^(−0.52x)
8
6.62
5.25
3.88
2.5
present
reduced
absent
log_e(Bit Score)
0.00
0.25
0.50
0.75
1.00
Distance from M. lignano
Bristle Status
present
reduced
absent

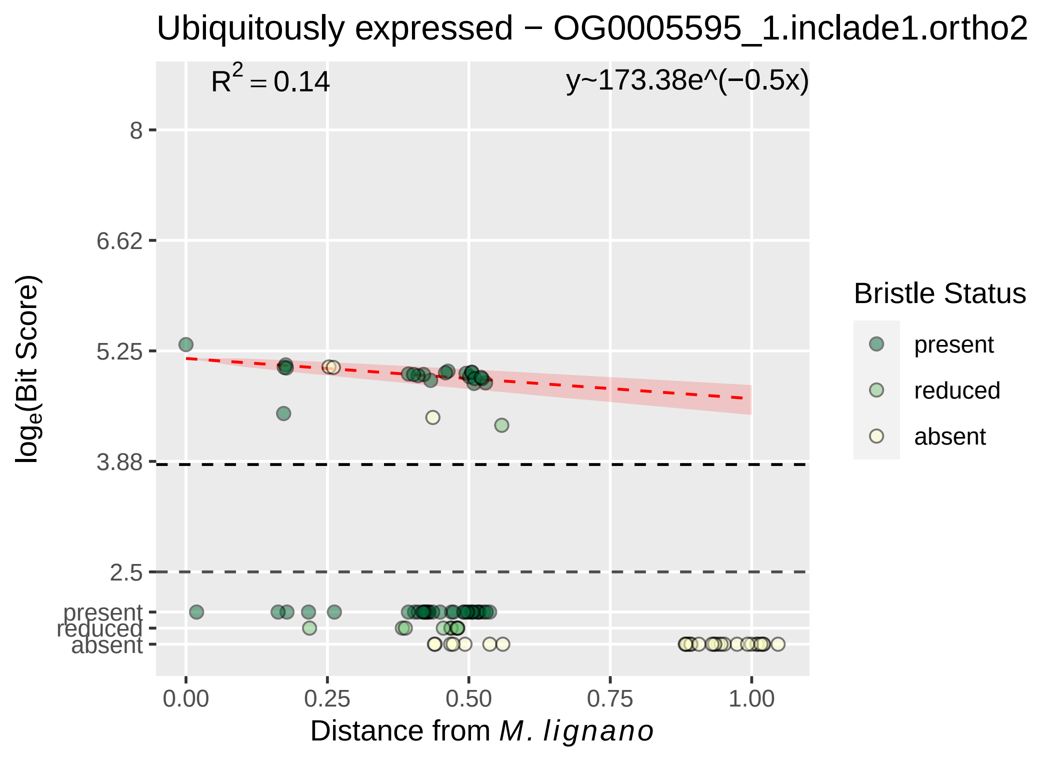

Ubiquitously expressed − OG0005595_1.inclade1.ortho2
R^2 = 0.14
y~173.38e^(−0.5x)
8
6.62
5.25
3.88
2.5
present
reduced
absent
loge(Bit Score)
0.00
0.25
0.50
0.75
1.00
Distance from M. lignano
Bristle Status
present
reduced
absent

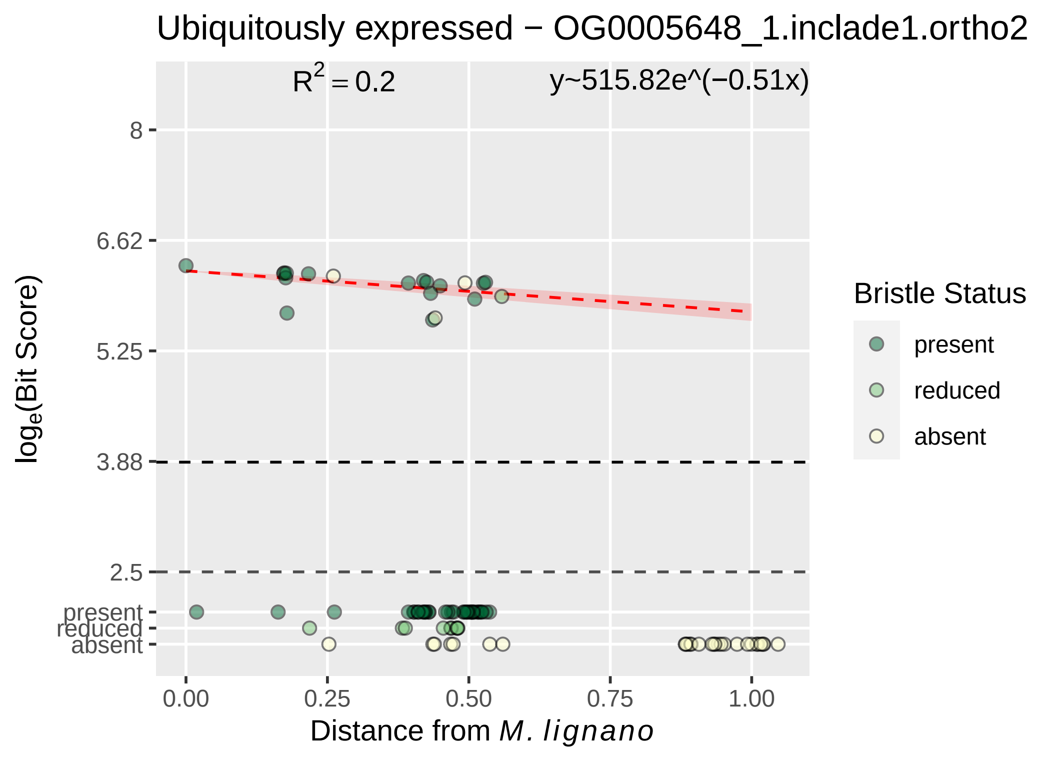
Ubiquitously expressed − OG0005648_1.inclade1.ortho2
R^2 = 0.2
y~515.82e^(−0.51x)
8
6.62
5.25
3.88
2.5
present
reduced
absent
log_e(Bit Score)
0.00
0.25
0.50
0.75
1.00
Distance from M. lignano
Bristle Status
present
reduced
absent

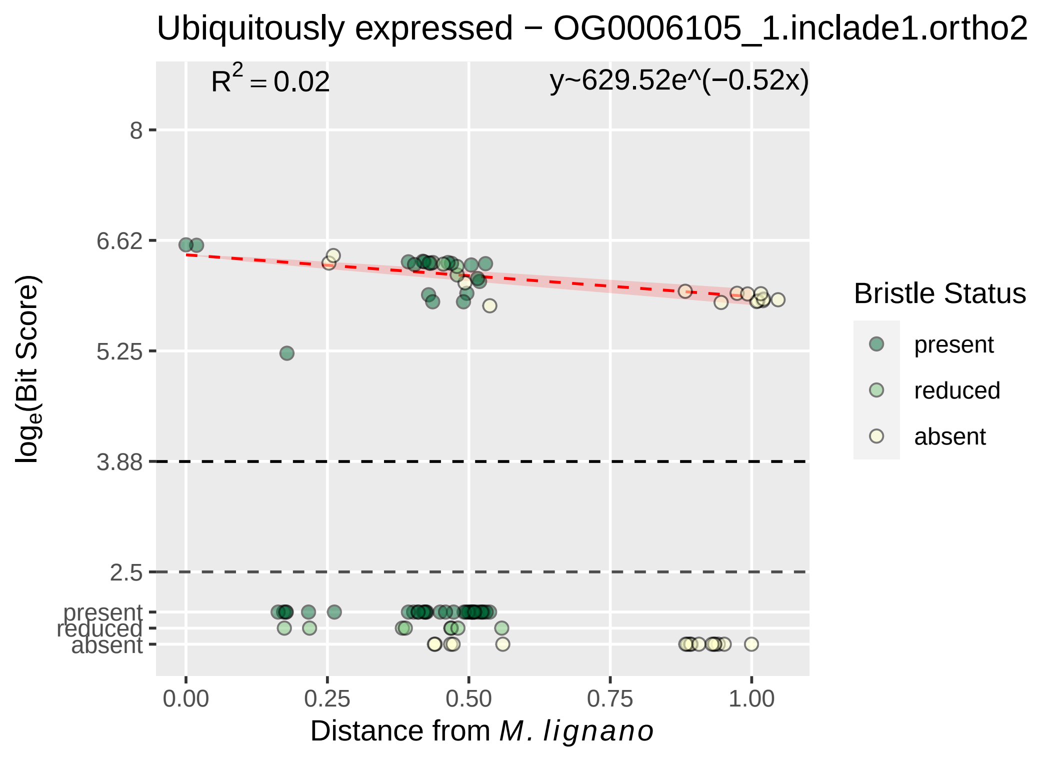

Ubiquitously expressed − OG0006105_1.inclade1.ortho2
$R^2 = 0.02$
y~629.52e^(−0.52x)
8
6.62
5.25
3.88
2.5
present
reduced
absent
0.00
0.25
0.50
0.75
1.00
$\log_e$(Bit Score)
Distance from *M. lignano*
Bristle Status
present
reduced
absent

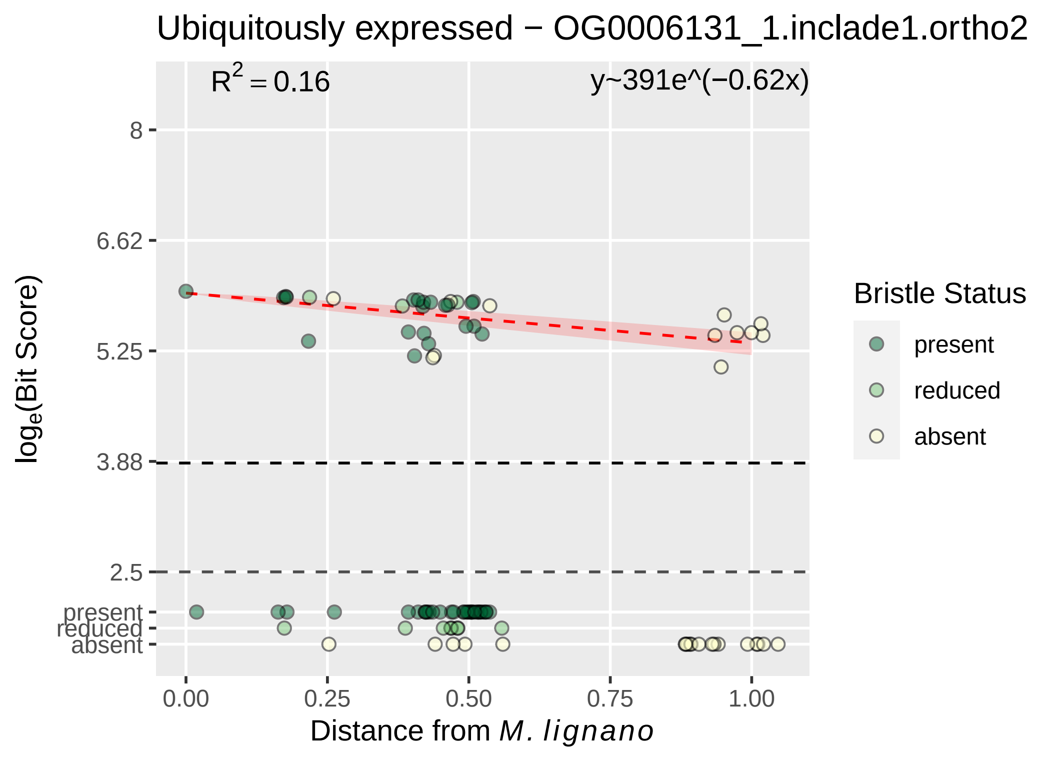
Ubiquitously expressed − OG0006131_1.inclade1.ortho2
R² = 0.16
y~391e^(−0.62x)
8
6.62
5.25
3.88
2.5
present
reduced
absent
log_e(Bit Score)
0.00
0.25
0.50
0.75
1.00
Distance from M. lignano
Bristle Status
present
reduced
absent

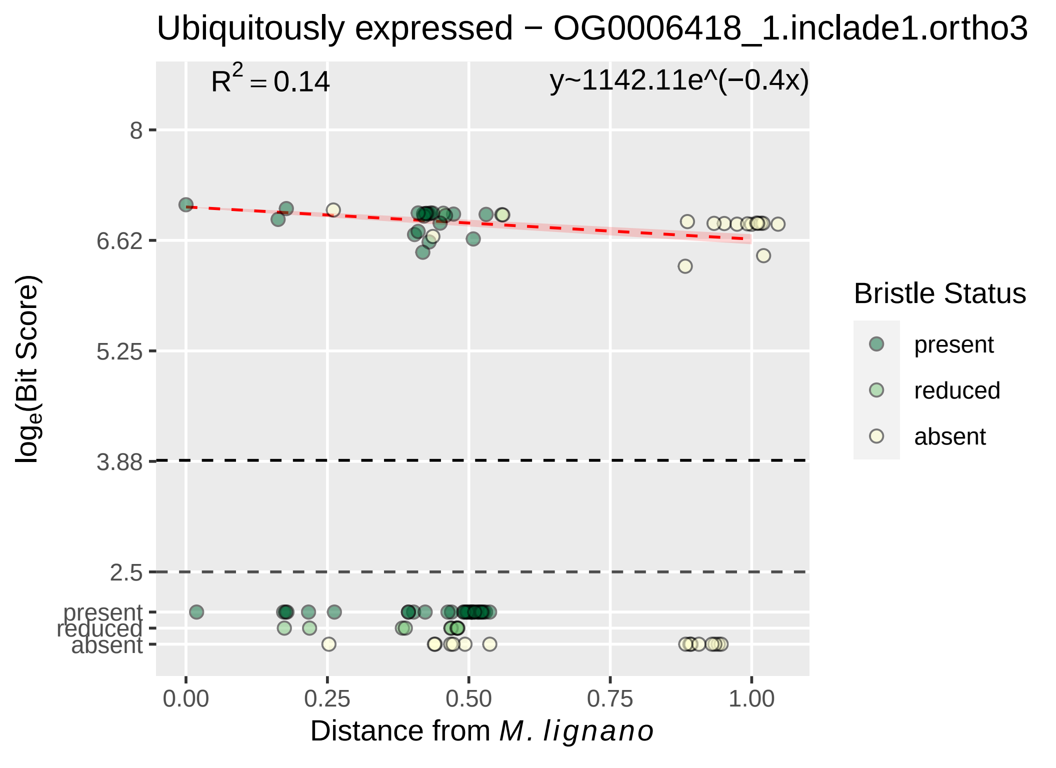

Ubiquitously expressed − OG0006418_1.inclade1.ortho3
R^2 = 0.14
y~1142.11e^(−0.4x)
8
6.62
5.25
3.88
2.5
present
reduced
absent
0.00
0.25
0.50
0.75
1.00
Distance from M. lignano
log_e(Bit Score)
Bristle Status
present
reduced
absent

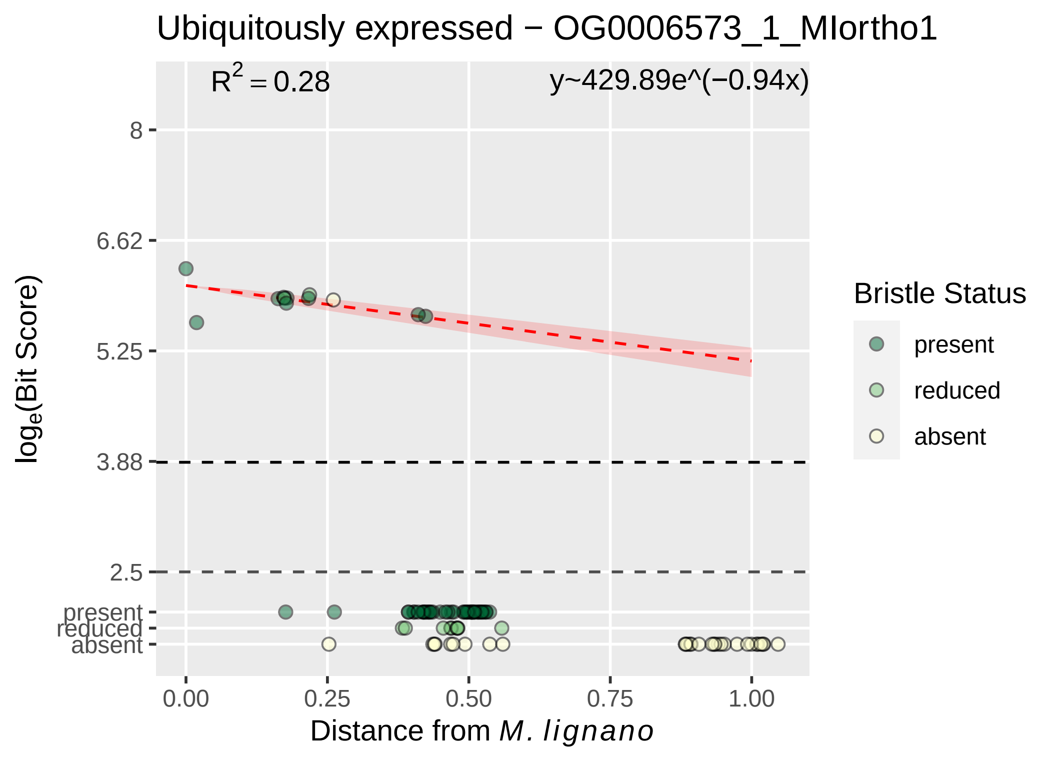
Ubiquitously expressed − OG0006573_1_MIortho1
$R^2 = 0.28$
y~429.89e^(−0.94x)
8
6.62
5.25
3.88
2.5
present
reduced
absent
0.00
0.25
0.50
0.75
1.00
$\log_e$(Bit Score)
Distance from *M. lignano*
Bristle Status
present
reduced
absent

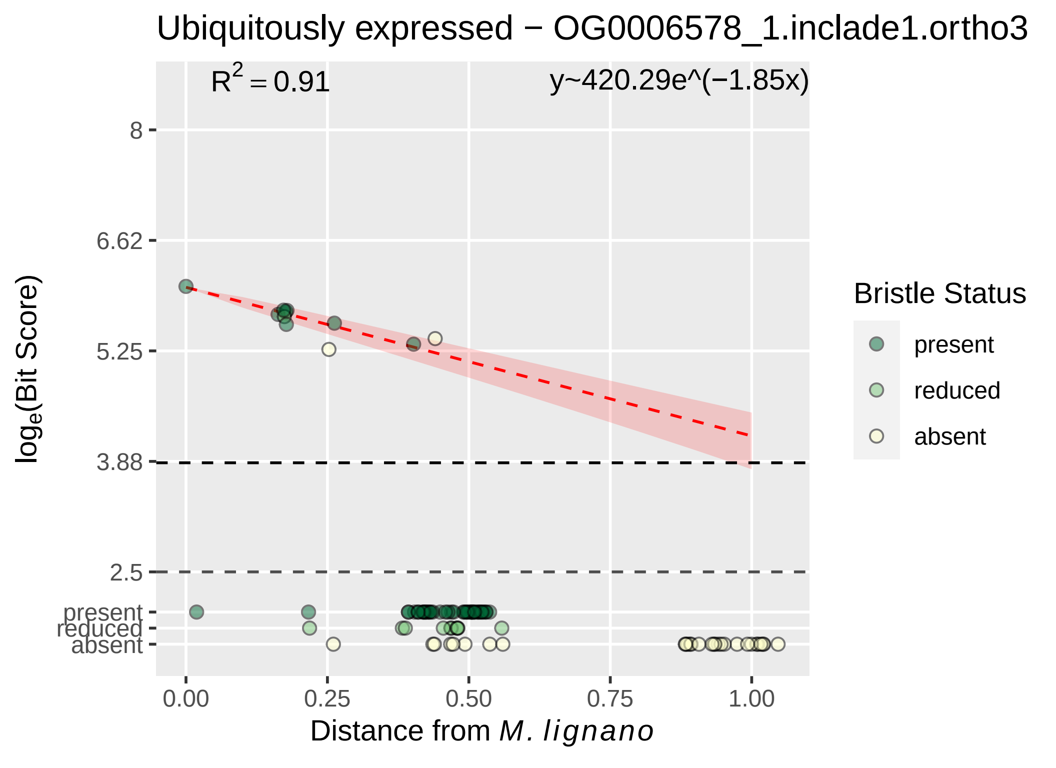

Ubiquitously expressed − OG0006578_1.inclade1.ortho3
R^2 = 0.91
y~420.29e^(−1.85x)
8
6.62
5.25
3.88
2.5
present
reduced
absent
log_e(Bit Score)
0.00
0.25
0.50
0.75
1.00
Distance from M. lignano
Bristle Status
present
reduced
absent

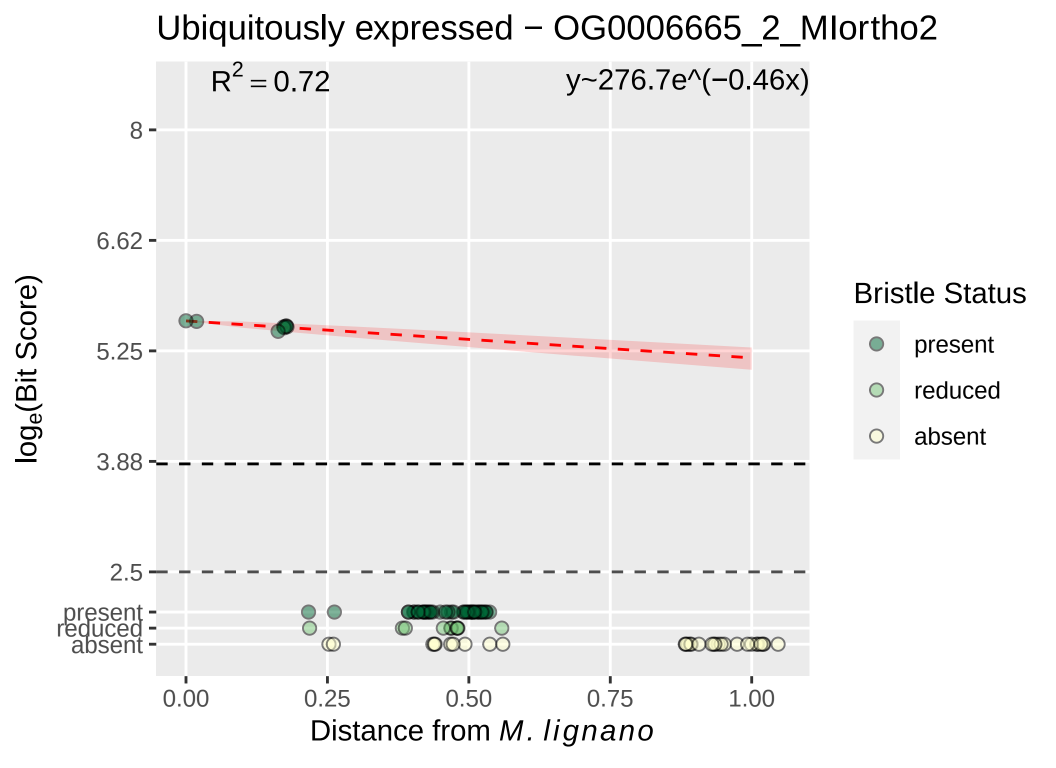
Ubiquitously expressed − OG0006665_2_MIortho2
R² = 0.72
y~276.7e^(−0.46x)
8
6.62
5.25
3.88
2.5
present
reduced
absent
log_e(Bit Score)
0.00
0.25
0.50
0.75
1.00
Distance from M. lignano
Bristle Status
present
reduced
absent

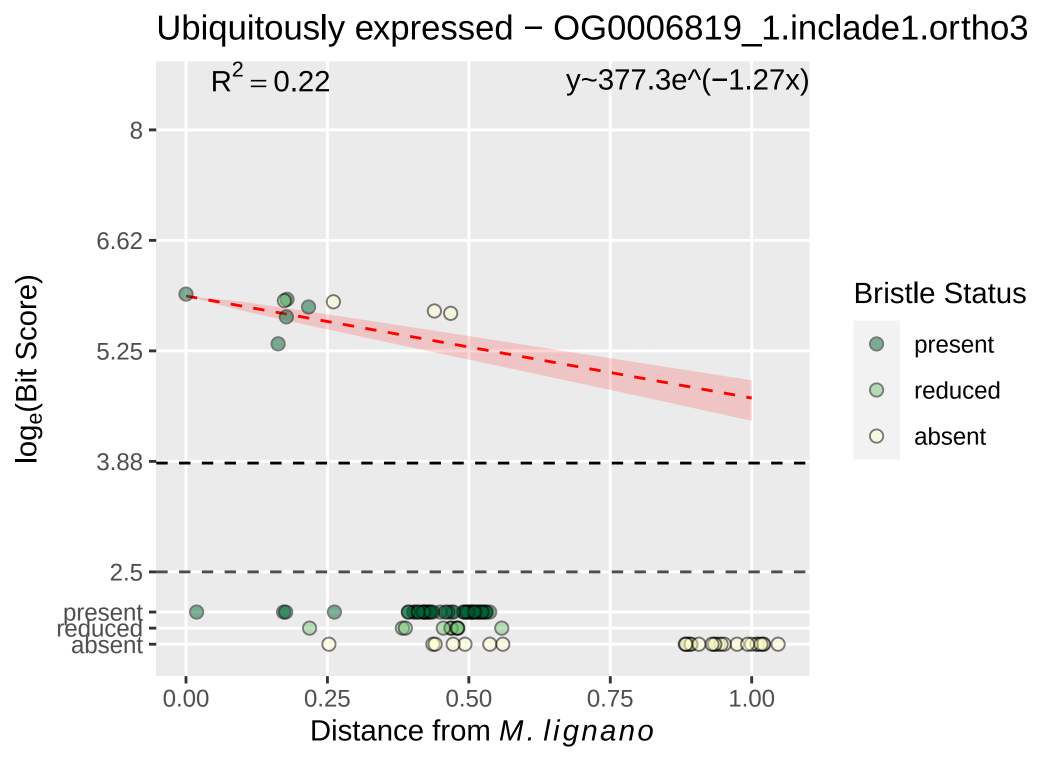

Ubiquitously expressed − OG0006819_1.inclade1.ortho3
R2 = 0.22
y~377.3e^(−1.27x)
8
6.62
5.25
3.88
2.5
present
reduced
absent
loge(Bit Score)
0.00
0.25
0.50
0.75
1.00
Distance from M. lignano
Bristle Status
present
reduced
absent

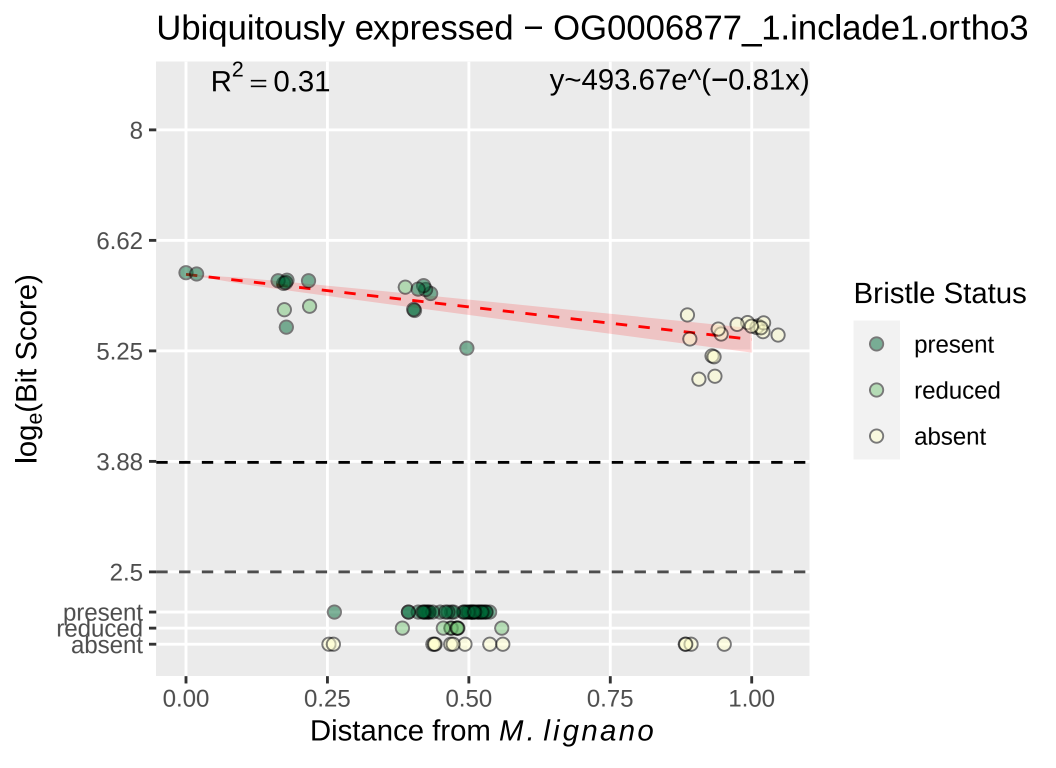

Ubiquitously expressed − OG0006877_1.inclade1.ortho3
R^2 = 0.31
y~493.67e^(−0.81x)
8
6.62
5.25
3.88
2.5
present
reduced
absent
log_e(Bit Score)
0.00
0.25
0.50
0.75
1.00
Distance from M. lignano
Bristle Status
present
reduced
absent

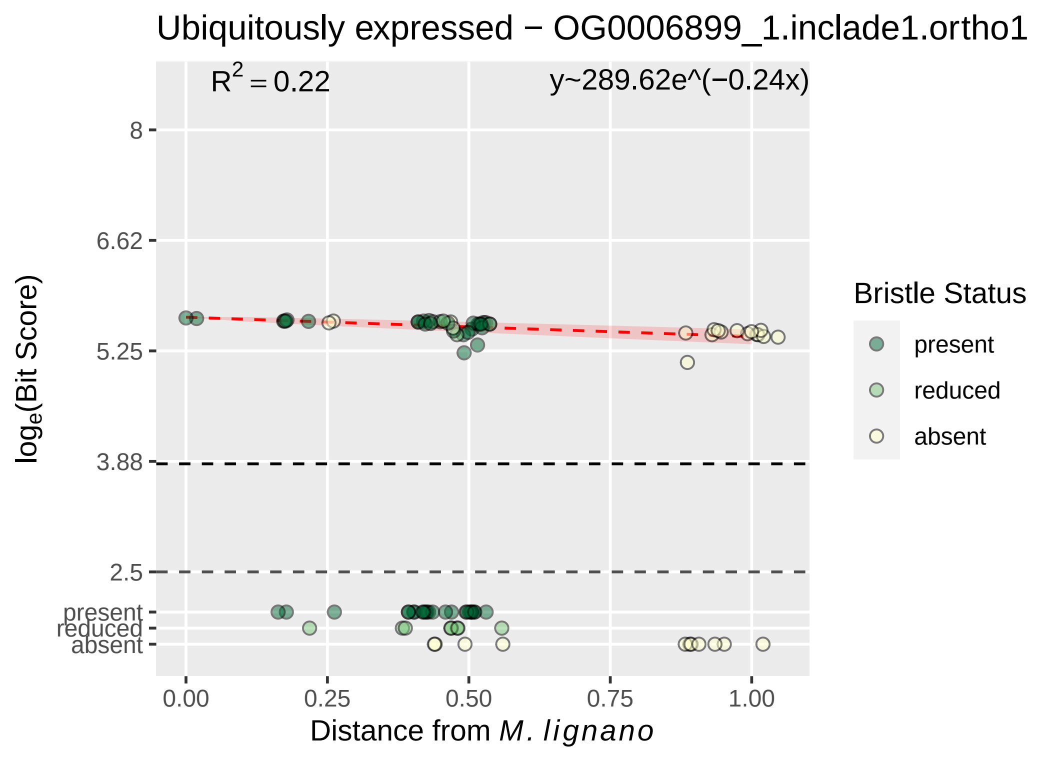

Ubiquitously expressed − OG0006899_1.inclade1.ortho1
$R^2 = 0.22$
y~289.62e^(−0.24x)
8
6.62
5.25
3.88
2.5
present
reduced
absent
$log_e$(Bit Score)
0.00
0.25
0.50
0.75
1.00
Distance from *M. lignano*
Bristle Status
present
reduced
absent

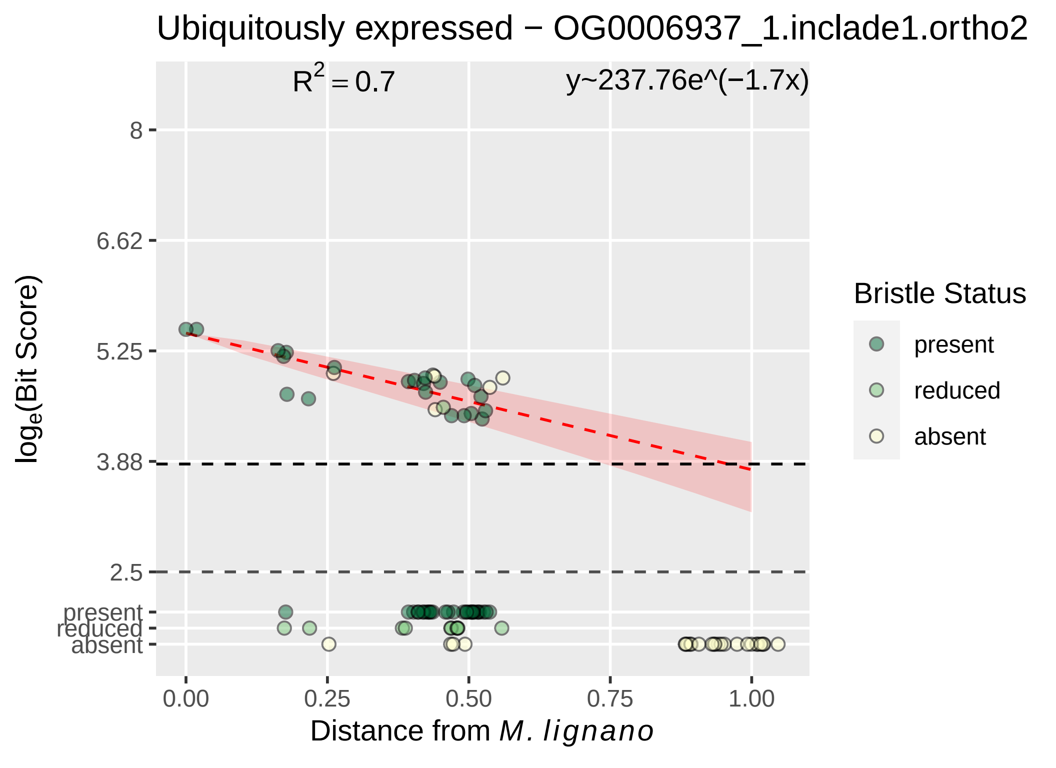

Ubiquitously expressed − OG0006937_1.inclade1.ortho2
$R^2 = 0.7$
y~237.76e^(−1.7x)
8
6.62
5.25
3.88
2.5
present
reduced
absent
$\log_e$(Bit Score)
0.00
0.25
0.50
0.75
1.00
Distance from *M. lignano*
Bristle Status
present
reduced
absent

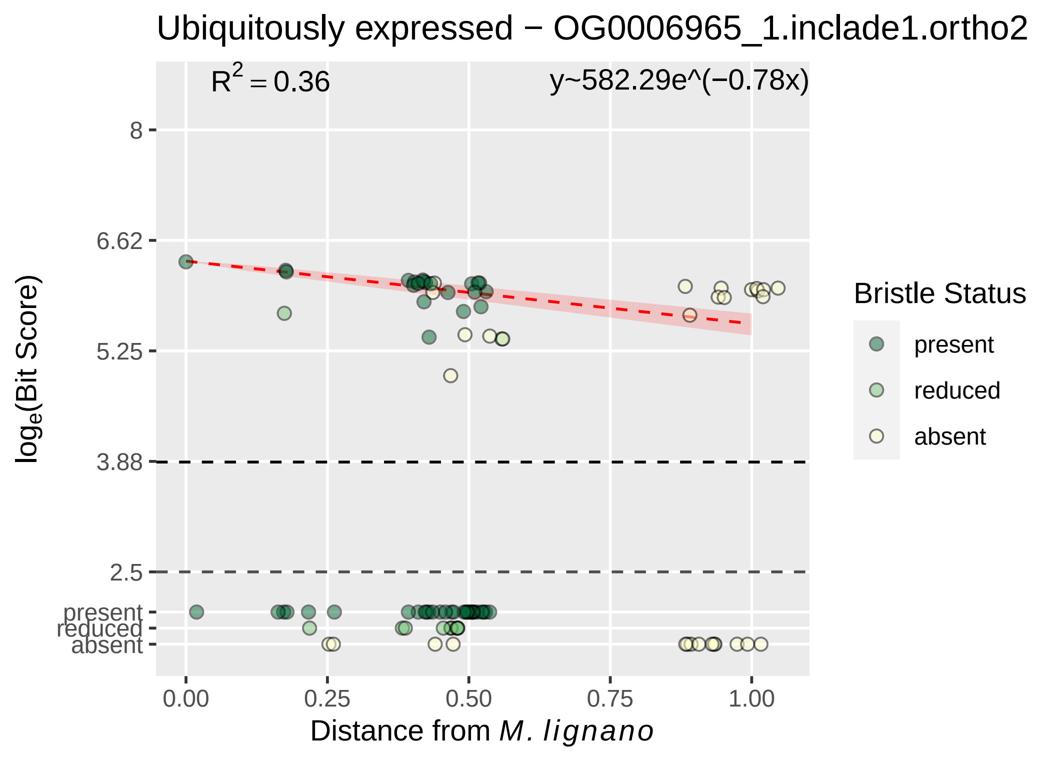

Ubiquitously expressed − OG0006965_1.inclade1.ortho2
$R^2 = 0.36$
y~582.29e^(−0.78x)
8
6.62
5.25
3.88
2.5
present
reduced
absent
0.00
0.25
0.50
0.75
1.00
$\log_e$(Bit Score)
Distance from *M. lignano*
Bristle Status
present
reduced
absent

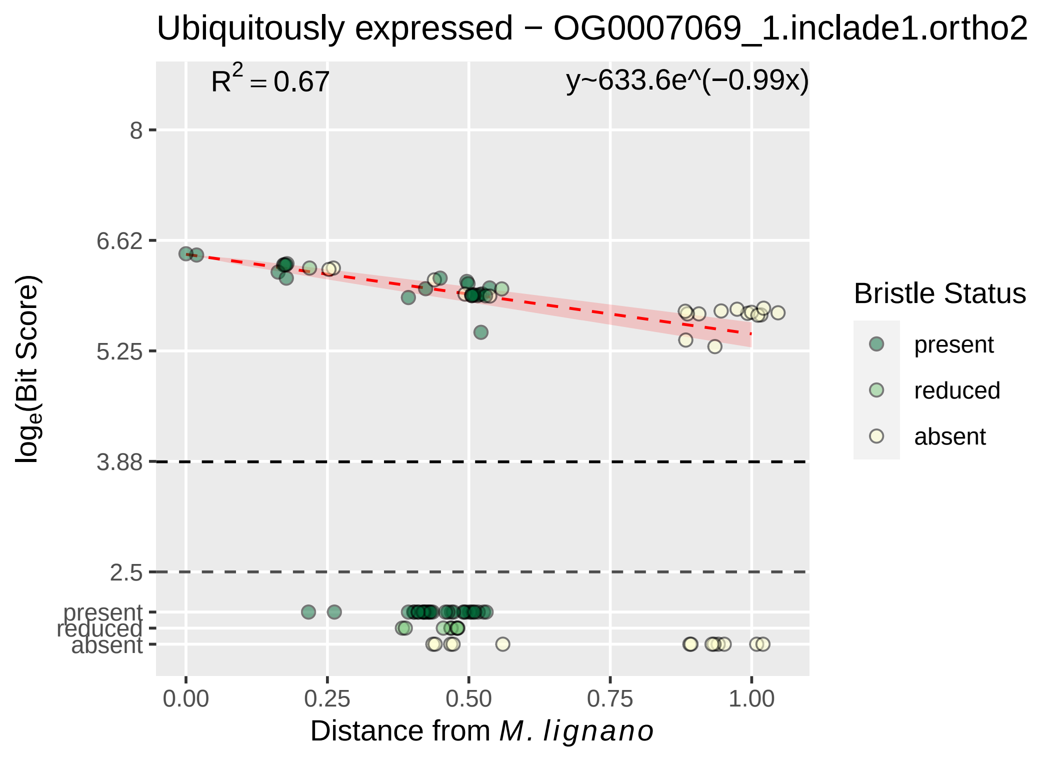
Ubiquitously expressed − OG0007069_1.inclade1.ortho2
R^2 = 0.67
y~633.6e^(−0.99x)
8
6.62
5.25
3.88
2.5
present
reduced
absent
0.00
0.25
0.50
0.75
1.00
loge(Bit Score)
Distance from M. lignano
Bristle Status
present
reduced
absent

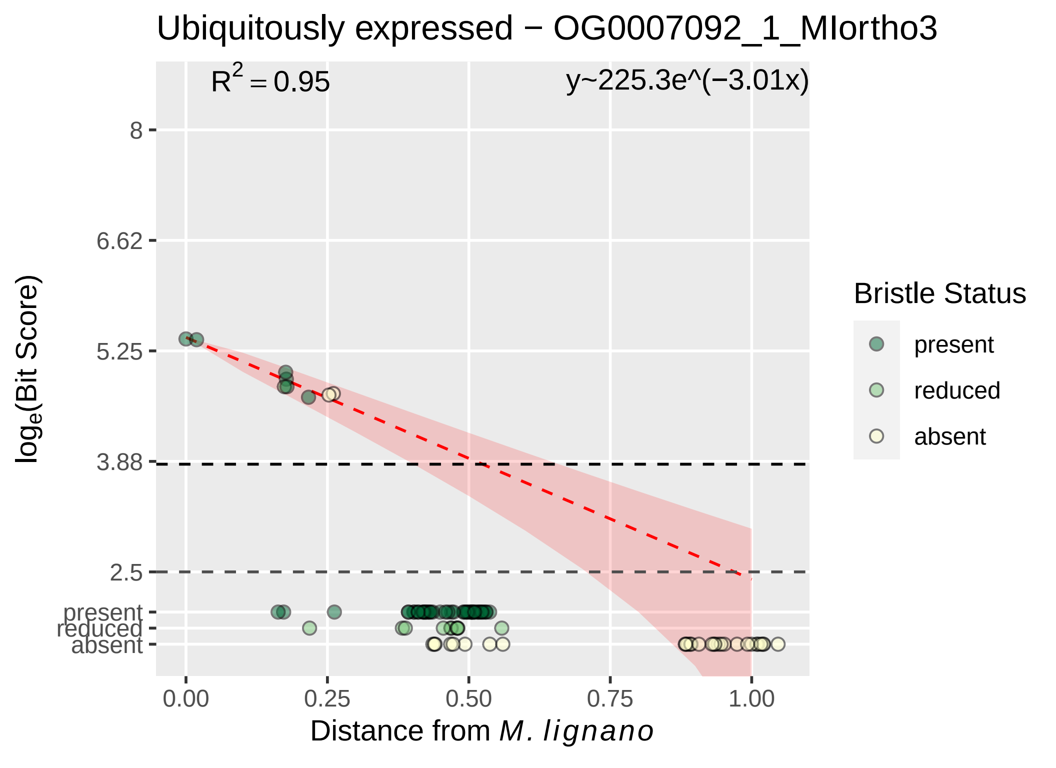

Ubiquitously expressed − OG0007092_1_MIortho3
R^2 = 0.95
y~225.3e^(−3.01x)
8
6.62
5.25
3.88
2.5
present
reduced
absent
loge(Bit Score)
0.00
0.25
0.50
0.75
1.00
Distance from M. lignano
Bristle Status
present
reduced
absent

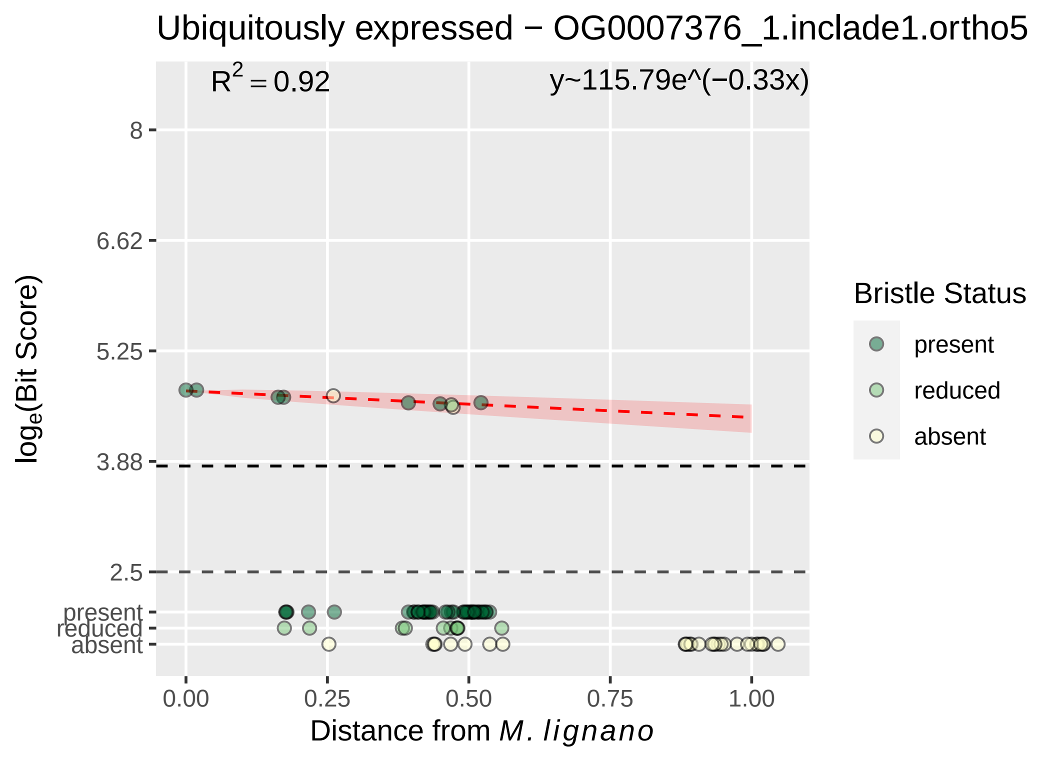

Ubiquitously expressed − OG0007376_1.inclade1.ortho5
R² = 0.92
y~115.79e^(−0.33x)
8
6.62
5.25
3.88
2.5
present
reduced
absent
log_e(Bit Score)
0.00
0.25
0.50
0.75
1.00
Distance from M. lignano
Bristle Status
present
reduced
absent

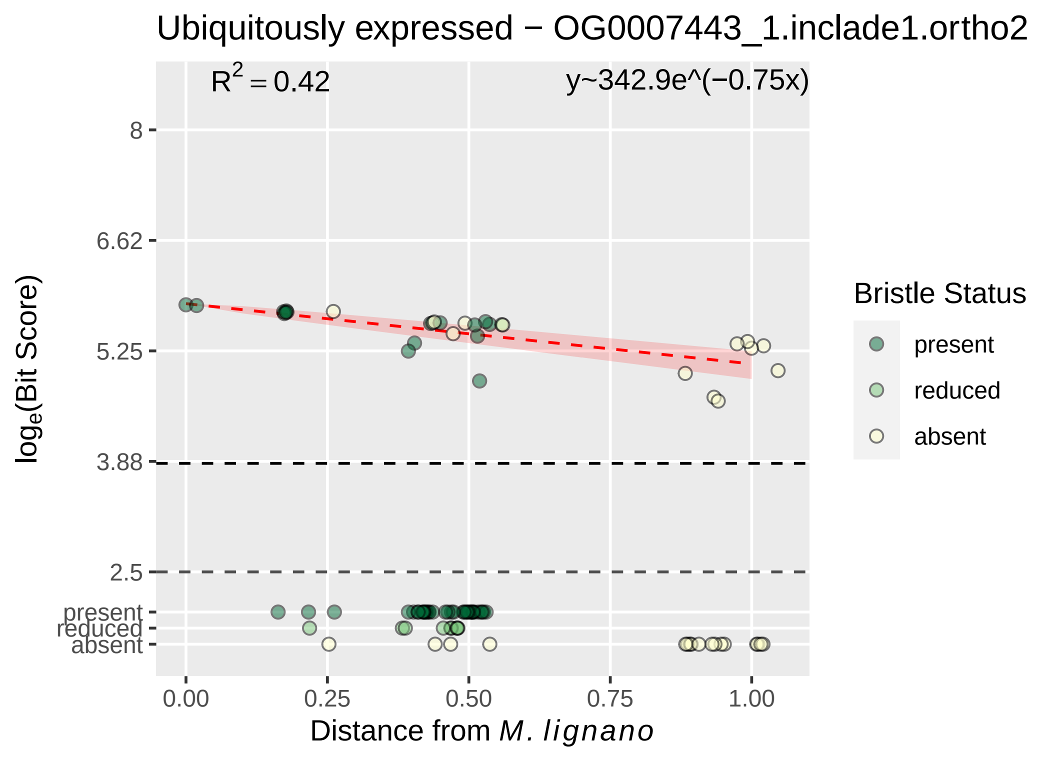
Ubiquitously expressed − OG0007443_1.inclade1.ortho2
R^2 = 0.42
y~342.9e^(−0.75x)
log_e(Bit Score)
8
6.62
5.25
3.88
2.5
present
reduced
absent
0.00
0.25
0.50
0.75
1.00
Distance from M. lignano
Bristle Status
present
reduced
absent

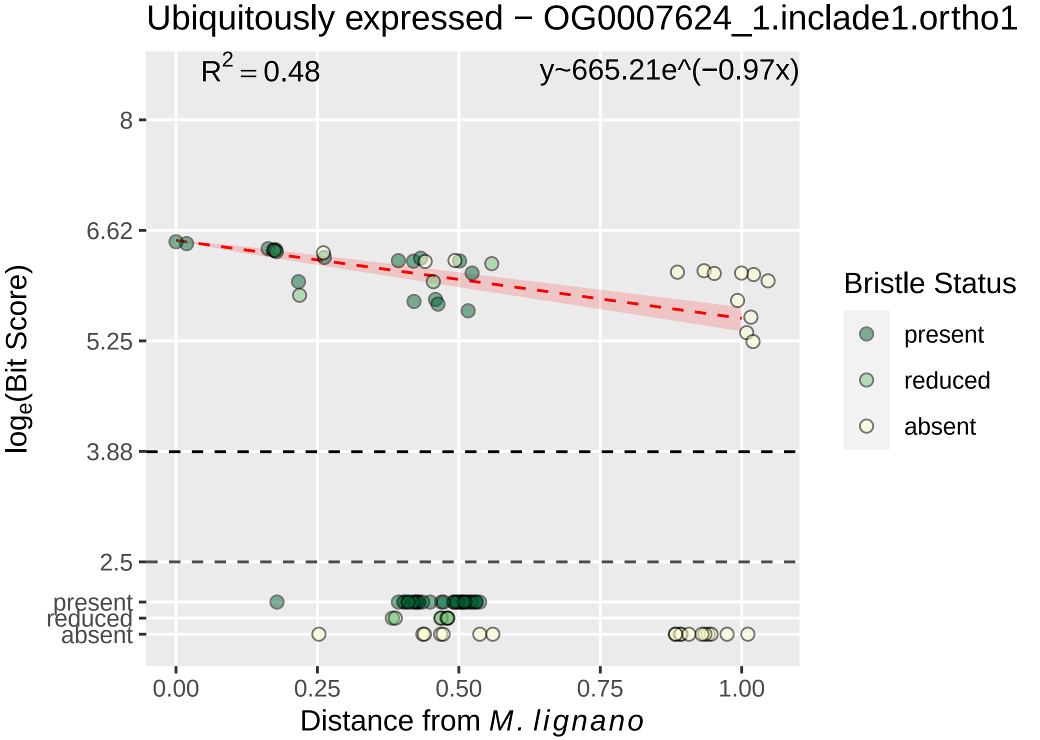

Ubiquitously expressed − OG0007624_1.inclade1.ortho1
R² = 0.48
y~665.21e^(−0.97x)
8
6.62
5.25
3.88
2.5
present
reduced
absent
0.00
0.25
0.50
0.75
1.00
loge(Bit Score)
Distance from M. lignano
Bristle Status
present
reduced
absent

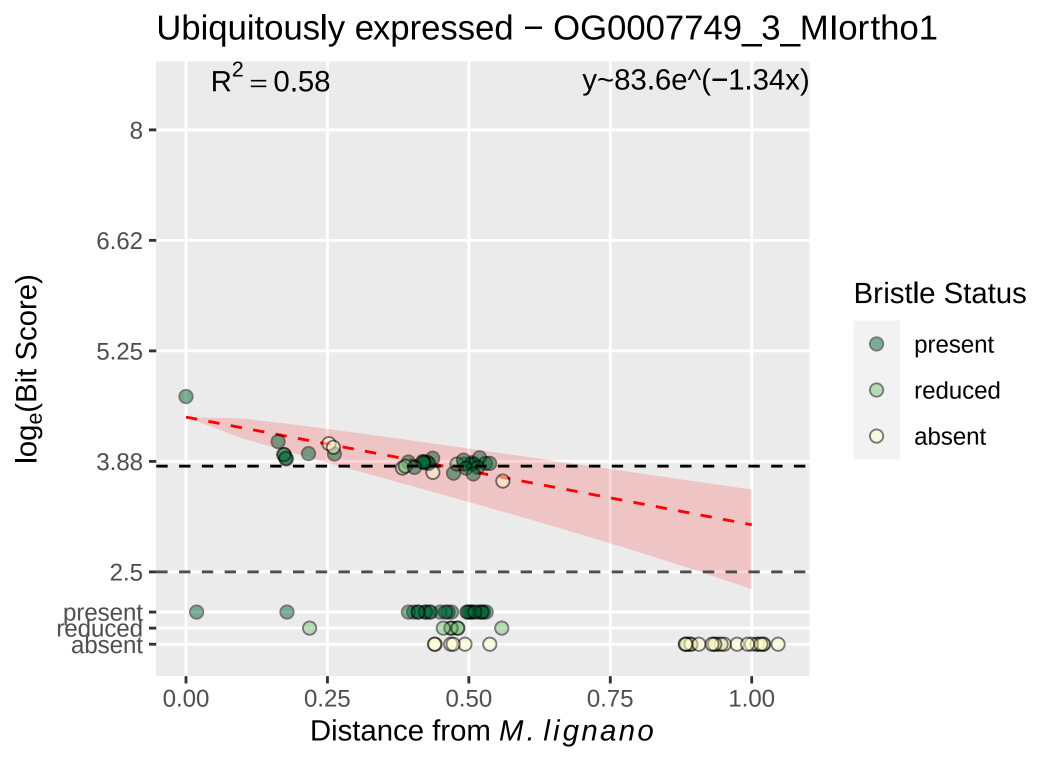

Ubiquitously expressed − OG0007749_3_MIortho1
$R^2 = 0.58$
y~83.6e^(−1.34x)
8
6.62
5.25
3.88
2.5
present
reduced
absent
$\log_e$(Bit Score)
0.00
0.25
0.50
0.75
1.00
Distance from *M. lignano*
Bristle Status
present
reduced
absent

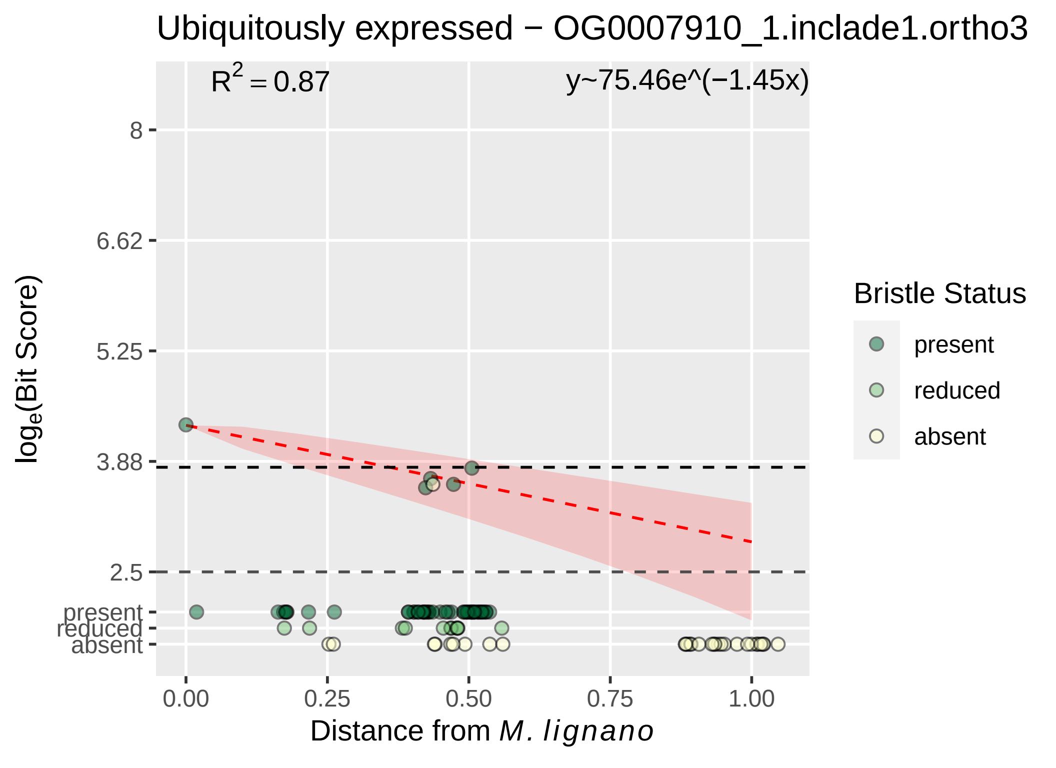

Ubiquitously expressed − OG0007910_1.inclade1.ortho3
$R^2 = 0.87$
y~75.46e^(−1.45x)
8
6.62
5.25
3.88
2.5
present
reduced
absent
0.00
0.25
0.50
0.75
1.00
Distance from M. lignano
log$_e$(Bit Score)
Bristle Status
present
reduced
absent

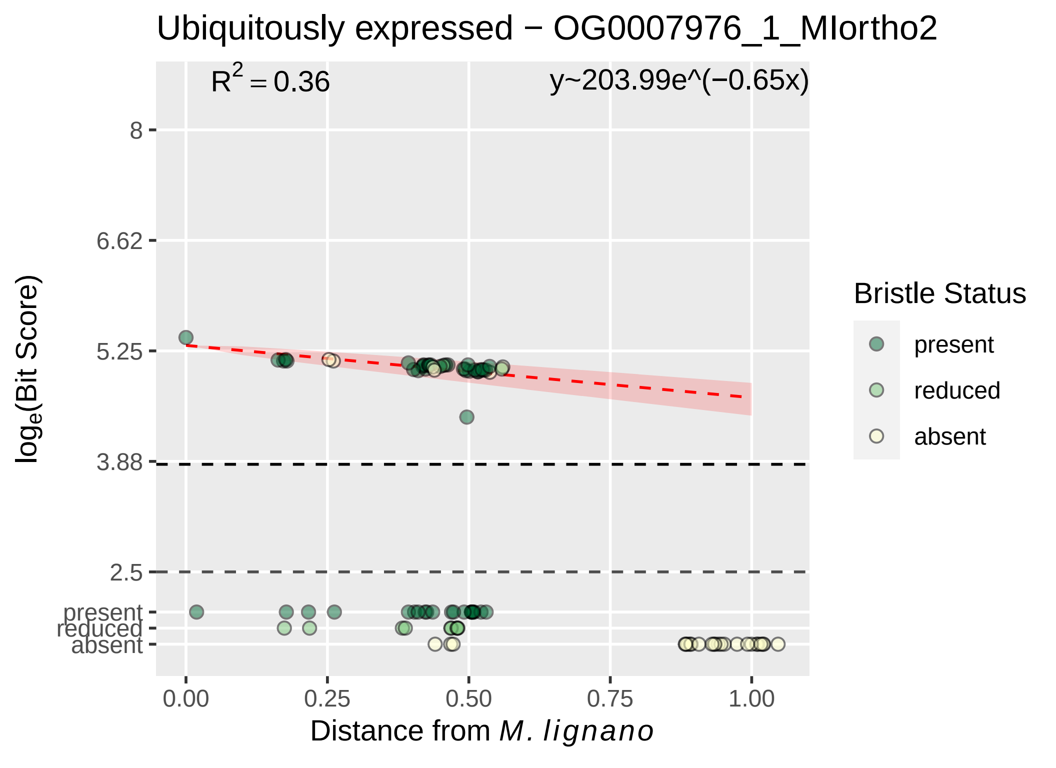

Ubiquitously expressed − OG0007976_1_MIortho2
R2 = 0.36
y~203.99e^(−0.65x)
8
6.62
5.25
3.88
2.5
present
reduced
absent
loge(Bit Score)
0.00
0.25
0.50
0.75
1.00
Distance from M. lignano
Bristle Status
present
reduced
absent

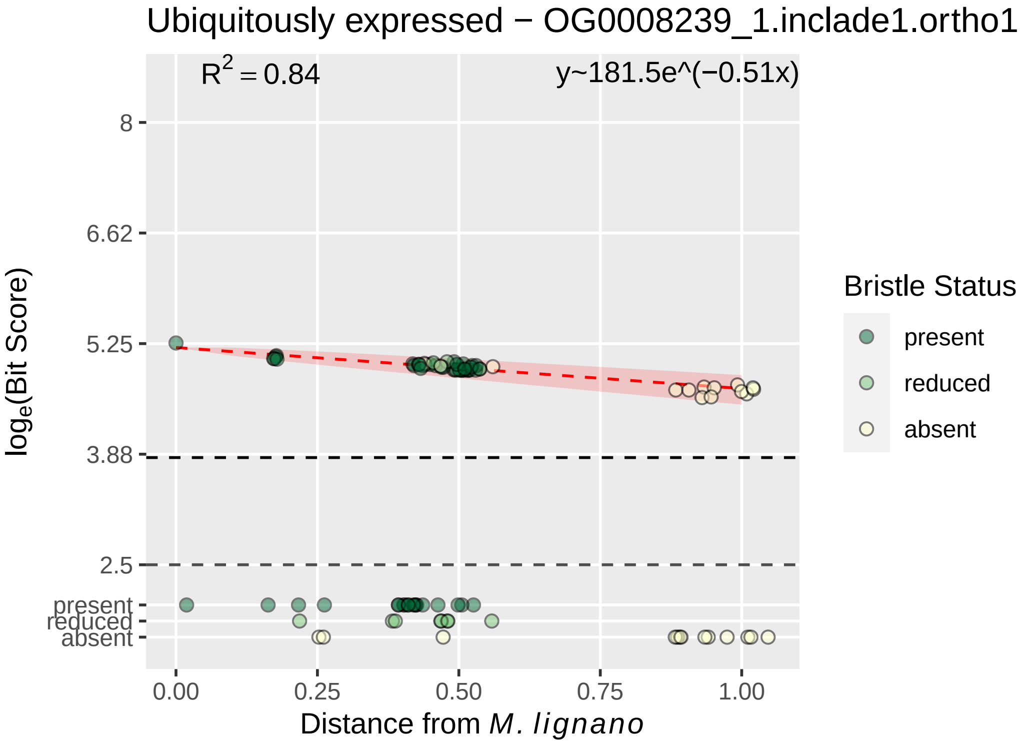

Ubiquitously expressed − OG0008239_1.inclade1.ortho1
$R^2 = 0.84$
y~181.5e^(−0.51x)
8
6.62
5.25
3.88
2.5
present
reduced
absent
$\log_e$(Bit Score)
0.00
0.25
0.50
0.75
1.00
Distance from *M. lignano*
Bristle Status
present
reduced
absent

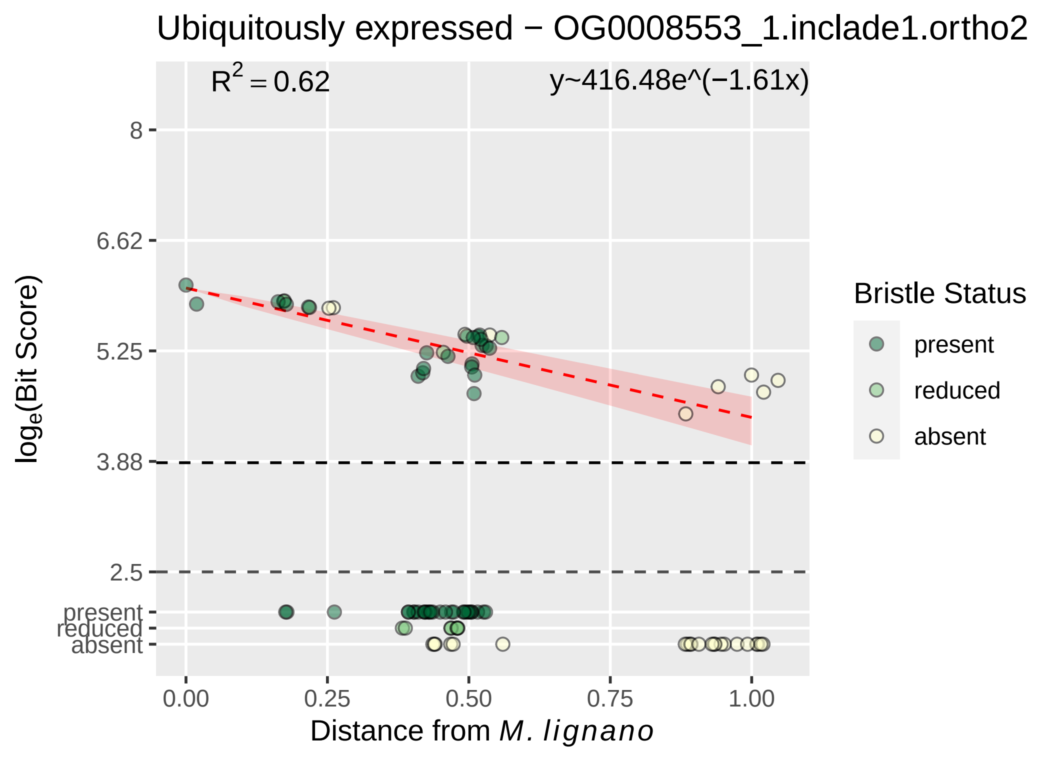

Ubiquitously expressed − OG0008553_1.inclade1.ortho2
$R^2 = 0.62$
y~416.48e^(−1.61x)
8
6.62
5.25
3.88
2.5
present
reduced
absent
0.00
0.25
0.50
0.75
1.00
Distance from M. lignano
loge(Bit Score)
Bristle Status
present
reduced
absent

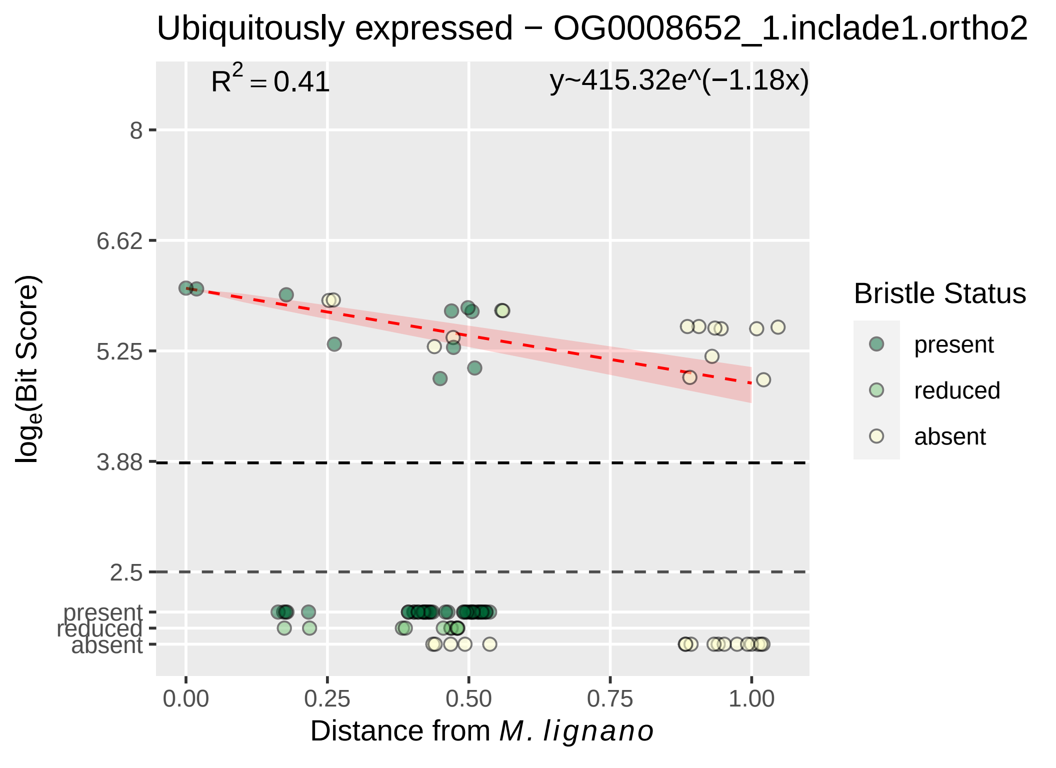

Ubiquitously expressed − OG0008652_1.inclade1.ortho2
R^2 = 0.41
y~415.32e^(−1.18x)
8
6.62
5.25
3.88
2.5
present
reduced
absent
log_e(Bit Score)
0.00
0.25
0.50
0.75
1.00
Distance from M. lignano
Bristle Status
present
reduced
absent

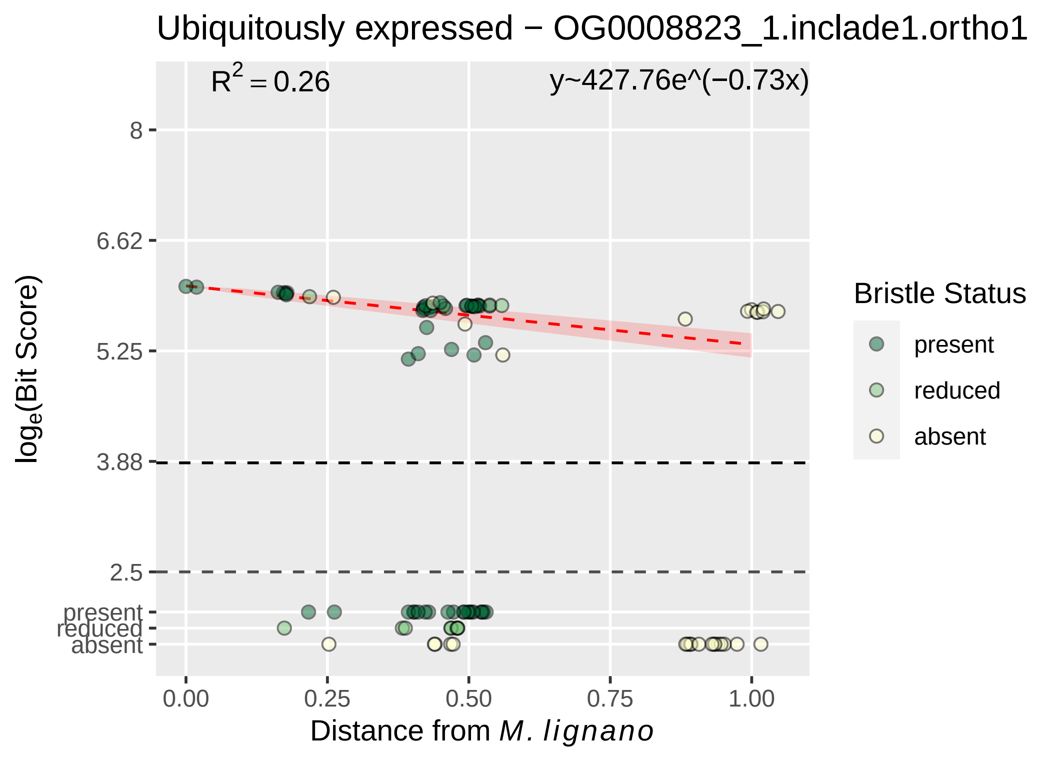

Ubiquitously expressed − OG0008823_1.inclade1.ortho1
R2 = 0.26
y~427.76e^(−0.73x)
8
6.62
5.25
3.88
2.5
present
reduced
absent
loge(Bit Score)
0.00
0.25
0.50
0.75
1.00
Distance from M. lignano
Bristle Status
present
reduced
absent

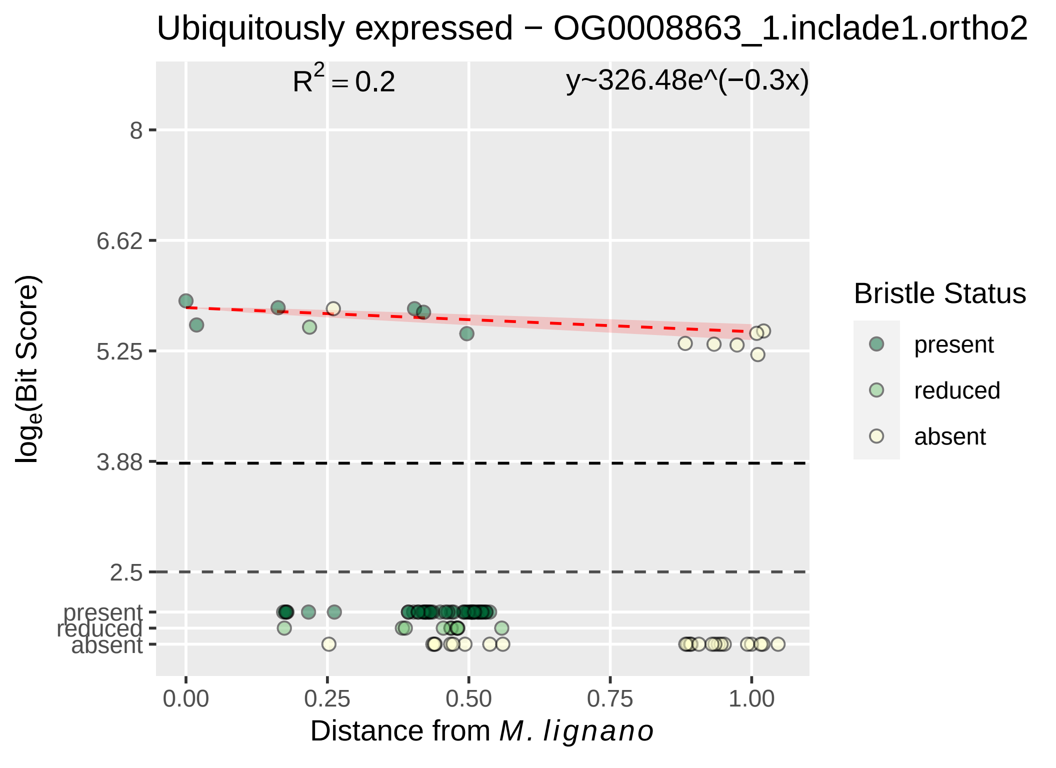
Ubiquitously expressed − OG0008863_1.inclade1.ortho2
R2 = 0.2
y~326.48e^(−0.3x)
8
6.62
5.25
3.88
2.5
present
reduced
absent
loge(Bit Score)
0.00
0.25
0.50
0.75
1.00
Distance from M. lignano
Bristle Status
present
reduced
absent

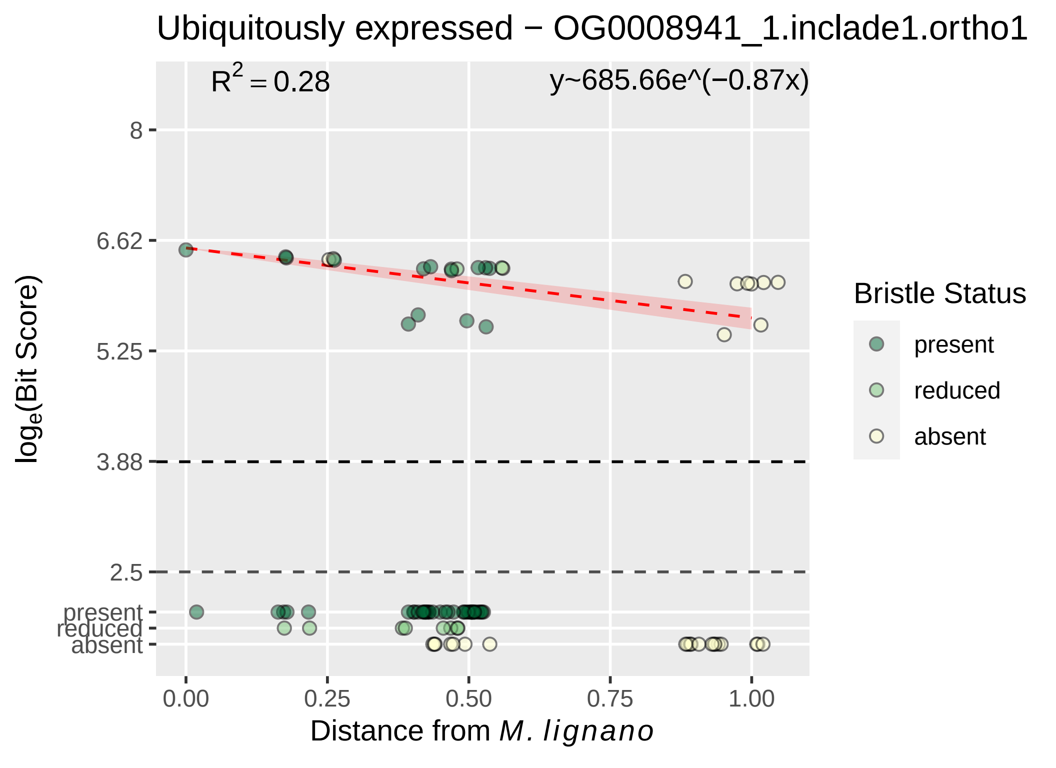

Ubiquitously expressed − OG0008941_1.inclade1.ortho1
$R^2 = 0.28$
y~685.66e^(−0.87x)
8
6.62
5.25
3.88
2.5
present
reduced
absent
log_e(Bit Score)
0.00
0.25
0.50
0.75
1.00
Distance from M. lignano
Bristle Status
present
reduced
absent

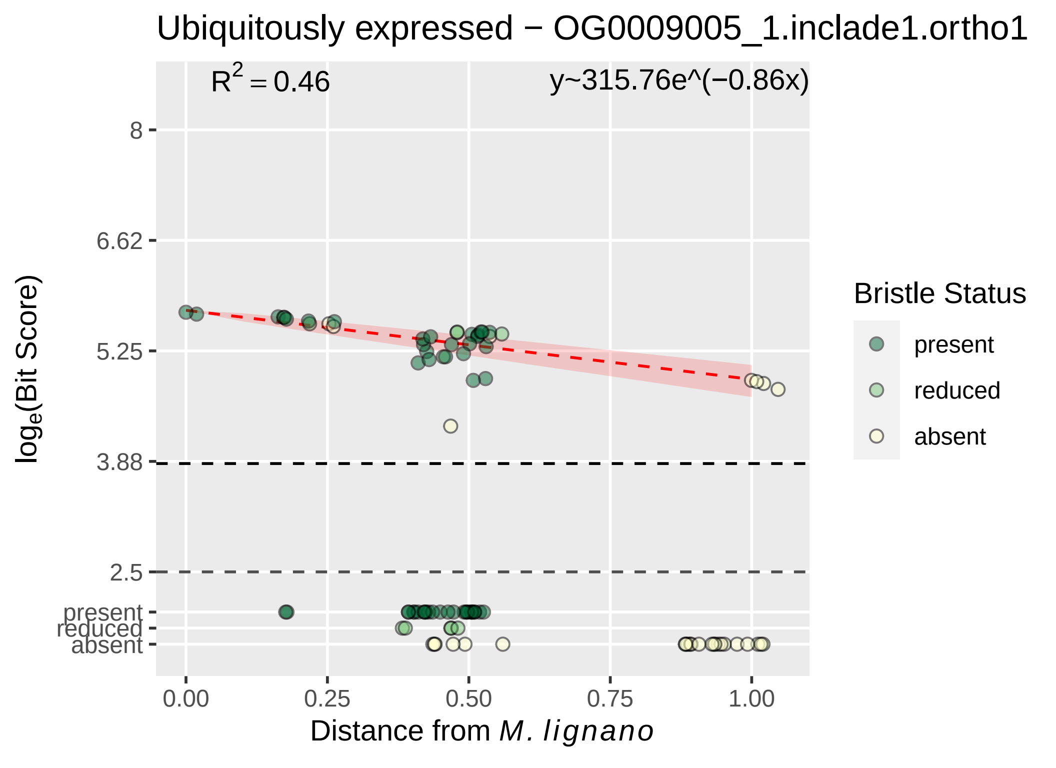
Ubiquitously expressed − OG0009005_1.inclade1.ortho1
$R^2 = 0.46$
y~315.76e^(−0.86x)
8
6.62
5.25
3.88
2.5
present
reduced
absent
$log_e$(Bit Score)
0.00
0.25
0.50
0.75
1.00
Distance from *M. lignano*
Bristle Status
present
reduced
absent

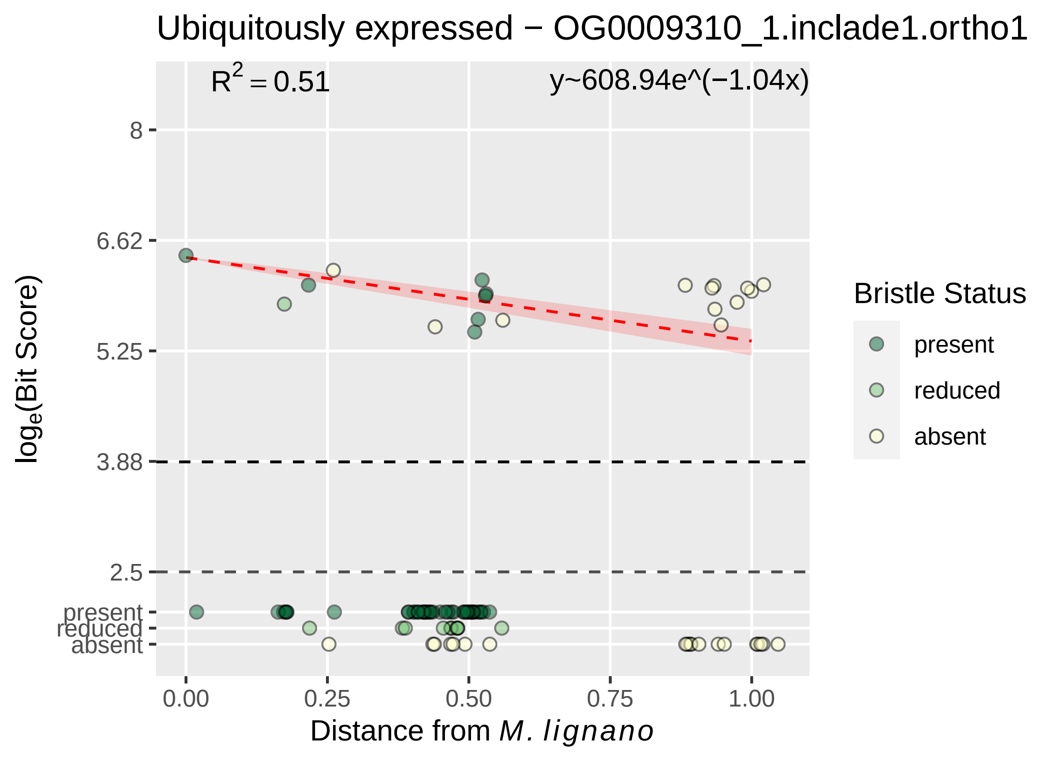

Ubiquitously expressed − OG0009310_1.inclade1.ortho1
R^2 = 0.51
y~608.94e^(−1.04x)
8
6.62
5.25
3.88
2.5
present
reduced
absent
log_e(Bit Score)
0.00
0.25
0.50
0.75
1.00
Distance from M. lignano
Bristle Status
present
reduced
absent

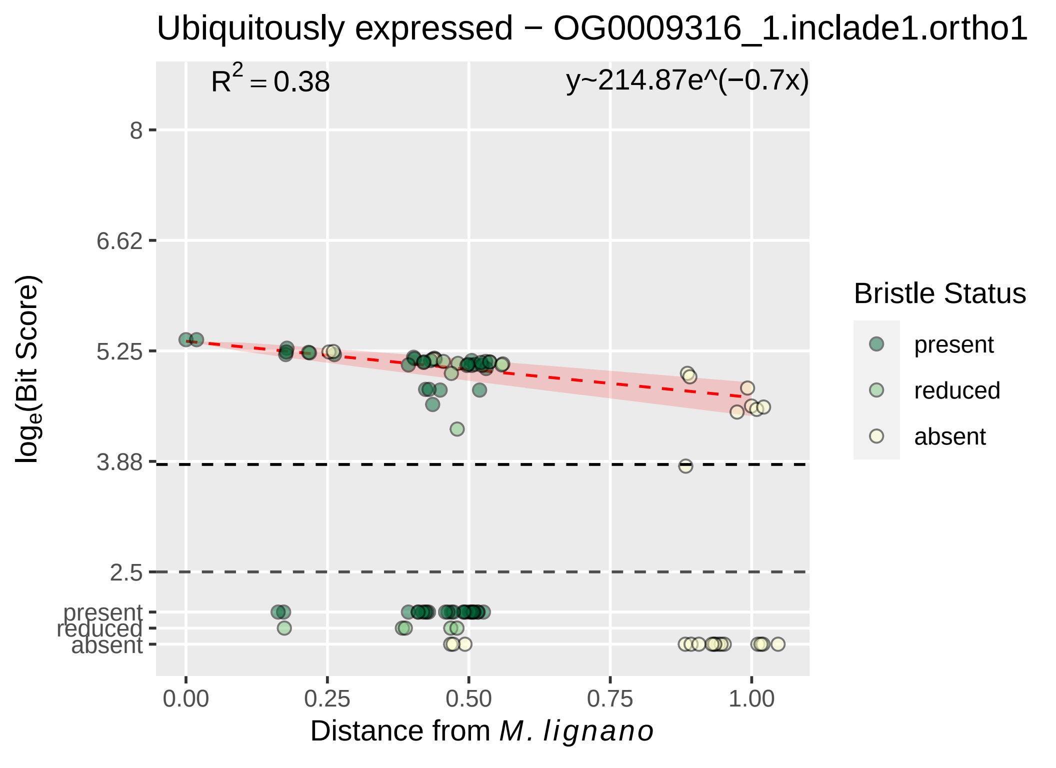

Ubiquitously expressed − OG0009316_1.inclade1.ortho1
R2 = 0.38
y~214.87e^(−0.7x)
8
6.62
5.25
3.88
2.5
present
reduced
absent
loge(Bit Score)
0.00
0.25
0.50
0.75
1.00
Distance from M. lignano
Bristle Status
present
reduced
absent

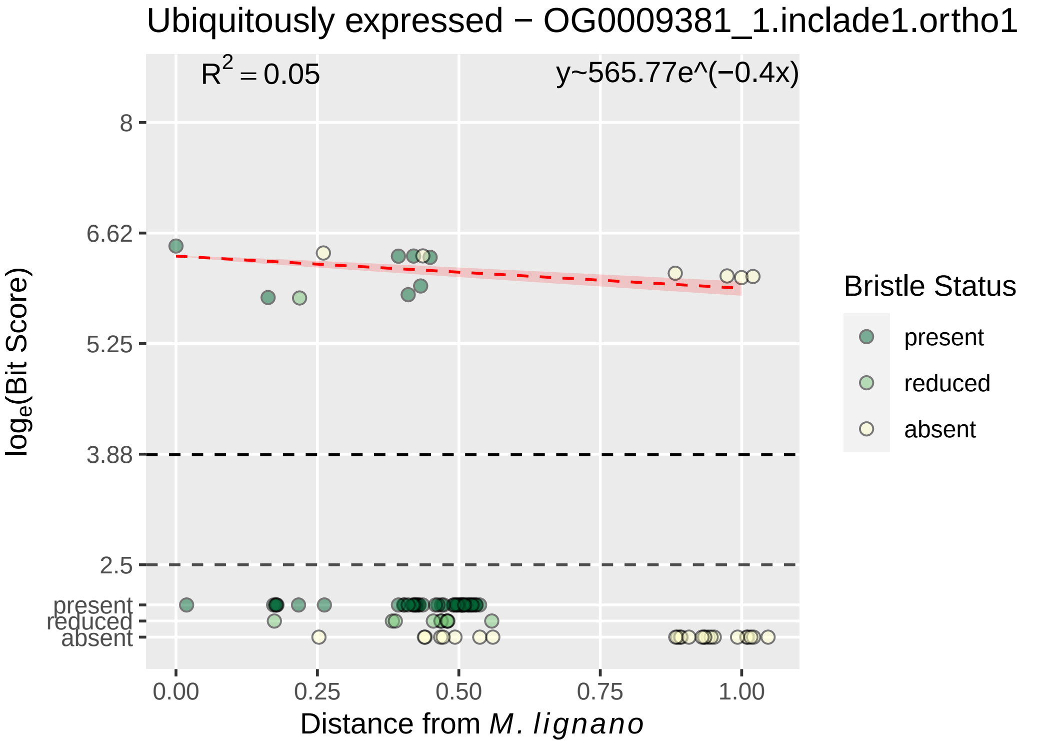

Ubiquitously expressed − OG0009381_1.inclade1.ortho1
R2 = 0.05
y~565.77e^(−0.4x)
8
6.62
5.25
3.88
2.5
present
reduced
absent
0.00
0.25
0.50
0.75
1.00
loge(Bit Score)
Distance from M. lignano
Bristle Status
present
reduced
absent

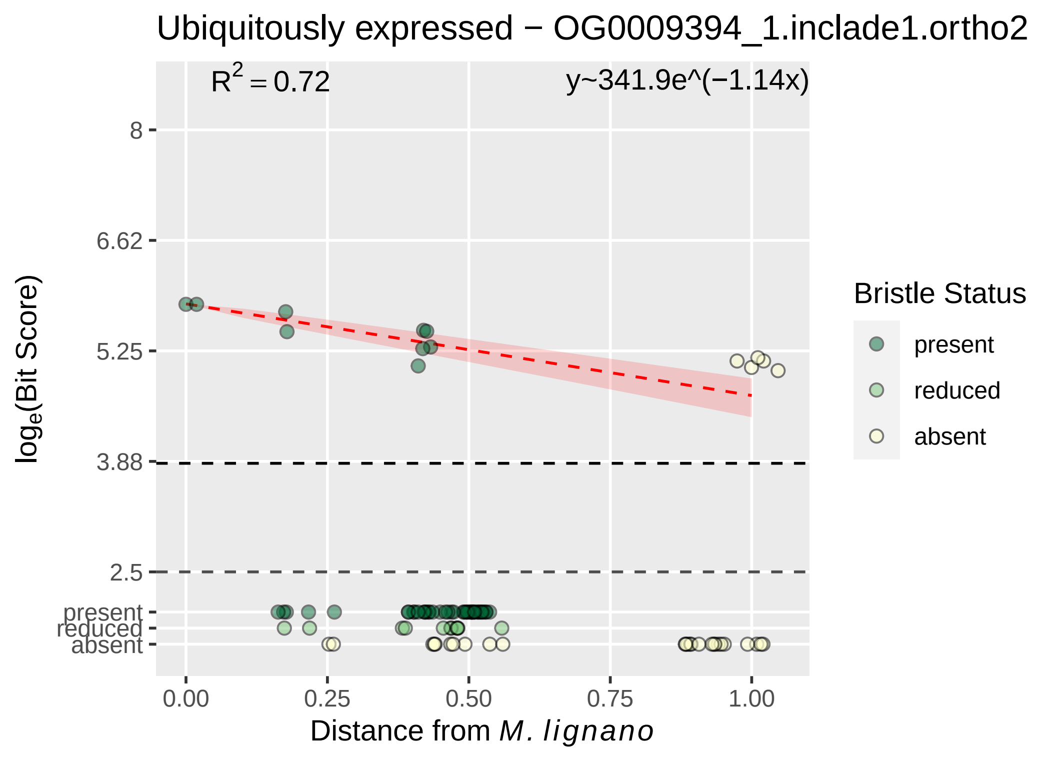

Ubiquitously expressed − OG0009394_1.inclade1.ortho2
R^2 = 0.72
y~341.9e^(−1.14x)
8
6.62
5.25
3.88
2.5
present
reduced
absent
0.00
0.25
0.50
0.75
1.00
log_e(Bit Score)
Distance from M. lignano
Bristle Status
present
reduced
absent

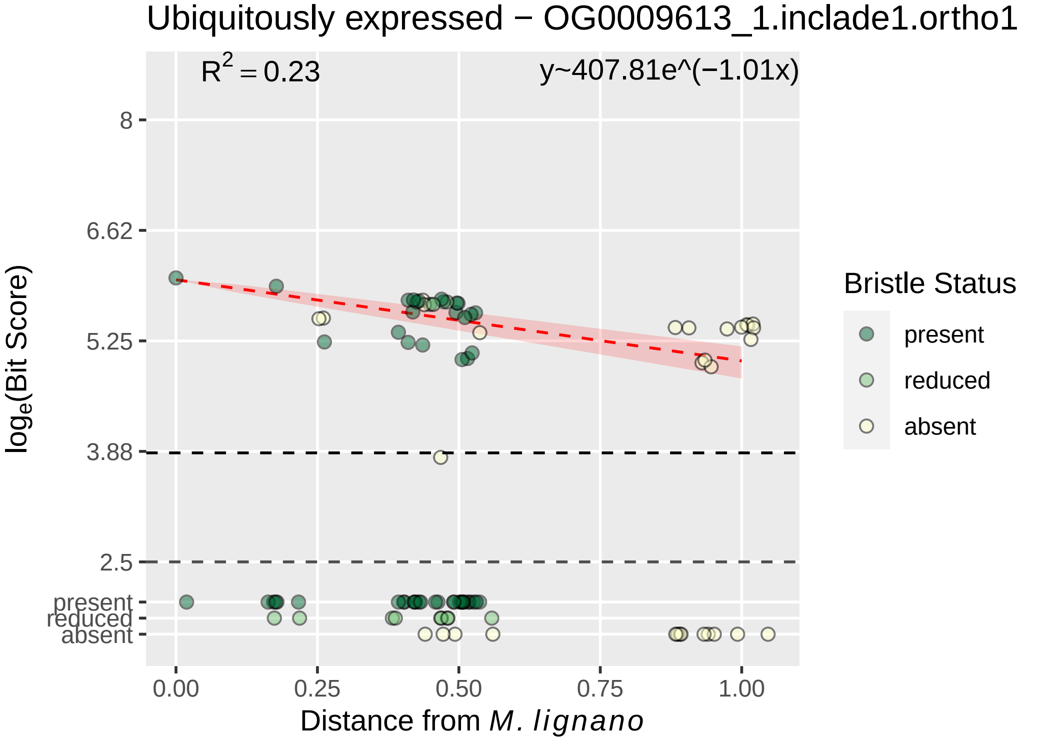

Ubiquitously expressed − OG0009613_1.inclade1.ortho1
$R^2 = 0.23$
y~407.81e^(−1.01x)
8
6.62
5.25
3.88
2.5
present
reduced
absent
$\log_e$(Bit Score)
0.00
0.25
0.50
0.75
1.00
Distance from *M. lignano*
Bristle Status
present
reduced
absent

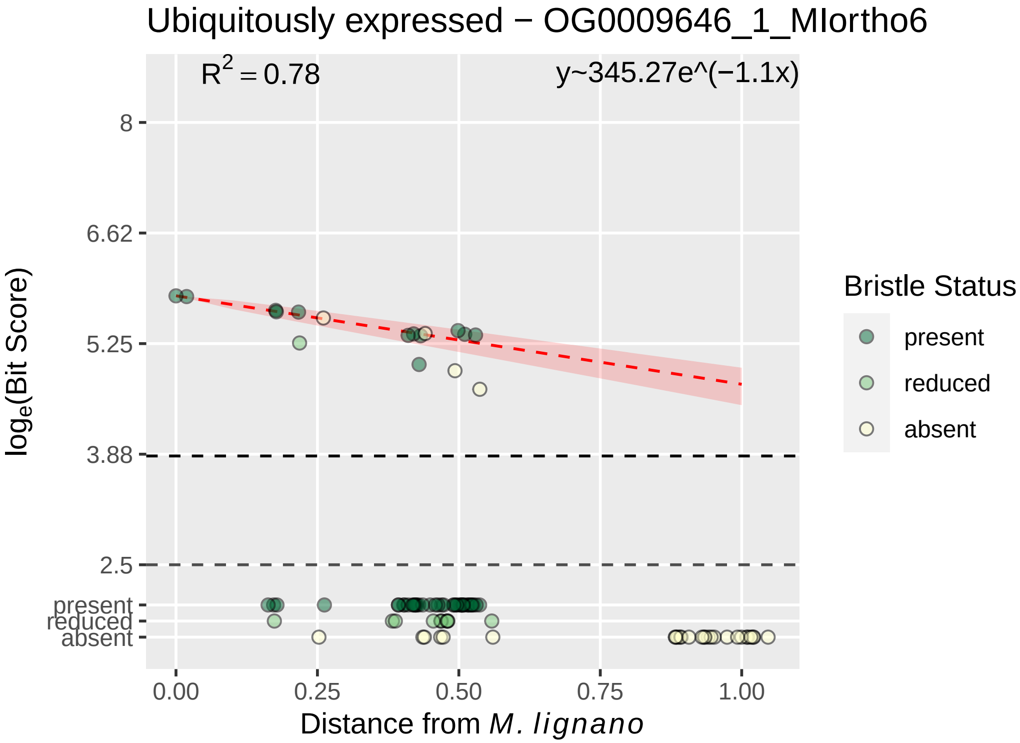
Ubiquitously expressed − OG0009646_1_MIortho6
R^2 = 0.78
y~345.27e^(−1.1x)
8
6.62
5.25
3.88
2.5
present
reduced
absent
log_e(Bit Score)
0.00
0.25
0.50
0.75
1.00
Distance from M. lignano
Bristle Status
present
reduced
absent

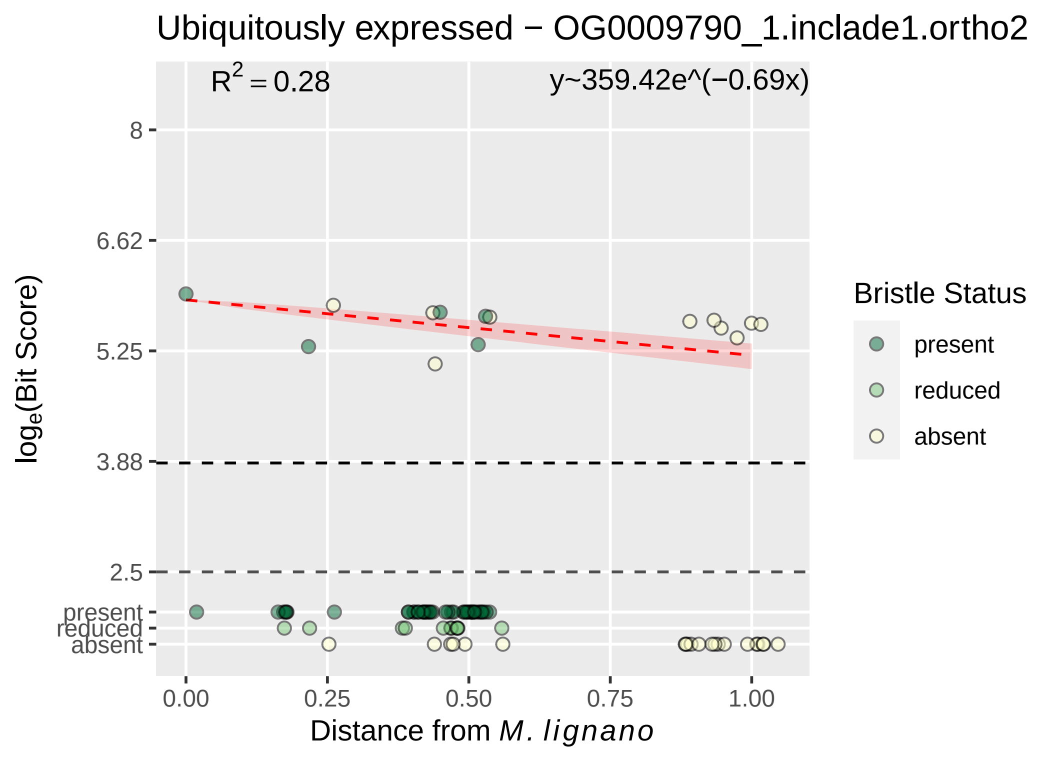

Ubiquitously expressed − OG0009790_1.inclade1.ortho2
R² = 0.28
y~359.42e^(−0.69x)
8
6.62
5.25
3.88
2.5
present
reduced
absent
0.00
0.25
0.50
0.75
1.00
Distance from M. lignano
loge(Bit Score)
Bristle Status
present
reduced
absent

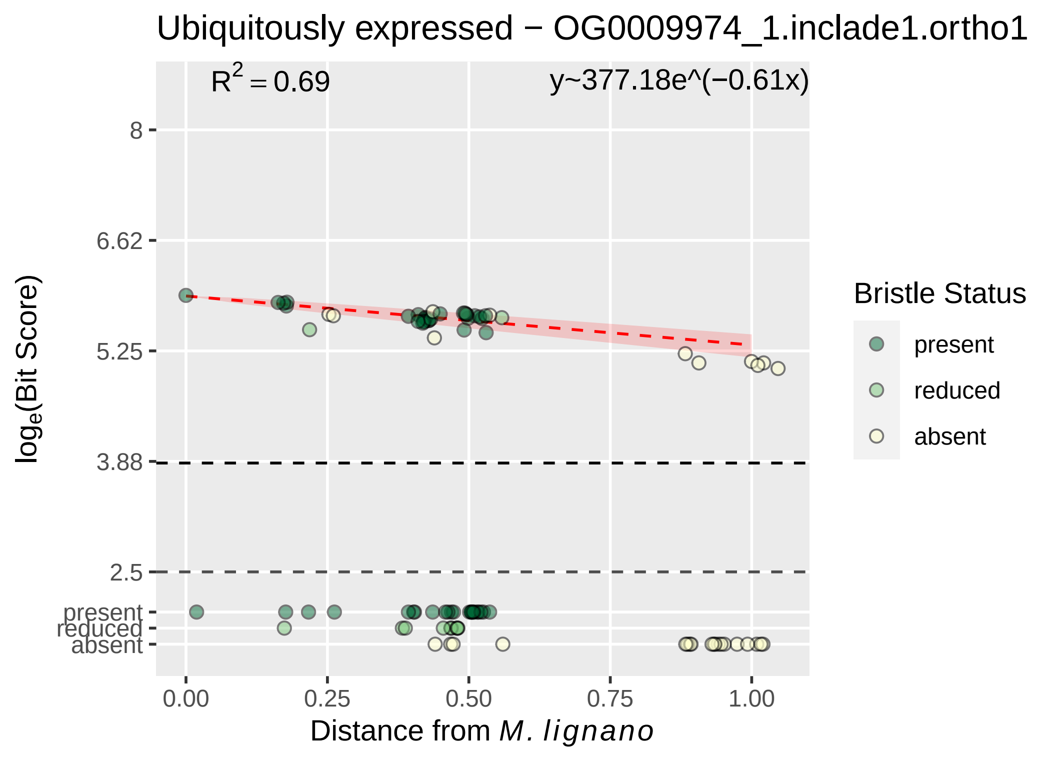

Ubiquitously expressed − OG0009974_1.inclade1.ortho1
$R^2 = 0.69$
y~377.18e^(−0.61x)
8
6.62
5.25
3.88
2.5
present
reduced
absent
0.00
0.25
0.50
0.75
1.00
Distance from M. lignano
loge(Bit Score)
Bristle Status
present
reduced
absent

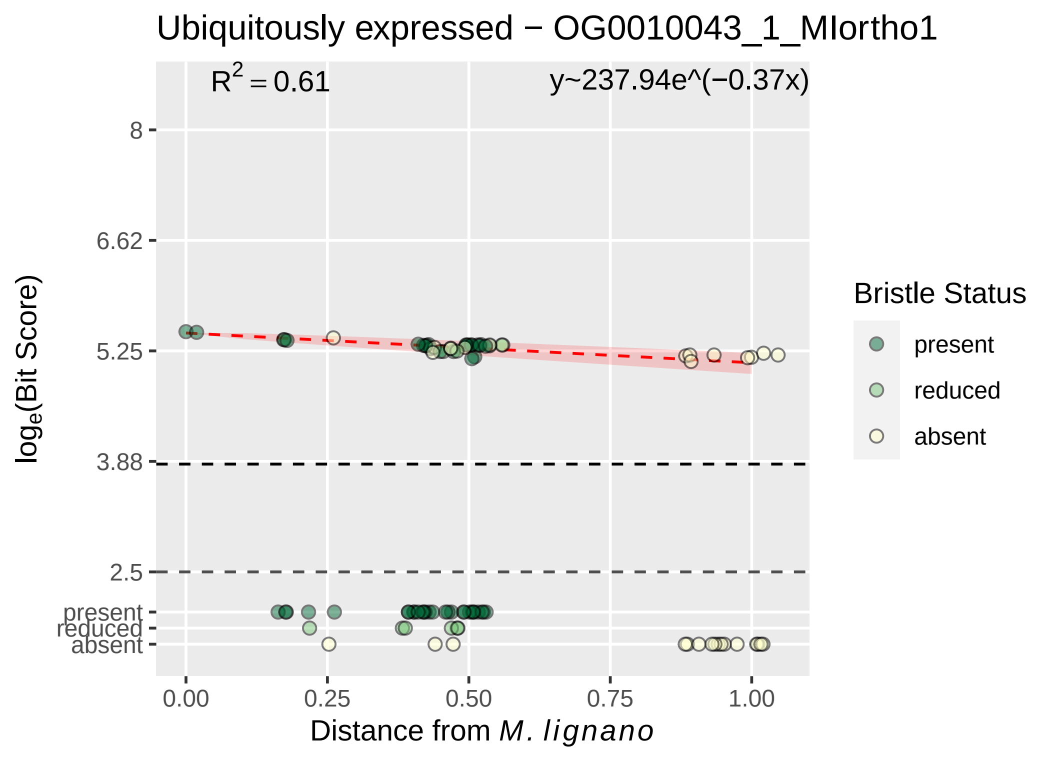

Ubiquitously expressed − OG0010043_1_MIortho1
$R^2 = 0.61$
y~237.94e^(−0.37x)
8
6.62
5.25
3.88
2.5
present
reduced
absent
$\log_e$(Bit Score)
0.00
0.25
0.50
0.75
1.00
Distance from *M. lignano*
Bristle Status
present
reduced
absent

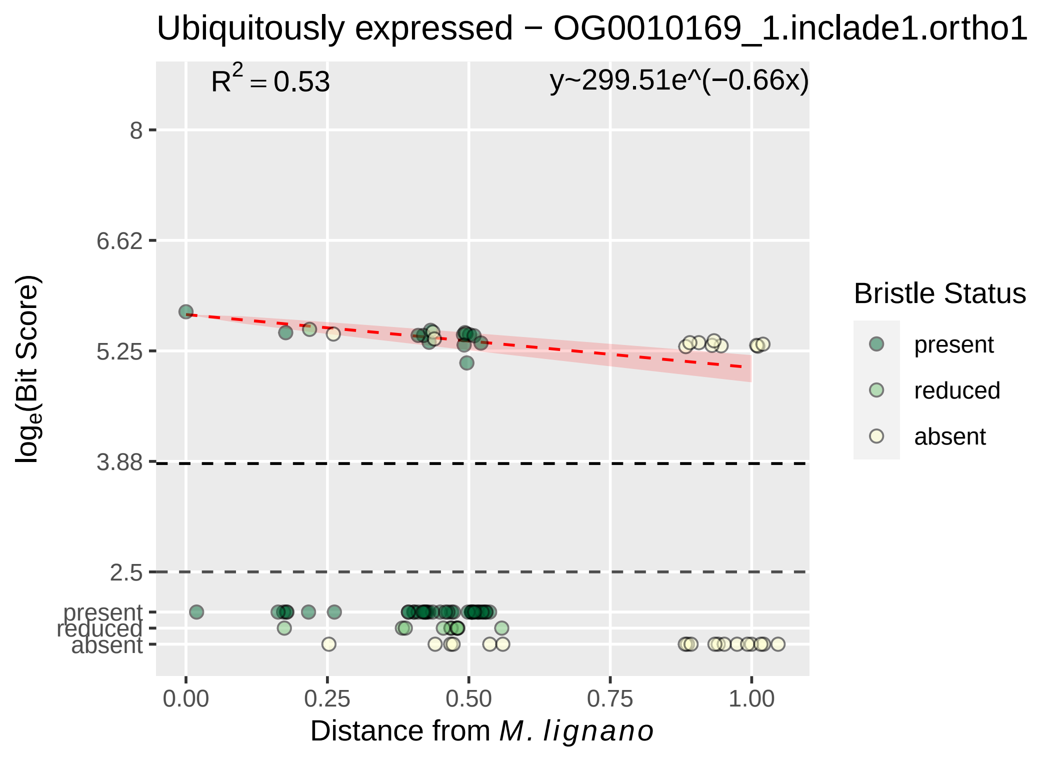
Ubiquitously expressed − OG0010169_1.inclade1.ortho1
R² = 0.53
y~299.51e^(−0.66x)
8
6.62
5.25
3.88
2.5
present
reduced
absent
log_e(Bit Score)
0.00
0.25
0.50
0.75
1.00
Distance from M. lignano
Bristle Status
present
reduced
absent

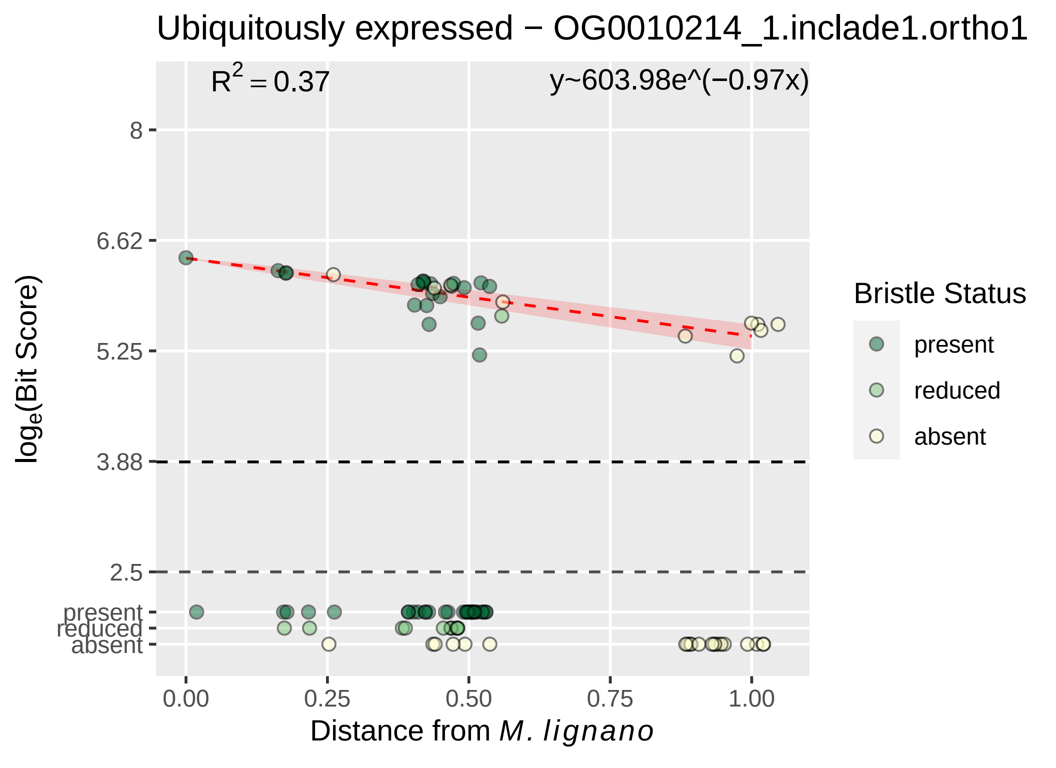
Ubiquitously expressed − OG0010214_1.inclade1.ortho1
R^2 = 0.37
y~603.98e^(−0.97x)
8
6.62
5.25
3.88
2.5
present
reduced
absent
0.00
0.25
0.50
0.75
1.00
Distance from M. lignano
log_e(Bit Score)
Bristle Status
present
reduced
absent

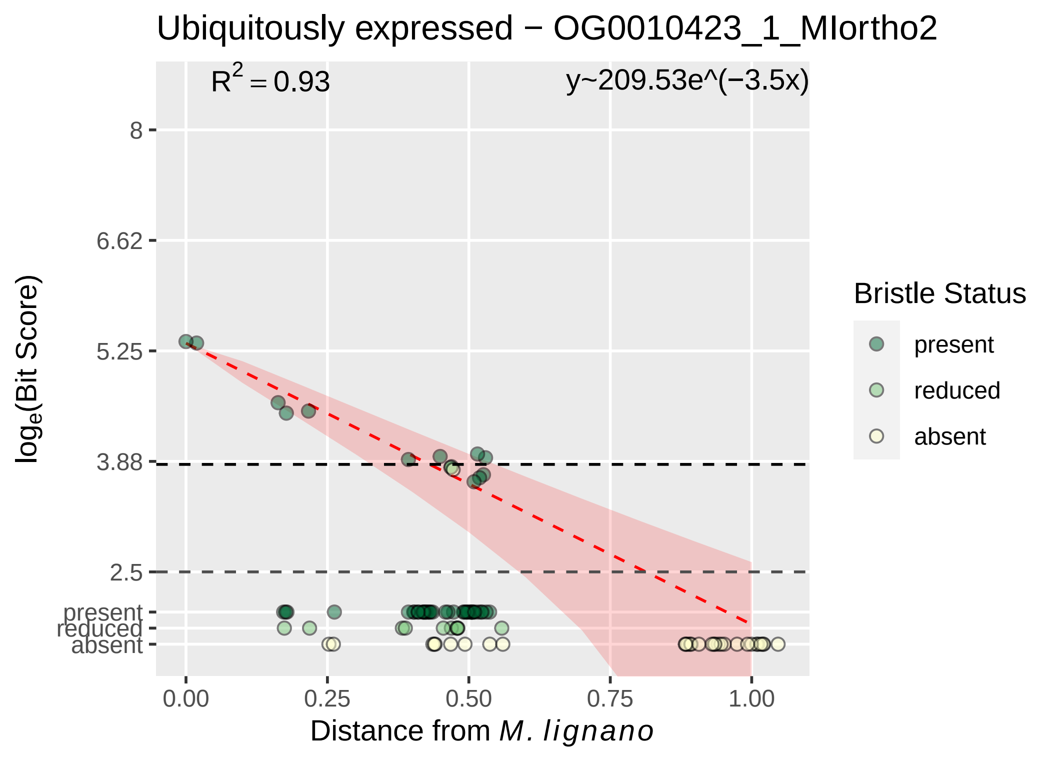

Ubiquitously expressed − OG0010423_1_MIortho2
R^2 = 0.93
y~209.53e^(−3.5x)
8
6.62
5.25
3.88
2.5
present
reduced
absent
0.00
0.25
0.50
0.75
1.00
Distance from M. lignano
loge(Bit Score)
Bristle Status
present
reduced
absent

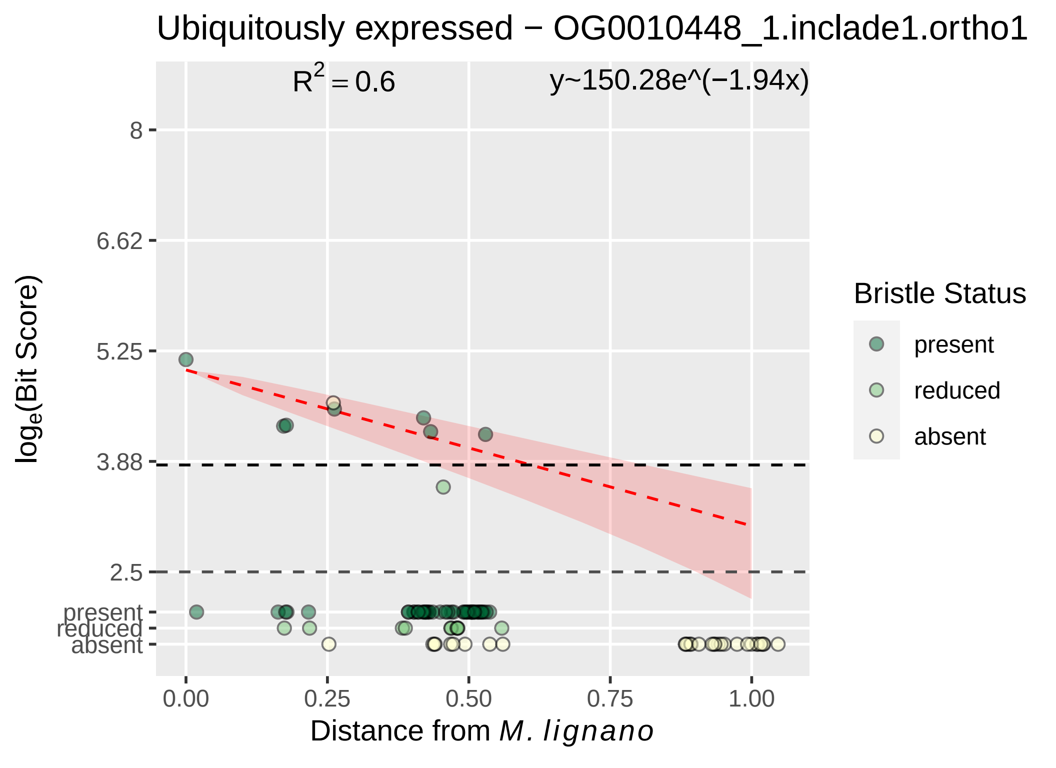

Ubiquitously expressed − OG0010448_1.inclade1.ortho1
$R^2 = 0.6$
y~150.28e^(−1.94x)
log$_e$(Bit Score)
8
6.62
5.25
3.88
2.5
present
reduced
absent
0.00
0.25
0.50
0.75
1.00
Distance from *M. lignano*
Bristle Status
present
reduced
absent

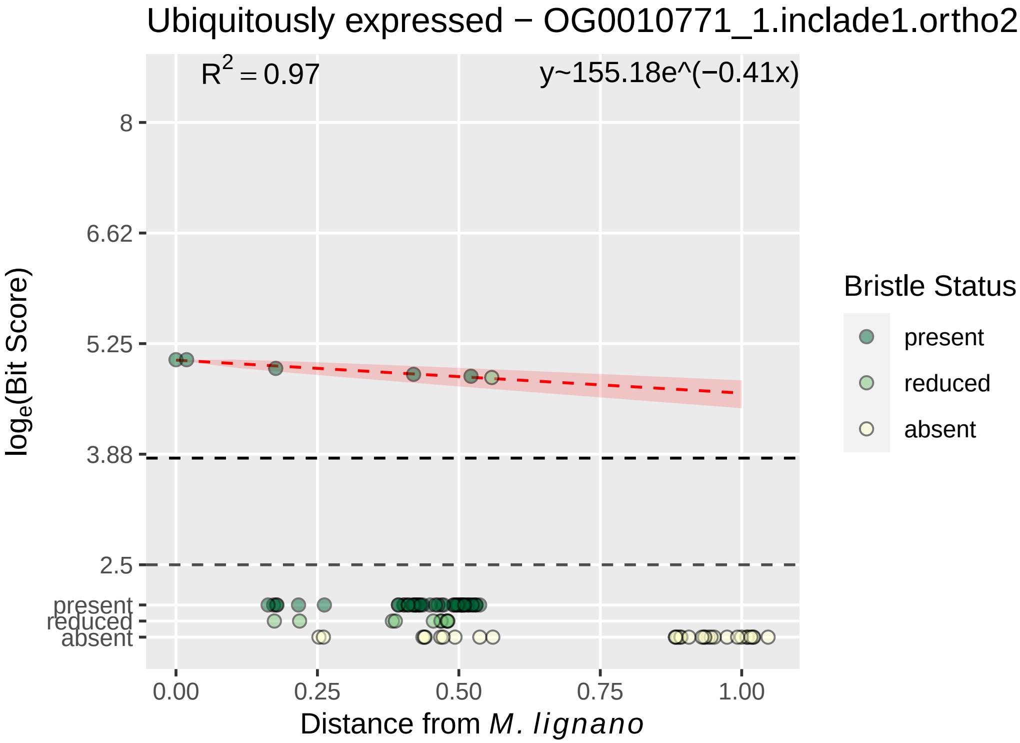

Ubiquitously expressed − OG0010771_1.inclade1.ortho2
R2 = 0.97
y~155.18e^(−0.41x)
8
6.62
5.25
3.88
2.5
present
reduced
absent
loge(Bit Score)
0.00
0.25
0.50
0.75
1.00
Distance from M. lignano
Bristle Status
present
reduced
absent

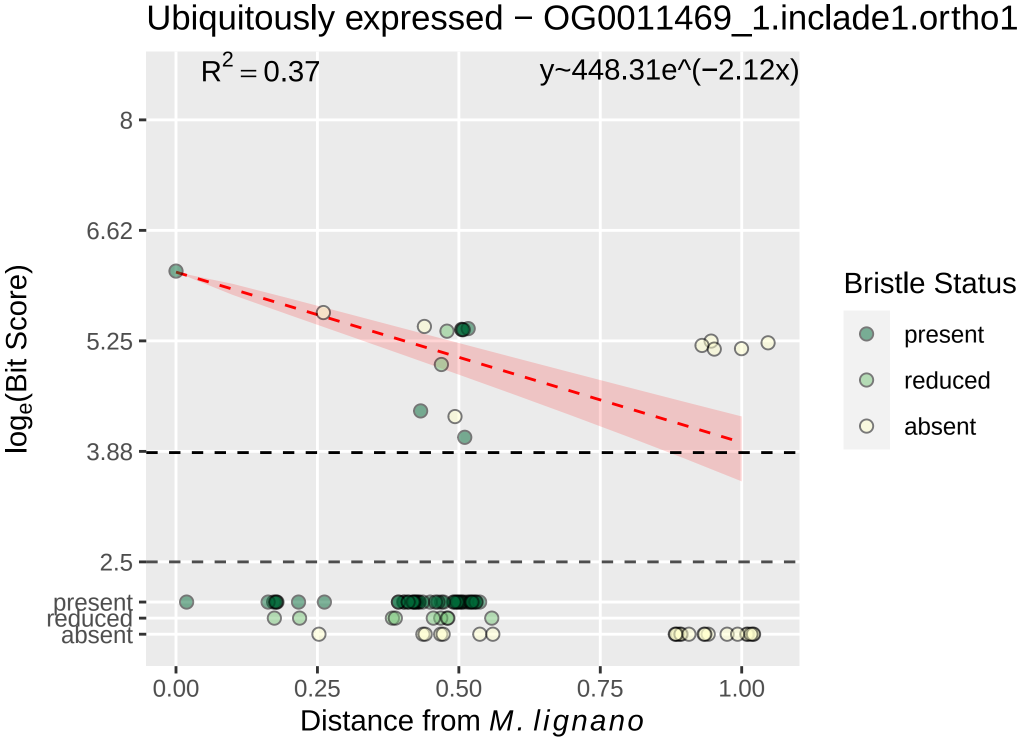

Ubiquitously expressed − OG0011469_1.inclade1.ortho1
$R^2 = 0.37$
y~448.31e^(−2.12x)
8
6.62
5.25
3.88
2.5
present
reduced
absent
0.00
0.25
0.50
0.75
1.00
Distance from *M. lignano*
$\log_e$(Bit Score)
Bristle Status
present
reduced
absent

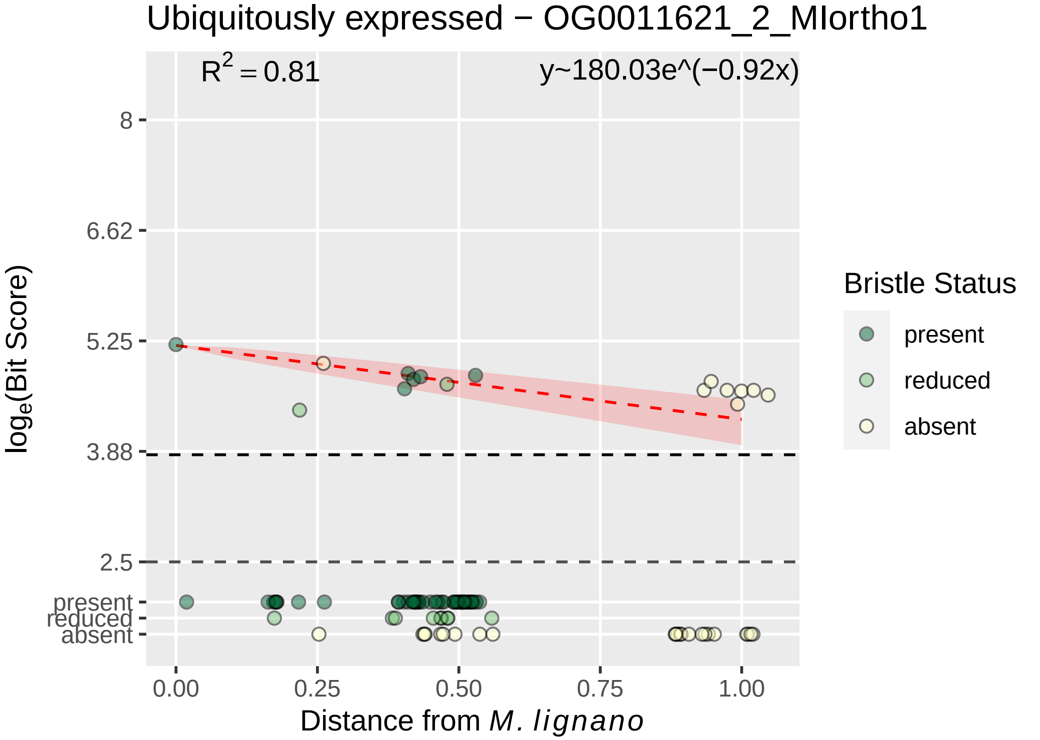

Ubiquitously expressed − OG0011621_2_MIortho1
$R^2 = 0.81$
y~180.03e^(−0.92x)
8
6.62
5.25
3.88
2.5
present
reduced
absent
$log_e$(Bit Score)
0.00
0.25
0.50
0.75
1.00
Distance from *M. lignano*
Bristle Status
present
reduced
absent

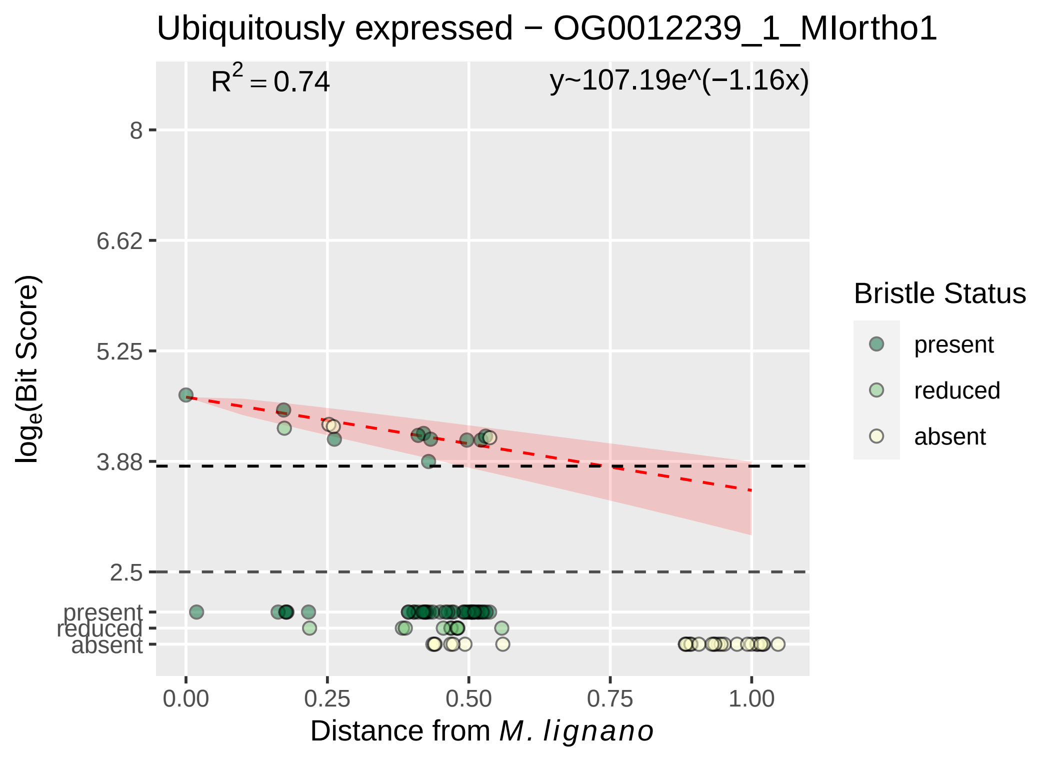

Ubiquitously expressed − OG0012239_1_MIortho1
$R^2 = 0.74$
y~107.19e^(−1.16x)
8
6.62
5.25
3.88
2.5
present
reduced
absent
0.00
0.25
0.50
0.75
1.00
log$_e$(Bit Score)
Distance from *M. lignano*
Bristle Status
present
reduced
absent

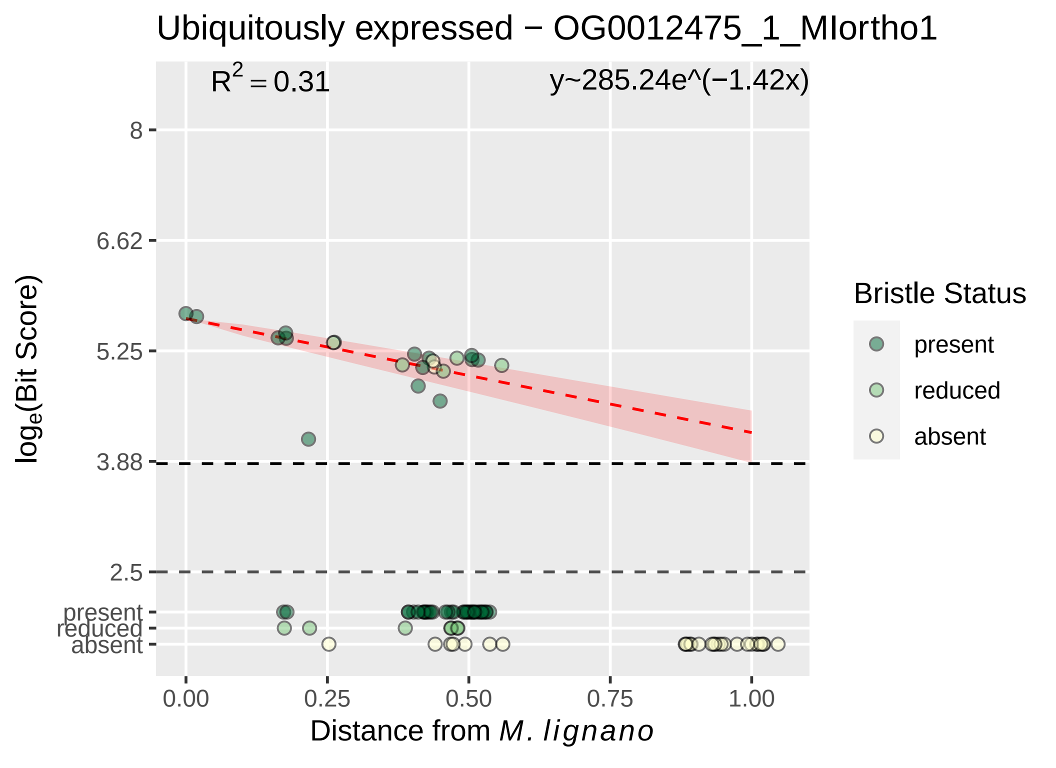

Ubiquitously expressed − OG0012475_1_MIortho1
R^2 = 0.31
y~285.24e^(−1.42x)
8
6.62
5.25
3.88
2.5
present
reduced
absent
log_e(Bit Score)
0.00
0.25
0.50
0.75
1.00
Distance from M. lignano
Bristle Status
present
reduced
absent

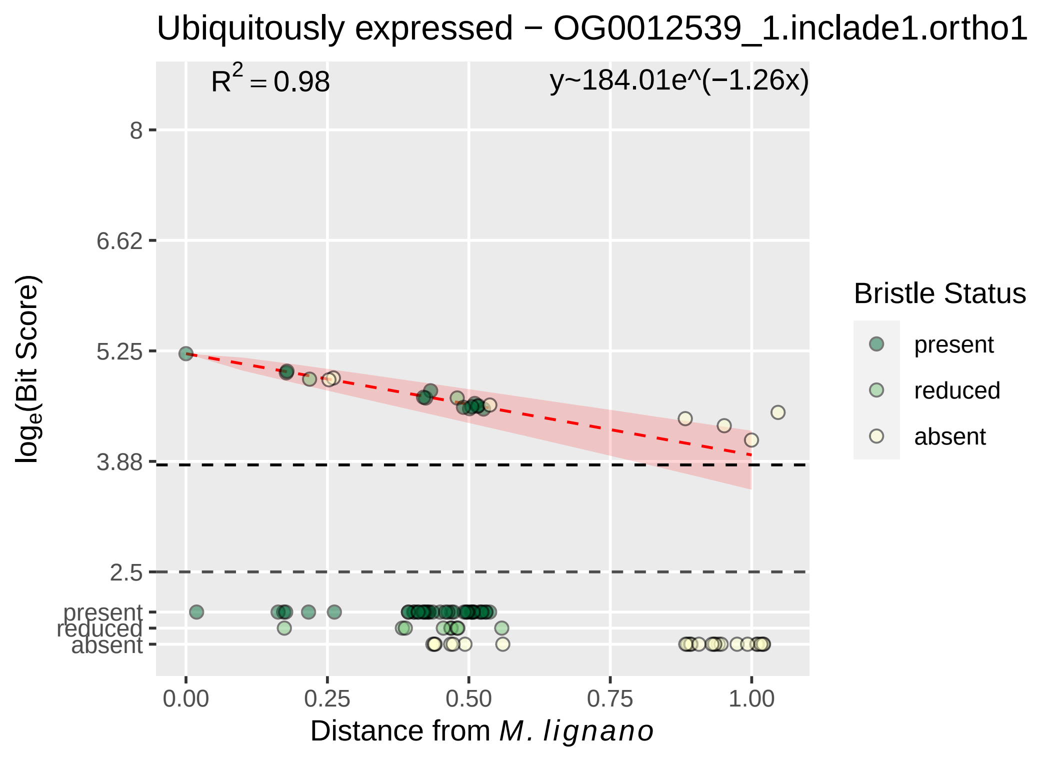

Ubiquitously expressed − OG0012539_1.inclade1.ortho1
R² = 0.98
y~184.01e^(−1.26x)
log_e(Bit Score)
8
6.62
5.25
3.88
2.5
present
reduced
absent
0.00
0.25
0.50
0.75
1.00
Distance from M. lignano
Bristle Status
present
reduced
absent

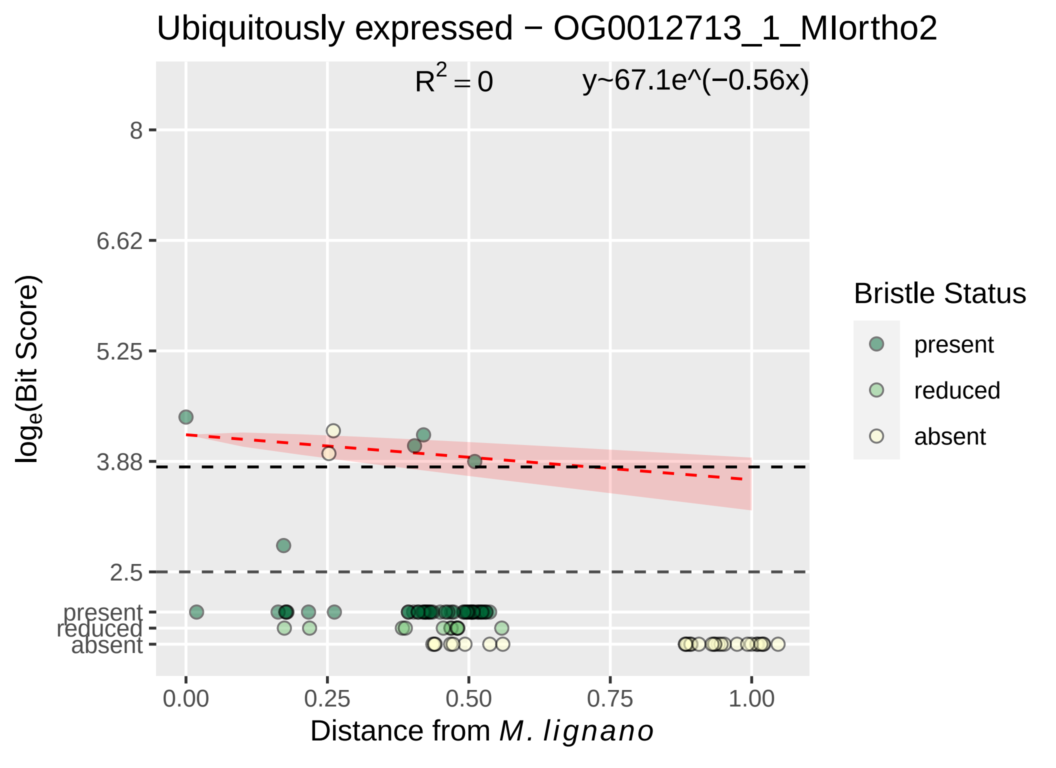

Ubiquitously expressed − OG0012713_1_MIortho2
R² = 0
y~67.1e^(−0.56x)
8
6.62
5.25
3.88
2.5
present
reduced
absent
log_e(Bit Score)
0.00
0.25
0.50
0.75
1.00
Distance from M. lignano
Bristle Status
present
reduced
absent

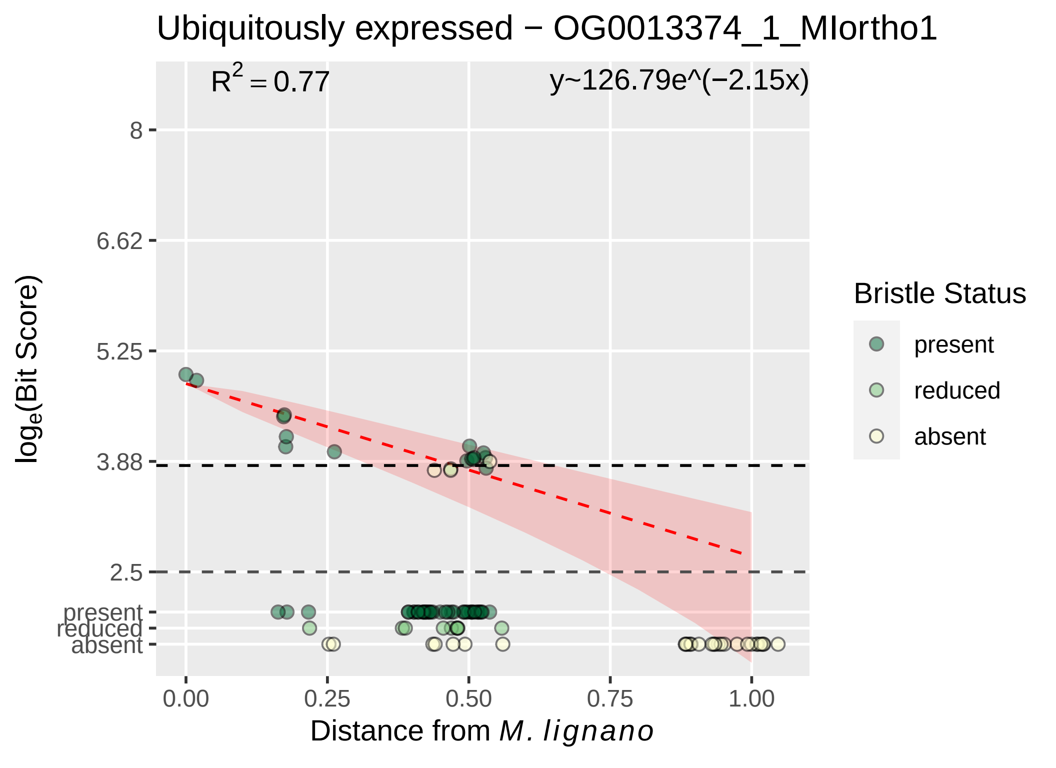

Ubiquitously expressed − OG0013374_1_MIortho1
$R^2 = 0.77$
y~126.79e^(−2.15x)
8
6.62
5.25
3.88
2.5
present
reduced
absent
0.00
0.25
0.50
0.75
1.00
Distance from M. lignano
log_e(Bit Score)
Bristle Status
present
reduced
absent

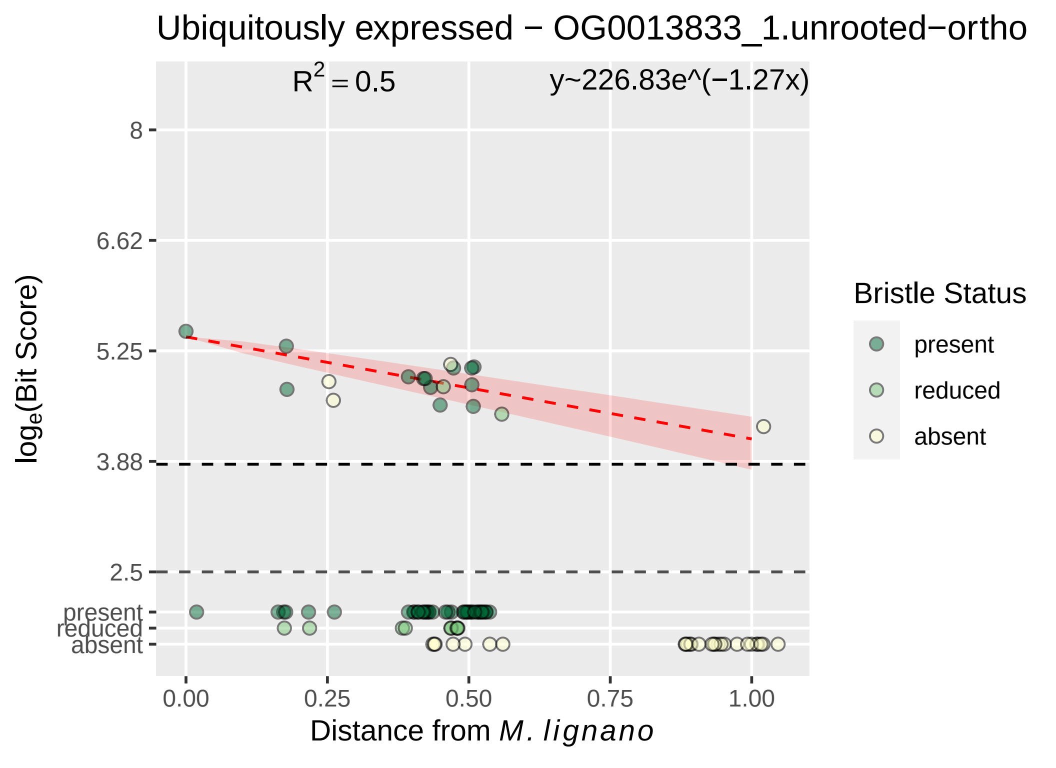
Ubiquitously expressed − OG0013833_1.unrooted−ortho
R² = 0.5
y~226.83e^(−1.27x)
8
6.62
5.25
3.88
2.5
present
reduced
absent
0.00
0.25
0.50
0.75
1.00
Distance from M. lignano
loge(Bit Score)
Bristle Status
present
reduced
absent

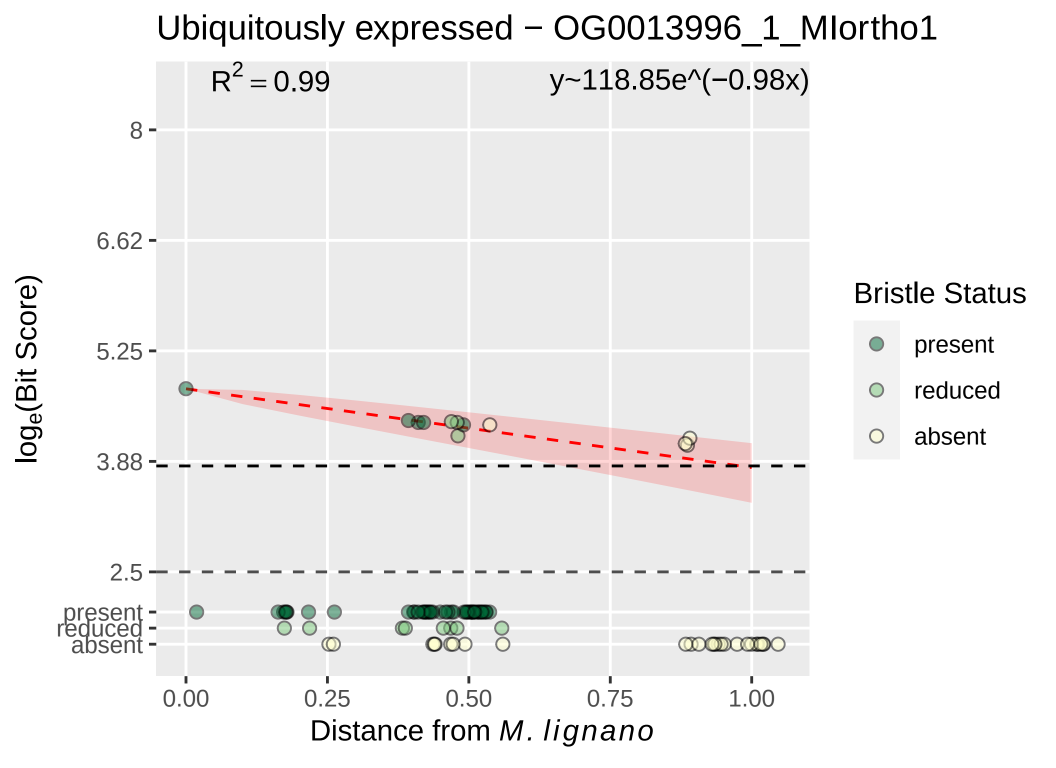

Ubiquitously expressed − OG0013996_1_MIortho1
R² = 0.99
y~118.85e^(−0.98x)
8
6.62
5.25
3.88
2.5
present
reduced
absent
0.00
0.25
0.50
0.75
1.00
log_e(Bit Score)
Distance from M. lignano
Bristle Status
present
reduced
absent

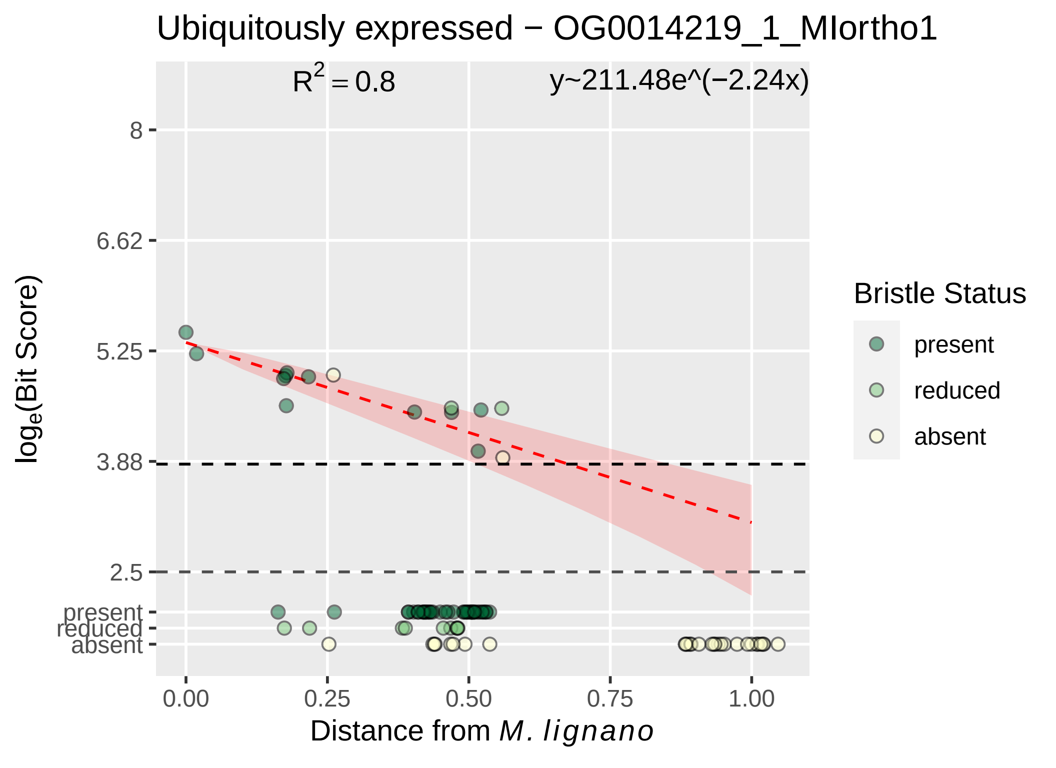

Ubiquitously expressed − OG0014219_1_MIortho1
R² = 0.8
y~211.48e^(−2.24x)
8
6.62
5.25
3.88
2.5
present
reduced
absent
loge(Bit Score)
0.00
0.25
0.50
0.75
1.00
Distance from M. lignano
Bristle Status
present
reduced
absent

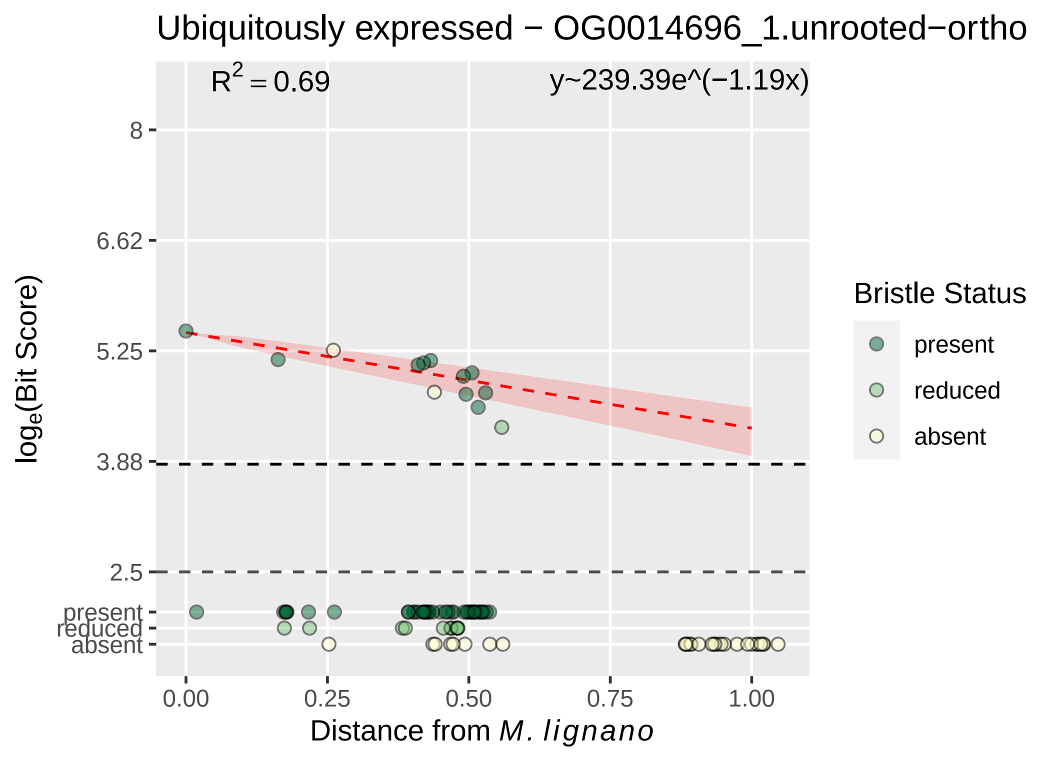

Ubiquitously expressed − OG0014696_1.unrooted−ortho
R2 = 0.69
y~239.39e^(−1.19x)
8
6.62
5.25
3.88
2.5
present
reduced
absent
0.00
0.25
0.50
0.75
1.00
loge(Bit Score)
Distance from M. lignano
Bristle Status
present
reduced
absent

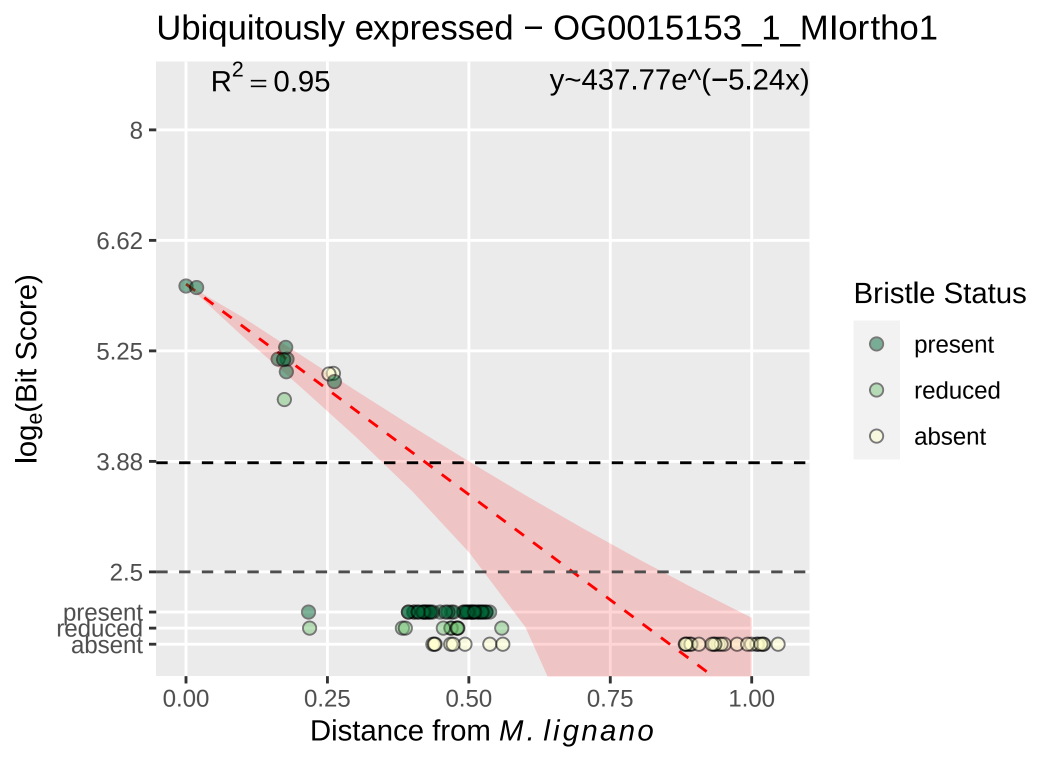

Ubiquitously expressed − OG0015153_1_MIortho1
R^2 = 0.95
y~437.77e^(−5.24x)
8
6.62
5.25
3.88
2.5
present
reduced
absent
log_e(Bit Score)
0.00
0.25
0.50
0.75
1.00
Distance from M. lignano
Bristle Status
present
reduced
absent

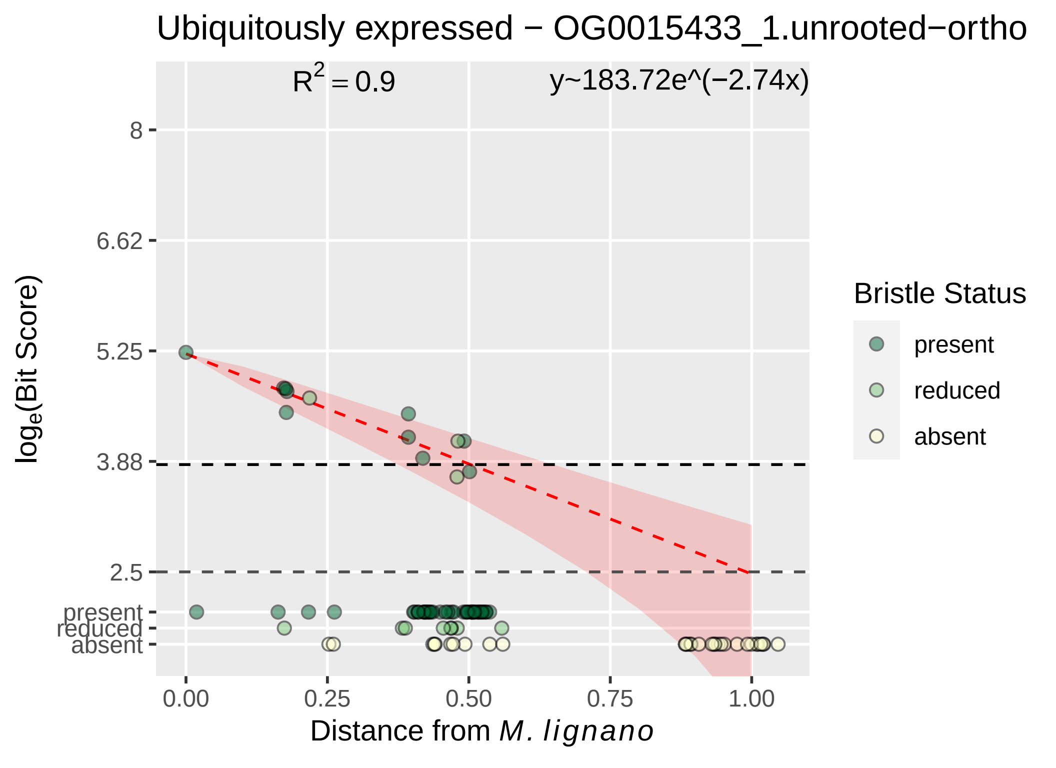

Ubiquitously expressed − OG0015433_1.unrooted−ortho
R² = 0.9
y~183.72e^(−2.74x)
8
6.62
5.25
3.88
2.5
present
reduced
absent
log_e(Bit Score)
0.00
0.25
0.50
0.75
1.00
Distance from M. lignano
Bristle Status
present
reduced
absent

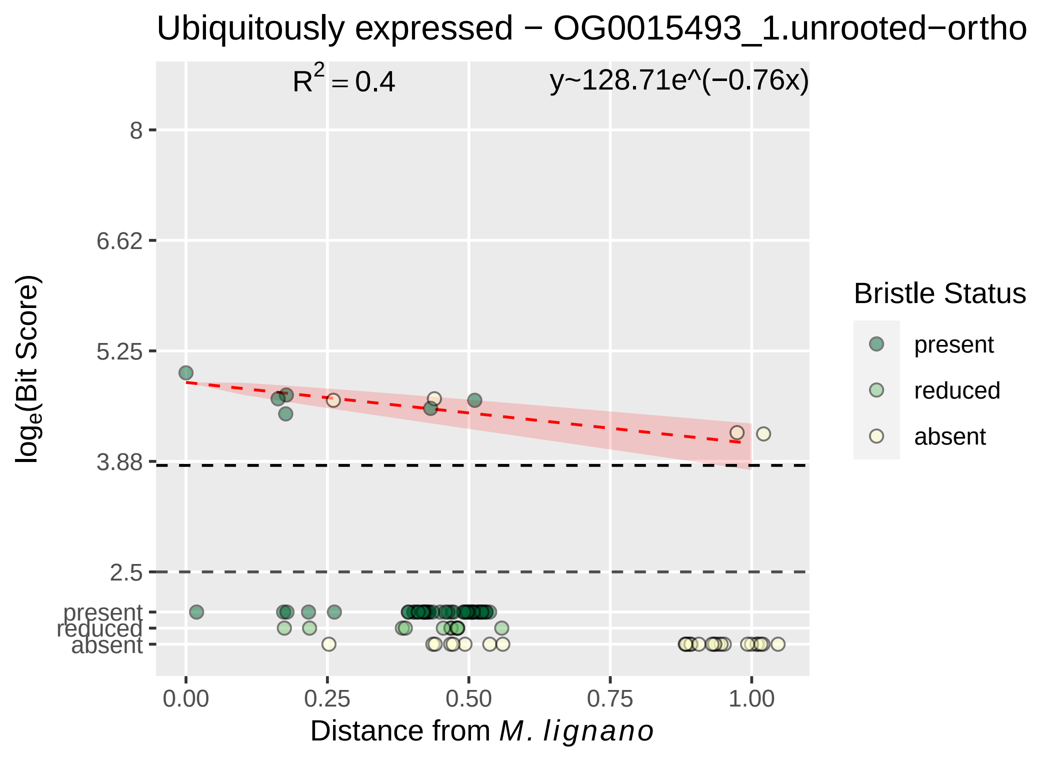

Ubiquitously expressed − OG0015493_1.unrooted−ortho
R^2 = 0.4
y~128.71e^(−0.76x)
8
6.62
5.25
3.88
2.5
present
reduced
absent
log_e(Bit Score)
0.00
0.25
0.50
0.75
1.00
Distance from M. lignano
Bristle Status
present
reduced
absent

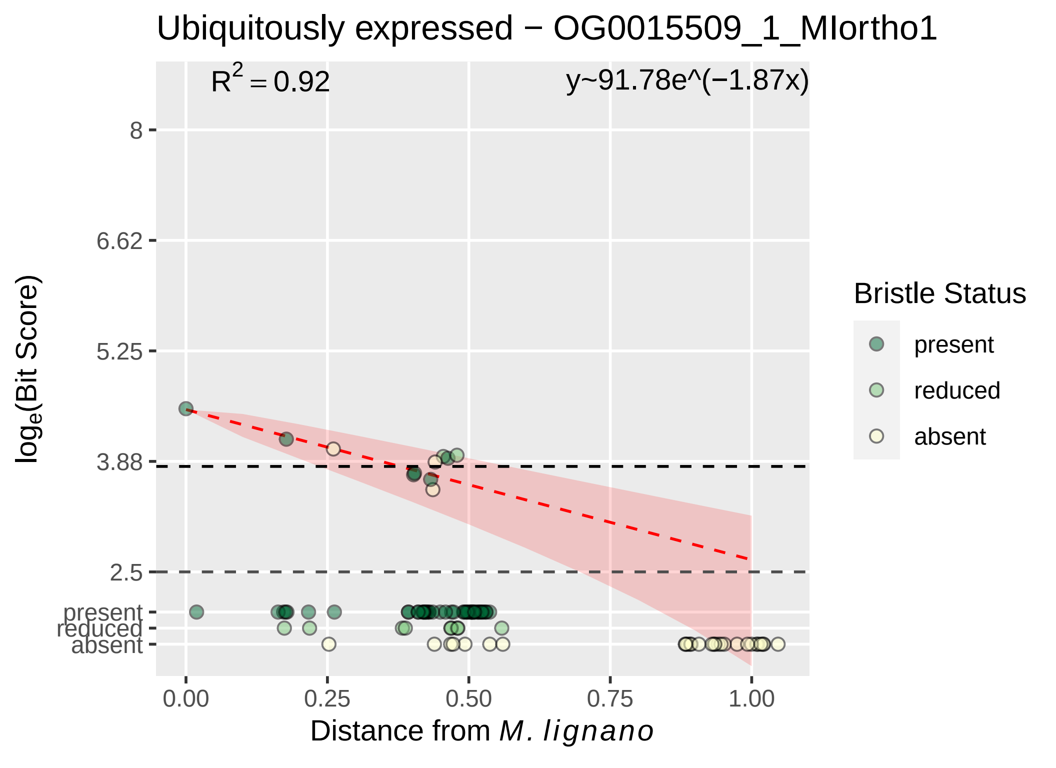
Ubiquitously expressed − OG0015509_1_MIortho1
R^2 = 0.92
y~91.78e^(−1.87x)
8
6.62
5.25
3.88
2.5
present
reduced
absent
0.00
0.25
0.50
0.75
1.00
log_e(Bit Score)
Distance from M. lignano
Bristle Status
present
reduced
absent

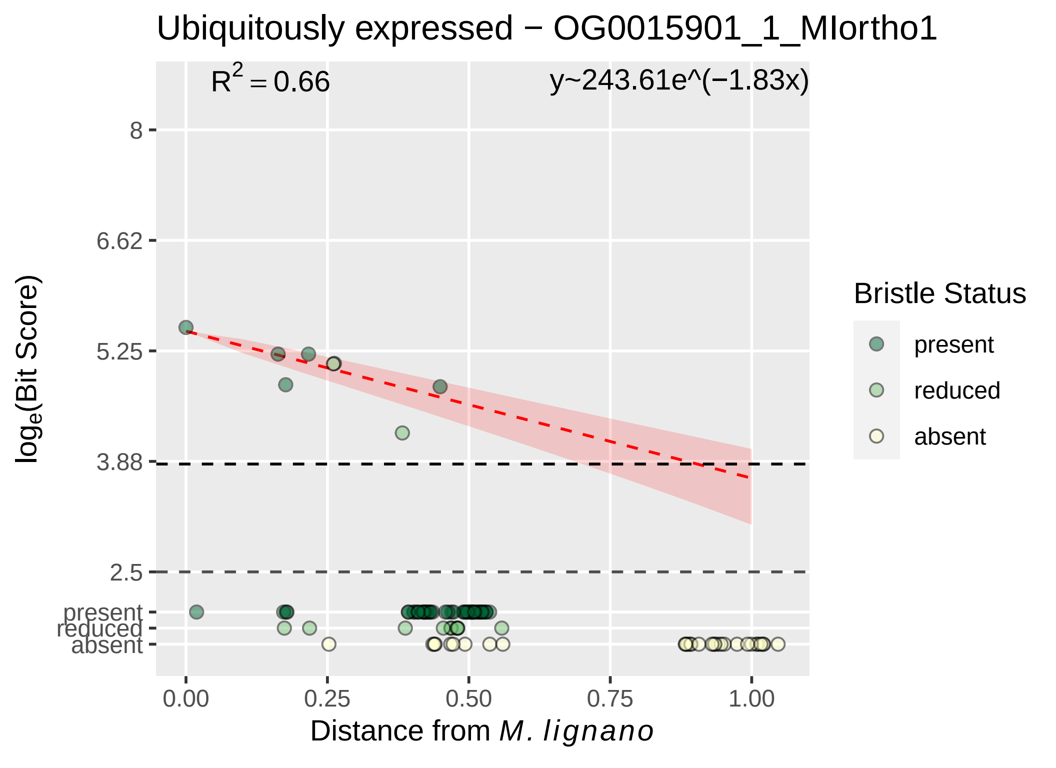

Ubiquitously expressed − OG0015901_1_MIortho1
R^2 = 0.66
y~243.61e^(−1.83x)
8
6.62
5.25
3.88
2.5
present
reduced
absent
0.00
0.25
0.50
0.75
1.00
Distance from M. lignano
loge(Bit Score)
Bristle Status
present
reduced
absent

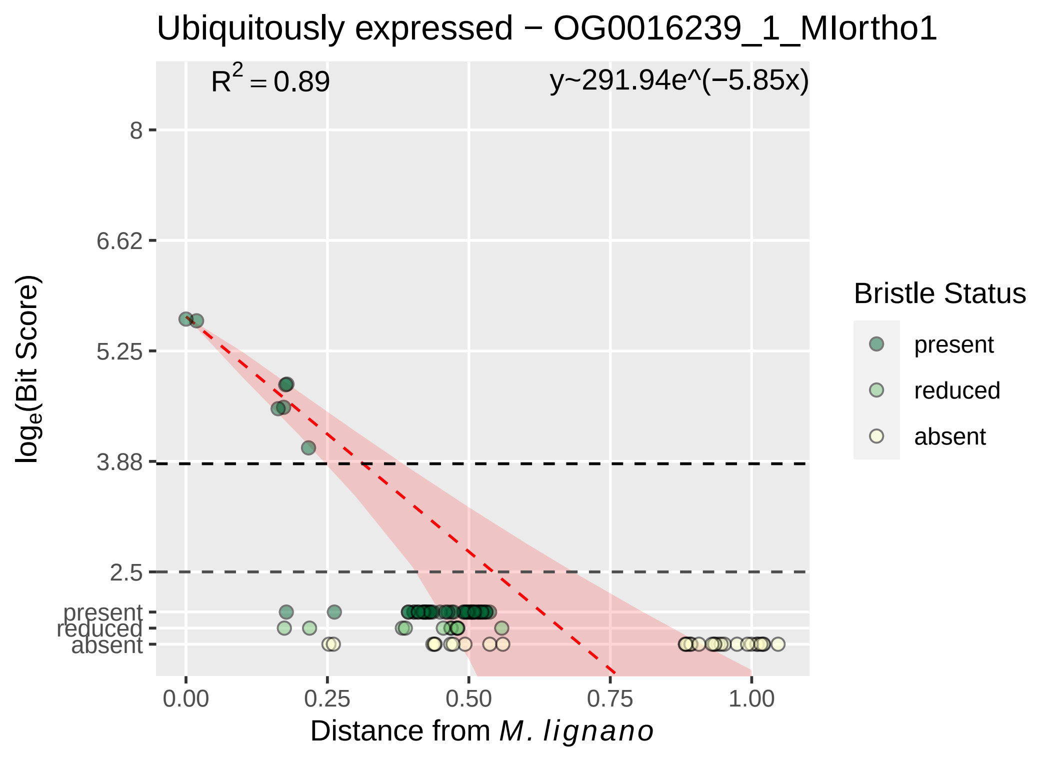

Ubiquitously expressed − OG0016239_1_MIortho1
$R^2 = 0.89$
y~291.94e^(−5.85x)
8
6.62
5.25
3.88
2.5
present
reduced
absent
0.00
0.25
0.50
0.75
1.00
Distance from *M. lignano*
$\log_e$(Bit Score)
Bristle Status
present
reduced
absent

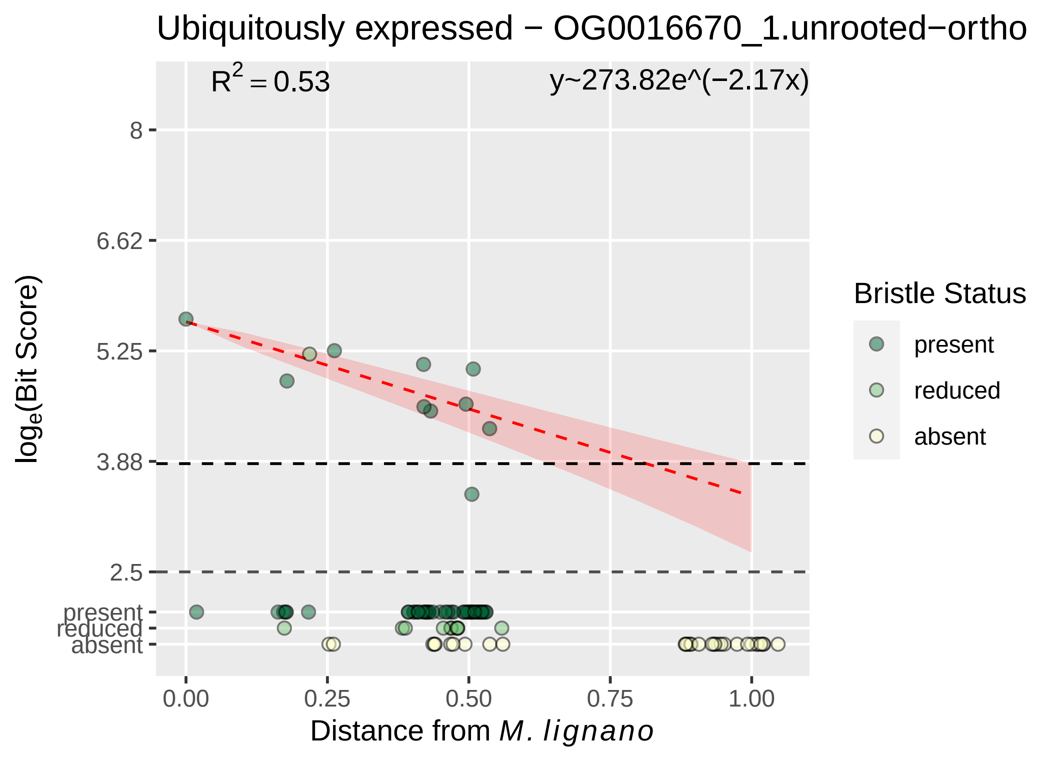

Ubiquitously expressed − OG0016670_1.unrooted−ortho
$R^2 = 0.53$
y~273.82e^(−2.17x)
8
6.62
5.25
3.88
2.5
present
reduced
absent
0.00
0.25
0.50
0.75
1.00
Distance from M. lignano
log$_e$(Bit Score)
Bristle Status
present
reduced
absent
