## Supplementary material for "Mating strategy predicts gene presence/absence patterns in a genus of simultaneously hermaphroditic flatworms": figure S5A

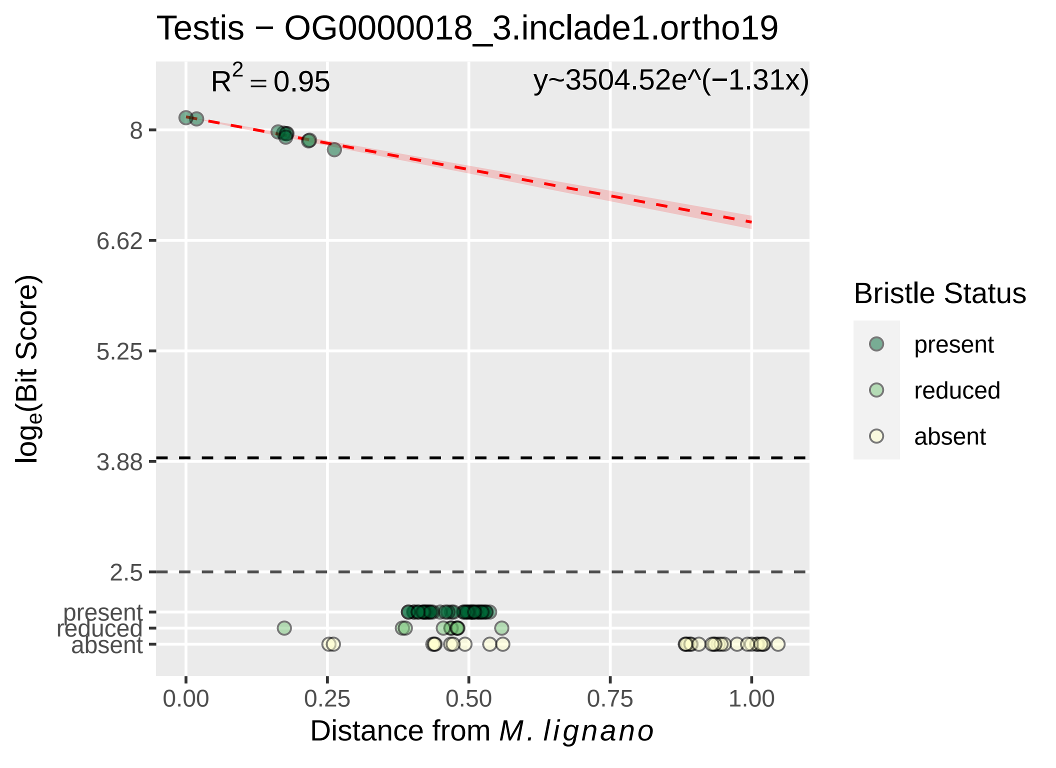

Testis − OG0000018_3.inclade1.ortho19
$R^2 = 0.95$
y~3504.52e^(−1.31x)
8
6.62
5.25
3.88
2.5
present
reduced
absent
$log_e$(Bit Score)
0.00
0.25
0.50
0.75
1.00
Distance from *M. lignano*
Bristle Status
present
reduced
absent

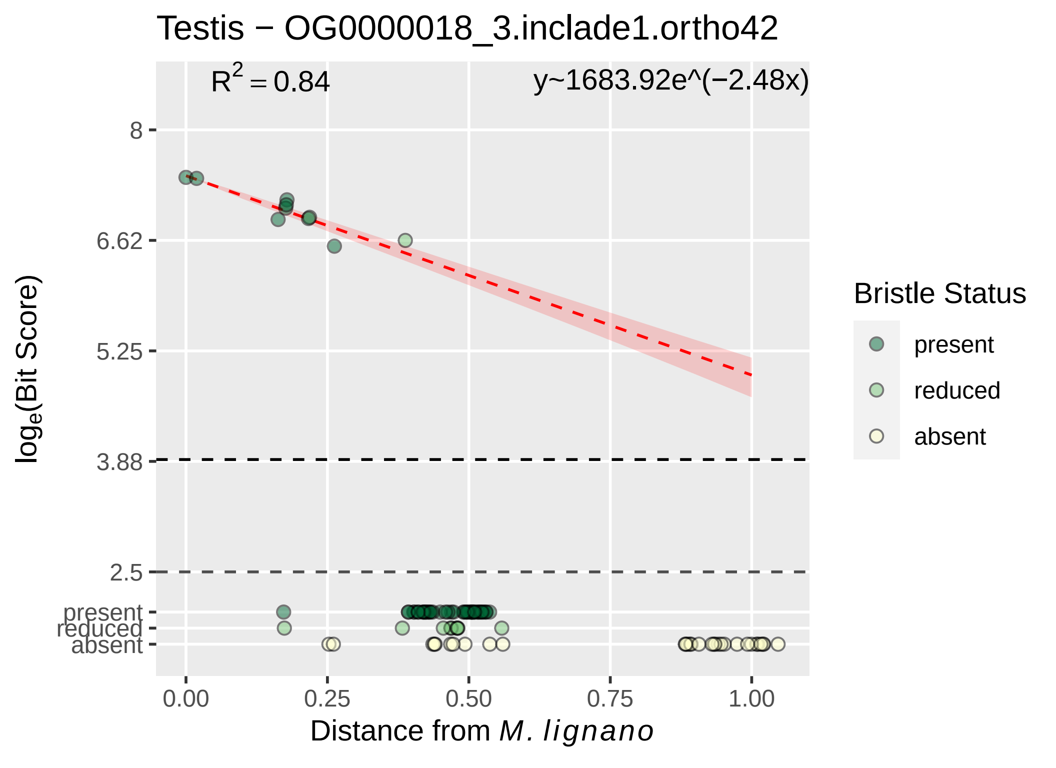
Testis − OG0000018_3.inclade1.ortho42
$R^2 = 0.84$
y~1683.92e^(−2.48x)
8
6.62
5.25
3.88
2.5
present
reduced
absent
0.00
0.25
0.50
0.75
1.00
Distance from *M. lignano*
$\log_e$(Bit Score)
Bristle Status
present
reduced
absent

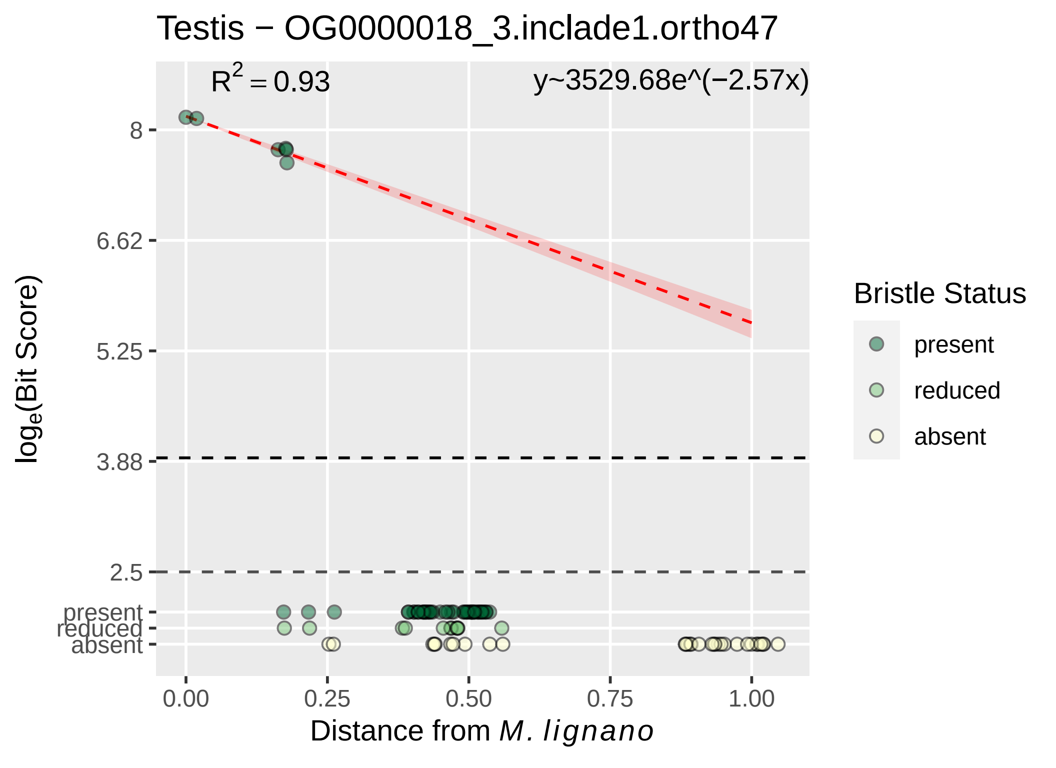

Testis − OG0000018_3.inclade1.ortho47
R^2 = 0.93
y~3529.68e^(−2.57x)
8
6.62
5.25
3.88
2.5
present
reduced
absent
log_e(Bit Score)
0.00
0.25
0.50
0.75
1.00
Distance from M. lignano
Bristle Status
present
reduced
absent

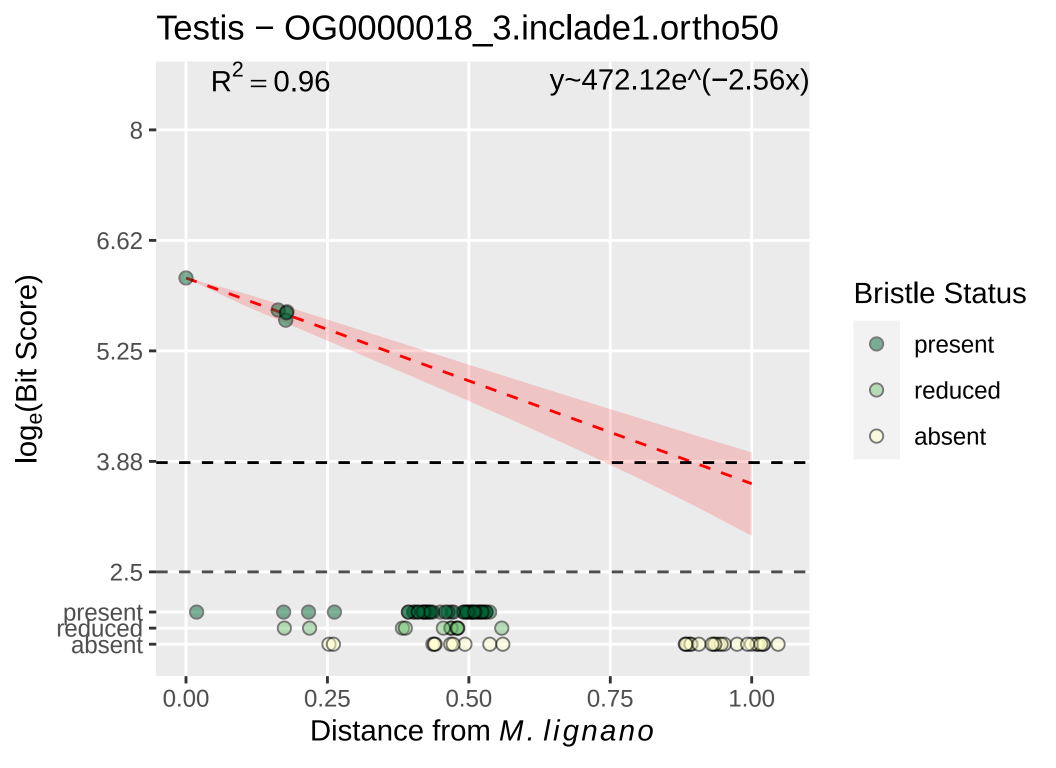

Testis − OG0000018_3.inclade1.ortho50
$R^2 = 0.96$
y~472.12e^(−2.56x)
8
6.62
5.25
3.88
2.5
present
reduced
absent
$\log_e$(Bit Score)
0.00
0.25
0.50
0.75
1.00
Distance from *M. lignano*
Bristle Status
present
reduced
absent

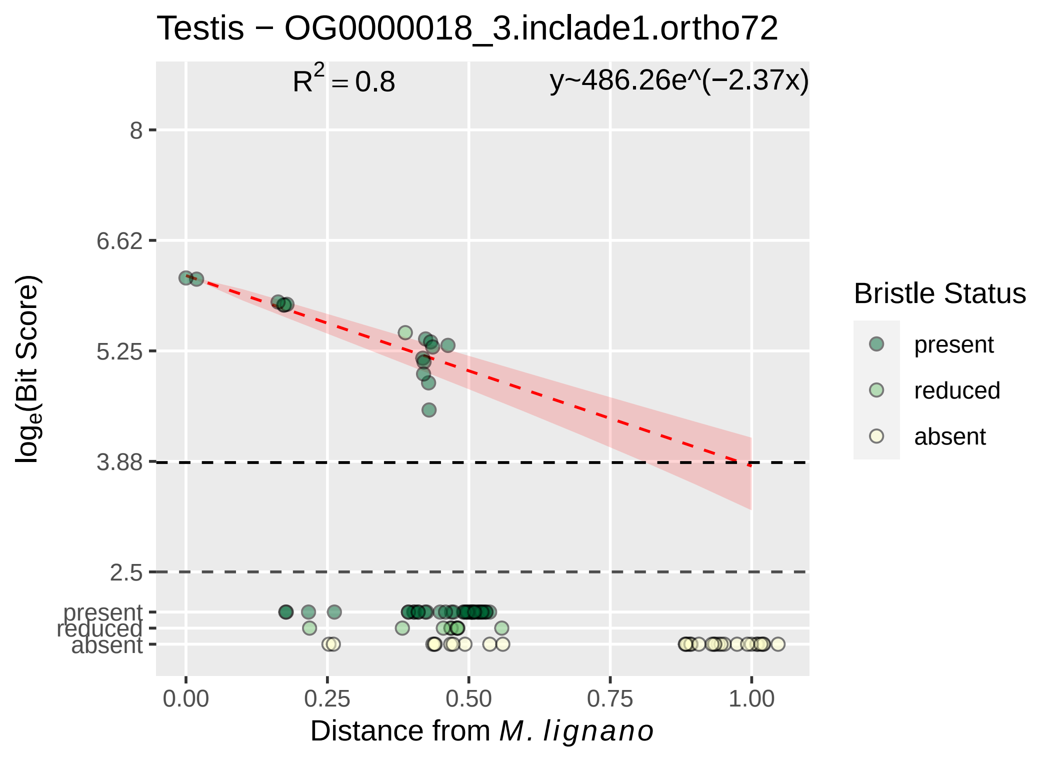

Testis − OG0000018_3.inclade1.ortho72
R^2 = 0.8
y~486.26e^(−2.37x)
8
6.62
5.25
3.88
2.5
present
reduced
absent
log_e(Bit Score)
0.00
0.25
0.50
0.75
1.00
Distance from M. lignano
Bristle Status
present
reduced
absent

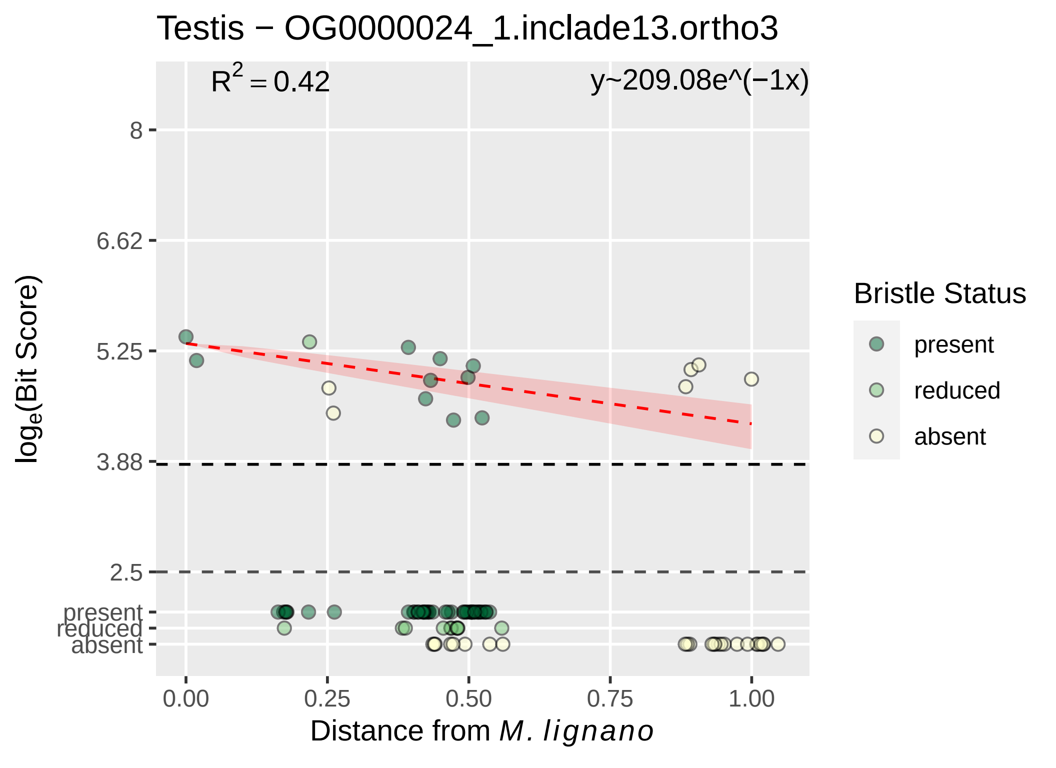

Testis − OG0000024_1.inclade13.ortho3
R2 = 0.42
y~209.08e^(−1x)
8
6.62
5.25
3.88
2.5
present
reduced
absent
0.00
0.25
0.50
0.75
1.00
Distance from M. lignano
loge(Bit Score)
Bristle Status
present
reduced
absent

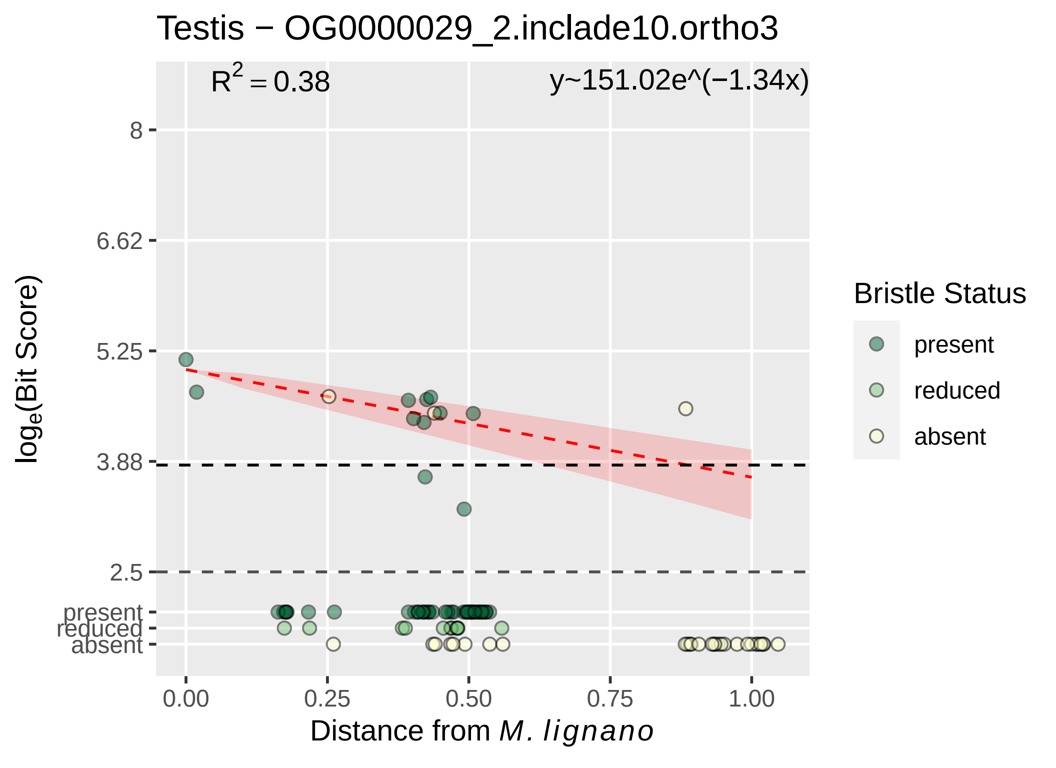

Testis − OG0000029_2.inclade10.ortho3
R² = 0.38
y~151.02e^(−1.34x)
8
6.62
5.25
3.88
2.5
present
reduced
absent
loge(Bit Score)
0.00
0.25
0.50
0.75
1.00
Distance from M. lignano
Bristle Status
present
reduced
absent

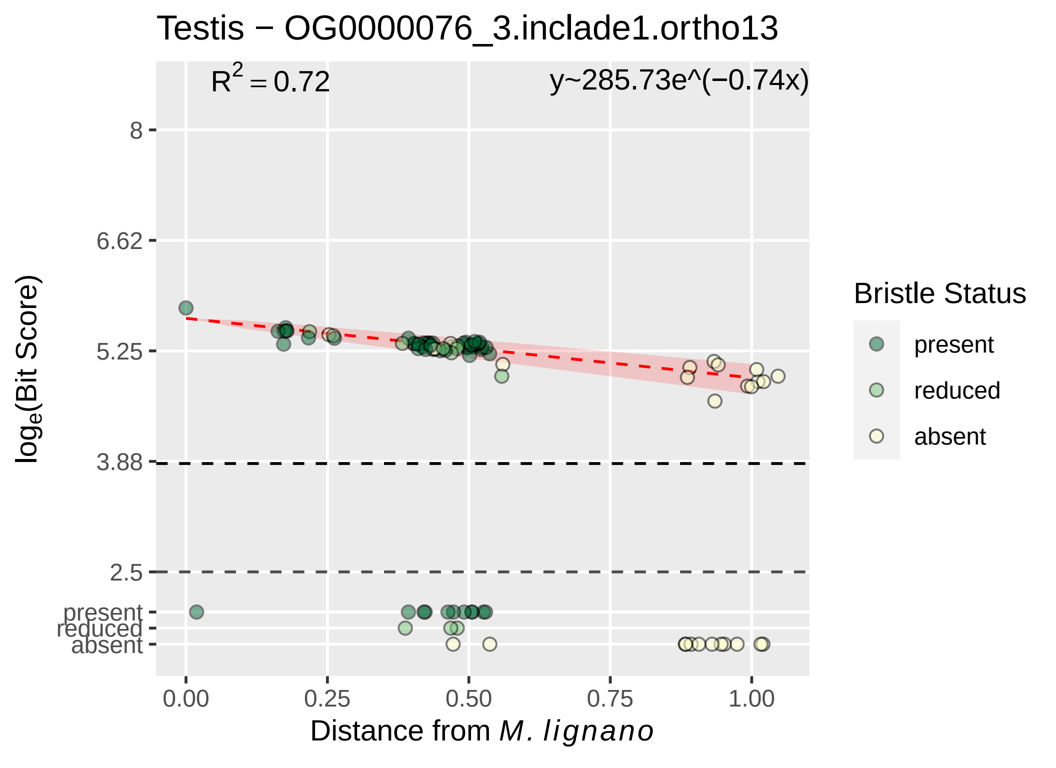

Testis − OG0000076_3.inclade1.ortho13
R^2 = 0.72
y~285.73e^(−0.74x)
8
6.62
5.25
3.88
2.5
present
reduced
absent
log_e(Bit Score)
0.00
0.25
0.50
0.75
1.00
Distance from M. lignano
Bristle Status
present
reduced
absent

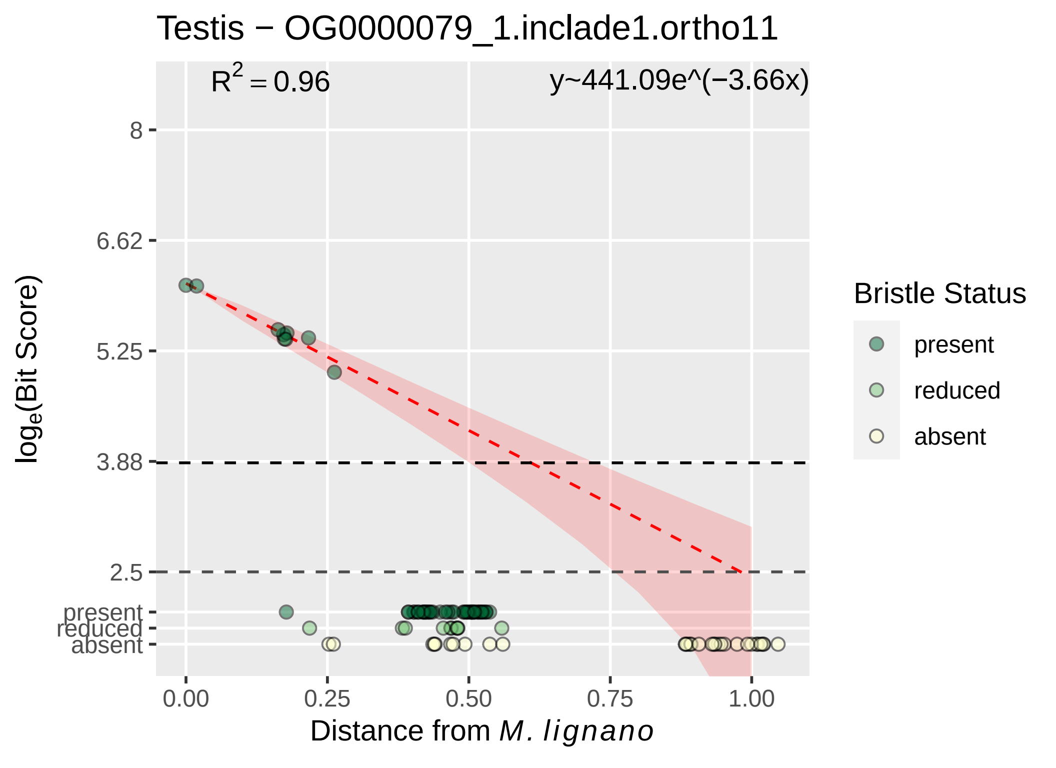

Testis − OG0000079_1.inclade1.ortho11
$R^2 = 0.96$
y~441.09e^(−3.66x)
8
6.62
5.25
3.88
2.5
present
reduced
absent
log$_e$(Bit Score)
0.00
0.25
0.50
0.75
1.00
Distance from *M. lignano*
Bristle Status
present
reduced
absent

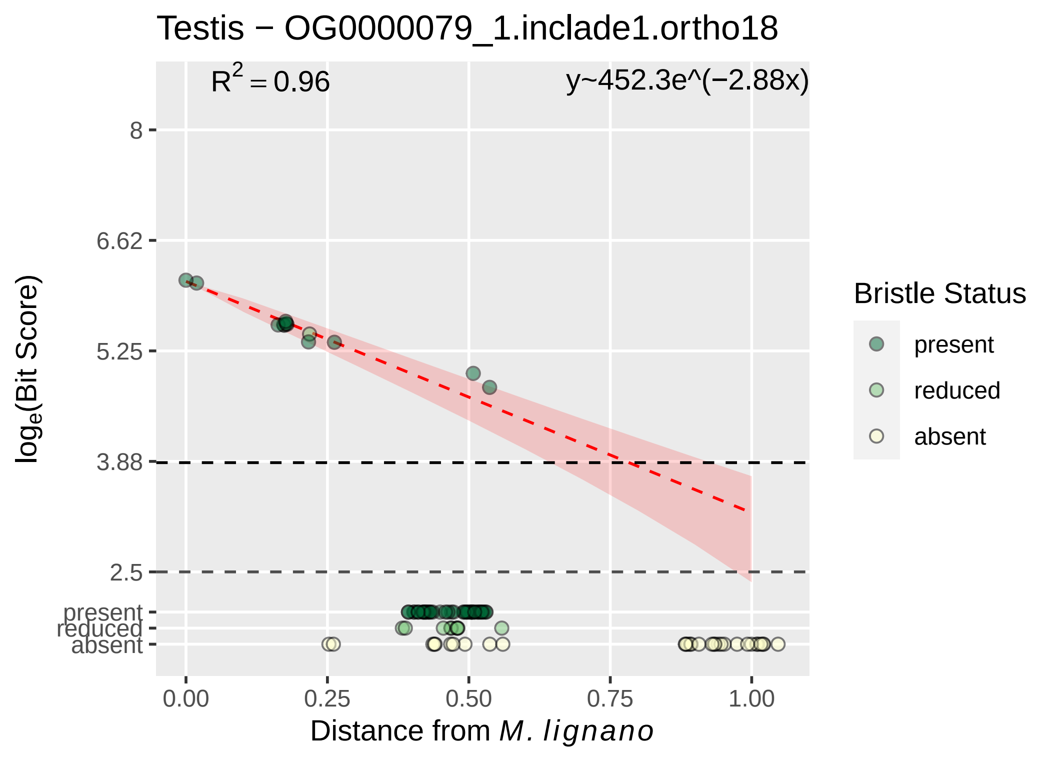
Testis − OG0000079_1.inclade1.ortho18
R^2 = 0.96
y~452.3e^(−2.88x)
8
6.62
5.25
3.88
2.5
present
reduced
absent
log_e(Bit Score)
0.00
0.25
0.50
0.75
1.00
Distance from M. lignano
Bristle Status
present
reduced
absent

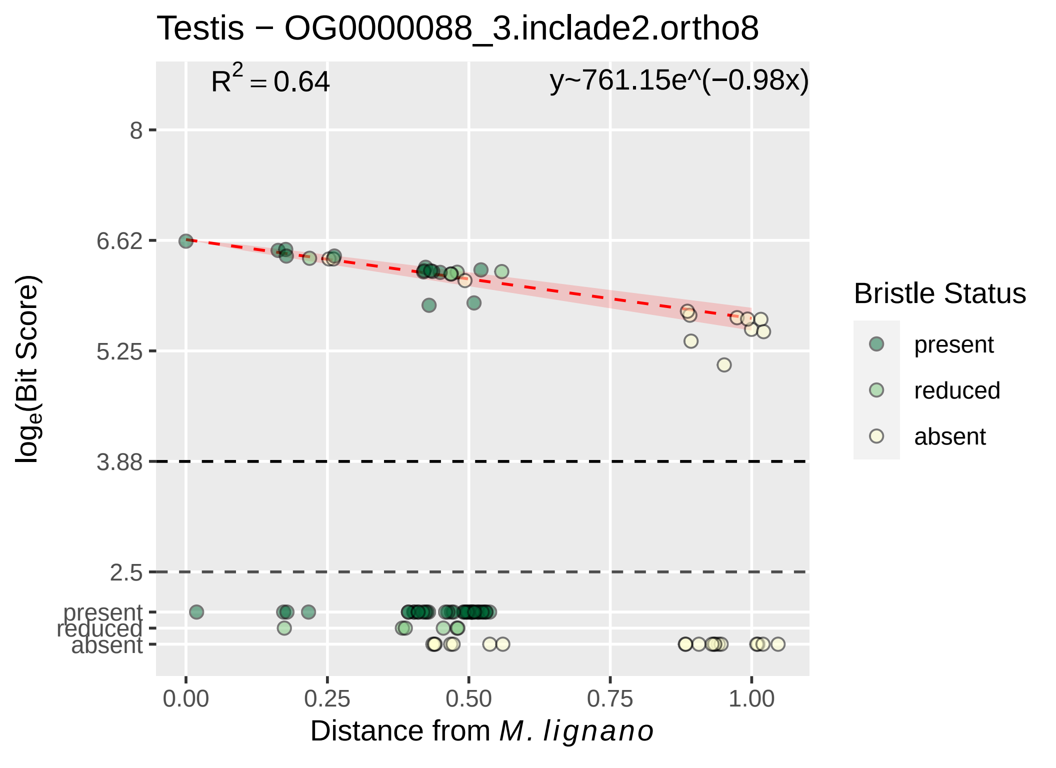

Testis − OG0000088_3.inclade2.ortho8
$R^2 = 0.64$
y~761.15e^(−0.98x)
8
6.62
5.25
3.88
2.5
present
reduced
absent
0.00
0.25
0.50
0.75
1.00
Distance from *M. lignano*
$log_e$(Bit Score)
Bristle Status
present
reduced
absent

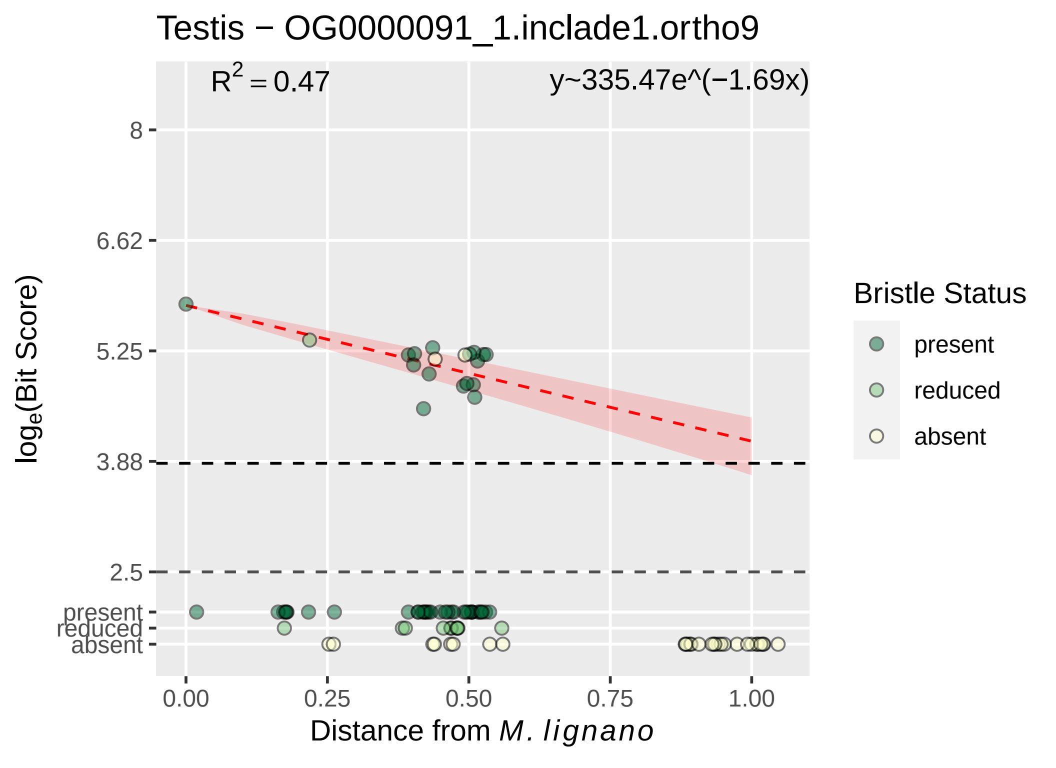
Testis − OG0000091_1.inclade1.ortho9
R² = 0.47
y~335.47e^(−1.69x)
8
6.62
5.25
3.88
2.5
present
reduced
absent
0.00
0.25
0.50
0.75
1.00
Distance from M. lignano
loge(Bit Score)
Bristle Status
present
reduced
absent

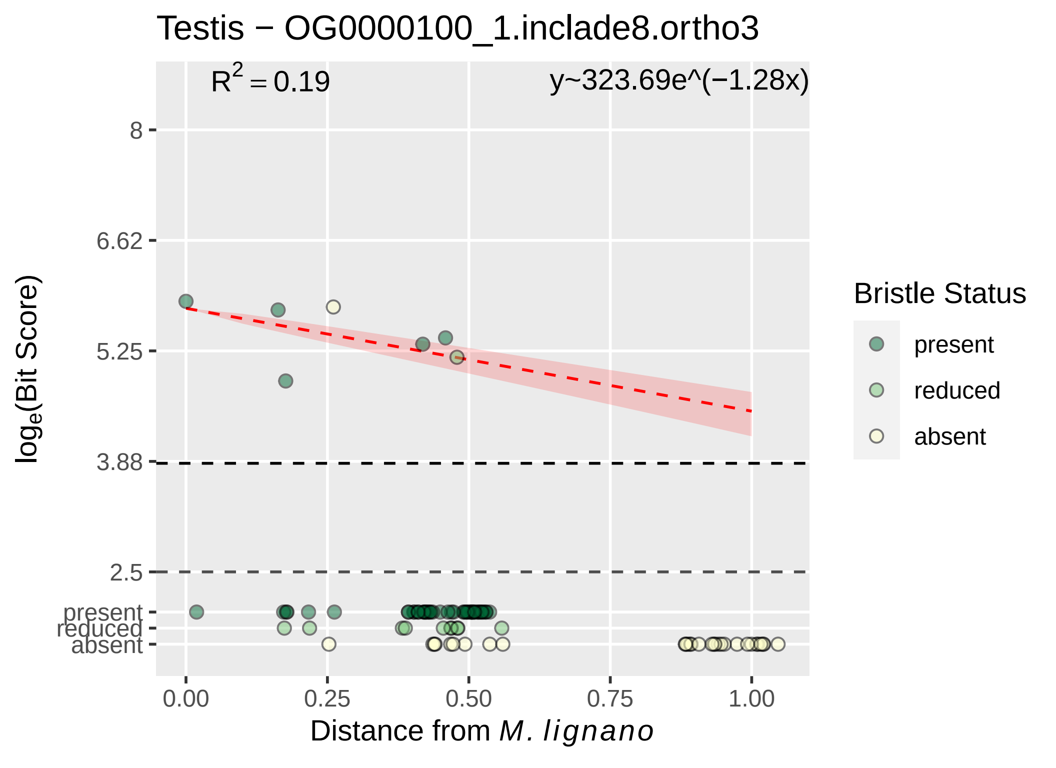

Testis − OG0000100_1.inclade8.ortho3
R^2 = 0.19
y~323.69e^(−1.28x)
8
6.62
5.25
3.88
2.5
present
reduced
absent
log_e(Bit Score)
0.00
0.25
0.50
0.75
1.00
Distance from M. lignano
Bristle Status
present
reduced
absent

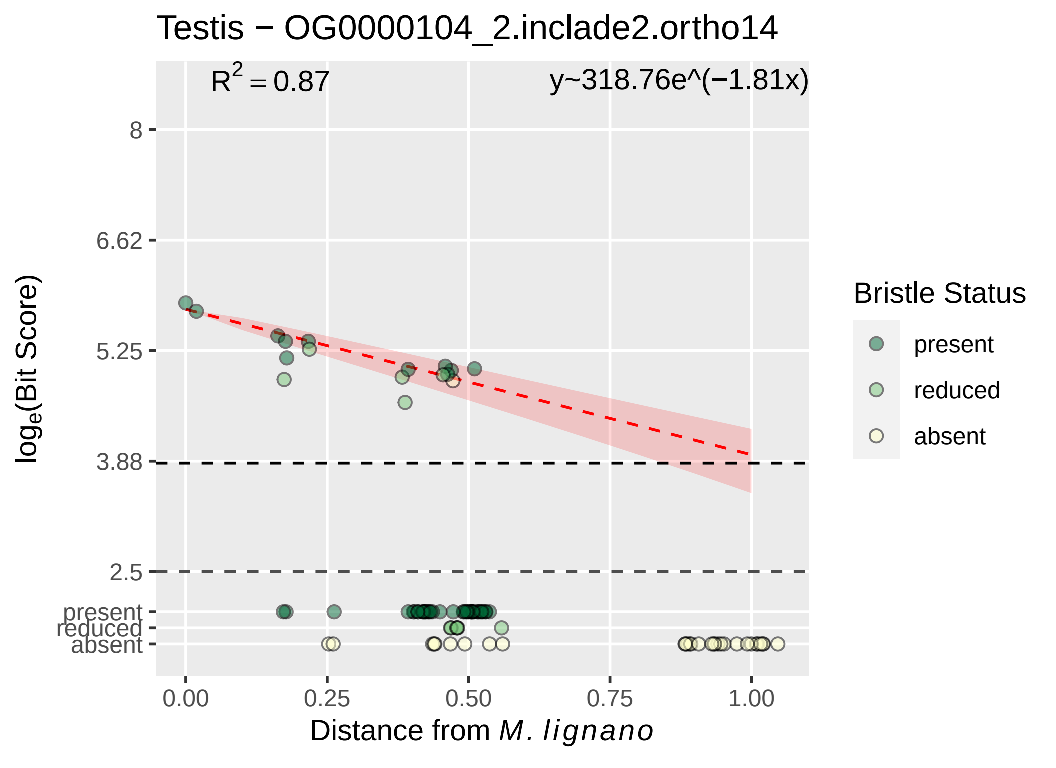

Testis − OG0000104_2.inclade2.ortho14
R^2 = 0.87
y~318.76e^(−1.81x)
8
6.62
5.25
3.88
2.5
present
reduced
absent
0.00
0.25
0.50
0.75
1.00
Distance from M. lignano
log_e(Bit Score)
Bristle Status
present
reduced
absent

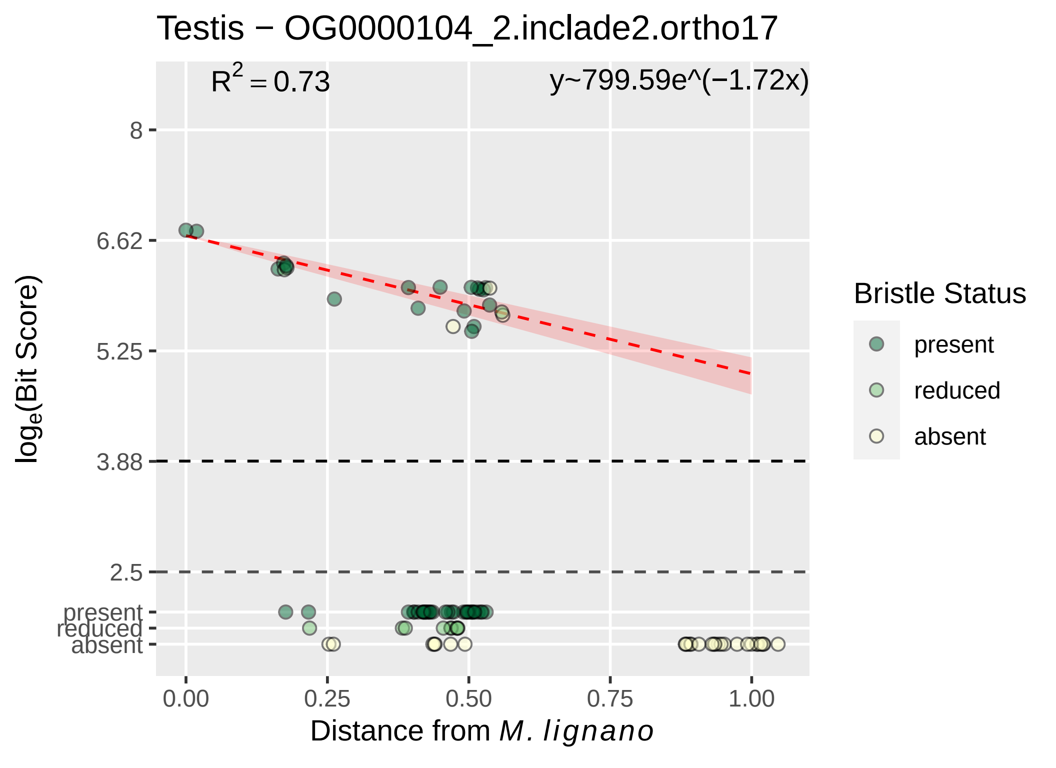

Testis − OG0000104_2.inclade2.ortho17
R^2 = 0.73
y~799.59e^(−1.72x)
8
6.62
5.25
3.88
2.5
present
reduced
absent
log_e(Bit Score)
0.00
0.25
0.50
0.75
1.00
Distance from M. lignano
Bristle Status
present
reduced
absent

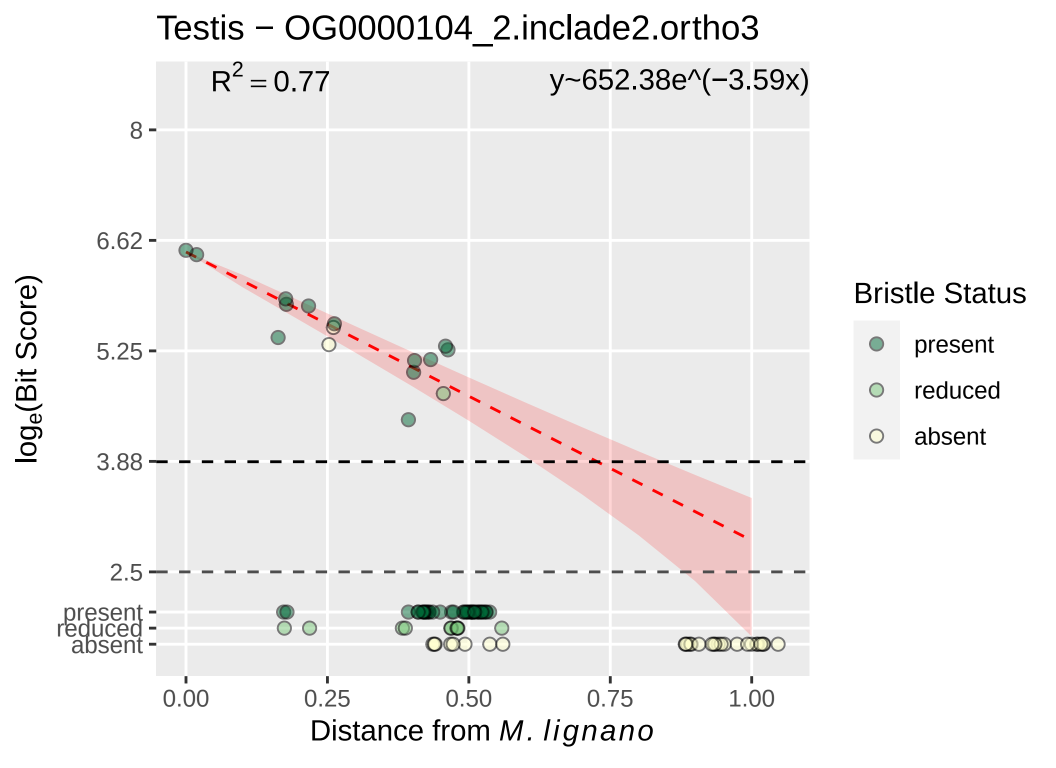

Testis − OG0000104_2.inclade2.ortho3
$R^2 = 0.77$
y~652.38e^(−3.59x)
8
6.62
5.25
3.88
2.5
present
reduced
absent
0.00
0.25
0.50
0.75
1.00
Distance from M. lignano
loge(Bit Score)
Bristle Status
present
reduced
absent

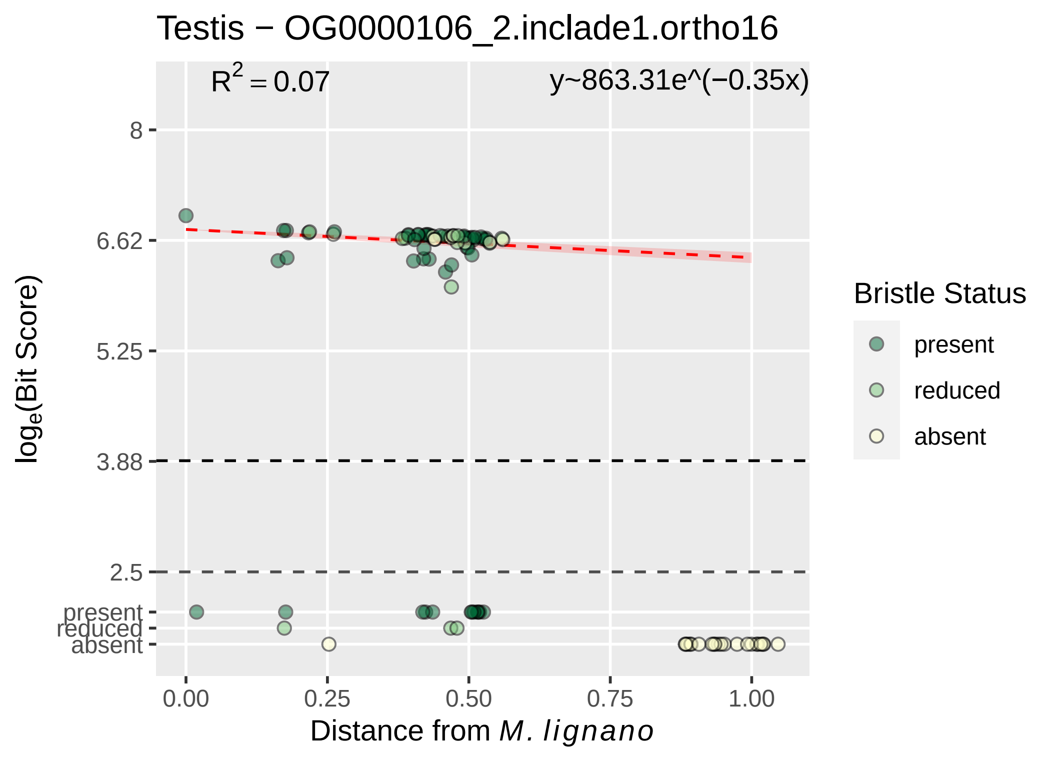

Testis − OG0000106_2.inclade1.ortho16
R^2 = 0.07
y~863.31e^(−0.35x)
8
6.62
5.25
3.88
2.5
present
reduced
absent
log_e(Bit Score)
0.00
0.25
0.50
0.75
1.00
Distance from M. lignano
Bristle Status
present
reduced
absent

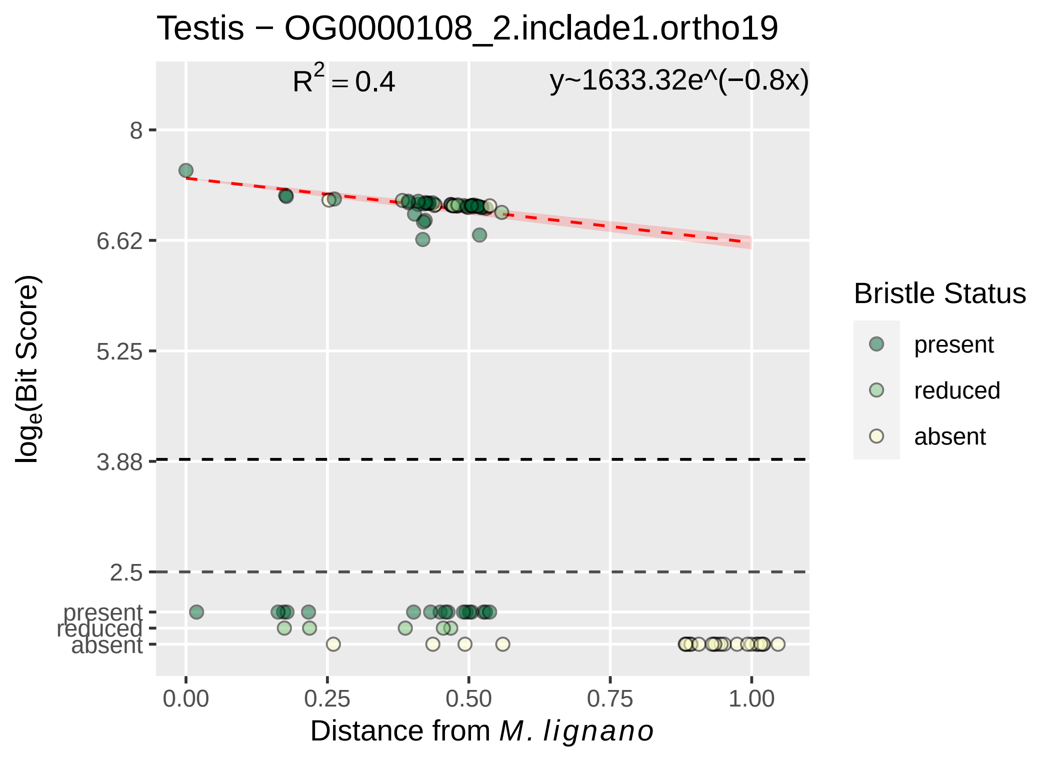
Testis − OG0000108_2.inclade1.ortho19
$R^2 = 0.4$
y~1633.32e^(−0.8x)
8
6.62
5.25
3.88
2.5
present
reduced
absent
$\log_e$(Bit Score)
0.00
0.25
0.50
0.75
1.00
Distance from *M. lignano*
Bristle Status
present
reduced
absent

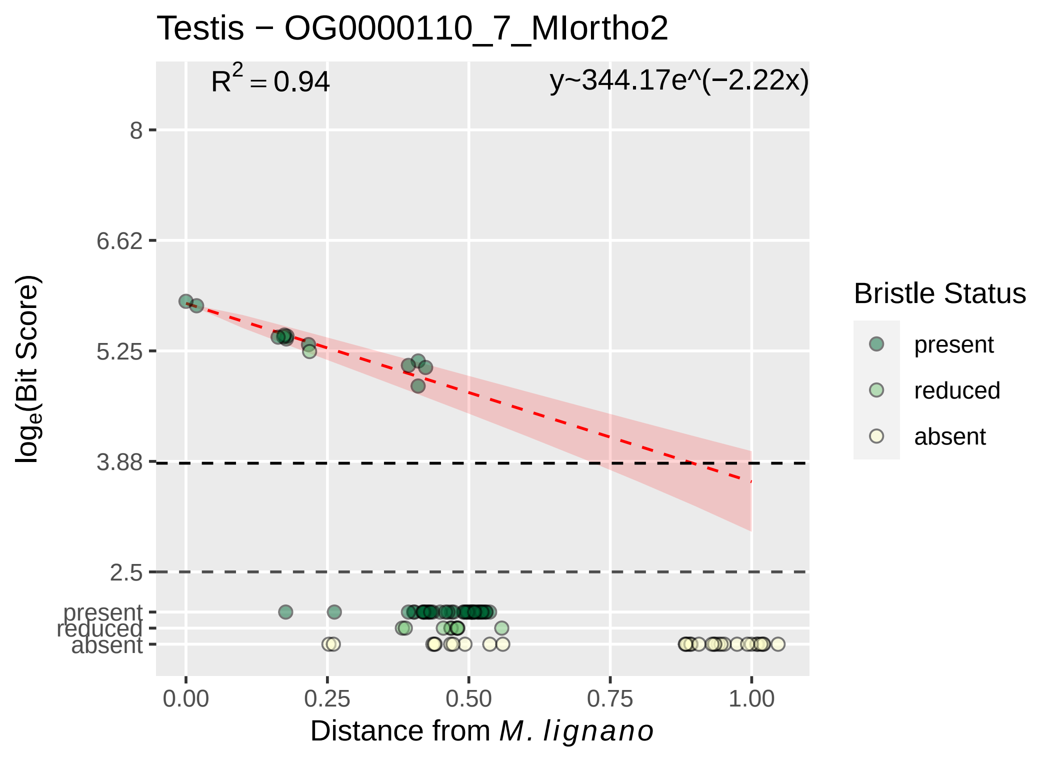

Testis − OG0000110_7_MIortho2
$R^2 = 0.94$
y~344.17e^(−2.22x)
8
6.62
5.25
3.88
2.5
present
reduced
absent
$\log_e$(Bit Score)
0.00
0.25
0.50
0.75
1.00
Distance from *M. lignano*
Bristle Status
present
reduced
absent

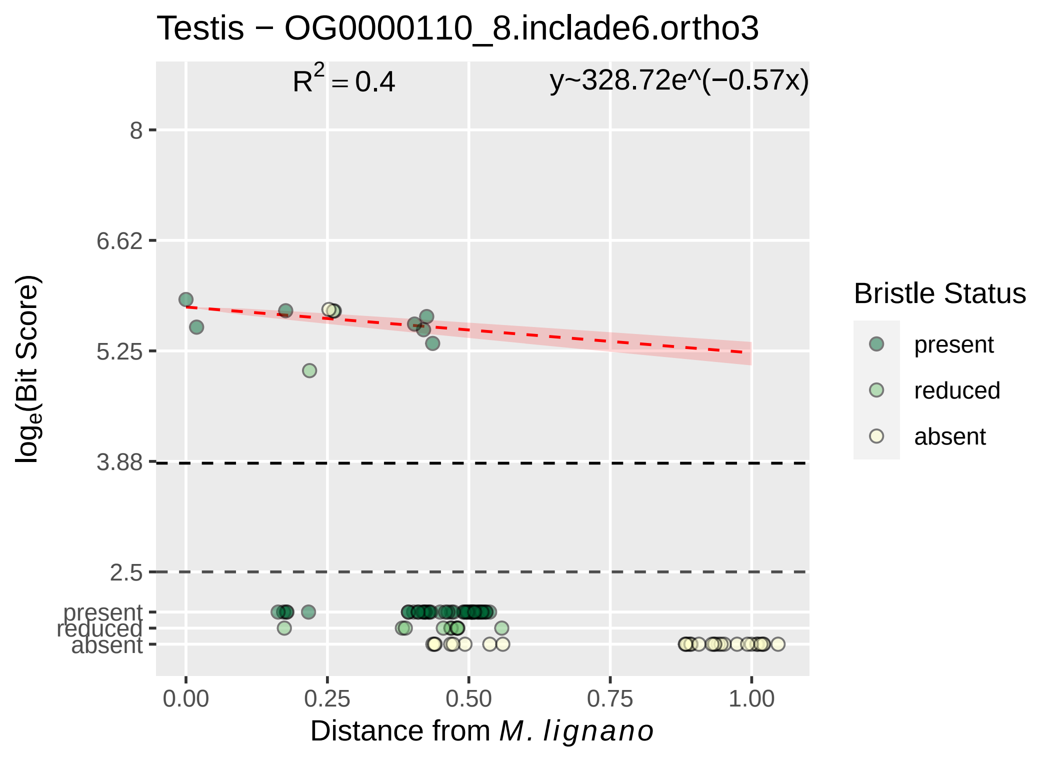

Testis − OG0000110_8.inclade6.ortho3
$R^2 = 0.4$
y~328.72e^(−0.57x)
8
6.62
5.25
3.88
2.5
present
reduced
absent
$\log_e$(Bit Score)
0.00
0.25
0.50
0.75
1.00
Distance from *M. lignano*
Bristle Status
present
reduced
absent

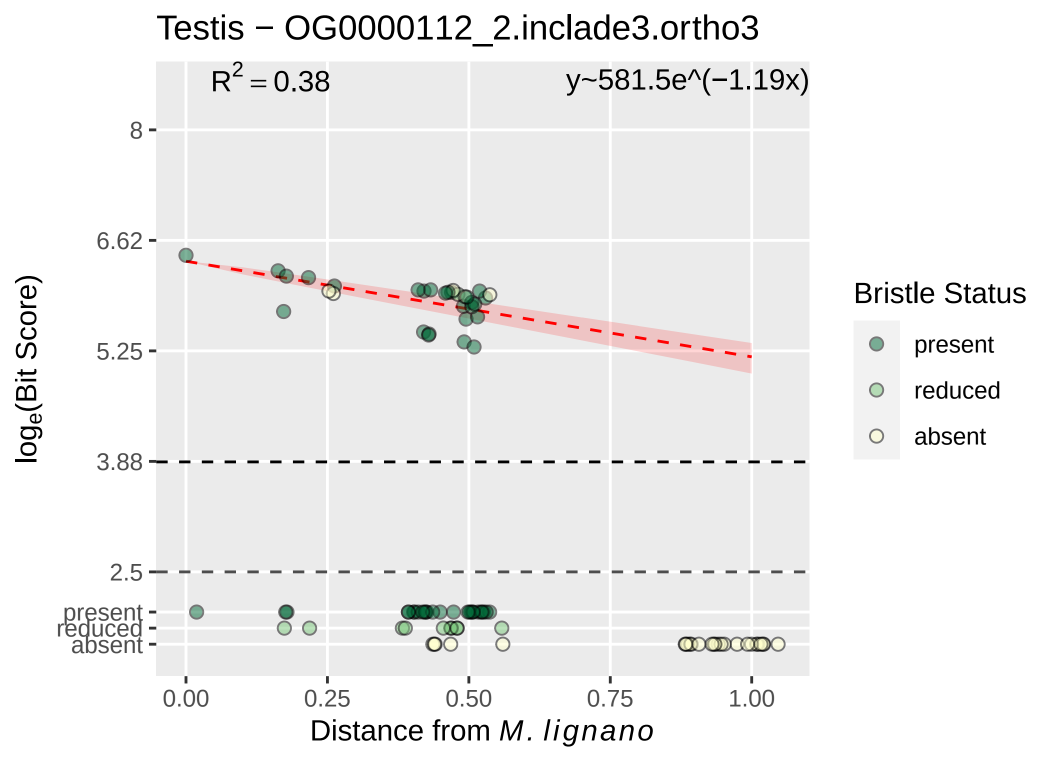

Testis − OG0000112_2.inclade3.ortho3
$R^2 = 0.38$
y~581.5e^(−1.19x)
8
6.62
5.25
3.88
2.5
present
reduced
absent
log$_e$(Bit Score)
0.00
0.25
0.50
0.75
1.00
Distance from *M. lignano*
Bristle Status
present
reduced
absent

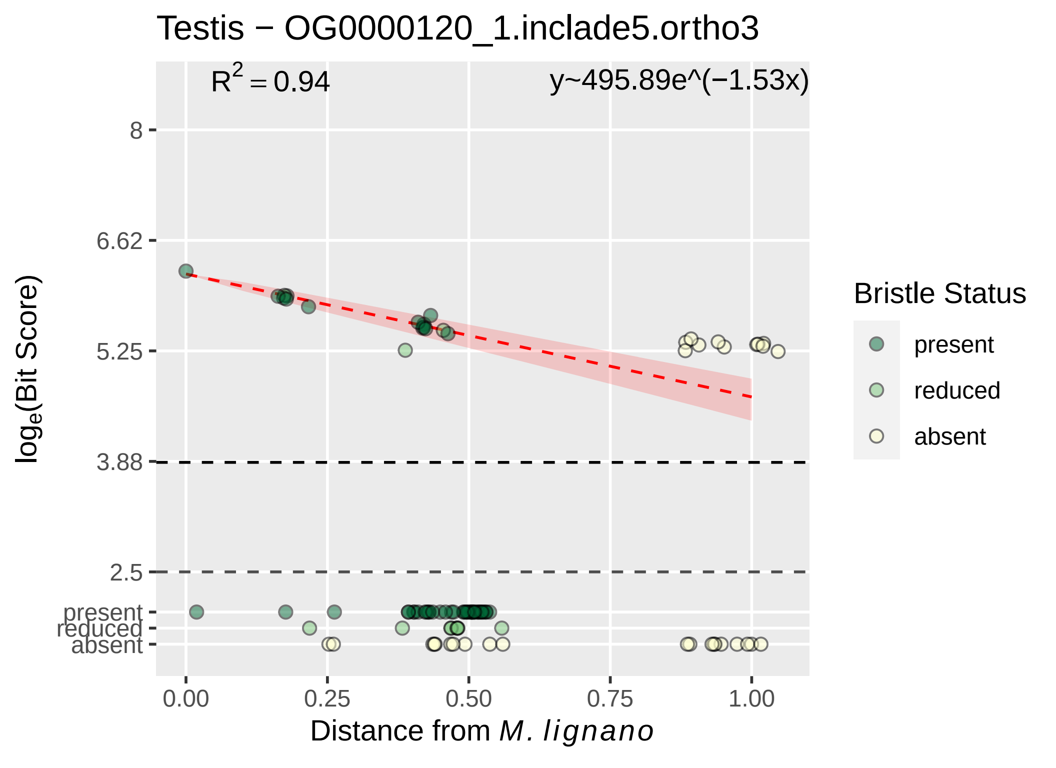

Testis − OG0000120_1.inclade5.ortho3
$R^2 = 0.94$
y~495.89e^(−1.53x)
8
6.62
5.25
3.88
2.5
present
reduced
absent
$log_e$(Bit Score)
0.00
0.25
0.50
0.75
1.00
Distance from *M. lignano*
Bristle Status
present
reduced
absent

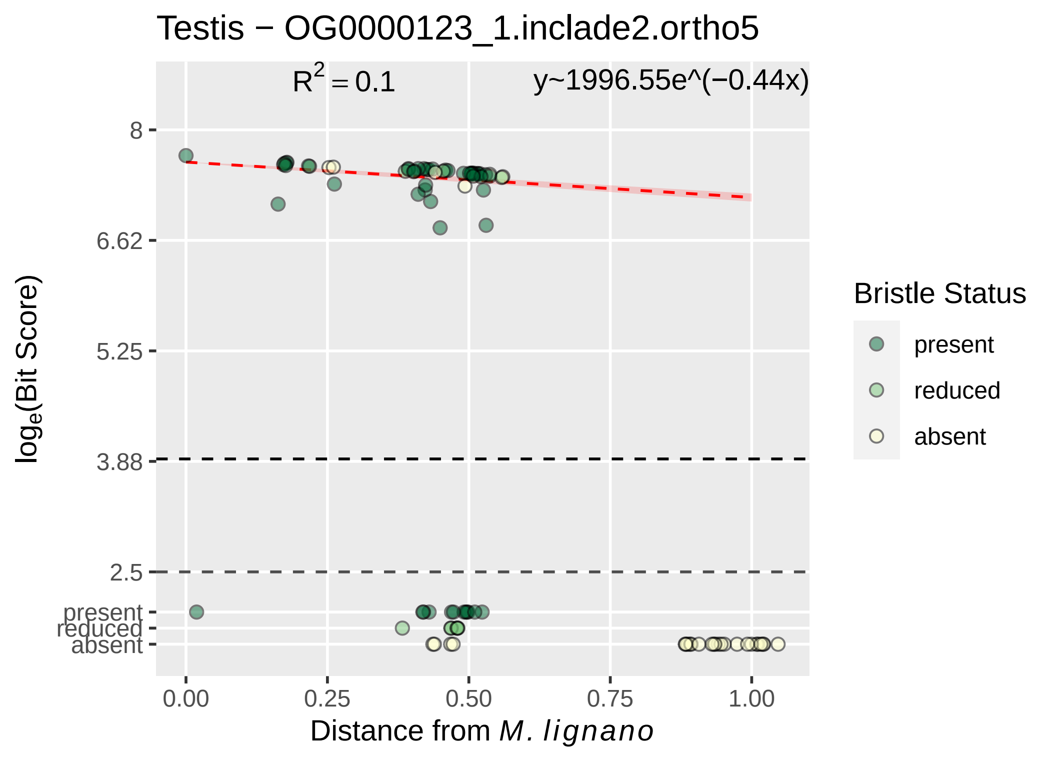

Testis − OG0000123_1.inclade2.ortho5
R² = 0.1
y~1996.55e^(−0.44x)
8
6.62
5.25
3.88
2.5
present
reduced
absent
log_e(Bit Score)
0.00
0.25
0.50
0.75
1.00
Distance from M. lignano
Bristle Status
present
reduced
absent

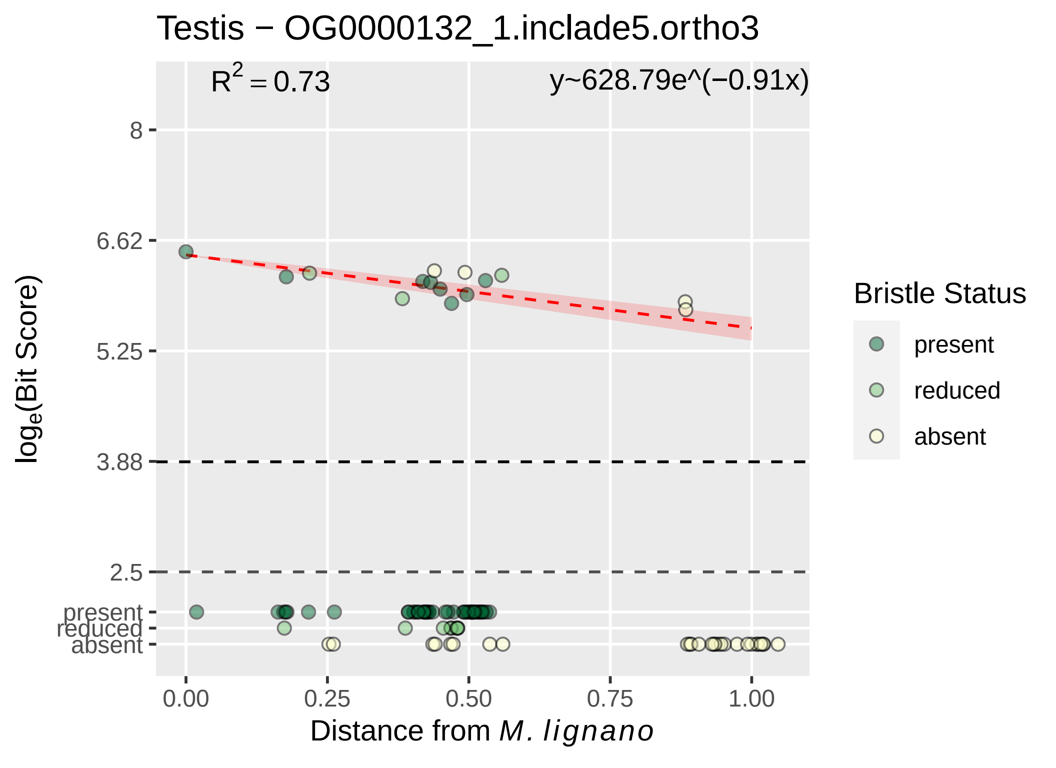

Testis − OG0000132_1.inclade5.ortho3
$R^2 = 0.73$
y~628.79e^(−0.91x)
8
6.62
5.25
3.88
2.5
present
reduced
absent
0.00
0.25
0.50
0.75
1.00
Distance from M. lignano
loge(Bit Score)
Bristle Status
present
reduced
absent

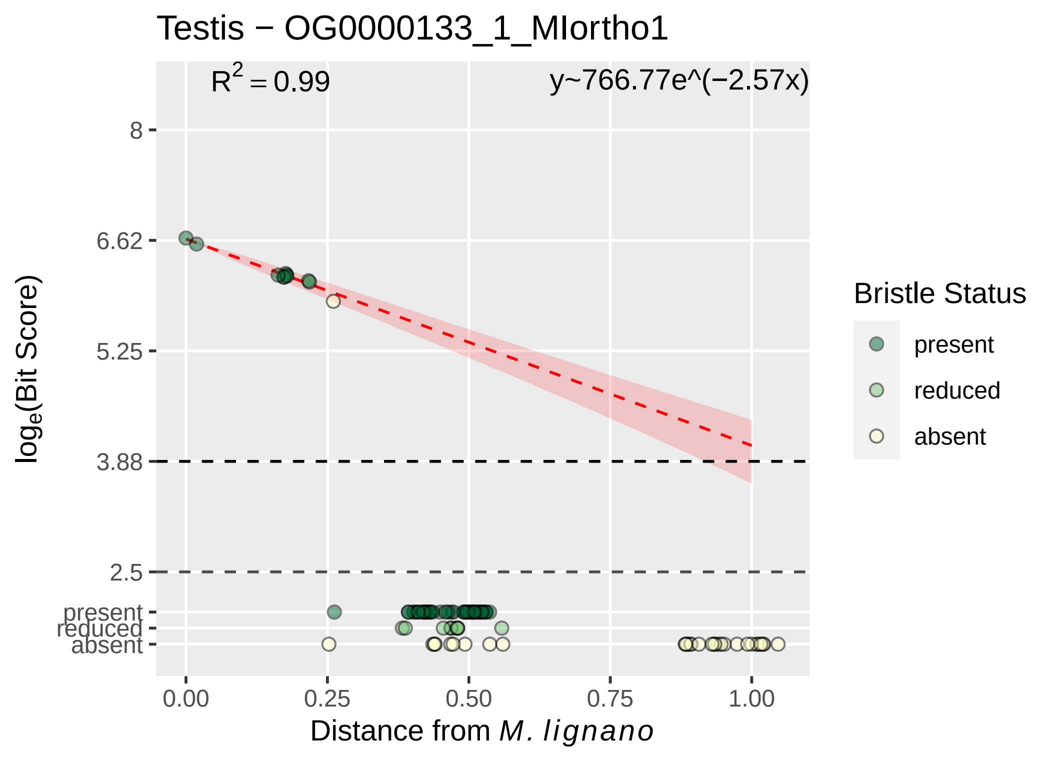
Testis − OG0000133_1_MIortho1
$R^2 = 0.99$
y~766.77e^(−2.57x)
8
6.62
5.25
3.88
2.5
present
reduced
absent
$\log_e$(Bit Score)
0.00
0.25
0.50
0.75
1.00
Distance from *M. lignano*
Bristle Status
present
reduced
absent

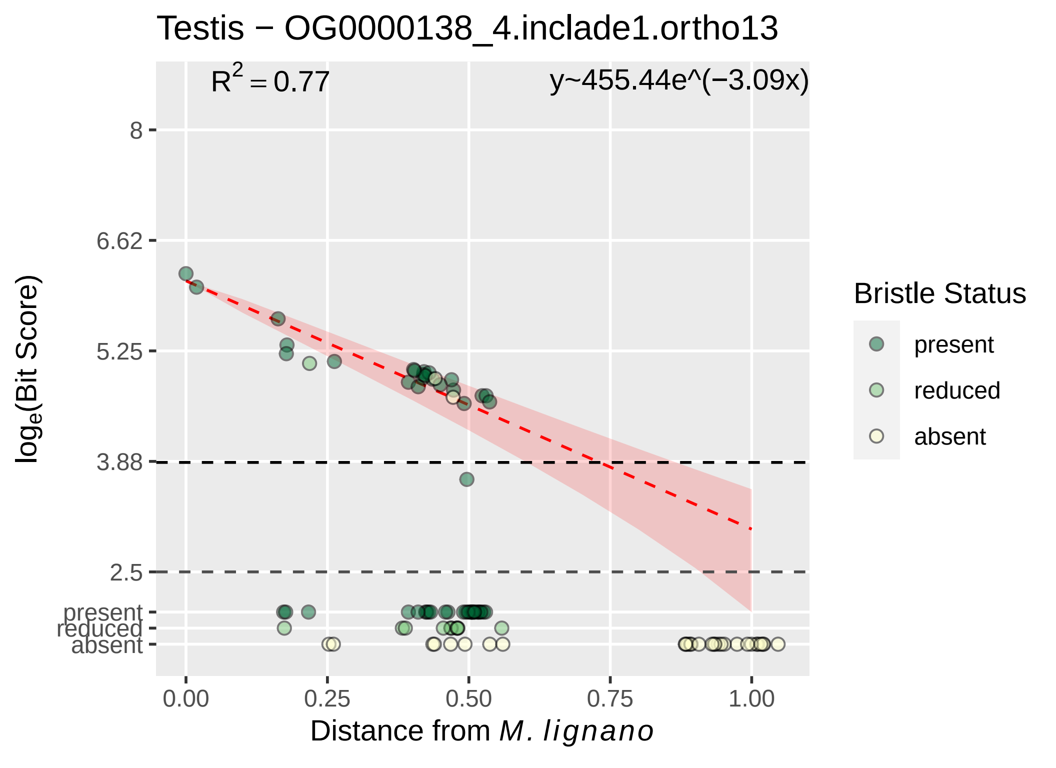

Testis − OG0000138_4.inclade1.ortho13
$R^2 = 0.77$
y~455.44e^(−3.09x)
8
6.62
5.25
3.88
2.5
present
reduced
absent
$log_e$(Bit Score)
0.00
0.25
0.50
0.75
1.00
Distance from M. lignano
Bristle Status
present
reduced
absent

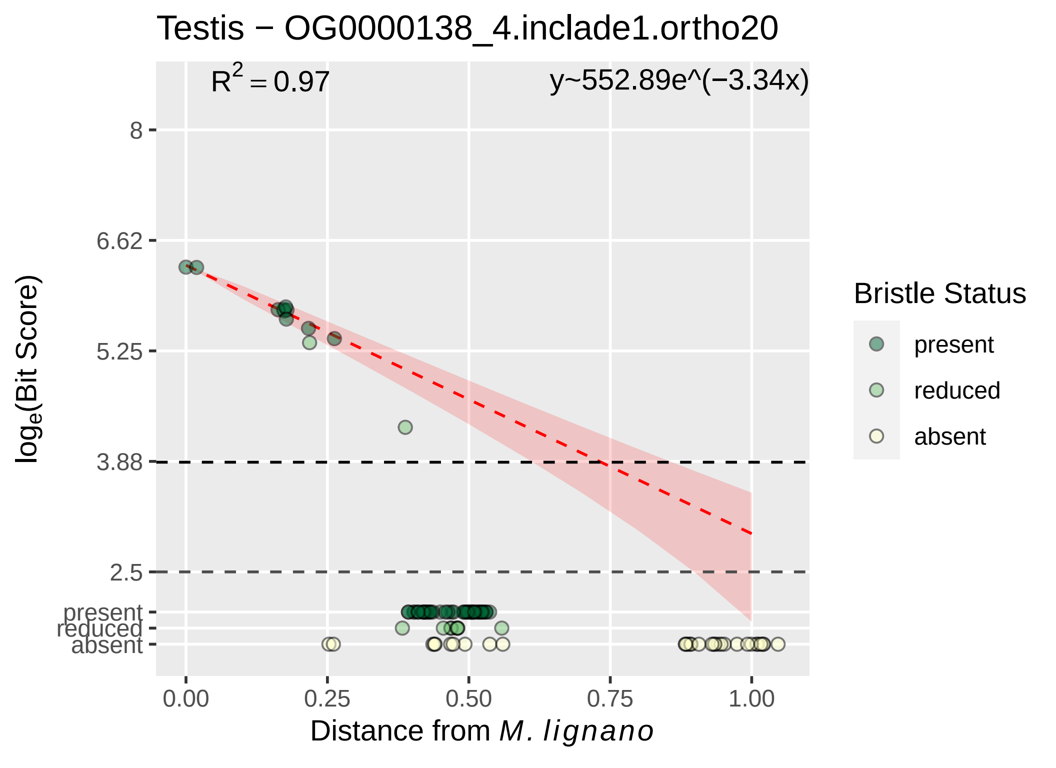
Testis − OG0000138_4.inclade1.ortho20
R² = 0.97
y~552.89e^(−3.34x)
8
6.62
5.25
3.88
2.5
present
reduced
absent
log_e(Bit Score)
0.00
0.25
0.50
0.75
1.00
Distance from M. lignano
Bristle Status
present
reduced
absent

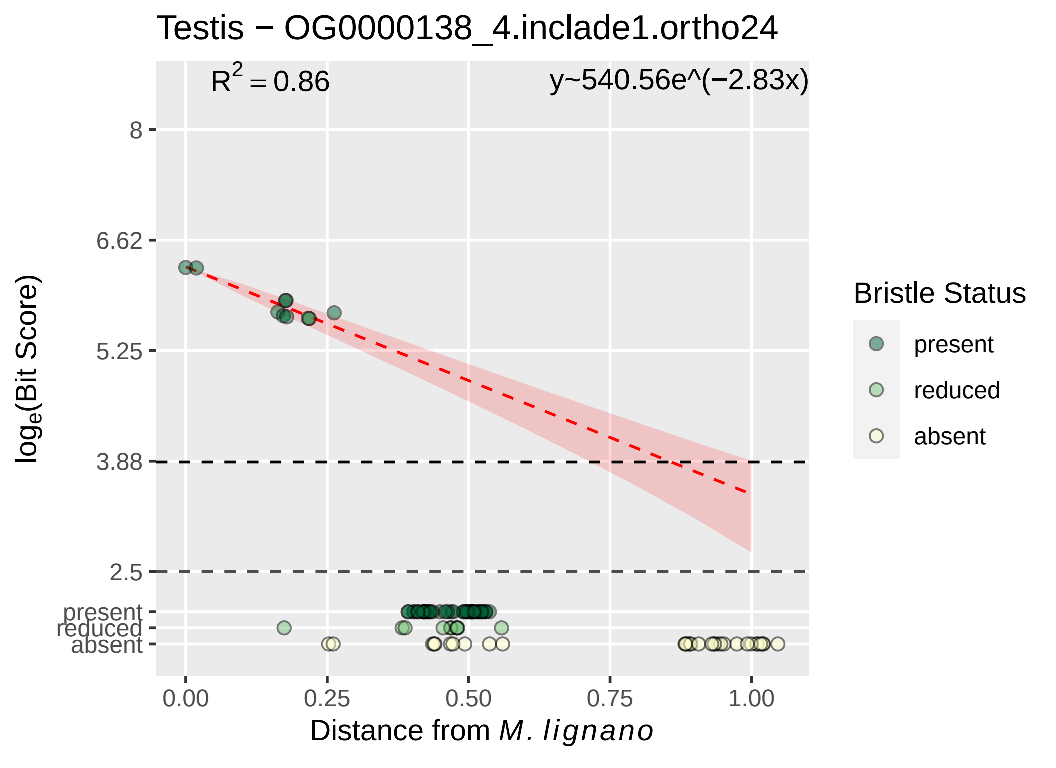
Testis − OG0000138_4.inclade1.ortho24
R2 = 0.86
y~540.56e^(−2.83x)
8
6.62
5.25
3.88
2.5
present
reduced
absent
loge(Bit Score)
0.00
0.25
0.50
0.75
1.00
Distance from M. lignano
Bristle Status
present
reduced
absent

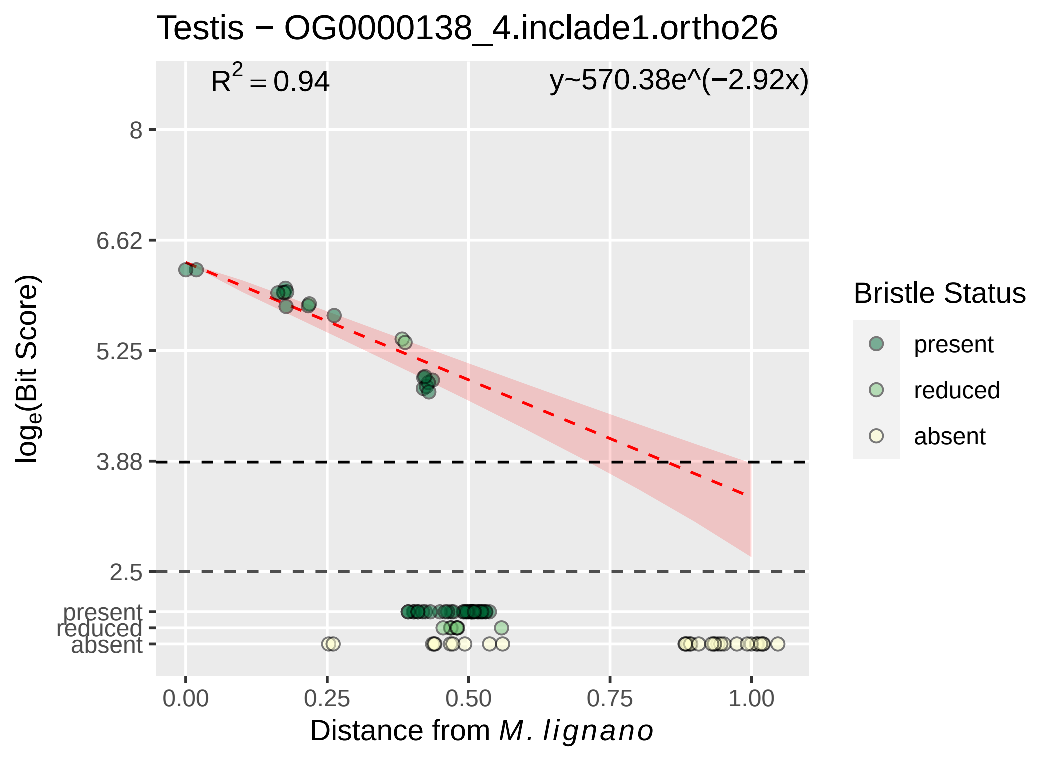

Testis − OG0000138_4.inclade1.ortho26
$R^2 = 0.94$
y~570.38e^(−2.92x)
8
6.62
5.25
3.88
2.5
present
reduced
absent
0.00
0.25
0.50
0.75
1.00
log_e(Bit Score)
Distance from M. lignano
Bristle Status
present
reduced
absent

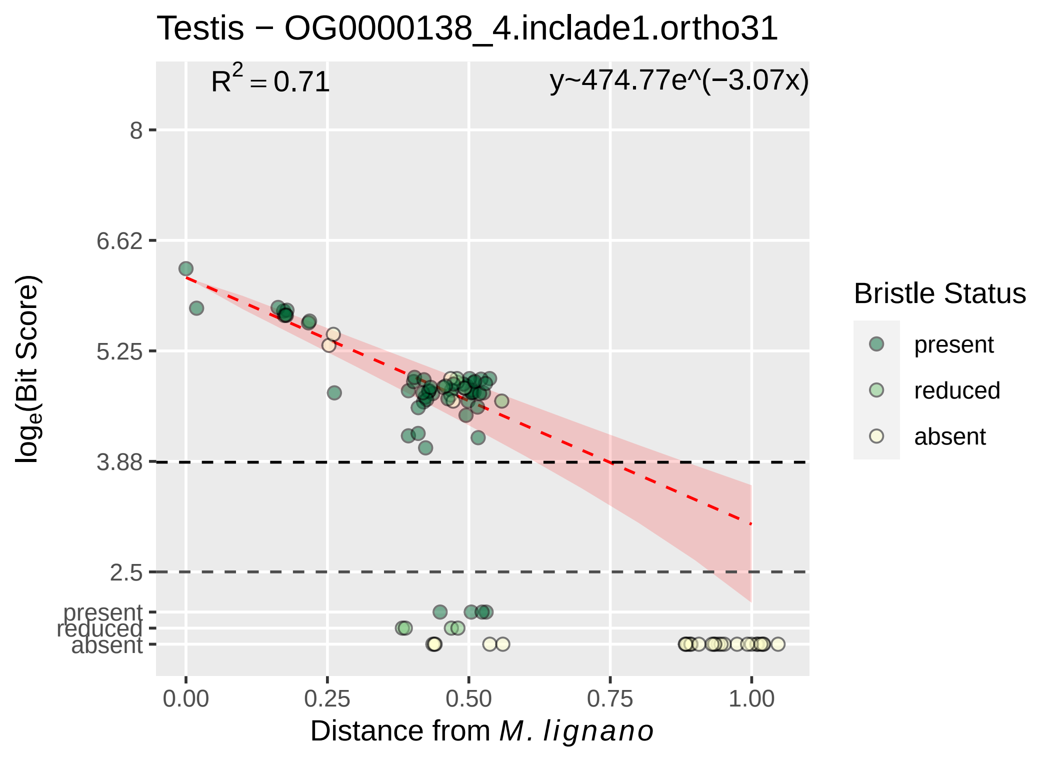

Testis − OG0000138_4.inclade1.ortho31
$R^2 = 0.71$
y~474.77e^(−3.07x)
log$_e$(Bit Score)
8
6.62
5.25
3.88
2.5
present
reduced
absent
0.00
0.25
0.50
0.75
1.00
Distance from *M. lignano*
Bristle Status
present
reduced
absent

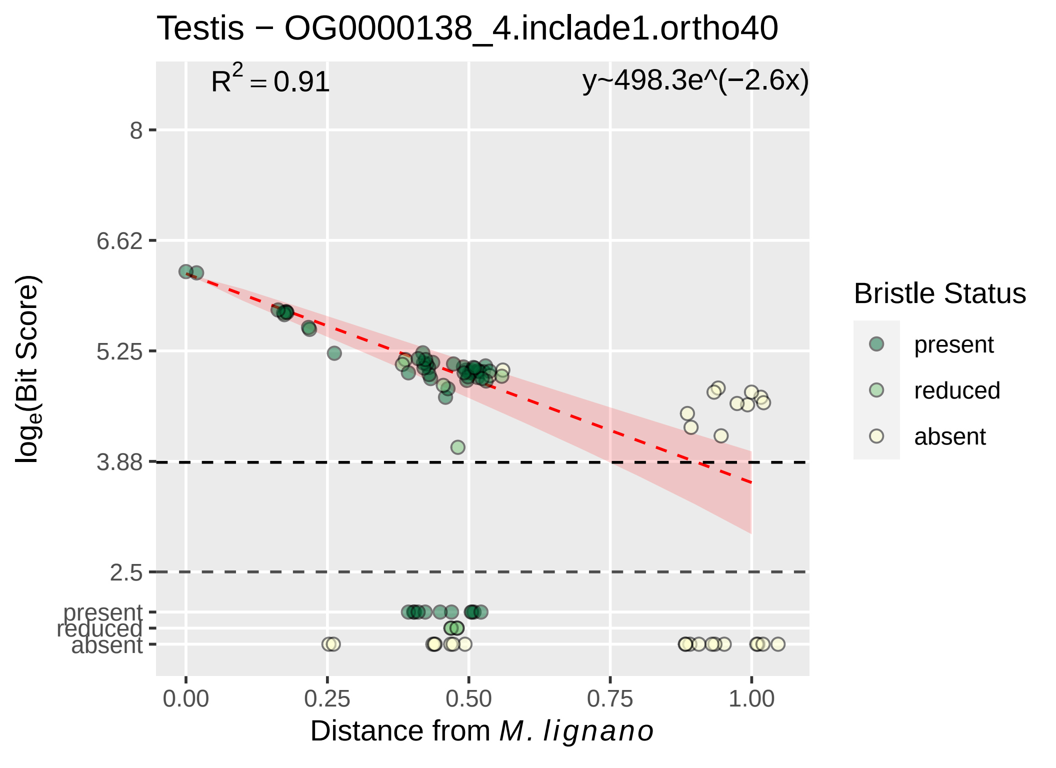

Testis − OG0000138_4.inclade1.ortho40
$R^2 = 0.91$
y~498.3e^(−2.6x)
8
6.62
5.25
3.88
2.5
present
reduced
absent
0.00
0.25
0.50
0.75
1.00
Distance from M. lignano
log_e(Bit Score)
Bristle Status
present
reduced
absent

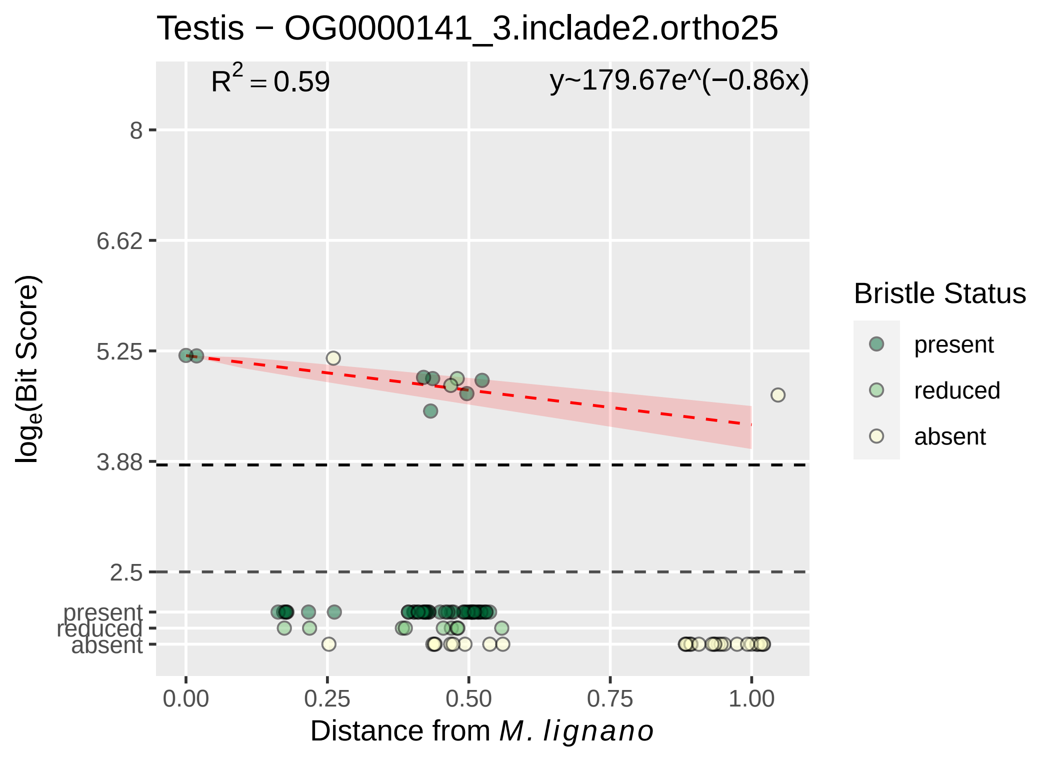

Testis − OG0000141_3.inclade2.ortho25
$R^2 = 0.59$
y~179.67e^(−0.86x)
8
6.62
5.25
3.88
2.5
present
reduced
absent
0.00
0.25
0.50
0.75
1.00
$log_e$(Bit Score)
Distance from *M. lignano*
Bristle Status
present
reduced
absent

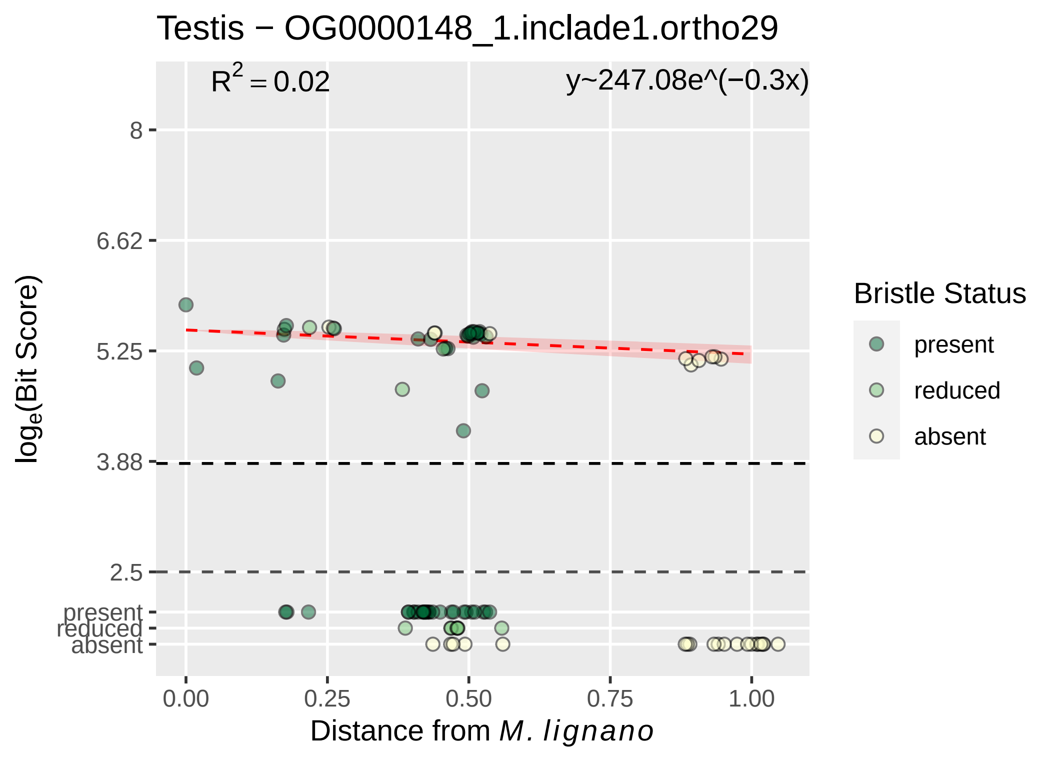

Testis − OG0000148_1.inclade1.ortho29
$R^2 = 0.02$
y~247.08e^(−0.3x)
8
6.62
5.25
3.88
2.5
present
reduced
absent
$\log_e$(Bit Score)
0.00
0.25
0.50
0.75
1.00
Distance from *M. lignano*
Bristle Status
present
reduced
absent

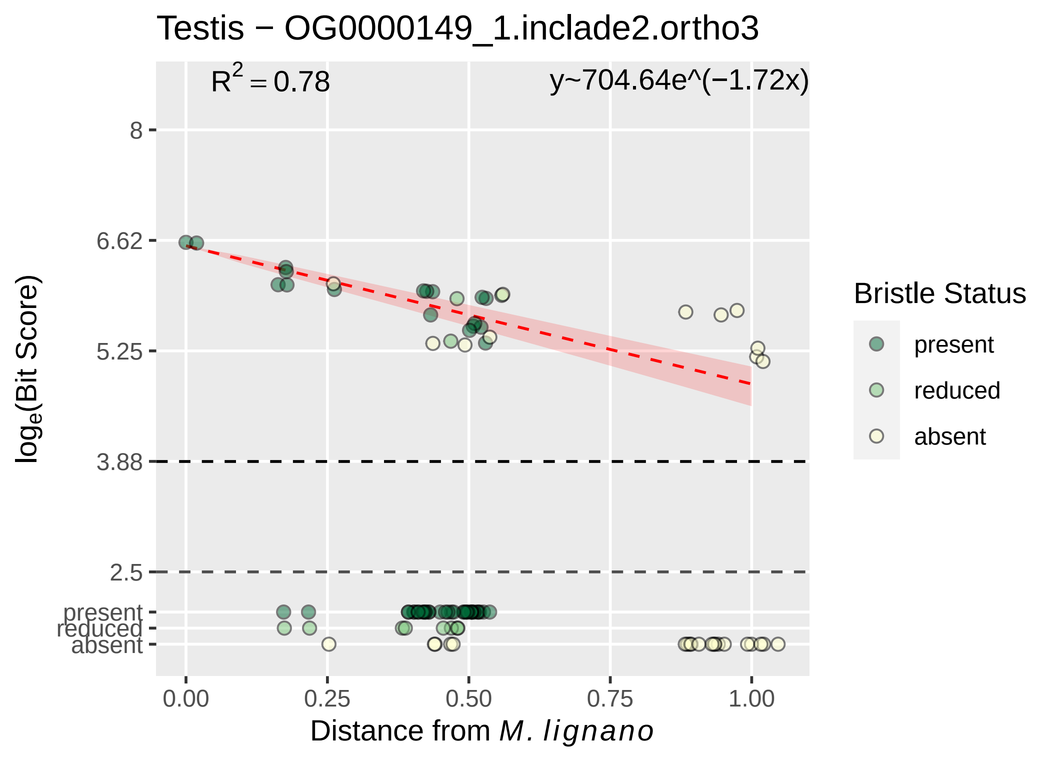

Testis − OG0000149_1.inclade2.ortho3
R2 = 0.78
y~704.64e^(−1.72x)
8
6.62
5.25
3.88
2.5
present
reduced
absent
loge(Bit Score)
0.00
0.25
0.50
0.75
1.00
Distance from M. lignano
Bristle Status
present
reduced
absent

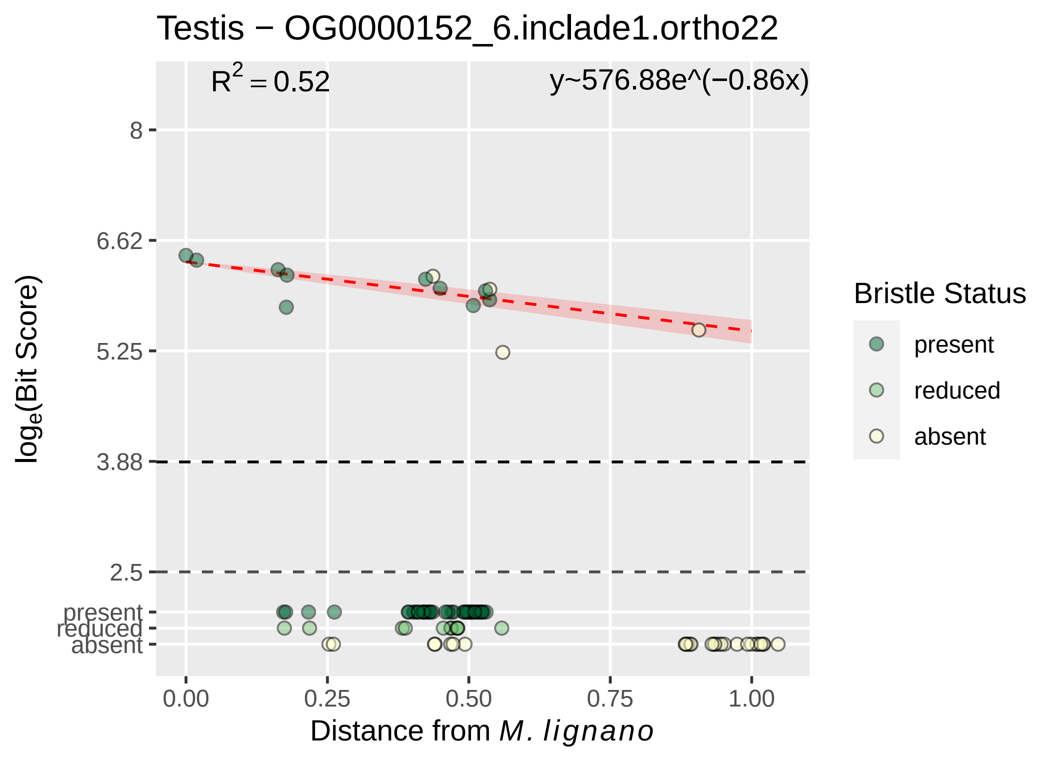

Testis − OG0000152_6.inclade1.ortho22
R2 = 0.52
y~576.88e^(−0.86x)
8
6.62
5.25
3.88
2.5
present
reduced
absent
loge(Bit Score)
0.00
0.25
0.50
0.75
1.00
Distance from M. lignano
Bristle Status
present
reduced
absent

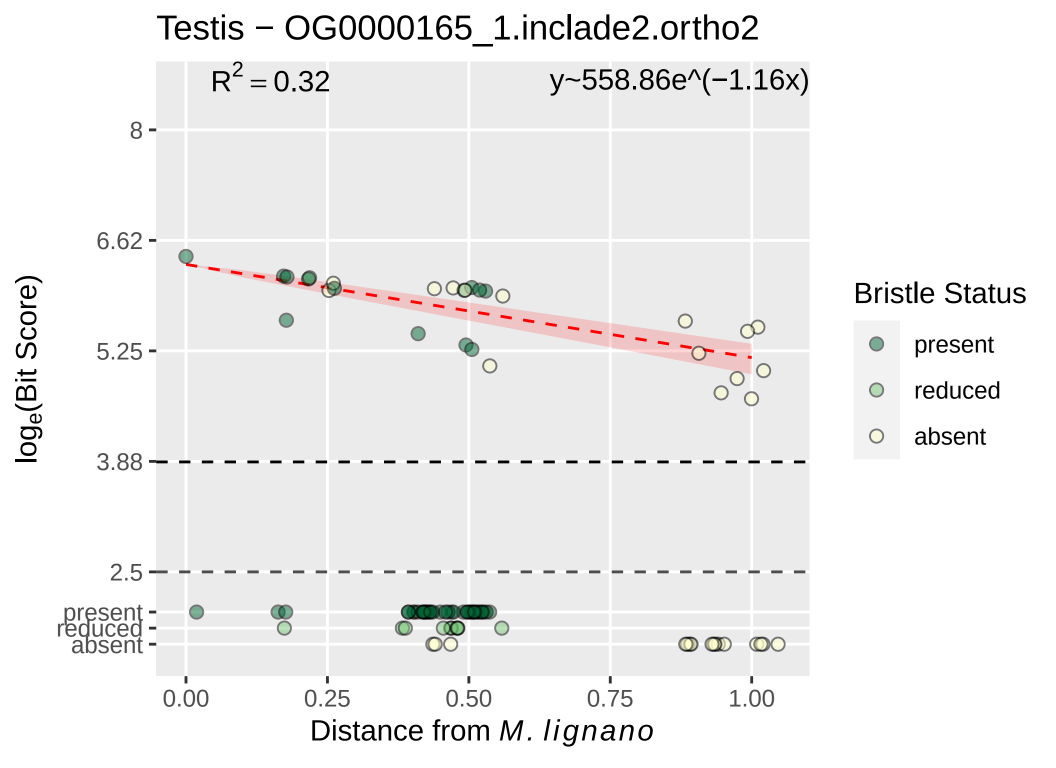

Testis − OG0000165_1.inclade2.ortho2
$R^2 = 0.32$
y~558.86e^(−1.16x)
8
6.62
5.25
3.88
2.5
present
reduced
absent
$log_e$(Bit Score)
0.00
0.25
0.50
0.75
1.00
Distance from *M. lignano*
Bristle Status
present
reduced
absent

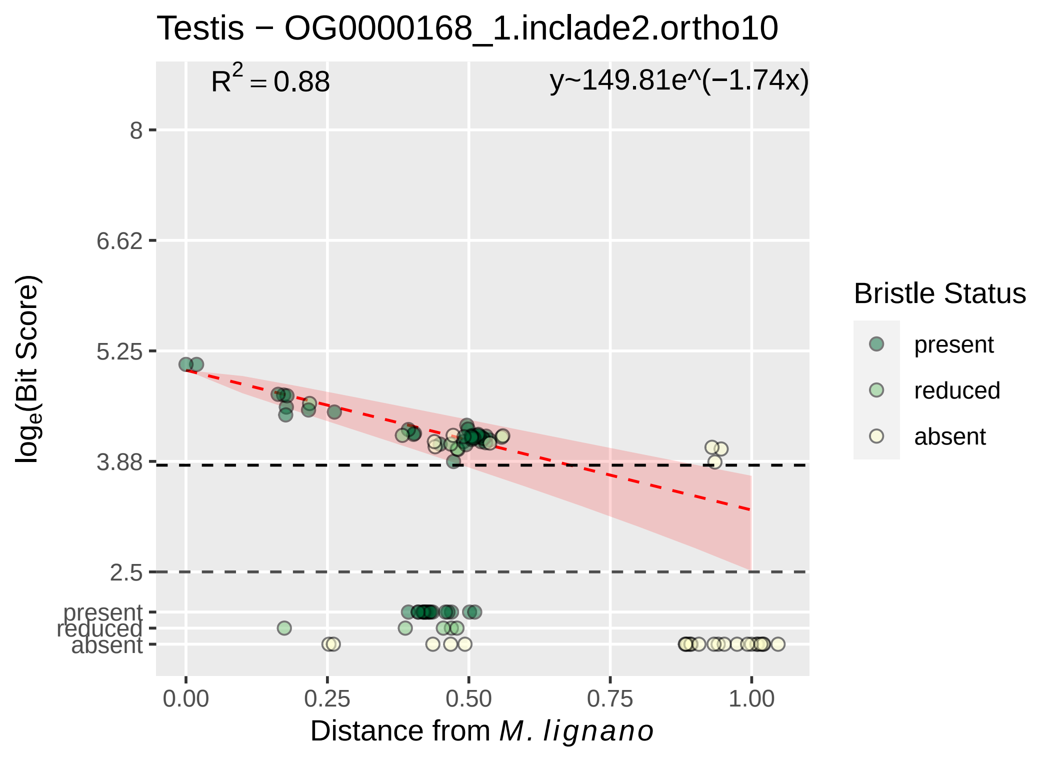

Testis − OG0000168_1.inclade2.ortho10
R^2 = 0.88
y~149.81e^(−1.74x)
8
6.62
5.25
3.88
2.5
present
reduced
absent
log_e(Bit Score)
0.00
0.25
0.50
0.75
1.00
Distance from M. lignano
Bristle Status
present
reduced
absent

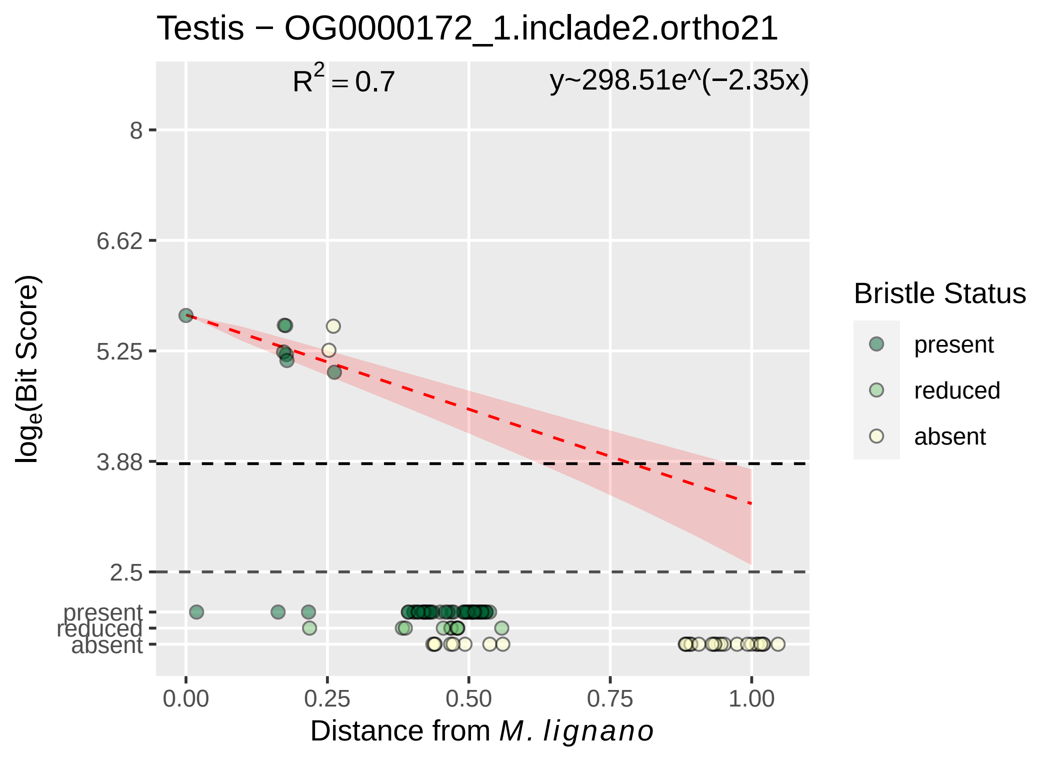

Testis − OG0000172_1.inclade2.ortho21
$R^2 = 0.7$
y~298.51e^(−2.35x)
8
6.62
5.25
3.88
2.5
present
reduced
absent
$\log_e$(Bit Score)
0.00
0.25
0.50
0.75
1.00
Distance from *M. lignano*
Bristle Status
present
reduced
absent

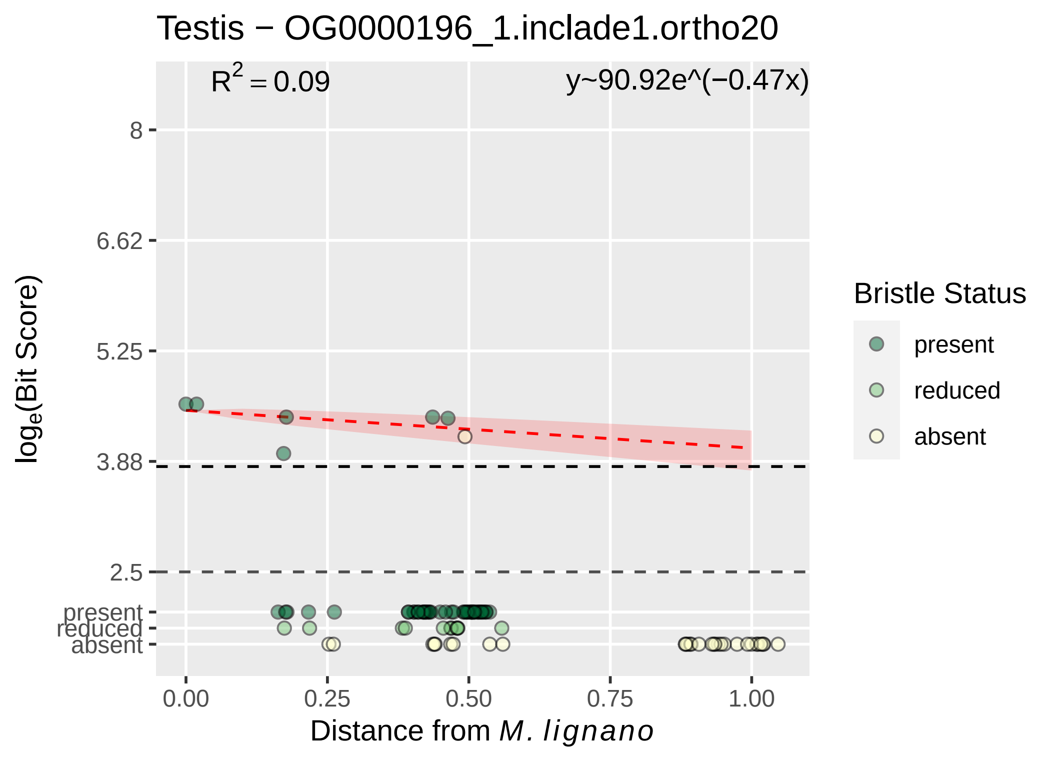

Testis − OG0000196_1.inclade1.ortho20
R^2 = 0.09
y~90.92e^(−0.47x)
8
6.62
5.25
3.88
2.5
present
reduced
absent
log_e(Bit Score)
0.00
0.25
0.50
0.75
1.00
Distance from M. lignano
Bristle Status
present
reduced
absent

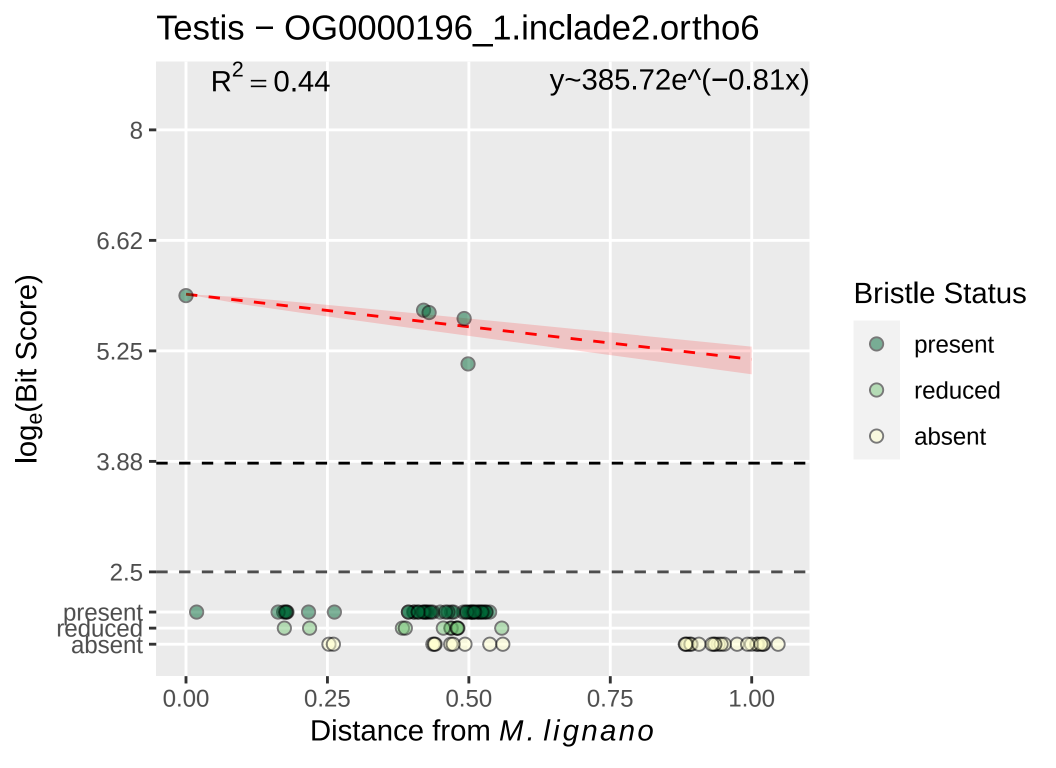

Testis − OG0000196_1.inclade2.ortho6
$R^2 = 0.44$
y~385.72e^(−0.81x)
8
6.62
5.25
3.88
2.5
present
reduced
absent
$\log_e$(Bit Score)
0.00
0.25
0.50
0.75
1.00
Distance from *M. lignano*
Bristle Status
present
reduced
absent

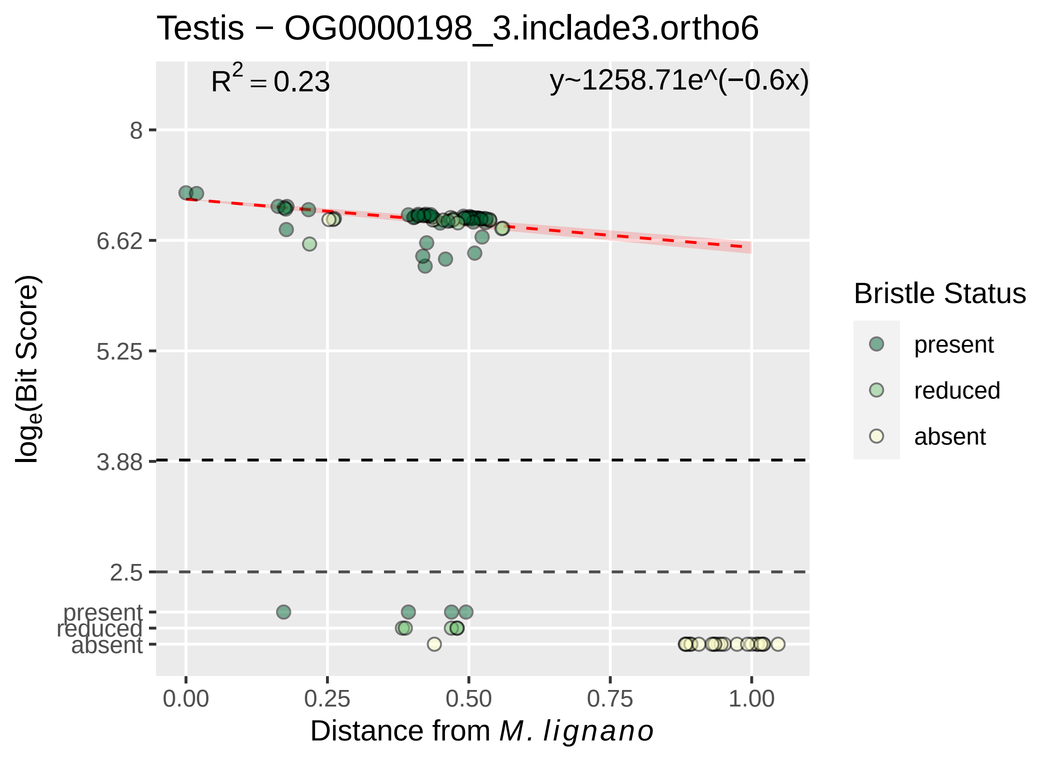

Testis − OG0000198_3.inclade3.ortho6
$R^2 = 0.23$
y~1258.71e^(−0.6x)
8
6.62
5.25
3.88
2.5
present
reduced
absent
0.00
0.25
0.50
0.75
1.00
log_e(Bit Score)
Distance from M. lignano
Bristle Status
present
reduced
absent

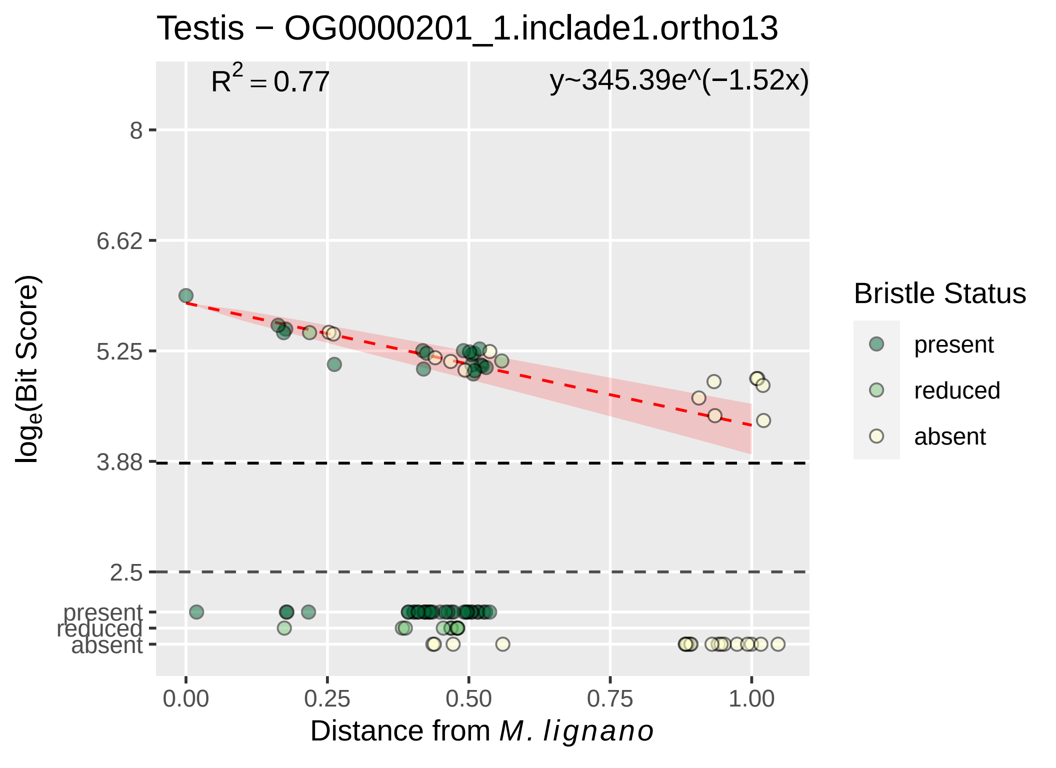

Testis − OG0000201_1.inclade1.ortho13
$R^2 = 0.77$
y~345.39e^(−1.52x)
8
6.62
5.25
3.88
2.5
present
reduced
absent
$\log_e$(Bit Score)
0.00
0.25
0.50
0.75
1.00
Distance from *M. lignano*
Bristle Status
present
reduced
absent

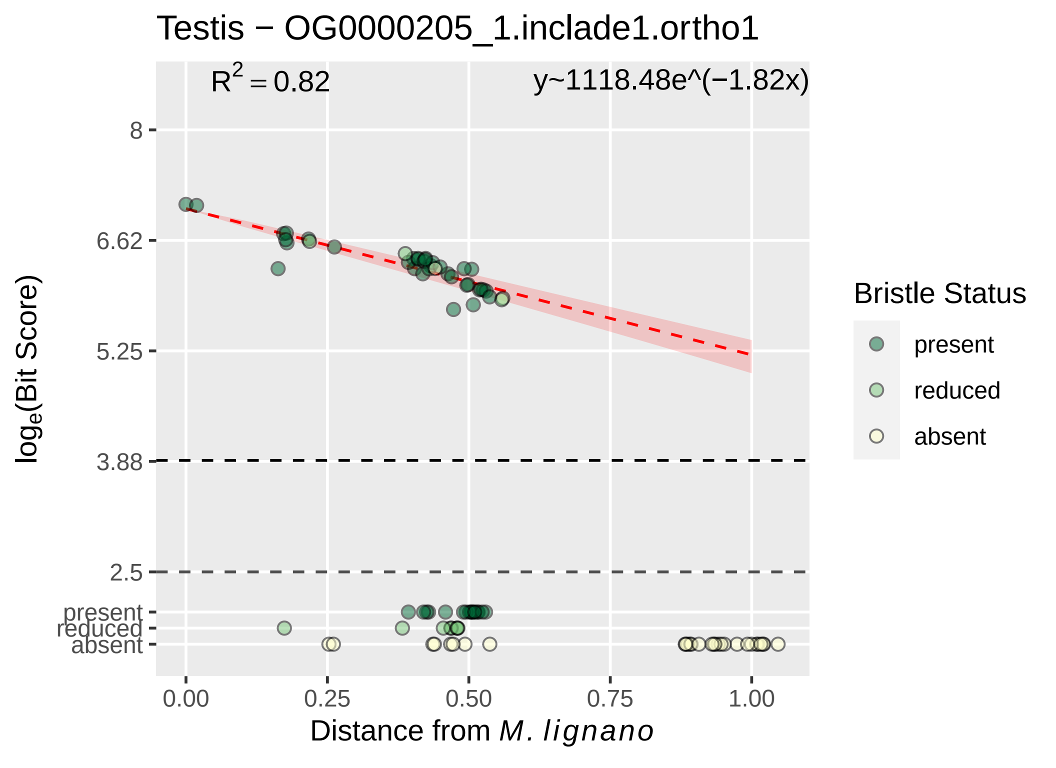

Testis − OG0000205_1.inclade1.ortho1
$R^2 = 0.82$
y~1118.48e^(−1.82x)
8
6.62
5.25
3.88
2.5
present
reduced
absent
0.00
0.25
0.50
0.75
1.00
Distance from *M. lignano*
$\log_e$(Bit Score)
Bristle Status
present
reduced
absent

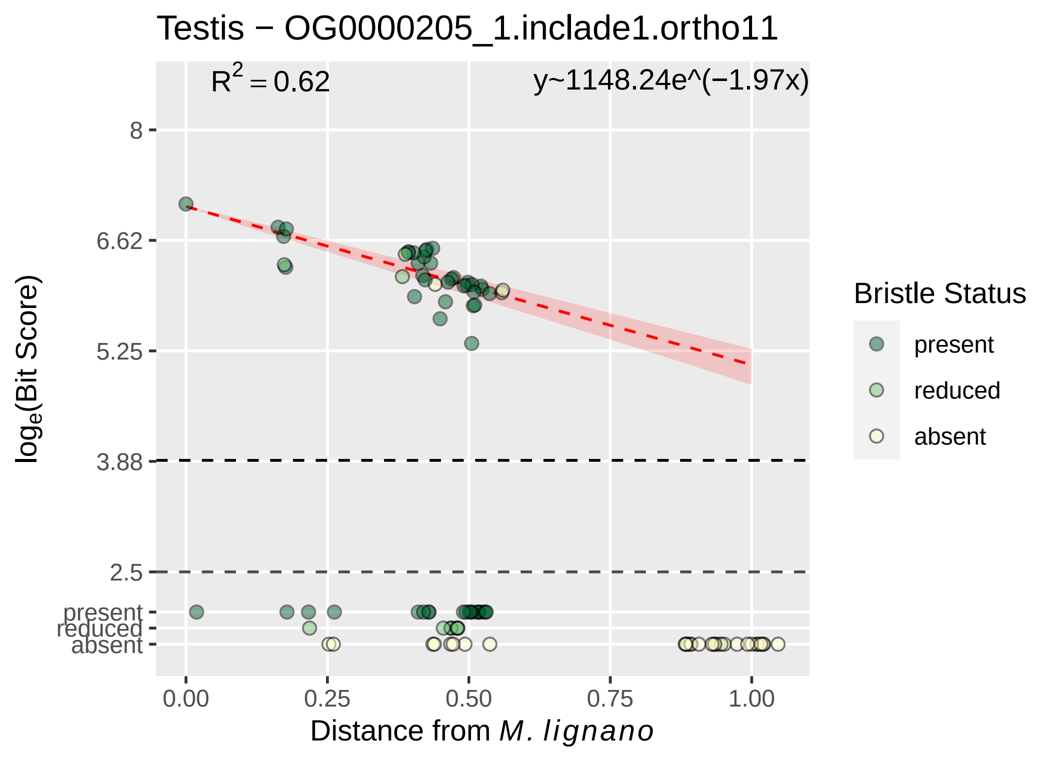

Testis − OG0000205_1.inclade1.ortho11
R2 = 0.62
y~1148.24e^(−1.97x)
8
6.62
5.25
3.88
2.5
present
reduced
absent
0.00
0.25
0.50
0.75
1.00
loge(Bit Score)
Distance from M. lignano
Bristle Status
present
reduced
absent

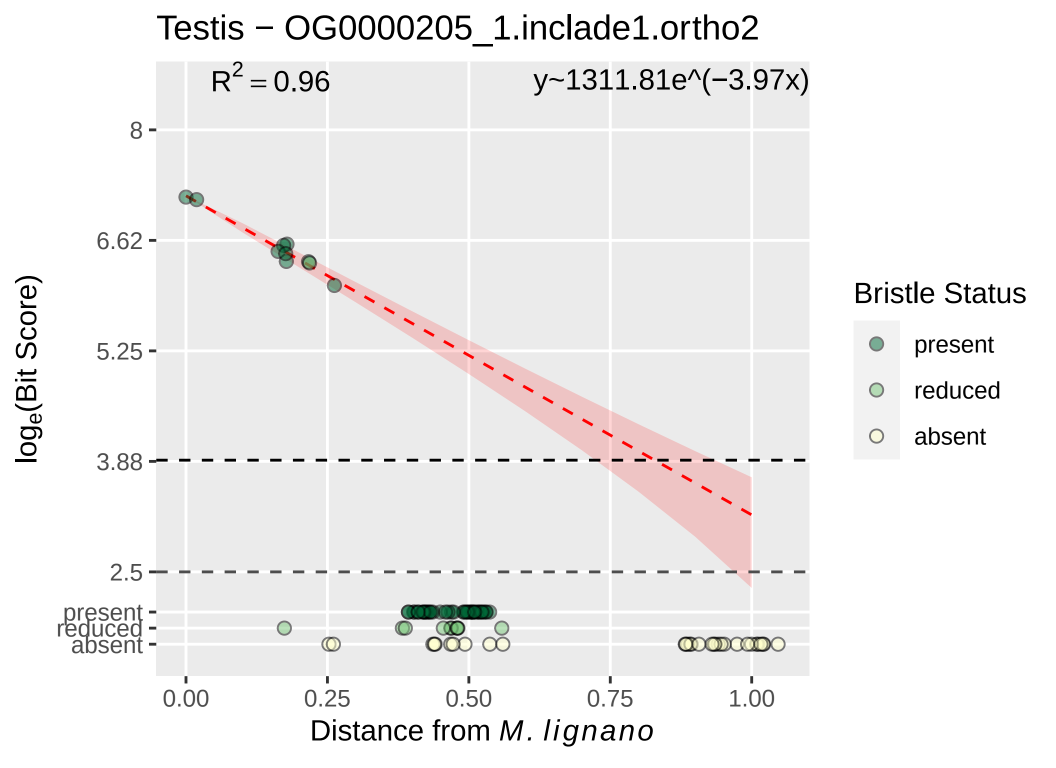

Testis − OG0000205_1.inclade1.ortho2
$R^2 = 0.96$
y~1311.81e^(−3.97x)
8
6.62
5.25
3.88
2.5
present
reduced
absent
log$_e$(Bit Score)
0.00
0.25
0.50
0.75
1.00
Distance from *M. lignano*
Bristle Status
present
reduced
absent

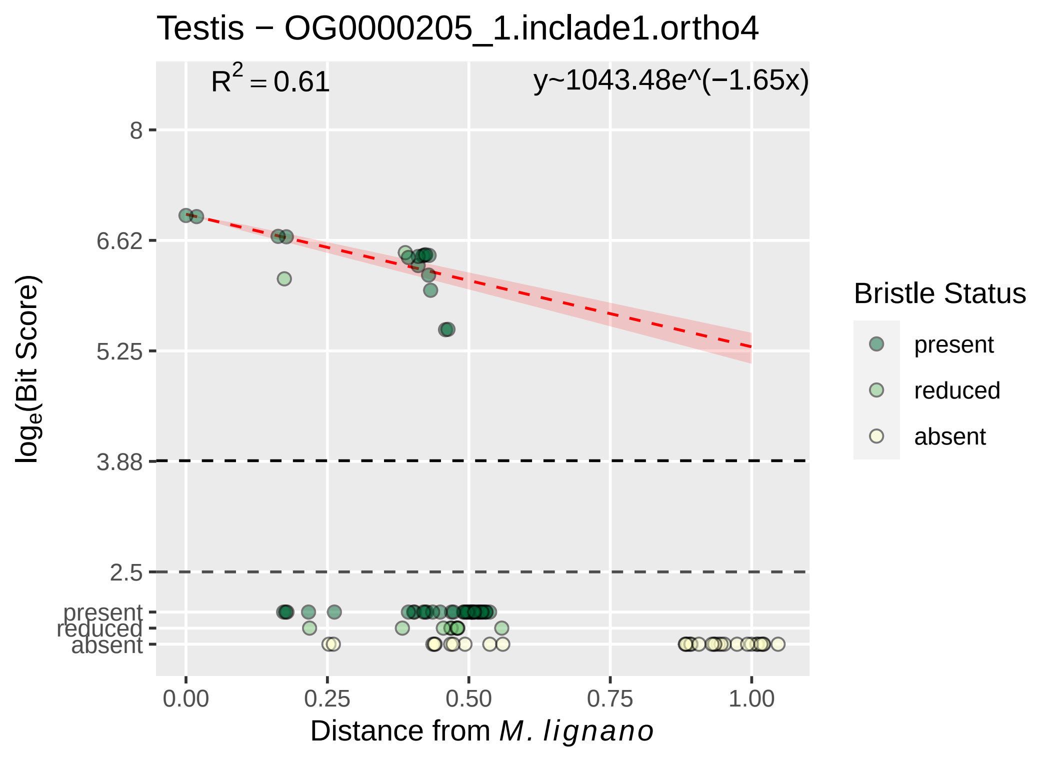
Testis − OG0000205_1.inclade1.ortho4
R^2 = 0.61
y~1043.48e^(−1.65x)
8
6.62
5.25
3.88
2.5
present
reduced
absent
log_e(Bit Score)
0.00
0.25
0.50
0.75
1.00
Distance from M. lignano
Bristle Status
present
reduced
absent

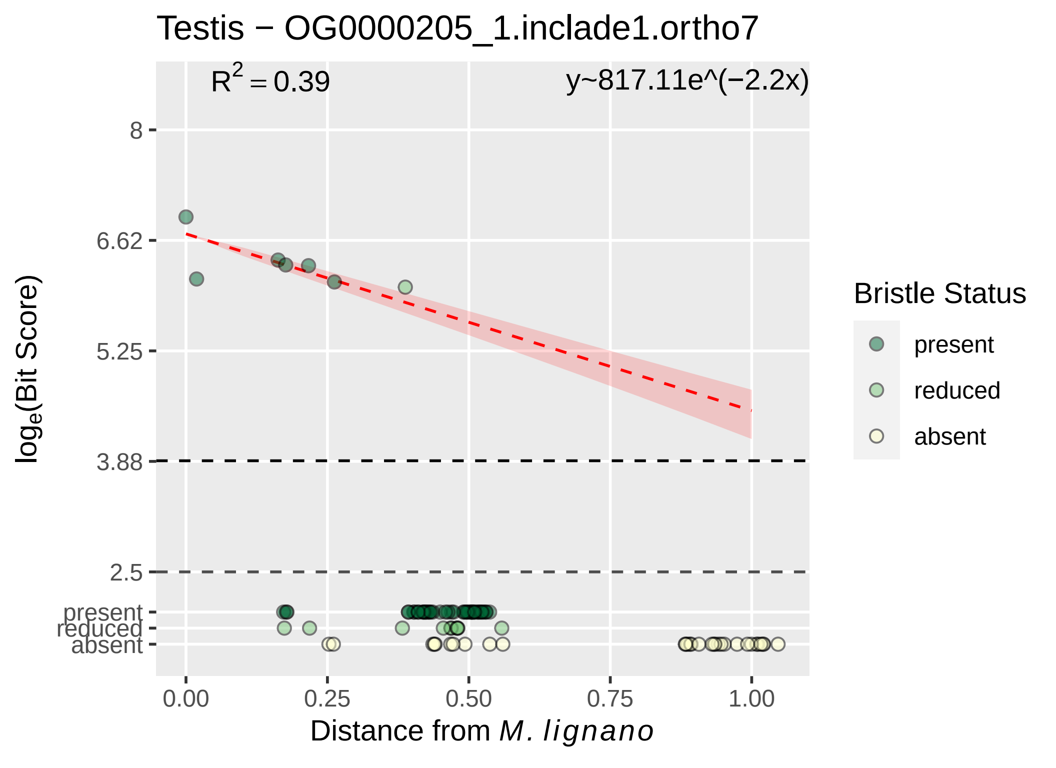

Testis − OG0000205_1.inclade1.ortho7
R^2 = 0.39
y~817.11e^(−2.2x)
8
6.62
5.25
3.88
2.5
present
reduced
absent
log_e(Bit Score)
0.00
0.25
0.50
0.75
1.00
Distance from M. lignano
Bristle Status
present
reduced
absent

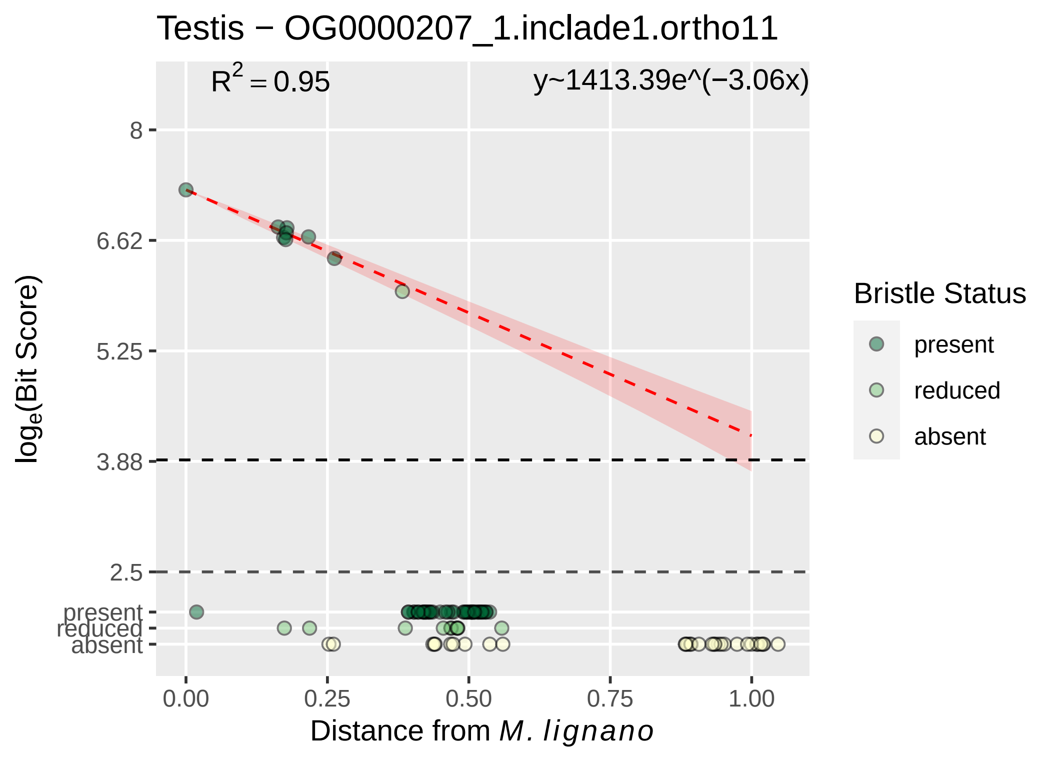
Testis − OG0000207_1.inclade1.ortho11
R^2 = 0.95
y~1413.39e^(−3.06x)
8
6.62
5.25
3.88
2.5
present
reduced
absent
log_e(Bit Score)
0.00
0.25
0.50
0.75
1.00
Distance from M. lignano
Bristle Status
present
reduced
absent

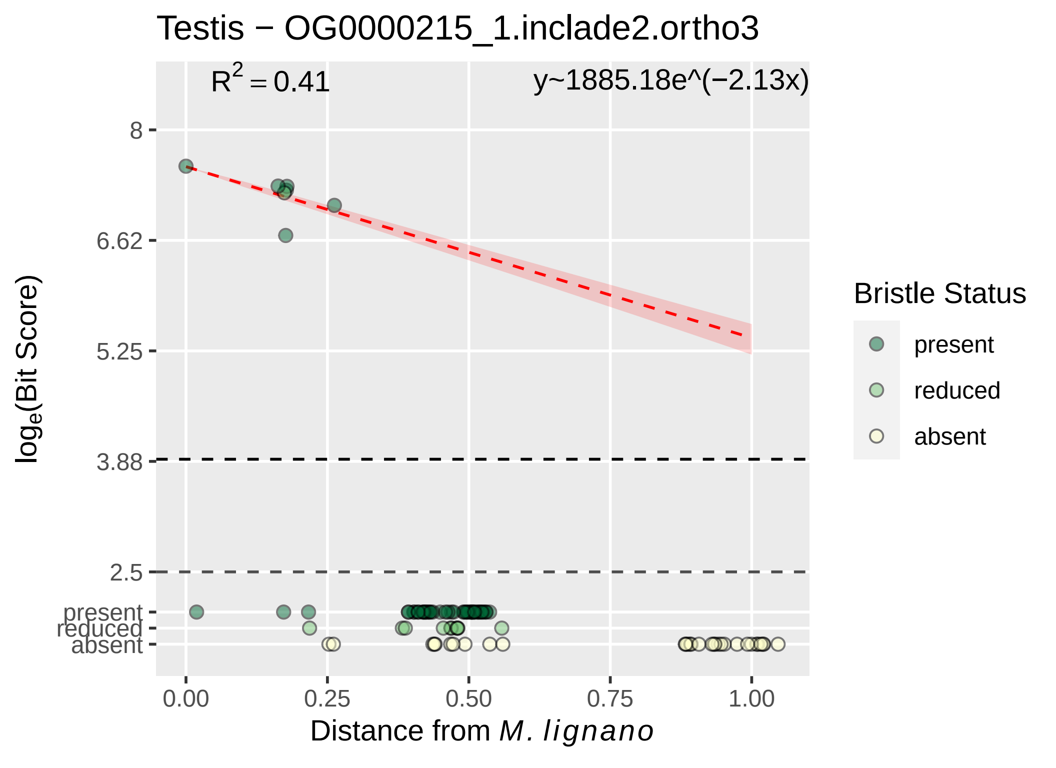

Testis − OG0000215_1.inclade2.ortho3
R2 = 0.41
y~1885.18e^(−2.13x)
8
6.62
5.25
3.88
2.5
present
reduced
absent
0.00
0.25
0.50
0.75
1.00
loge(Bit Score)
Distance from M. lignano
Bristle Status
present
reduced
absent

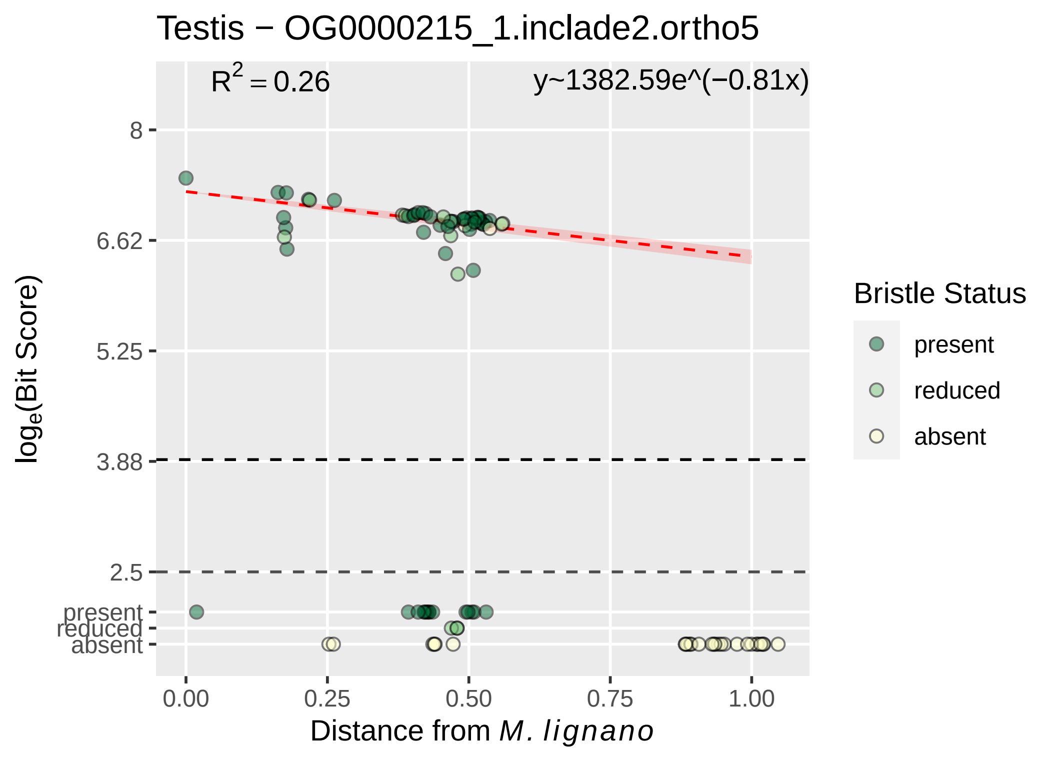
Testis − OG0000215_1.inclade2.ortho5
R² = 0.26
y~1382.59e^(−0.81x)
8
6.62
5.25
3.88
2.5
present
reduced
absent
0.00
0.25
0.50
0.75
1.00
loge(Bit Score)
Distance from M. lignano
Bristle Status
present
reduced
absent

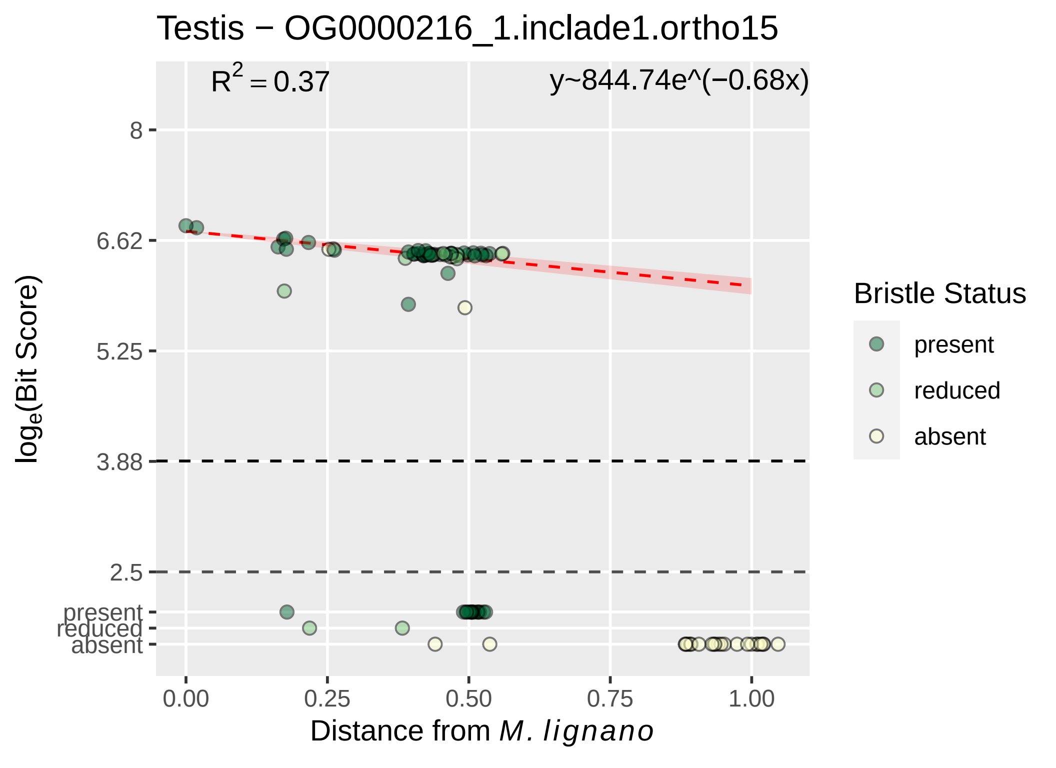
Testis − OG0000216_1.inclade1.ortho15
$R^2 = 0.37$
y~844.74e^(−0.68x)
8
6.62
5.25
3.88
2.5
present
reduced
absent
0.00
0.25
0.50
0.75
1.00
$log_e$(Bit Score)
Distance from *M. lignano*
Bristle Status
present
reduced
absent

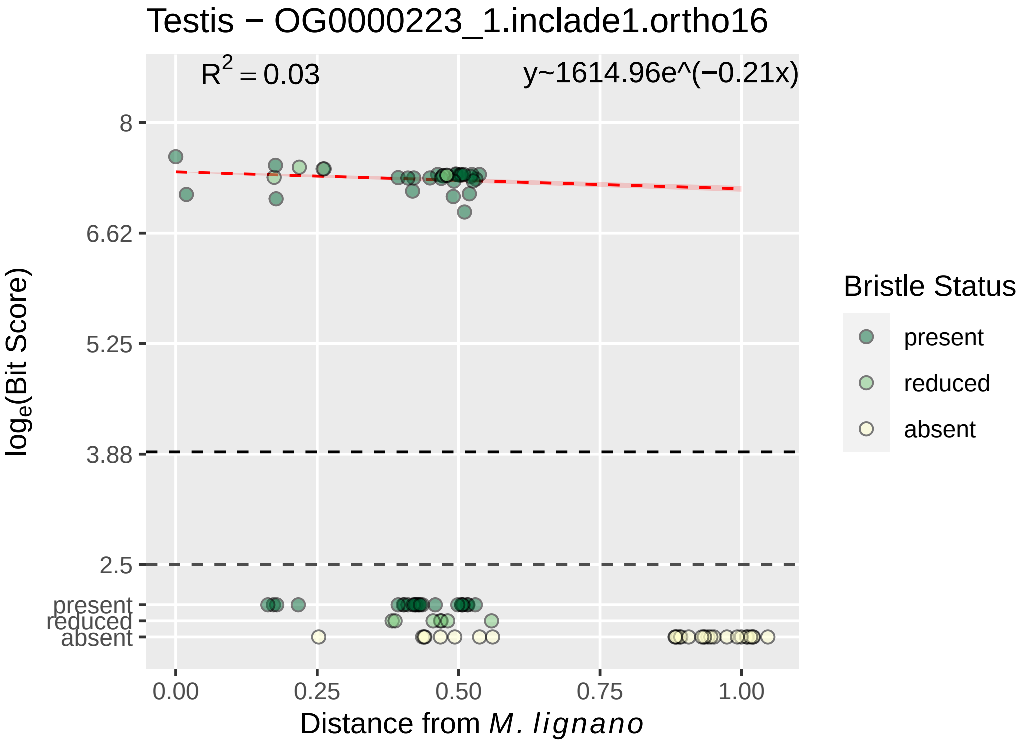

Testis − OG0000223_1.inclade1.ortho16
$R^2 = 0.03$
y~1614.96e^(−0.21x)
8
6.62
5.25
3.88
2.5
present
reduced
absent
$\log_e$(Bit Score)
0.00
0.25
0.50
0.75
1.00
Distance from *M. lignano*
Bristle Status
present
reduced
absent

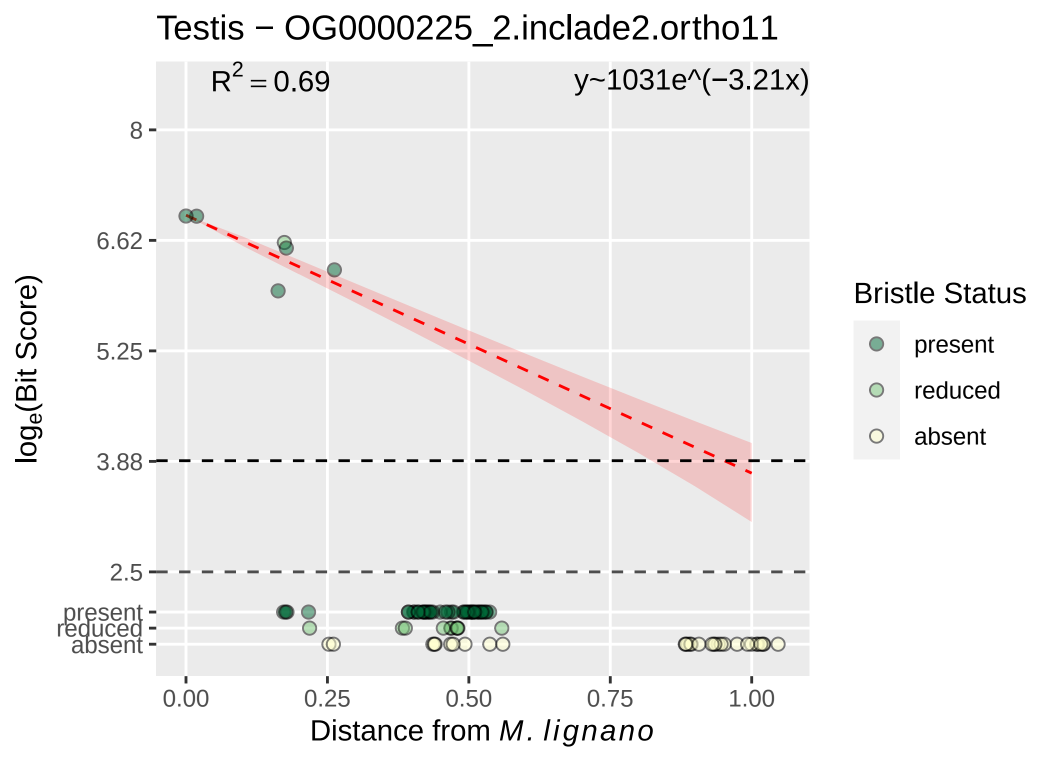

Testis − OG0000225_2.inclade2.ortho11
$R^2 = 0.69$
y~1031e^(−3.21x)
8
6.62
5.25
3.88
2.5
present
reduced
absent
$log_e$(Bit Score)
0.00
0.25
0.50
0.75
1.00
Distance from *M. lignano*
Bristle Status
present
reduced
absent

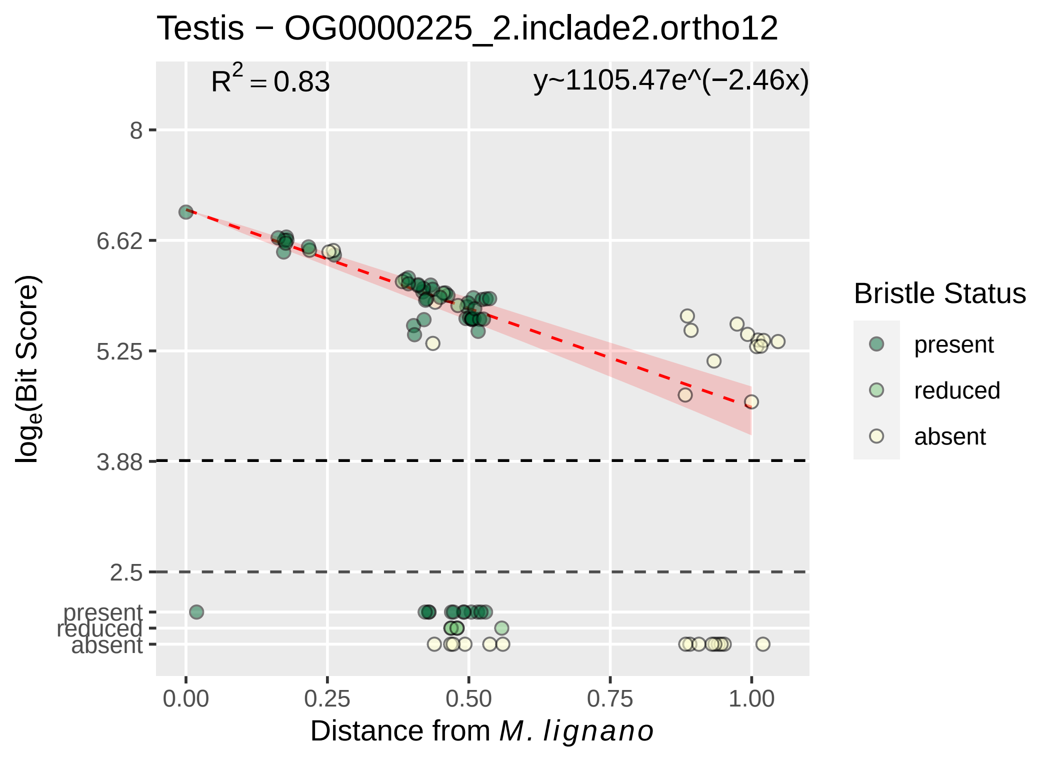

Testis − OG0000225_2.inclade2.ortho12
R^2 = 0.83
y~1105.47e^(−2.46x)
8
6.62
5.25
3.88
2.5
present
reduced
absent
log_e(Bit Score)
0.00
0.25
0.50
0.75
1.00
Distance from M. lignano
Bristle Status
present
reduced
absent

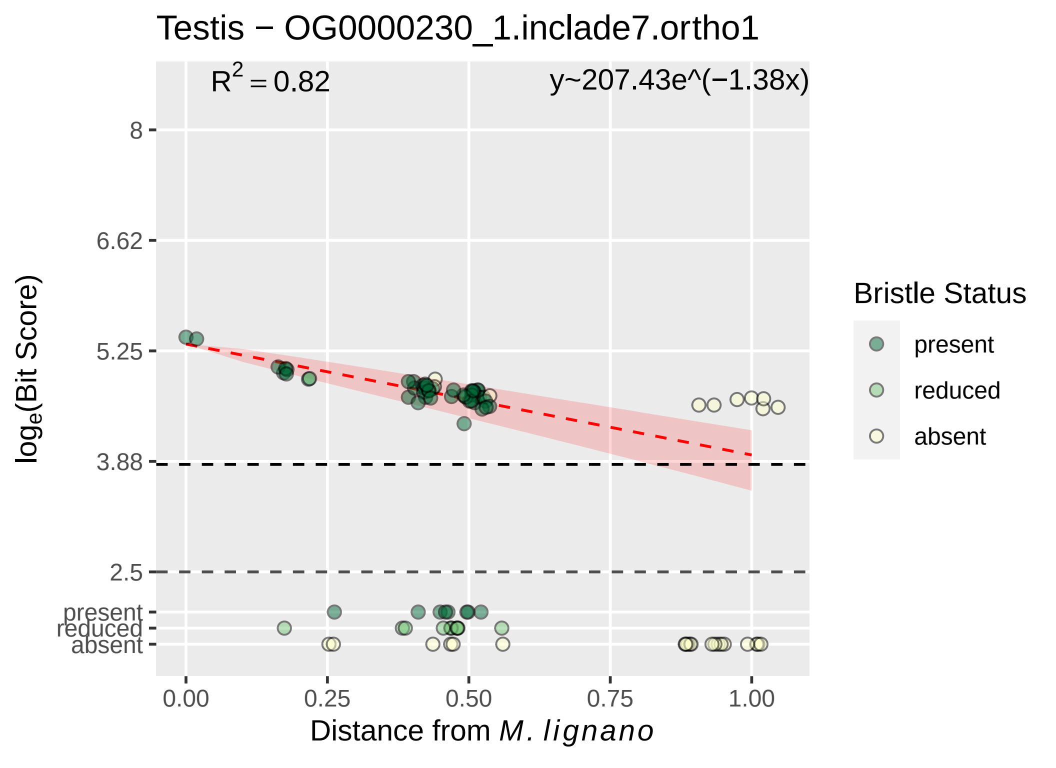
Testis − OG0000230_1.inclade7.ortho1
$R^2 = 0.82$
y~207.43e^(−1.38x)
8
6.62
5.25
3.88
2.5
present
reduced
absent
$log_e$(Bit Score)
0.00
0.25
0.50
0.75
1.00
Distance from *M. lignano*
Bristle Status
present
reduced
absent

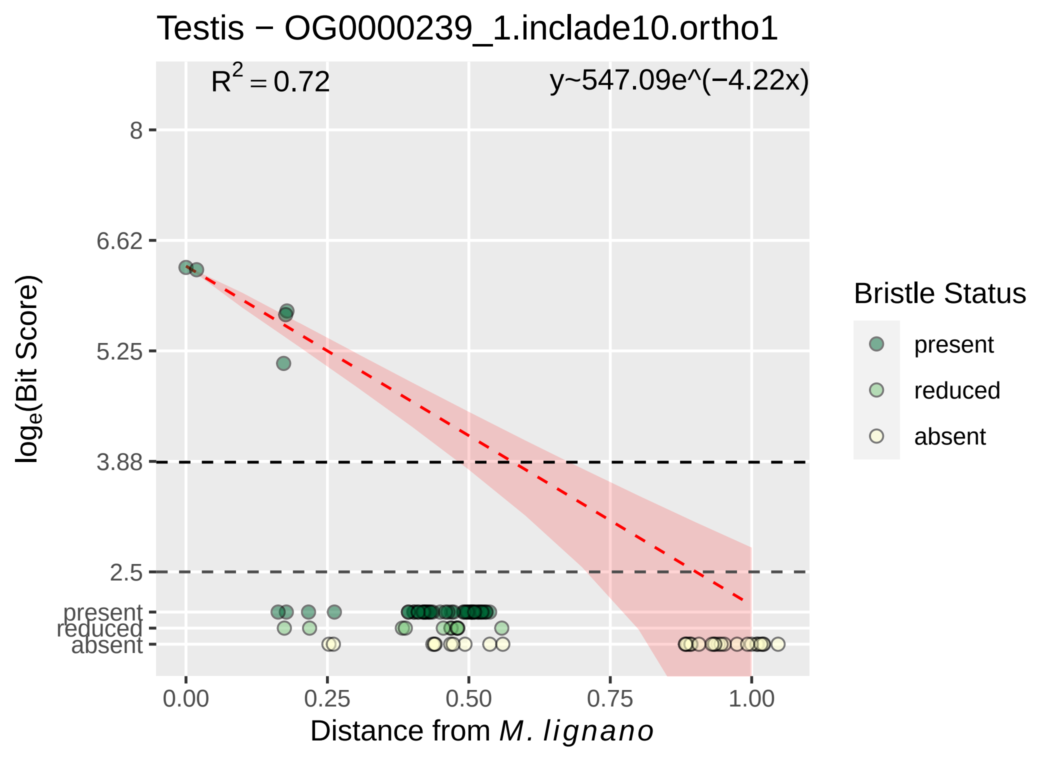

Testis − OG0000239_1.inclade10.ortho1
R^2 = 0.72
y~547.09e^(−4.22x)
8
6.62
5.25
3.88
2.5
present
reduced
absent
0.00
0.25
0.50
0.75
1.00
Distance from M. lignano
loge(Bit Score)
Bristle Status
present
reduced
absent

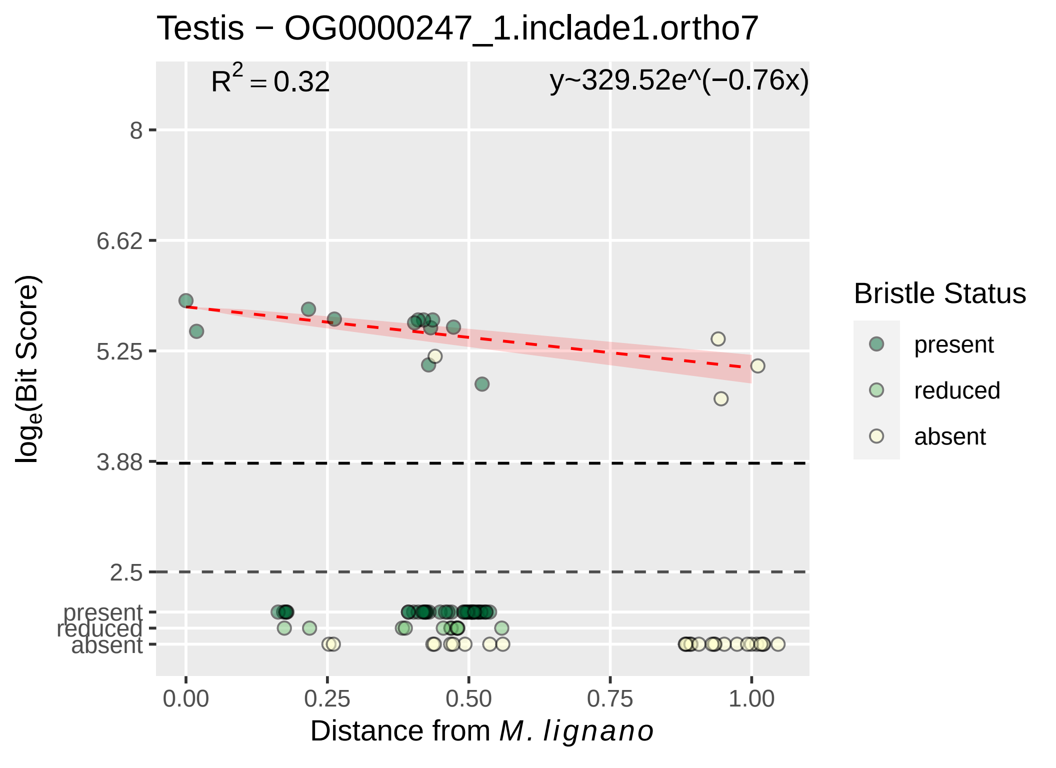

Testis − OG0000247_1.inclade1.ortho7
R^2 = 0.32
y~329.52e^(−0.76x)
8
6.62
5.25
3.88
2.5
present
reduced
absent
log_e(Bit Score)
0.00
0.25
0.50
0.75
1.00
Distance from M. lignano
Bristle Status
present
reduced
absent

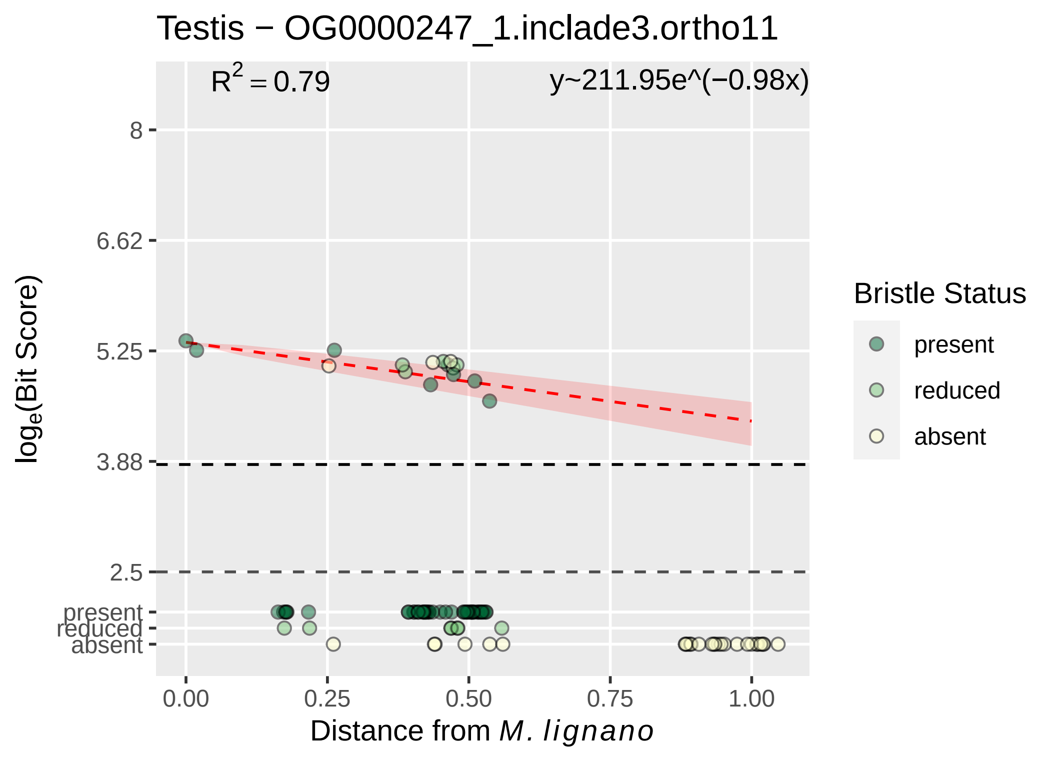

Testis − OG0000247_1.inclade3.ortho11
$R^2 = 0.79$
y~211.95e^(−0.98x)
8
6.62
5.25
3.88
2.5
present
reduced
absent
log$_e$(Bit Score)
0.00
0.25
0.50
0.75
1.00
Distance from *M. lignano*
Bristle Status
present
reduced
absent

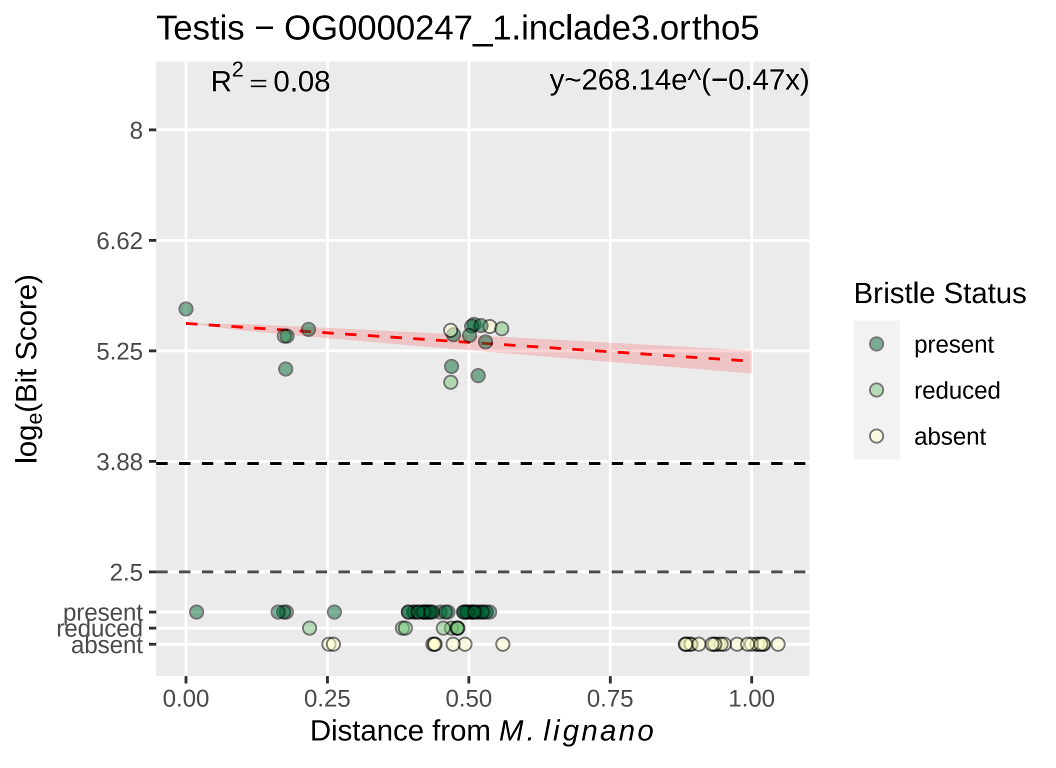

Testis − OG0000247_1.inclade3.ortho5
R^2 = 0.08
y~268.14e^(−0.47x)
8
6.62
5.25
3.88
2.5
present
reduced
absent
log_e(Bit Score)
0.00
0.25
0.50
0.75
1.00
Distance from M. lignano
Bristle Status
present
reduced
absent

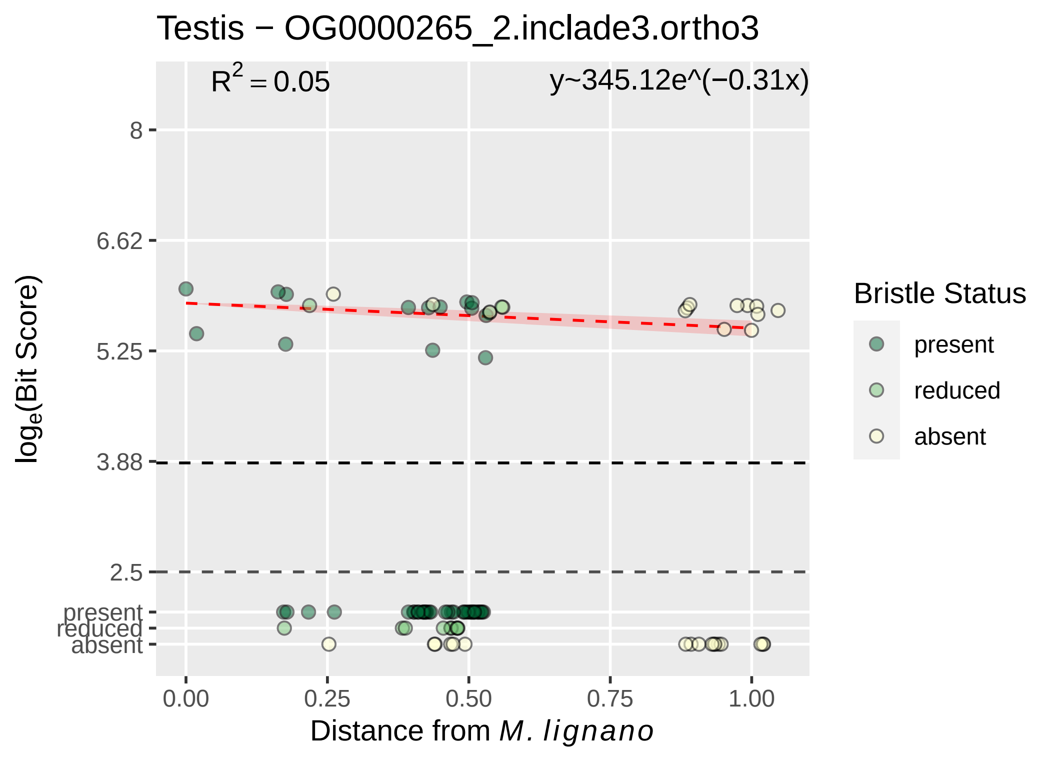
Testis − OG0000265_2.inclade3.ortho3
R^2 = 0.05
y~345.12e^(−0.31x)
log_e(Bit Score)
8
6.62
5.25
3.88
2.5
present
reduced
absent
0.00
0.25
0.50
0.75
1.00
Distance from M. lignano
Bristle Status
present
reduced
absent

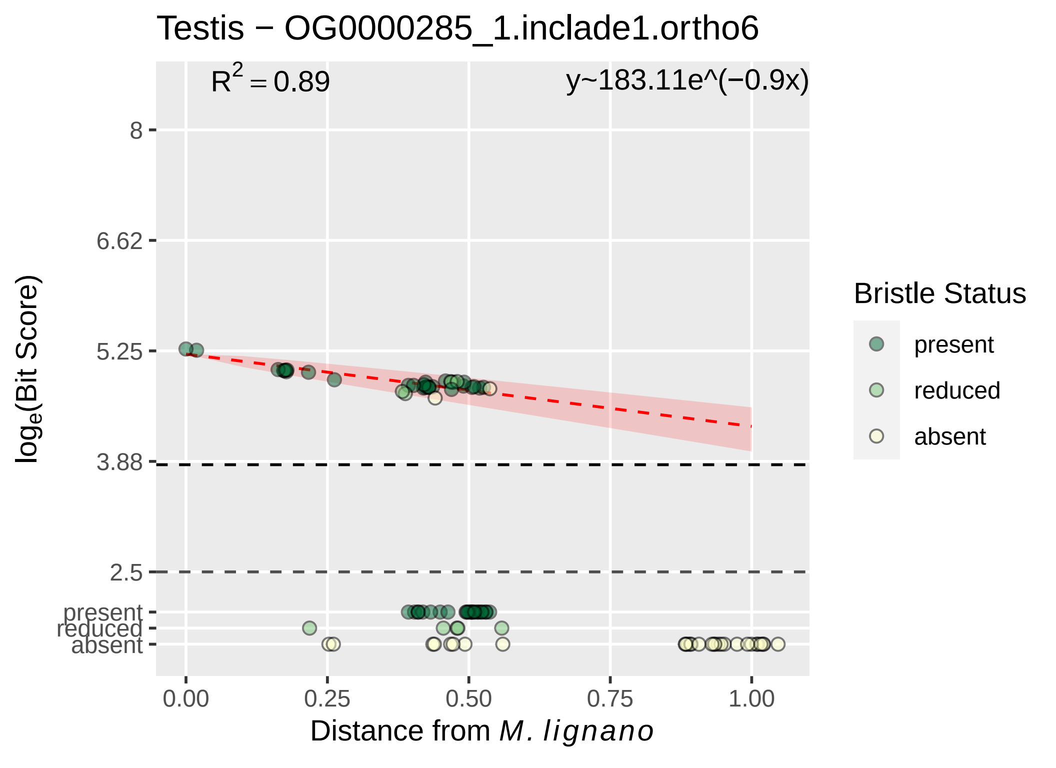

Testis − OG0000285_1.inclade1.ortho6
$R^2 = 0.89$
y~183.11e^(−0.9x)
8
6.62
5.25
3.88
2.5
present
reduced
absent
0.00
0.25
0.50
0.75
1.00
$\log_e$(Bit Score)
Distance from *M. lignano*
Bristle Status
present
reduced
absent

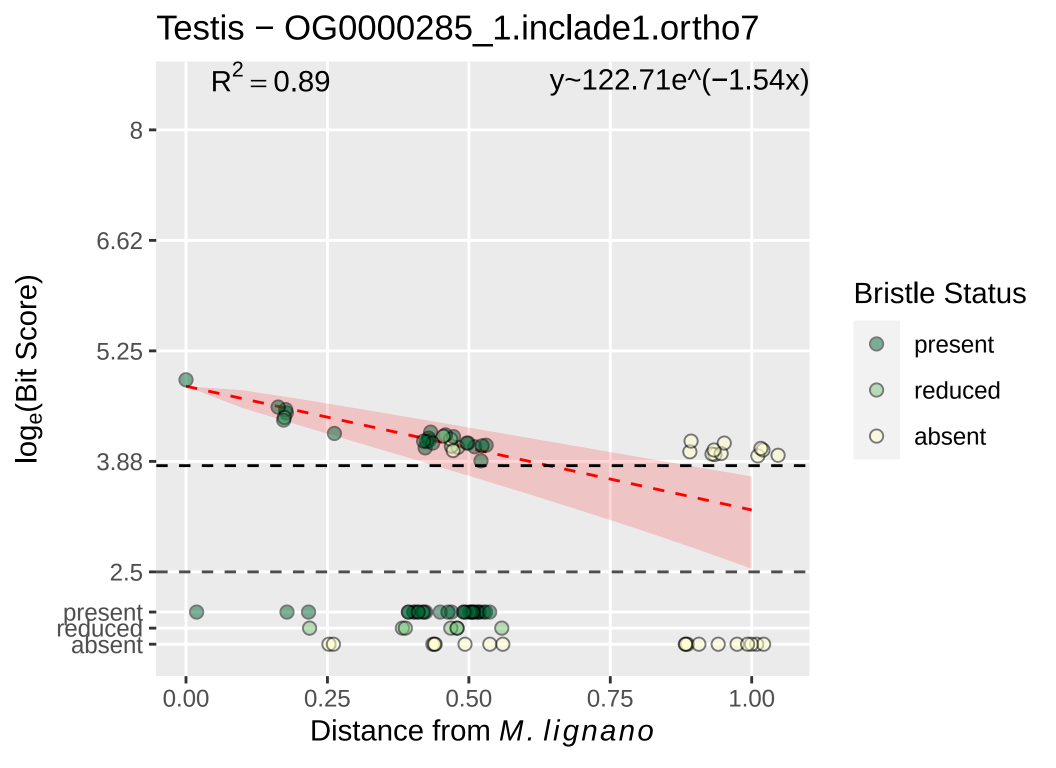

Testis − OG0000285_1.inclade1.ortho7
$R^2 = 0.89$
y~122.71e^(−1.54x)
8
6.62
5.25
3.88
2.5
present
reduced
absent
$\log_e$(Bit Score)
0.00
0.25
0.50
0.75
1.00
Distance from *M. lignano*
Bristle Status
present
reduced
absent

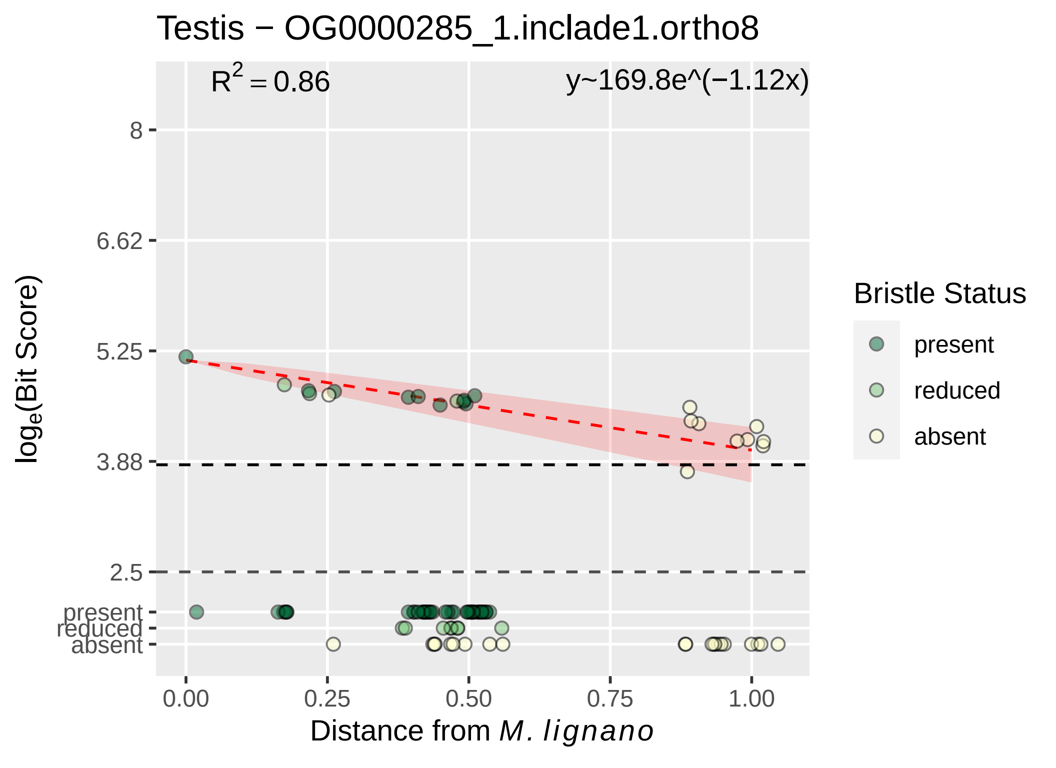

Testis − OG0000285_1.inclade1.ortho8
$R^2 = 0.86$
y~169.8e^(−1.12x)
8
6.62
5.25
3.88
2.5
present
reduced
absent
$\log_e$(Bit Score)
0.00
0.25
0.50
0.75
1.00
Distance from *M. lignano*
Bristle Status
present
reduced
absent

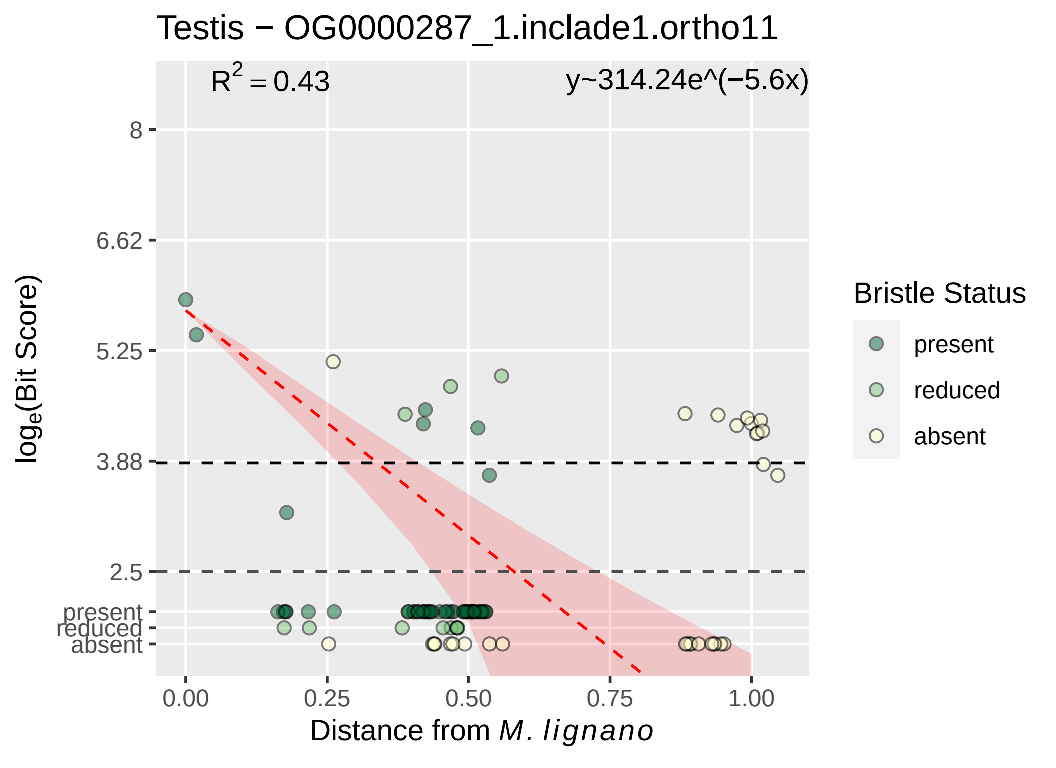

Testis − OG0000287_1.inclade1.ortho11
R² = 0.43
y~314.24e^(−5.6x)
8
6.62
5.25
3.88
2.5
present
reduced
absent
loge(Bit Score)
0.00
0.25
0.50
0.75
1.00
Distance from M. lignano
Bristle Status
present
reduced
absent

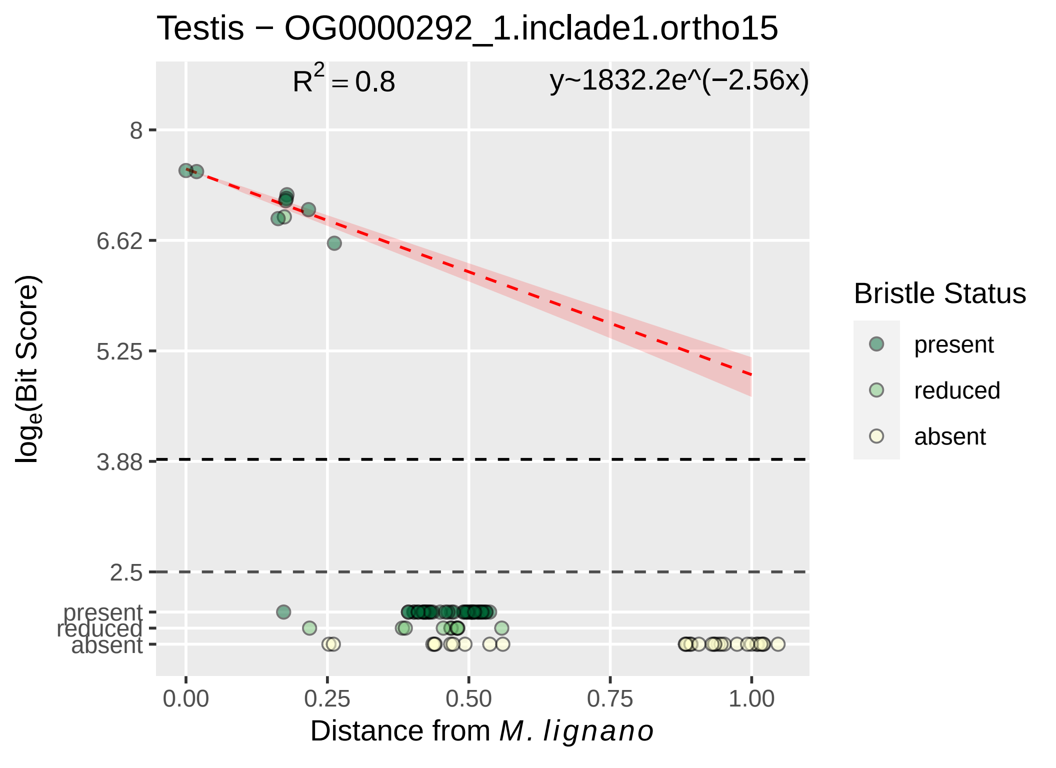

Testis − OG0000292_1.inclade1.ortho15
$R^2 = 0.8$
y~1832.2e^(−2.56x)
8
6.62
5.25
3.88
2.5
present
reduced
absent
0.00
0.25
0.50
0.75
1.00
$\log_e$(Bit Score)
Distance from *M. lignano*
Bristle Status
present
reduced
absent

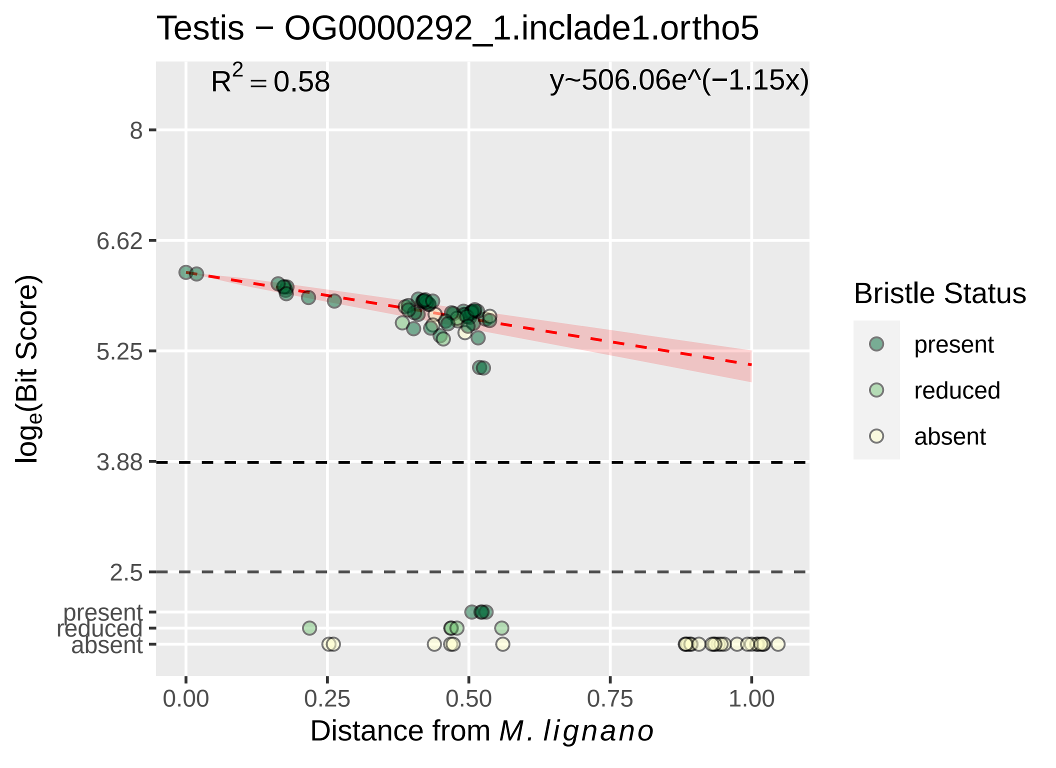

Testis − OG0000292_1.inclade1.ortho5
$R^2 = 0.58$
y~506.06e^(−1.15x)
8
6.62
5.25
3.88
2.5
present
reduced
absent
$\log_e$(Bit Score)
0.00
0.25
0.50
0.75
1.00
Distance from *M. lignano*
Bristle Status
present
reduced
absent

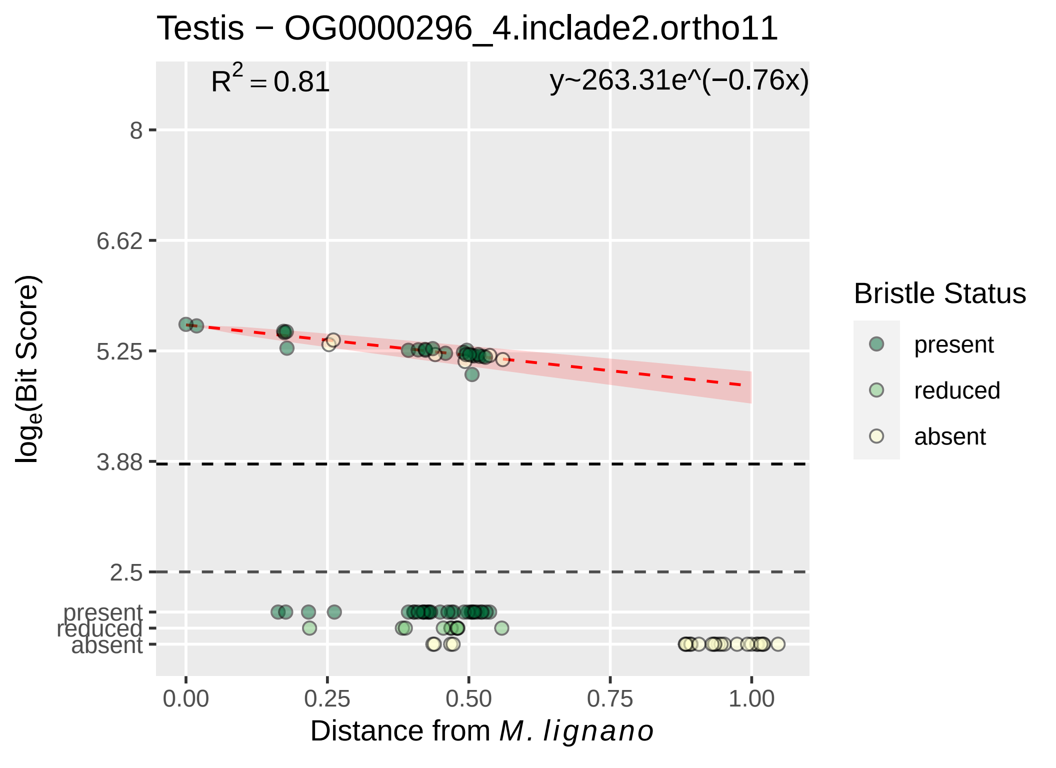

Testis − OG0000296_4.inclade2.ortho11
$R^2 = 0.81$
y~263.31e^(−0.76x)
8
6.62
5.25
3.88
2.5
present
reduced
absent
$log_e$(Bit Score)
0.00
0.25
0.50
0.75
1.00
Distance from *M. lignano*
Bristle Status
present
reduced
absent

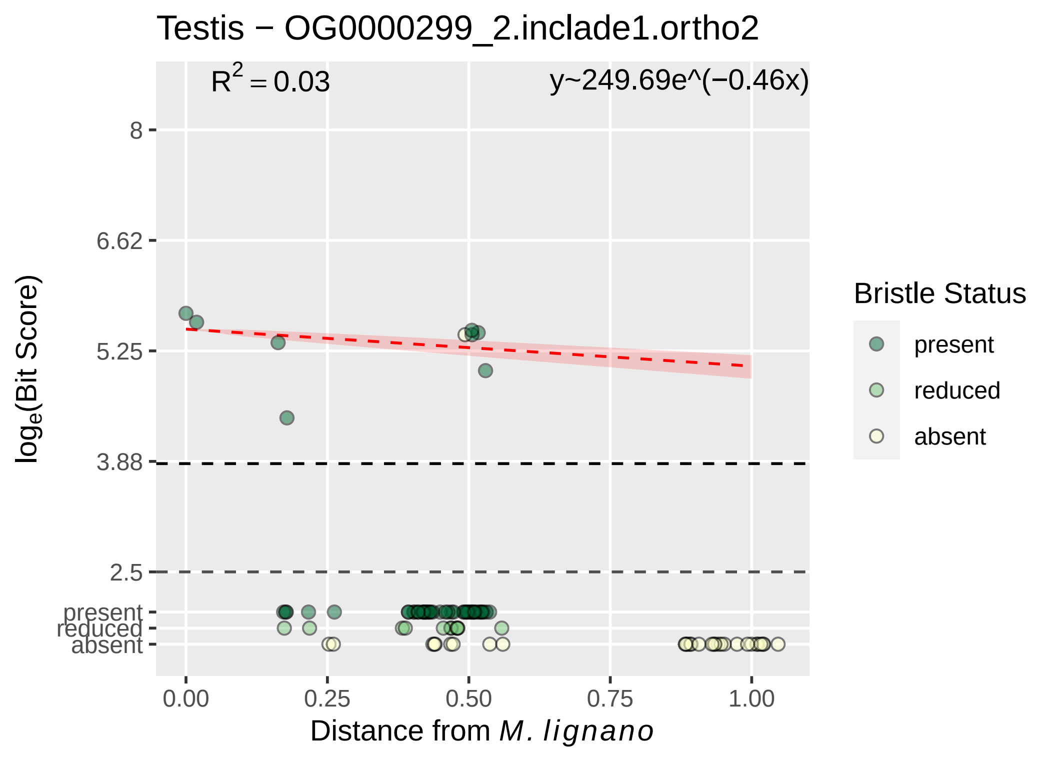

Testis − OG0000299_2.inclade1.ortho2
R2 = 0.03
y~249.69e^(−0.46x)
8
6.62
5.25
3.88
2.5
present
reduced
absent
loge(Bit Score)
0.00
0.25
0.50
0.75
1.00
Distance from M. lignano
Bristle Status
present
reduced
absent

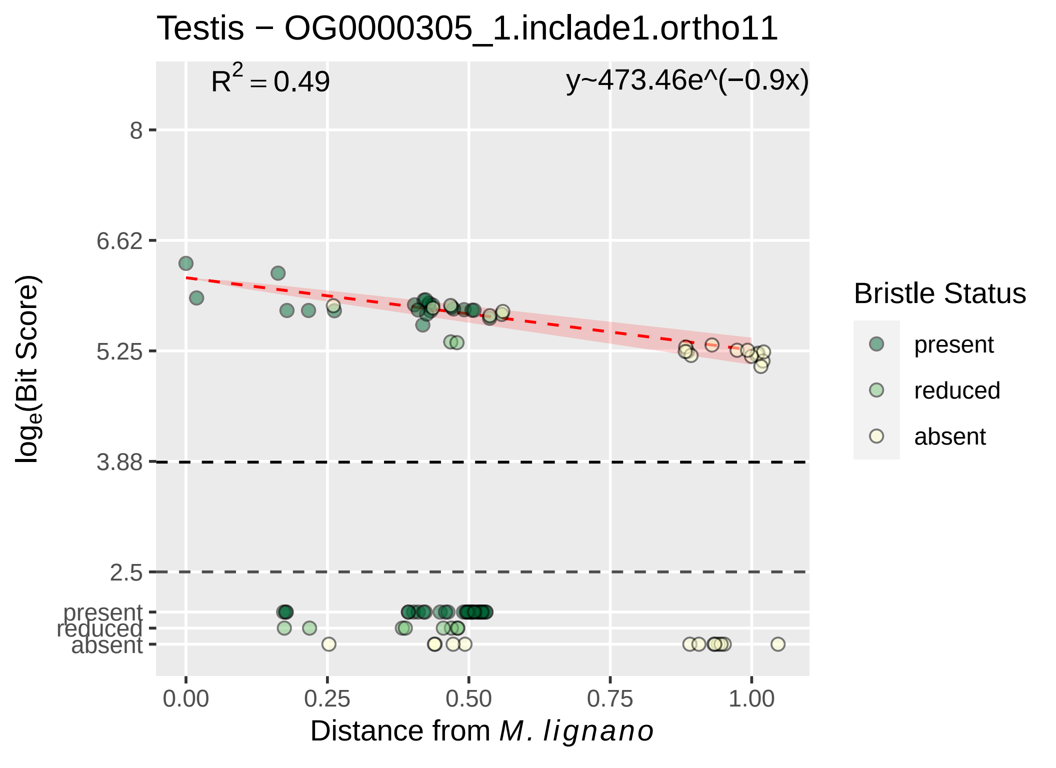

Testis − OG0000305_1.inclade1.ortho11
R^2 = 0.49
y~473.46e^(−0.9x)
8
6.62
5.25
3.88
2.5
present
reduced
absent
loge(Bit Score)
0.00
0.25
0.50
0.75
1.00
Distance from M. lignano
Bristle Status
present
reduced
absent

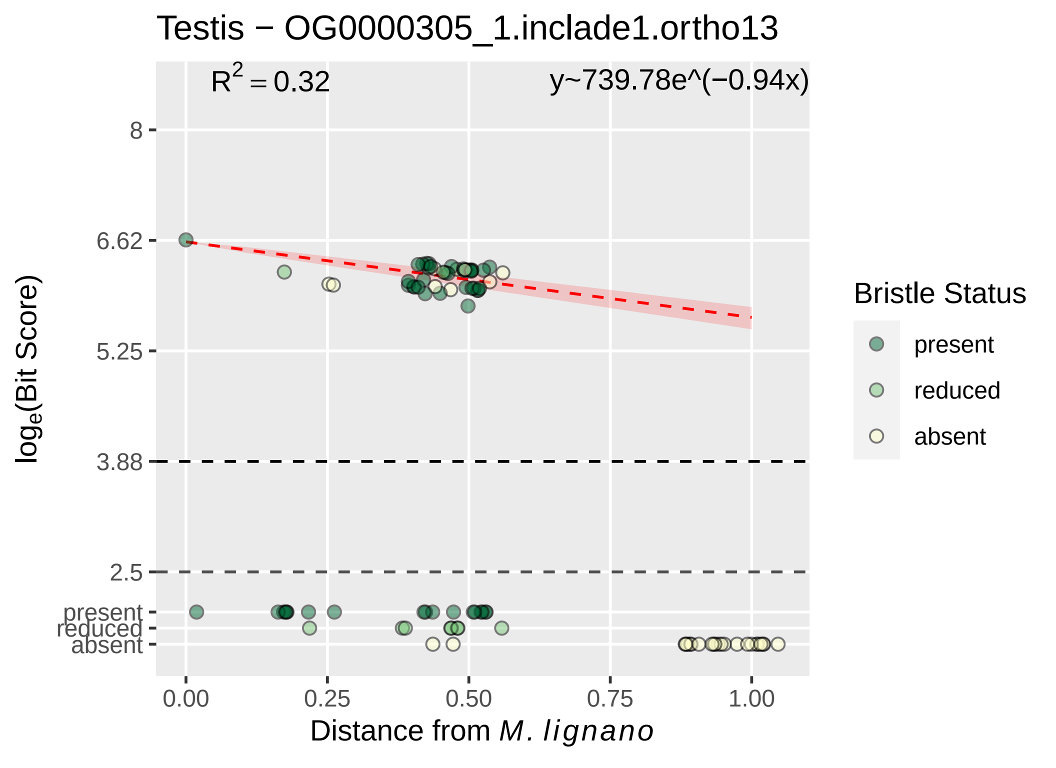

Testis − OG0000305_1.inclade1.ortho13
$R^2 = 0.32$
y~739.78e^(−0.94x)
8
6.62
5.25
3.88
2.5
present
reduced
absent
0.00
0.25
0.50
0.75
1.00
Distance from M. lignano
loge(Bit Score)
Bristle Status
present
reduced
absent

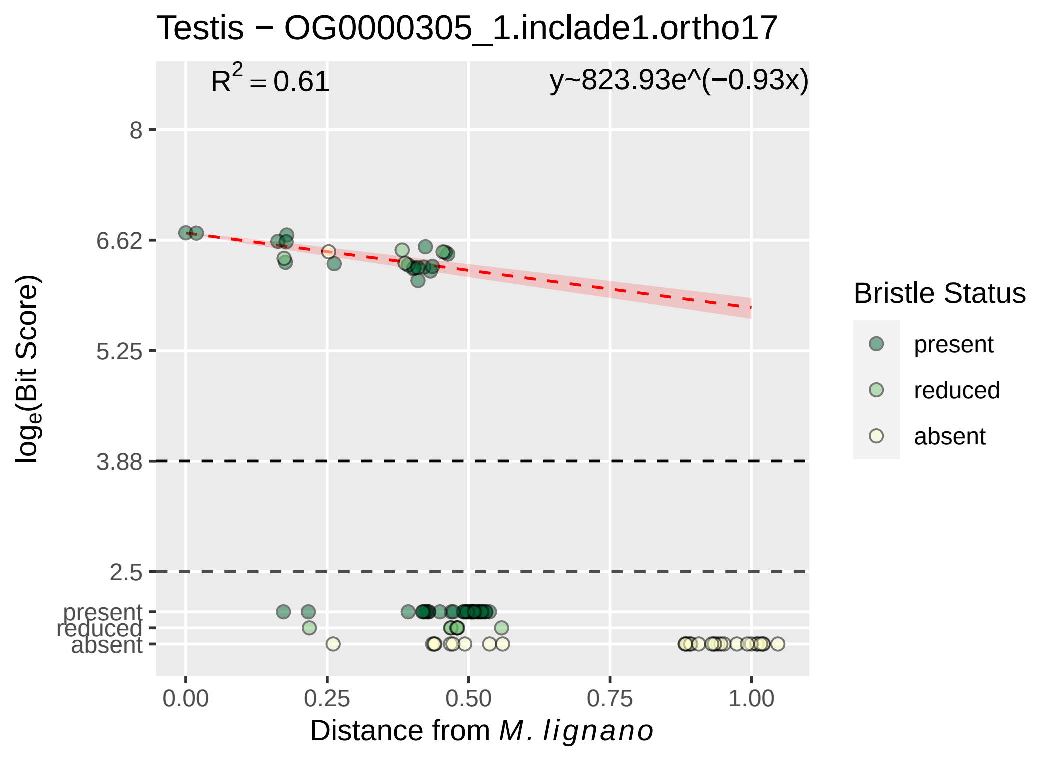
Testis − OG0000305_1.inclade1.ortho17
R^2 = 0.61
y~823.93e^(−0.93x)
8
6.62
5.25
3.88
2.5
present
reduced
absent
log_e(Bit Score)
0.00
0.25
0.50
0.75
1.00
Distance from M. lignano
Bristle Status
present
reduced
absent

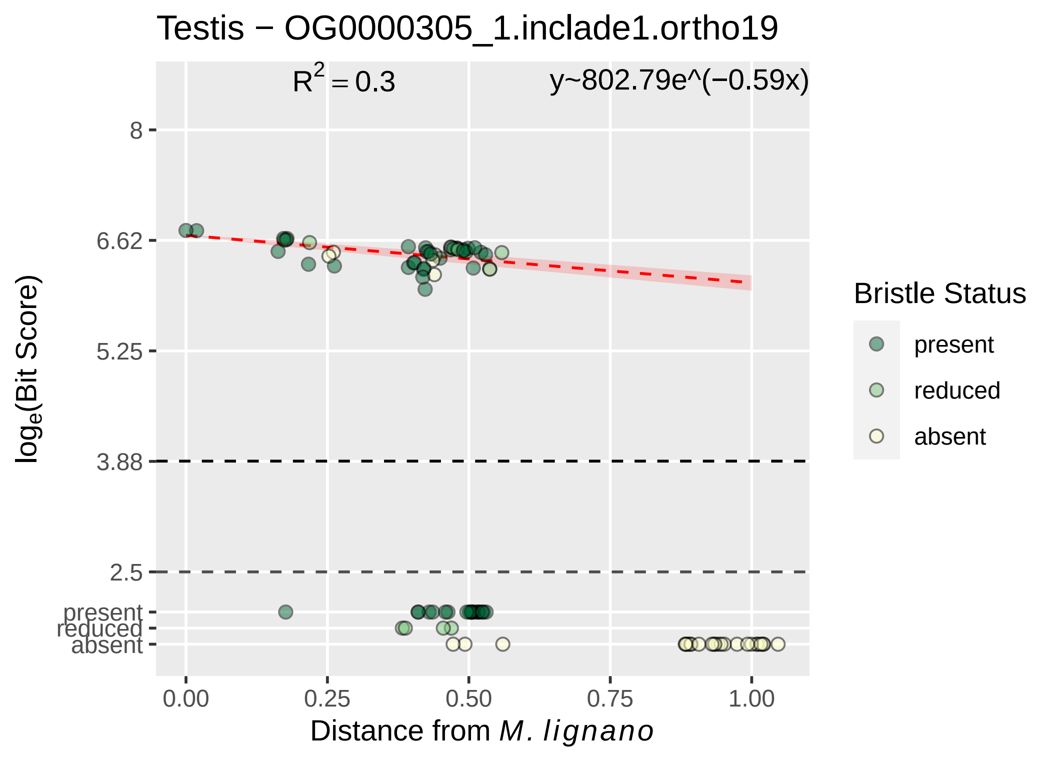

Testis − OG0000305_1.inclade1.ortho19
$R^2 = 0.3$
y~802.79e^(−0.59x)
8
6.62
5.25
3.88
2.5
present
reduced
absent
0.00
0.25
0.50
0.75
1.00
Distance from M. lignano
log$_e$(Bit Score)
Bristle Status
present
reduced
absent

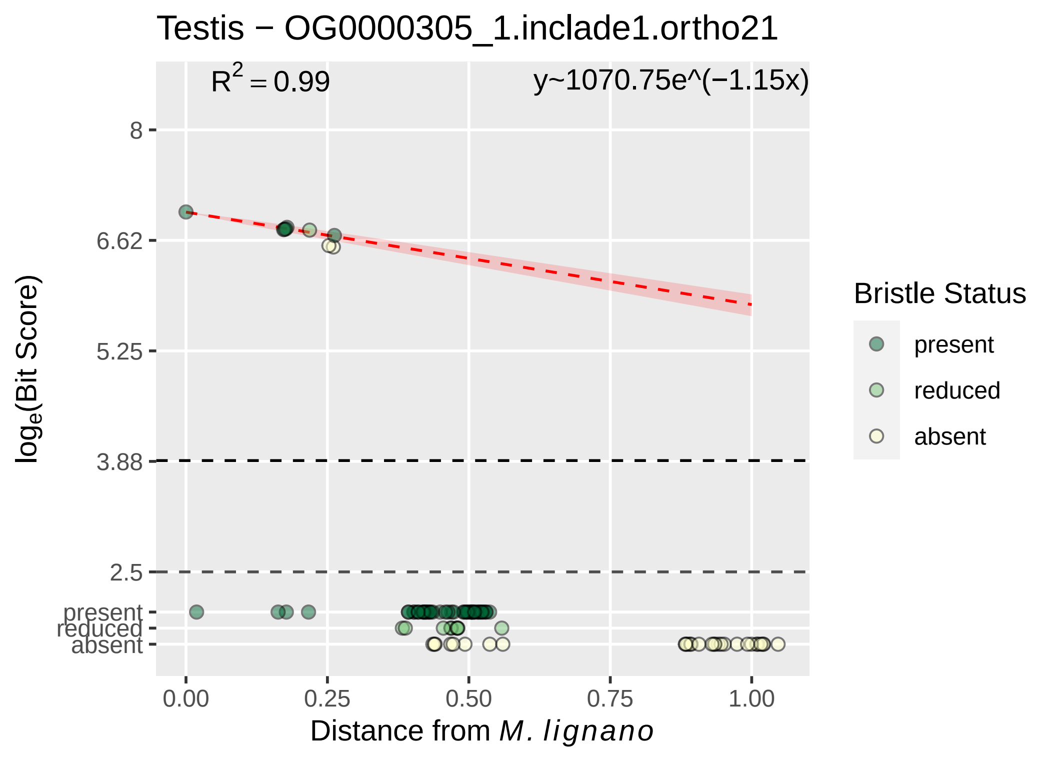

Testis − OG0000305_1.inclade1.ortho21
$R^2 = 0.99$
y~1070.75e^(−1.15x)
8
6.62
5.25
3.88
2.5
present
reduced
absent
0.00
0.25
0.50
0.75
1.00
Distance from *M. lignano*
$\log_e$(Bit Score)
Bristle Status
present
reduced
absent

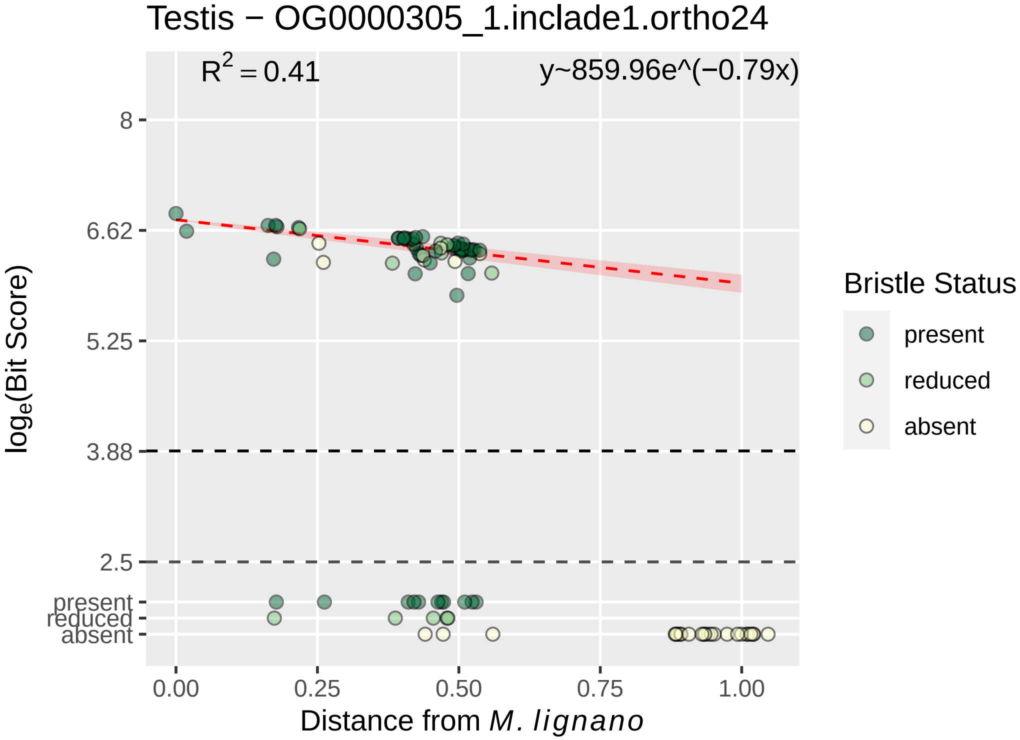

Testis − OG0000305_1.inclade1.ortho24
$R^2 = 0.41$
y~859.96e^(−0.79x)
8
6.62
5.25
3.88
2.5
present
reduced
absent
log$_e$(Bit Score)
0.00
0.25
0.50
0.75
1.00
Distance from *M. lignano*
Bristle Status
present
reduced
absent

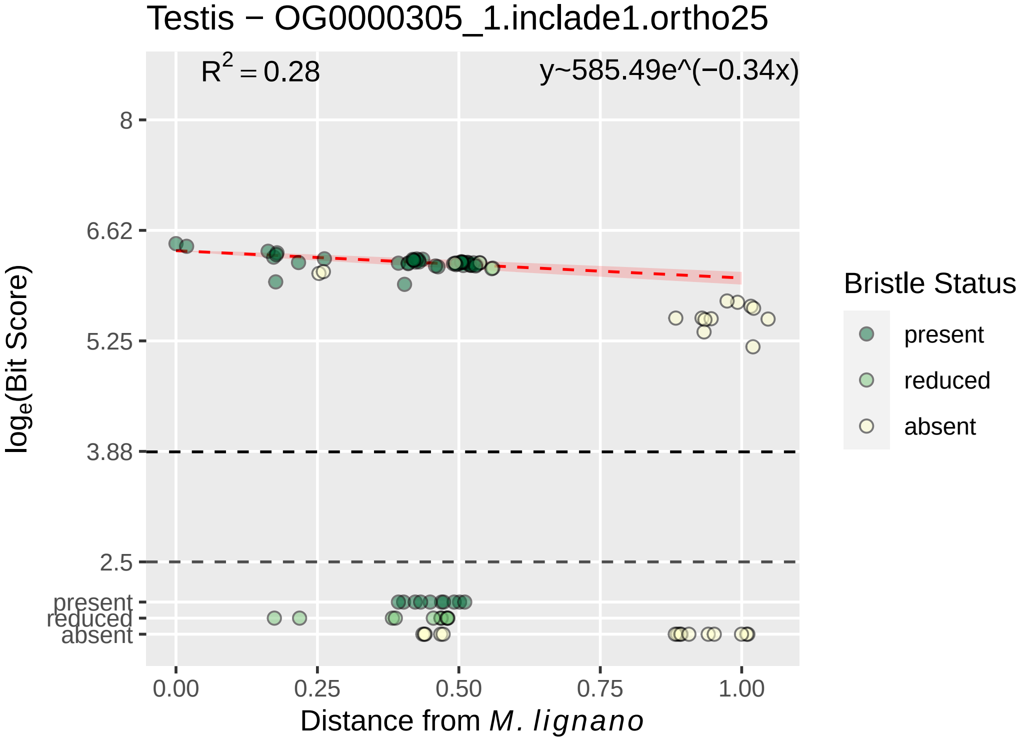

Testis − OG0000305_1.inclade1.ortho25
$R^2 = 0.28$
y~585.49e^(−0.34x)
8
6.62
5.25
3.88
2.5
present
reduced
absent
$log_e$(Bit Score)
0.00
0.25
0.50
0.75
1.00
Distance from *M. lignano*
Bristle Status
present
reduced
absent

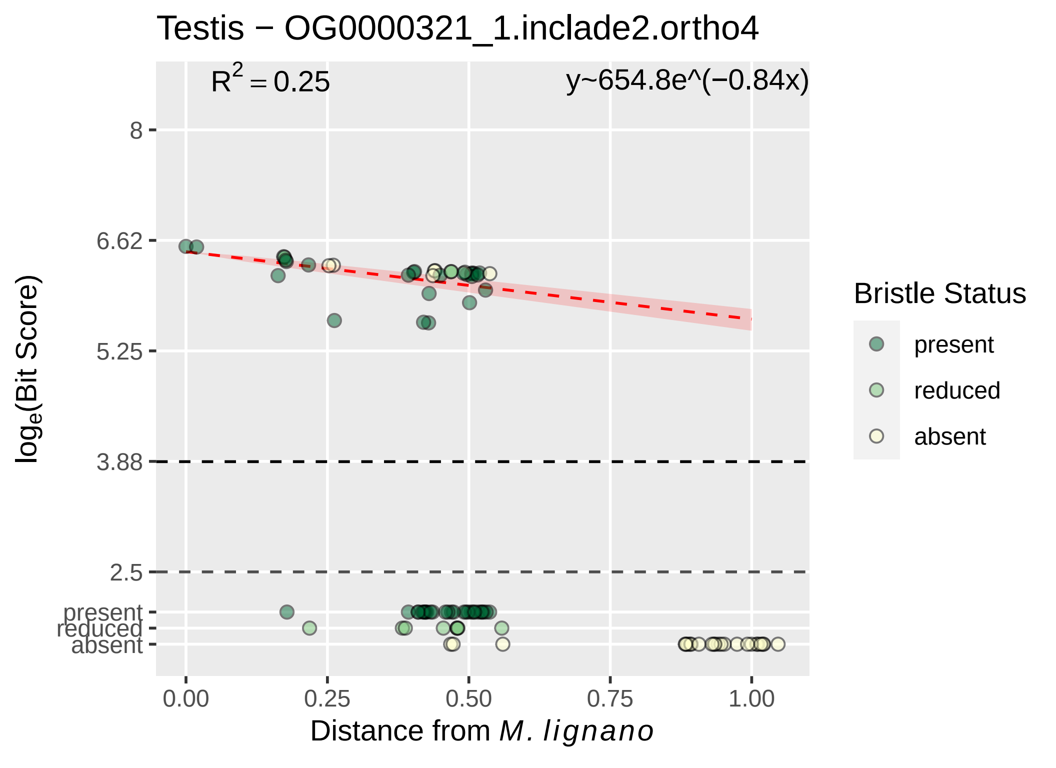

Testis − OG0000321_1.inclade2.ortho4
R^2 = 0.25
y~654.8e^(−0.84x)
8
6.62
5.25
3.88
2.5
present
reduced
absent
log_e(Bit Score)
0.00
0.25
0.50
0.75
1.00
Distance from M. lignano
Bristle Status
present
reduced
absent

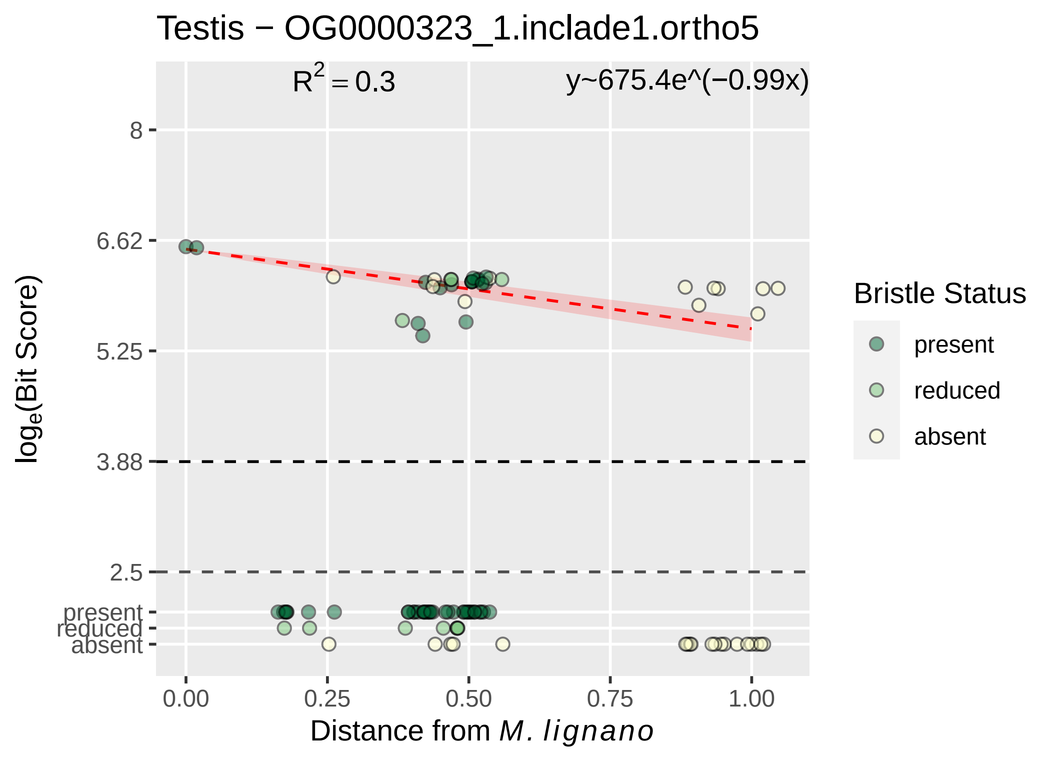
Testis − OG0000323_1.inclade1.ortho5
$R^2 = 0.3$
y~675.4e^(−0.99x)
8
6.62
5.25
3.88
2.5
present
reduced
absent
$\log_e$(Bit Score)
0.00
0.25
0.50
0.75
1.00
Distance from *M. lignano*
Bristle Status
present
reduced
absent

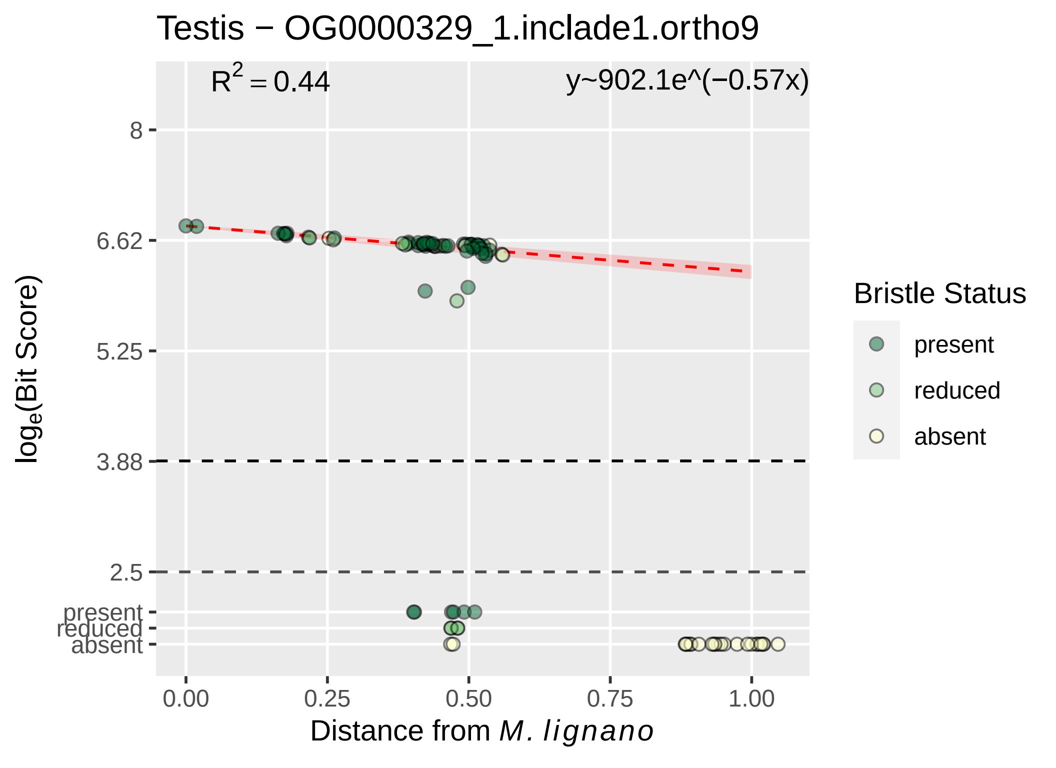

Testis − OG0000329_1.inclade1.ortho9
R^2 = 0.44
y~902.1e^(−0.57x)
8
6.62
5.25
3.88
2.5
present
reduced
absent
log_e(Bit Score)
0.00
0.25
0.50
0.75
1.00
Distance from M. lignano
Bristle Status
present
reduced
absent

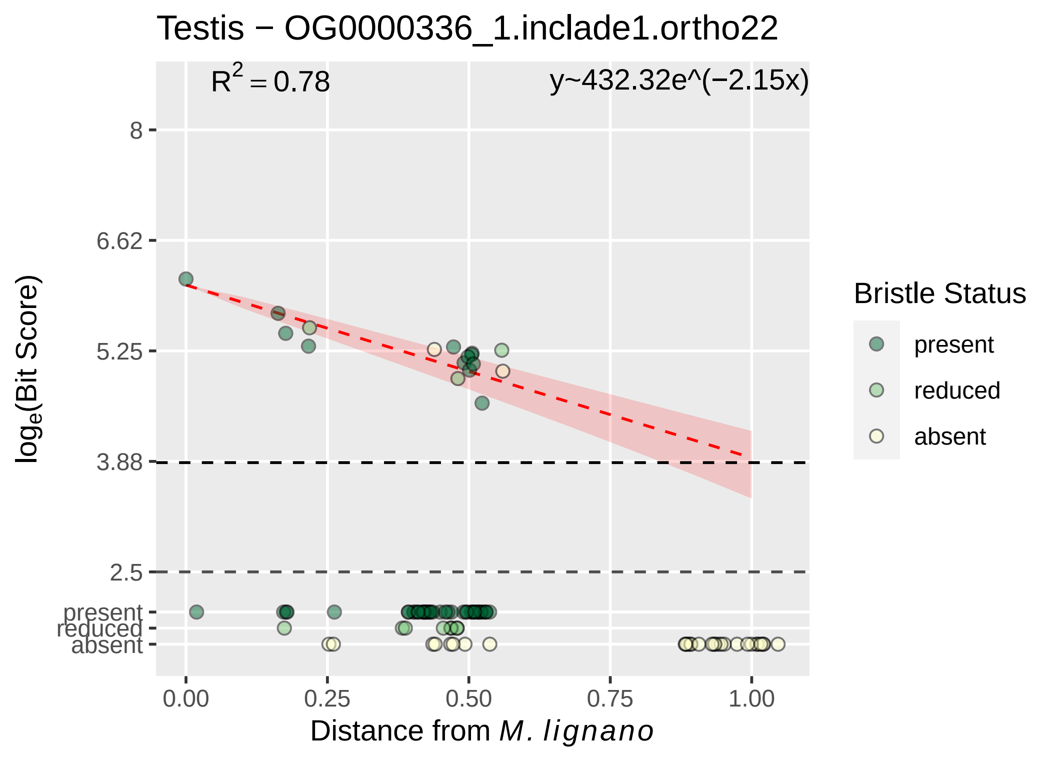

Testis − OG0000336_1.inclade1.ortho22
$R^2 = 0.78$
y~432.32e^(−2.15x)
8
6.62
5.25
3.88
2.5
present
reduced
absent
loge(Bit Score)
0.00
0.25
0.50
0.75
1.00
Distance from M. lignano
Bristle Status
present
reduced
absent

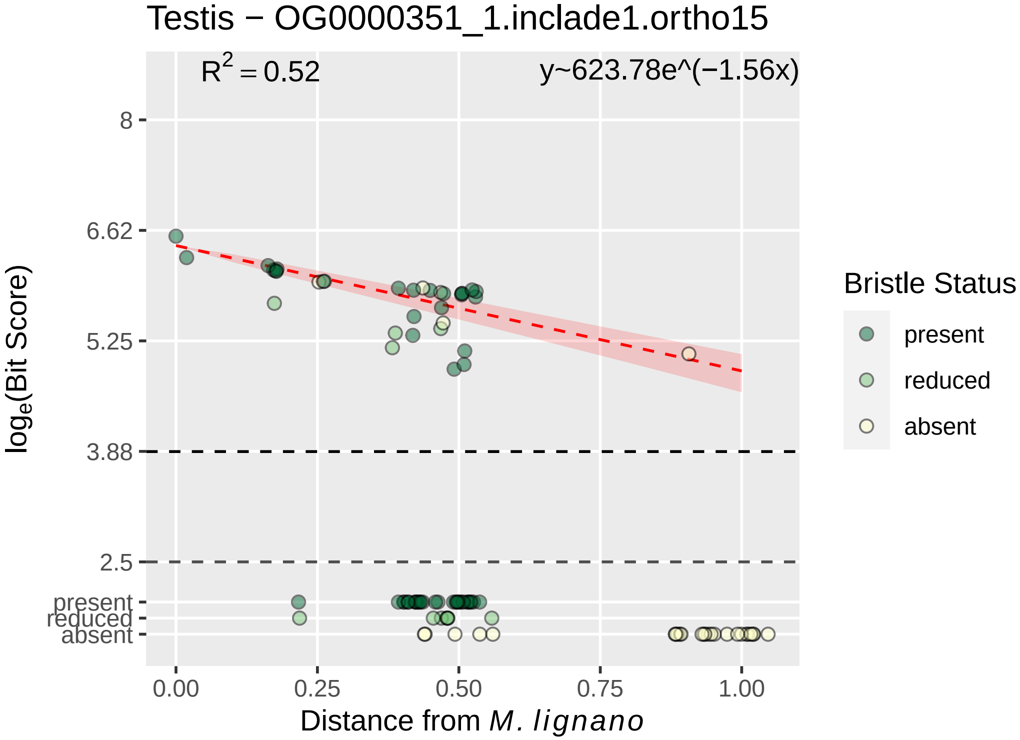
Testis − OG0000351_1.inclade1.ortho15
$R^2 = 0.52$
y~623.78e^(−1.56x)
8
6.62
5.25
3.88
2.5
present
reduced
absent
0.00
0.25
0.50
0.75
1.00
Distance from *M. lignano*
$\log_e$(Bit Score)
Bristle Status
present
reduced
absent

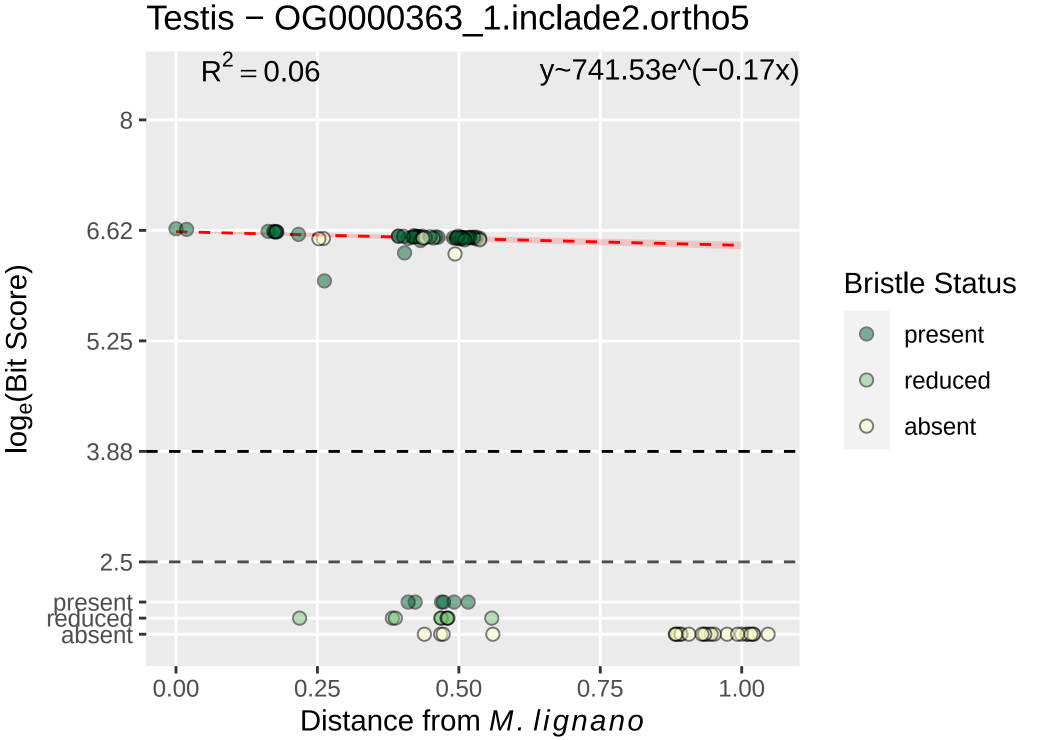

Testis − OG0000363_1.inclade2.ortho5
R2 = 0.06
y~741.53e^(−0.17x)
8
6.62
5.25
3.88
2.5
present
reduced
absent
loge(Bit Score)
0.00
0.25
0.50
0.75
1.00
Distance from M. lignano
Bristle Status
present
reduced
absent

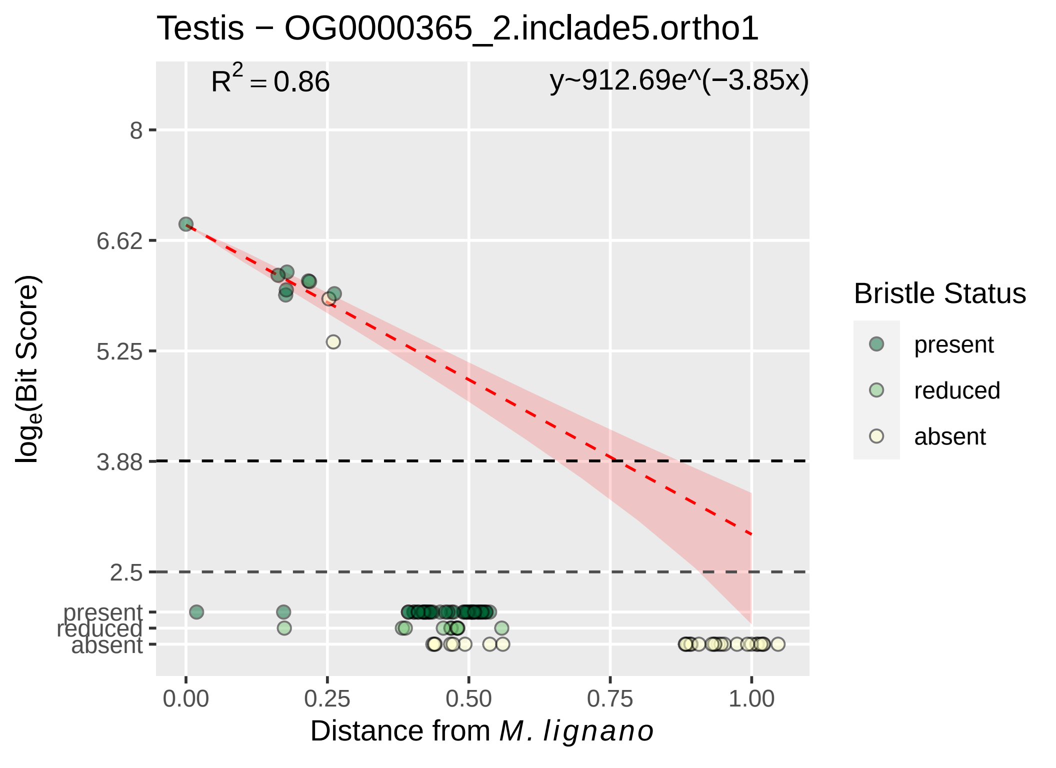

Testis − OG0000365_2.inclade5.ortho1
$R^2 = 0.86$
y~912.69e^(−3.85x)
8
6.62
5.25
3.88
2.5
present
reduced
absent
0.00
0.25
0.50
0.75
1.00
$\log_e$(Bit Score)
Distance from M. lignano
Bristle Status
present
reduced
absent

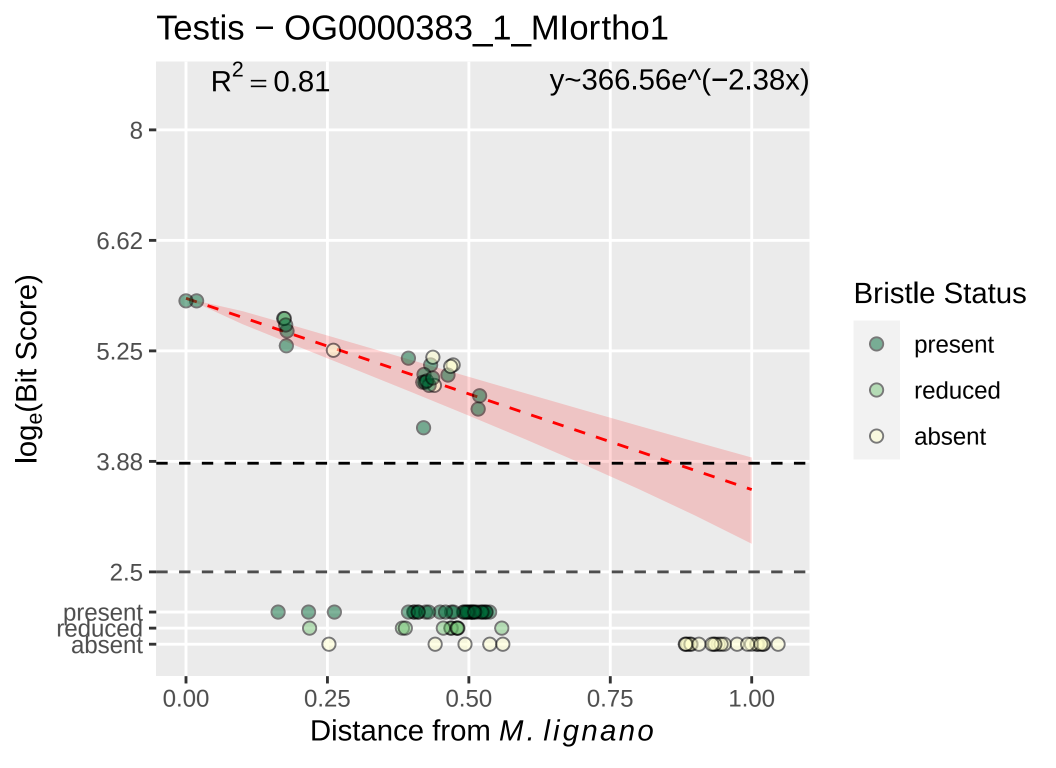

Testis − OG0000383_1_MIortho1
$R^2 = 0.81$
y~366.56e^(−2.38x)
8
6.62
5.25
3.88
2.5
present
reduced
absent
$\log_e$(Bit Score)
0.00
0.25
0.50
0.75
1.00
Distance from *M. lignano*
Bristle Status
present
reduced
absent

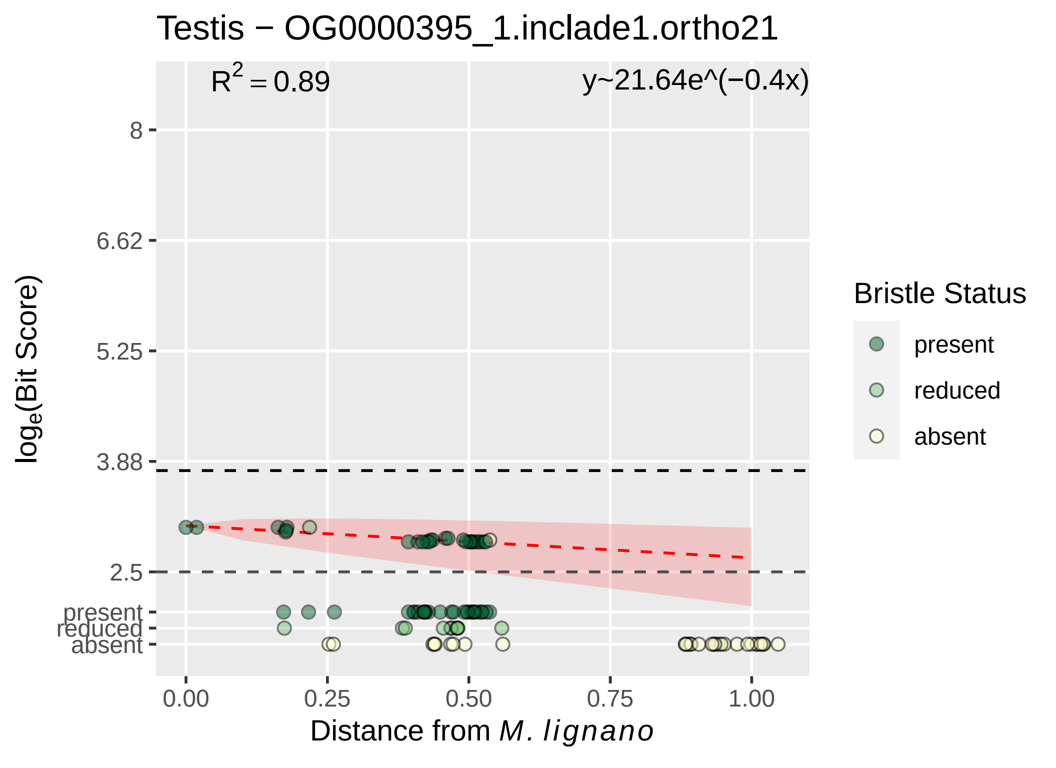
Testis − OG0000395_1.inclade1.ortho21
R² = 0.89
y~21.64e^(−0.4x)
8
6.62
5.25
3.88
2.5
present
reduced
absent
log_e(Bit Score)
0.00
0.25
0.50
0.75
1.00
Distance from M. lignano
Bristle Status
present
reduced
absent

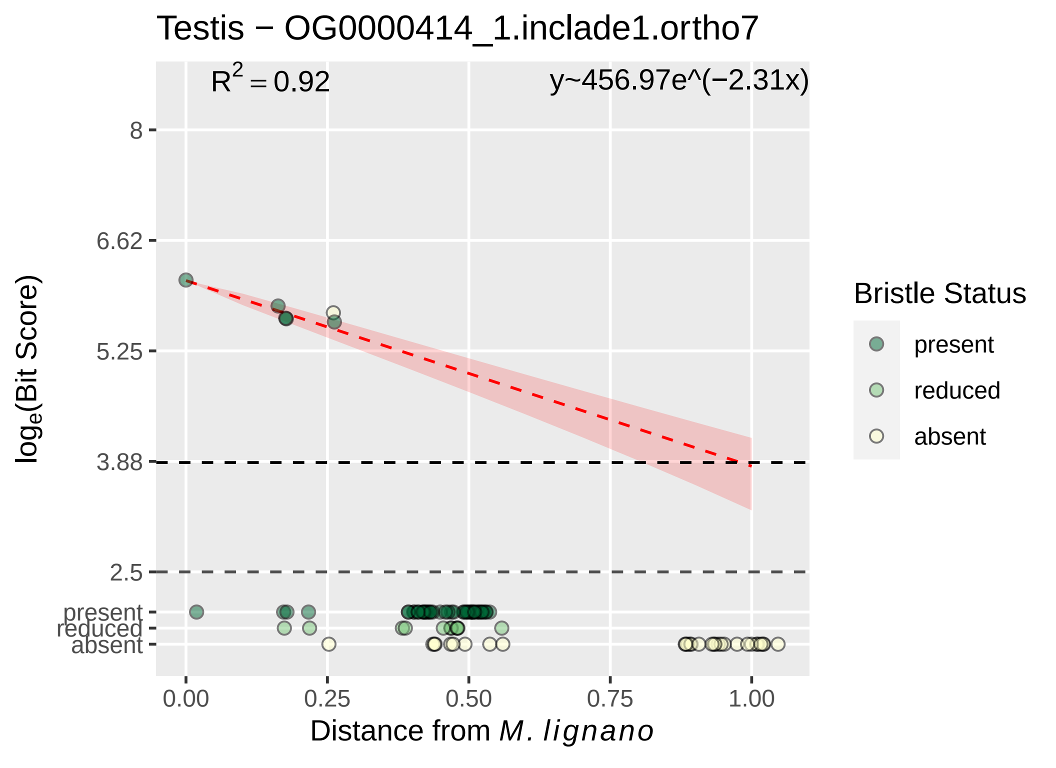

Testis − OG0000414_1.inclade1.ortho7
R^2 = 0.92
y~456.97e^(−2.31x)
8
6.62
5.25
3.88
2.5
present
reduced
absent
0.00
0.25
0.50
0.75
1.00
Distance from M. lignano
log_e(Bit Score)
Bristle Status
present
reduced
absent

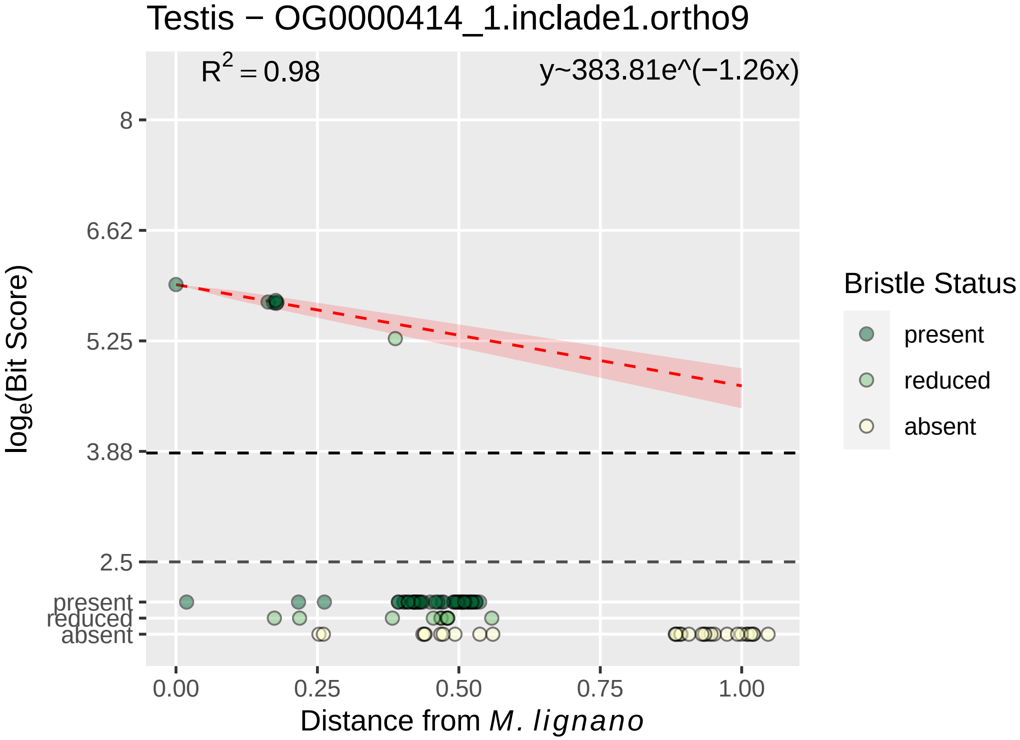
Testis − OG0000414_1.inclade1.ortho9
$R^2 = 0.98$
y~383.81e^(−1.26x)
8
6.62
5.25
3.88
2.5
present
reduced
absent
$\log_e$(Bit Score)
0.00
0.25
0.50
0.75
1.00
Distance from *M. lignano*
Bristle Status
present
reduced
absent

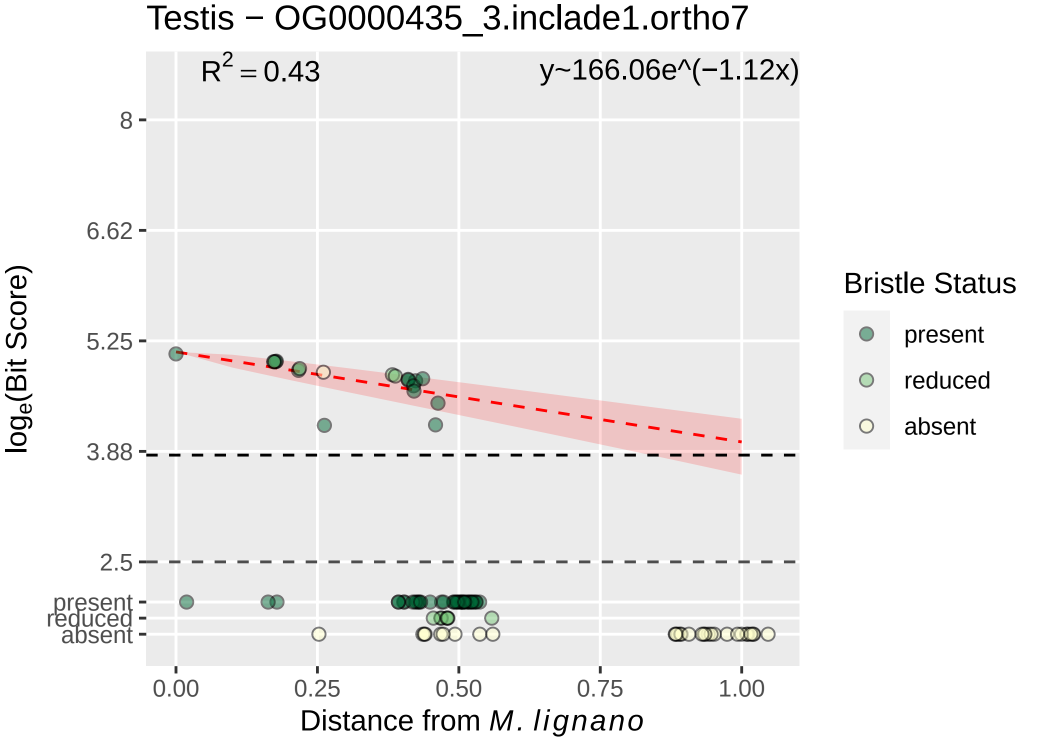
Testis − OG0000435_3.inclade1.ortho7
$R^2 = 0.43$
y~166.06e^(−1.12x)
8
6.62
5.25
3.88
2.5
present
reduced
absent
$\log_e$(Bit Score)
0.00
0.25
0.50
0.75
1.00
Distance from *M. lignano*
Bristle Status
present
reduced
absent

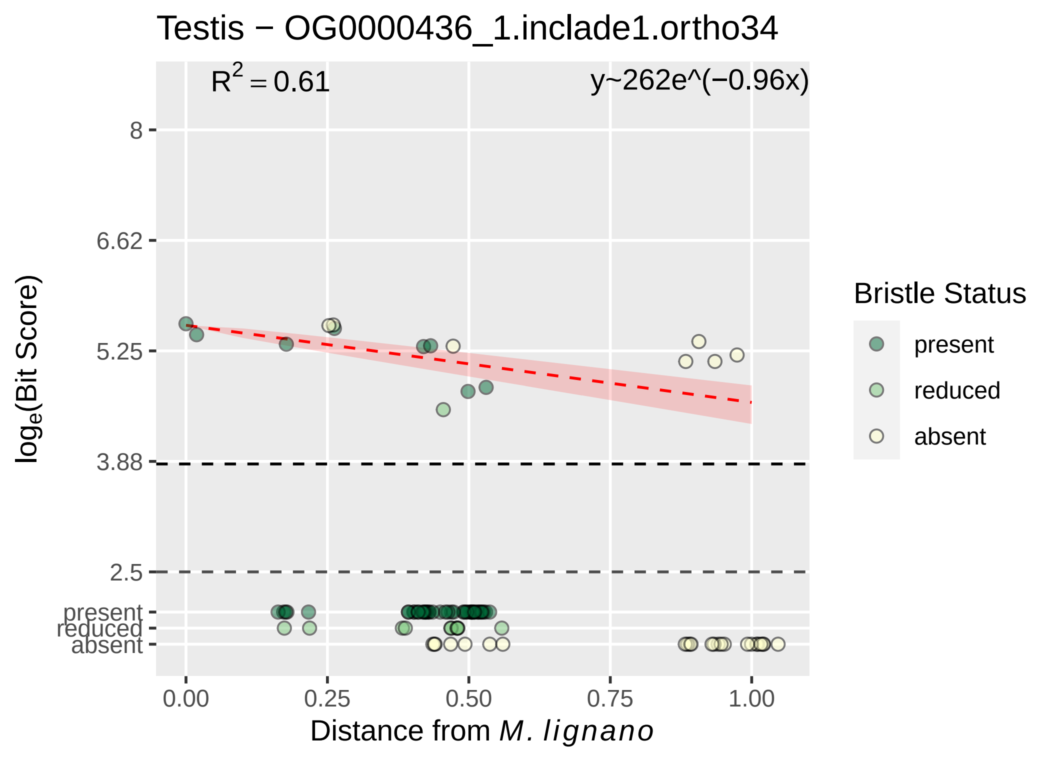

Testis − OG0000436_1.inclade1.ortho34
R^2 = 0.61
y~262e^(−0.96x)
8
6.62
5.25
3.88
2.5
present
reduced
absent
log_e(Bit Score)
0.00
0.25
0.50
0.75
1.00
Distance from M. lignano
Bristle Status
present
reduced
absent

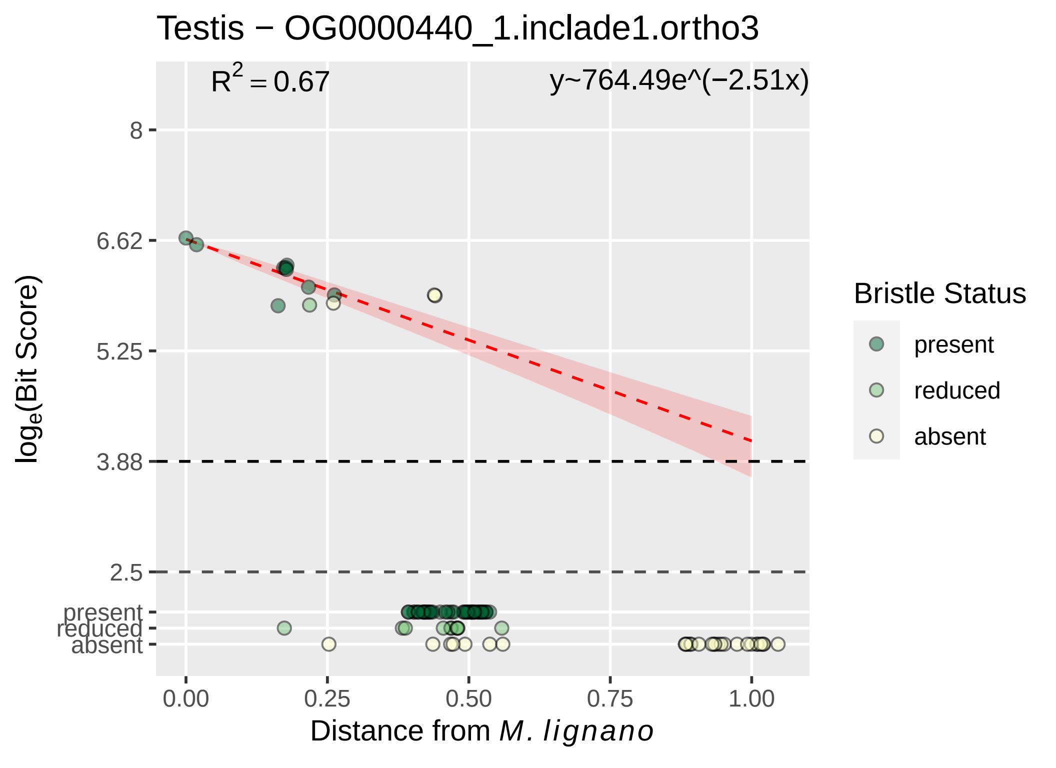

Testis − OG0000440_1.inclade1.ortho3
$R^2 = 0.67$
y~764.49e^(−2.51x)
8
6.62
5.25
3.88
2.5
present
reduced
absent
0.00
0.25
0.50
0.75
1.00
Distance from *M. lignano*
$log_e$(Bit Score)
Bristle Status
present
reduced
absent

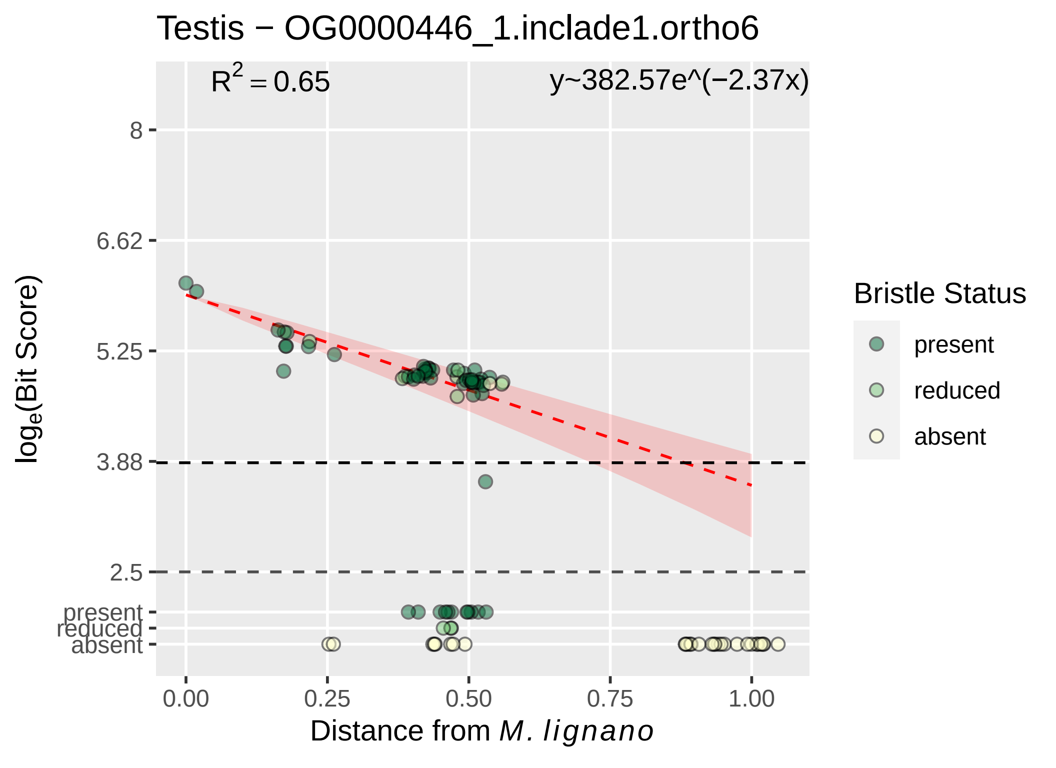

Testis − OG0000446_1.inclade1.ortho6
R2 = 0.65
y~382.57e^(−2.37x)
8
6.62
5.25
3.88
2.5
present
reduced
absent
loge(Bit Score)
0.00
0.25
0.50
0.75
1.00
Distance from M. lignano
Bristle Status
present
reduced
absent

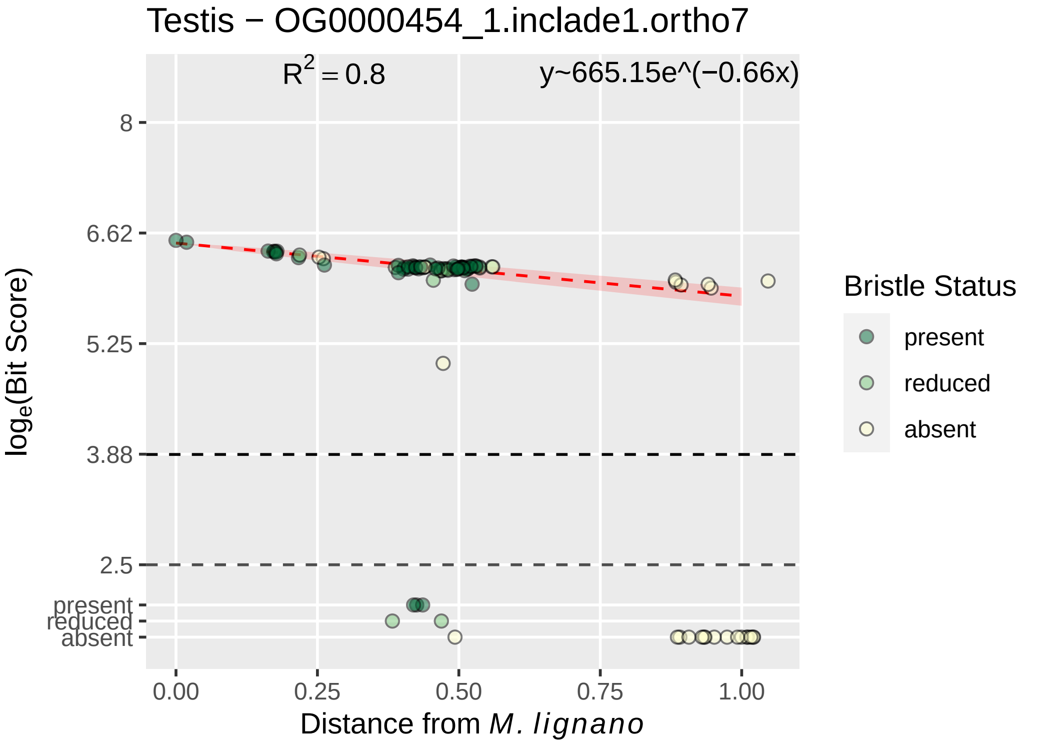
Testis − OG0000454_1.inclade1.ortho7
R^2 = 0.8
y~665.15e^(−0.66x)
8
6.62
5.25
3.88
2.5
present
reduced
absent
log_e(Bit Score)
0.00
0.25
0.50
0.75
1.00
Distance from M. lignano
Bristle Status
present
reduced
absent

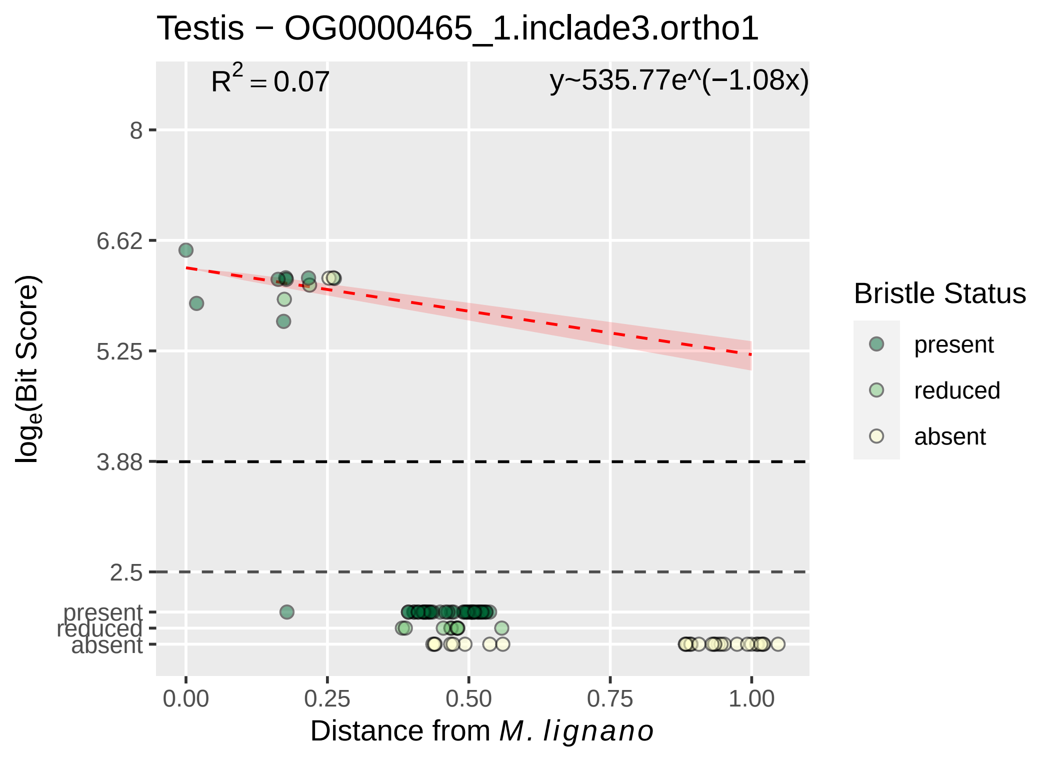

Testis − OG0000465_1.inclade3.ortho1
R^2 = 0.07
y~535.77e^(−1.08x)
8
6.62
5.25
3.88
2.5
present
reduced
absent
log_e(Bit Score)
0.00
0.25
0.50
0.75
1.00
Distance from M. lignano
Bristle Status
present
reduced
absent

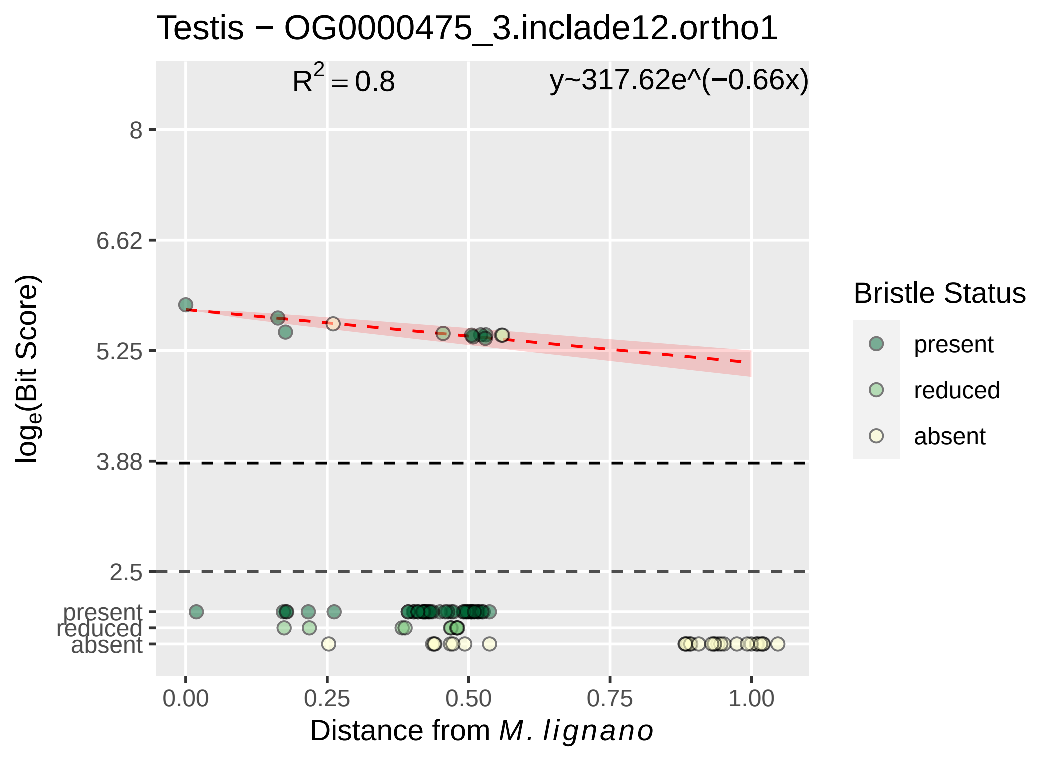

Testis − OG0000475_3.inclade12.ortho1
$R^2 = 0.8$
y~317.62e^(−0.66x)
8
6.62
5.25
3.88
2.5
present
reduced
absent
0.00
0.25
0.50
0.75
1.00
$log_e$(Bit Score)
Distance from *M. lignano*
Bristle Status
present
reduced
absent

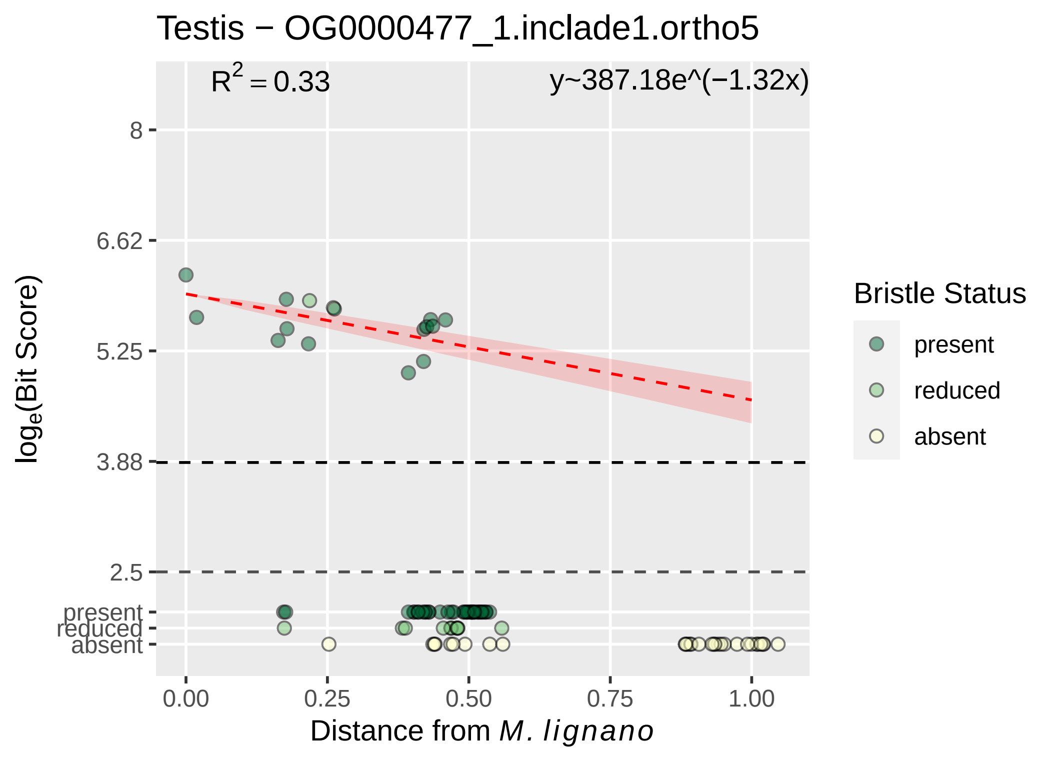

Testis − OG0000477_1.inclade1.ortho5
R^2 = 0.33
y~387.18e^(−1.32x)
8
6.62
5.25
3.88
2.5
present
reduced
absent
log_e(Bit Score)
0.00
0.25
0.50
0.75
1.00
Distance from M. lignano
Bristle Status
present
reduced
absent

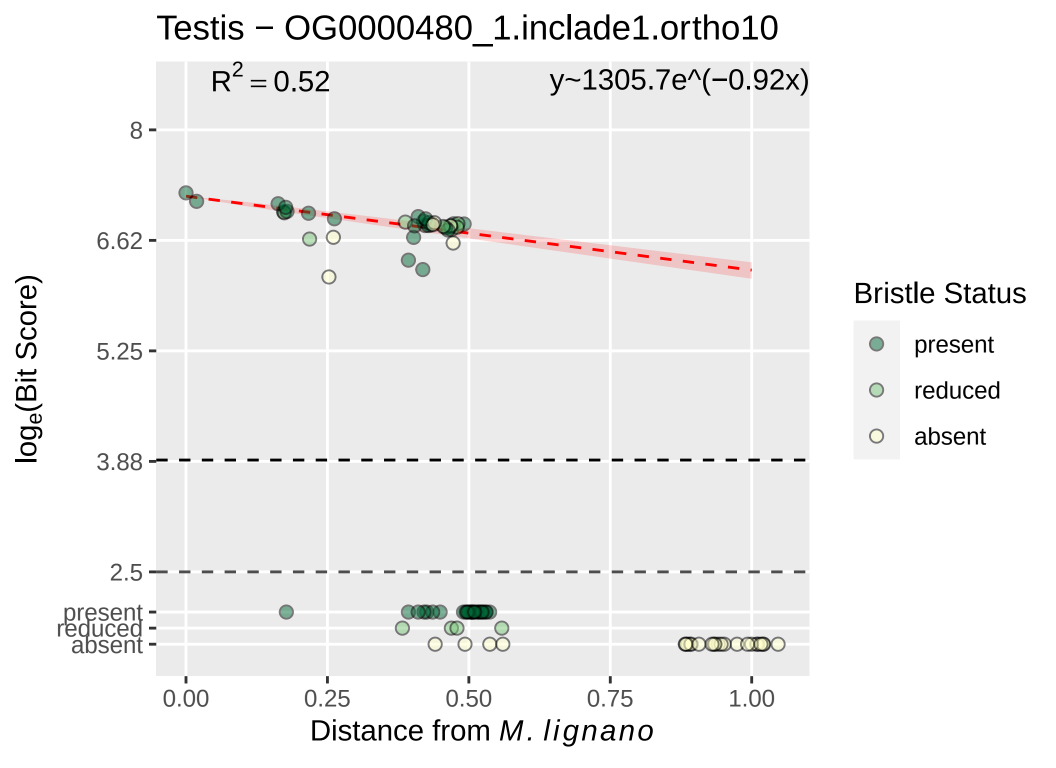

Testis − OG0000480_1.inclade1.ortho10
$R^2 = 0.52$
y~1305.7e^(−0.92x)
8
6.62
5.25
3.88
2.5
present
reduced
absent
0.00
0.25
0.50
0.75
1.00
$\log_e$(Bit Score)
Distance from *M. lignano*
Bristle Status
present
reduced
absent

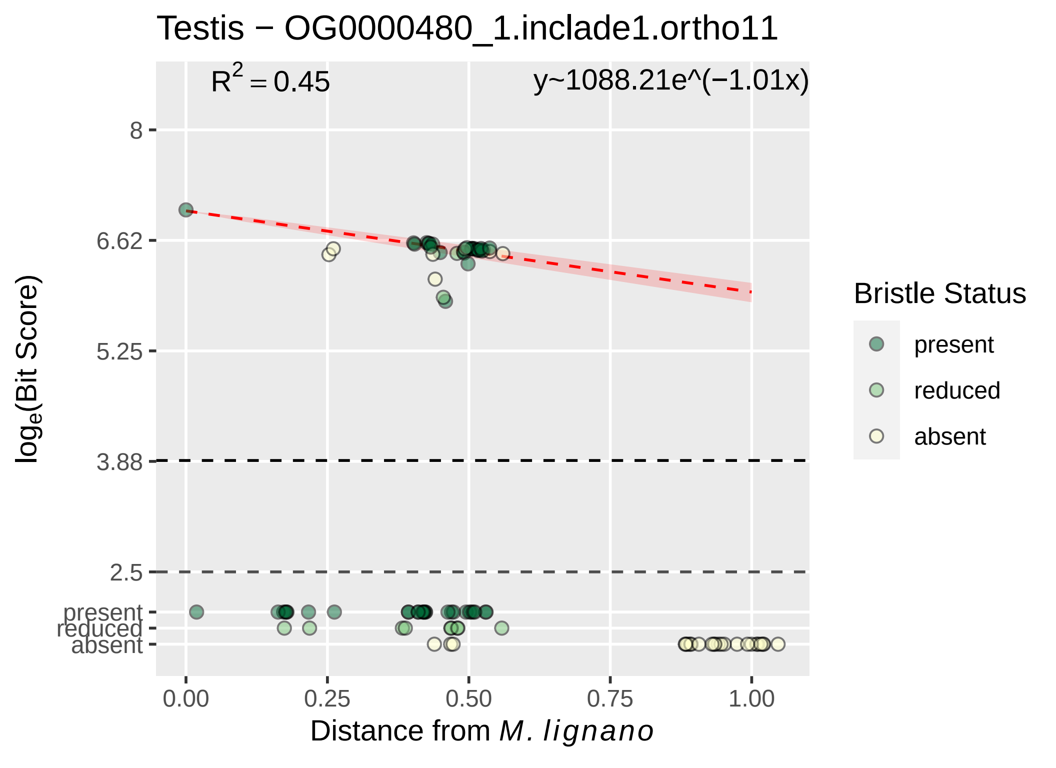
Testis − OG0000480_1.inclade1.ortho11
$R^2 = 0.45$
y~1088.21e^(−1.01x)
8
6.62
5.25
3.88
2.5
present
reduced
absent
0.00
0.25
0.50
0.75
1.00
$\log_e$(Bit Score)
Distance from M. lignano
Bristle Status
present
reduced
absent

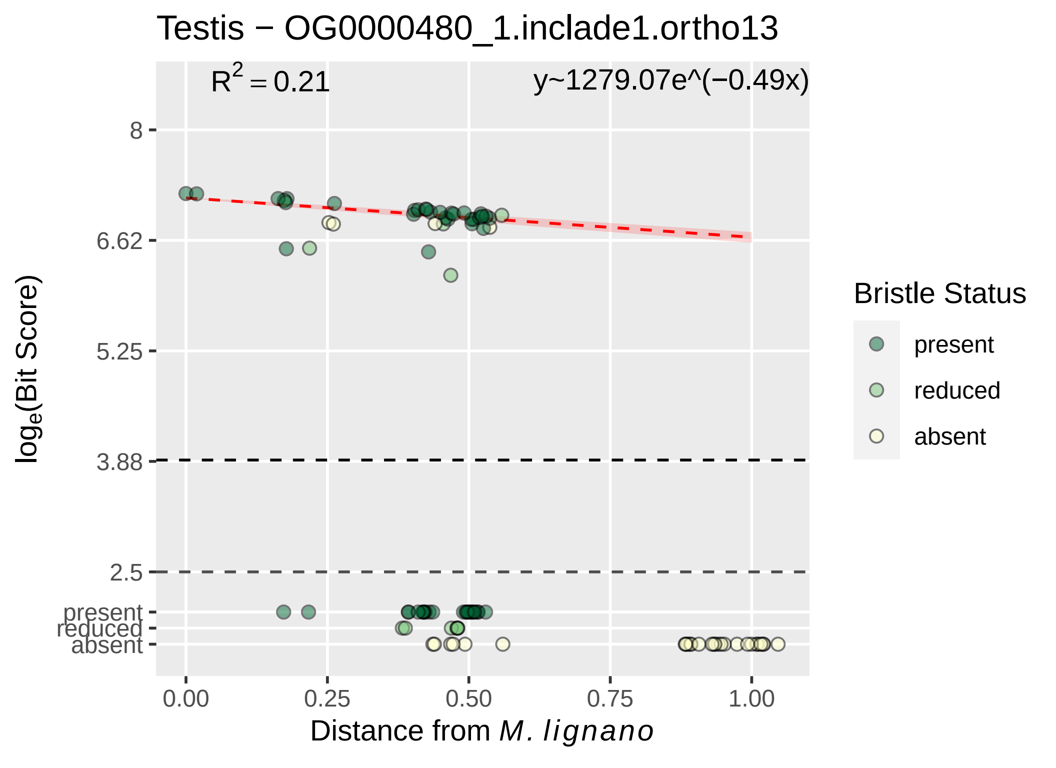
Testis − OG0000480_1.inclade1.ortho13
$R^2 = 0.21$
y~1279.07e^(−0.49x)
8
6.62
5.25
3.88
2.5
present
reduced
absent
0.00
0.25
0.50
0.75
1.00
Distance from M. lignano
$\log_e$(Bit Score)
Bristle Status
present
reduced
absent

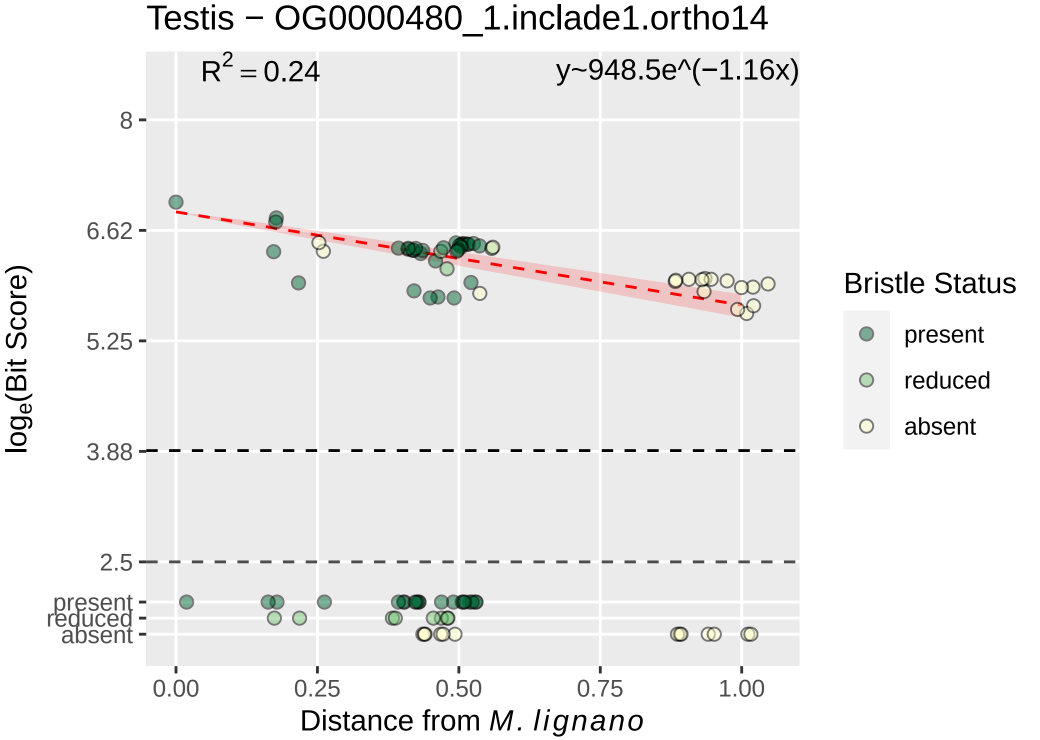

Testis − OG0000480_1.inclade1.ortho14
$R^2 = 0.24$
y~948.5e^(−1.16x)
8
6.62
5.25
3.88
2.5
present
reduced
absent
0.00
0.25
0.50
0.75
1.00
log$_e$(Bit Score)
Distance from *M. lignano*
Bristle Status
present
reduced
absent

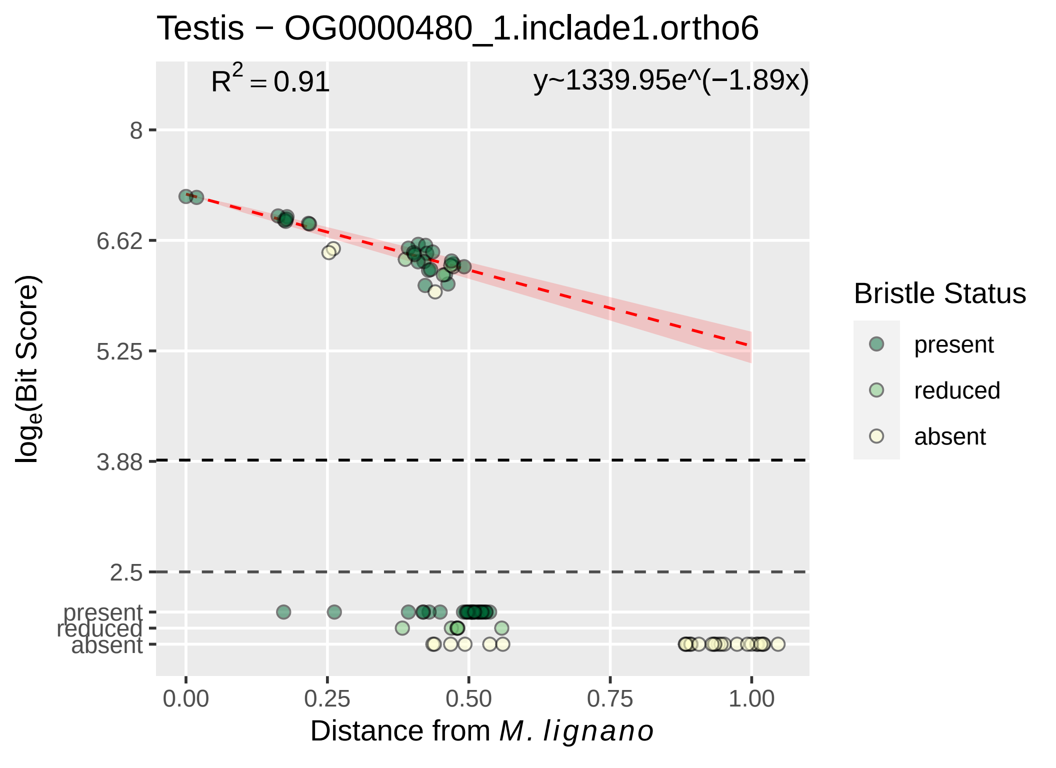
Testis − OG0000480_1.inclade1.ortho6
$R^2 = 0.91$
y~1339.95e^(−1.89x)
8
6.62
5.25
3.88
2.5
present
reduced
absent
$\log_e$(Bit Score)
0.00
0.25
0.50
0.75
1.00
Distance from *M. lignano*
Bristle Status
present
reduced
absent

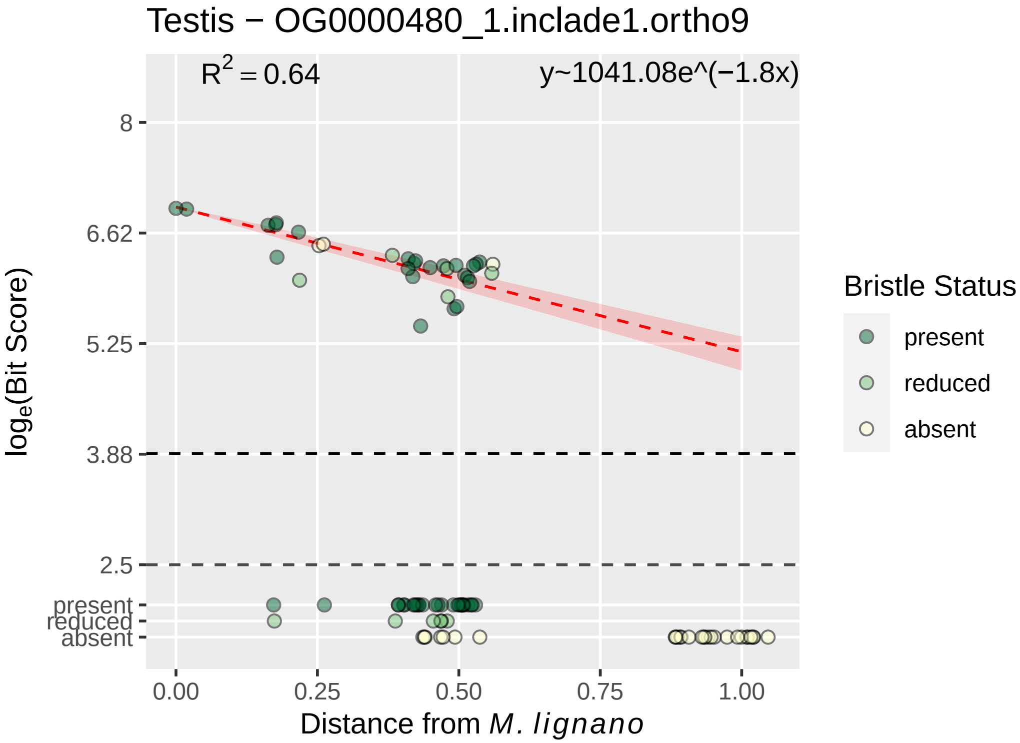
Testis − OG0000480_1.inclade1.ortho9
R^2 = 0.64
y~1041.08e^(−1.8x)
8
6.62
5.25
3.88
2.5
present
reduced
absent
loge(Bit Score)
0.00
0.25
0.50
0.75
1.00
Distance from M. lignano
Bristle Status
present
reduced
absent

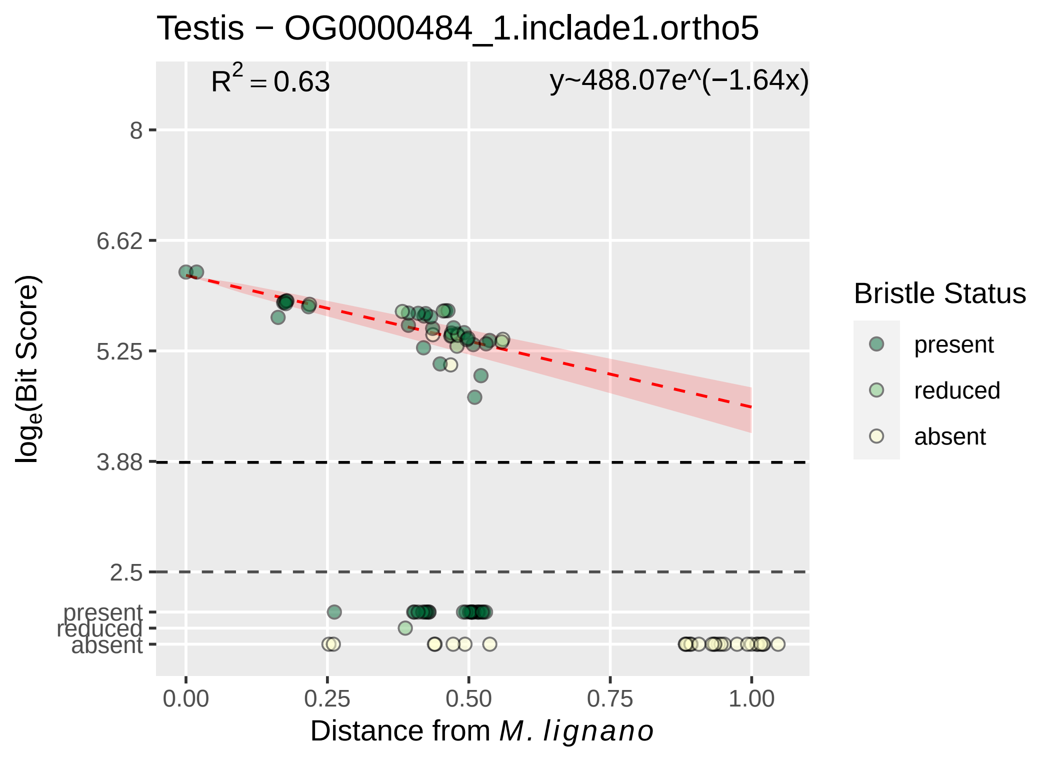

Testis − OG0000484_1.inclade1.ortho5
$R^2 = 0.63$
y~488.07e^(−1.64x)
8
6.62
5.25
3.88
2.5
present
reduced
absent
$log_e$(Bit Score)
0.00
0.25
0.50
0.75
1.00
Distance from *M. lignano*
Bristle Status
present
reduced
absent

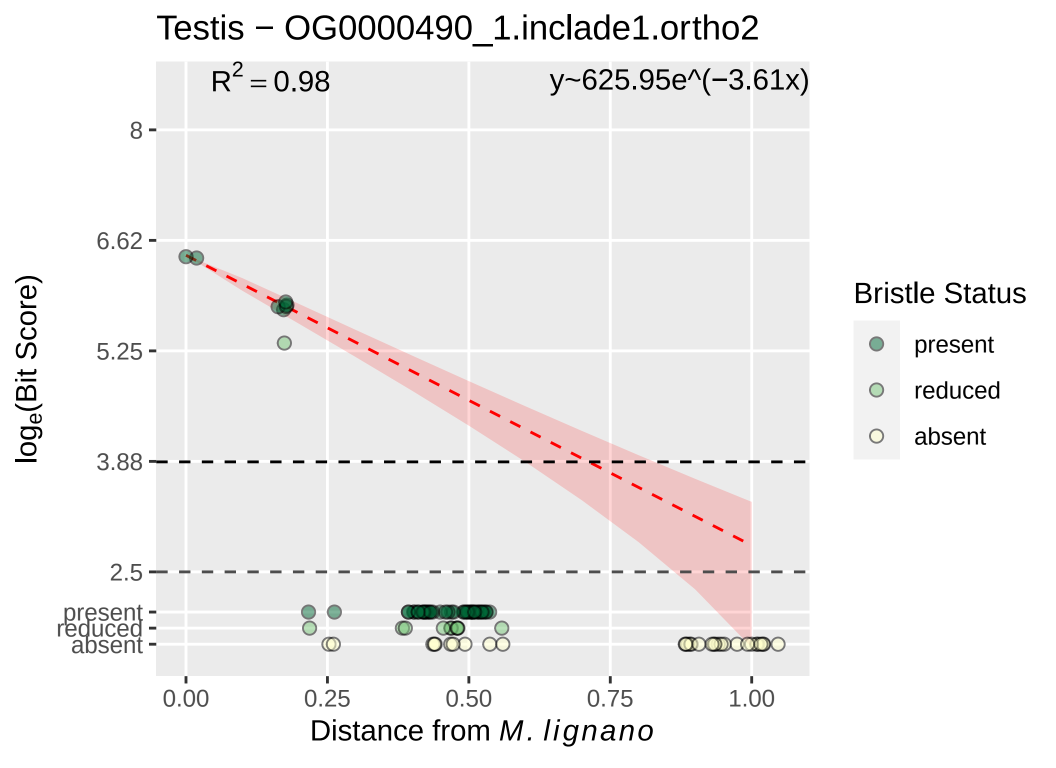
Testis − OG0000490_1.inclade1.ortho2
$R^2 = 0.98$
y~625.95e^(−3.61x)
8
6.62
5.25
3.88
2.5
present
reduced
absent
$\log_e$(Bit Score)
0.00
0.25
0.50
0.75
1.00
Distance from *M. lignano*
Bristle Status
present
reduced
absent

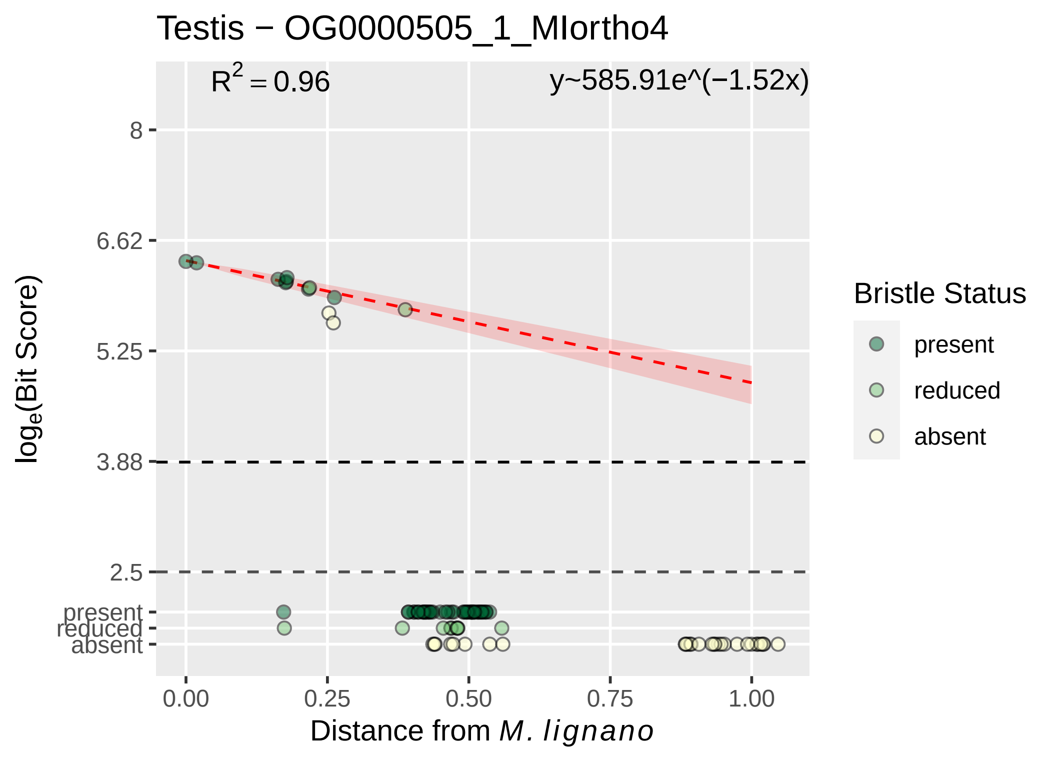
Testis − OG0000505_1_MIortho4
R² = 0.96
y~585.91e^(−1.52x)
8
6.62
5.25
3.88
2.5
present
reduced
absent
loge(Bit Score)
0.00
0.25
0.50
0.75
1.00
Distance from M. lignano
Bristle Status
present
reduced
absent

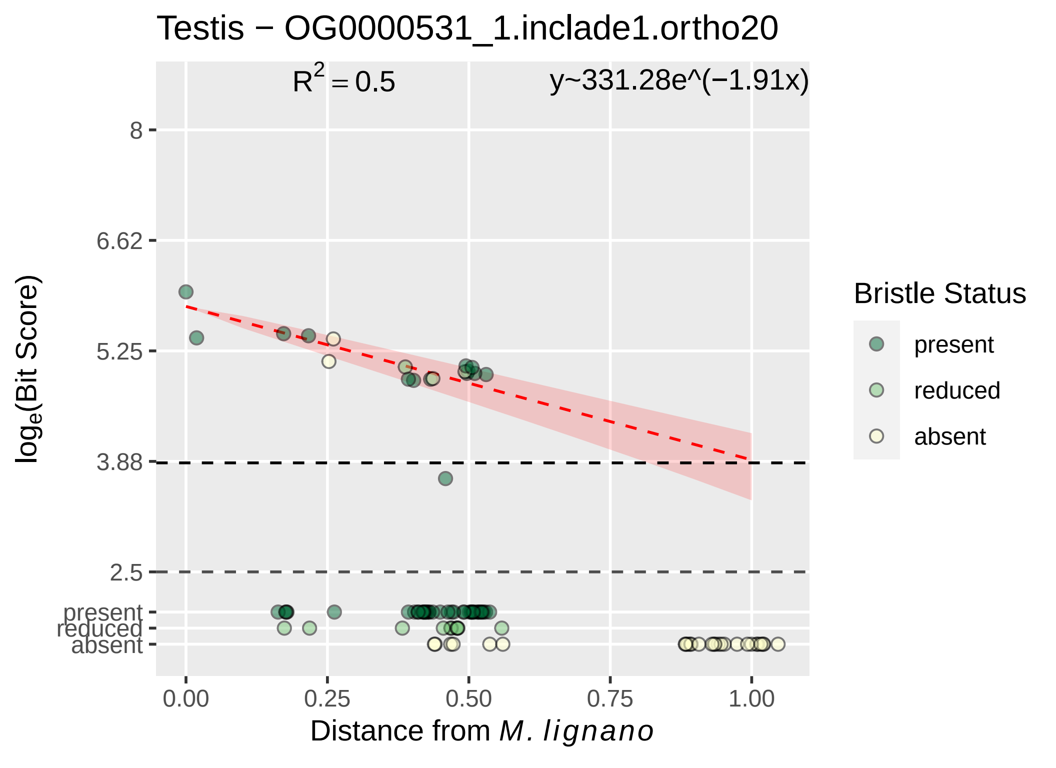

Testis − OG0000531_1.inclade1.ortho20
$R^2 = 0.5$
y~331.28e^(−1.91x)
8
6.62
5.25
3.88
2.5
present
reduced
absent
$log_e$(Bit Score)
0.00
0.25
0.50
0.75
1.00
Distance from *M. lignano*
Bristle Status
present
reduced
absent

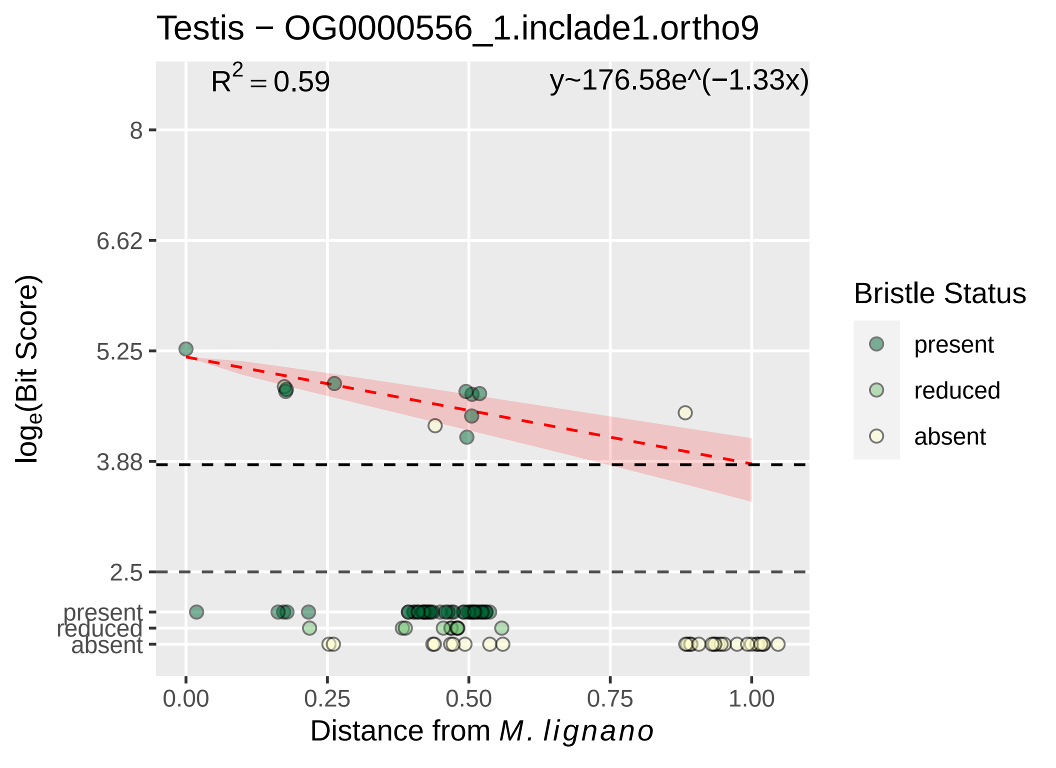

Testis − OG0000556_1.inclade1.ortho9
R^2 = 0.59
y~176.58e^(−1.33x)
8
6.62
5.25
3.88
2.5
present
reduced
absent
log_e(Bit Score)
0.00
0.25
0.50
0.75
1.00
Distance from M. lignano
Bristle Status
present
reduced
absent

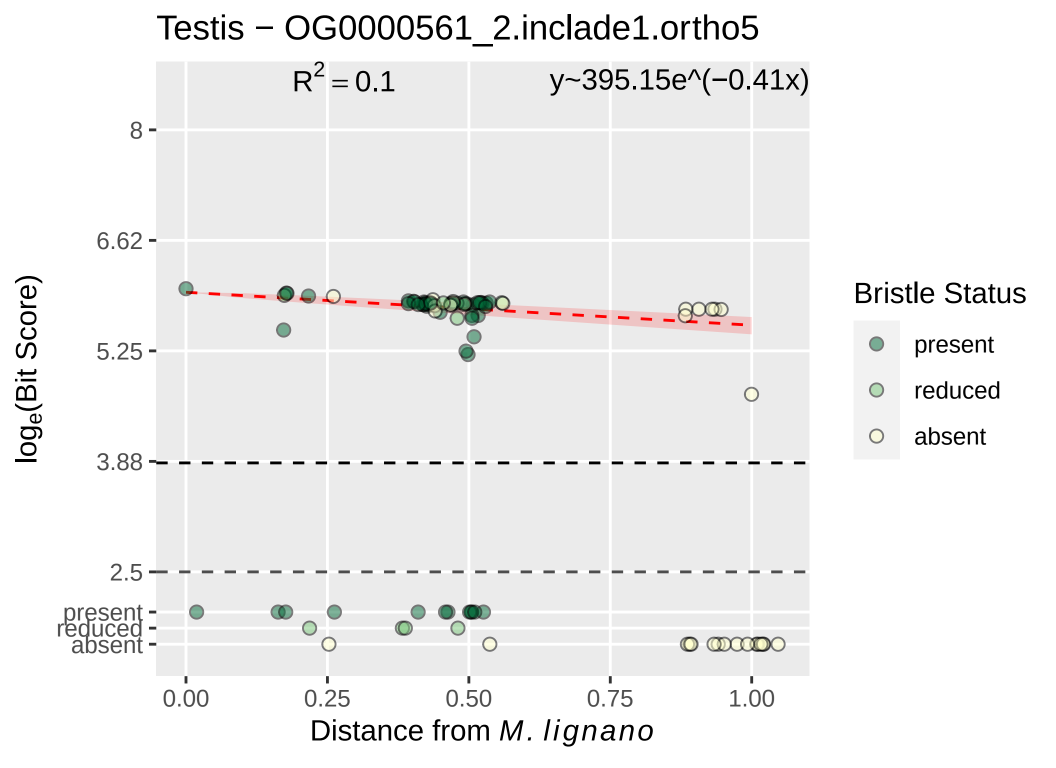
Testis − OG0000561_2.inclade1.ortho5
$R^2 = 0.1$
y~395.15e^(−0.41x)
8
6.62
5.25
3.88
2.5
present
reduced
absent
$log_e$(Bit Score)
0.00
0.25
0.50
0.75
1.00
Distance from M. lignano
Bristle Status
present
reduced
absent

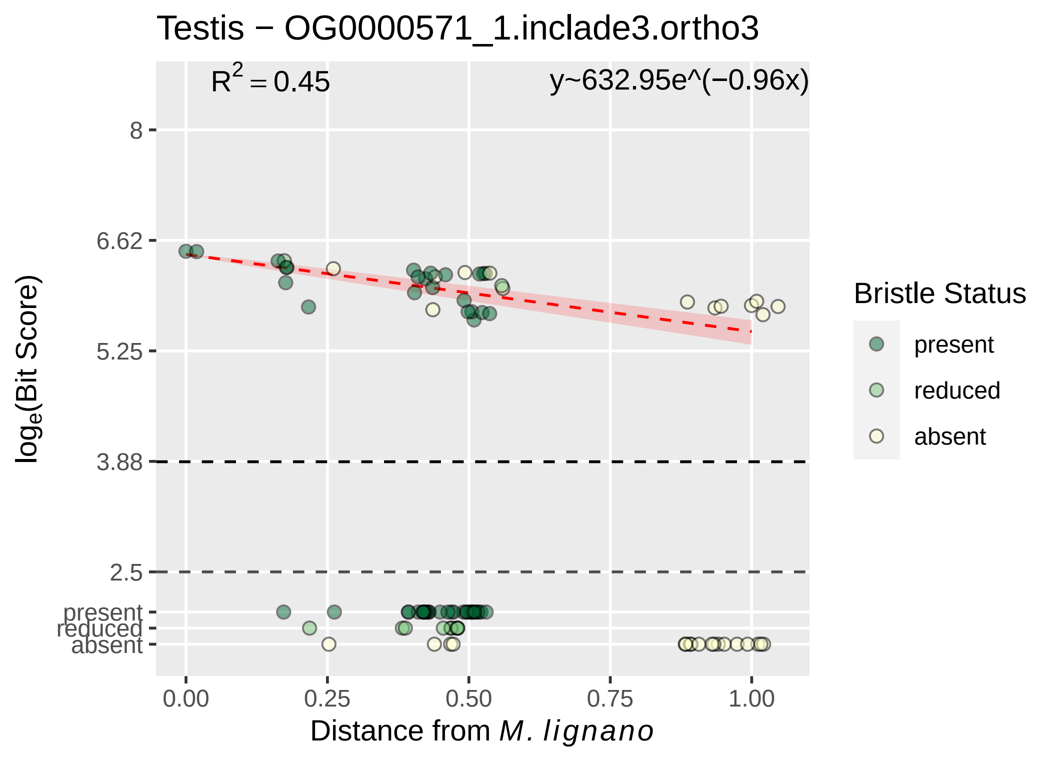

Testis − OG0000571_1.inclade3.ortho3
R2 = 0.45
y~632.95e^(−0.96x)
8
6.62
5.25
3.88
2.5
present
reduced
absent
loge(Bit Score)
0.00
0.25
0.50
0.75
1.00
Distance from M. lignano
Bristle Status
present
reduced
absent

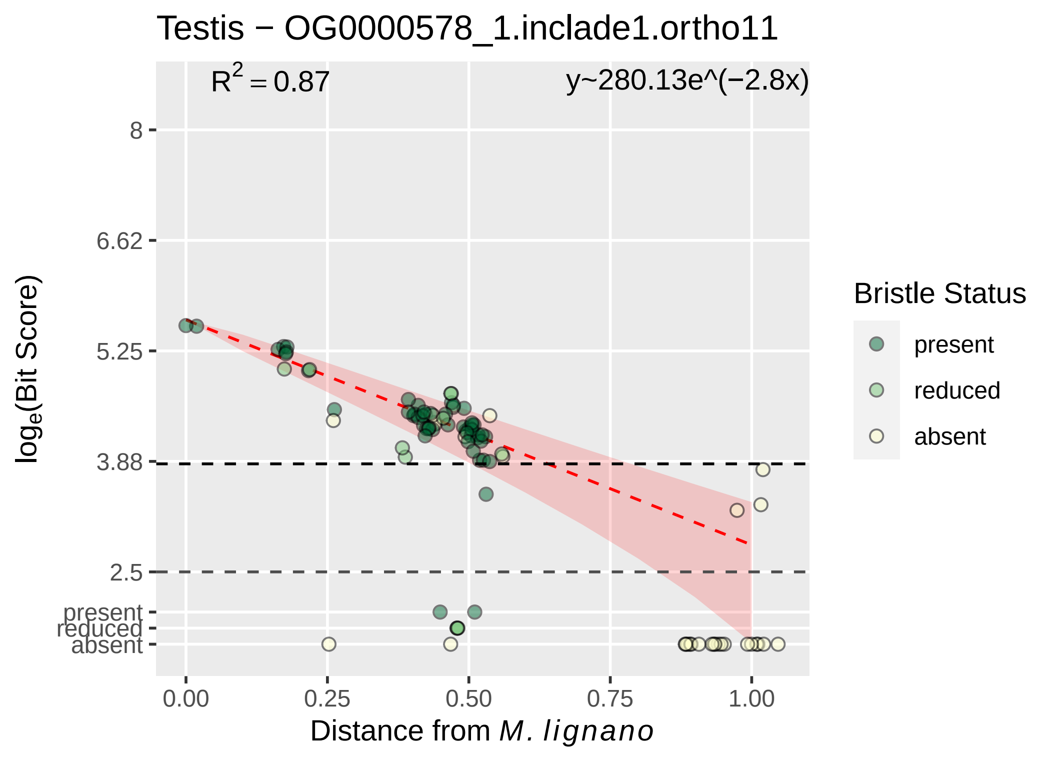

Testis − OG0000578_1.inclade1.ortho11
$R^2 = 0.87$
y~280.13e^(−2.8x)
8
6.62
5.25
3.88
2.5
present
reduced
absent
0.00
0.25
0.50
0.75
1.00
$\log_e$(Bit Score)
Distance from *M. lignano*
Bristle Status
present
reduced
absent

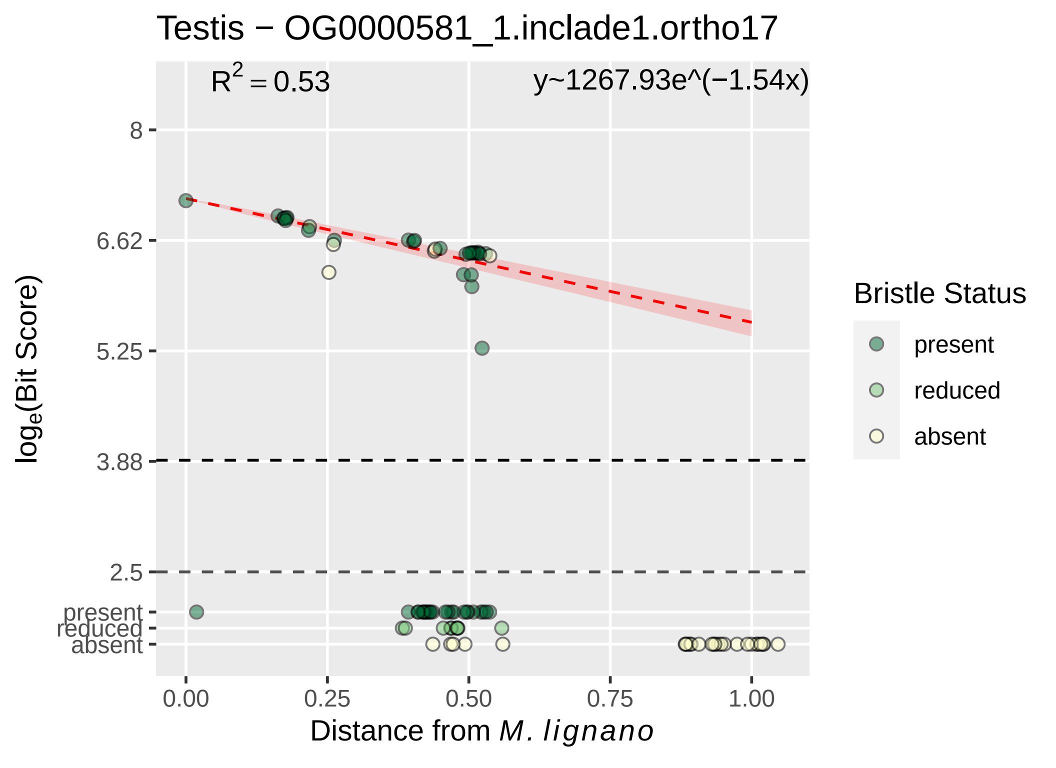

Testis − OG0000581_1.inclade1.ortho17
$R^2 = 0.53$
y~1267.93e^(−1.54x)
8
6.62
5.25
3.88
2.5
present
reduced
absent
$log_e$(Bit Score)
0.00
0.25
0.50
0.75
1.00
Distance from *M. lignano*
Bristle Status
present
reduced
absent

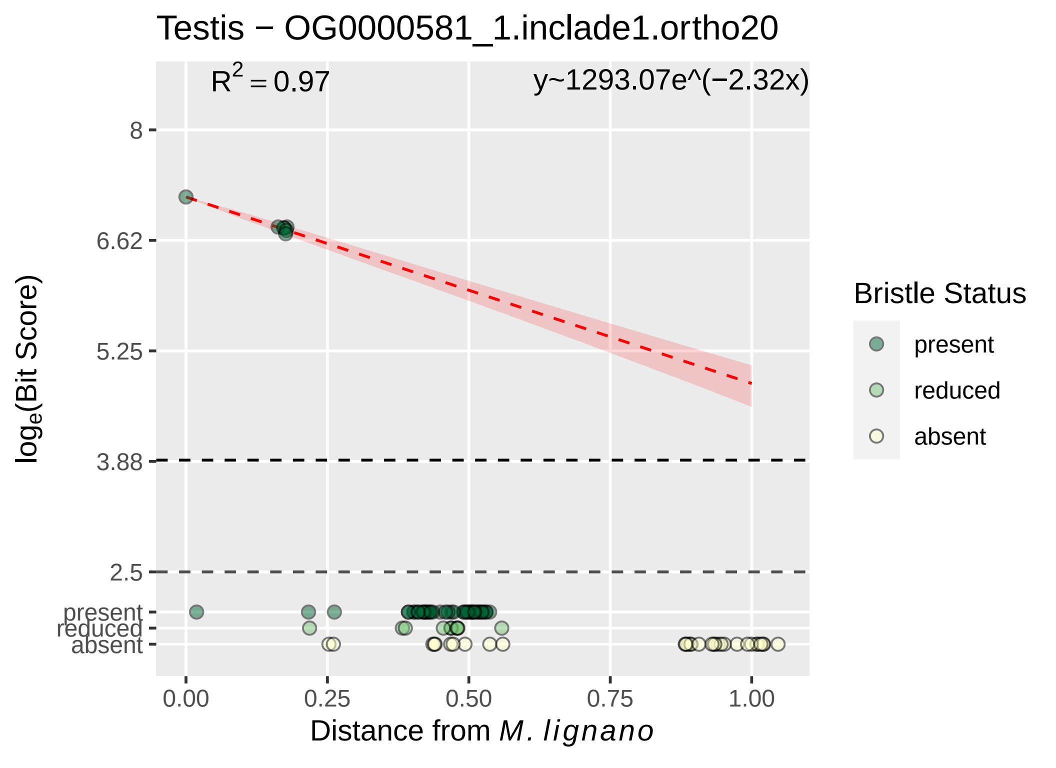

Testis − OG0000581_1.inclade1.ortho20
R^2 = 0.97
y~1293.07e^(−2.32x)
8
6.62
5.25
3.88
2.5
present
reduced
absent
0.00
0.25
0.50
0.75
1.00
Distance from M. lignano
loge(Bit Score)
Bristle Status
present
reduced
absent

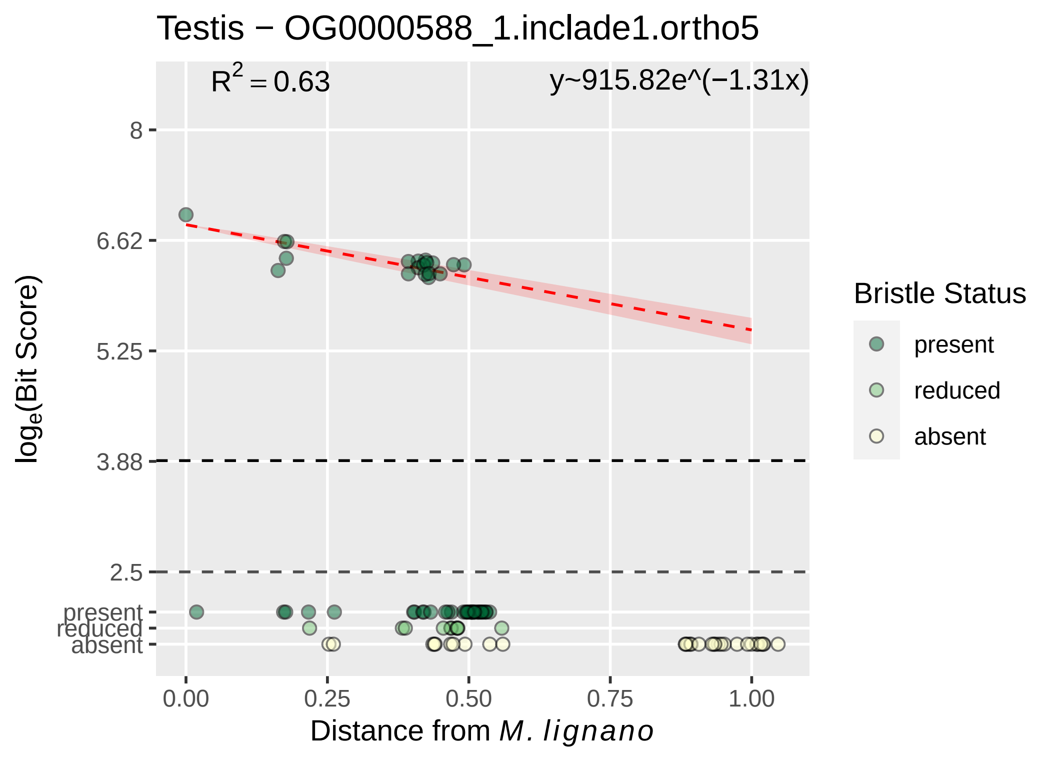

Testis − OG0000588_1.inclade1.ortho5
$R^2 = 0.63$
y~915.82e^(−1.31x)
8
6.62
5.25
3.88
2.5
present
reduced
absent
0.00
0.25
0.50
0.75
1.00
$\log_e$(Bit Score)
Distance from *M. lignano*
Bristle Status
present
reduced
absent

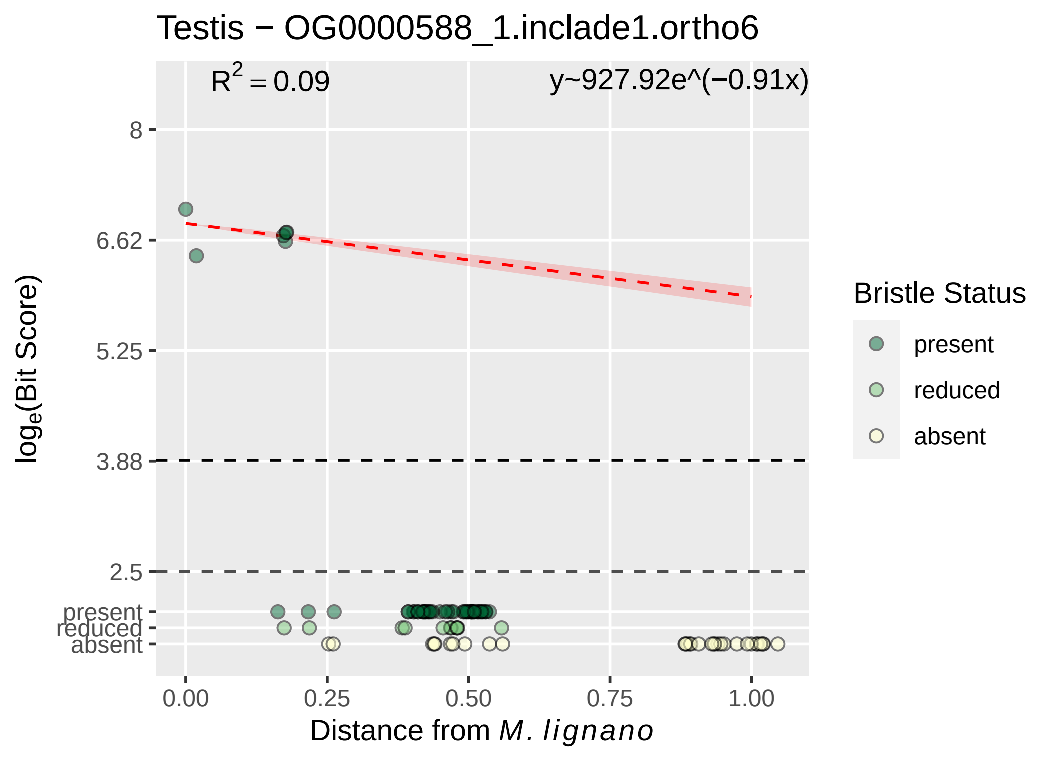

Testis − OG0000588_1.inclade1.ortho6
R2 = 0.09
y~927.92e^(−0.91x)
8
6.62
5.25
3.88
2.5
present
reduced
absent
loge(Bit Score)
0.00
0.25
0.50
0.75
1.00
Distance from M. lignano
Bristle Status
present
reduced
absent

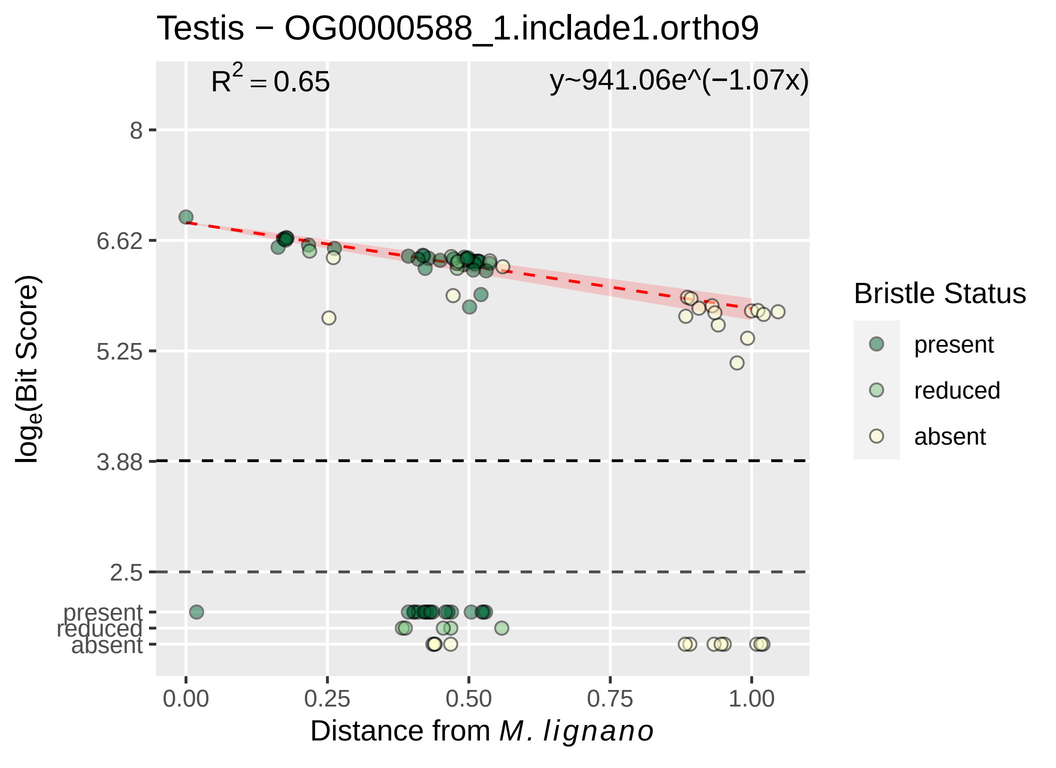

Testis − OG0000588_1.inclade1.ortho9
$R^2 = 0.65$
y~941.06e^(−1.07x)
8
6.62
5.25
3.88
2.5
present
reduced
absent
$\log_e$(Bit Score)
0.00
0.25
0.50
0.75
1.00
Distance from *M. lignano*
Bristle Status
present
reduced
absent

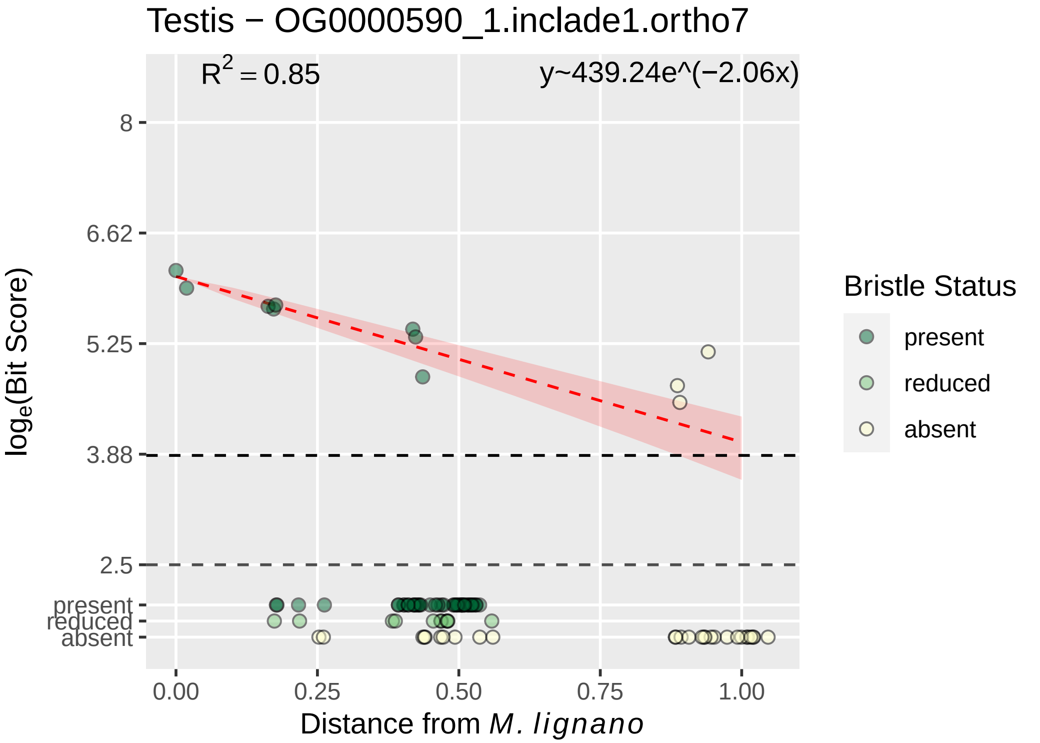

Testis − OG0000590_1.inclade1.ortho7
R^2 = 0.85
y~439.24e^(−2.06x)
8
6.62
5.25
3.88
2.5
present
reduced
absent
log_e(Bit Score)
0.00
0.25
0.50
0.75
1.00
Distance from M. lignano
Bristle Status
present
reduced
absent

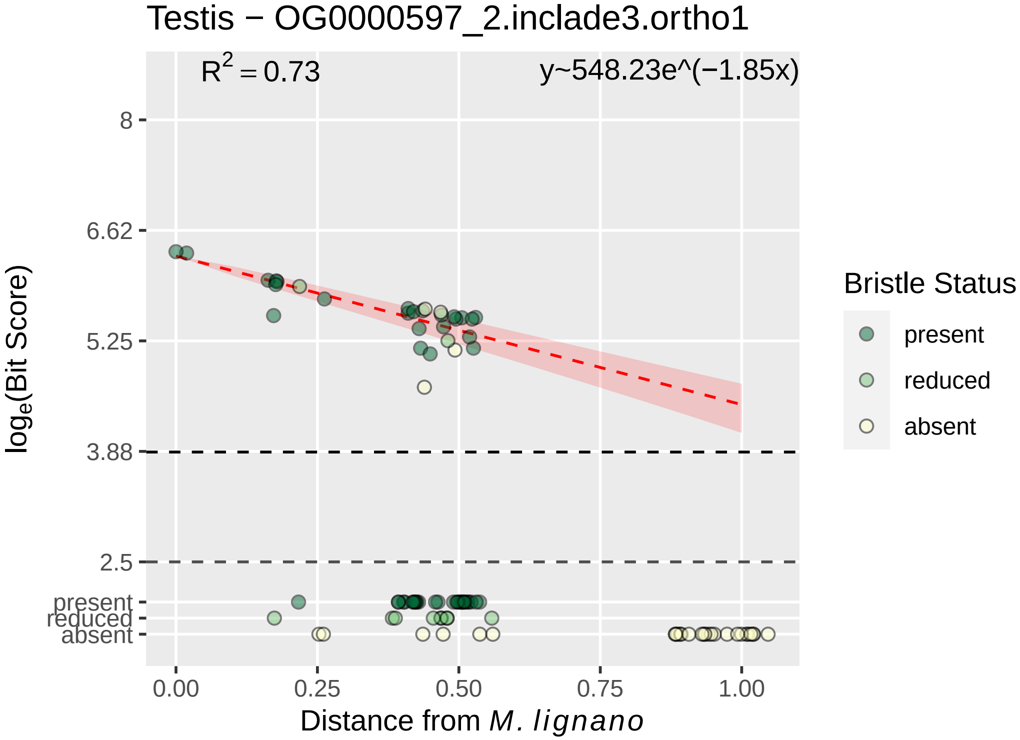

Testis − OG0000597_2.inclade3.ortho1
R² = 0.73
y~548.23e^(−1.85x)
8
6.62
5.25
3.88
2.5
present
reduced
absent
log_e(Bit Score)
0.00
0.25
0.50
0.75
1.00
Distance from M. lignano
Bristle Status
present
reduced
absent

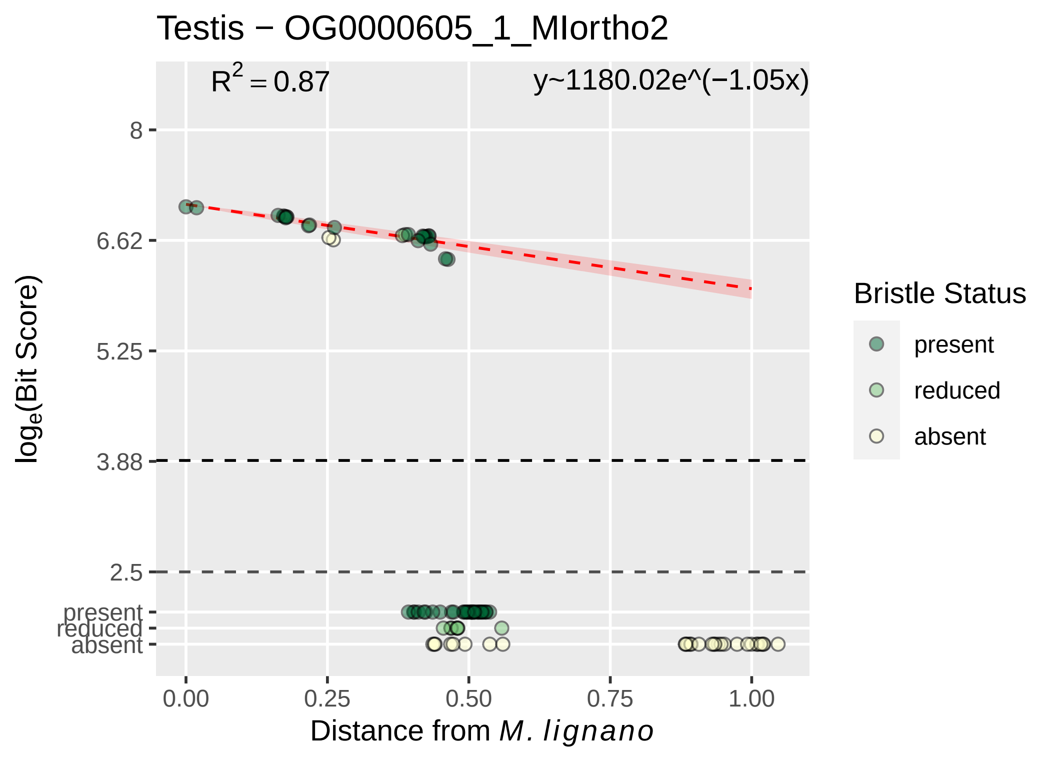
Testis − OG0000605_1_MIortho2
R^2 = 0.87
y~1180.02e^(−1.05x)
8
6.62
5.25
3.88
2.5
present
reduced
absent
0.00
0.25
0.50
0.75
1.00
Distance from M. lignano
loge(Bit Score)
Bristle Status
present
reduced
absent

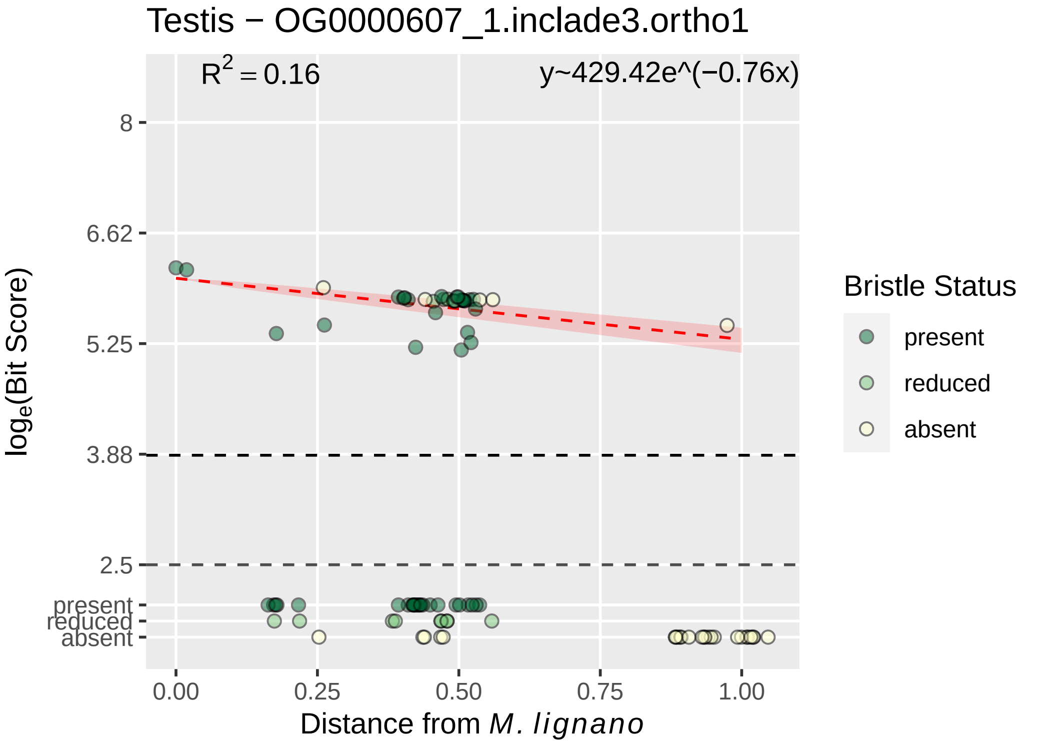
Testis − OG0000607_1.inclade3.ortho1
$R^2 = 0.16$
y~429.42e^(−0.76x)
8
6.62
5.25
3.88
2.5
present
reduced
absent
0.00
0.25
0.50
0.75
1.00
$\log_e$(Bit Score)
Distance from M. lignano
Bristle Status
present
reduced
absent

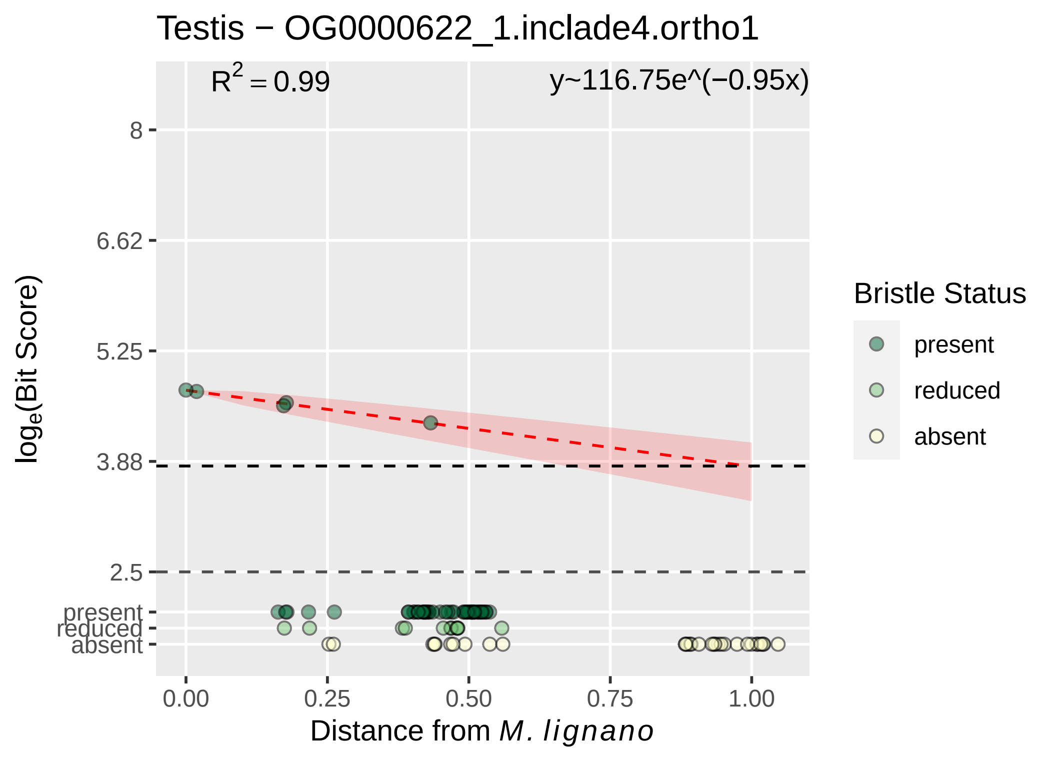

Testis − OG0000622_1.inclade4.ortho1
R² = 0.99
y~116.75e^(−0.95x)
8
6.62
5.25
3.88
2.5
present
reduced
absent
0.00
0.25
0.50
0.75
1.00
Distance from M. lignano
loge(Bit Score)
Bristle Status
present
reduced
absent

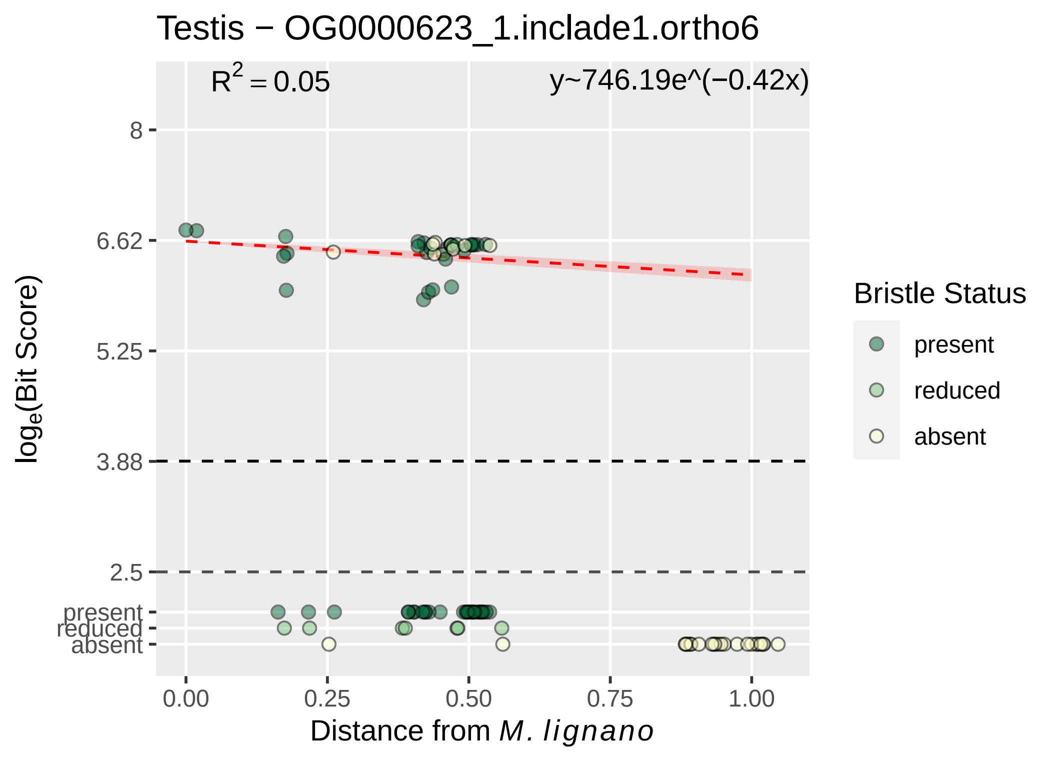

Testis − OG0000623_1.inclade1.ortho6
$R^2 = 0.05$
y~746.19e^(−0.42x)
8
6.62
5.25
3.88
2.5
present
reduced
absent
0.00
0.25
0.50
0.75
1.00
Distance from *M. lignano*
$\log_e$(Bit Score)
Bristle Status
present
reduced
absent

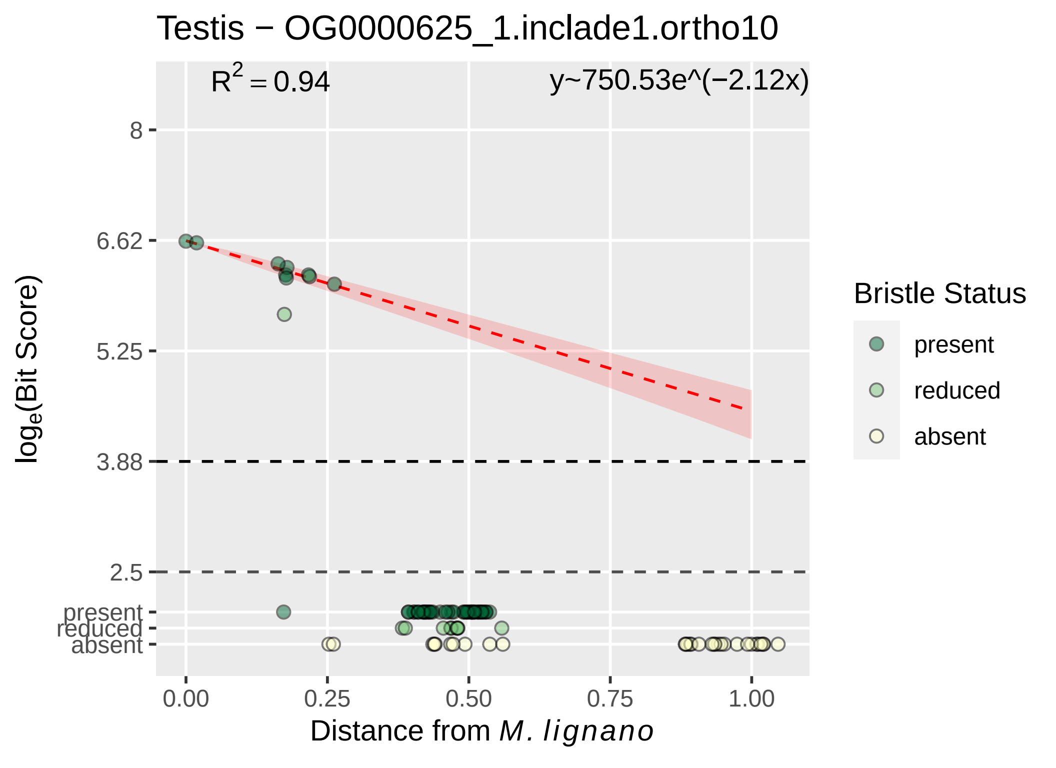

Testis − OG0000625_1.inclade1.ortho10
R² = 0.94
y~750.53e^(−2.12x)
8
6.62
5.25
3.88
2.5
present
reduced
absent
log_e(Bit Score)
0.00
0.25
0.50
0.75
1.00
Distance from M. lignano
Bristle Status
present
reduced
absent

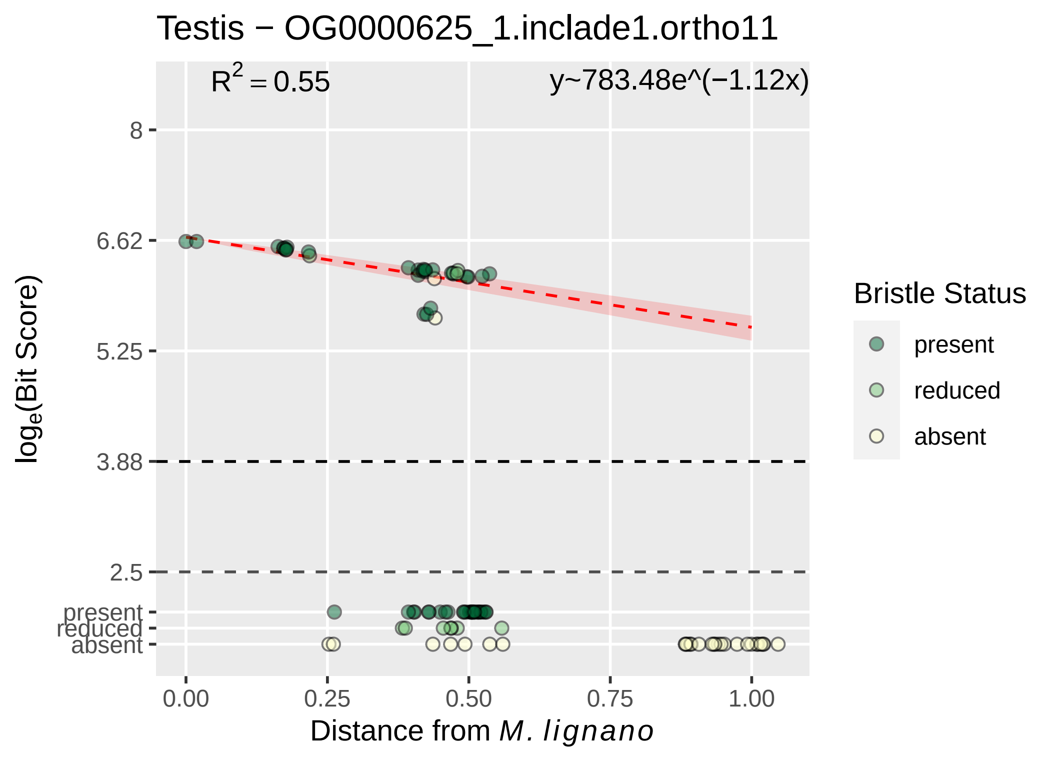

Testis − OG0000625_1.inclade1.ortho11
R^2 = 0.55
y~783.48e^(−1.12x)
8
6.62
5.25
3.88
2.5
present
reduced
absent
log_e(Bit Score)
0.00
0.25
0.50
0.75
1.00
Distance from M. lignano
Bristle Status
present
reduced
absent

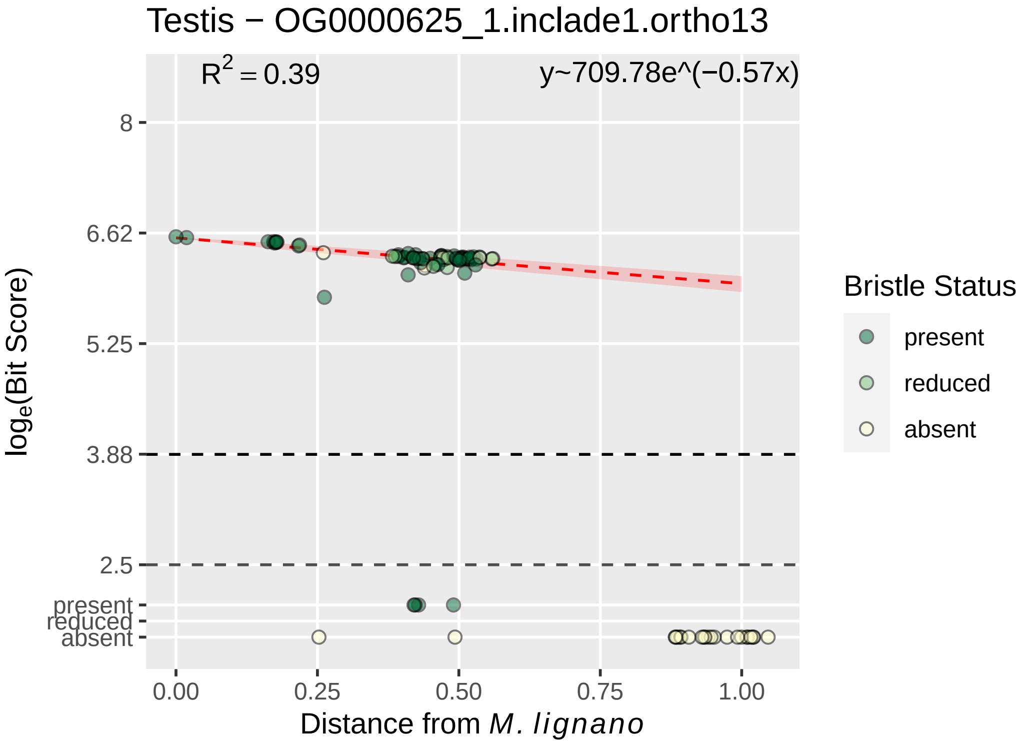

Testis − OG0000625_1.inclade1.ortho13
R^2 = 0.39
y~709.78e^(−0.57x)
8
6.62
5.25
3.88
2.5
present
reduced
absent
log_e(Bit Score)
0.00
0.25
0.50
0.75
1.00
Distance from M. lignano
Bristle Status
present
reduced
absent

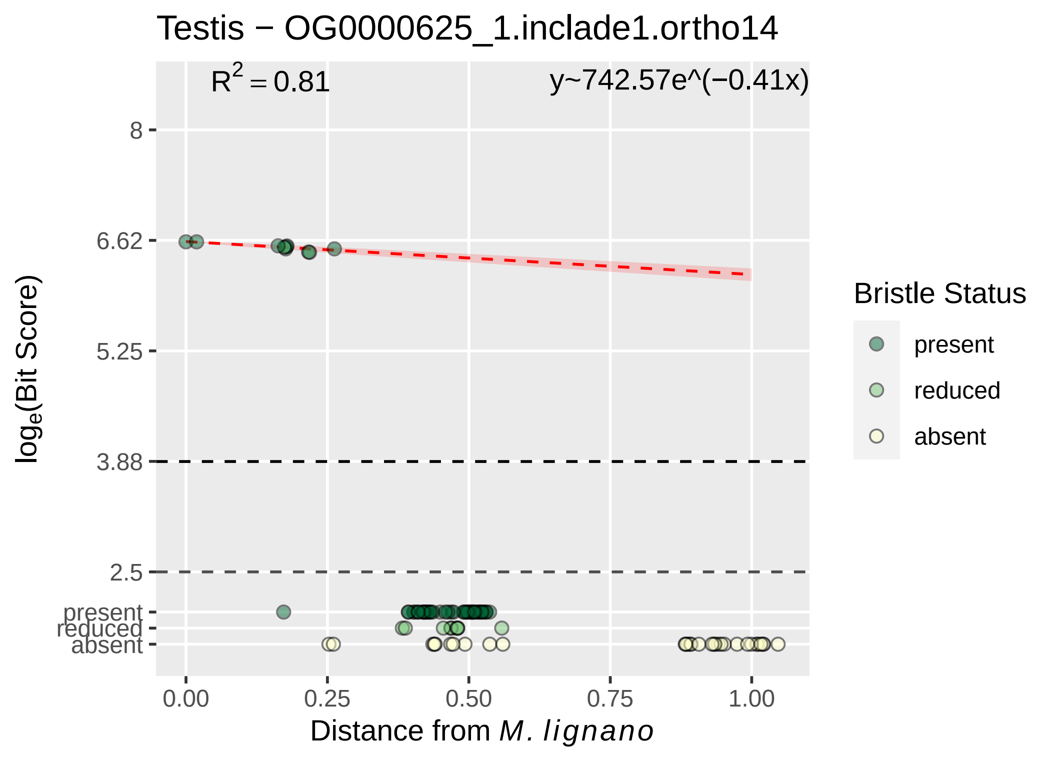

Testis − OG0000625_1.inclade1.ortho14
R^2 = 0.81
y~742.57e^(−0.41x)
8
6.62
5.25
3.88
2.5
present
reduced
absent
loge(Bit Score)
0.00
0.25
0.50
0.75
1.00
Distance from M. lignano
Bristle Status
present
reduced
absent

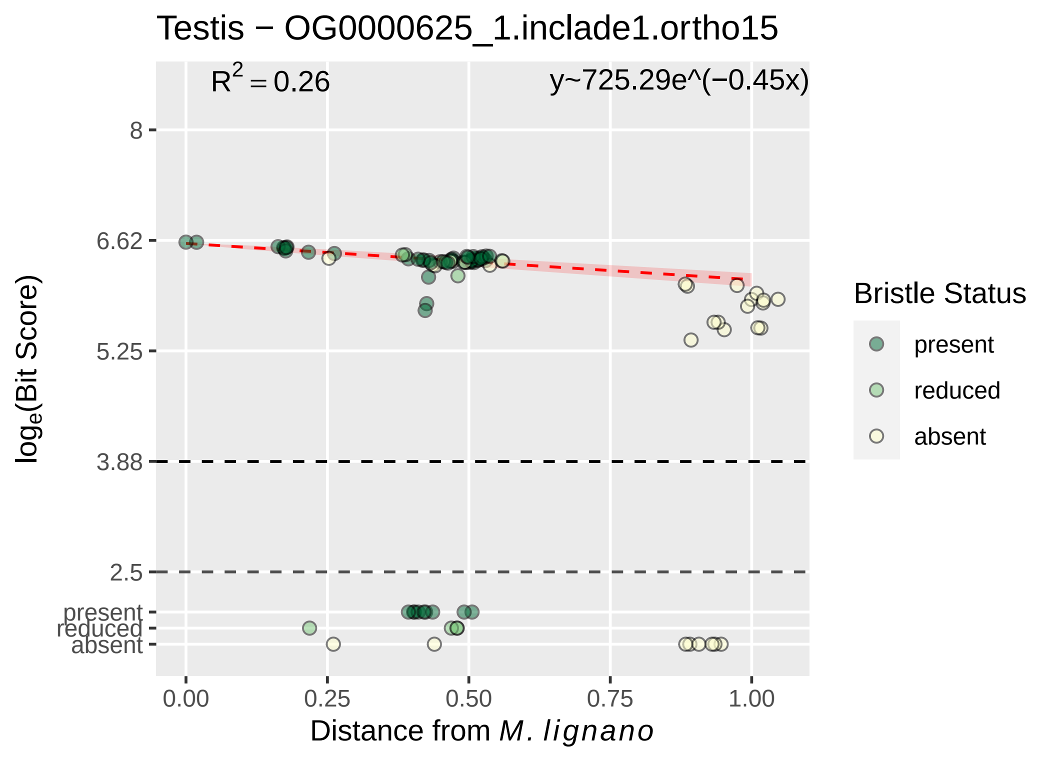

Testis − OG0000625_1.inclade1.ortho15
$R^2 = 0.26$
y~725.29e^(−0.45x)
8
6.62
5.25
3.88
2.5
present
reduced
absent
0.00
0.25
0.50
0.75
1.00
Distance from M. lignano
$\log_e$(Bit Score)
Bristle Status
present
reduced
absent

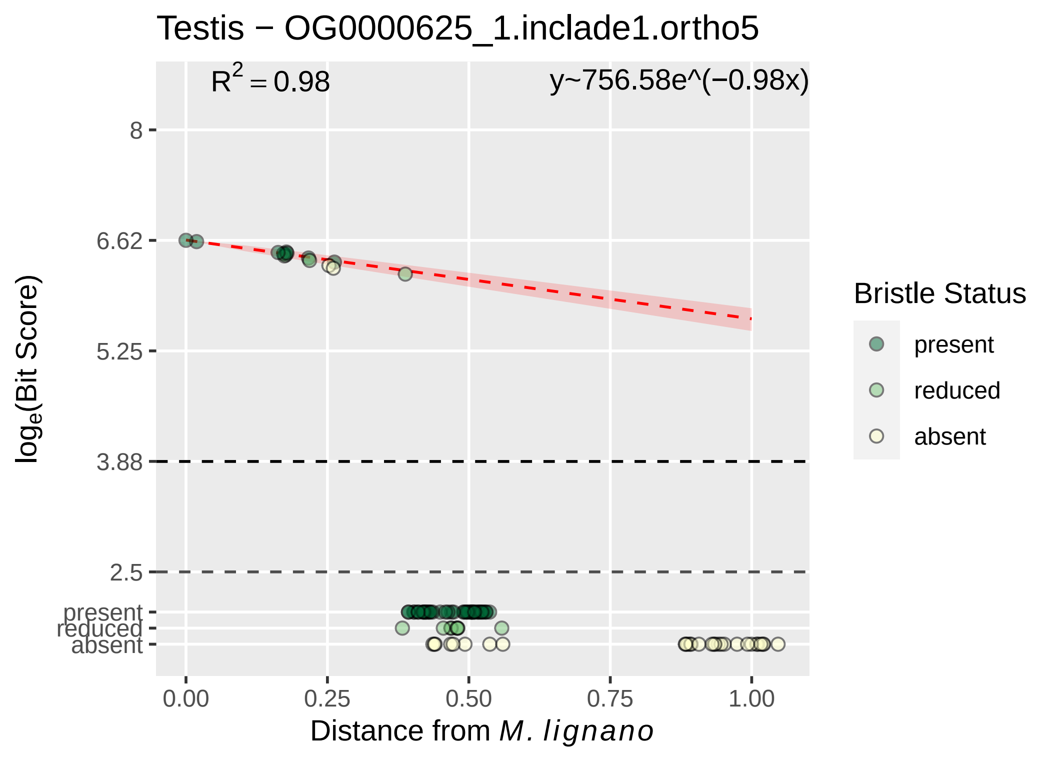

Testis − OG0000625_1.inclade1.ortho5
$R^2 = 0.98$
y~756.58e^(−0.98x)
8
6.62
5.25
3.88
2.5
present
reduced
absent
$\log_e$(Bit Score)
0.00
0.25
0.50
0.75
1.00
Distance from *M. lignano*
Bristle Status
present
reduced
absent

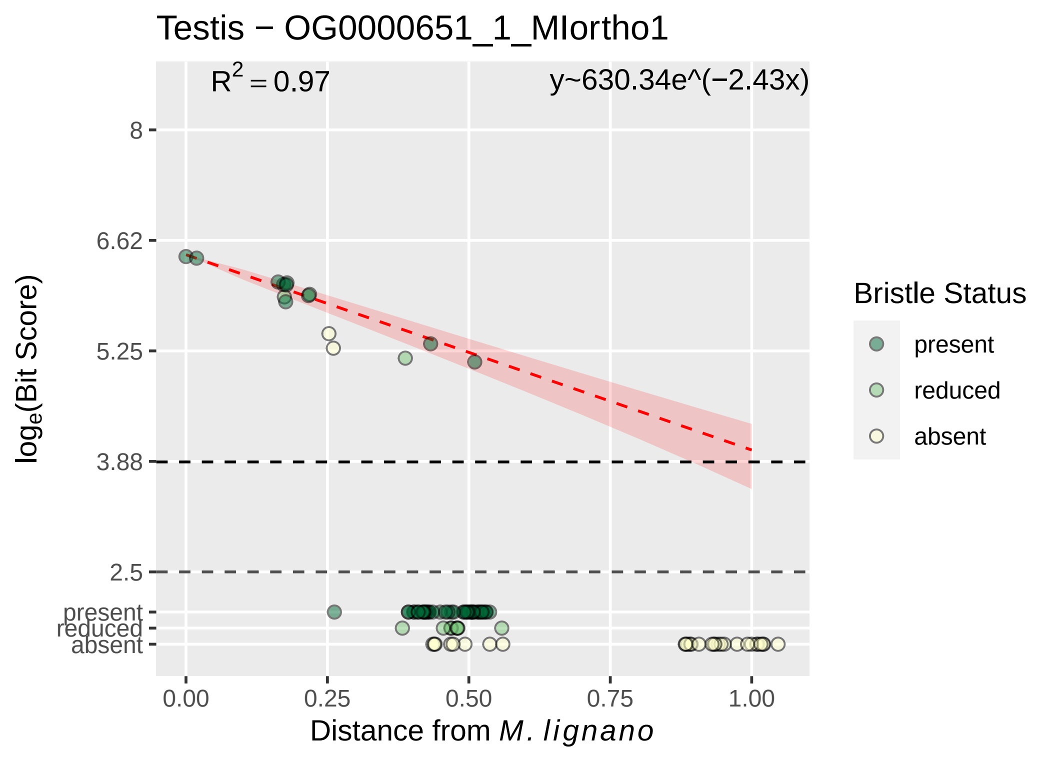
Testis − OG0000651_1_MIortho1
$R^2 = 0.97$
y~630.34e^(−2.43x)
8
6.62
5.25
3.88
2.5
present
reduced
absent
$\log_e$(Bit Score)
0.00
0.25
0.50
0.75
1.00
Distance from *M. lignano*
Bristle Status
present
reduced
absent

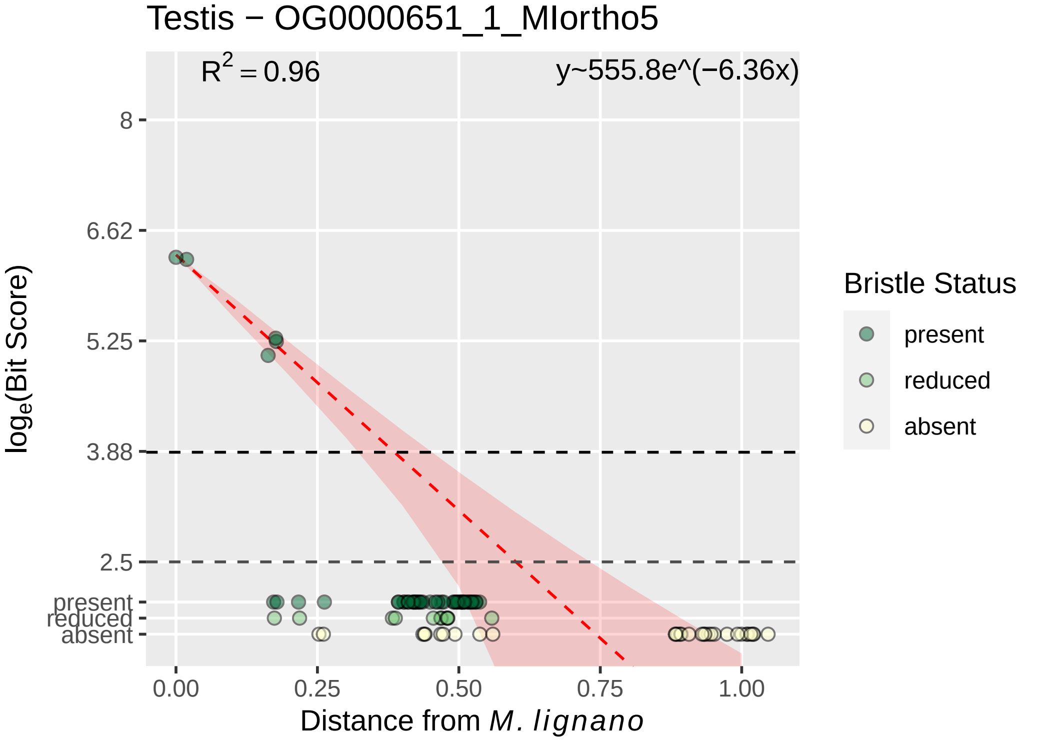
Testis − OG0000651_1_MIortho5
R^2 = 0.96
y~555.8e^(−6.36x)
8
6.62
5.25
3.88
2.5
present
reduced
absent
log_e(Bit Score)
0.00
0.25
0.50
0.75
1.00
Distance from M. lignano
Bristle Status
present
reduced
absent

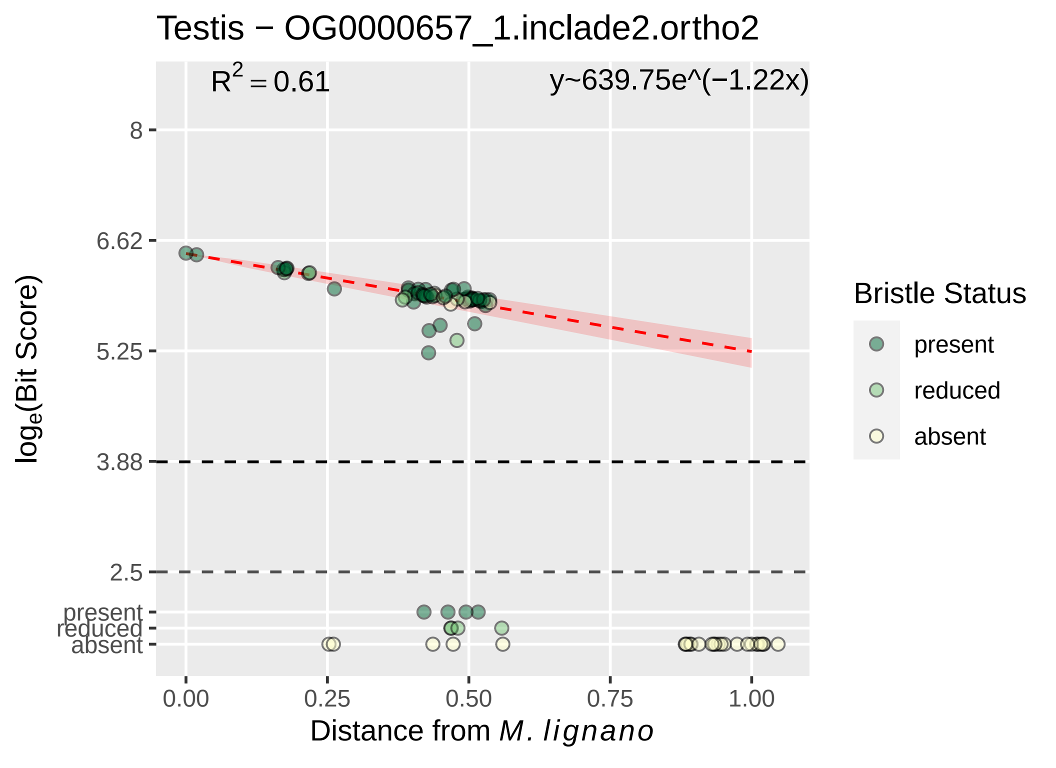

Testis − OG0000657_1.inclade2.ortho2
$R^2 = 0.61$
y~639.75e^(−1.22x)
8
6.62
5.25
3.88
2.5
present
reduced
absent
0.00
0.25
0.50
0.75
1.00
Distance from M. lignano
log$_e$(Bit Score)
Bristle Status
present
reduced
absent

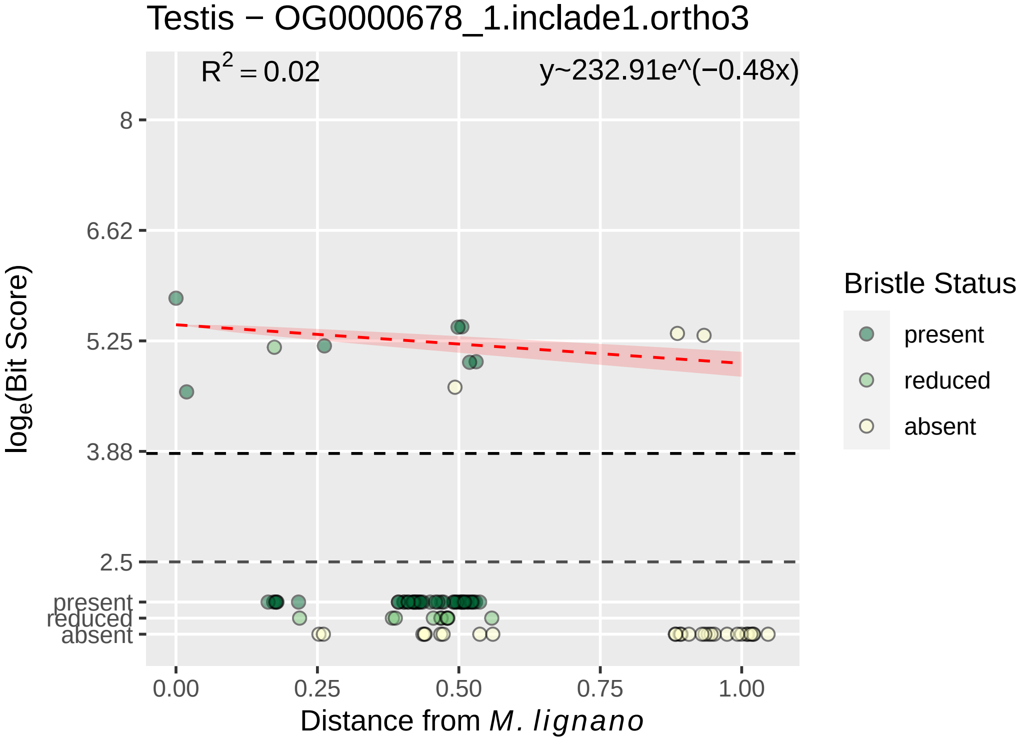

Testis − OG0000678_1.inclade1.ortho3
$R^2 = 0.02$
y~232.91e^(−0.48x)
8
6.62
5.25
3.88
2.5
present
reduced
absent
log_e(Bit Score)
0.00
0.25
0.50
0.75
1.00
Distance from M. lignano
Bristle Status
present
reduced
absent

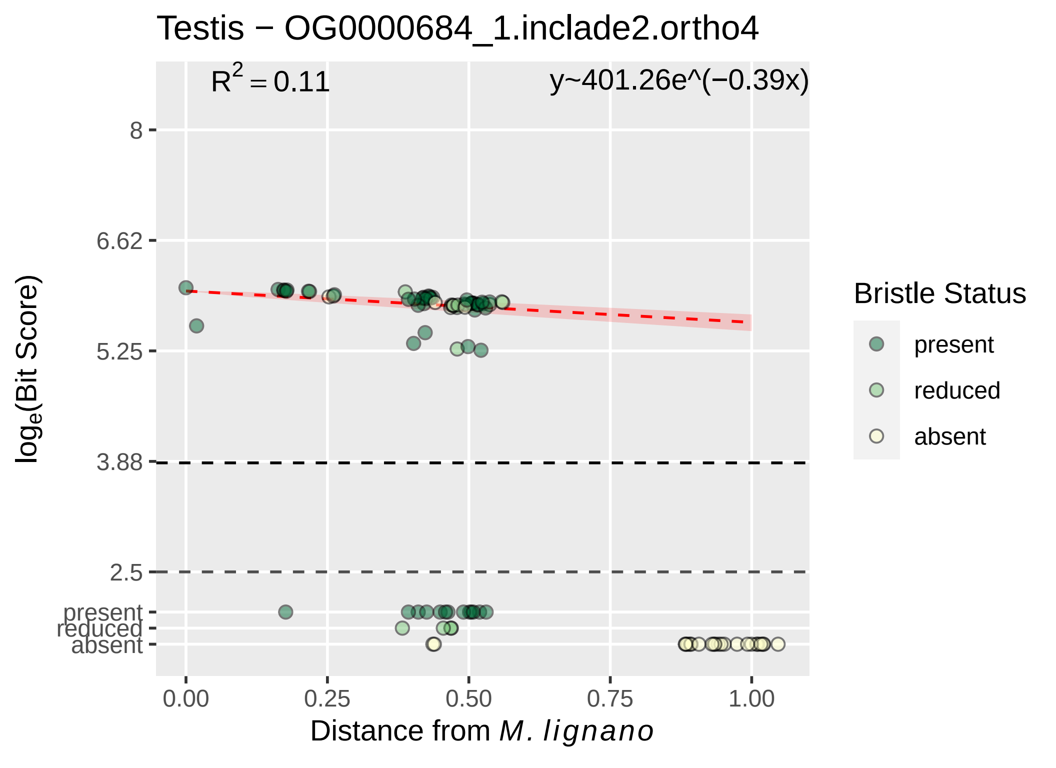

Testis − OG0000684_1.inclade2.ortho4
$R^2 = 0.11$
y~401.26e^(−0.39x)
8
6.62
5.25
3.88
2.5
present
reduced
absent
0.00
0.25
0.50
0.75
1.00
log$_e$(Bit Score)
Distance from *M. lignano*
Bristle Status
present
reduced
absent

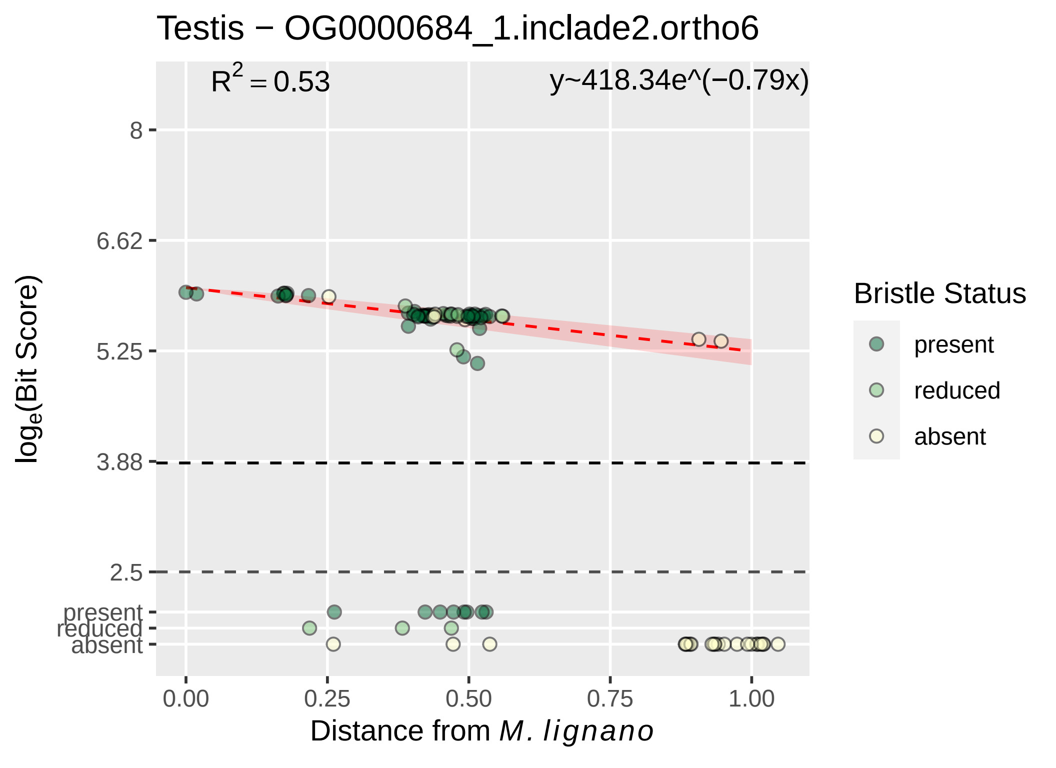
Testis − OG0000684_1.inclade2.ortho6
$R^2 = 0.53$
y~418.34e^(−0.79x)
8
6.62
5.25
3.88
2.5
present
reduced
absent
0.00
0.25
0.50
0.75
1.00
log$_e$(Bit Score)
Distance from M. lignano
Bristle Status
present
reduced
absent

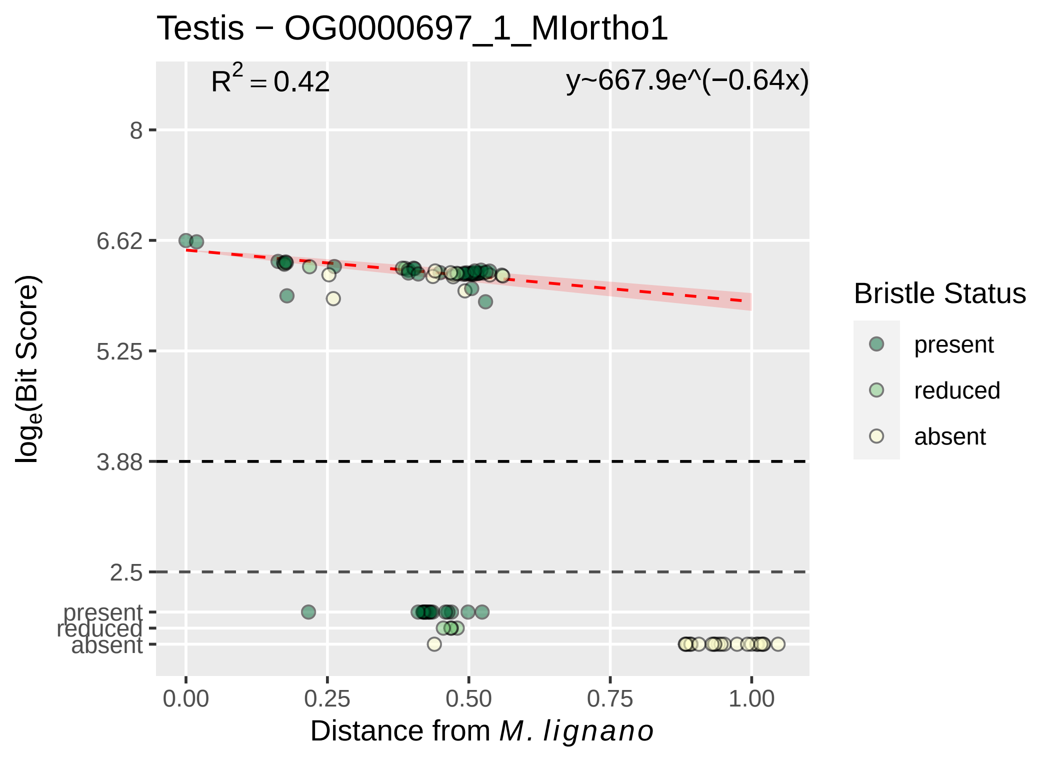

Testis − OG0000697_1_MIortho1
$R^2 = 0.42$
y~667.9e^(−0.64x)
8
6.62
5.25
3.88
2.5
present
reduced
absent
0.00
0.25
0.50
0.75
1.00
Distance from *M. lignano*
$\log_e$(Bit Score)
Bristle Status
present
reduced
absent

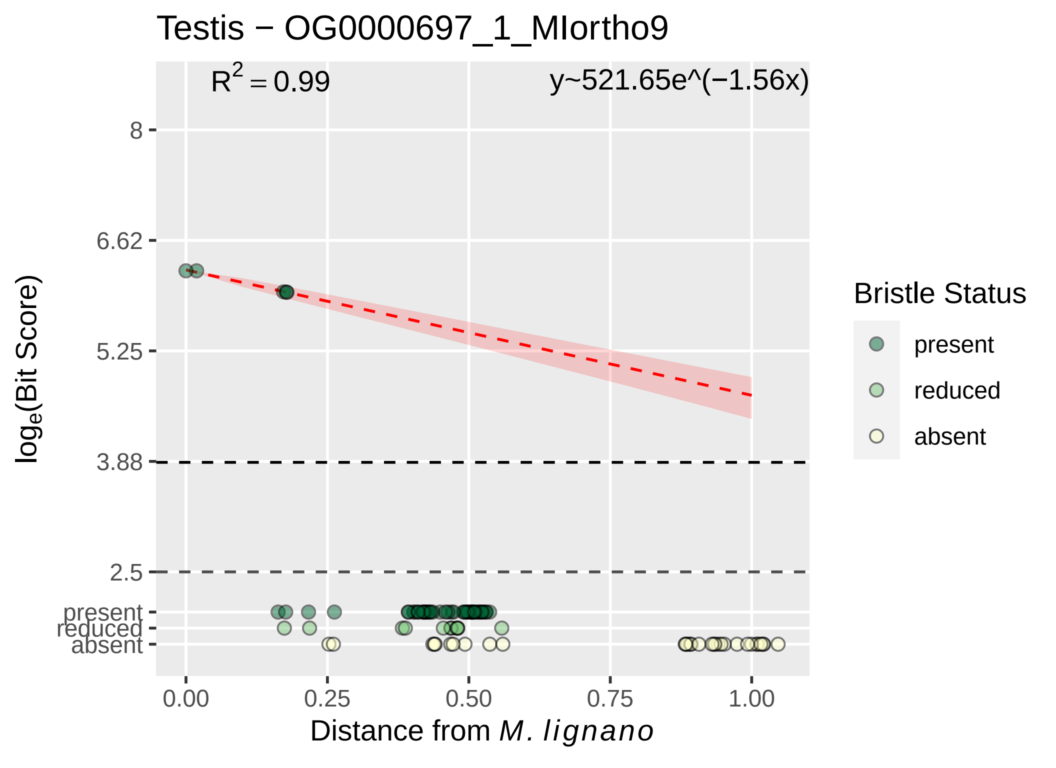

Testis − OG0000697_1_MIortho9
R2 = 0.99
y~521.65e^(−1.56x)
8
6.62
5.25
3.88
2.5
present
reduced
absent
loge(Bit Score)
0.00
0.25
0.50
0.75
1.00
Distance from M. lignano
Bristle Status
present
reduced
absent

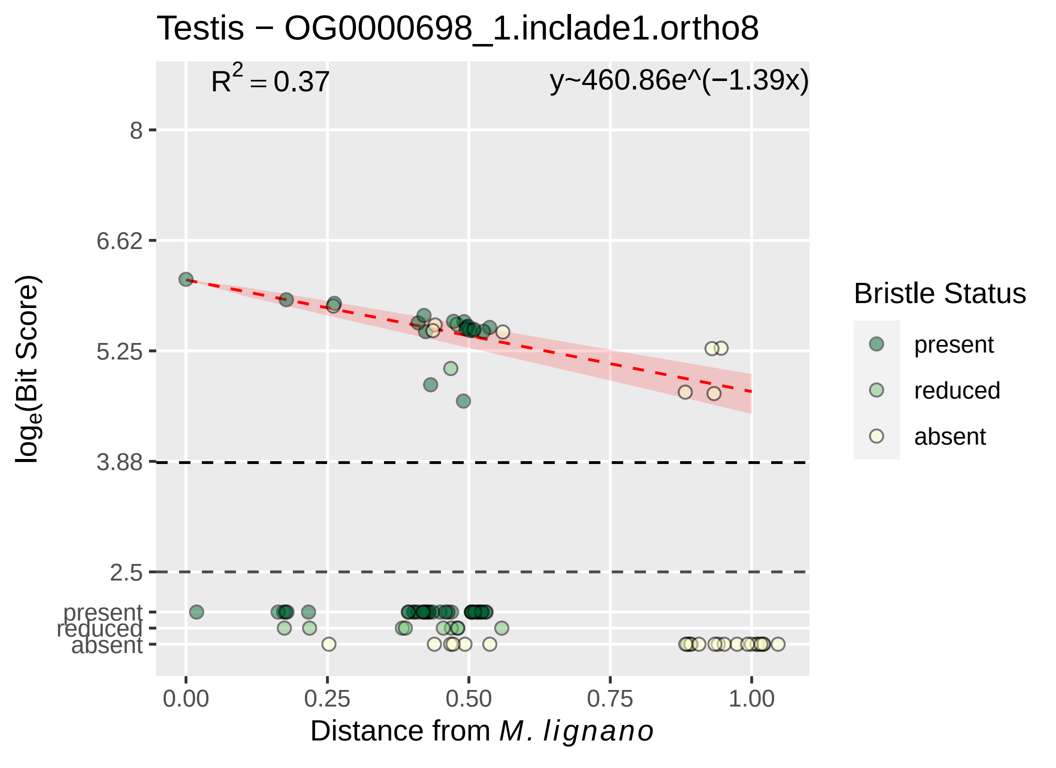

Testis − OG0000698_1.inclade1.ortho8
R^2 = 0.37
y~460.86e^(−1.39x)
8
6.62
5.25
3.88
2.5
present
reduced
absent
log_e(Bit Score)
0.00
0.25
0.50
0.75
1.00
Distance from M. lignano
Bristle Status
present
reduced
absent

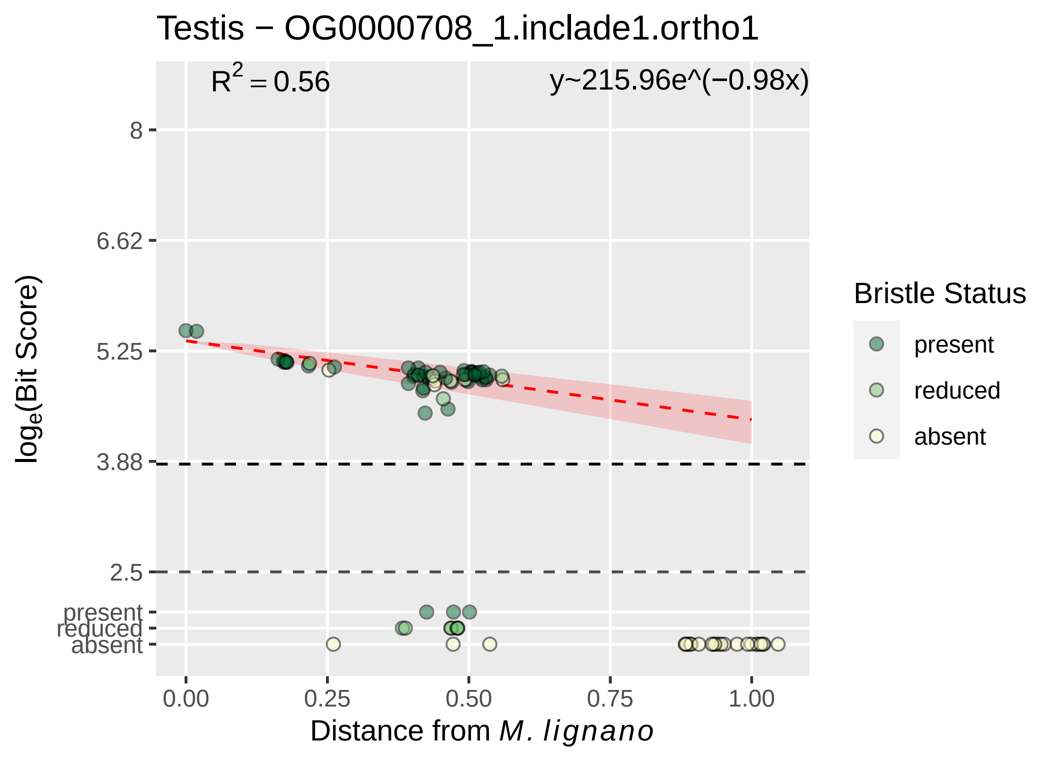

Testis − OG0000708_1.inclade1.ortho1
R^2 = 0.56
y~215.96e^(−0.98x)
8
6.62
5.25
3.88
2.5
present
reduced
absent
log_e(Bit Score)
0.00
0.25
0.50
0.75
1.00
Distance from M. lignano
Bristle Status
present
reduced
absent

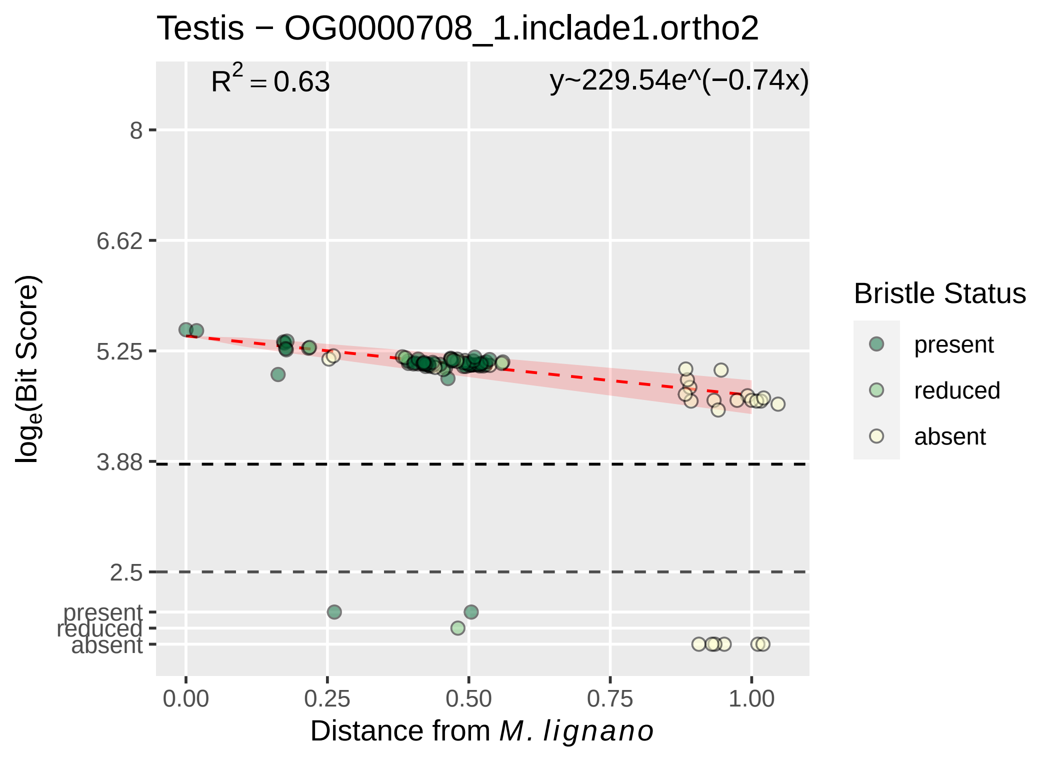
Testis − OG0000708_1.inclade1.ortho2
$R^2 = 0.63$
y~229.54e^(−0.74x)
8
6.62
5.25
3.88
2.5
present
reduced
absent
$\log_e$(Bit Score)
0.00
0.25
0.50
0.75
1.00
Distance from *M. lignano*
Bristle Status
present
reduced
absent

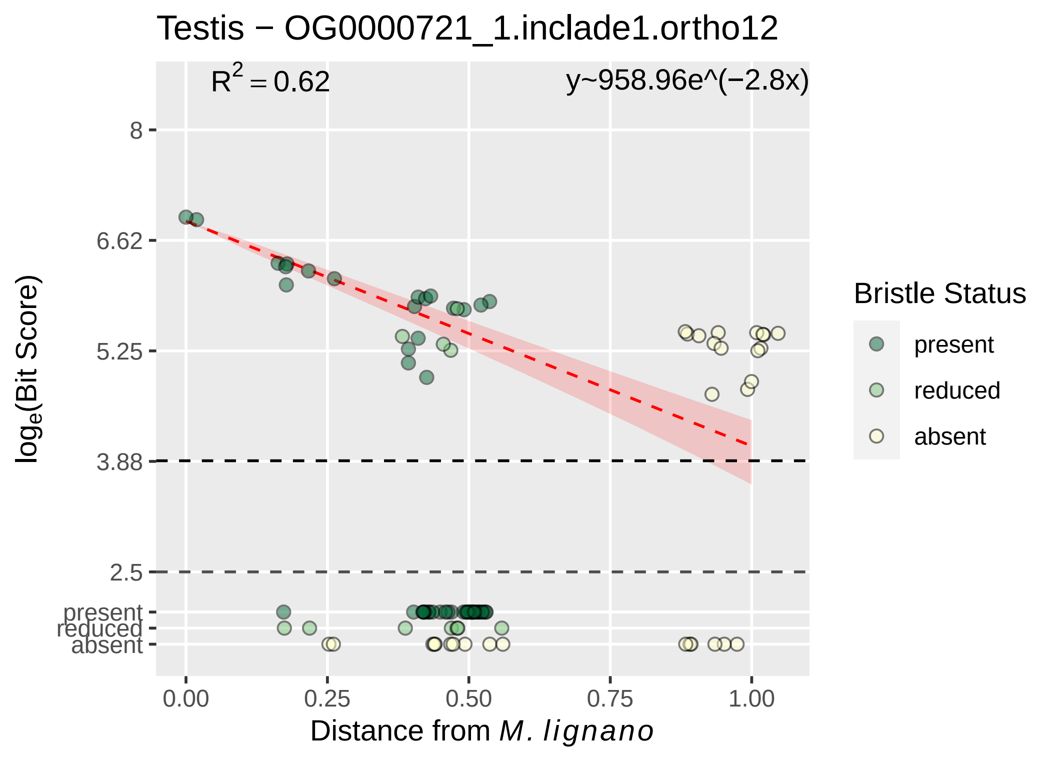

Testis − OG0000721_1.inclade1.ortho12
R² = 0.62
y~958.96e^(−2.8x)
8
6.62
5.25
3.88
2.5
present
reduced
absent
log_e(Bit Score)
0.00
0.25
0.50
0.75
1.00
Distance from M. lignano
Bristle Status
present
reduced
absent

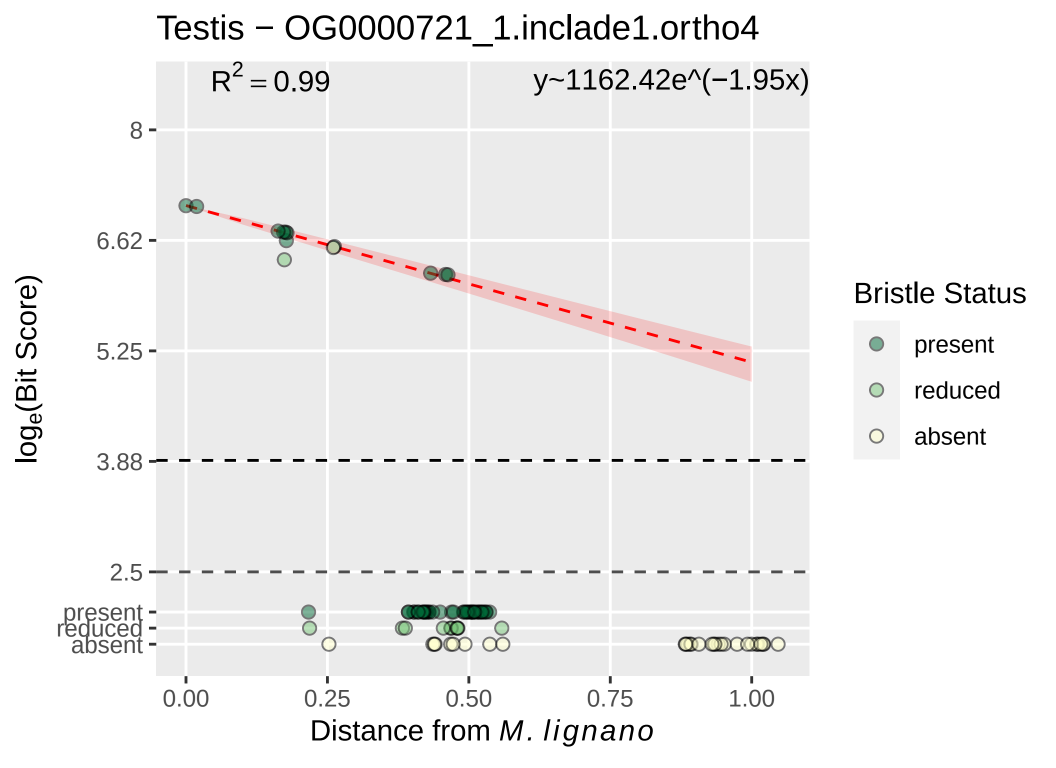

Testis − OG0000721_1.inclade1.ortho4
R2 = 0.99
y~1162.42e^(−1.95x)
8
6.62
5.25
3.88
2.5
present
reduced
absent
loge(Bit Score)
0.00
0.25
0.50
0.75
1.00
Distance from M. lignano
Bristle Status
present
reduced
absent

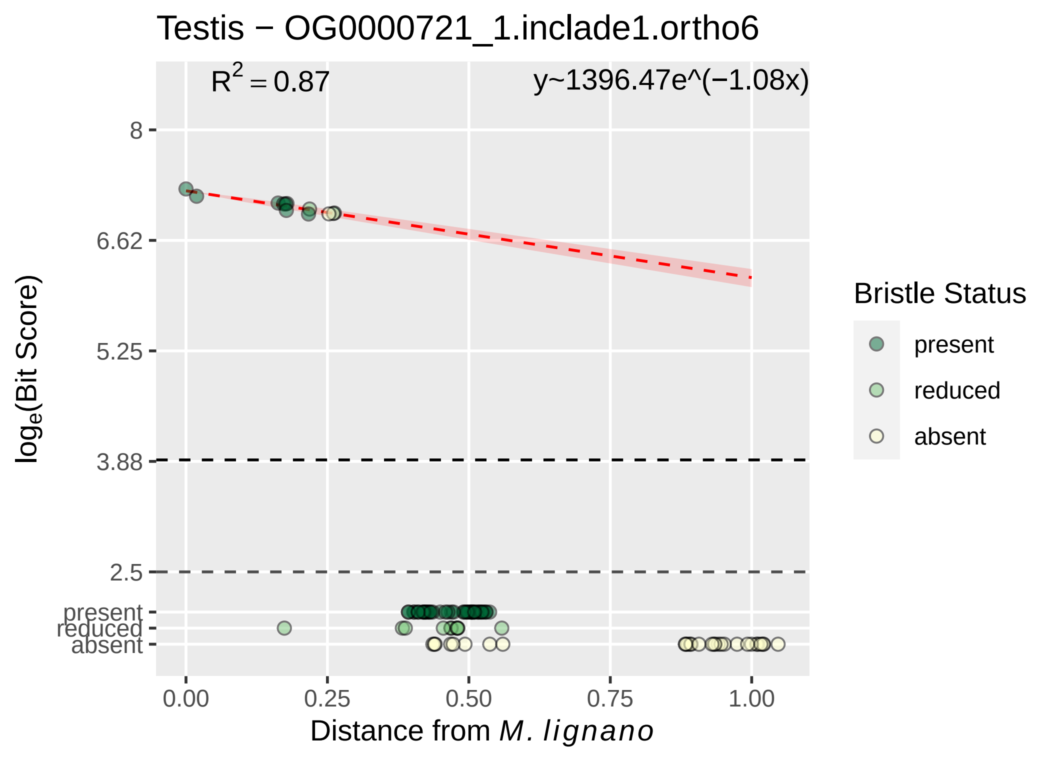
Testis − OG0000721_1.inclade1.ortho6
$R^2 = 0.87$
y~1396.47e^(−1.08x)
8
6.62
5.25
3.88
2.5
present
reduced
absent
$\log_e$(Bit Score)
0.00
0.25
0.50
0.75
1.00
Distance from *M. lignano*
Bristle Status
present
reduced
absent

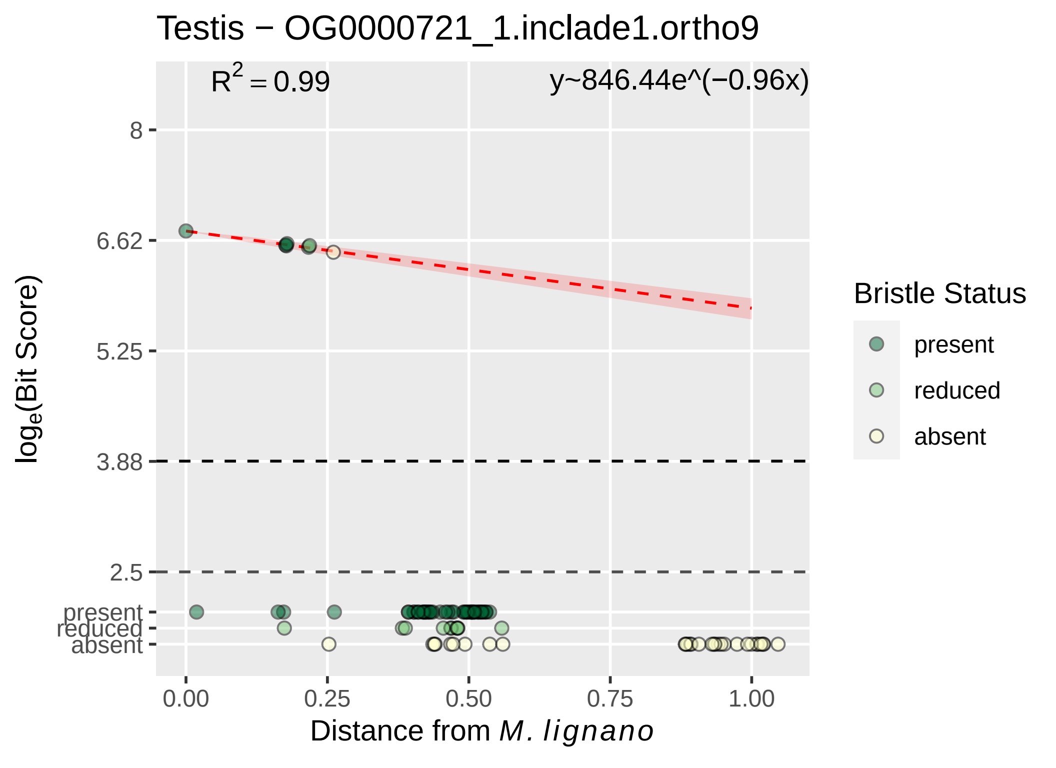

Testis − OG0000721_1.inclade1.ortho9
$R^2 = 0.99$
y~846.44e^(−0.96x)
8
6.62
5.25
3.88
2.5
present
reduced
absent
0.00
0.25
0.50
0.75
1.00
log$_e$(Bit Score)
Distance from M. lignano
Bristle Status
present
reduced
absent

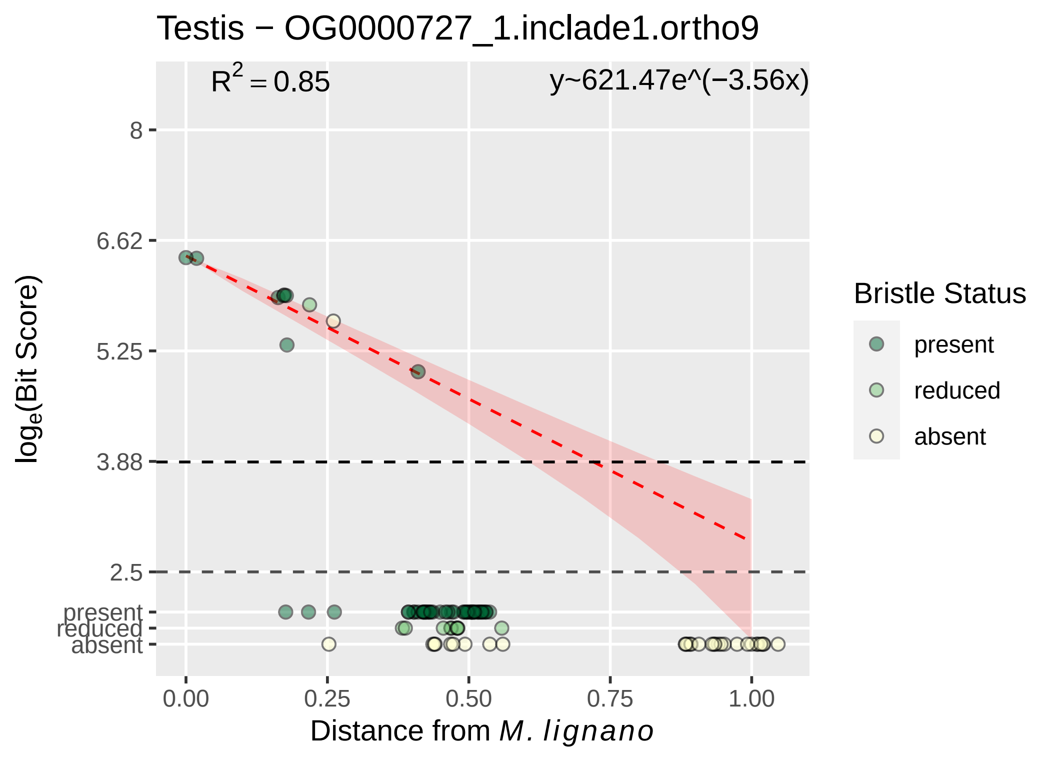

Testis − OG0000727_1.inclade1.ortho9
R^2 = 0.85
y~621.47e^(−3.56x)
8
6.62
5.25
3.88
2.5
present
reduced
absent
0.00
0.25
0.50
0.75
1.00
Distance from M. lignano
log_e(Bit Score)
Bristle Status
present
reduced
absent

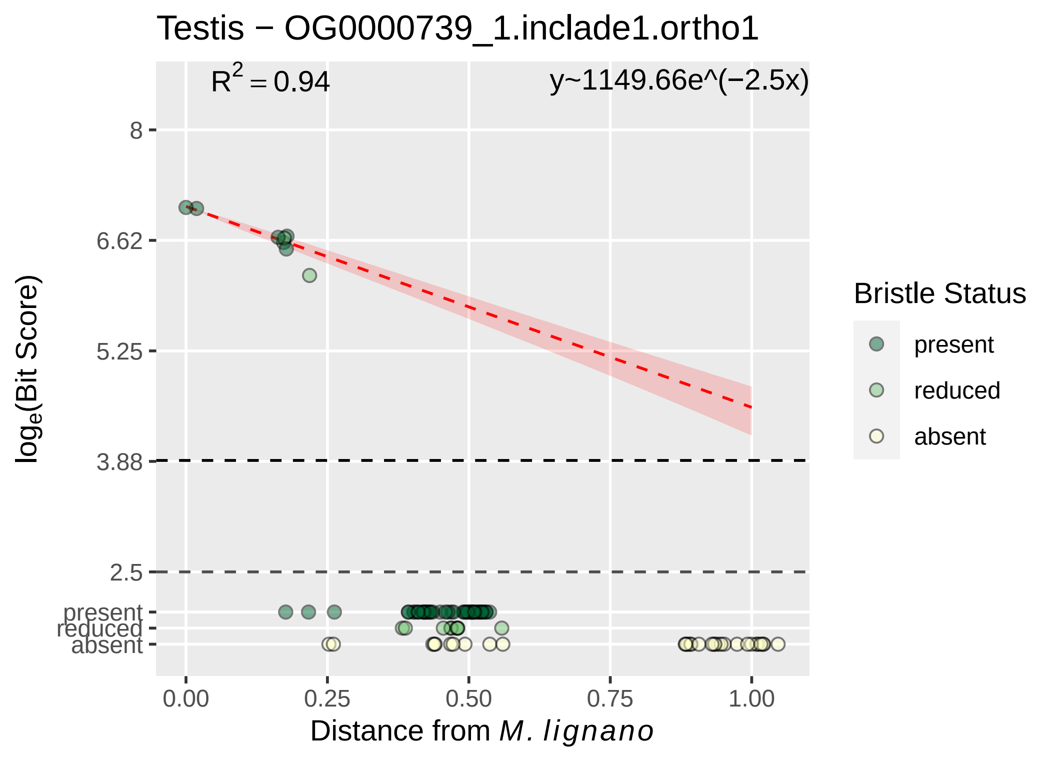

Testis − OG0000739_1.inclade1.ortho1
R2 = 0.94
y~1149.66e^(−2.5x)
8
6.62
5.25
3.88
2.5
present
reduced
absent
0.00
0.25
0.50
0.75
1.00
loge(Bit Score)
Distance from M. lignano
Bristle Status
present
reduced
absent

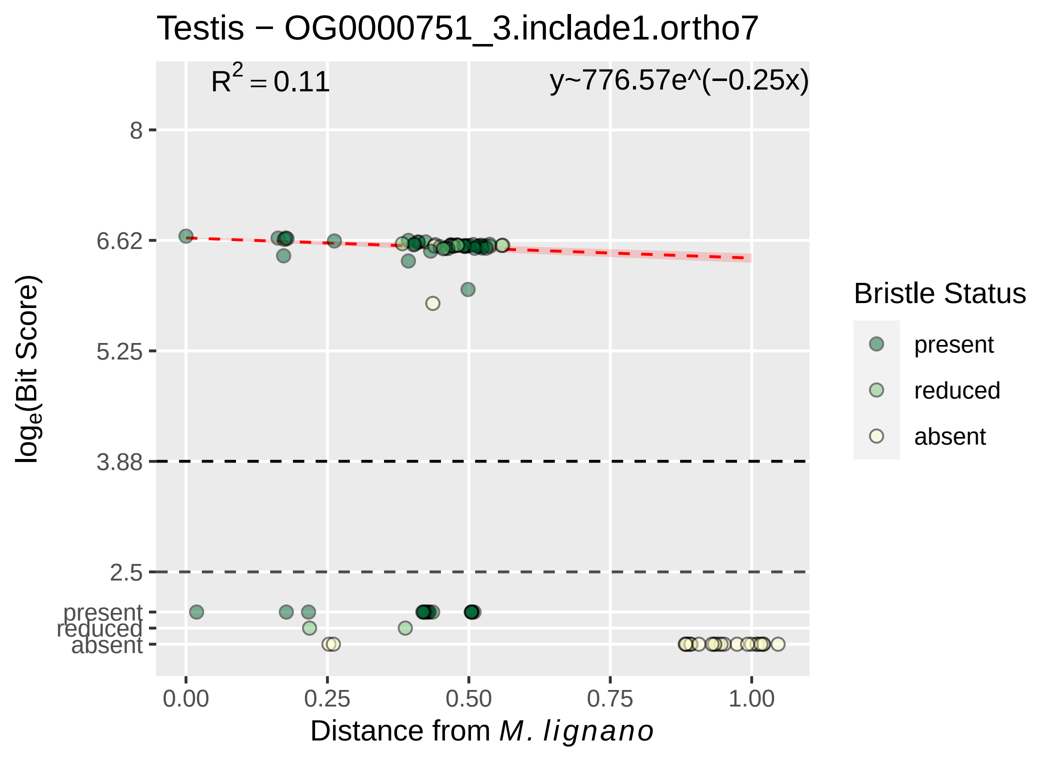
Testis − OG0000751_3.inclade1.ortho7
$R^2 = 0.11$
y~776.57e^(−0.25x)
8
6.62
5.25
3.88
2.5
present
reduced
absent
log$_e$(Bit Score)
0.00
0.25
0.50
0.75
1.00
Distance from *M. lignano*
Bristle Status
present
reduced
absent

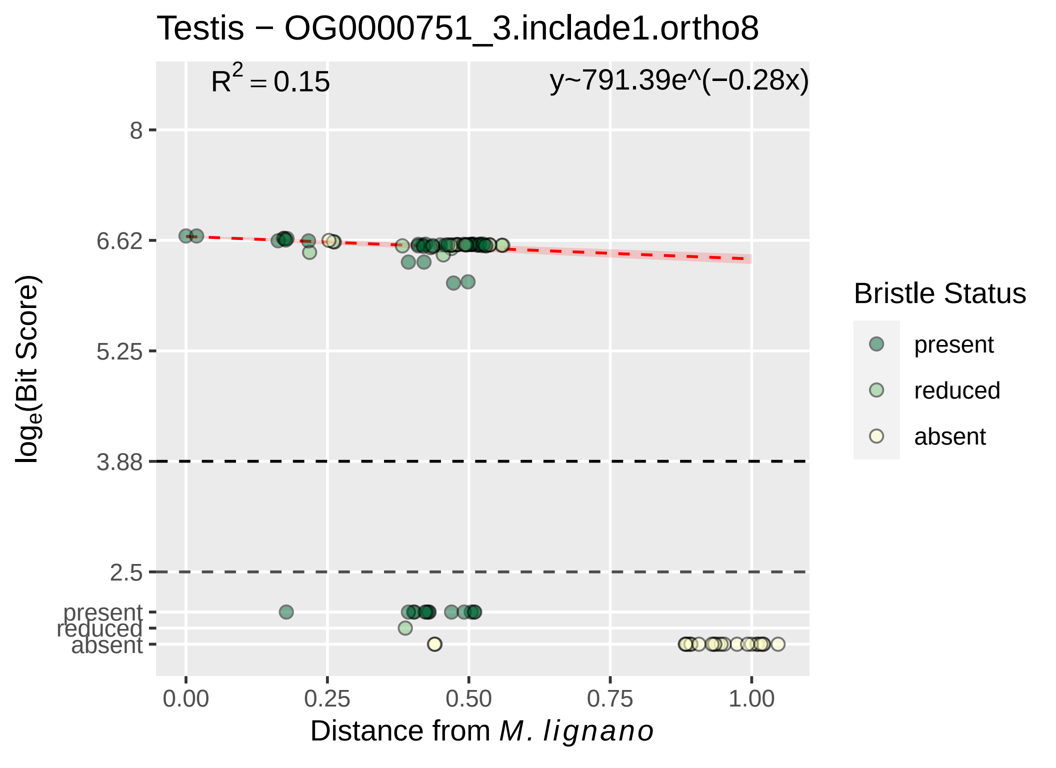

Testis − OG0000751_3.inclade1.ortho8
R^2 = 0.15
y~791.39e^(−0.28x)
8
6.62
5.25
3.88
2.5
present
reduced
absent
0.00
0.25
0.50
0.75
1.00
log_e(Bit Score)
Distance from M. lignano
Bristle Status
present
reduced
absent

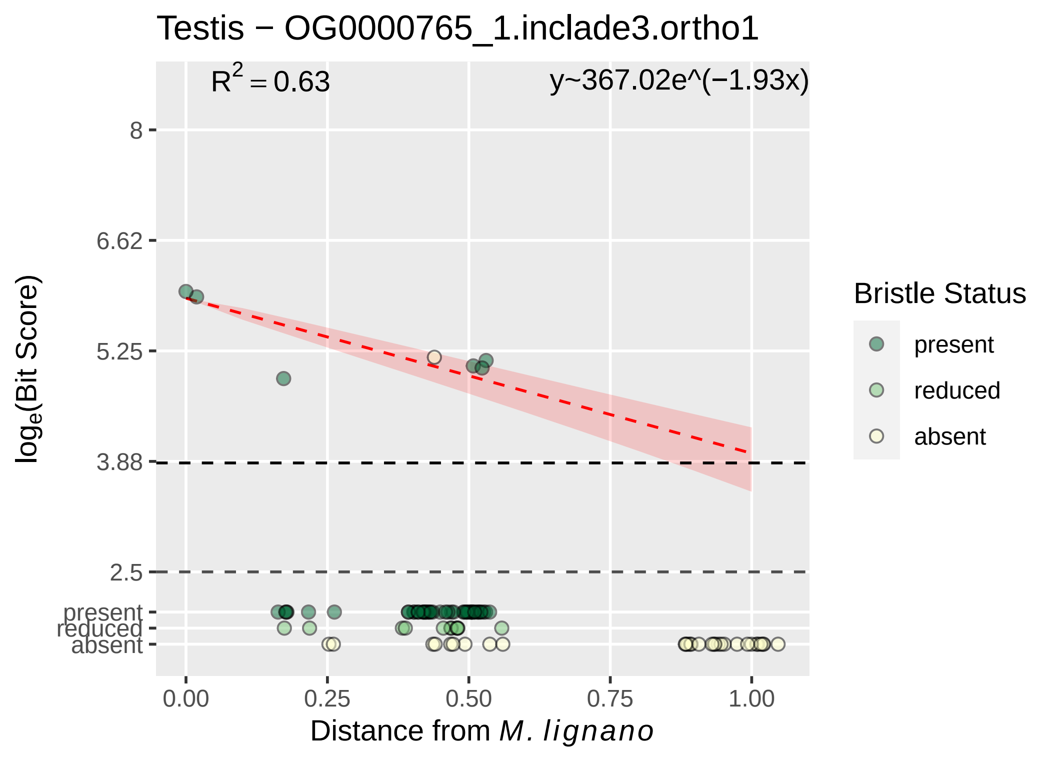
Testis − OG0000765_1.inclade3.ortho1
$R^2 = 0.63$
y~367.02e^(−1.93x)
8
6.62
5.25
3.88
2.5
present
reduced
absent
$log_e$(Bit Score)
0.00
0.25
0.50
0.75
1.00
Distance from *M. lignano*
Bristle Status
present
reduced
absent

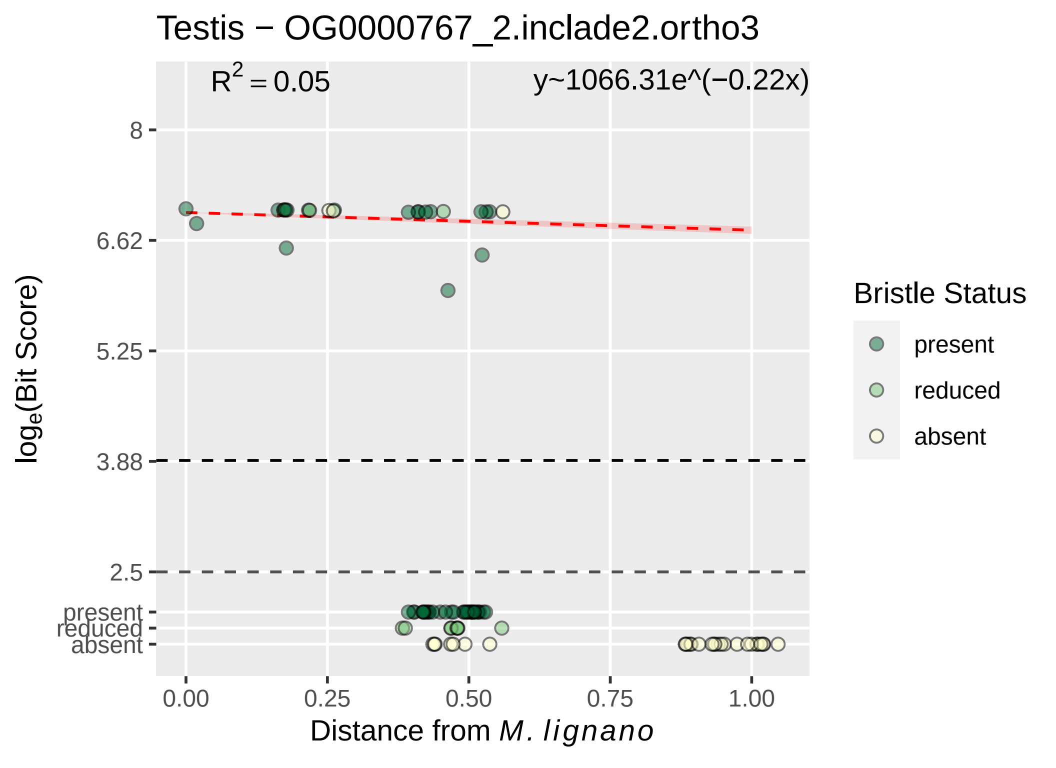

Testis − OG0000767_2.inclade2.ortho3
$R^2 = 0.05$
y~1066.31e^(−0.22x)
8
6.62
5.25
3.88
2.5
present
reduced
absent
$\log_e$(Bit Score)
0.00
0.25
0.50
0.75
1.00
Distance from *M. lignano*
Bristle Status
present
reduced
absent

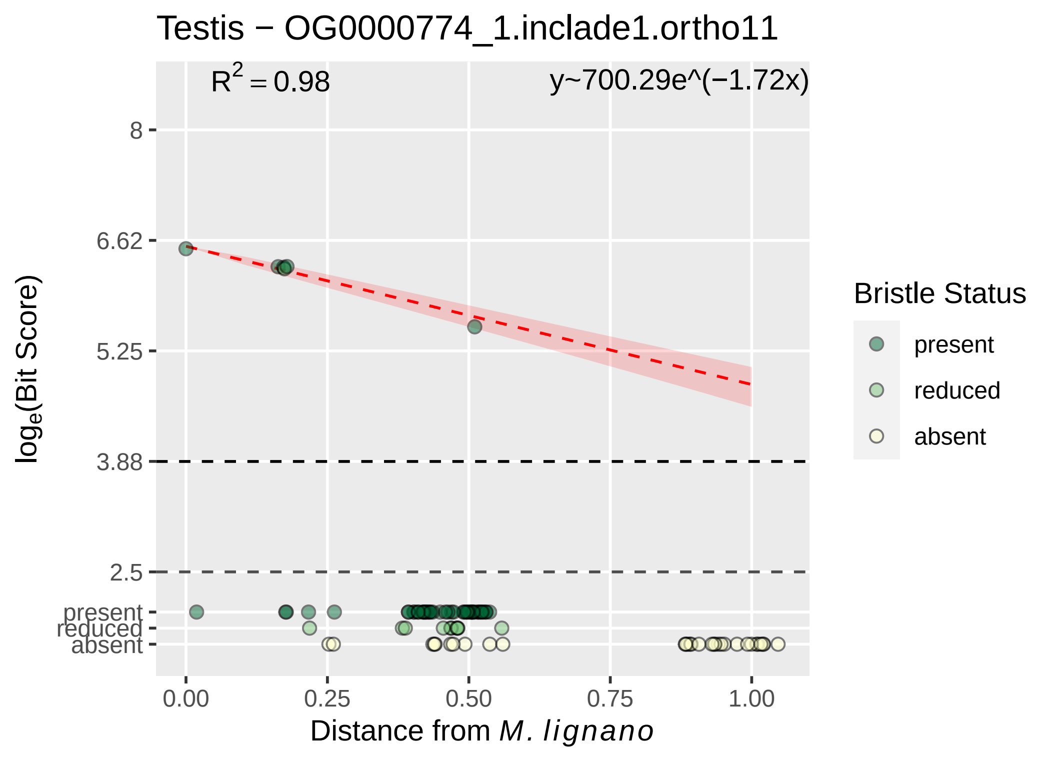

Testis − OG0000774_1.inclade1.ortho11
R^2 = 0.98
y~700.29e^(−1.72x)
8
6.62
5.25
3.88
2.5
present
reduced
absent
log_e(Bit Score)
0.00
0.25
0.50
0.75
1.00
Distance from M. lignano
Bristle Status
present
reduced
absent

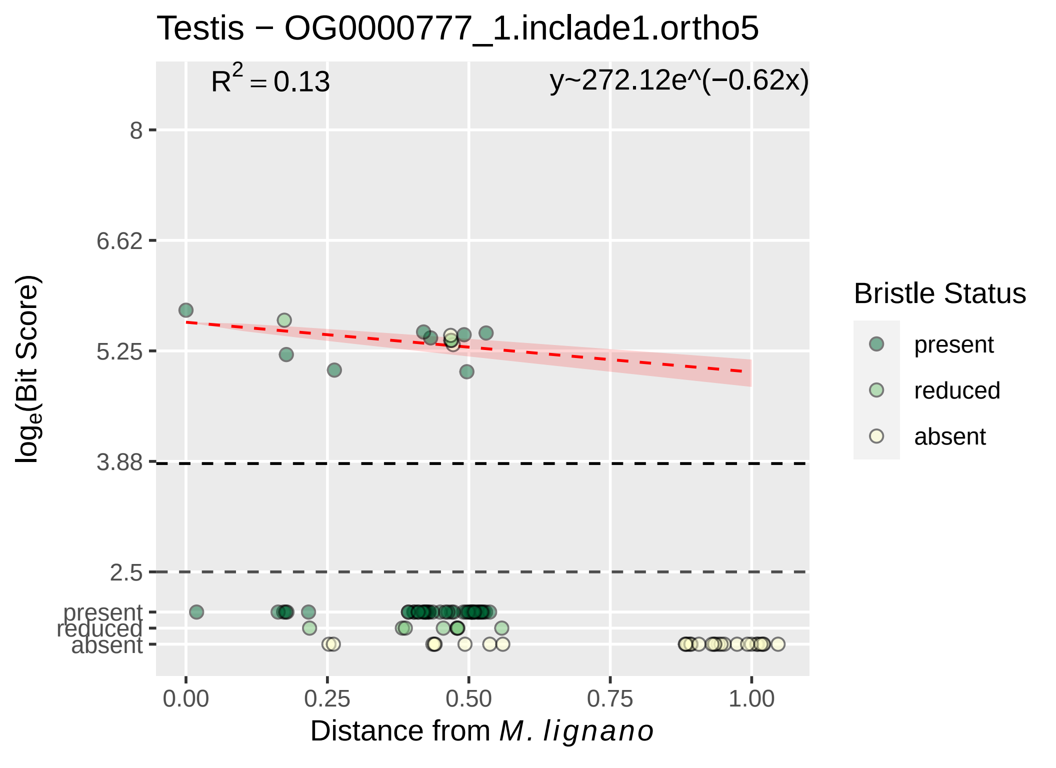
Testis − OG0000777_1.inclade1.ortho5
$R^2 = 0.13$
y~272.12e^(−0.62x)
8
6.62
5.25
3.88
2.5
present
reduced
absent
$log_e$(Bit Score)
0.00
0.25
0.50
0.75
1.00
Distance from *M. lignano*
Bristle Status
present
reduced
absent

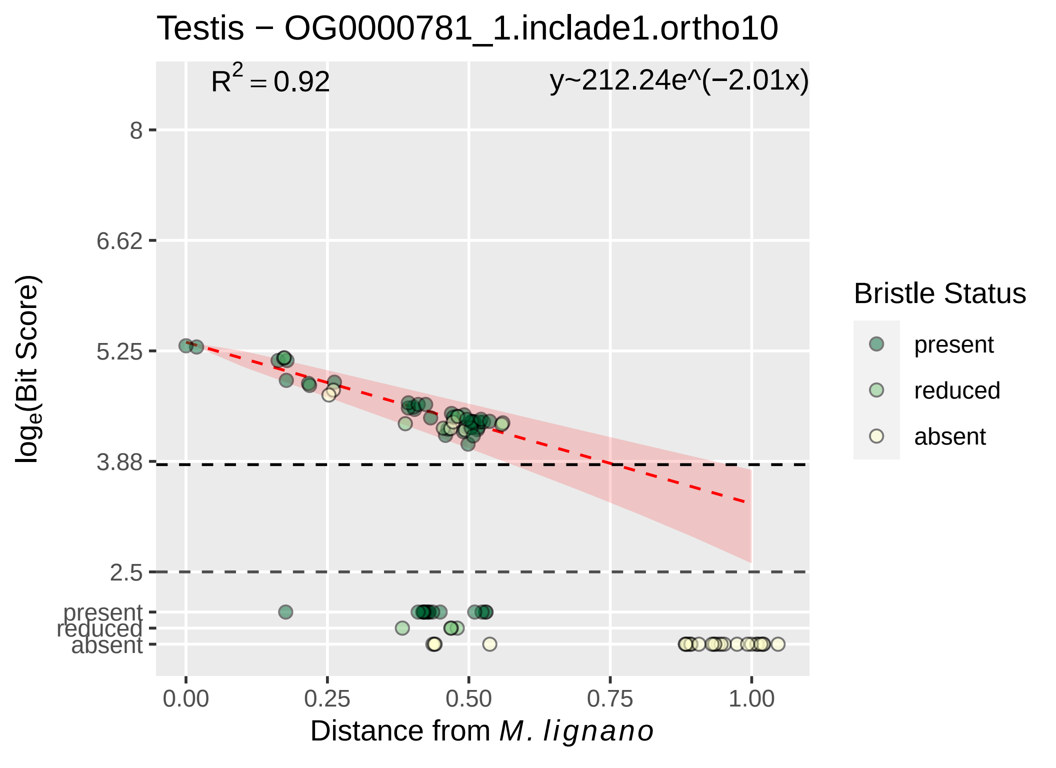

Testis − OG0000781_1.inclade1.ortho10
R^2 = 0.92
y~212.24e^(−2.01x)
8
6.62
5.25
3.88
2.5
present
reduced
absent
loge(Bit Score)
0.00
0.25
0.50
0.75
1.00
Distance from M. lignano
Bristle Status
present
reduced
absent

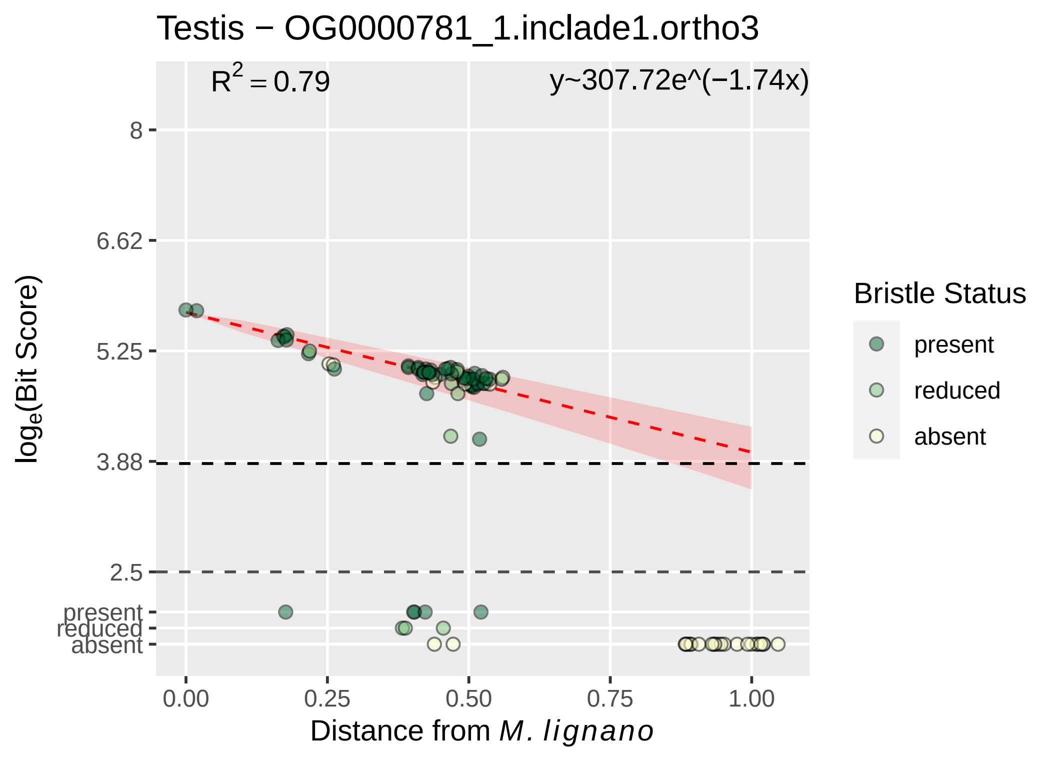

Testis − OG0000781_1.inclade1.ortho3
$R^2 = 0.79$
y~307.72e^(−1.74x)
8
6.62
5.25
3.88
2.5
present
reduced
absent
$log_e$(Bit Score)
0.00
0.25
0.50
0.75
1.00
Distance from *M. lignano*
Bristle Status
present
reduced
absent

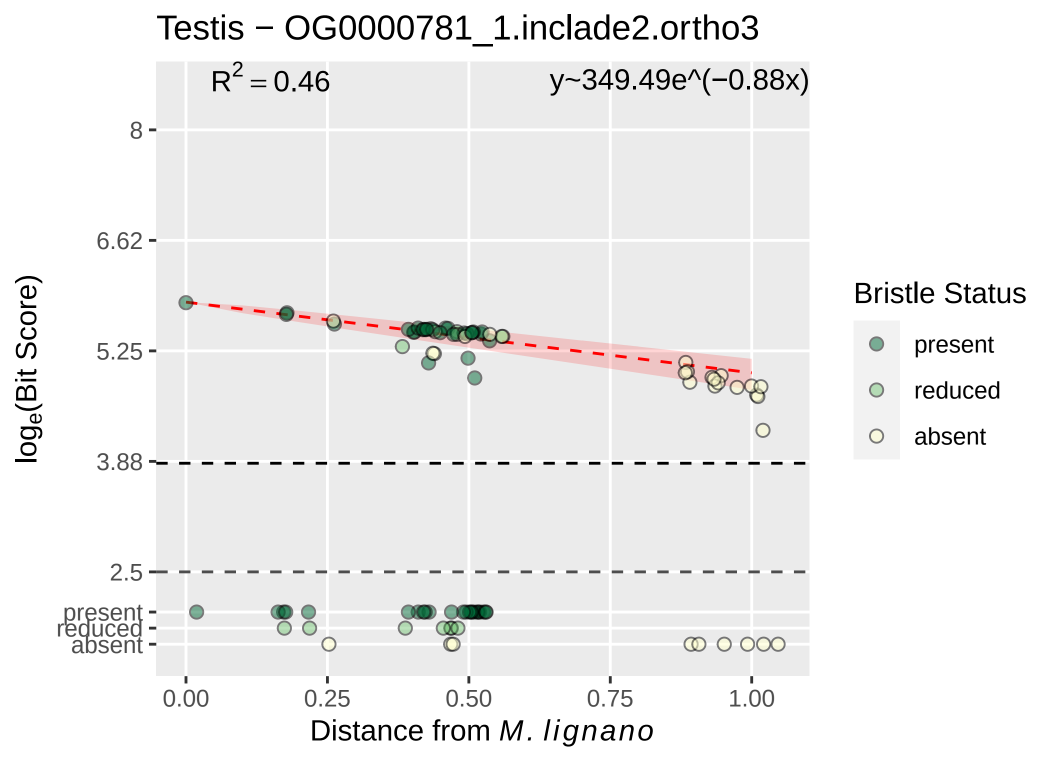

Testis − OG0000781_1.inclade2.ortho3
$R^2 = 0.46$
y~349.49e^(−0.88x)
8
6.62
5.25
3.88
2.5
present
reduced
absent
$\log_e$(Bit Score)
0.00
0.25
0.50
0.75
1.00
Distance from *M. lignano*
Bristle Status
present
reduced
absent

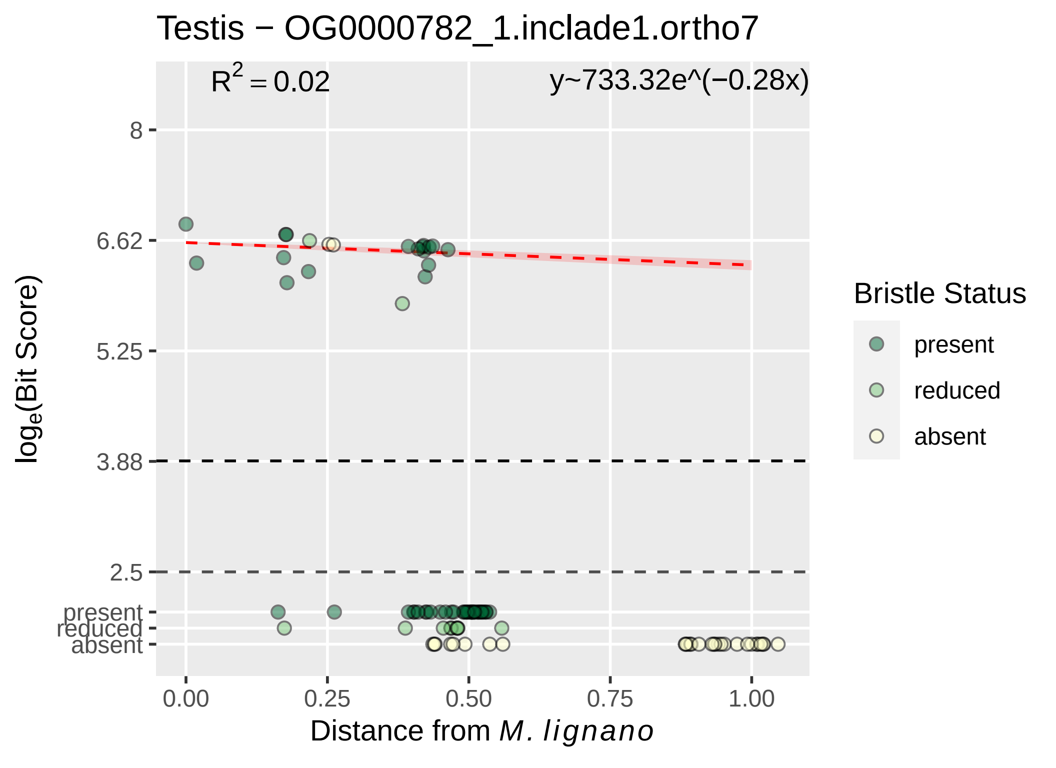

Testis − OG0000782_1.inclade1.ortho7
R^2 = 0.02
y~733.32e^(−0.28x)
8
6.62
5.25
3.88
2.5
present
reduced
absent
log_e(Bit Score)
0.00
0.25
0.50
0.75
1.00
Distance from M. lignano
Bristle Status
present
reduced
absent

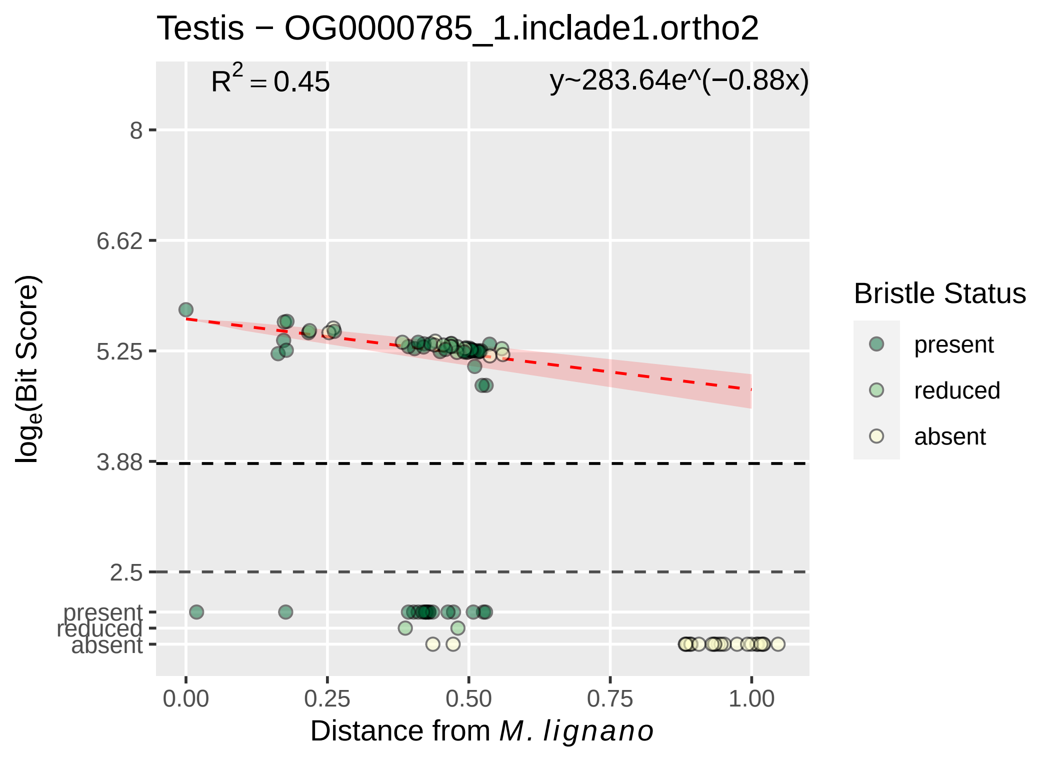

Testis − OG0000785_1.inclade1.ortho2
$R^2 = 0.45$
y~283.64e^(−0.88x)
8
6.62
5.25
3.88
2.5
present
reduced
absent
0.00
0.25
0.50
0.75
1.00
$log_e$(Bit Score)
Distance from *M. lignano*
Bristle Status
present
reduced
absent

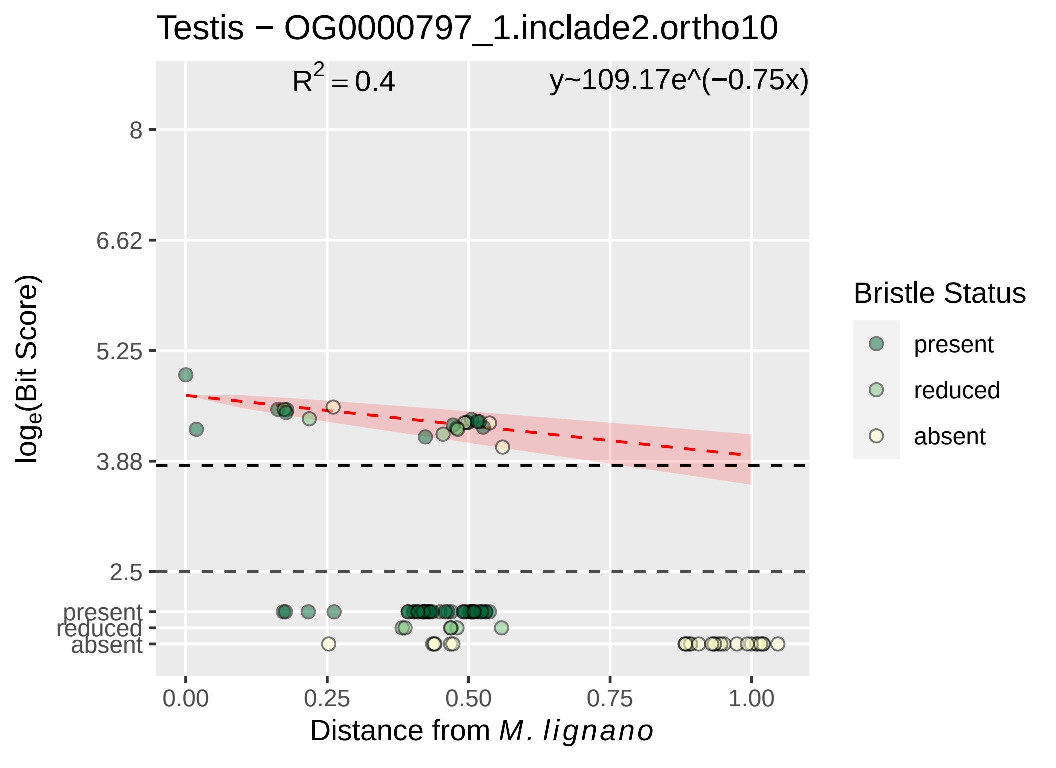

Testis − OG0000797_1.inclade2.ortho10
$R^2 = 0.4$
y~109.17e^(−0.75x)
8
6.62
5.25
3.88
2.5
present
reduced
absent
$log_e$(Bit Score)
0.00
0.25
0.50
0.75
1.00
Distance from *M. lignano*
Bristle Status
present
reduced
absent

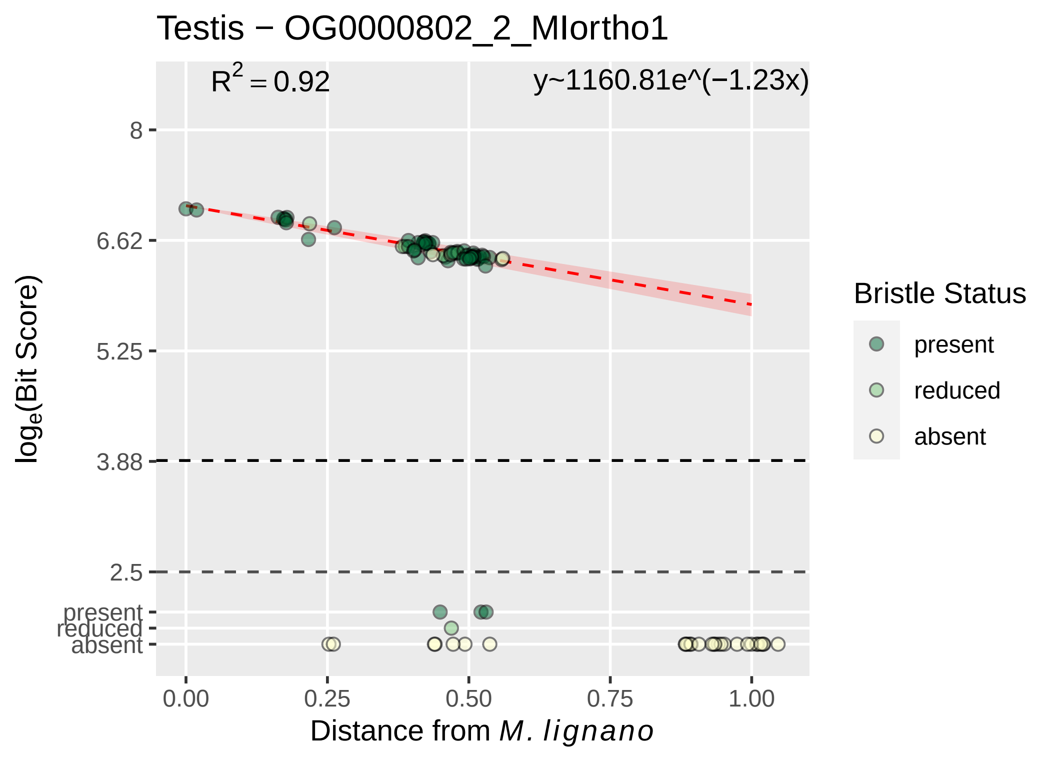
Testis − OG0000802_2_MIortho1
$R^2 = 0.92$
y~1160.81e^(−1.23x)
8
6.62
5.25
3.88
2.5
present
reduced
absent
0.00
0.25
0.50
0.75
1.00
log$_e$(Bit Score)
Distance from *M. lignano*
Bristle Status
present
reduced
absent

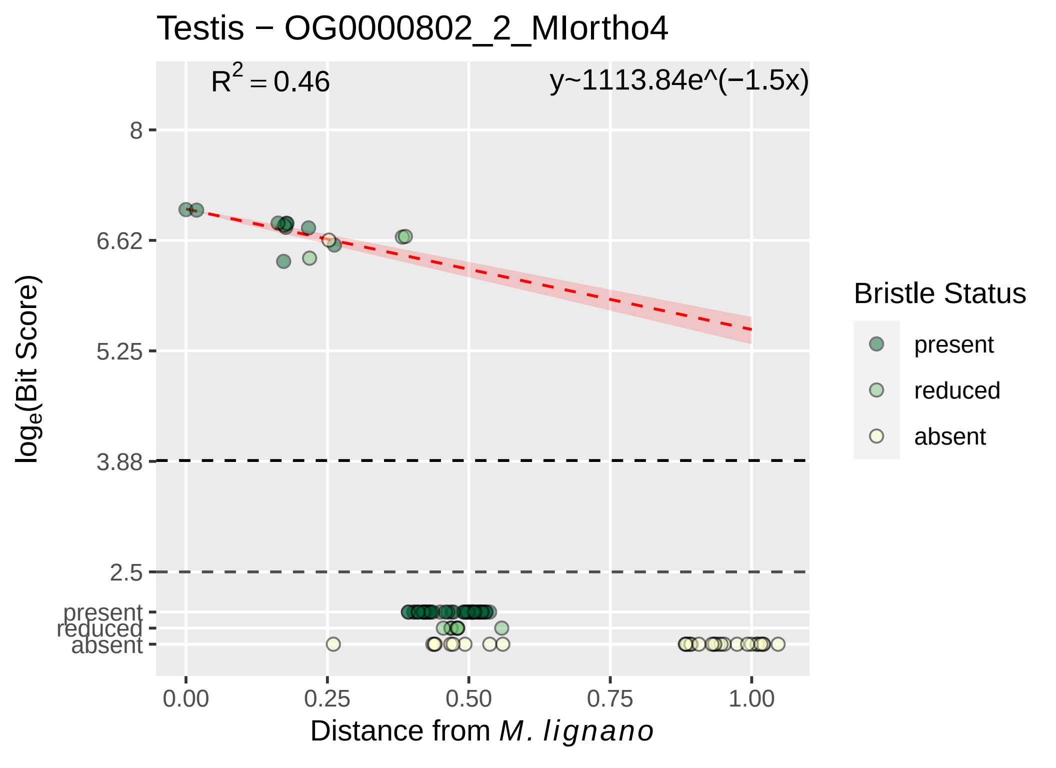
Testis − OG0000802_2_MIortho4
R^2 = 0.46
y~1113.84e^(−1.5x)
8
6.62
5.25
3.88
2.5
present
reduced
absent
log_e(Bit Score)
0.00
0.25
0.50
0.75
1.00
Distance from M. lignano
Bristle Status
present
reduced
absent

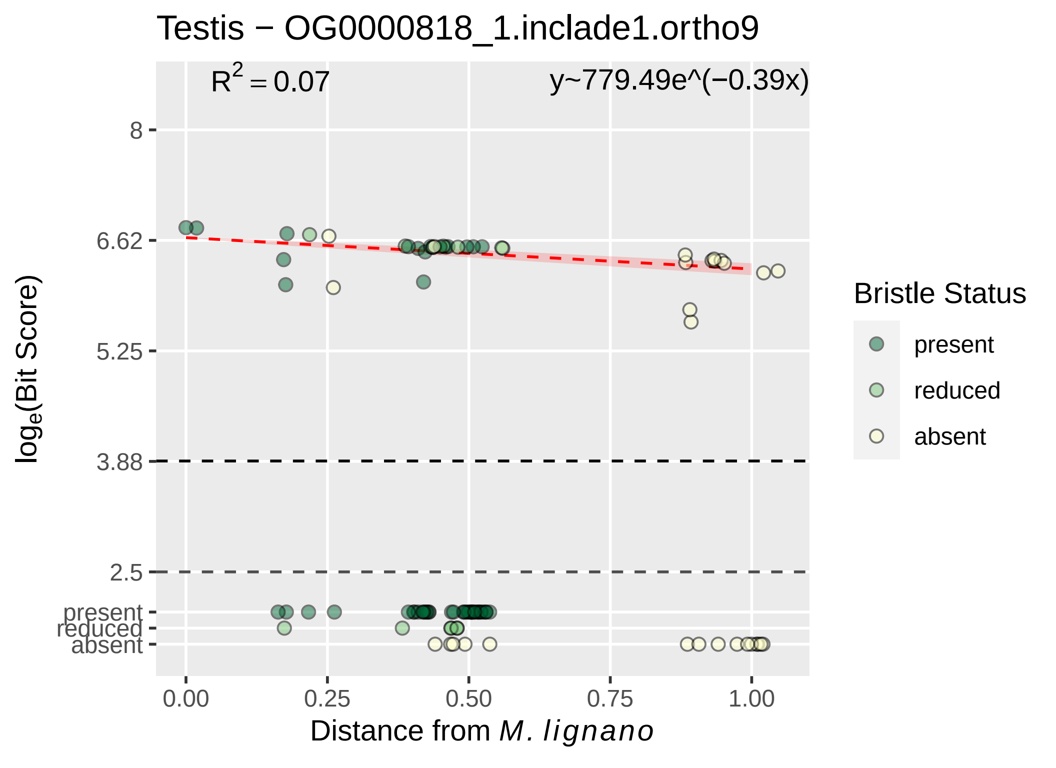

Testis − OG0000818_1.inclade1.ortho9
$R^2 = 0.07$
y~779.49e^(−0.39x)
8
6.62
5.25
3.88
2.5
present
reduced
absent
0.00
0.25
0.50
0.75
1.00
Distance from M. lignano
log$_e$(Bit Score)
Bristle Status
present
reduced
absent

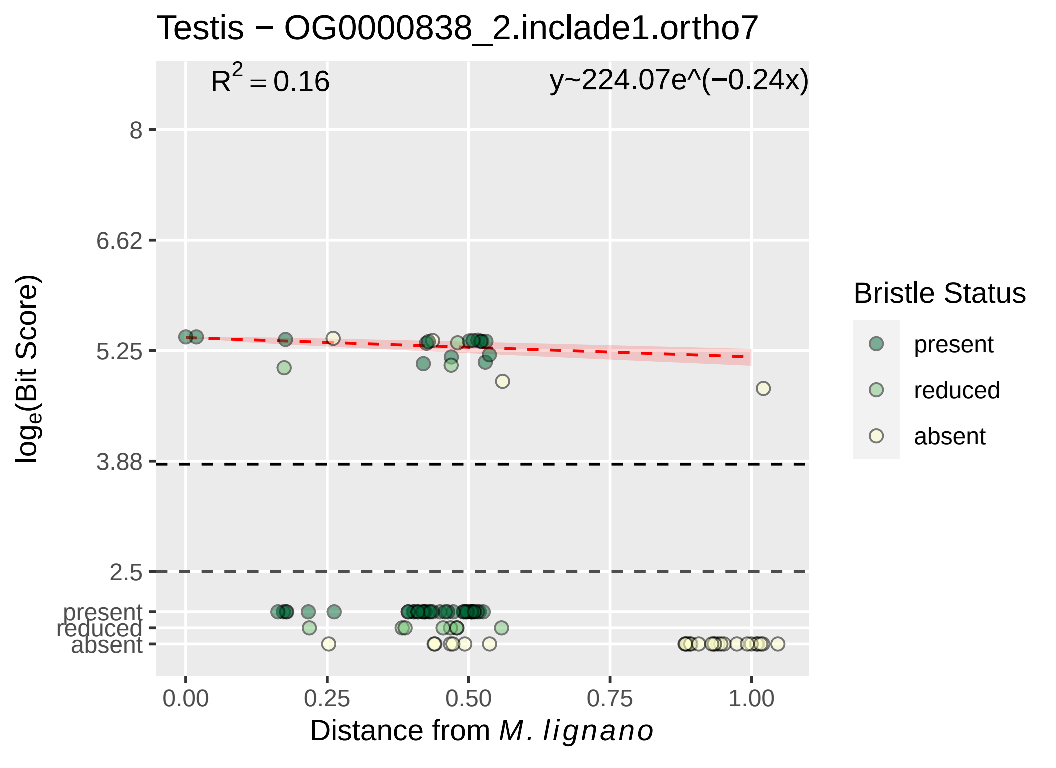

Testis − OG0000838_2.inclade1.ortho7
$R^2 = 0.16$
y~224.07e^(−0.24x)
8
6.62
5.25
3.88
2.5
present
reduced
absent
$\log_e$(Bit Score)
0.00
0.25
0.50
0.75
1.00
Distance from *M. lignano*
Bristle Status
present
reduced
absent

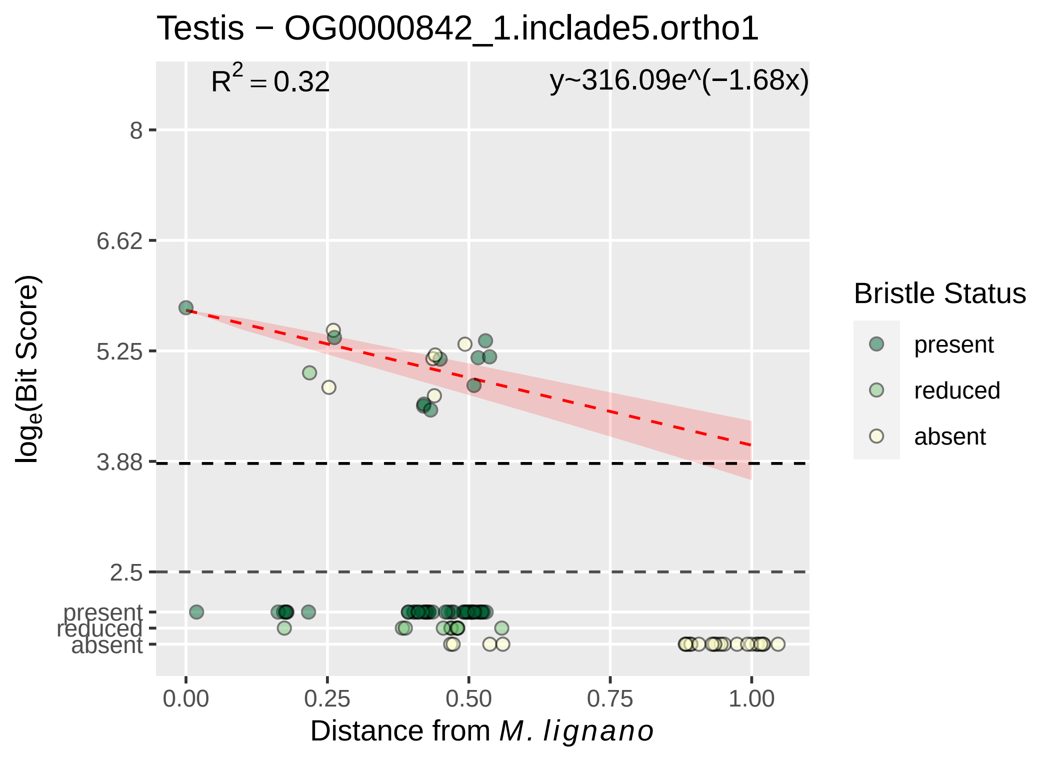

Testis − OG0000842_1.inclade5.ortho1
R^2 = 0.32
y~316.09e^(−1.68x)
8
6.62
5.25
3.88
2.5
present
reduced
absent
loge(Bit Score)
0.00
0.25
0.50
0.75
1.00
Distance from M. lignano
Bristle Status
present
reduced
absent

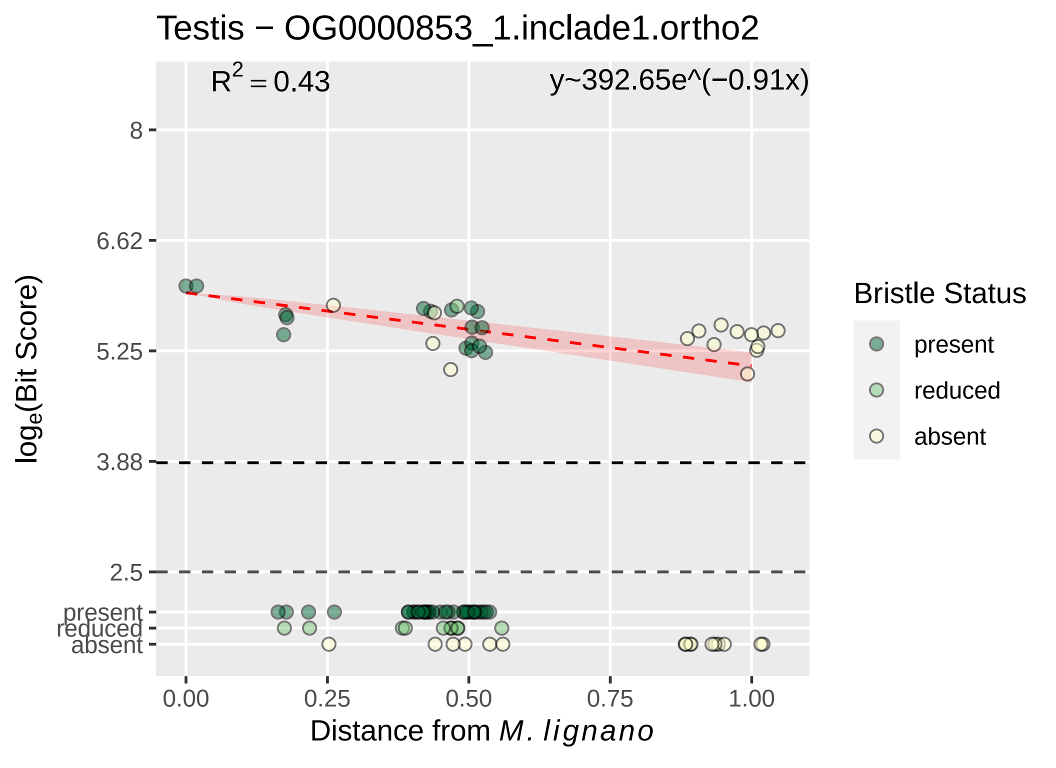

Testis − OG0000853_1.inclade1.ortho2
$R^2 = 0.43$
y~392.65e^(−0.91x)
8
6.62
5.25
3.88
2.5
present
reduced
absent
0.00
0.25
0.50
0.75
1.00
Distance from M. lignano
log$_e$(Bit Score)
Bristle Status
present
reduced
absent

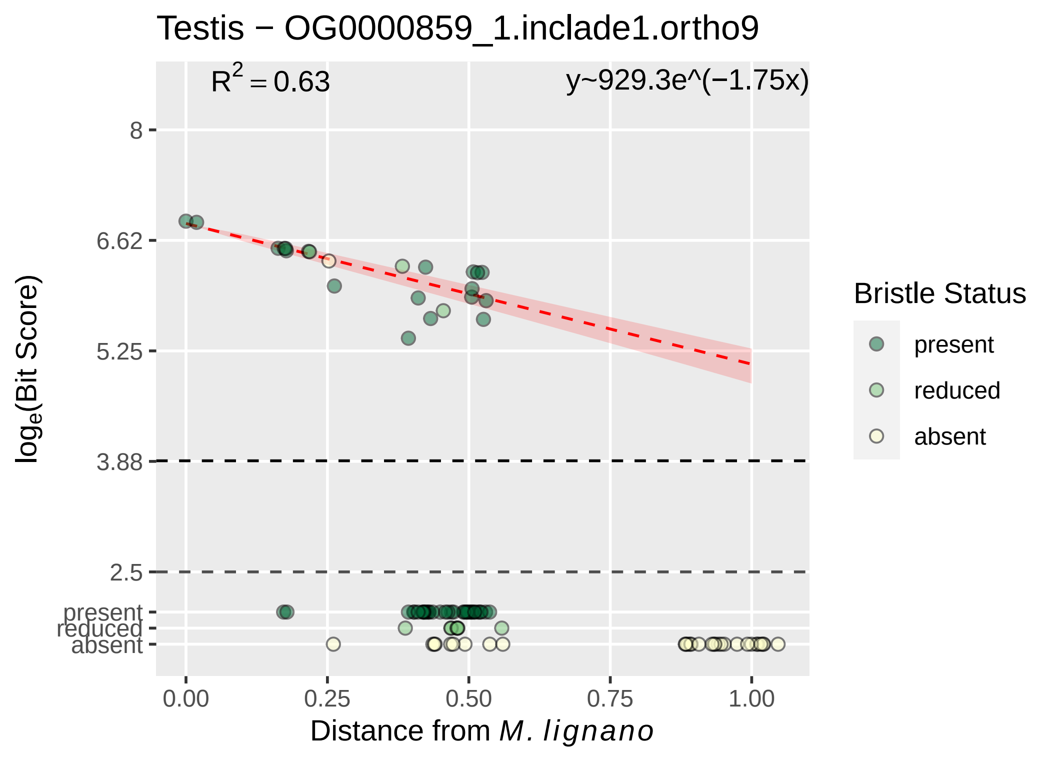

Testis − OG0000859_1.inclade1.ortho9
R^2 = 0.63
y~929.3e^(−1.75x)
8
6.62
5.25
3.88
2.5
present
reduced
absent
0.00
0.25
0.50
0.75
1.00
Distance from M. lignano
log_e(Bit Score)
Bristle Status
present
reduced
absent

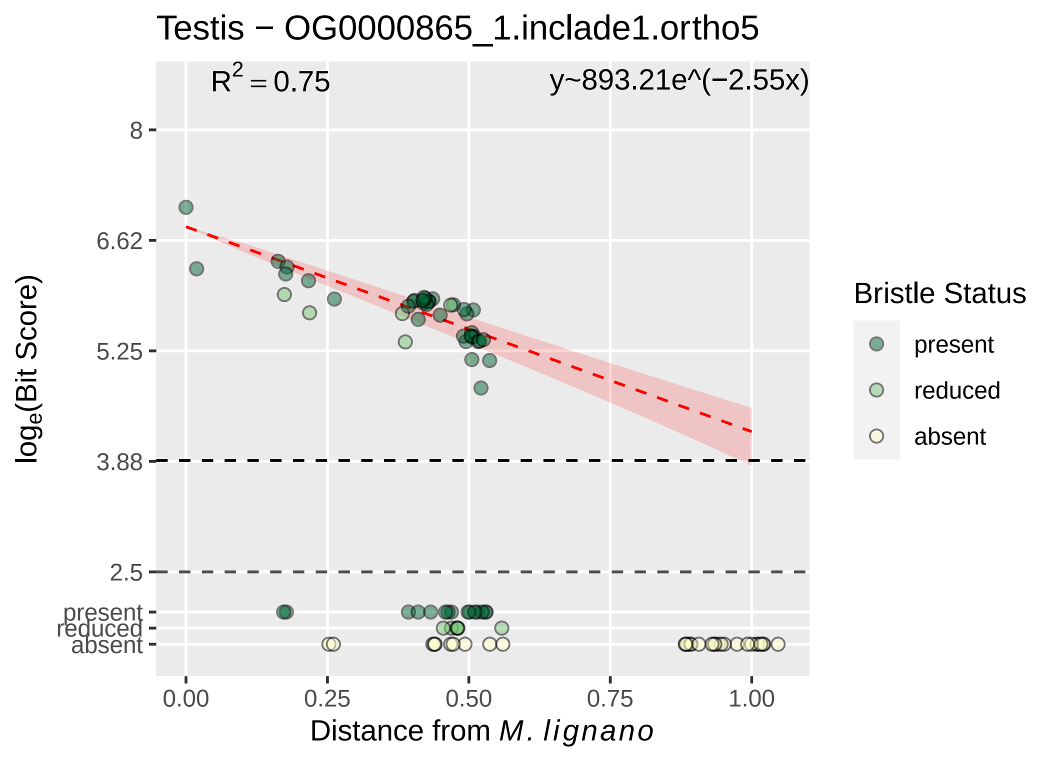

Testis − OG0000865_1.inclade1.ortho5
R^2 = 0.75
y~893.21e^(−2.55x)
8
6.62
5.25
3.88
2.5
present
reduced
absent
log_e(Bit Score)
0.00
0.25
0.50
0.75
1.00
Distance from M. lignano
Bristle Status
present
reduced
absent

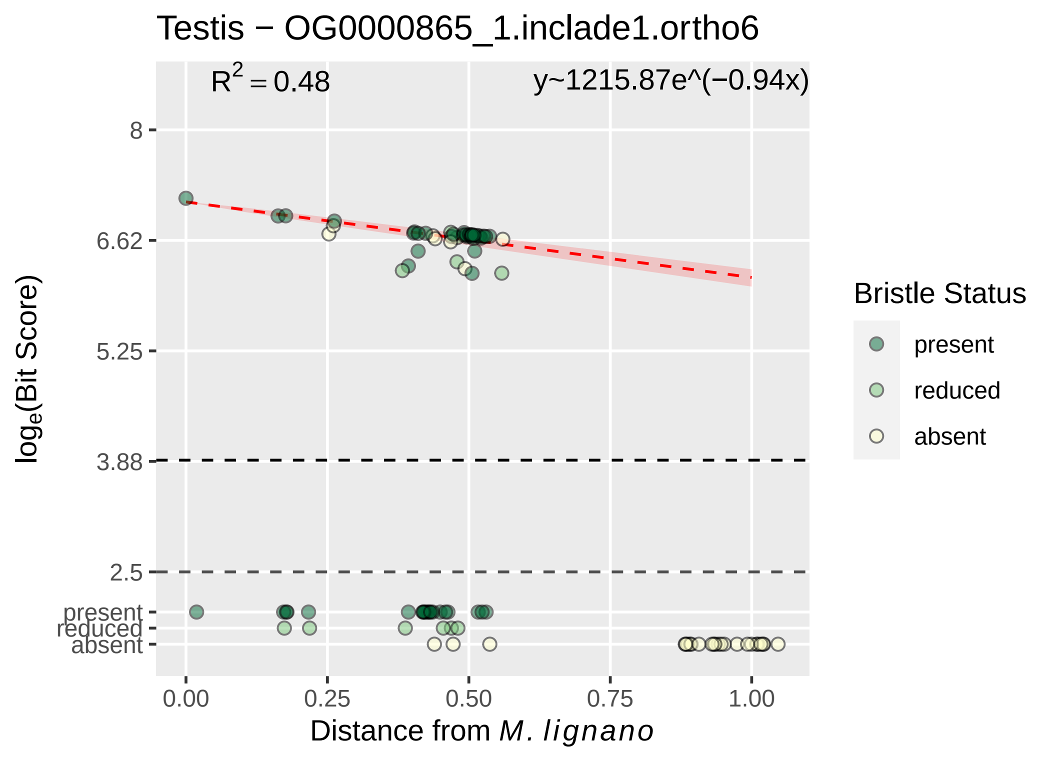

Testis − OG0000865_1.inclade1.ortho6
R2 = 0.48
y~1215.87e^(−0.94x)
8
6.62
5.25
3.88
2.5
present
reduced
absent
0.00
0.25
0.50
0.75
1.00
Distance from M. lignano
loge(Bit Score)
Bristle Status
present
reduced
absent

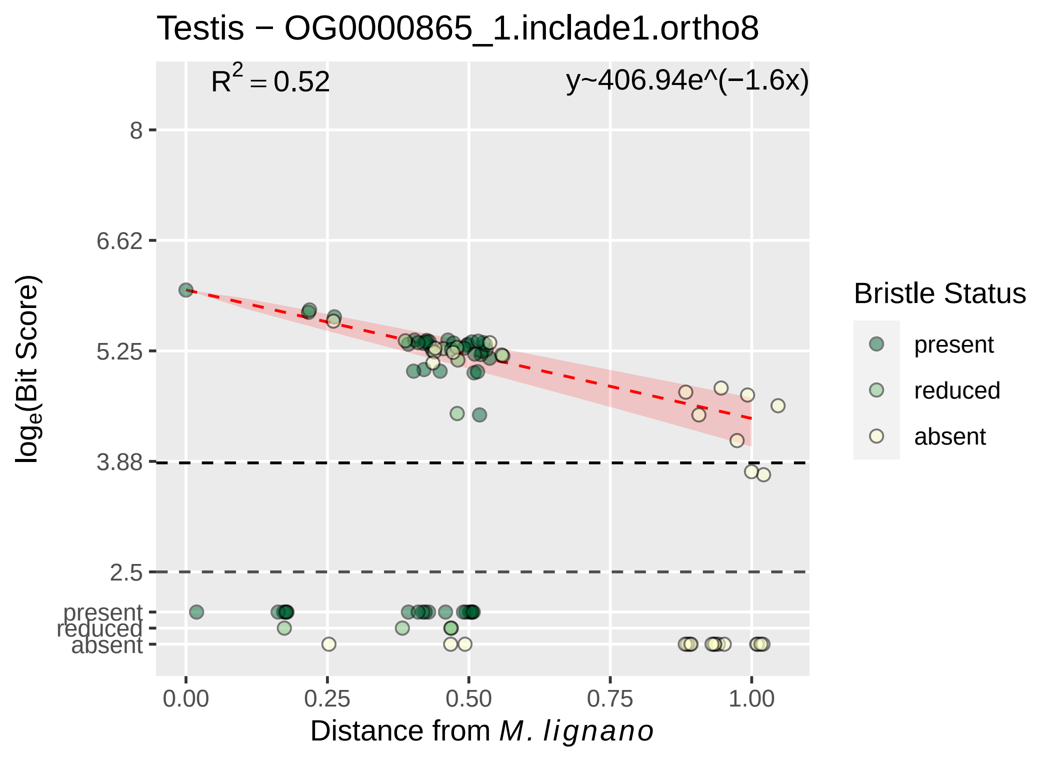

Testis − OG0000865_1.inclade1.ortho8
R^2 = 0.52
y~406.94e^(−1.6x)
8
6.62
5.25
3.88
2.5
present
reduced
absent
0.00
0.25
0.50
0.75
1.00
log_e(Bit Score)
Distance from M. lignano
Bristle Status
present
reduced
absent

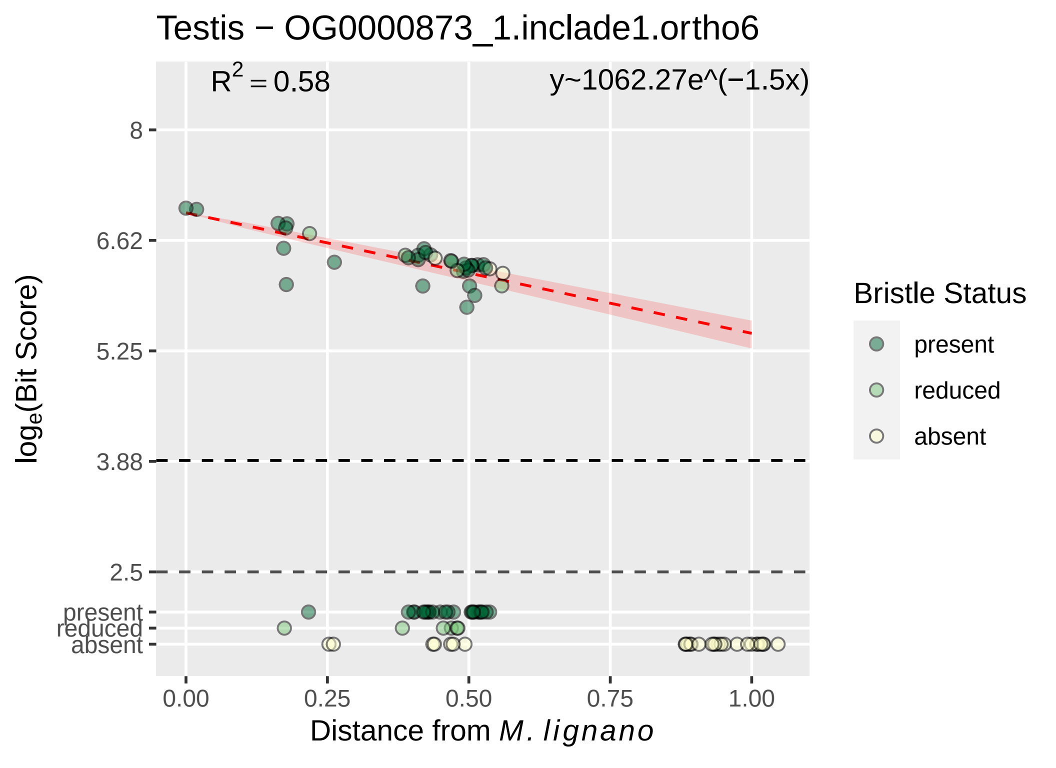

Testis − OG0000873_1.inclade1.ortho6
R^2 = 0.58
y~1062.27e^(−1.5x)
8
6.62
5.25
3.88
2.5
present
reduced
absent
0.00
0.25
0.50
0.75
1.00
log_e(Bit Score)
Distance from M. lignano
Bristle Status
present
reduced
absent

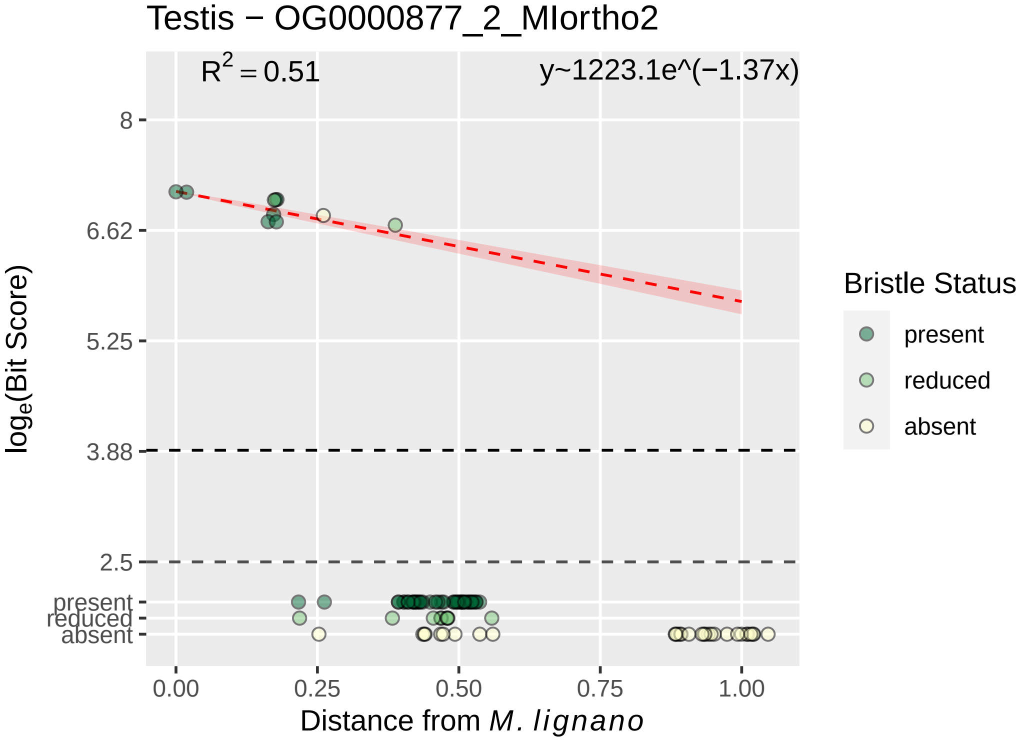

Testis − OG0000877_2_MIortho2
$R^2 = 0.51$
y~1223.1e^(−1.37x)
8
6.62
5.25
3.88
2.5
present
reduced
absent
0.00
0.25
0.50
0.75
1.00
Distance from *M. lignano*
$\log_e$(Bit Score)
Bristle Status
present
reduced
absent

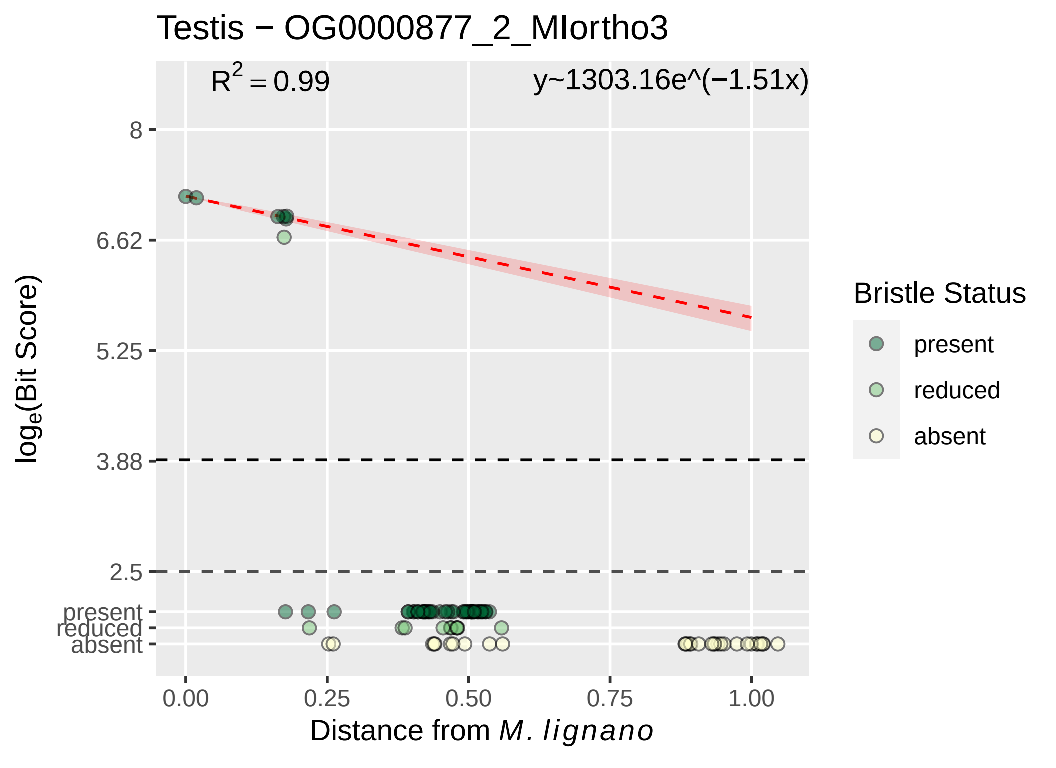

Testis − OG0000877_2_MIortho3
R^2 = 0.99
y~1303.16e^(−1.51x)
8
6.62
5.25
3.88
2.5
present
reduced
absent
0.00
0.25
0.50
0.75
1.00
Distance from M. lignano
loge(Bit Score)
Bristle Status
present
reduced
absent

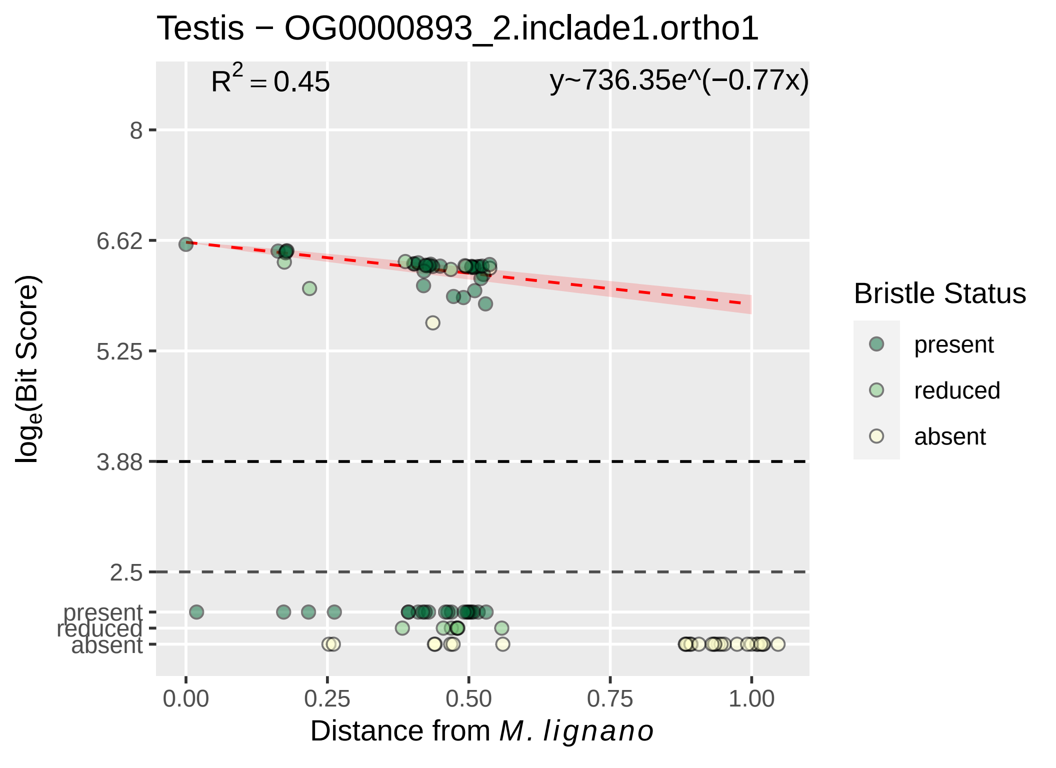

Testis − OG0000893_2.inclade1.ortho1
R^2 = 0.45
y~736.35e^(−0.77x)
8
6.62
5.25
3.88
2.5
present
reduced
absent
0.00
0.25
0.50
0.75
1.00
loge(Bit Score)
Distance from M. lignano
Bristle Status
present
reduced
absent

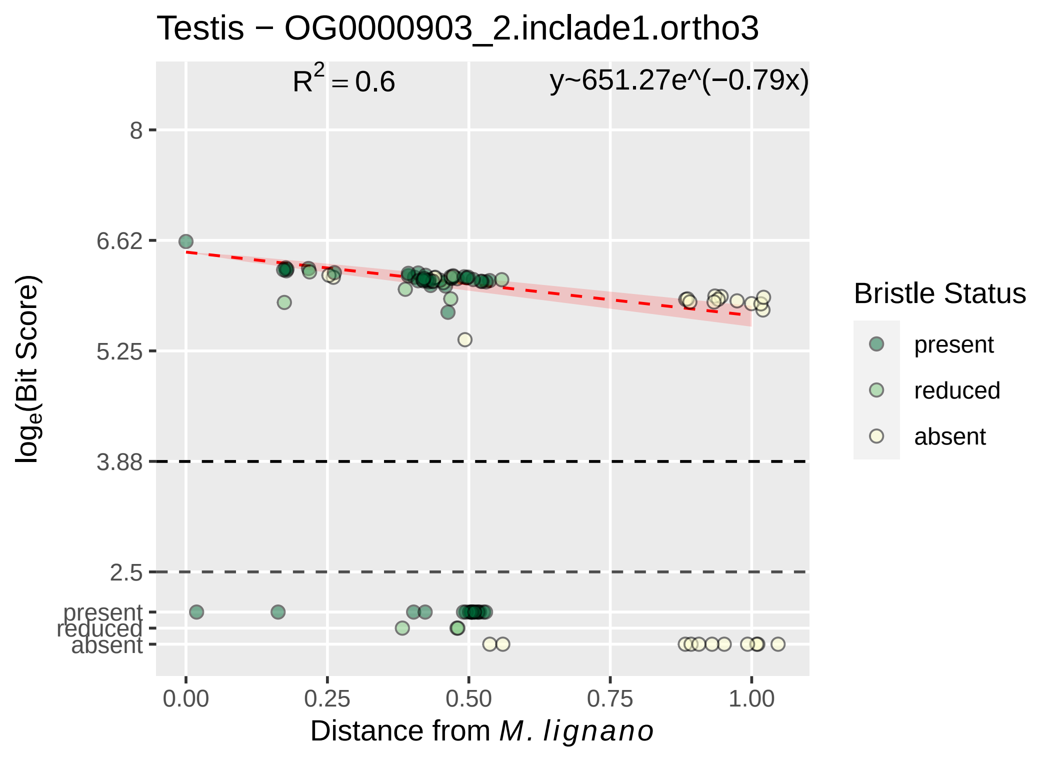

Testis − OG0000903_2.inclade1.ortho3
$R^2 = 0.6$
y~651.27e^(−0.79x)
8
6.62
5.25
3.88
2.5
present
reduced
absent
log_e(Bit Score)
0.00
0.25
0.50
0.75
1.00
Distance from M. lignano
Bristle Status
present
reduced
absent

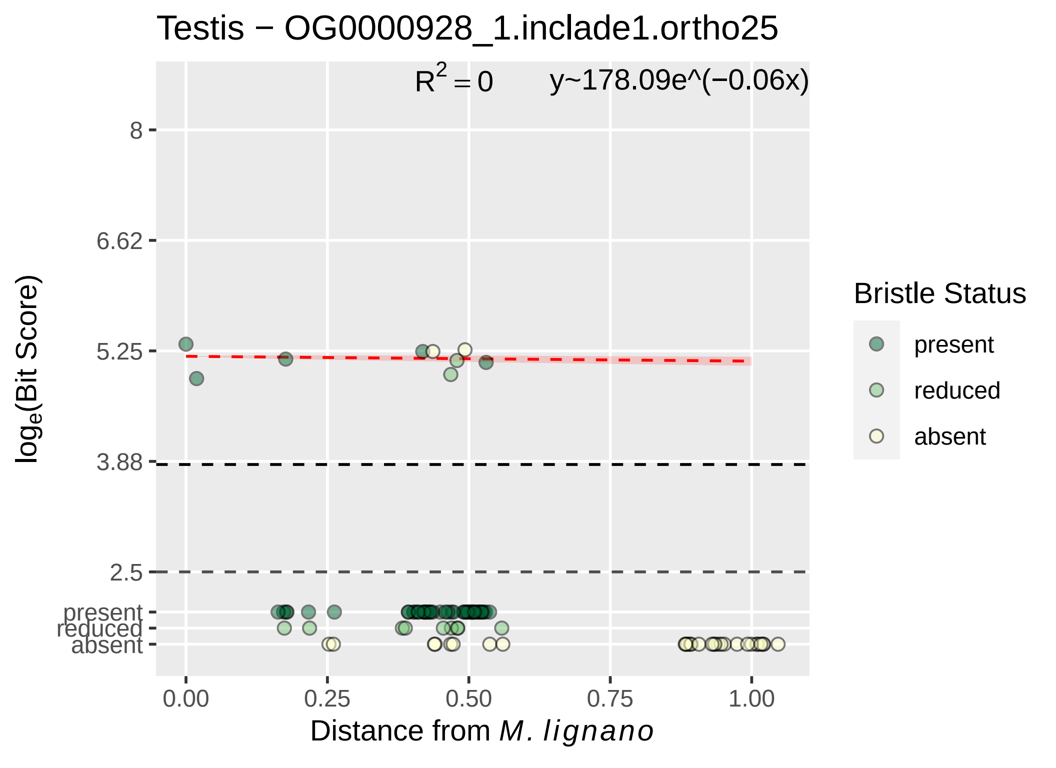

Testis − OG0000928_1.inclade1.ortho25
$R^2 = 0$
y~178.09e^(−0.06x)
8
6.62
5.25
3.88
2.5
present
reduced
absent
0.00
0.25
0.50
0.75
1.00
$\log_e$(Bit Score)
Distance from *M. lignano*
Bristle Status
present
reduced
absent

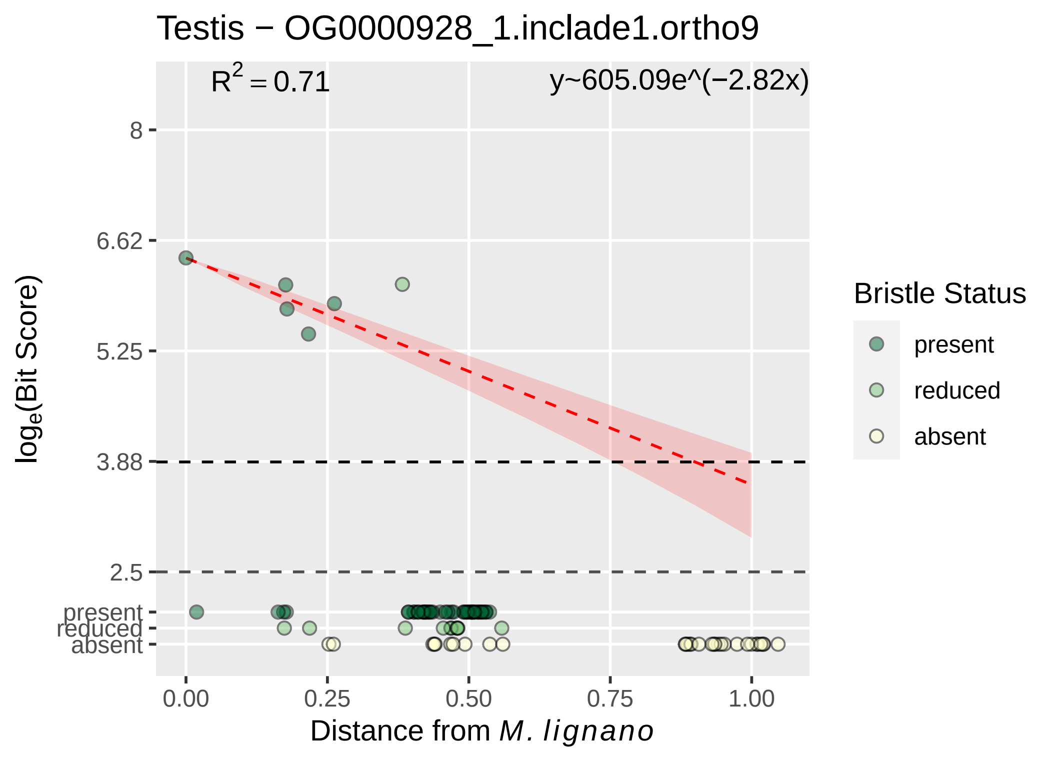

Testis − OG0000928_1.inclade1.ortho9
$R^2 = 0.71$
y~605.09e^(−2.82x)
8
6.62
5.25
3.88
2.5
present
reduced
absent
$\log_e$(Bit Score)
0.00
0.25
0.50
0.75
1.00
Distance from *M. lignano*
Bristle Status
present
reduced
absent

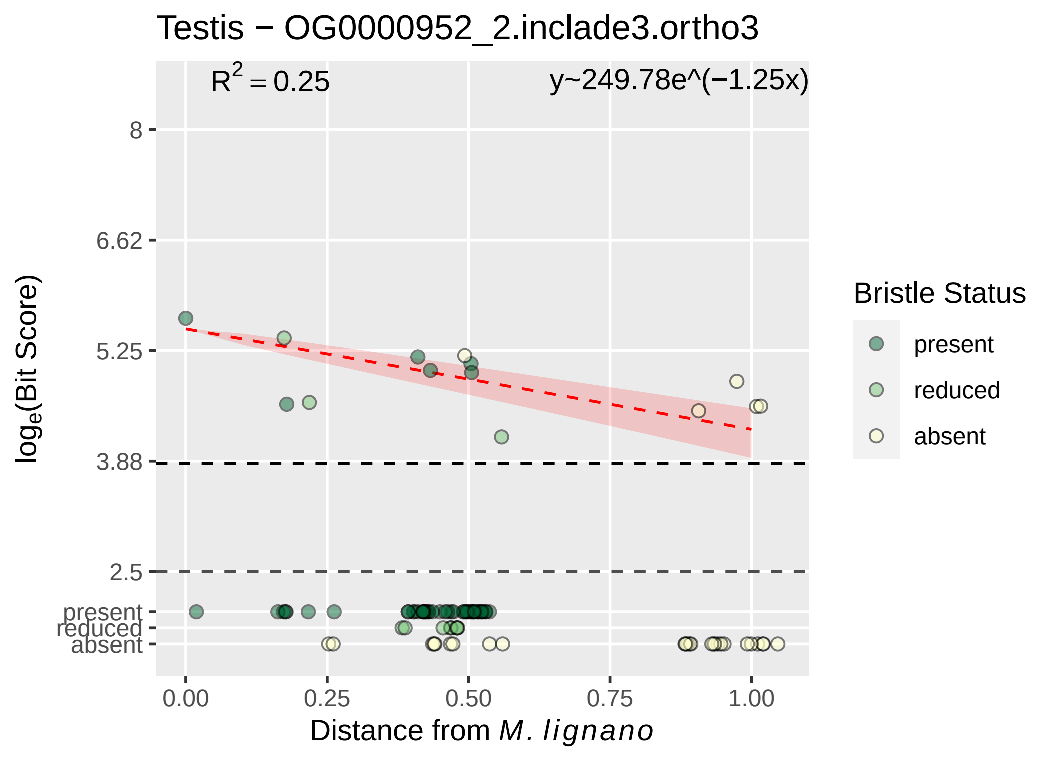

Testis − OG0000952_2.inclade3.ortho3
R^2 = 0.25
y~249.78e^(−1.25x)
8
6.62
5.25
3.88
2.5
present
reduced
absent
log_e(Bit Score)
0.00
0.25
0.50
0.75
1.00
Distance from M. lignano
Bristle Status
present
reduced
absent

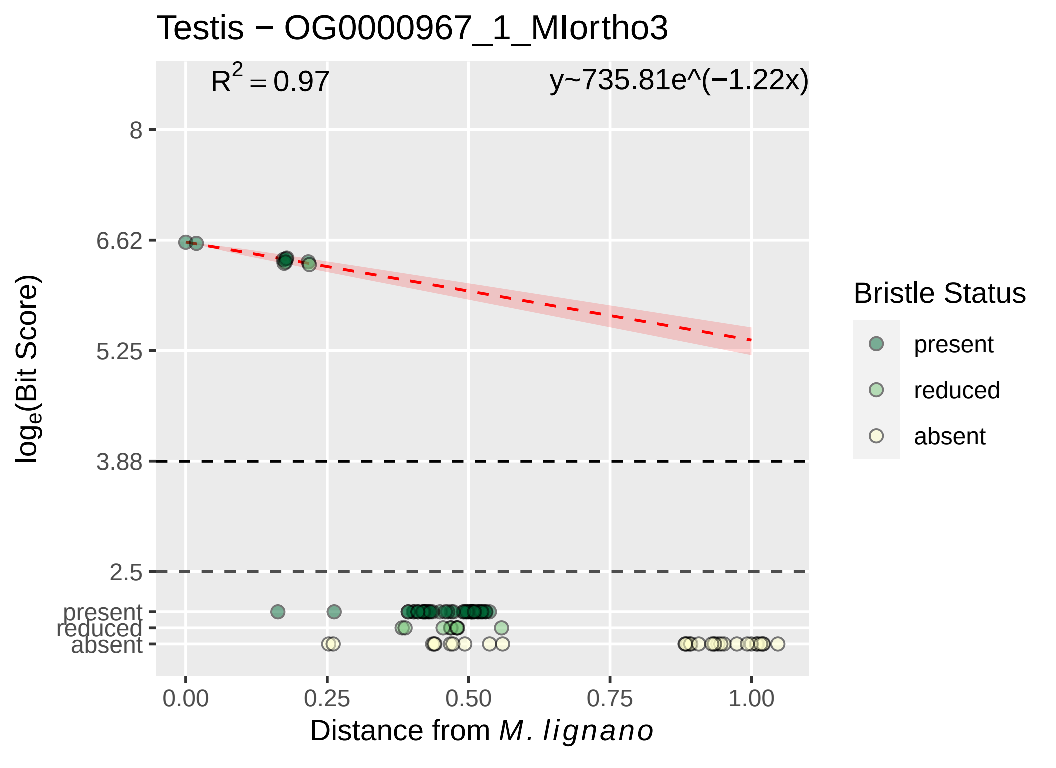

Testis − OG0000967_1_MIortho3
R^2 = 0.97
y~735.81e^(−1.22x)
8
6.62
5.25
3.88
2.5
present
reduced
absent
log_e(Bit Score)
0.00
0.25
0.50
0.75
1.00
Distance from M. lignano
Bristle Status
present
reduced
absent

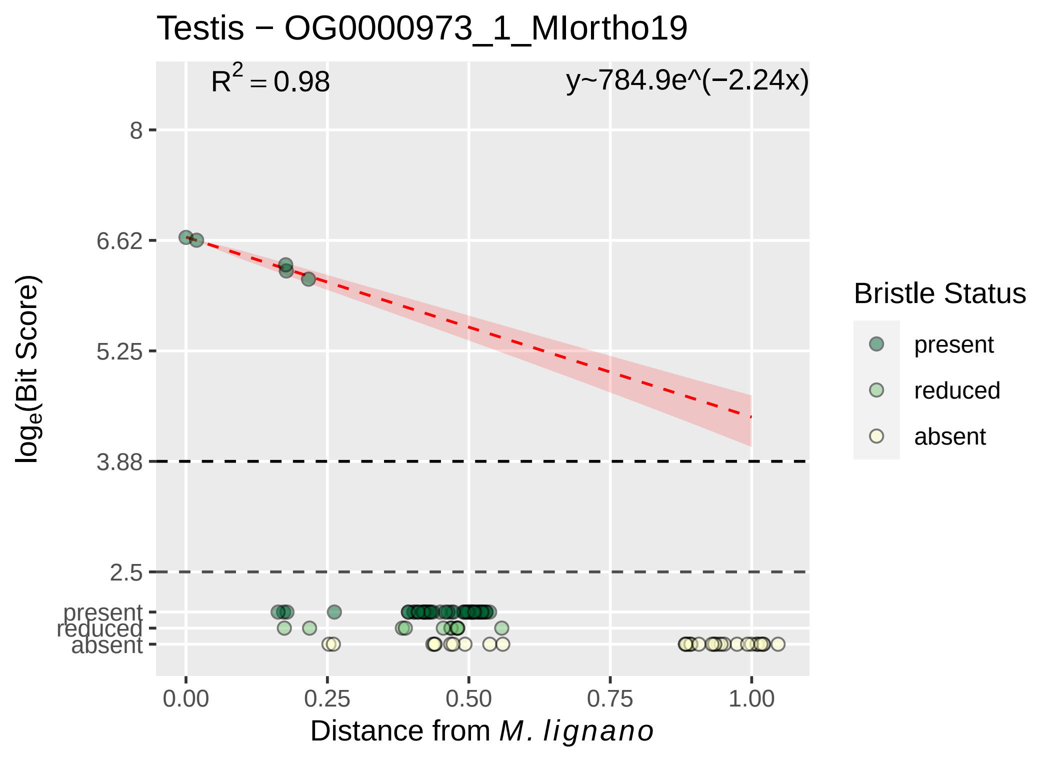

Testis − OG0000973_1_MIortho19
$R^2 = 0.98$
y~784.9e^(−2.24x)
8
6.62
5.25
3.88
2.5
present
reduced
absent
$log_e$(Bit Score)
0.00
0.25
0.50
0.75
1.00
Distance from *M. lignano*
Bristle Status
present
reduced
absent

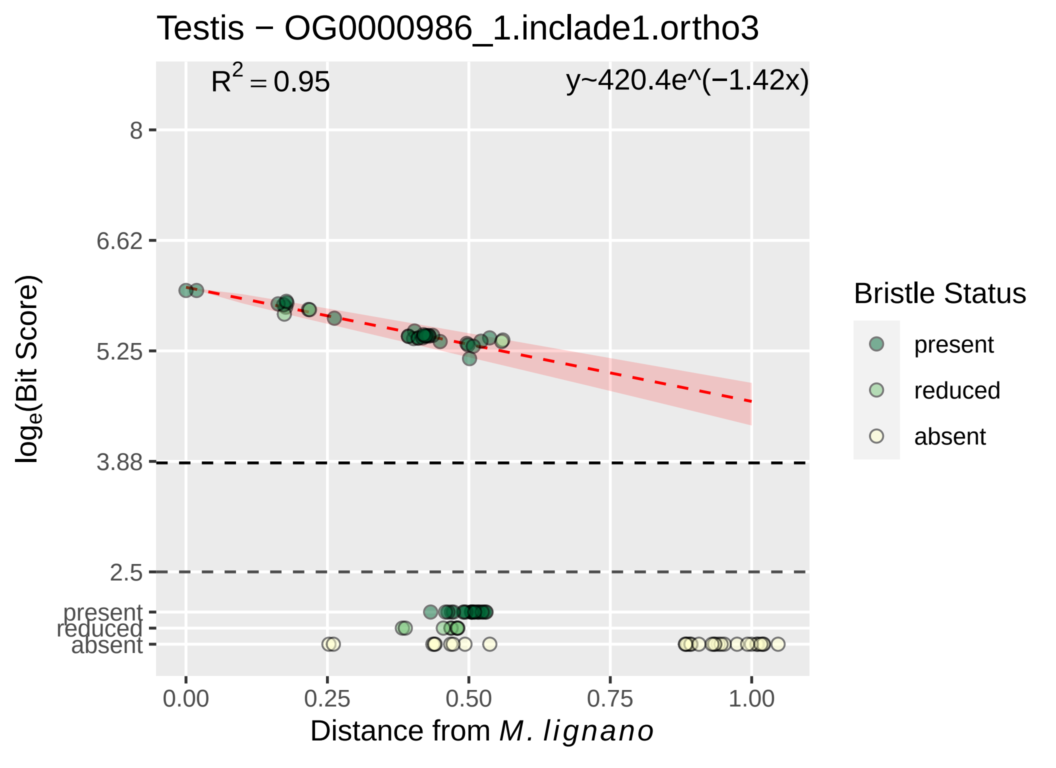
Testis − OG0000986_1.inclade1.ortho3
$R^2 = 0.95$
y~420.4e^(−1.42x)
8
6.62
5.25
3.88
2.5
present
reduced
absent
$log_e$(Bit Score)
0.00
0.25
0.50
0.75
1.00
Distance from M. lignano
Bristle Status
present
reduced
absent

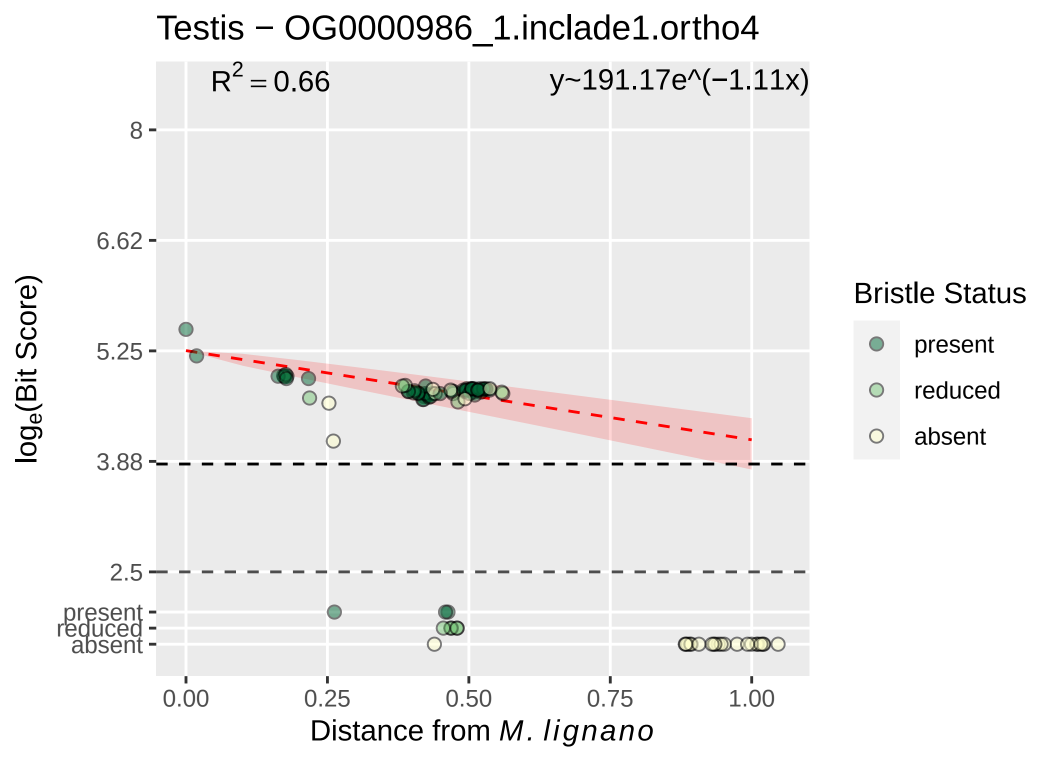

Testis − OG0000986_1.inclade1.ortho4
$R^2 = 0.66$
y~191.17e^(−1.11x)
8
6.62
5.25
3.88
2.5
present
reduced
absent
0.00
0.25
0.50
0.75
1.00
$\log_e$(Bit Score)
Distance from *M. lignano*
Bristle Status
present
reduced
absent

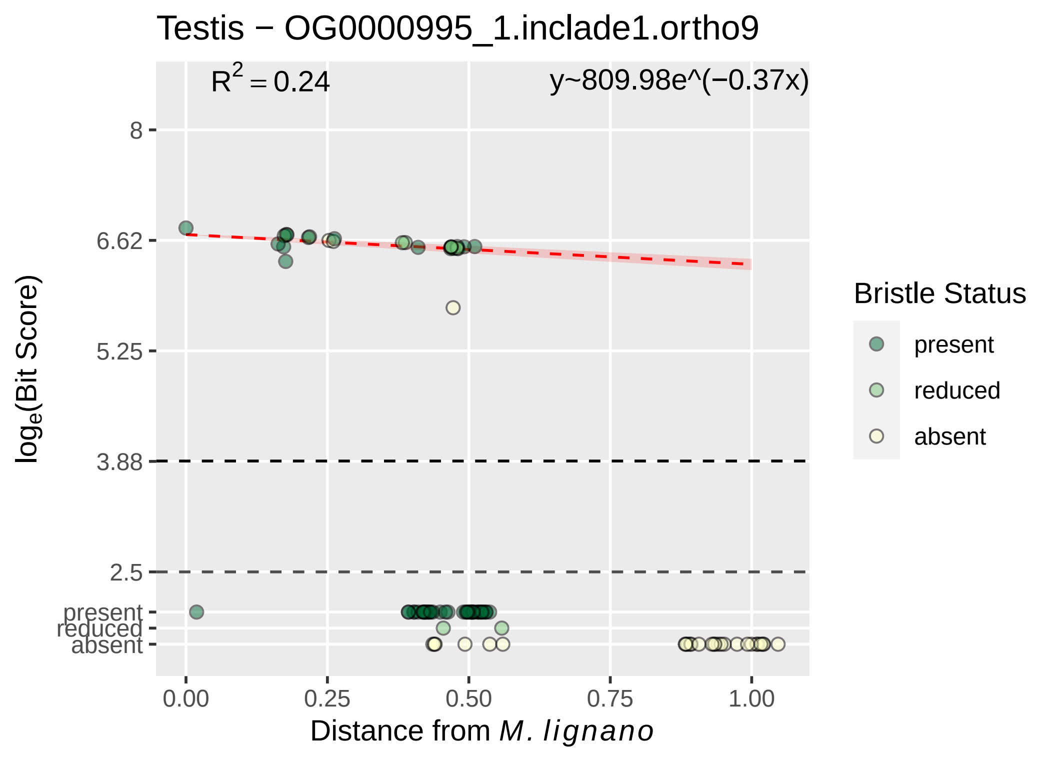

Testis − OG0000995_1.inclade1.ortho9
R^2 = 0.24
y~809.98e^(−0.37x)
8
6.62
5.25
3.88
2.5
present
reduced
absent
0.00
0.25
0.50
0.75
1.00
log_e(Bit Score)
Distance from M. lignano
Bristle Status
present
reduced
absent

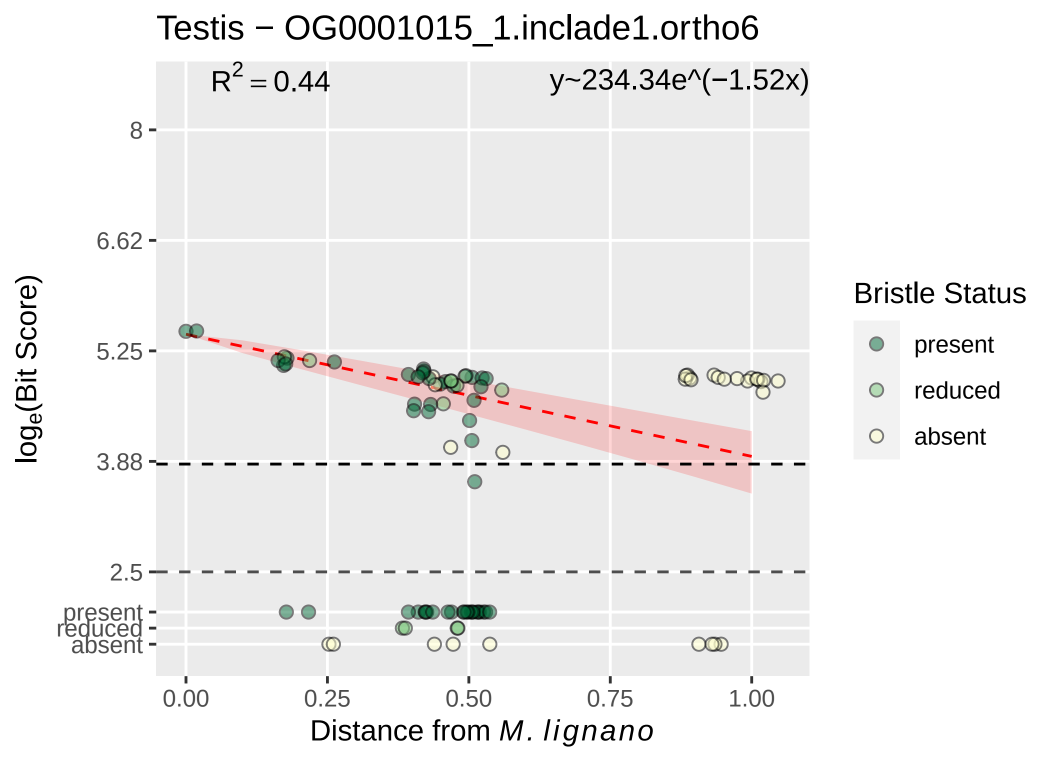
Testis − OG0001015_1.inclade1.ortho6
R2 = 0.44
y~234.34e^(−1.52x)
8
6.62
5.25
3.88
2.5
present
reduced
absent
loge(Bit Score)
0.00
0.25
0.50
0.75
1.00
Distance from M. lignano
Bristle Status
present
reduced
absent

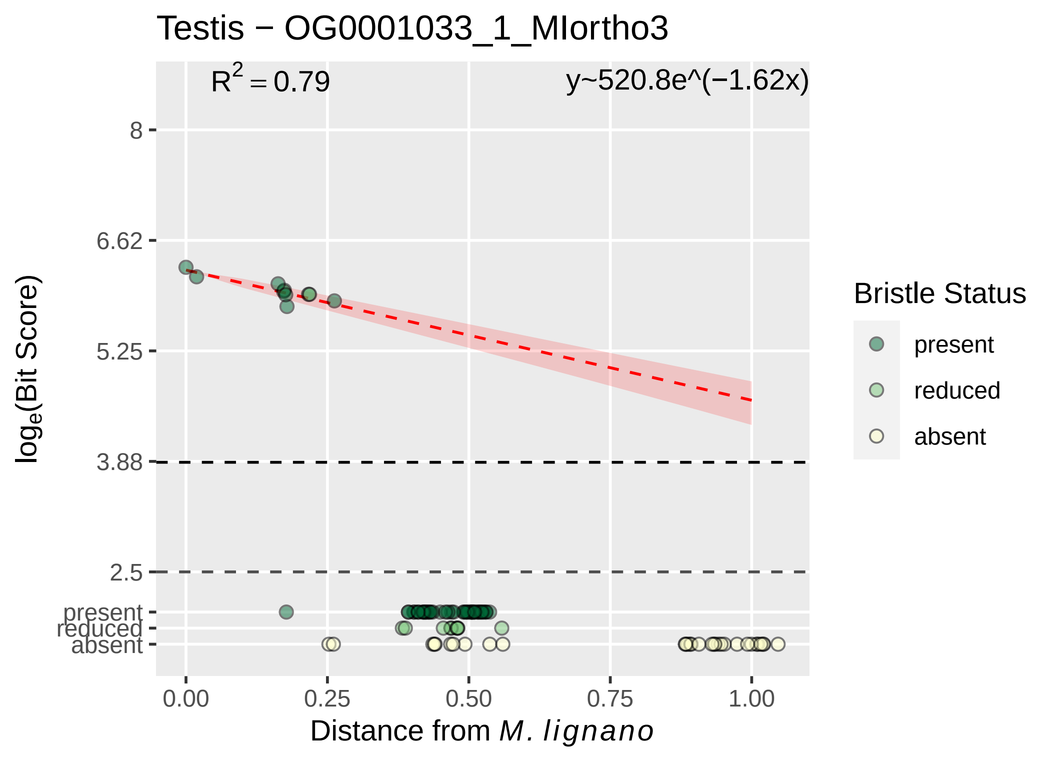

Testis − OG0001033_1_MIortho3
$R^2 = 0.79$
y~520.8e^(−1.62x)
8
6.62
5.25
3.88
2.5
present
reduced
absent
$\log_e$(Bit Score)
0.00
0.25
0.50
0.75
1.00
Distance from *M. lignano*
Bristle Status
present
reduced
absent

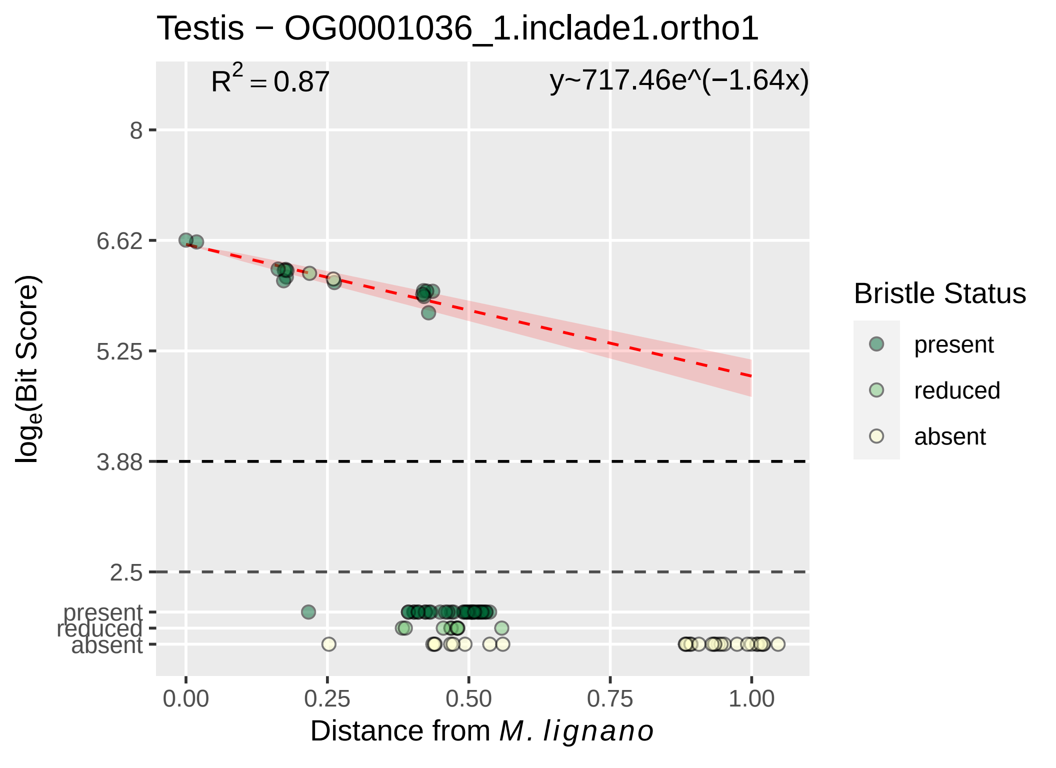
Testis − OG0001036_1.inclade1.ortho1
$R^2 = 0.87$
y~717.46e^(−1.64x)
8
6.62
5.25
3.88
2.5
present
reduced
absent
0.00
0.25
0.50
0.75
1.00
$log_e$(Bit Score)
Distance from *M. lignano*
Bristle Status
present
reduced
absent

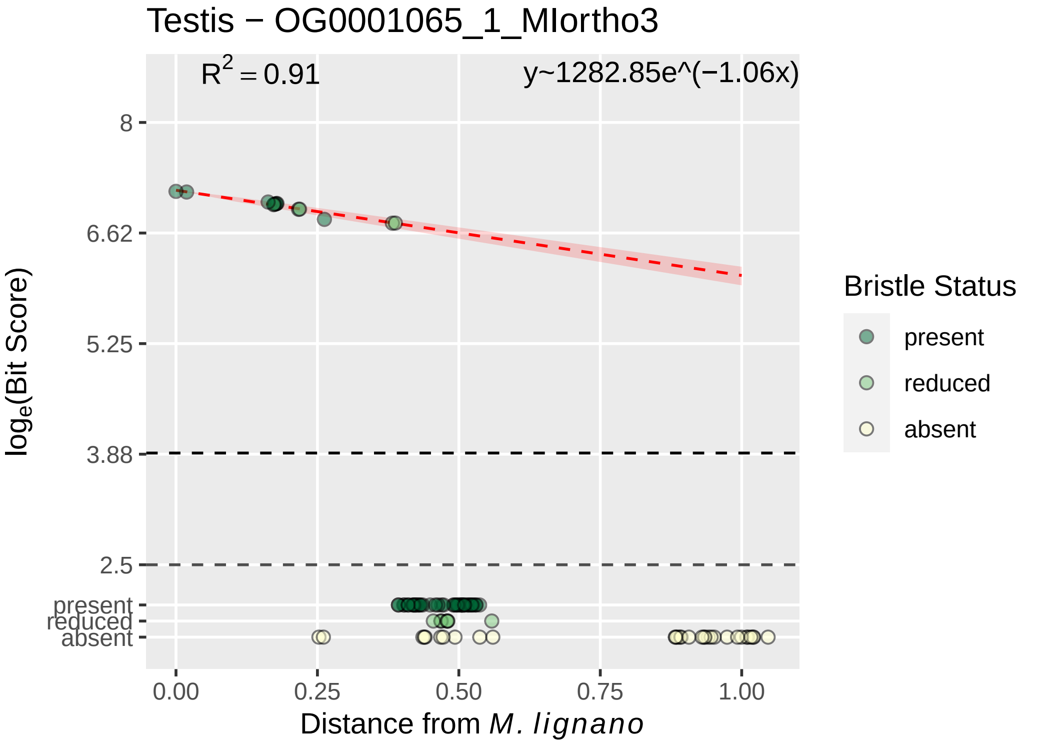
Testis − OG0001065_1_MIortho3
R^2 = 0.91
y~1282.85e^(−1.06x)
log_e(Bit Score)
8
6.62
5.25
3.88
2.5
present
reduced
absent
0.00
0.25
0.50
0.75
1.00
Distance from M. lignano
Bristle Status
present
reduced
absent

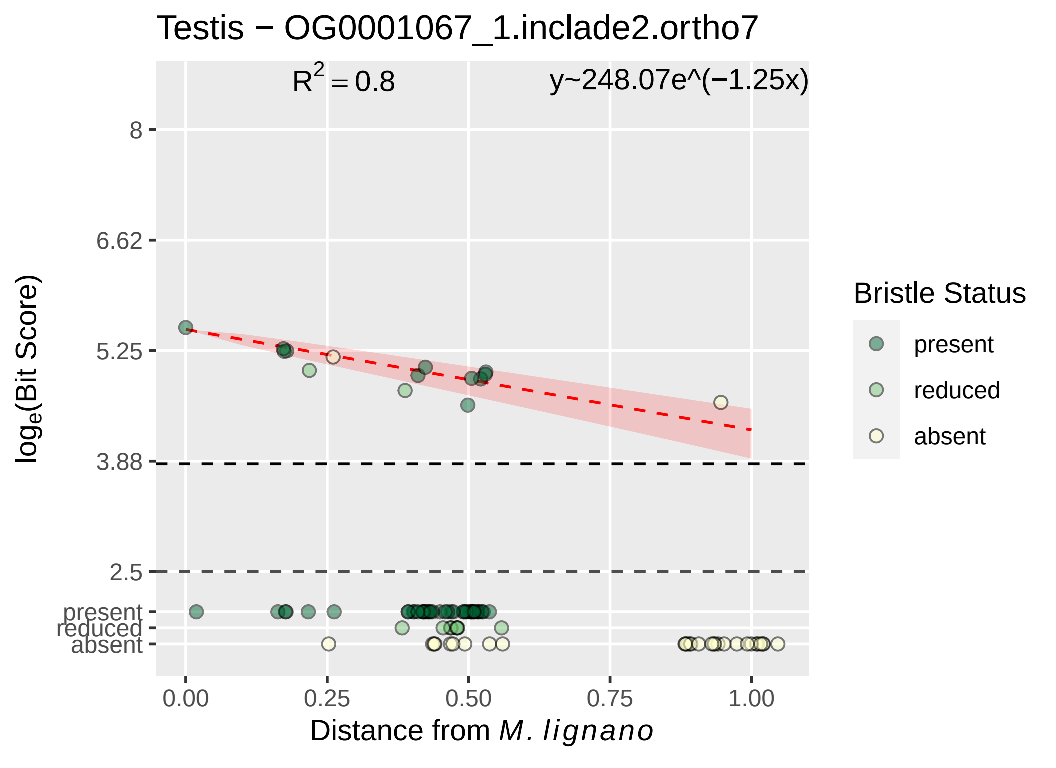

Testis − OG0001067_1.inclade2.ortho7
$R^2 = 0.8$
y~248.07e^(−1.25x)
8
6.62
5.25
3.88
2.5
present
reduced
absent
0.00
0.25
0.50
0.75
1.00
Distance from *M. lignano*
$log_e$(Bit Score)
Bristle Status
present
reduced
absent

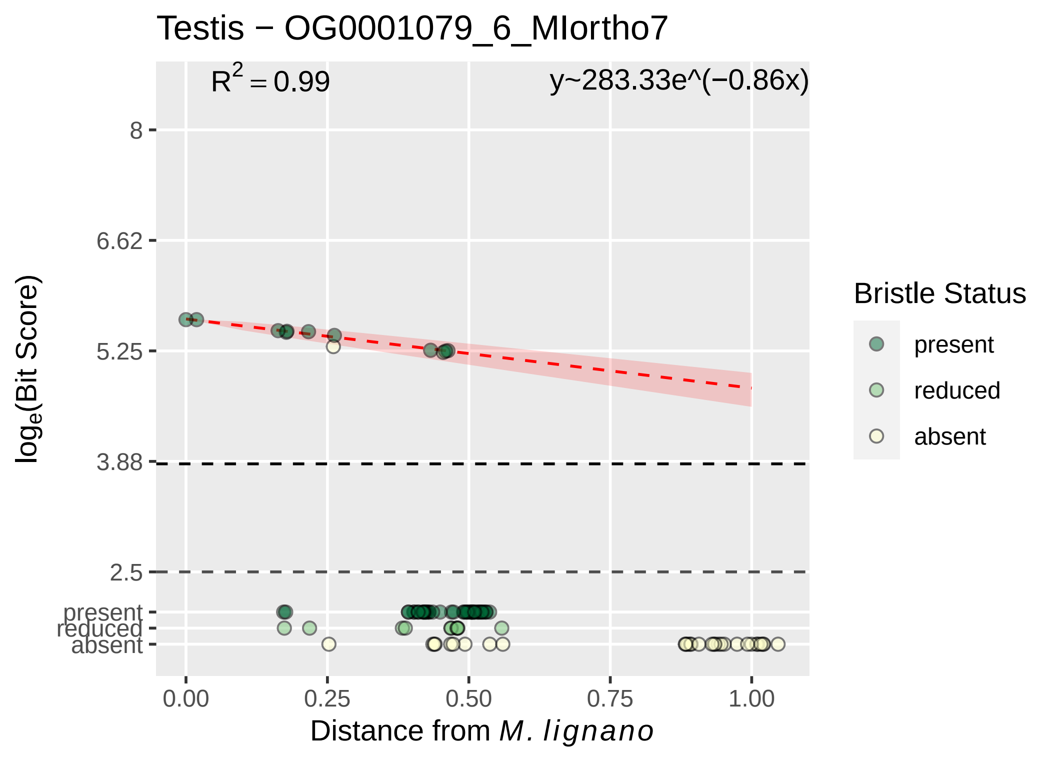
Testis − OG0001079_6_MIortho7
$R^2 = 0.99$
y~283.33e^(−0.86x)
8
6.62
5.25
3.88
2.5
present
reduced
absent
0.00
0.25
0.50
0.75
1.00
$\log_e$(Bit Score)
Distance from *M. lignano*
Bristle Status
present
reduced
absent

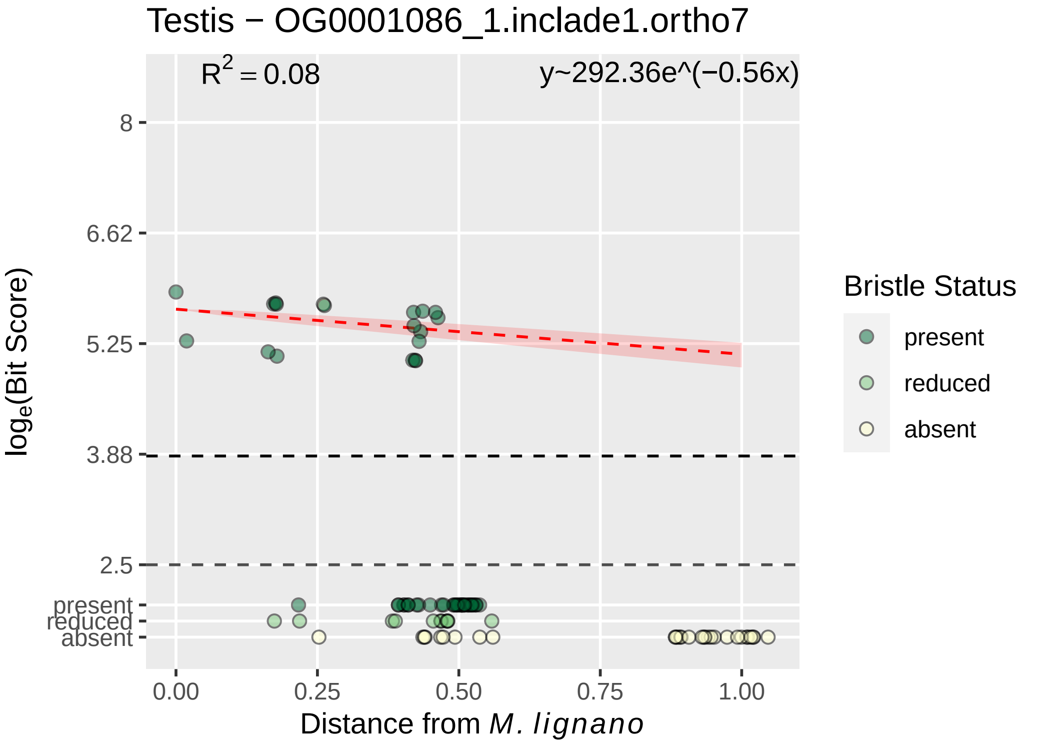

Testis − OG0001086_1.inclade1.ortho7
$R^2 = 0.08$
y~292.36e^(−0.56x)
8
6.62
5.25
3.88
2.5
present
reduced
absent
log$_e$(Bit Score)
0.00
0.25
0.50
0.75
1.00
Distance from *M. lignano*
Bristle Status
present
reduced
absent

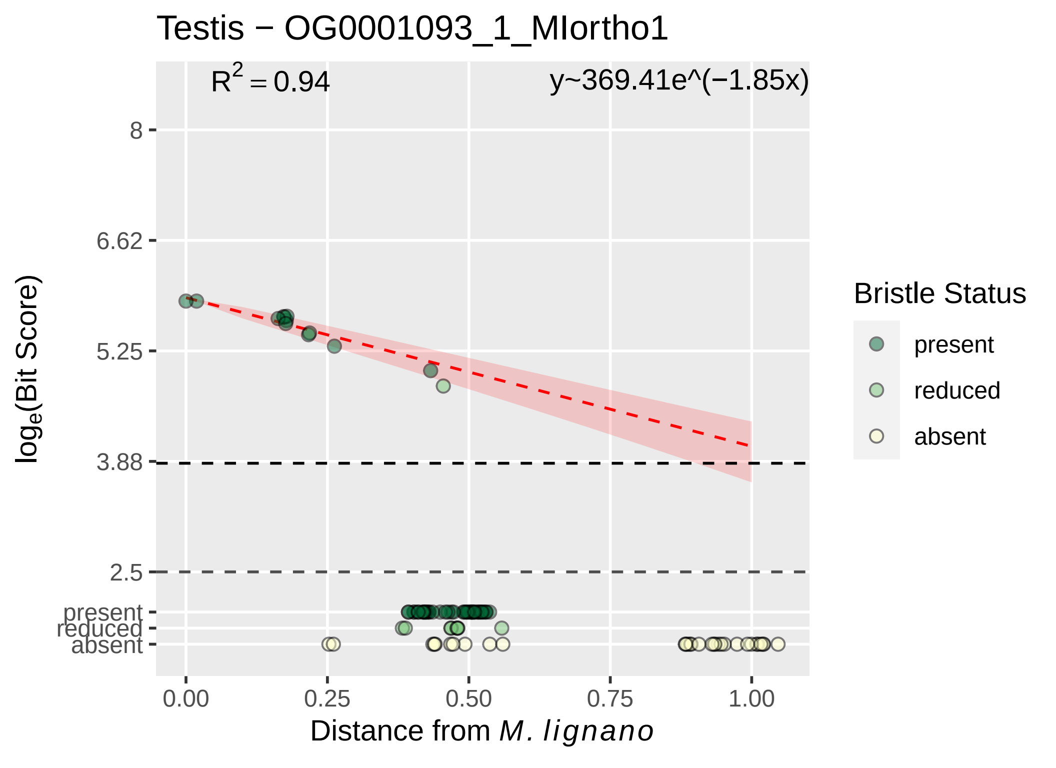

Testis − OG0001093_1_MIortho1
R2 = 0.94
y~369.41e^(−1.85x)
8
6.62
5.25
3.88
2.5
present
reduced
absent
log_e(Bit Score)
0.00
0.25
0.50
0.75
1.00
Distance from M. lignano
Bristle Status
present
reduced
absent

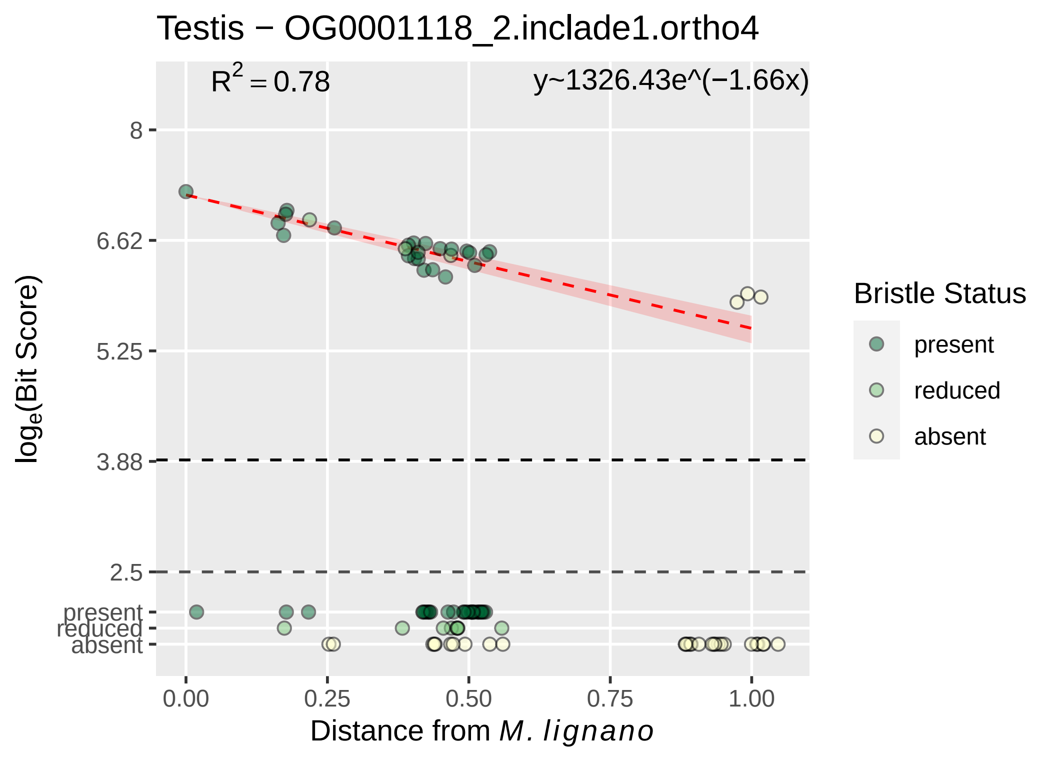
Testis − OG0001118_2.inclade1.ortho4
R2 = 0.78
y~1326.43e^(−1.66x)
8
6.62
5.25
3.88
2.5
present
reduced
absent
loge(Bit Score)
0.00
0.25
0.50
0.75
1.00
Distance from M. lignano
Bristle Status
present
reduced
absent

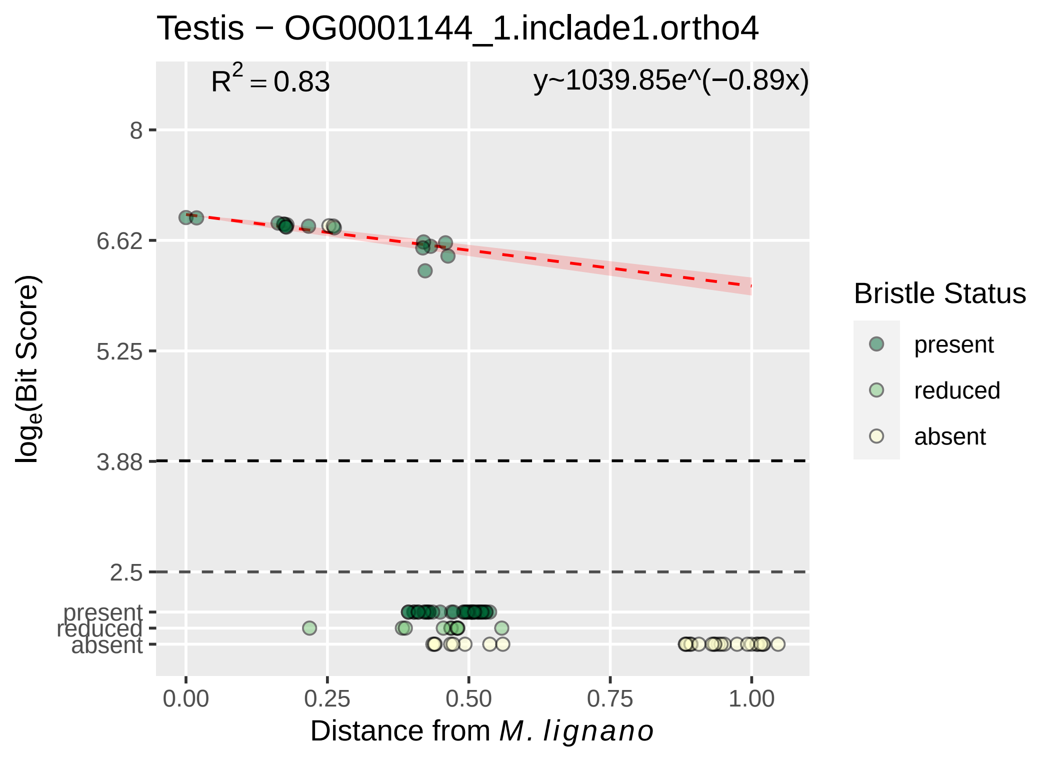

Testis − OG0001144_1.inclade1.ortho4
$R^2 = 0.83$
y~1039.85e^(−0.89x)
8
6.62
5.25
3.88
2.5
present
reduced
absent
$\log_e$(Bit Score)
0.00
0.25
0.50
0.75
1.00
Distance from *M. lignano*
Bristle Status
present
reduced
absent

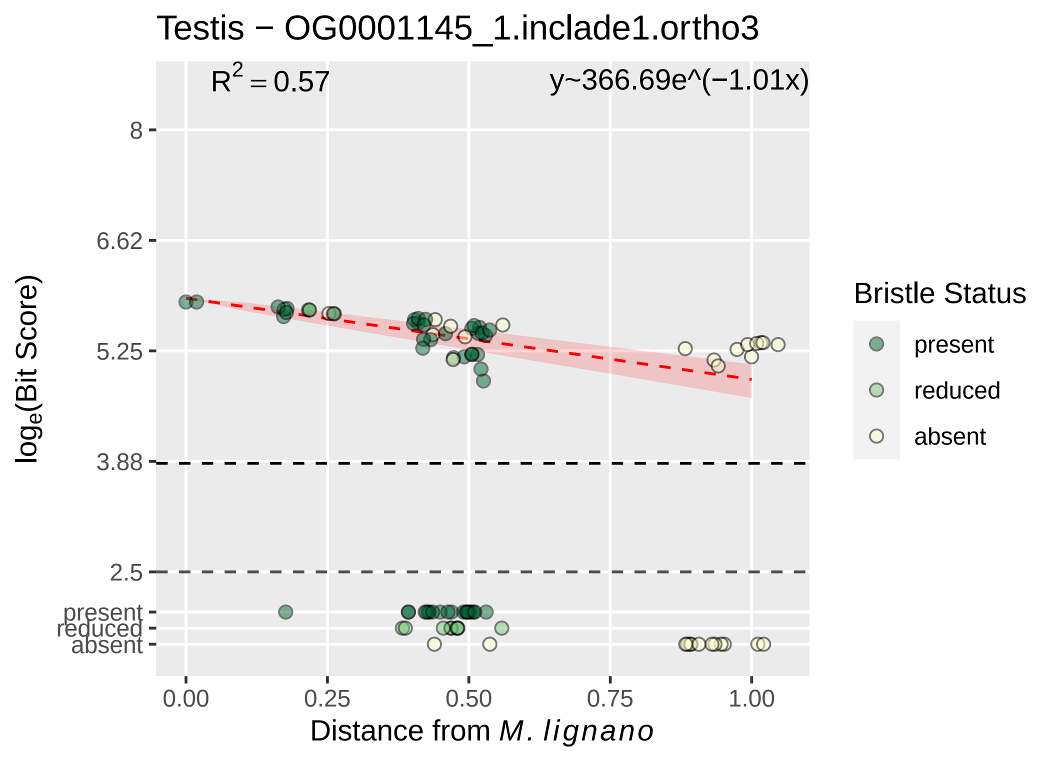

Testis − OG0001145_1.inclade1.ortho3
$R^2 = 0.57$
y~366.69e^(−1.01x)
8
6.62
5.25
3.88
2.5
present
reduced
absent
0.00
0.25
0.50
0.75
1.00
Distance from M. lignano
log_e(Bit Score)
Bristle Status
present
reduced
absent

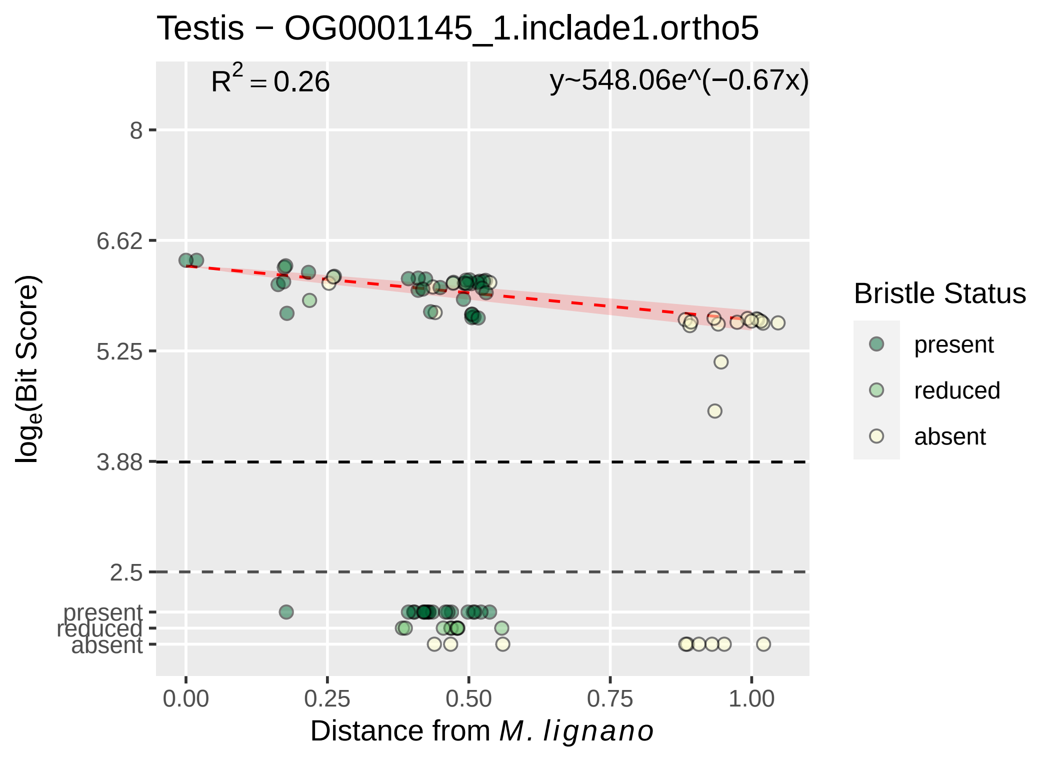

Testis − OG0001145_1.inclade1.ortho5
R^2 = 0.26
y~548.06e^(−0.67x)
8
6.62
5.25
3.88
2.5
present
reduced
absent
0.00
0.25
0.50
0.75
1.00
log_e(Bit Score)
Distance from M. lignano
Bristle Status
present
reduced
absent

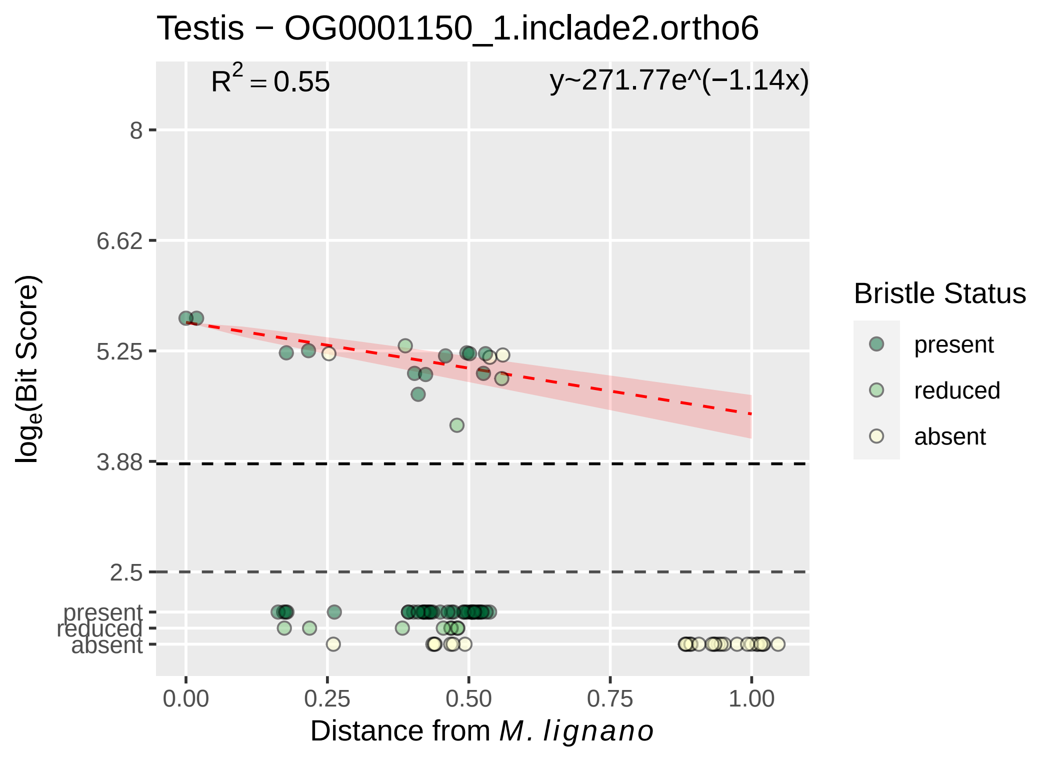

Testis − OG0001150_1.inclade2.ortho6
$R^2 = 0.55$
y~271.77e^(−1.14x)
8
6.62
5.25
3.88
2.5
present
reduced
absent
0.00
0.25
0.50
0.75
1.00
Distance from *M. lignano*
$\log_e$(Bit Score)
Bristle Status
present
reduced
absent

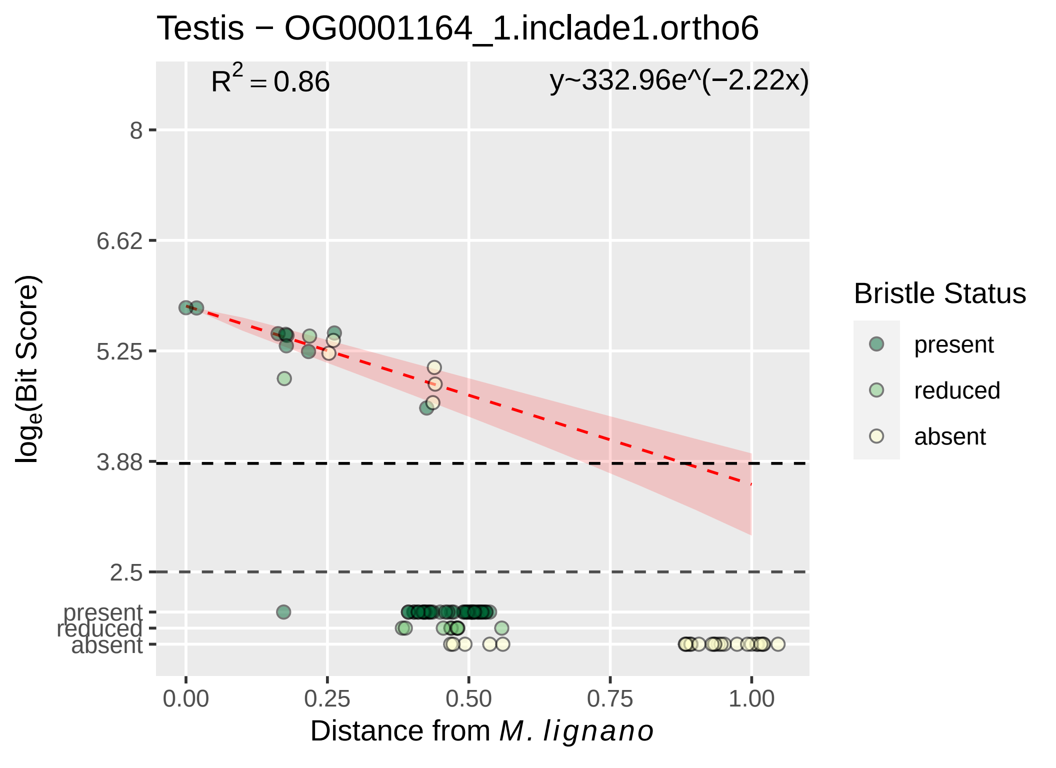

Testis − OG0001164_1.inclade1.ortho6
$R^2 = 0.86$
y~332.96e^(−2.22x)
8
6.62
5.25
3.88
2.5
present
reduced
absent
0.00
0.25
0.50
0.75
1.00
$log_e$(Bit Score)
Distance from *M. lignano*
Bristle Status
present
reduced
absent

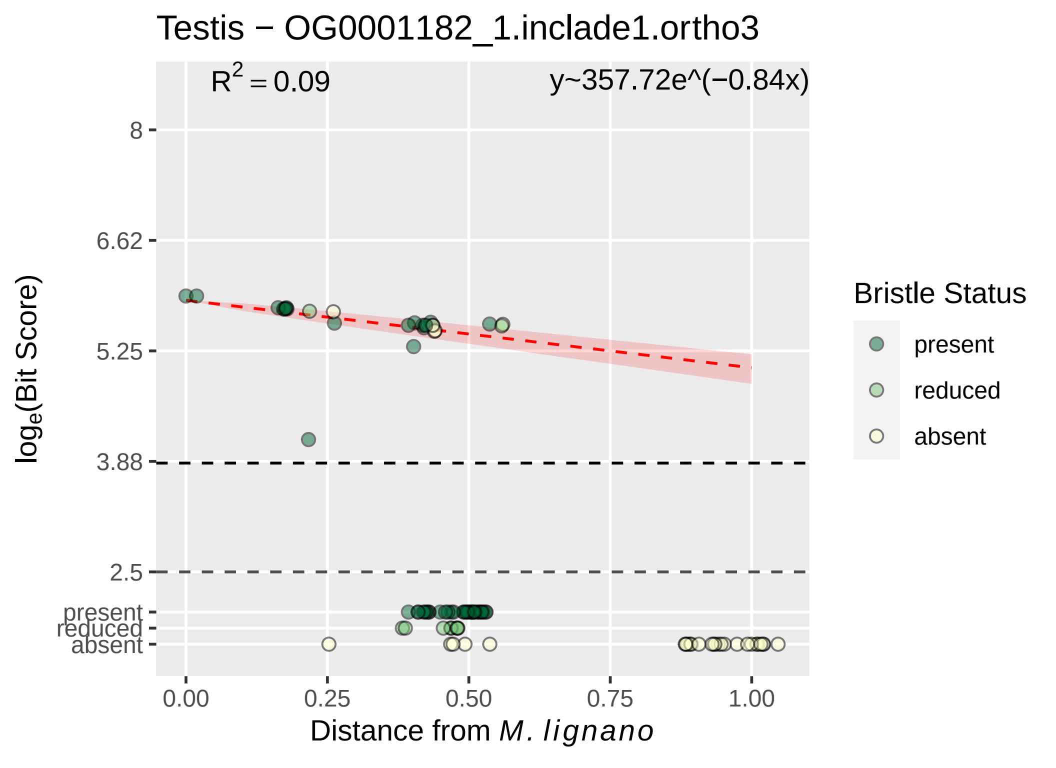

Testis − OG0001182_1.inclade1.ortho3
$R^2 = 0.09$
y~357.72e^(−0.84x)
8
6.62
5.25
3.88
2.5
present
reduced
absent
0.00
0.25
0.50
0.75
1.00
$\log_e$(Bit Score)
Distance from *M. lignano*
Bristle Status
present
reduced
absent

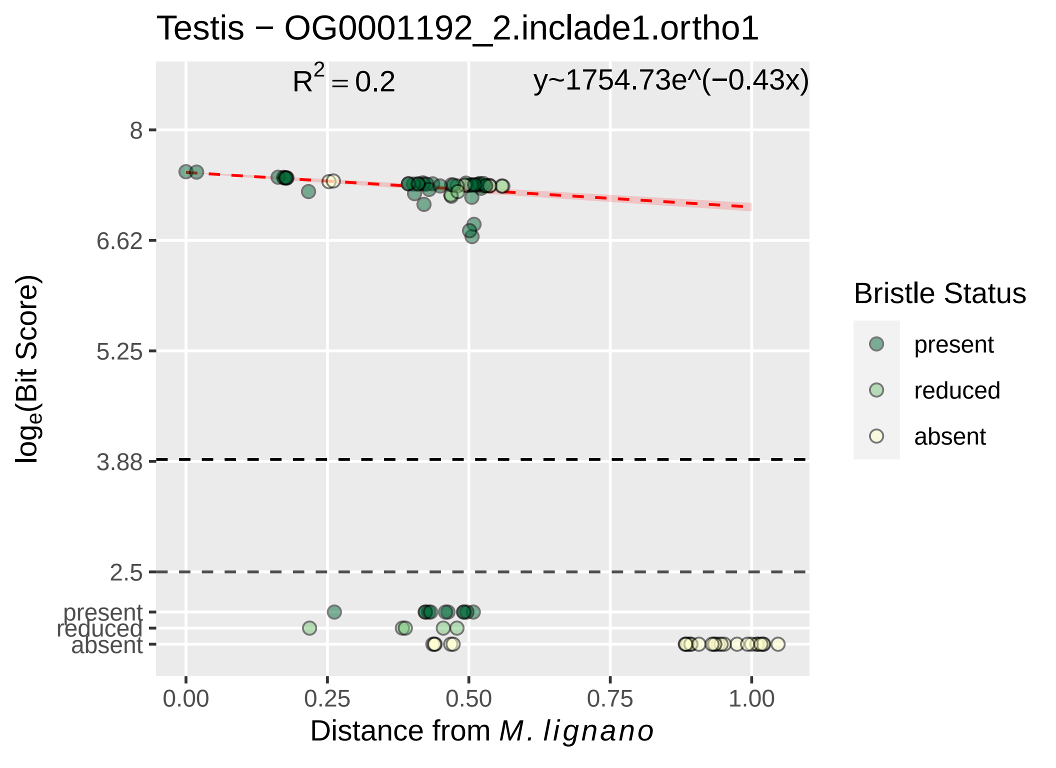

Testis − OG0001192_2.inclade1.ortho1
$R^2 = 0.2$
y~1754.73e^(−0.43x)
8
6.62
5.25
3.88
2.5
present
reduced
absent
0.00
0.25
0.50
0.75
1.00
$log_e$(Bit Score)
Distance from *M. lignano*
Bristle Status
present
reduced
absent

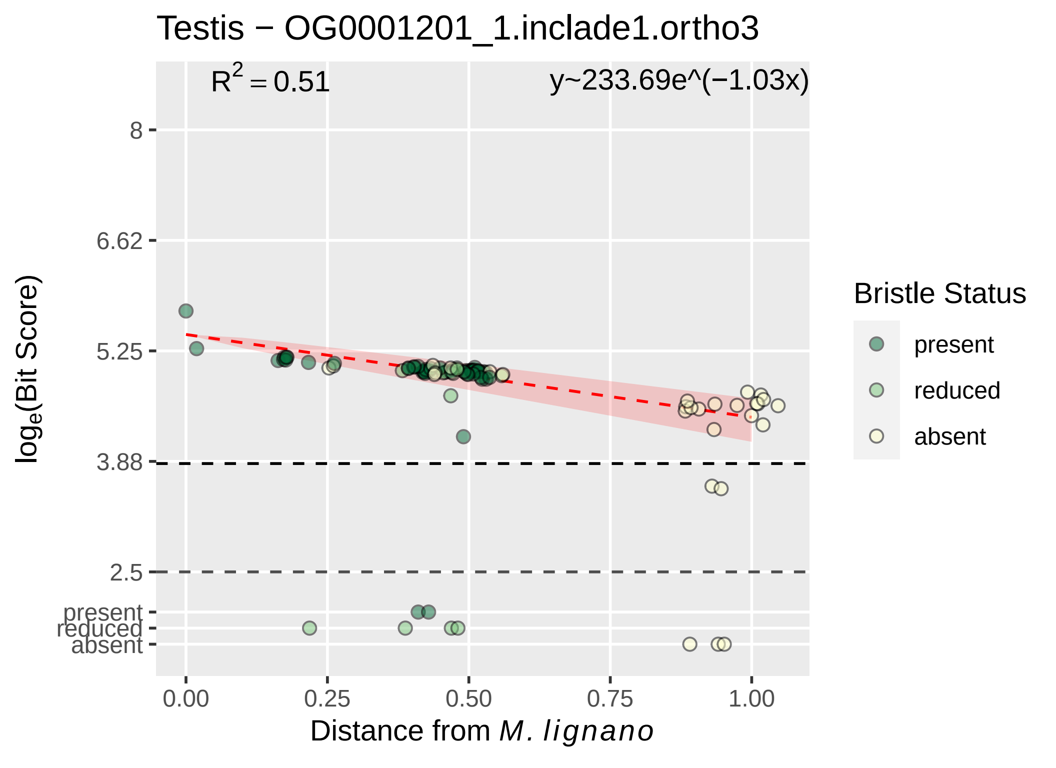

Testis − OG0001201_1.inclade1.ortho3
R^2 = 0.51
y~233.69e^(−1.03x)
8
6.62
5.25
3.88
2.5
present
reduced
absent
0.00
0.25
0.50
0.75
1.00
log_e(Bit Score)
Distance from M. lignano
Bristle Status
present
reduced
absent

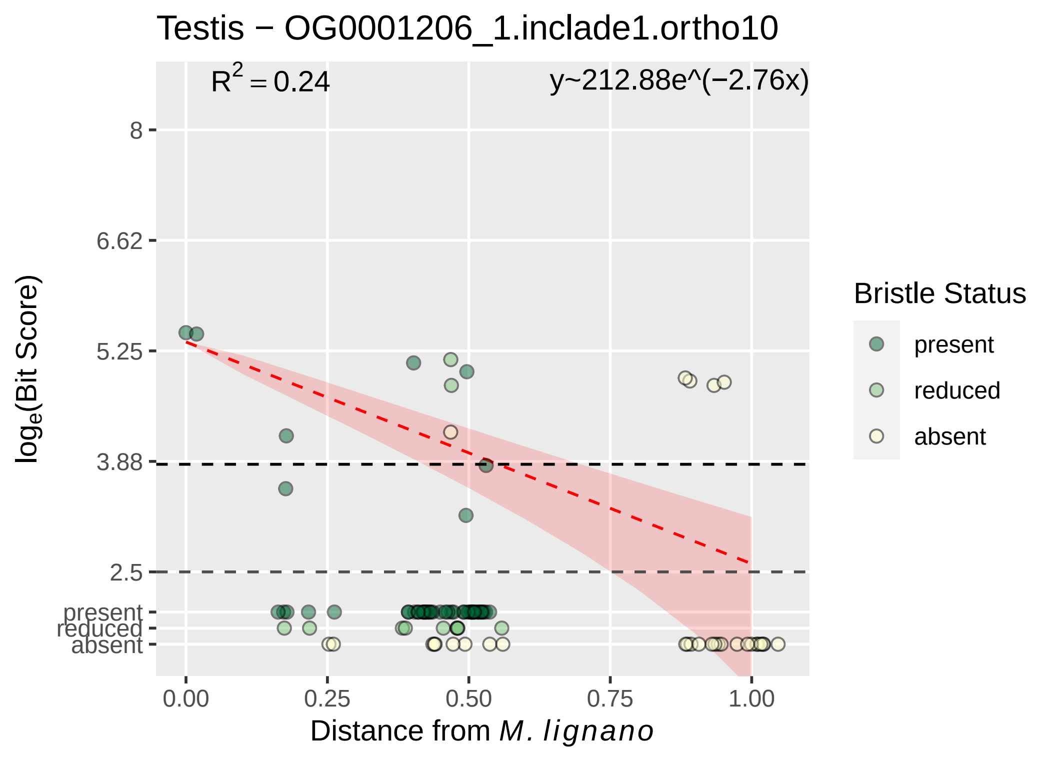

Testis − OG0001206_1.inclade1.ortho10
R^2 = 0.24
y~212.88e^(−2.76x)
8
6.62
5.25
3.88
2.5
present
reduced
absent
log_e(Bit Score)
0.00
0.25
0.50
0.75
1.00
Distance from M. lignano
Bristle Status
present
reduced
absent

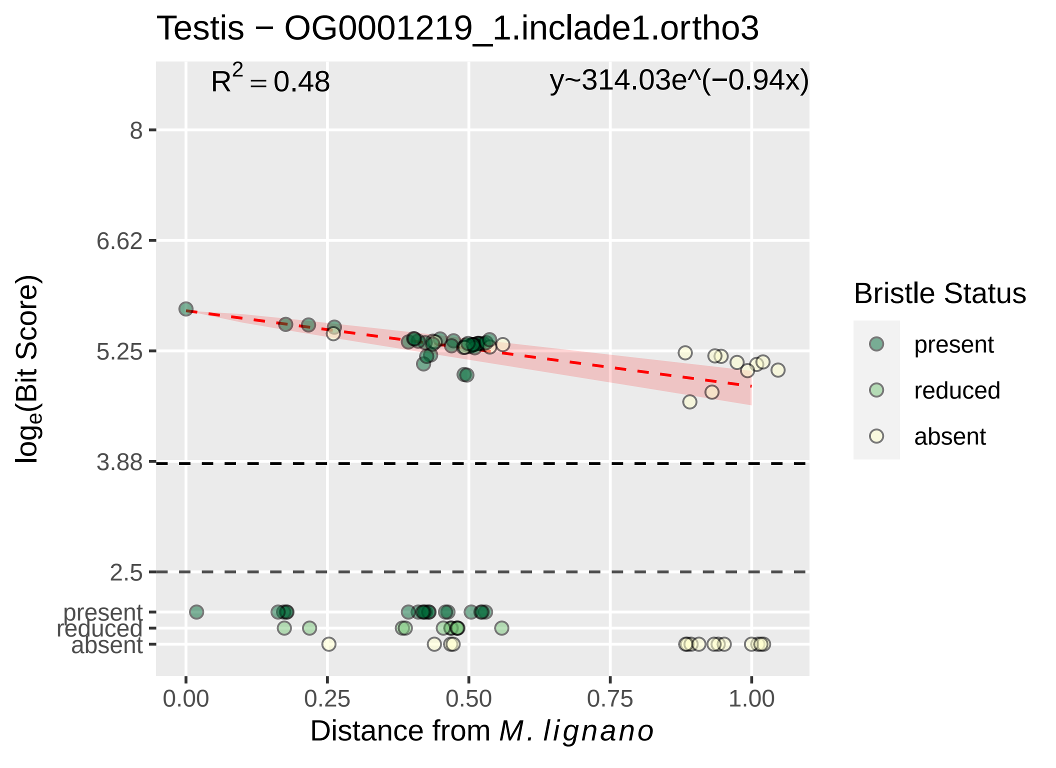

Testis − OG0001219_1.inclade1.ortho3
R^2 = 0.48
y~314.03e^(−0.94x)
8
6.62
5.25
3.88
2.5
present
reduced
absent
log_e(Bit Score)
0.00
0.25
0.50
0.75
1.00
Distance from M. lignano
Bristle Status
present
reduced
absent

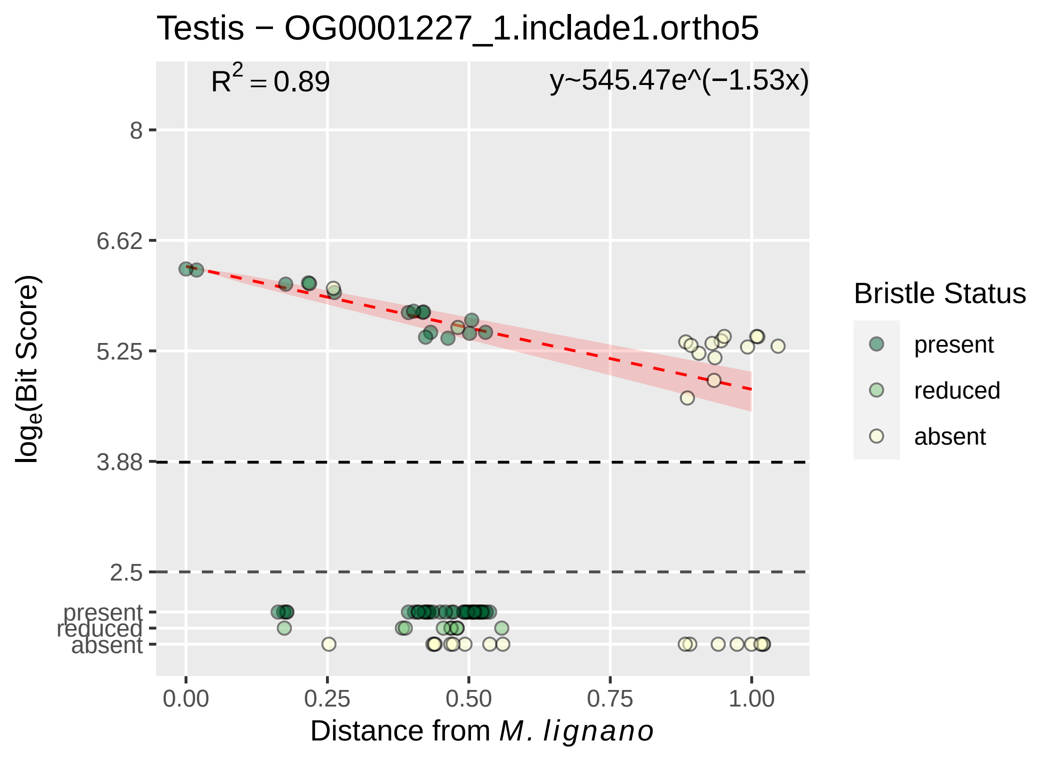

Testis − OG0001227_1.inclade1.ortho5
R² = 0.89
y~545.47e^(−1.53x)
8
6.62
5.25
3.88
2.5
present
reduced
absent
loge(Bit Score)
0.00
0.25
0.50
0.75
1.00
Distance from M. lignano
Bristle Status
present
reduced
absent

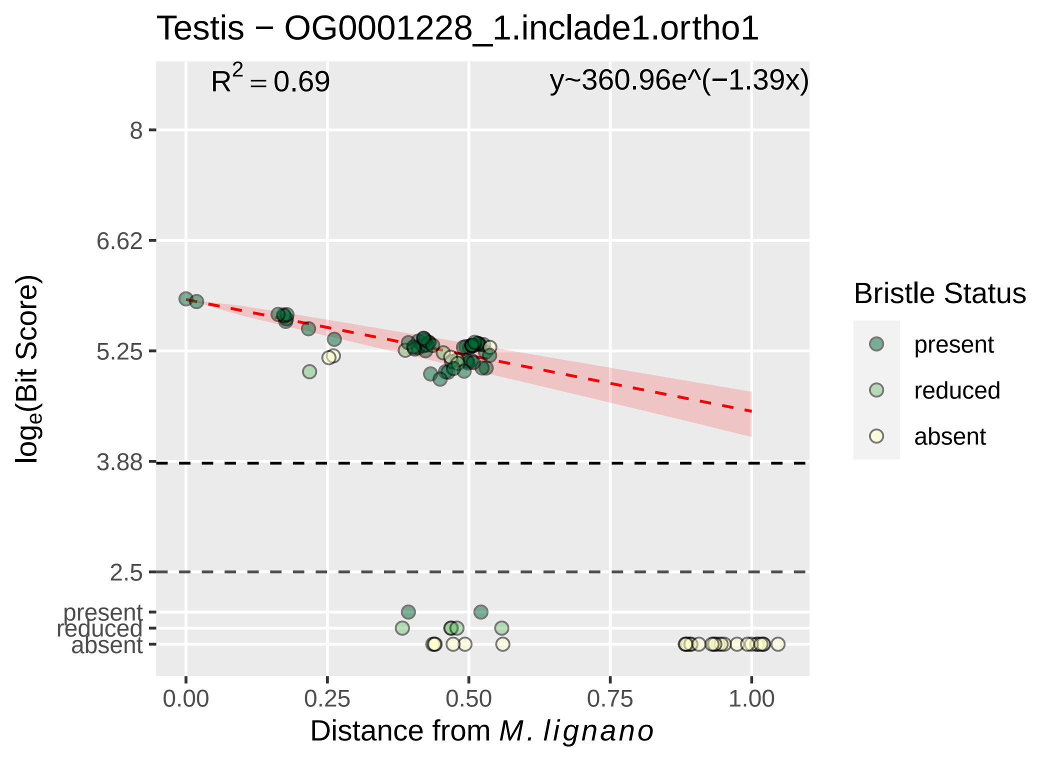
Testis − OG0001228_1.inclade1.ortho1
$R^2 = 0.69$
y~360.96e^(−1.39x)
8
6.62
5.25
3.88
2.5
present
reduced
absent
$\log_e$(Bit Score)
0.00
0.25
0.50
0.75
1.00
Distance from *M. lignano*
Bristle Status
present
reduced
absent

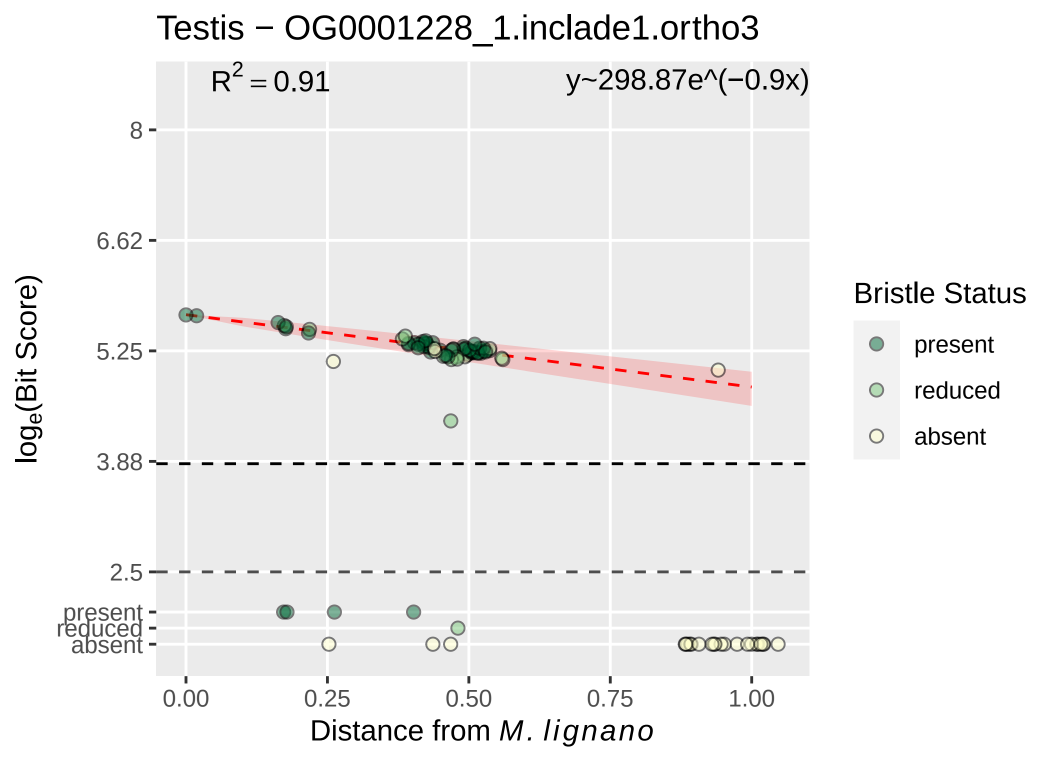

Testis − OG0001228_1.inclade1.ortho3
R^2 = 0.91
y~298.87e^(−0.9x)
8
6.62
5.25
3.88
2.5
present
reduced
absent
log_e(Bit Score)
0.00
0.25
0.50
0.75
1.00
Distance from M. lignano
Bristle Status
present
reduced
absent

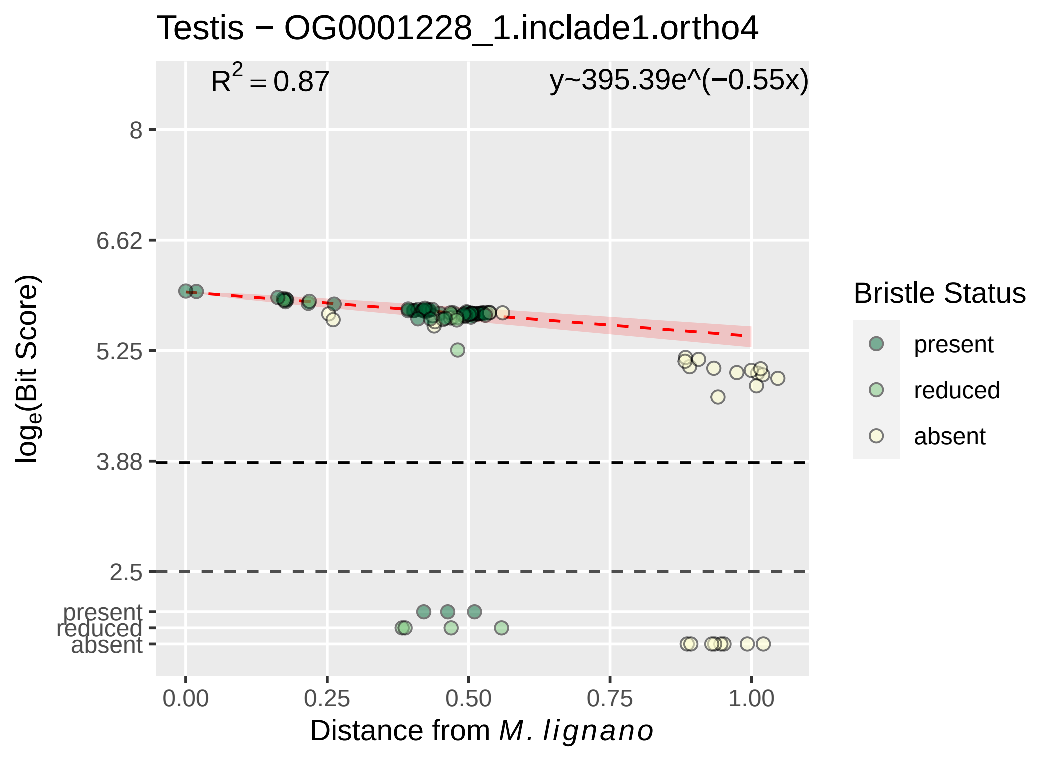

Testis − OG0001228_1.inclade1.ortho4
R2 = 0.87
y~395.39e^(−0.55x)
8
6.62
5.25
3.88
2.5
present
reduced
absent
loge(Bit Score)
0.00
0.25
0.50
0.75
1.00
Distance from M. lignano
Bristle Status
present
reduced
absent

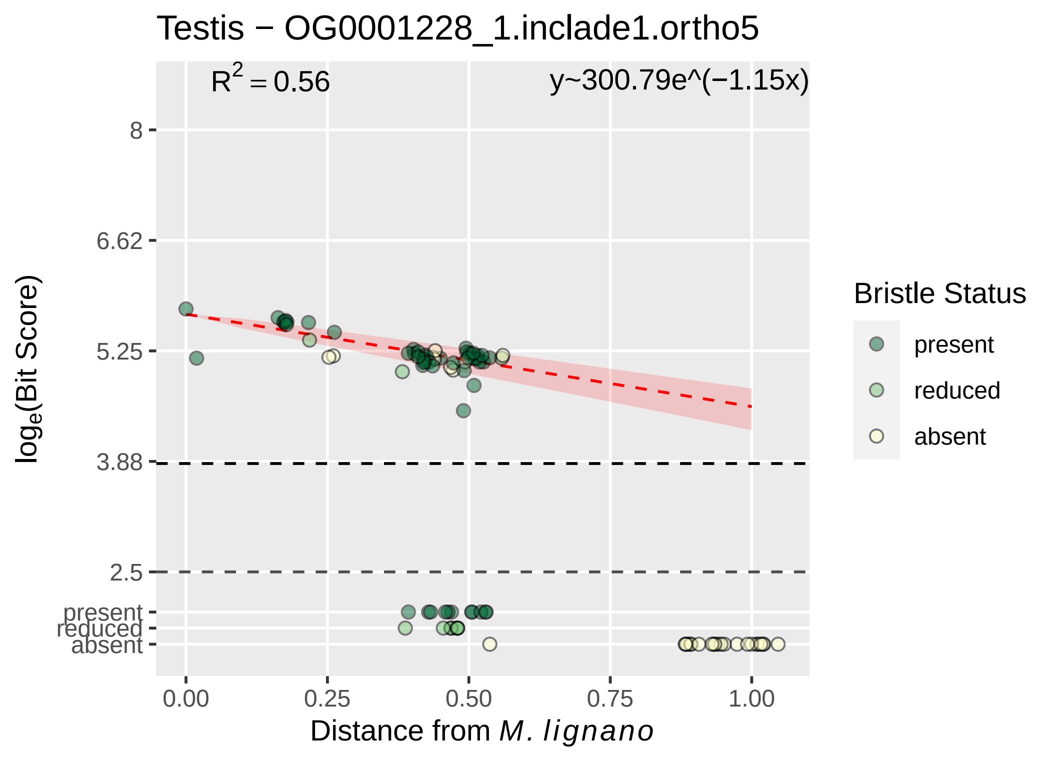

Testis − OG0001228_1.inclade1.ortho5
$R^2 = 0.56$
y~300.79e^(−1.15x)
8
6.62
5.25
3.88
2.5
present
reduced
absent
$\log_e$(Bit Score)
0.00
0.25
0.50
0.75
1.00
Distance from *M. lignano*
Bristle Status
present
reduced
absent

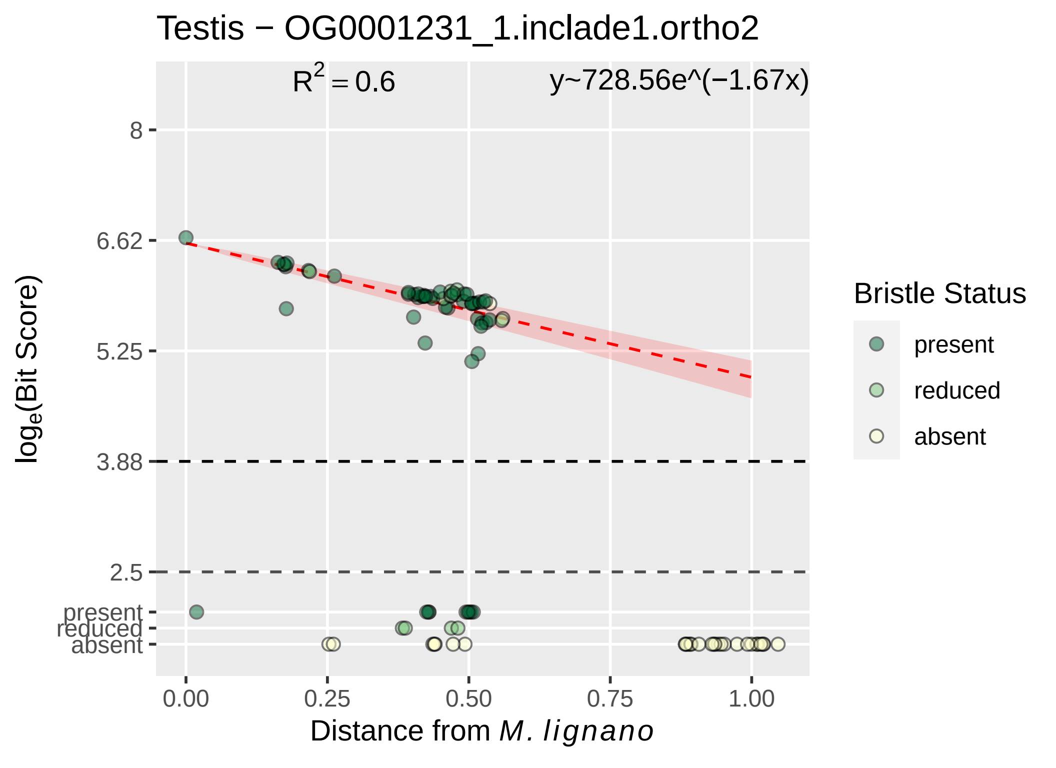
Testis − OG0001231_1.inclade1.ortho2
R^2 = 0.6
y~728.56e^(−1.67x)
8
6.62
5.25
3.88
2.5
present
reduced
absent
0.00
0.25
0.50
0.75
1.00
Distance from M. lignano
loge(Bit Score)
Bristle Status
present
reduced
absent

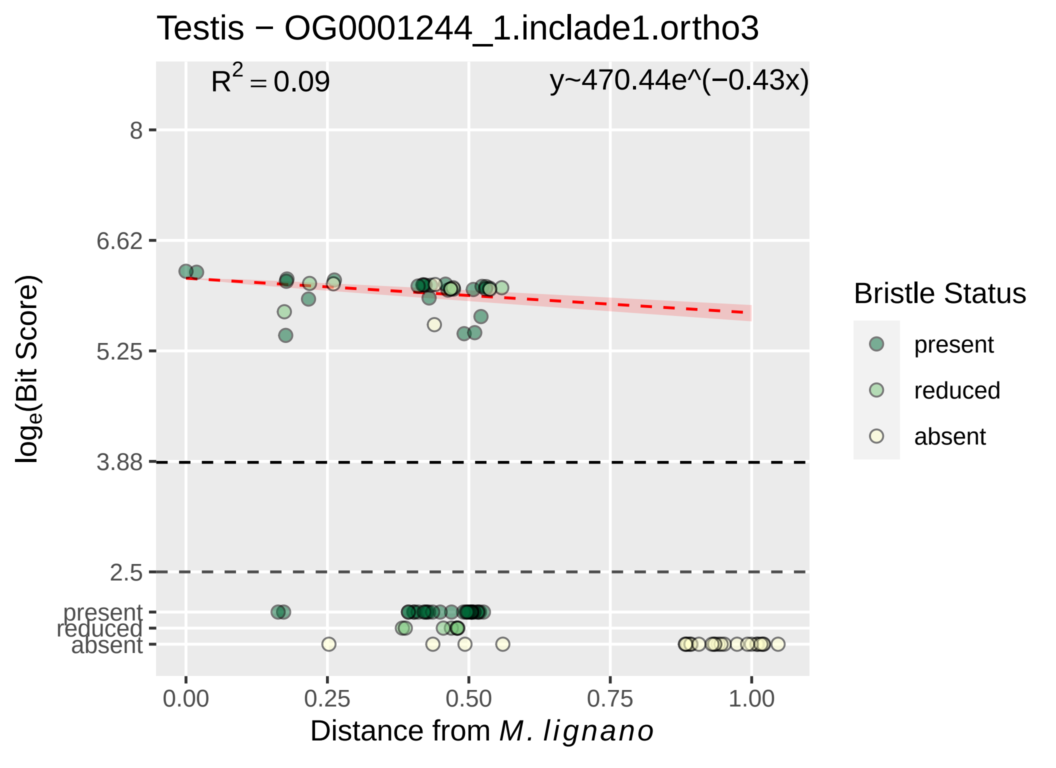
Testis − OG0001244_1.inclade1.ortho3
$R^2 = 0.09$
y~470.44e^(−0.43x)
8
6.62
5.25
3.88
2.5
present
reduced
absent
$\log_e$(Bit Score)
0.00
0.25
0.50
0.75
1.00
Distance from *M. lignano*
Bristle Status
present
reduced
absent

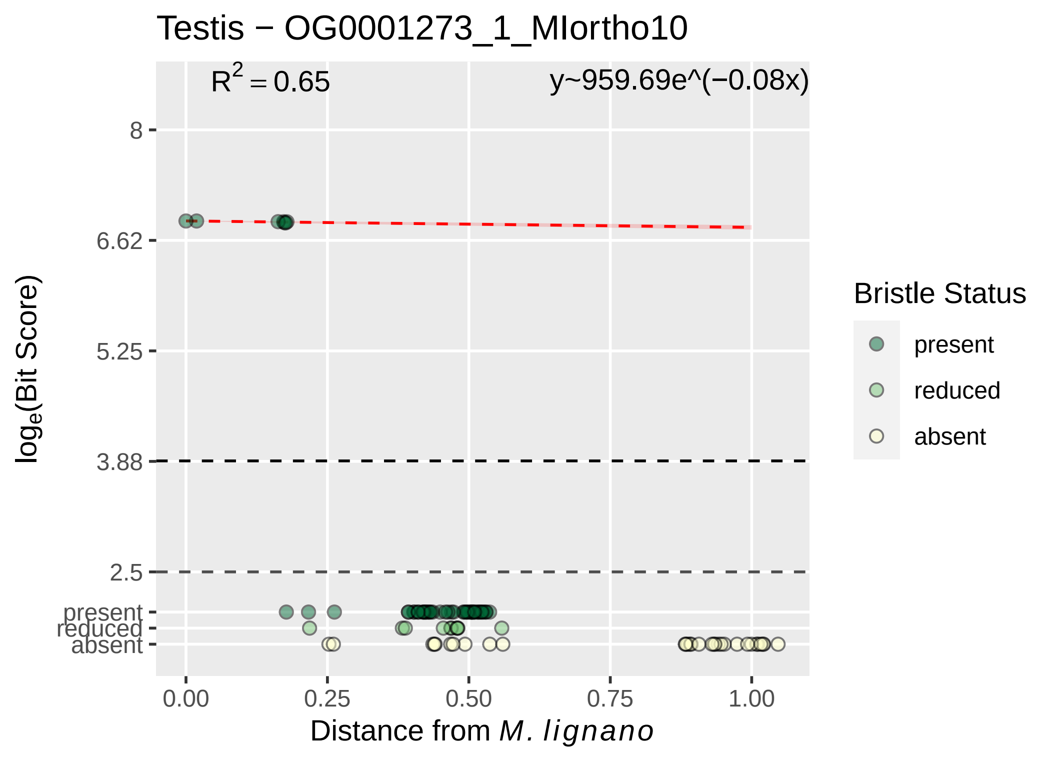

Testis − OG0001273_1_MIortho10
R^2 = 0.65
y~959.69e^(−0.08x)
8
6.62
5.25
3.88
2.5
present
reduced
absent
log_e(Bit Score)
0.00
0.25
0.50
0.75
1.00
Distance from M. lignano
Bristle Status
present
reduced
absent

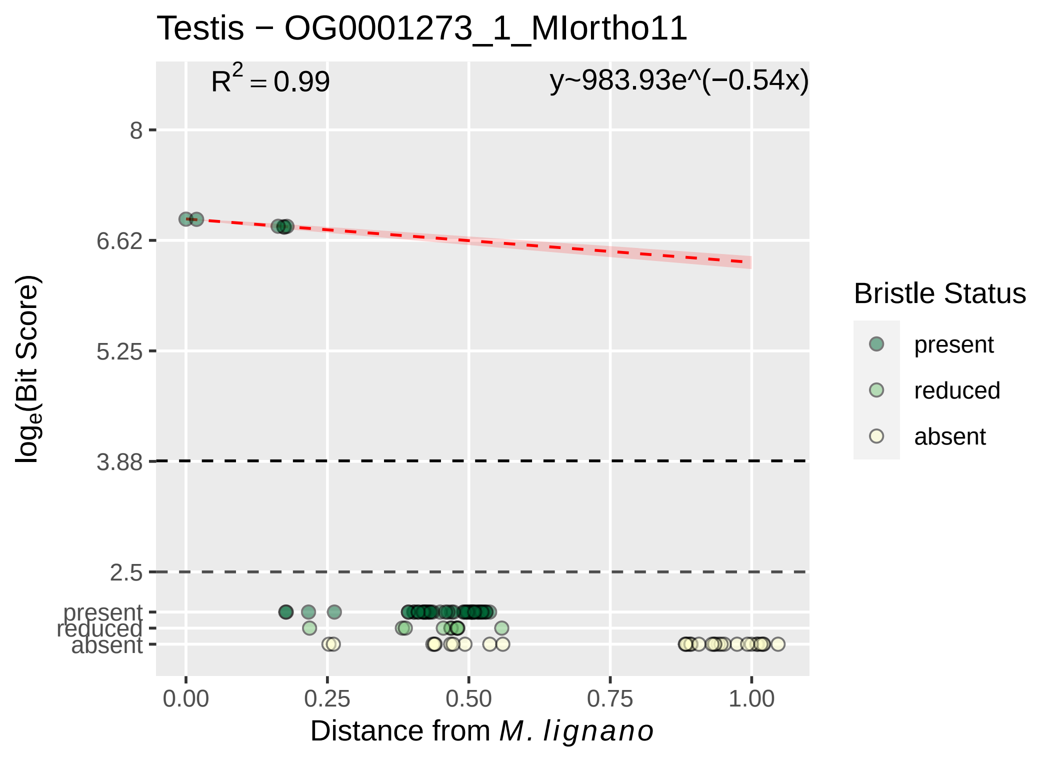
Testis − OG0001273_1_MIortho11
R² = 0.99
y~983.93e^(−0.54x)
8
6.62
5.25
3.88
2.5
present
reduced
absent
log_e(Bit Score)
0.00
0.25
0.50
0.75
1.00
Distance from M. lignano
Bristle Status
present
reduced
absent

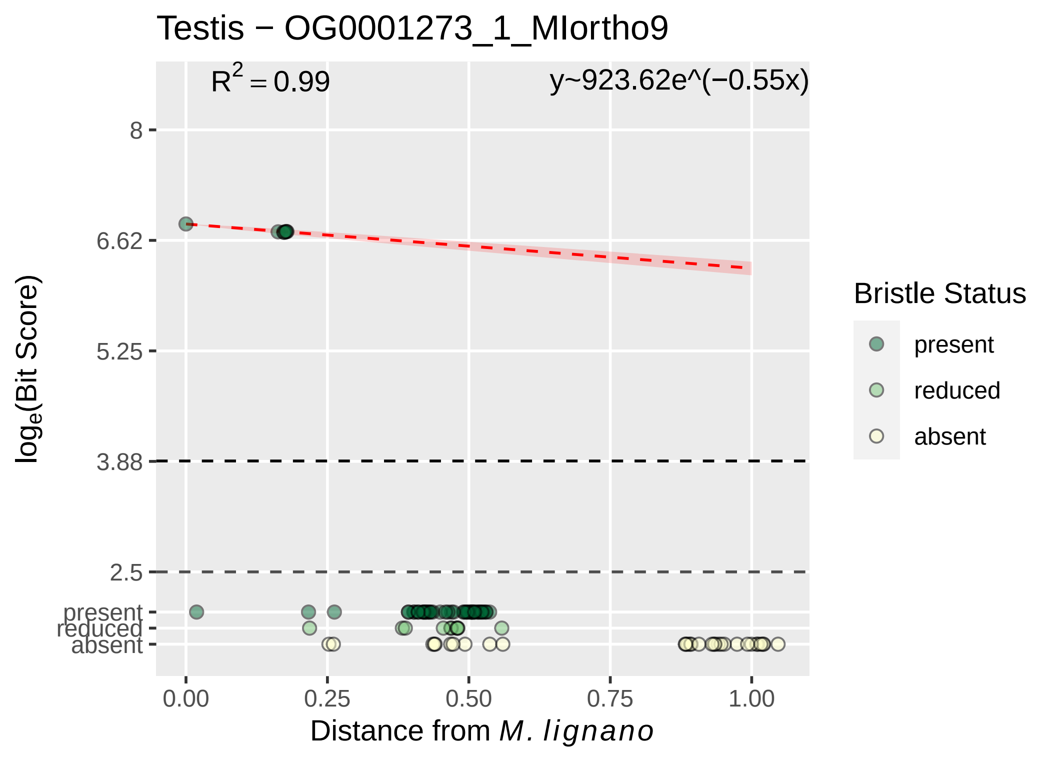

Testis − OG0001273_1_MIortho9
R^2 = 0.99
y~923.62e^(−0.55x)
8
6.62
5.25
3.88
2.5
present
reduced
absent
loge(Bit Score)
0.00
0.25
0.50
0.75
1.00
Distance from M. lignano
Bristle Status
present
reduced
absent

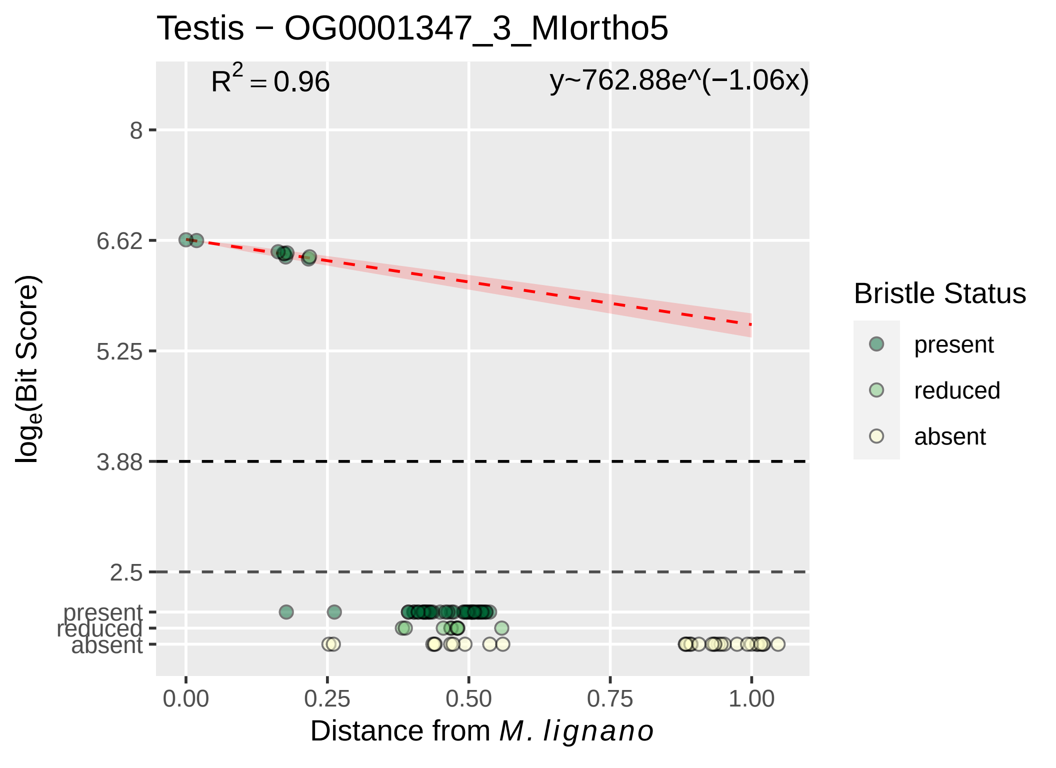

Testis − OG0001347_3_MIortho5
$R^2 = 0.96$
y~762.88e^(−1.06x)
8
6.62
5.25
3.88
2.5
present
reduced
absent
0.00
0.25
0.50
0.75
1.00
$\log_e$(Bit Score)
Distance from *M. lignano*
Bristle Status
present
reduced
absent

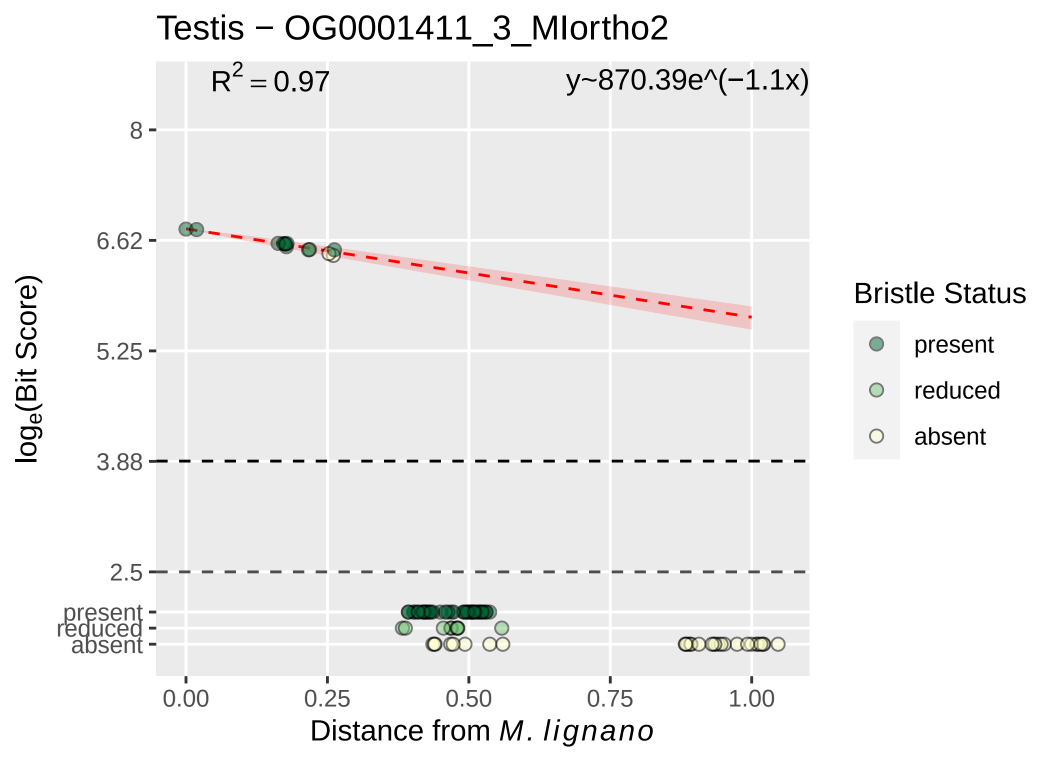

Testis − OG0001411_3_MIortho2
R^2 = 0.97
y~870.39e^(−1.1x)
8
6.62
5.25
3.88
2.5
present
reduced
absent
0.00
0.25
0.50
0.75
1.00
Distance from M. lignano
log_e(Bit Score)
Bristle Status
present
reduced
absent

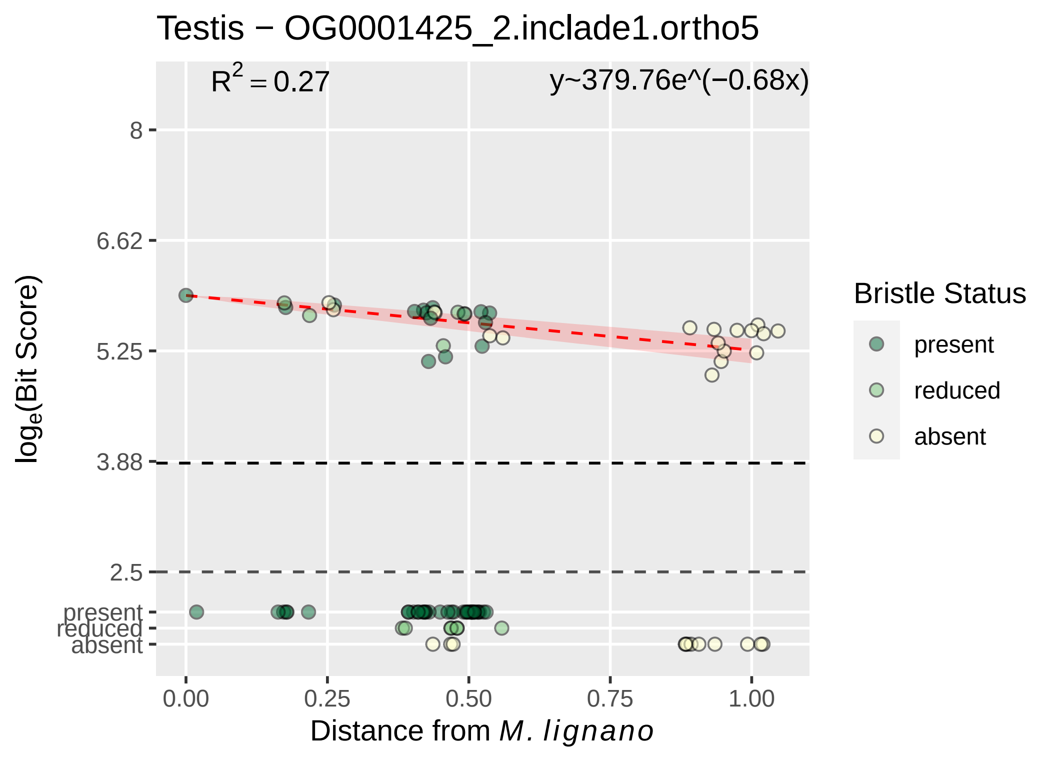

Testis − OG0001425_2.inclade1.ortho5
$R^2 = 0.27$
y~379.76e^(−0.68x)
8
6.62
5.25
3.88
2.5
present
reduced
absent
0.00
0.25
0.50
0.75
1.00
Distance from *M. lignano*
$\log_e$(Bit Score)
Bristle Status
present
reduced
absent

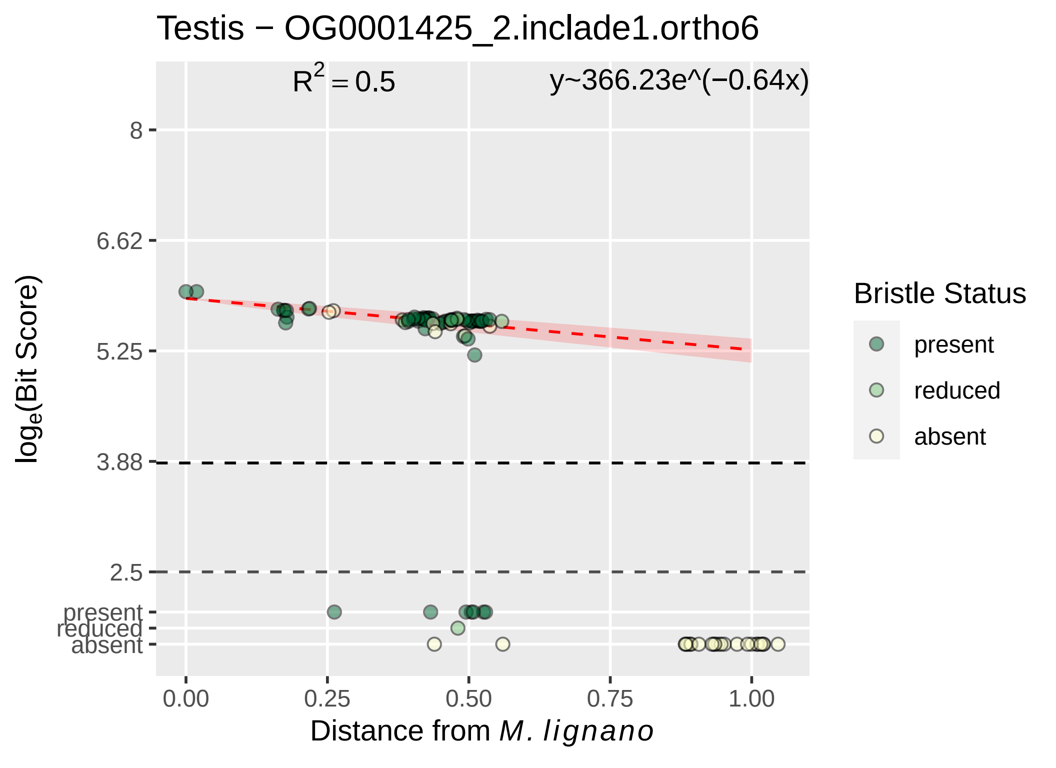

Testis − OG0001425_2.inclade1.ortho6
$R^2 = 0.5$
y~366.23e^(−0.64x)
8
6.62
5.25
3.88
2.5
present
reduced
absent
0.00
0.25
0.50
0.75
1.00
$\log_e$(Bit Score)
Distance from *M. lignano*
Bristle Status
present
reduced
absent

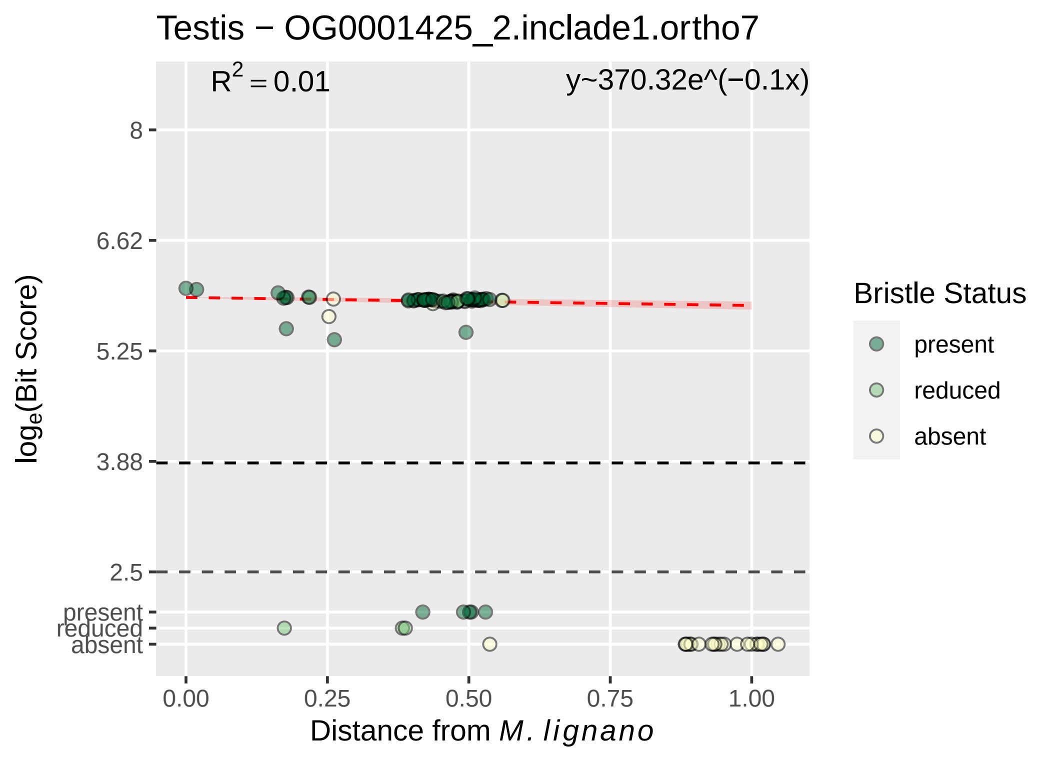

Testis − OG0001425_2.inclade1.ortho7
R^2 = 0.01
y~370.32e^(−0.1x)
8
6.62
5.25
3.88
2.5
present
reduced
absent
log_e(Bit Score)
0.00
0.25
0.50
0.75
1.00
Distance from M. lignano
Bristle Status
present
reduced
absent

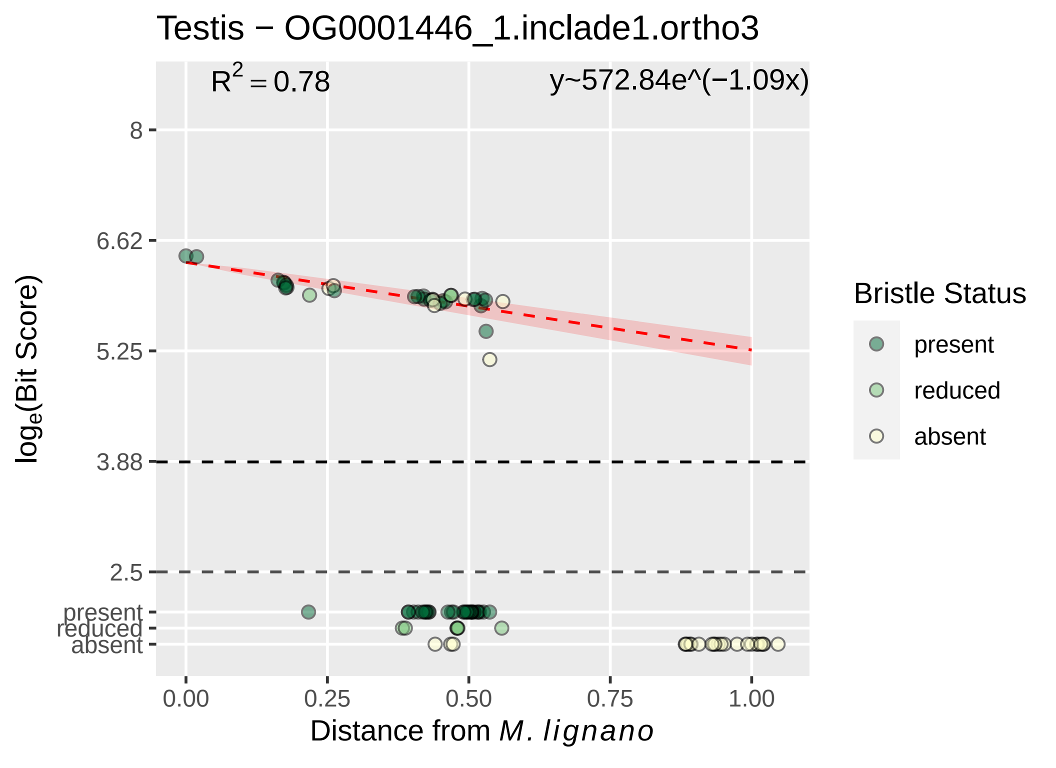

Testis − OG0001446_1.inclade1.ortho3
R^2 = 0.78
y~572.84e^(−1.09x)
8
6.62
5.25
3.88
2.5
present
reduced
absent
log_e(Bit Score)
0.00
0.25
0.50
0.75
1.00
Distance from M. lignano
Bristle Status
present
reduced
absent

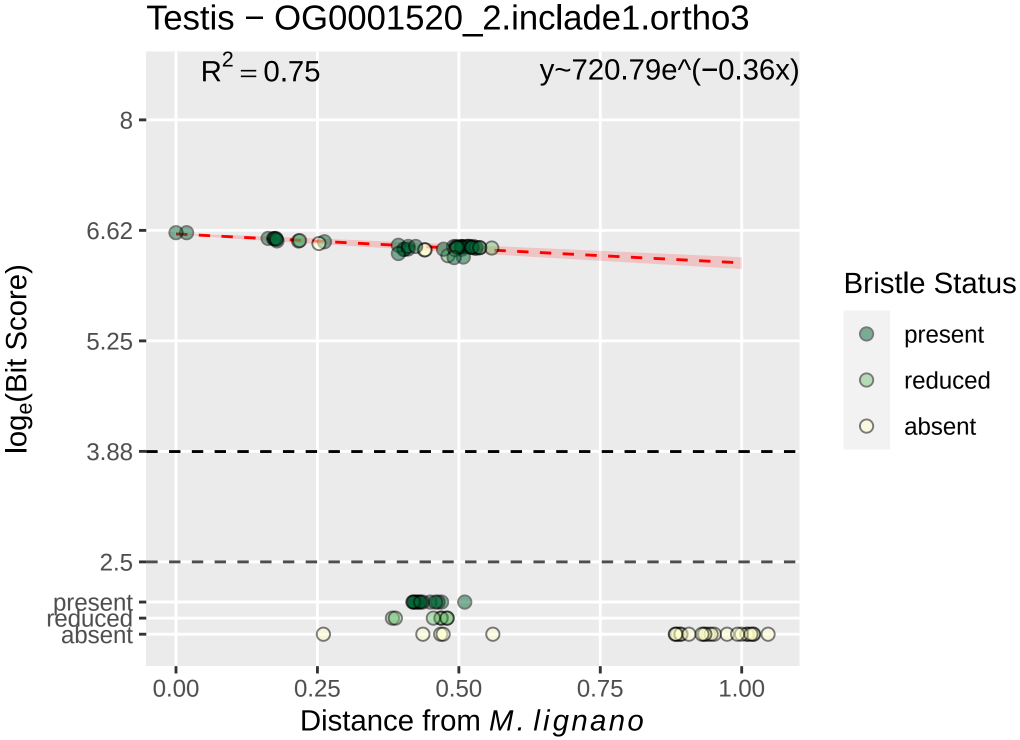

Testis − OG0001520_2.inclade1.ortho3
R^2 = 0.75
y~720.79e^(−0.36x)
8
6.62
5.25
3.88
2.5
present
reduced
absent
0.00
0.25
0.50
0.75
1.00
Distance from M. lignano
log_e(Bit Score)
Bristle Status
present
reduced
absent

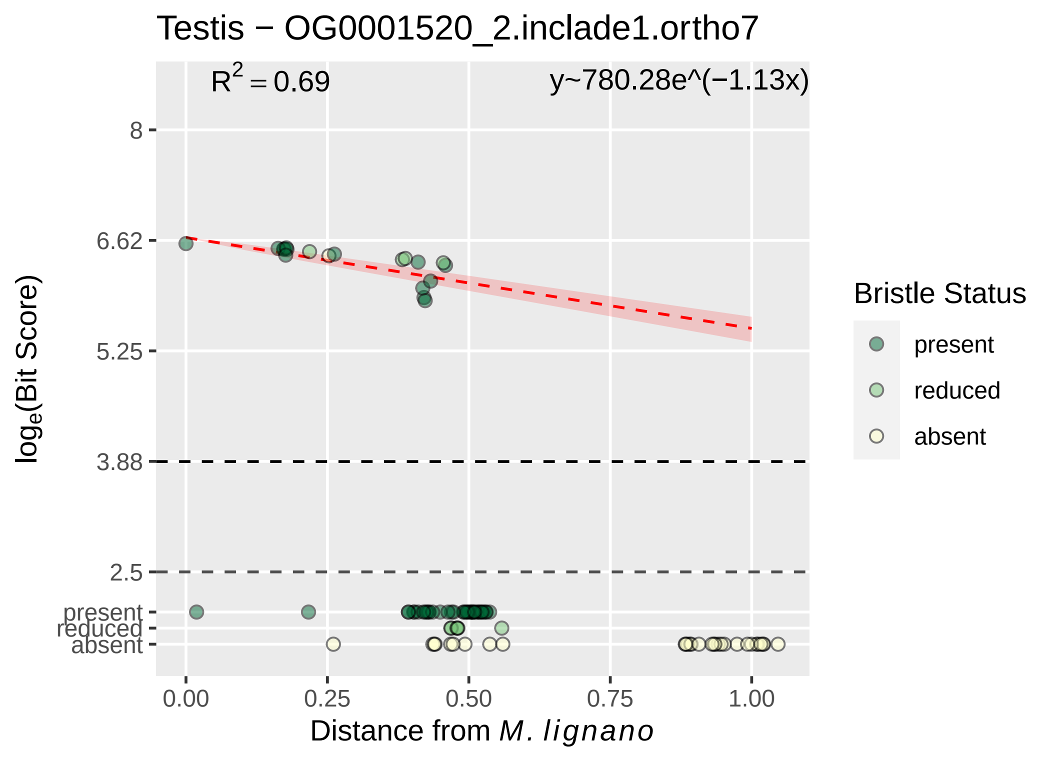

Testis − OG0001520_2.inclade1.ortho7
R² = 0.69
y~780.28e^(−1.13x)
8
6.62
5.25
3.88
2.5
present
reduced
absent
log_e(Bit Score)
0.00
0.25
0.50
0.75
1.00
Distance from M. lignano
Bristle Status
present
reduced
absent

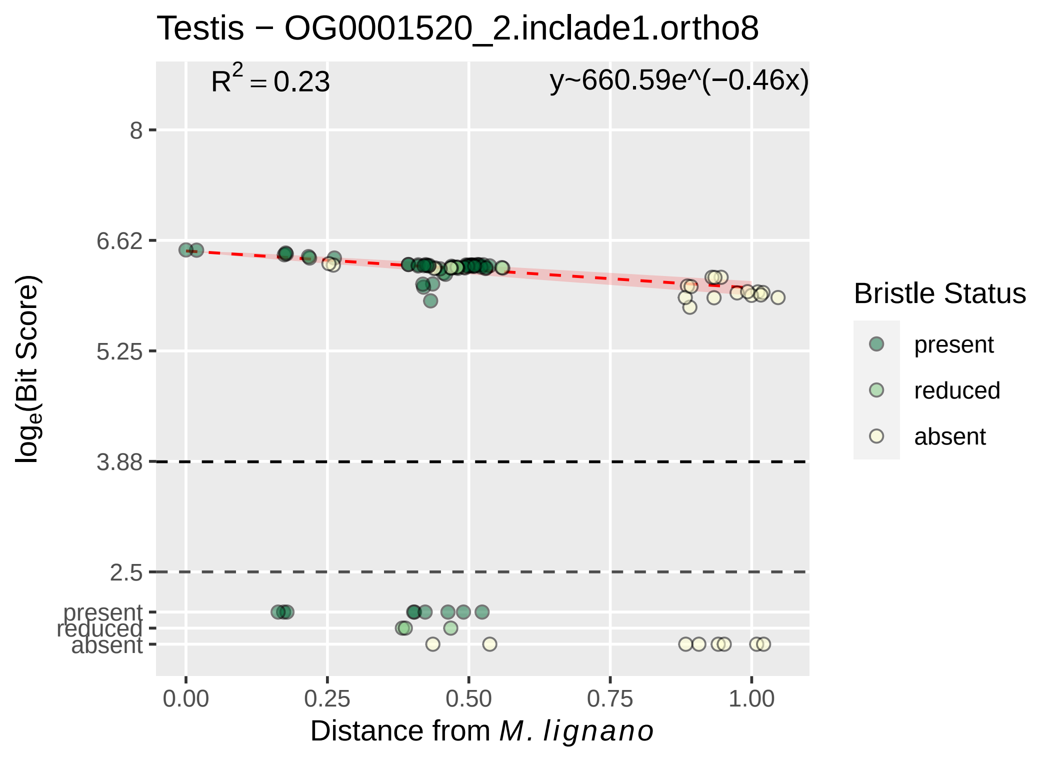

Testis − OG0001520_2.inclade1.ortho8
$R^2 = 0.23$
y~660.59e^(−0.46x)
8
6.62
5.25
3.88
2.5
present
reduced
absent
log_e(Bit Score)
0.00
0.25
0.50
0.75
1.00
Distance from M. lignano
Bristle Status
present
reduced
absent

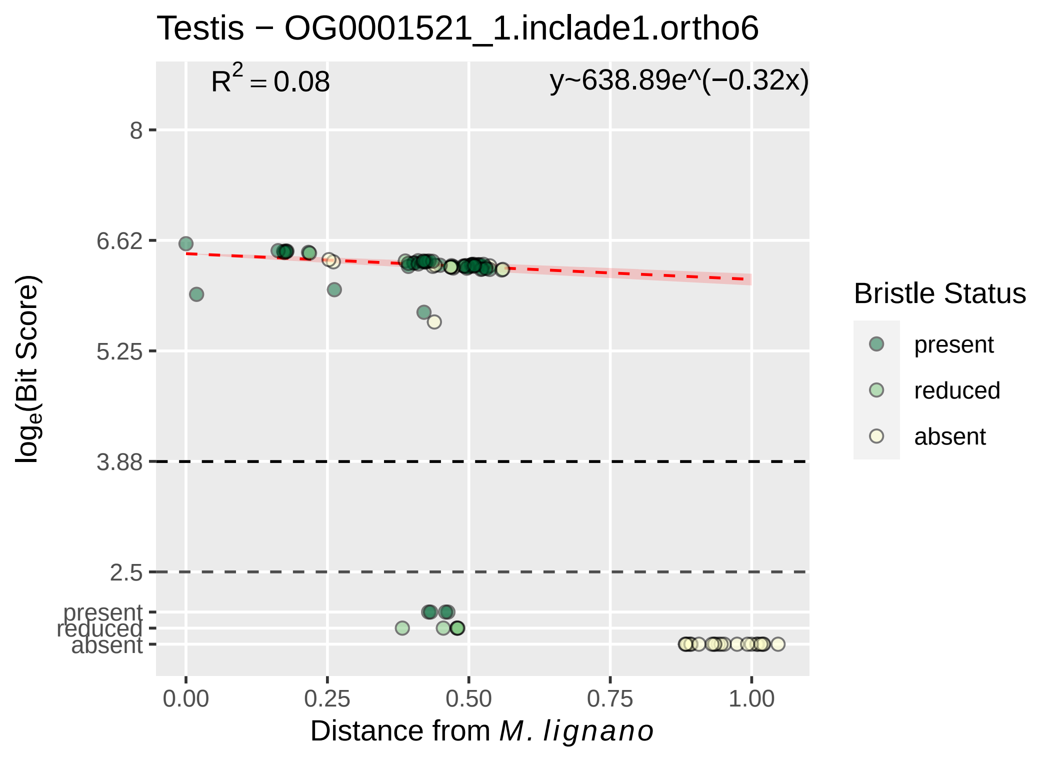

Testis − OG0001521_1.inclade1.ortho6
R^2 = 0.08
y~638.89e^(−0.32x)
8
6.62
5.25
3.88
2.5
present
reduced
absent
loge(Bit Score)
0.00
0.25
0.50
0.75
1.00
Distance from M. lignano
Bristle Status
present
reduced
absent

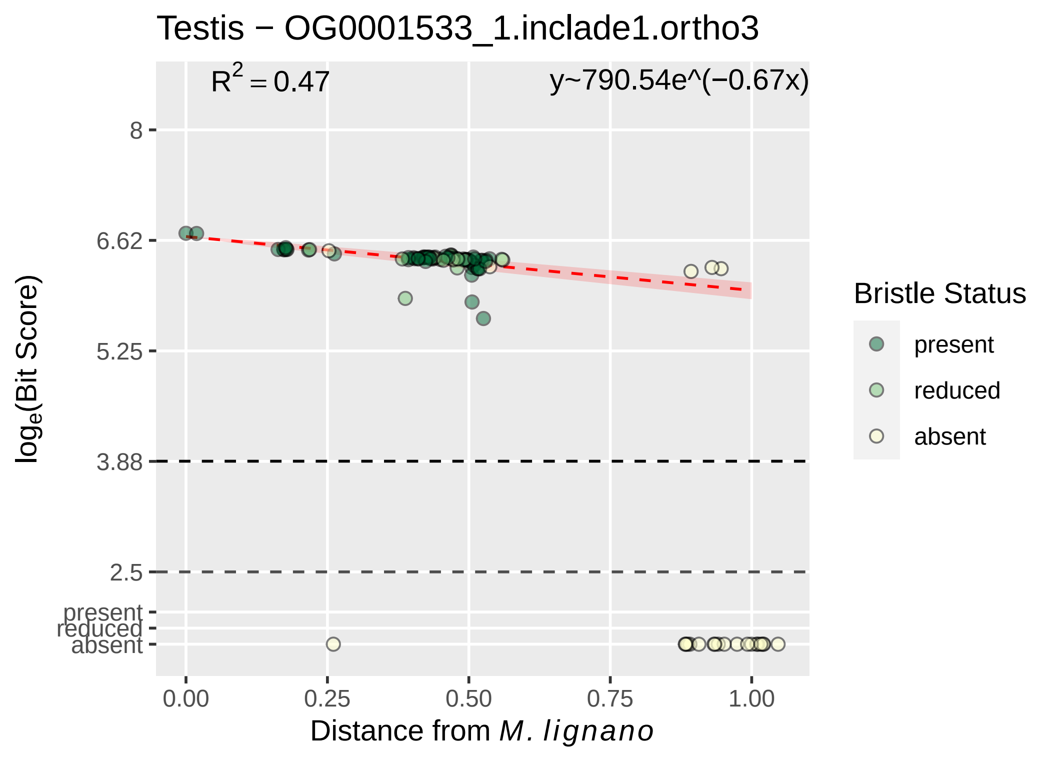
Testis − OG0001533_1.inclade1.ortho3
R^2 = 0.47
y~790.54e^(−0.67x)
8
6.62
5.25
3.88
2.5
present
reduced
absent
log_e(Bit Score)
0.00
0.25
0.50
0.75
1.00
Distance from M. lignano
Bristle Status
present
reduced
absent

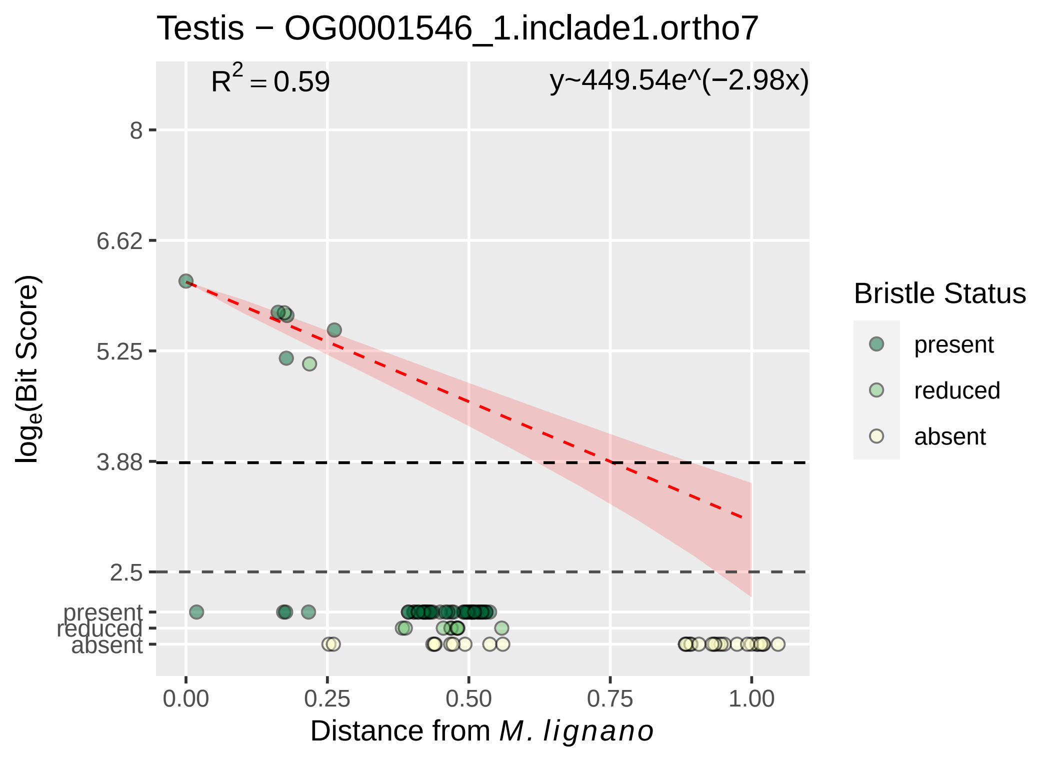

Testis − OG0001546_1.inclade1.ortho7
$R^2 = 0.59$
y~449.54e^(−2.98x)
8
6.62
5.25
3.88
2.5
present
reduced
absent
0.00
0.25
0.50
0.75
1.00
$\log_e$(Bit Score)
Distance from *M. lignano*
Bristle Status
present
reduced
absent

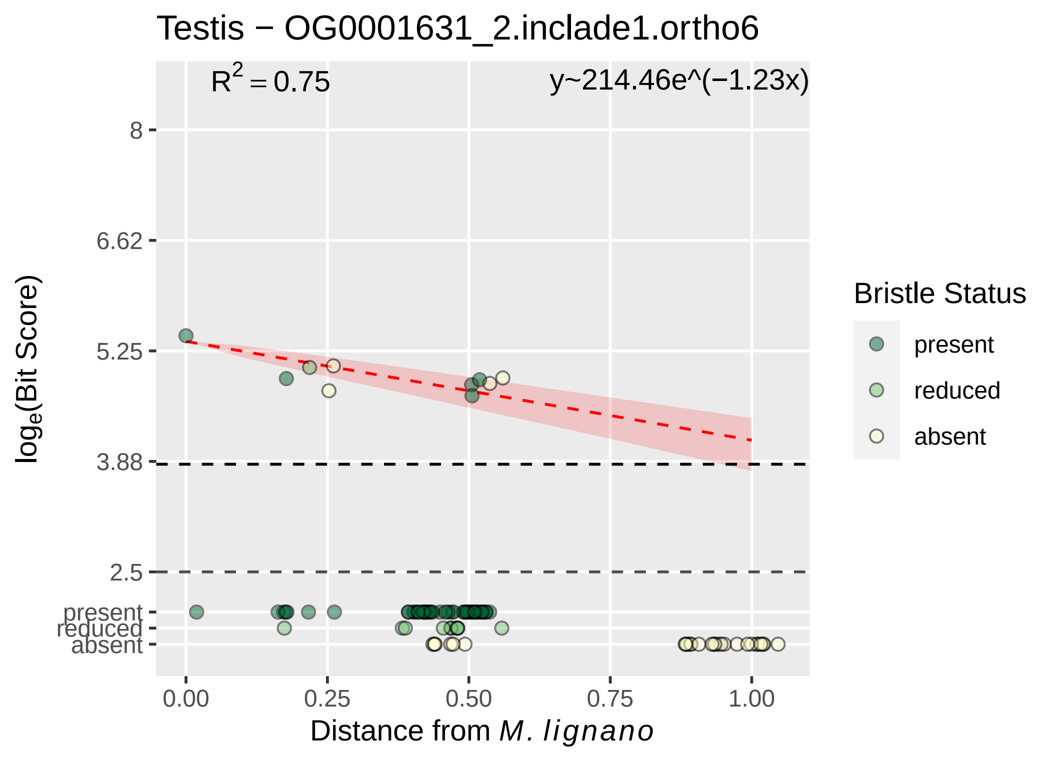

Testis − OG0001631_2.inclade1.ortho6
$R^2 = 0.75$
y~214.46e^(−1.23x)
8
6.62
5.25
3.88
2.5
present
reduced
absent
$\log_e$(Bit Score)
0.00
0.25
0.50
0.75
1.00
Distance from *M. lignano*
Bristle Status
present
reduced
absent

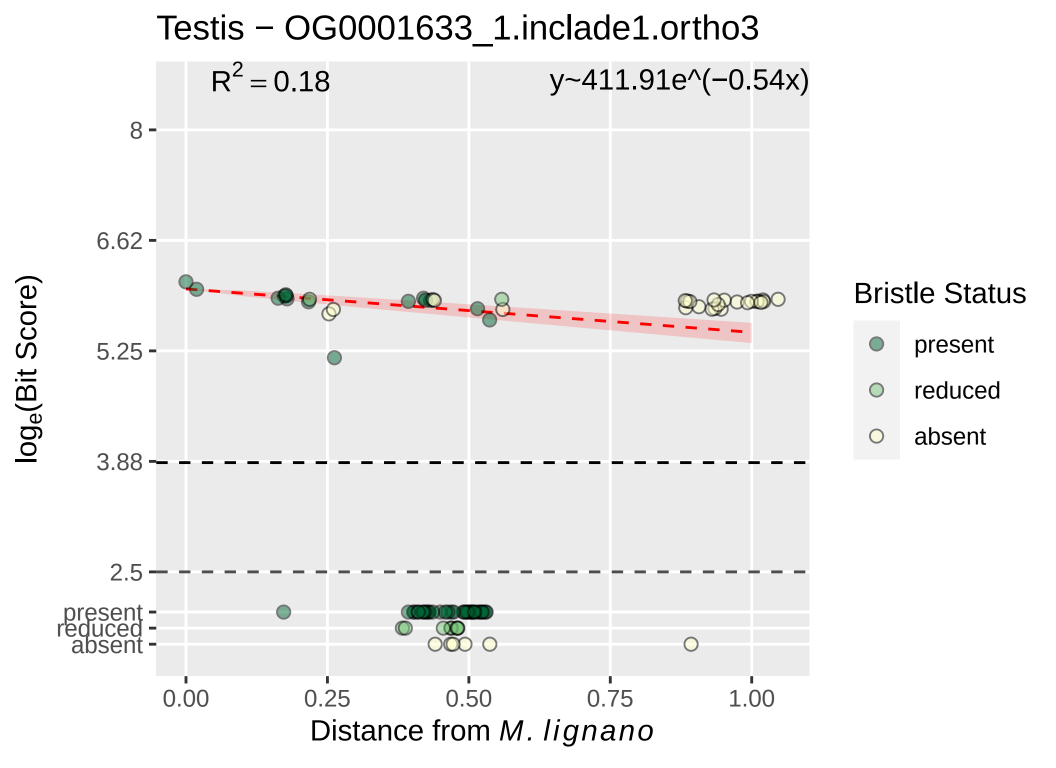

Testis − OG0001633_1.inclade1.ortho3
R^2 = 0.18
y~411.91e^(−0.54x)
8
6.62
5.25
3.88
2.5
present
reduced
absent
loge(Bit Score)
0.00
0.25
0.50
0.75
1.00
Distance from M. lignano
Bristle Status
present
reduced
absent

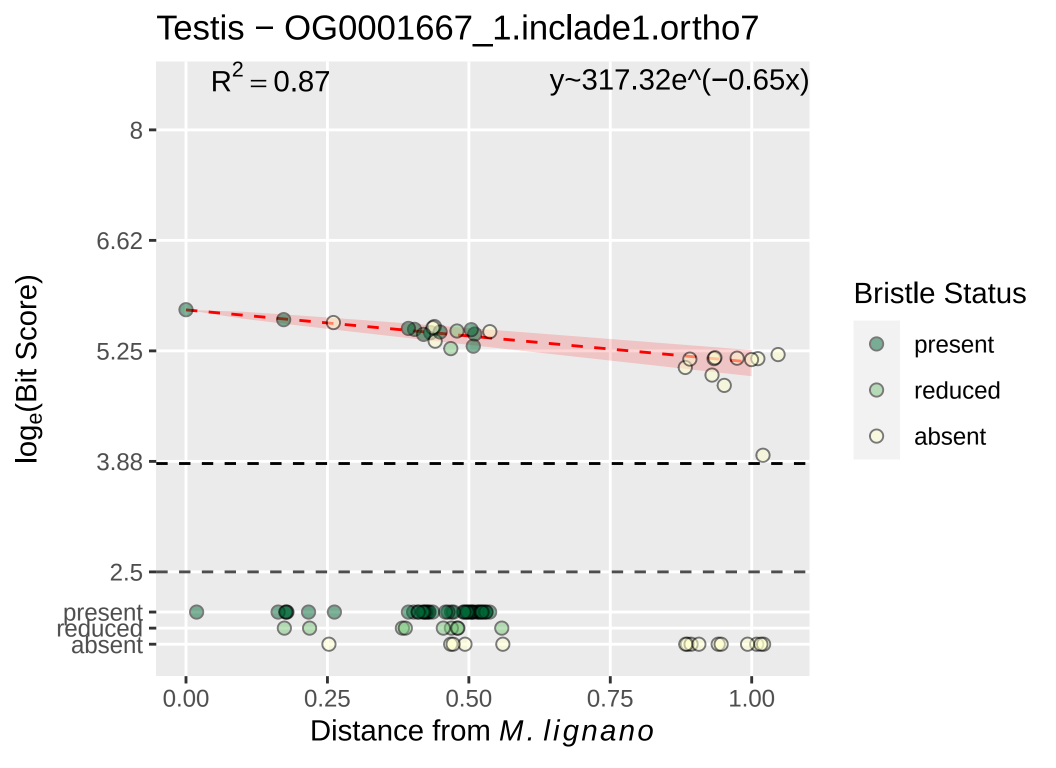

Testis − OG0001667_1.inclade1.ortho7
R^2 = 0.87
y~317.32e^(−0.65x)
loge(Bit Score)
8
6.62
5.25
3.88
2.5
present
reduced
absent
0.00
0.25
0.50
0.75
1.00
Distance from M. lignano
Bristle Status
present
reduced
absent

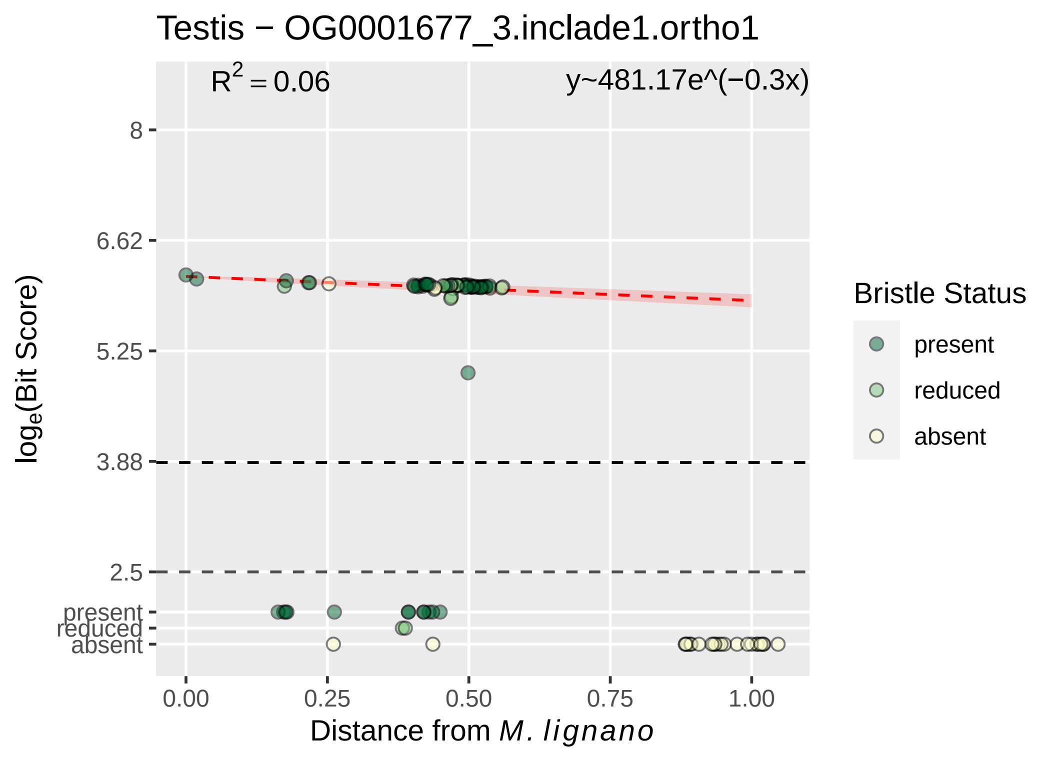

Testis − OG0001677_3.inclade1.ortho1
R2 = 0.06
y~481.17e^(−0.3x)
8
6.62
5.25
3.88
2.5
present
reduced
absent
loge(Bit Score)
0.00
0.25
0.50
0.75
1.00
Distance from M. lignano
Bristle Status
present
reduced
absent

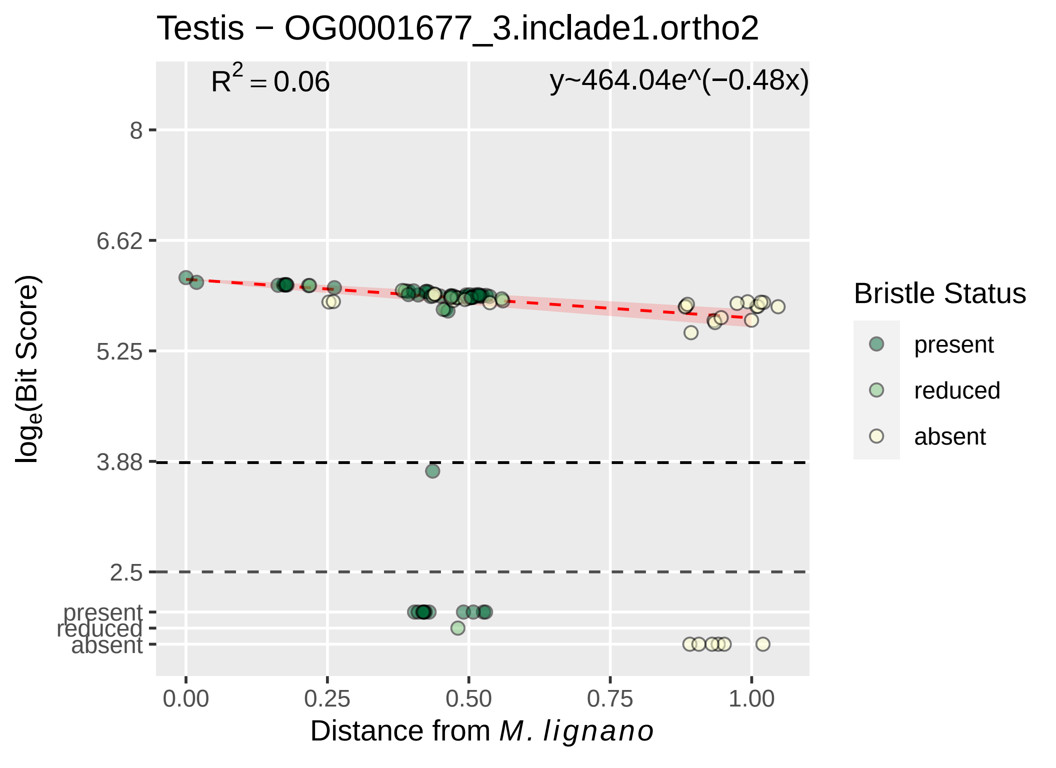

Testis − OG0001677_3.inclade1.ortho2
$R^2 = 0.06$
y~464.04e^(−0.48x)
8
6.62
5.25
3.88
2.5
present
reduced
absent
$\log_e$(Bit Score)
0.00
0.25
0.50
0.75
1.00
Distance from *M. lignano*
Bristle Status
present
reduced
absent

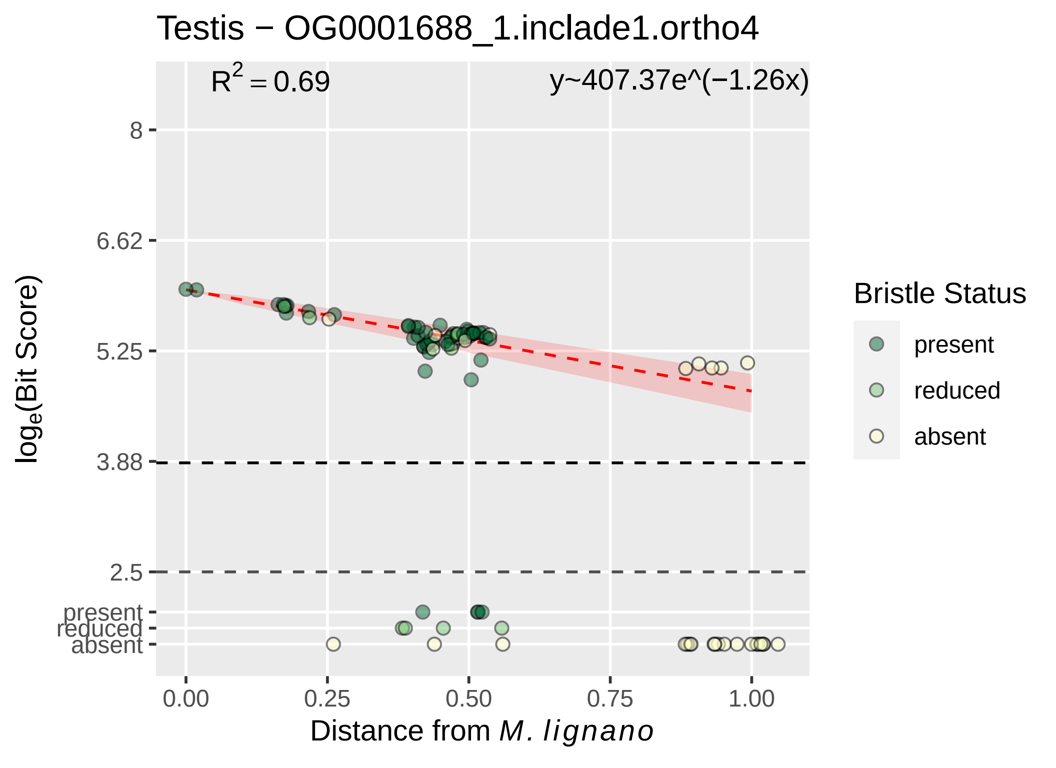

Testis − OG0001688_1.inclade1.ortho4
$R^2 = 0.69$
y~407.37e^(−1.26x)
8
6.62
5.25
3.88
2.5
present
reduced
absent
0.00
0.25
0.50
0.75
1.00
Distance from M. lignano
loge(Bit Score)
Bristle Status
present
reduced
absent

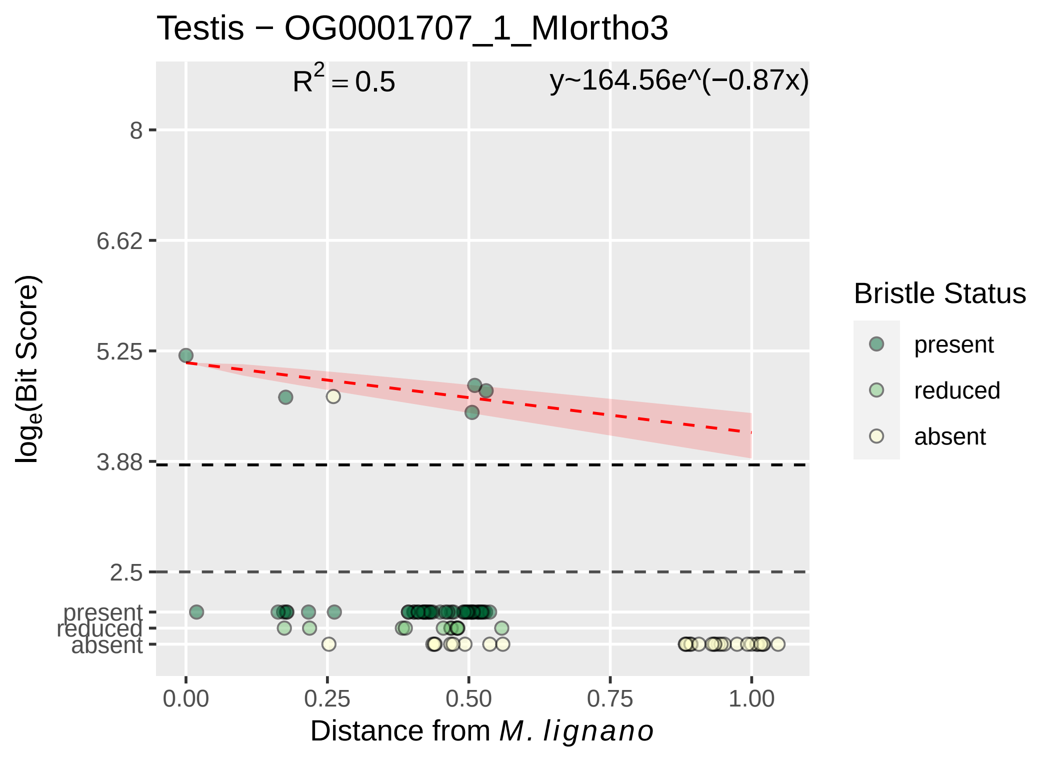
Testis − OG0001707_1_MIortho3
$R^2 = 0.5$
y~164.56e^(−0.87x)
8
6.62
5.25
3.88
2.5
present
reduced
absent
$log_e$(Bit Score)
0.00
0.25
0.50
0.75
1.00
Distance from *M. lignano*
Bristle Status
present
reduced
absent

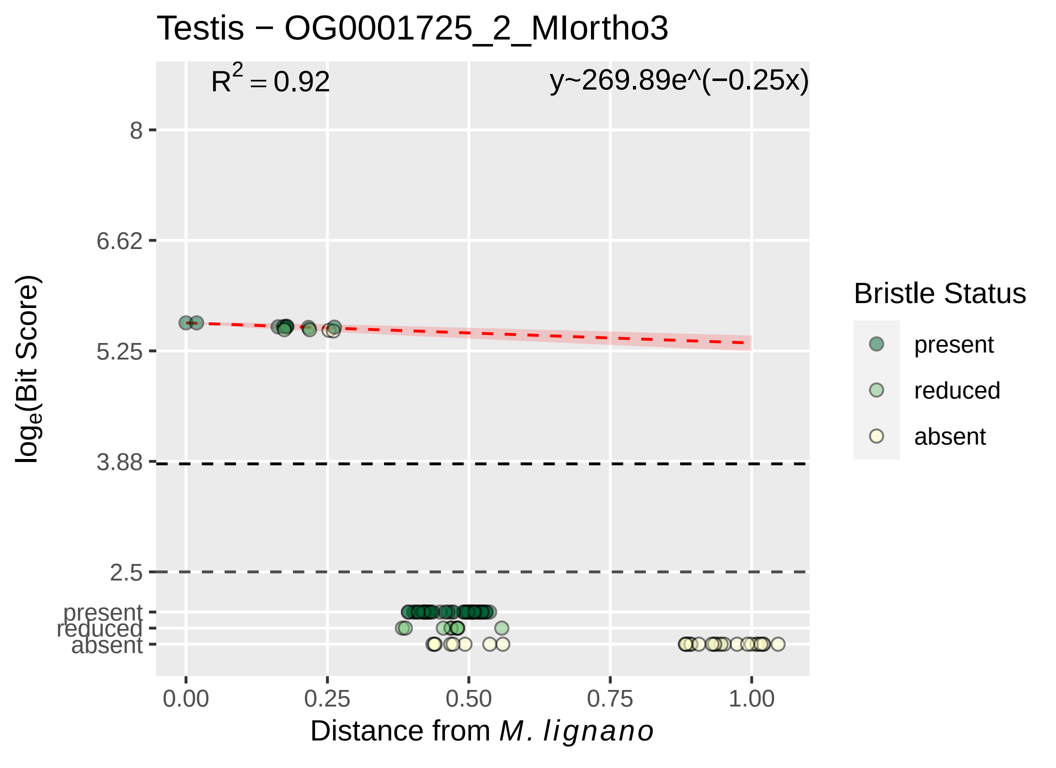

Testis − OG0001725_2_MIortho3
$R^2 = 0.92$
y~269.89e^(−0.25x)
8
6.62
5.25
3.88
2.5
present
reduced
absent
$log_e$(Bit Score)
0.00
0.25
0.50
0.75
1.00
Distance from *M. lignano*
Bristle Status
present
reduced
absent

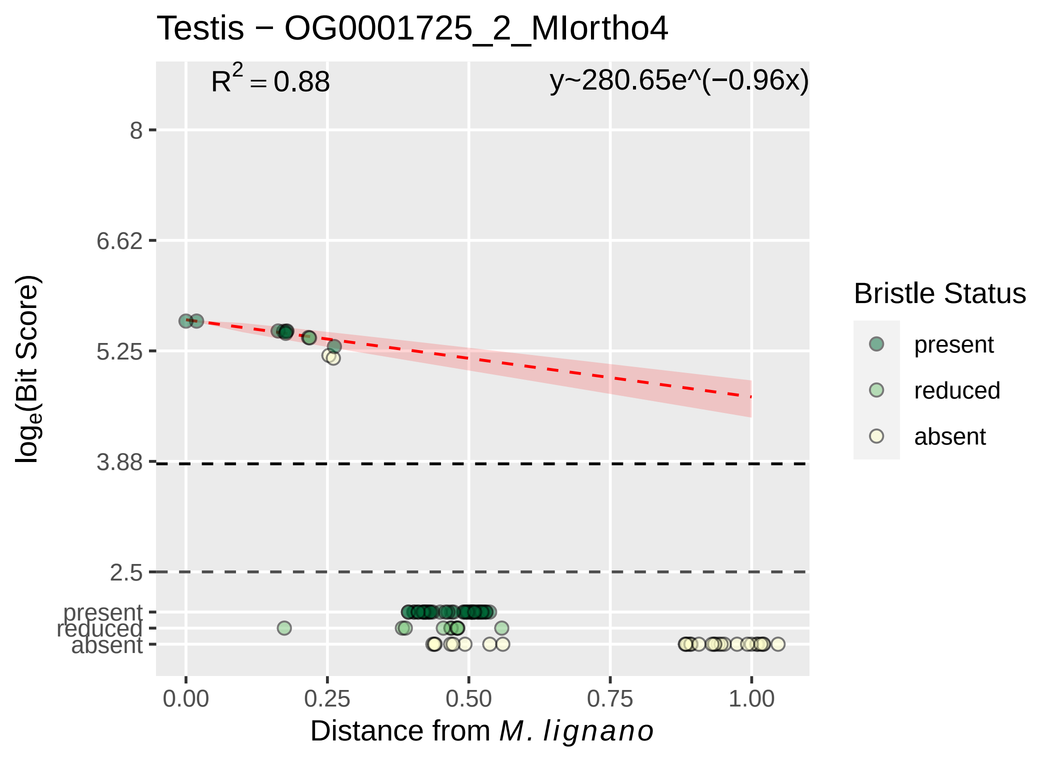

Testis − OG0001725_2_MIortho4
$R^2 = 0.88$
y~280.65e^(−0.96x)
8
6.62
5.25
3.88
2.5
present
reduced
absent
log$_e$(Bit Score)
0.00
0.25
0.50
0.75
1.00
Distance from *M. lignano*
Bristle Status
present
reduced
absent

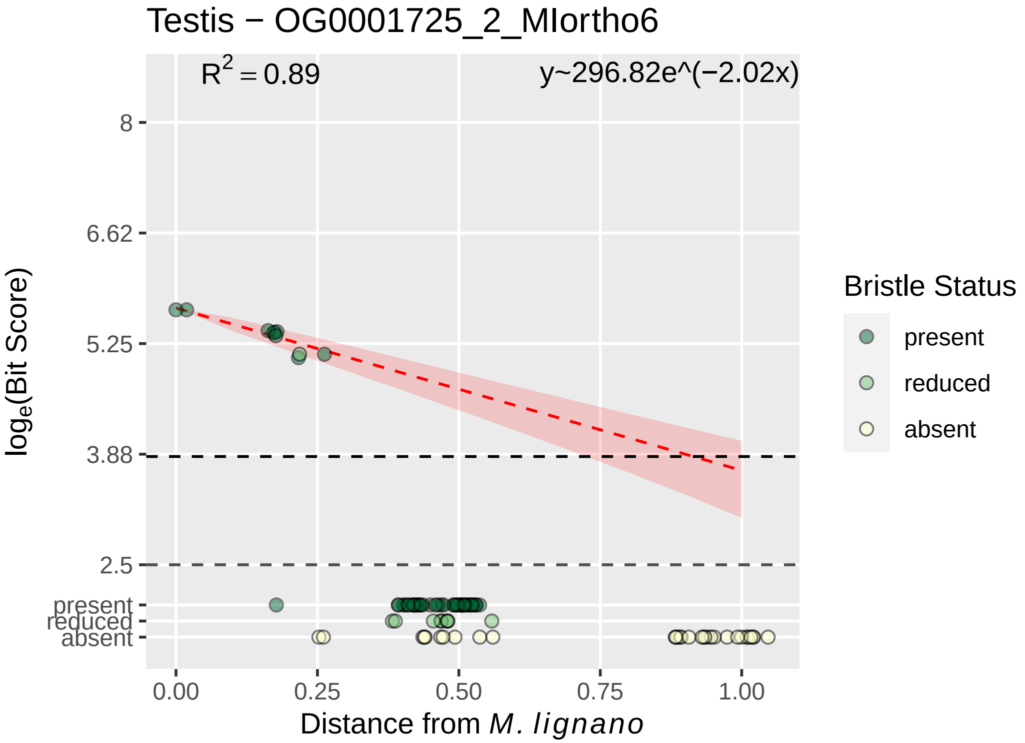
Testis − OG0001725_2_MIortho6
R2 = 0.89
y~296.82e^(−2.02x)
8
6.62
5.25
3.88
2.5
present
reduced
absent
log_e(Bit Score)
0.00
0.25
0.50
0.75
1.00
Distance from M. lignano
Bristle Status
present
reduced
absent

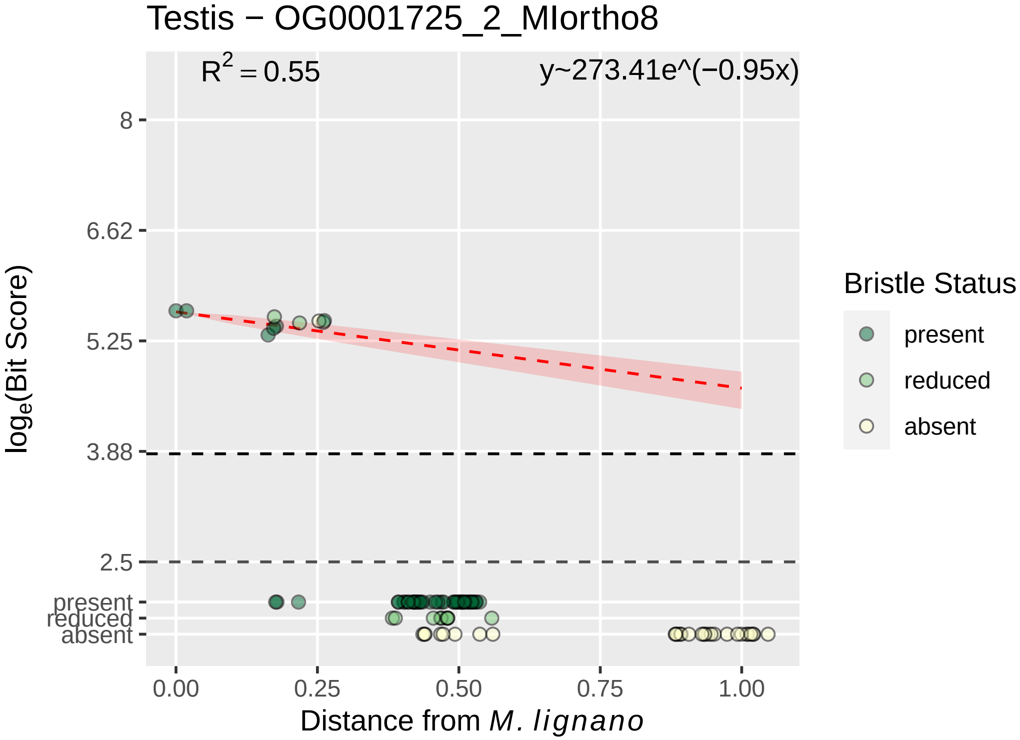

Testis − OG0001725_2_MIortho8
$R^2 = 0.55$
y~273.41e^(−0.95x)
8
6.62
5.25
3.88
2.5
present
reduced
absent
$log_e$(Bit Score)
0.00
0.25
0.50
0.75
1.00
Distance from *M. lignano*
Bristle Status
present
reduced
absent

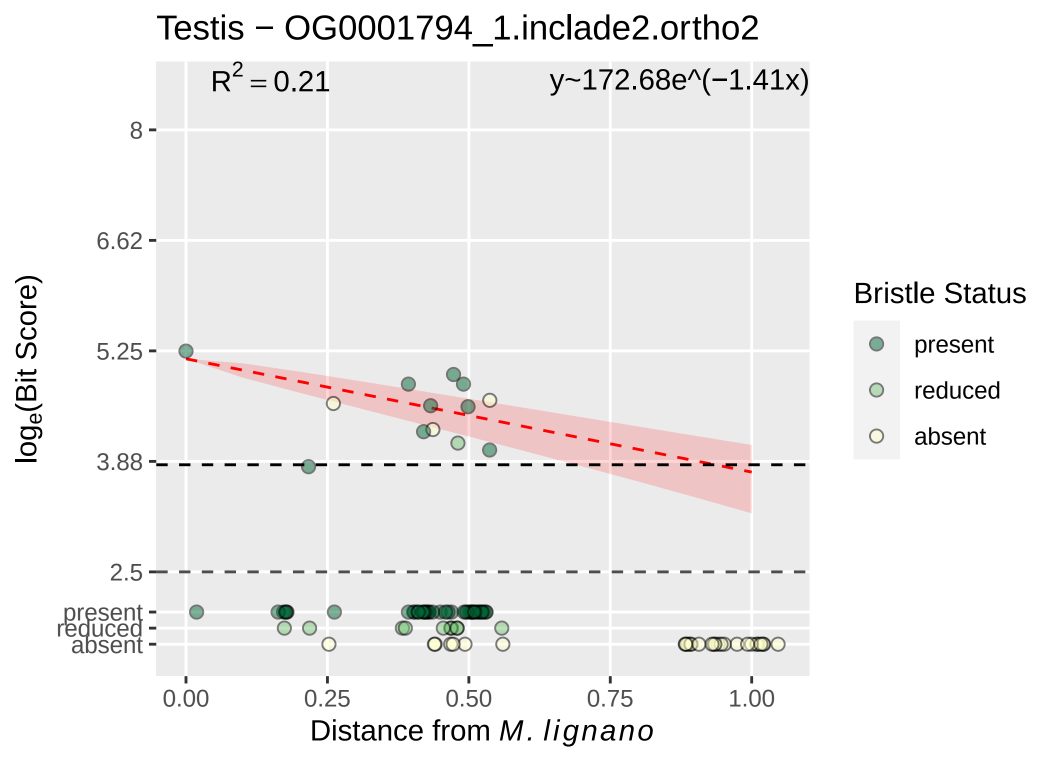
Testis − OG0001794_1.inclade2.ortho2
$R^2 = 0.21$
y~172.68e^(−1.41x)
8
6.62
5.25
3.88
2.5
present
reduced
absent
0.00
0.25
0.50
0.75
1.00
$log_e$(Bit Score)
Distance from *M. lignano*
Bristle Status
present
reduced
absent

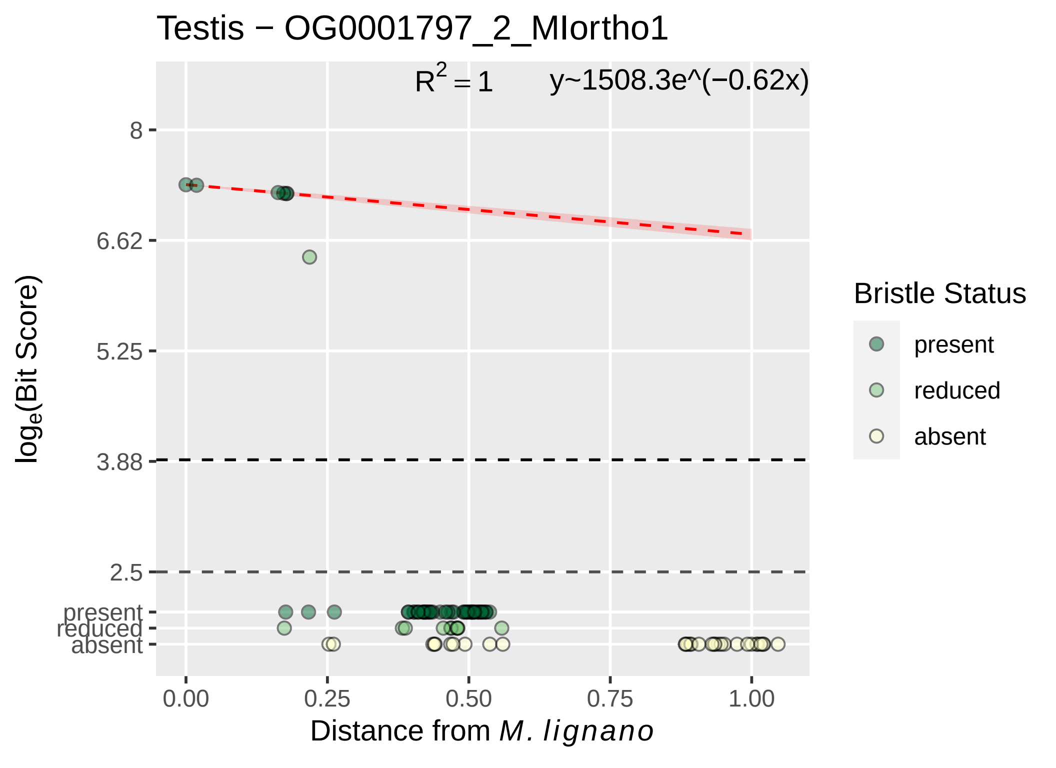

Testis − OG0001797_2_MIortho1
$R^2 = 1$
y~1508.3e^(−0.62x)
8
6.62
5.25
3.88
2.5
present
reduced
absent
$\log_e$(Bit Score)
0.00
0.25
0.50
0.75
1.00
Distance from M. lignano
Bristle Status
present
reduced
absent

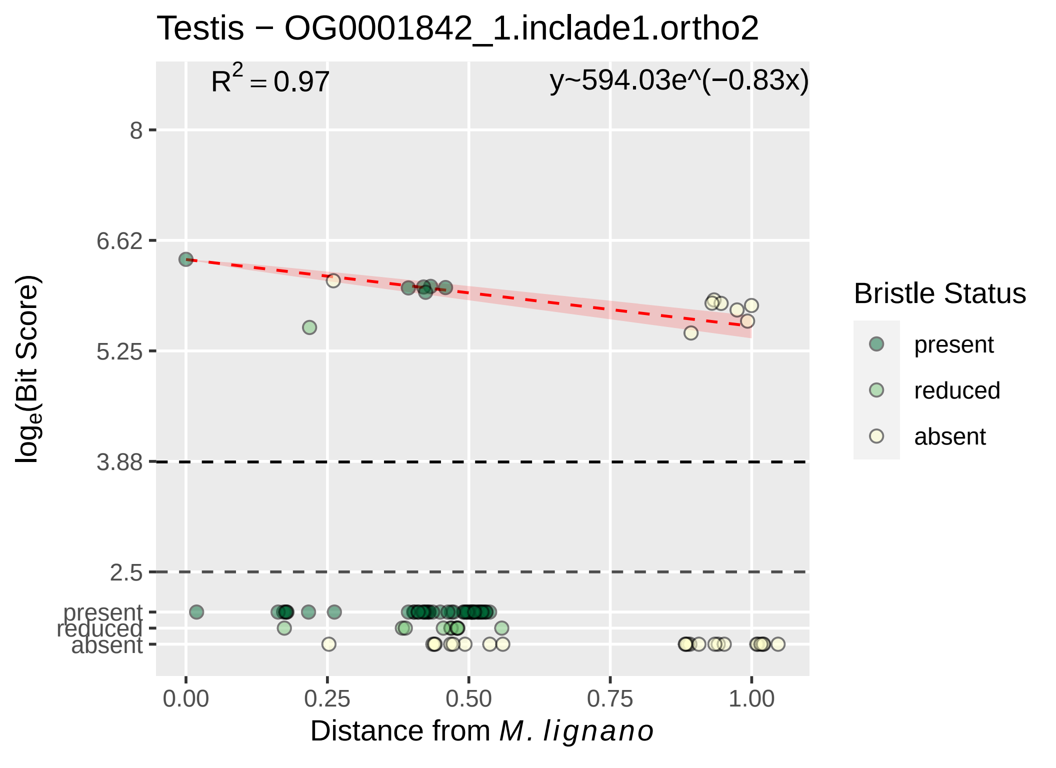
Testis − OG0001842_1.inclade1.ortho2
R2 = 0.97
y~594.03e^(−0.83x)
8
6.62
5.25
3.88
2.5
present
reduced
absent
0.00
0.25
0.50
0.75
1.00
loge(Bit Score)
Distance from M. lignano
Bristle Status
present
reduced
absent

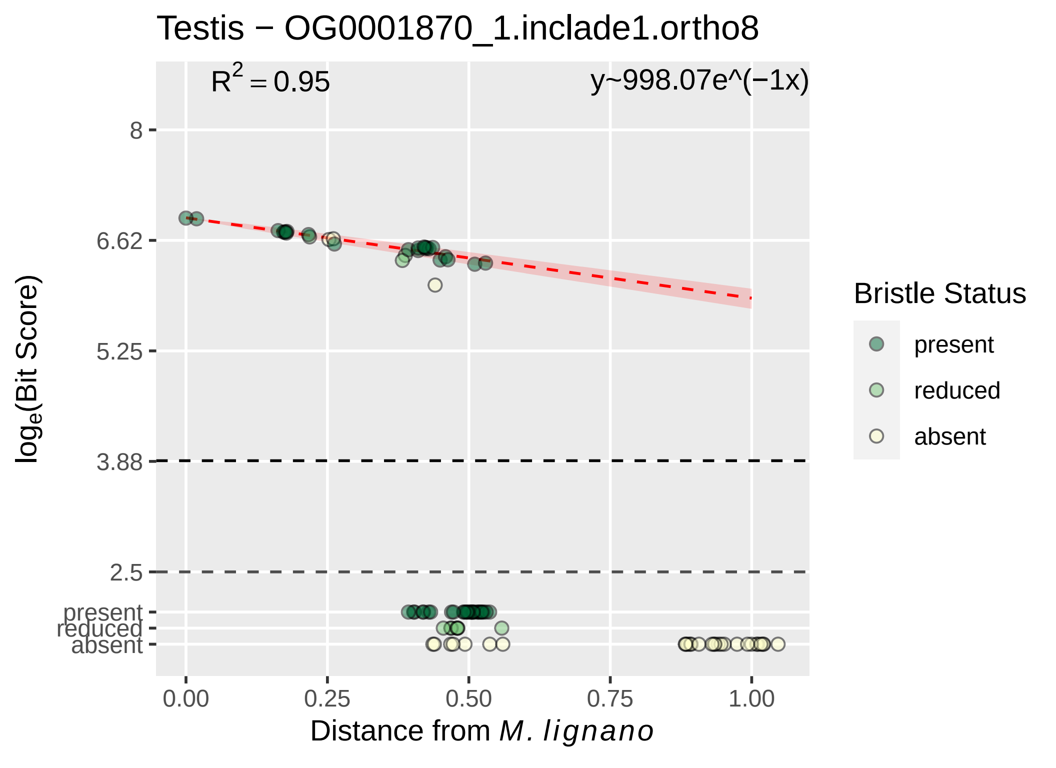

Testis − OG0001870_1.inclade1.ortho8
$R^2 = 0.95$
y~998.07e^(−1x)
8
6.62
5.25
3.88
2.5
present
reduced
absent
0.00
0.25
0.50
0.75
1.00
Distance from *M. lignano*
$\log_e$(Bit Score)
Bristle Status
present
reduced
absent

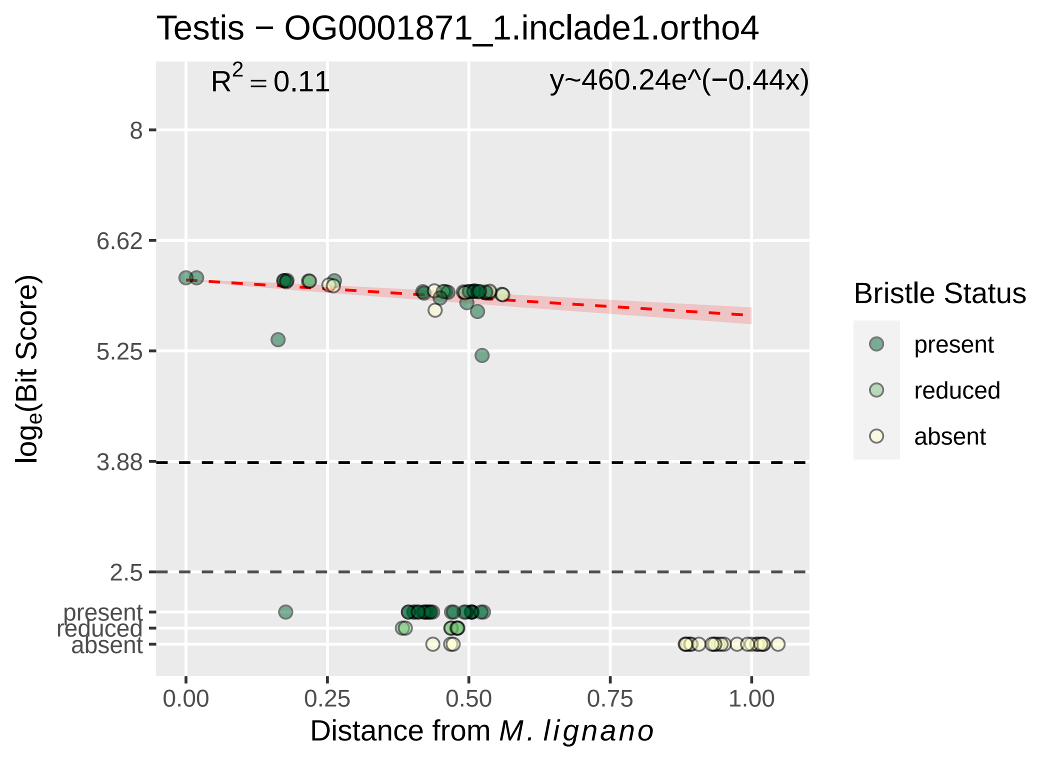

Testis − OG0001871_1.inclade1.ortho4
$R^2 = 0.11$
y~460.24e^(−0.44x)
8
6.62
5.25
3.88
2.5
present
reduced
absent
0.00
0.25
0.50
0.75
1.00
Distance from M. lignano
log_e(Bit Score)
Bristle Status
present
reduced
absent

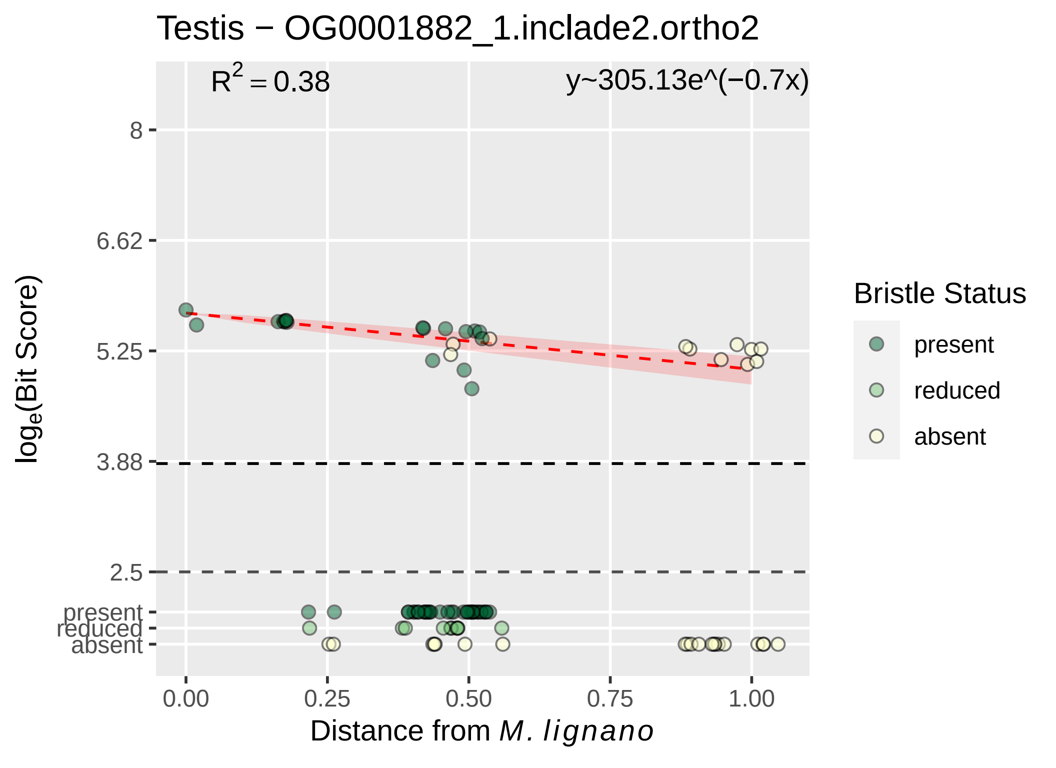

Testis − OG0001882_1.inclade2.ortho2
R^2 = 0.38
y~305.13e^(−0.7x)
8
6.62
5.25
3.88
2.5
present
reduced
absent
log_e(Bit Score)
0.00
0.25
0.50
0.75
1.00
Distance from M. lignano
Bristle Status
present
reduced
absent

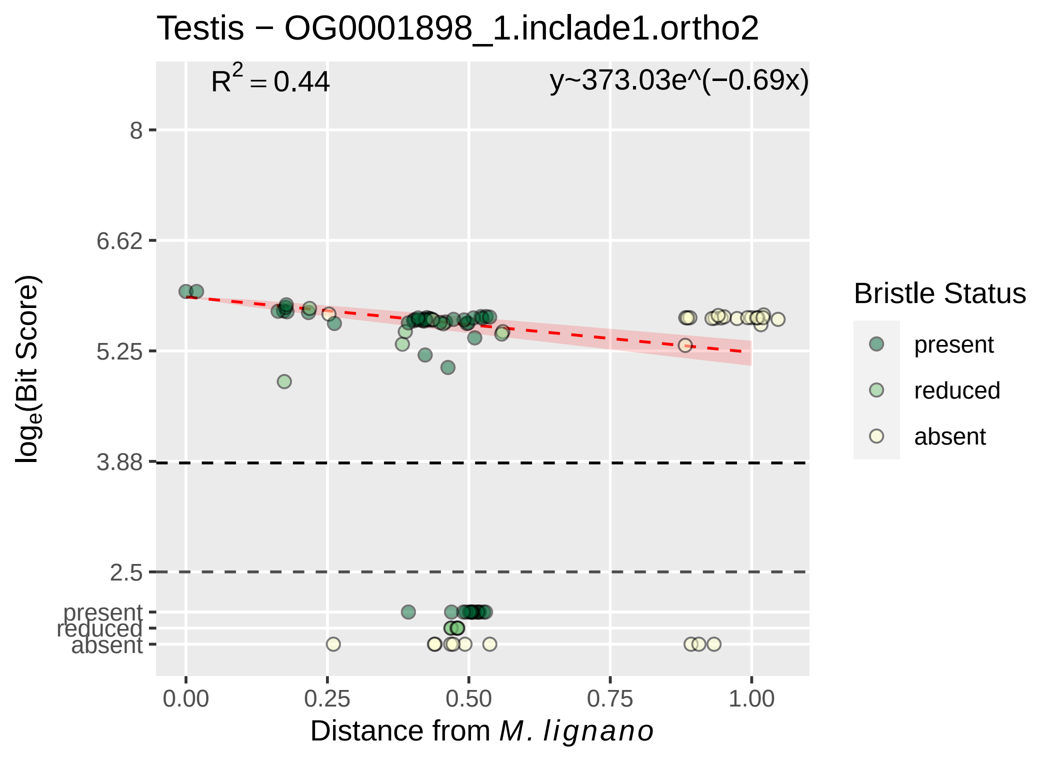

Testis − OG0001898_1.inclade1.ortho2
$R^2 = 0.44$
y~373.03e^(−0.69x)
8
6.62
5.25
3.88
2.5
present
reduced
absent
$\log_e$(Bit Score)
0.00
0.25
0.50
0.75
1.00
Distance from *M. lignano*
Bristle Status
present
reduced
absent

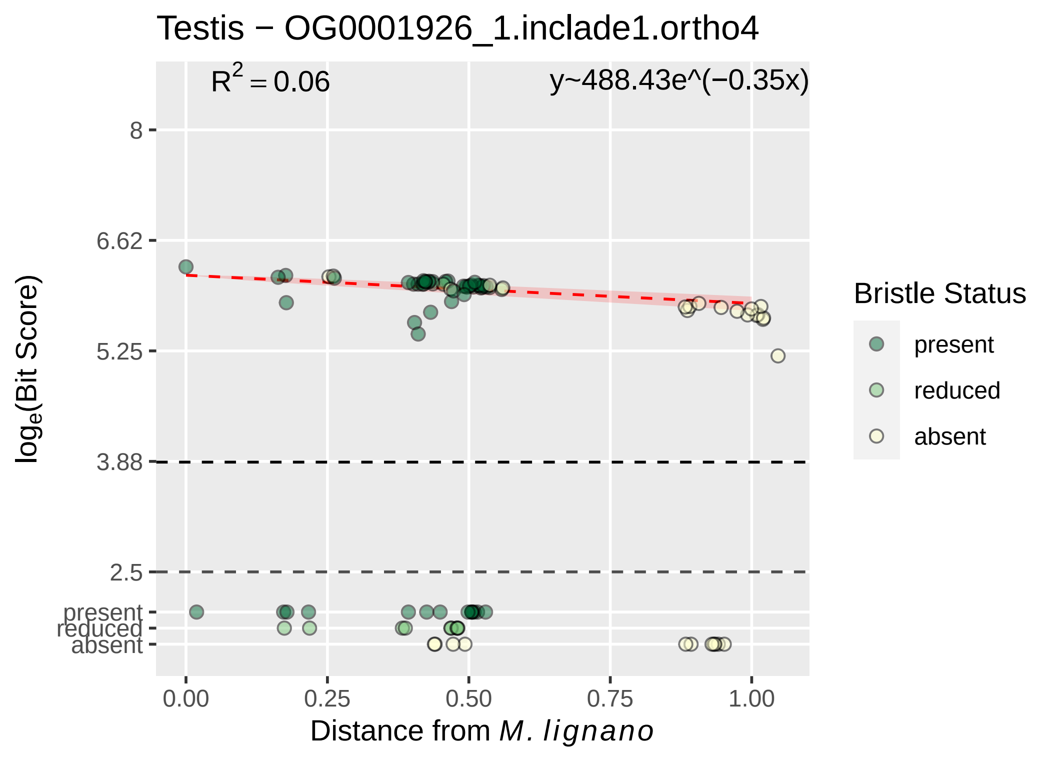
Testis − OG0001926_1.inclade1.ortho4
R2 = 0.06
y~488.43e^(−0.35x)
8
6.62
5.25
3.88
2.5
present
reduced
absent
loge(Bit Score)
0.00
0.25
0.50
0.75
1.00
Distance from M. lignano
Bristle Status
present
reduced
absent

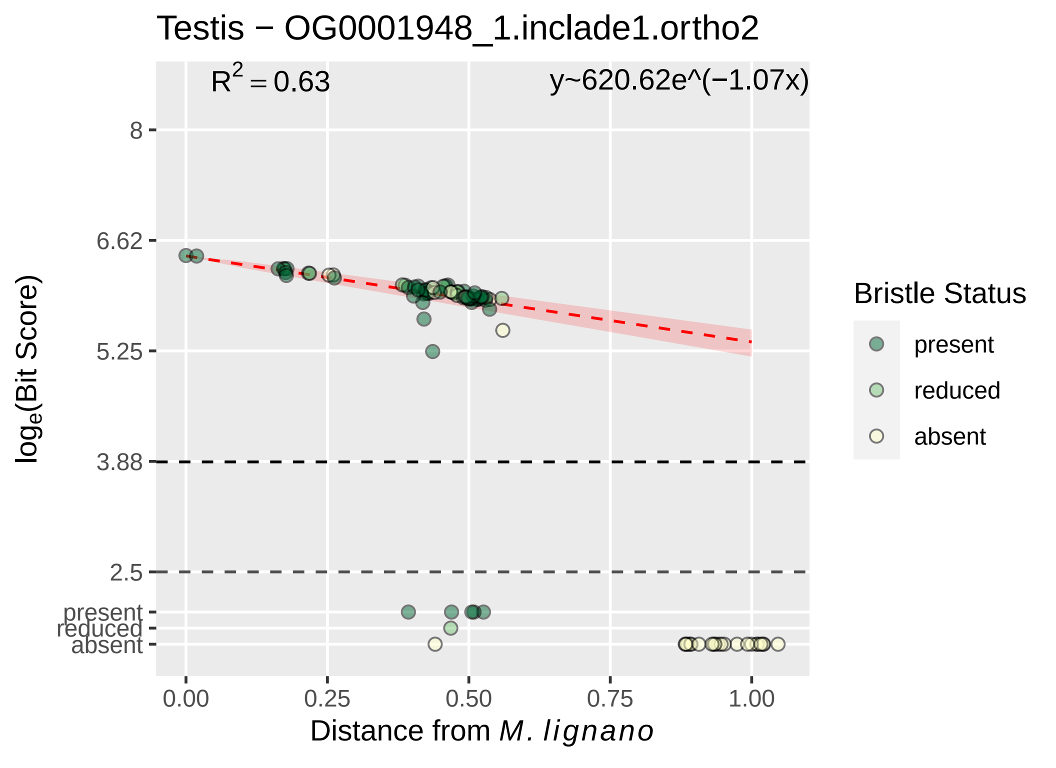

Testis − OG0001948_1.inclade1.ortho2
R² = 0.63
y~620.62e^(−1.07x)
8
6.62
5.25
3.88
2.5
present
reduced
absent
log_e(Bit Score)
0.00
0.25
0.50
0.75
1.00
Distance from M. lignano
Bristle Status
present
reduced
absent

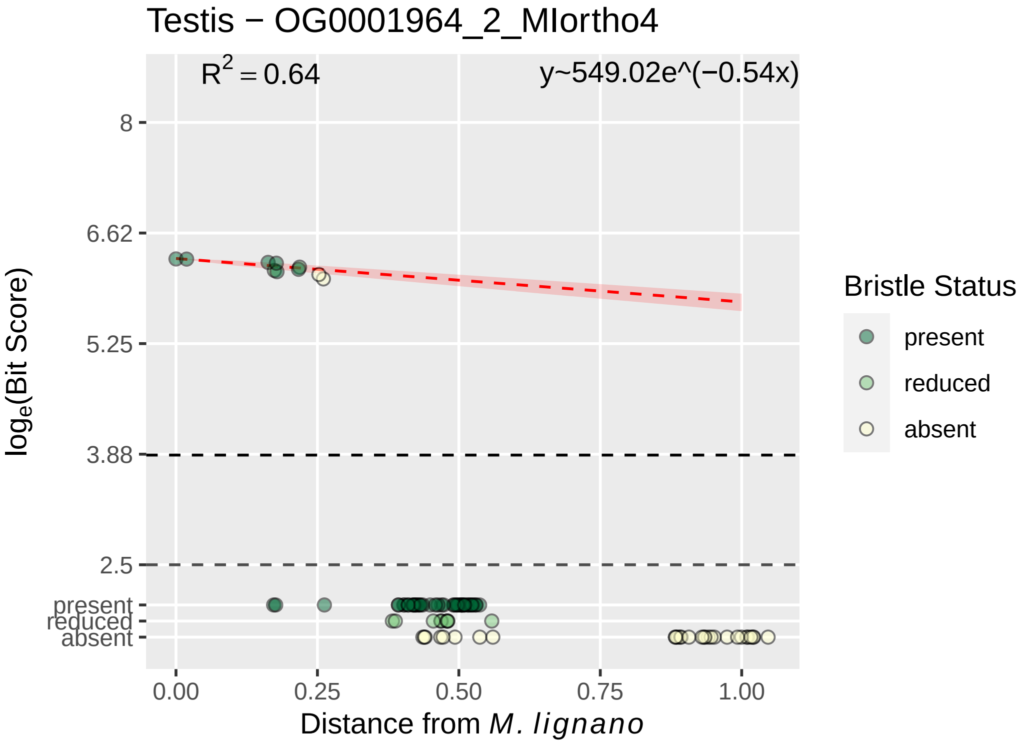

Testis − OG0001964_2_MIortho4
$R^2 = 0.64$
y~549.02e^(−0.54x)
8
6.62
5.25
3.88
2.5
present
reduced
absent
$\log_e$(Bit Score)
0.00
0.25
0.50
0.75
1.00
Distance from *M. lignano*
Bristle Status
present
reduced
absent

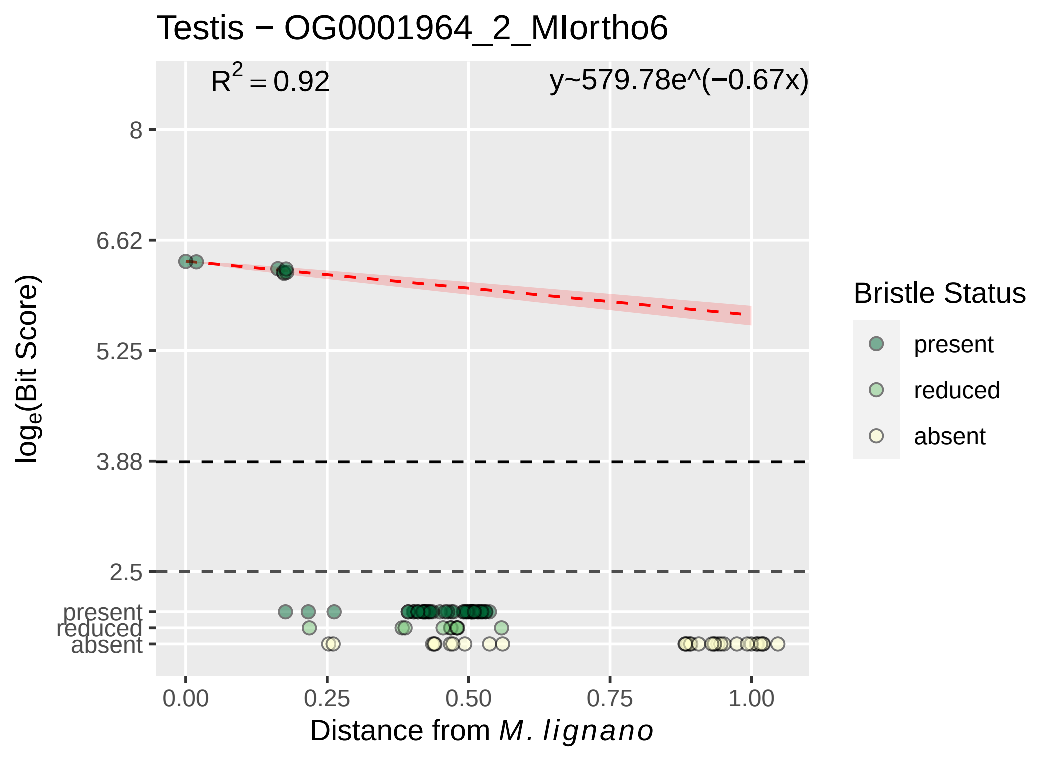
Testis − OG0001964_2_MIortho6
$R^2 = 0.92$
y~579.78e^(−0.67x)
8
6.62
5.25
3.88
2.5
present
reduced
absent
$\log_e$(Bit Score)
0.00
0.25
0.50
0.75
1.00
Distance from *M. lignano*
Bristle Status
present
reduced
absent

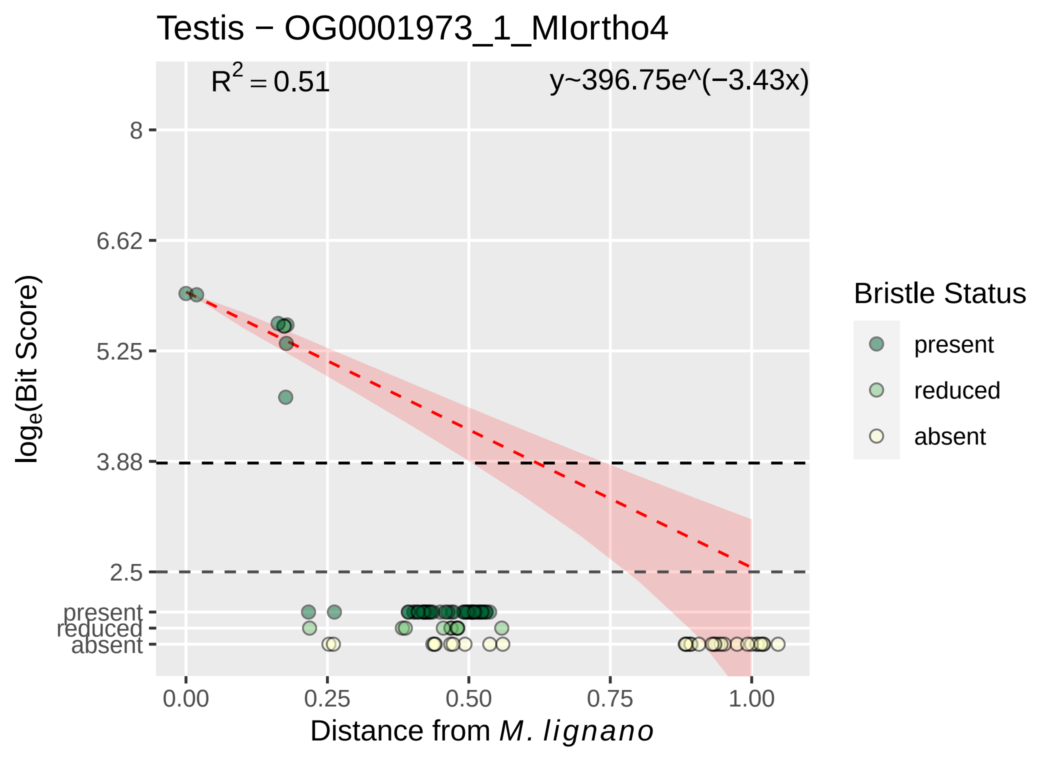

Testis − OG0001973_1_MIortho4
$R^2 = 0.51$
y~396.75e^(−3.43x)
8
6.62
5.25
3.88
2.5
present
reduced
absent
$\log_e$(Bit Score)
0.00
0.25
0.50
0.75
1.00
Distance from *M. lignano*
Bristle Status
present
reduced
absent

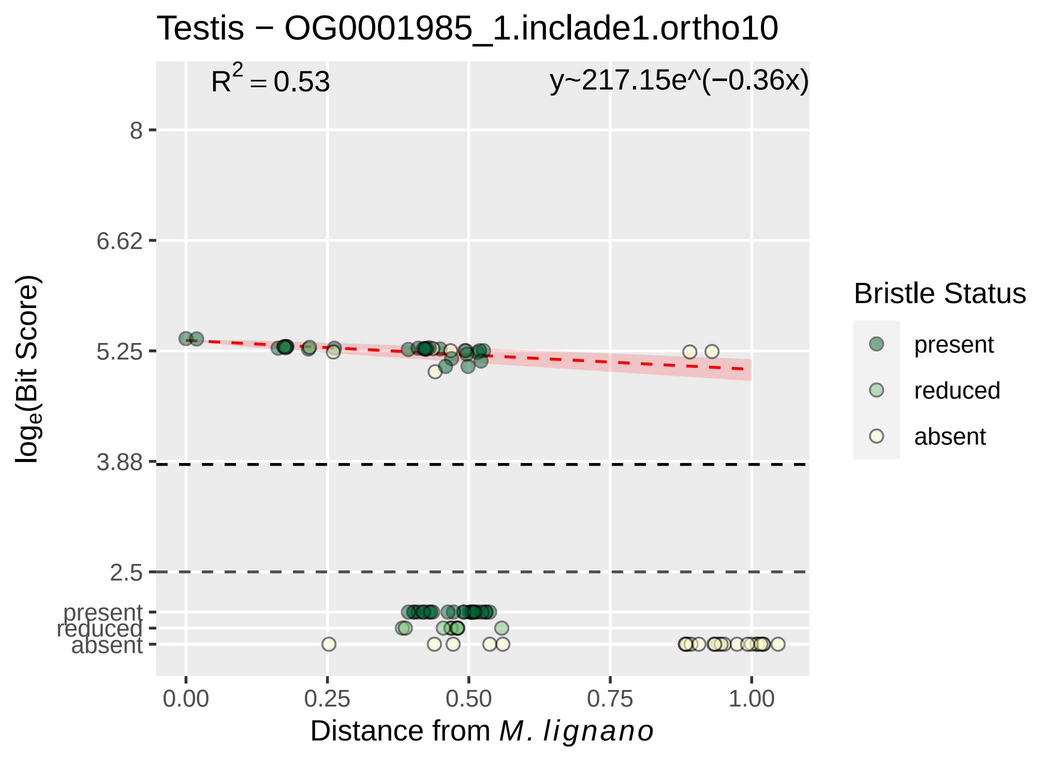
Testis − OG0001985_1.inclade1.ortho10
R² = 0.53
y~217.15e^(−0.36x)
8
6.62
5.25
3.88
2.5
present
reduced
absent
log_e(Bit Score)
0.00
0.25
0.50
0.75
1.00
Distance from M. lignano
Bristle Status
present
reduced
absent

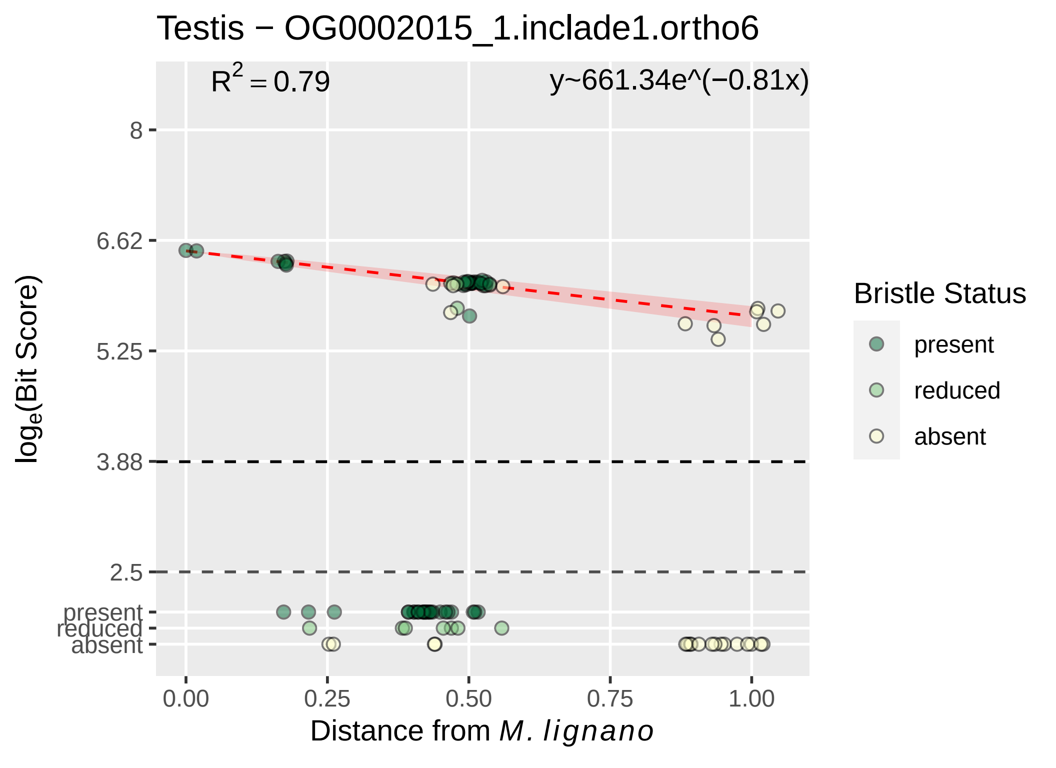
Testis − OG0002015_1.inclade1.ortho6
$R^2 = 0.79$
y~661.34e^(−0.81x)
8
6.62
5.25
3.88
2.5
present
reduced
absent
$log_e$(Bit Score)
0.00
0.25
0.50
0.75
1.00
Distance from *M. lignano*
Bristle Status
present
reduced
absent

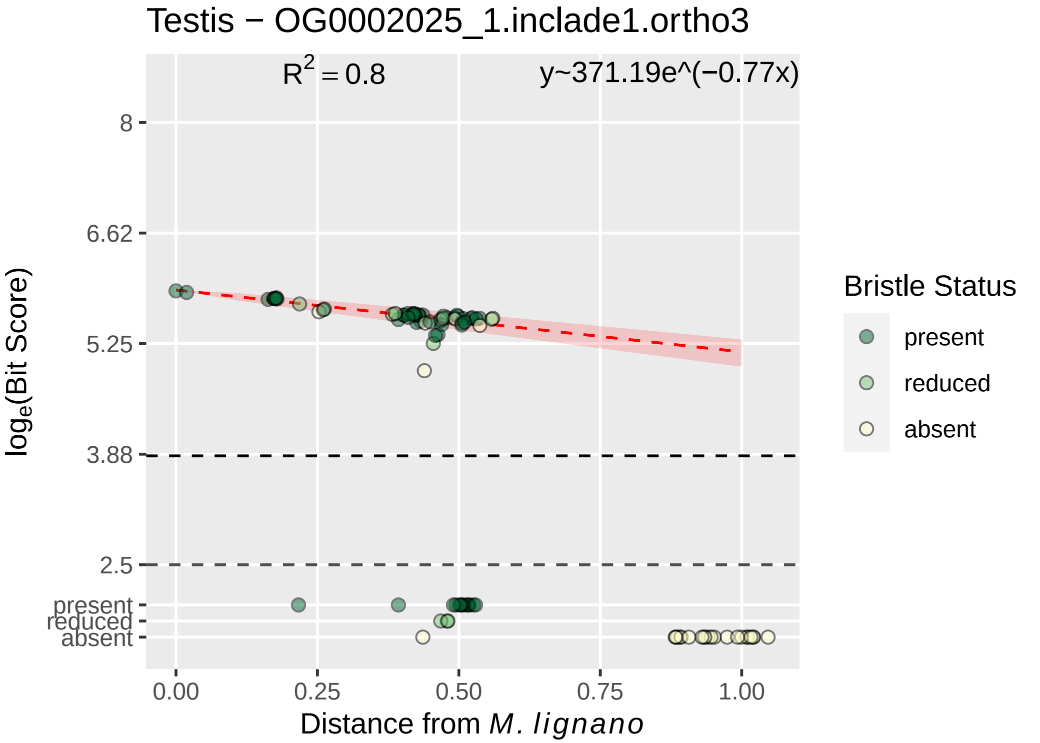
Testis − OG0002025_1.inclade1.ortho3
$R^2 = 0.8$
y~371.19e^(−0.77x)
8
6.62
5.25
3.88
2.5
present
reduced
absent
$\log_e$(Bit Score)
0.00
0.25
0.50
0.75
1.00
Distance from *M. lignano*
Bristle Status
present
reduced
absent

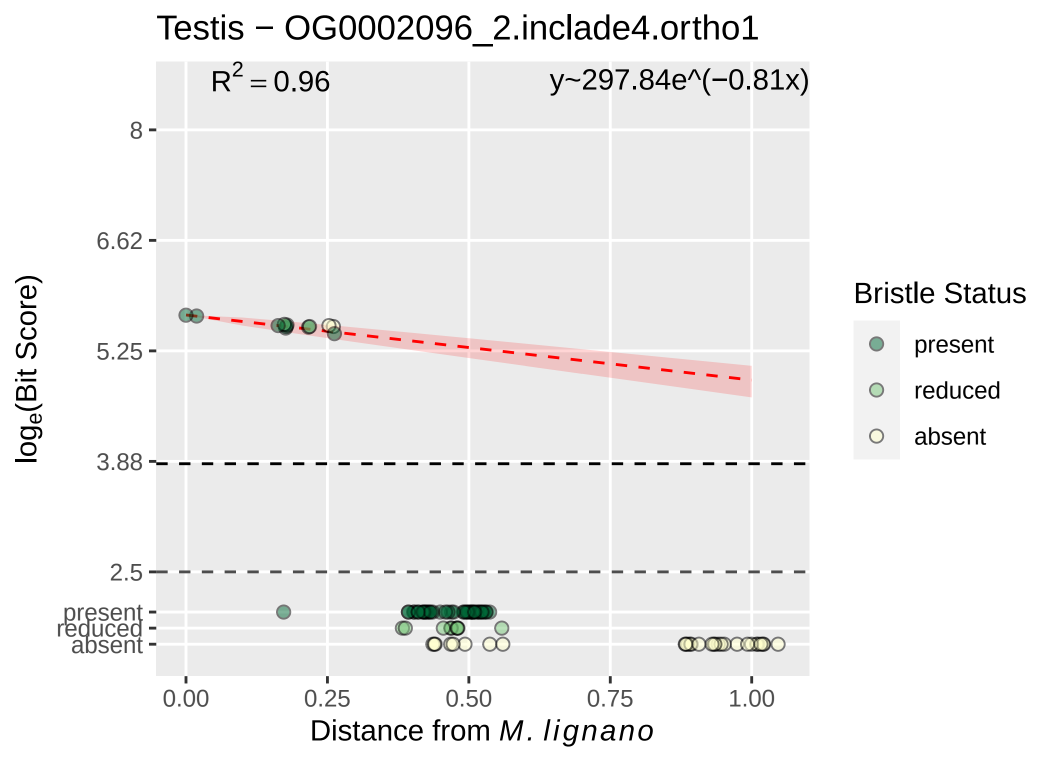

Testis − OG0002096_2.inclade4.ortho1
$R^2 = 0.96$
y~297.84e^(−0.81x)
8
6.62
5.25
3.88
2.5
present
reduced
absent
$\log_e$(Bit Score)
0.00
0.25
0.50
0.75
1.00
Distance from *M. lignano*
Bristle Status
present
reduced
absent

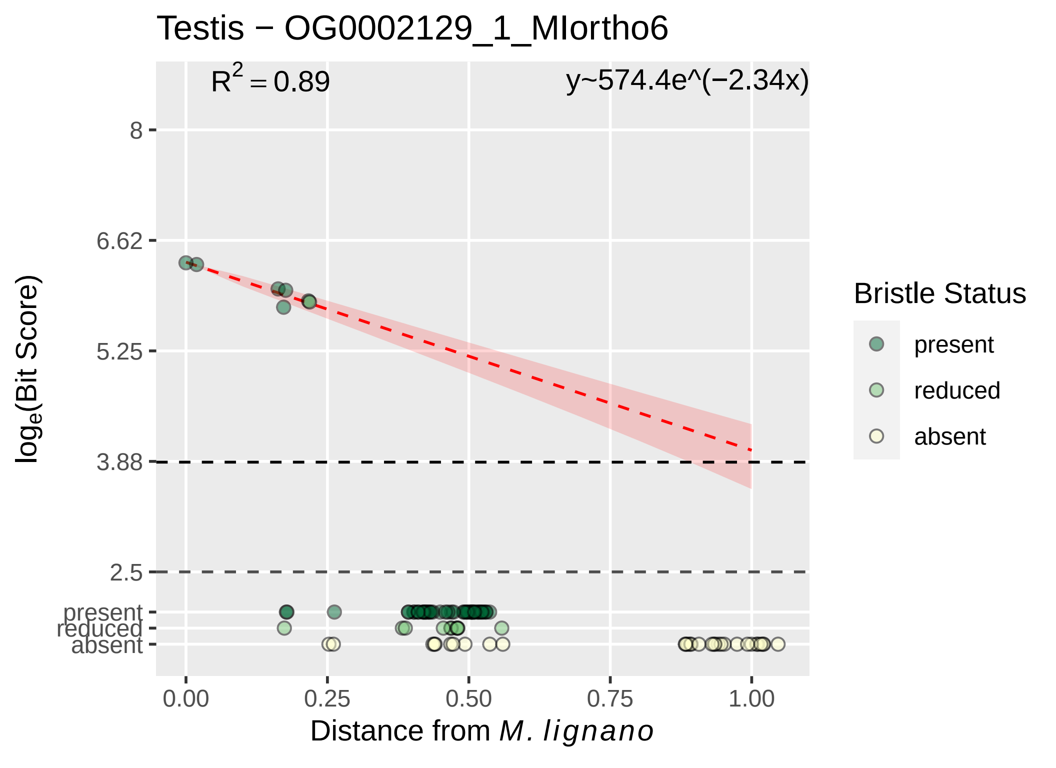

Testis − OG0002129_1_MIortho6
R2 = 0.89
y~574.4e^(−2.34x)
8
6.62
5.25
3.88
2.5
present
reduced
absent
0.00
0.25
0.50
0.75
1.00
loge(Bit Score)
Distance from M. lignano
Bristle Status
present
reduced
absent

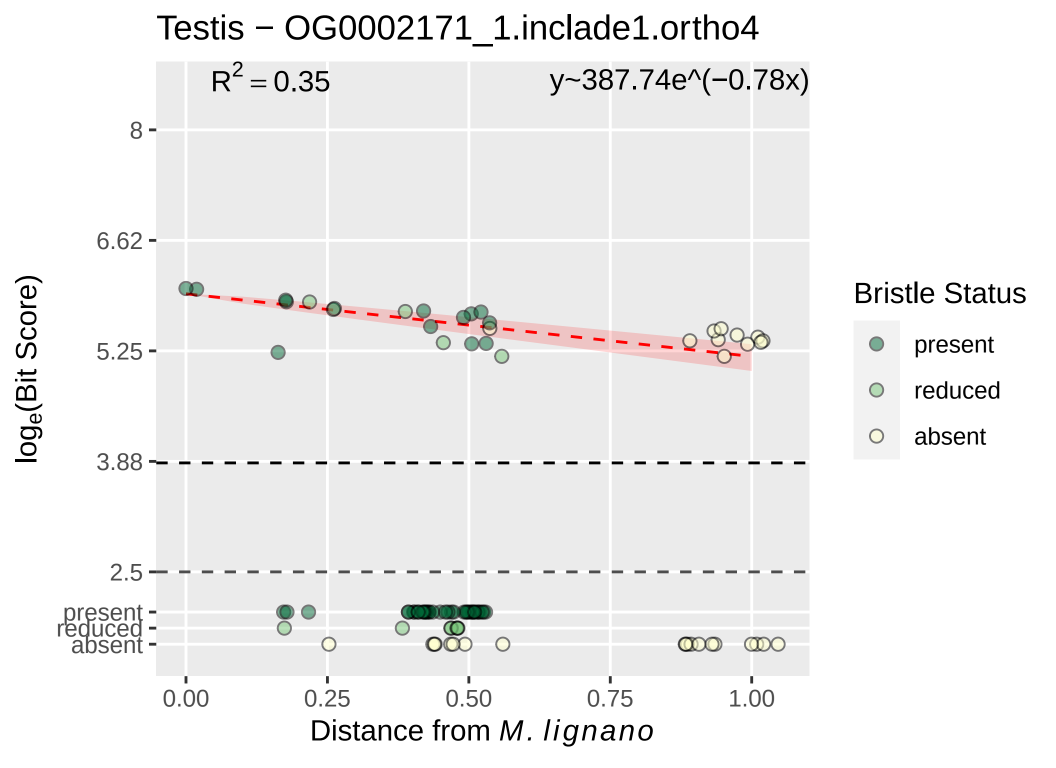

Testis − OG0002171_1.inclade1.ortho4
$R^2 = 0.35$
y~387.74e^(−0.78x)
8
6.62
5.25
3.88
2.5
present
reduced
absent
0.00
0.25
0.50
0.75
1.00
$\log_e$(Bit Score)
Distance from *M. lignano*
Bristle Status
present
reduced
absent

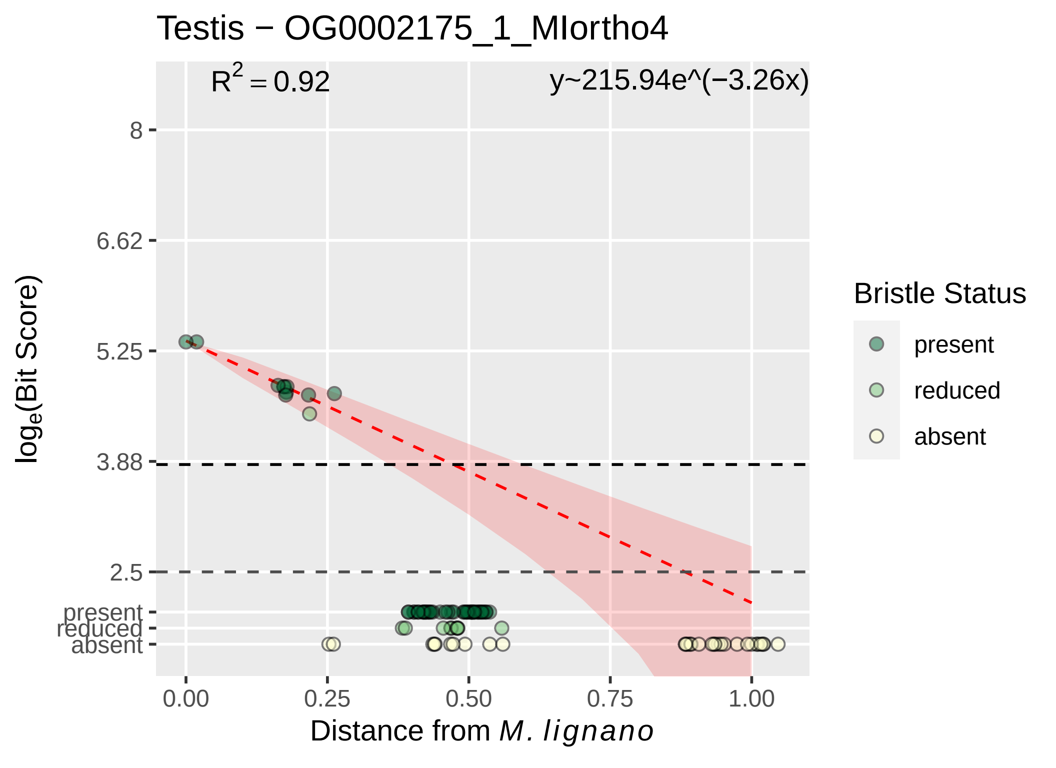

Testis − OG0002175_1_MIortho4
$R^2 = 0.92$
y~215.94e^(−3.26x)
8
6.62
5.25
3.88
2.5
present
reduced
absent
0.00
0.25
0.50
0.75
1.00
Distance from M. lignano
log$_e$(Bit Score)
Bristle Status
present
reduced
absent

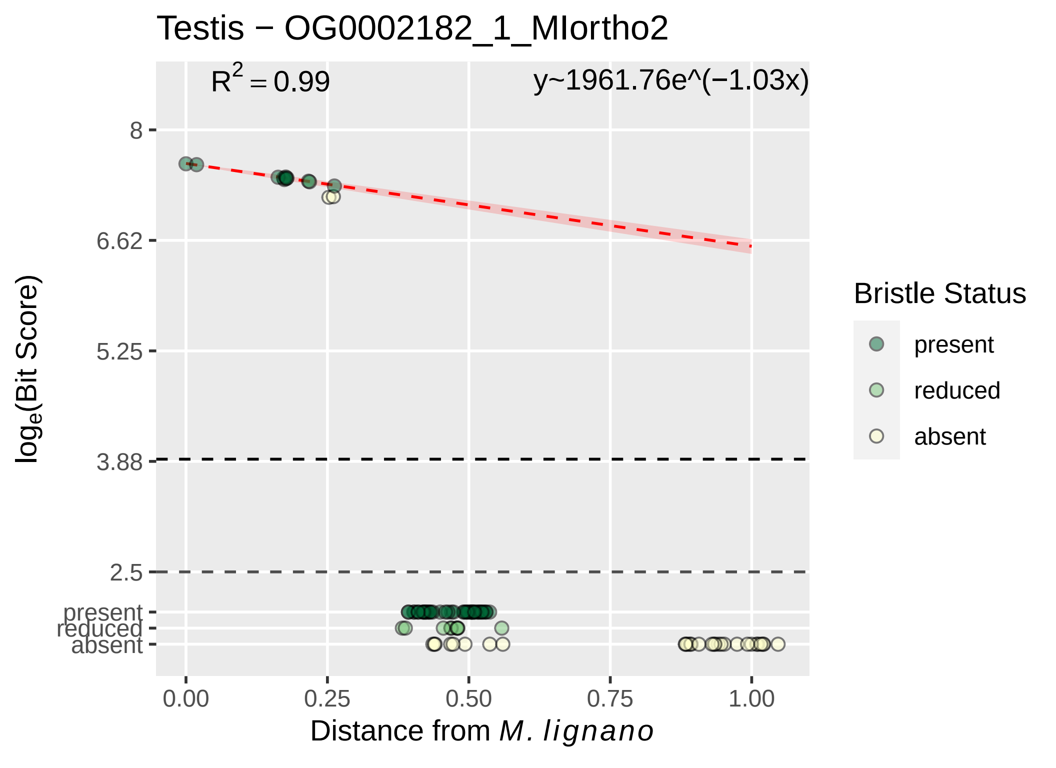

Testis − OG0002182_1_MIortho2
R2 = 0.99
y~1961.76e^(−1.03x)
8
6.62
5.25
3.88
2.5
present
reduced
absent
0.00
0.25
0.50
0.75
1.00
Distance from M. lignano
loge(Bit Score)
Bristle Status
present
reduced
absent

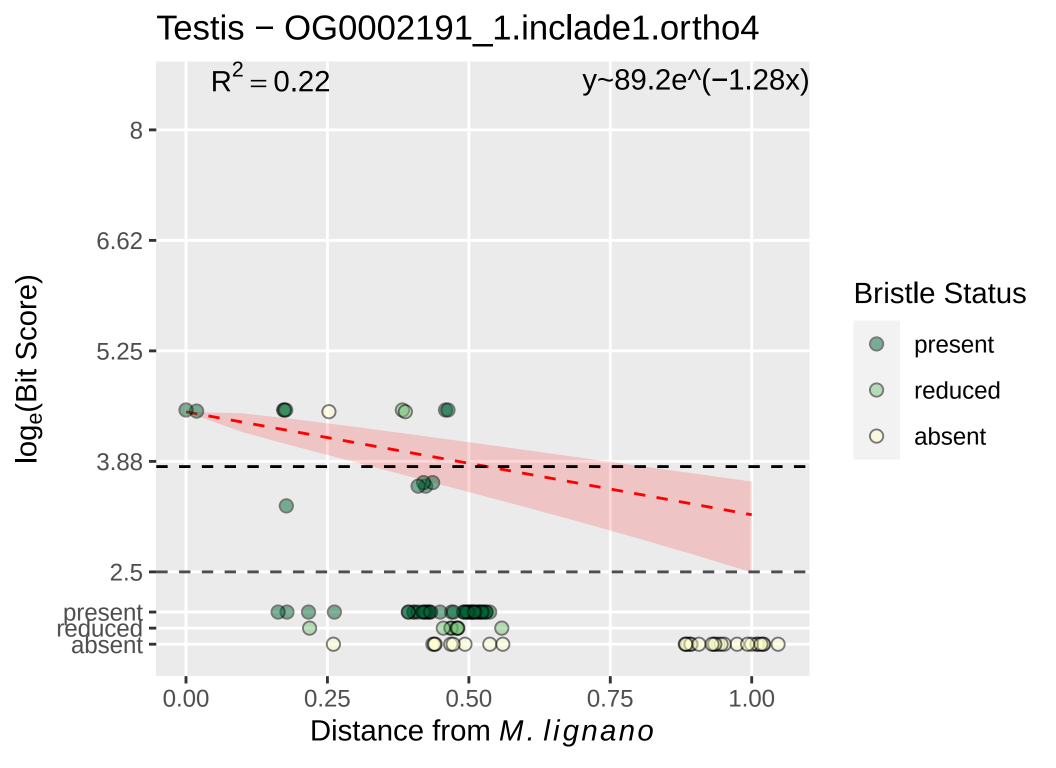

Testis − OG0002191_1.inclade1.ortho4
R^2 = 0.22
y~89.2e^(−1.28x)
8
6.62
5.25
3.88
2.5
present
reduced
absent
0.00
0.25
0.50
0.75
1.00
Distance from M. lignano
log_e(Bit Score)
Bristle Status
present
reduced
absent

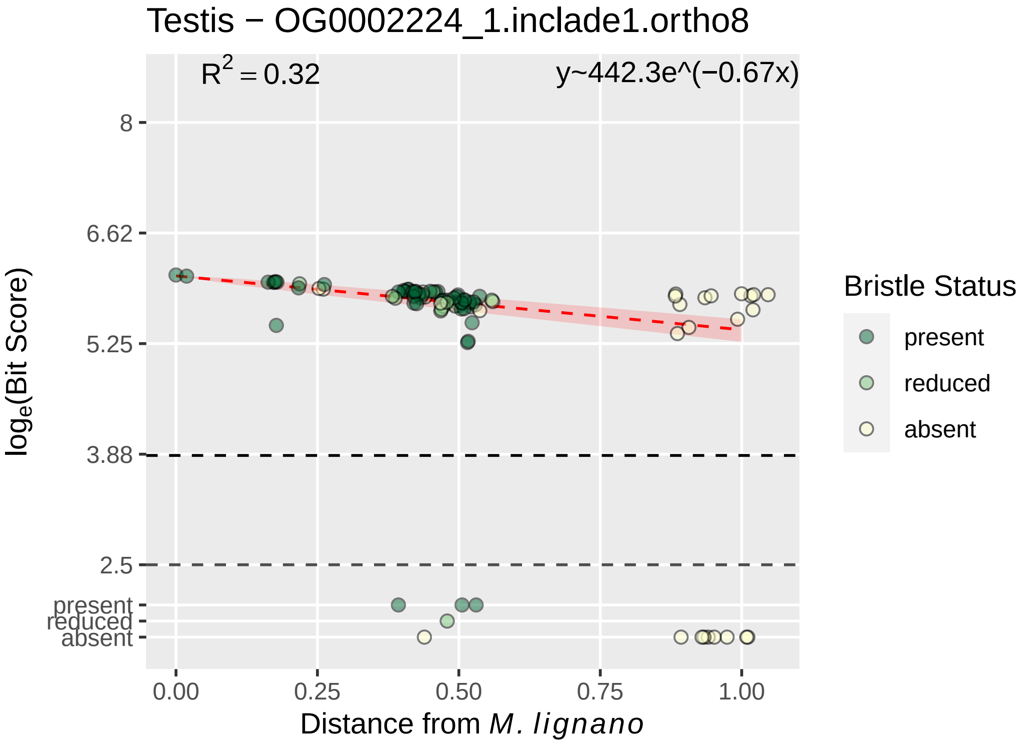

Testis − OG0002224_1.inclade1.ortho8
$R^2 = 0.32$
y~442.3e^(−0.67x)
8
6.62
5.25
3.88
2.5
present
reduced
absent
$\log_e$(Bit Score)
0.00
0.25
0.50
0.75
1.00
Distance from *M. lignano*
Bristle Status
present
reduced
absent

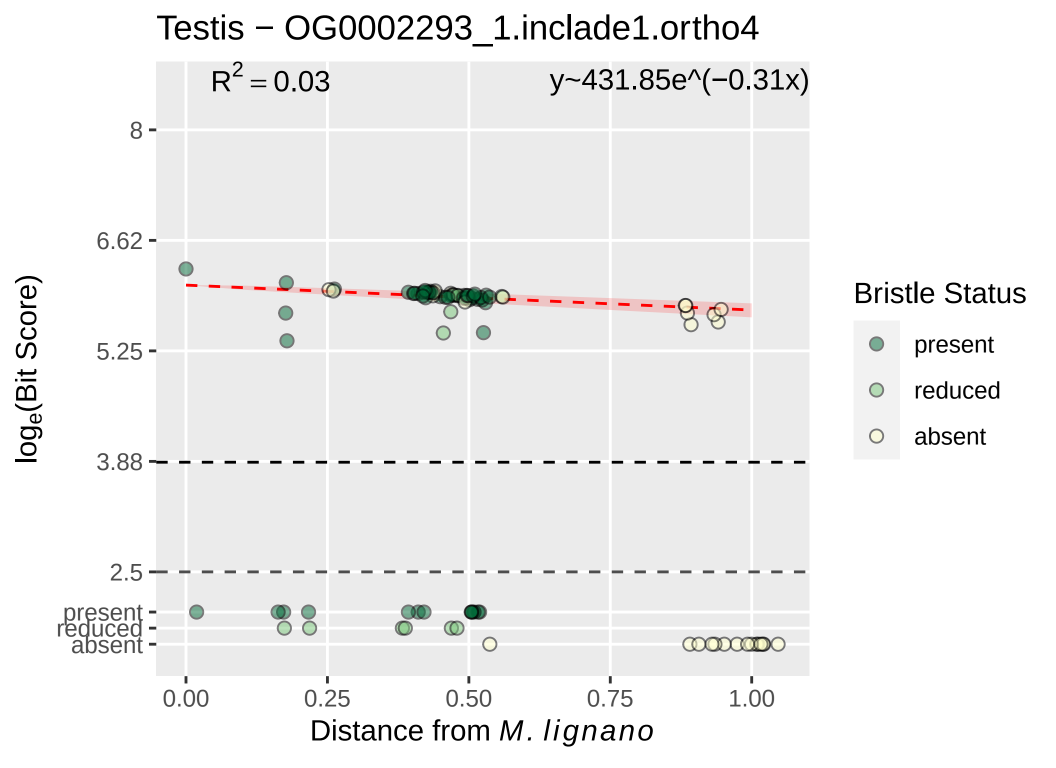

Testis − OG0002293_1.inclade1.ortho4
$R^2 = 0.03$
y~431.85e^(−0.31x)
8
6.62
5.25
3.88
2.5
present
reduced
absent
$log_e$(Bit Score)
0.00
0.25
0.50
0.75
1.00
Distance from *M. lignano*
Bristle Status
present
reduced
absent

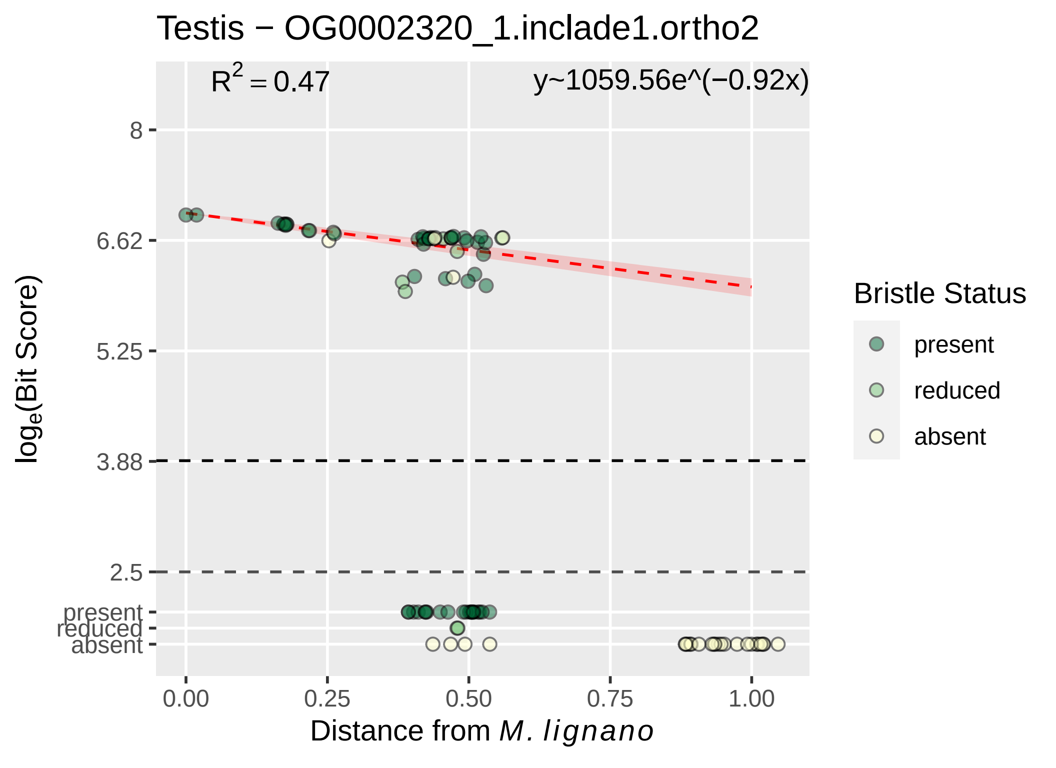

Testis − OG0002320_1.inclade1.ortho2
$R^2 = 0.47$
y~1059.56e^(−0.92x)
8
6.62
5.25
3.88
2.5
present
reduced
absent
$\log_e$(Bit Score)
0.00
0.25
0.50
0.75
1.00
Distance from *M. lignano*
Bristle Status
present
reduced
absent

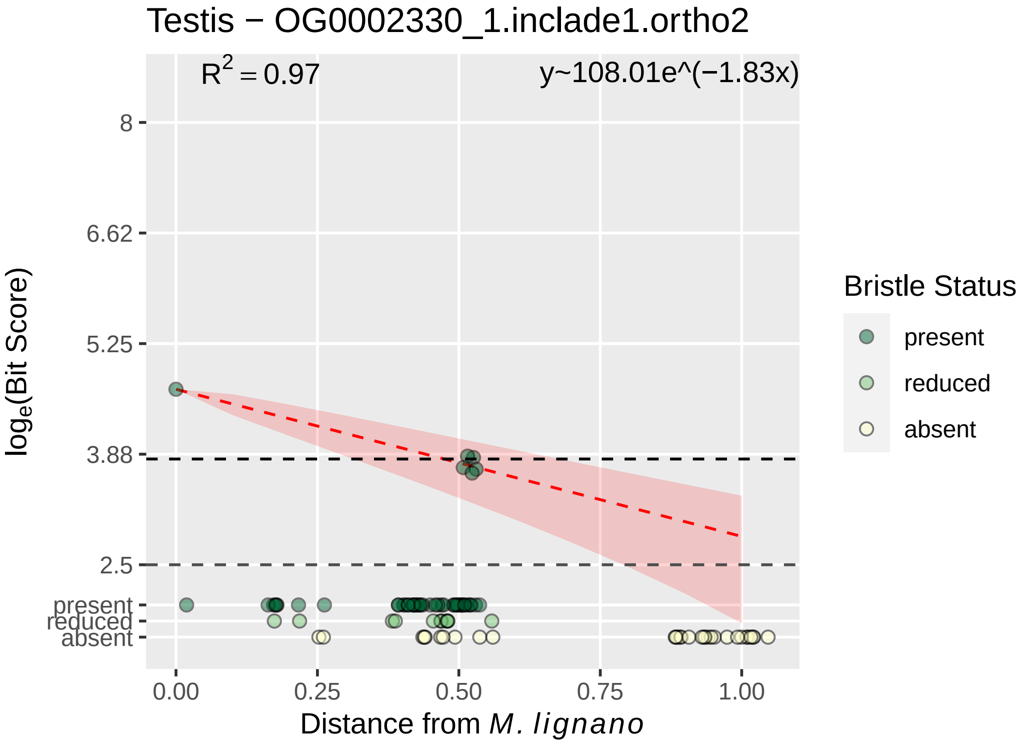

Testis − OG0002330_1.inclade1.ortho2
R^2 = 0.97
y~108.01e^(−1.83x)
8
6.62
5.25
3.88
2.5
present
reduced
absent
log_e(Bit Score)
0.00
0.25
0.50
0.75
1.00
Distance from M. lignano
Bristle Status
present
reduced
absent

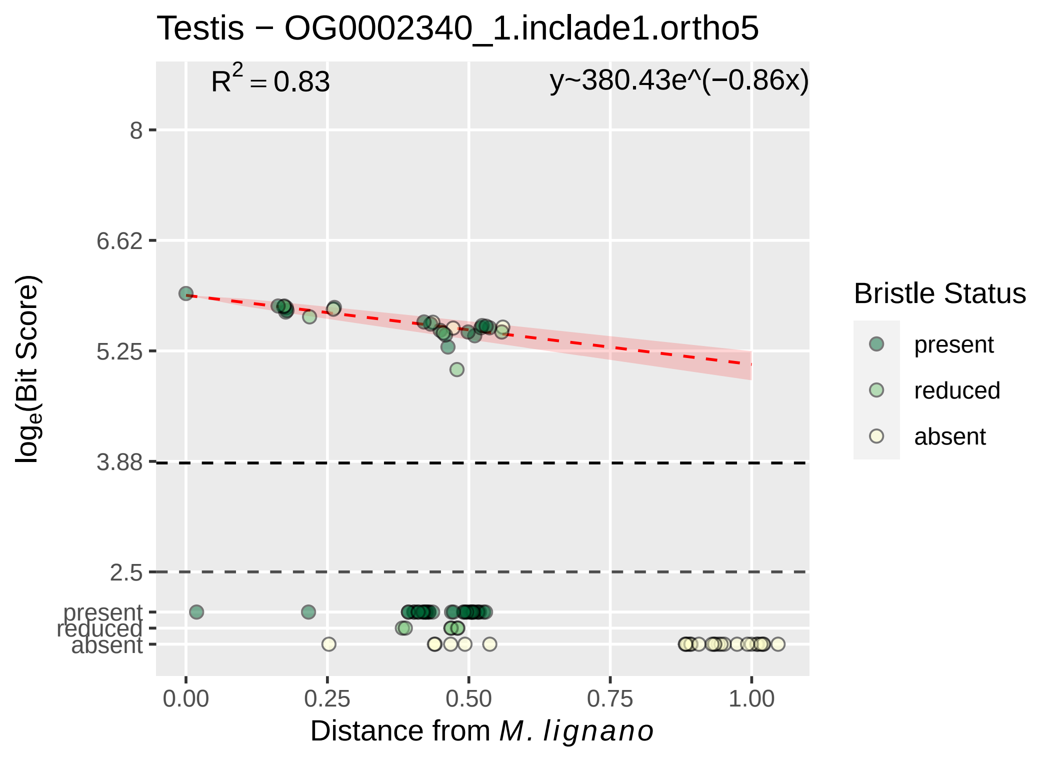
Testis − OG0002340_1.inclade1.ortho5
R² = 0.83
y~380.43e^(−0.86x)
8
6.62
5.25
3.88
2.5
present
reduced
absent
0.00
0.25
0.50
0.75
1.00
log_e(Bit Score)
Distance from M. lignano
Bristle Status
present
reduced
absent

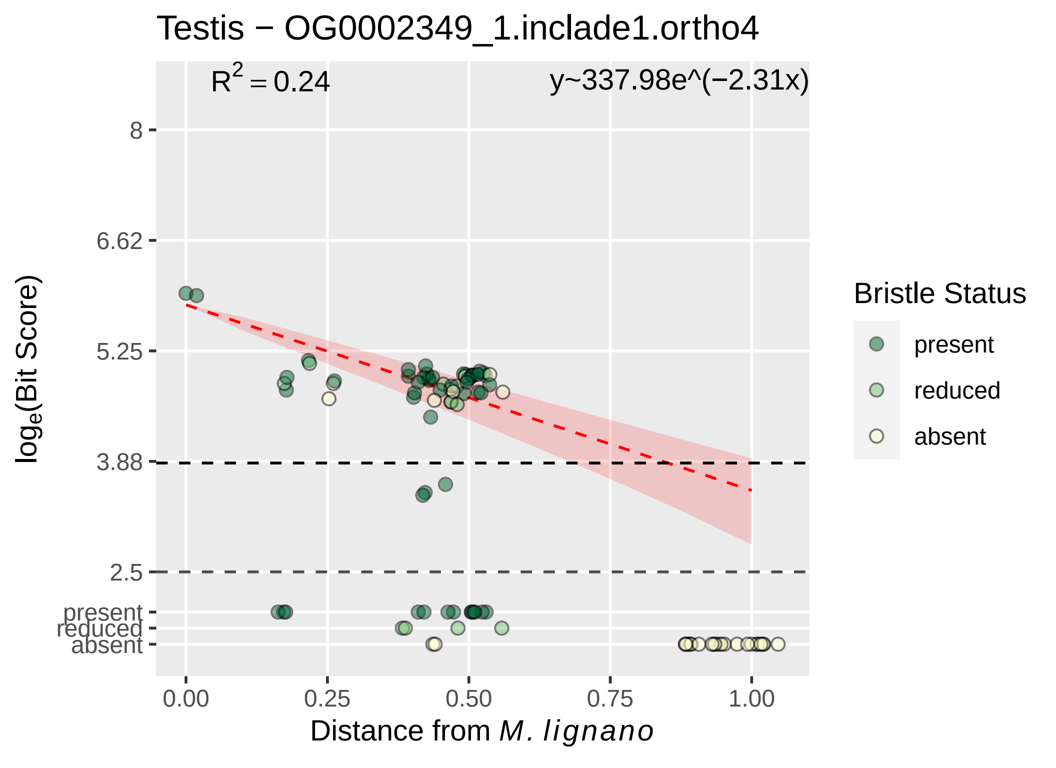

Testis − OG0002349_1.inclade1.ortho4
R2 = 0.24
y~337.98e^(−2.31x)
8
6.62
5.25
3.88
2.5
present
reduced
absent
0.00
0.25
0.50
0.75
1.00
loge(Bit Score)
Distance from M. lignano
Bristle Status
present
reduced
absent

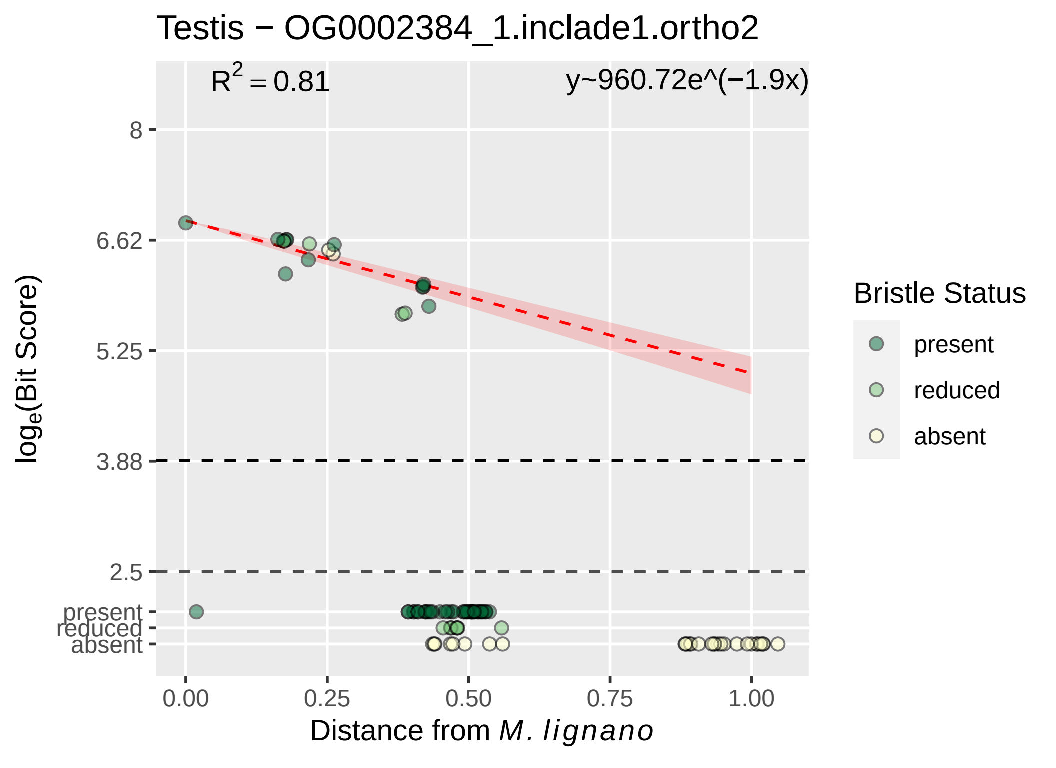

Testis − OG0002384_1.inclade1.ortho2
$R^2 = 0.81$
y~960.72e^(−1.9x)
8
6.62
5.25
3.88
2.5
present
reduced
absent
$\log_e$(Bit Score)
0.00
0.25
0.50
0.75
1.00
Distance from *M. lignano*
Bristle Status
present
reduced
absent

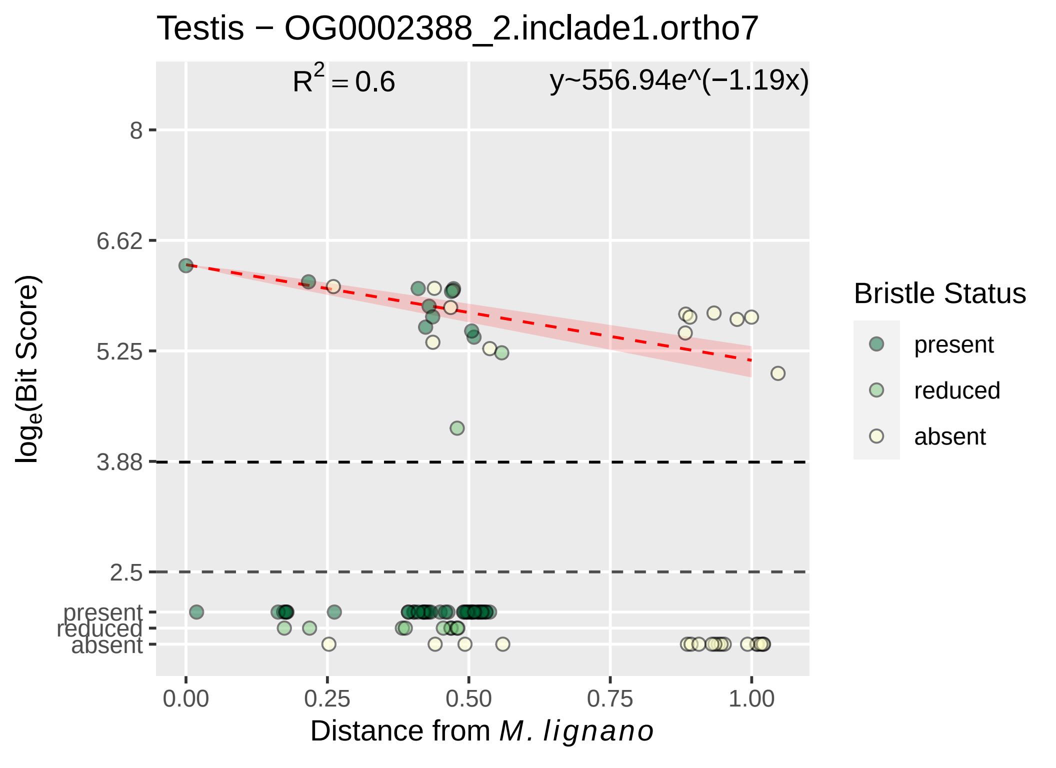

Testis − OG0002388_2.inclade1.ortho7
$R^2 = 0.6$
y~556.94e^(−1.19x)
8
6.62
5.25
3.88
2.5
present
reduced
absent
0.00
0.25
0.50
0.75
1.00
Distance from M. lignano
$\log_e$(Bit Score)
Bristle Status
present
reduced
absent

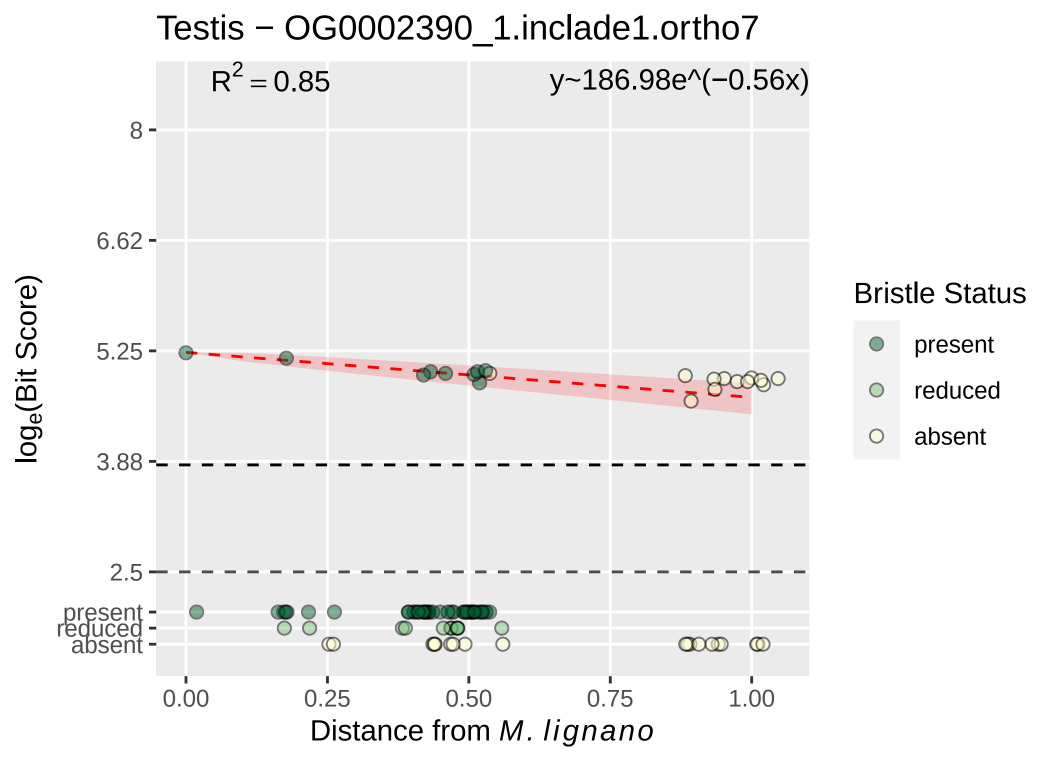

Testis − OG0002390_1.inclade1.ortho7
$R^2 = 0.85$
y~186.98e^(−0.56x)
8
6.62
5.25
3.88
2.5
present
reduced
absent
$\log_e$(Bit Score)
0.00
0.25
0.50
0.75
1.00
Distance from *M. lignano*
Bristle Status
present
reduced
absent

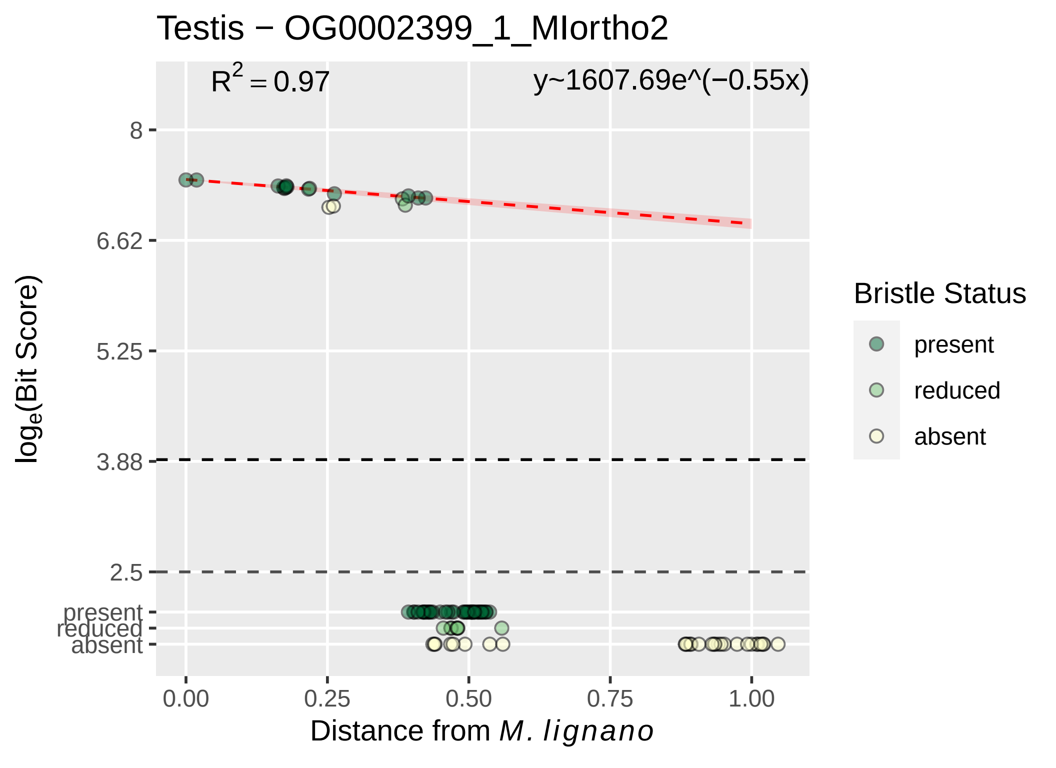

Testis − OG0002399_1_MIortho2
$R^2 = 0.97$
y~1607.69e^(−0.55x)
8
6.62
5.25
3.88
2.5
present
reduced
absent
0.00
0.25
0.50
0.75
1.00
$\log_e$(Bit Score)
Distance from *M. lignano*
Bristle Status
present
reduced
absent

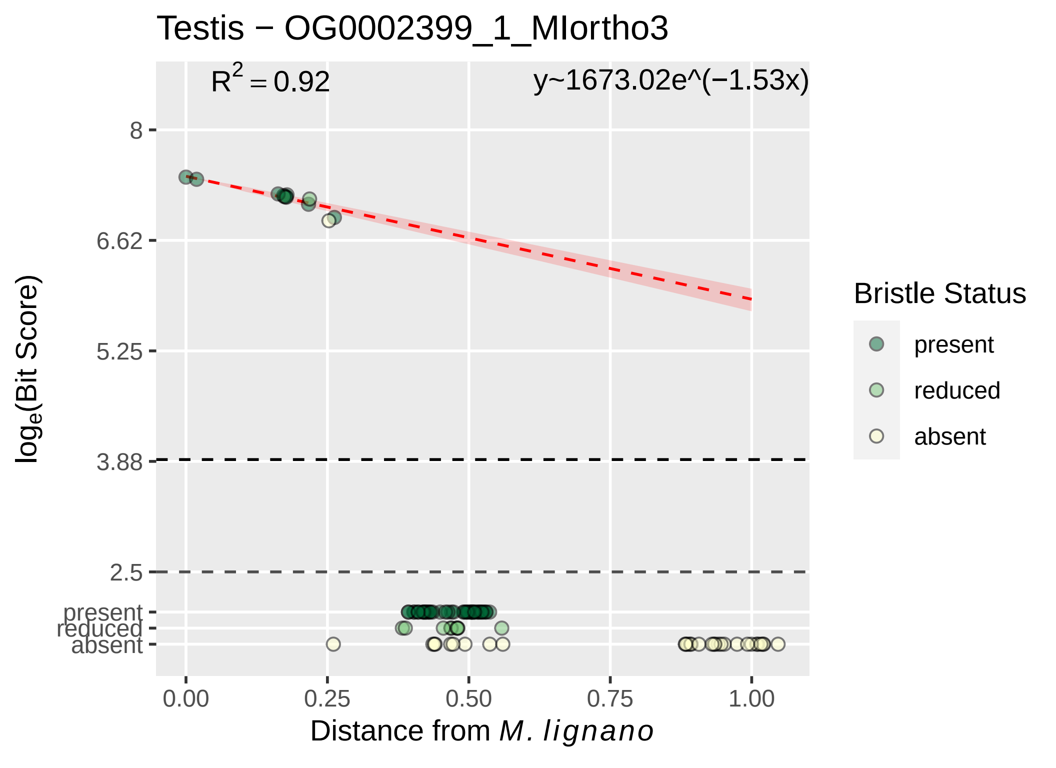

Testis − OG0002399_1_MIortho3
$R^2 = 0.92$
y~1673.02e^(−1.53x)
8
6.62
5.25
3.88
2.5
present
reduced
absent
$\log_e$(Bit Score)
0.00
0.25
0.50
0.75
1.00
Distance from M. lignano
Bristle Status
present
reduced
absent

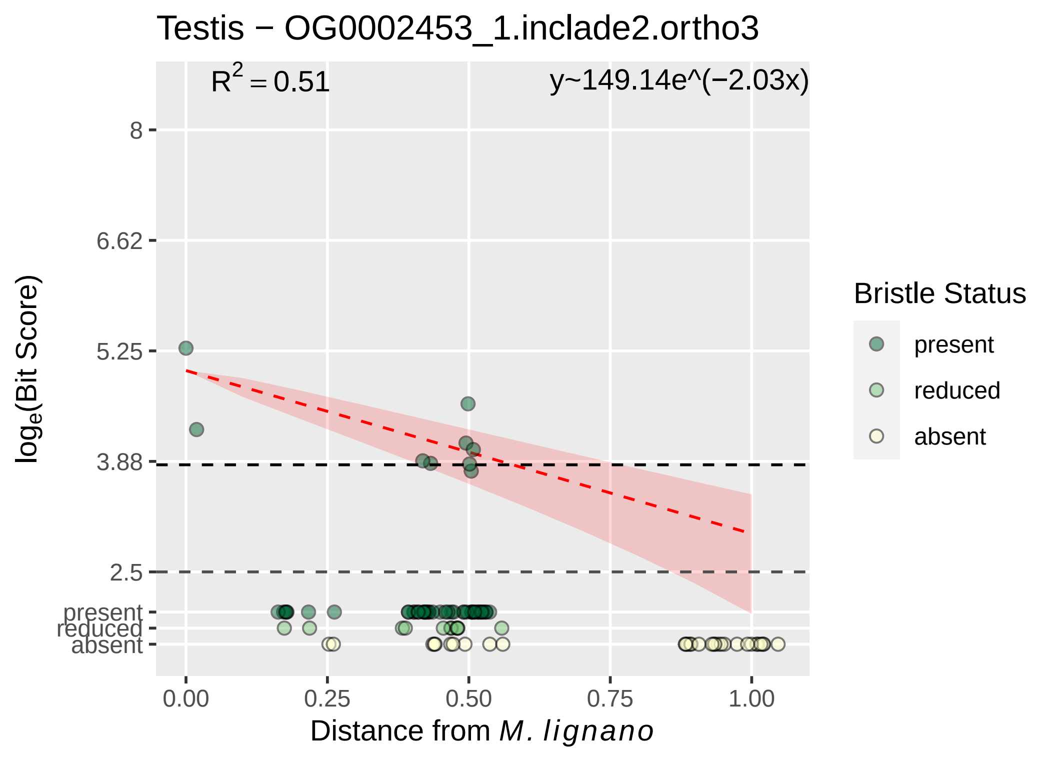

Testis − OG0002453_1.inclade2.ortho3
$R^2 = 0.51$
y~149.14e^(−2.03x)
8
6.62
5.25
3.88
2.5
present
reduced
absent
log_e(Bit Score)
0.00
0.25
0.50
0.75
1.00
Distance from M. lignano
Bristle Status
present
reduced
absent

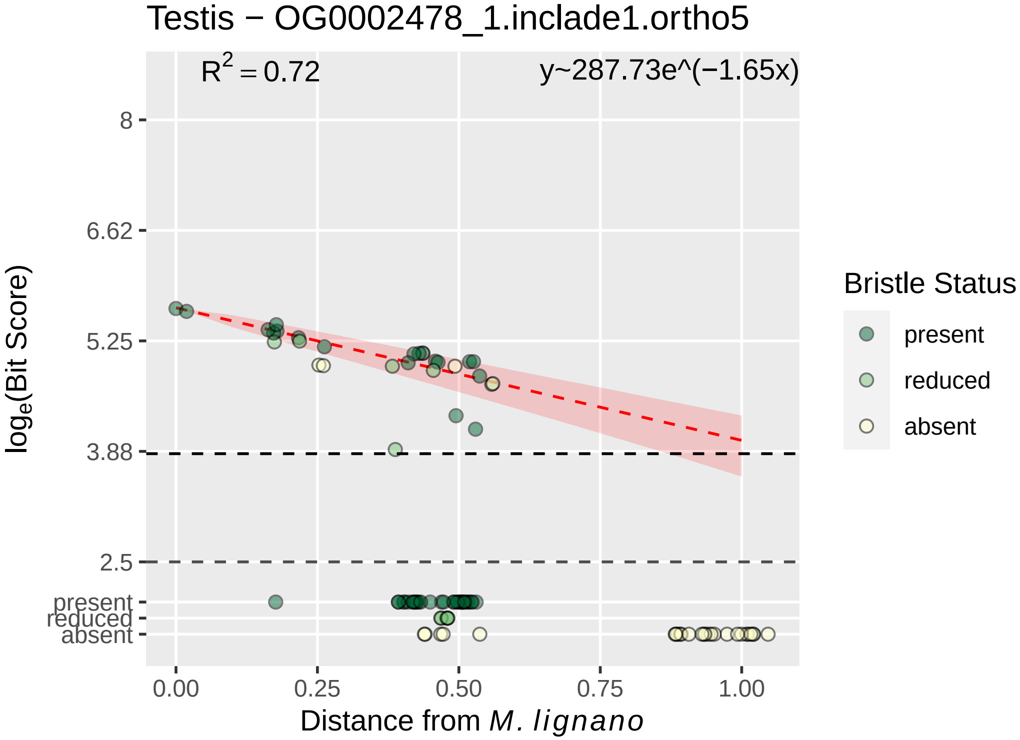

Testis − OG0002478_1.inclade1.ortho5
$R^2 = 0.72$
y~287.73e^(−1.65x)
8
6.62
5.25
3.88
2.5
present
reduced
absent
0.00
0.25
0.50
0.75
1.00
Distance from M. lignano
log_e(Bit Score)
Bristle Status
present
reduced
absent

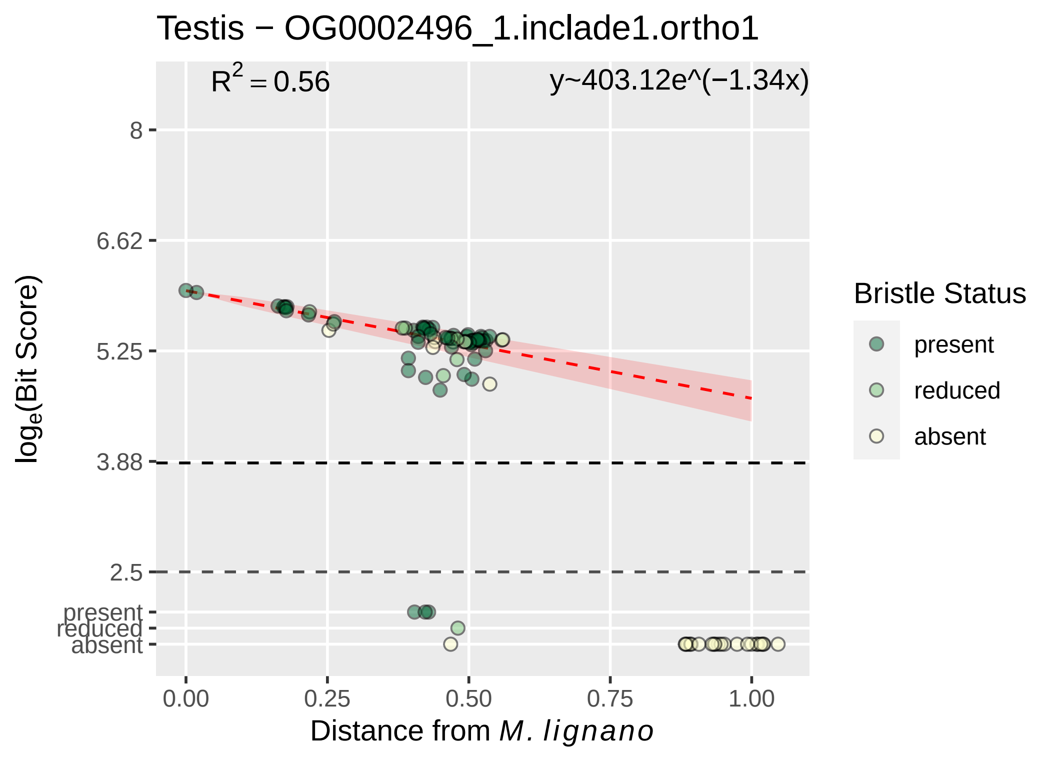
Testis − OG0002496_1.inclade1.ortho1
R2 = 0.56
y~403.12e^(−1.34x)
8
6.62
5.25
3.88
2.5
present
reduced
absent
loge(Bit Score)
0.00
0.25
0.50
0.75
1.00
Distance from M. lignano
Bristle Status
present
reduced
absent

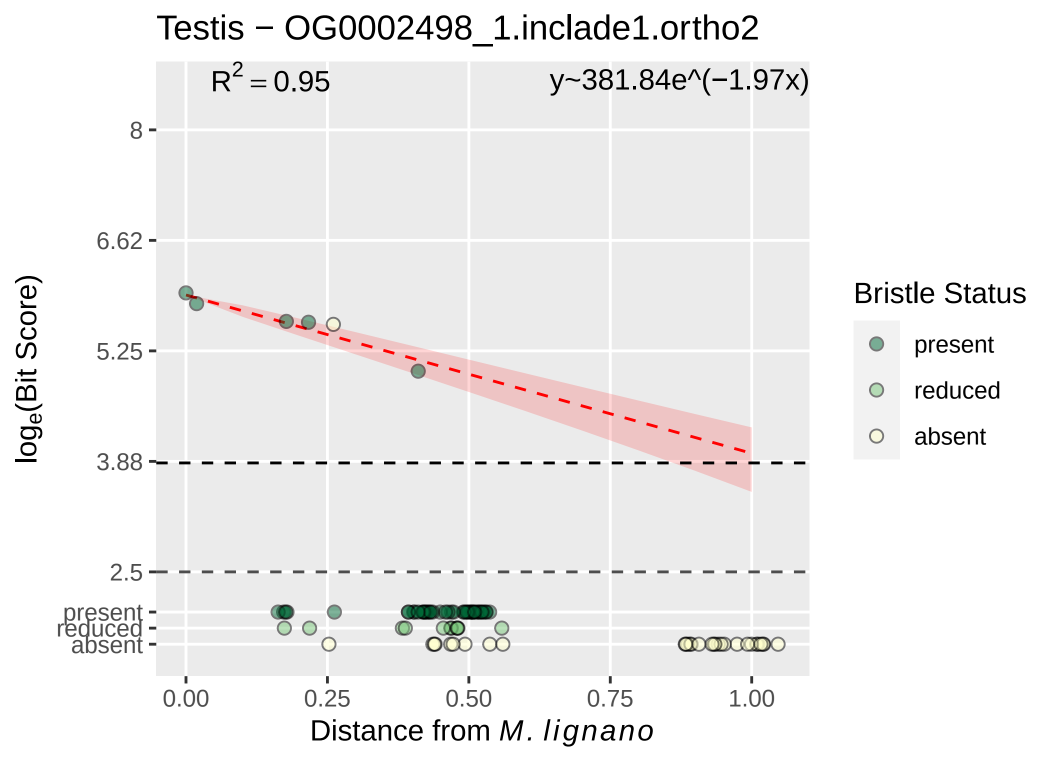

Testis − OG0002498_1.inclade1.ortho2
R^2 = 0.95
y~381.84e^(−1.97x)
8
6.62
5.25
3.88
2.5
present
reduced
absent
loge(Bit Score)
0.00
0.25
0.50
0.75
1.00
Distance from M. lignano
Bristle Status
present
reduced
absent

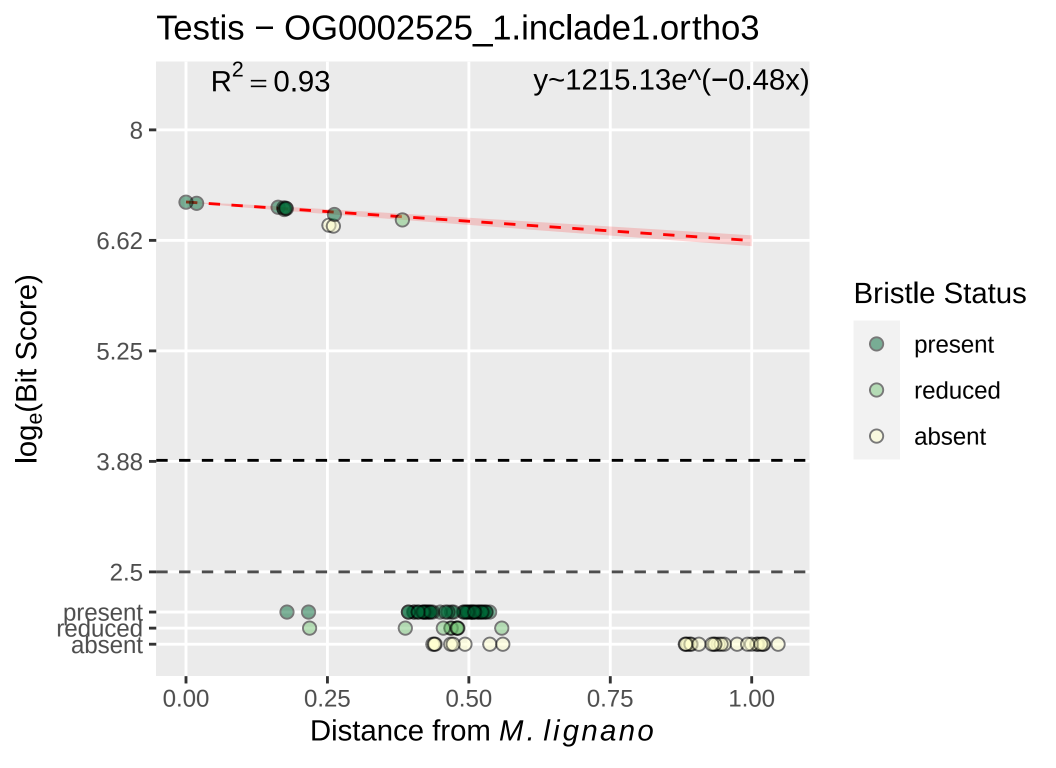
Testis − OG0002525_1.inclade1.ortho3
$R^2 = 0.93$
y~1215.13e^(−0.48x)
8
6.62
5.25
3.88
2.5
present
reduced
absent
0.00
0.25
0.50
0.75
1.00
log$_e$(Bit Score)
Distance from *M. lignano*
Bristle Status
present
reduced
absent

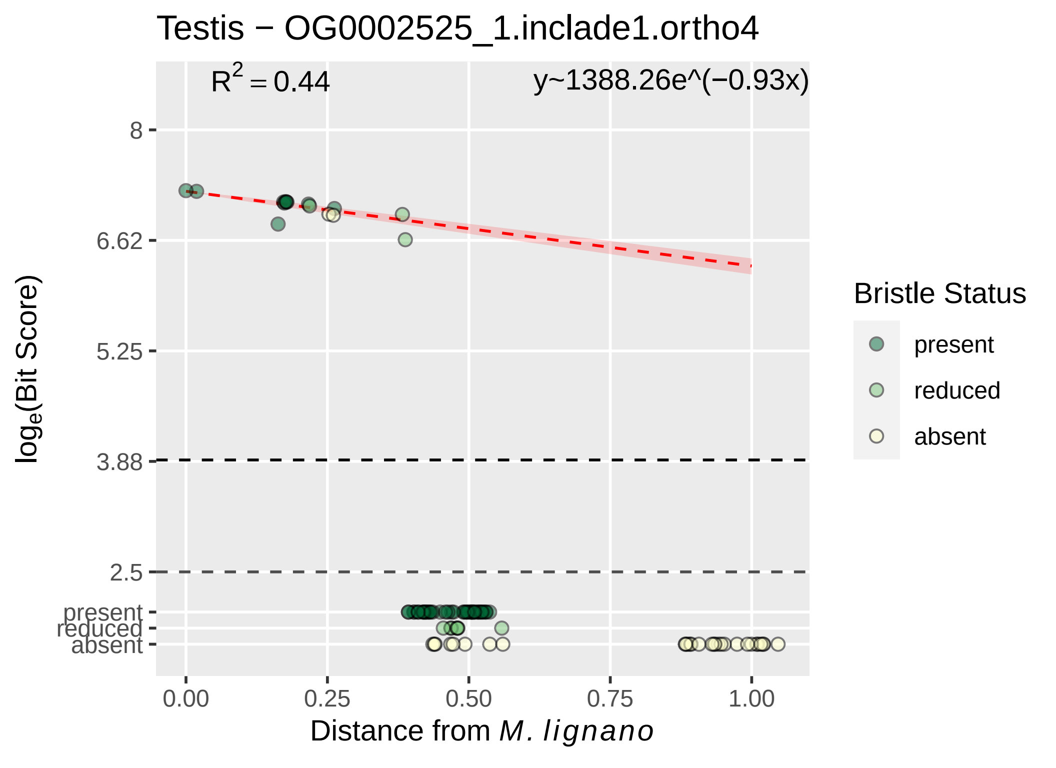

Testis − OG0002525_1.inclade1.ortho4
R² = 0.44
y~1388.26e^(−0.93x)
8
6.62
5.25
3.88
2.5
present
reduced
absent
loge(Bit Score)
0.00
0.25
0.50
0.75
1.00
Distance from M. lignano
Bristle Status
present
reduced
absent

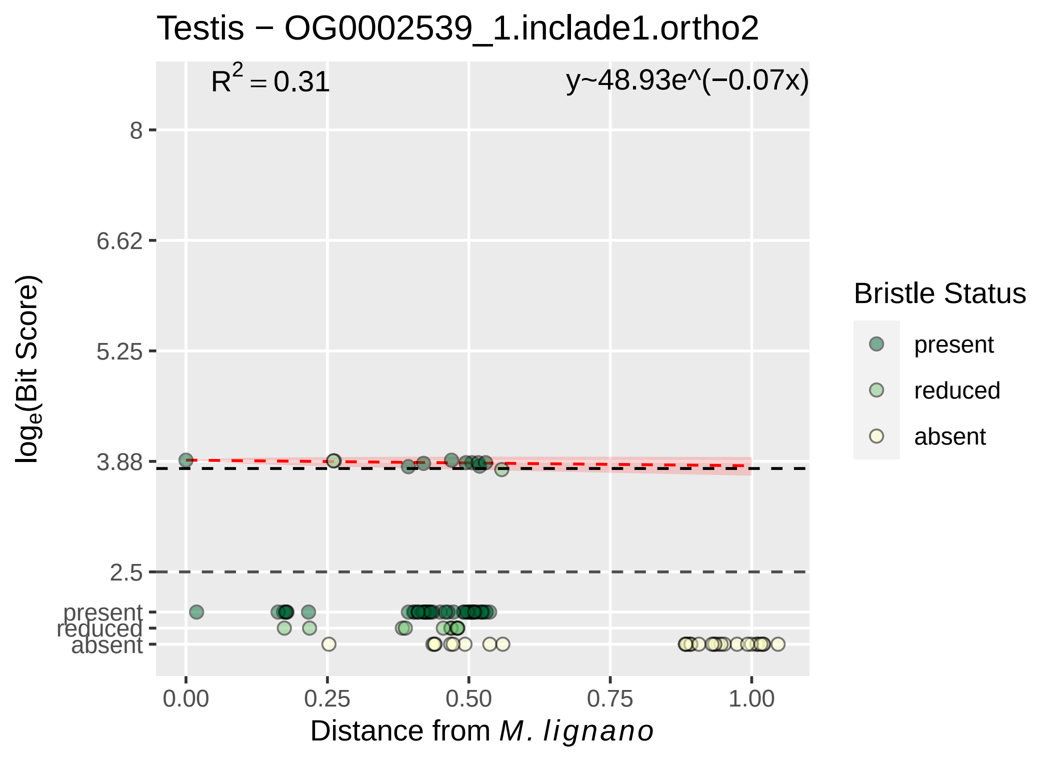

Testis − OG0002539_1.inclade1.ortho2
R^2 = 0.31
y~48.93e^(−0.07x)
8
6.62
5.25
3.88
2.5
present
reduced
absent
0.00
0.25
0.50
0.75
1.00
Distance from M. lignano
log_e(Bit Score)
Bristle Status
present
reduced
absent

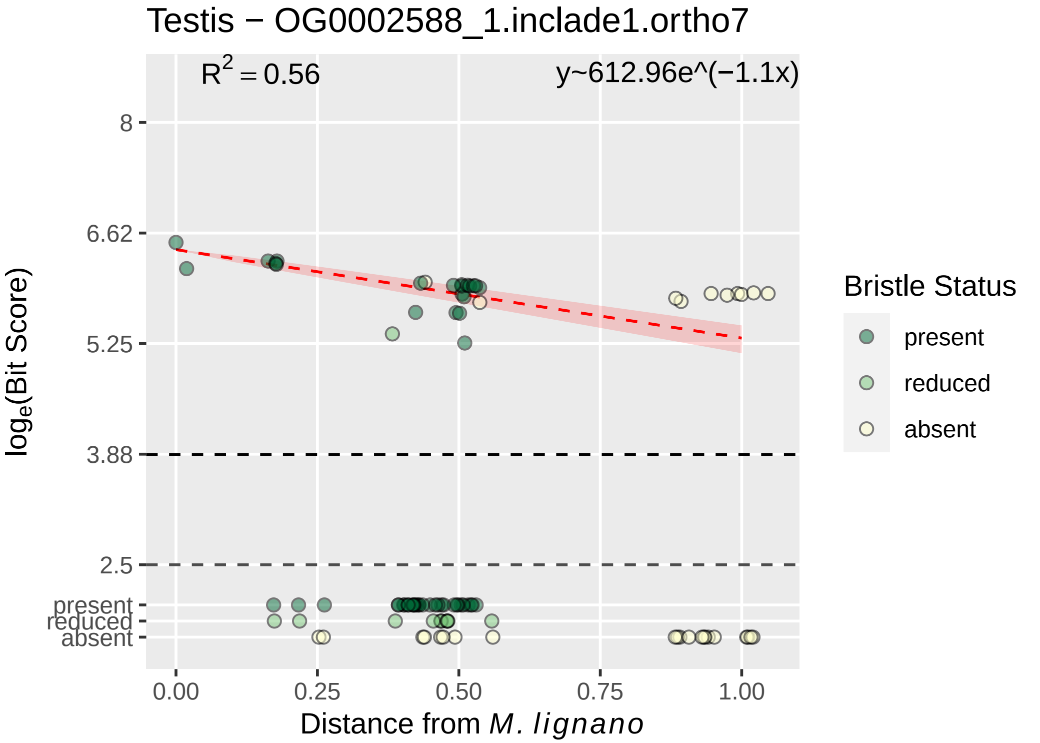

Testis − OG0002588_1.inclade1.ortho7
R^2 = 0.56
y~612.96e^(−1.1x)
8
6.62
5.25
3.88
2.5
present
reduced
absent
log_e(Bit Score)
0.00
0.25
0.50
0.75
1.00
Distance from M. lignano
Bristle Status
present
reduced
absent

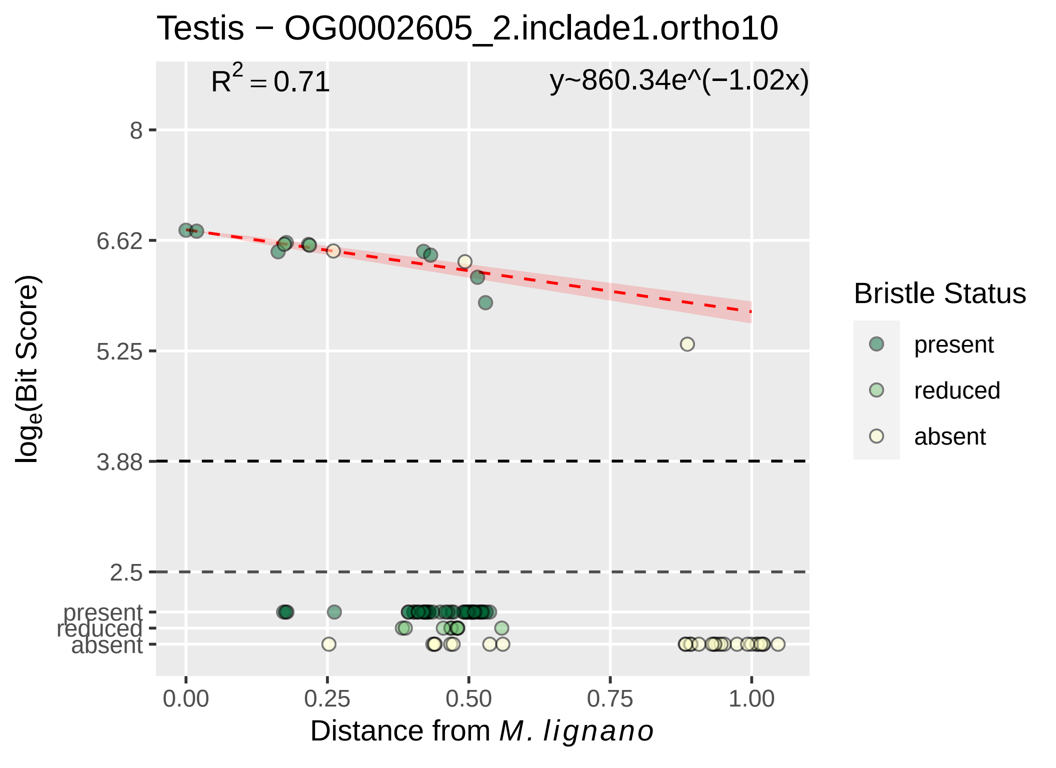

Testis − OG0002605_2.inclade1.ortho10
$R^2 = 0.71$
y~860.34e^(−1.02x)
8
6.62
5.25
3.88
2.5
present
reduced
absent
$\log_e$(Bit Score)
0.00
0.25
0.50
0.75
1.00
Distance from *M. lignano*
Bristle Status
present
reduced
absent

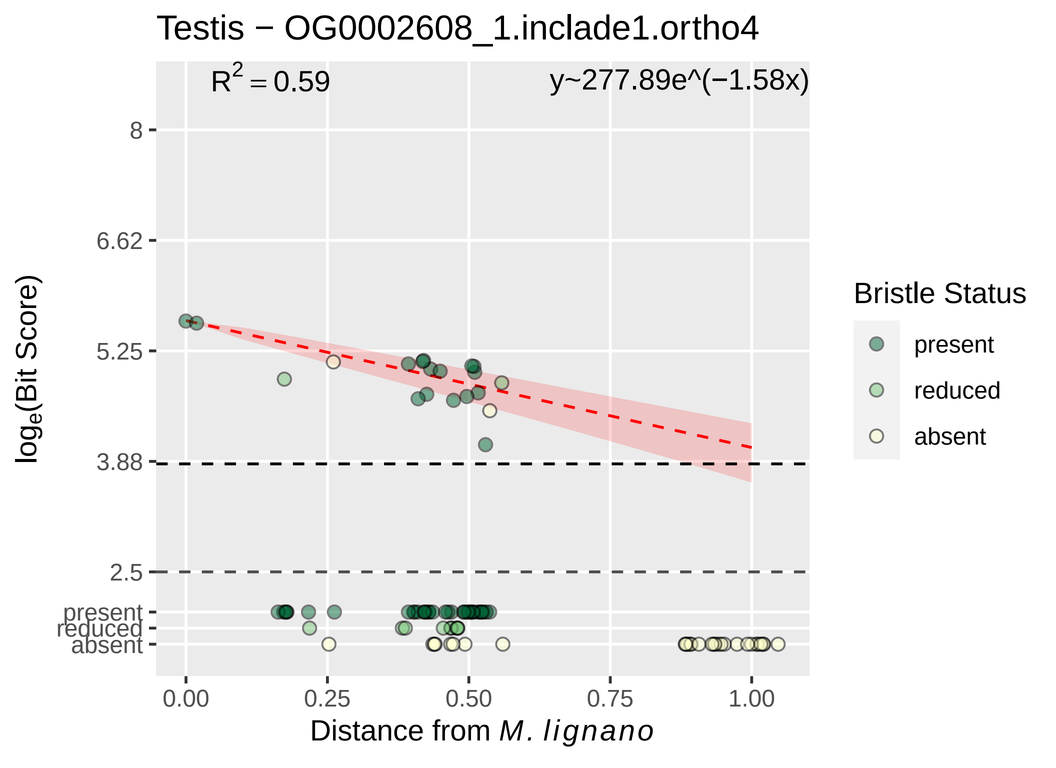

Testis − OG0002608_1.inclade1.ortho4
R^2 = 0.59
y~277.89e^(−1.58x)
8
6.62
5.25
3.88
2.5
present
reduced
absent
log_e(Bit Score)
0.00
0.25
0.50
0.75
1.00
Distance from M. lignano
Bristle Status
present
reduced
absent

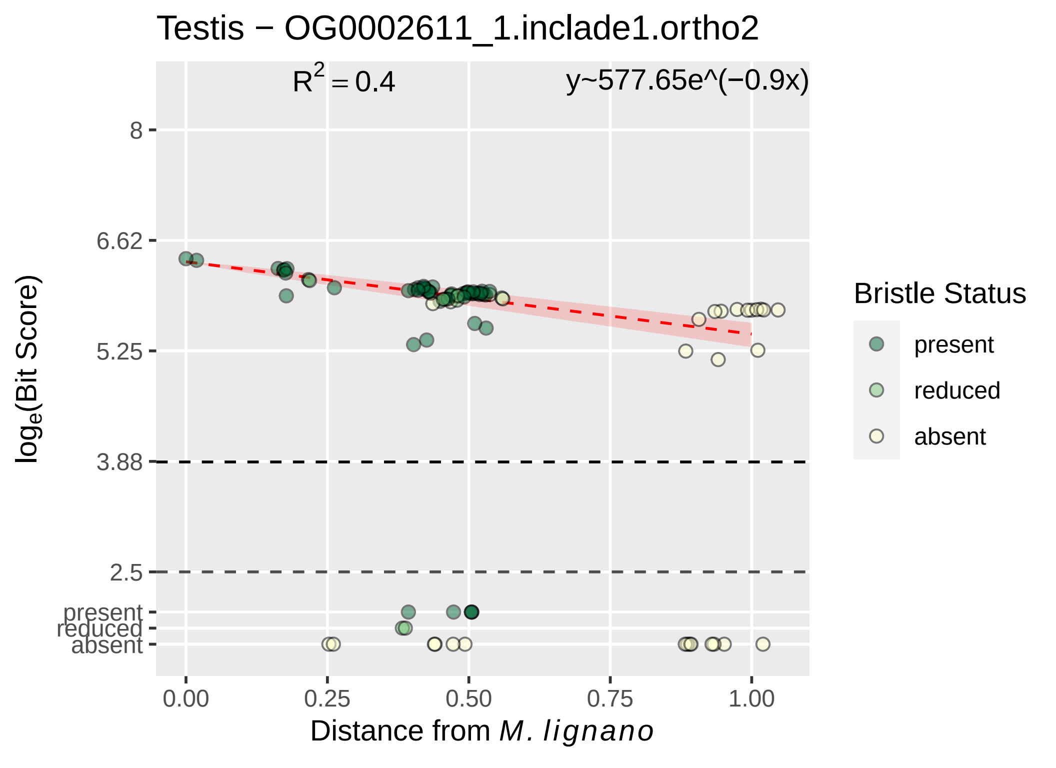

Testis − OG0002611_1.inclade1.ortho2
$R^2 = 0.4$
y~577.65e^(−0.9x)
8
6.62
5.25
3.88
2.5
present
reduced
absent
0.00
0.25
0.50
0.75
1.00
$log_e$(Bit Score)
Distance from *M. lignano*
Bristle Status
present
reduced
absent

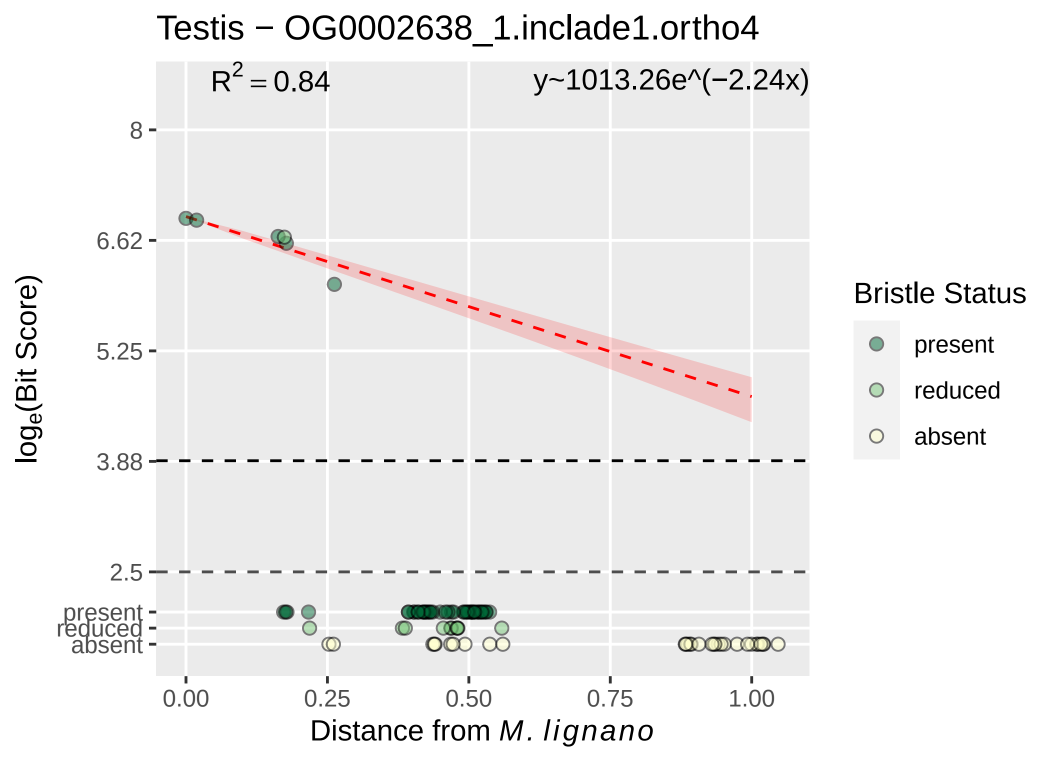
Testis − OG0002638_1.inclade1.ortho4
$R^2 = 0.84$
y~1013.26e^(−2.24x)
8
6.62
5.25
3.88
2.5
present
reduced
absent
0.00
0.25
0.50
0.75
1.00
Distance from M. lignano
loge(Bit Score)
Bristle Status
present
reduced
absent

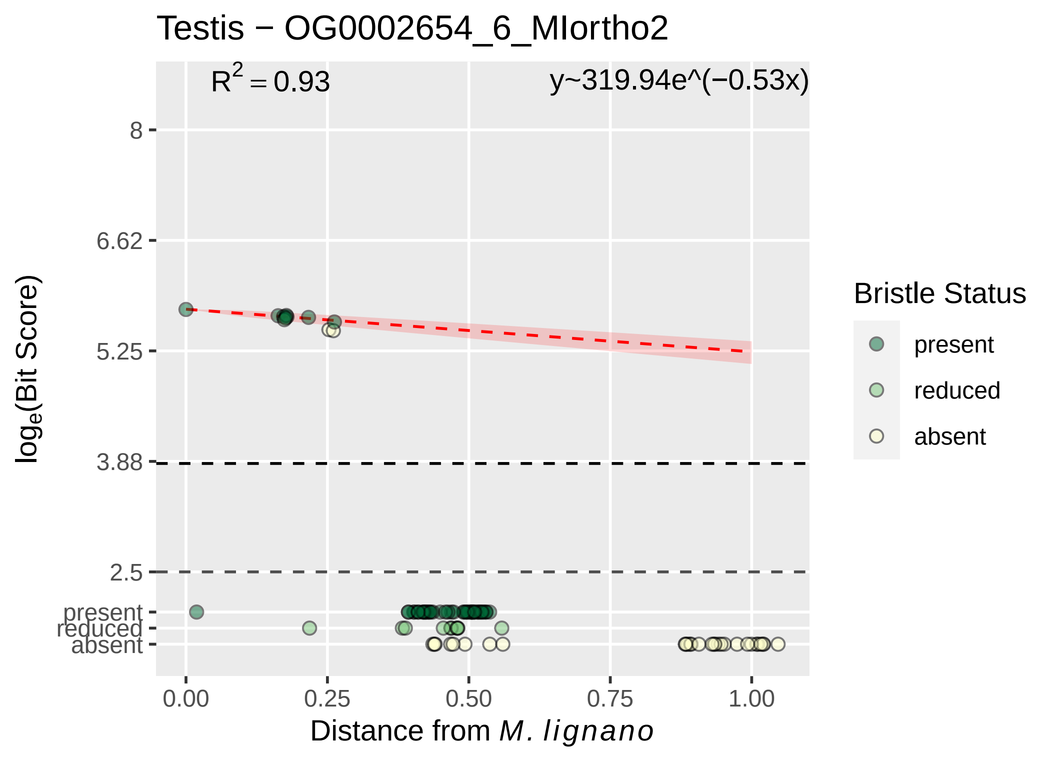

Testis − OG0002654_6_MIortho2
$R^2 = 0.93$
y~319.94e^(−0.53x)
8
6.62
5.25
3.88
2.5
present
reduced
absent
$\log_e$(Bit Score)
0.00
0.25
0.50
0.75
1.00
Distance from *M. lignano*
Bristle Status
present
reduced
absent

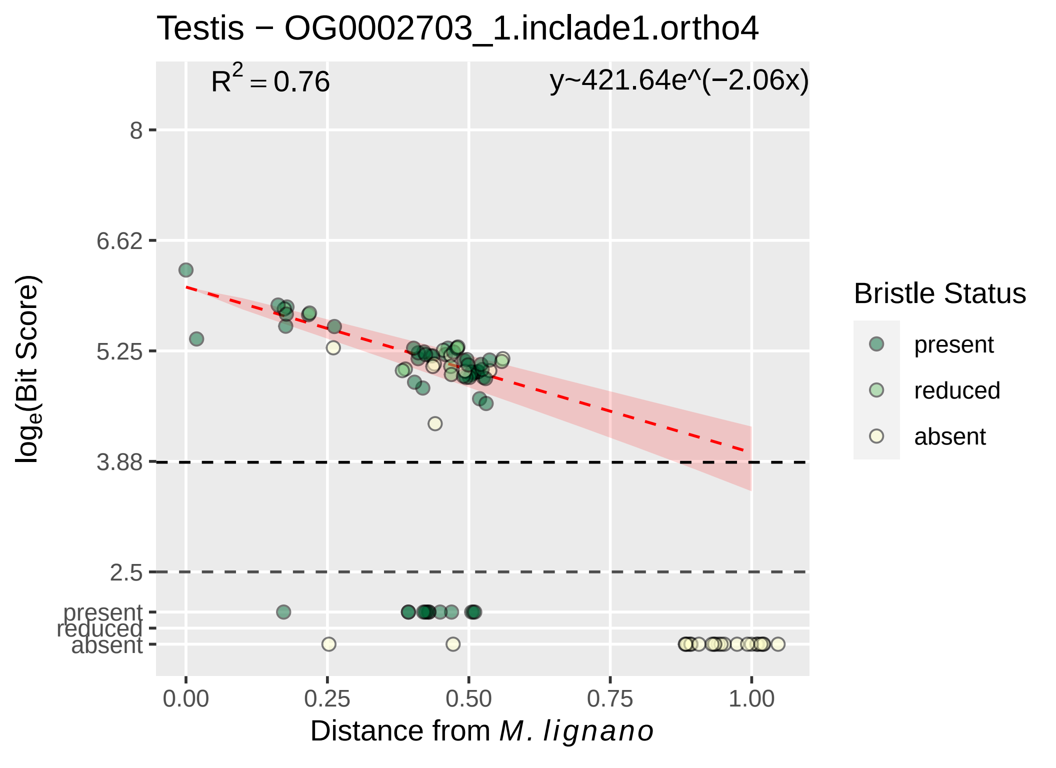

Testis − OG0002703_1.inclade1.ortho4
$R^2 = 0.76$
y~421.64e^(−2.06x)
8
6.62
5.25
3.88
2.5
present
reduced
absent
0.00
0.25
0.50
0.75
1.00
Distance from *M. lignano*
$\log_e$(Bit Score)
Bristle Status
present
reduced
absent

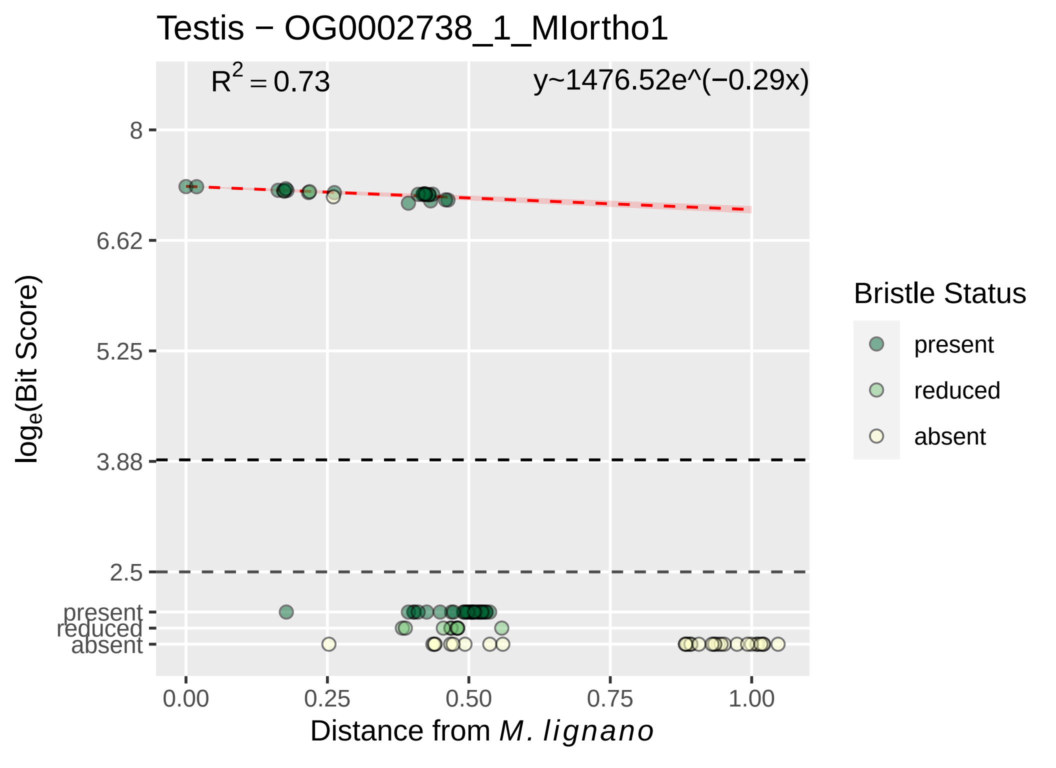
Testis − OG0002738_1_MIortho1
R^2 = 0.73
y~1476.52e^(−0.29x)
8
6.62
5.25
3.88
2.5
present
reduced
absent
log_e(Bit Score)
0.00
0.25
0.50
0.75
1.00
Distance from M. lignano
Bristle Status
present
reduced
absent

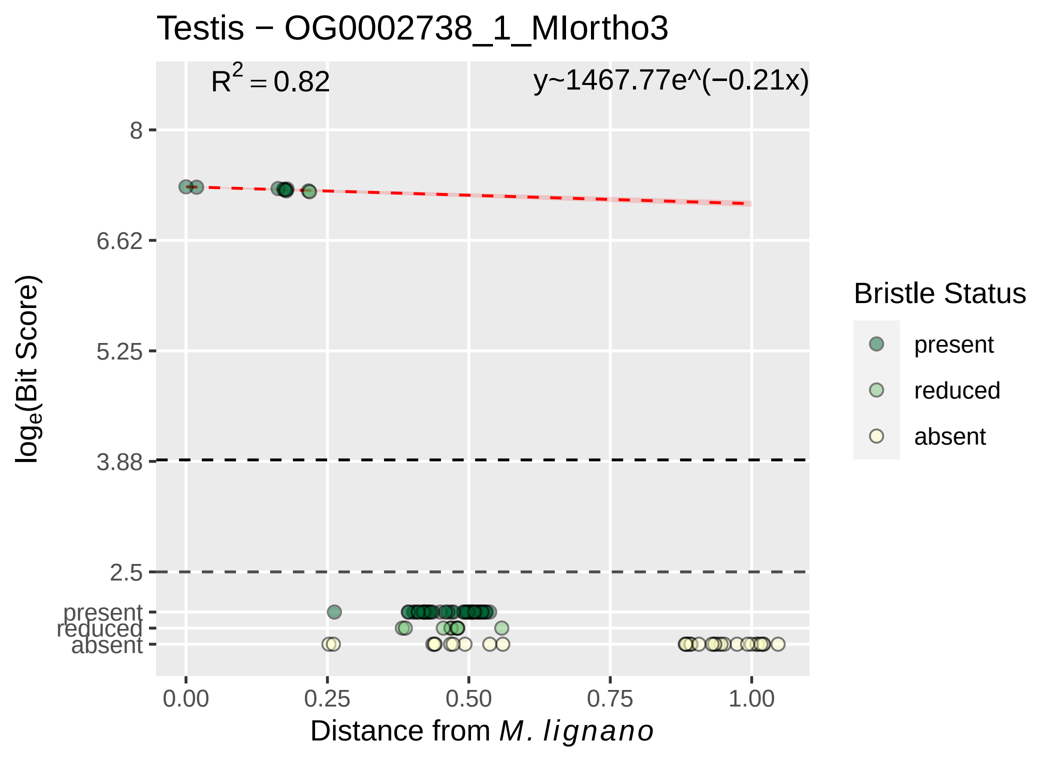

Testis − OG0002738_1_MIortho3
R^2 = 0.82
y~1467.77e^(−0.21x)
8
6.62
5.25
3.88
2.5
present
reduced
absent
log_e(Bit Score)
0.00
0.25
0.50
0.75
1.00
Distance from M. lignano
Bristle Status
present
reduced
absent

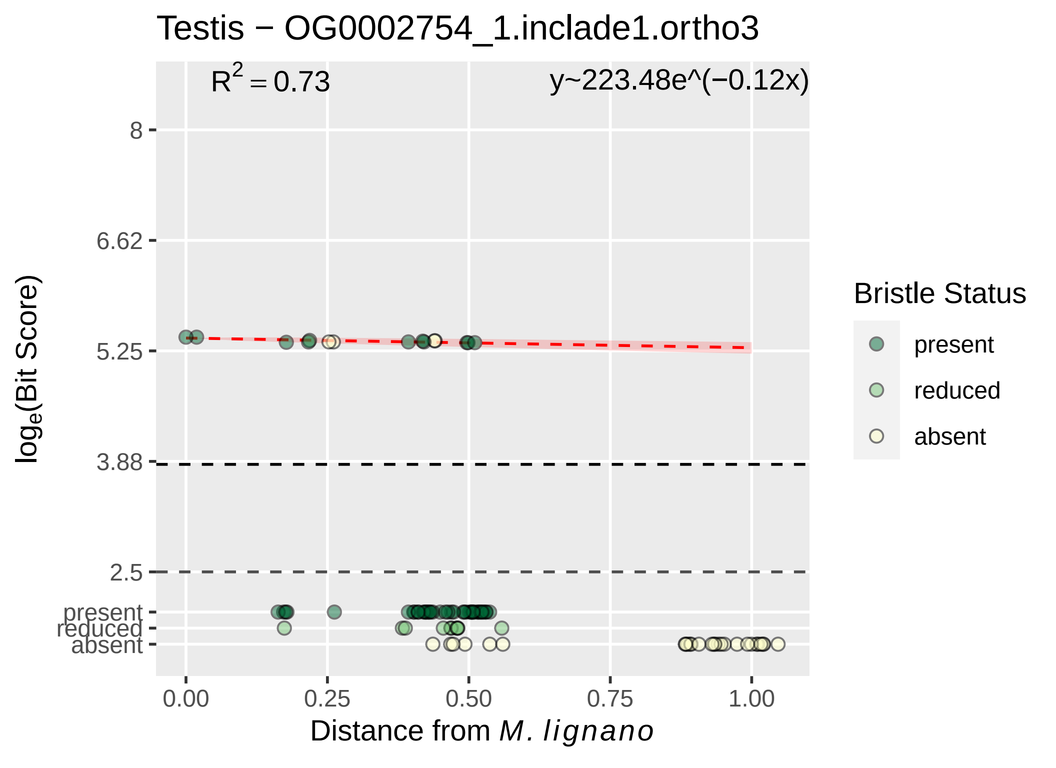

Testis − OG0002754_1.inclade1.ortho3
R2 = 0.73
y~223.48e^(−0.12x)
8
6.62
5.25
3.88
2.5
present
reduced
absent
0.00
0.25
0.50
0.75
1.00
Distance from M. lignano
loge(Bit Score)
Bristle Status
present
reduced
absent

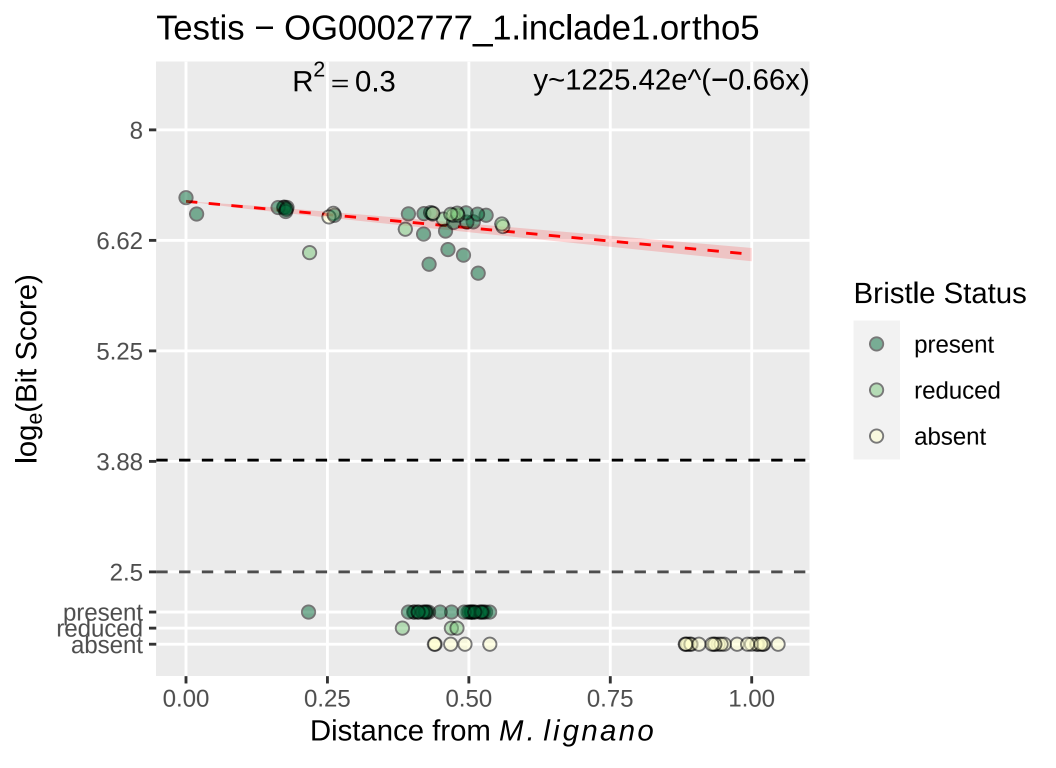

Testis − OG0002777_1.inclade1.ortho5
$R^2 = 0.3$
y~1225.42e^(−0.66x)
8
6.62
5.25
3.88
2.5
present
reduced
absent
0.00
0.25
0.50
0.75
1.00
Distance from *M. lignano*
$\log_e$(Bit Score)
Bristle Status
present
reduced
absent

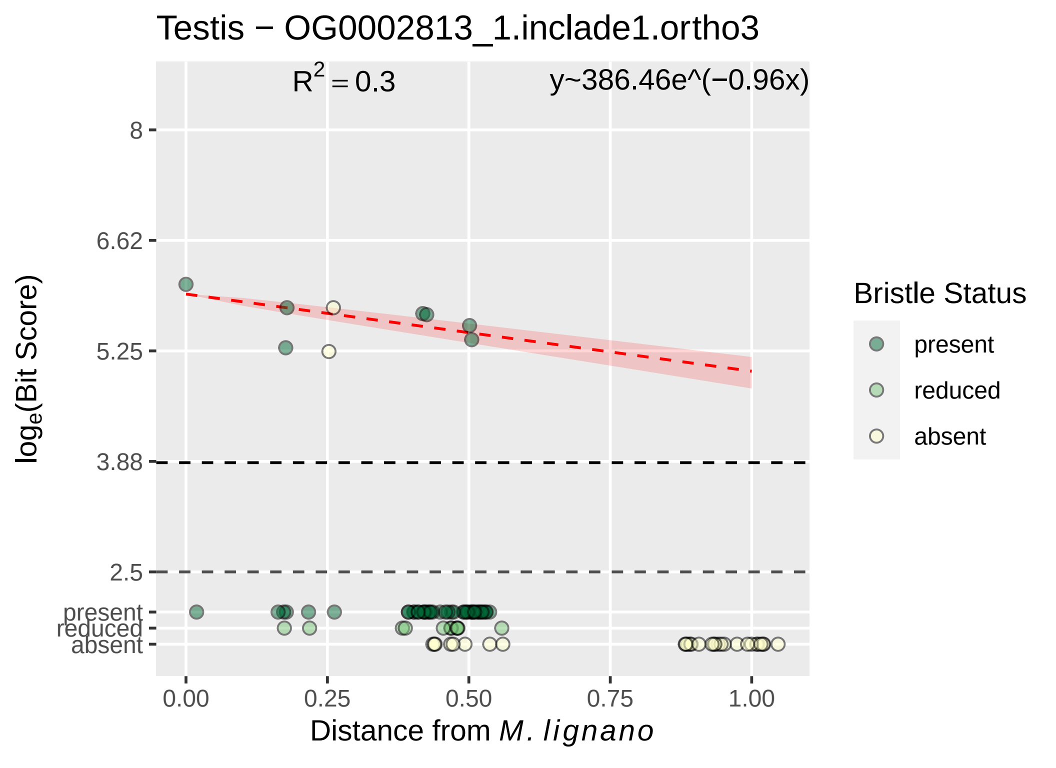

Testis − OG0002813_1.inclade1.ortho3
R^2 = 0.3
y~386.46e^(−0.96x)
8
6.62
5.25
3.88
2.5
present
reduced
absent
log_e(Bit Score)
0.00
0.25
0.50
0.75
1.00
Distance from M. lignano
Bristle Status
present
reduced
absent

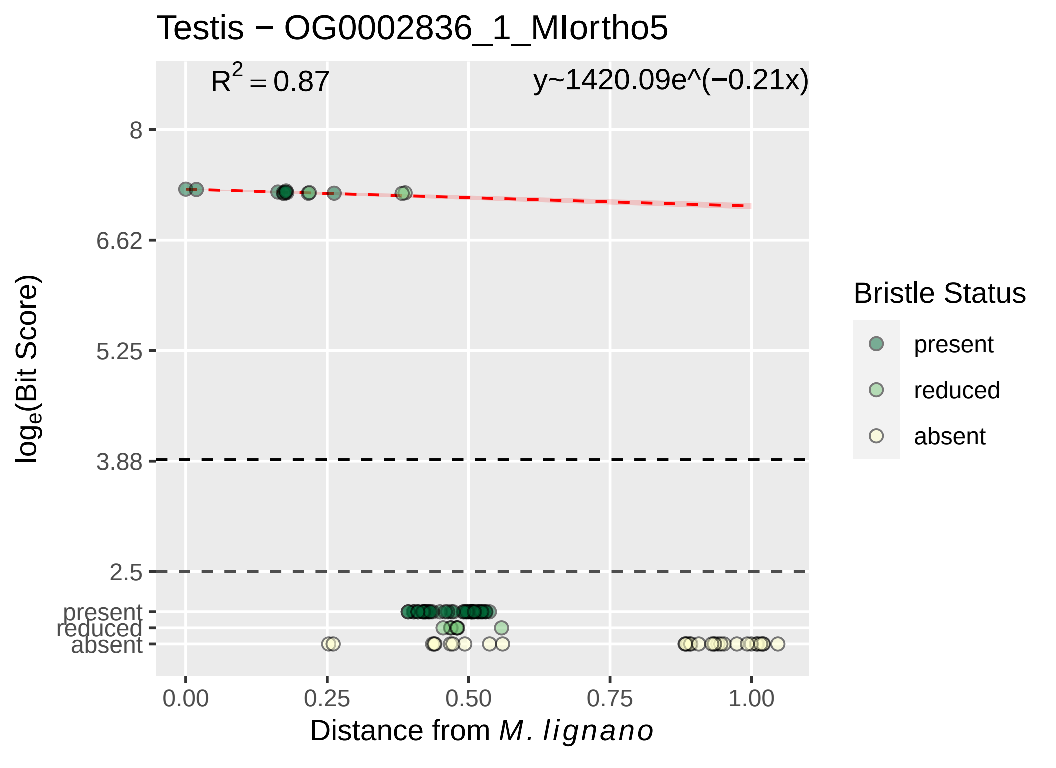

Testis − OG0002836_1_MIortho5
R2 = 0.87
y~1420.09e^(−0.21x)
8
6.62
5.25
3.88
2.5
present
reduced
absent
0.00
0.25
0.50
0.75
1.00
Distance from M. lignano
loge(Bit Score)
Bristle Status
present
reduced
absent

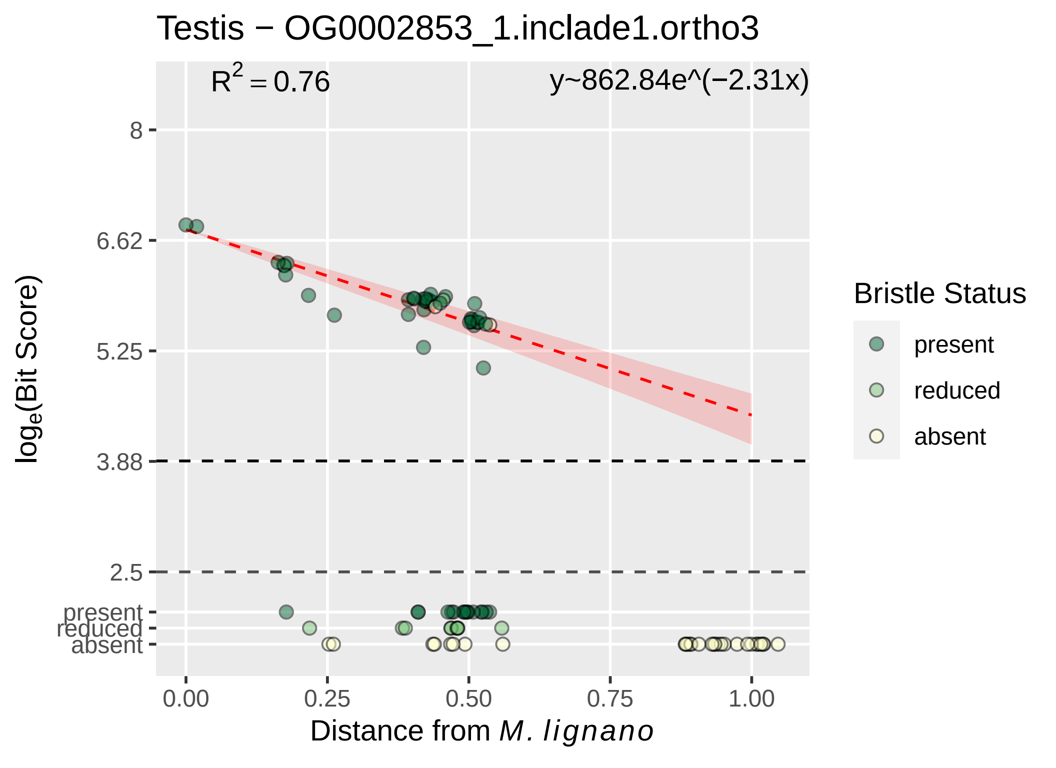

Testis − OG0002853_1.inclade1.ortho3
$R^2 = 0.76$
y~862.84e^(−2.31x)
8
6.62
5.25
3.88
2.5
present
reduced
absent
$\log_e$(Bit Score)
0.00
0.25
0.50
0.75
1.00
Distance from *M. lignano*
Bristle Status
present
reduced
absent

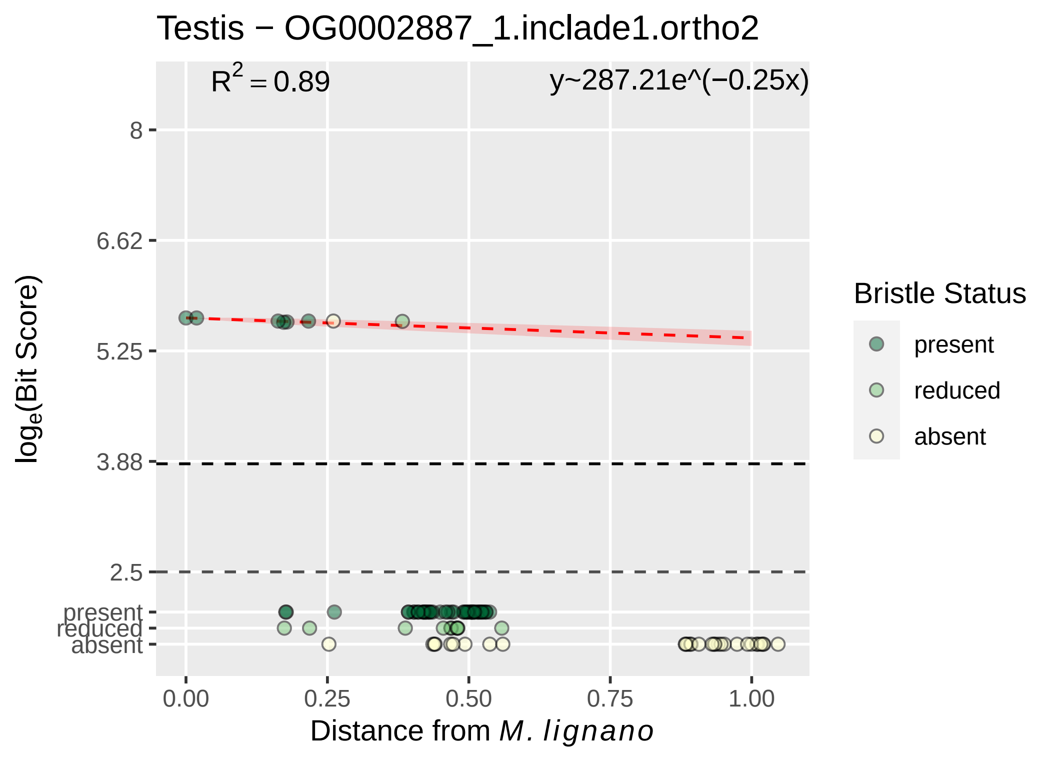
Testis − OG0002887_1.inclade1.ortho2
$R^2 = 0.89$
y~287.21e^(−0.25x)
8
6.62
5.25
3.88
2.5
present
reduced
absent
$\log_e$(Bit Score)
0.00
0.25
0.50
0.75
1.00
Distance from *M. lignano*
Bristle Status
present
reduced
absent

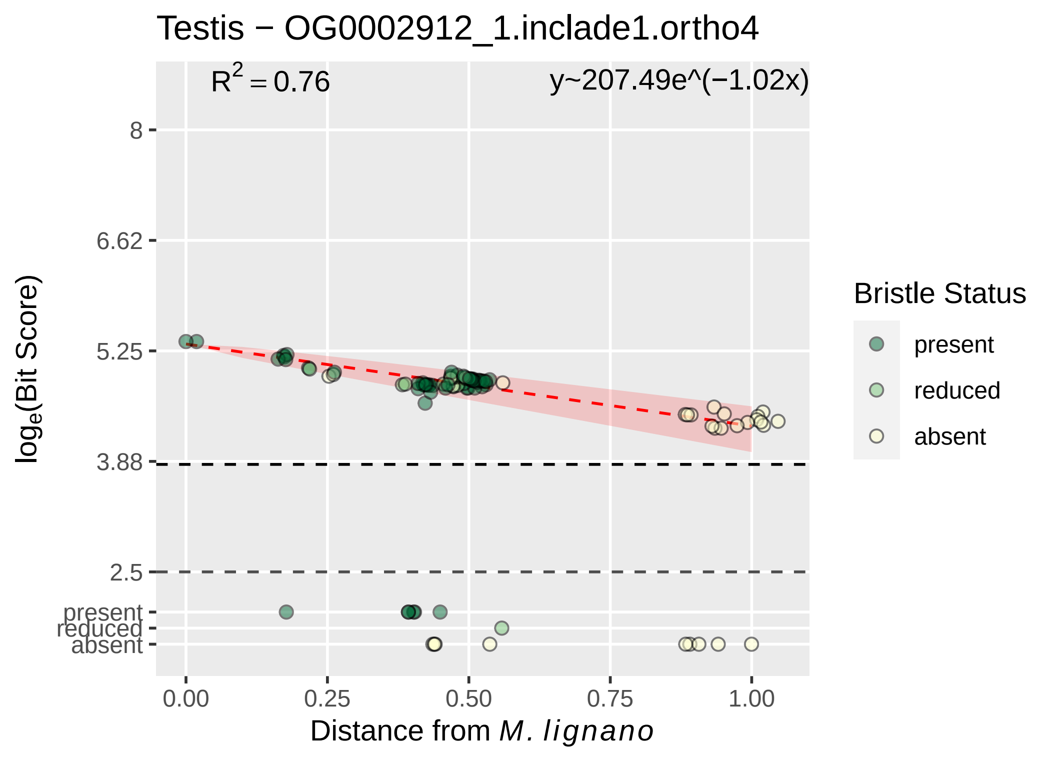

Testis − OG0002912_1.inclade1.ortho4
R² = 0.76
y~207.49e^(−1.02x)
8
6.62
5.25
3.88
2.5
present
reduced
absent
log_e(Bit Score)
0.00
0.25
0.50
0.75
1.00
Distance from M. lignano
Bristle Status
present
reduced
absent

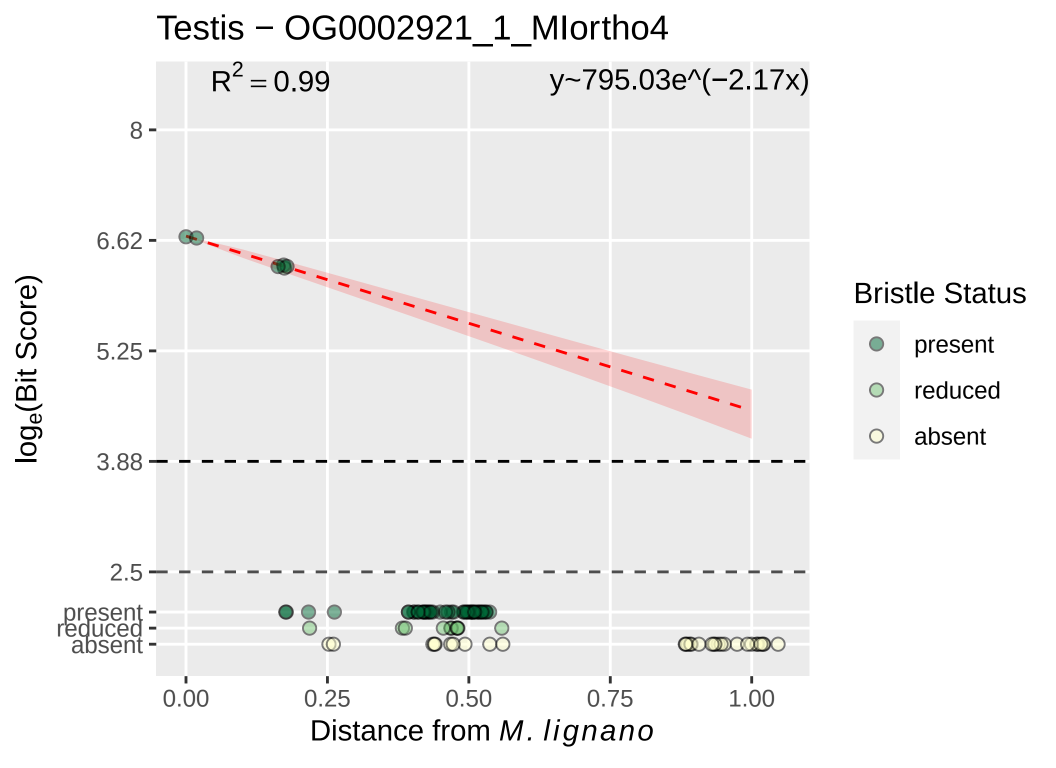

Testis − OG0002921_1_MIortho4
$R^2 = 0.99$
y~795.03e^(−2.17x)
8
6.62
5.25
3.88
2.5
present
reduced
absent
0.00
0.25
0.50
0.75
1.00
Distance from *M. lignano*
$\log_e$(Bit Score)
Bristle Status
present
reduced
absent

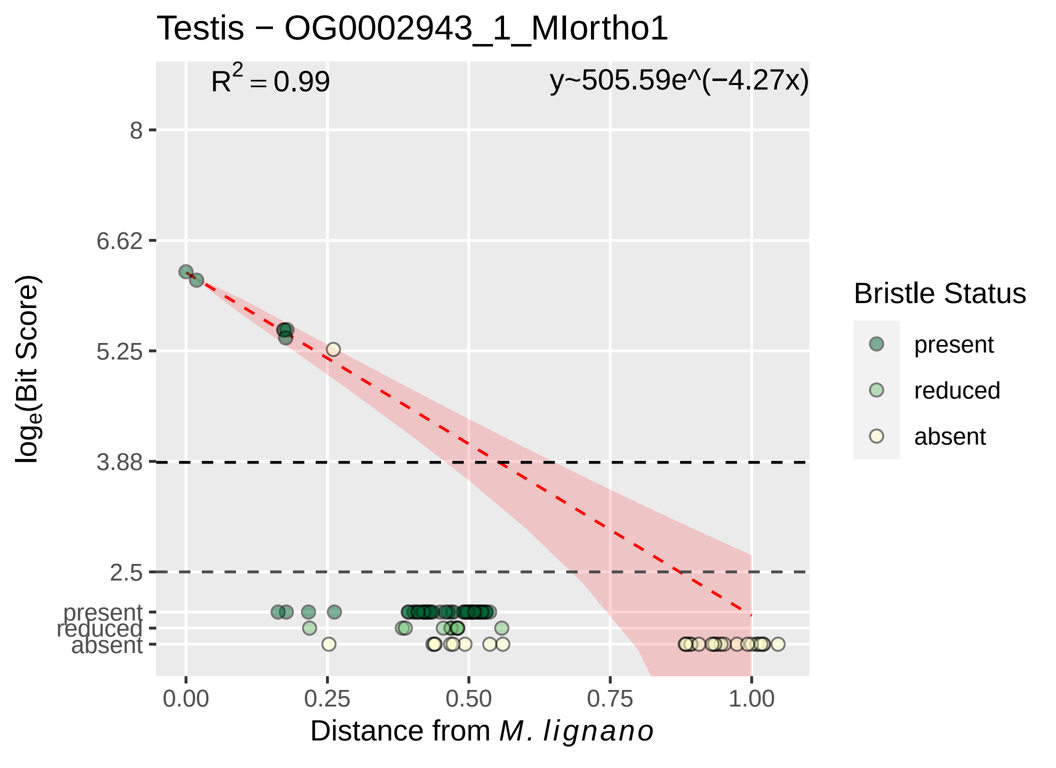

Testis − OG0002943_1_MIortho1
R^2 = 0.99
y~505.59e^(−4.27x)
8
6.62
5.25
3.88
2.5
present
reduced
absent
log_e(Bit Score)
0.00
0.25
0.50
0.75
1.00
Distance from M. lignano
Bristle Status
present
reduced
absent

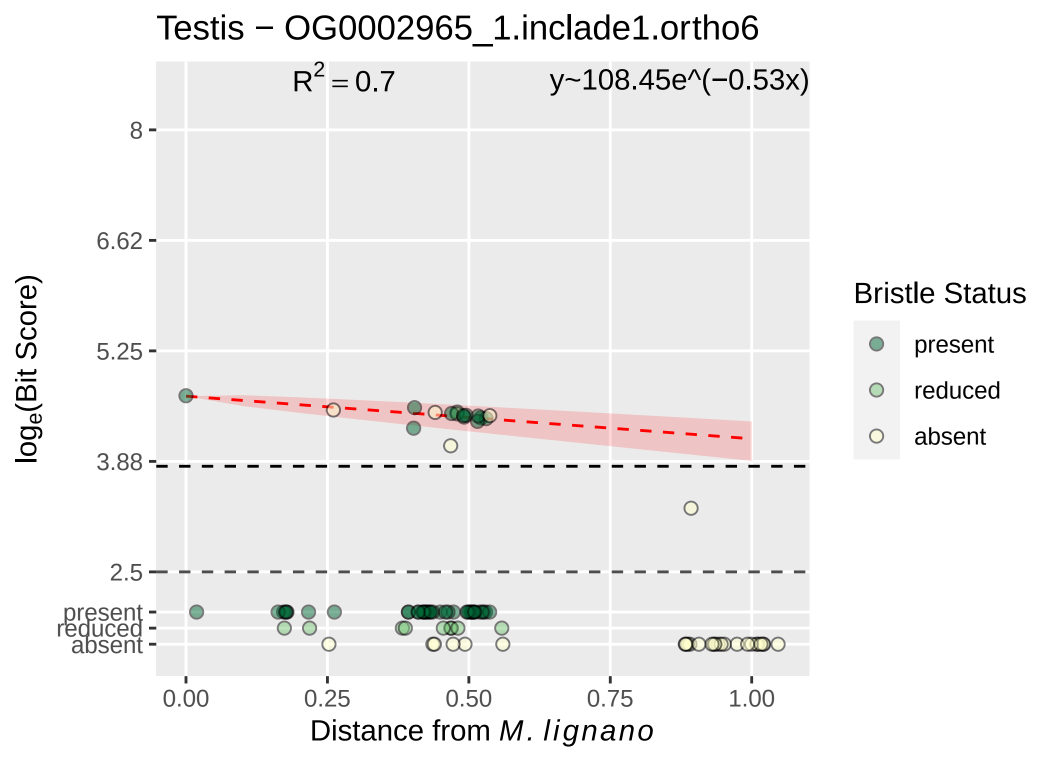

Testis − OG0002965_1.inclade1.ortho6
$R^2 = 0.7$
y~108.45e^(−0.53x)
log_e(Bit Score)
8
6.62
5.25
3.88
2.5
present
reduced
absent
0.00
0.25
0.50
0.75
1.00
Distance from M. lignano
Bristle Status
present
reduced
absent

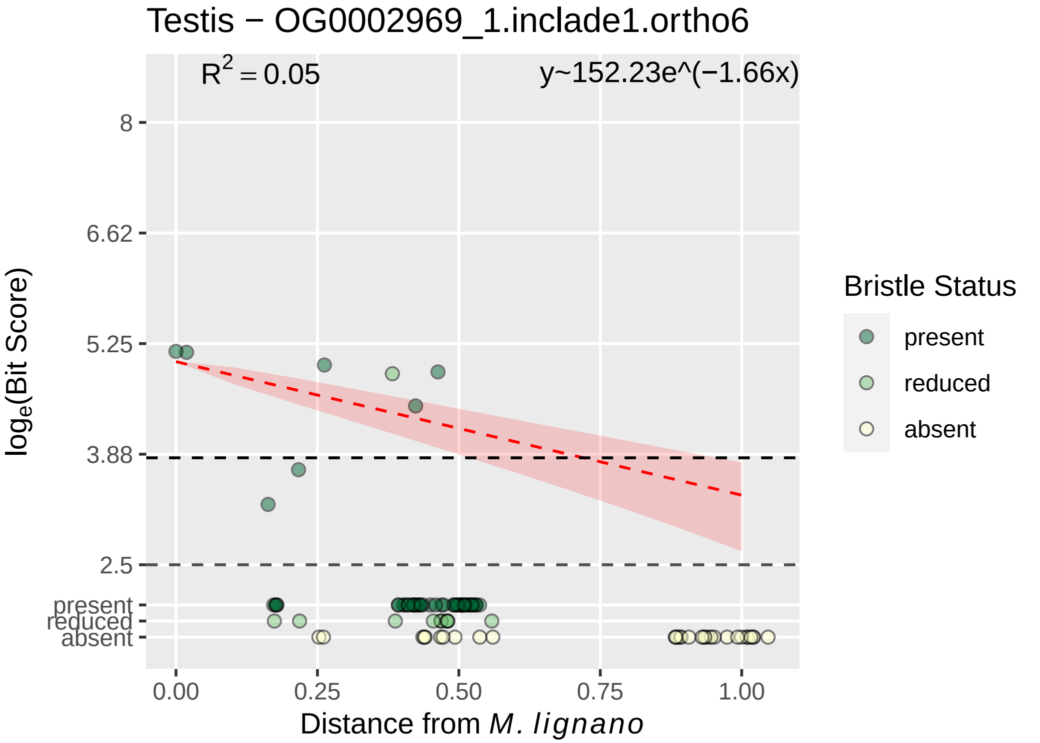

Testis − OG0002969_1.inclade1.ortho6
$R^2 = 0.05$
y~152.23e^(−1.66x)
8
6.62
5.25
3.88
2.5
present
reduced
absent
log$_e$(Bit Score)
0.00
0.25
0.50
0.75
1.00
Distance from *M. lignano*
Bristle Status
present
reduced
absent

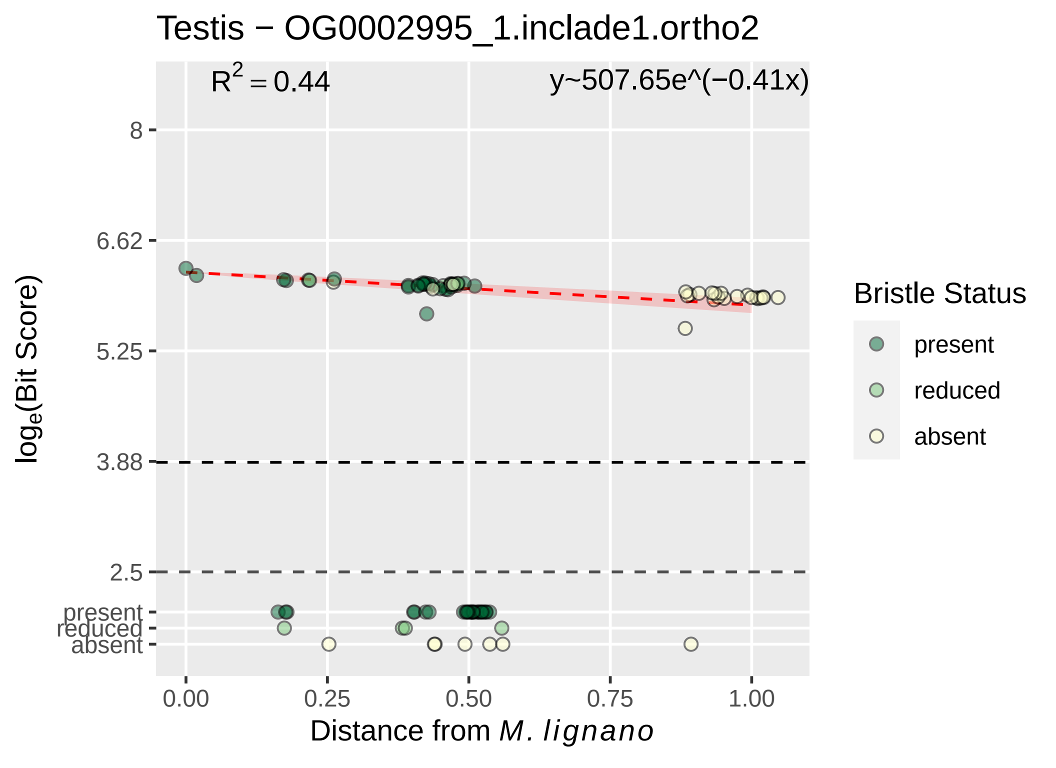

Testis − OG0002995_1.inclade1.ortho2
R^2 = 0.44
y~507.65e^(−0.41x)
8
6.62
5.25
3.88
2.5
present
reduced
absent
log_e(Bit Score)
0.00
0.25
0.50
0.75
1.00
Distance from M. lignano
Bristle Status
present
reduced
absent

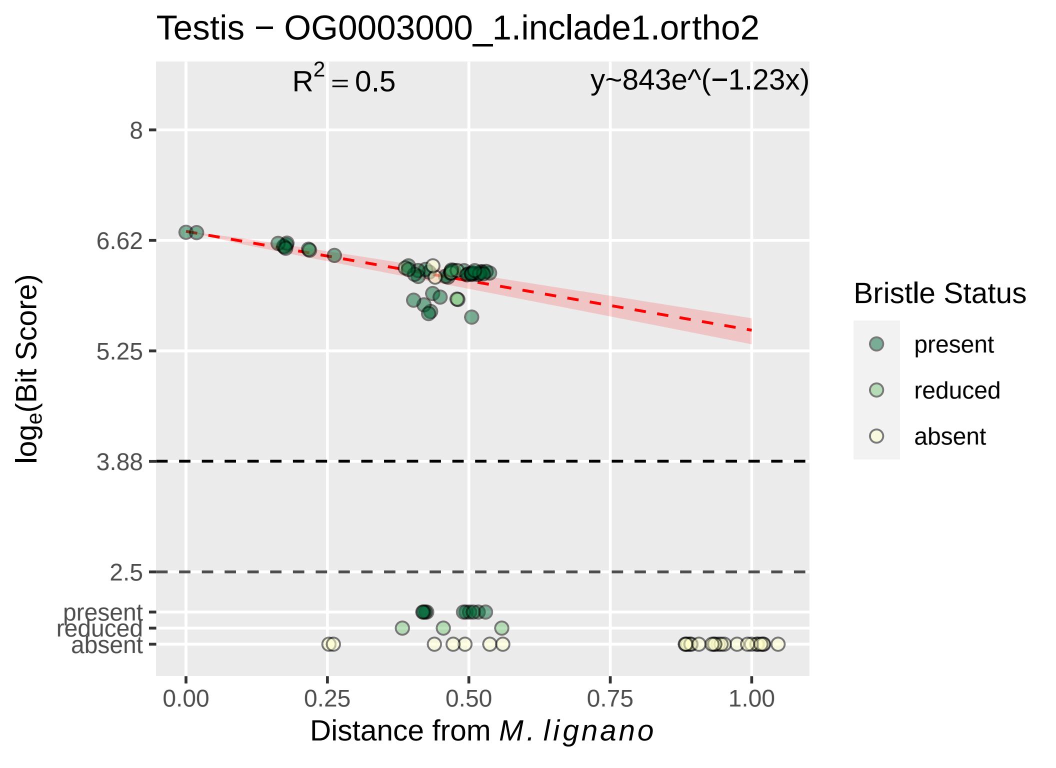

Testis − OG0003000_1.inclade1.ortho2
$R^2 = 0.5$
y~843e^(−1.23x)
8
6.62
5.25
3.88
2.5
present
reduced
absent
$log_e$(Bit Score)
0.00
0.25
0.50
0.75
1.00
Distance from *M. lignano*
Bristle Status
present
reduced
absent

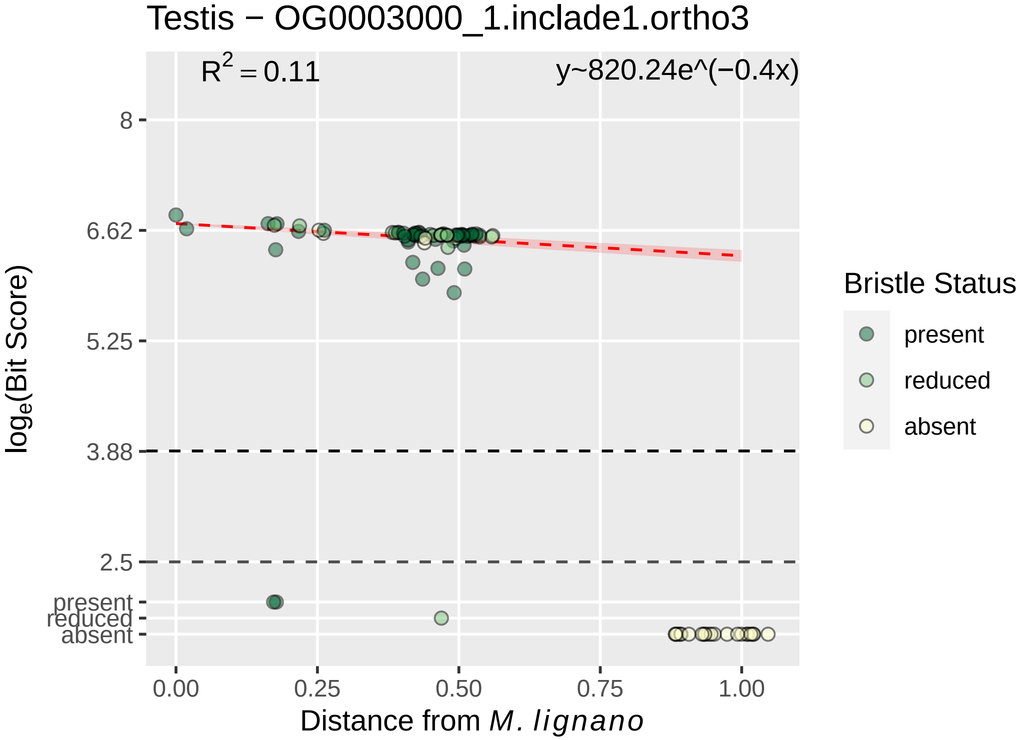

Testis − OG0003000_1.inclade1.ortho3
R^2 = 0.11
y~820.24e^(−0.4x)
8
6.62
5.25
3.88
2.5
present
reduced
absent
log_e(Bit Score)
0.00
0.25
0.50
0.75
1.00
Distance from M. lignano
Bristle Status
present
reduced
absent

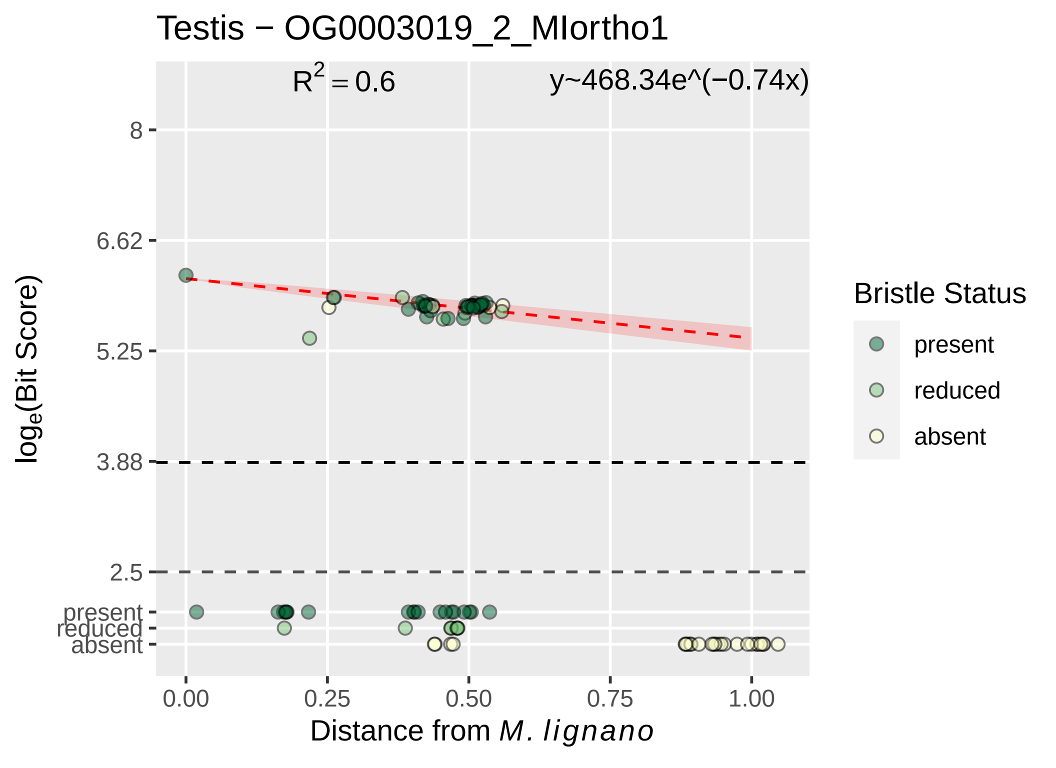

Testis − OG0003019_2_MIortho1
R² = 0.6
y~468.34e^(−0.74x)
8
6.62
5.25
3.88
2.5
present
reduced
absent
loge(Bit Score)
0.00
0.25
0.50
0.75
1.00
Distance from M. lignano
Bristle Status
present
reduced
absent

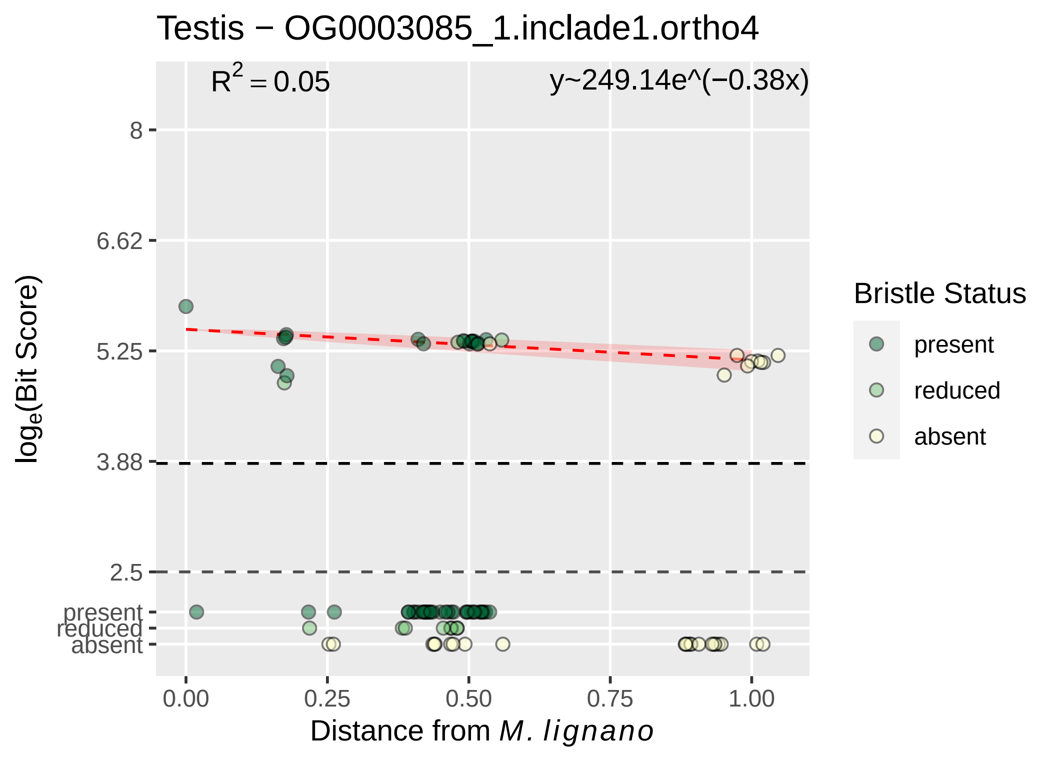
Testis − OG0003085_1.inclade1.ortho4
R2 = 0.05
y~249.14e^(−0.38x)
8
6.62
5.25
3.88
2.5
present
reduced
absent
loge(Bit Score)
0.00
0.25
0.50
0.75
1.00
Distance from M. lignano
Bristle Status
present
reduced
absent

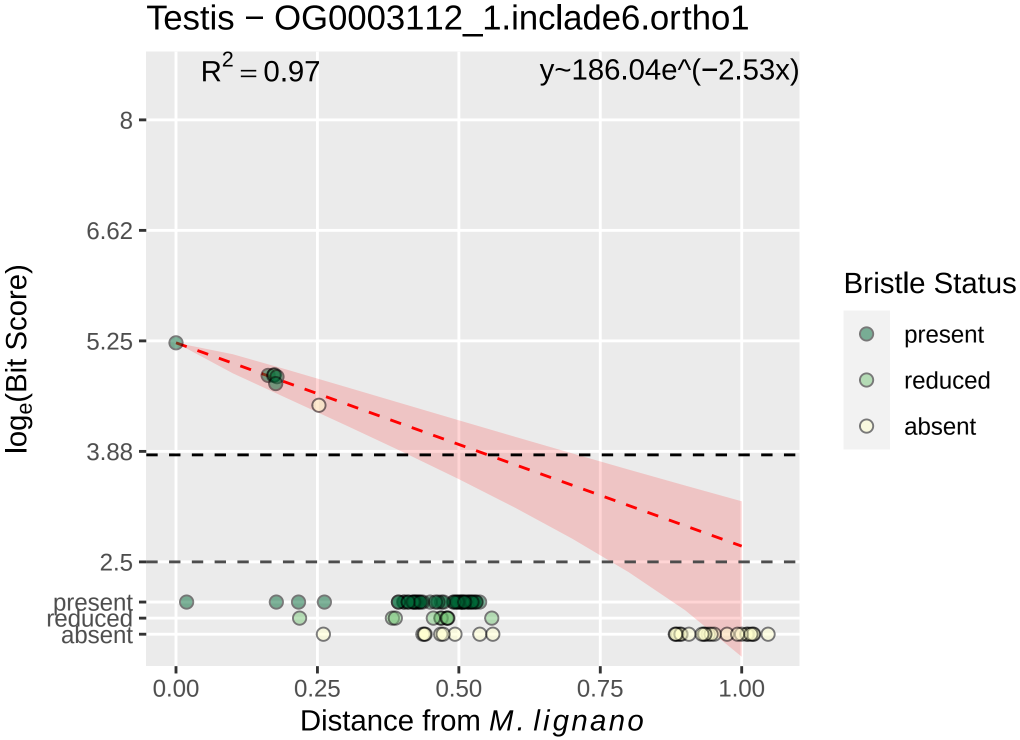

Testis − OG0003112_1.inclade6.ortho1
$R^2 = 0.97$
y~186.04e^(−2.53x)
8
6.62
5.25
3.88
2.5
present
reduced
absent
0.00
0.25
0.50
0.75
1.00
Distance from *M. lignano*
$\log_e$(Bit Score)
Bristle Status
present
reduced
absent

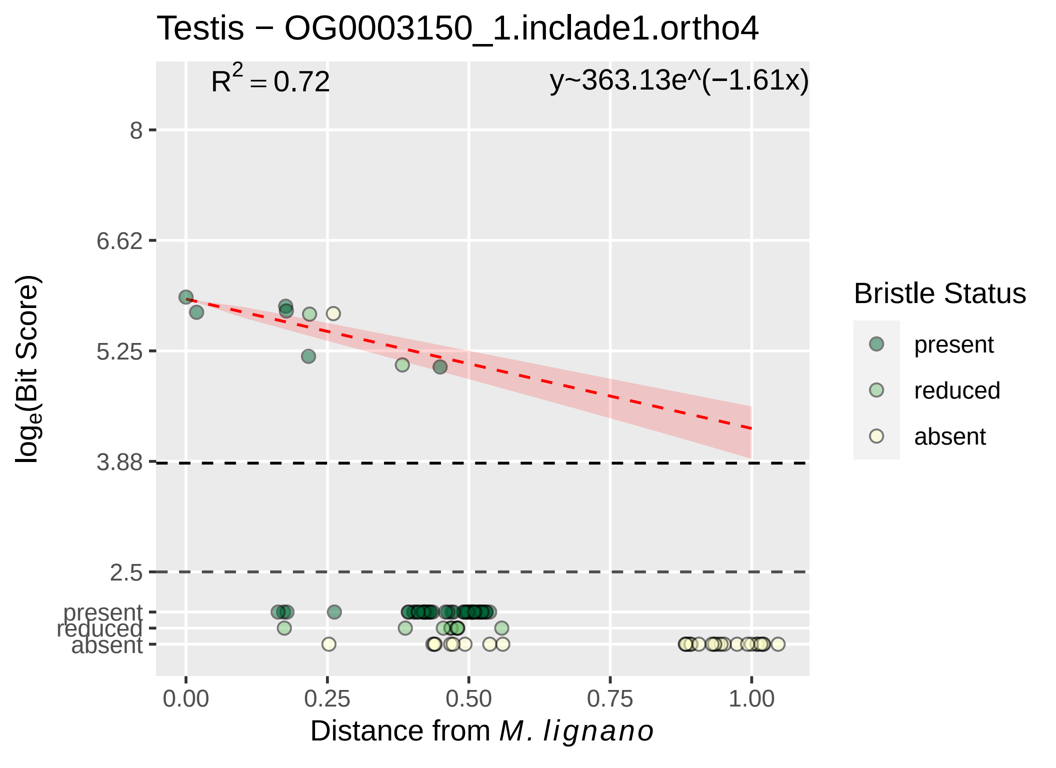

Testis − OG0003150_1.inclade1.ortho4
$R^2 = 0.72$
y~363.13e^(−1.61x)
8
6.62
5.25
3.88
2.5
present
reduced
absent
$\log_e$(Bit Score)
0.00
0.25
0.50
0.75
1.00
Distance from *M. lignano*
Bristle Status
present
reduced
absent

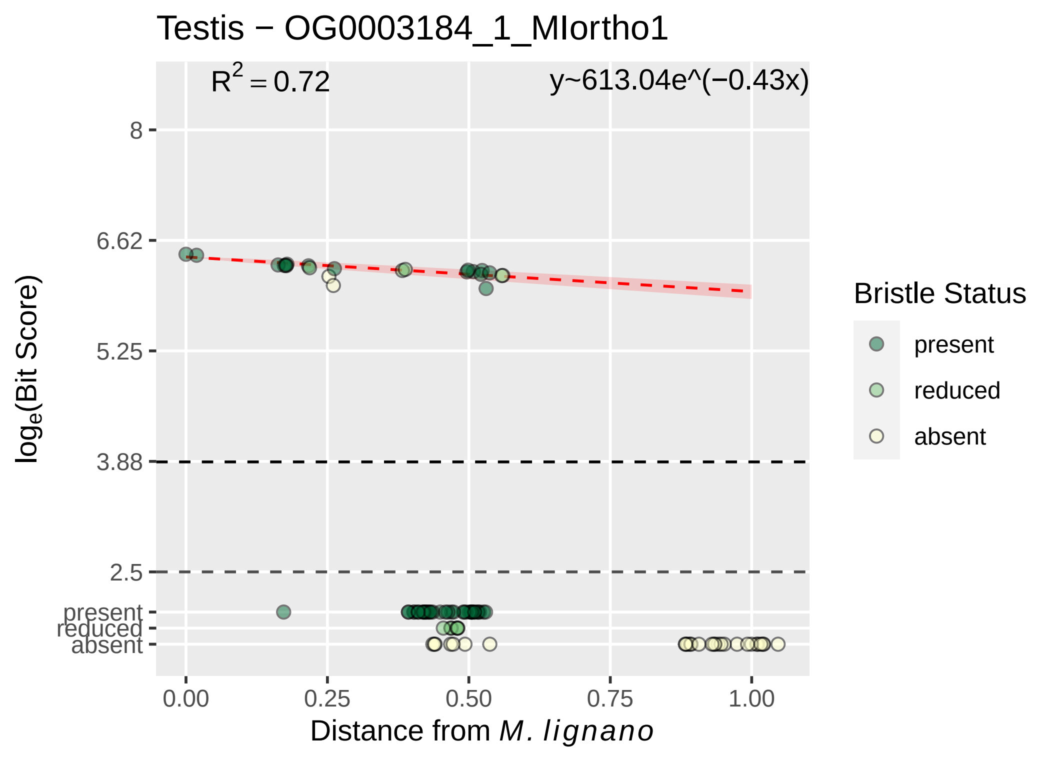

Testis − OG0003184_1_MIortho1
R^2 = 0.72
y~613.04e^(−0.43x)
8
6.62
5.25
3.88
2.5
present
reduced
absent
log_e(Bit Score)
0.00
0.25
0.50
0.75
1.00
Distance from M. lignano
Bristle Status
present
reduced
absent

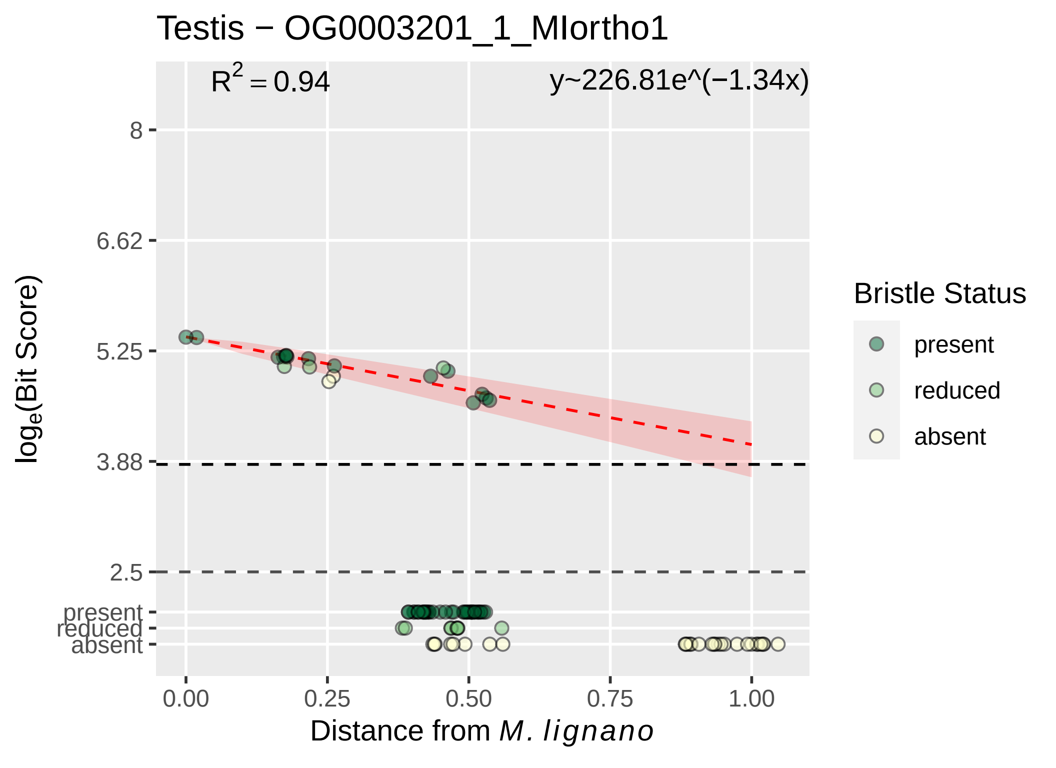

Testis − OG0003201_1_MIortho1
$R^2 = 0.94$
y~226.81e^(−1.34x)
8
6.62
5.25
3.88
2.5
present
reduced
absent
$\log_e$(Bit Score)
0.00
0.25
0.50
0.75
1.00
Distance from *M. lignano*
Bristle Status
present
reduced
absent

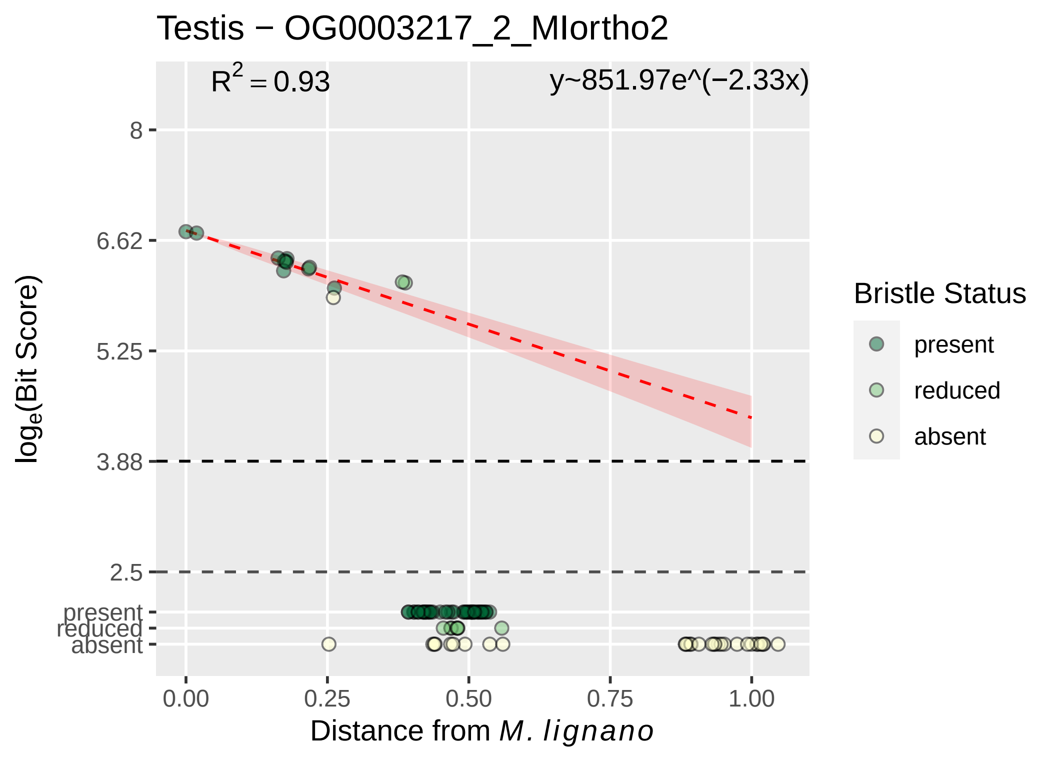

Testis − OG0003217_2_MIortho2
R² = 0.93
y~851.97e^(−2.33x)
8
6.62
5.25
3.88
2.5
present
reduced
absent
log_e(Bit Score)
0.00
0.25
0.50
0.75
1.00
Distance from M. lignano
Bristle Status
present
reduced
absent

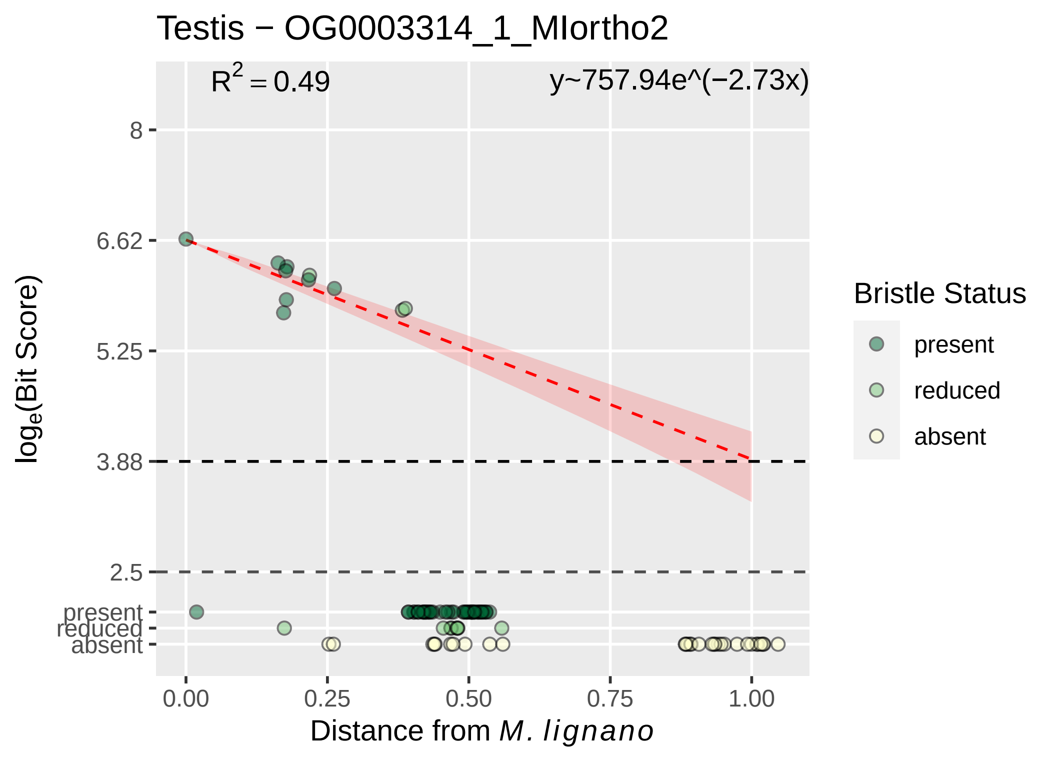

Testis − OG0003314_1_MIortho2
$R^2 = 0.49$
y~757.94e^(−2.73x)
8
6.62
5.25
3.88
2.5
present
reduced
absent
log$_e$(Bit Score)
0.00
0.25
0.50
0.75
1.00
Distance from *M. lignano*
Bristle Status
present
reduced
absent

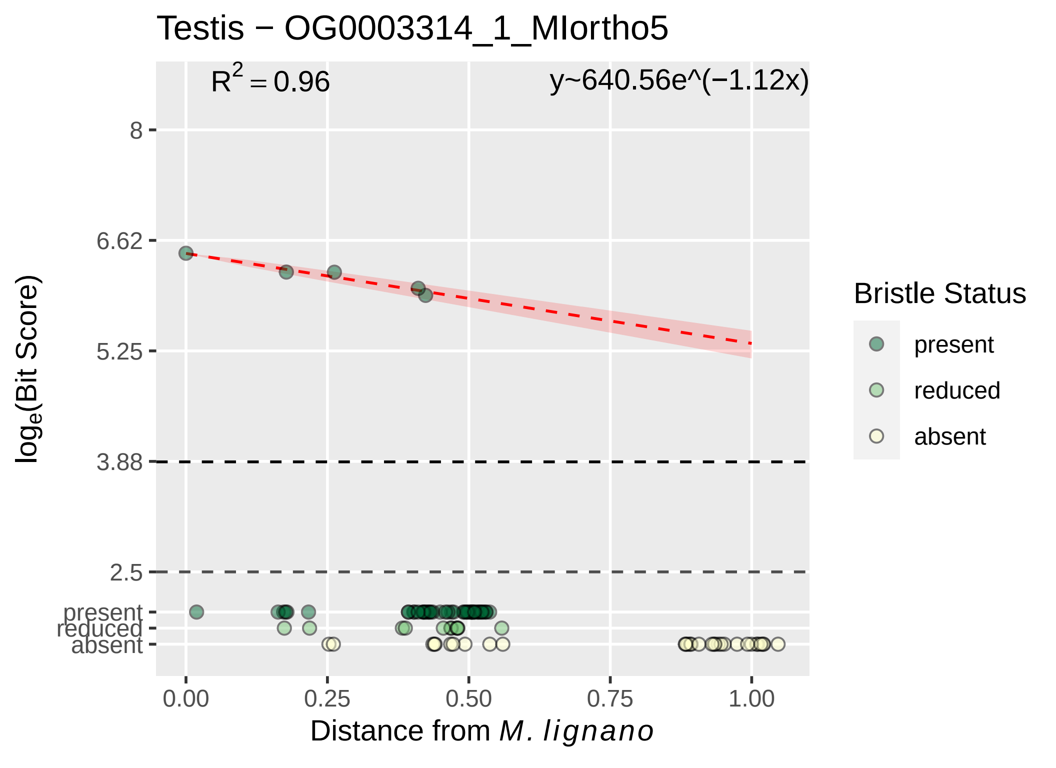

Testis − OG0003314_1_MIortho5
R2 = 0.96
y~640.56e^(−1.12x)
8
6.62
5.25
3.88
2.5
present
reduced
absent
loge(Bit Score)
0.00
0.25
0.50
0.75
1.00
Distance from M. lignano
Bristle Status
present
reduced
absent

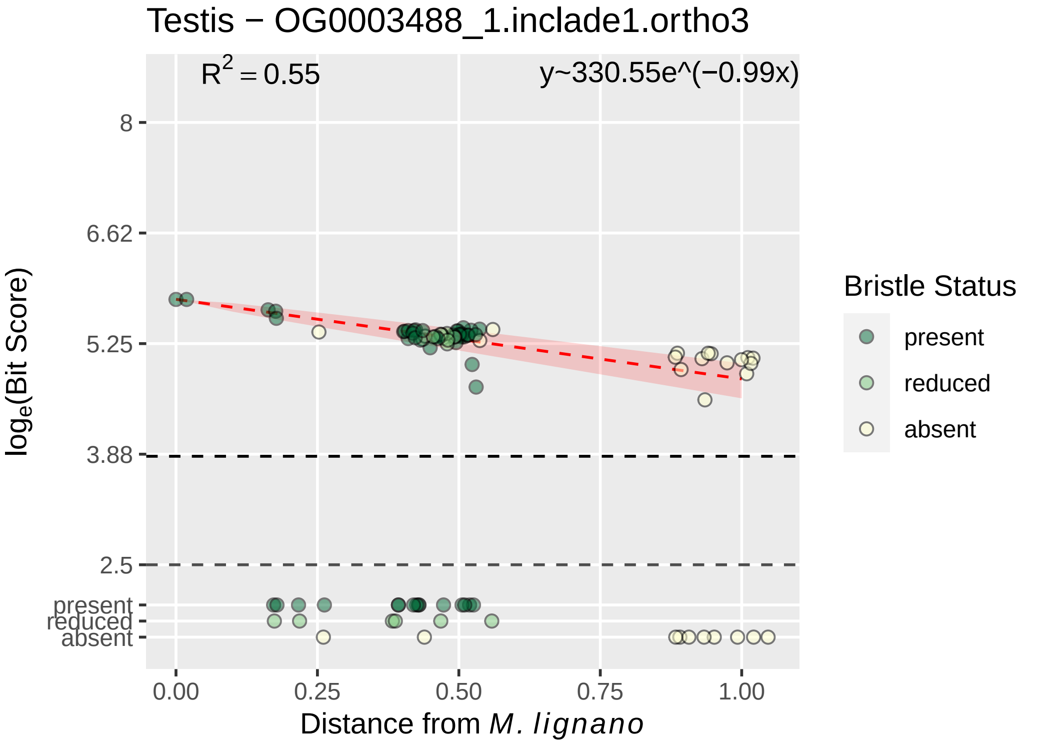

Testis − OG0003488_1.inclade1.ortho3
$R^2 = 0.55$
y~330.55e^(−0.99x)
8
6.62
5.25
3.88
2.5
present
reduced
absent
0.00
0.25
0.50
0.75
1.00
Distance from M. lignano
loge(Bit Score)
Bristle Status
present
reduced
absent

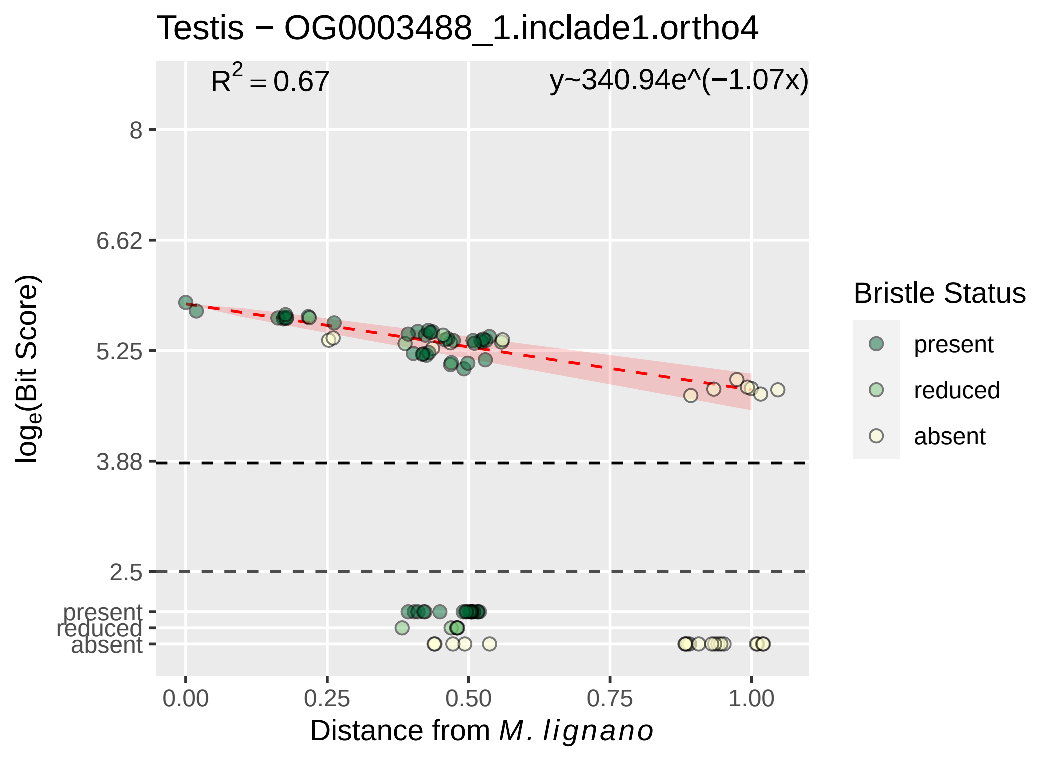
Testis − OG0003488_1.inclade1.ortho4
$R^2 = 0.67$
y~340.94e^(−1.07x)
8
6.62
5.25
3.88
2.5
present
reduced
absent
$\log_e$(Bit Score)
0.00
0.25
0.50
0.75
1.00
Distance from M. lignano
Bristle Status
present
reduced
absent

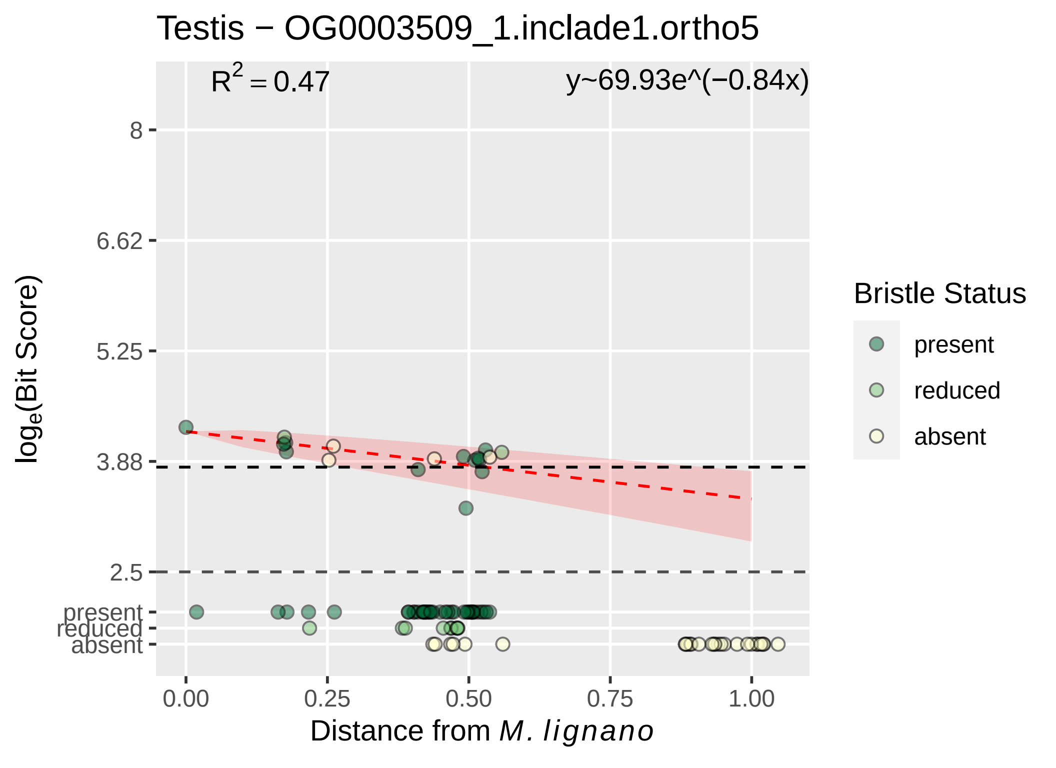

Testis − OG0003509_1.inclade1.ortho5
$R^2 = 0.47$
y~69.93e^(−0.84x)
8
6.62
5.25
3.88
2.5
present
reduced
absent
0.00
0.25
0.50
0.75
1.00
Distance from *M. lignano*
$\log_e$(Bit Score)
Bristle Status
present
reduced
absent

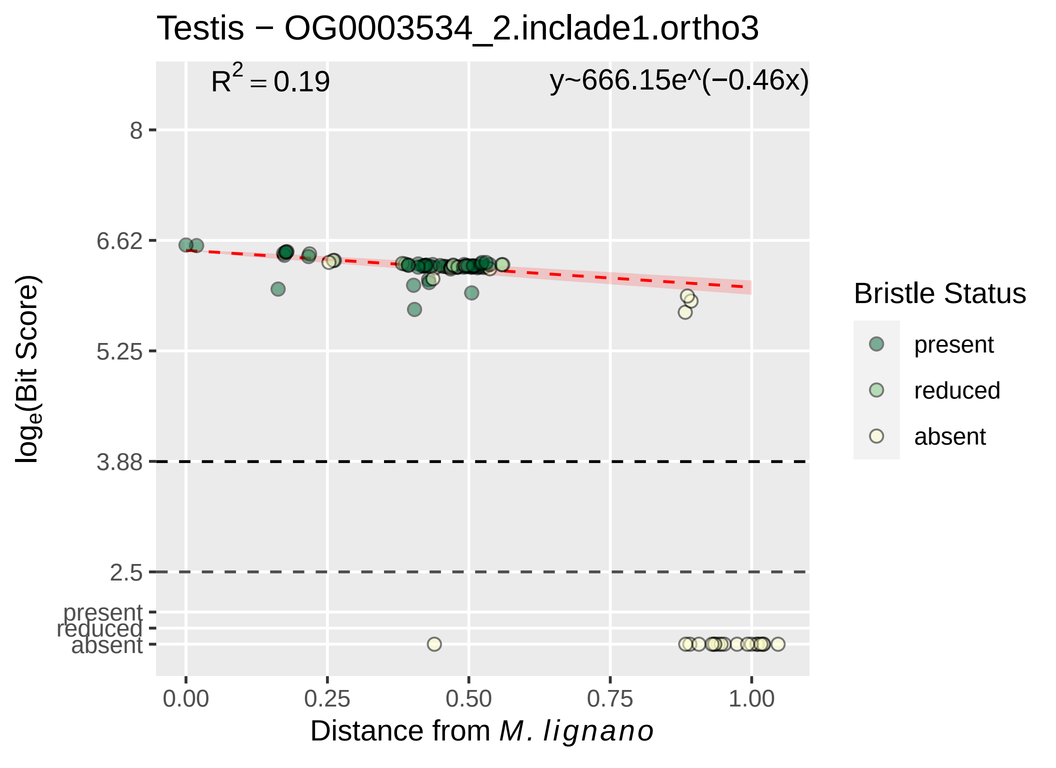
Testis − OG0003534_2.inclade1.ortho3
R2 = 0.19
y~666.15e^(−0.46x)
8
6.62
5.25
3.88
2.5
present
reduced
absent
0.00
0.25
0.50
0.75
1.00
loge(Bit Score)
Distance from M. lignano
Bristle Status
present
reduced
absent

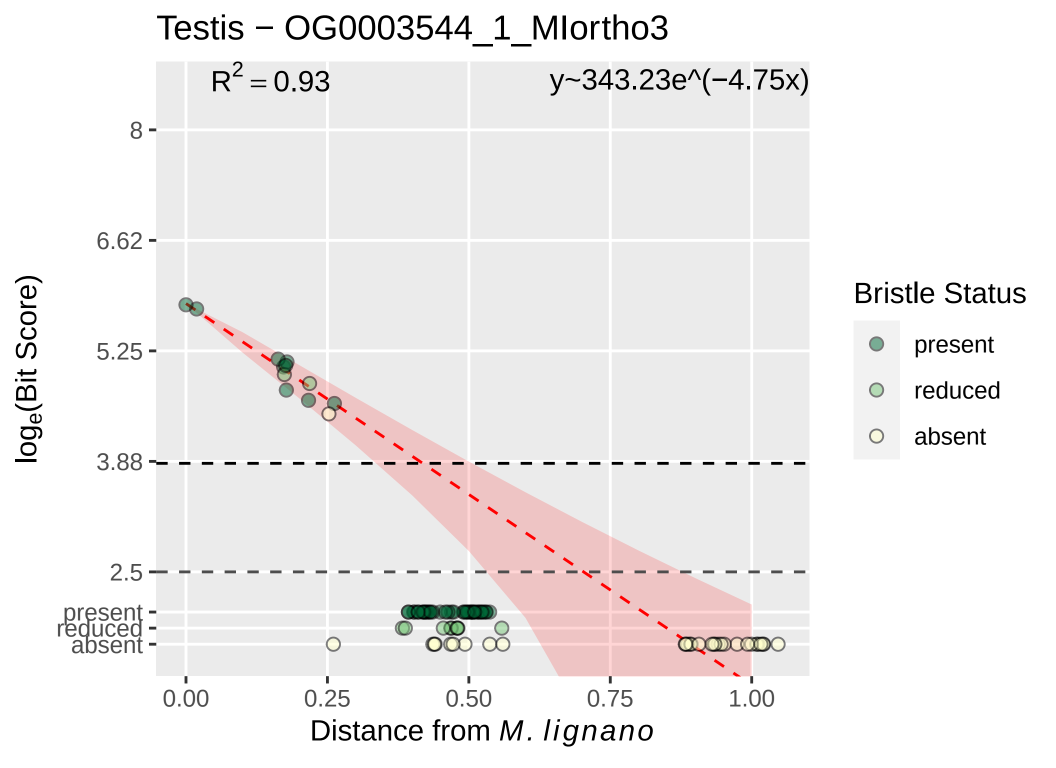

Testis − OG0003544_1_MIortho3
$R^2 = 0.93$
y~343.23e^(−4.75x)
8
6.62
5.25
3.88
2.5
present
reduced
absent
0.00
0.25
0.50
0.75
1.00
Distance from M. lignano
log$_e$(Bit Score)
Bristle Status
present
reduced
absent

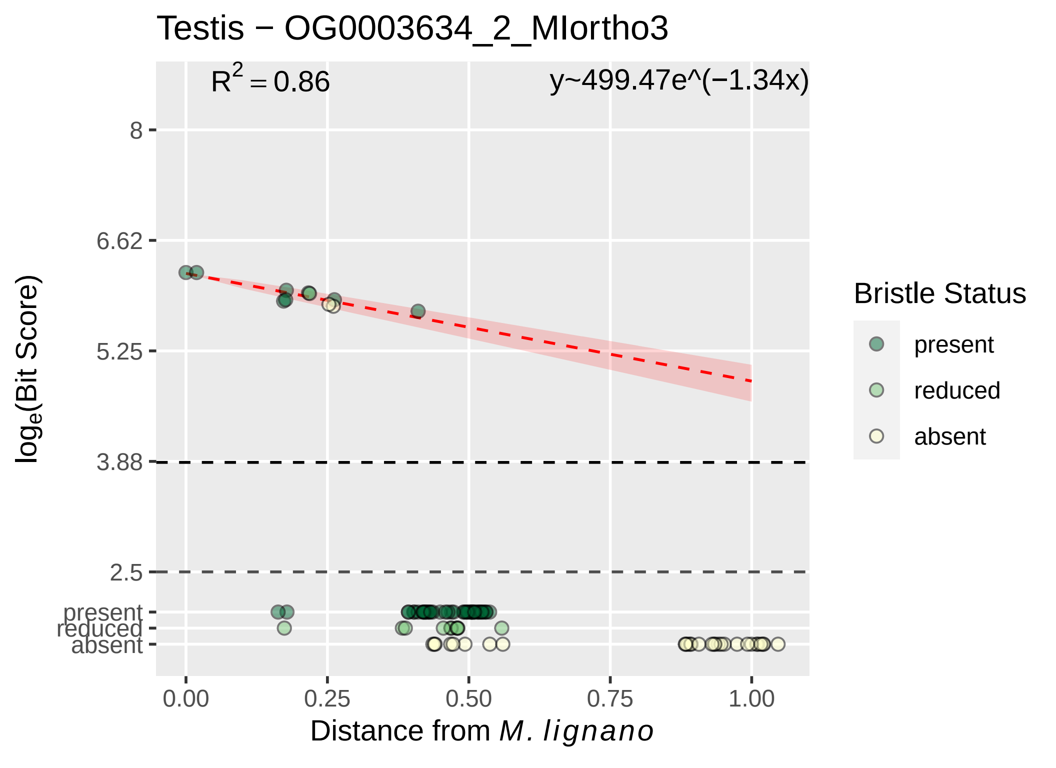

Testis − OG0003634_2_MIortho3
$R^2 = 0.86$
y~499.47e^(−1.34x)
8
6.62
5.25
3.88
2.5
present
reduced
absent
$\log_e$(Bit Score)
0.00
0.25
0.50
0.75
1.00
Distance from *M. lignano*
Bristle Status
present
reduced
absent

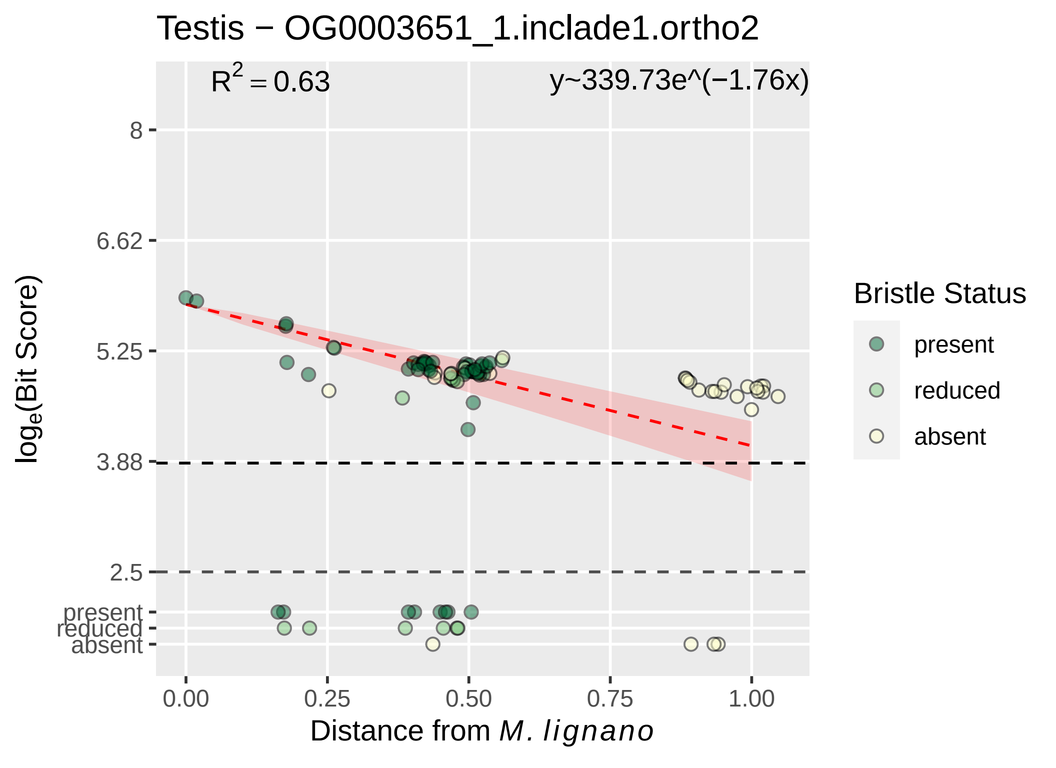

Testis − OG0003651_1.inclade1.ortho2
$R^2 = 0.63$
y~339.73e^(−1.76x)
8
6.62
5.25
3.88
2.5
present
reduced
absent
$\log_e$(Bit Score)
0.00
0.25
0.50
0.75
1.00
Distance from *M. lignano*
Bristle Status
present
reduced
absent

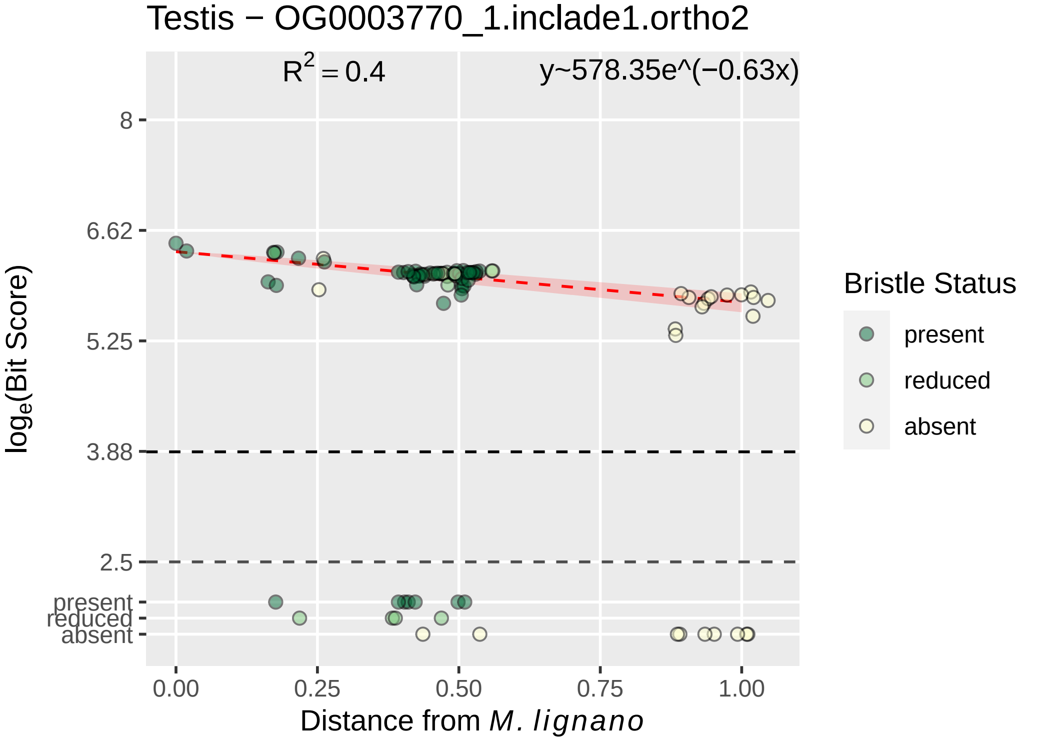
Testis − OG0003770_1.inclade1.ortho2
$R^2 = 0.4$
y~578.35e^(−0.63x)
8
6.62
5.25
3.88
2.5
present
reduced
absent
$\log_e$(Bit Score)
0.00
0.25
0.50
0.75
1.00
Distance from *M. lignano*
Bristle Status
present
reduced
absent

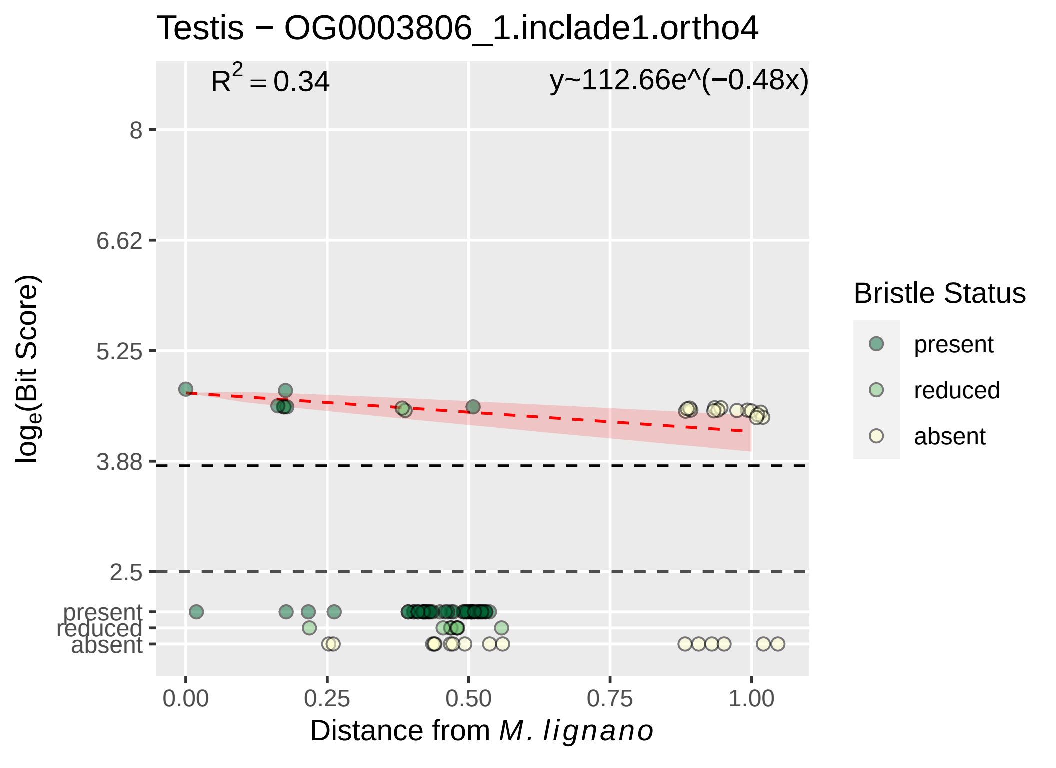

Testis − OG0003806_1.inclade1.ortho4
R2 = 0.34
y~112.66e^(−0.48x)
8
6.62
5.25
3.88
2.5
present
reduced
absent
0.00
0.25
0.50
0.75
1.00
Distance from M. lignano
loge(Bit Score)
Bristle Status
present
reduced
absent

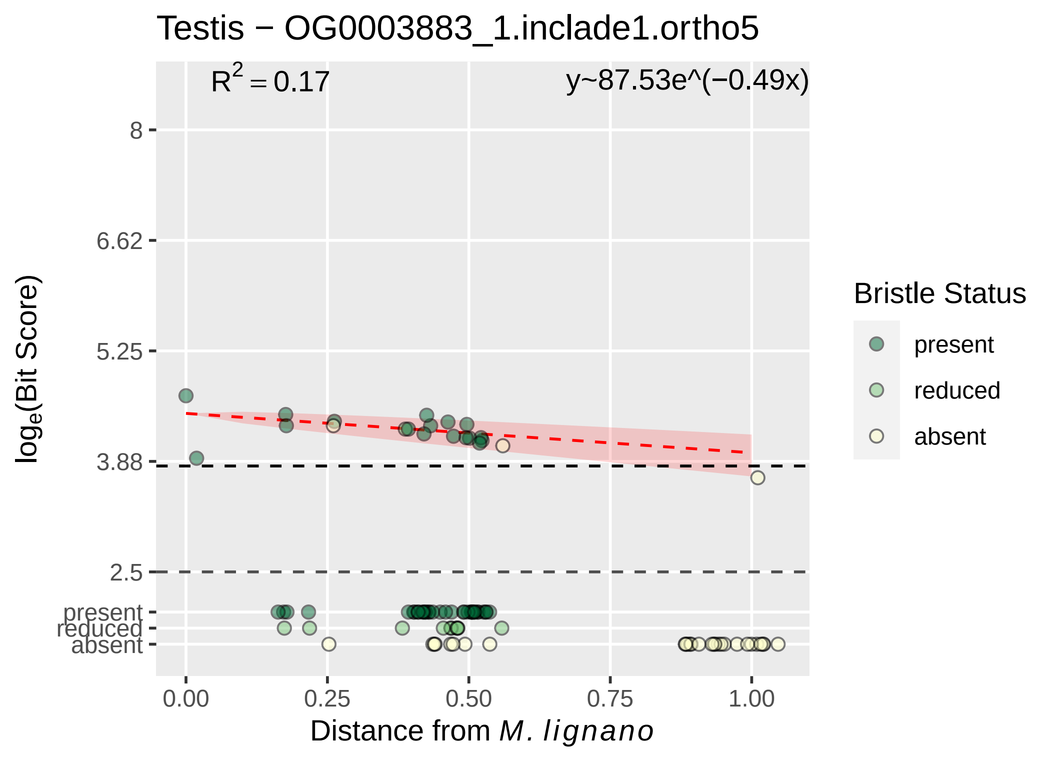

Testis − OG0003883_1.inclade1.ortho5
R² = 0.17
y~87.53e^(−0.49x)
8
6.62
5.25
3.88
2.5
present
reduced
absent
0.00
0.25
0.50
0.75
1.00
Distance from M. lignano
loge(Bit Score)
Bristle Status
present
reduced
absent

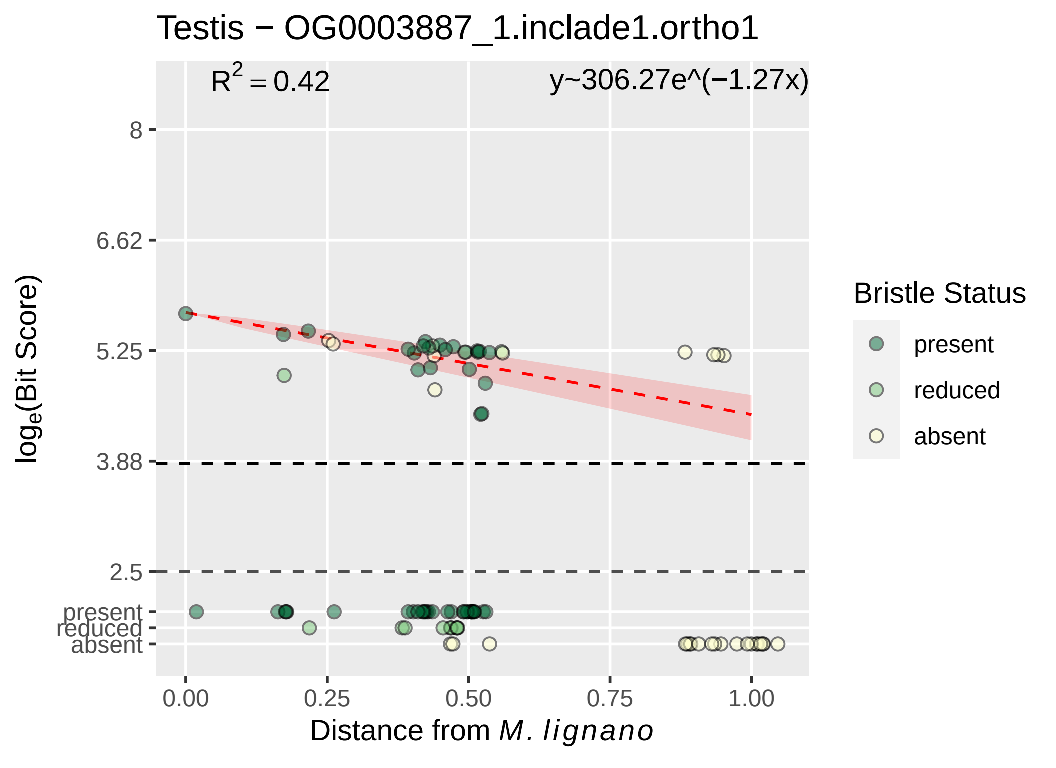

Testis − OG0003887_1.inclade1.ortho1
$R^2 = 0.42$
y~306.27e^(−1.27x)
8
6.62
5.25
3.88
2.5
present
reduced
absent
0.00
0.25
0.50
0.75
1.00
$\log_e$(Bit Score)
Distance from *M. lignano*
Bristle Status
present
reduced
absent

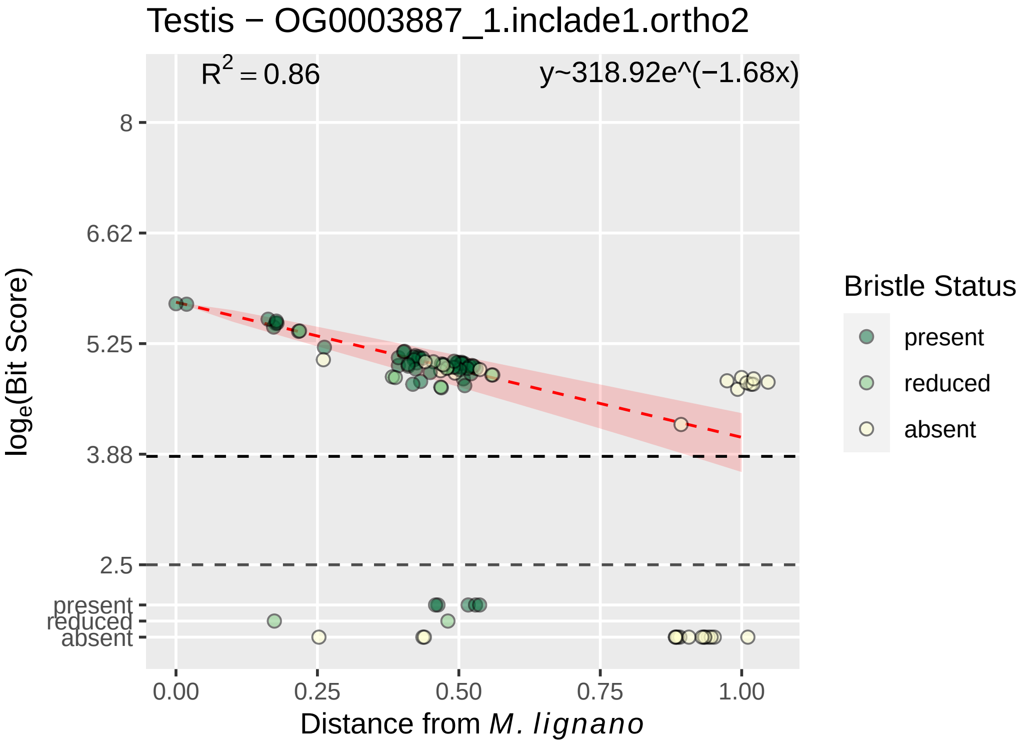

Testis − OG0003887_1.inclade1.ortho2
$R^2 = 0.86$
y~318.92e^(−1.68x)
8
6.62
5.25
3.88
2.5
present
reduced
absent
$\log_e$(Bit Score)
0.00
0.25
0.50
0.75
1.00
Distance from *M. lignano*
Bristle Status
present
reduced
absent

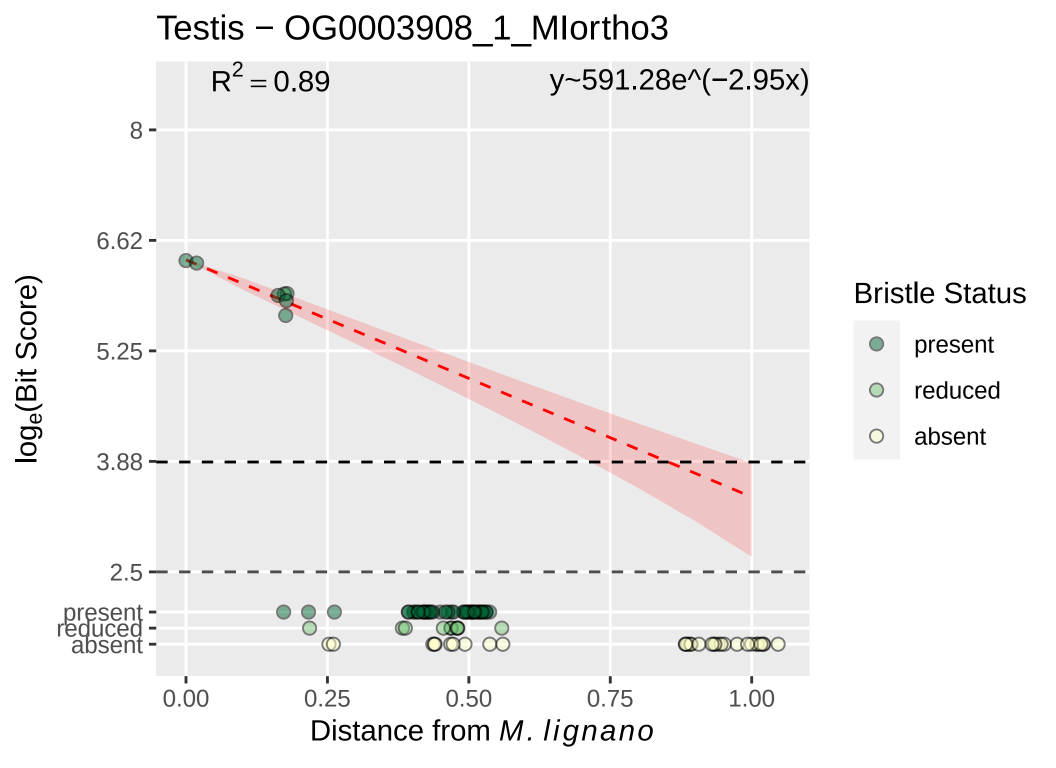

Testis − OG0003908_1_MIortho3
$R^2 = 0.89$
y~591.28e^(−2.95x)
8
6.62
5.25
3.88
2.5
present
reduced
absent
$\log_e$(Bit Score)
0.00
0.25
0.50
0.75
1.00
Distance from *M. lignano*
Bristle Status
present
reduced
absent

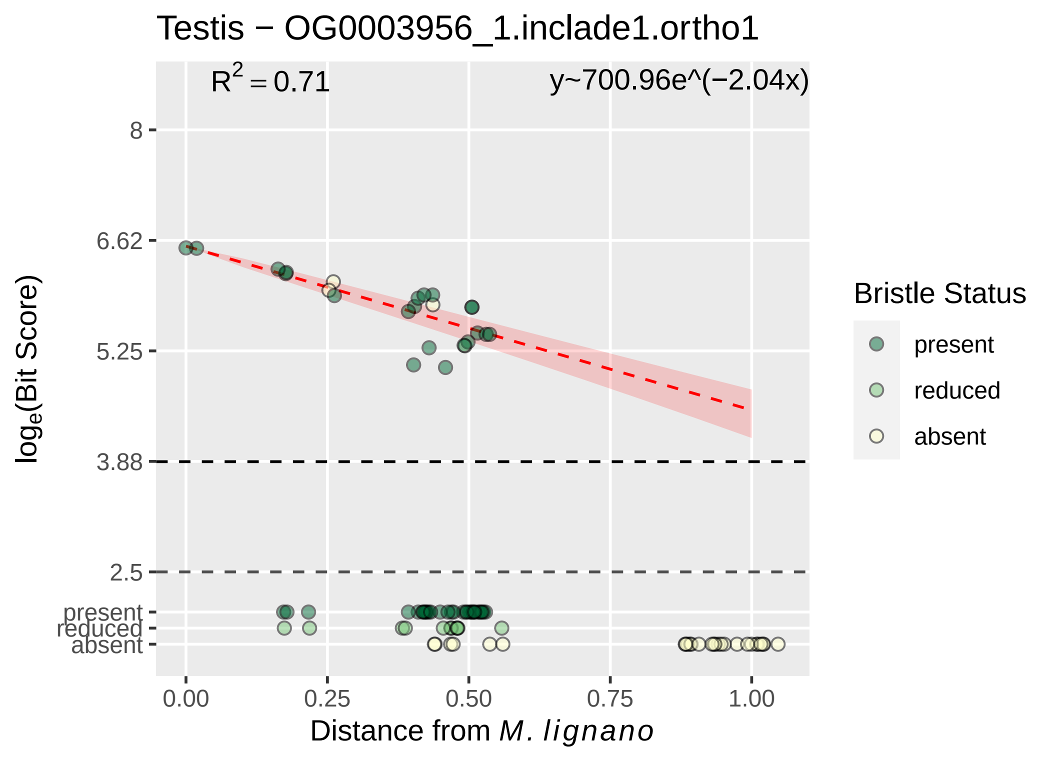

Testis − OG0003956_1.inclade1.ortho1
R^2 = 0.71
y~700.96e^(−2.04x)
8
6.62
5.25
3.88
2.5
present
reduced
absent
log_e(Bit Score)
0.00
0.25
0.50
0.75
1.00
Distance from M. lignano
Bristle Status
present
reduced
absent

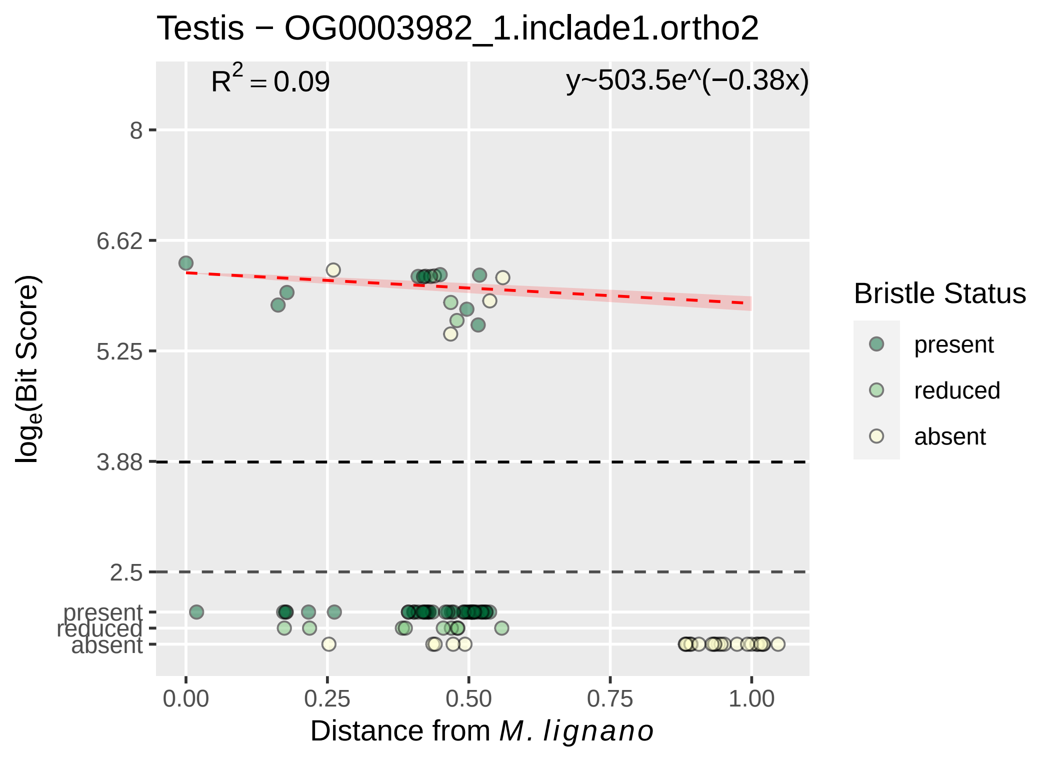

Testis − OG0003982_1.inclade1.ortho2
R2 = 0.09
y~503.5e^(−0.38x)
8
6.62
5.25
3.88
2.5
present
reduced
absent
loge(Bit Score)
0.00
0.25
0.50
0.75
1.00
Distance from M. lignano
Bristle Status
present
reduced
absent

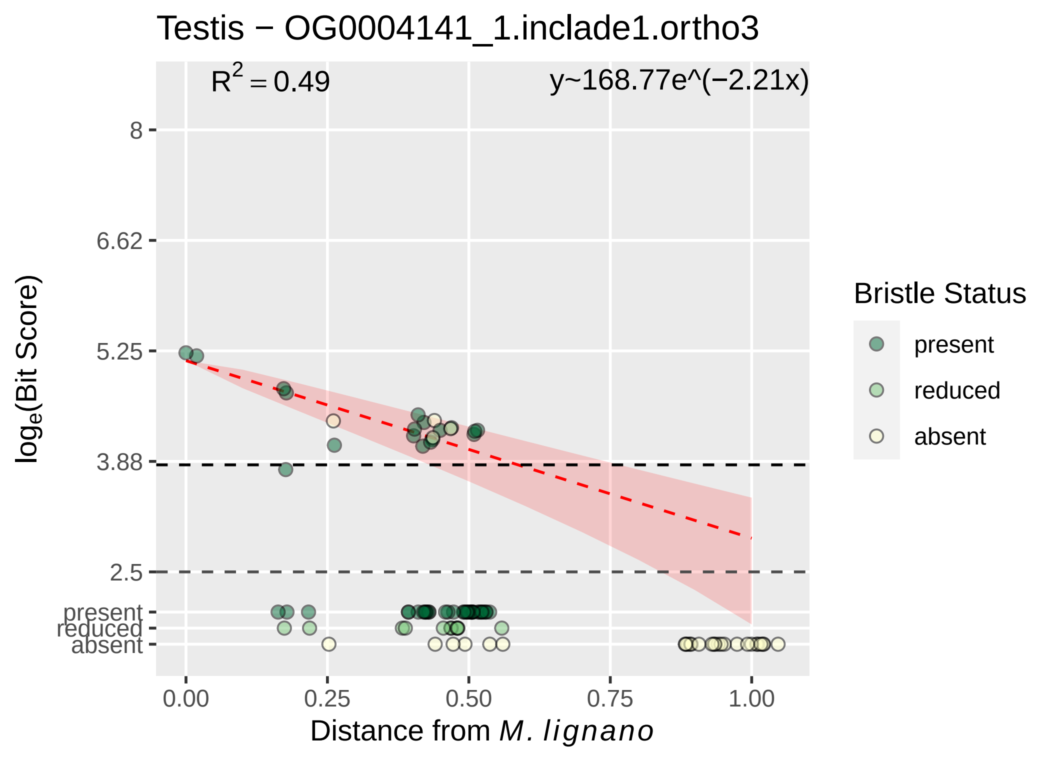

Testis − OG0004141_1.inclade1.ortho3
$R^2 = 0.49$
y~168.77e^(−2.21x)
8
6.62
5.25
3.88
2.5
present
reduced
absent
0.00
0.25
0.50
0.75
1.00
$\log_e$(Bit Score)
Distance from *M. lignano*
Bristle Status
present
reduced
absent

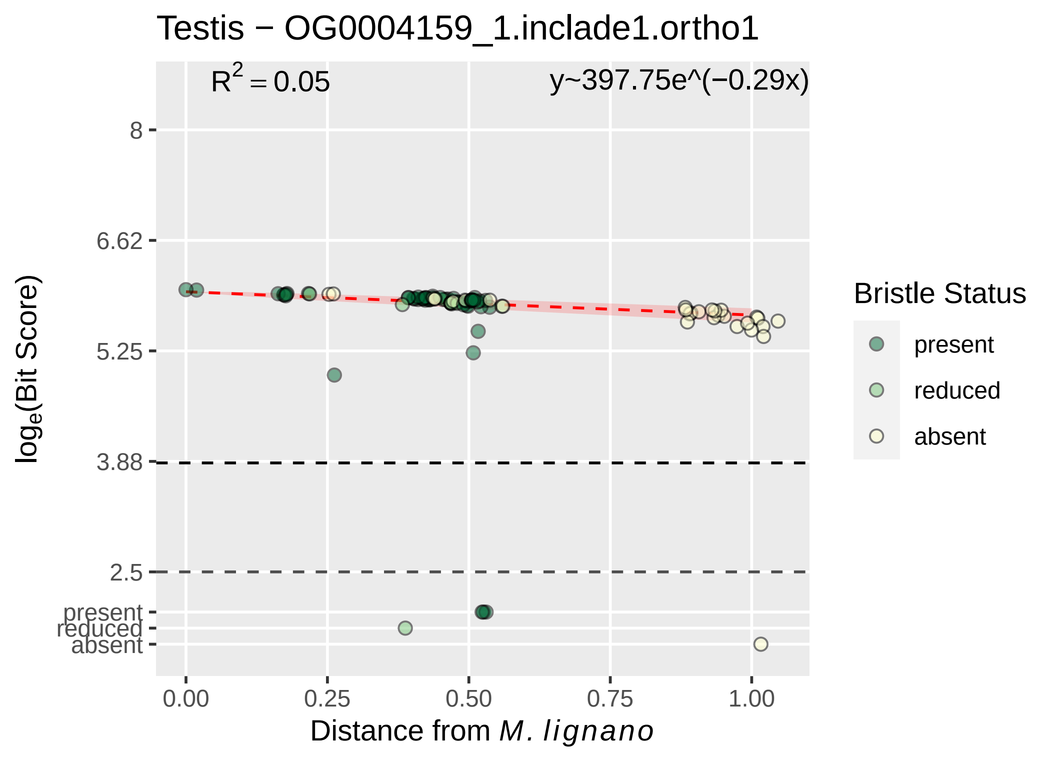
Testis − OG0004159_1.inclade1.ortho1
R^2 = 0.05
y~397.75e^(−0.29x)
8
6.62
5.25
3.88
2.5
present
reduced
absent
loge(Bit Score)
0.00
0.25
0.50
0.75
1.00
Distance from M. lignano
Bristle Status
present
reduced
absent

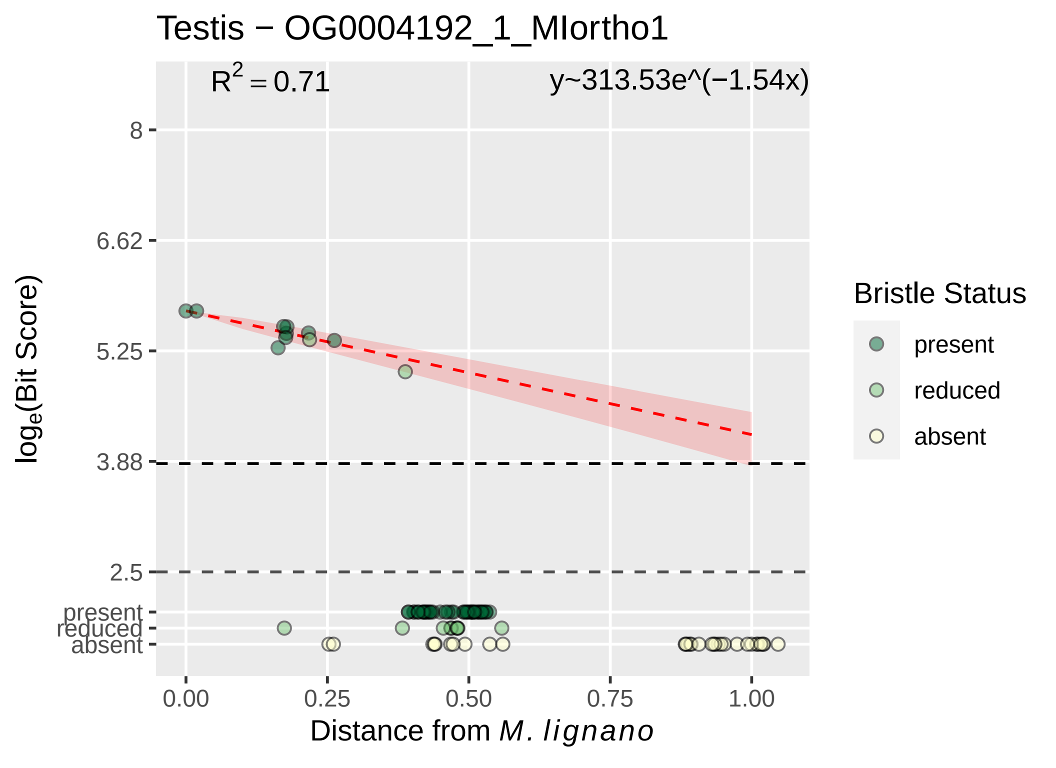

Testis − OG0004192_1_MIortho1
$R^2 = 0.71$
y~313.53e^(−1.54x)
8
6.62
5.25
3.88
2.5
present
reduced
absent
$\log_e$(Bit Score)
0.00
0.25
0.50
0.75
1.00
Distance from *M. lignano*
Bristle Status
present
reduced
absent

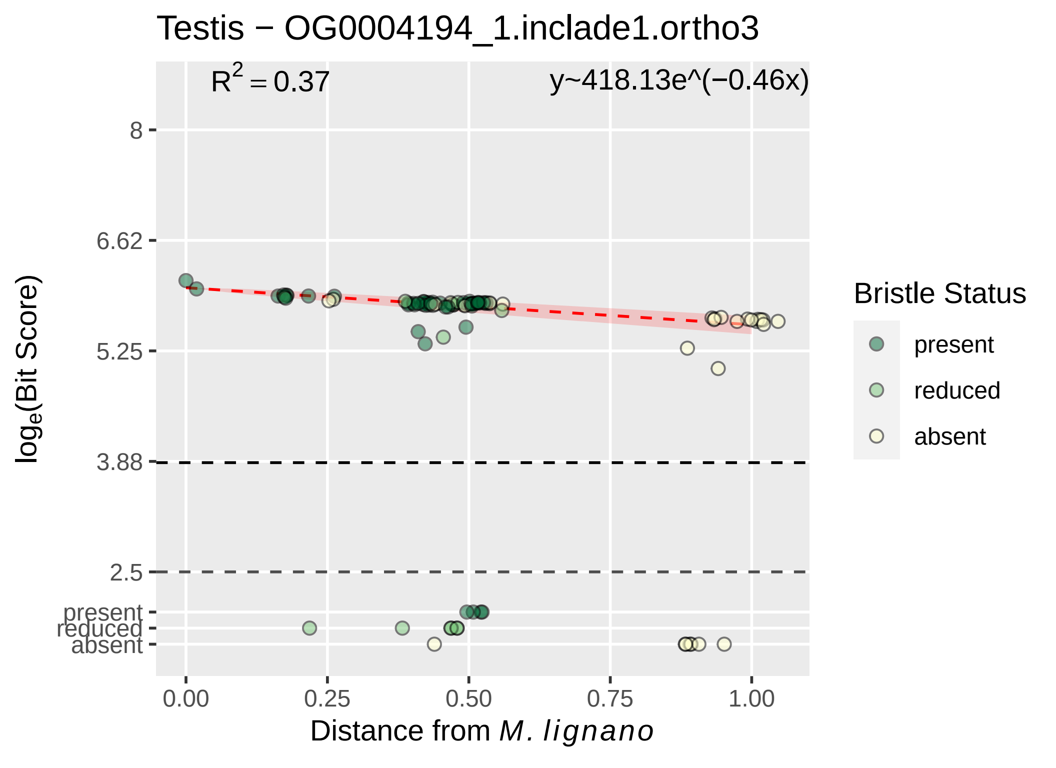

Testis − OG0004194_1.inclade1.ortho3
$R^2 = 0.37$
y~418.13e^(−0.46x)
8
6.62
5.25
3.88
2.5
present
reduced
absent
$log_e$(Bit Score)
0.00
0.25
0.50
0.75
1.00
Distance from M. lignano
Bristle Status
present
reduced
absent

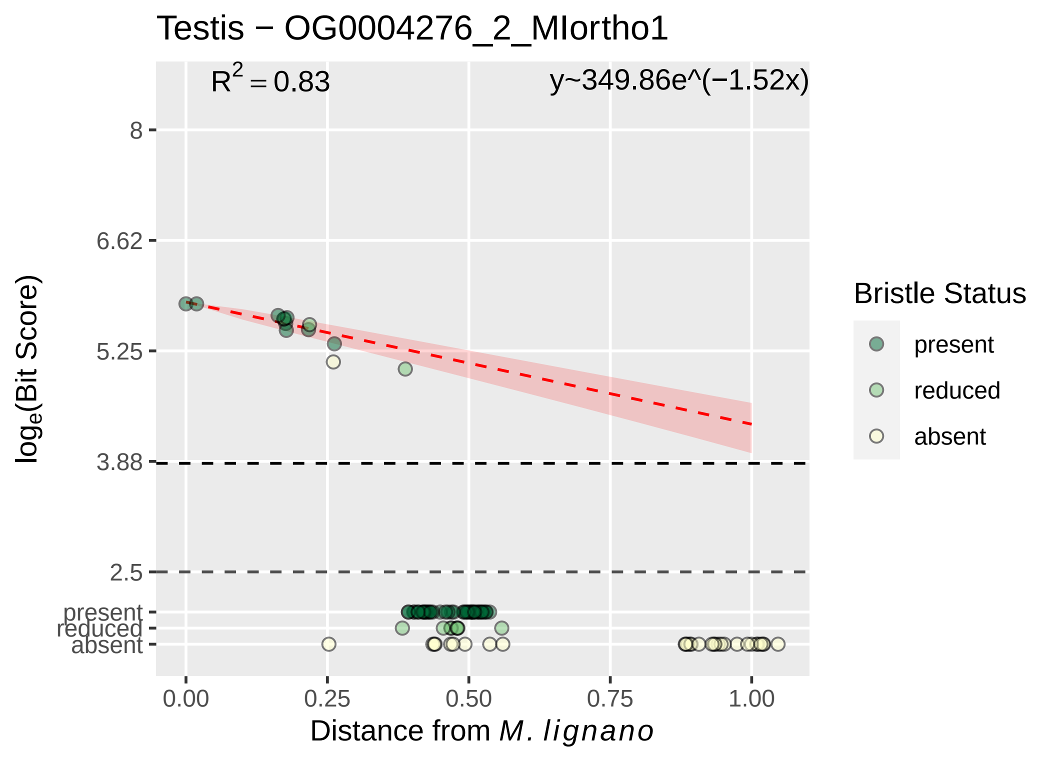
Testis − OG0004276_2_MIortho1
R^2 = 0.83
y~349.86e^(−1.52x)
8
6.62
5.25
3.88
2.5
present
reduced
absent
log_e(Bit Score)
0.00
0.25
0.50
0.75
1.00
Distance from M. lignano
Bristle Status
present
reduced
absent

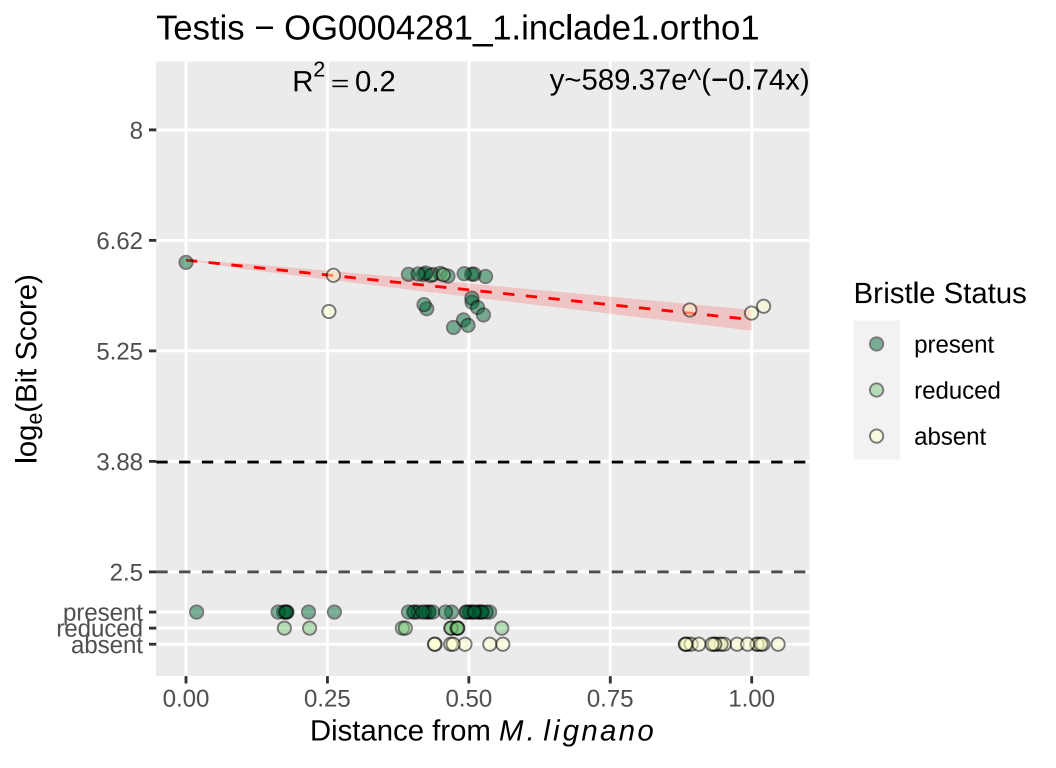
Testis − OG0004281_1.inclade1.ortho1
$R^2 = 0.2$
y~589.37e^(−0.74x)
8
6.62
5.25
3.88
2.5
present
reduced
absent
$log_e$(Bit Score)
0.00
0.25
0.50
0.75
1.00
Distance from *M. lignano*
Bristle Status
present
reduced
absent

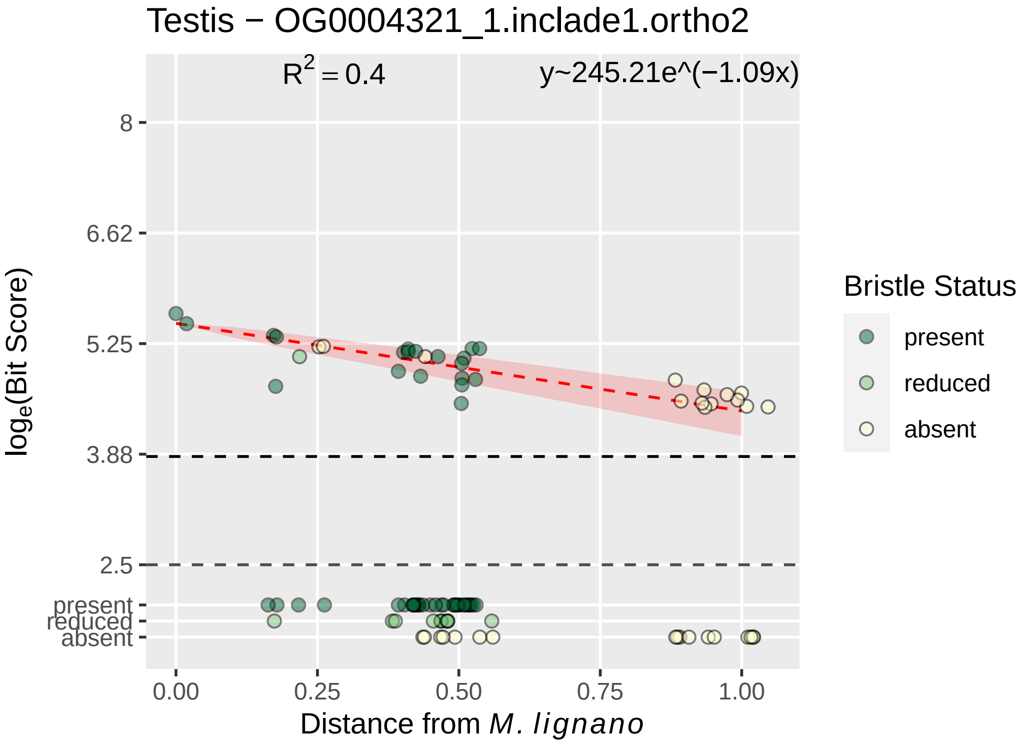

Testis − OG0004321_1.inclade1.ortho2
$R^2 = 0.4$
y~245.21e^(−1.09x)
8
6.62
5.25
3.88
2.5
present
reduced
absent
0.00
0.25
0.50
0.75
1.00
$\log_e$(Bit Score)
Distance from *M. lignano*
Bristle Status
present
reduced
absent

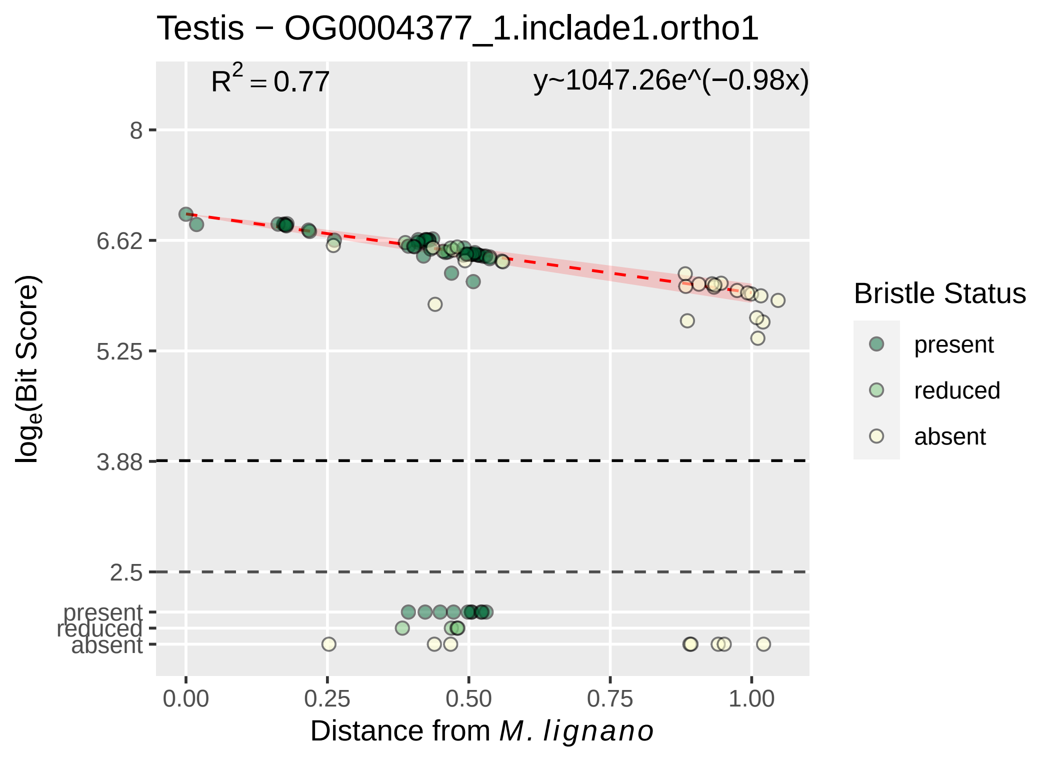

Testis − OG0004377_1.inclade1.ortho1
$R^2 = 0.77$
y~1047.26e^(−0.98x)
8
6.62
5.25
3.88
2.5
present
reduced
absent
0.00
0.25
0.50
0.75
1.00
$log_e$(Bit Score)
Distance from *M. lignano*
Bristle Status
present
reduced
absent

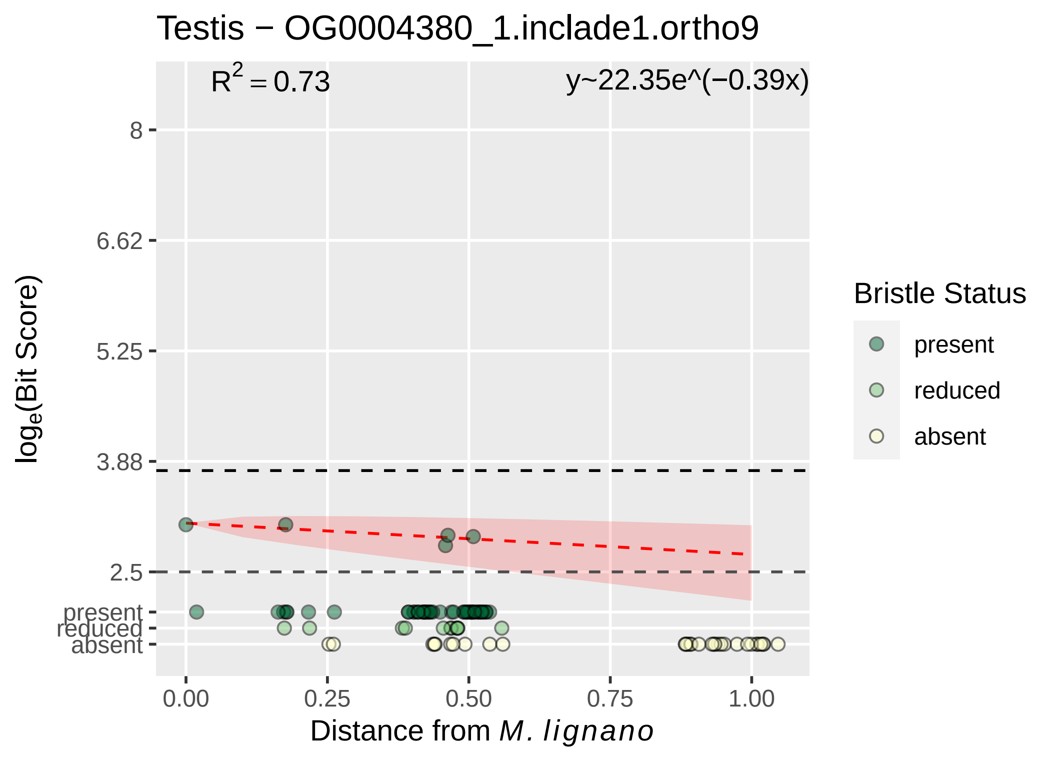
Testis − OG0004380_1.inclade1.ortho9
R^2 = 0.73
y~22.35e^(−0.39x)
8
6.62
5.25
3.88
2.5
present
reduced
absent
0.00
0.25
0.50
0.75
1.00
Distance from M. lignano
log_e(Bit Score)
Bristle Status
present
reduced
absent

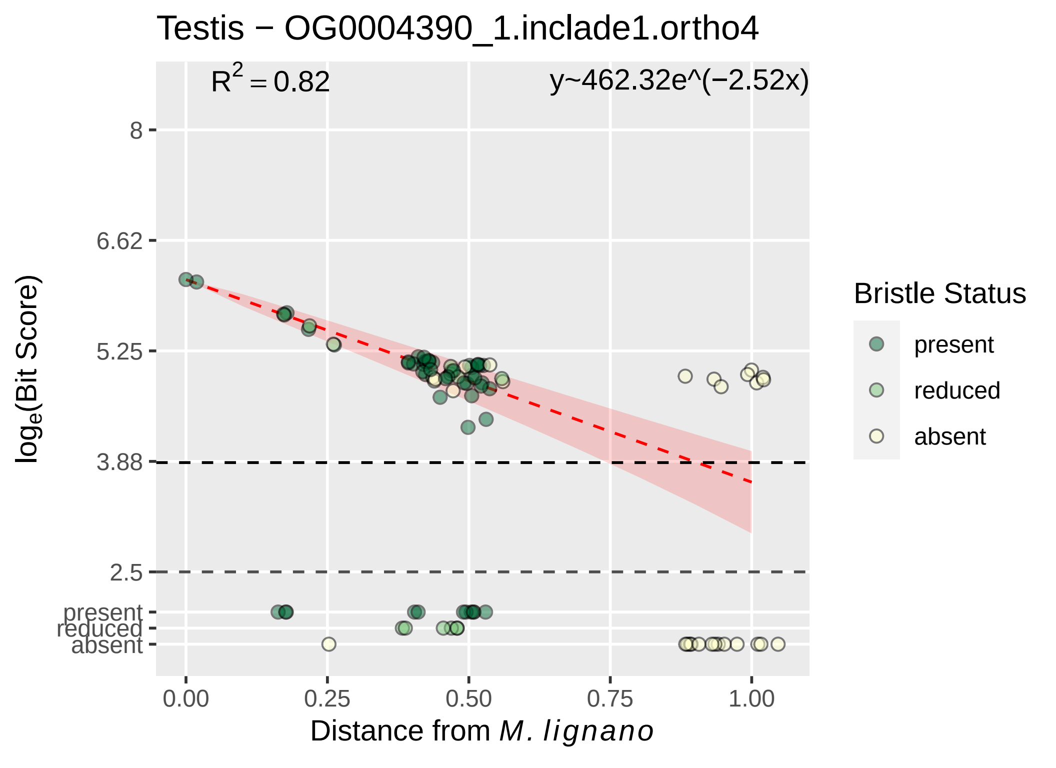

Testis − OG0004390_1.inclade1.ortho4
$R^2 = 0.82$
y~462.32e^(−2.52x)
8
6.62
5.25
3.88
2.5
present
reduced
absent
0.00
0.25
0.50
0.75
1.00
Distance from M. lignano
$log_e$(Bit Score)
Bristle Status
present
reduced
absent

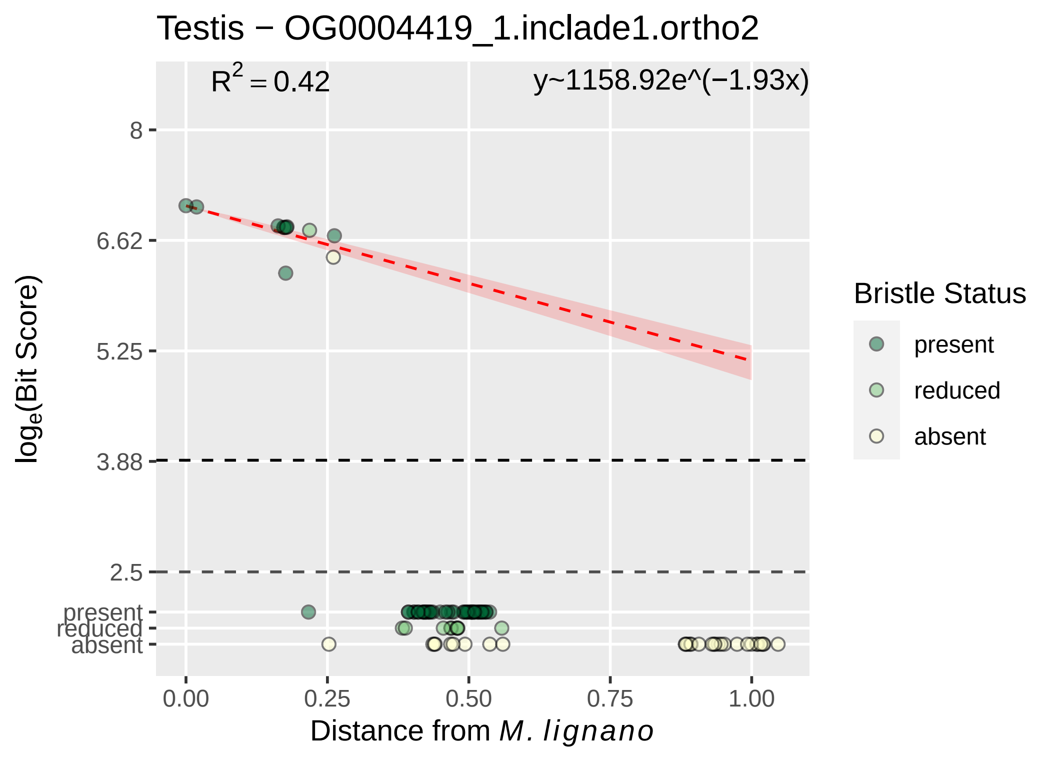

Testis − OG0004419_1.inclade1.ortho2
$R^2 = 0.42$
y~1158.92e^(−1.93x)
8
6.62
5.25
3.88
2.5
present
reduced
absent
0.00
0.25
0.50
0.75
1.00
Distance from M. lignano
loge(Bit Score)
Bristle Status
present
reduced
absent

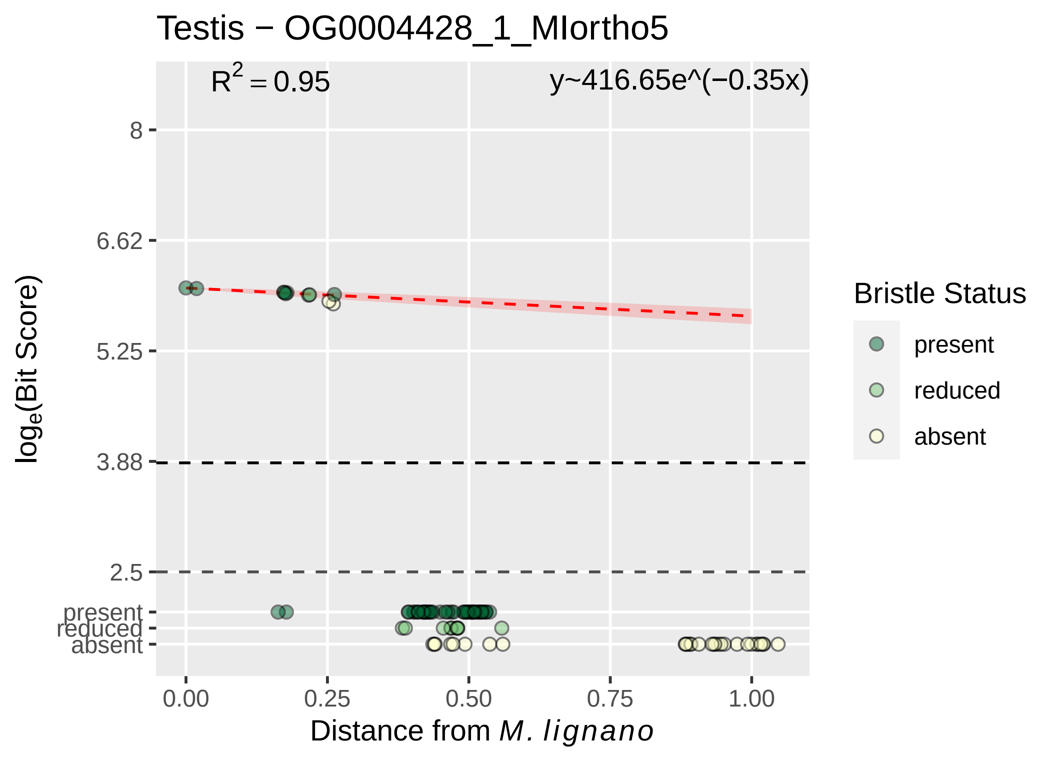

Testis − OG0004428_1_MIortho5
R2 = 0.95
y~416.65e^(−0.35x)
8
6.62
5.25
3.88
2.5
present
reduced
absent
0.00
0.25
0.50
0.75
1.00
Distance from M. lignano
loge(Bit Score)
Bristle Status
present
reduced
absent

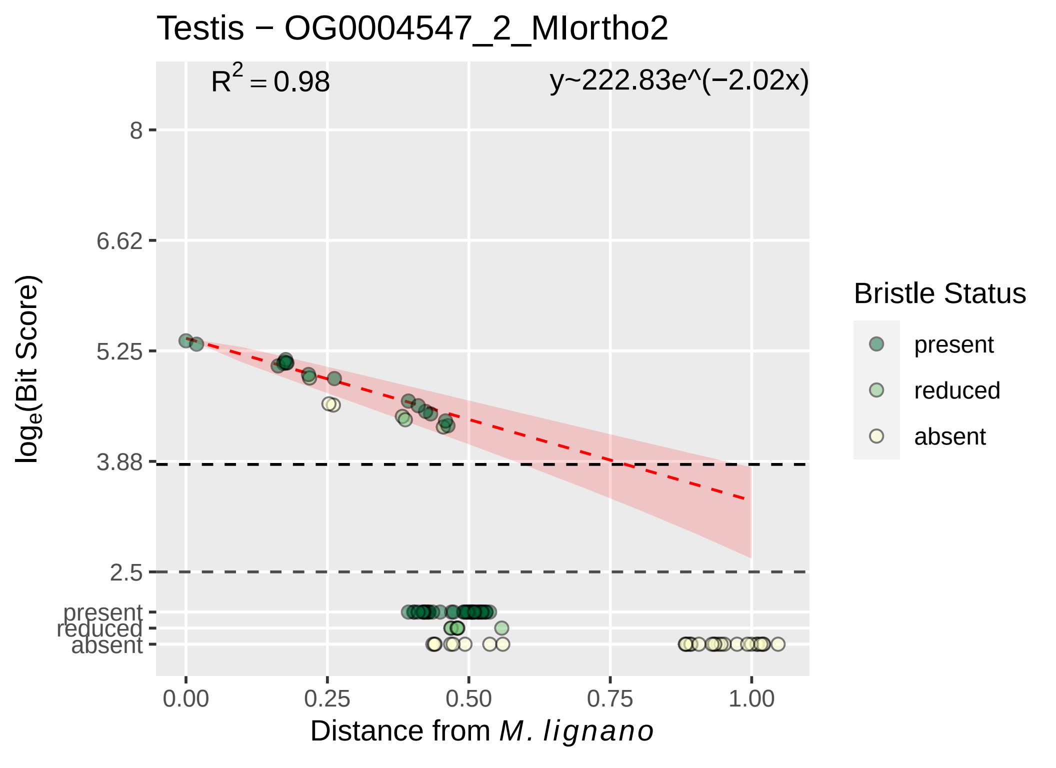

Testis − OG0004547_2_MIortho2
$R^2 = 0.98$
y~222.83e^(−2.02x)
8
6.62
5.25
3.88
2.5
present
reduced
absent
$\log_e$(Bit Score)
0.00
0.25
0.50
0.75
1.00
Distance from *M. lignano*
Bristle Status
present
reduced
absent

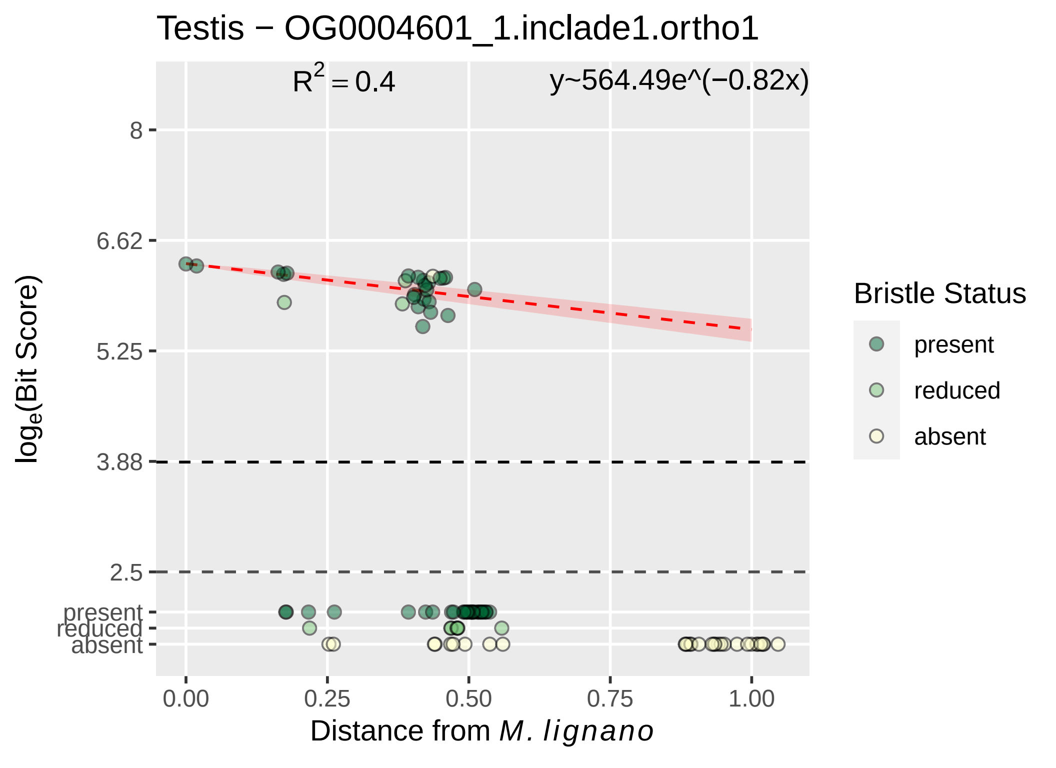

Testis − OG0004601_1.inclade1.ortho1
R2 = 0.4
y~564.49e^(−0.82x)
8
6.62
5.25
3.88
2.5
present
reduced
absent
0.00
0.25
0.50
0.75
1.00
Distance from M. lignano
loge(Bit Score)
Bristle Status
present
reduced
absent

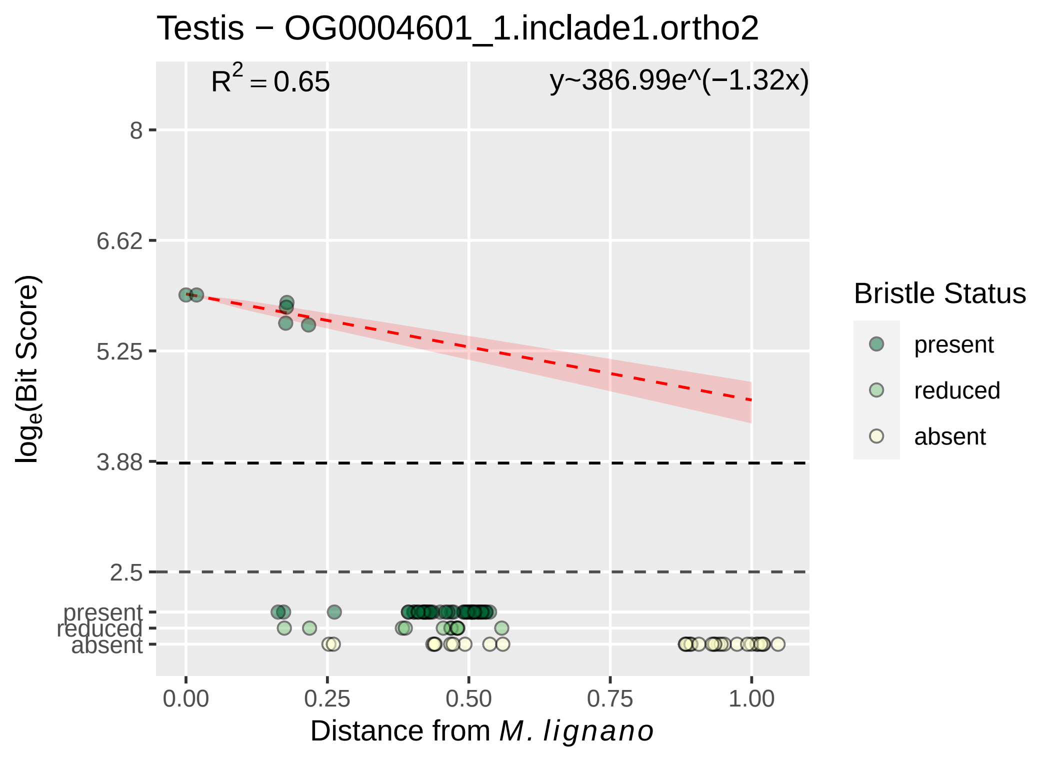

Testis − OG0004601_1.inclade1.ortho2
$R^2 = 0.65$
y~386.99e^(−1.32x)
8
6.62
5.25
3.88
2.5
present
reduced
absent
$log_e$(Bit Score)
0.00
0.25
0.50
0.75
1.00
Distance from *M. lignano*
Bristle Status
present
reduced
absent

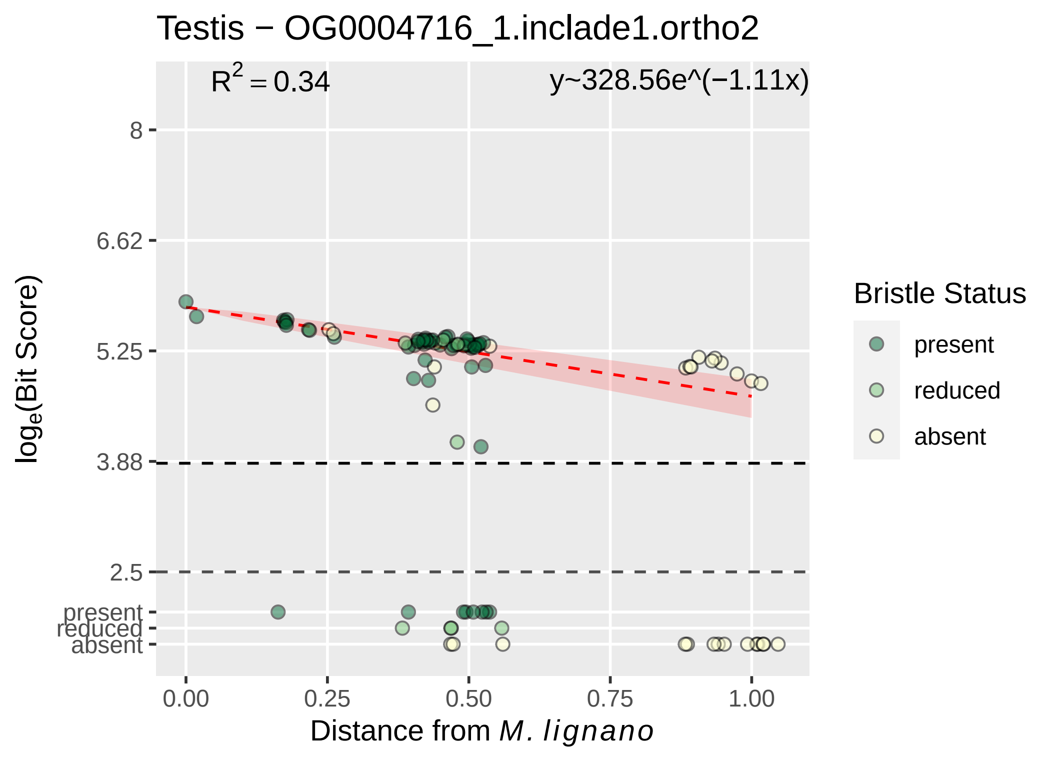

Testis − OG0004716_1.inclade1.ortho2
$R^2 = 0.34$
y~328.56e^(−1.11x)
8
6.62
5.25
3.88
2.5
present
reduced
absent
loge(Bit Score)
0.00
0.25
0.50
0.75
1.00
Distance from M. lignano
Bristle Status
present
reduced
absent

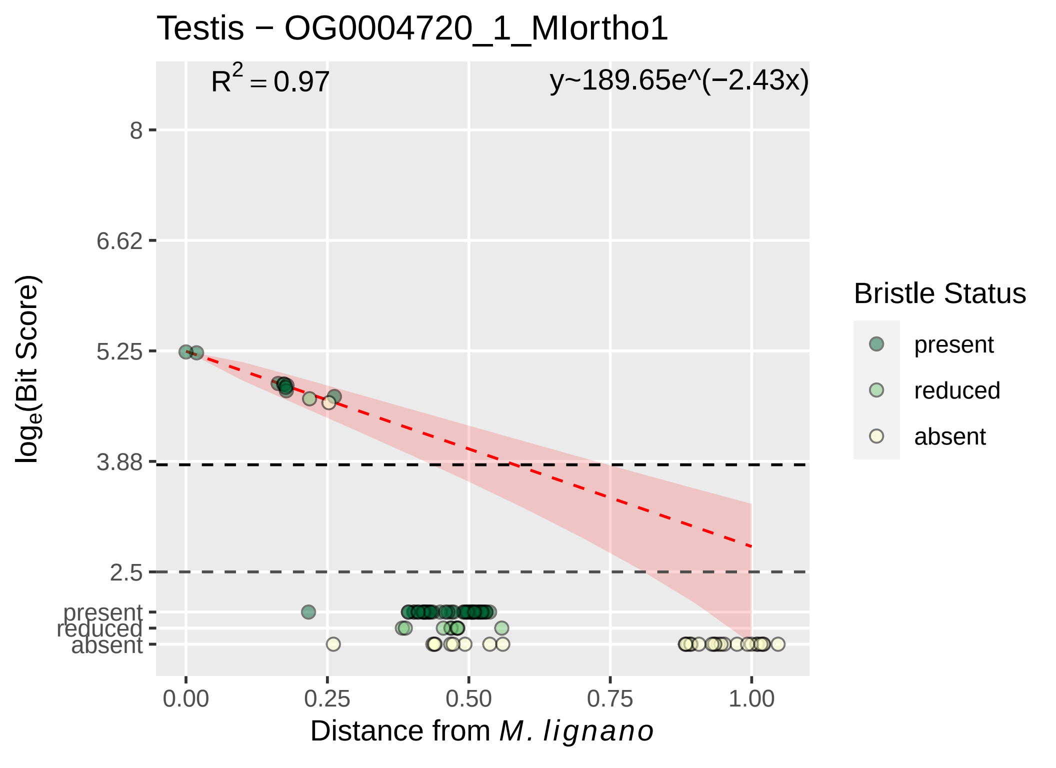

Testis − OG0004720_1_MIortho1
$R^2 = 0.97$
y~189.65e^(−2.43x)
8
6.62
5.25
3.88
2.5
present
reduced
absent
$\log_e$(Bit Score)
0.00
0.25
0.50
0.75
1.00
Distance from *M. lignano*
Bristle Status
present
reduced
absent

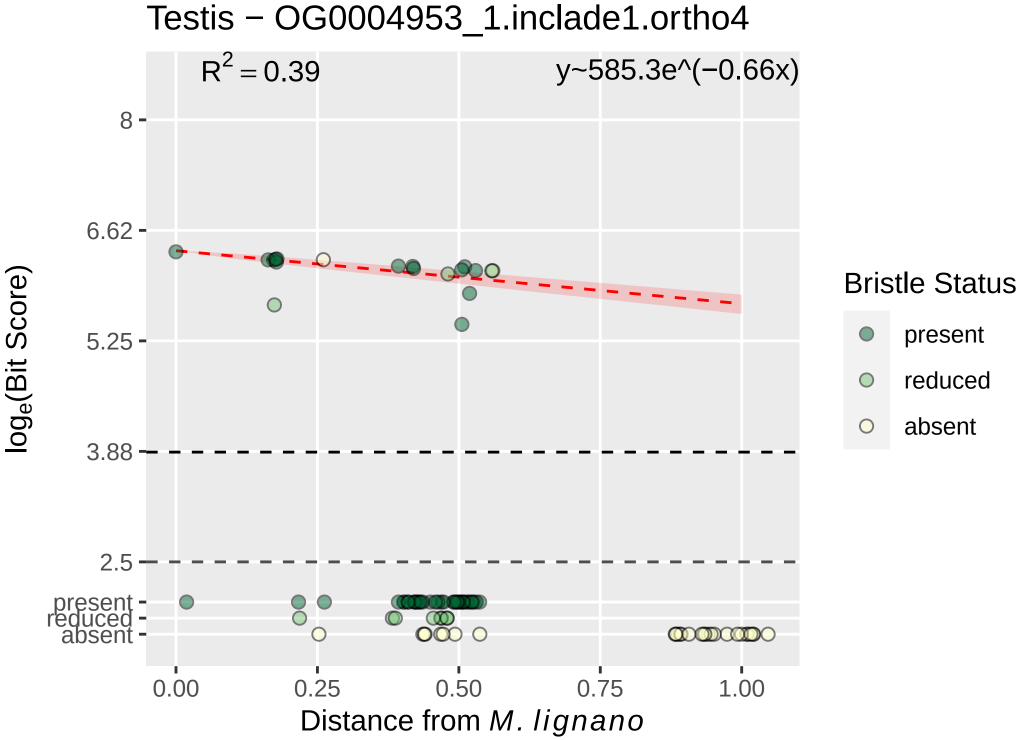

Testis − OG0004953_1.inclade1.ortho4
R2 = 0.39
y~585.3e^(−0.66x)
8
6.62
5.25
3.88
2.5
present
reduced
absent
loge(Bit Score)
0.00
0.25
0.50
0.75
1.00
Distance from M. lignano
Bristle Status
present
reduced
absent

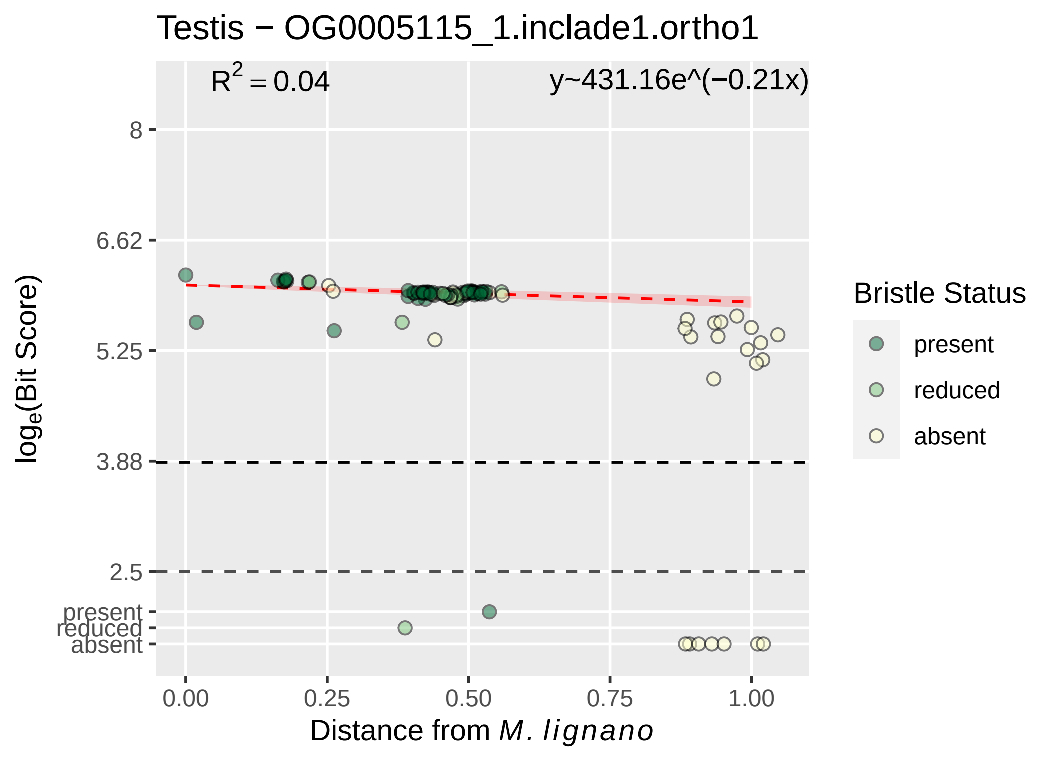
Testis − OG0005115_1.inclade1.ortho1
R² = 0.04
y~431.16e^(−0.21x)
8
6.62
5.25
3.88
2.5
present
reduced
absent
0.00
0.25
0.50
0.75
1.00
Distance from M. lignano
loge(Bit Score)
Bristle Status
present
reduced
absent

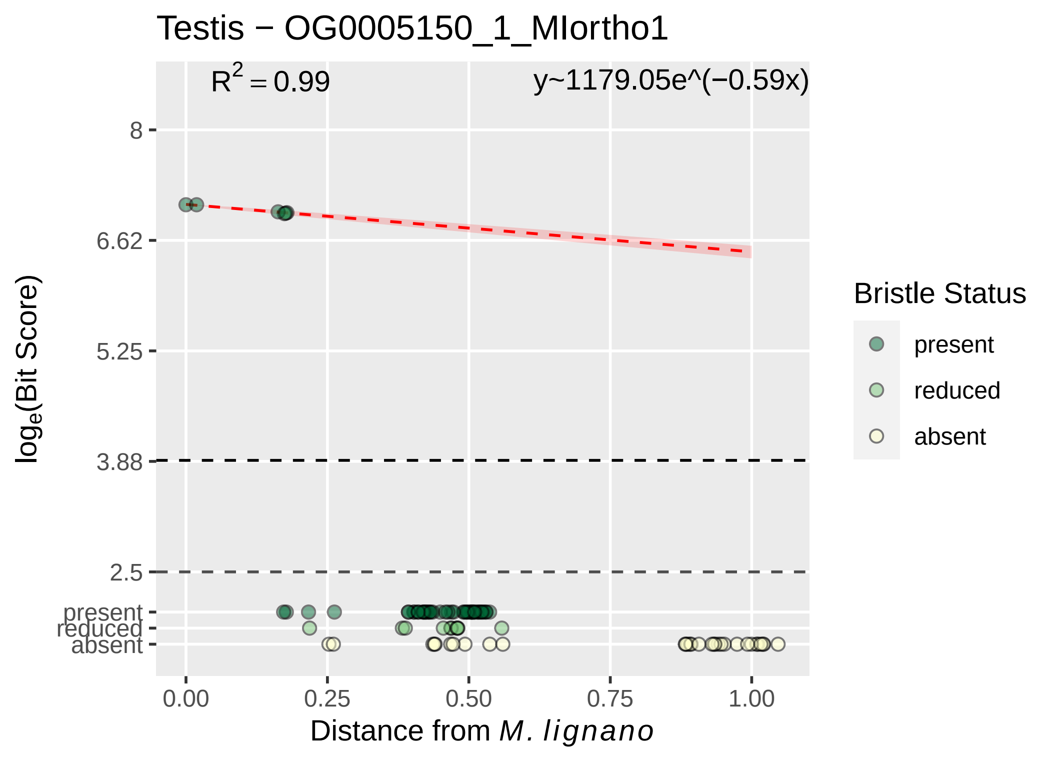

Testis − OG0005150_1_MIortho1
R^2 = 0.99
y~1179.05e^(−0.59x)
8
6.62
5.25
3.88
2.5
present
reduced
absent
0.00
0.25
0.50
0.75
1.00
Distance from M. lignano
log_e(Bit Score)
Bristle Status
present
reduced
absent

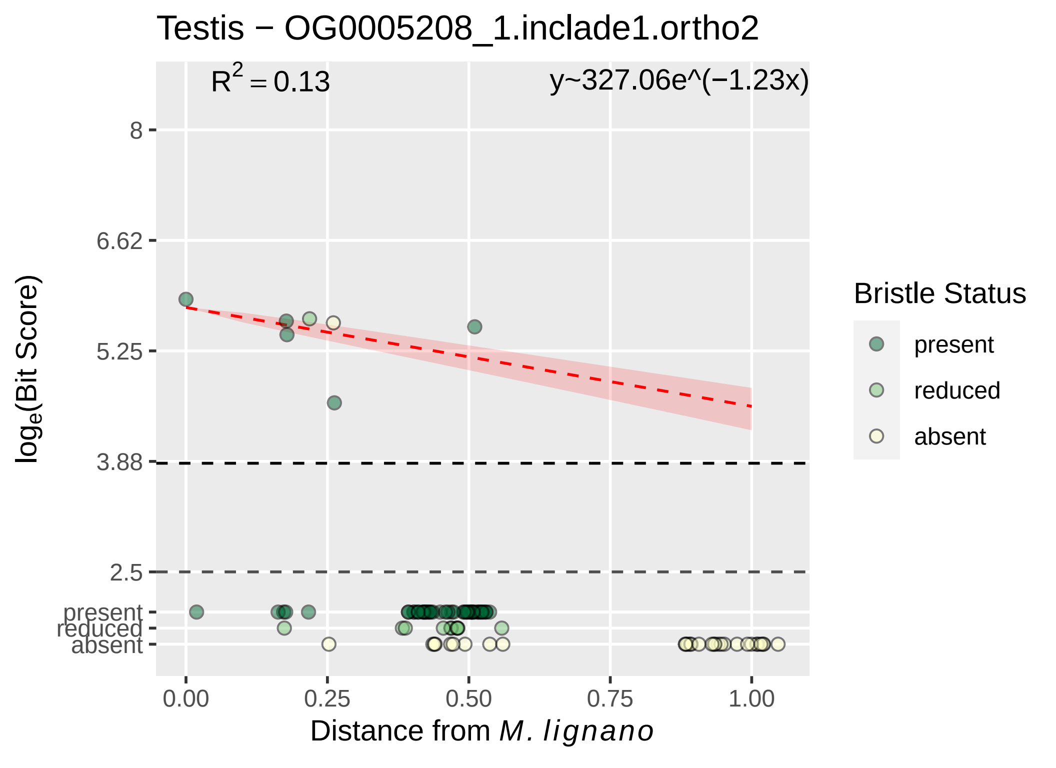

Testis − OG0005208_1.inclade1.ortho2
R^2 = 0.13
y~327.06e^(−1.23x)
8
6.62
5.25
3.88
2.5
present
reduced
absent
0.00
0.25
0.50
0.75
1.00
Distance from M. lignano
loge(Bit Score)
Bristle Status
present
reduced
absent

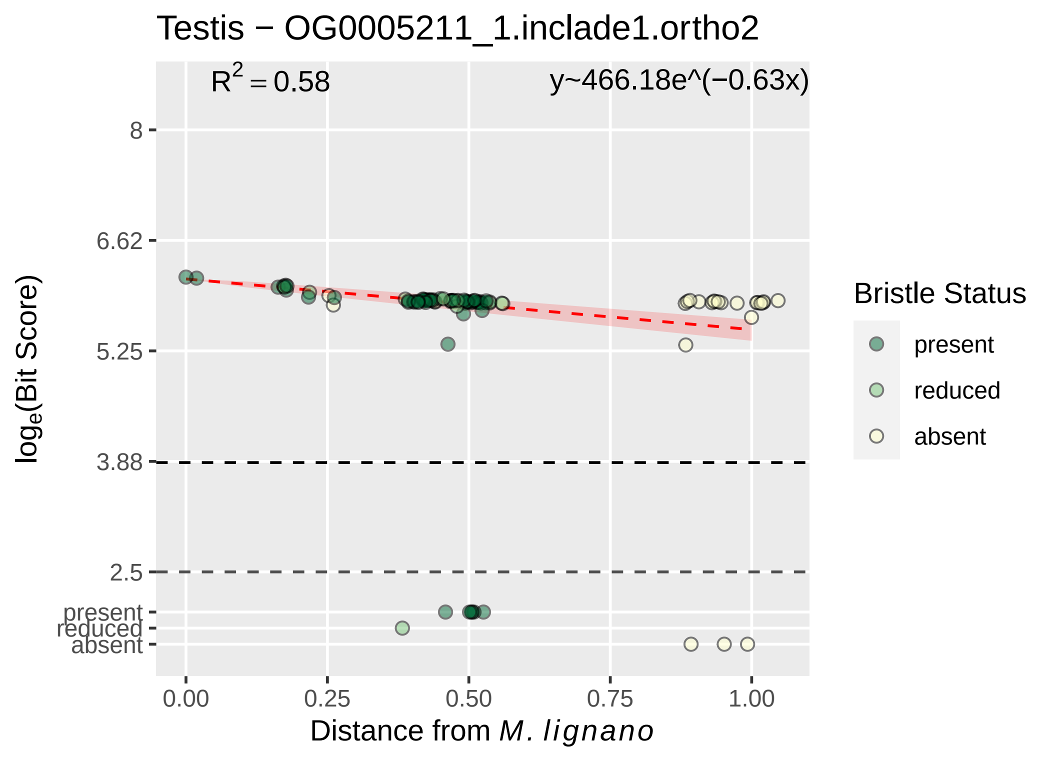

Testis − OG0005211_1.inclade1.ortho2
$R^2 = 0.58$
y~466.18e^(−0.63x)
8
6.62
5.25
3.88
2.5
present
reduced
absent
$\log_e$(Bit Score)
0.00
0.25
0.50
0.75
1.00
Distance from *M. lignano*
Bristle Status
present
reduced
absent

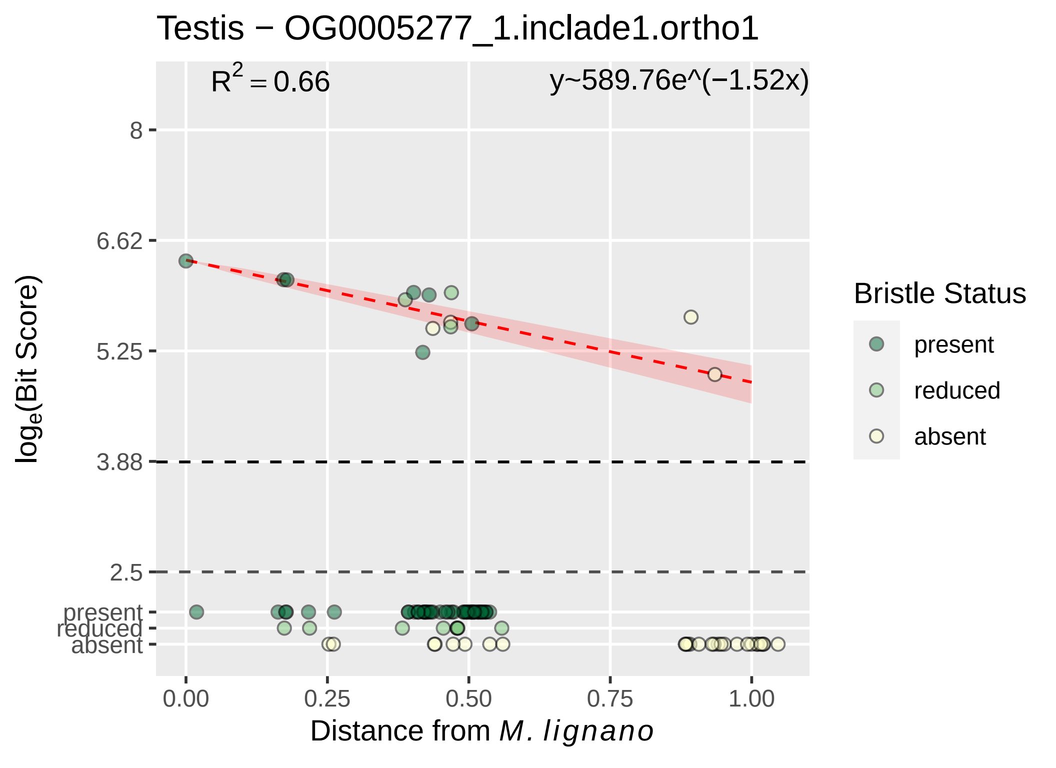

Testis − OG0005277_1.inclade1.ortho1
$R^2 = 0.66$
y~589.76e^(−1.52x)
8
6.62
5.25
3.88
2.5
present
reduced
absent
0.00
0.25
0.50
0.75
1.00
$\log_e$(Bit Score)
Distance from *M. lignano*
Bristle Status
present
reduced
absent

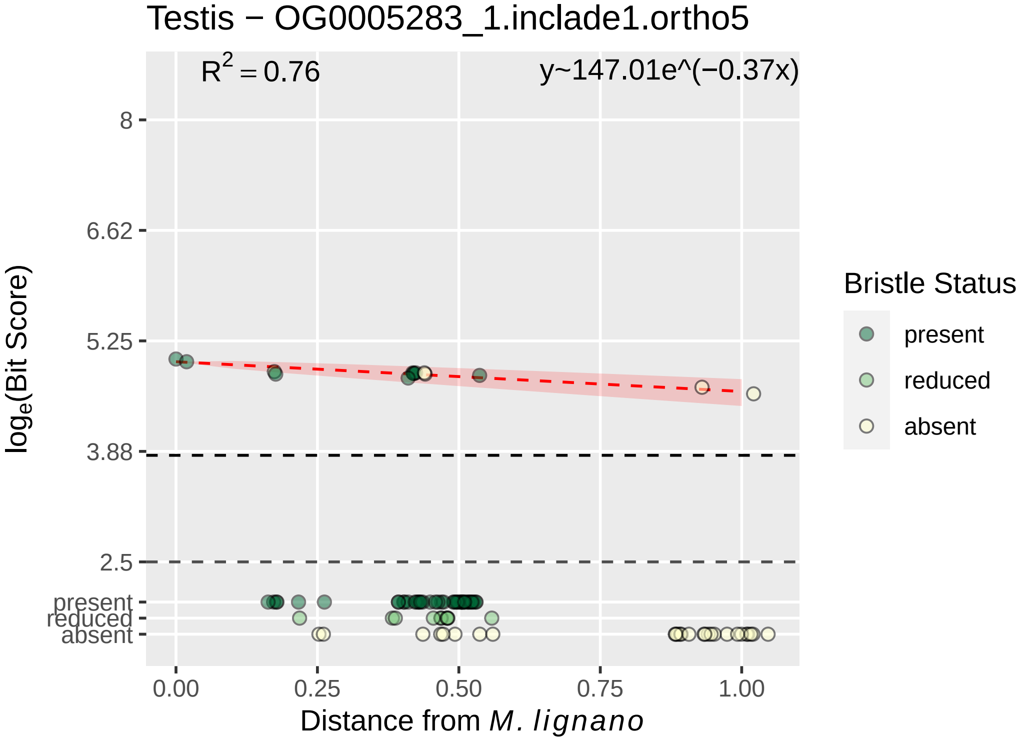

Testis − OG0005283_1.inclade1.ortho5
$R^2 = 0.76$
y~147.01e^(−0.37x)
8
6.62
5.25
3.88
2.5
present
reduced
absent
log_e(Bit Score)
0.00
0.25
0.50
0.75
1.00
Distance from M. lignano
Bristle Status
present
reduced
absent

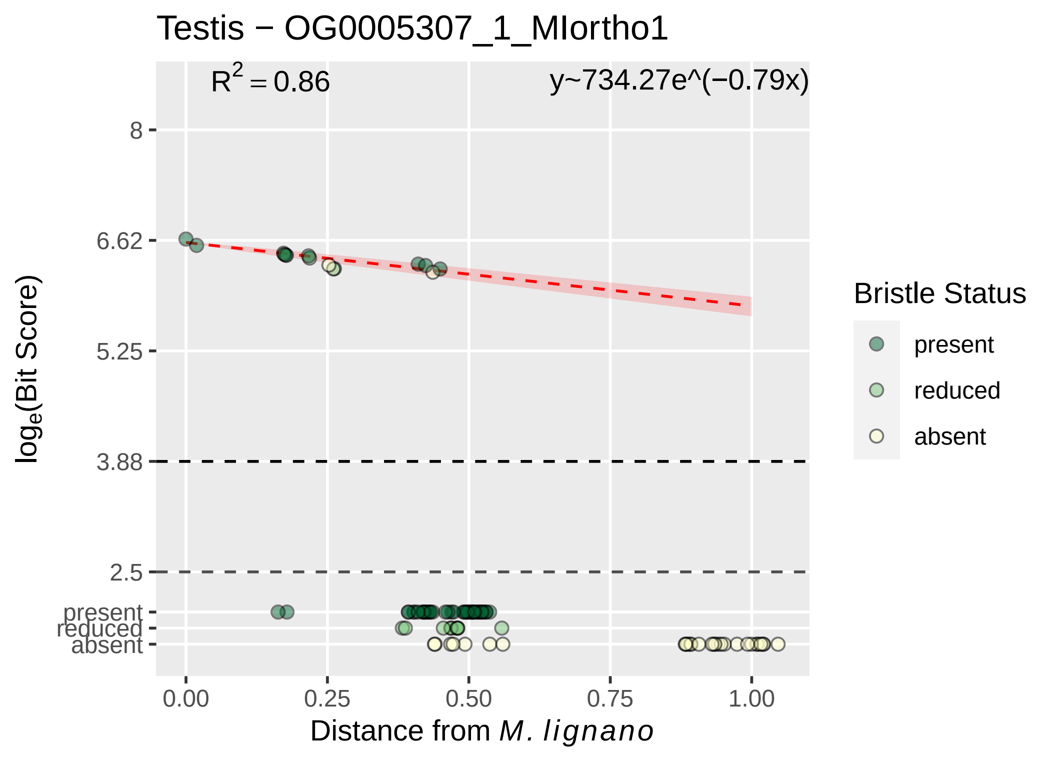

Testis − OG0005307_1_MIortho1
R^2 = 0.86
y~734.27e^(−0.79x)
8
6.62
5.25
3.88
2.5
present
reduced
absent
0.00
0.25
0.50
0.75
1.00
Distance from M. lignano
log_e(Bit Score)
Bristle Status
present
reduced
absent

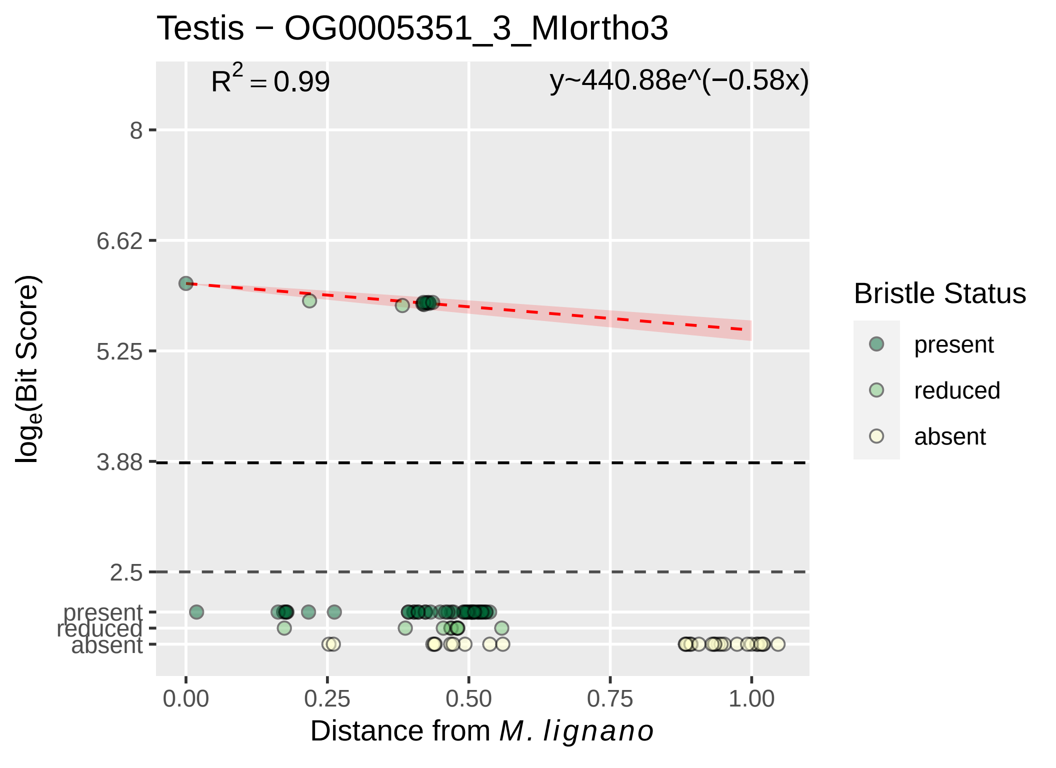

Testis − OG0005351_3_MIortho3
R² = 0.99
y~440.88e^(−0.58x)
8
6.62
5.25
3.88
2.5
present
reduced
absent
log_e(Bit Score)
0.00
0.25
0.50
0.75
1.00
Distance from M. lignano
Bristle Status
present
reduced
absent

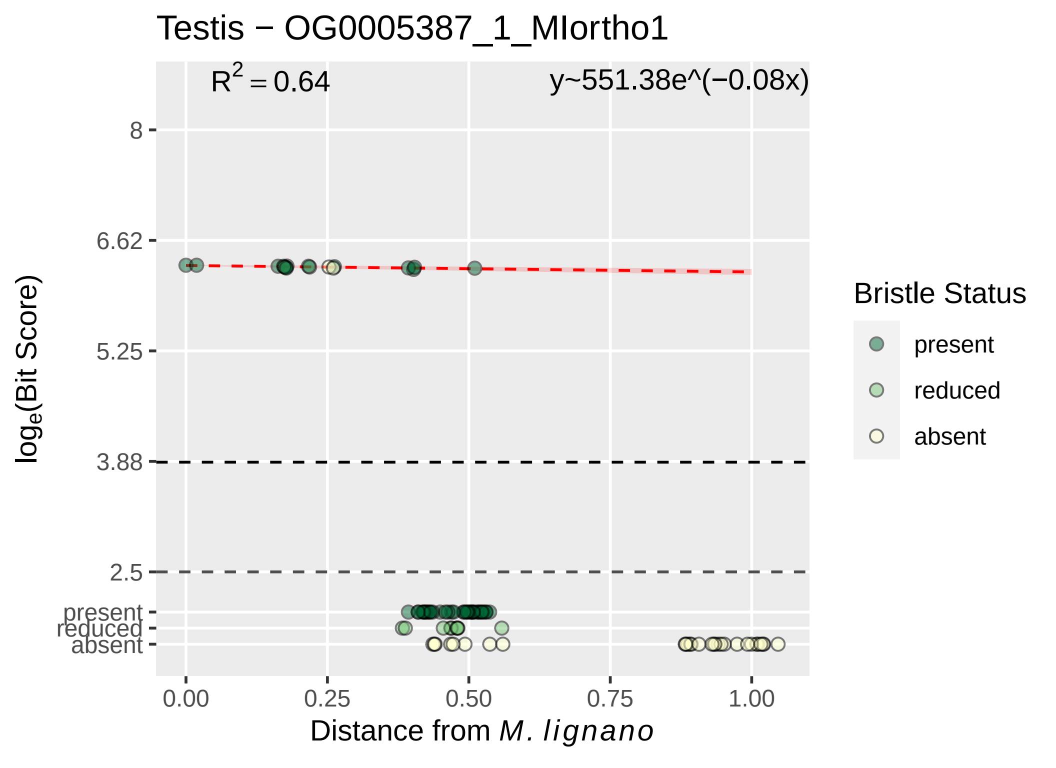

Testis − OG0005387_1_MIortho1
$R^2 = 0.64$
y~551.38e^(−0.08x)
8
6.62
5.25
3.88
2.5
present
reduced
absent
$\log_e$(Bit Score)
0.00
0.25
0.50
0.75
1.00
Distance from *M. lignano*
Bristle Status
present
reduced
absent

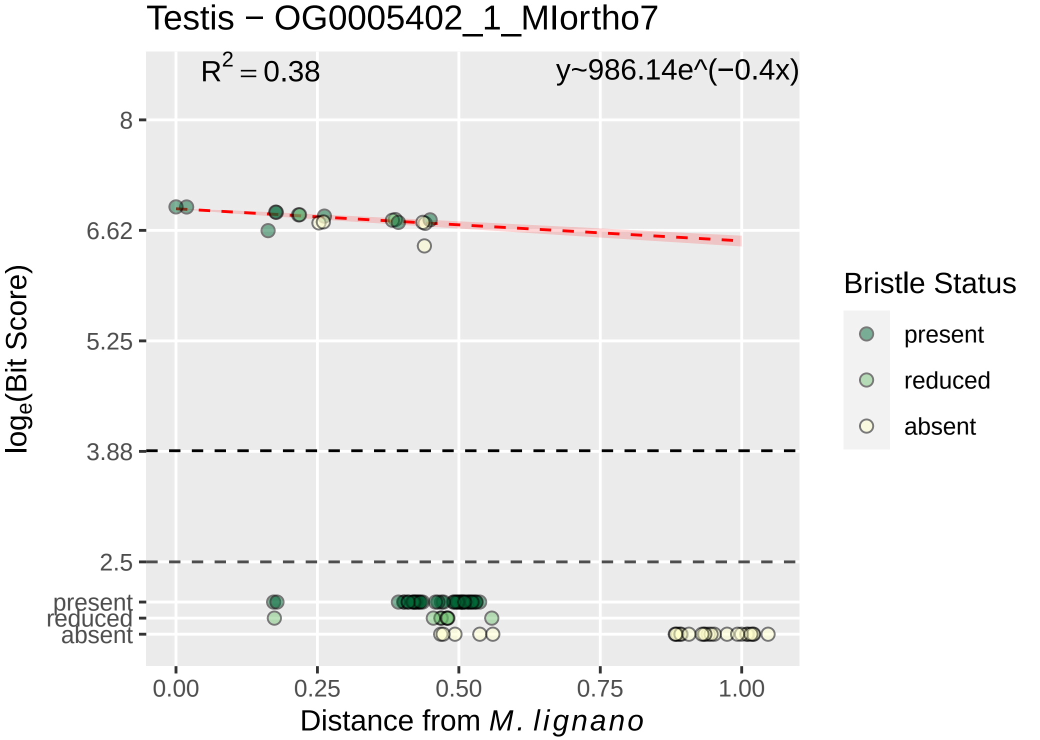

Testis − OG0005402_1_MIortho7
$R^2 = 0.38$
y~986.14e^(−0.4x)
8
6.62
5.25
3.88
2.5
present
reduced
absent
$\log_e$(Bit Score)
0.00
0.25
0.50
0.75
1.00
Distance from *M. lignano*
Bristle Status
present
reduced
absent

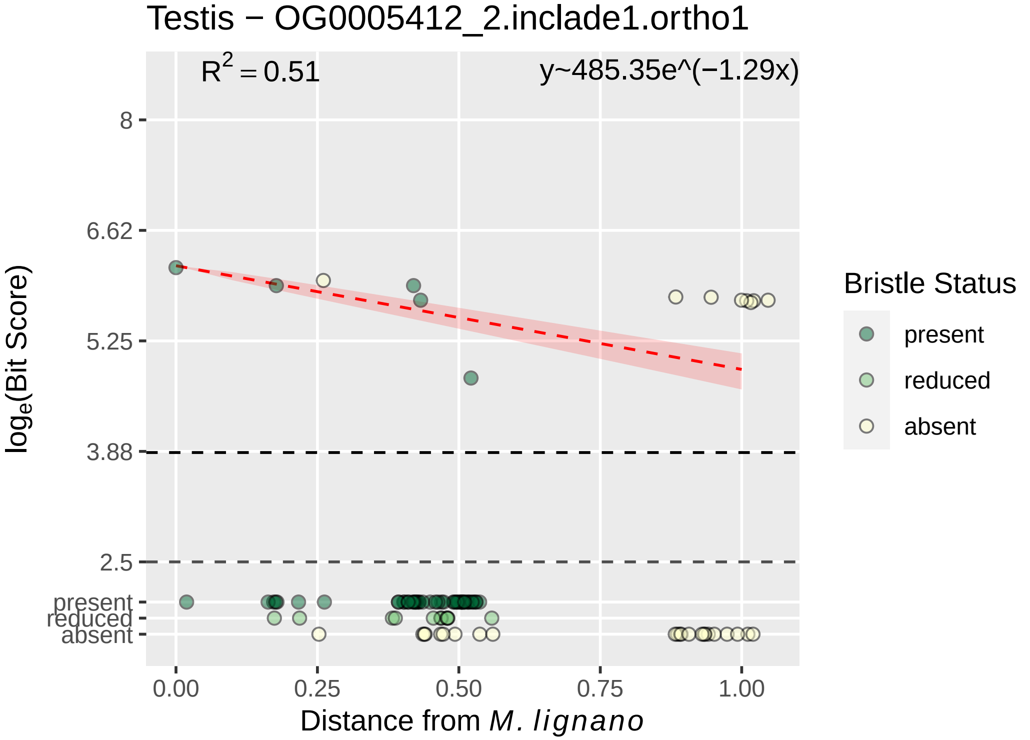

Testis − OG0005412_2.inclade1.ortho1
$R^2 = 0.51$
y~485.35e^(−1.29x)
8
6.62
5.25
3.88
2.5
present
reduced
absent
0.00
0.25
0.50
0.75
1.00
Distance from *M. lignano*
$\log_e$(Bit Score)
Bristle Status
present
reduced
absent

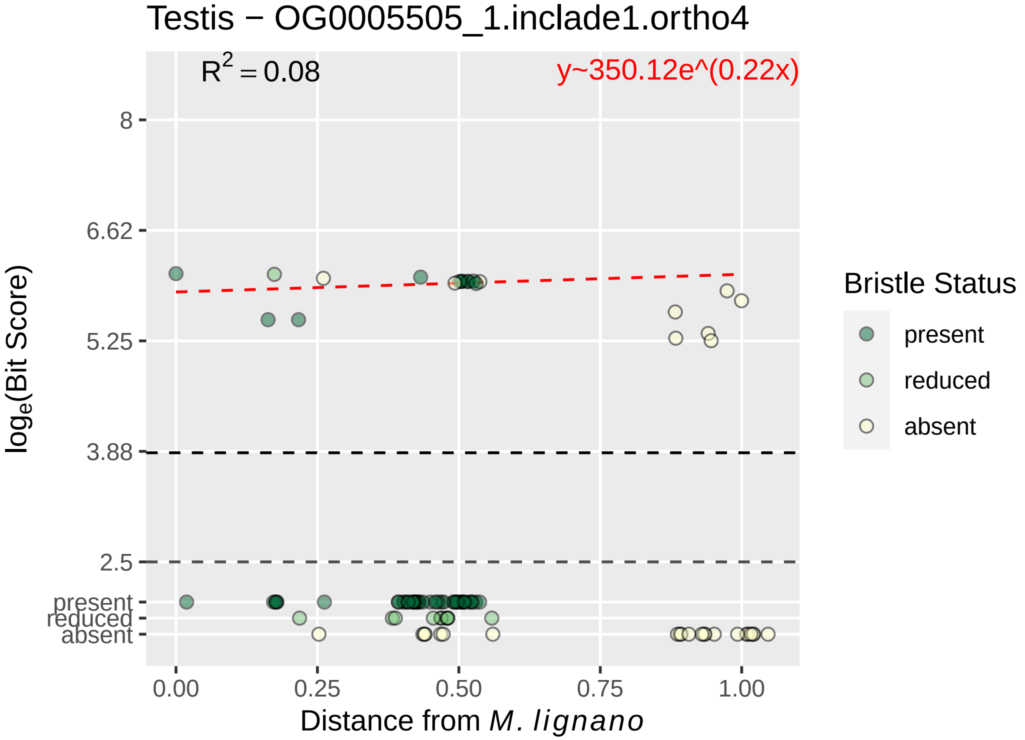
Testis − OG0005505_1.inclade1.ortho4
$R^2 = 0.08$
y~350.12e^(0.22x)
8
6.62
5.25
3.88
2.5
present
reduced
absent
0.00
0.25
0.50
0.75
1.00
Distance from M. lignano
log$_e$(Bit Score)
Bristle Status
present
reduced
absent

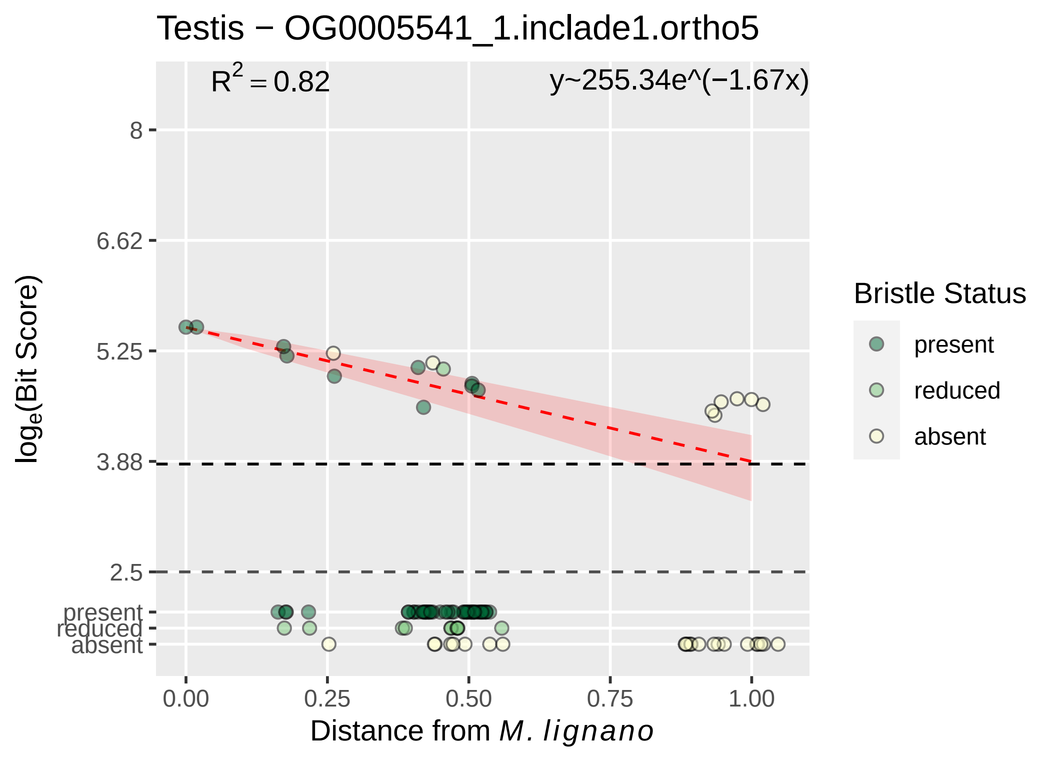
Testis − OG0005541_1.inclade1.ortho5
R^2 = 0.82
y~255.34e^(−1.67x)
8
6.62
5.25
3.88
2.5
present
reduced
absent
log_e(Bit Score)
0.00
0.25
0.50
0.75
1.00
Distance from M. lignano
Bristle Status
present
reduced
absent

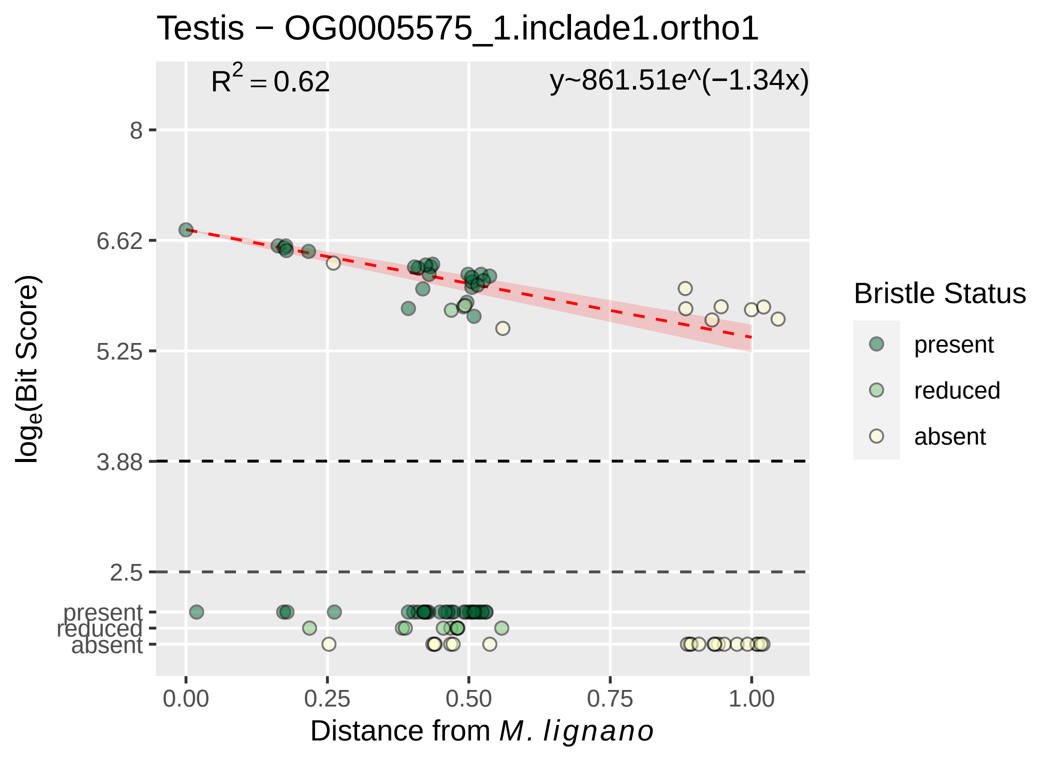

Testis − OG0005575_1.inclade1.ortho1
R^2 = 0.62
y~861.51e^(−1.34x)
8
6.62
5.25
3.88
2.5
present
reduced
absent
log_e(Bit Score)
0.00
0.25
0.50
0.75
1.00
Distance from M. lignano
Bristle Status
present
reduced
absent

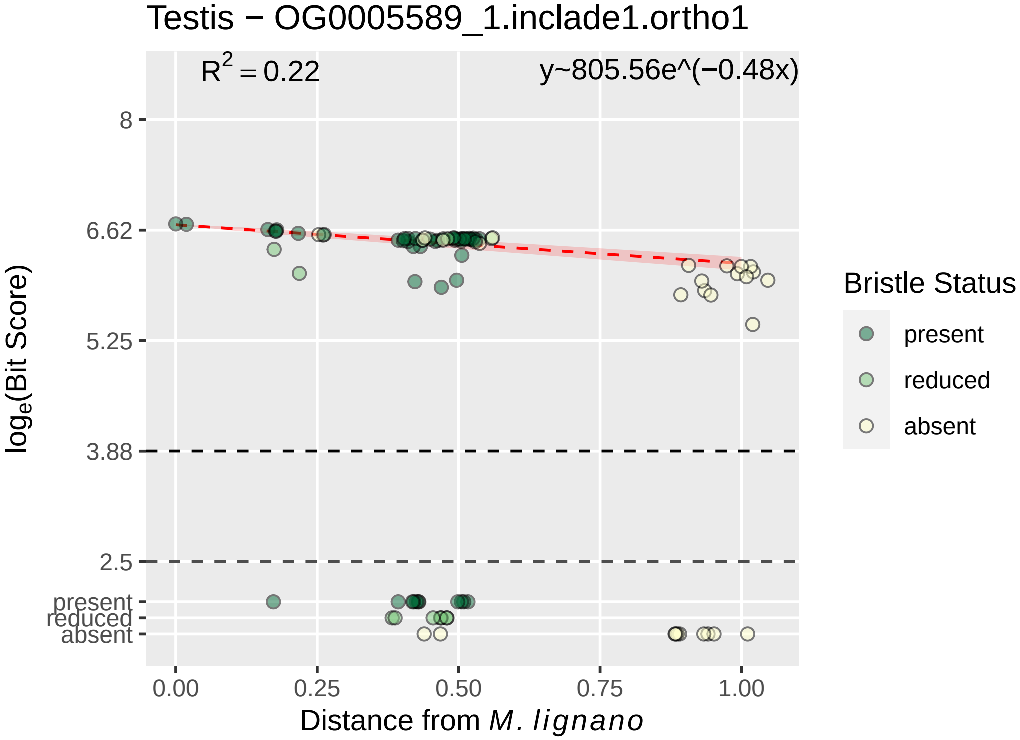

Testis − OG0005589_1.inclade1.ortho1
$R^2 = 0.22$
y~805.56e^(−0.48x)
8
6.62
5.25
3.88
2.5
present
reduced
absent
0.00
0.25
0.50
0.75
1.00
log$_e$(Bit Score)
Distance from M. lignano
Bristle Status
present
reduced
absent

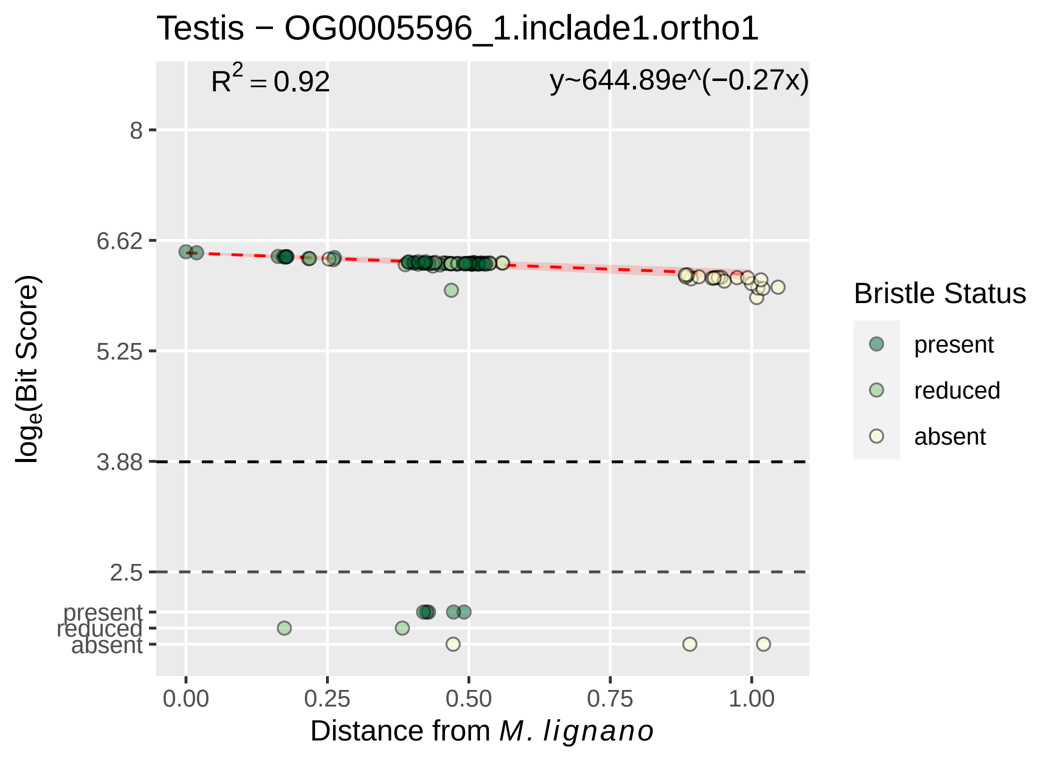

Testis − OG0005596_1.inclade1.ortho1
$R^2 = 0.92$
y~644.89e^(−0.27x)
8
6.62
5.25
3.88
2.5
present
reduced
absent
0.00
0.25
0.50
0.75
1.00
Distance from *M. lignano*
$log_e$(Bit Score)
Bristle Status
present
reduced
absent

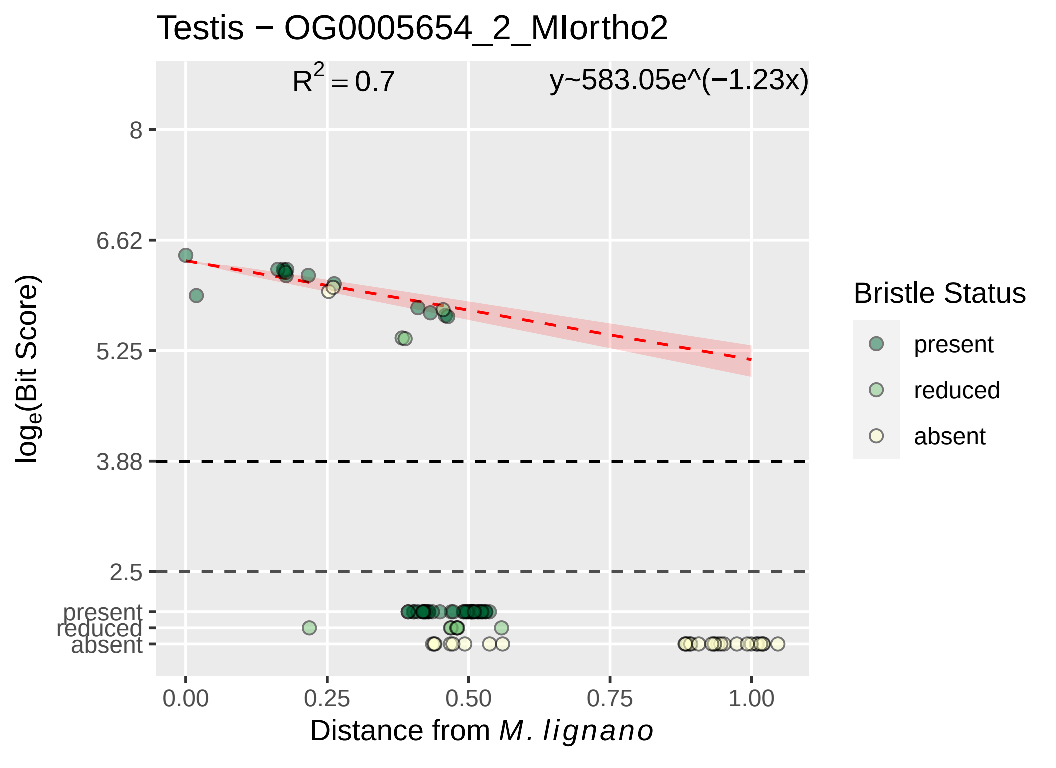

Testis − OG0005654_2_MIortho2
$R^2 = 0.7$
y~583.05e^(−1.23x)
8
6.62
5.25
3.88
2.5
present
reduced
absent
$\log_e$(Bit Score)
0.00
0.25
0.50
0.75
1.00
Distance from *M. lignano*
Bristle Status
present
reduced
absent

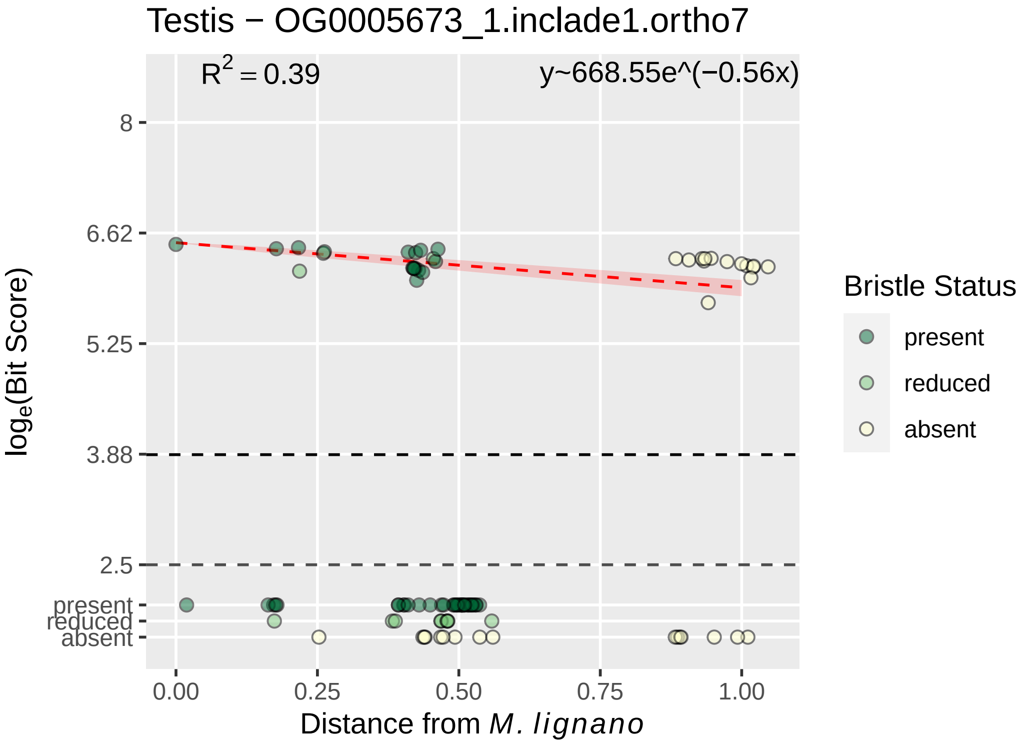

Testis − OG0005673_1.inclade1.ortho7
R^2 = 0.39
y~668.55e^(−0.56x)
8
6.62
5.25
3.88
2.5
present
reduced
absent
0.00
0.25
0.50
0.75
1.00
Distance from M. lignano
loge(Bit Score)
Bristle Status
present
reduced
absent

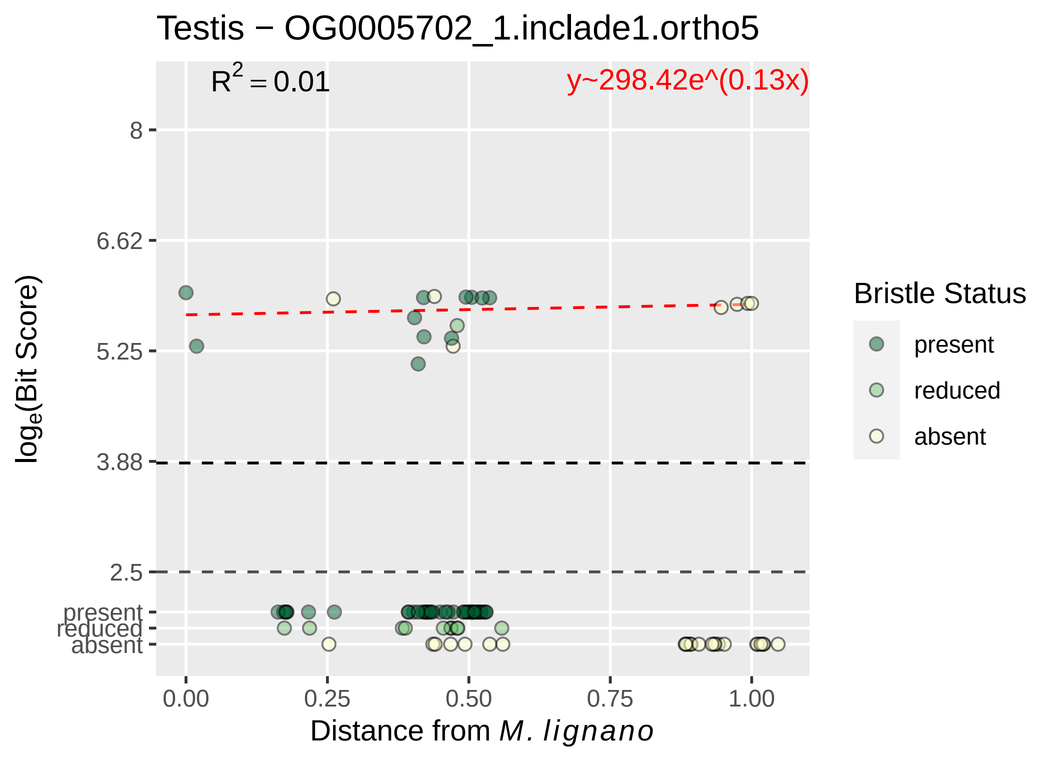

Testis − OG0005702_1.inclade1.ortho5
R$^2$ = 0.01
y~298.42e^(0.13x)
8
6.62
5.25
3.88
2.5
present
reduced
absent
0.00
0.25
0.50
0.75
1.00
log$_e$(Bit Score)
Distance from *M. lignano*
Bristle Status
present
reduced
absent

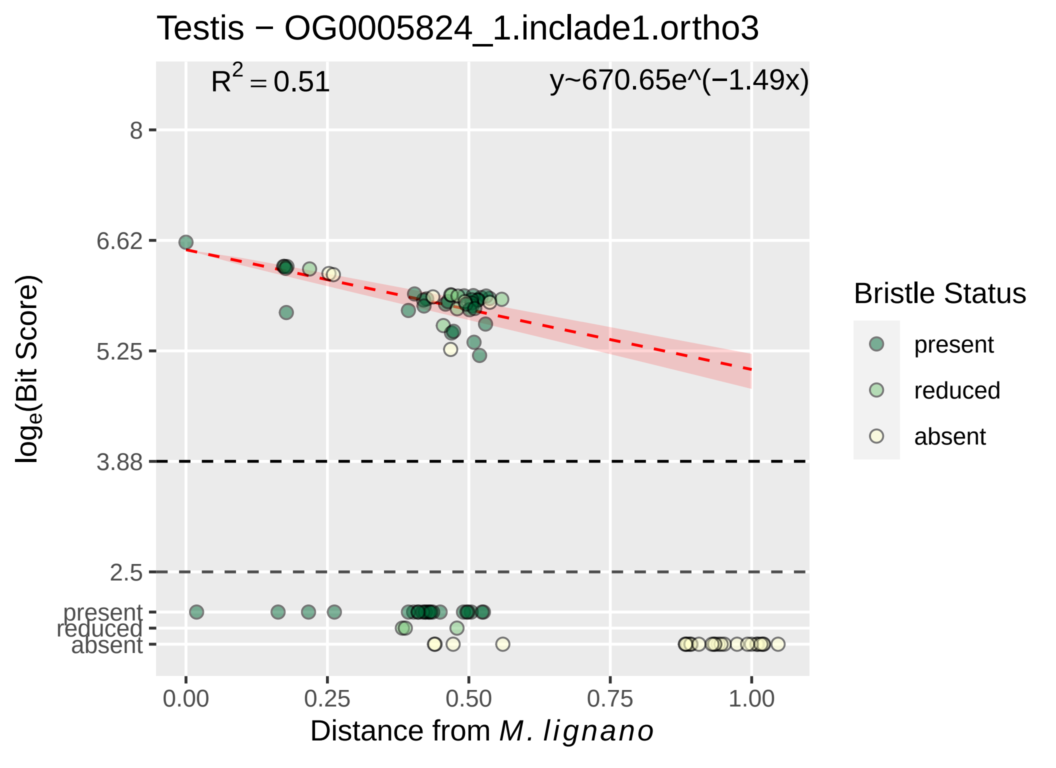

Testis − OG0005824_1.inclade1.ortho3
$R^2 = 0.51$
y~670.65e^(−1.49x)
8
6.62
5.25
3.88
2.5
present
reduced
absent
$log_e$(Bit Score)
0.00
0.25
0.50
0.75
1.00
Distance from *M. lignano*
Bristle Status
present
reduced
absent

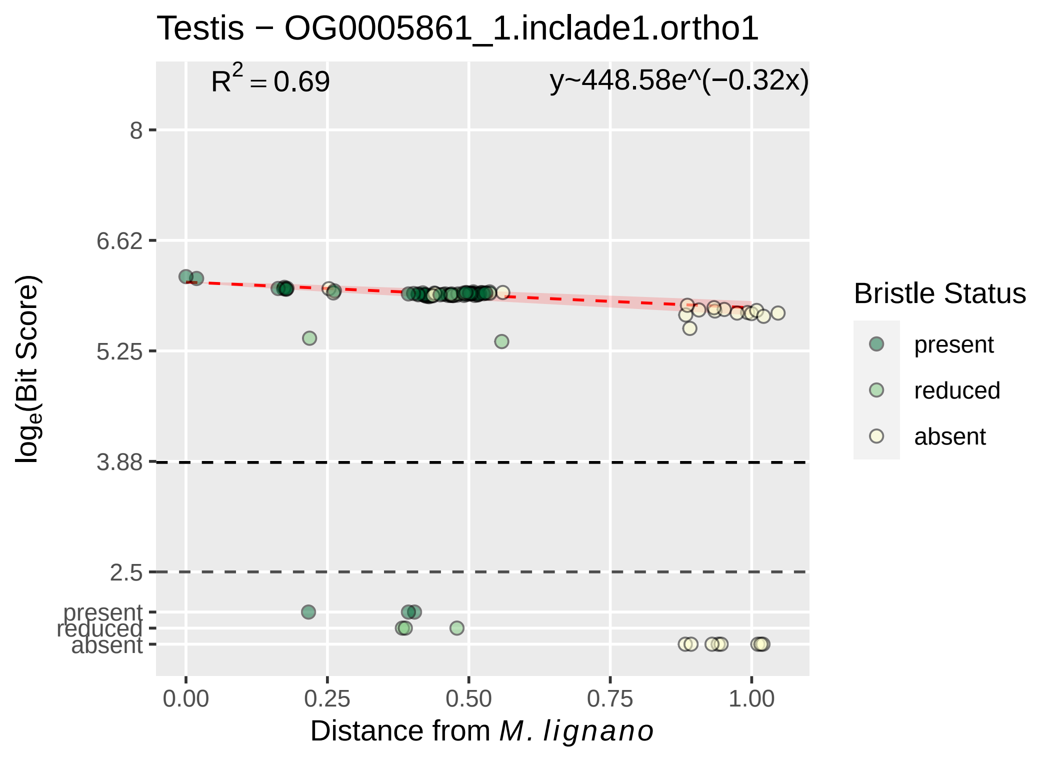

Testis − OG0005861_1.inclade1.ortho1
$R^2 = 0.69$
y~448.58e^(−0.32x)
8
6.62
5.25
3.88
2.5
present
reduced
absent
0.00
0.25
0.50
0.75
1.00
$\log_e$(Bit Score)
Distance from *M. lignano*
Bristle Status
present
reduced
absent

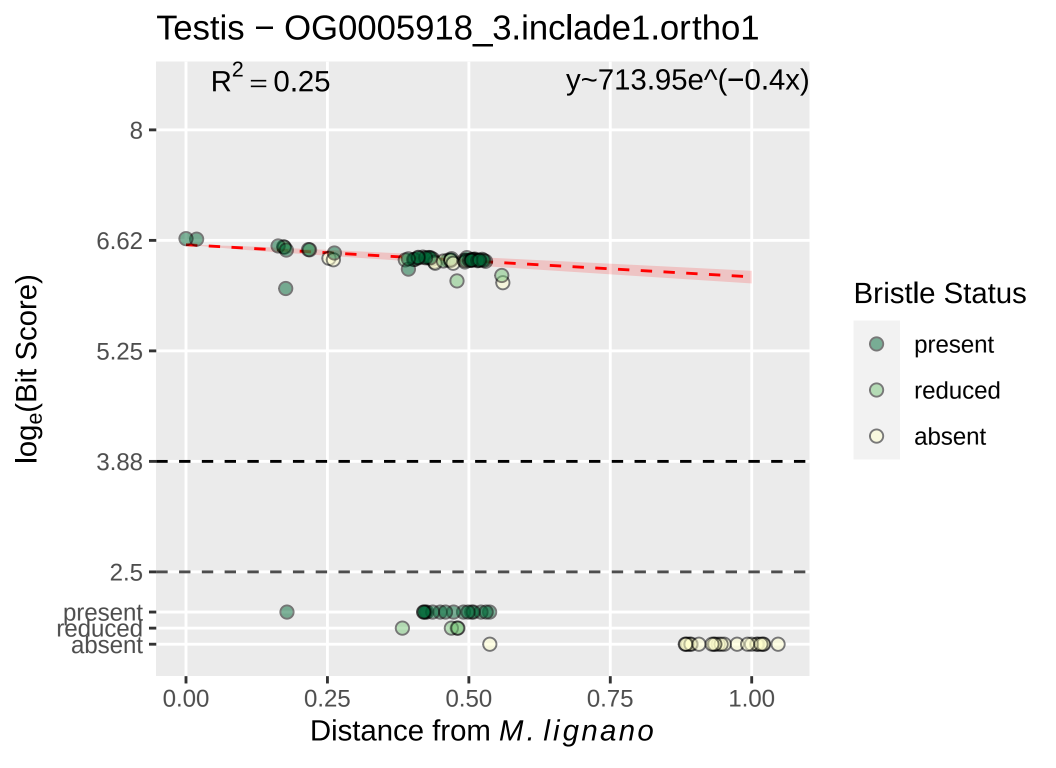

Testis − OG0005918_3.inclade1.ortho1
$R^2 = 0.25$
y~713.95e^(−0.4x)
8
6.62
5.25
3.88
2.5
present
reduced
absent
log$_e$(Bit Score)
0.00
0.25
0.50
0.75
1.00
Distance from *M. lignano*
Bristle Status
present
reduced
absent

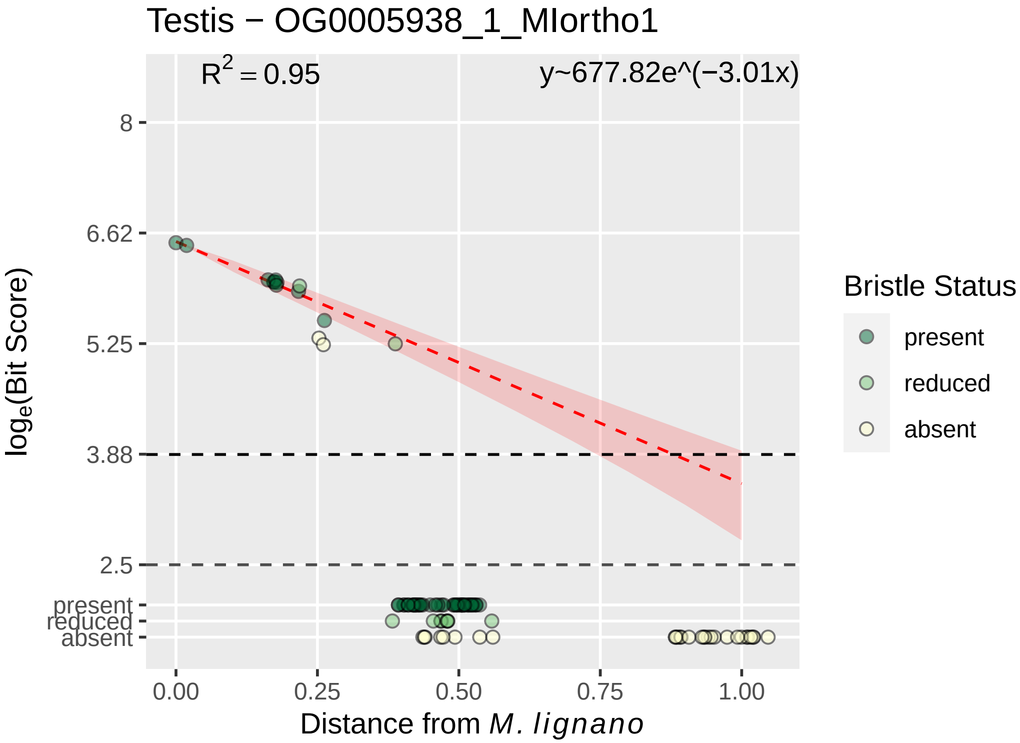

Testis − OG0005938_1_MIortho1
R^2 = 0.95
y~677.82e^(−3.01x)
8
6.62
5.25
3.88
2.5
present
reduced
absent
log_e(Bit Score)
0.00
0.25
0.50
0.75
1.00
Distance from M. lignano
Bristle Status
present
reduced
absent

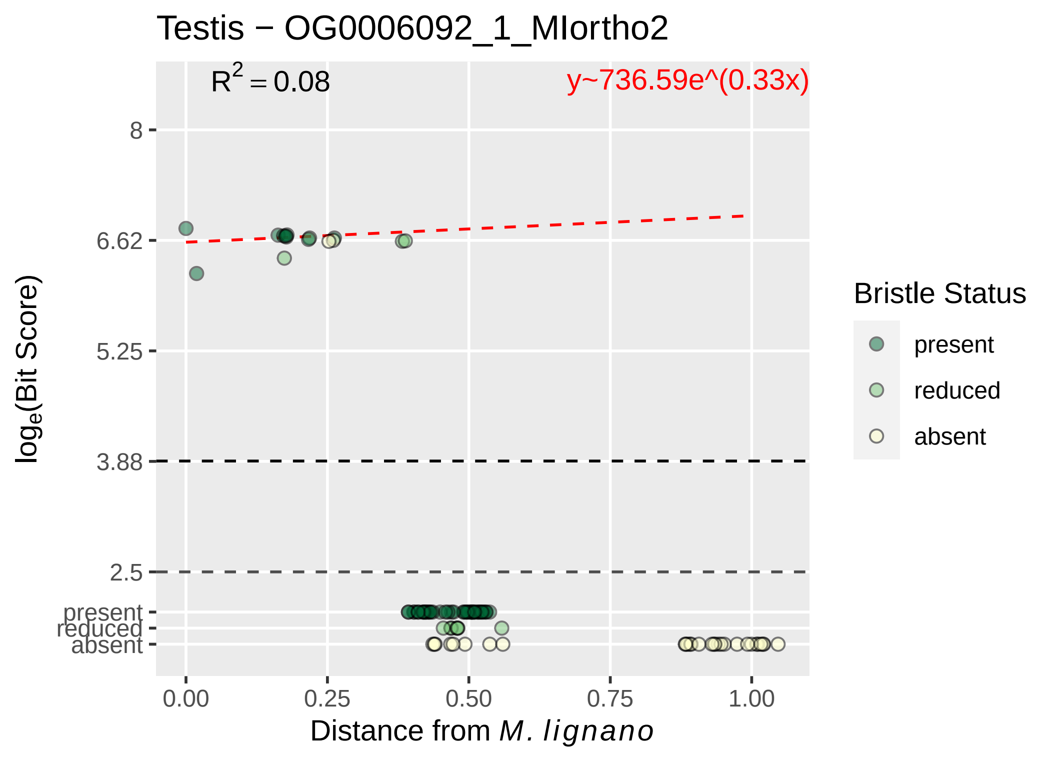

Testis − OG0006092_1_MIortho2
$R^2 = 0.08$
y~736.59e^(0.33x)
8
6.62
5.25
3.88
2.5
present
reduced
absent
0.00
0.25
0.50
0.75
1.00
Distance from *M. lignano*
$\log_e$(Bit Score)
Bristle Status
present
reduced
absent

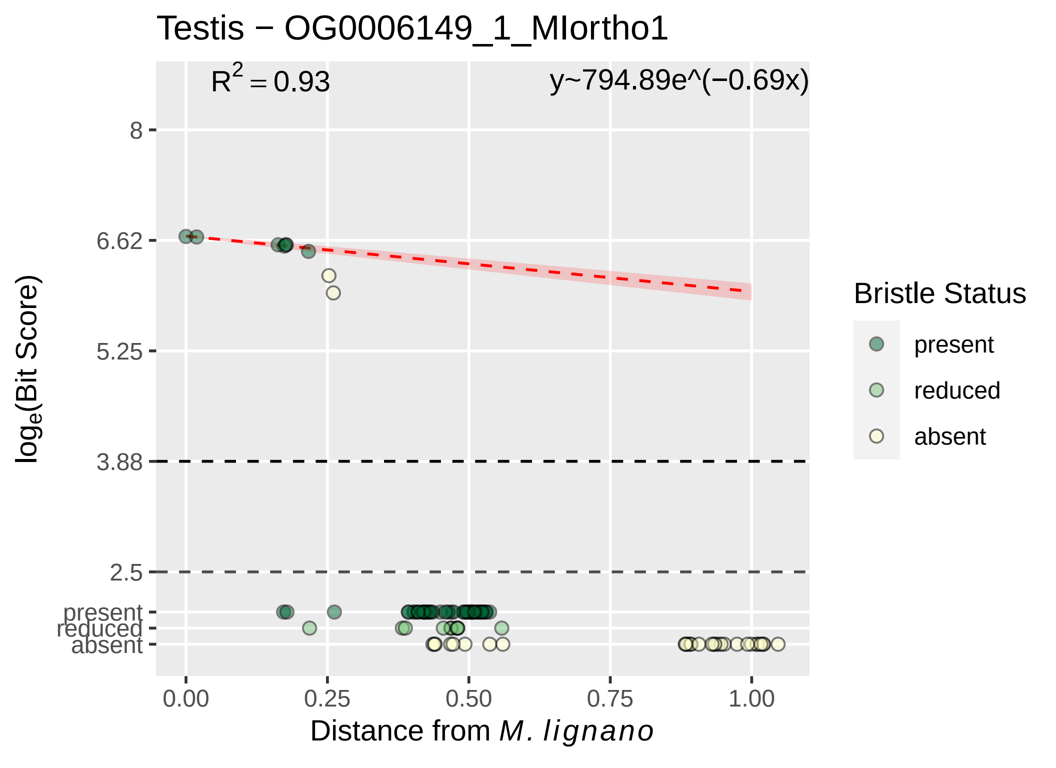

Testis − OG0006149_1_MIortho1
$R^2 = 0.93$
y~794.89e^(−0.69x)
8
6.62
5.25
3.88
2.5
present
reduced
absent
$\log_e$(Bit Score)
0.00
0.25
0.50
0.75
1.00
Distance from *M. lignano*
Bristle Status
present
reduced
absent

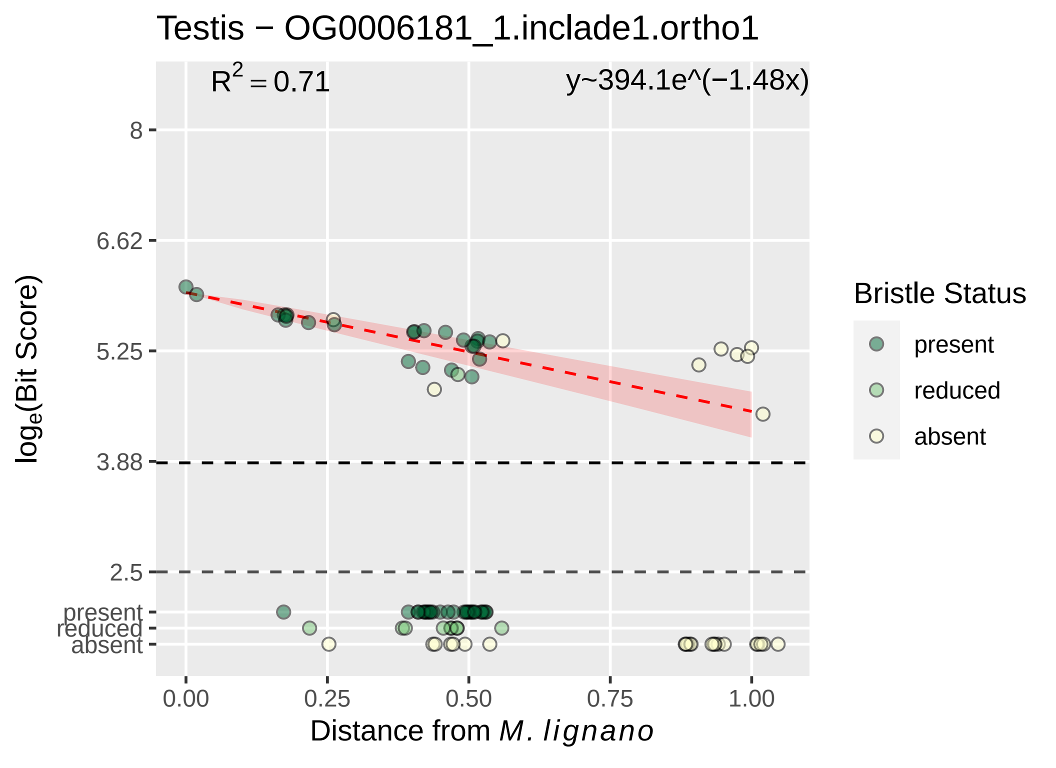

Testis − OG0006181_1.inclade1.ortho1
R² = 0.71
y~394.1e^(−1.48x)
8
6.62
5.25
3.88
2.5
present
reduced
absent
loge(Bit Score)
0.00
0.25
0.50
0.75
1.00
Distance from M. lignano
Bristle Status
present
reduced
absent

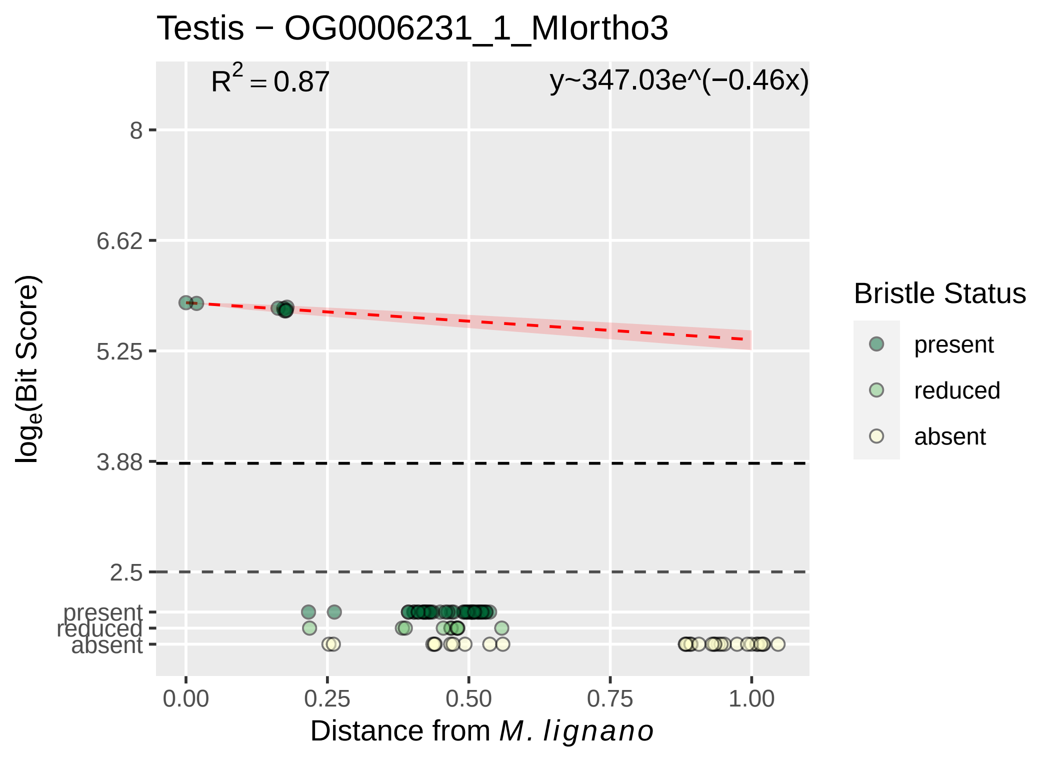

Testis − OG0006231_1_MIortho3
$R^2 = 0.87$
y~347.03e^(−0.46x)
8
6.62
5.25
3.88
2.5
present
reduced
absent
$\log_e$(Bit Score)
0.00
0.25
0.50
0.75
1.00
Distance from *M. lignano*
Bristle Status
present
reduced
absent

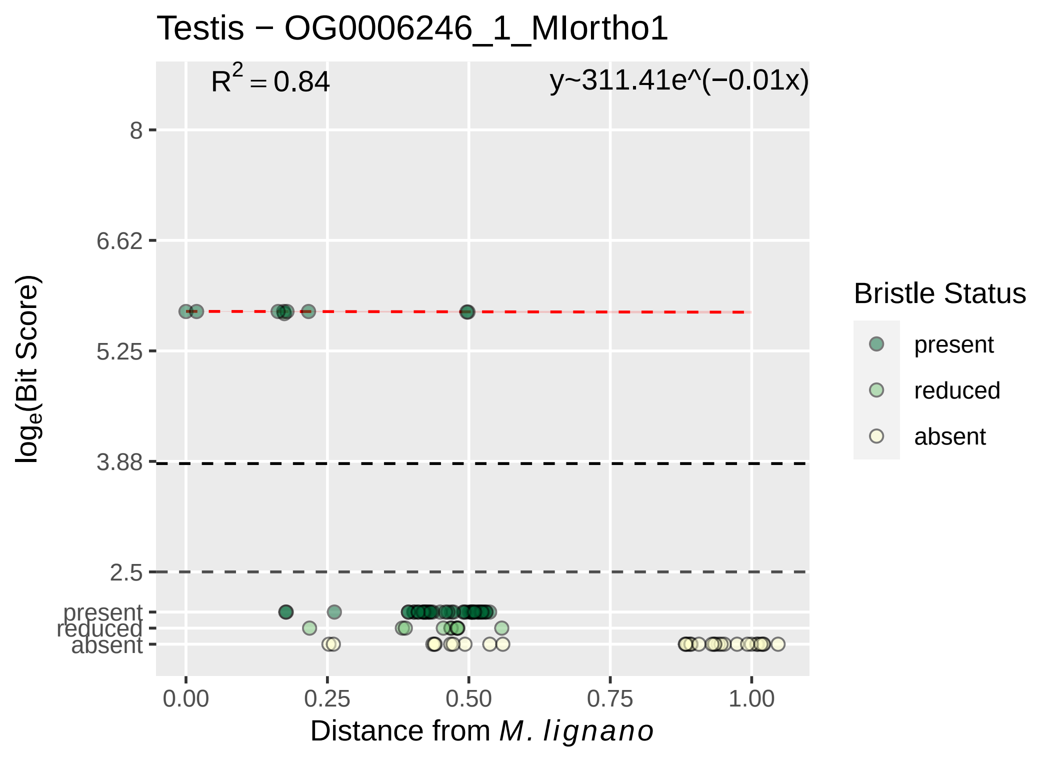
Testis − OG0006246_1_MIortho1
$R^2 = 0.84$
y~311.41e^(−0.01x)
8
6.62
5.25
3.88
2.5
present
reduced
absent
0.00
0.25
0.50
0.75
1.00
Distance from M. lignano
$log_e$(Bit Score)
Bristle Status
present
reduced
absent

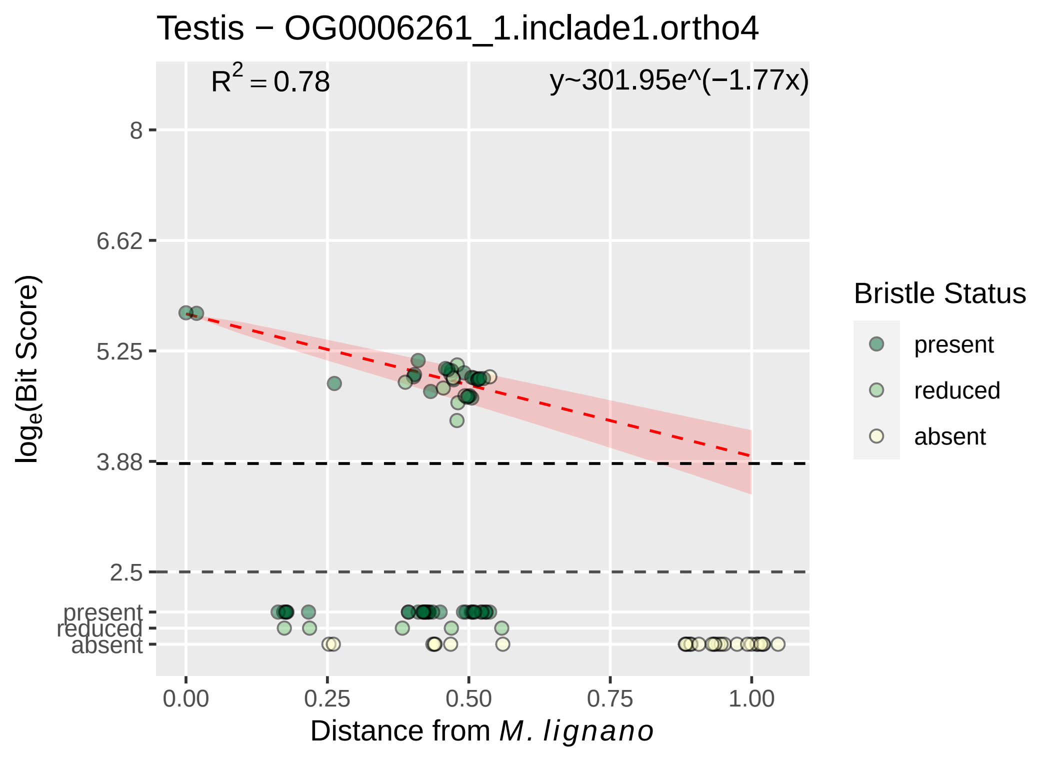

Testis − OG0006261_1.inclade1.ortho4
$R^2 = 0.78$
y~301.95e^(−1.77x)
8
6.62
5.25
3.88
2.5
present
reduced
absent
0.00
0.25
0.50
0.75
1.00
Distance from M. lignano
$\log_e$(Bit Score)
Bristle Status
present
reduced
absent

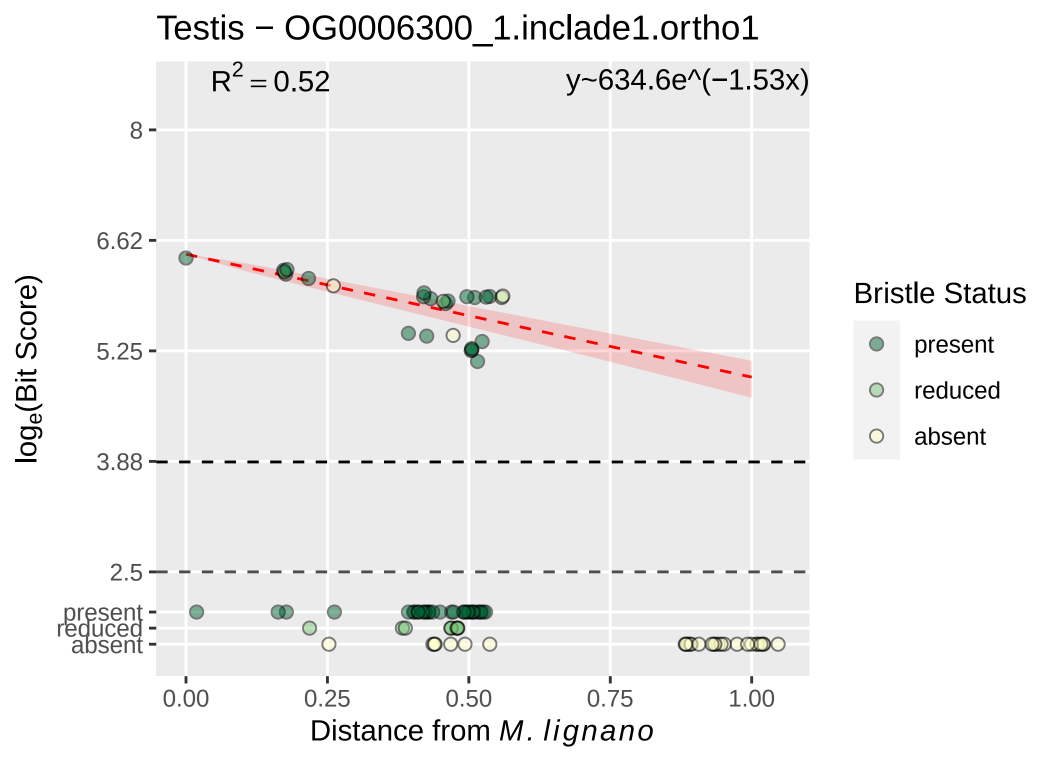
Testis − OG0006300_1.inclade1.ortho1
$R^2 = 0.52$
y~634.6e^(−1.53x)
8
6.62
5.25
3.88
2.5
present
reduced
absent
$\log_e$(Bit Score)
0.00
0.25
0.50
0.75
1.00
Distance from *M. lignano*
Bristle Status
present
reduced
absent

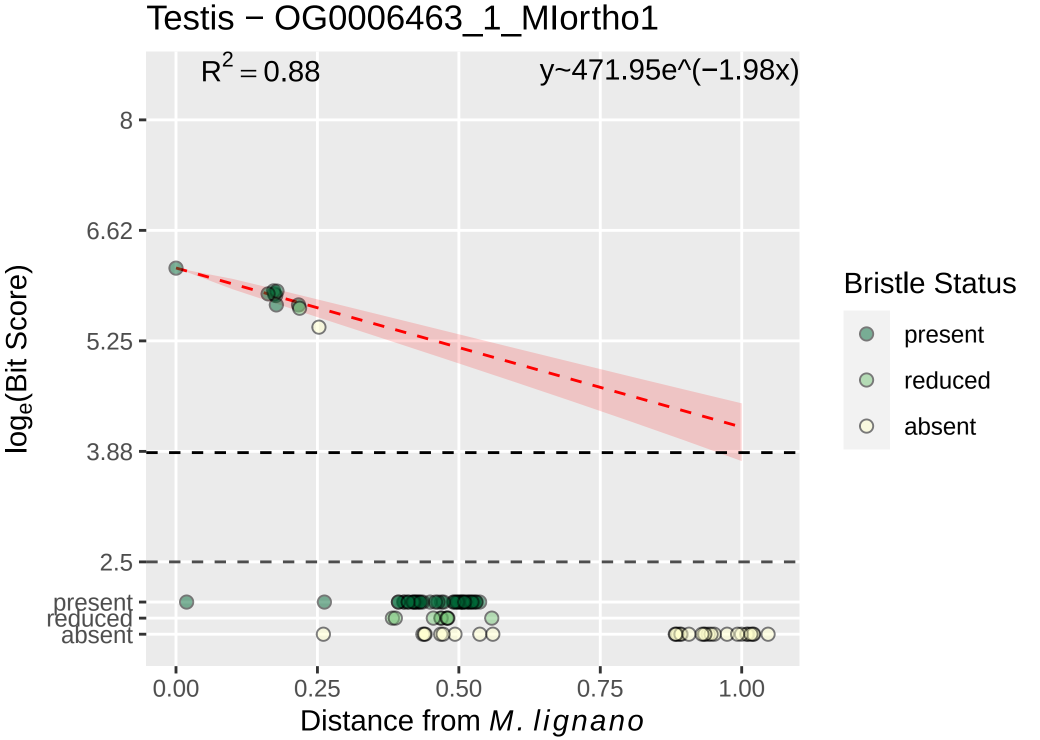

Testis − OG0006463_1_MIortho1
$R^2 = 0.88$
y~471.95e^(−1.98x)
8
6.62
5.25
3.88
2.5
present
reduced
absent
$\log_e$(Bit Score)
0.00
0.25
0.50
0.75
1.00
Distance from *M. lignano*
Bristle Status
present
reduced
absent

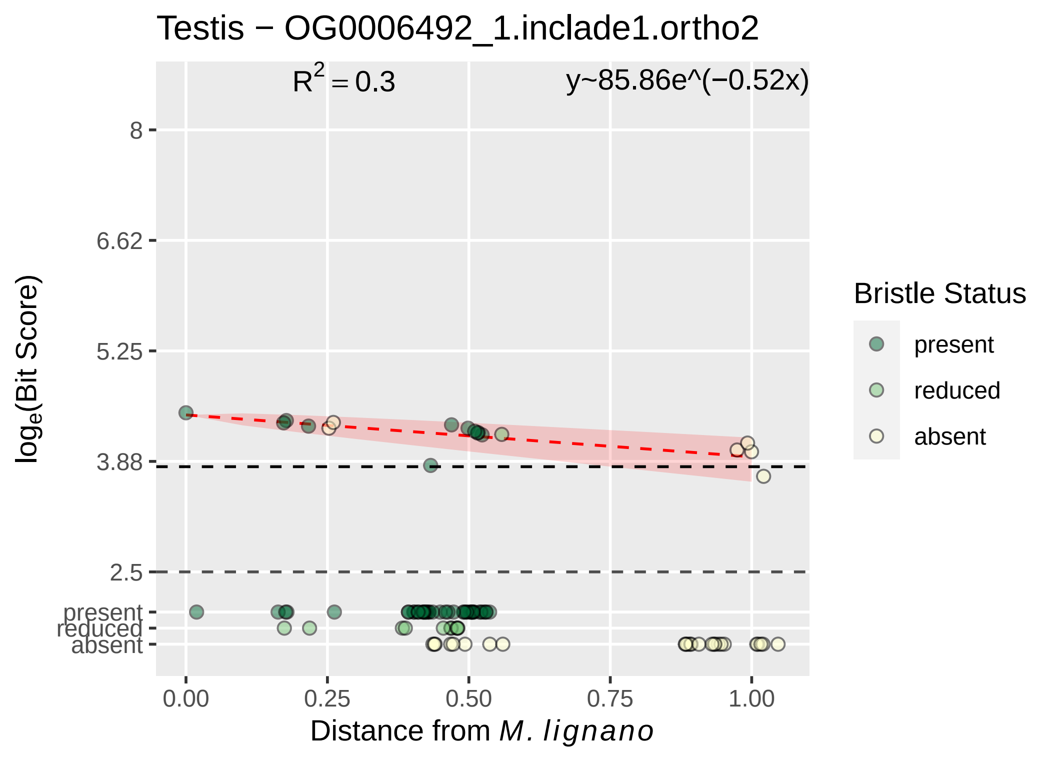
Testis − OG0006492_1.inclade1.ortho2
R2 = 0.3
y~85.86e^(−0.52x)
8
6.62
5.25
3.88
2.5
present
reduced
absent
loge(Bit Score)
0.00
0.25
0.50
0.75
1.00
Distance from M. lignano
Bristle Status
present
reduced
absent

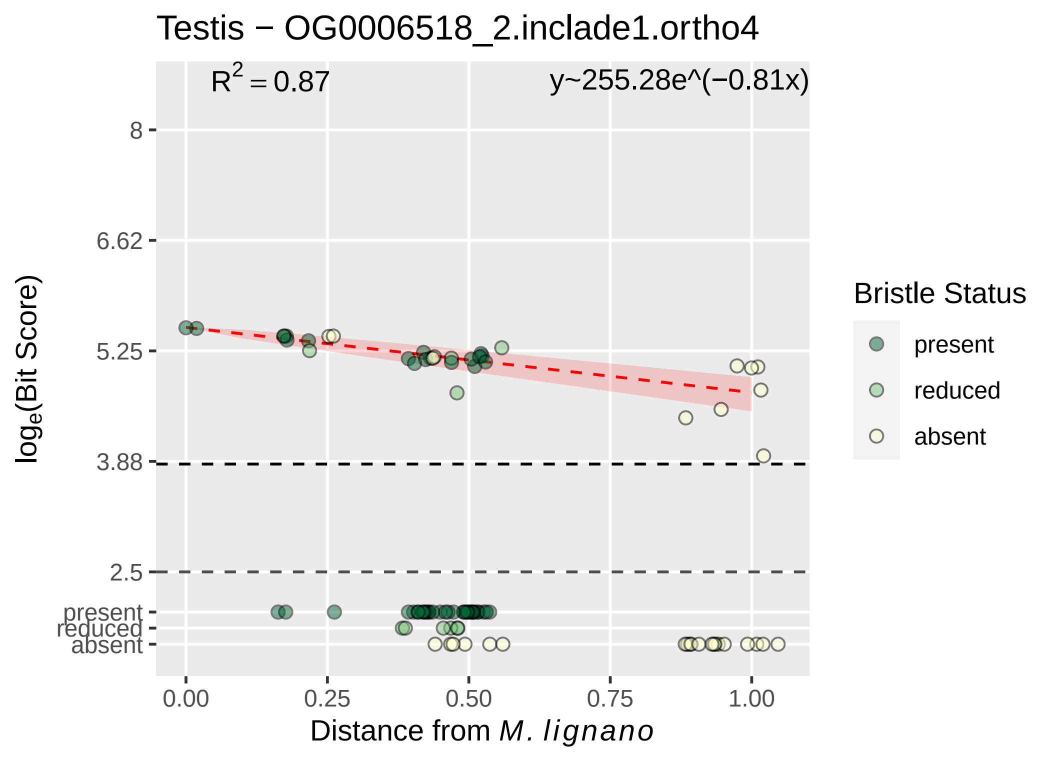

Testis − OG0006518_2.inclade1.ortho4
$R^2 = 0.87$
y~255.28e^(−0.81x)
8
6.62
5.25
3.88
2.5
present
reduced
absent
log$_e$(Bit Score)
0.00
0.25
0.50
0.75
1.00
Distance from *M. lignano*
Bristle Status
present
reduced
absent

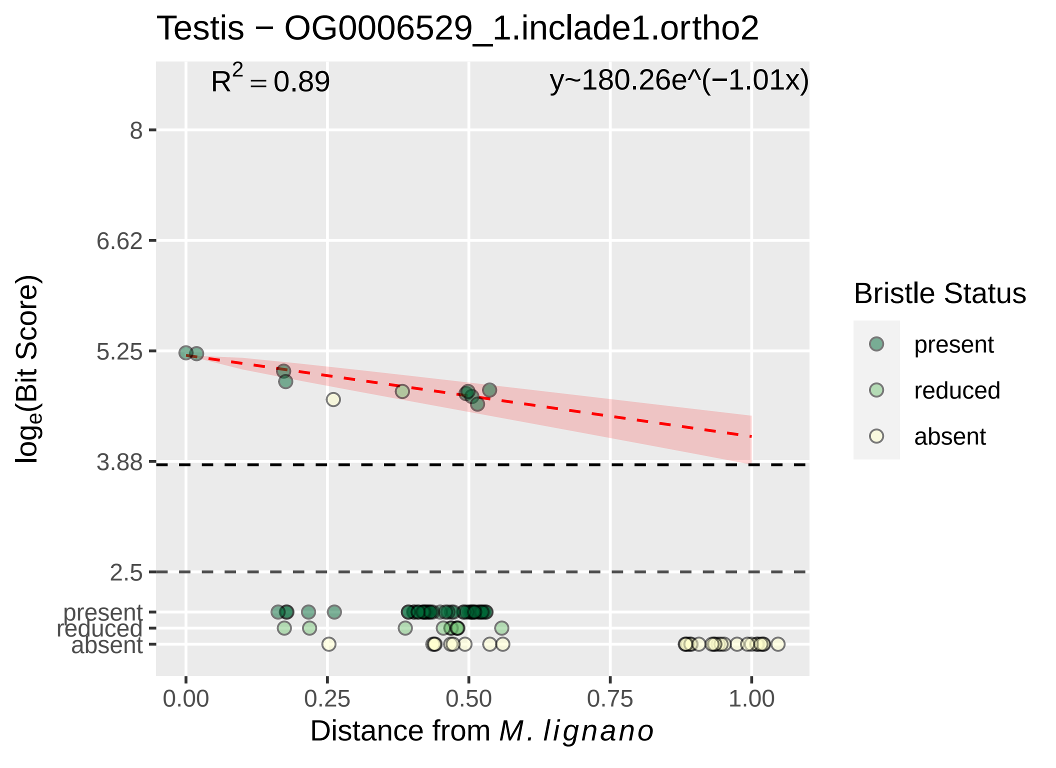

Testis − OG0006529_1.inclade1.ortho2
R2 = 0.89
y~180.26e^(−1.01x)
8
6.62
5.25
3.88
2.5
present
reduced
absent
loge(Bit Score)
0.00
0.25
0.50
0.75
1.00
Distance from M. lignano
Bristle Status
present
reduced
absent

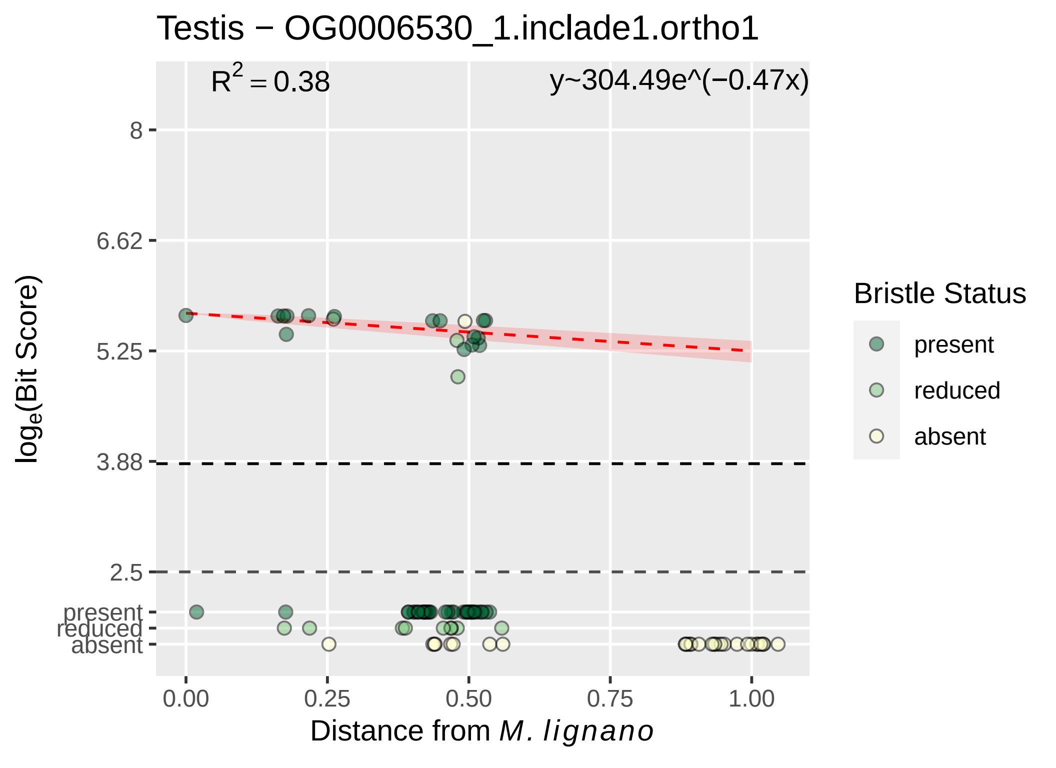

Testis − OG0006530_1.inclade1.ortho1
R² = 0.38
y~304.49e^(−0.47x)
8
6.62
5.25
3.88
2.5
present
reduced
absent
loge(Bit Score)
0.00
0.25
0.50
0.75
1.00
Distance from M. lignano
Bristle Status
present
reduced
absent

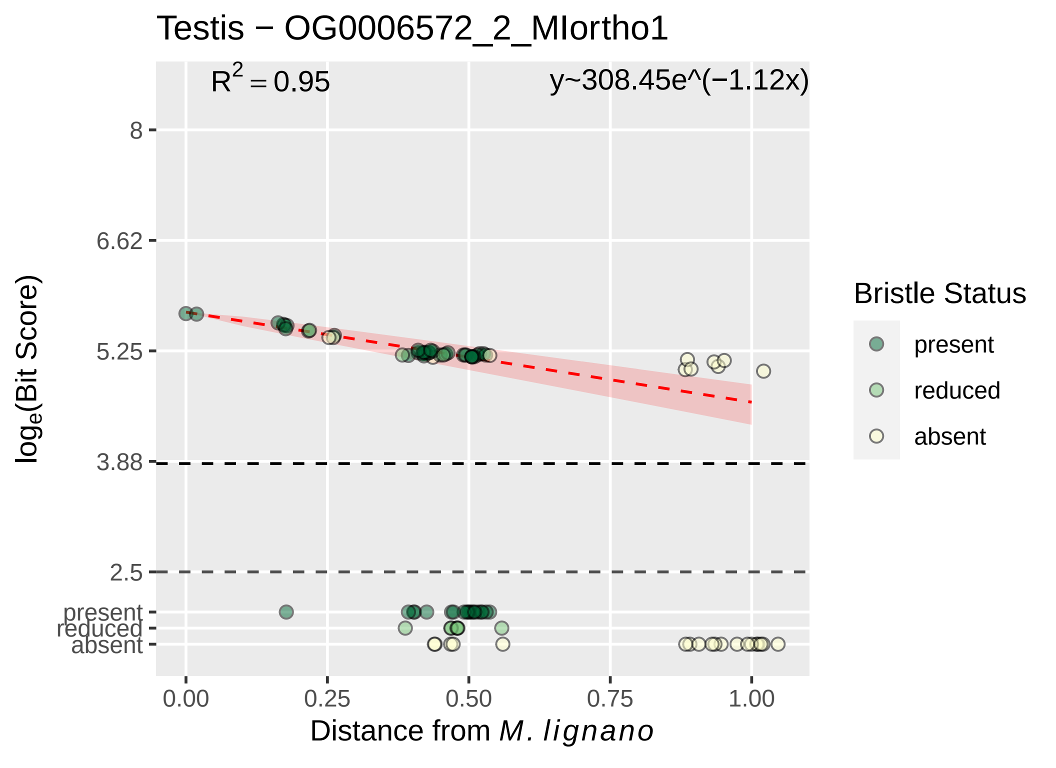
Testis − OG0006572_2_MIortho1
$R^2 = 0.95$
y~308.45e^(−1.12x)
8
6.62
5.25
3.88
2.5
present
reduced
absent
$\log_e$(Bit Score)
0.00
0.25
0.50
0.75
1.00
Distance from *M. lignano*
Bristle Status
present
reduced
absent

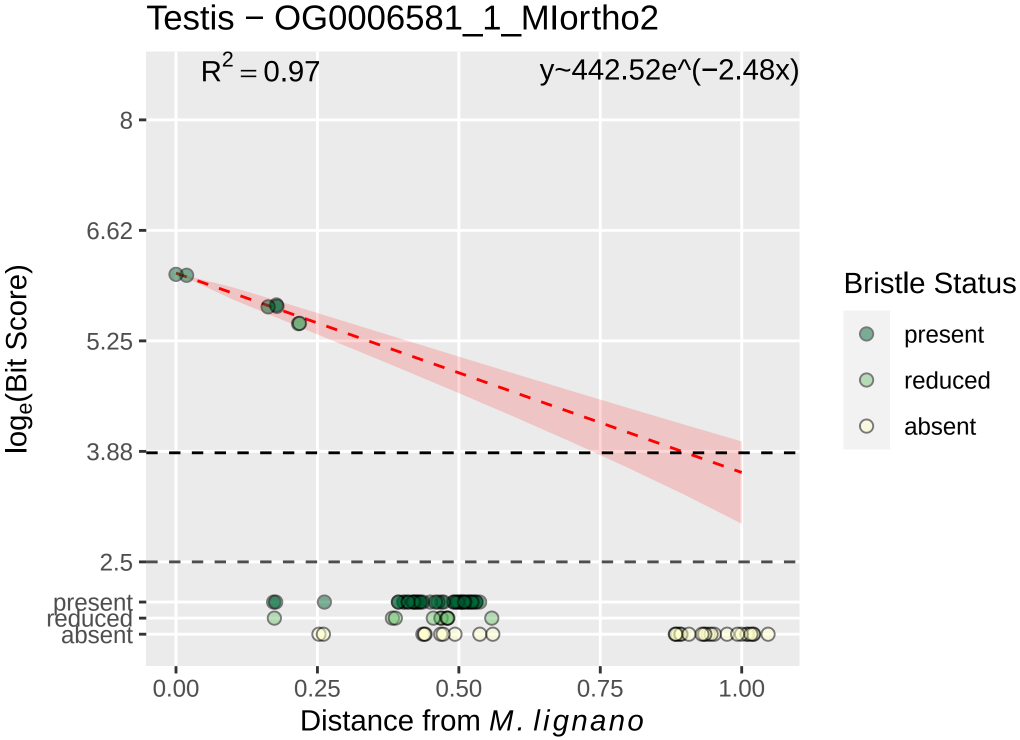

Testis − OG0006581_1_MIortho2
R^2 = 0.97
y~442.52e^(−2.48x)
8
6.62
5.25
3.88
2.5
present
reduced
absent
log_e(Bit Score)
0.00
0.25
0.50
0.75
1.00
Distance from M. lignano
Bristle Status
present
reduced
absent

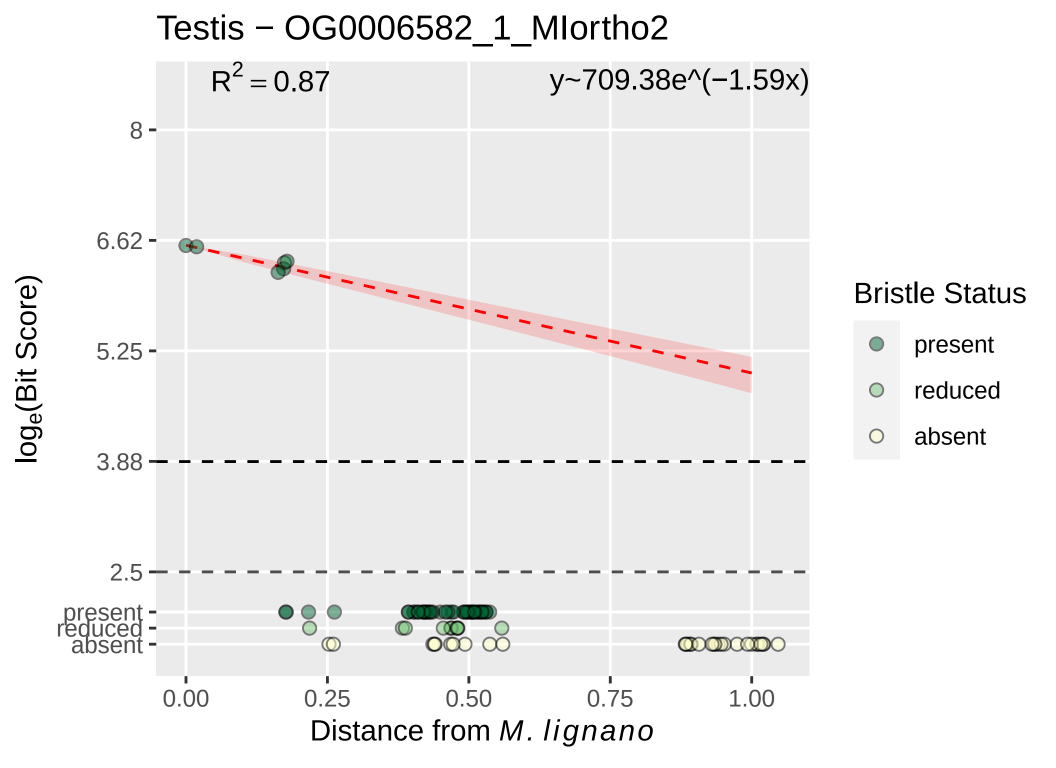

Testis − OG0006582_1_MIortho2
$R^2 = 0.87$
y~709.38e^(−1.59x)
8
6.62
5.25
3.88
2.5
present
reduced
absent
$\log_e$(Bit Score)
0.00
0.25
0.50
0.75
1.00
Distance from *M. lignano*
Bristle Status
present
reduced
absent

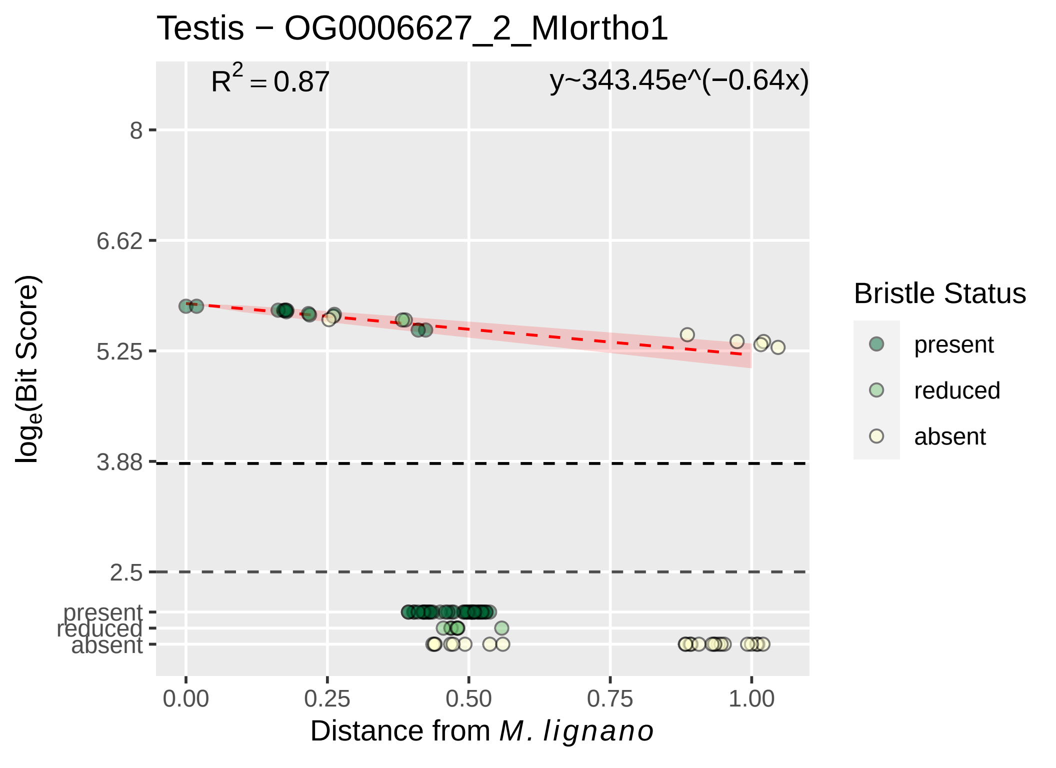

Testis − OG0006627_2_MIortho1
R2 = 0.87
y~343.45e^(−0.64x)
loge(Bit Score)
8
6.62
5.25
3.88
2.5
present
reduced
absent
0.00
0.25
0.50
0.75
1.00
Distance from M. lignano
Bristle Status
present
reduced
absent

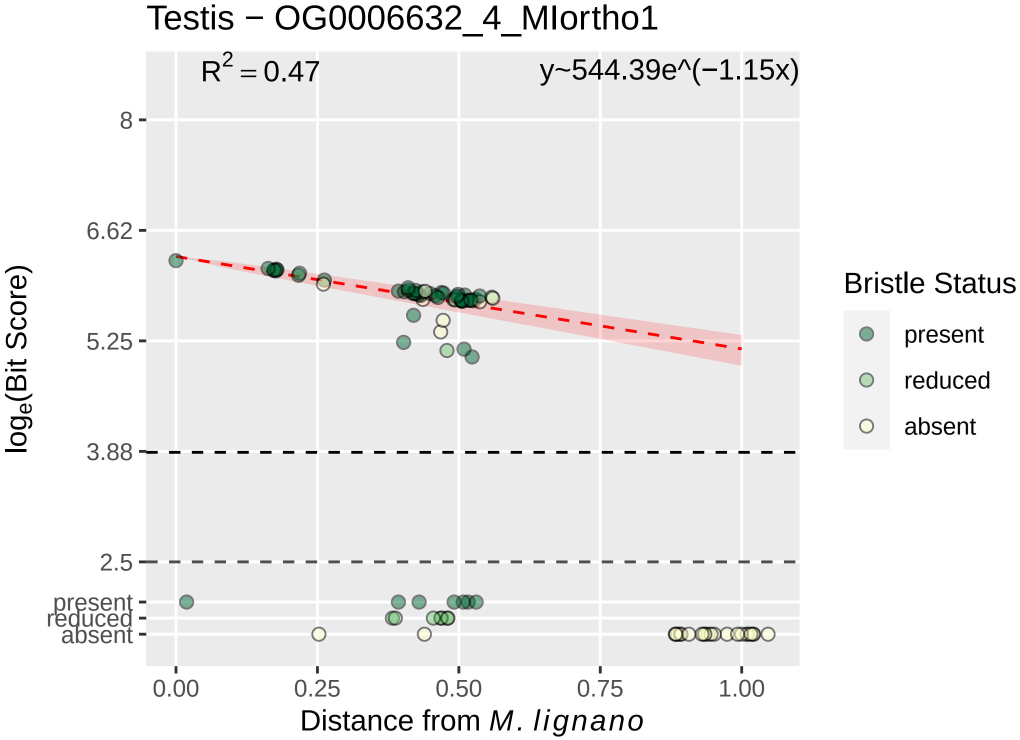

Testis − OG0006632_4_MIortho1
$R^2 = 0.47$
y~544.39e^(−1.15x)
8
6.62
5.25
3.88
2.5
present
reduced
absent
0.00
0.25
0.50
0.75
1.00
Distance from *M. lignano*
$\log_e$(Bit Score)
Bristle Status
present
reduced
absent

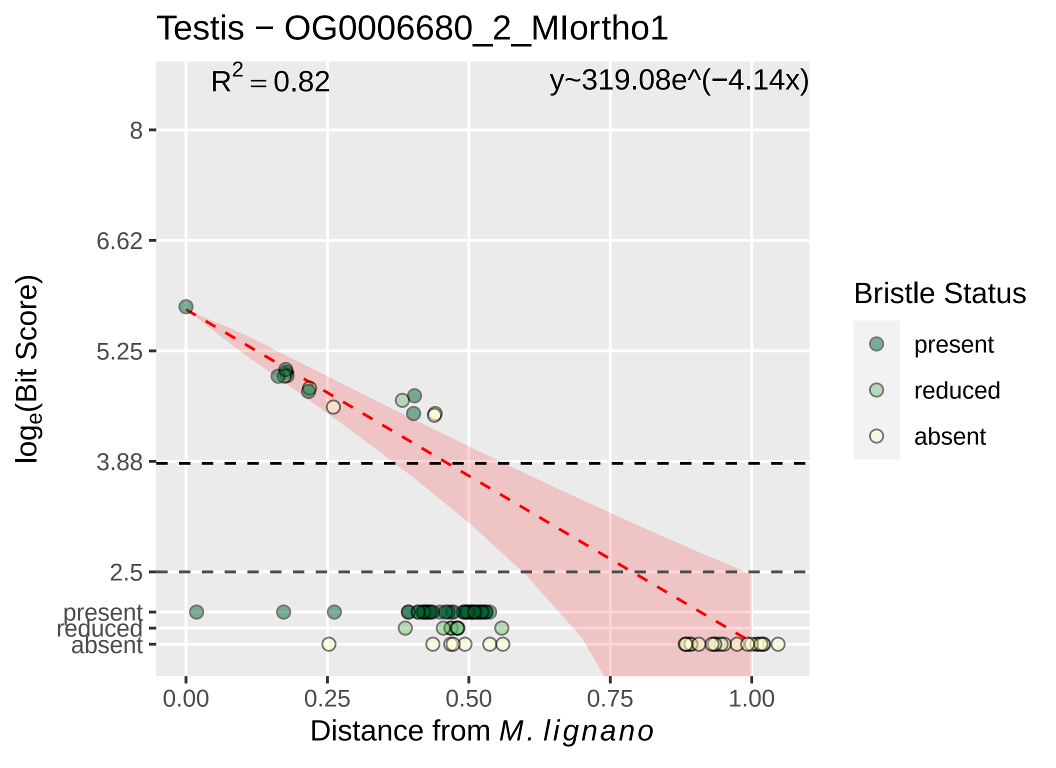

Testis − OG0006680_2_MIortho1
$R^2 = 0.82$
y~319.08e^(−4.14x)
8
6.62
5.25
3.88
2.5
present
reduced
absent
$\log_e$(Bit Score)
0.00
0.25
0.50
0.75
1.00
Distance from *M. lignano*
Bristle Status
present
reduced
absent

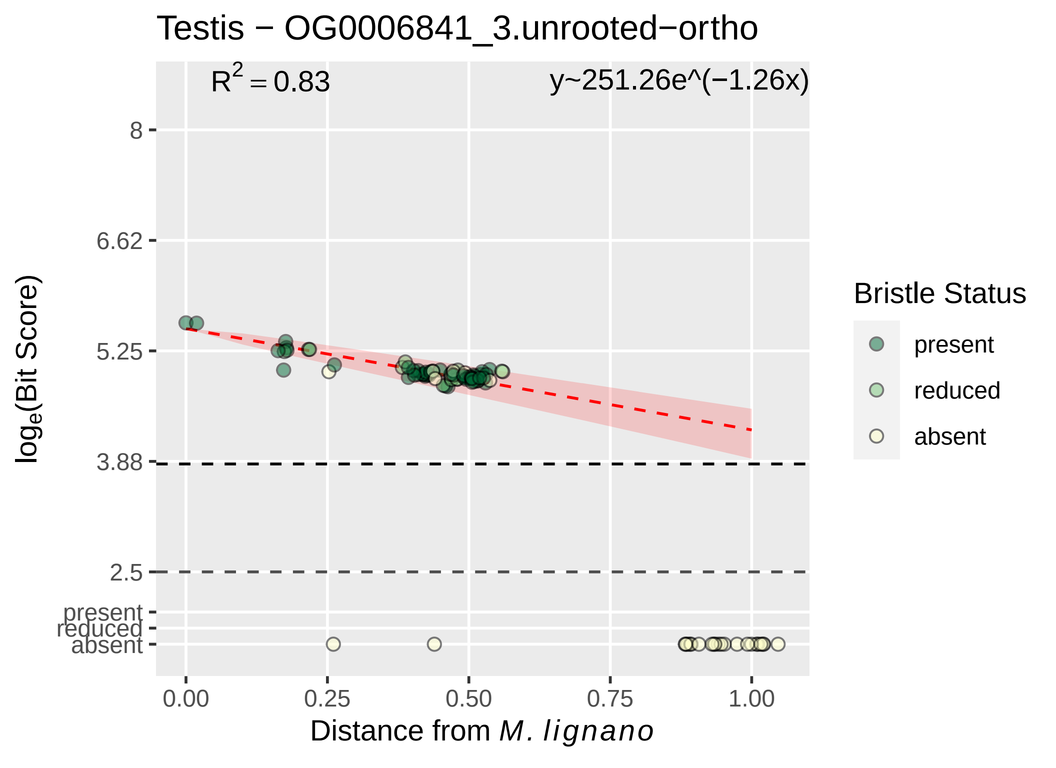

Testis − OG0006841_3.unrooted−ortho
$R^2 = 0.83$
y~251.26e^(−1.26x)
8
6.62
5.25
3.88
2.5
present
reduced
absent
$log_e$(Bit Score)
0.00
0.25
0.50
0.75
1.00
Distance from *M. lignano*
Bristle Status
present
reduced
absent

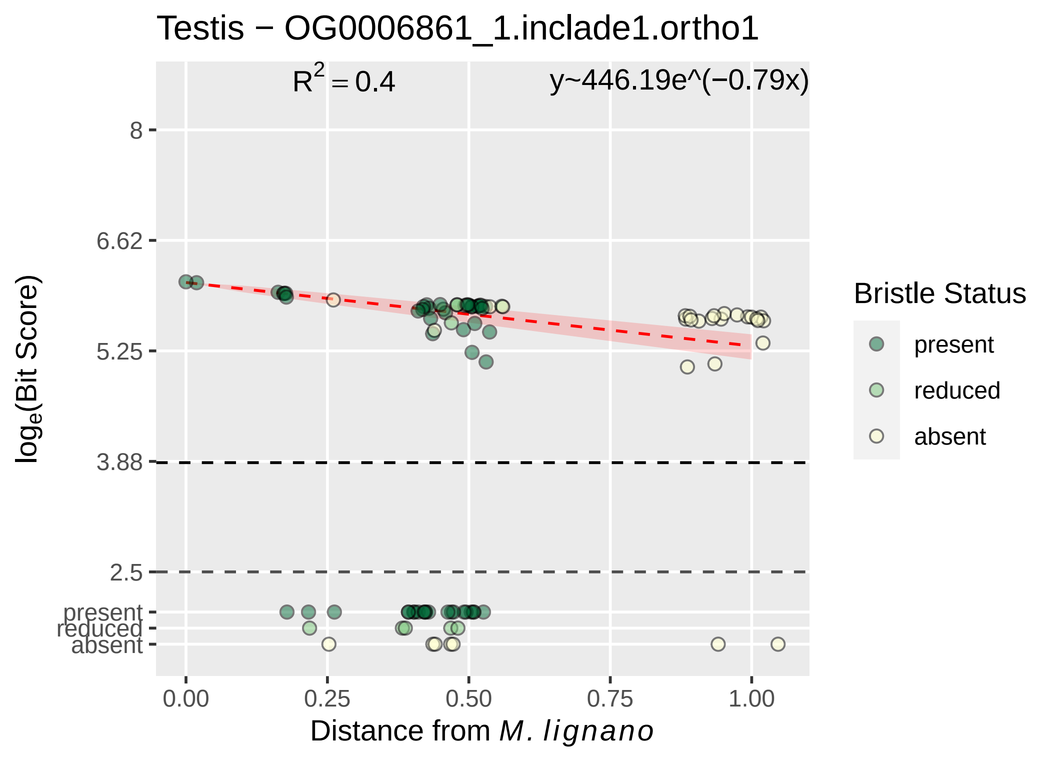

Testis − OG0006861_1.inclade1.ortho1
R^2 = 0.4
y~446.19e^(−0.79x)
8
6.62
5.25
3.88
2.5
present
reduced
absent
0.00
0.25
0.50
0.75
1.00
Distance from M. lignano
loge(Bit Score)
Bristle Status
present
reduced
absent

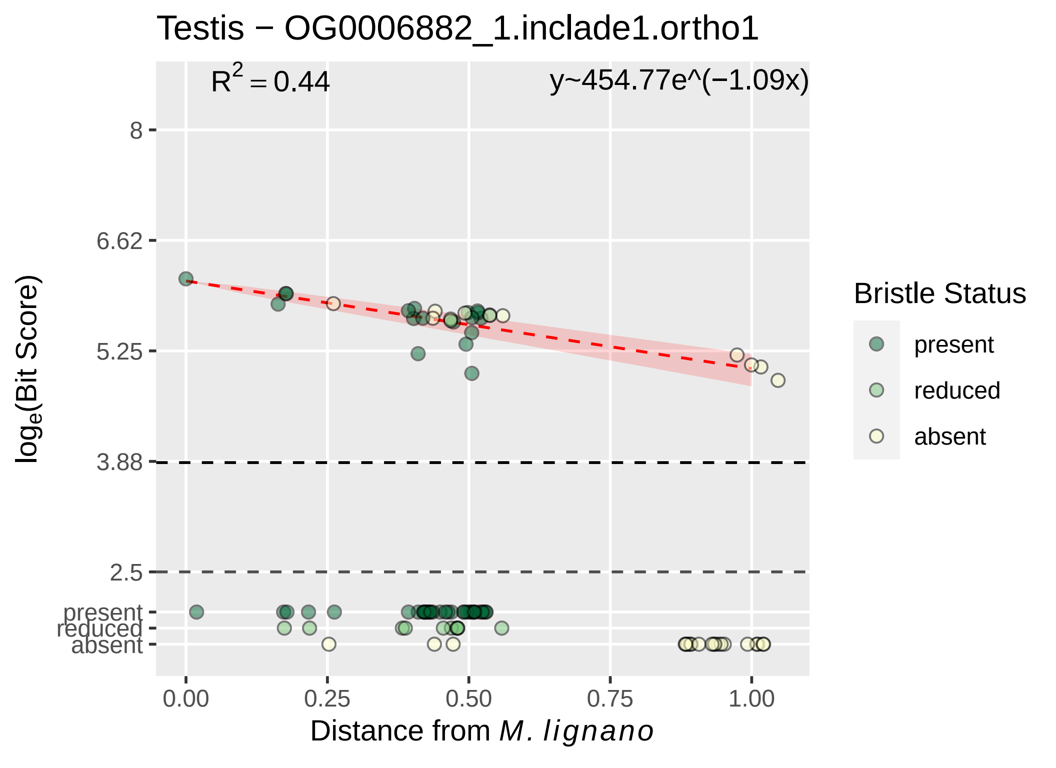

Testis − OG0006882_1.inclade1.ortho1
$R^2 = 0.44$
y~454.77e^(−1.09x)
8
6.62
5.25
3.88
2.5
present
reduced
absent
$\log_e$(Bit Score)
0.00
0.25
0.50
0.75
1.00
Distance from *M. lignano*
Bristle Status
present
reduced
absent

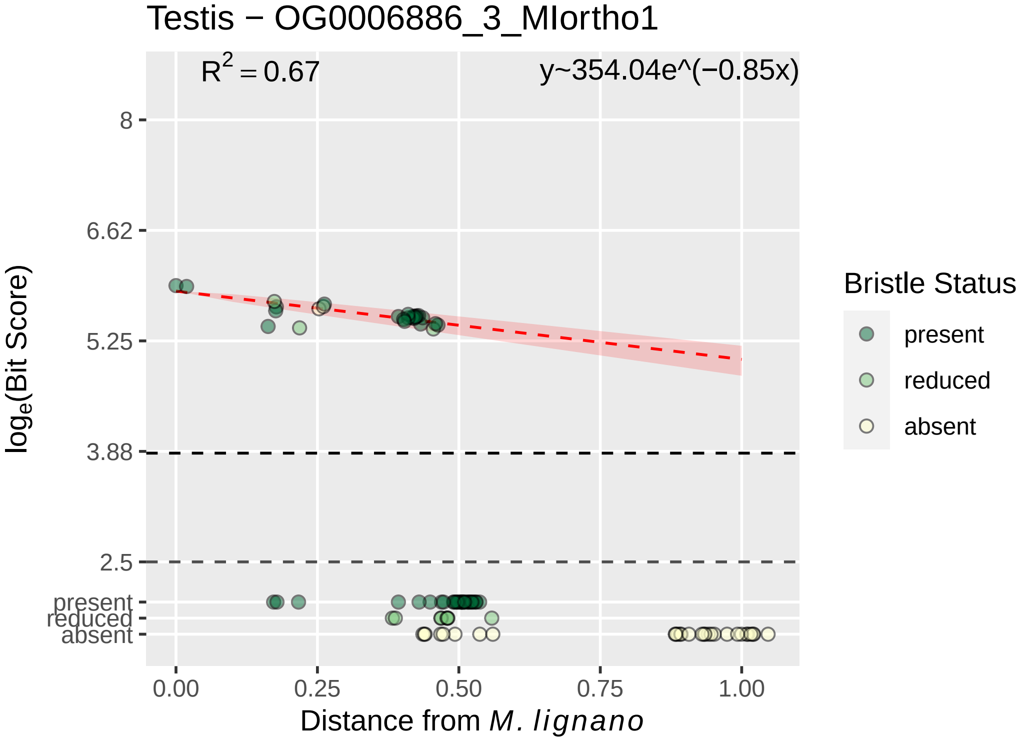

Testis − OG0006886_3_MIortho1
R^2 = 0.67
y~354.04e^(−0.85x)
8
6.62
5.25
3.88
2.5
present
reduced
absent
log_e(Bit Score)
0.00
0.25
0.50
0.75
1.00
Distance from M. lignano
Bristle Status
present
reduced
absent

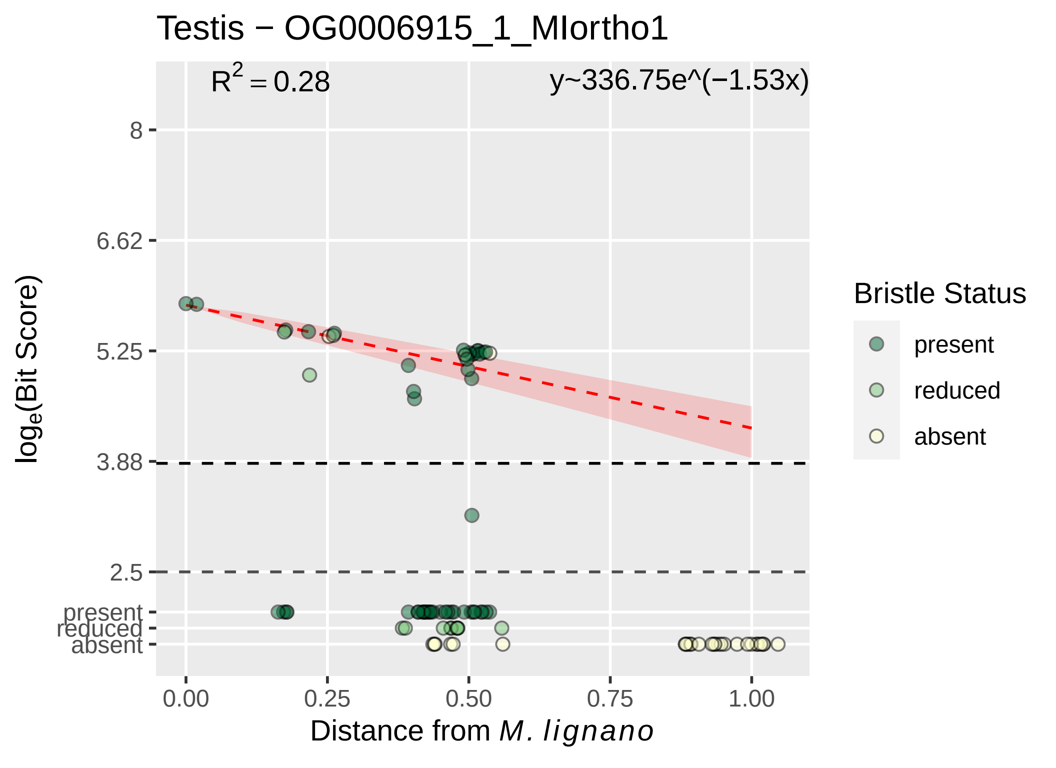
Testis − OG0006915_1_MIortho1
$R^2 = 0.28$
y~336.75e^(−1.53x)
8
6.62
5.25
3.88
2.5
present
reduced
absent
$log_e$(Bit Score)
0.00
0.25
0.50
0.75
1.00
Distance from *M. lignano*
Bristle Status
present
reduced
absent

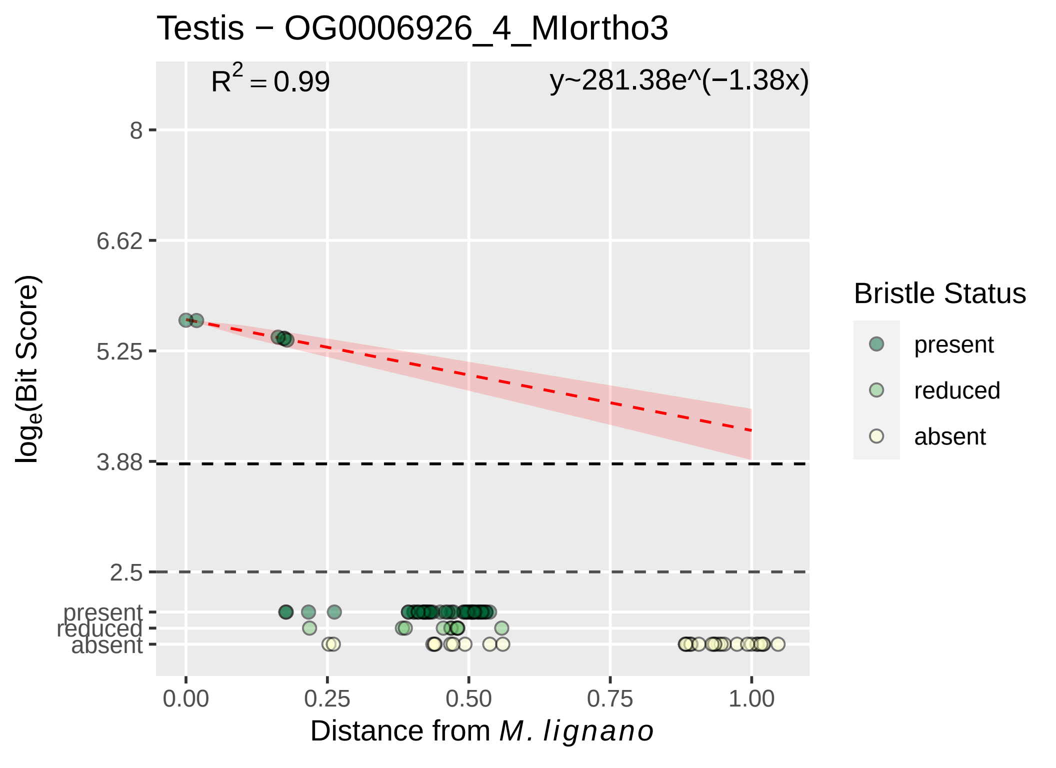
Testis − OG0006926_4_MIortho3
R2 = 0.99
y~281.38e^(−1.38x)
loge(Bit Score)
8
6.62
5.25
3.88
2.5
present
reduced
absent
0.00
0.25
0.50
0.75
1.00
Distance from M. lignano
Bristle Status
present
reduced
absent

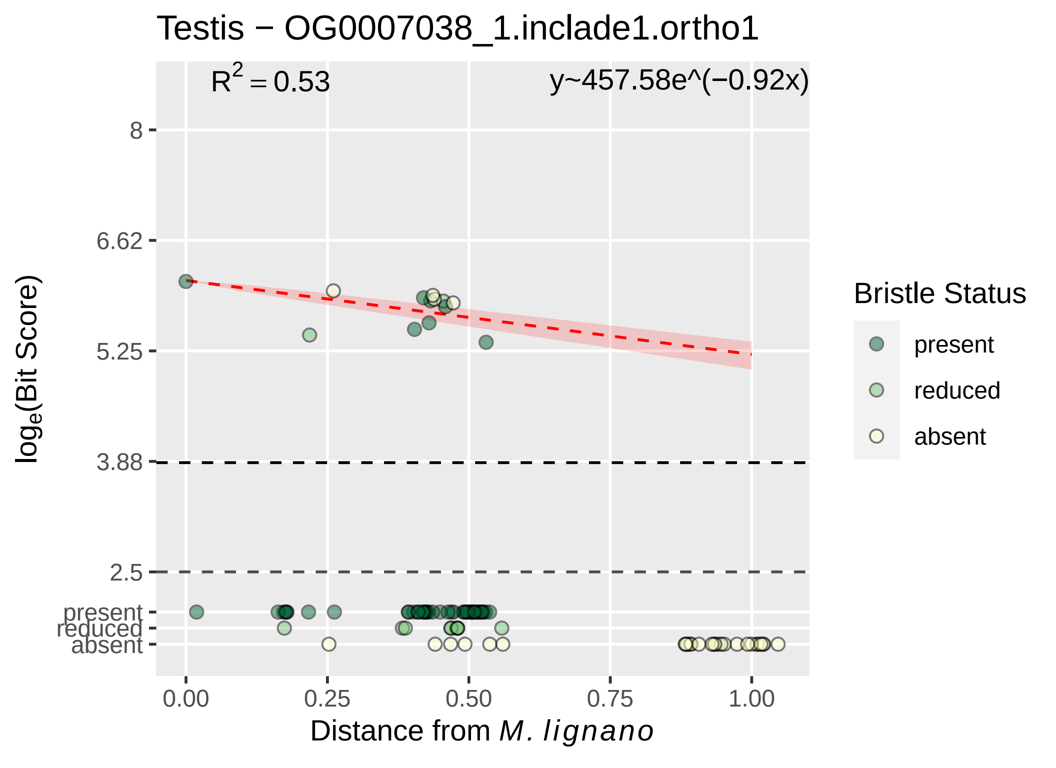

Testis − OG0007038_1.inclade1.ortho1
$R^2 = 0.53$
y~457.58e^(−0.92x)
8
6.62
5.25
3.88
2.5
present
reduced
absent
$\log_e$(Bit Score)
0.00
0.25
0.50
0.75
1.00
Distance from *M. lignano*
Bristle Status
present
reduced
absent

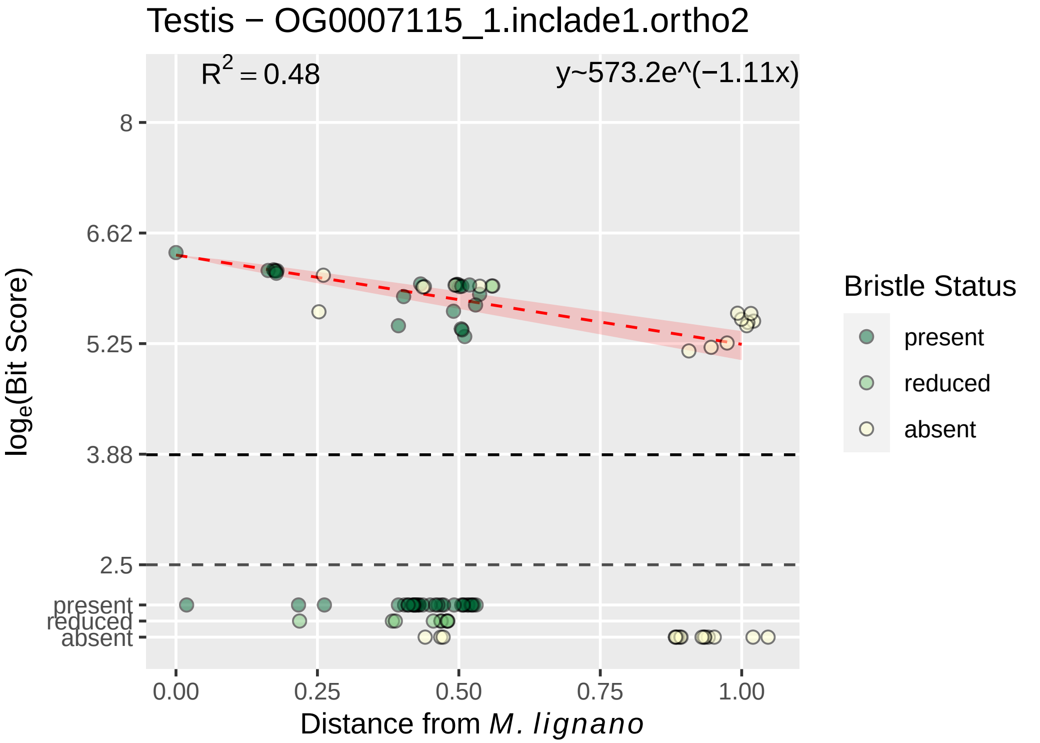

Testis − OG0007115_1.inclade1.ortho2
$R^2 = 0.48$
y~573.2e^(−1.11x)
8
6.62
5.25
3.88
2.5
present
reduced
absent
$\log_e$(Bit Score)
0.00
0.25
0.50
0.75
1.00
Distance from *M. lignano*
Bristle Status
present
reduced
absent

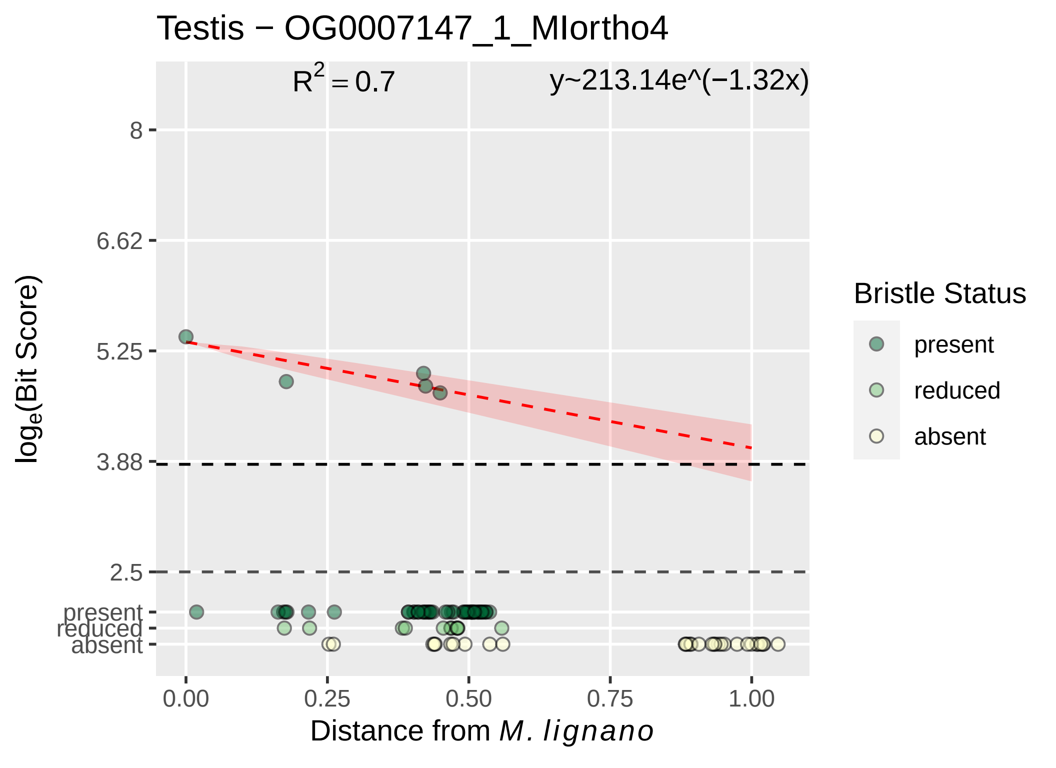

Testis − OG0007147_1_MIortho4
$R^2 = 0.7$
y~213.14e^(−1.32x)
8
6.62
5.25
3.88
2.5
present
reduced
absent
$\log_e$(Bit Score)
0.00
0.25
0.50
0.75
1.00
Distance from *M. lignano*
Bristle Status
present
reduced
absent

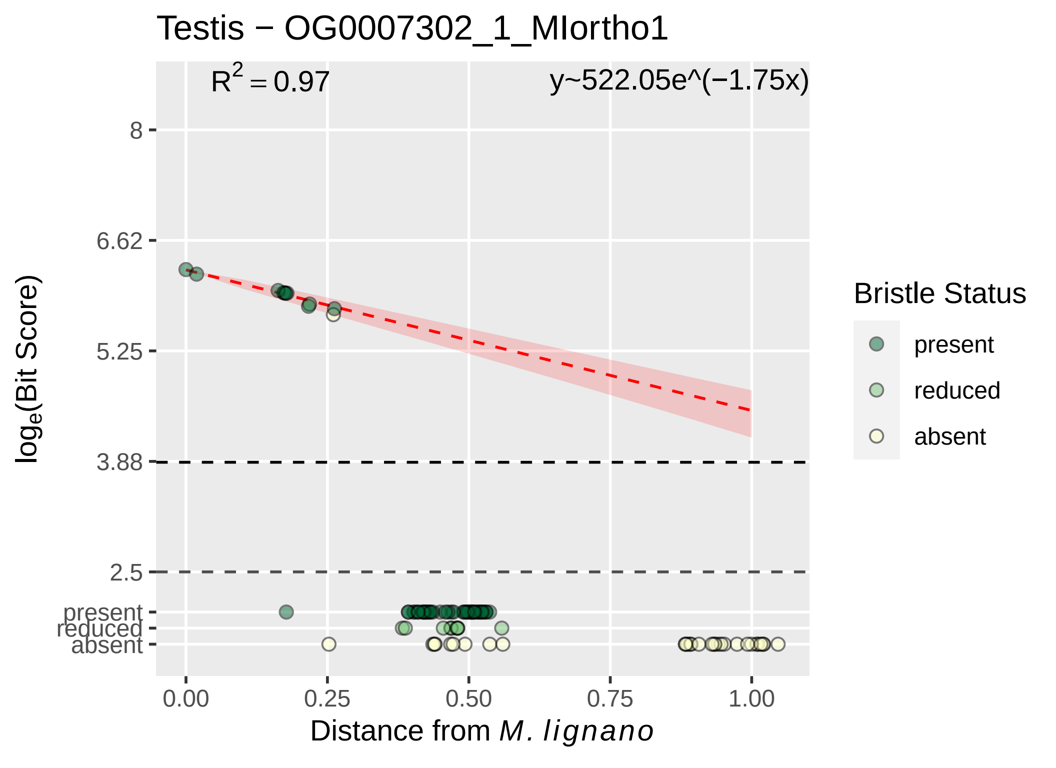

Testis − OG0007302_1_MIortho1
R^2 = 0.97
y~522.05e^(−1.75x)
8
6.62
5.25
3.88
2.5
present
reduced
absent
0.00
0.25
0.50
0.75
1.00
log_e(Bit Score)
Distance from M. lignano
Bristle Status
present
reduced
absent

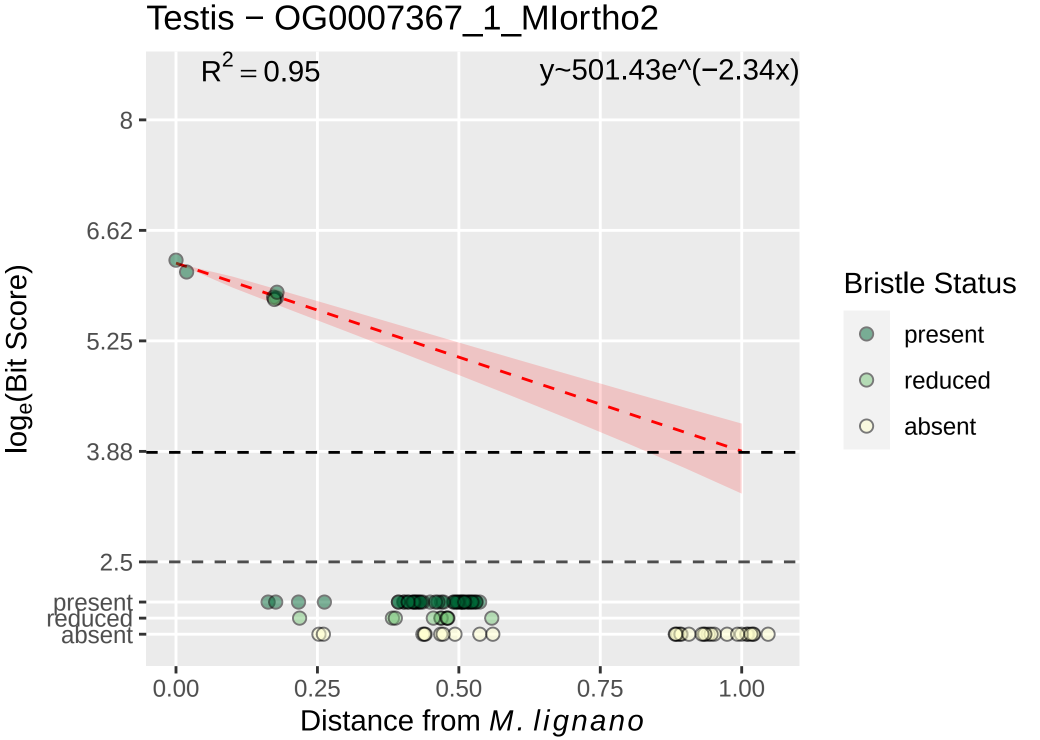

Testis − OG0007367_1_MIortho2
$R^2 = 0.95$
y~501.43e^(−2.34x)
8
6.62
5.25
3.88
2.5
present
reduced
absent
loge(Bit Score)
0.00
0.25
0.50
0.75
1.00
Distance from M. lignano
Bristle Status
present
reduced
absent

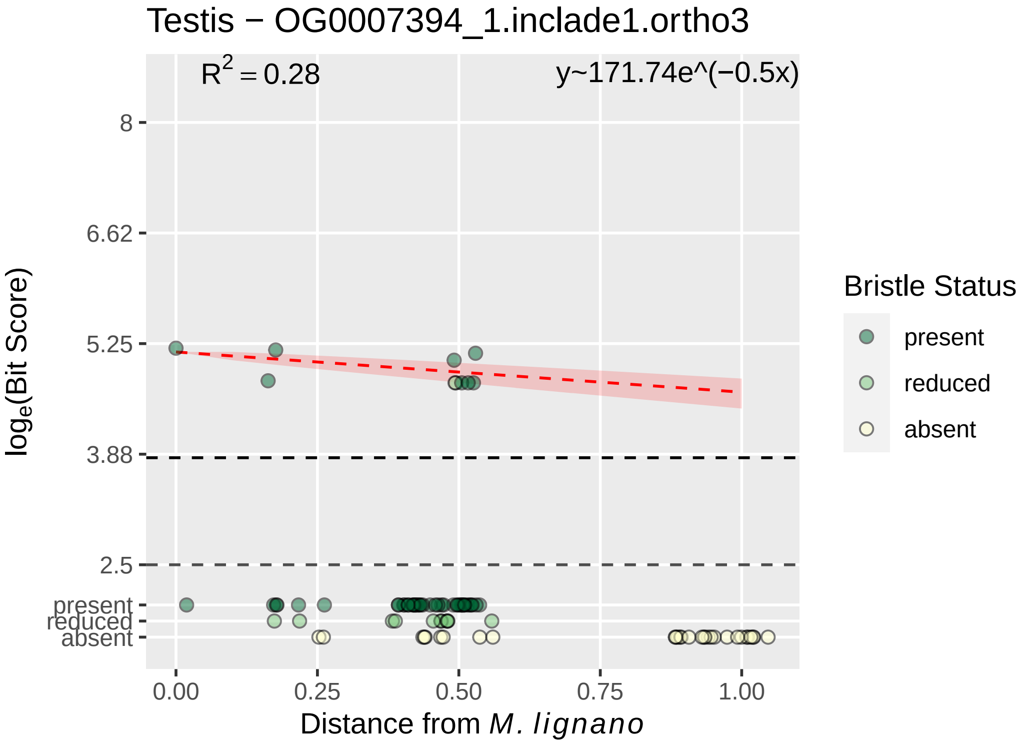

Testis − OG0007394_1.inclade1.ortho3
$R^2 = 0.28$
y~171.74e^(−0.5x)
8
6.62
5.25
3.88
2.5
present
reduced
absent
$log_e$(Bit Score)
0.00
0.25
0.50
0.75
1.00
Distance from *M. lignano*
Bristle Status
present
reduced
absent

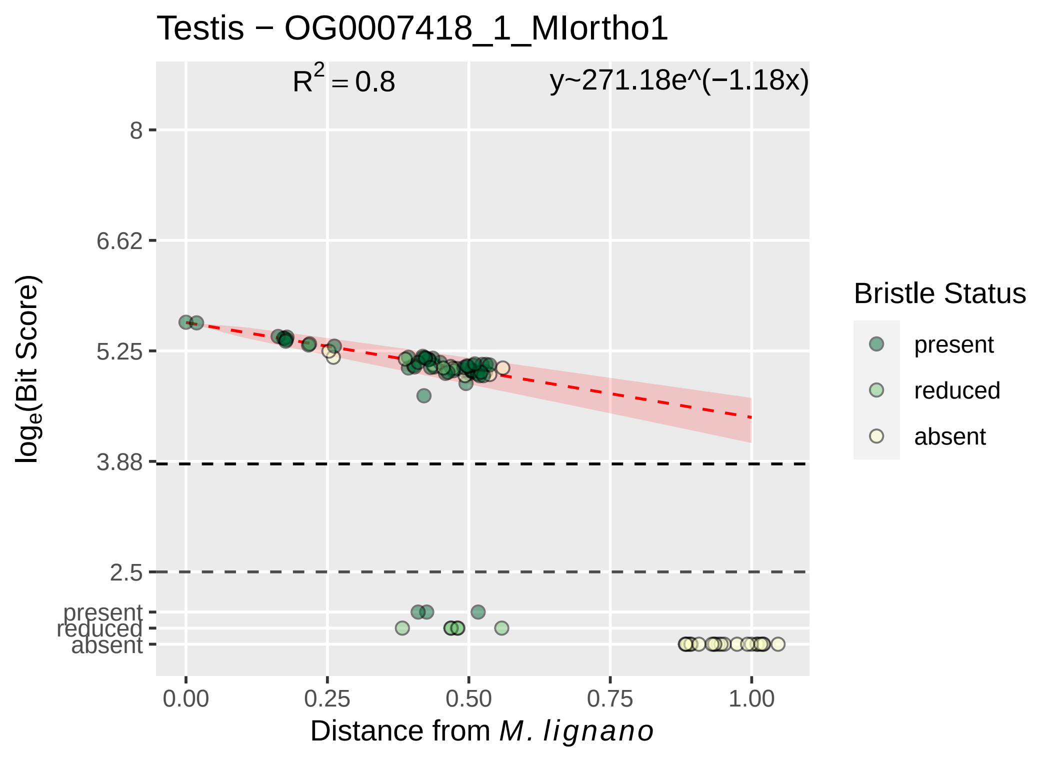

Testis − OG0007418_1_MIortho1
$R^2 = 0.8$
y~271.18e^(−1.18x)
8
6.62
5.25
3.88
2.5
present
reduced
absent
$\log_e$(Bit Score)
0.00
0.25
0.50
0.75
1.00
Distance from *M. lignano*
Bristle Status
present
reduced
absent

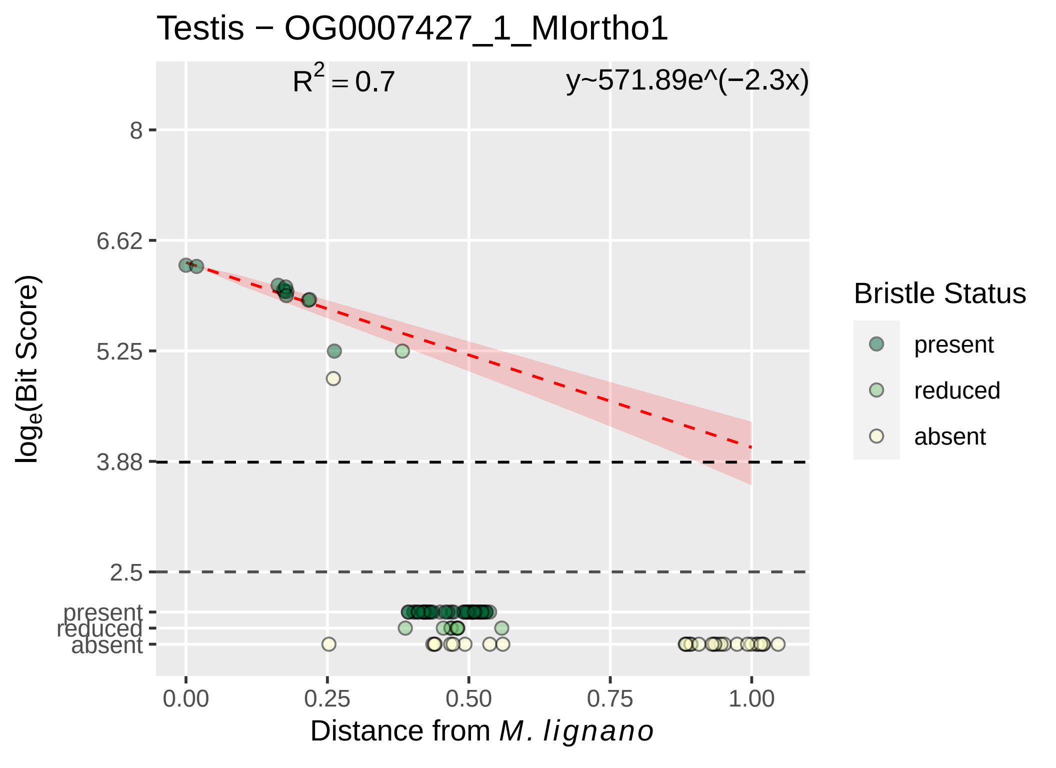

Testis − OG0007427_1_MIortho1
$R^2 = 0.7$
y~571.89e^(−2.3x)
8
6.62
5.25
3.88
2.5
present
reduced
absent
$\log_e$(Bit Score)
0.00
0.25
0.50
0.75
1.00
Distance from *M. lignano*
Bristle Status
present
reduced
absent

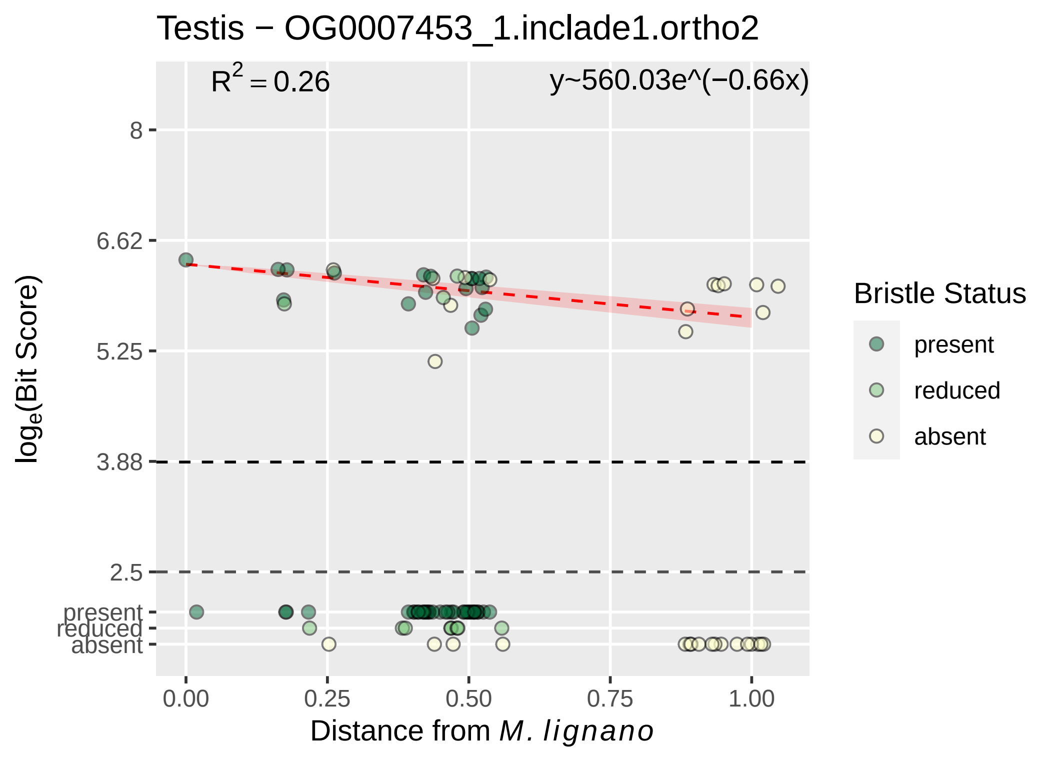

Testis − OG0007453_1.inclade1.ortho2
R^2 = 0.26
y~560.03e^(−0.66x)
8
6.62
5.25
3.88
2.5
present
reduced
absent
log_e(Bit Score)
0.00
0.25
0.50
0.75
1.00
Distance from M. lignano
Bristle Status
present
reduced
absent

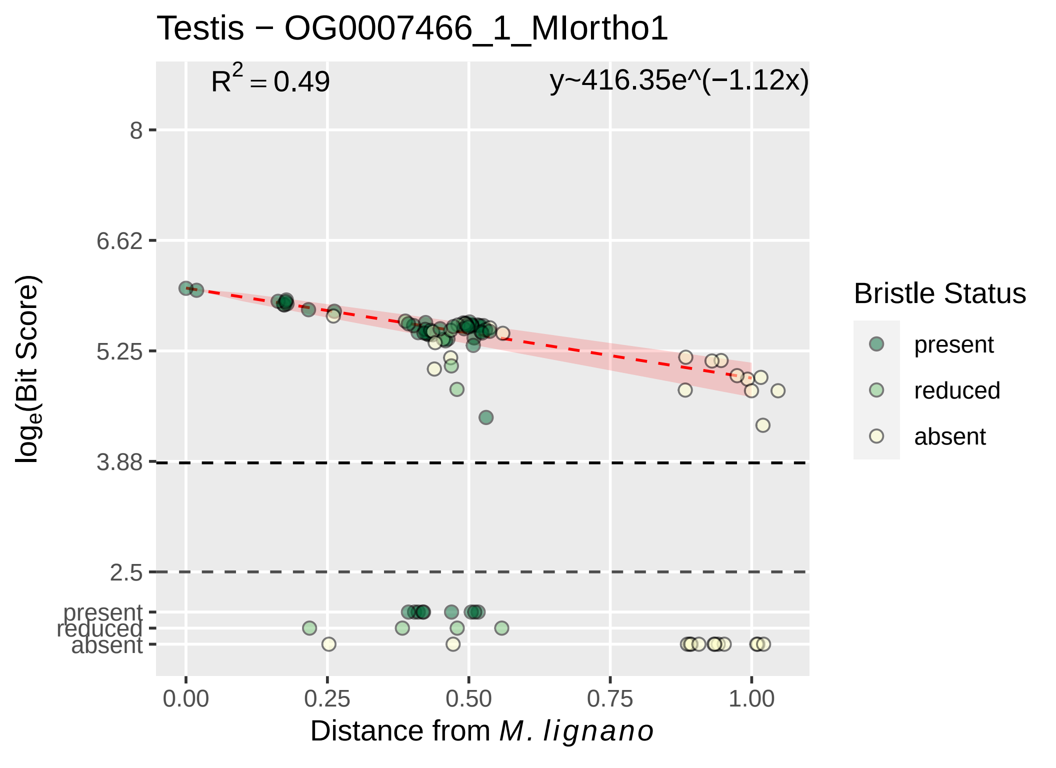
Testis − OG0007466_1_MIortho1
R^2 = 0.49
y~416.35e^(−1.12x)
8
6.62
5.25
3.88
2.5
present
reduced
absent
log_e(Bit Score)
0.00
0.25
0.50
0.75
1.00
Distance from M. lignano
Bristle Status
present
reduced
absent

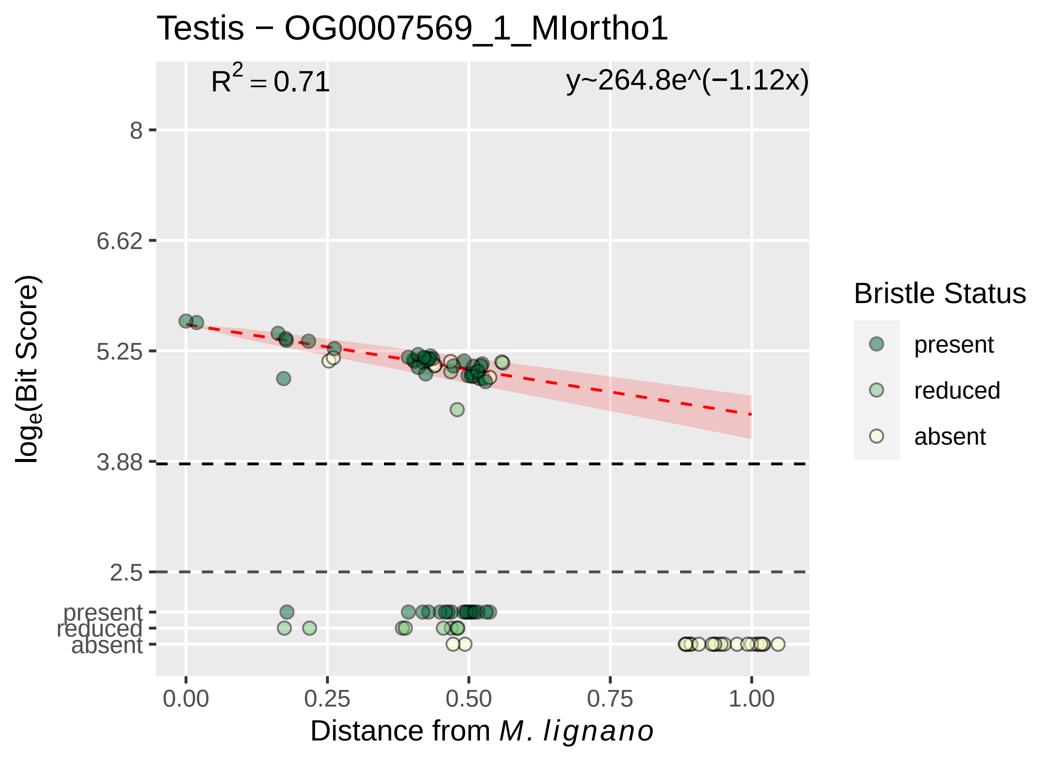

Testis − OG0007569_1_MIortho1
R² = 0.71
y~264.8e^(−1.12x)
8
6.62
5.25
3.88
2.5
present
reduced
absent
log_e(Bit Score)
0.00
0.25
0.50
0.75
1.00
Distance from M. lignano
Bristle Status
present
reduced
absent

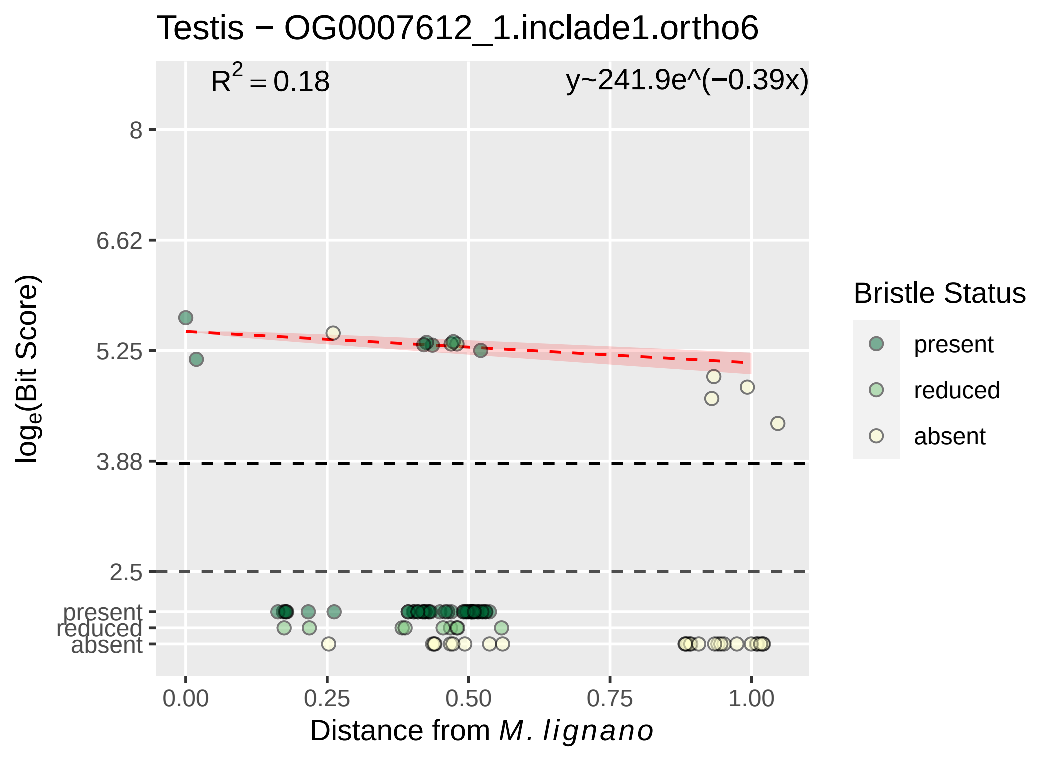

Testis − OG0007612_1.inclade1.ortho6
R^2 = 0.18
y~241.9e^(−0.39x)
8
6.62
5.25
3.88
2.5
present
reduced
absent
log_e(Bit Score)
0.00
0.25
0.50
0.75
1.00
Distance from M. lignano
Bristle Status
present
reduced
absent

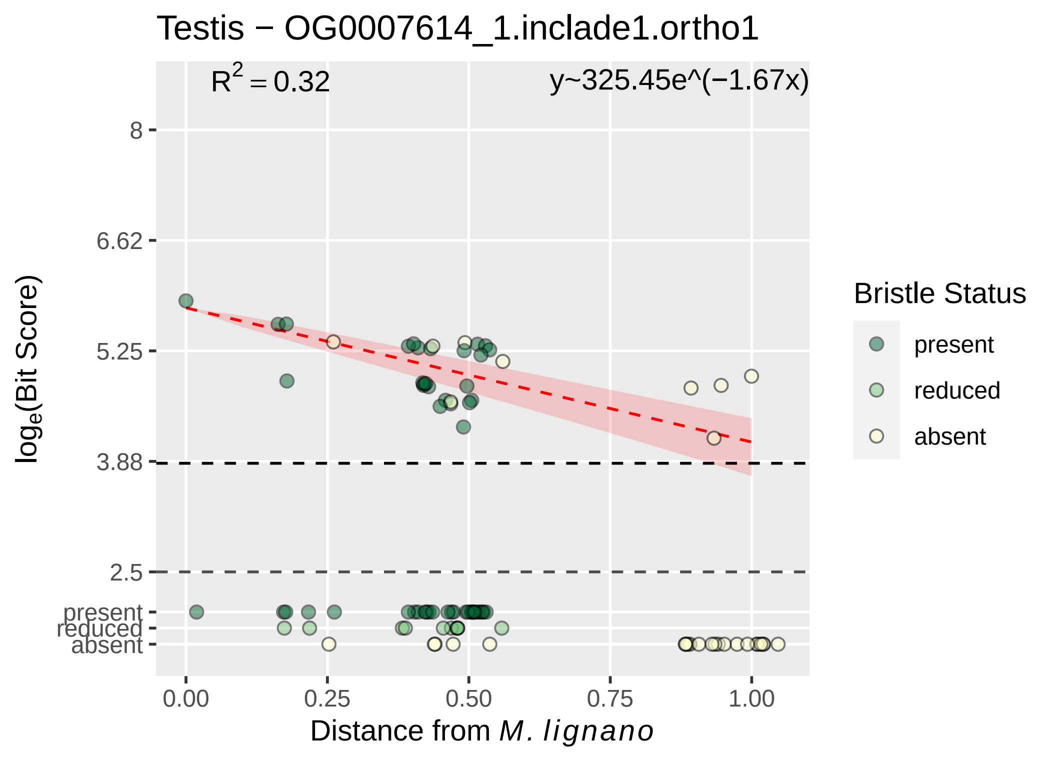

Testis − OG0007614_1.inclade1.ortho1
$R^2 = 0.32$
y~325.45e^(−1.67x)
8
6.62
5.25
3.88
2.5
present
reduced
absent
0.00
0.25
0.50
0.75
1.00
Distance from M. lignano
log_e(Bit Score)
Bristle Status
present
reduced
absent

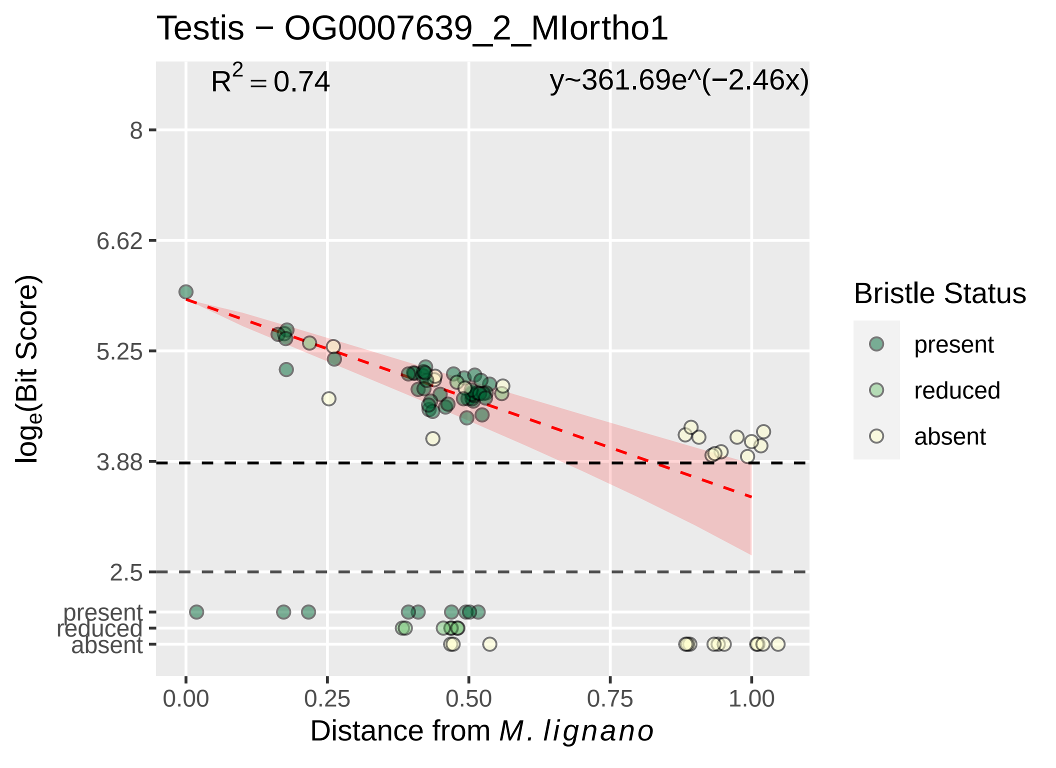

Testis − OG0007639_2_MIortho1
$R^2 = 0.74$
y~361.69e^(−2.46x)
8
6.62
5.25
3.88
2.5
present
reduced
absent
$\log_e$(Bit Score)
0.00
0.25
0.50
0.75
1.00
Distance from *M. lignano*
Bristle Status
present
reduced
absent

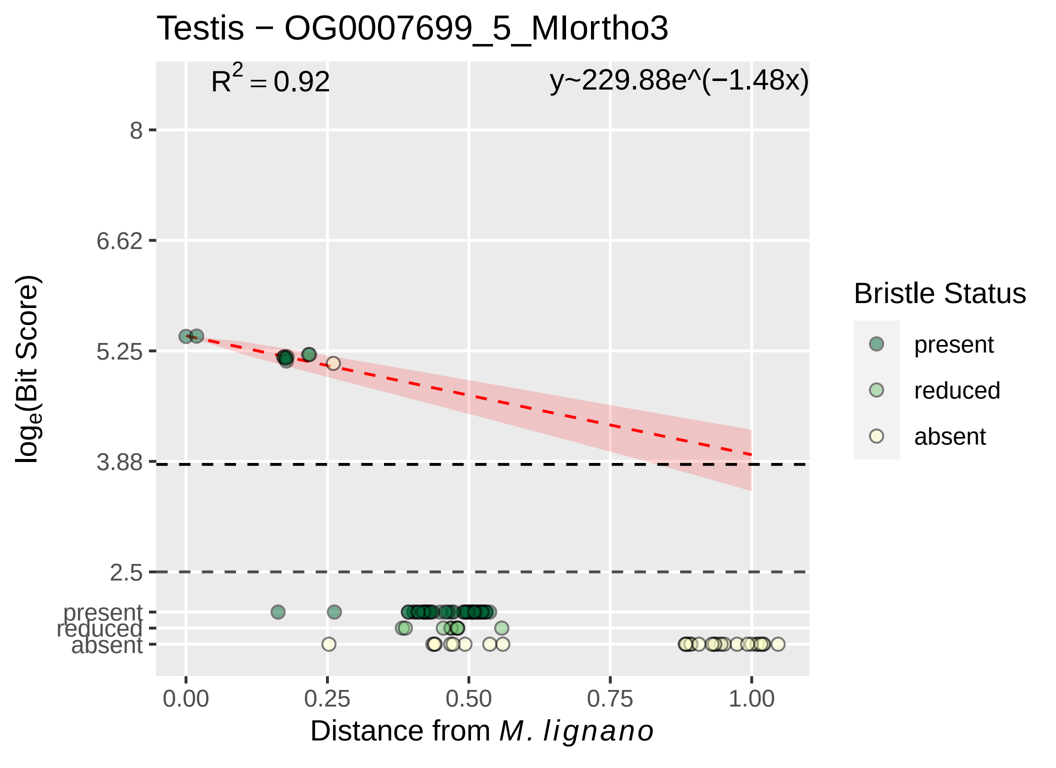

Testis − OG0007699_5_MIortho3
$R^2 = 0.92$
y~229.88e^(−1.48x)
8
6.62
5.25
3.88
2.5
present
reduced
absent
0.00
0.25
0.50
0.75
1.00
$\log_e$(Bit Score)
Distance from *M. lignano*
Bristle Status
present
reduced
absent

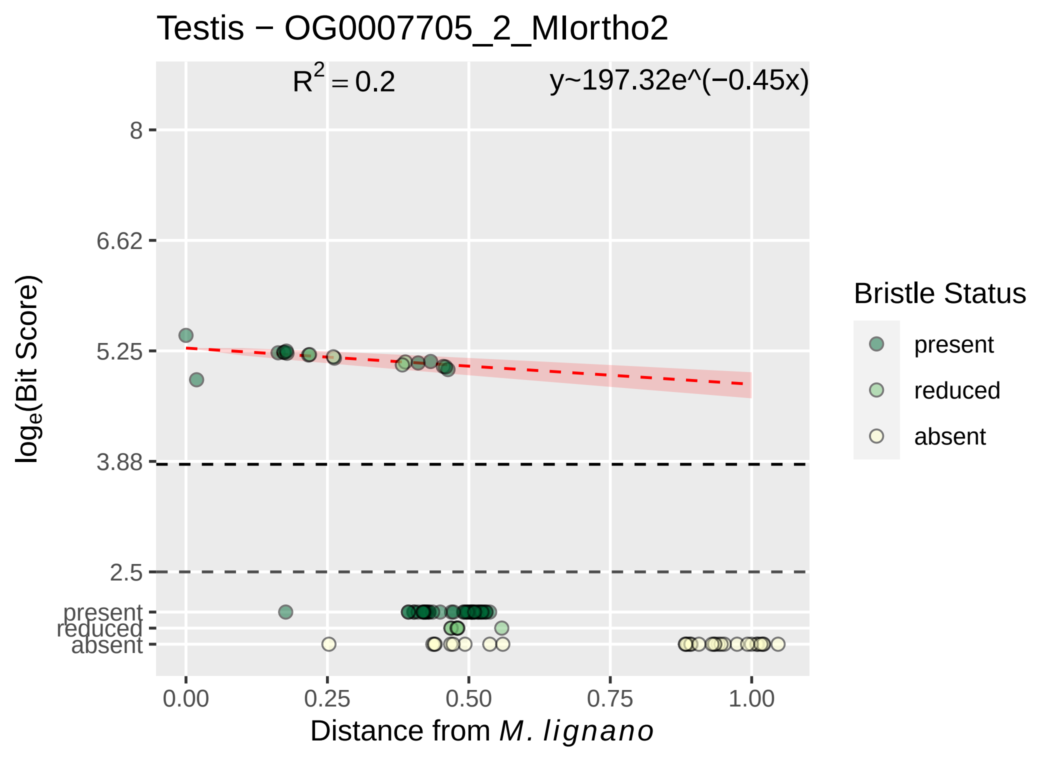

Testis − OG0007705_2_MIortho2
$R^2 = 0.2$
y~197.32e^(−0.45x)
8
6.62
5.25
3.88
2.5
present
reduced
absent
$\log_e$(Bit Score)
0.00
0.25
0.50
0.75
1.00
Distance from *M. lignano*
Bristle Status
present
reduced
absent

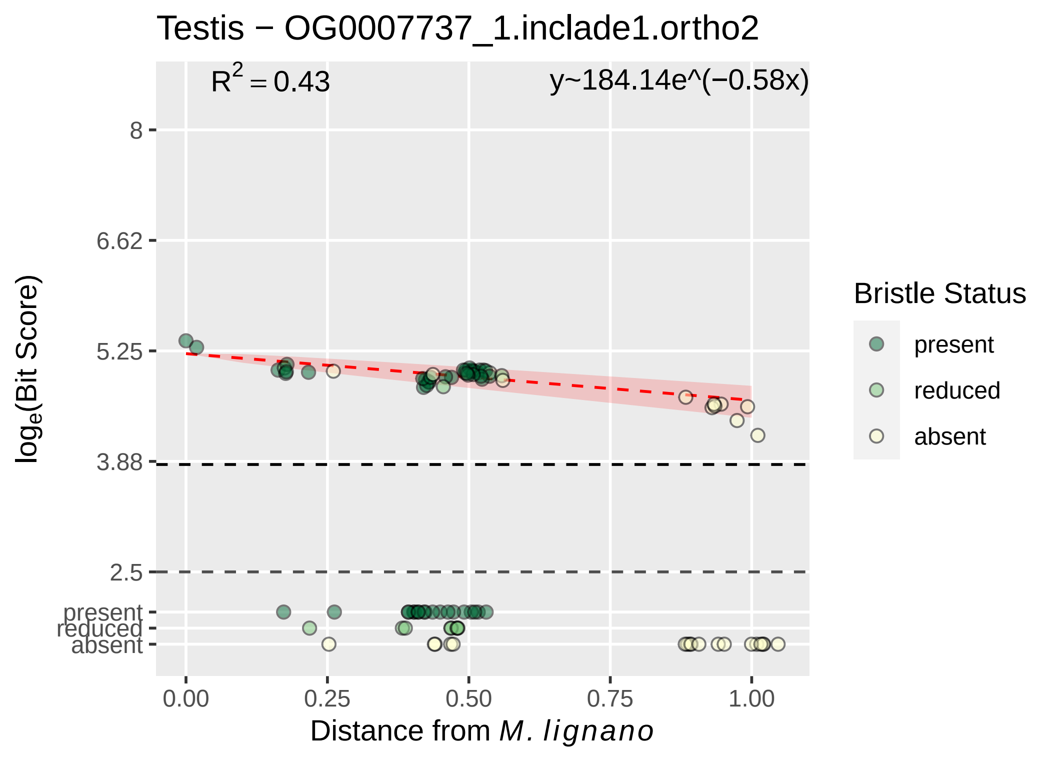

Testis − OG0007737_1.inclade1.ortho2
$R^2 = 0.43$
y~184.14e^(−0.58x)
8
6.62
5.25
3.88
2.5
present
reduced
absent
$\log_e$(Bit Score)
0.00
0.25
0.50
0.75
1.00
Distance from *M. lignano*
Bristle Status
present
reduced
absent

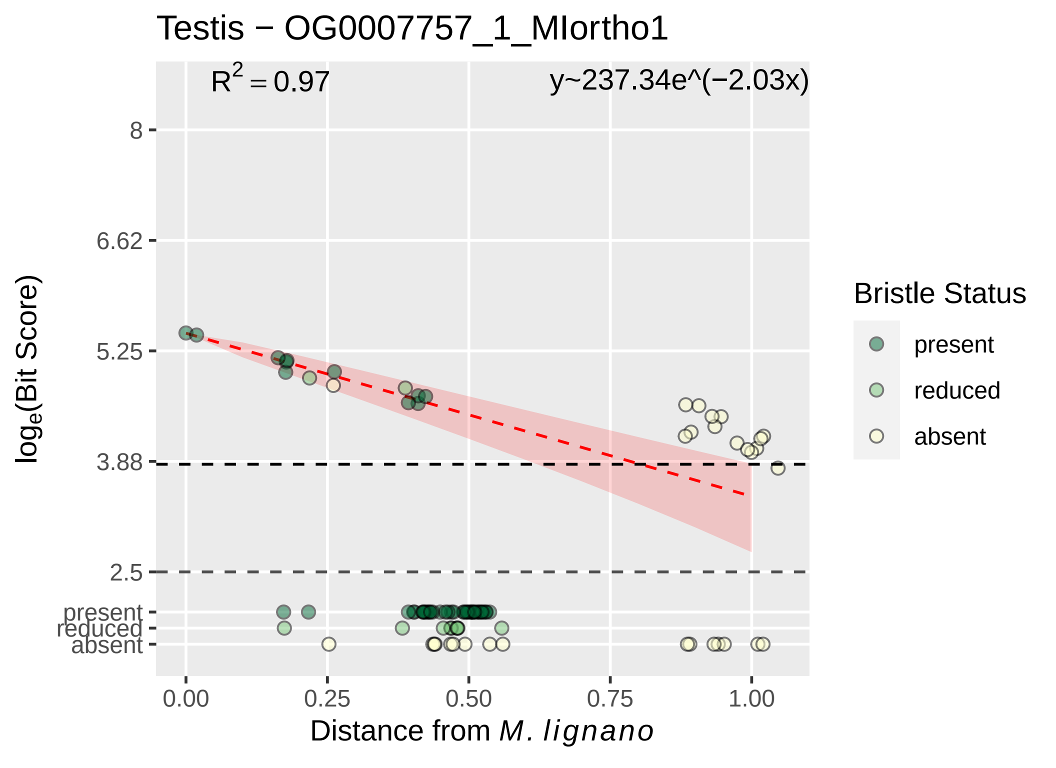

Testis − OG0007757_1_MIortho1
$R^2 = 0.97$
y~237.34e^(−2.03x)
8
6.62
5.25
3.88
2.5
present
reduced
absent
$\log_e$(Bit Score)
0.00
0.25
0.50
0.75
1.00
Distance from *M. lignano*
Bristle Status
present
reduced
absent

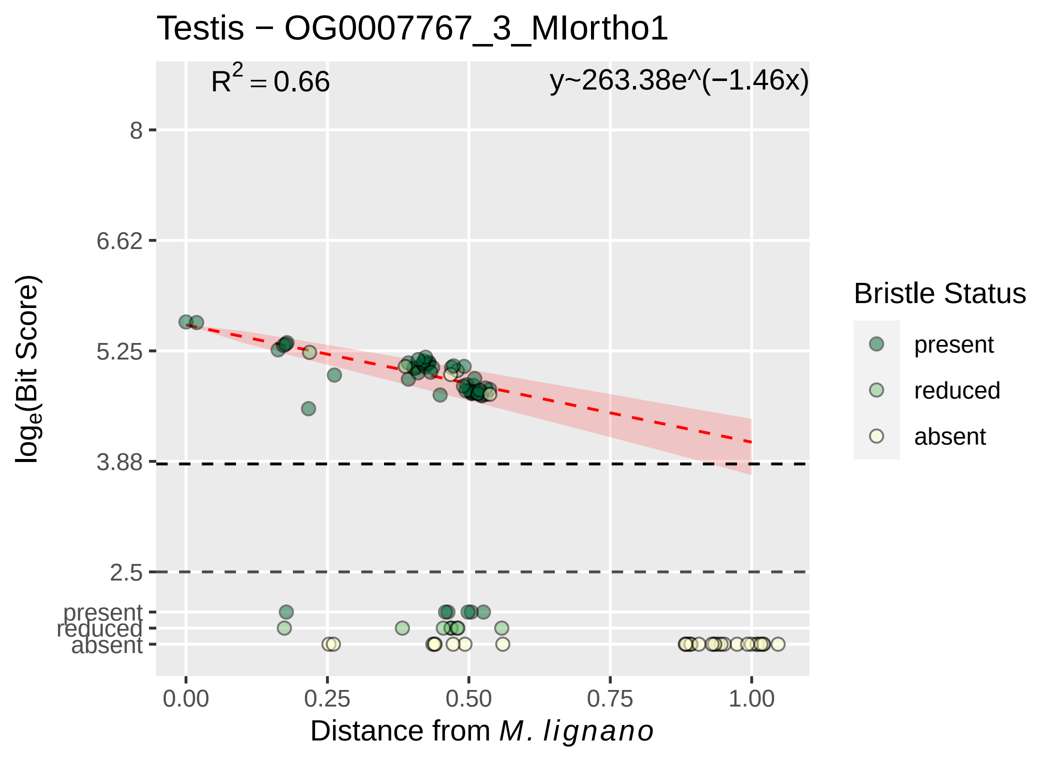

Testis − OG0007767_3_MIortho1
R^2 = 0.66
y~263.38e^(−1.46x)
8
6.62
5.25
3.88
2.5
present
reduced
absent
log_e(Bit Score)
0.00
0.25
0.50
0.75
1.00
Distance from M. lignano
Bristle Status
present
reduced
absent

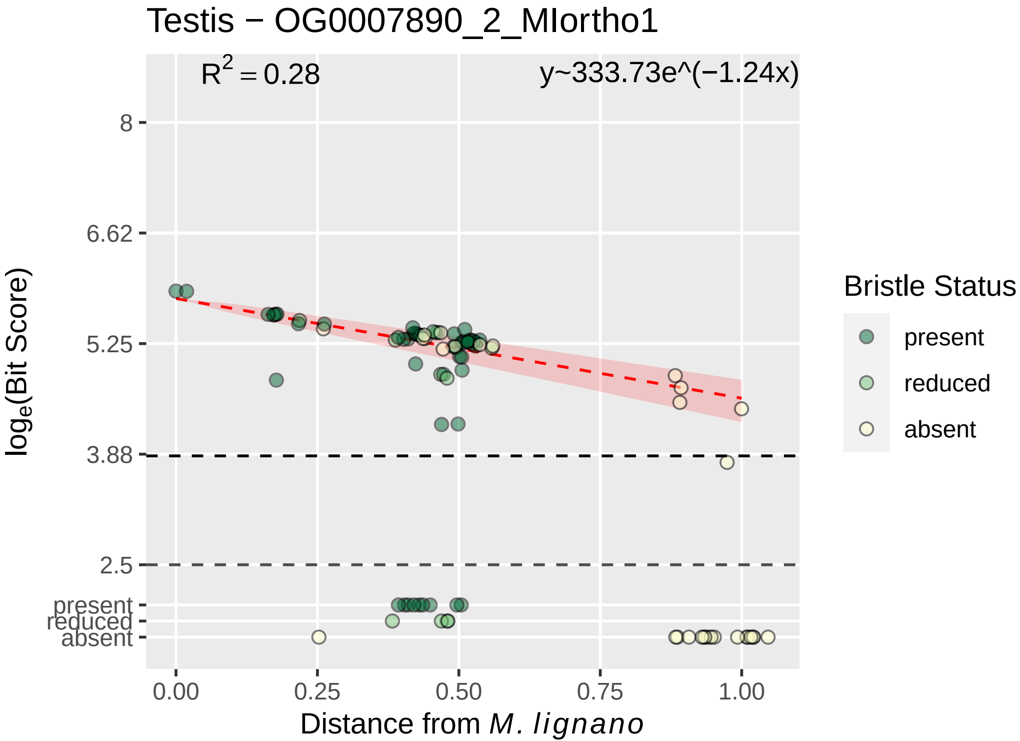
Testis − OG0007890_2_MIortho1
$R^2 = 0.28$
y~333.73e^(−1.24x)
8
6.62
5.25
3.88
2.5
present
reduced
absent
$log_e$(Bit Score)
0.00
0.25
0.50
0.75
1.00
Distance from *M. lignano*
Bristle Status
present
reduced
absent

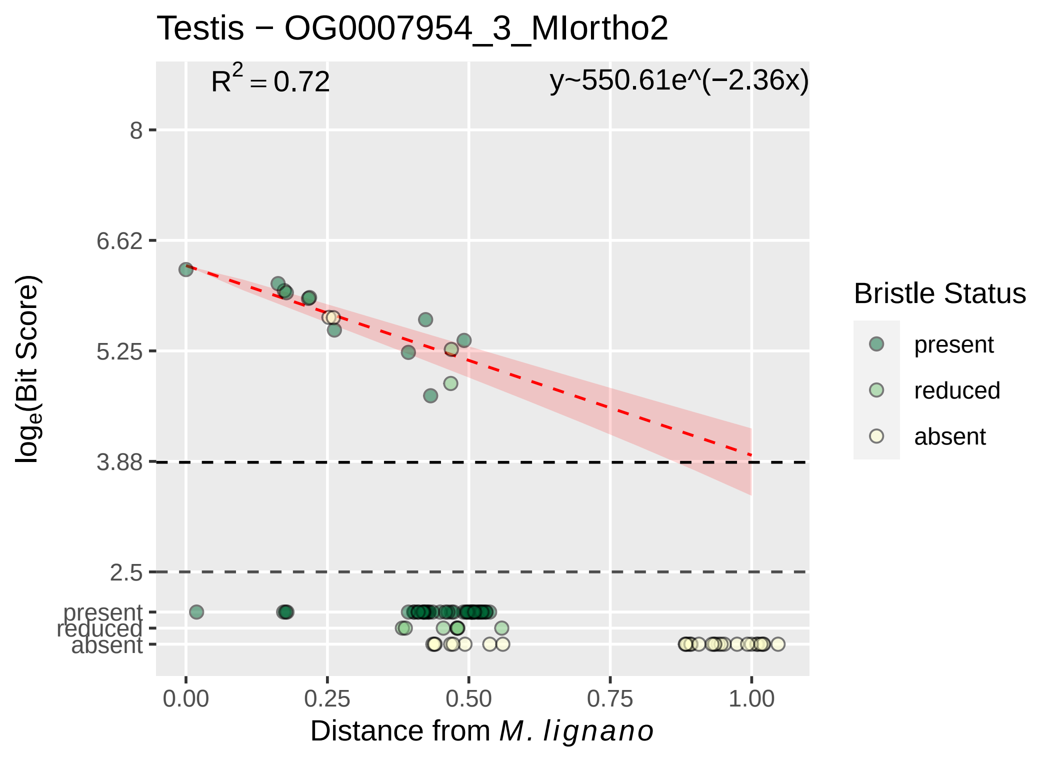

Testis − OG0007954_3_MIortho2
$R^2 = 0.72$
y~550.61e^(−2.36x)
8
6.62
5.25
3.88
2.5
present
reduced
absent
$\log_e$(Bit Score)
0.00
0.25
0.50
0.75
1.00
Distance from *M. lignano*
Bristle Status
present
reduced
absent

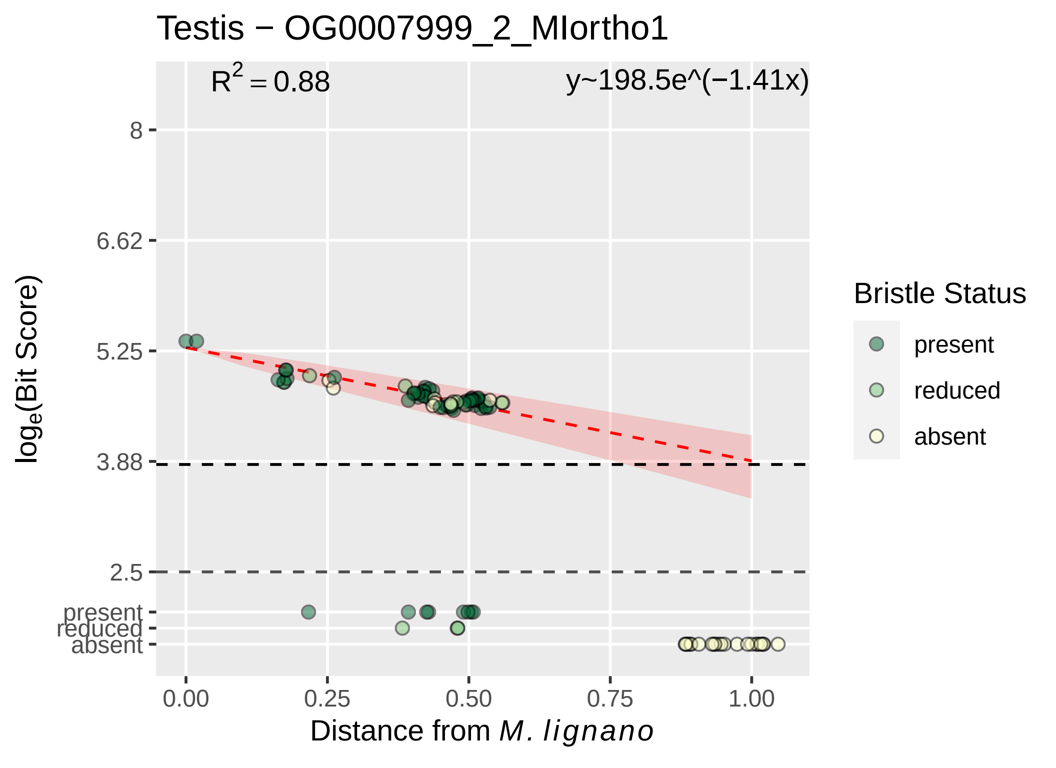

Testis − OG0007999_2_MIortho1
R² = 0.88
y~198.5e^(−1.41x)
8
6.62
5.25
3.88
2.5
present
reduced
absent
log_e(Bit Score)
0.00
0.25
0.50
0.75
1.00
Distance from M. lignano
Bristle Status
present
reduced
absent

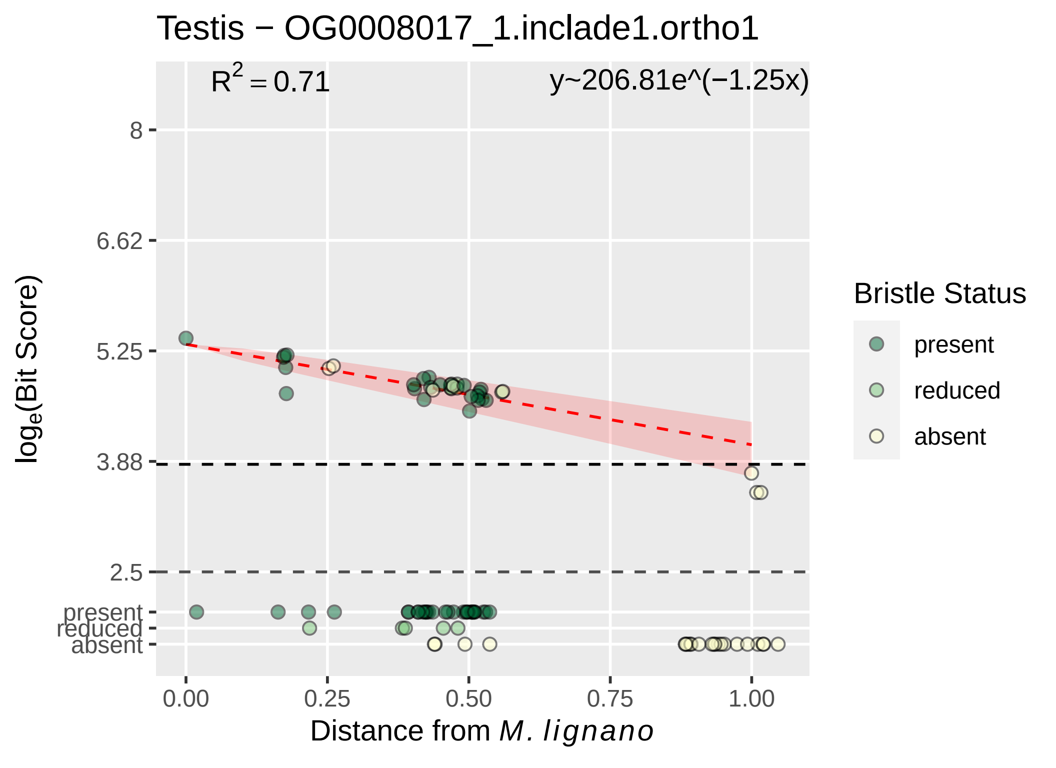

Testis − OG0008017_1.inclade1.ortho1
R² = 0.71
y~206.81e^(−1.25x)
8
6.62
5.25
3.88
2.5
present
reduced
absent
loge(Bit Score)
0.00
0.25
0.50
0.75
1.00
Distance from M. lignano
Bristle Status
present
reduced
absent

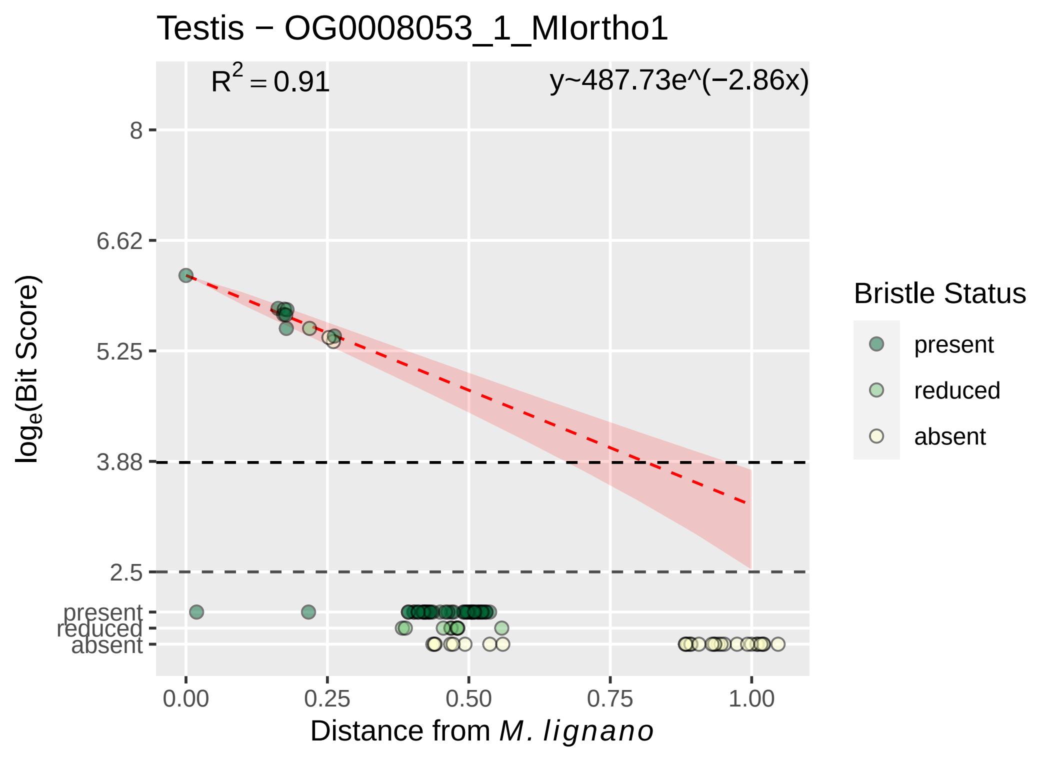

Testis − OG0008053_1_MIortho1
R^2 = 0.91
y~487.73e^(−2.86x)
8
6.62
5.25
3.88
2.5
present
reduced
absent
0.00
0.25
0.50
0.75
1.00
Distance from M. lignano
log_e(Bit Score)
Bristle Status
present
reduced
absent

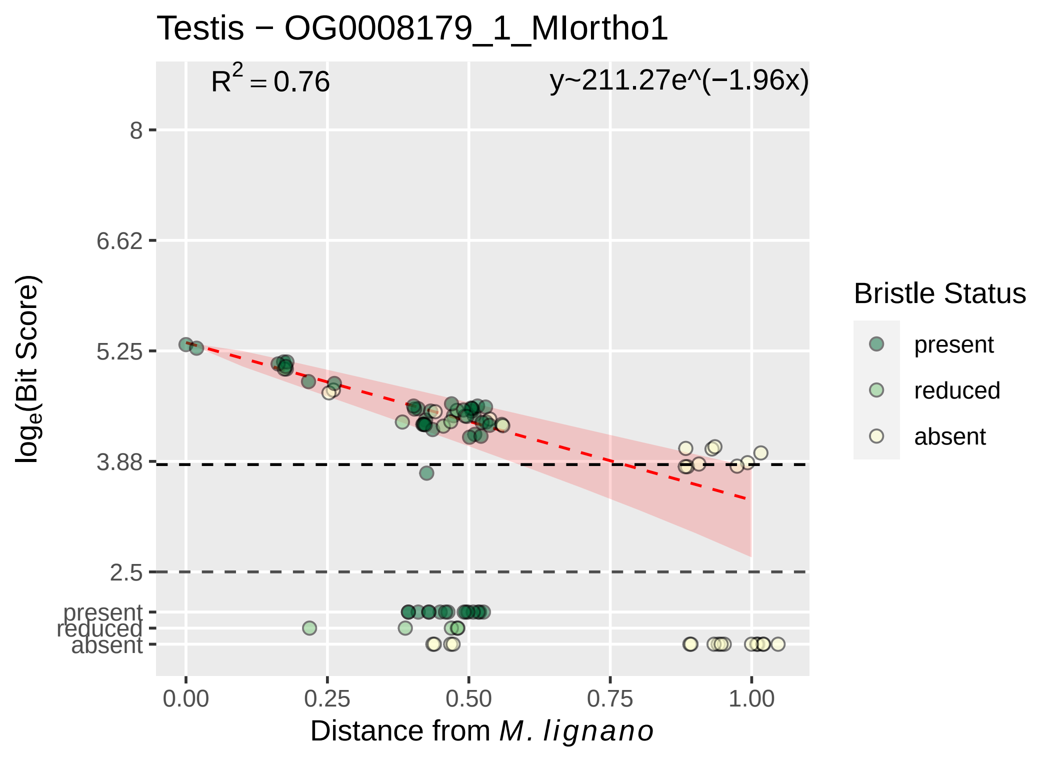

Testis − OG0008179_1_MIortho1
R^2 = 0.76
y~211.27e^(−1.96x)
8
6.62
5.25
3.88
2.5
present
reduced
absent
log_e(Bit Score)
0.00
0.25
0.50
0.75
1.00
Distance from M. lignano
Bristle Status
present
reduced
absent

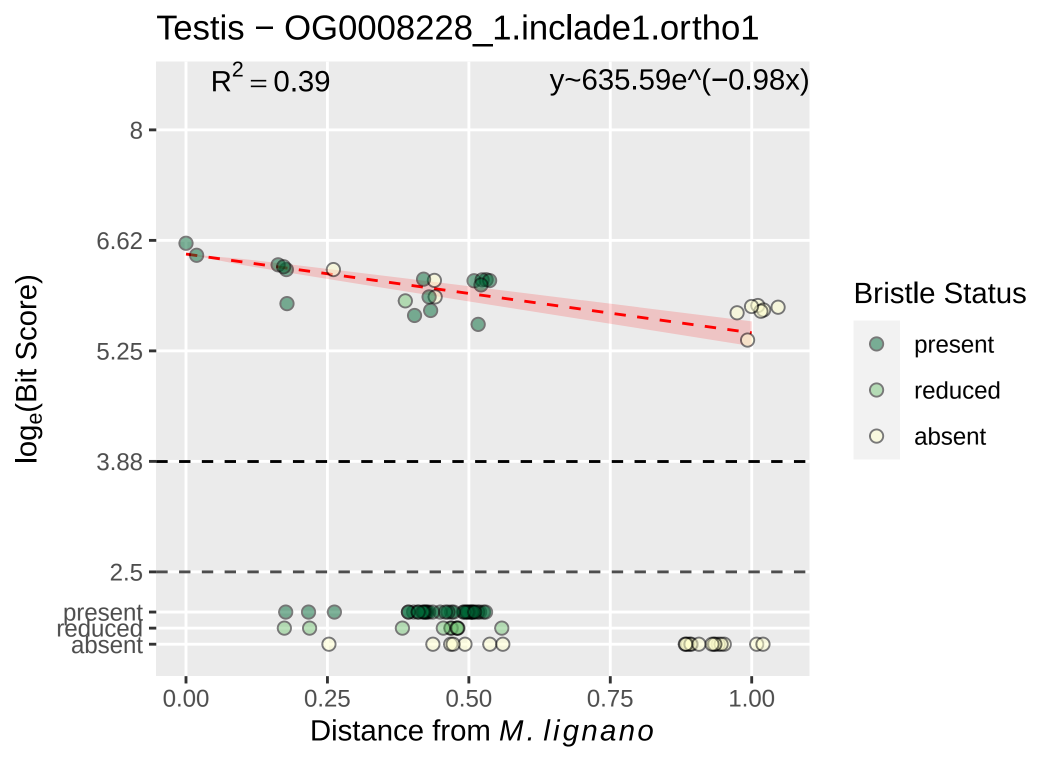

Testis − OG0008228_1.inclade1.ortho1
R² = 0.39
y~635.59e^(−0.98x)
8
6.62
5.25
3.88
2.5
present
reduced
absent
log_e(Bit Score)
0.00
0.25
0.50
0.75
1.00
Distance from M. lignano
Bristle Status
present
reduced
absent

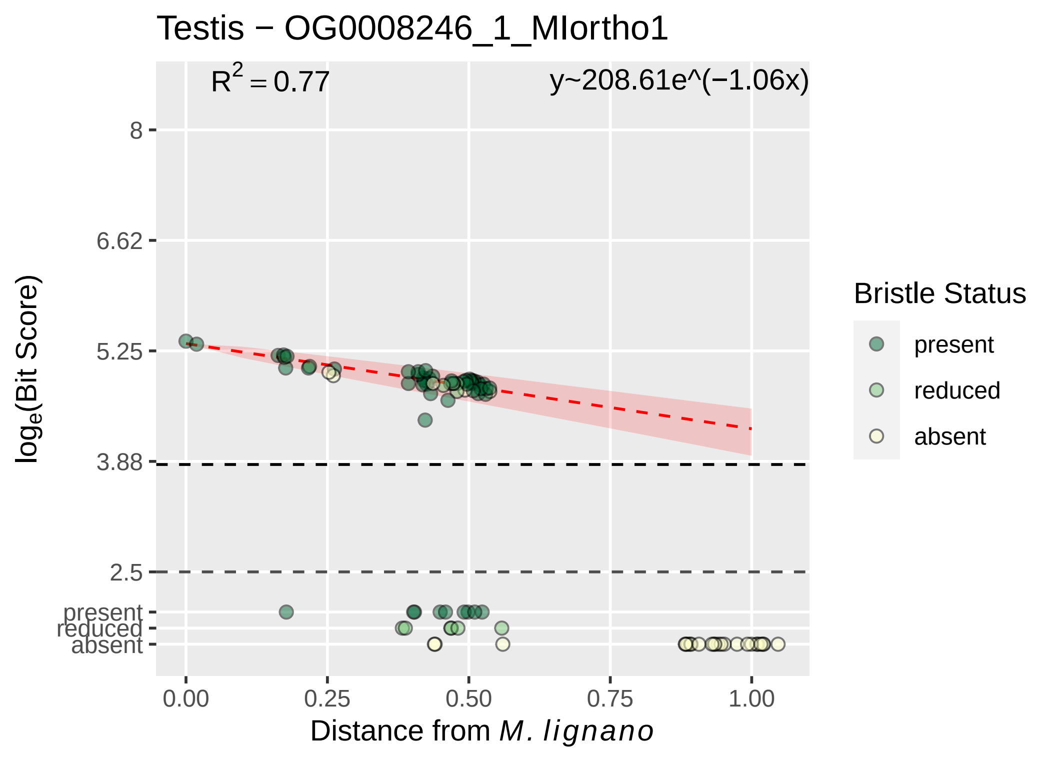

Testis − OG0008246_1_MIortho1
$R^2 = 0.77$
y~208.61e^(−1.06x)
8
6.62
5.25
3.88
2.5
present
reduced
absent
$\log_e$(Bit Score)
0.00
0.25
0.50
0.75
1.00
Distance from *M. lignano*
Bristle Status
present
reduced
absent

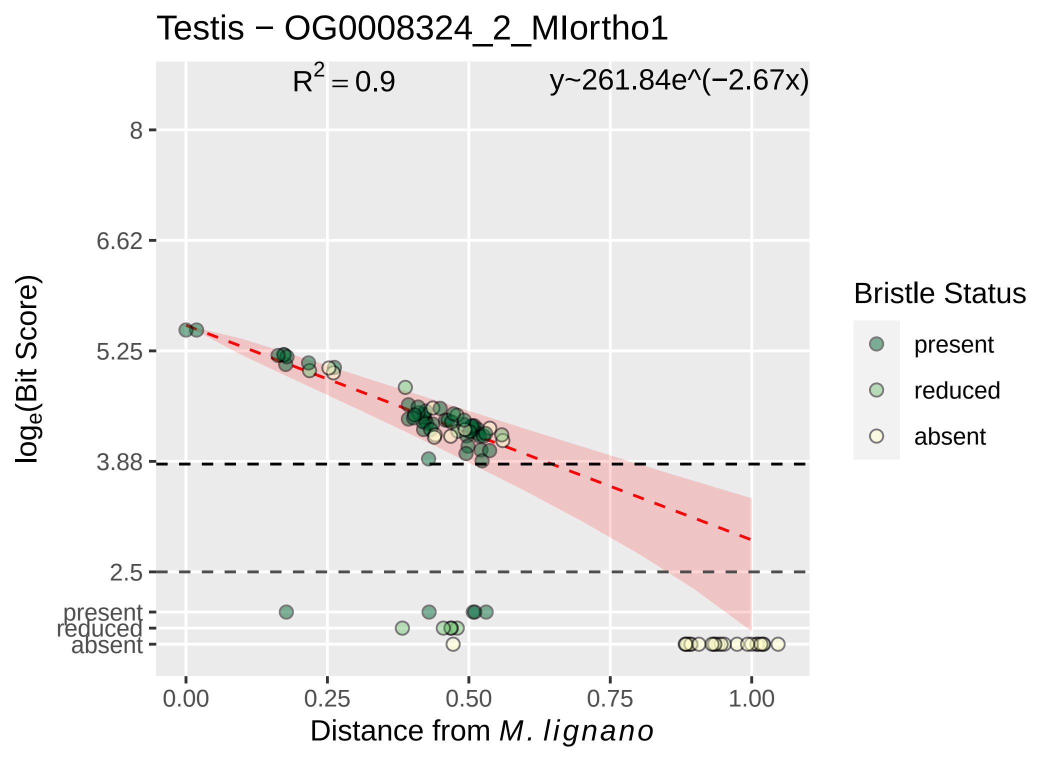

Testis − OG0008324_2_MIortho1
R2 = 0.9
y~261.84e^(−2.67x)
8
6.62
5.25
3.88
2.5
present
reduced
absent
loge(Bit Score)
0.00
0.25
0.50
0.75
1.00
Distance from M. lignano
Bristle Status
present
reduced
absent

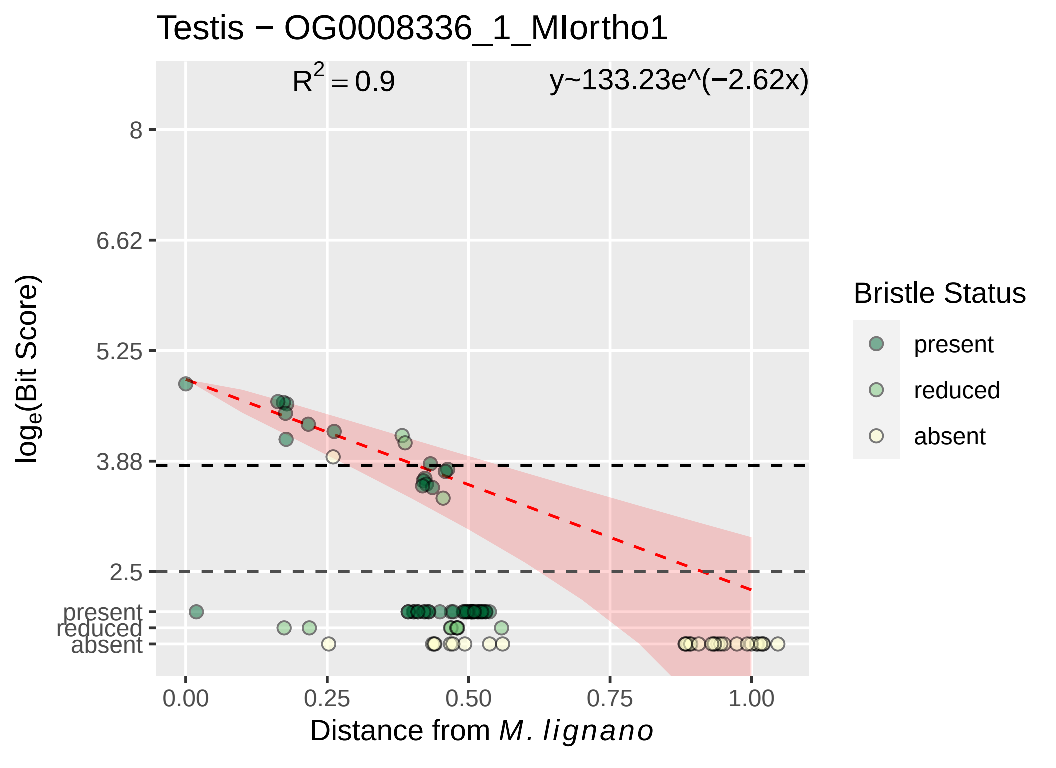

Testis − OG0008336_1_MIortho1
$R^2 = 0.9$
y~133.23e^(−2.62x)
8
6.62
5.25
3.88
2.5
present
reduced
absent
$\log_e$(Bit Score)
0.00
0.25
0.50
0.75
1.00
Distance from *M. lignano*
Bristle Status
present
reduced
absent

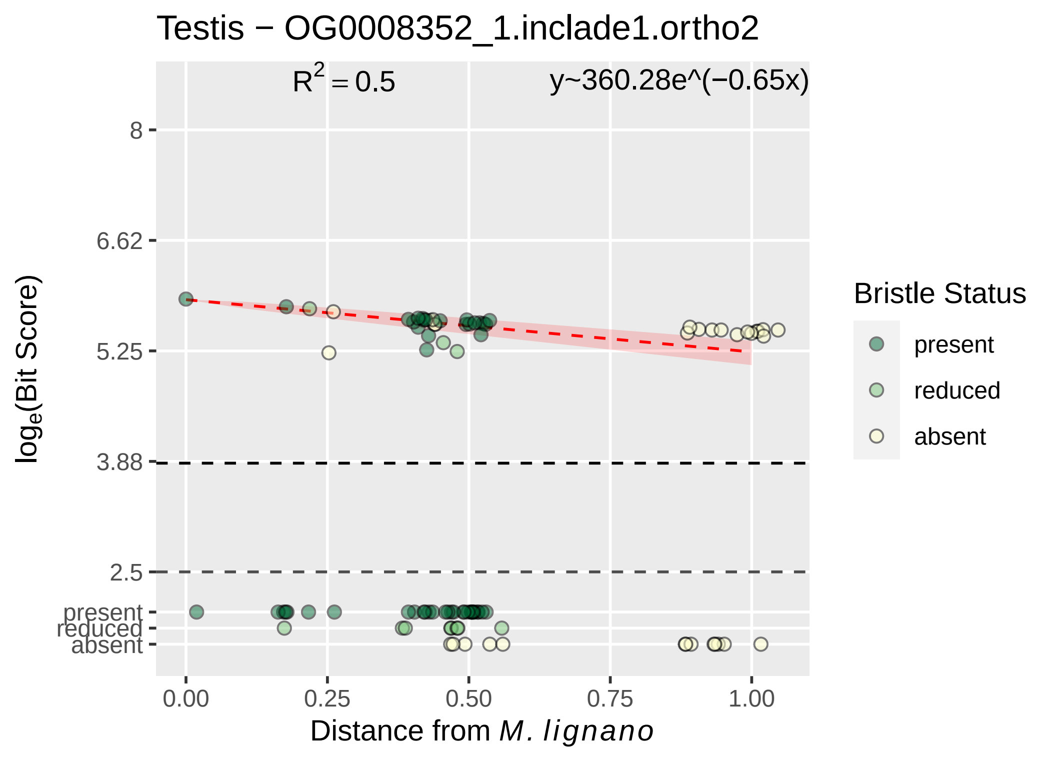

Testis − OG0008352_1.inclade1.ortho2
R2 = 0.5
y~360.28e^(−0.65x)
8
6.62
5.25
3.88
2.5
present
reduced
absent
log_e(Bit Score)
0.00
0.25
0.50
0.75
1.00
Distance from M. lignano
Bristle Status
present
reduced
absent

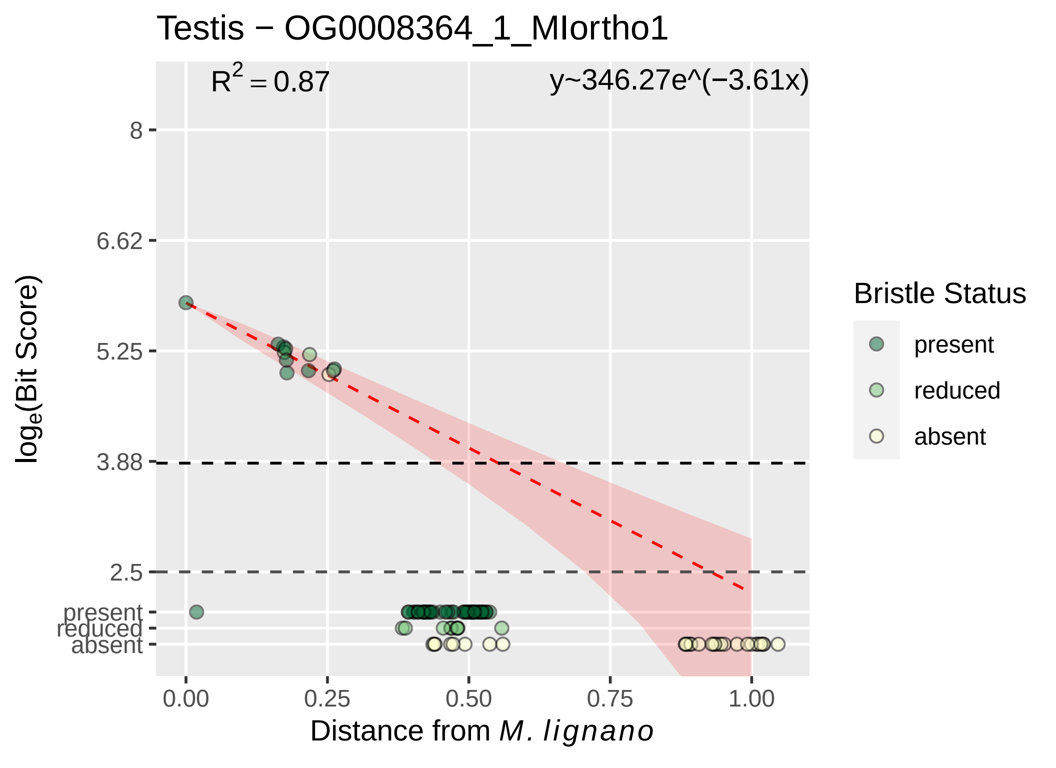

Testis − OG0008364_1_MIortho1
R2 = 0.87
y~346.27e^(−3.61x)
8
6.62
5.25
3.88
2.5
present
reduced
absent
0.00
0.25
0.50
0.75
1.00
Distance from M. lignano
loge(Bit Score)
Bristle Status
present
reduced
absent

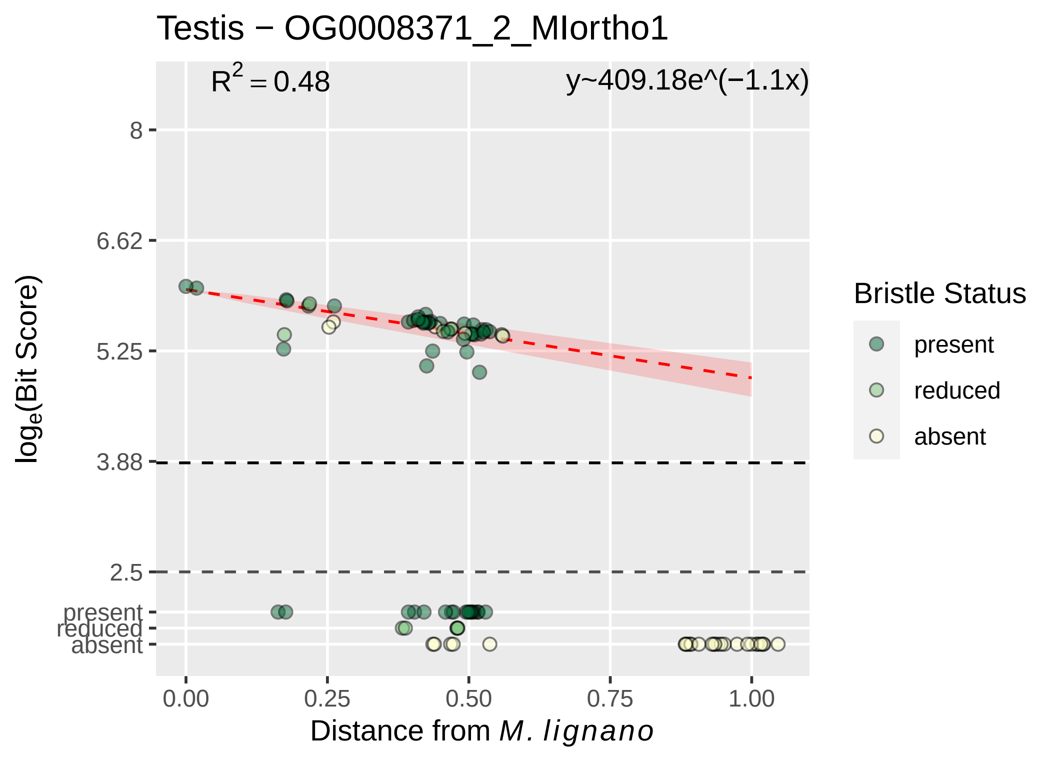

Testis − OG0008371_2_MIortho1
$R^2 = 0.48$
y~409.18e^(−1.1x)
8
6.62
5.25
3.88
2.5
present
reduced
absent
$\log_e$(Bit Score)
0.00
0.25
0.50
0.75
1.00
Distance from *M. lignano*
Bristle Status
present
reduced
absent

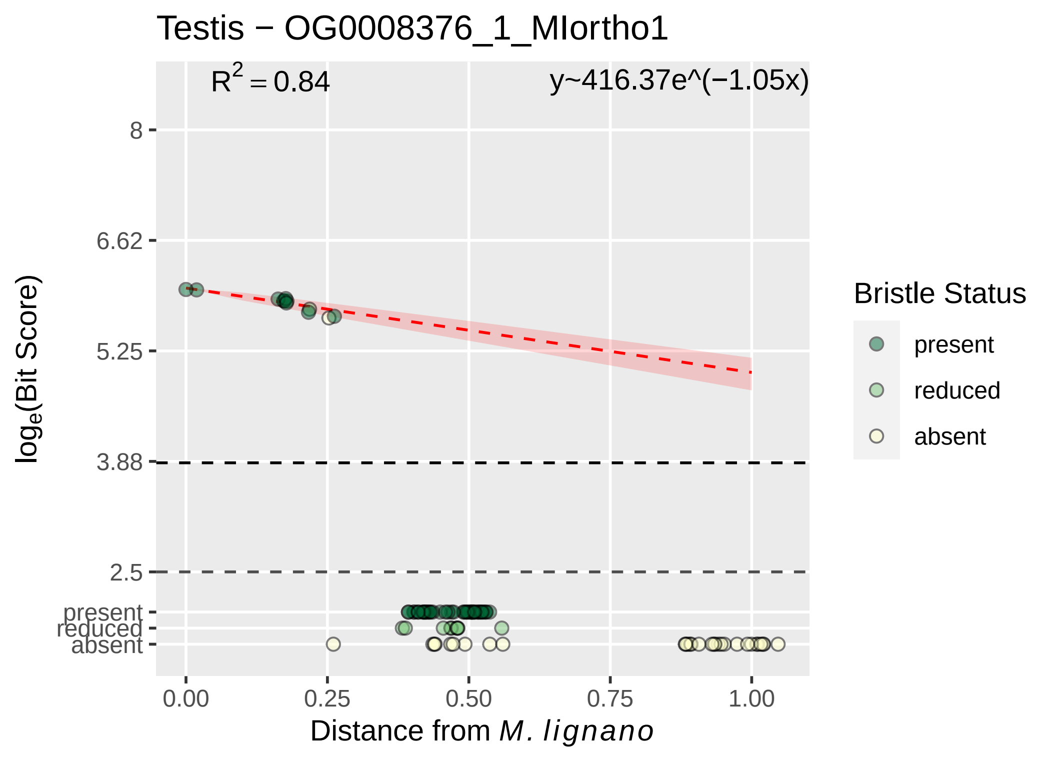

Testis − OG0008376_1_MIortho1
$R^2 = 0.84$
y~416.37e^(−1.05x)
8
6.62
5.25
3.88
2.5
present
reduced
absent
$\log_e$(Bit Score)
0.00
0.25
0.50
0.75
1.00
Distance from *M. lignano*
Bristle Status
present
reduced
absent

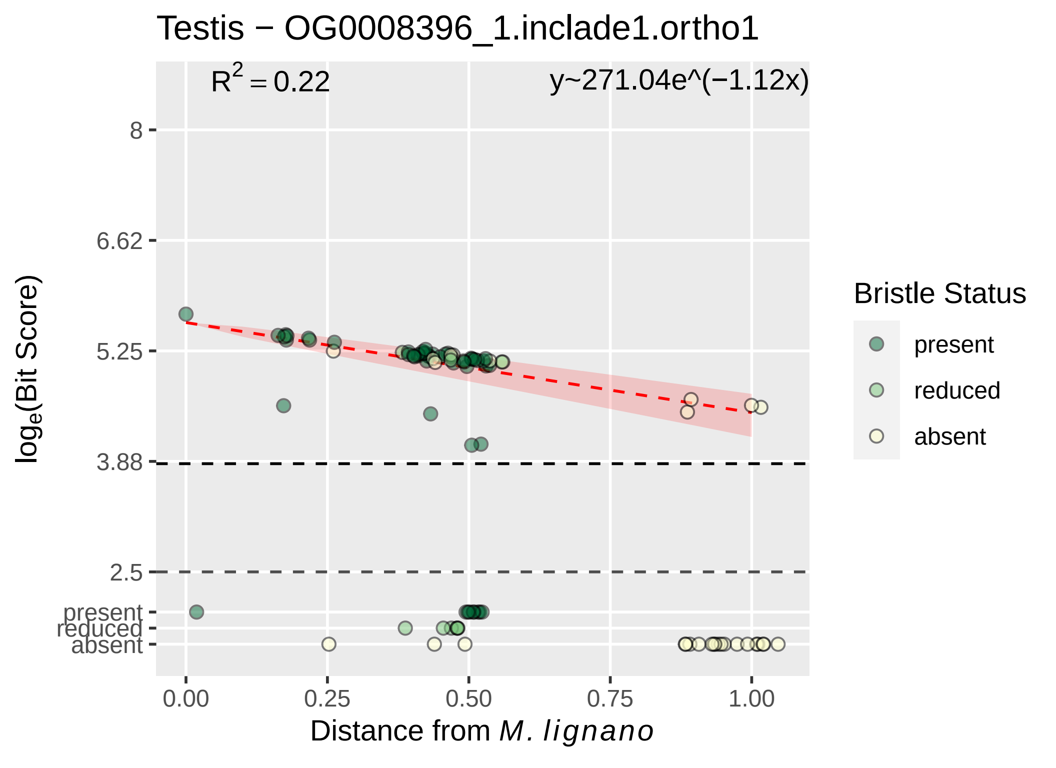

Testis − OG0008396_1.inclade1.ortho1
$R^2 = 0.22$
y~271.04e^(−1.12x)
8
6.62
5.25
3.88
2.5
present
reduced
absent
$\log_e$(Bit Score)
0.00
0.25
0.50
0.75
1.00
Distance from *M. lignano*
Bristle Status
present
reduced
absent

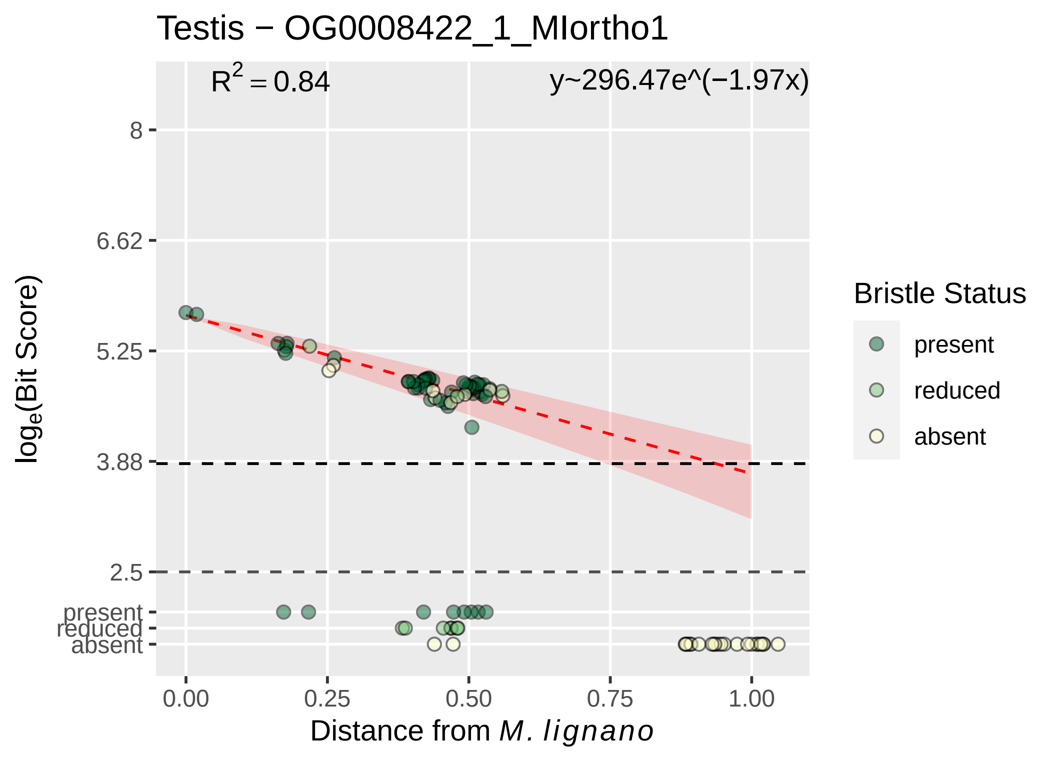

Testis − OG0008422_1_MIortho1
R^2 = 0.84
y~296.47e^(−1.97x)
8
6.62
5.25
3.88
2.5
present
reduced
absent
log_e(Bit Score)
0.00
0.25
0.50
0.75
1.00
Distance from M. lignano
Bristle Status
present
reduced
absent

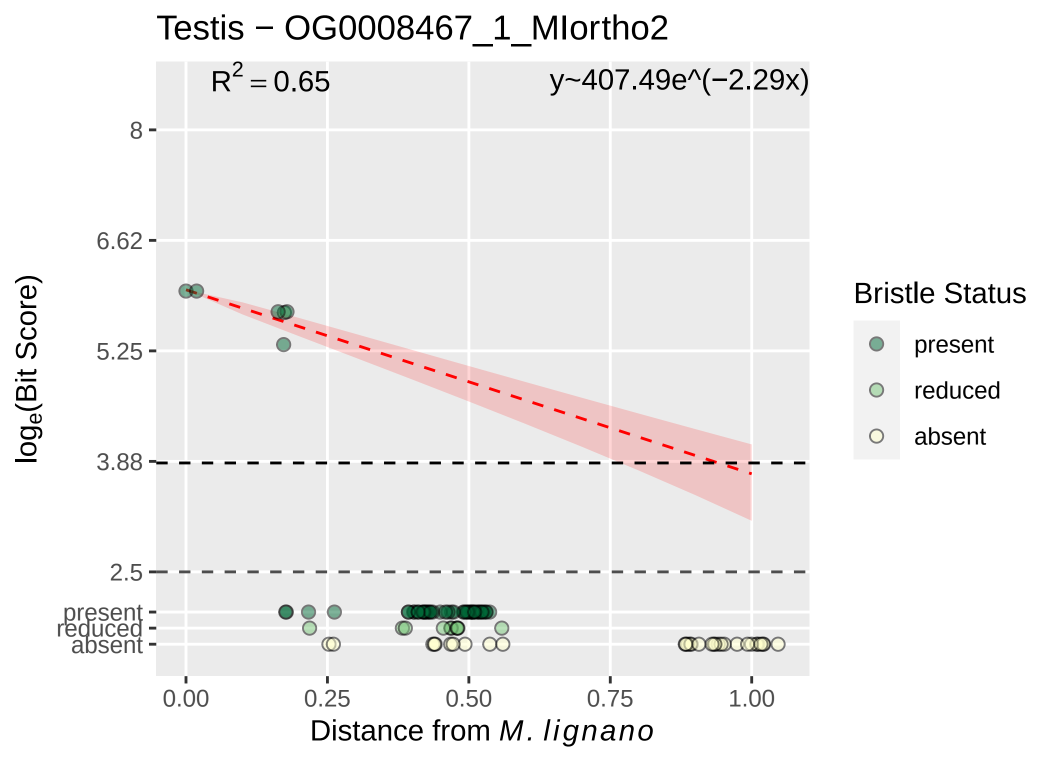

Testis − OG0008467_1_MIortho2
$R^2 = 0.65$
y~407.49e^(−2.29x)
8
6.62
5.25
3.88
2.5
present
reduced
absent
$\log_e$(Bit Score)
0.00
0.25
0.50
0.75
1.00
Distance from *M. lignano*
Bristle Status
present
reduced
absent

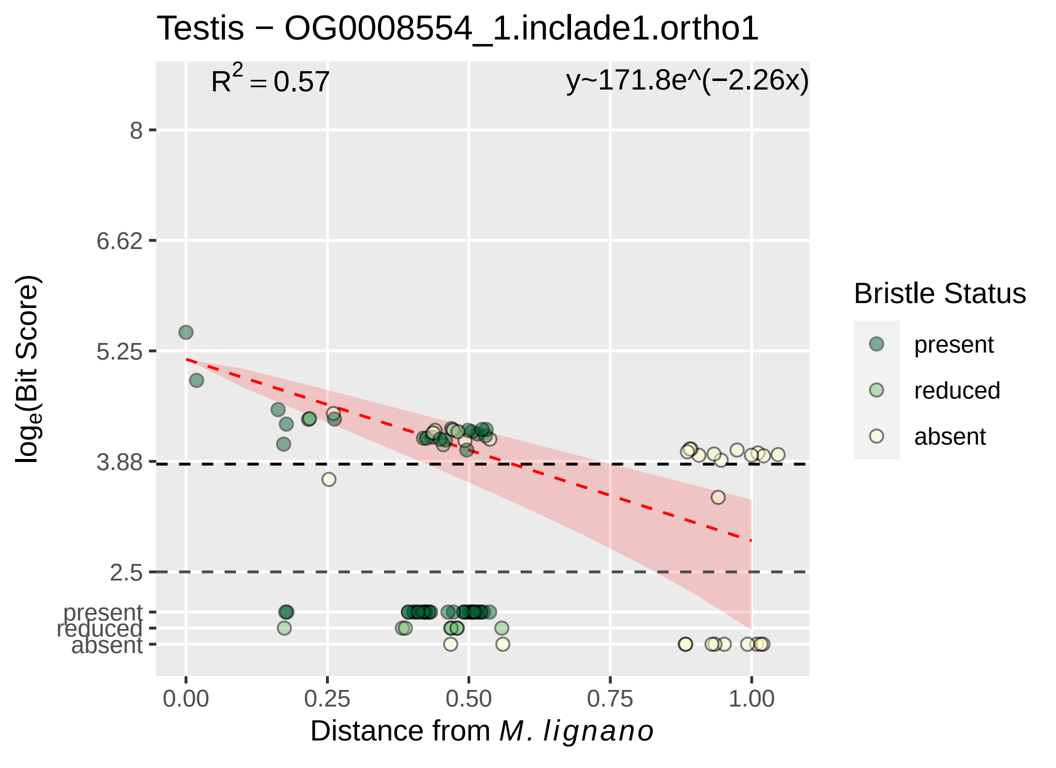

Testis − OG0008554_1.inclade1.ortho1
$R^2 = 0.57$
y~171.8e^(−2.26x)
8
6.62
5.25
3.88
2.5
present
reduced
absent
0.00
0.25
0.50
0.75
1.00
$log_e$(Bit Score)
Distance from *M. lignano*
Bristle Status
present
reduced
absent

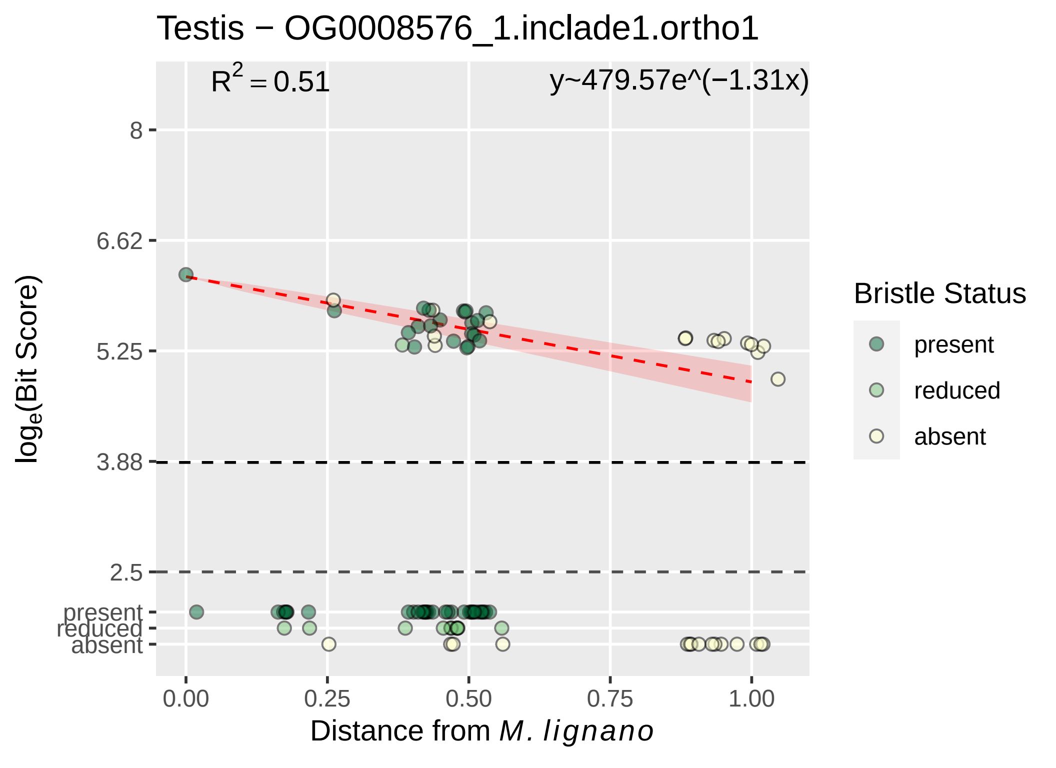

Testis − OG0008576_1.inclade1.ortho1
$R^2 = 0.51$
y~479.57e^(−1.31x)
8
6.62
5.25
3.88
2.5
present
reduced
absent
0.00
0.25
0.50
0.75
1.00
Distance from M. lignano
log$_e$(Bit Score)
Bristle Status
present
reduced
absent

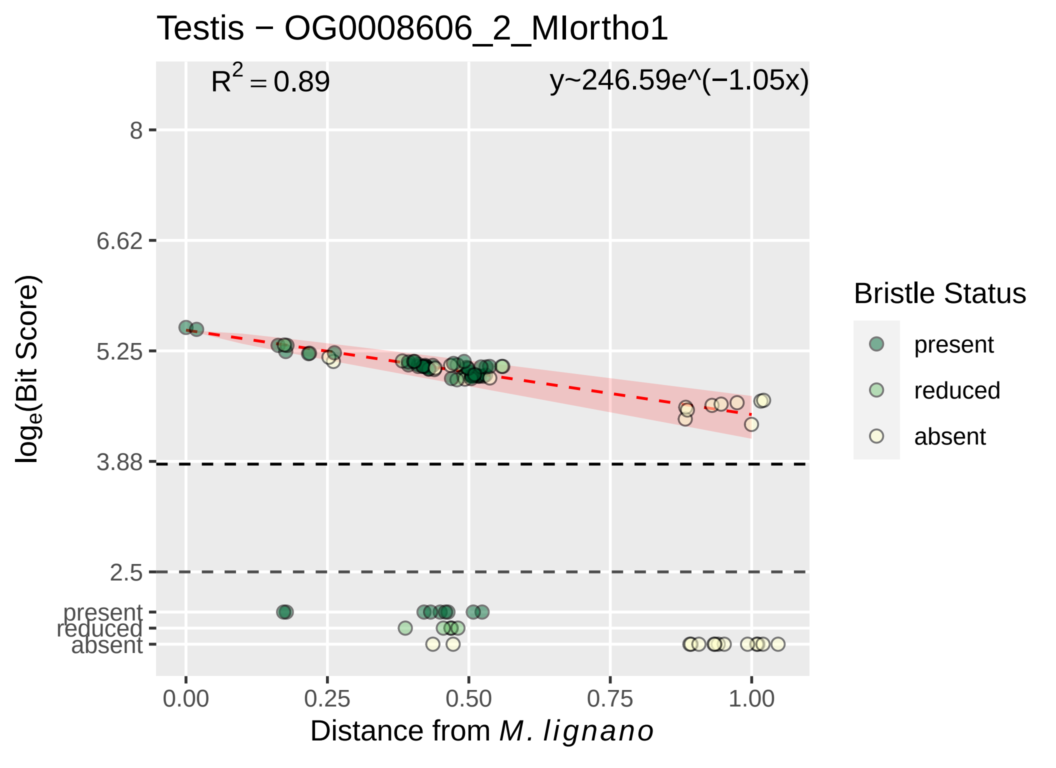

Testis − OG0008606_2_MIortho1
$R^2 = 0.89$
y~246.59e^(−1.05x)
8
6.62
5.25
3.88
2.5
present
reduced
absent
log_e(Bit Score)
0.00
0.25
0.50
0.75
1.00
Distance from M. lignano
Bristle Status
present
reduced
absent

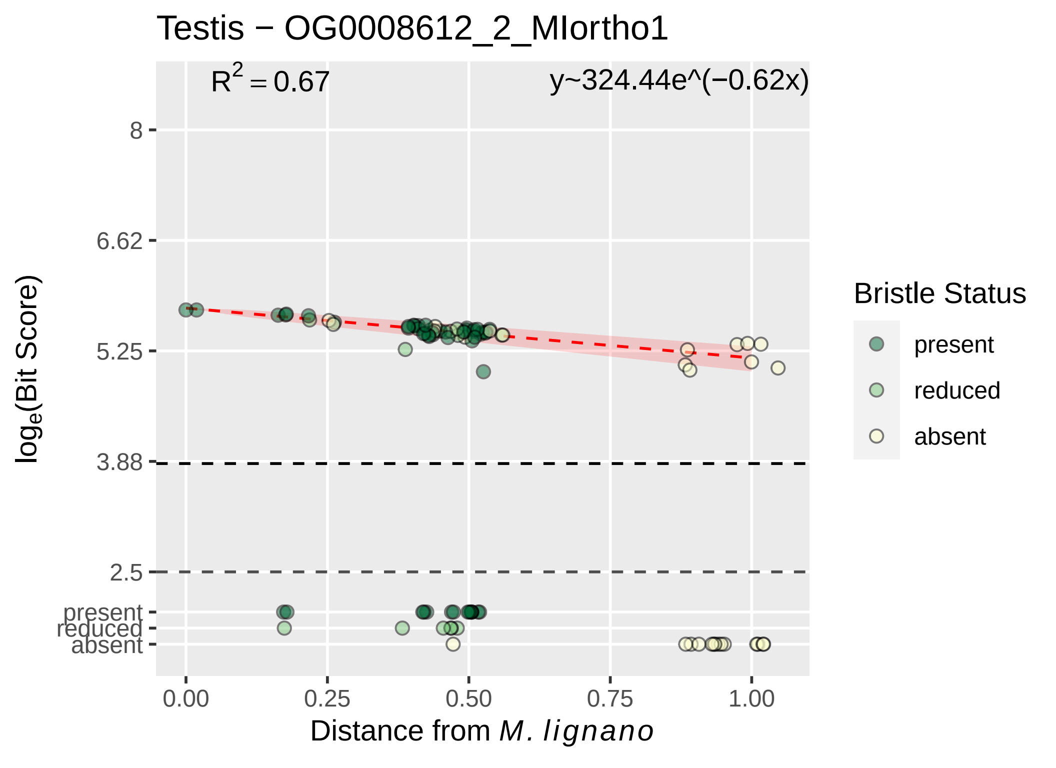

Testis − OG0008612_2_MIortho1
$R^2 = 0.67$
y~324.44e^(−0.62x)
8
6.62
5.25
3.88
2.5
present
reduced
absent
$\log_e$(Bit Score)
0.00
0.25
0.50
0.75
1.00
Distance from *M. lignano*
Bristle Status
present
reduced
absent

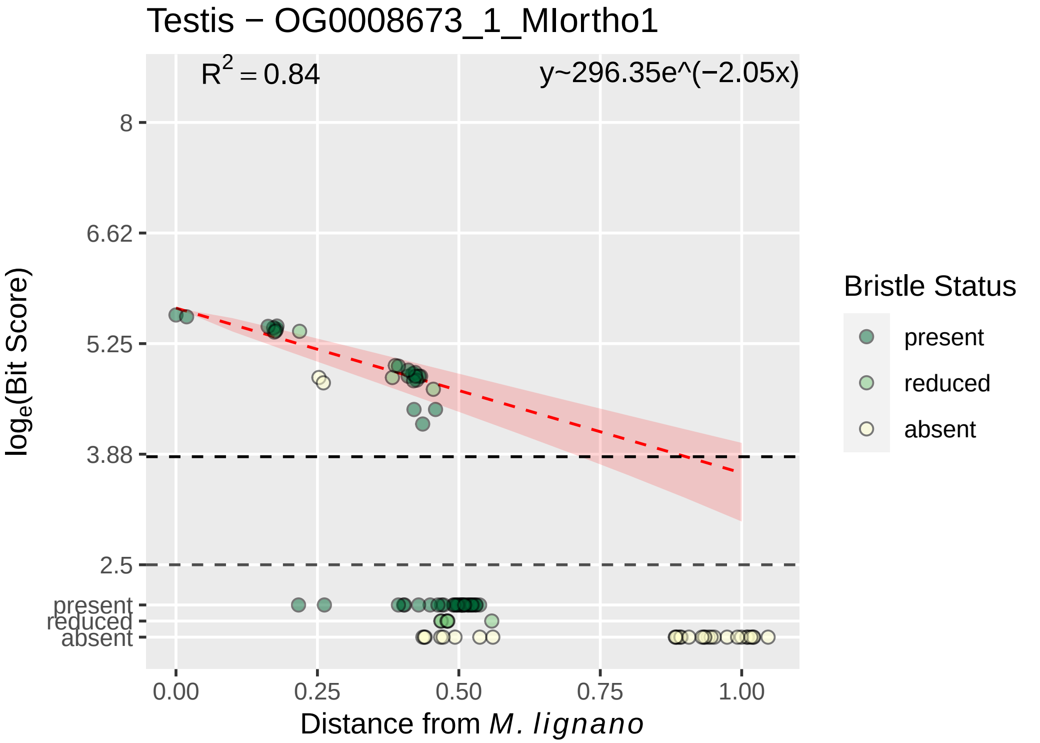
Testis − OG0008673_1_MIortho1
R^2 = 0.84
y~296.35e^(−2.05x)
8
6.62
5.25
3.88
2.5
present
reduced
absent
log_e(Bit Score)
0.00
0.25
0.50
0.75
1.00
Distance from M. lignano
Bristle Status
present
reduced
absent

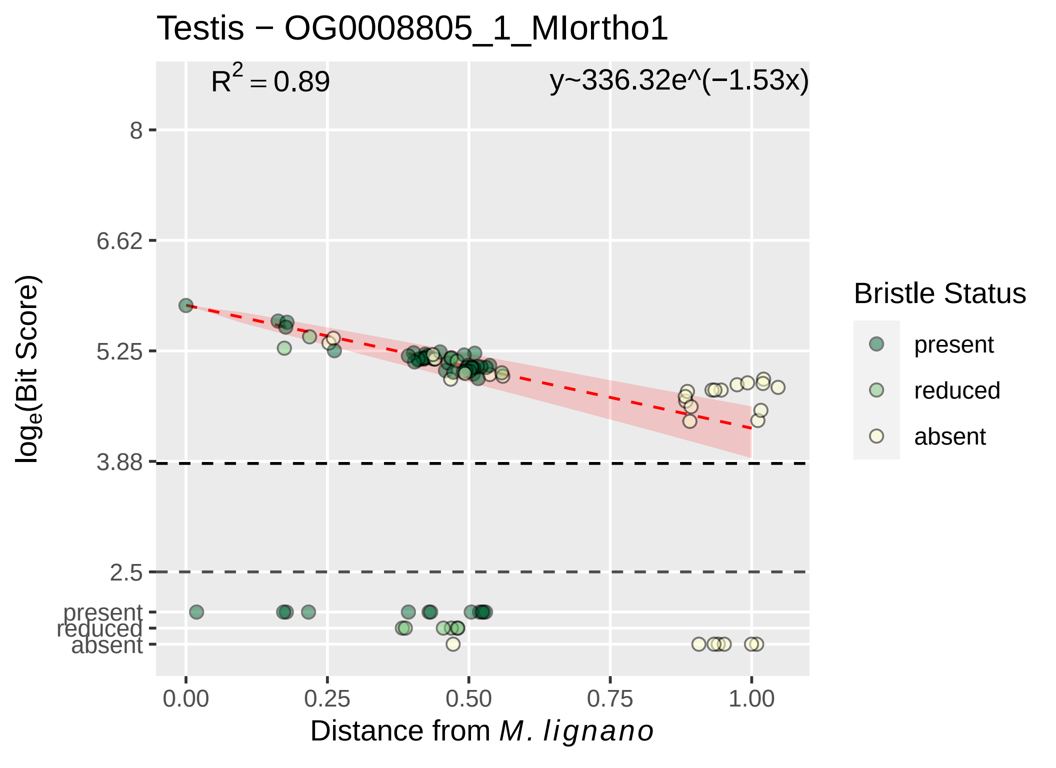

Testis − OG0008805_1_MIortho1
$R^2 = 0.89$
y~336.32e^(−1.53x)
8
6.62
5.25
3.88
2.5
present
reduced
absent
$log_e$(Bit Score)
0.00
0.25
0.50
0.75
1.00
Distance from *M. lignano*
Bristle Status
present
reduced
absent

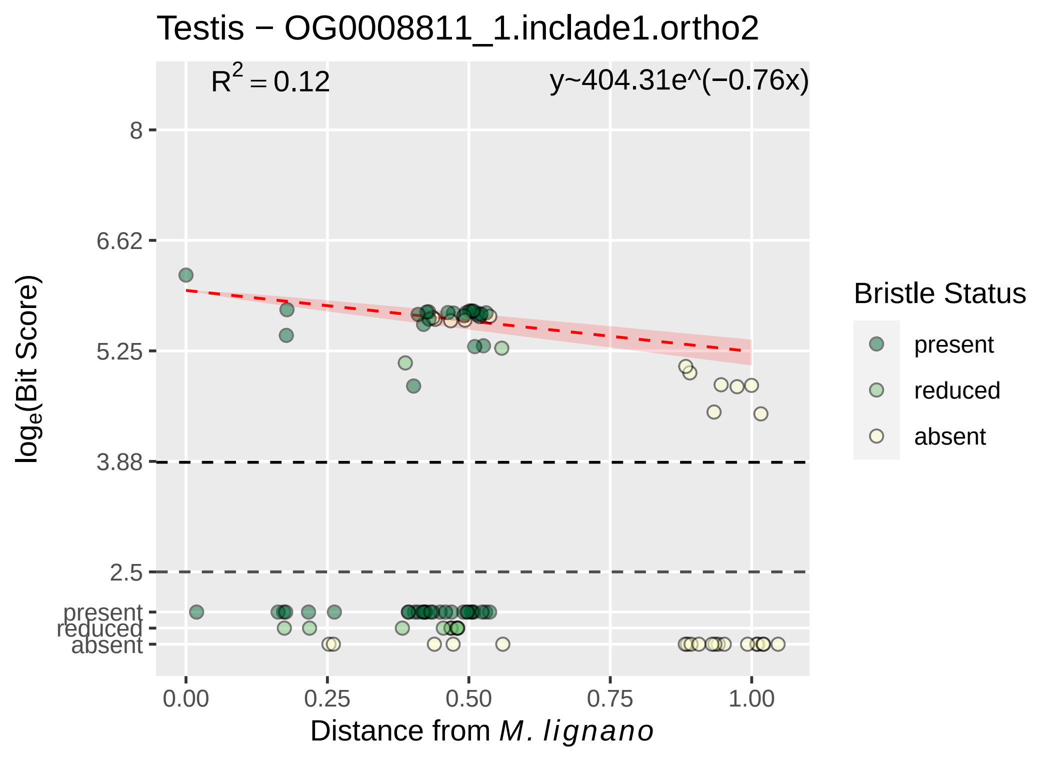

Testis − OG0008811_1.inclade1.ortho2
$R^2 = 0.12$
y~404.31e^(−0.76x)
8
6.62
5.25
3.88
2.5
present
reduced
absent
$\log_e$(Bit Score)
0.00
0.25
0.50
0.75
1.00
Distance from *M. lignano*
Bristle Status
present
reduced
absent

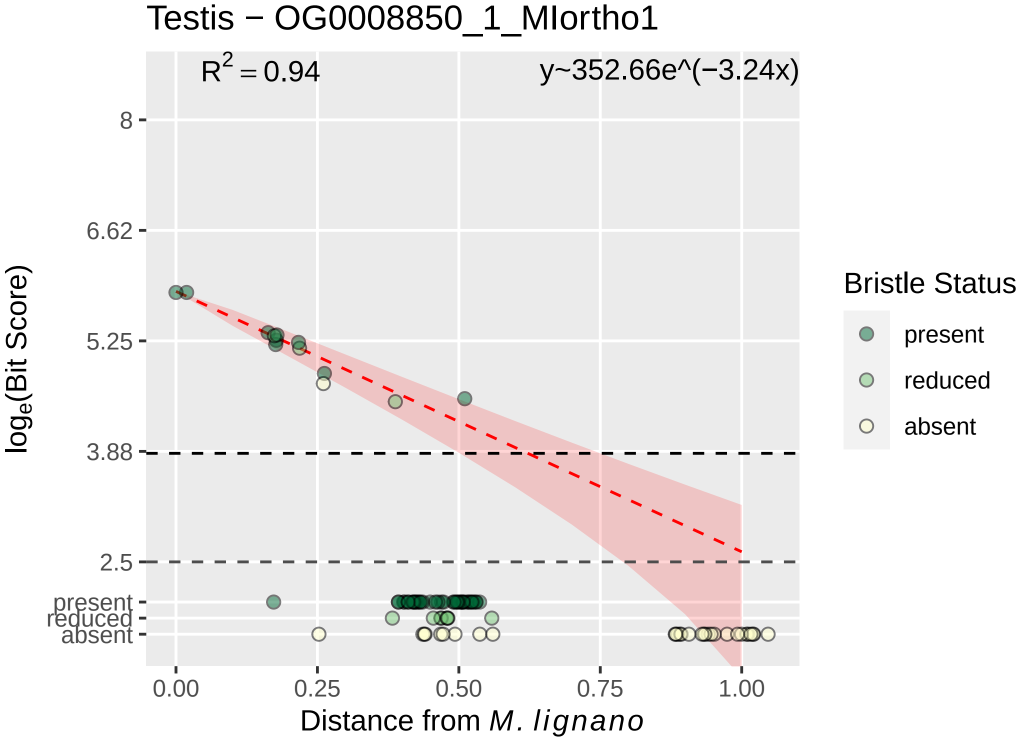

Testis − OG0008850_1_MIortho1
R2 = 0.94
y~352.66e^(−3.24x)
8
6.62
5.25
3.88
2.5
present
reduced
absent
log_e(Bit Score)
0.00
0.25
0.50
0.75
1.00
Distance from M. lignano
Bristle Status
present
reduced
absent

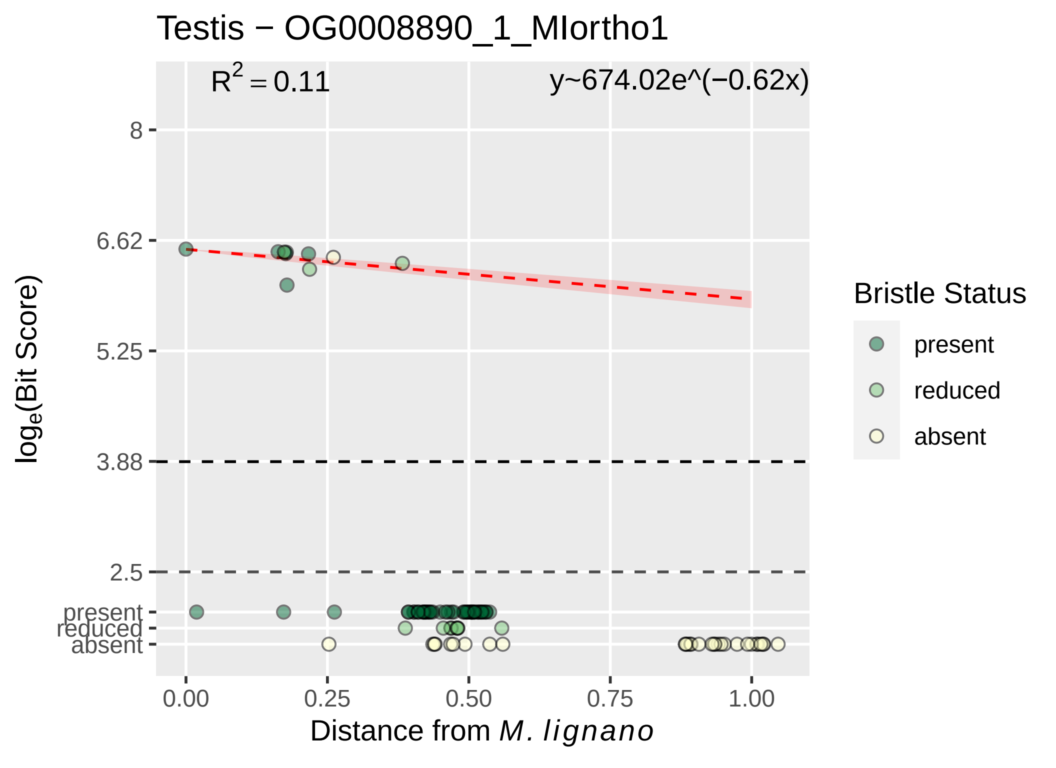

Testis − OG0008890_1_MIortho1
$R^2 = 0.11$
y~674.02e^(−0.62x)
8
6.62
5.25
3.88
2.5
present
reduced
absent
0.00
0.25
0.50
0.75
1.00
Distance from M. lignano
$\log_e$(Bit Score)
Bristle Status
present
reduced
absent

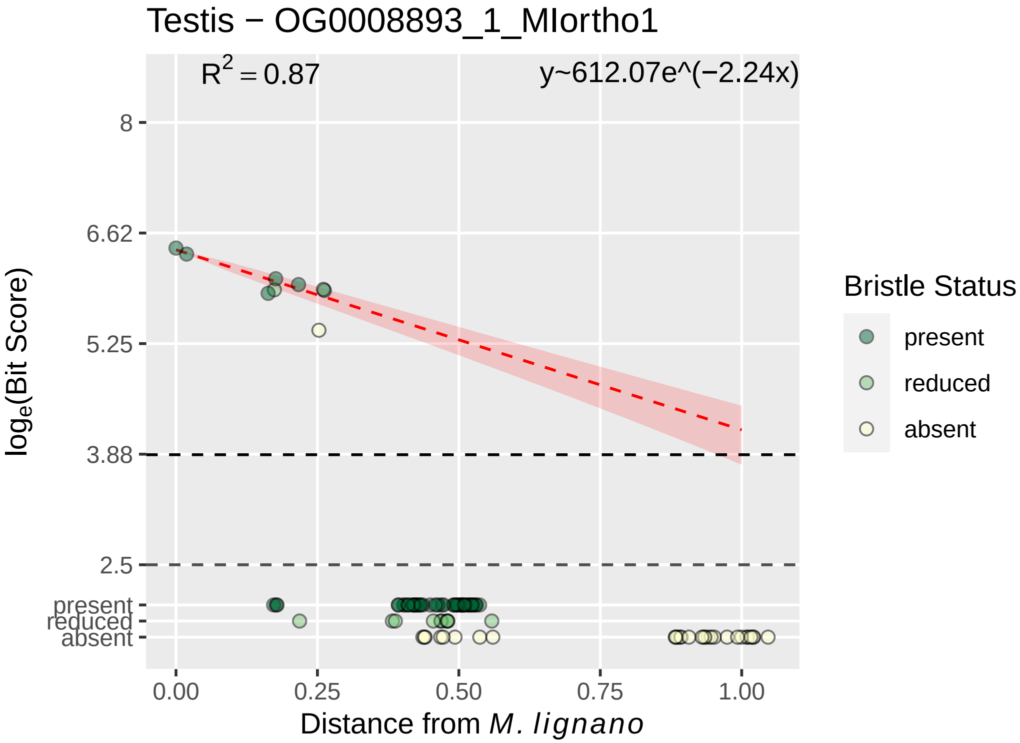

Testis − OG0008893_1_MIortho1
$R^2 = 0.87$
y~612.07e^(−2.24x)
8
6.62
5.25
3.88
2.5
present
reduced
absent
$\log_e$(Bit Score)
0.00
0.25
0.50
0.75
1.00
Distance from *M. lignano*
Bristle Status
present
reduced
absent

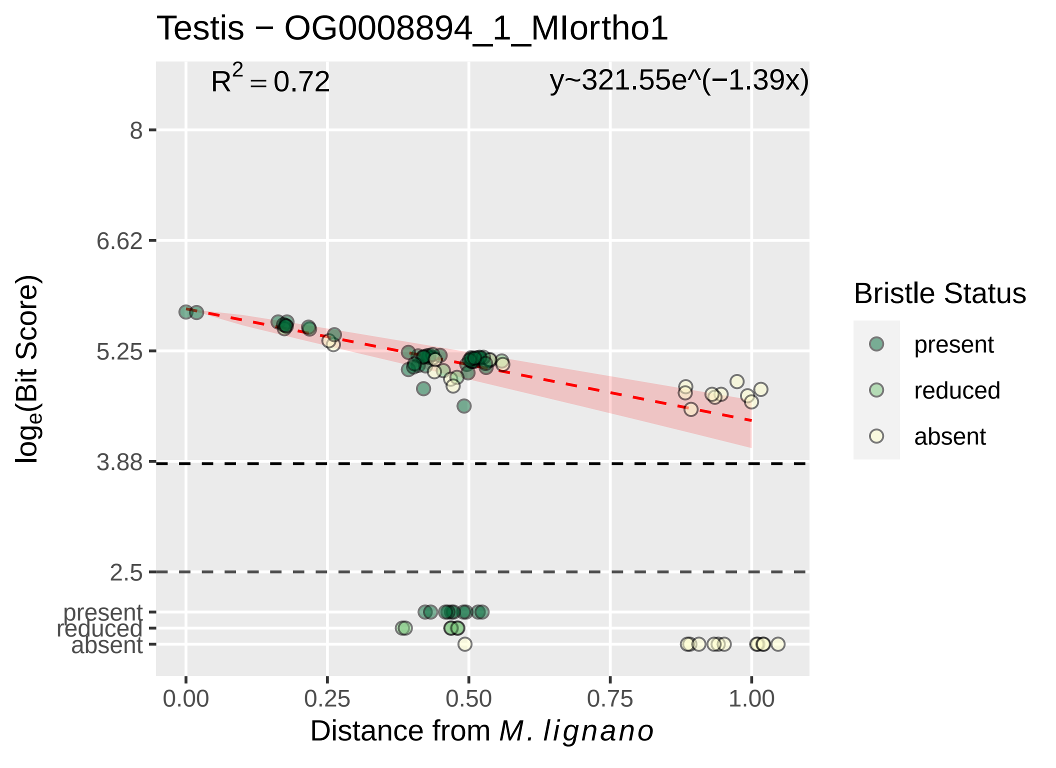

Testis − OG0008894_1_MIortho1
R2 = 0.72
y~321.55e^(−1.39x)
8
6.62
5.25
3.88
2.5
present
reduced
absent
loge(Bit Score)
0.00
0.25
0.50
0.75
1.00
Distance from M. lignano
Bristle Status
present
reduced
absent

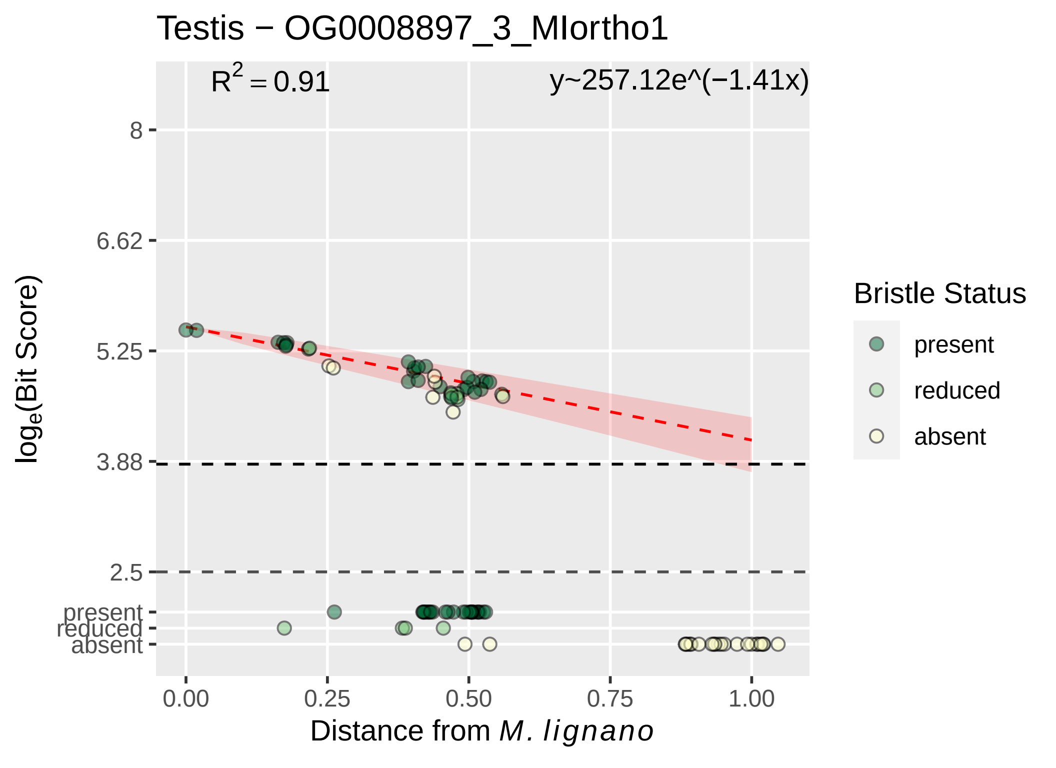

Testis − OG0008897_3_MIortho1
R^2 = 0.91
y~257.12e^(−1.41x)
8
6.62
5.25
3.88
2.5
present
reduced
absent
loge(Bit Score)
0.00
0.25
0.50
0.75
1.00
Distance from M. lignano
Bristle Status
present
reduced
absent

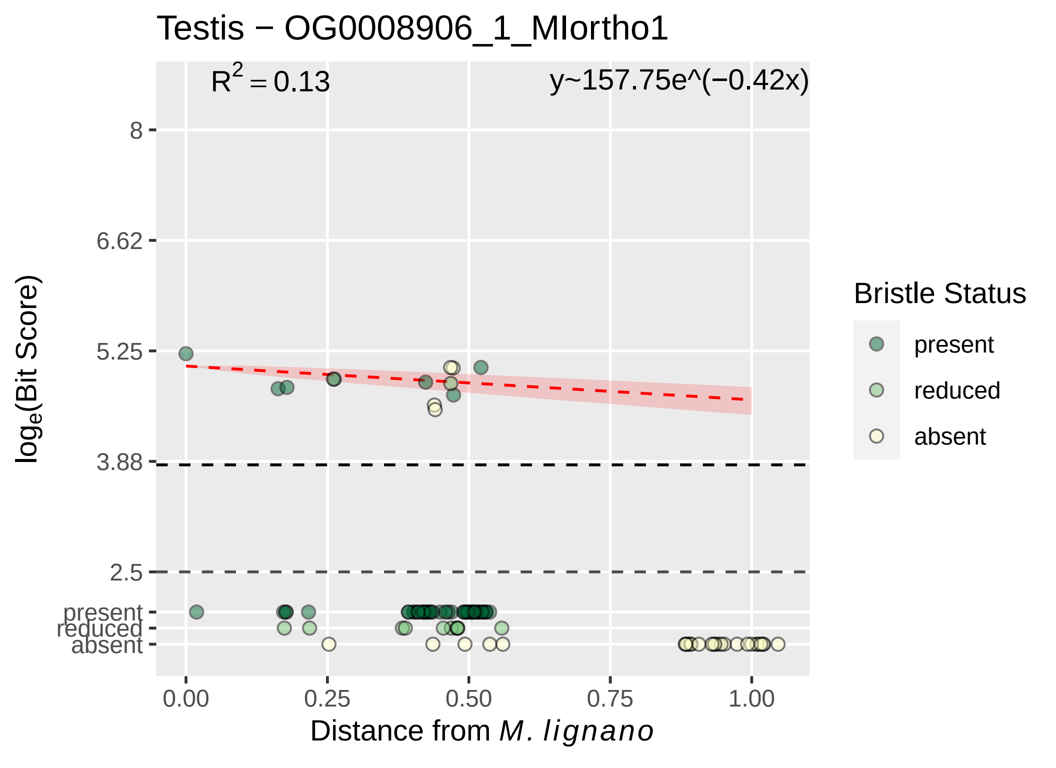
Testis − OG0008906_1_MIortho1
$R^2 = 0.13$
y~157.75e^(−0.42x)
8
6.62
5.25
3.88
2.5
present
reduced
absent
$\log_e$(Bit Score)
0.00
0.25
0.50
0.75
1.00
Distance from *M. lignano*
Bristle Status
present
reduced
absent

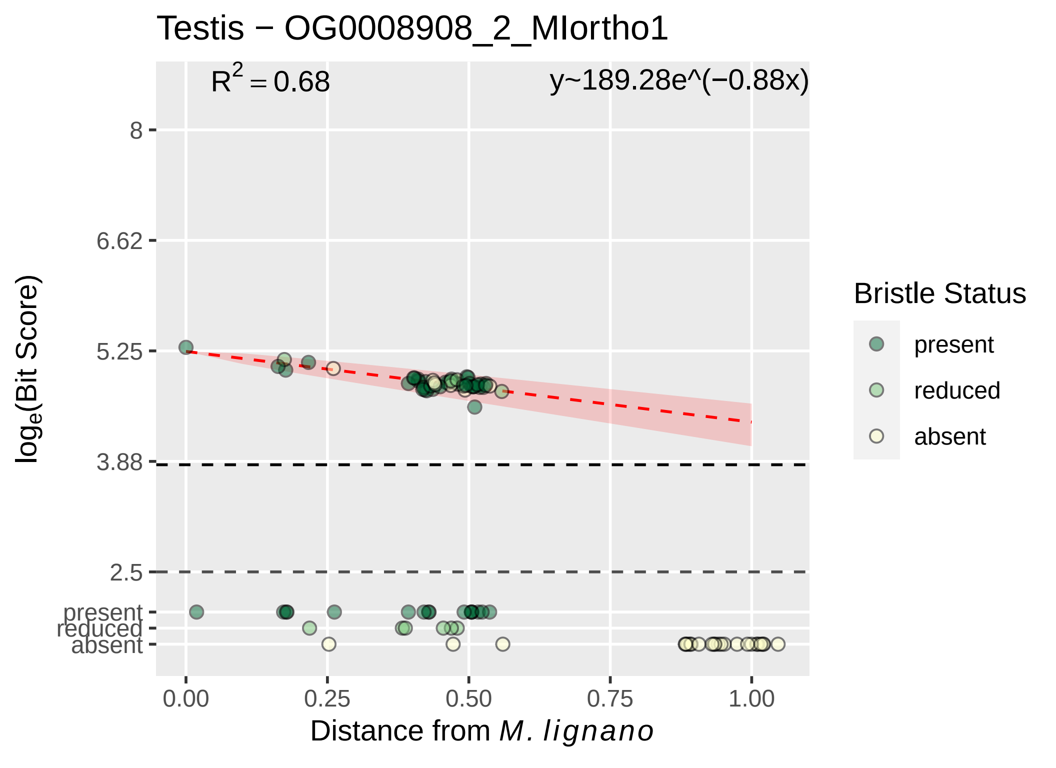

Testis − OG0008908_2_MIortho1
R^2 = 0.68
y~189.28e^(−0.88x)
8
6.62
5.25
3.88
2.5
present
reduced
absent
log_e(Bit Score)
0.00
0.25
0.50
0.75
1.00
Distance from M. lignano
Bristle Status
present
reduced
absent

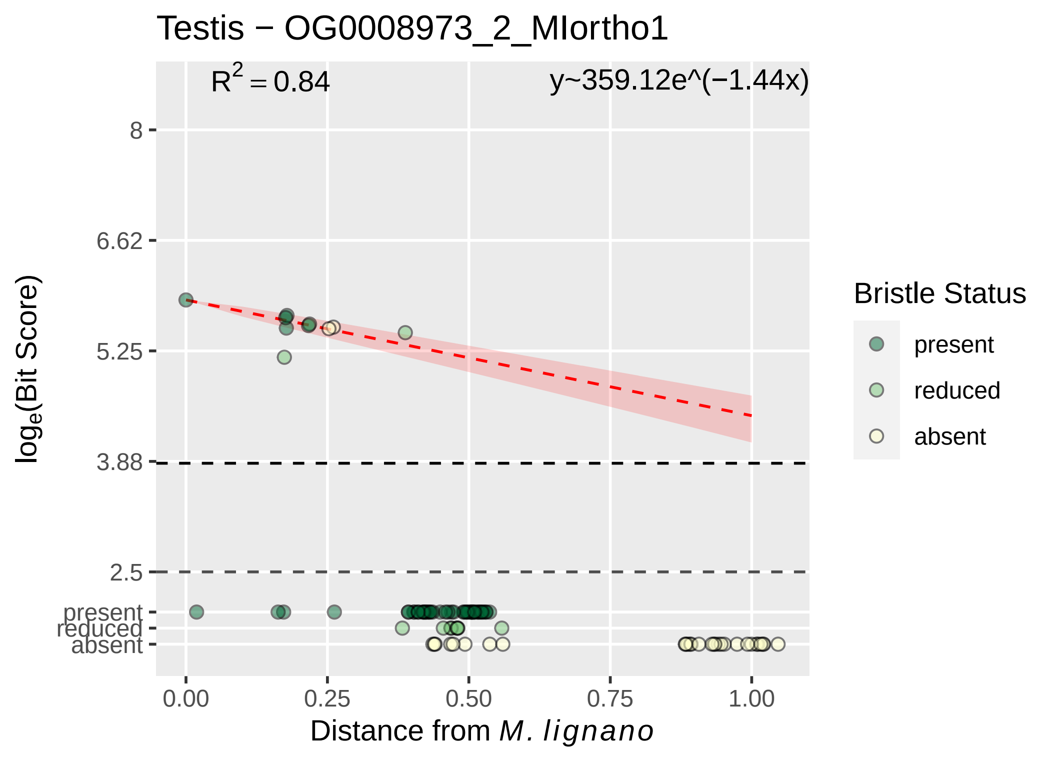

Testis − OG0008973_2_MIortho1
$R^2 = 0.84$
y~359.12e^(−1.44x)
8
6.62
5.25
3.88
2.5
present
reduced
absent
0.00
0.25
0.50
0.75
1.00
Distance from *M. lignano*
$\log_e$(Bit Score)
Bristle Status
present
reduced
absent

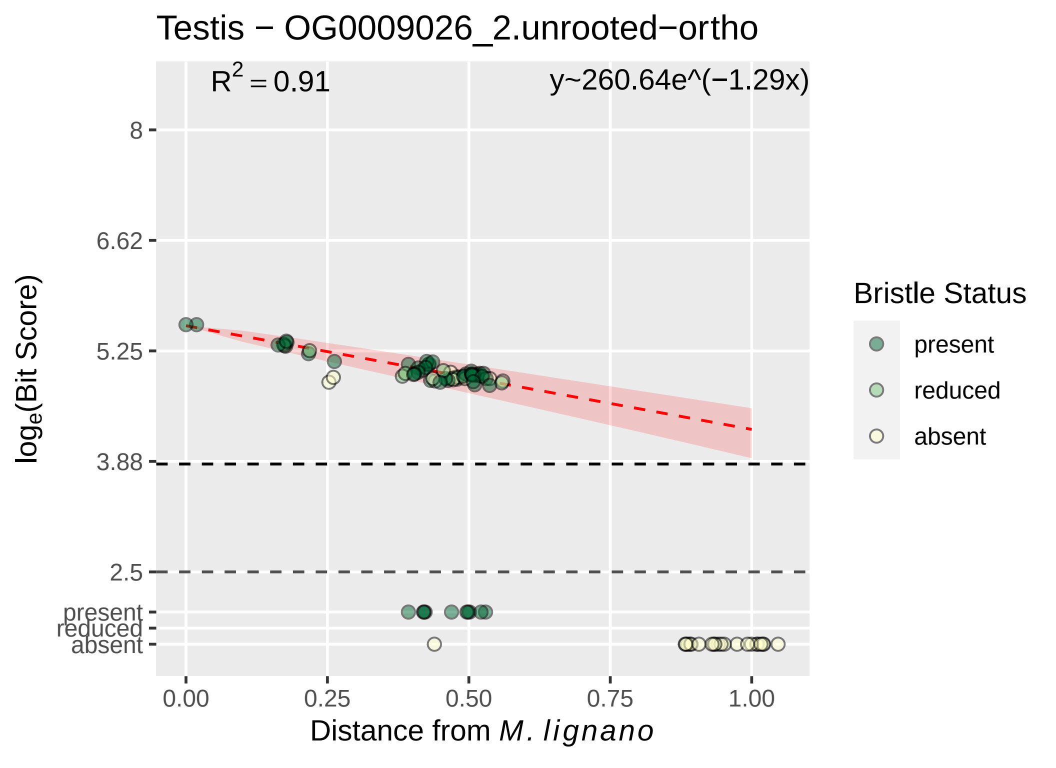

Testis − OG0009026_2.unrooted−ortho
R2 = 0.91
y~260.64e^(−1.29x)
8
6.62
5.25
3.88
2.5
present
reduced
absent
loge(Bit Score)
0.00
0.25
0.50
0.75
1.00
Distance from M. lignano
Bristle Status
present
reduced
absent

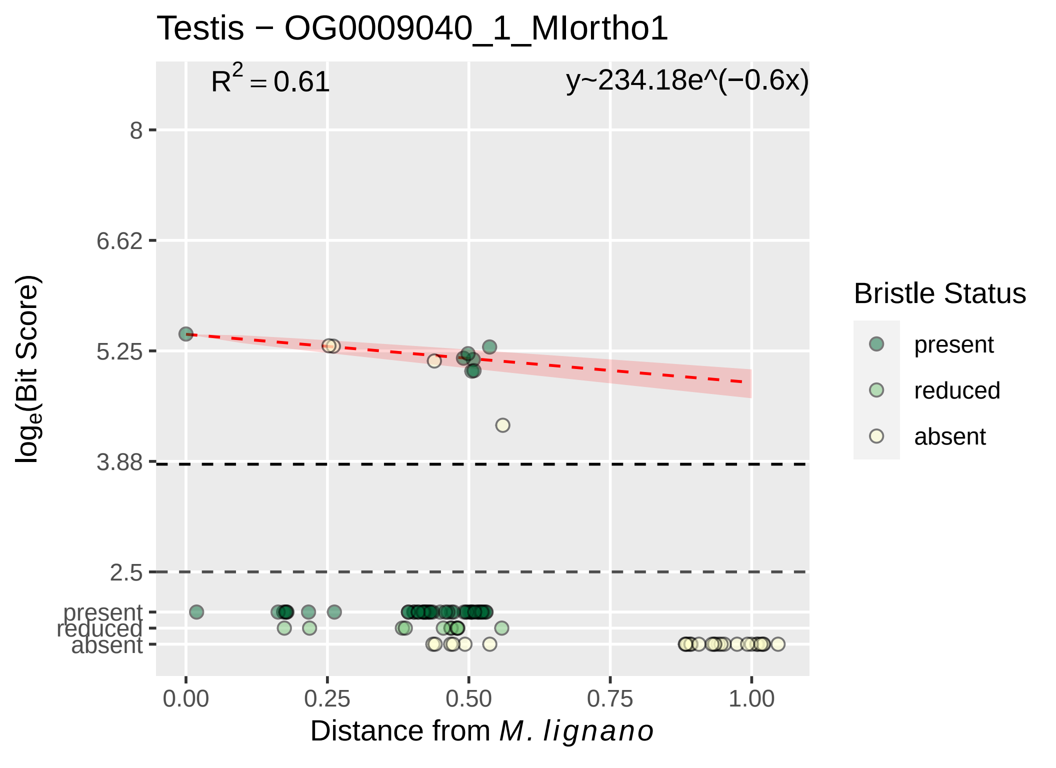

Testis − OG0009040_1_MIortho1
$R^2 = 0.61$
y~234.18e^(−0.6x)
8
6.62
5.25
3.88
2.5
present
reduced
absent
$\log_e$(Bit Score)
0.00
0.25
0.50
0.75
1.00
Distance from *M. lignano*
Bristle Status
present
reduced
absent

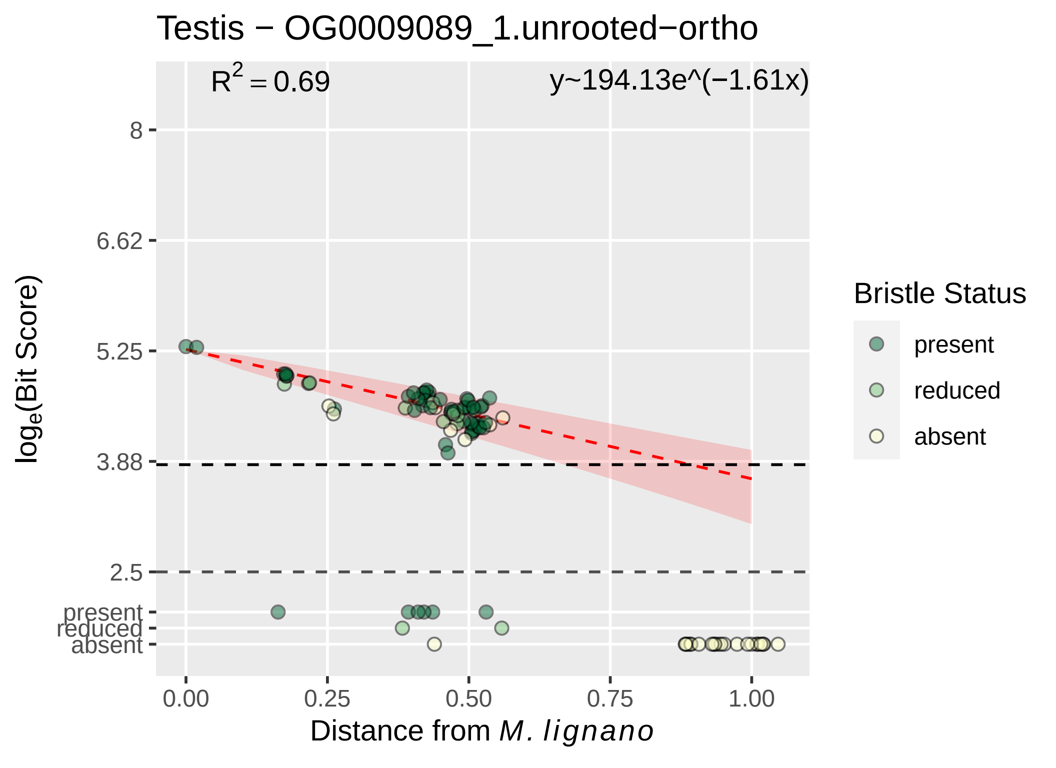

Testis − OG0009089_1.unrooted−ortho
$R^2 = 0.69$
y~194.13e^(−1.61x)
8
6.62
5.25
3.88
2.5
present
reduced
absent
$\log_e$(Bit Score)
0.00
0.25
0.50
0.75
1.00
Distance from *M. lignano*
Bristle Status
present
reduced
absent

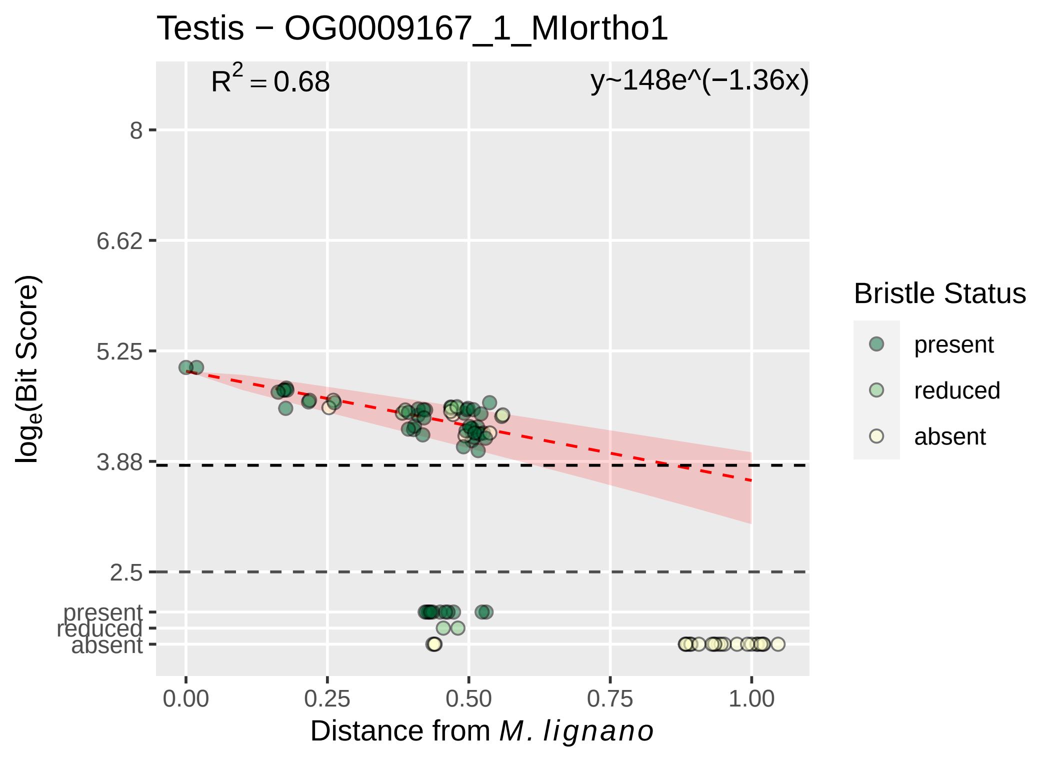

Testis − OG0009167_1_MIortho1
R^2 = 0.68
y~148e^(−1.36x)
8
6.62
5.25
3.88
2.5
present
reduced
absent
0.00
0.25
0.50
0.75
1.00
Distance from M. lignano
log_e(Bit Score)
Bristle Status
present
reduced
absent

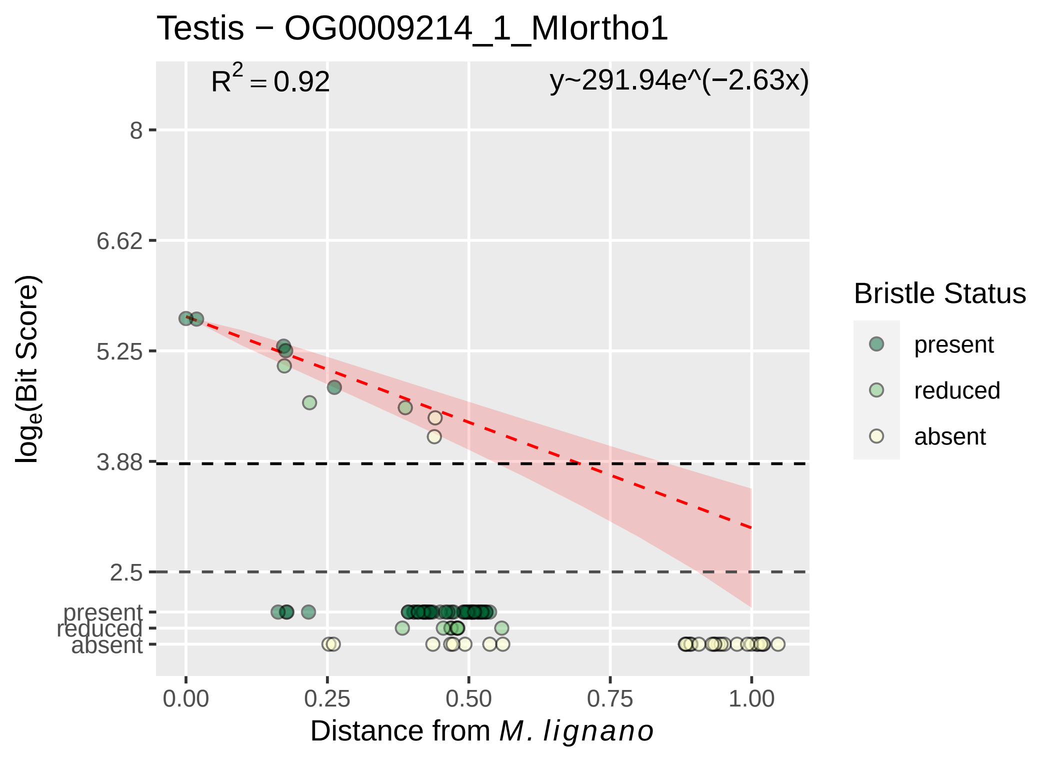

Testis − OG0009214_1_MIortho1
$R^2 = 0.92$
y~291.94e^(−2.63x)
8
6.62
5.25
3.88
2.5
present
reduced
absent
0.00
0.25
0.50
0.75
1.00
Distance from *M. lignano*
$\log_e$(Bit Score)
Bristle Status
present
reduced
absent

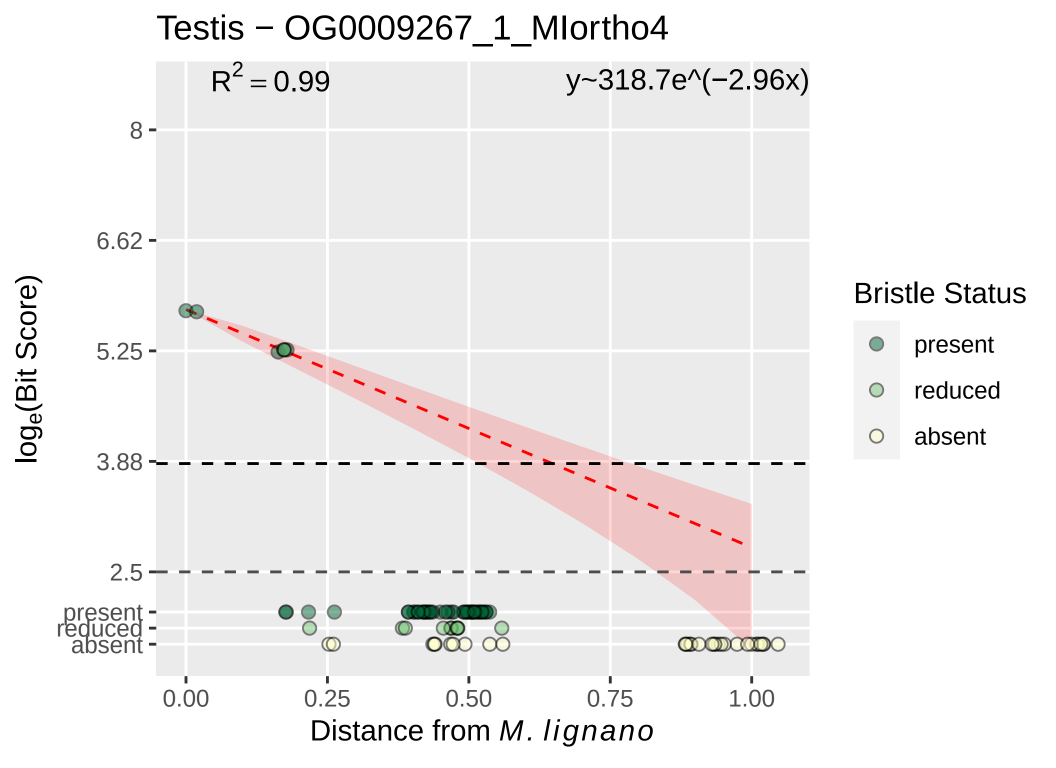

Testis − OG0009267_1_MIortho4
R$^2$ = 0.99
y~318.7e^(−2.96x)
8
6.62
5.25
3.88
2.5
present
reduced
absent
0.00
0.25
0.50
0.75
1.00
log$_e$(Bit Score)
Distance from M. lignano
Bristle Status
present
reduced
absent

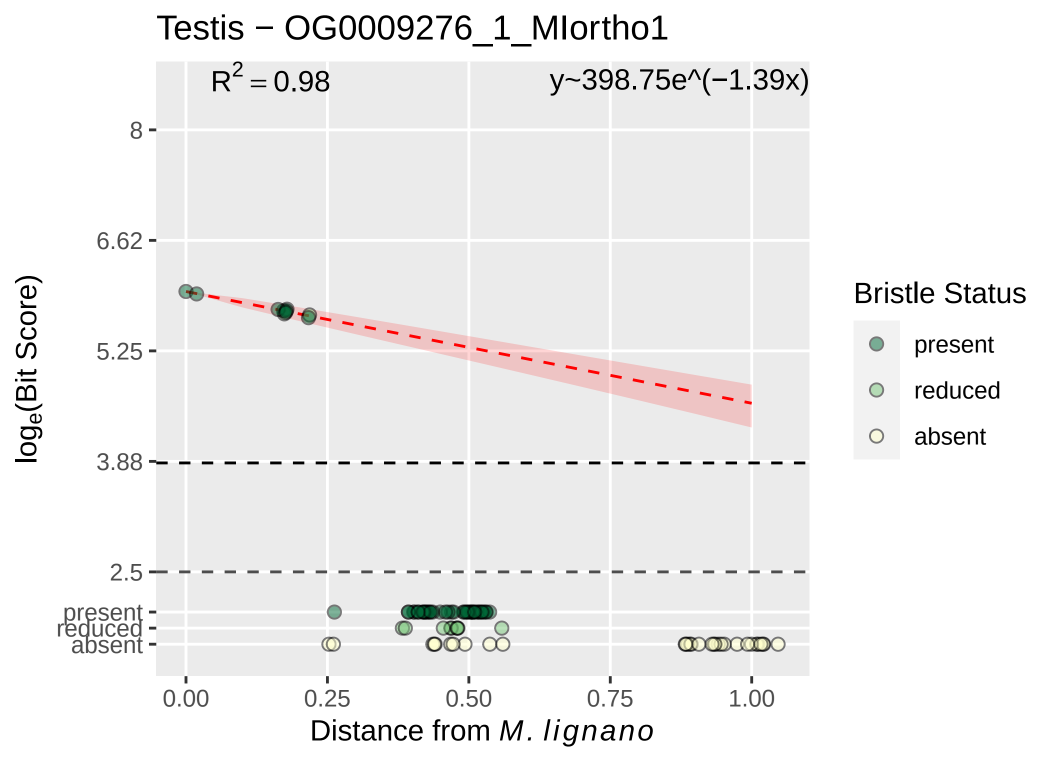

Testis − OG0009276_1_MIortho1
$R^2 = 0.98$
y~398.75e^(−1.39x)
8
6.62
5.25
3.88
2.5
present
reduced
absent
$log_e$(Bit Score)
0.00
0.25
0.50
0.75
1.00
Distance from *M. lignano*
Bristle Status
present
reduced
absent

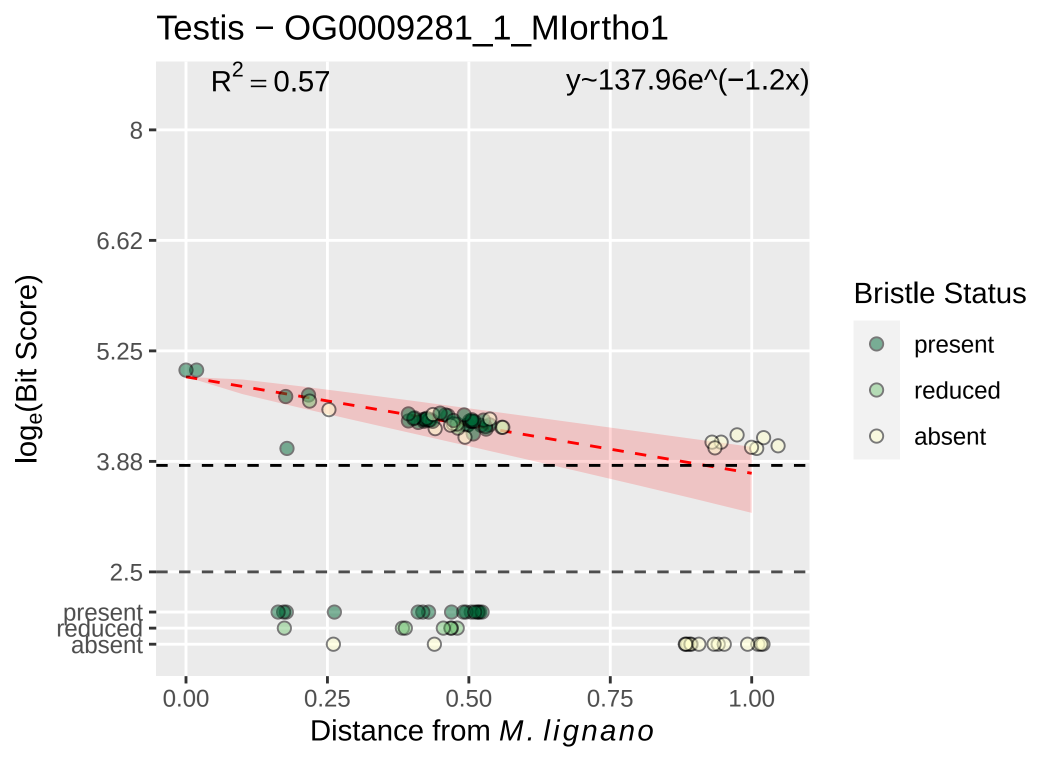

Testis − OG0009281_1_MIortho1
R^2 = 0.57
y~137.96e^(−1.2x)
8
6.62
5.25
3.88
2.5
present
reduced
absent
log_e(Bit Score)
0.00
0.25
0.50
0.75
1.00
Distance from M. lignano
Bristle Status
present
reduced
absent

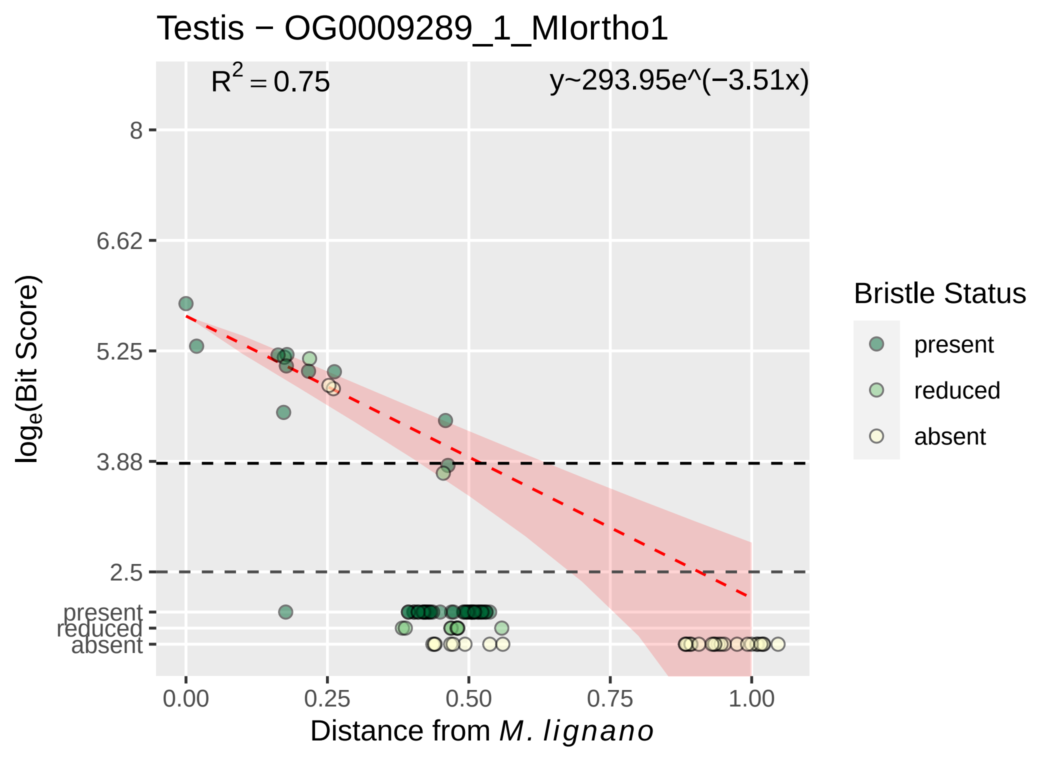
Testis − OG0009289_1_MIortho1
R^2 = 0.75
y~293.95e^(−3.51x)
8
6.62
5.25
3.88
2.5
present
reduced
absent
log_e(Bit Score)
0.00
0.25
0.50
0.75
1.00
Distance from M. lignano
Bristle Status
present
reduced
absent

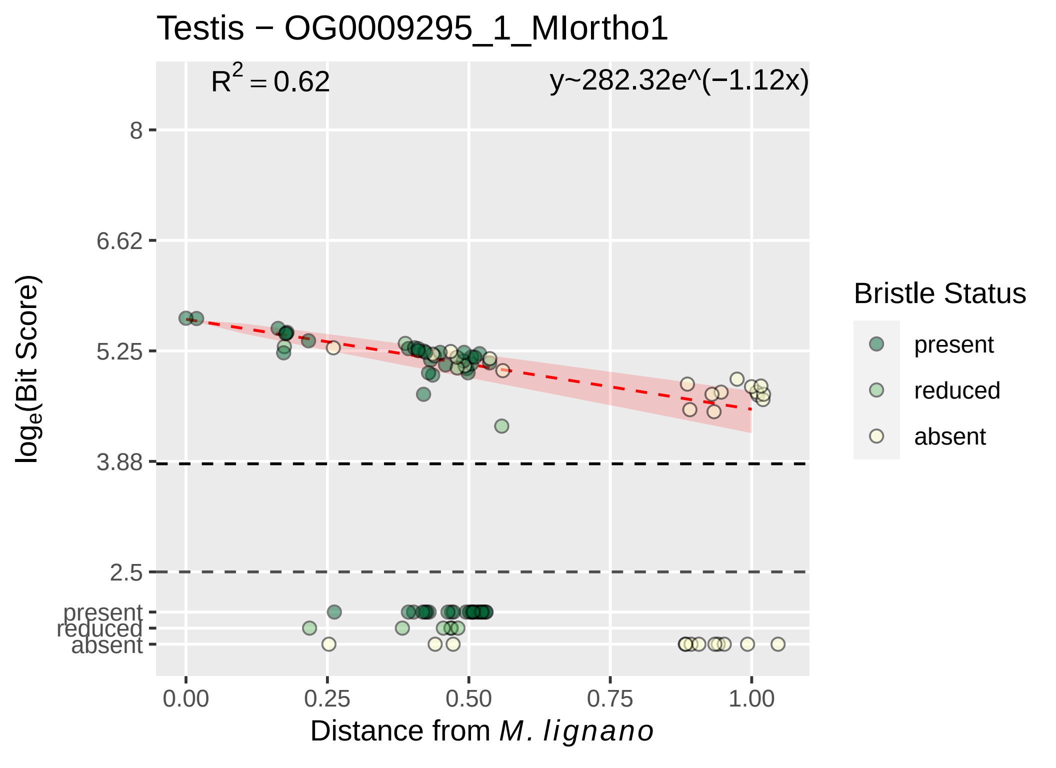
Testis − OG0009295_1_MIortho1
R² = 0.62
y~282.32e^(−1.12x)
8
6.62
5.25
3.88
2.5
present
reduced
absent
loge(Bit Score)
0.00
0.25
0.50
0.75
1.00
Distance from M. lignano
Bristle Status
present
reduced
absent

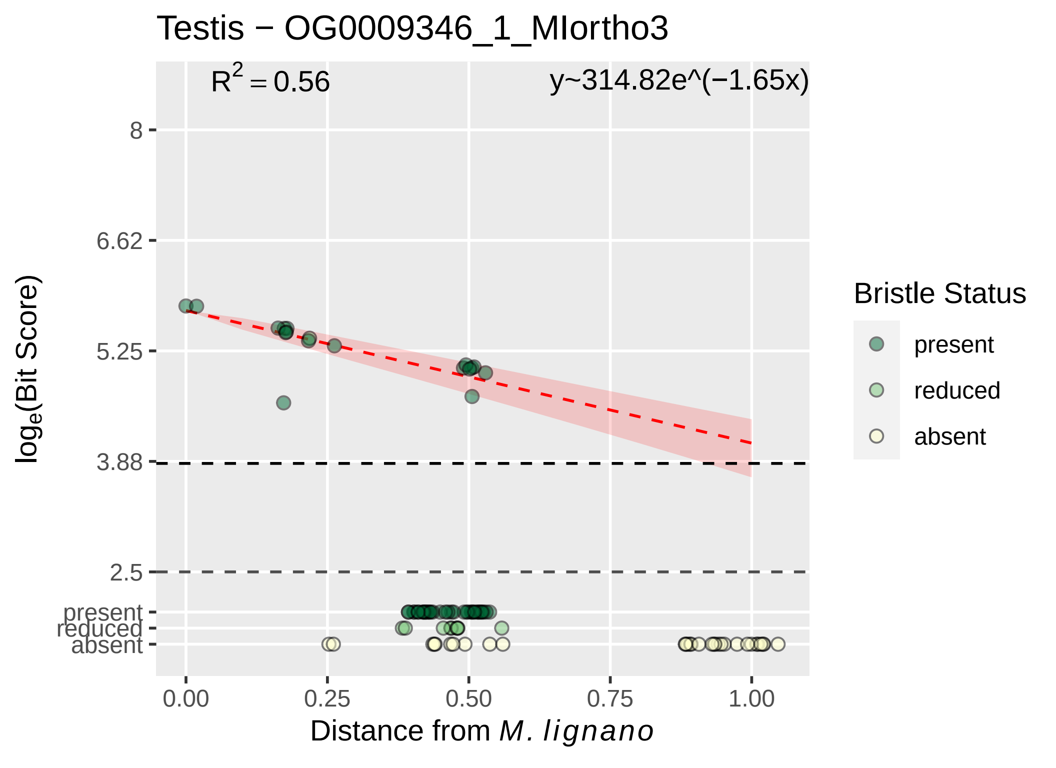

Testis − OG0009346_1_MIortho3
$R^2 = 0.56$
y~314.82e^(−1.65x)
8
6.62
5.25
3.88
2.5
present
reduced
absent
$log_e$(Bit Score)
0.00
0.25
0.50
0.75
1.00
Distance from *M. lignano*
Bristle Status
present
reduced
absent

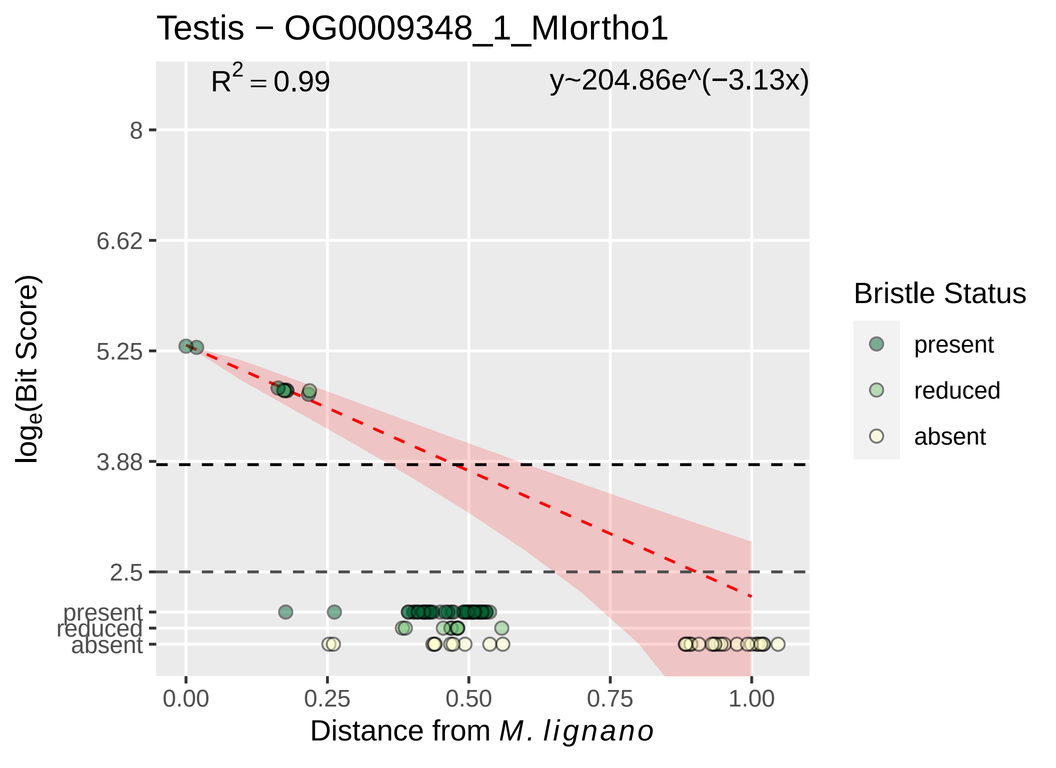

Testis − OG0009348_1_MIortho1
R2 = 0.99
y~204.86e^(−3.13x)
8
6.62
5.25
3.88
2.5
present
reduced
absent
0.00
0.25
0.50
0.75
1.00
loge(Bit Score)
Distance from M. lignano
Bristle Status
present
reduced
absent

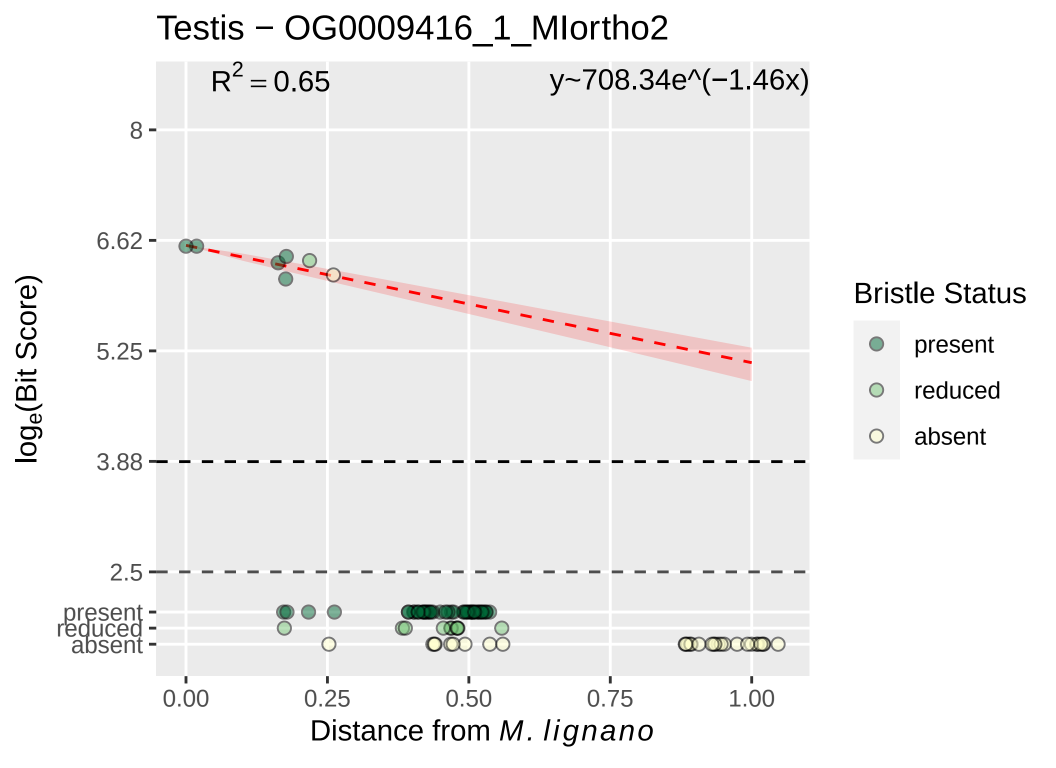
Testis − OG0009416_1_MIortho2
R2 = 0.65
y~708.34e^(−1.46x)
8
6.62
5.25
3.88
2.5
present
reduced
absent
loge(Bit Score)
0.00
0.25
0.50
0.75
1.00
Distance from M. lignano
Bristle Status
present
reduced
absent

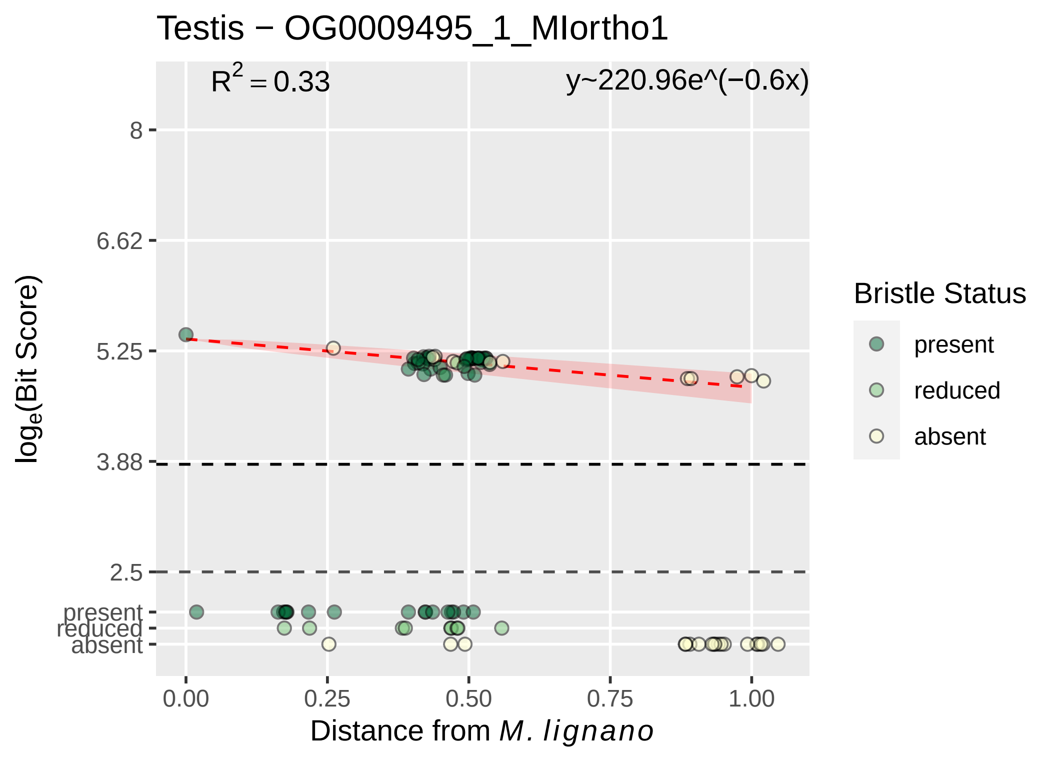

Testis − OG0009495_1_MIortho1
$R^2 = 0.33$
y~220.96e^(−0.6x)
8
6.62
5.25
3.88
2.5
present
reduced
absent
$\log_e$(Bit Score)
0.00
0.25
0.50
0.75
1.00
Distance from *M. lignano*
Bristle Status
present
reduced
absent

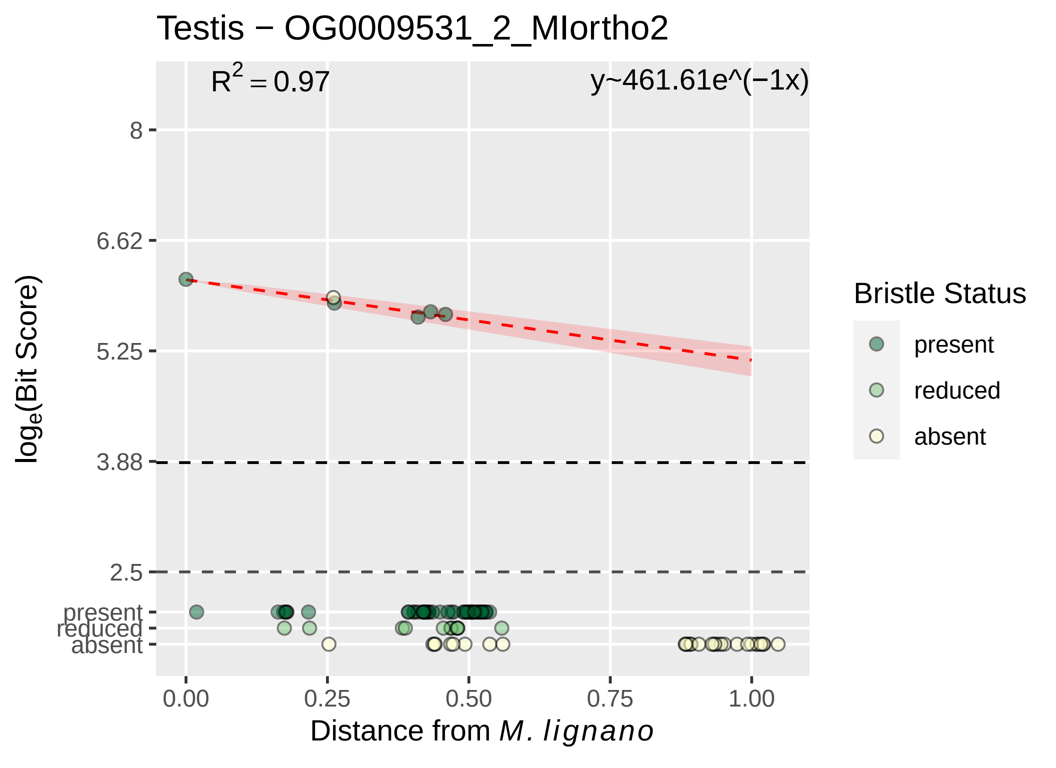
Testis − OG0009531_2_MIortho2
R² = 0.97
y~461.61e^(−1x)
8
6.62
5.25
3.88
2.5
present
reduced
absent
log_e(Bit Score)
0.00
0.25
0.50
0.75
1.00
Distance from M. lignano
Bristle Status
present
reduced
absent

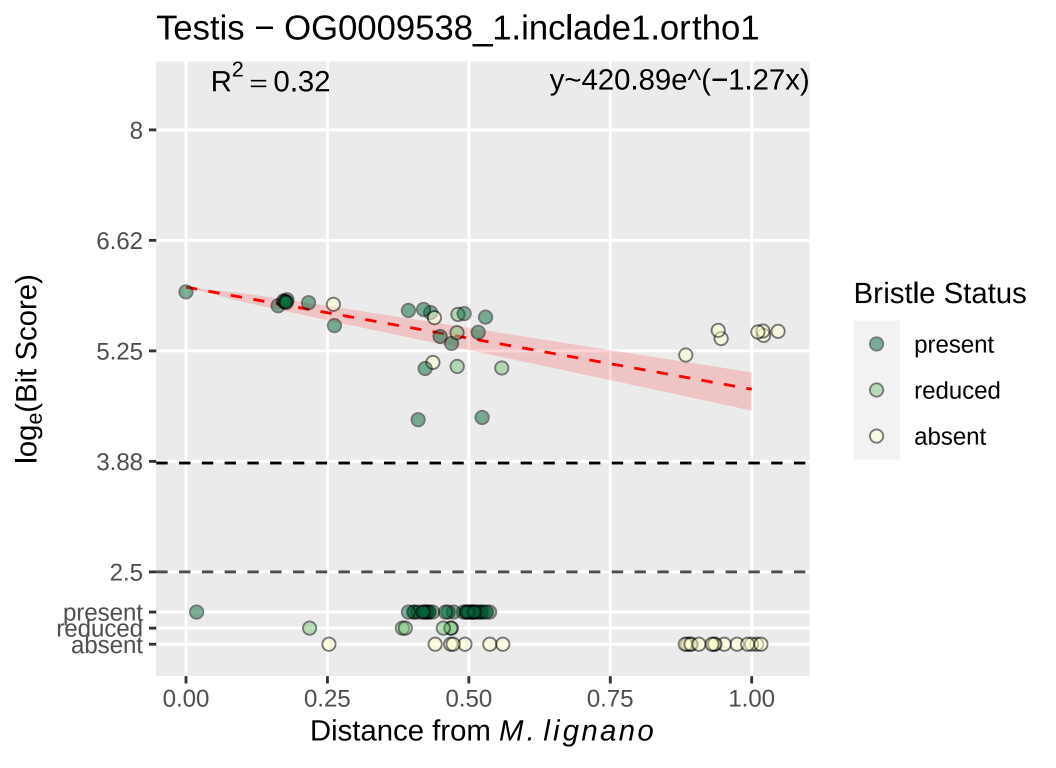
Testis − OG0009538_1.inclade1.ortho1
$R^2 = 0.32$
y~420.89e^(−1.27x)
8
6.62
5.25
3.88
2.5
present
reduced
absent
0.00
0.25
0.50
0.75
1.00
log$_e$(Bit Score)
Distance from *M. lignano*
Bristle Status
present
reduced
absent

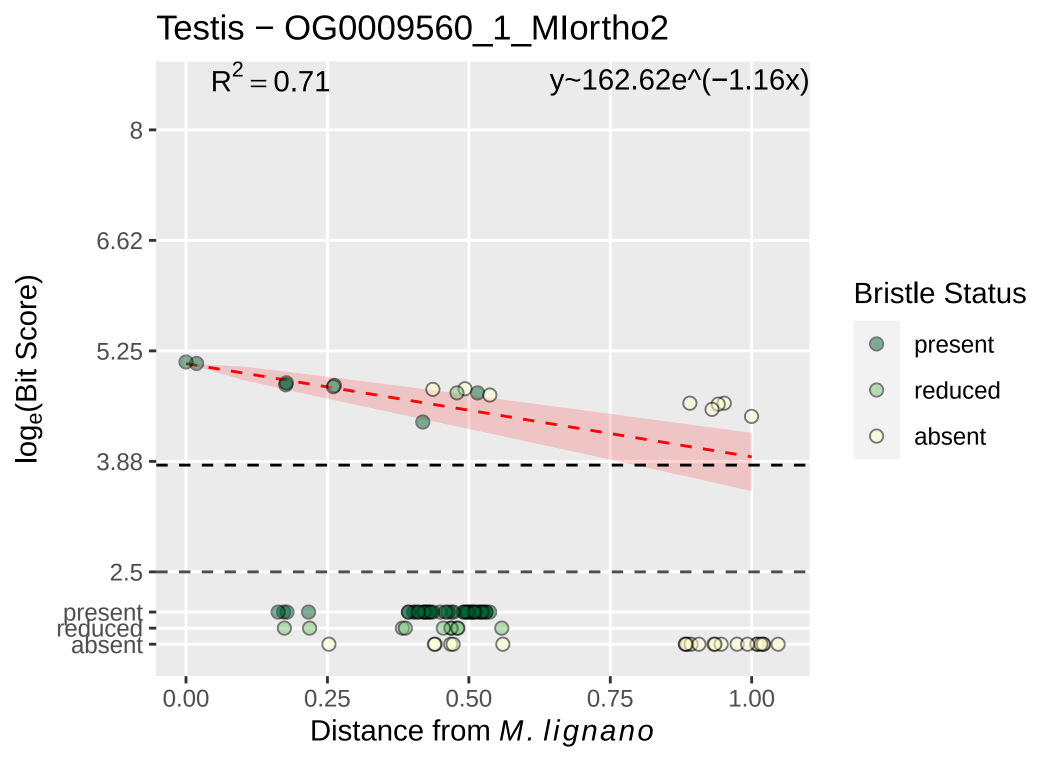

Testis − OG0009560_1_MIortho2
R^2 = 0.71
y~162.62e^(−1.16x)
8
6.62
5.25
3.88
2.5
present
reduced
absent
log_e(Bit Score)
0.00
0.25
0.50
0.75
1.00
Distance from M. lignano
Bristle Status
present
reduced
absent

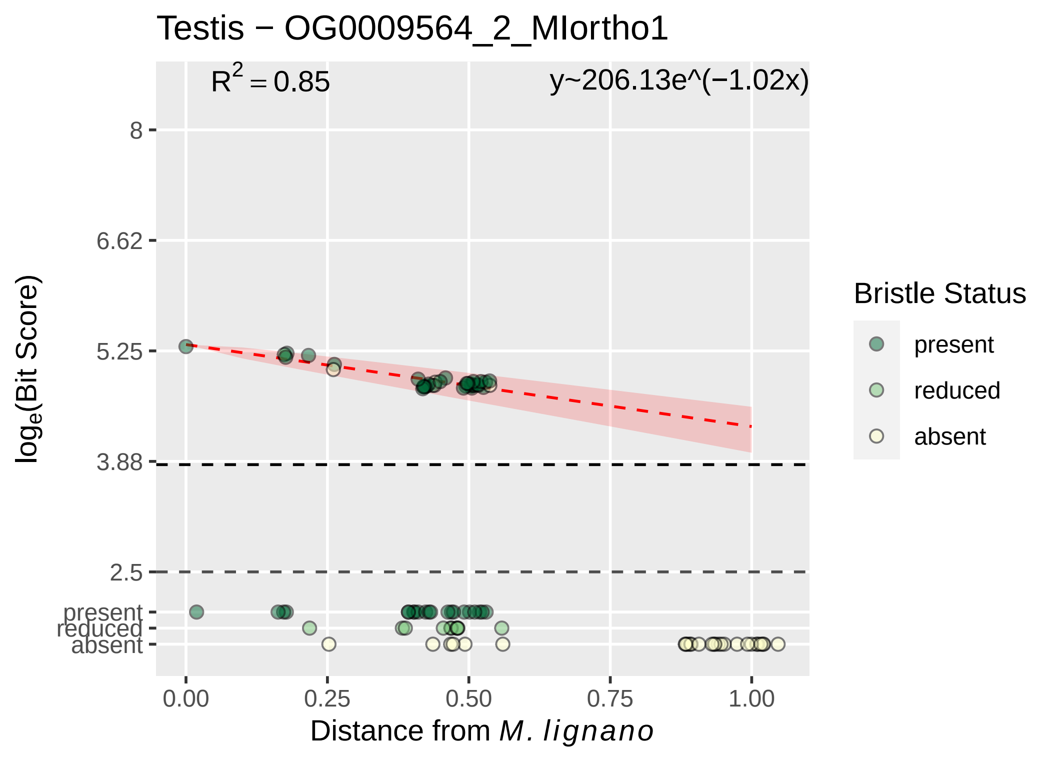

Testis − OG0009564_2_MIortho1
$R^2 = 0.85$
y~206.13e^(−1.02x)
8
6.62
5.25
3.88
2.5
present
reduced
absent
$\log_e$(Bit Score)
0.00
0.25
0.50
0.75
1.00
Distance from *M. lignano*
Bristle Status
present
reduced
absent

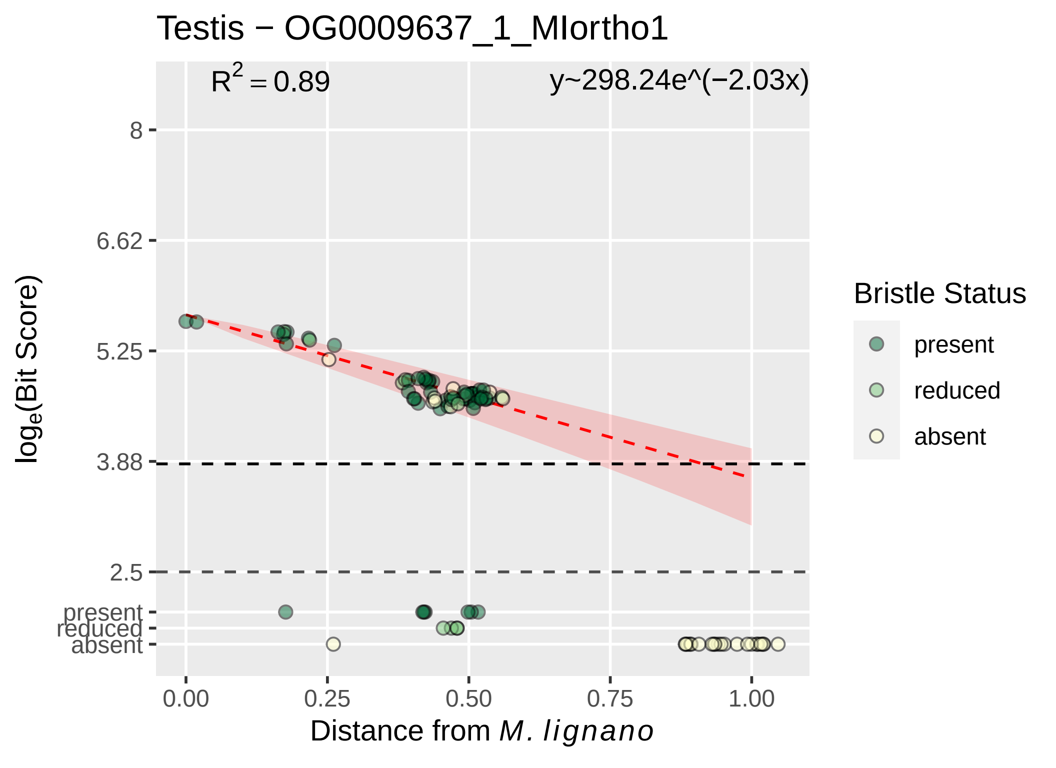
Testis − OG0009637_1_MIortho1
R^2 = 0.89
y~298.24e^(−2.03x)
8
6.62
5.25
3.88
2.5
present
reduced
absent
log_e(Bit Score)
0.00
0.25
0.50
0.75
1.00
Distance from M. lignano
Bristle Status
present
reduced
absent

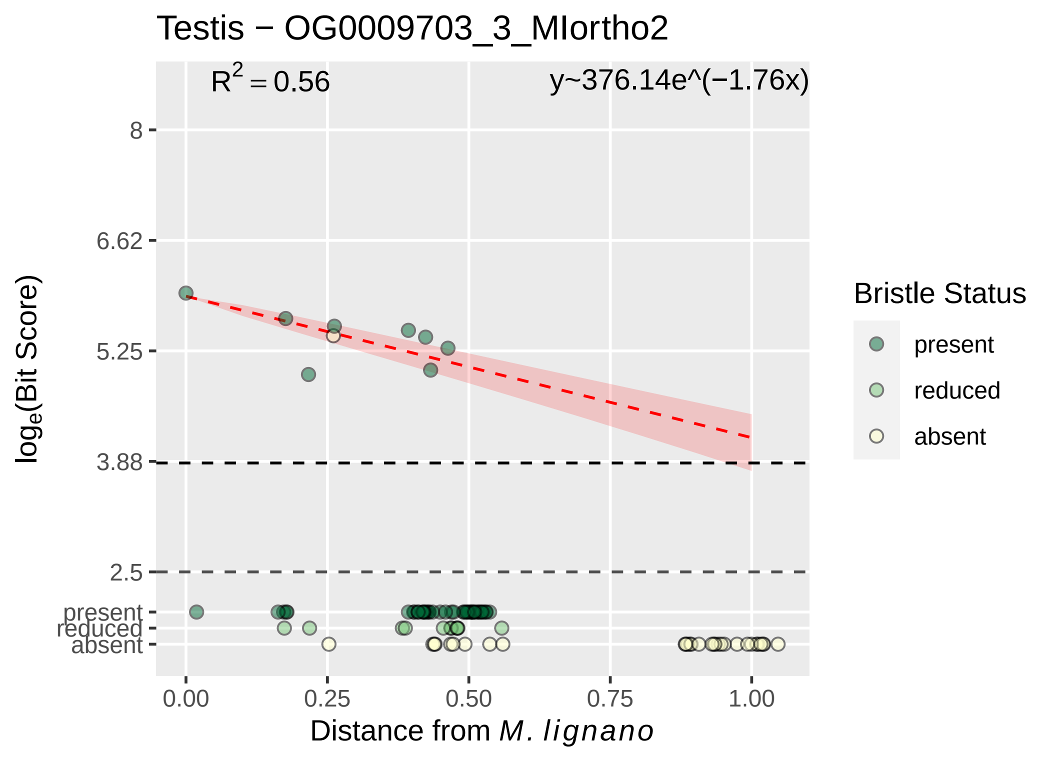

Testis − OG0009703_3_MIortho2
$R^2 = 0.56$
y~376.14e^(−1.76x)
8
6.62
5.25
3.88
2.5
present
reduced
absent
0.00
0.25
0.50
0.75
1.00
log$_e$(Bit Score)
Distance from *M. lignano*
Bristle Status
present
reduced
absent

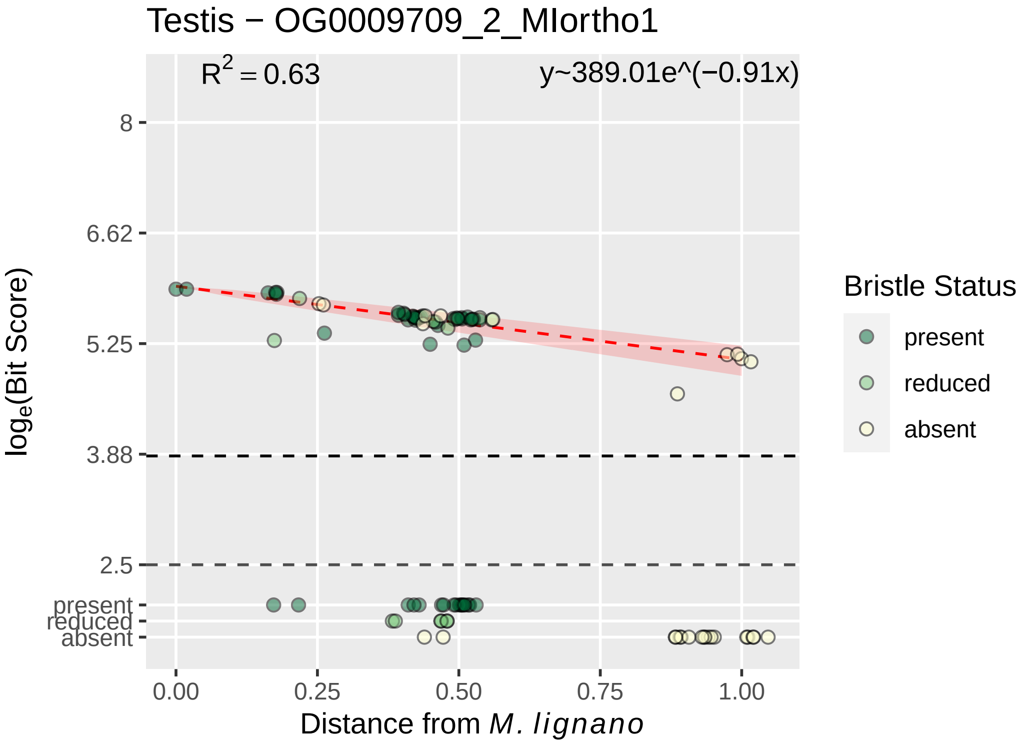
Testis − OG0009709_2_MIortho1
$R^2 = 0.63$
y~389.01e^(−0.91x)
8
6.62
5.25
3.88
2.5
present
reduced
absent
0.00
0.25
0.50
0.75
1.00
Distance from *M. lignano*
$\log_e$(Bit Score)
Bristle Status
present
reduced
absent

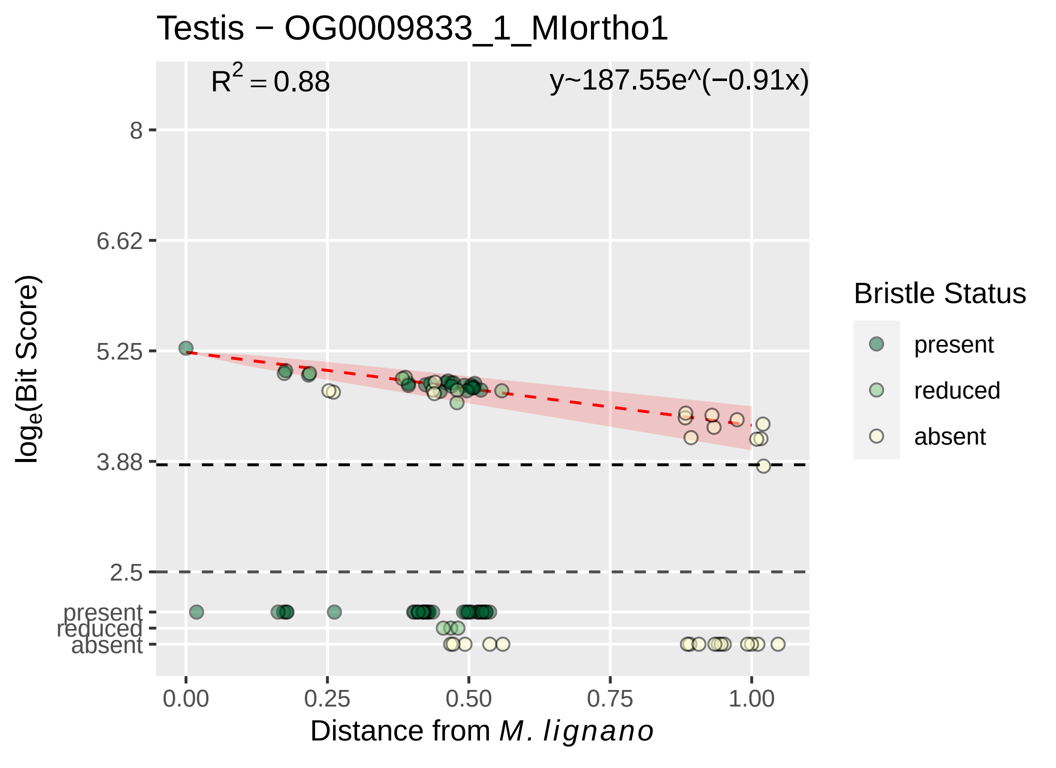

Testis − OG0009833_1_MIortho1
R2 = 0.88
y~187.55e^(−0.91x)
loge(Bit Score)
8
6.62
5.25
3.88
2.5
present
reduced
absent
0.00
0.25
0.50
0.75
1.00
Distance from M. lignano
Bristle Status
present
reduced
absent

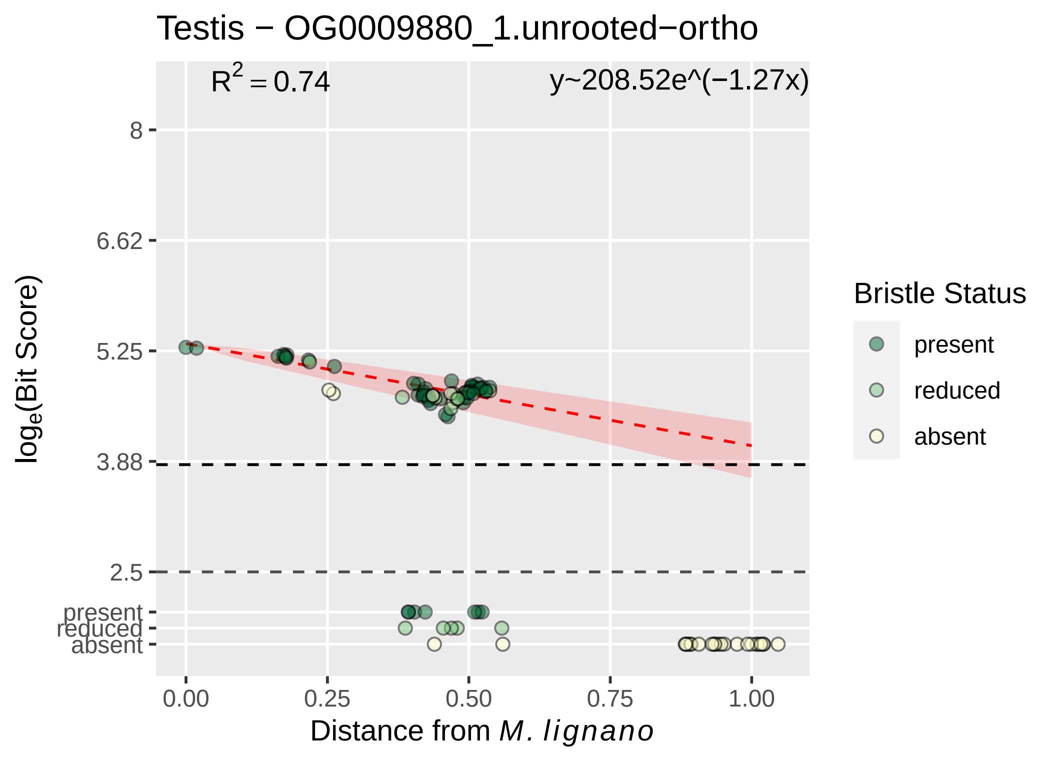

Testis − OG0009880_1.unrooted−ortho
$R^2 = 0.74$
y~208.52e^(−1.27x)
8
6.62
5.25
3.88
2.5
present
reduced
absent
0.00
0.25
0.50
0.75
1.00
Distance from M. lignano
$\log_e$(Bit Score)
Bristle Status
present
reduced
absent

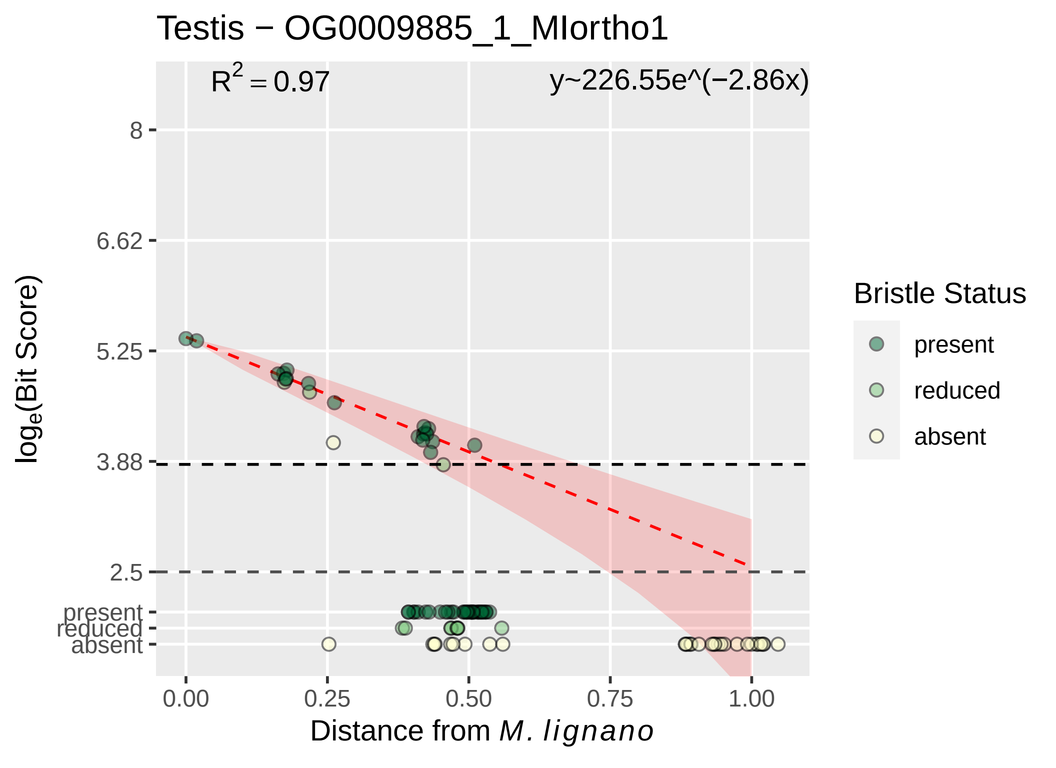
Testis − OG0009885_1_MIortho1
$R^2 = 0.97$
y~226.55e^(−2.86x)
8
6.62
5.25
3.88
2.5
present
reduced
absent
$\log_e$(Bit Score)
0.00
0.25
0.50
0.75
1.00
Distance from *M. lignano*
Bristle Status
present
reduced
absent

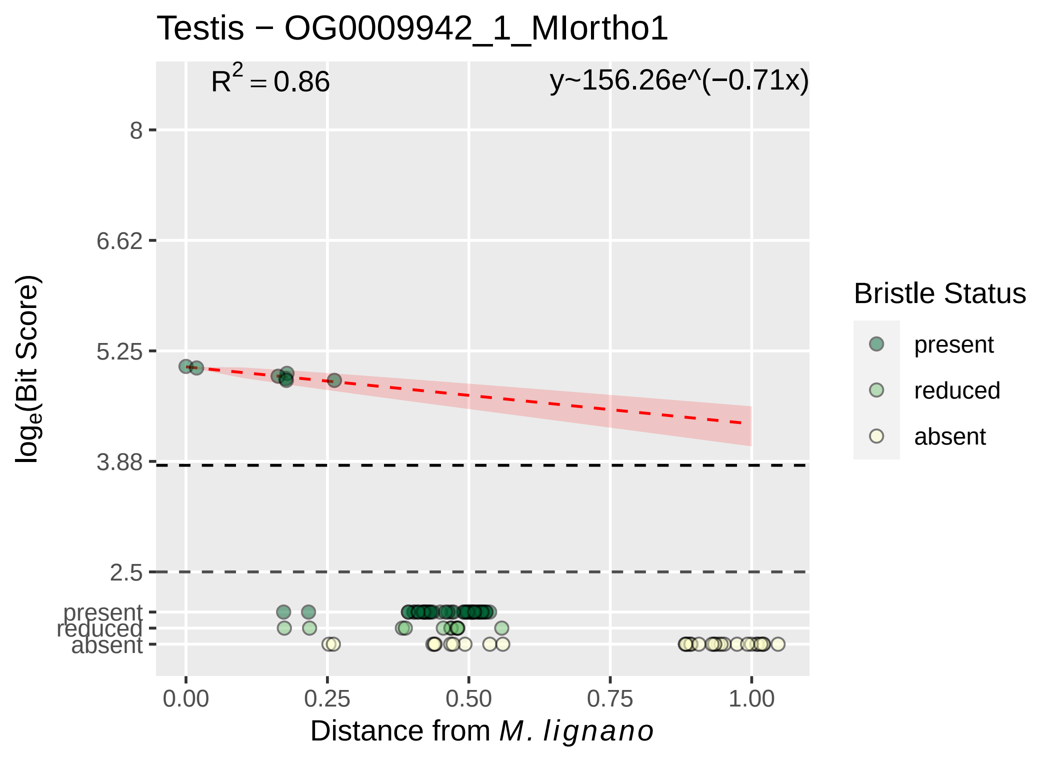

Testis − OG0009942_1_MIortho1
R2 = 0.86
y~156.26e^(−0.71x)
8
6.62
5.25
3.88
2.5
present
reduced
absent
loge(Bit Score)
0.00
0.25
0.50
0.75
1.00
Distance from M. lignano
Bristle Status
present
reduced
absent

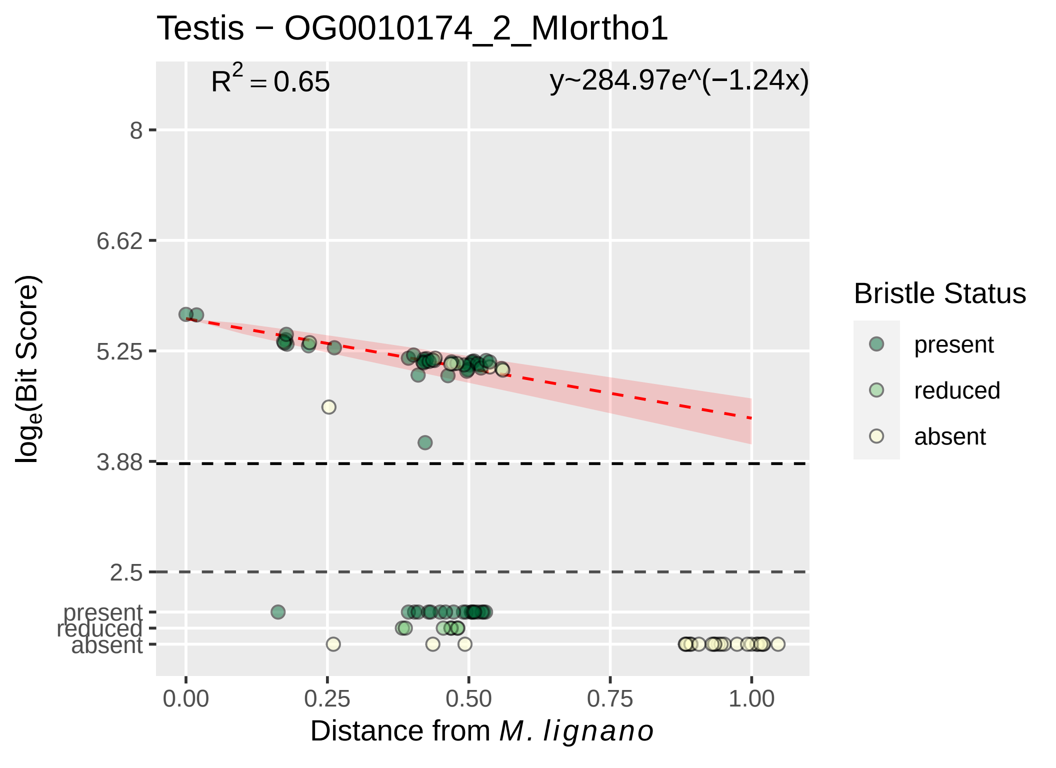

Testis − OG0010174_2_MIortho1
$R^2 = 0.65$
y~284.97e^(−1.24x)
log_e(Bit Score)
8
6.62
5.25
3.88
2.5
present
reduced
absent
0.00
0.25
0.50
0.75
1.00
Distance from M. lignano
Bristle Status
present
reduced
absent

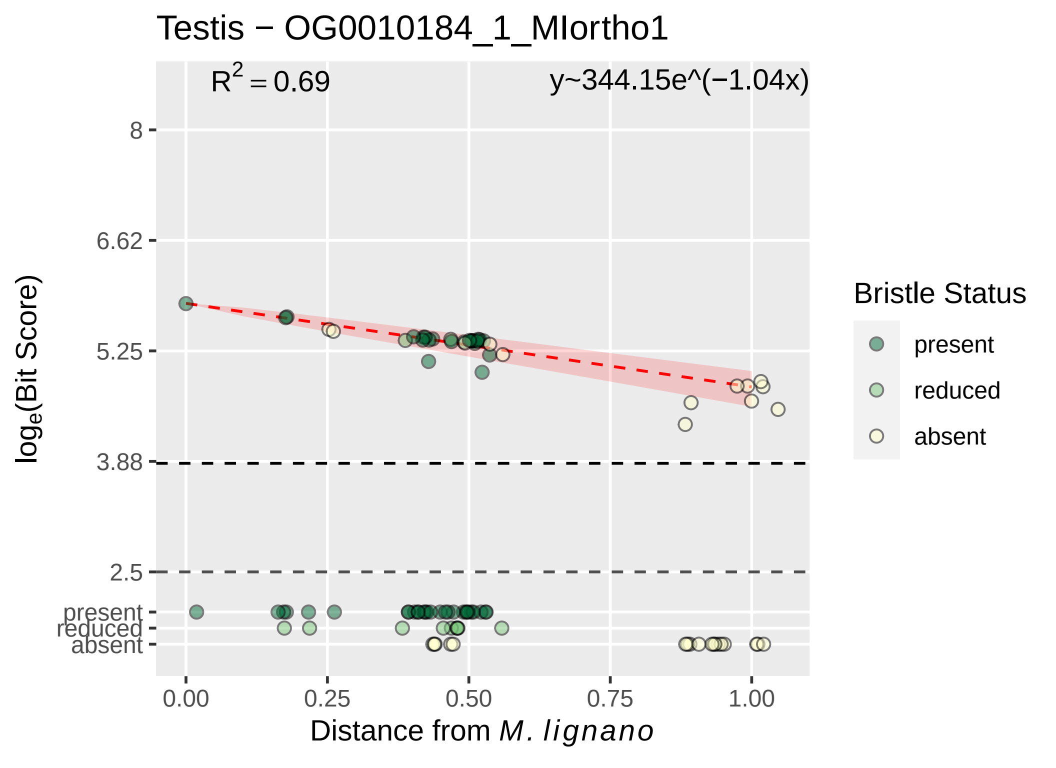

Testis − OG0010184_1_MIortho1
R² = 0.69
y~344.15e^(−1.04x)
8
6.62
5.25
3.88
2.5
present
reduced
absent
log_e(Bit Score)
0.00
0.25
0.50
0.75
1.00
Distance from M. lignano
Bristle Status
present
reduced
absent

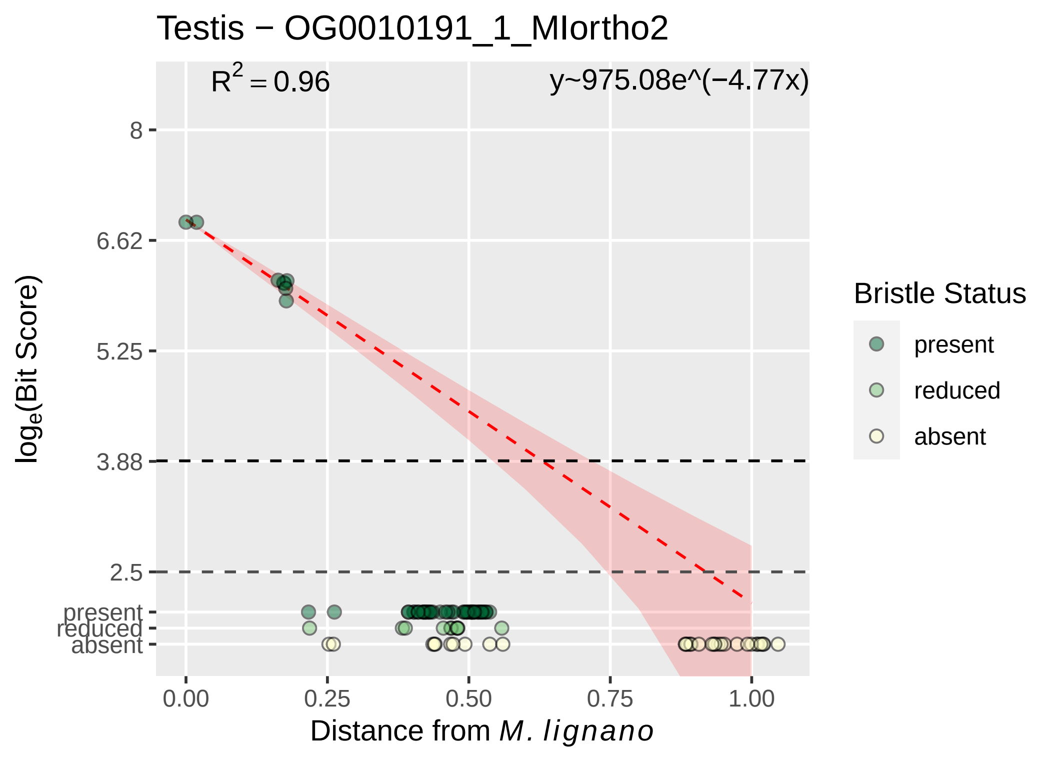

Testis − OG0010191_1_MIortho2
$R^2 = 0.96$
y~975.08e^(−4.77x)
8
6.62
5.25
3.88
2.5
present
reduced
absent
log$_e$(Bit Score)
0.00
0.25
0.50
0.75
1.00
Distance from *M. lignano*
Bristle Status
present
reduced
absent

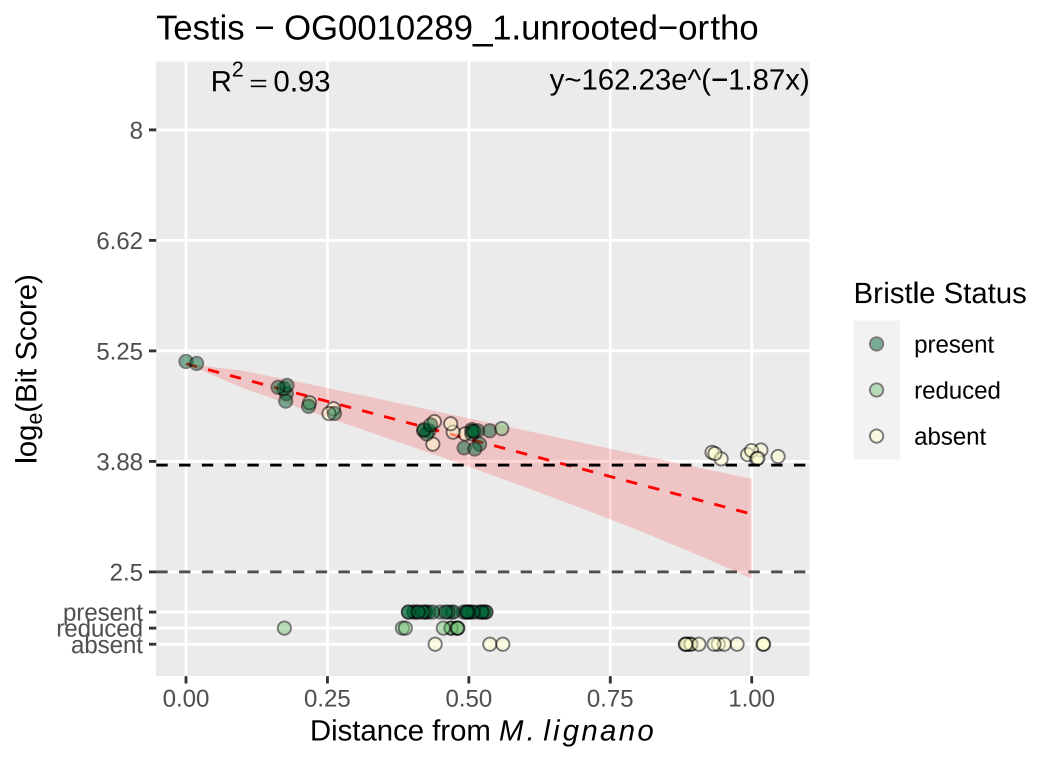

Testis − OG0010289_1.unrooted−ortho
$R^2 = 0.93$
y~162.23e^(−1.87x)
8
6.62
5.25
3.88
2.5
present
reduced
absent
log_e(Bit Score)
0.00
0.25
0.50
0.75
1.00
Distance from M. lignano
Bristle Status
present
reduced
absent

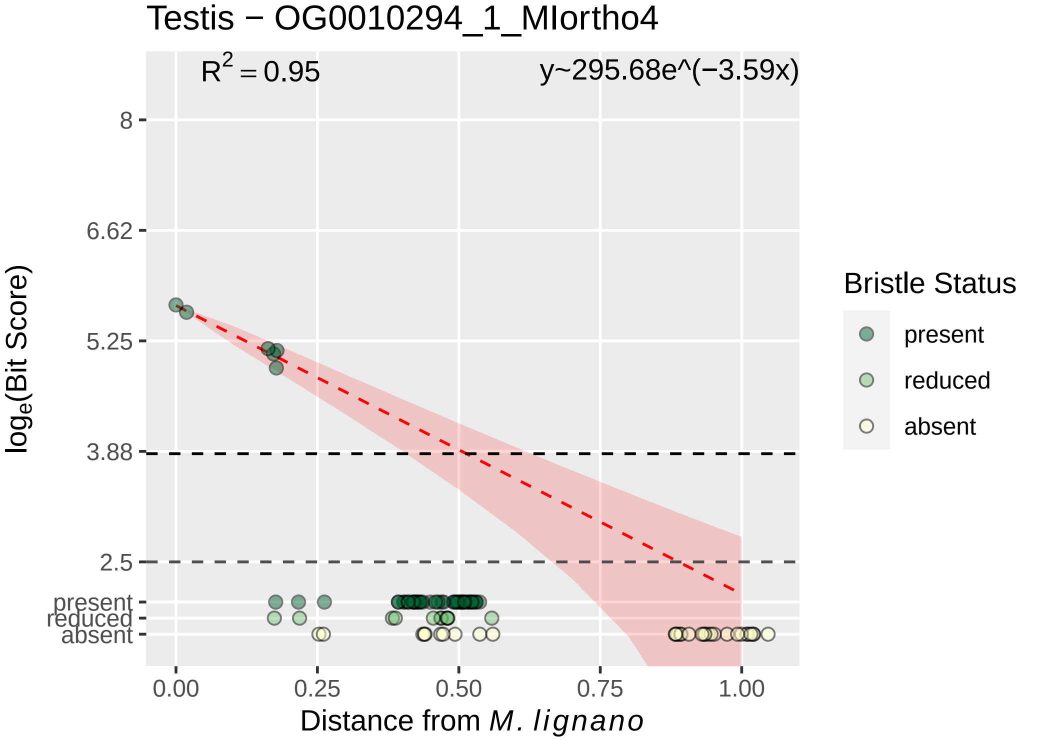

Testis − OG0010294_1_MIortho4
$R^2 = 0.95$
y~295.68e^(−3.59x)
log_e(Bit Score)
8
6.62
5.25
3.88
2.5
present
reduced
absent
0.00
0.25
0.50
0.75
1.00
Distance from M. lignano
Bristle Status
present
reduced
absent

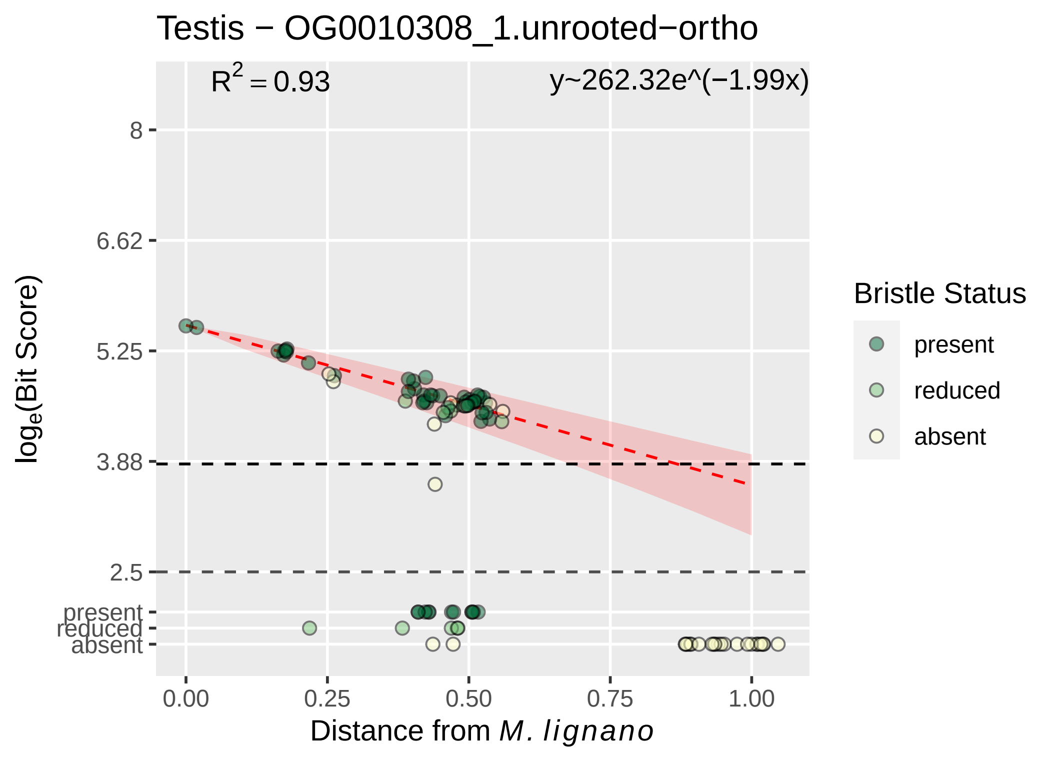
Testis − OG0010308_1.unrooted−ortho
R2 = 0.93
y~262.32e^(−1.99x)
8
6.62
5.25
3.88
2.5
present
reduced
absent
loge(Bit Score)
0.00
0.25
0.50
0.75
1.00
Distance from M. lignano
Bristle Status
present
reduced
absent

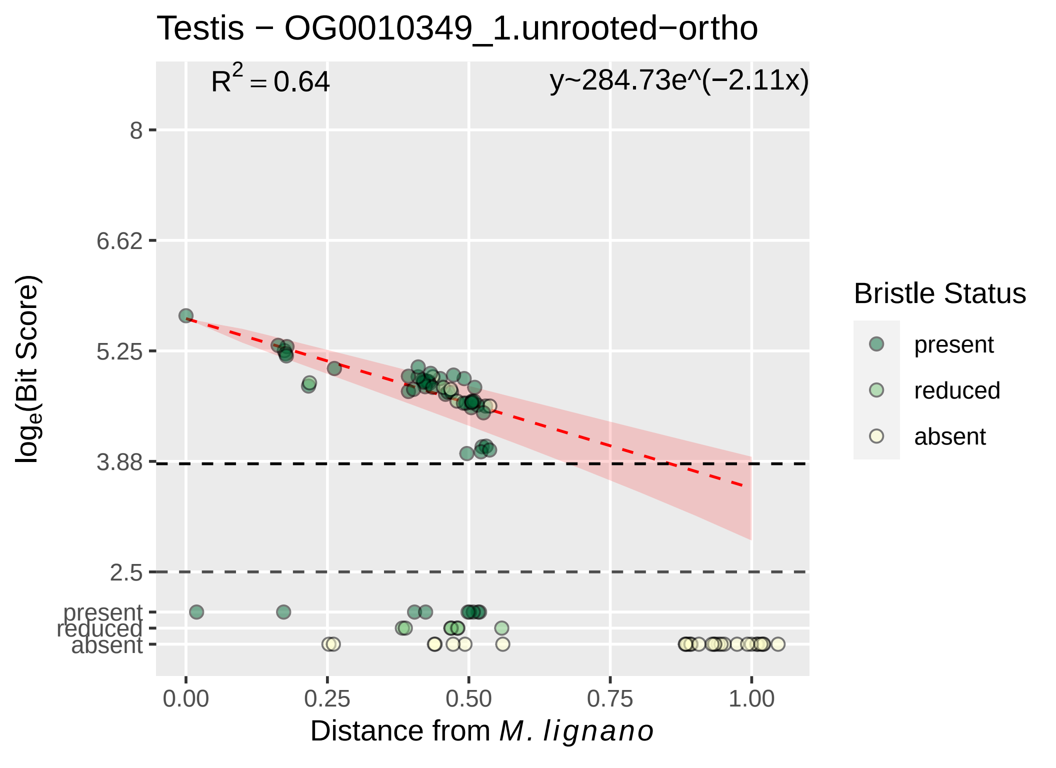

Testis − OG0010349_1.unrooted−ortho
$R^2 = 0.64$
y~284.73e^(−2.11x)
8
6.62
5.25
3.88
2.5
present
reduced
absent
$\log_e$(Bit Score)
0.00
0.25
0.50
0.75
1.00
Distance from *M. lignano*
Bristle Status
present
reduced
absent

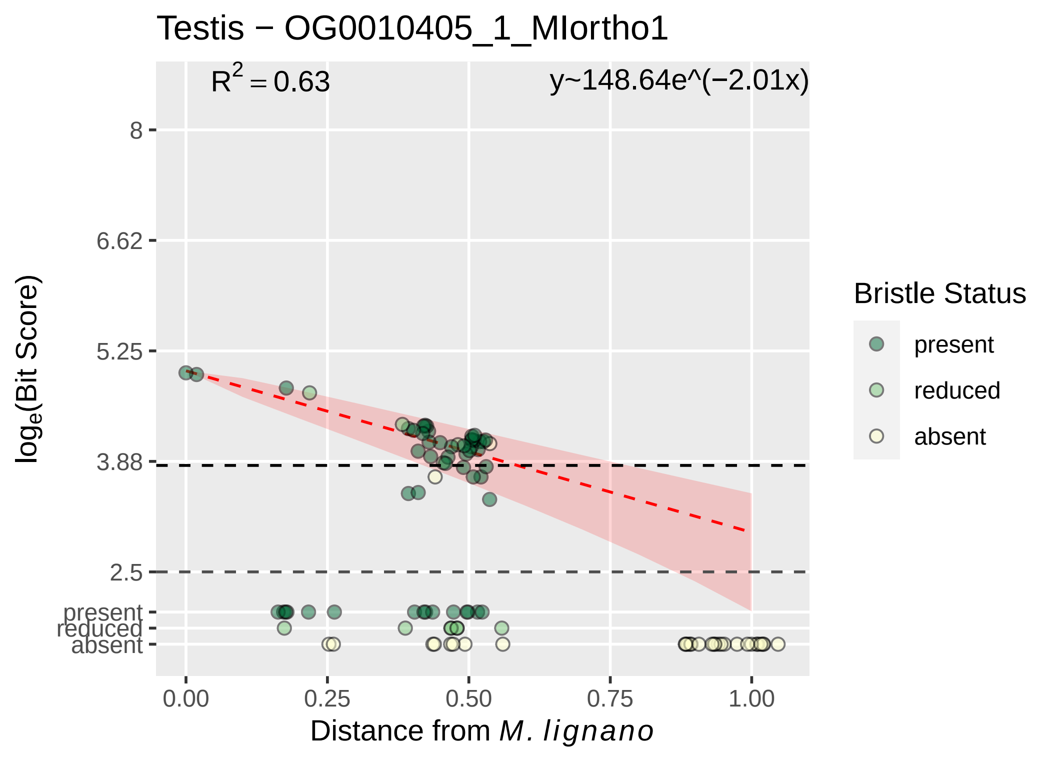

Testis − OG0010405_1_MIortho1
$R^2 = 0.63$
y~148.64e^(−2.01x)
8
6.62
5.25
3.88
2.5
present
reduced
absent
0.00
0.25
0.50
0.75
1.00
Distance from *M. lignano*
$\log_e$(Bit Score)
Bristle Status
present
reduced
absent

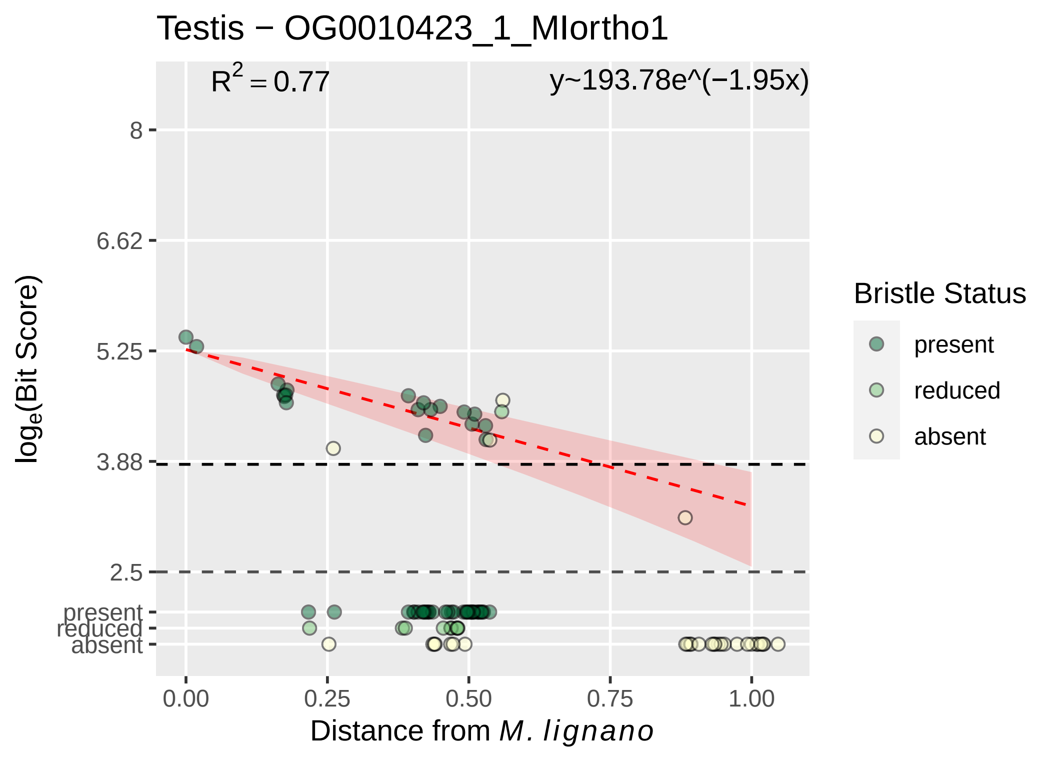

Testis − OG0010423_1_MIortho1
R² = 0.77
y~193.78e^(−1.95x)
8
6.62
5.25
3.88
2.5
present
reduced
absent
log_e(Bit Score)
0.00
0.25
0.50
0.75
1.00
Distance from M. lignano
Bristle Status
present
reduced
absent

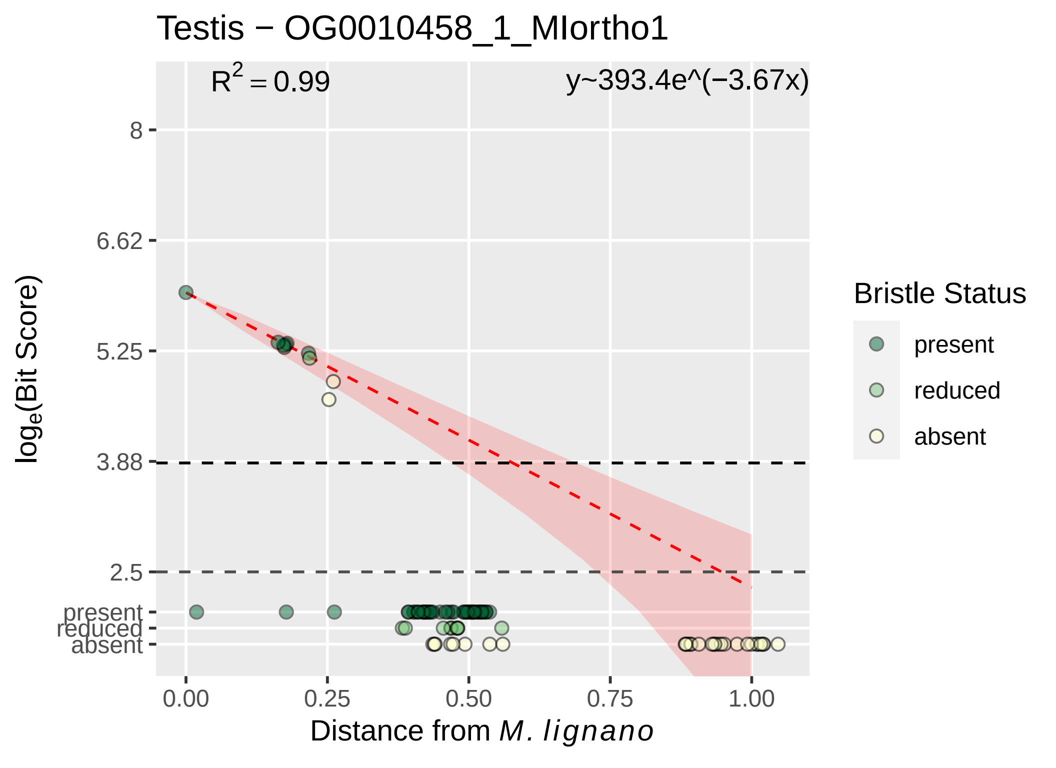

Testis − OG0010458_1_MIortho1
$R^2 = 0.99$
y~393.4e^(−3.67x)
8
6.62
5.25
3.88
2.5
present
reduced
absent
0.00
0.25
0.50
0.75
1.00
log$_e$(Bit Score)
Distance from *M. lignano*
Bristle Status
present
reduced
absent

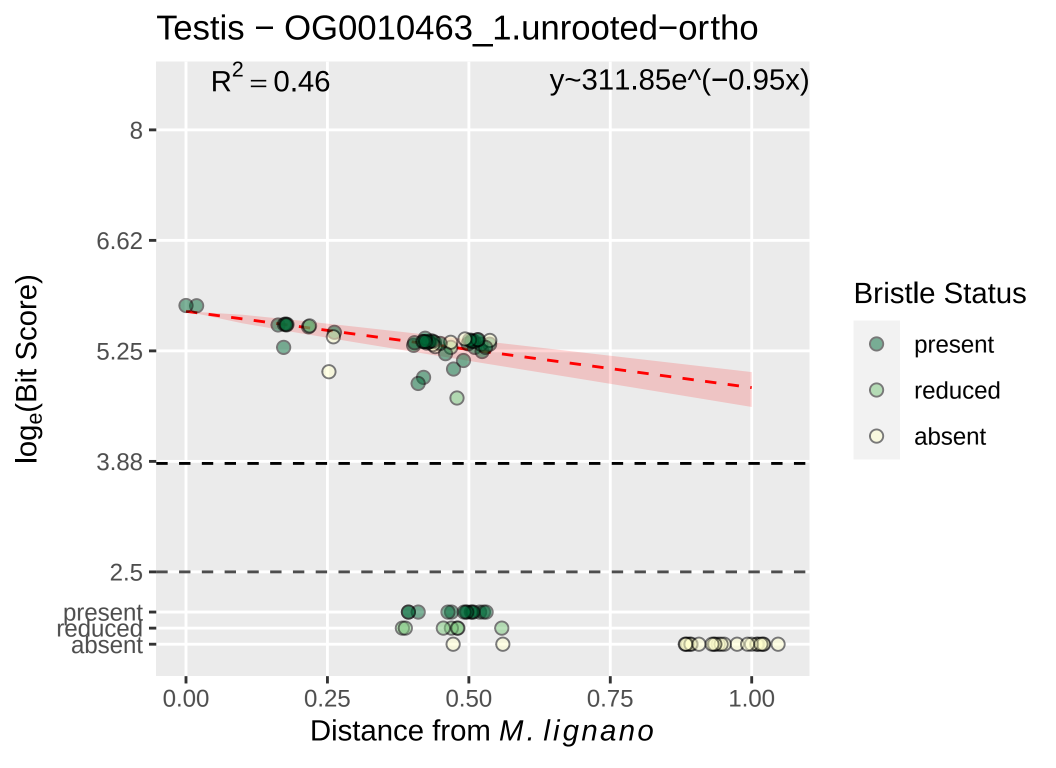
Testis − OG0010463_1.unrooted−ortho
$R^2 = 0.46$
y~311.85e^(−0.95x)
8
6.62
5.25
3.88
2.5
present
reduced
absent
$log_e$(Bit Score)
0.00
0.25
0.50
0.75
1.00
Distance from *M. lignano*
Bristle Status
present
reduced
absent

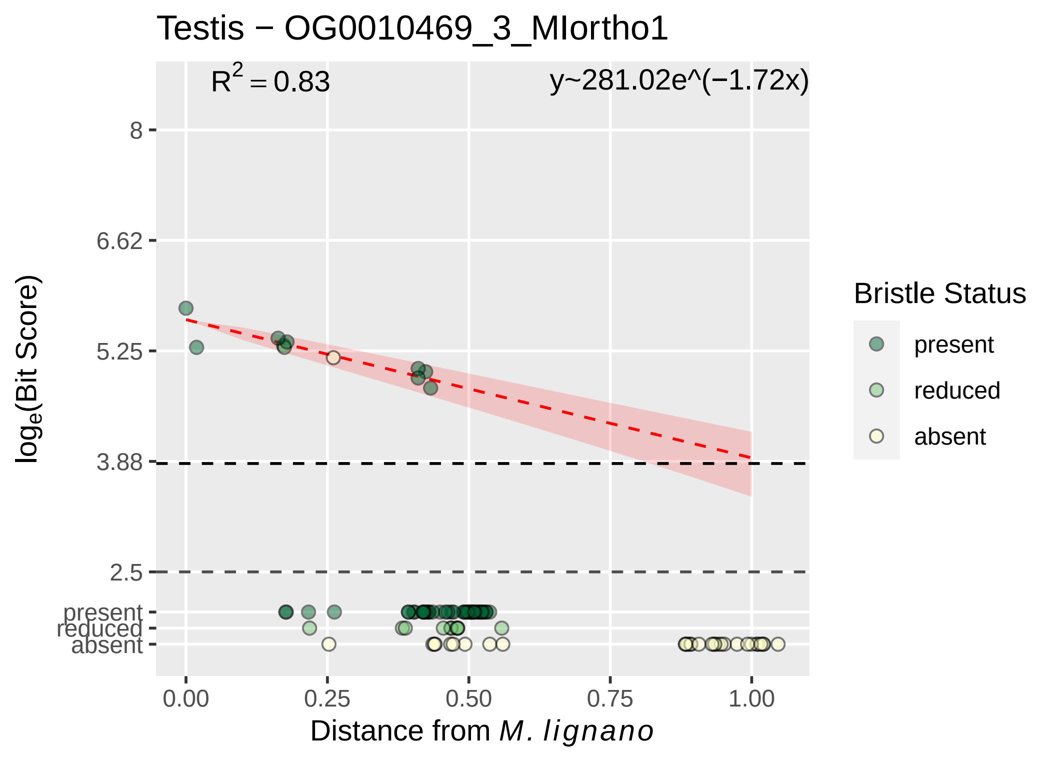

Testis − OG0010469_3_MIortho1
R2 = 0.83
y~281.02e^(−1.72x)
8
6.62
5.25
3.88
2.5
present
reduced
absent
0.00
0.25
0.50
0.75
1.00
Distance from M. lignano
loge(Bit Score)
Bristle Status
present
reduced
absent

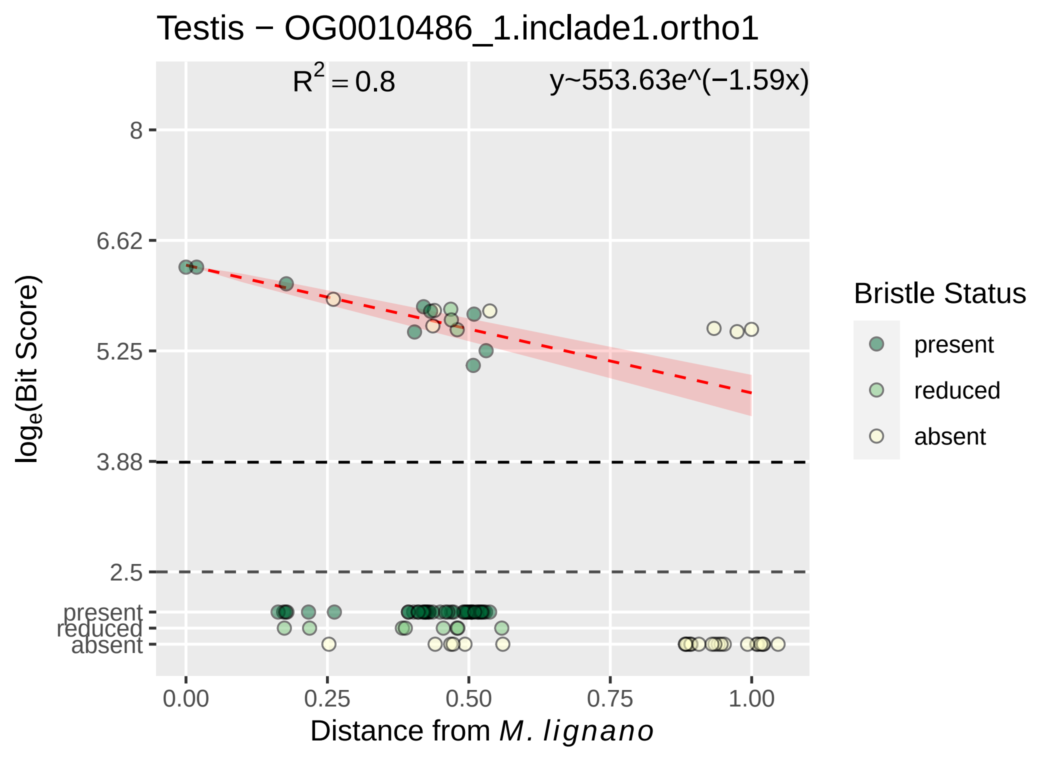

Testis − OG0010486_1.inclade1.ortho1
R² = 0.8
y~553.63e^(−1.59x)
8
6.62
5.25
3.88
2.5
present
reduced
absent
0.00
0.25
0.50
0.75
1.00
Distance from M. lignano
loge(Bit Score)
Bristle Status
present
reduced
absent

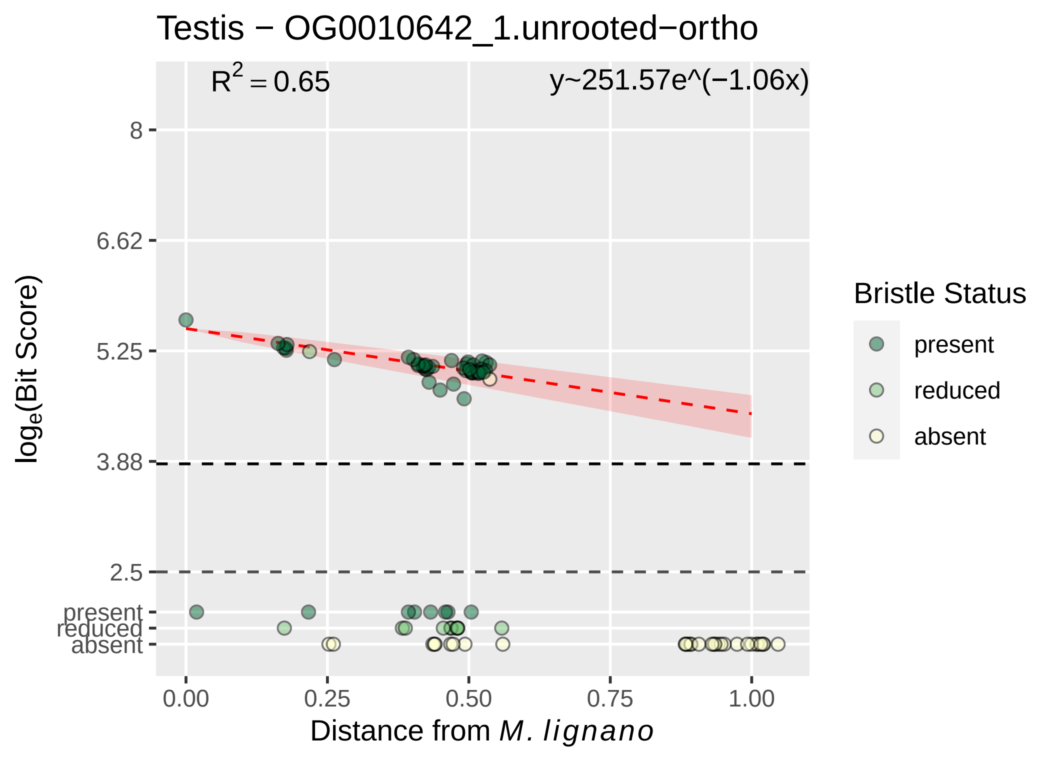
Testis − OG0010642_1.unrooted−ortho
$R^2 = 0.65$
y~251.57e^(−1.06x)
8
6.62
5.25
3.88
2.5
present
reduced
absent
$\log_e$(Bit Score)
0.00
0.25
0.50
0.75
1.00
Distance from *M. lignano*
Bristle Status
present
reduced
absent

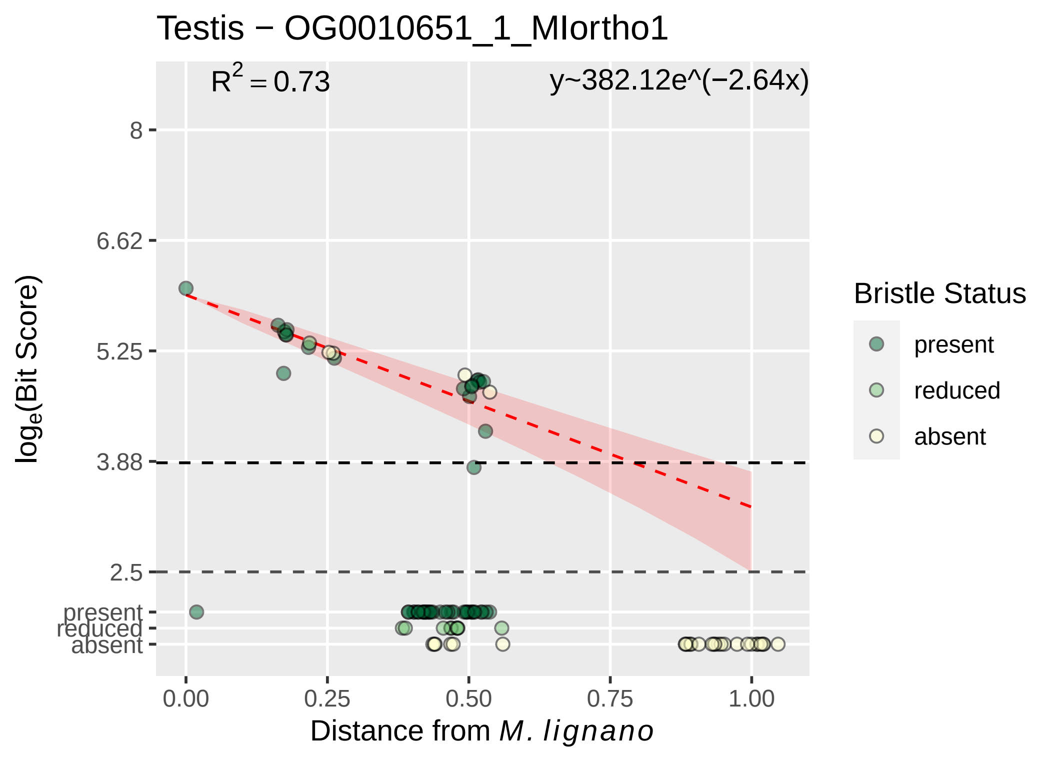

Testis − OG0010651_1_MIortho1
$R^2 = 0.73$
y~382.12e^(−2.64x)
8
6.62
5.25
3.88
2.5
present
reduced
absent
log$_e$(Bit Score)
0.00
0.25
0.50
0.75
1.00
Distance from *M. lignano*
Bristle Status
present
reduced
absent

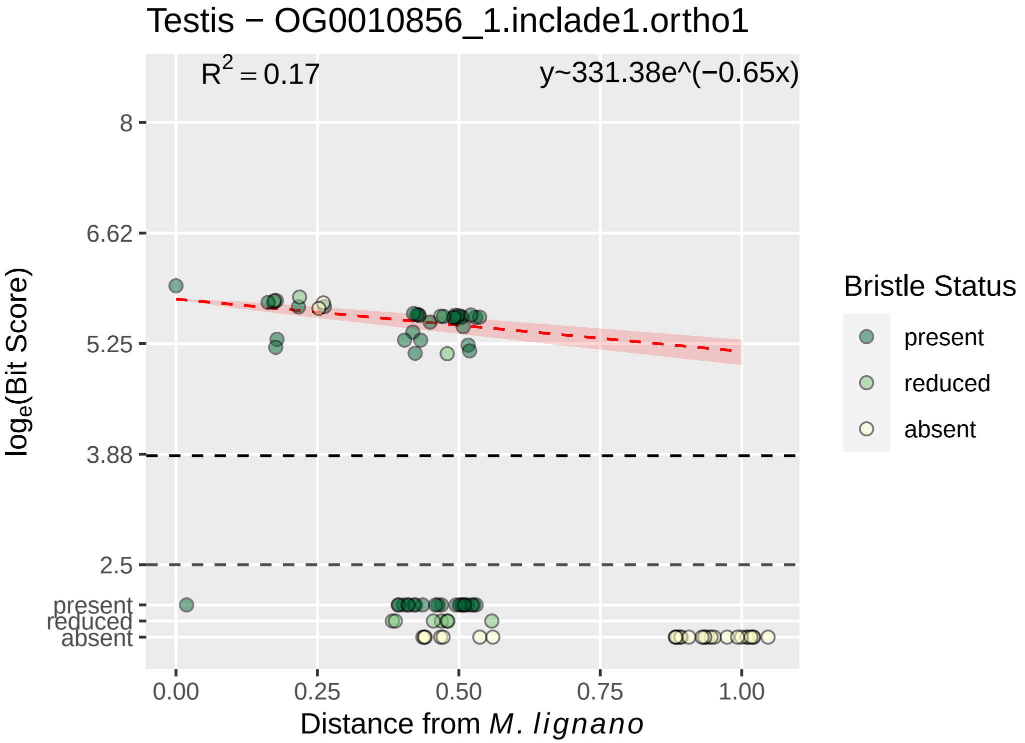

Testis − OG0010856_1.inclade1.ortho1
$R^2 = 0.17$
y~331.38e^(−0.65x)
8
6.62
5.25
3.88
2.5
present
reduced
absent
$log_e$(Bit Score)
0.00
0.25
0.50
0.75
1.00
Distance from *M. lignano*
Bristle Status
present
reduced
absent

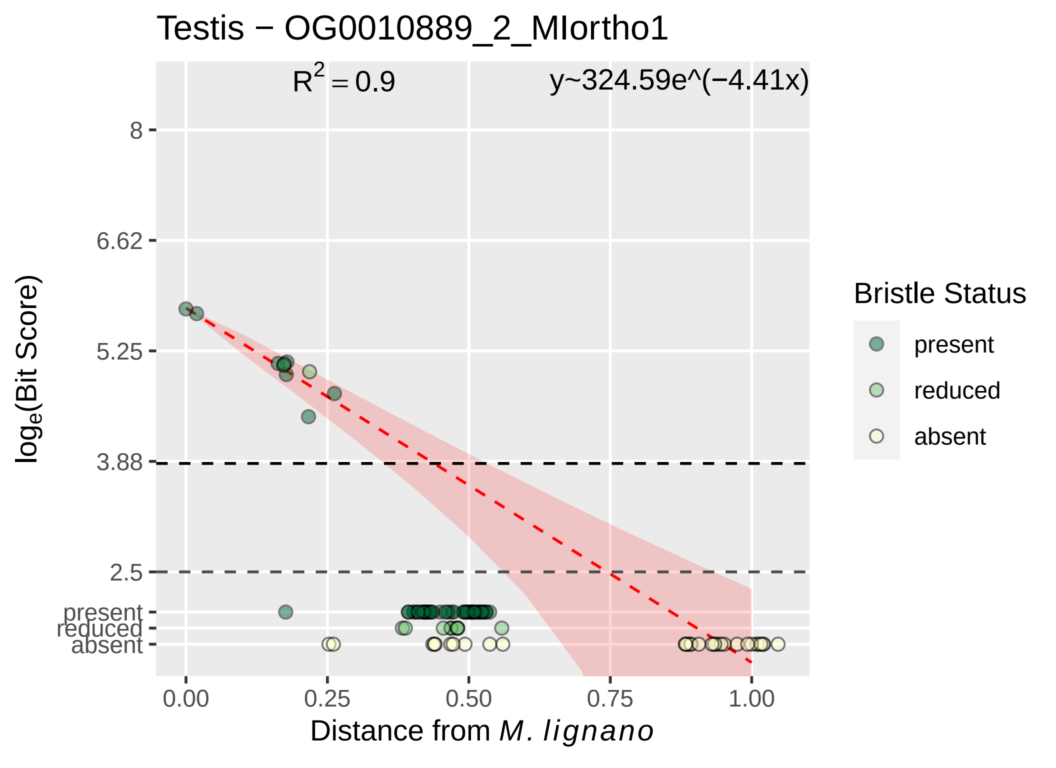
Testis − OG0010889_2_MIortho1
R^2 = 0.9
y~324.59e^(−4.41x)
8
6.62
5.25
3.88
2.5
present
reduced
absent
log_e(Bit Score)
0.00
0.25
0.50
0.75
1.00
Distance from M. lignano
Bristle Status
present
reduced
absent

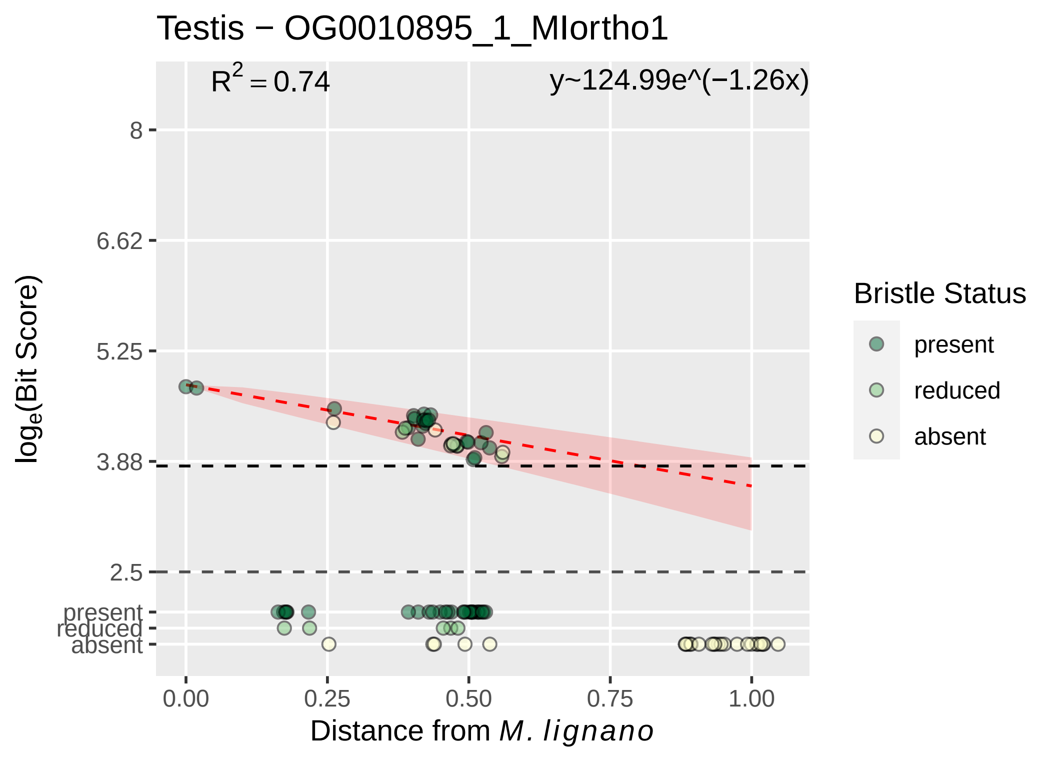

Testis − OG0010895_1_MIortho1
R2 = 0.74
y~124.99e^(−1.26x)
8
6.62
5.25
3.88
2.5
present
reduced
absent
0.00
0.25
0.50
0.75
1.00
Distance from M. lignano
loge(Bit Score)
Bristle Status
present
reduced
absent

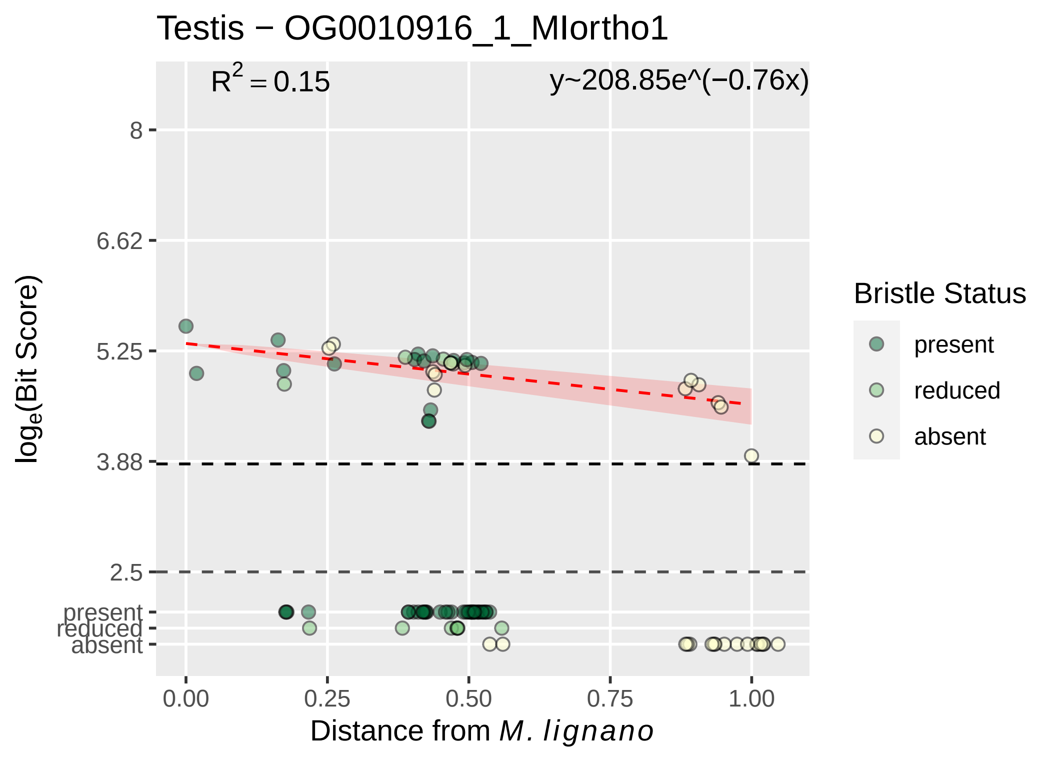
Testis − OG0010916_1_MIortho1
$R^2 = 0.15$
y~208.85e^(−0.76x)
8
6.62
5.25
3.88
2.5
present
reduced
absent
0.00
0.25
0.50
0.75
1.00
Distance from M. lignano
log$_e$(Bit Score)
Bristle Status
present
reduced
absent

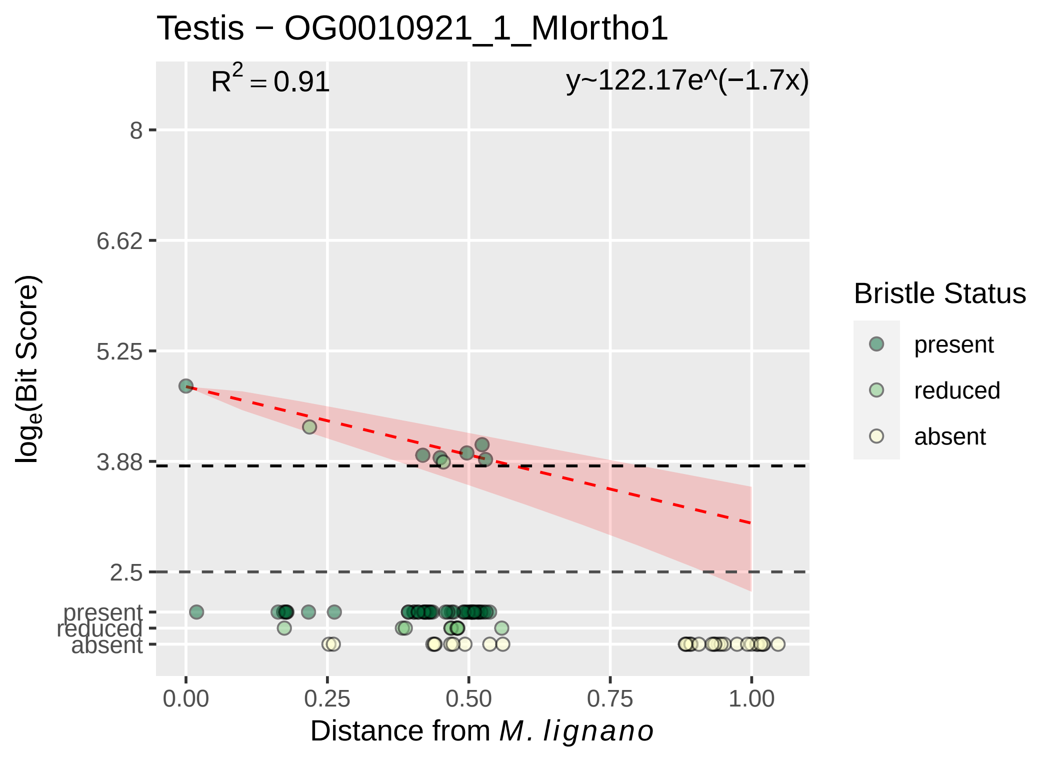

Testis − OG0010921_1_MIortho1
R2 = 0.91
y~122.17e^(−1.7x)
8
6.62
5.25
3.88
2.5
present
reduced
absent
loge(Bit Score)
0.00
0.25
0.50
0.75
1.00
Distance from M. lignano
Bristle Status
present
reduced
absent

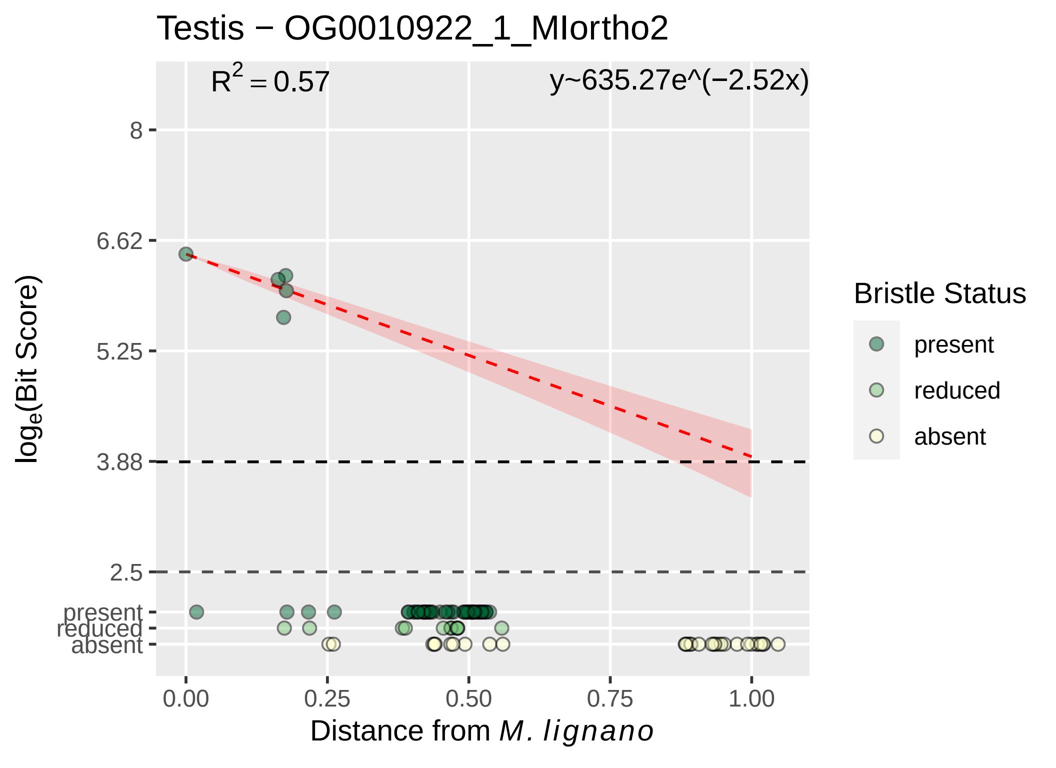

Testis − OG0010922_1_MIortho2
R2 = 0.57
y~635.27e^(−2.52x)
8
6.62
5.25
3.88
2.5
present
reduced
absent
0.00
0.25
0.50
0.75
1.00
Distance from M. lignano
loge(Bit Score)
Bristle Status
present
reduced
absent

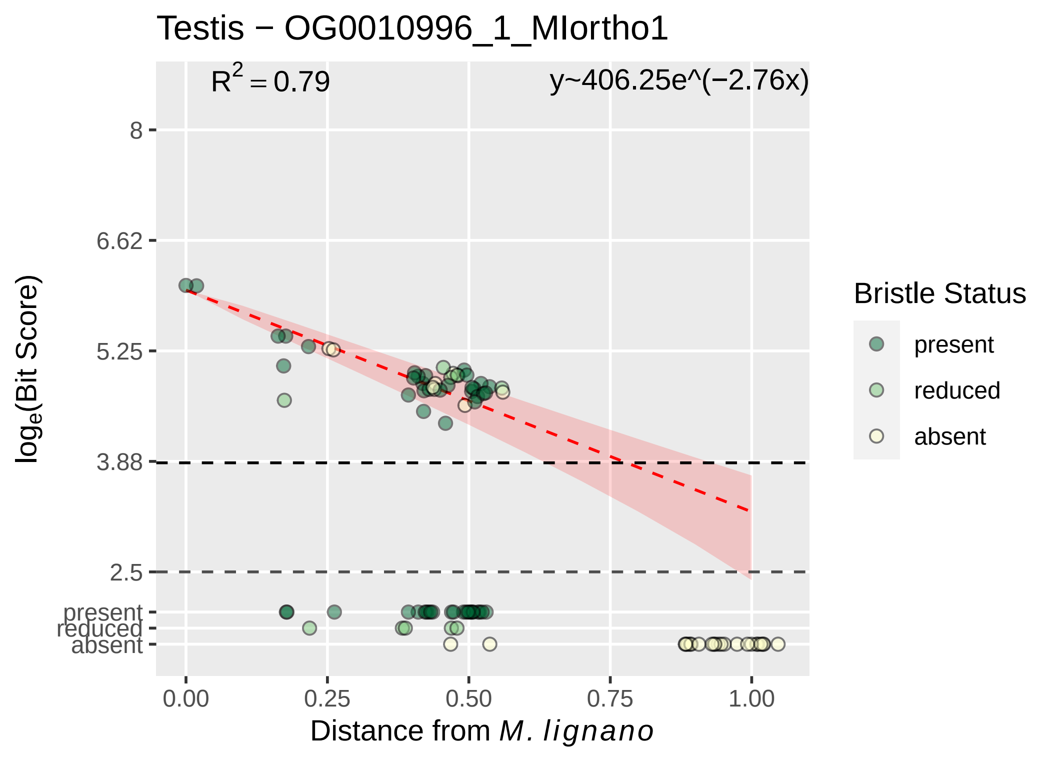

Testis − OG0010996_1_MIortho1
$R^2 = 0.79$
y~406.25e^(−2.76x)
8
6.62
5.25
3.88
2.5
present
reduced
absent
0.00
0.25
0.50
0.75
1.00
Distance from *M. lignano*
$\log_e$(Bit Score)
Bristle Status
present
reduced
absent

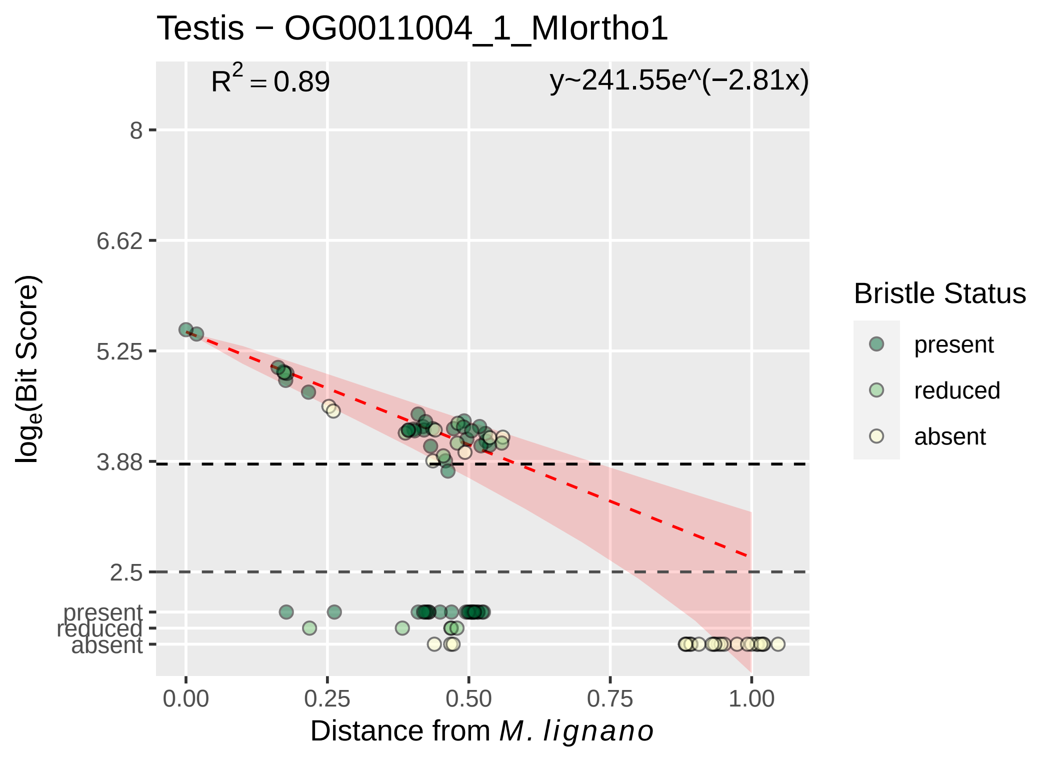
Testis − OG0011004_1_MIortho1
$R^2 = 0.89$
y~241.55e^(−2.81x)
8
6.62
5.25
3.88
2.5
present
reduced
absent
$\log_e$(Bit Score)
0.00
0.25
0.50
0.75
1.00
Distance from *M. lignano*
Bristle Status
present
reduced
absent

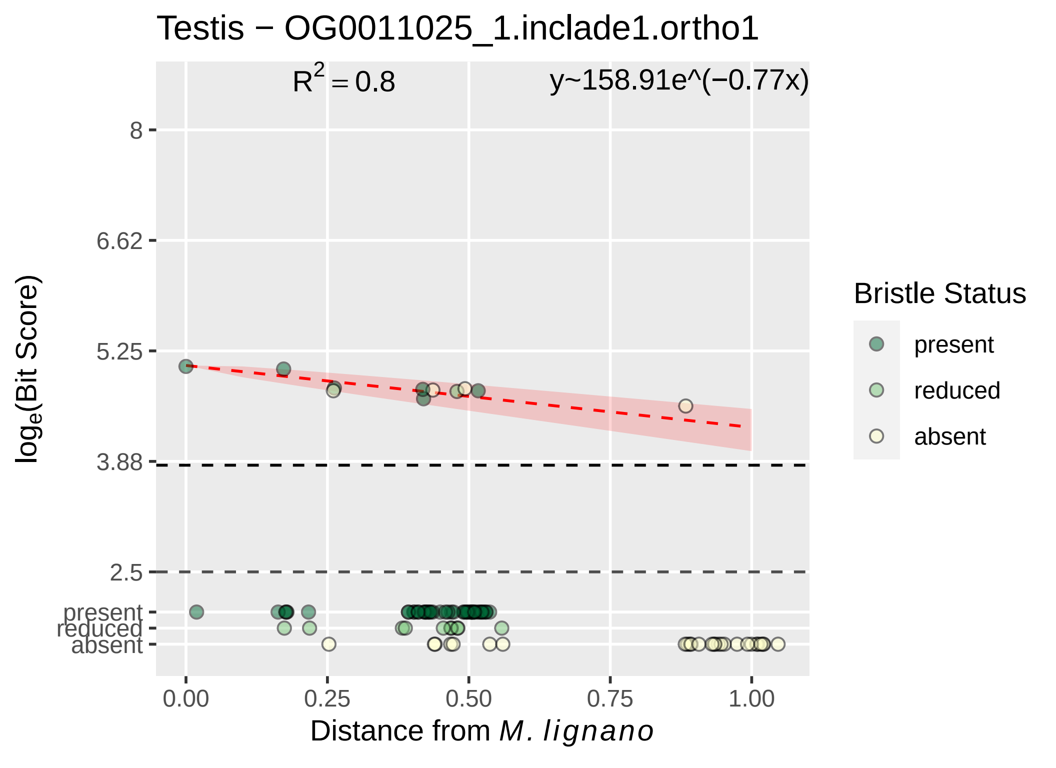

Testis − OG0011025_1.inclade1.ortho1
R^2 = 0.8
y~158.91e^(−0.77x)
8
6.62
5.25
3.88
2.5
present
reduced
absent
log_e(Bit Score)
0.00
0.25
0.50
0.75
1.00
Distance from M. lignano
Bristle Status
present
reduced
absent

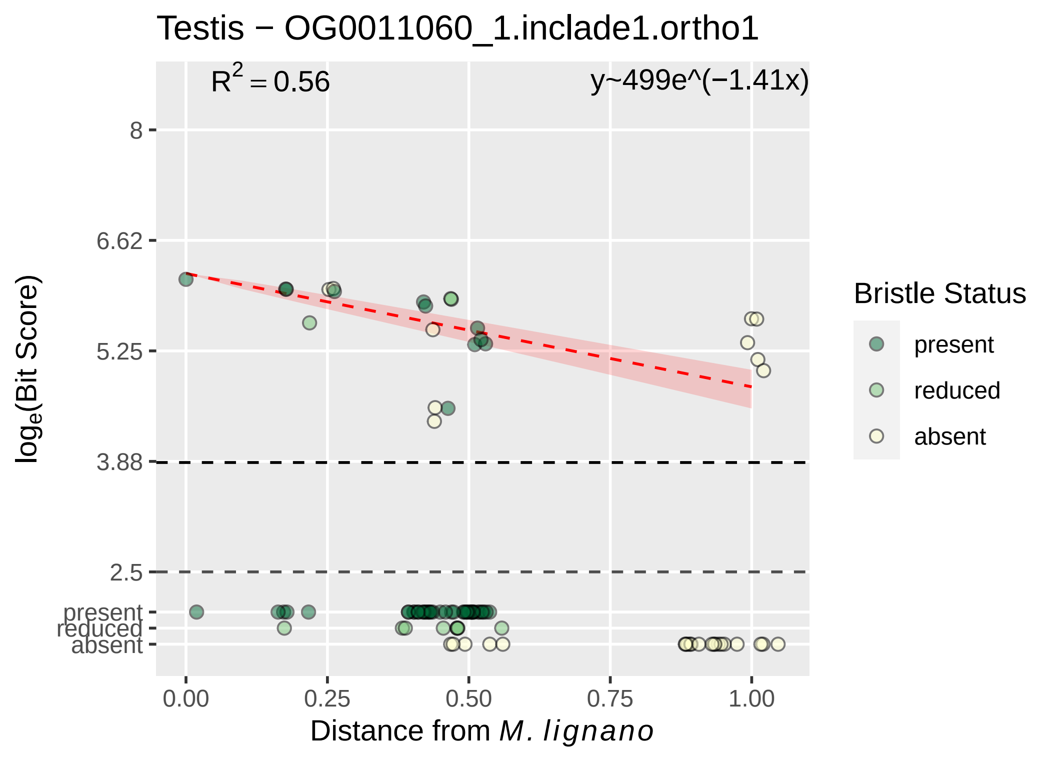

Testis − OG0011060_1.inclade1.ortho1
R^2 = 0.56
y~499e^(−1.41x)
8
6.62
5.25
3.88
2.5
present
reduced
absent
log_e(Bit Score)
0.00
0.25
0.50
0.75
1.00
Distance from M. lignano
Bristle Status
present
reduced
absent

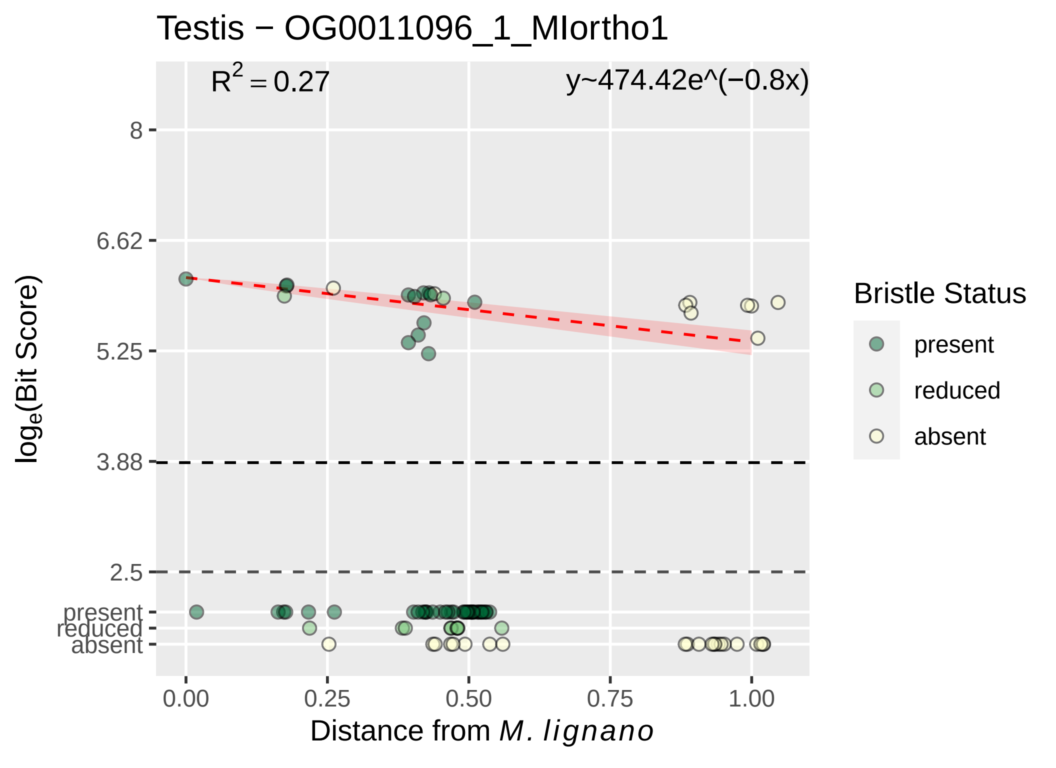

Testis − OG0011096_1_MIortho1
$R^2 = 0.27$
y~474.42e^(−0.8x)
8
6.62
5.25
3.88
2.5
present
reduced
absent
0.00
0.25
0.50
0.75
1.00
Distance from *M. lignano*
$\log_e$(Bit Score)
Bristle Status
present
reduced
absent

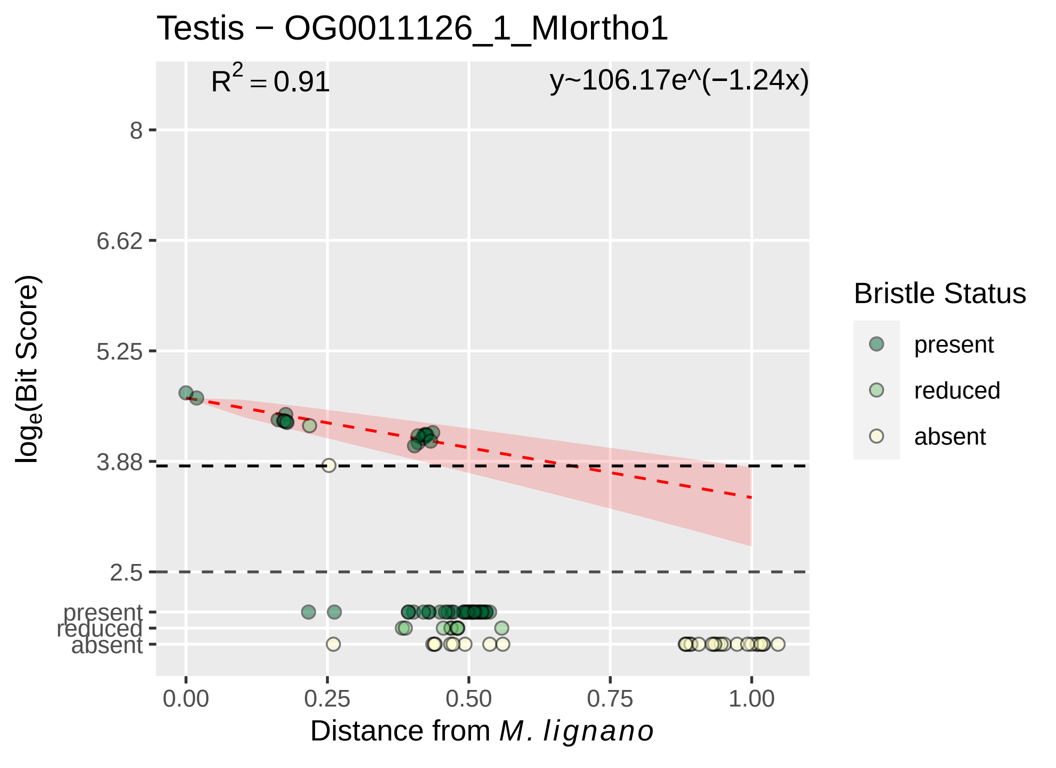

Testis − OG0011126_1_MIortho1
$R^2 = 0.91$
y~106.17e^(−1.24x)
8
6.62
5.25
3.88
2.5
present
reduced
absent
$\log_e$(Bit Score)
0.00
0.25
0.50
0.75
1.00
Distance from *M. lignano*
Bristle Status
present
reduced
absent

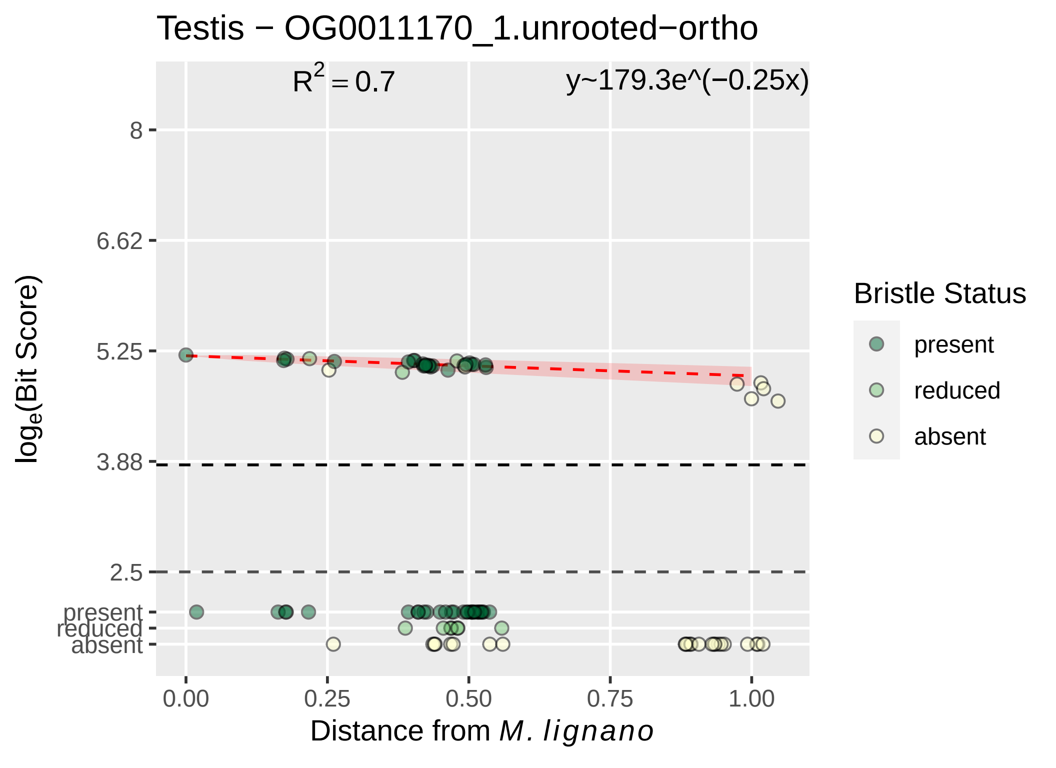

Testis − OG0011170_1.unrooted−ortho
R^2 = 0.7
y~179.3e^(−0.25x)
8
6.62
5.25
3.88
2.5
present
reduced
absent
log_e(Bit Score)
0.00
0.25
0.50
0.75
1.00
Distance from M. lignano
Bristle Status
present
reduced
absent

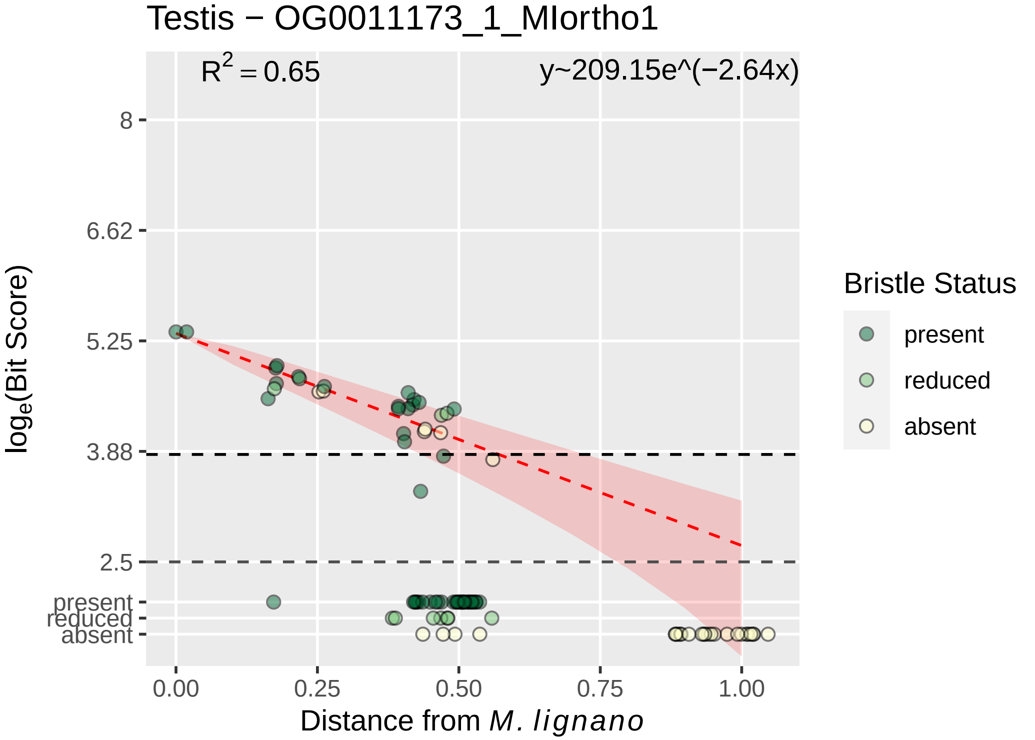
Testis − OG0011173_1_MIortho1
$R^2 = 0.65$
y~209.15e^(−2.64x)
8
6.62
5.25
3.88
2.5
present
reduced
absent
0.00
0.25
0.50
0.75
1.00
$log_e$(Bit Score)
Distance from *M. lignano*
Bristle Status
present
reduced
absent

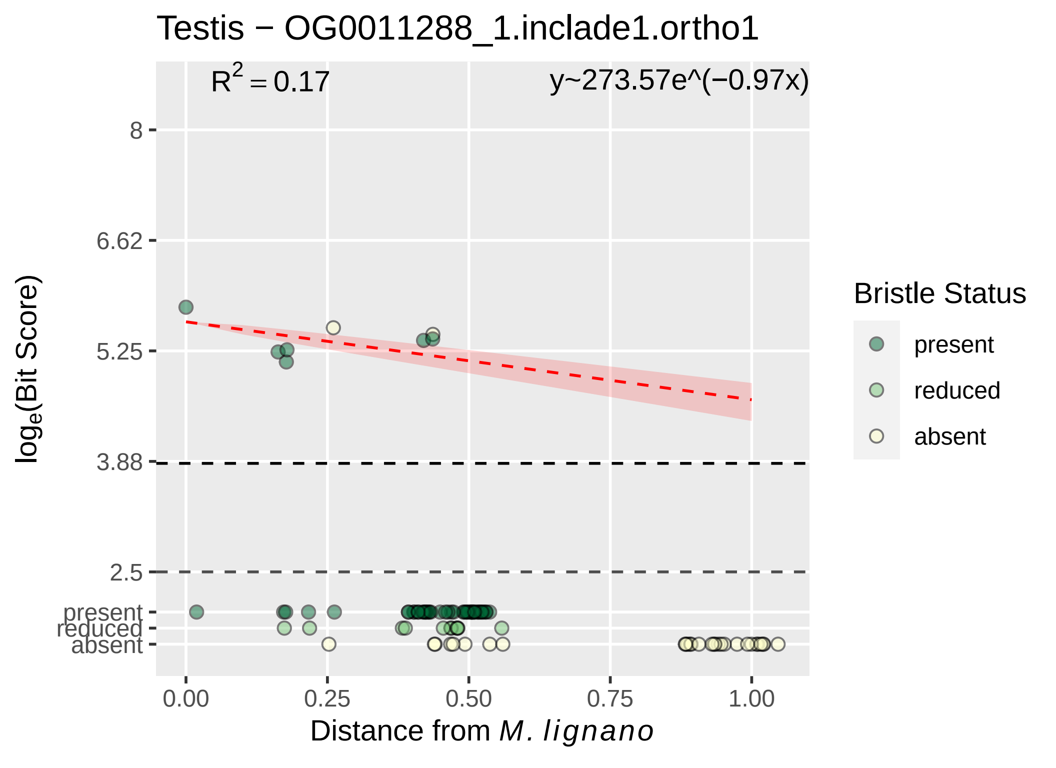
Testis − OG0011288_1.inclade1.ortho1
$R^2 = 0.17$
y~273.57e^(−0.97x)
8
6.62
5.25
3.88
2.5
present
reduced
absent
$\log_e$(Bit Score)
0.00
0.25
0.50
0.75
1.00
Distance from *M. lignano*
Bristle Status
present
reduced
absent

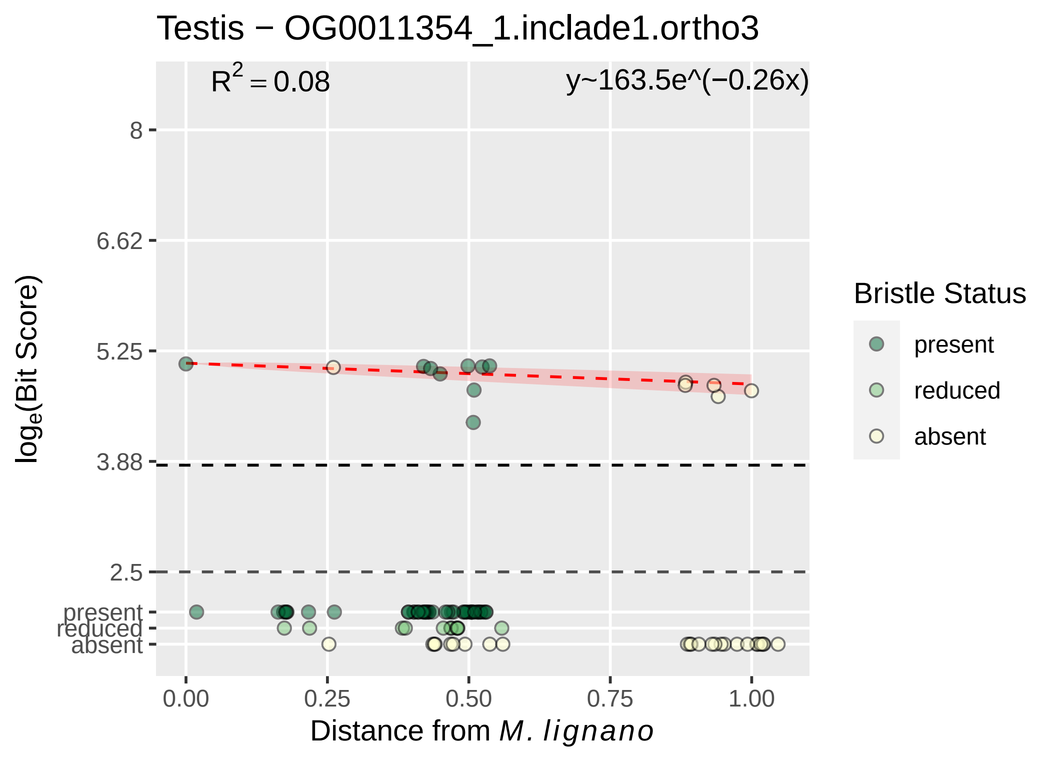
Testis − OG0011354_1.inclade1.ortho3
$R^2 = 0.08$
y~163.5e^(−0.26x)
8
6.62
5.25
3.88
2.5
present
reduced
absent
$log_e$(Bit Score)
0.00
0.25
0.50
0.75
1.00
Distance from *M. lignano*
Bristle Status
present
reduced
absent

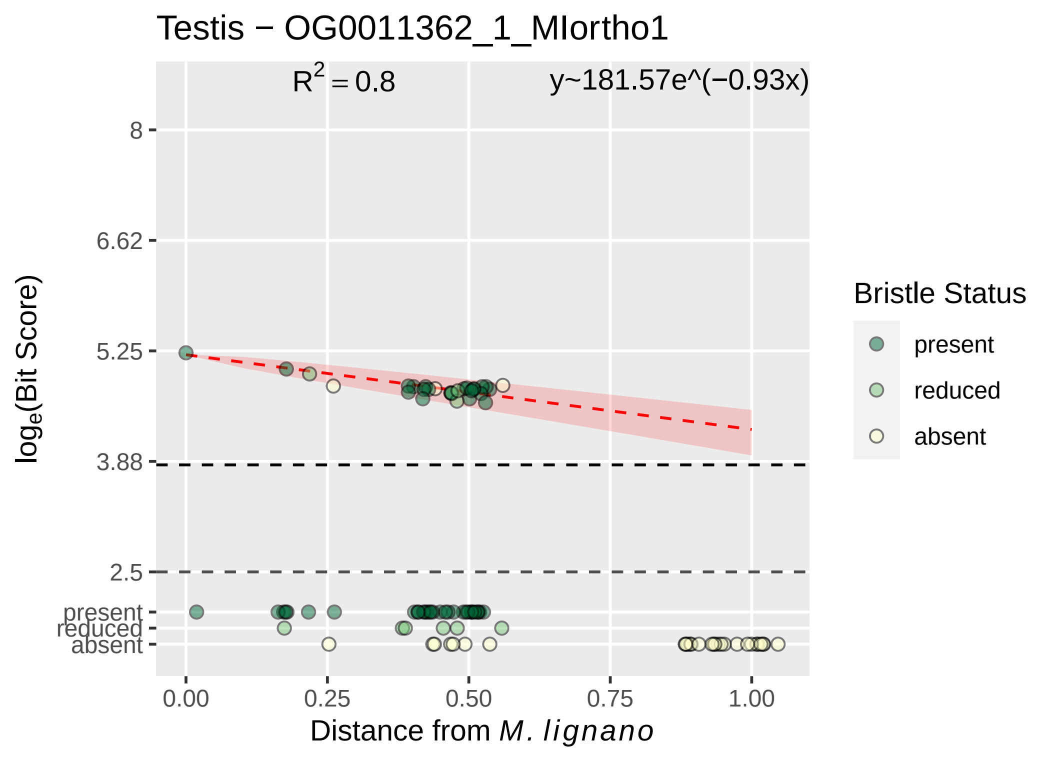

Testis − OG0011362_1_MIortho1
R^2 = 0.8
y~181.57e^(−0.93x)
8
6.62
5.25
3.88
2.5
present
reduced
absent
log_e(Bit Score)
0.00
0.25
0.50
0.75
1.00
Distance from M. lignano
Bristle Status
present
reduced
absent

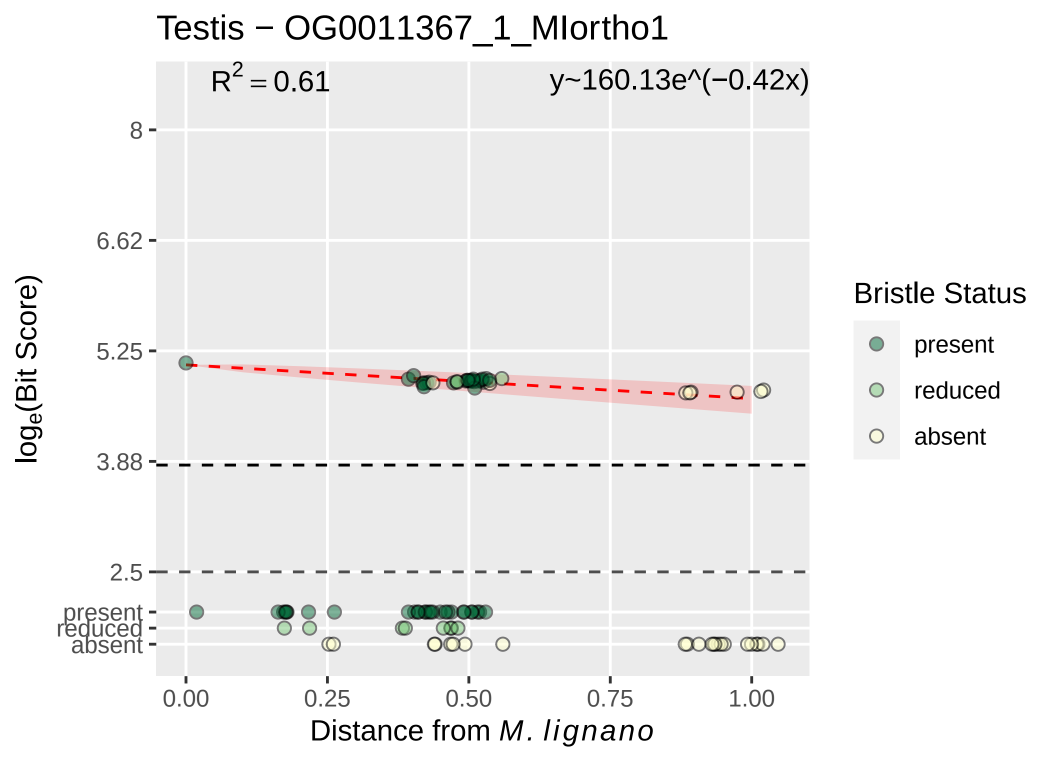
Testis − OG0011367_1_MIortho1
$R^2 = 0.61$
y~160.13e^(−0.42x)
8
6.62
5.25
3.88
2.5
present
reduced
absent
0.00
0.25
0.50
0.75
1.00
log_e(Bit Score)
Distance from M. lignano
Bristle Status
present
reduced
absent

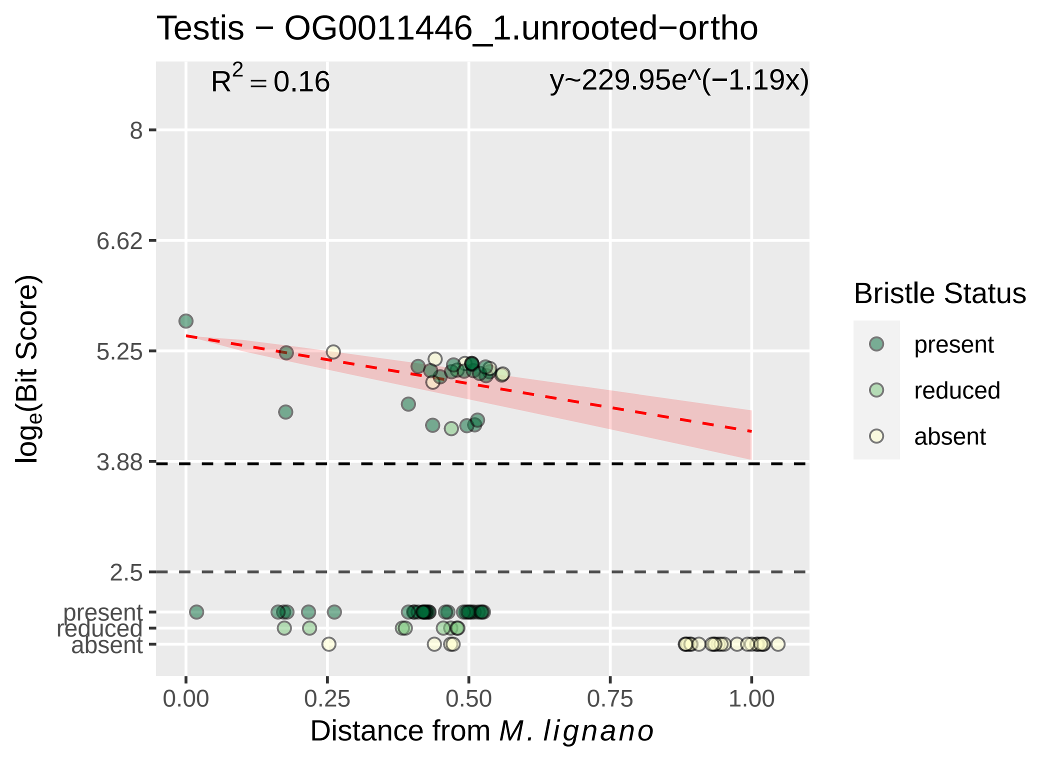

Testis − OG0011446_1.unrooted−ortho
$R^2 = 0.16$
y~229.95e^(−1.19x)
8
6.62
5.25
3.88
2.5
present
reduced
absent
log$_e$(Bit Score)
0.00
0.25
0.50
0.75
1.00
Distance from *M. lignano*
Bristle Status
present
reduced
absent

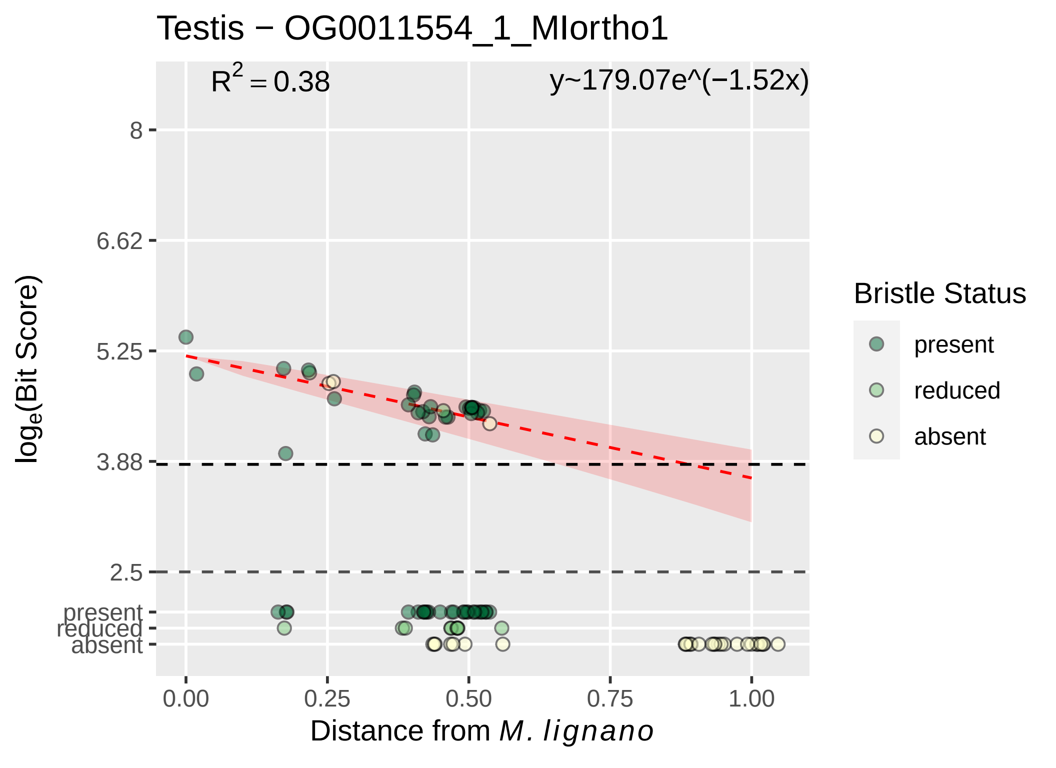

Testis − OG0011554_1_MIortho1
$R^2 = 0.38$
y~179.07e^(−1.52x)
8
6.62
5.25
3.88
2.5
present
reduced
absent
0.00
0.25
0.50
0.75
1.00
$\log_e$(Bit Score)
Distance from *M. lignano*
Bristle Status
present
reduced
absent

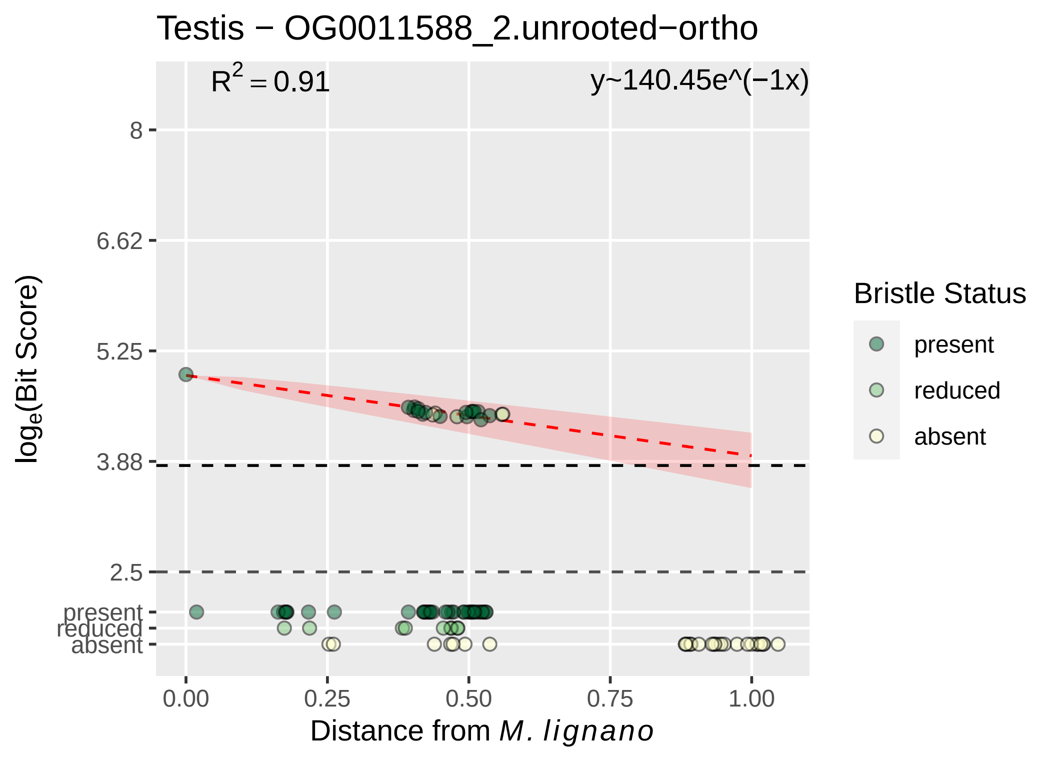
Testis − OG0011588_2.unrooted−ortho
$R^2 = 0.91$
y~140.45e^(−1x)
8
6.62
5.25
3.88
2.5
present
reduced
absent
0.00
0.25
0.50
0.75
1.00
Distance from M. lignano
log$_e$(Bit Score)
Bristle Status
present
reduced
absent

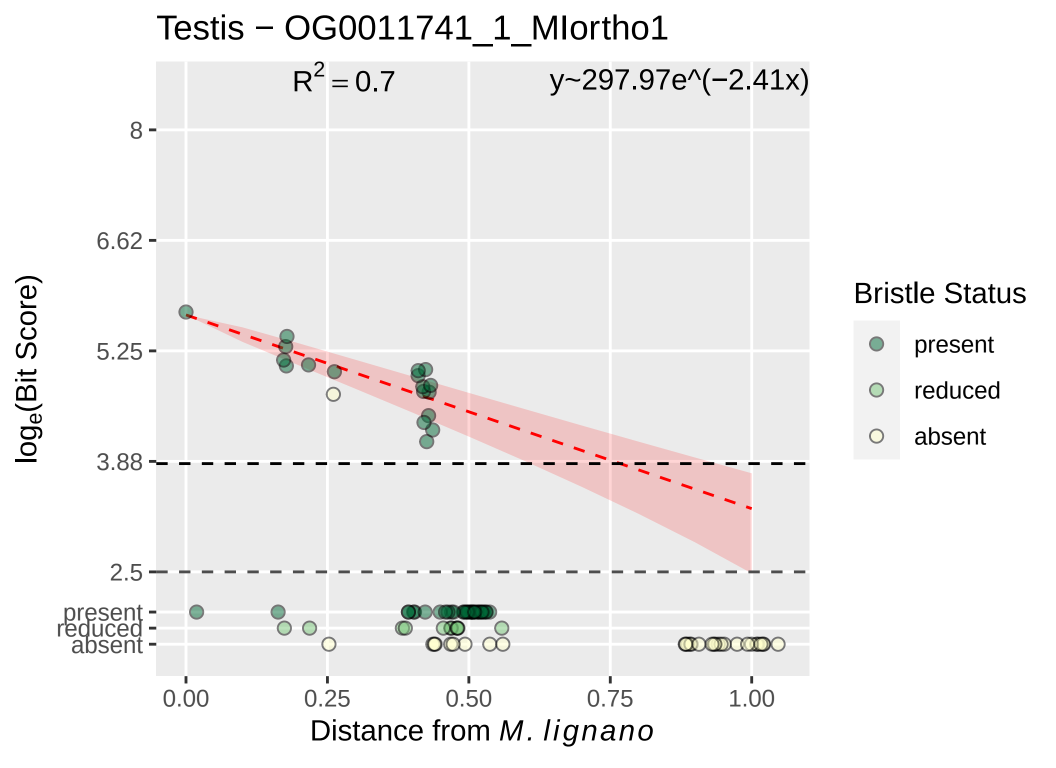

Testis − OG0011741_1_MIortho1
R2 = 0.7
y~297.97e^(−2.41x)
8
6.62
5.25
3.88
2.5
present
reduced
absent
loge(Bit Score)
0.00
0.25
0.50
0.75
1.00
Distance from M. lignano
Bristle Status
present
reduced
absent

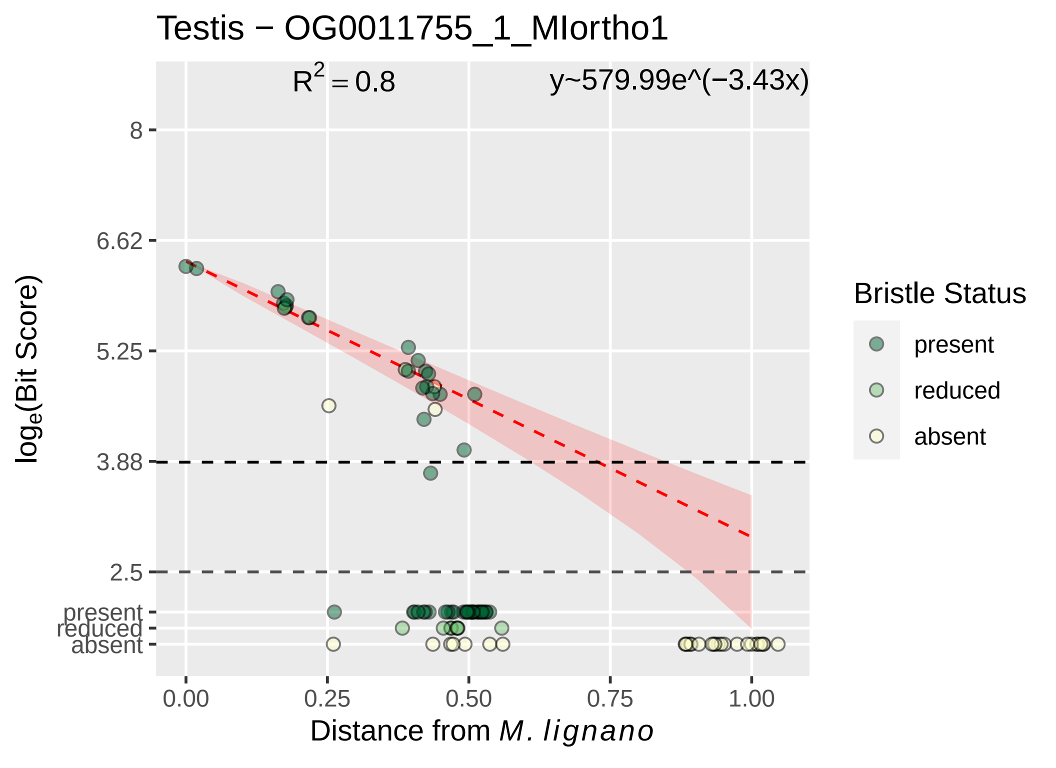

Testis − OG0011755_1_MIortho1
$R^2 = 0.8$
y~579.99e^(−3.43x)
8
6.62
5.25
3.88
2.5
present
reduced
absent
0.00
0.25
0.50
0.75
1.00
Distance from *M. lignano*
$log_e$(Bit Score)
Bristle Status
present
reduced
absent

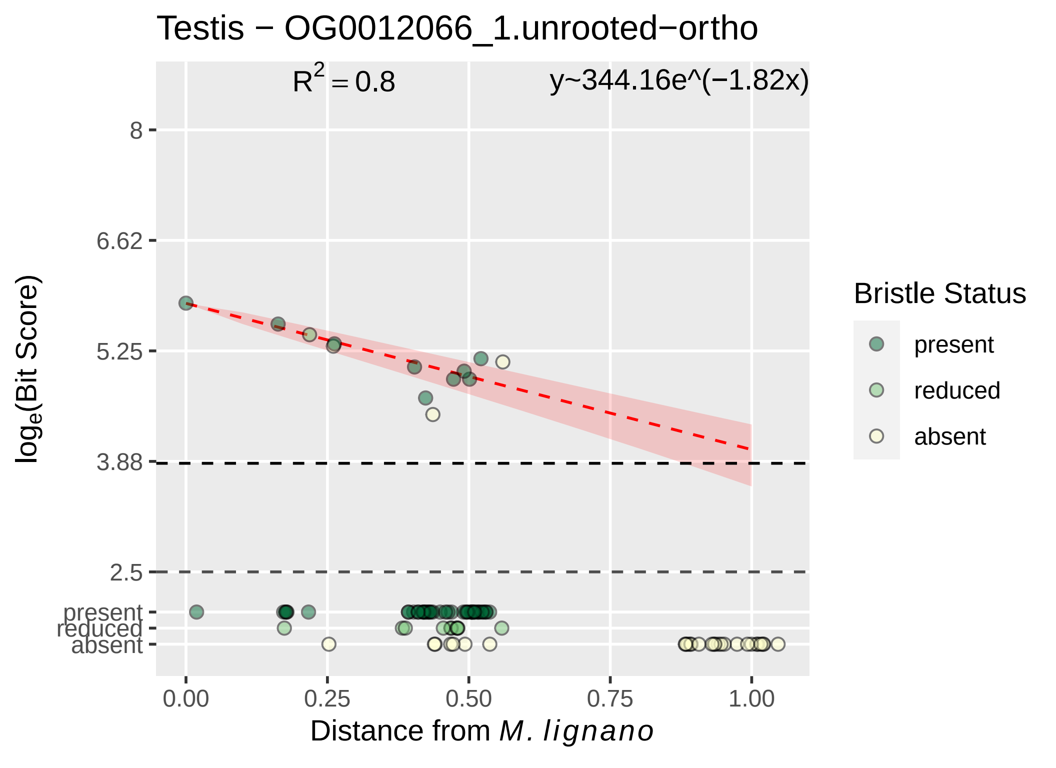
Testis − OG0012066_1.unrooted−ortho
R^2 = 0.8
y~344.16e^(−1.82x)
8
6.62
5.25
3.88
2.5
present
reduced
absent
loge(Bit Score)
0.00
0.25
0.50
0.75
1.00
Distance from M. lignano
Bristle Status
present
reduced
absent

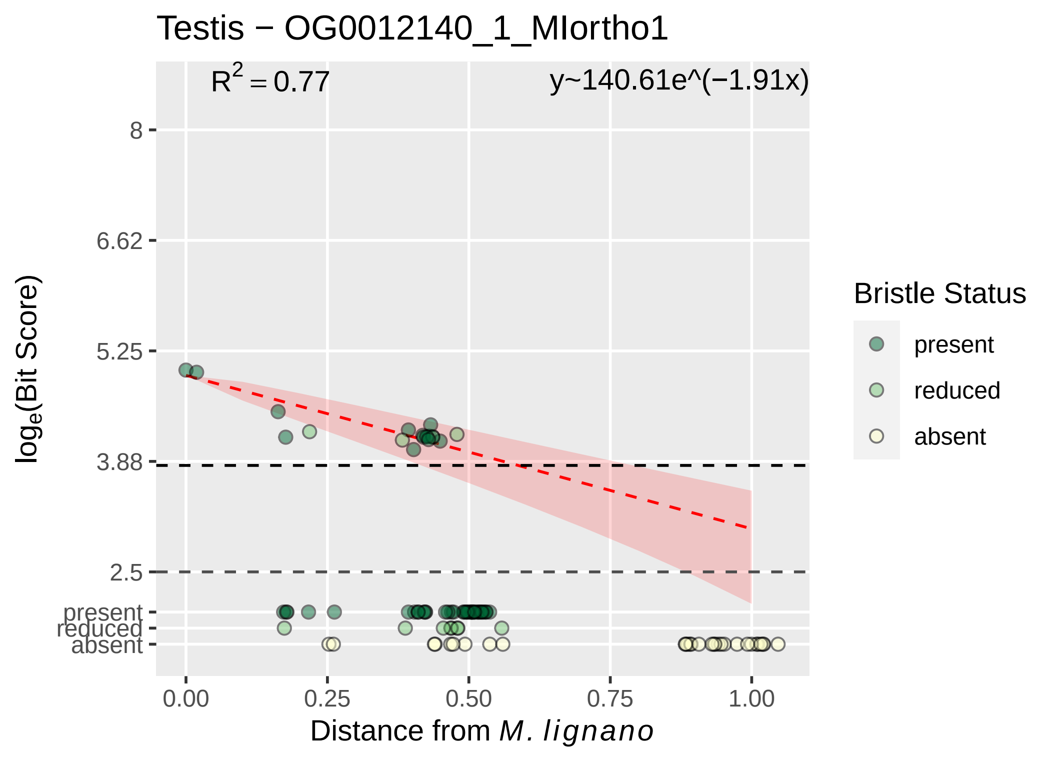

Testis − OG0012140_1_MIortho1
$R^2 = 0.77$
y~140.61e^(−1.91x)
8
6.62
5.25
3.88
2.5
present
reduced
absent
$\log_e$(Bit Score)
0.00
0.25
0.50
0.75
1.00
Distance from *M. lignano*
Bristle Status
present
reduced
absent

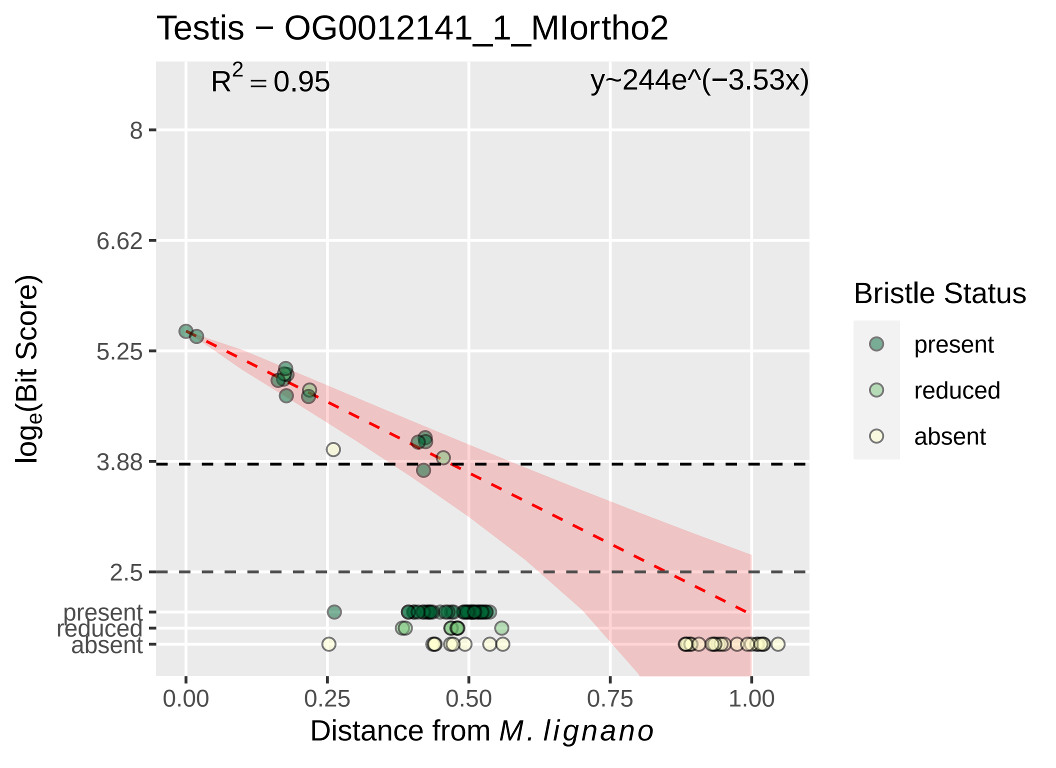

Testis − OG0012141_1_MIortho2
R2 = 0.95
y~244e^(−3.53x)
8
6.62
5.25
3.88
2.5
present
reduced
absent
0.00
0.25
0.50
0.75
1.00
Distance from M. lignano
loge(Bit Score)
Bristle Status
present
reduced
absent

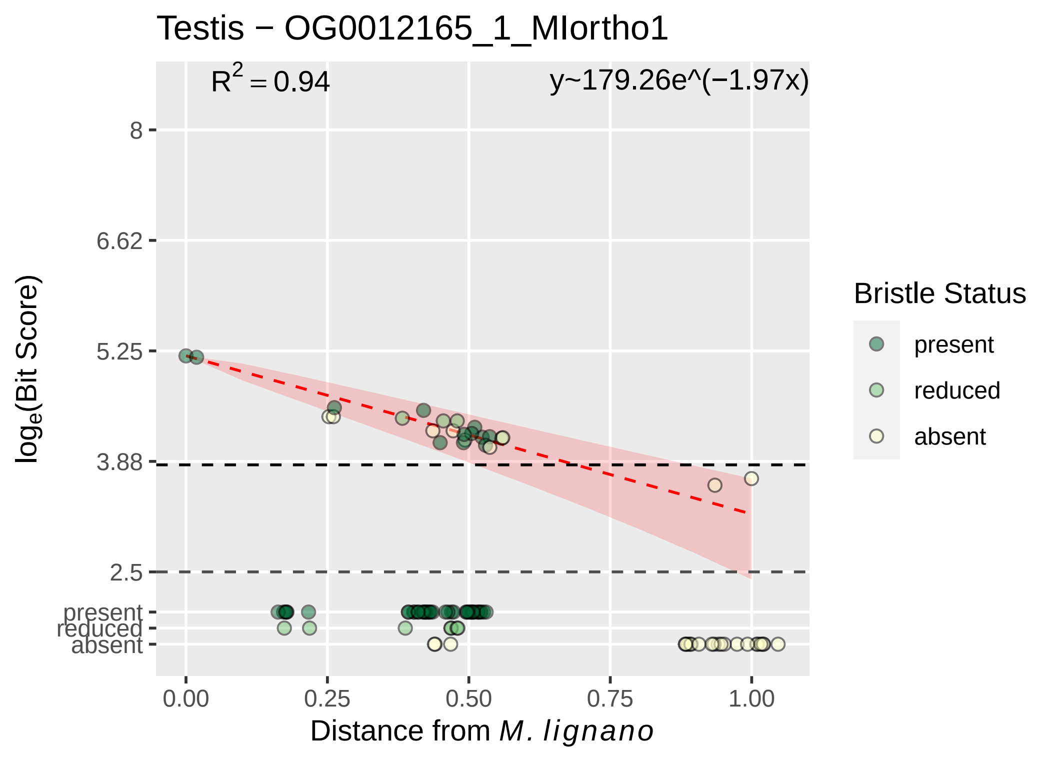

Testis − OG0012165_1_MIortho1
$R^2 = 0.94$
y~179.26e^(−1.97x)
8
6.62
5.25
3.88
2.5
present
reduced
absent
0.00
0.25
0.50
0.75
1.00
Distance from *M. lignano*
$log_e$(Bit Score)
Bristle Status
present
reduced
absent

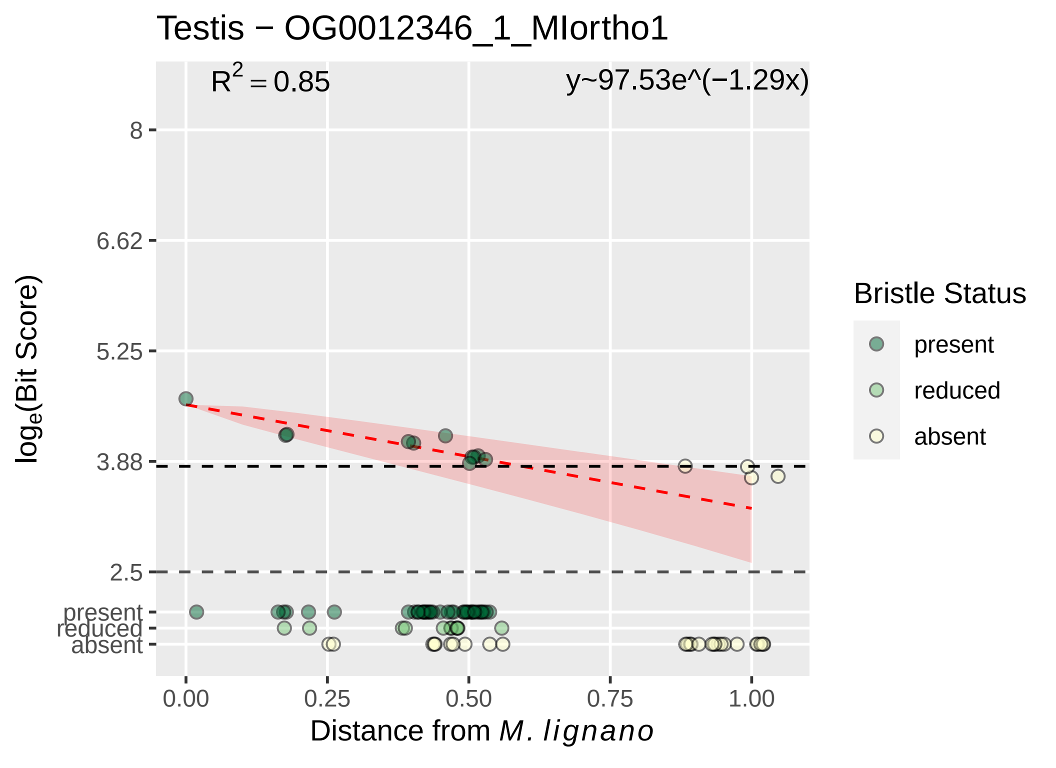
Testis − OG0012346_1_MIortho1
R^2 = 0.85
y~97.53e^(−1.29x)
8
6.62
5.25
3.88
2.5
present
reduced
absent
log_e(Bit Score)
0.00
0.25
0.50
0.75
1.00
Distance from M. lignano
Bristle Status
present
reduced
absent

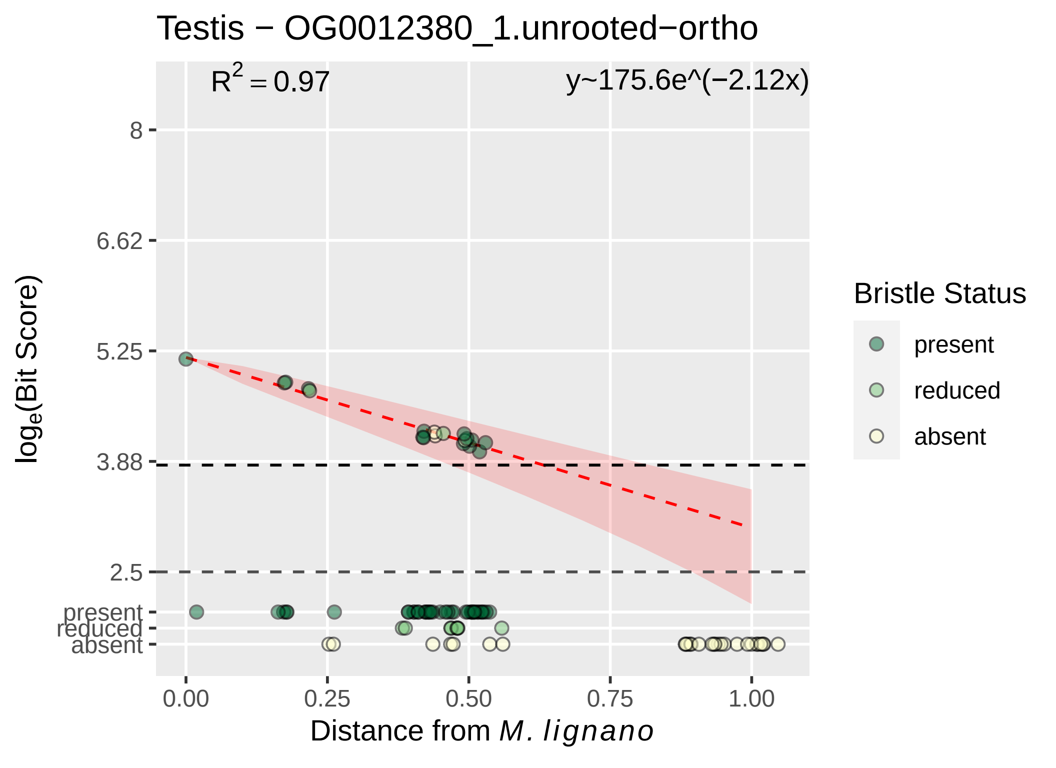

Testis − OG0012380_1.unrooted−ortho
$R^2 = 0.97$
y~175.6e^(−2.12x)
$log_e$(Bit Score)
8
6.62
5.25
3.88
2.5
present
reduced
absent
0.00
0.25
0.50
0.75
1.00
Distance from M. lignano
Bristle Status
present
reduced
absent

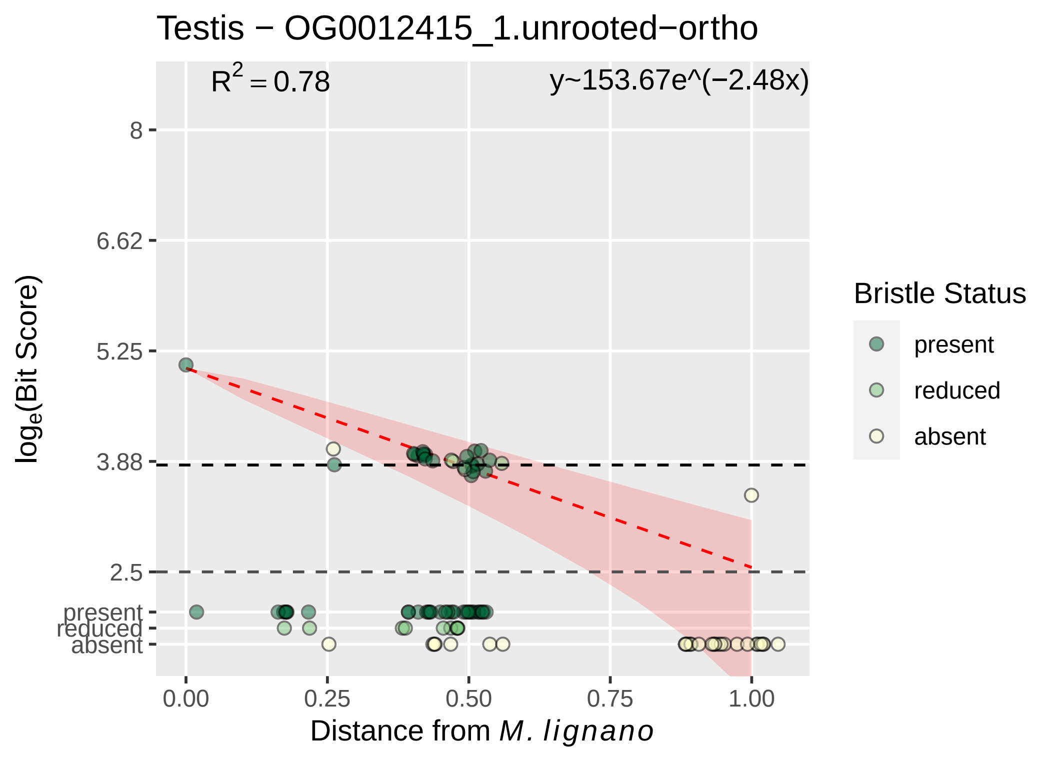

Testis − OG0012415_1.unrooted−ortho
$R^2 = 0.78$
y~153.67e^(−2.48x)
8
6.62
5.25
3.88
2.5
present
reduced
absent
0.00
0.25
0.50
0.75
1.00
Distance from M. lignano
log$_e$(Bit Score)
Bristle Status
present
reduced
absent

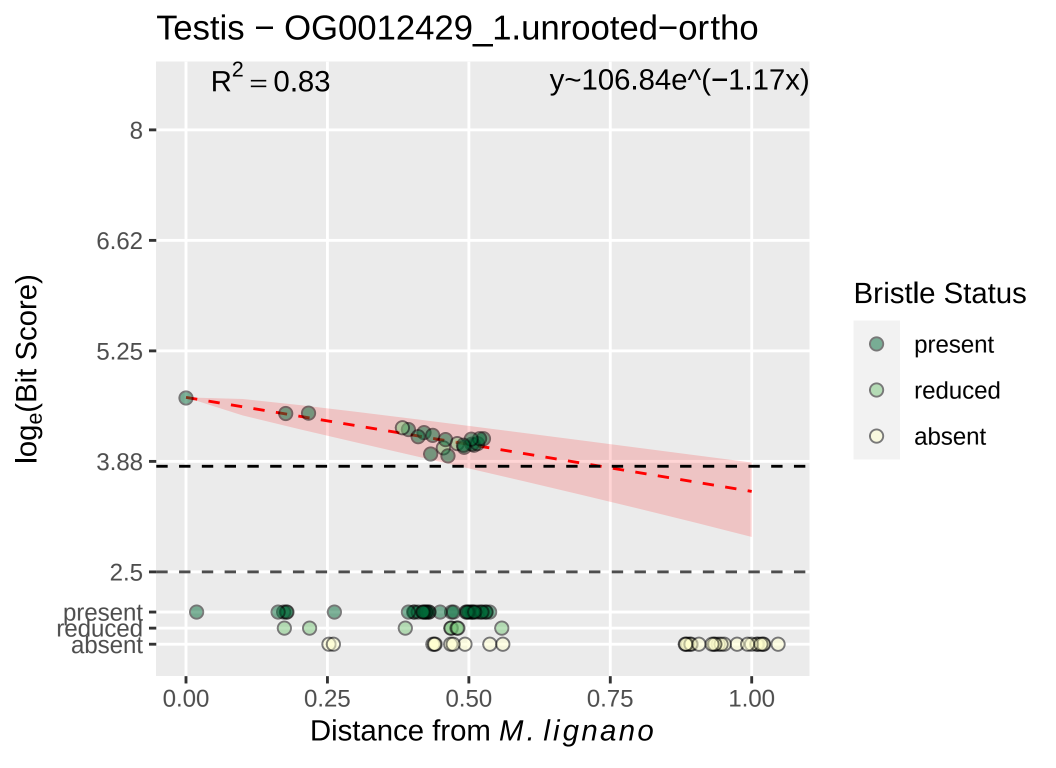

Testis − OG0012429_1.unrooted−ortho
$R^2 = 0.83$
y~106.84e^(−1.17x)
8
6.62
5.25
3.88
2.5
present
reduced
absent
0.00
0.25
0.50
0.75
1.00
Distance from *M. lignano*
$\log_e$(Bit Score)
Bristle Status
present
reduced
absent

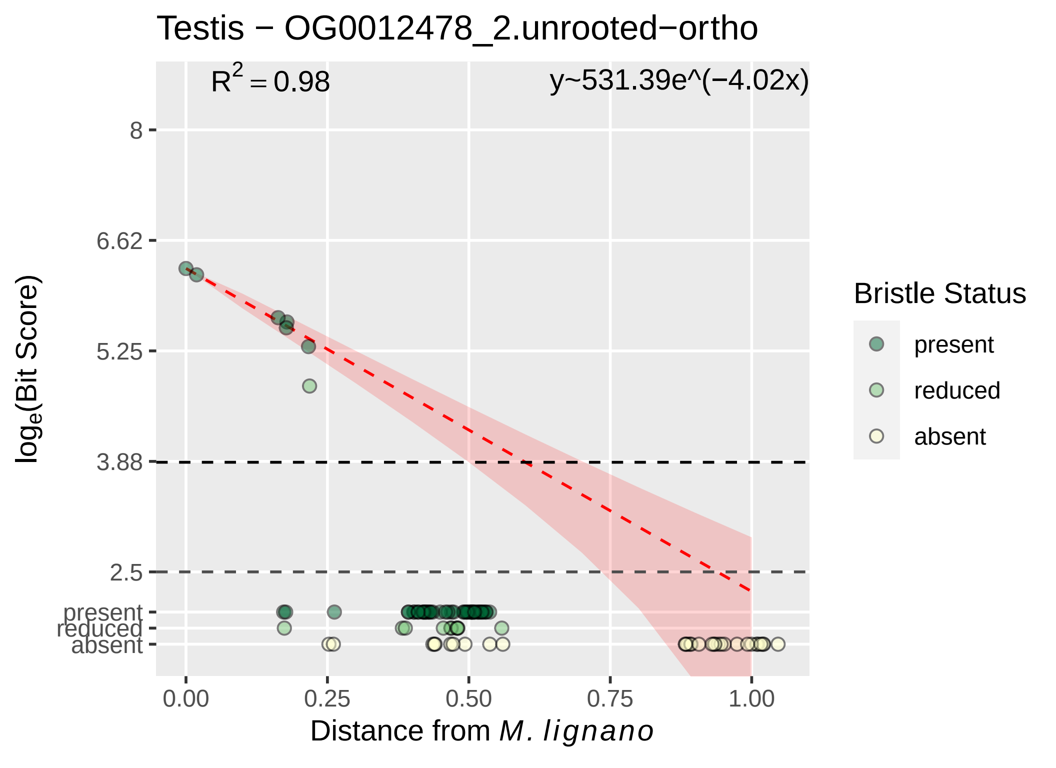

Testis − OG0012478_2.unrooted−ortho
$R^2 = 0.98$
y~531.39e^(−4.02x)
8
6.62
5.25
3.88
2.5
present
reduced
absent
0.00
0.25
0.50
0.75
1.00
Distance from M. lignano
log_e(Bit Score)
Bristle Status
present
reduced
absent

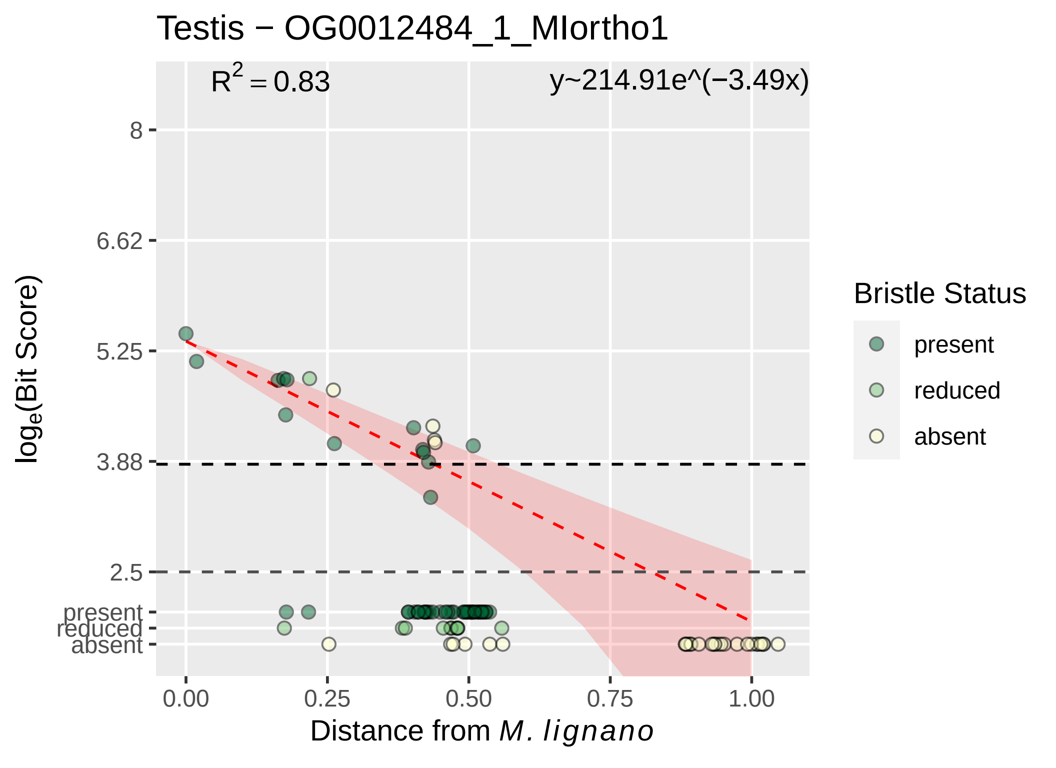

Testis − OG0012484_1_MIortho1
R^2 = 0.83
y~214.91e^(−3.49x)
8
6.62
5.25
3.88
2.5
present
reduced
absent
log_e(Bit Score)
0.00
0.25
0.50
0.75
1.00
Distance from M. lignano
Bristle Status
present
reduced
absent

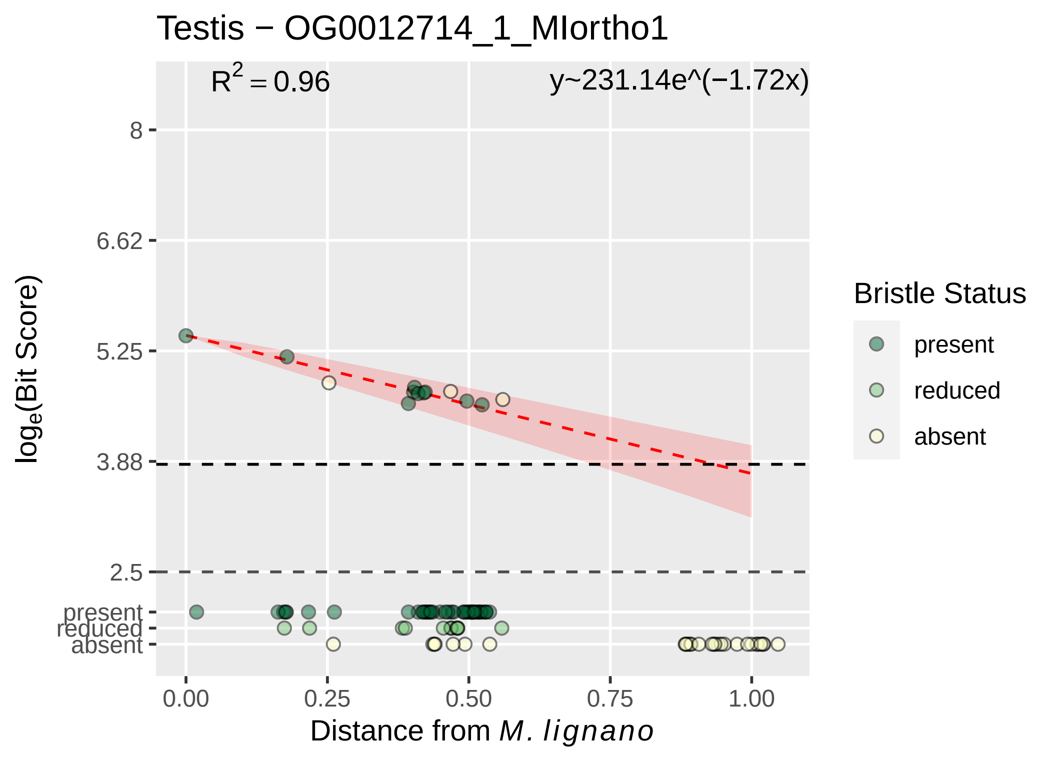
Testis − OG0012714_1_MIortho1
R^2 = 0.96
y~231.14e^(−1.72x)
8
6.62
5.25
3.88
2.5
present
reduced
absent
log_e(Bit Score)
0.00
0.25
0.50
0.75
1.00
Distance from M. lignano
Bristle Status
present
reduced
absent

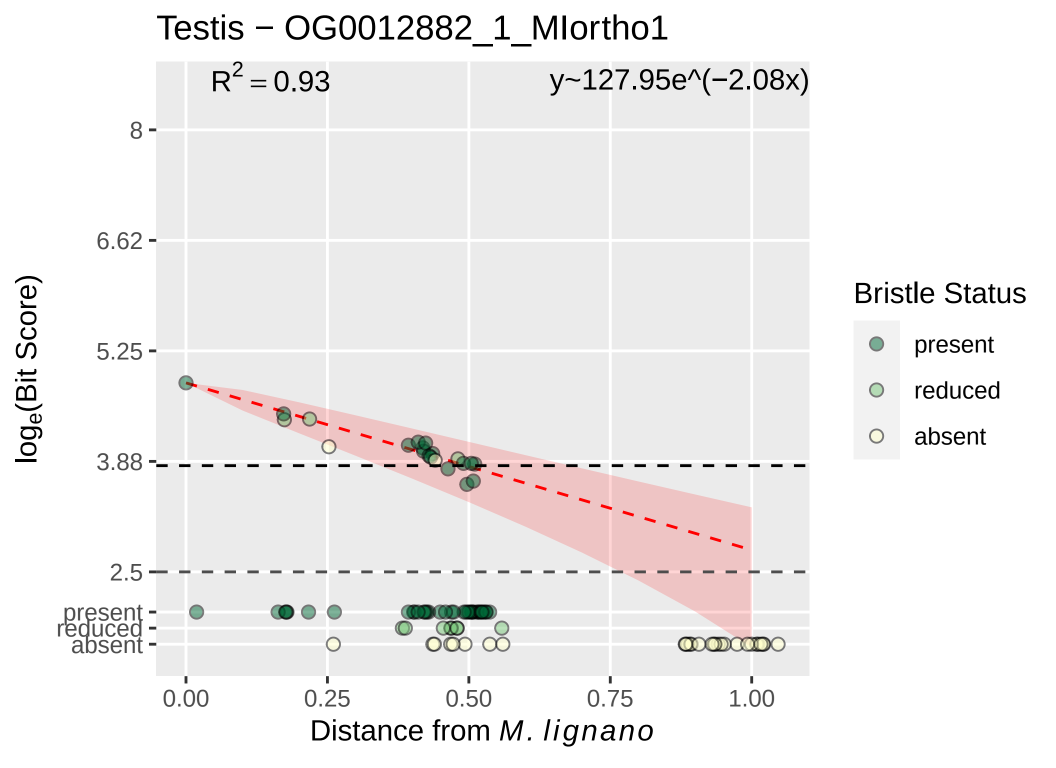
Testis − OG0012882_1_MIortho1
$R^2 = 0.93$
y~127.95e^(−2.08x)
8
6.62
5.25
3.88
2.5
present
reduced
absent
0.00
0.25
0.50
0.75
1.00
Distance from M. lignano
$\log_e$(Bit Score)
Bristle Status
present
reduced
absent

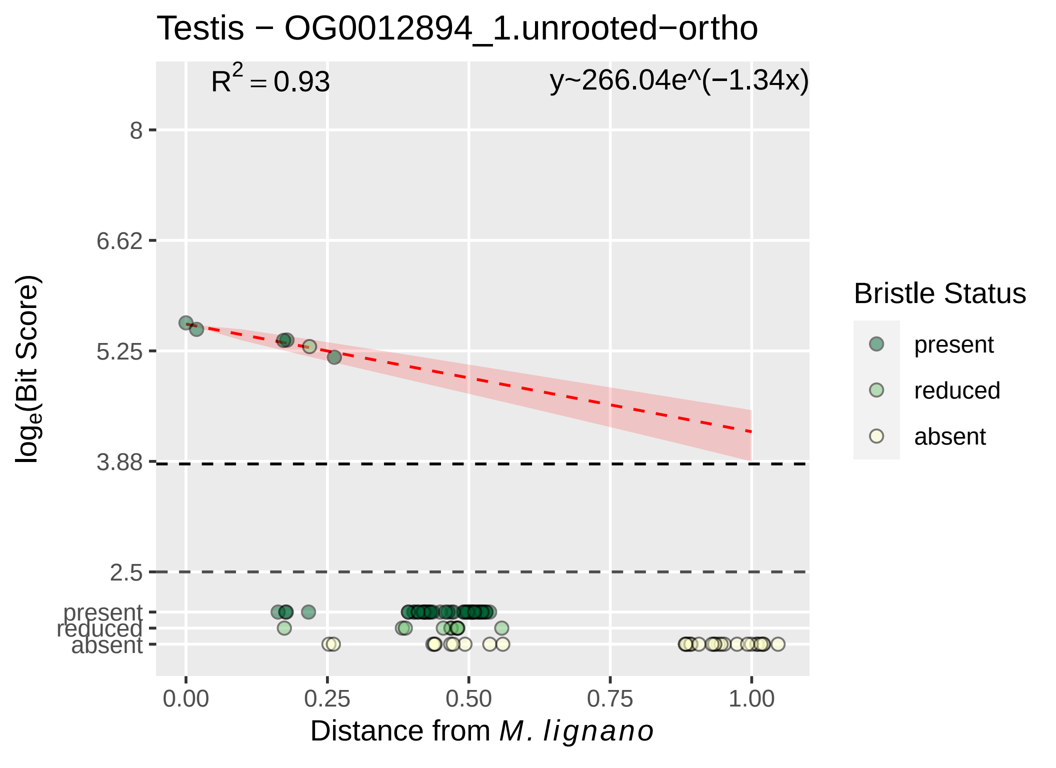

Testis − OG0012894_1.unrooted−ortho
$R^2 = 0.93$
y~266.04e^(−1.34x)
8
6.62
5.25
3.88
2.5
present
reduced
absent
log$_e$(Bit Score)
0.00
0.25
0.50
0.75
1.00
Distance from *M. lignano*
Bristle Status
present
reduced
absent

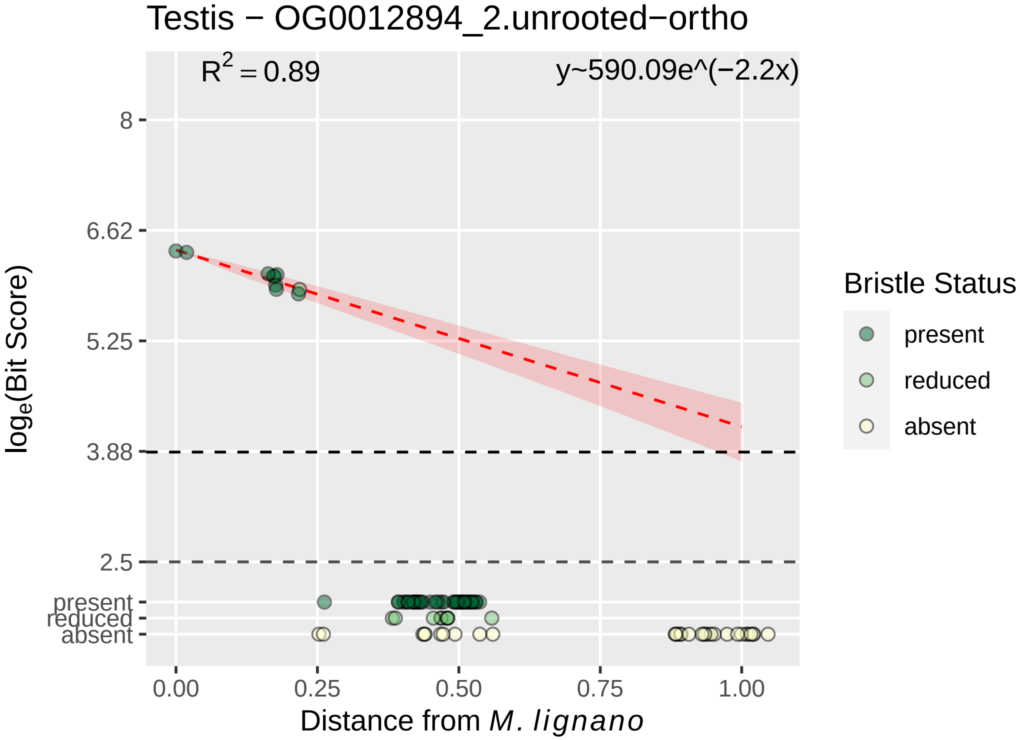
Testis − OG0012894_2.unrooted−ortho
$R^2 = 0.89$
y~590.09e^(−2.2x)
8
6.62
5.25
3.88
2.5
present
reduced
absent
$\log_e$(Bit Score)
0.00
0.25
0.50
0.75
1.00
Distance from *M. lignano*
Bristle Status
present
reduced
absent

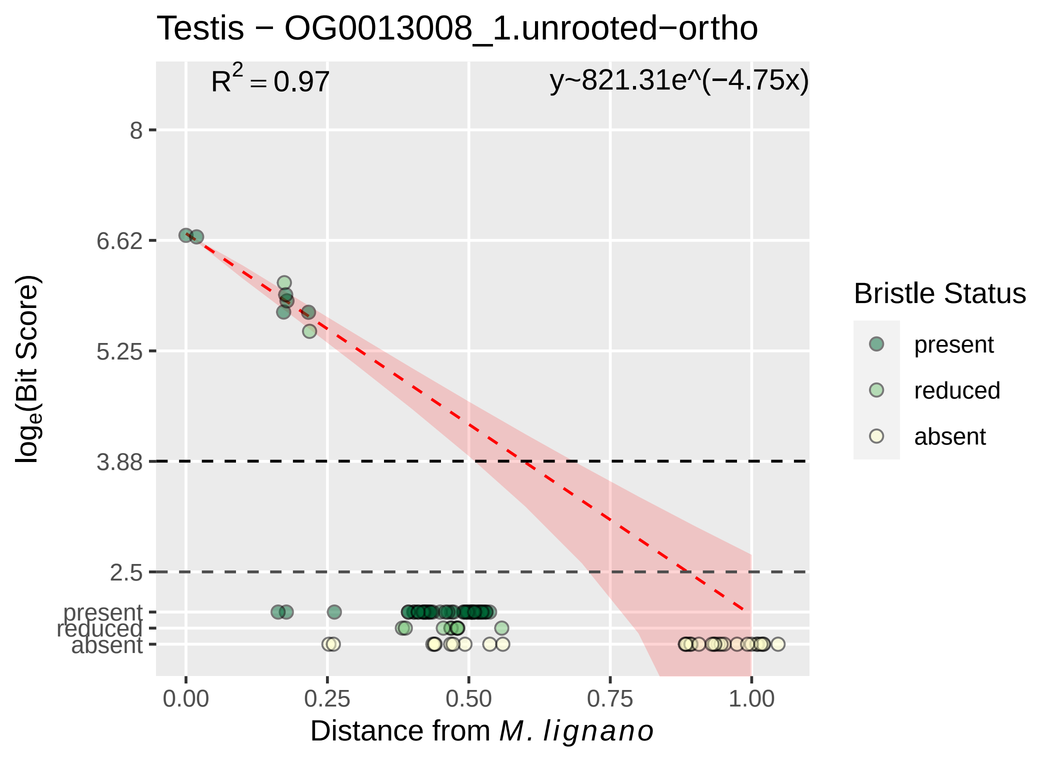

Testis − OG0013008_1.unrooted−ortho
$R^2 = 0.97$
y~821.31e^(−4.75x)
$\log_e$(Bit Score)
8
6.62
5.25
3.88
2.5
present
reduced
absent
0.00
0.25
0.50
0.75
1.00
Distance from *M. lignano*
Bristle Status
present
reduced
absent

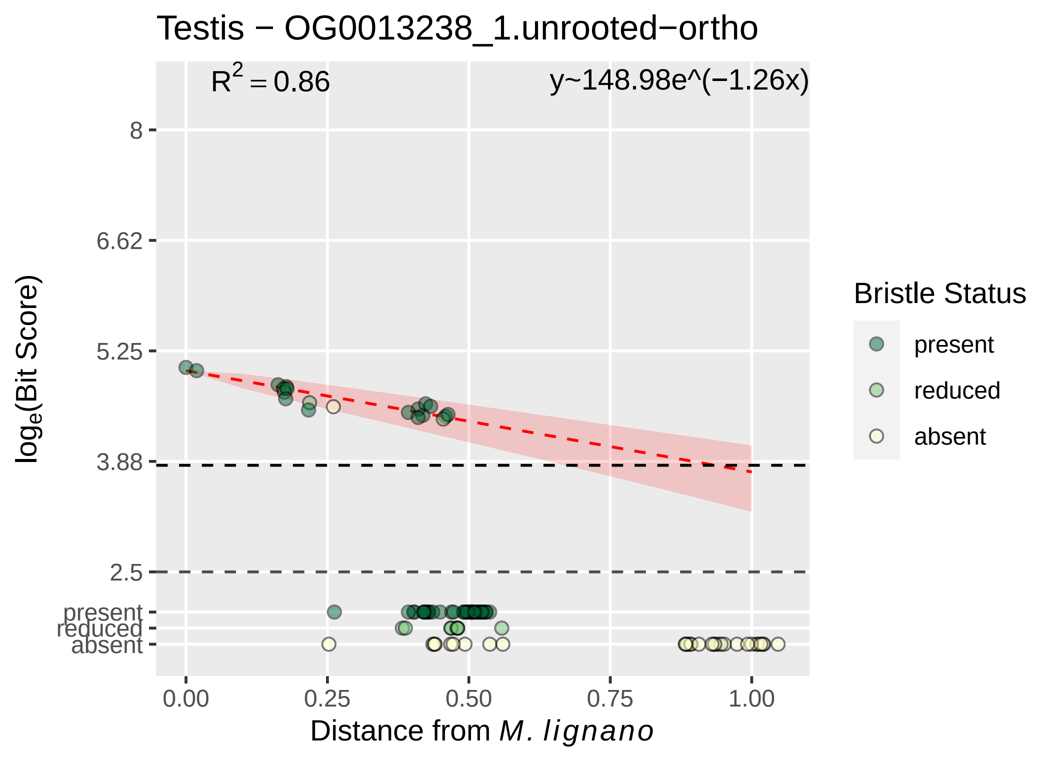

Testis − OG0013238_1.unrooted−ortho
$R^2 = 0.86$
y~148.98e^(−1.26x)
8
6.62
5.25
3.88
2.5
present
reduced
absent
0.00
0.25
0.50
0.75
1.00
Distance from *M. lignano*
$\log_e$(Bit Score)
Bristle Status
present
reduced
absent

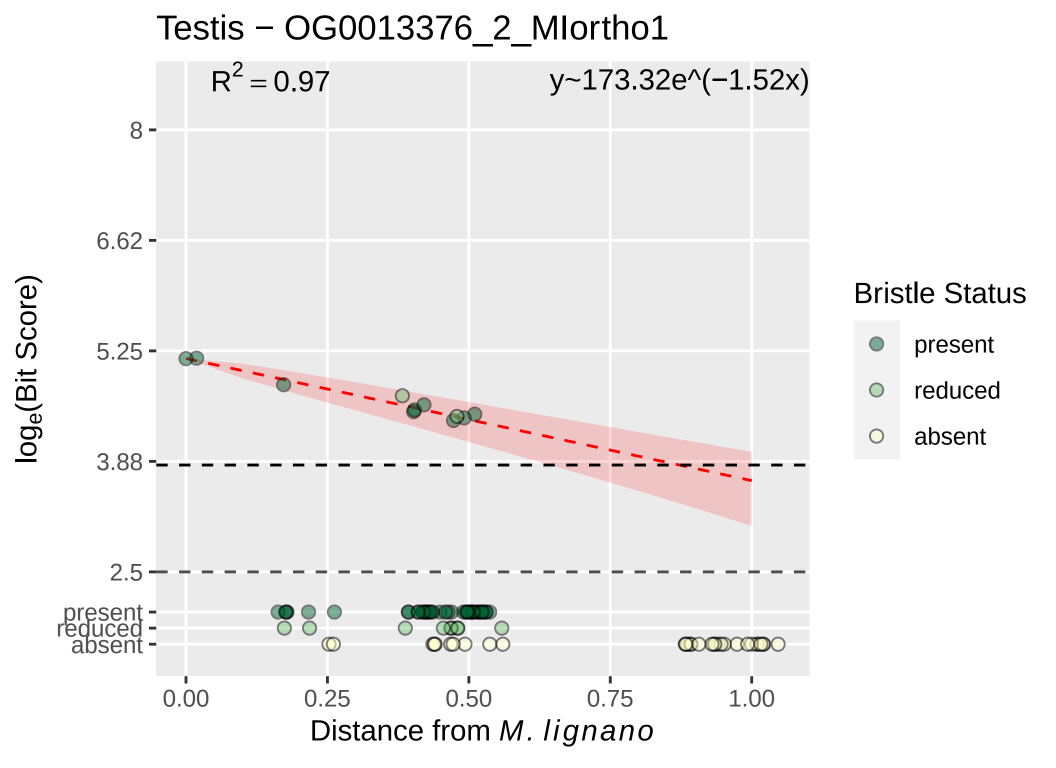

Testis − OG0013376_2_MIortho1
R^2 = 0.97
y~173.32e^(−1.52x)
log_e(Bit Score)
8
6.62
5.25
3.88
2.5
present
reduced
absent
0.00
0.25
0.50
0.75
1.00
Distance from M. lignano
Bristle Status
present
reduced
absent

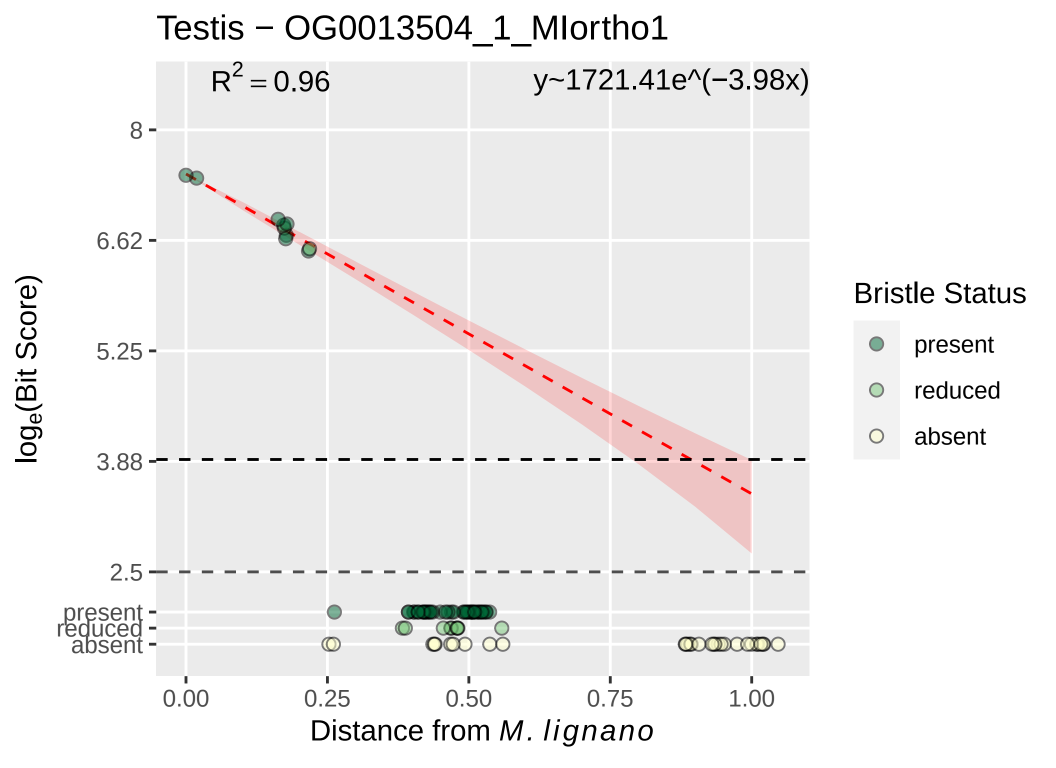

Testis − OG0013504_1_MIortho1
$R^2 = 0.96$
y~1721.41e^(−3.98x)
8
6.62
5.25
3.88
2.5
present
reduced
absent
0.00
0.25
0.50
0.75
1.00
Distance from *M. lignano*
$log_e$(Bit Score)
Bristle Status
present
reduced
absent

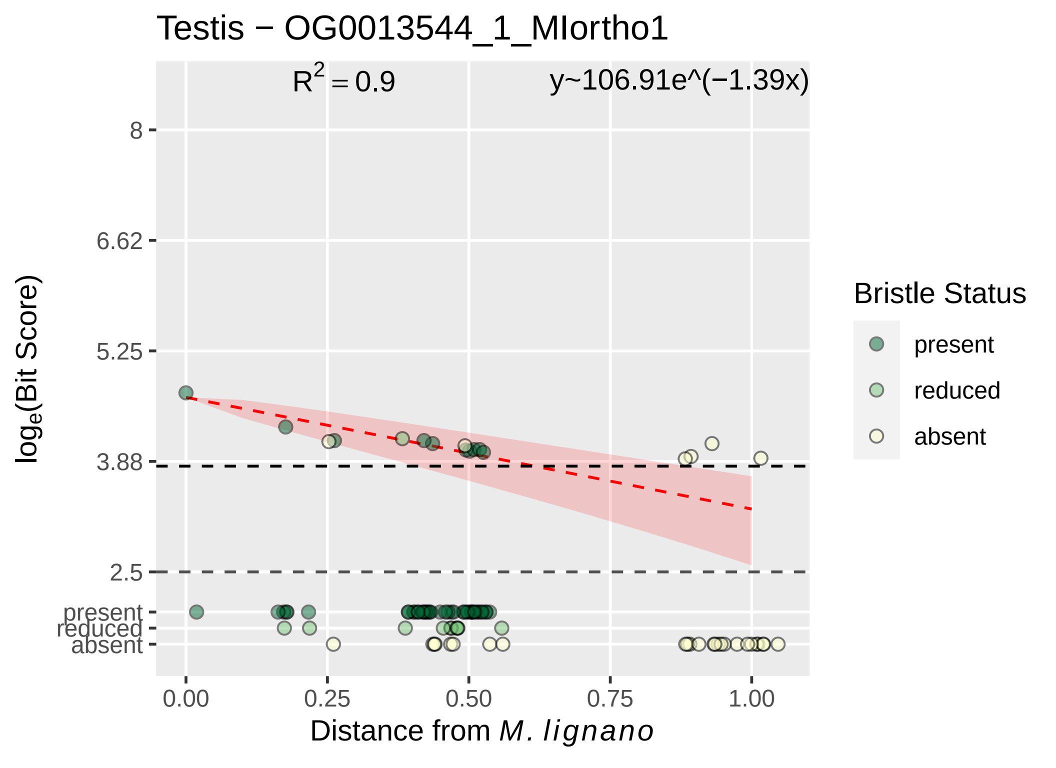

Testis − OG0013544_1_MIortho1
$R^2 = 0.9$
y~106.91e^(−1.39x)
8
6.62
5.25
3.88
2.5
present
reduced
absent
log$_e$(Bit Score)
0.00
0.25
0.50
0.75
1.00
Distance from *M. lignano*
Bristle Status
present
reduced
absent

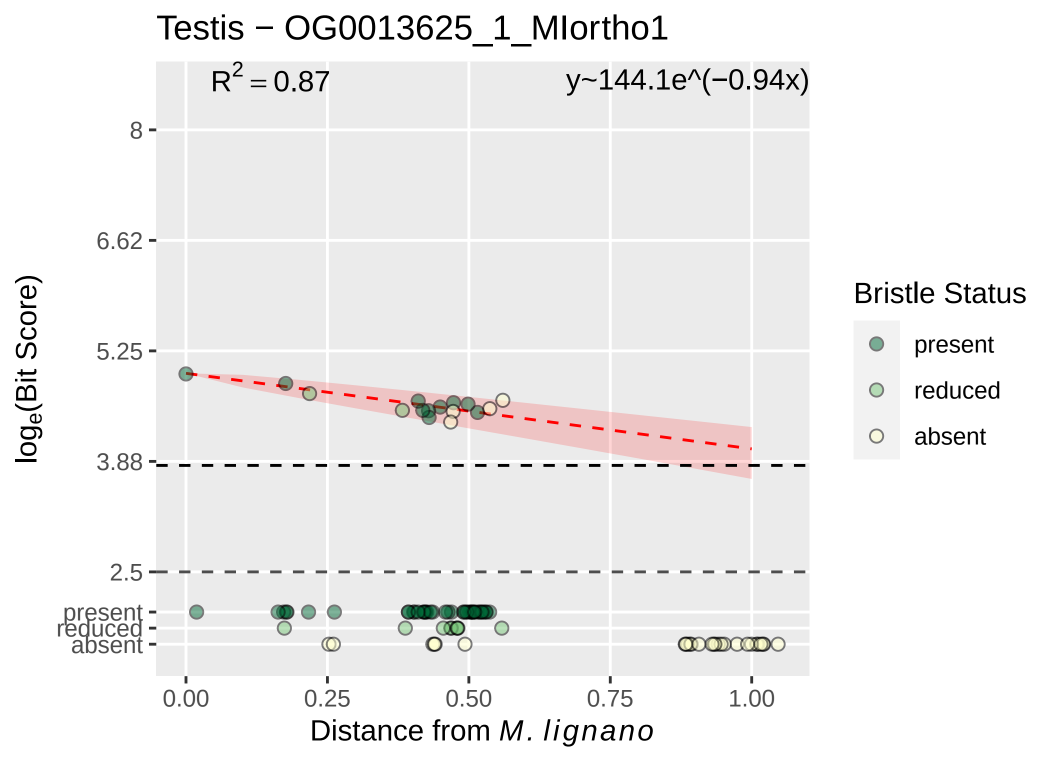

Testis − OG0013625_1_MIortho1
R2 = 0.87
y~144.1e^(−0.94x)
8
6.62
5.25
3.88
2.5
present
reduced
absent
log e(Bit Score)
0.00
0.25
0.50
0.75
1.00
Distance from M. lignano
Bristle Status
present
reduced
absent

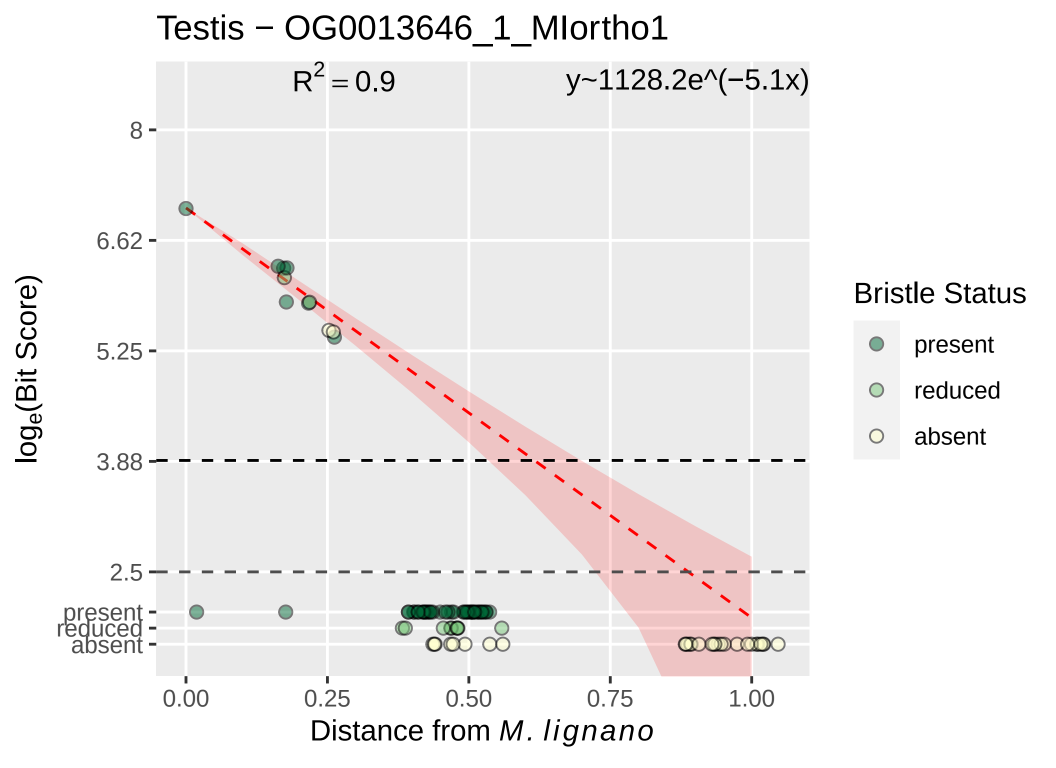

Testis − OG0013646_1_MIortho1
R^2 = 0.9
y~1128.2e^(−5.1x)
8
6.62
5.25
3.88
2.5
present
reduced
absent
0.00
0.25
0.50
0.75
1.00
log_e(Bit Score)
Distance from M. lignano
Bristle Status
present
reduced
absent

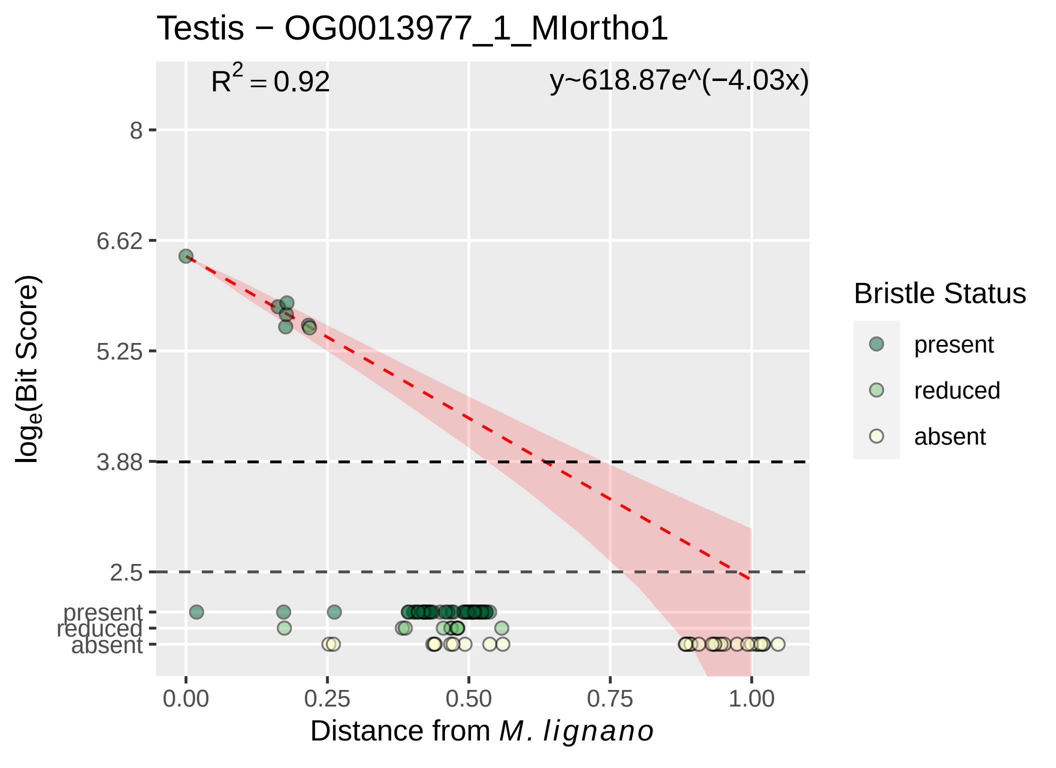

Testis − OG0013977_1_MIortho1
R^2 = 0.92
y~618.87e^(−4.03x)
8
6.62
5.25
3.88
2.5
present
reduced
absent
log_e(Bit Score)
0.00
0.25
0.50
0.75
1.00
Distance from M. lignano
Bristle Status
present
reduced
absent

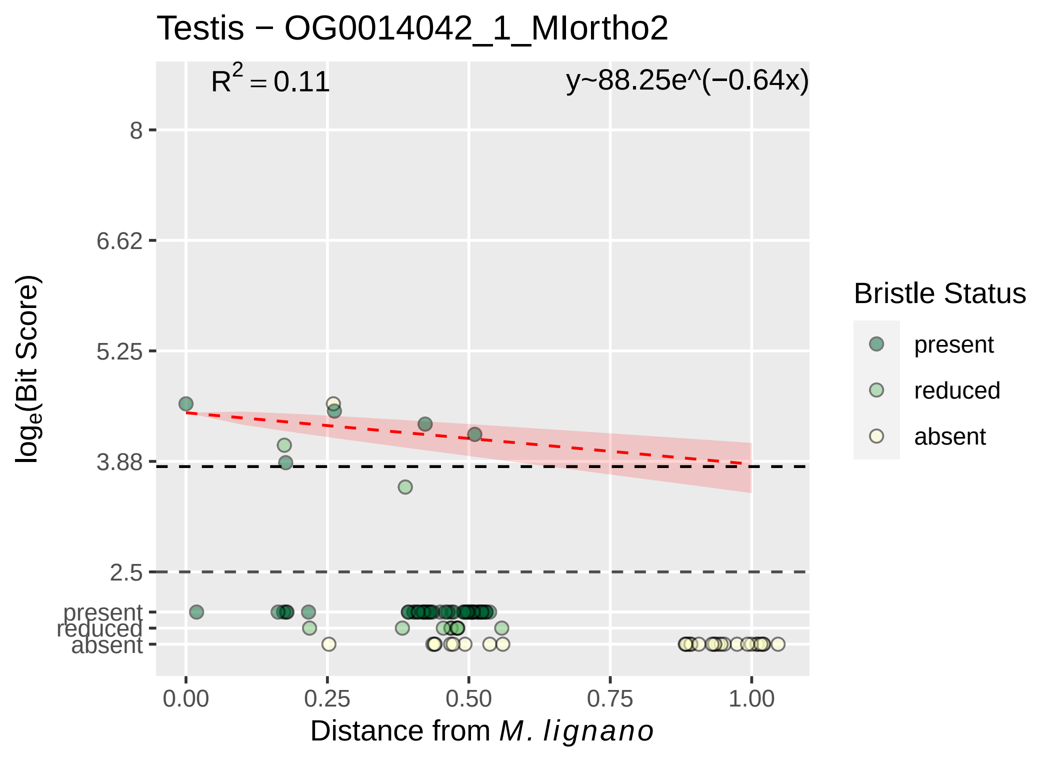

Testis − OG0014042_1_MIortho2
$R^2 = 0.11$
y~88.25e^(−0.64x)
8
6.62
5.25
3.88
2.5
present
reduced
absent
$\log_e$(Bit Score)
0.00
0.25
0.50
0.75
1.00
Distance from *M. lignano*
Bristle Status
present
reduced
absent

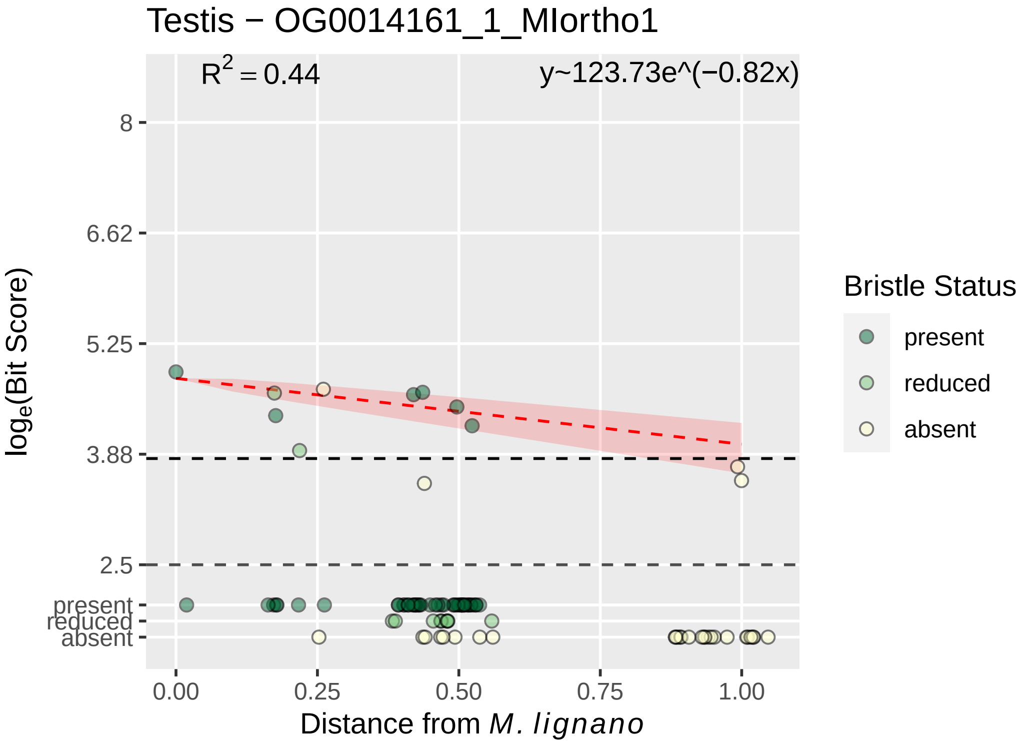

Testis − OG0014161_1_MIortho1
$R^2 = 0.44$
y~123.73e^(−0.82x)
8
6.62
5.25
3.88
2.5
present
reduced
absent
0.00
0.25
0.50
0.75
1.00
Distance from *M. lignano*
$\log_e$(Bit Score)
Bristle Status
present
reduced
absent

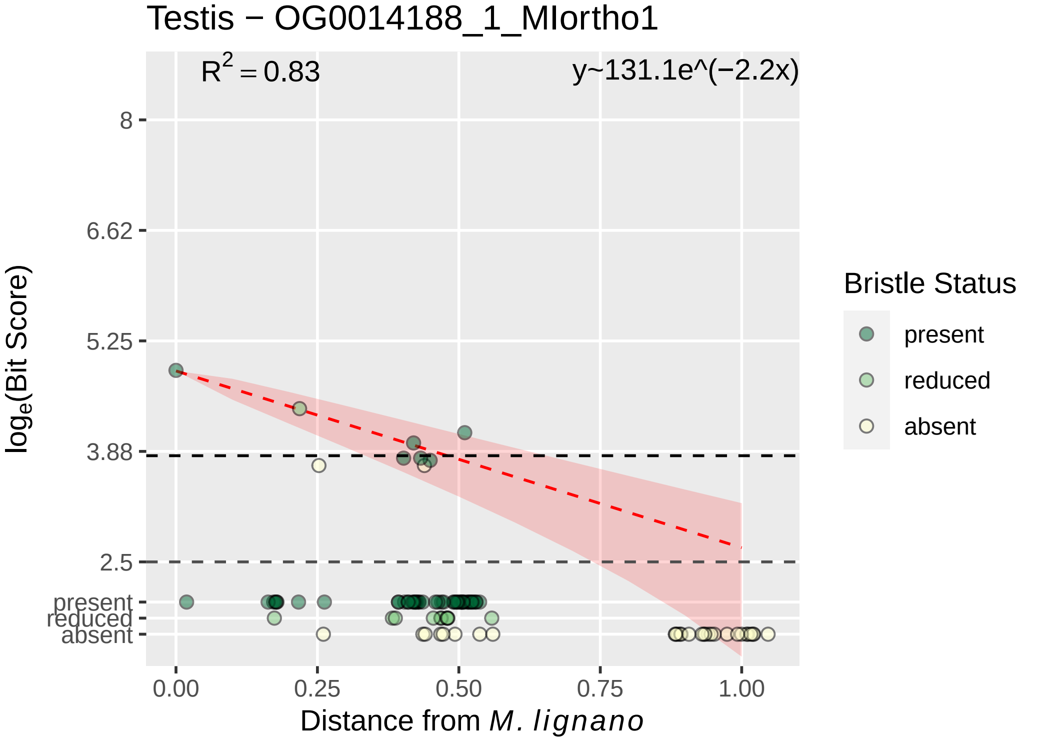

Testis − OG0014188_1_MIortho1
$R^2 = 0.83$
y~131.1e^(−2.2x)
8
6.62
5.25
3.88
2.5
present
reduced
absent
$log_e$(Bit Score)
0.00
0.25
0.50
0.75
1.00
Distance from M. lignano
Bristle Status
present
reduced
absent

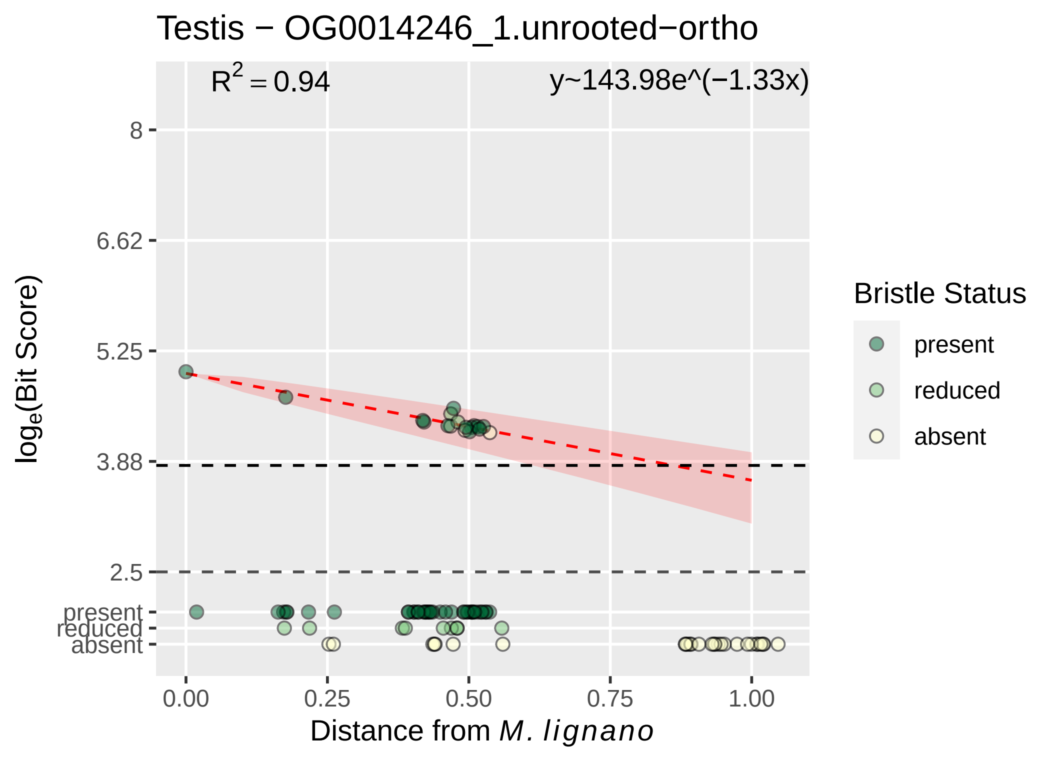

Testis − OG0014246_1.unrooted−ortho
$R^2 = 0.94$
y~143.98e^(−1.33x)
8
6.62
5.25
3.88
2.5
present
reduced
absent
log_e(Bit Score)
0.00
0.25
0.50
0.75
1.00
Distance from M. lignano
Bristle Status
present
reduced
absent

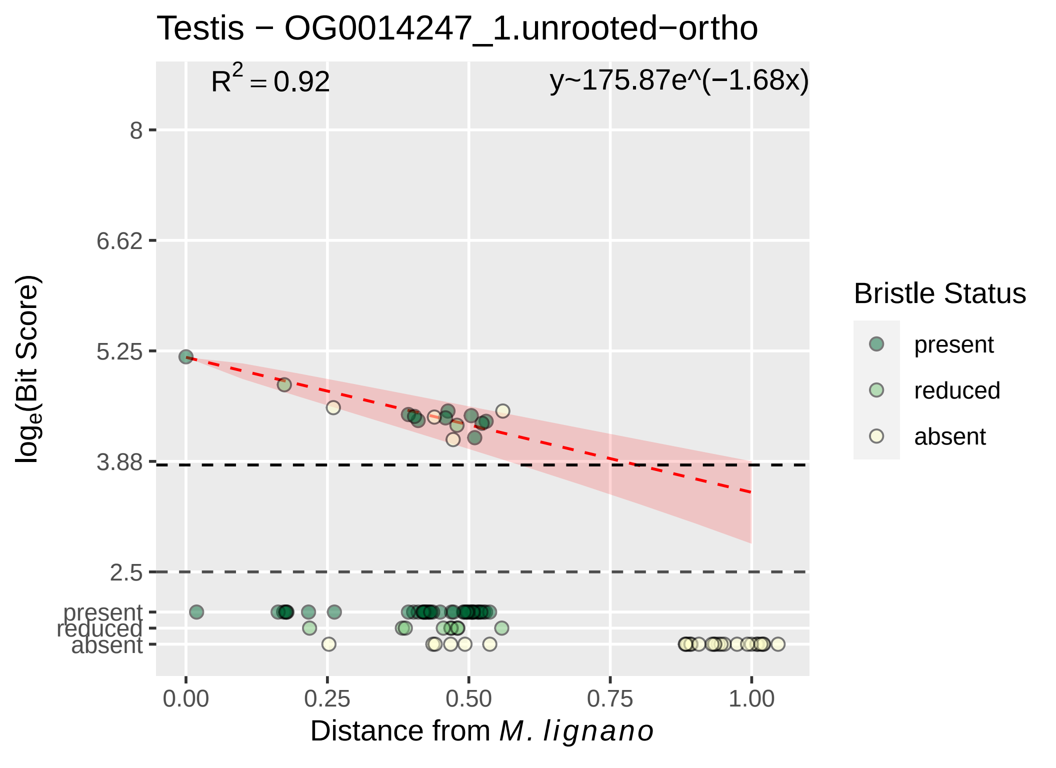

Testis − OG0014247_1.unrooted−ortho
$R^2 = 0.92$
y~175.87e^(−1.68x)
8
6.62
5.25
3.88
2.5
present
reduced
absent
$\log_e$(Bit Score)
0.00
0.25
0.50
0.75
1.00
Distance from *M. lignano*
Bristle Status
present
reduced
absent

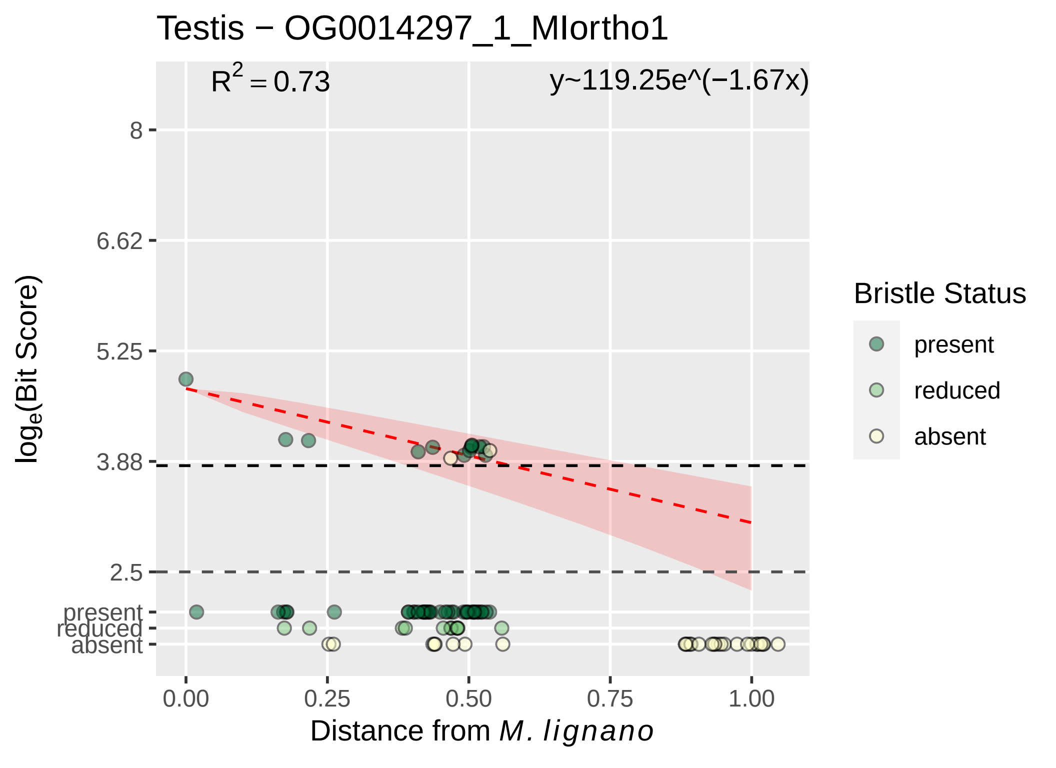

Testis − OG0014297_1_MIortho1
$R^2 = 0.73$
y~119.25e^(−1.67x)
8
6.62
5.25
3.88
2.5
present
reduced
absent
log$_e$(Bit Score)
0.00
0.25
0.50
0.75
1.00
Distance from *M. lignano*
Bristle Status
present
reduced
absent

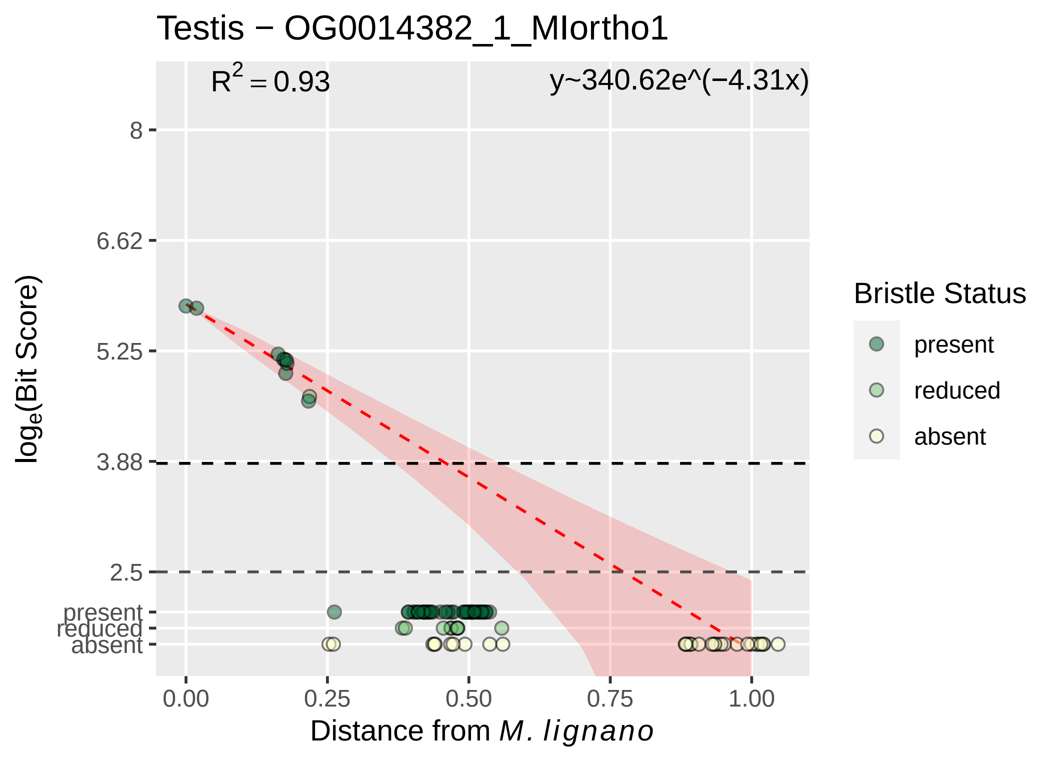

Testis − OG0014382_1_MIortho1
R² = 0.93
y~340.62e^(−4.31x)
8
6.62
5.25
3.88
2.5
present
reduced
absent
0.00
0.25
0.50
0.75
1.00
Distance from M. lignano
loge(Bit Score)
Bristle Status
present
reduced
absent

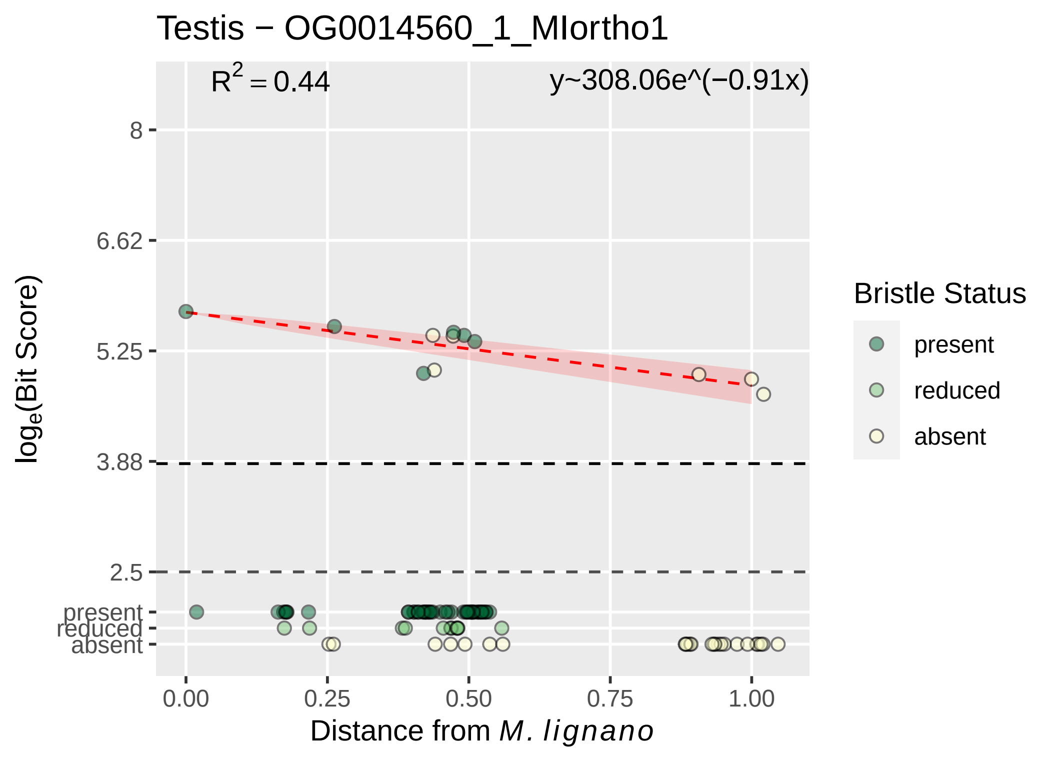
Testis − OG0014560_1_MIortho1
R^2 = 0.44
y~308.06e^(−0.91x)
8
6.62
5.25
3.88
2.5
present
reduced
absent
log_e(Bit Score)
0.00
0.25
0.50
0.75
1.00
Distance from M. lignano
Bristle Status
present
reduced
absent

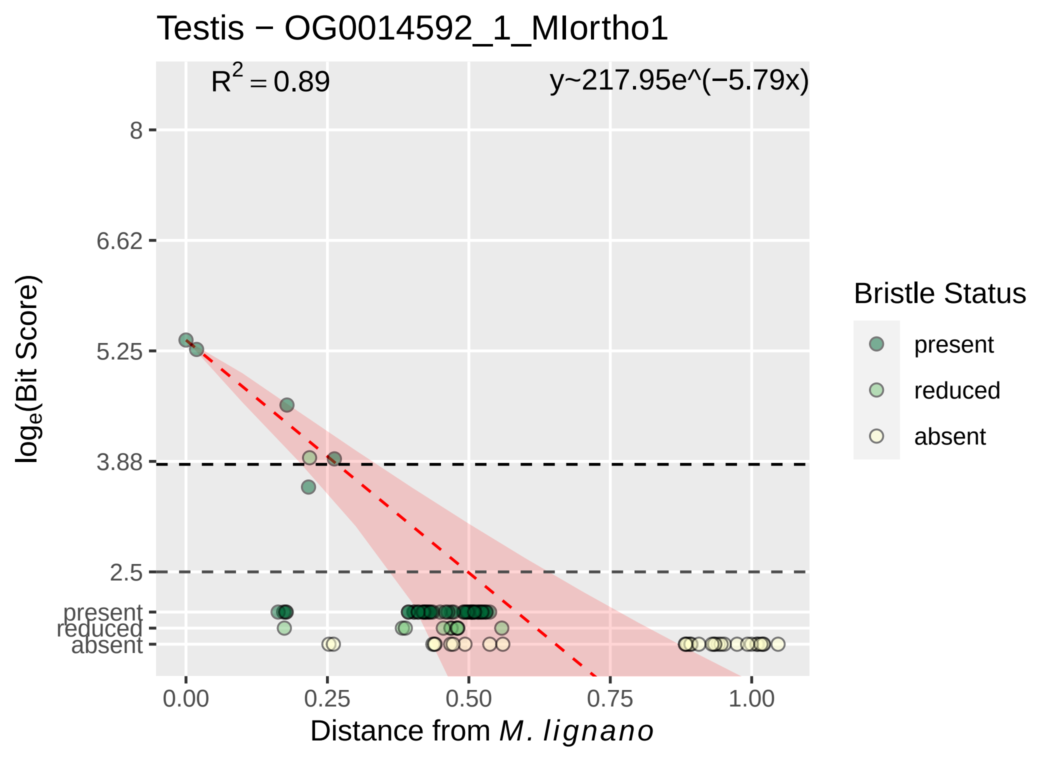
Testis − OG0014592_1_MIortho1
$R^2 = 0.89$
y~217.95e^(−5.79x)
8
6.62
5.25
3.88
2.5
present
reduced
absent
$\log_e$(Bit Score)
0.00
0.25
0.50
0.75
1.00
Distance from *M. lignano*
Bristle Status
present
reduced
absent

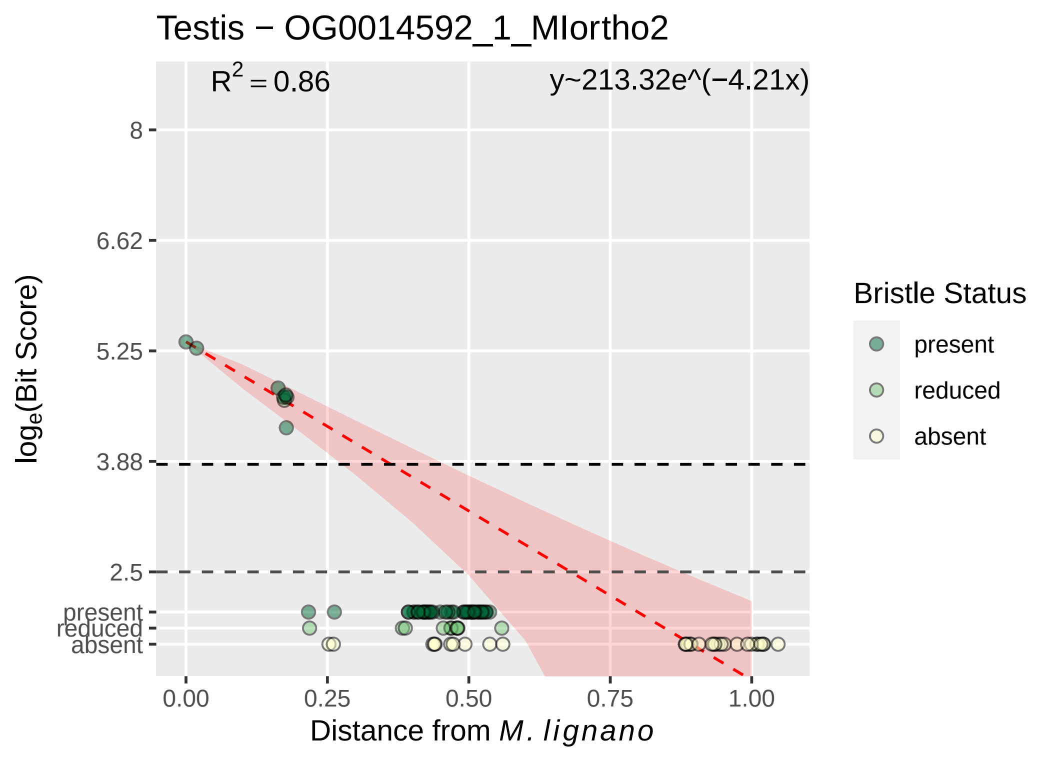

Testis − OG0014592_1_MIortho2
R^2 = 0.86
y~213.32e^(−4.21x)
8
6.62
5.25
3.88
2.5
present
reduced
absent
0.00
0.25
0.50
0.75
1.00
Distance from M. lignano
loge(Bit Score)
Bristle Status
present
reduced
absent

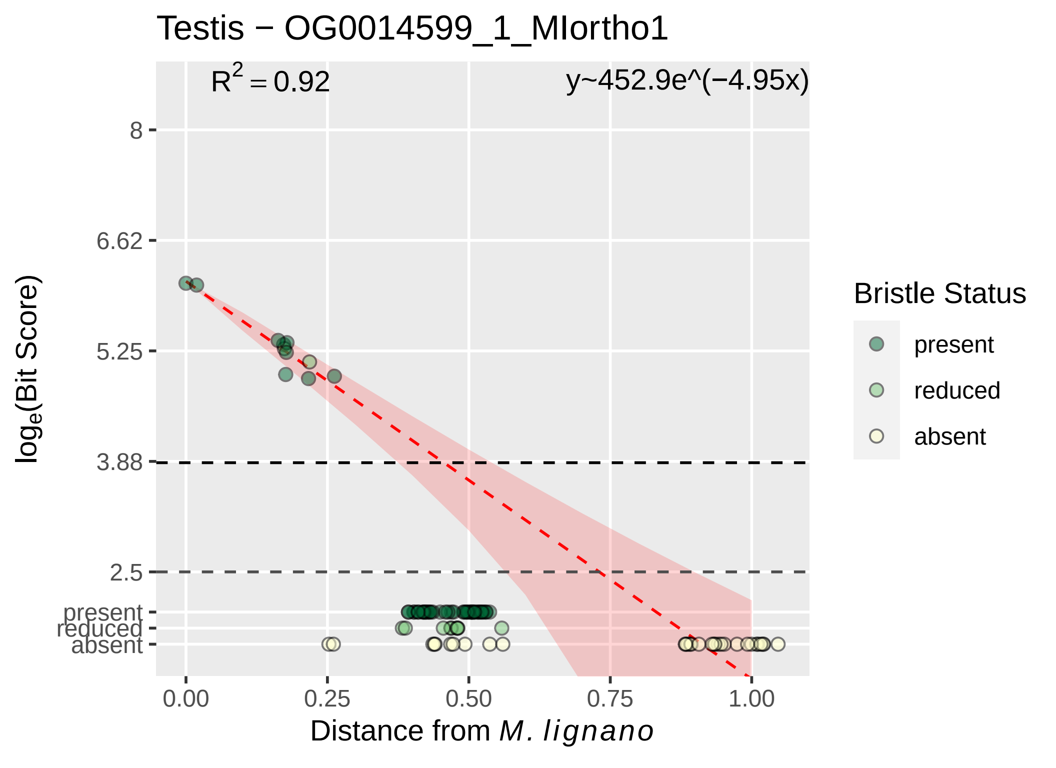

Testis − OG0014599_1_MIortho1
$R^2 = 0.92$
y~452.9e^(−4.95x)
8
6.62
5.25
3.88
2.5
present
reduced
absent
$\log_e$(Bit Score)
0.00
0.25
0.50
0.75
1.00
Distance from *M. lignano*
Bristle Status
present
reduced
absent

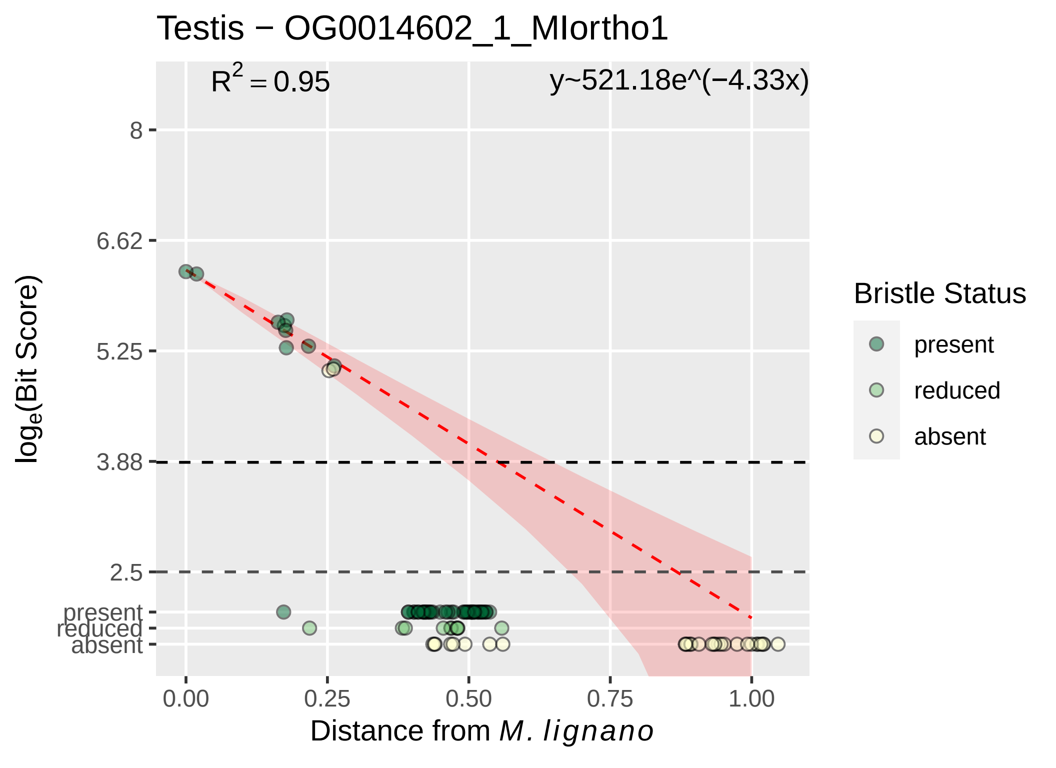

Testis − OG0014602_1_MIortho1
$R^2 = 0.95$
y~521.18e^(−4.33x)
8
6.62
5.25
3.88
2.5
present
reduced
absent
$\log_e$(Bit Score)
0.00
0.25
0.50
0.75
1.00
Distance from *M. lignano*
Bristle Status
present
reduced
absent

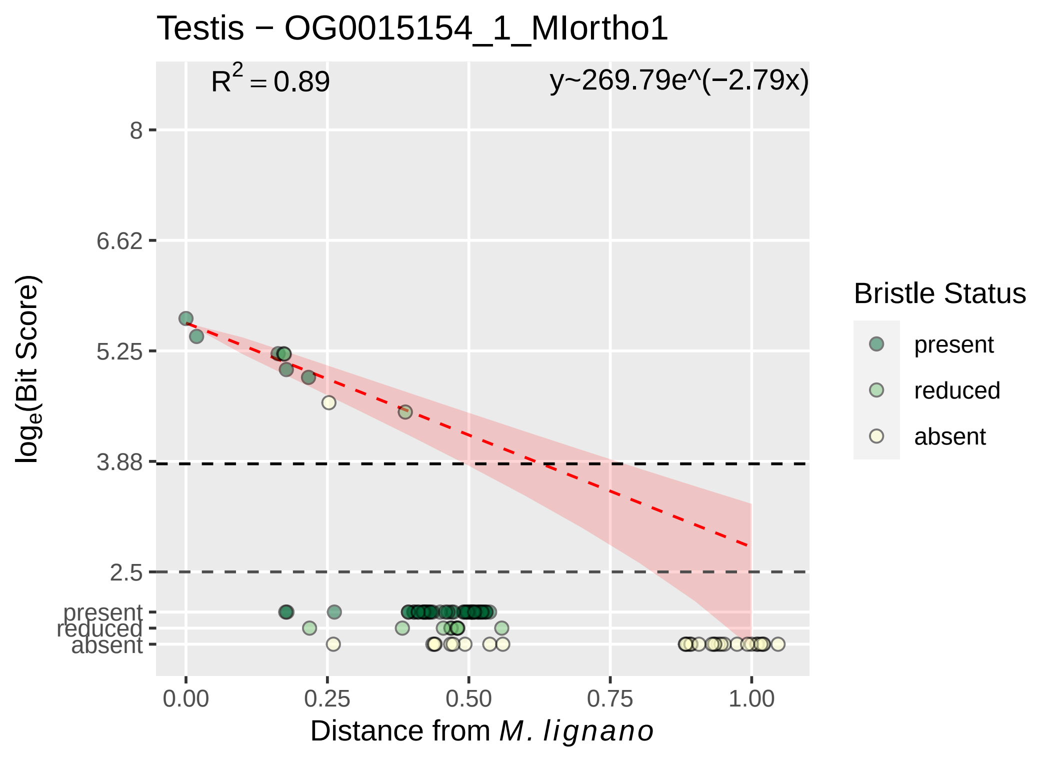

Testis − OG0015154_1_MIortho1
$R^2 = 0.89$
y~269.79e^(−2.79x)
8
6.62
5.25
3.88
2.5
present
reduced
absent
log$_e$(Bit Score)
0.00
0.25
0.50
0.75
1.00
Distance from *M. lignano*
Bristle Status
present
reduced
absent

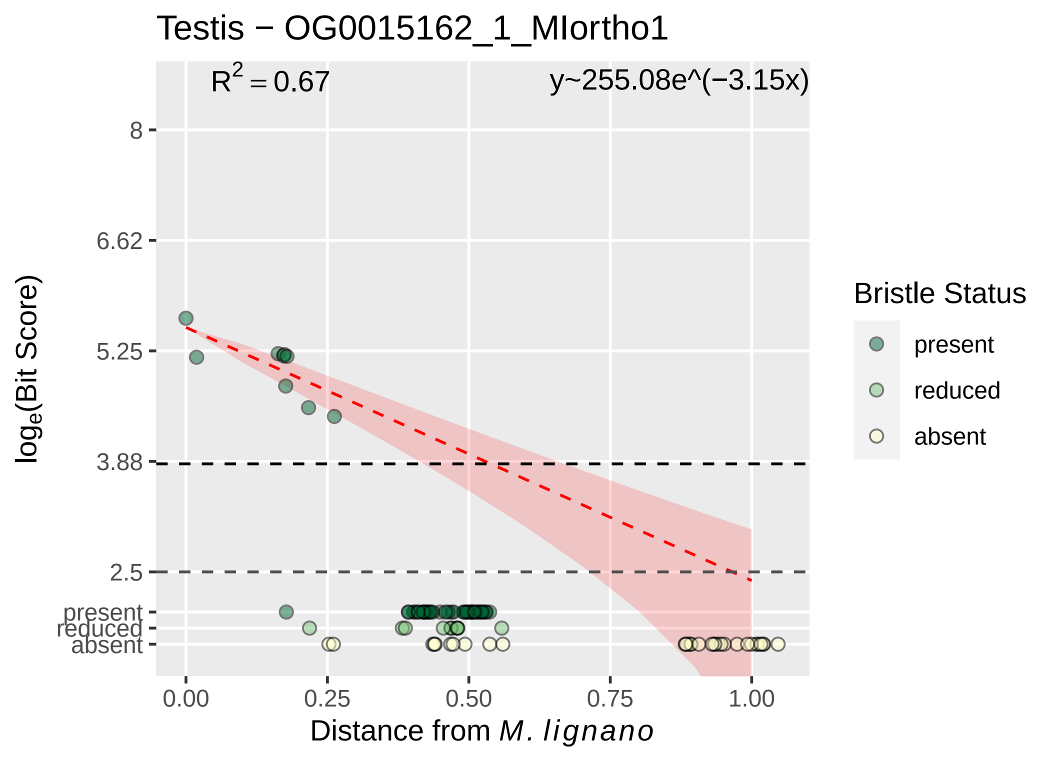

Testis − OG0015162_1_MIortho1
R² = 0.67
y~255.08e^(−3.15x)
8
6.62
5.25
3.88
2.5
present
reduced
absent
log_e(Bit Score)
0.00
0.25
0.50
0.75
1.00
Distance from M. lignano
Bristle Status
present
reduced
absent

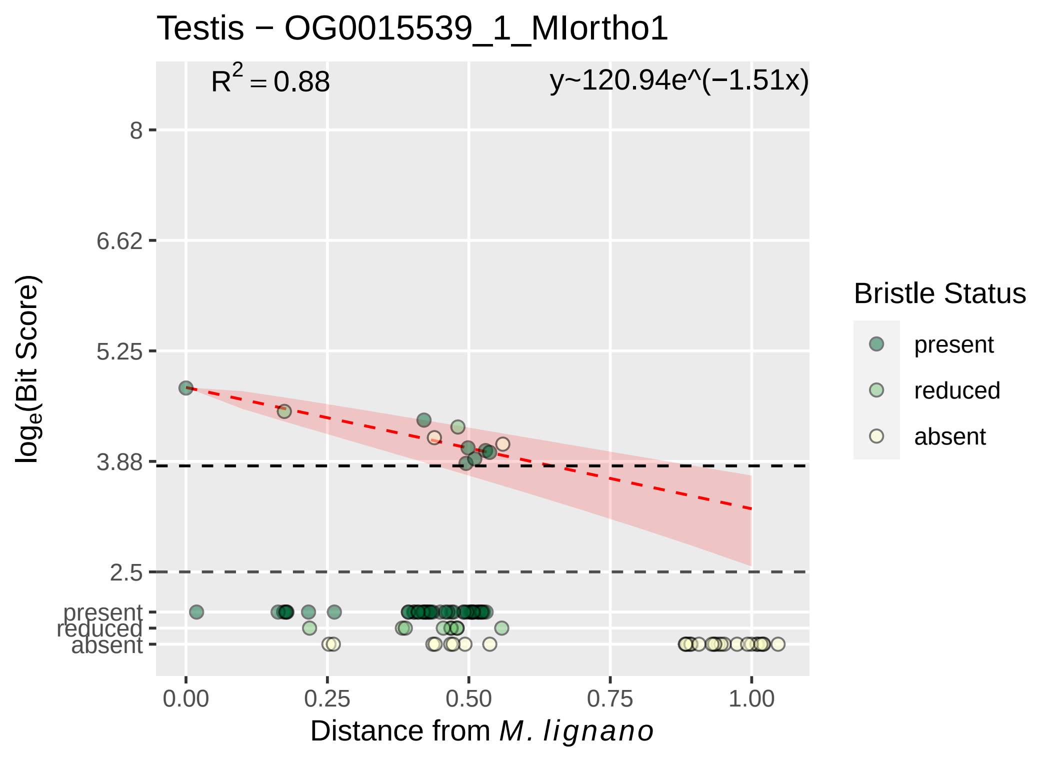

Testis − OG0015539_1_MIortho1
R² = 0.88
y~120.94e^(−1.51x)
8
6.62
5.25
3.88
2.5
present
reduced
absent
log_e(Bit Score)
0.00
0.25
0.50
0.75
1.00
Distance from M. lignano
Bristle Status
present
reduced
absent

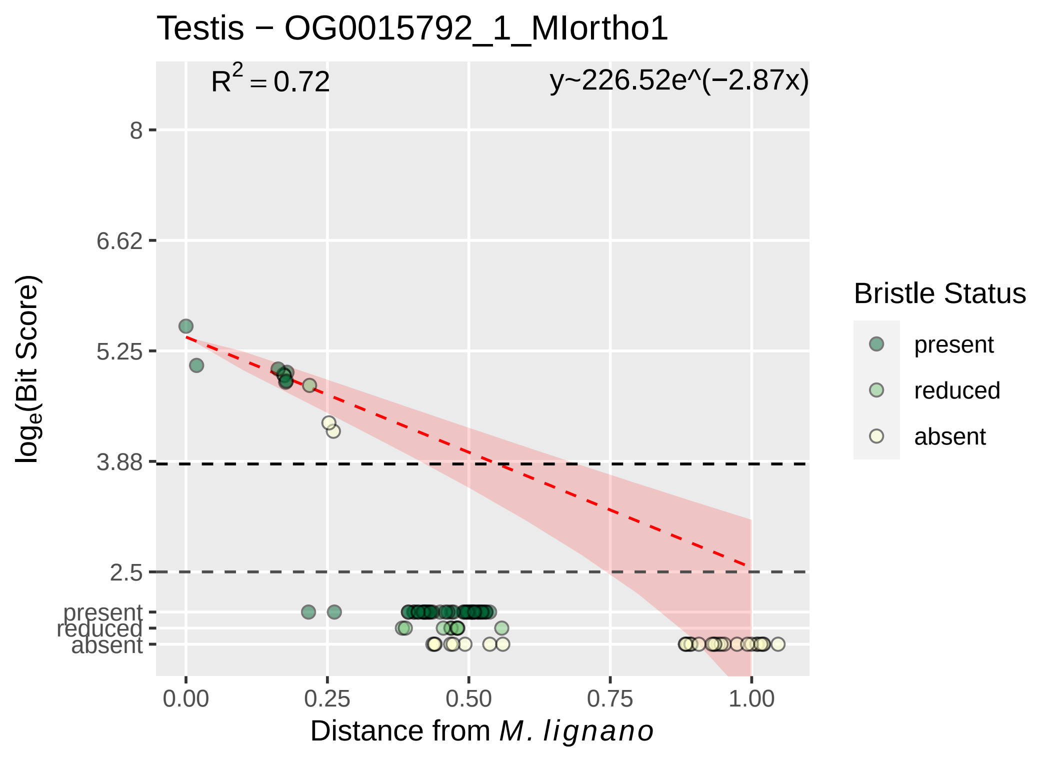

Testis − OG0015792_1_MIortho1
$R^2 = 0.72$
y~226.52e^(−2.87x)
8
6.62
5.25
3.88
2.5
present
reduced
absent
log$_e$(Bit Score)
0.00
0.25
0.50
0.75
1.00
Distance from *M. lignano*
Bristle Status
present
reduced
absent

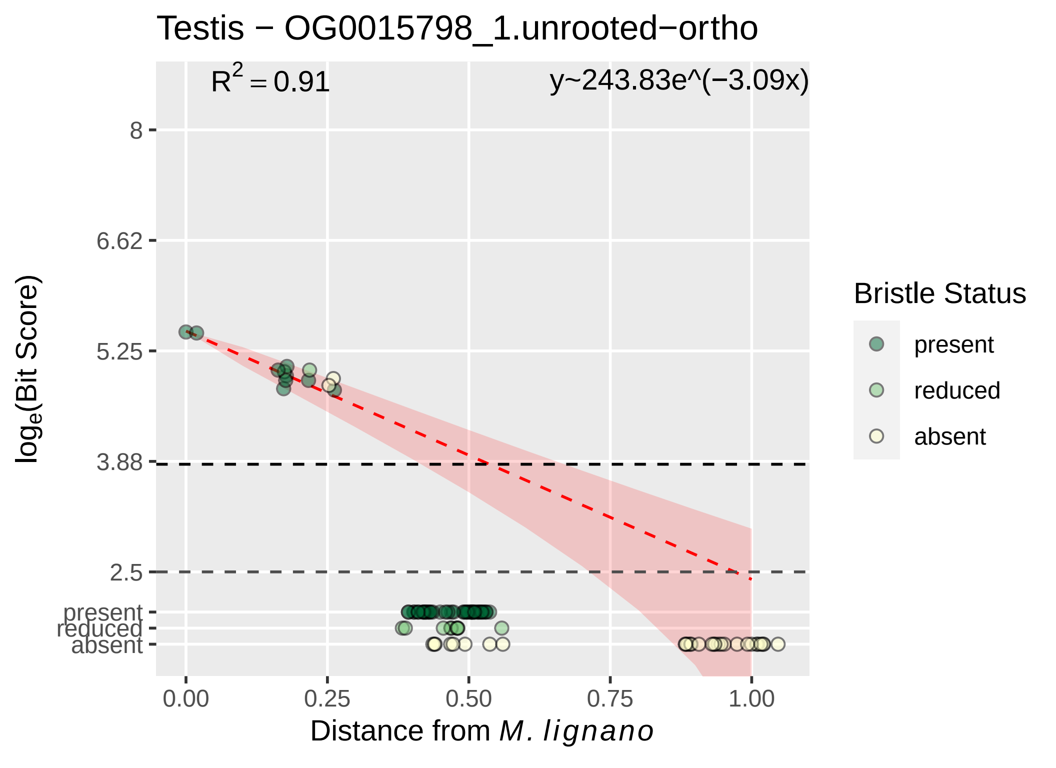

Testis − OG0015798_1.unrooted−ortho
$R^2 = 0.91$
y~243.83e^(−3.09x)
8
6.62
5.25
3.88
2.5
present
reduced
absent
$\log_e$(Bit Score)
0.00
0.25
0.50
0.75
1.00
Distance from *M. lignano*
Bristle Status
present
reduced
absent

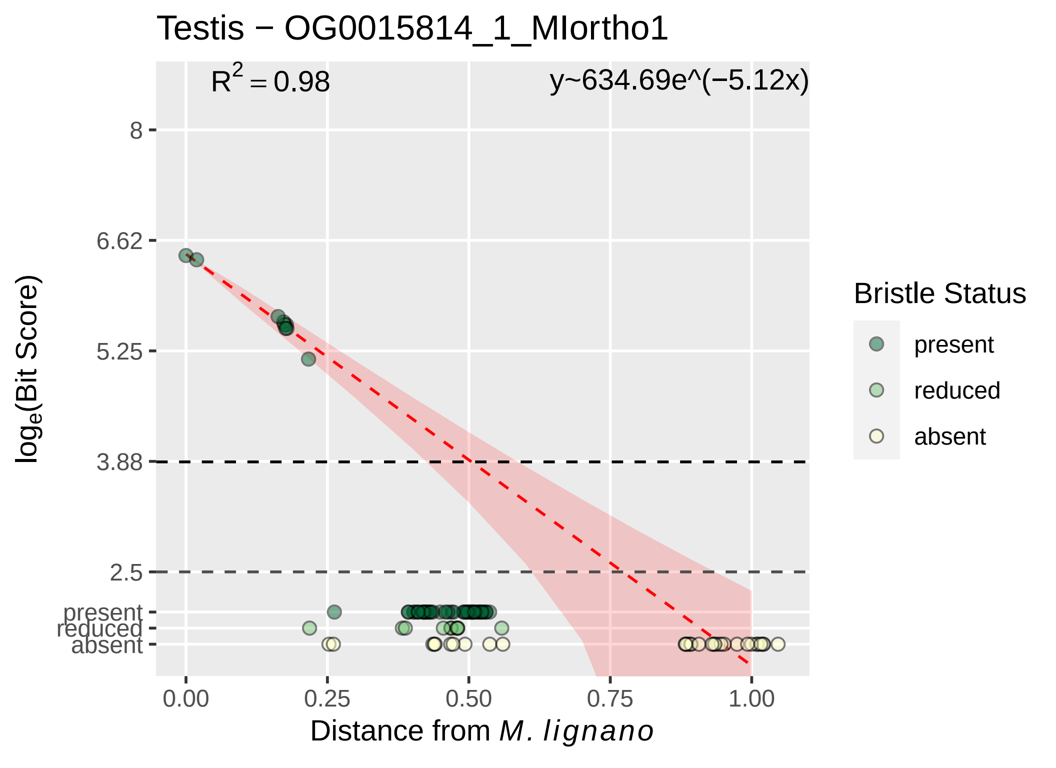

Testis − OG0015814_1_MIortho1
R^2 = 0.98
y~634.69e^(−5.12x)
8
6.62
5.25
3.88
2.5
present
reduced
absent
log_e(Bit Score)
0.00
0.25
0.50
0.75
1.00
Distance from M. lignano
Bristle Status
present
reduced
absent

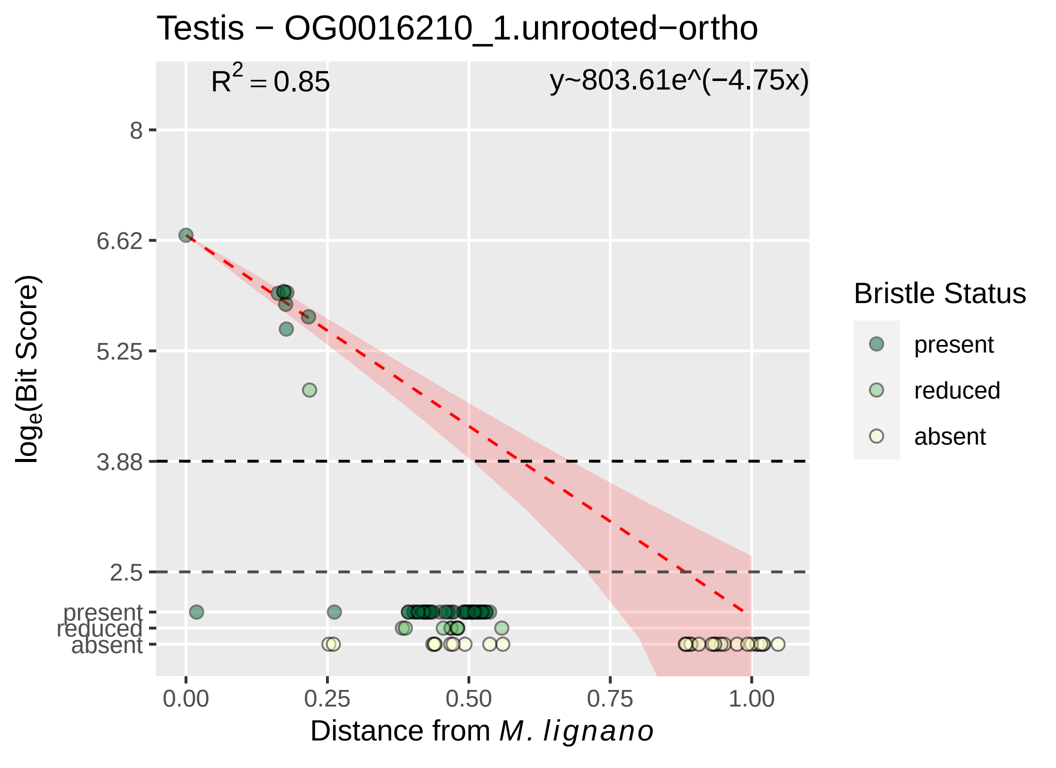

Testis − OG0016210_1.unrooted−ortho
$R^2 = 0.85$
y~803.61e^(−4.75x)
8
6.62
5.25
3.88
2.5
present
reduced
absent
0.00
0.25
0.50
0.75
1.00
Distance from M. lignano
loge(Bit Score)
Bristle Status
present
reduced
absent

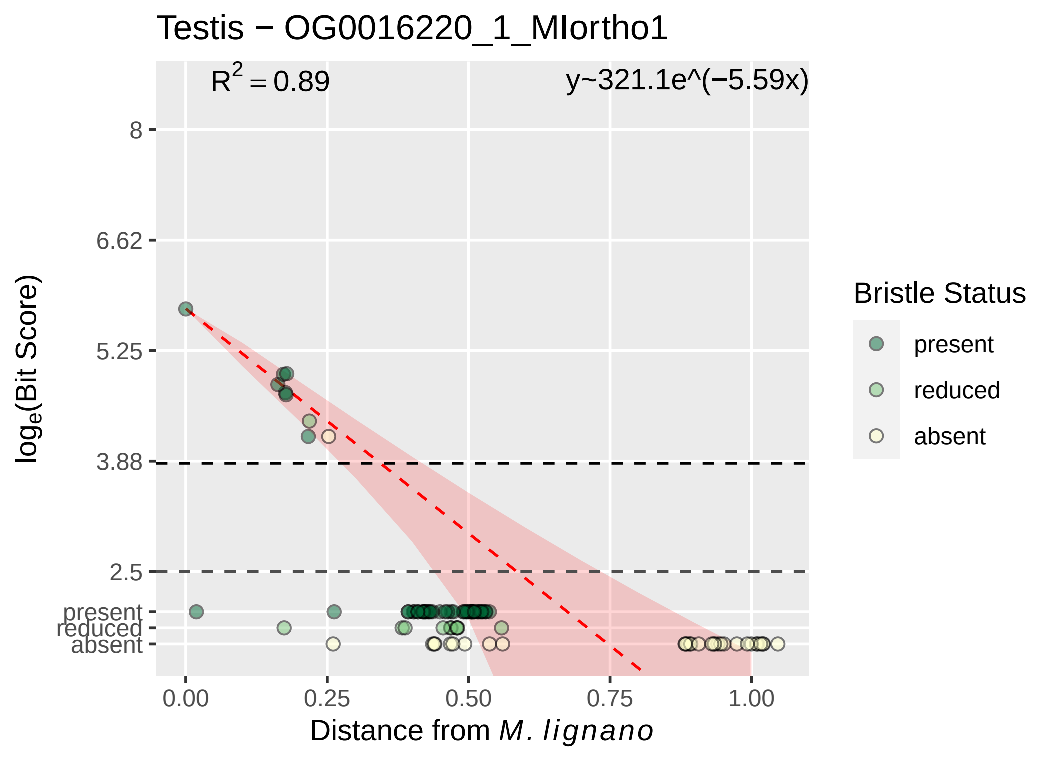

Testis − OG0016220_1_MIortho1
$R^2 = 0.89$
y~321.1e^(−5.59x)
8
6.62
5.25
3.88
2.5
present
reduced
absent
$\log_e$(Bit Score)
0.00
0.25
0.50
0.75
1.00
Distance from *M. lignano*
Bristle Status
present
reduced
absent

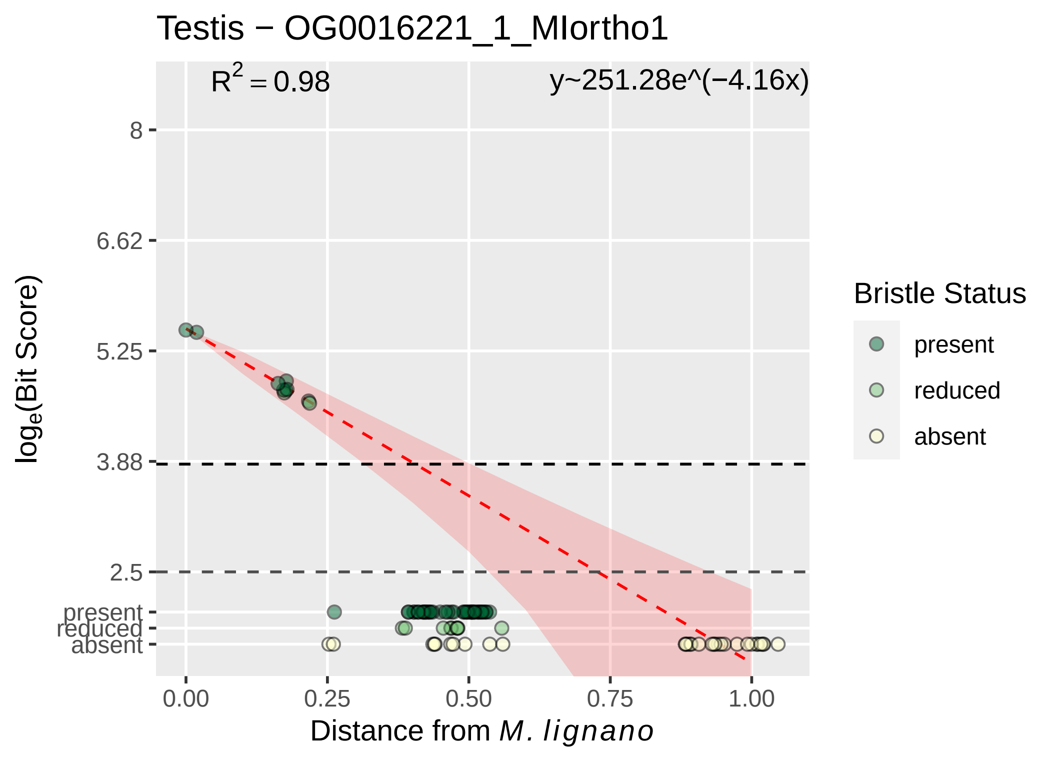

Testis − OG0016221_1_MIortho1
$R^2 = 0.98$
y~251.28e^(−4.16x)
8
6.62
5.25
3.88
2.5
present
reduced
absent
log_e(Bit Score)
0.00
0.25
0.50
0.75
1.00
Distance from M. lignano
Bristle Status
present
reduced
absent

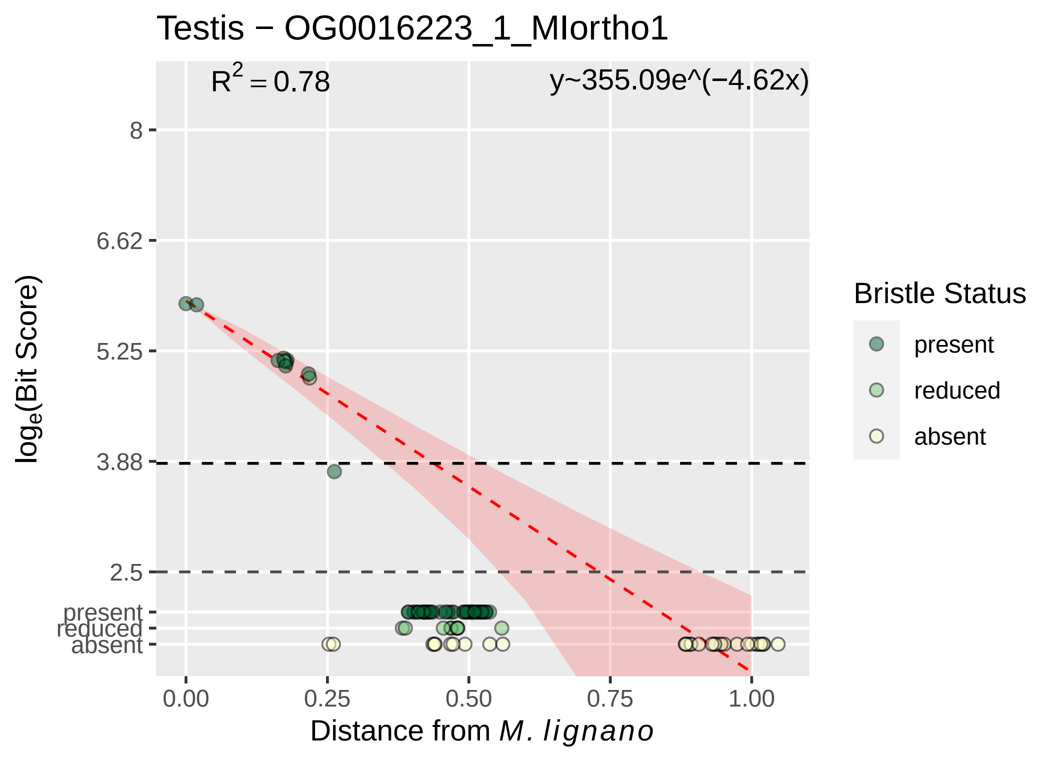

Testis − OG0016223_1_MIortho1
$R^2 = 0.78$
y~355.09e^(−4.62x)
8
6.62
5.25
3.88
2.5
present
reduced
absent
0.00
0.25
0.50
0.75
1.00
$\log_e$(Bit Score)
Distance from *M. lignano*
Bristle Status
present
reduced
absent

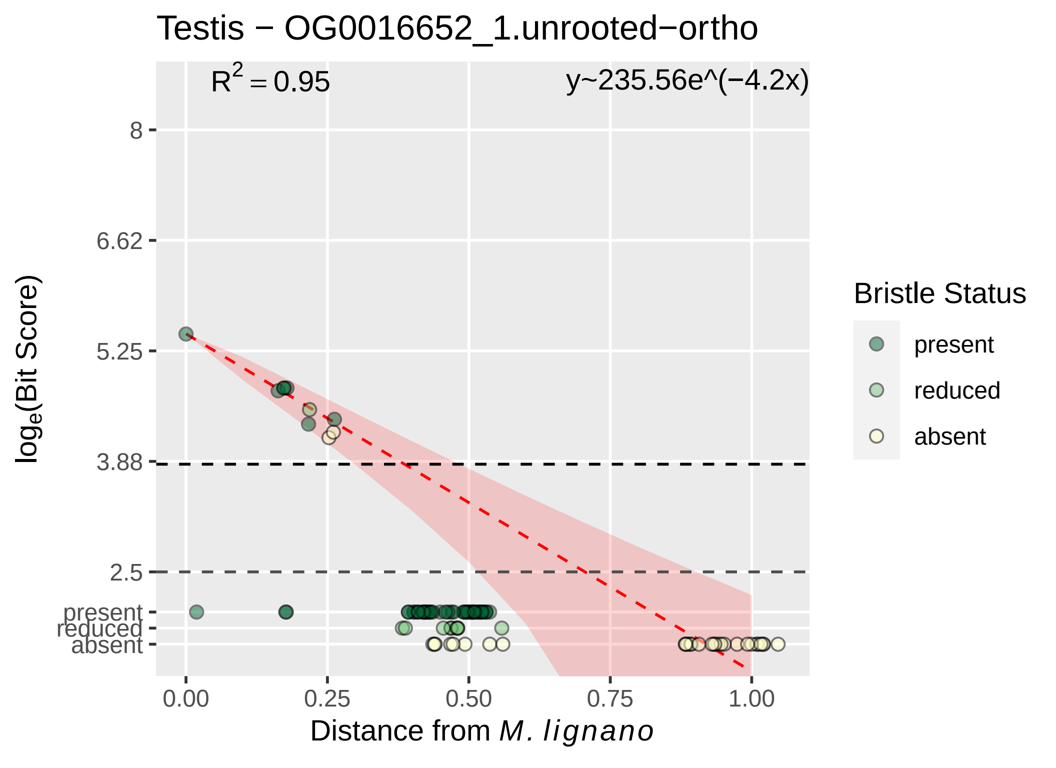

Testis − OG0016652_1.unrooted−ortho
$R^2 = 0.95$
y~235.56e^(−4.2x)
8
6.62
5.25
3.88
2.5
present
reduced
absent
0.00
0.25
0.50
0.75
1.00
$\log_e$(Bit Score)
Distance from *M. lignano*
Bristle Status
present
reduced
absent

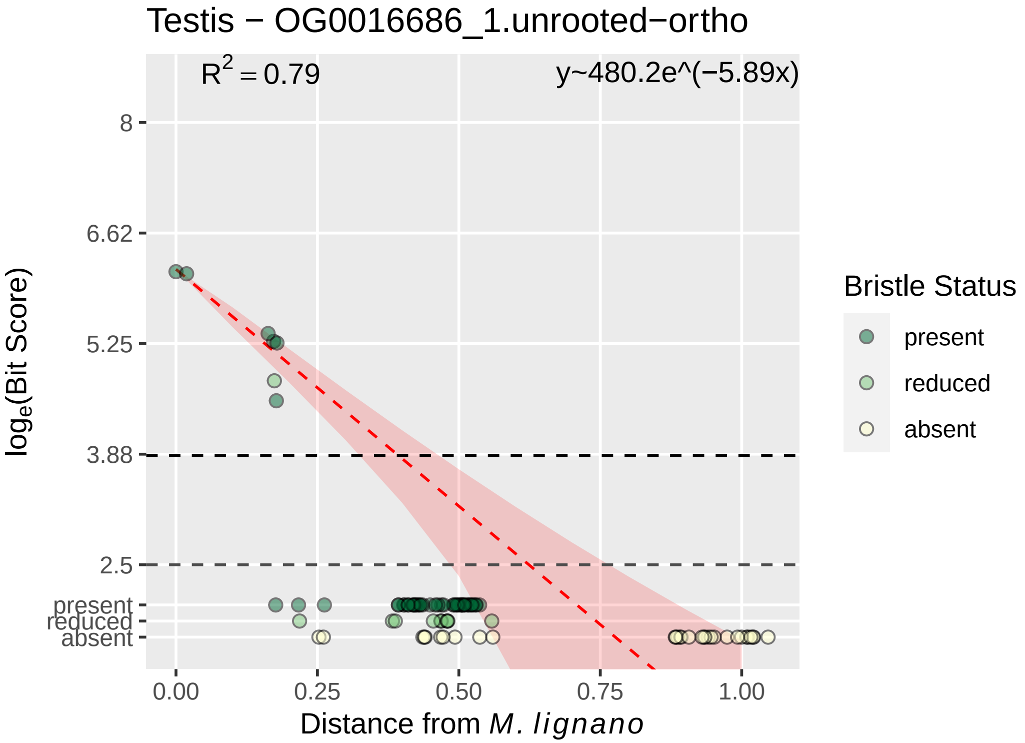

Testis − OG0016686_1.unrooted−ortho
$R^2 = 0.79$
y~480.2e^(−5.89x)
8
6.62
5.25
3.88
2.5
present
reduced
absent
log_e(Bit Score)
0.00
0.25
0.50
0.75
1.00
Distance from M. lignano
Bristle Status
present
reduced
absent

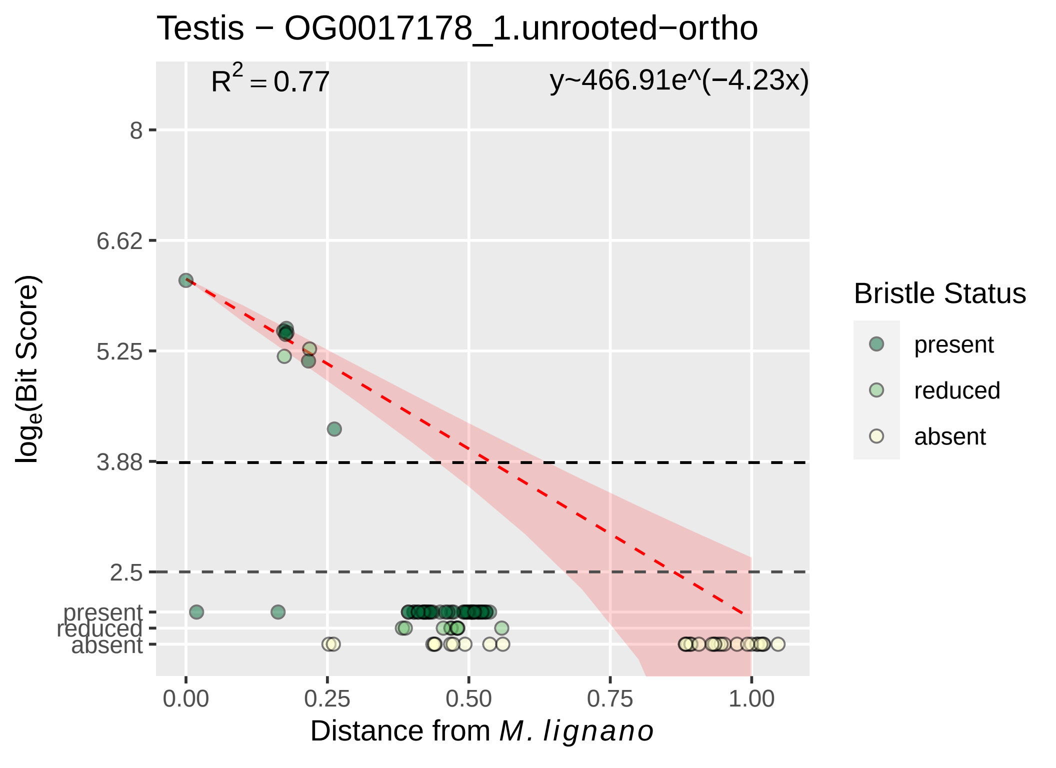

Testis − OG0017178_1.unrooted−ortho
$R^2 = 0.77$
y~466.91e^(−4.23x)
8
6.62
5.25
3.88
2.5
present
reduced
absent
0.00
0.25
0.50
0.75
1.00
Distance from *M. lignano*
$\log_e$(Bit Score)
Bristle Status
present
reduced
absent

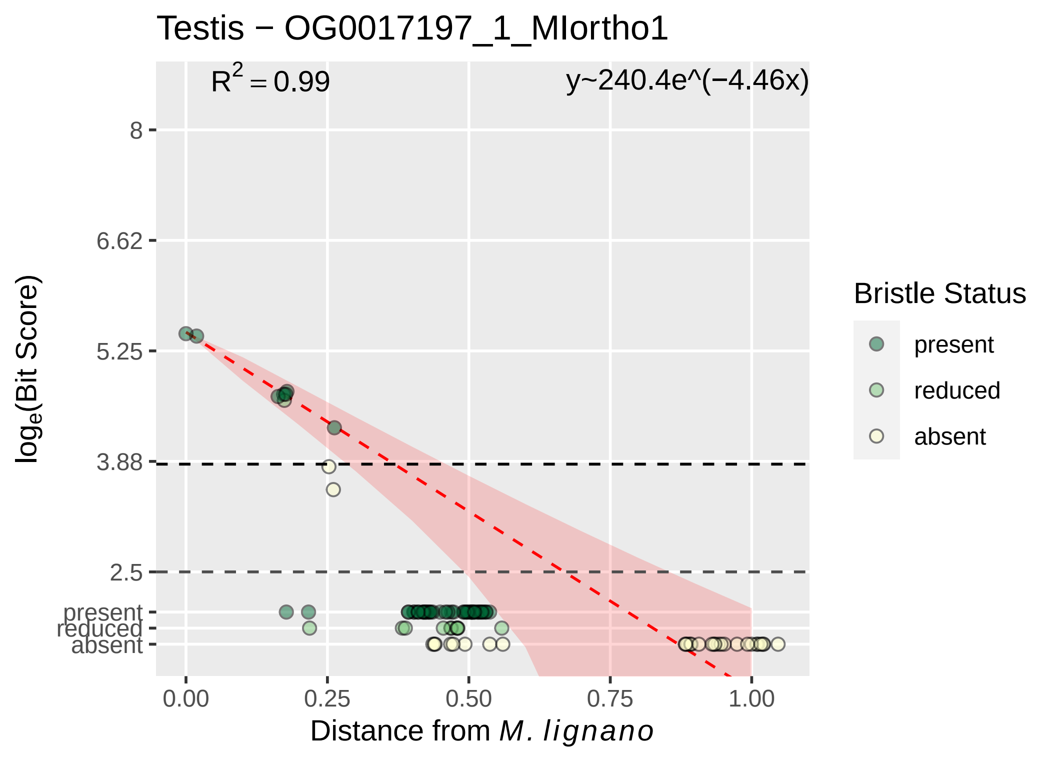

Testis − OG0017197_1_MIortho1
$R^2 = 0.99$
y~240.4e^(−4.46x)
8
6.62
5.25
3.88
2.5
present
reduced
absent
$log_e$(Bit Score)
0.00
0.25
0.50
0.75
1.00
Distance from *M. lignano*
Bristle Status
present
reduced
absent

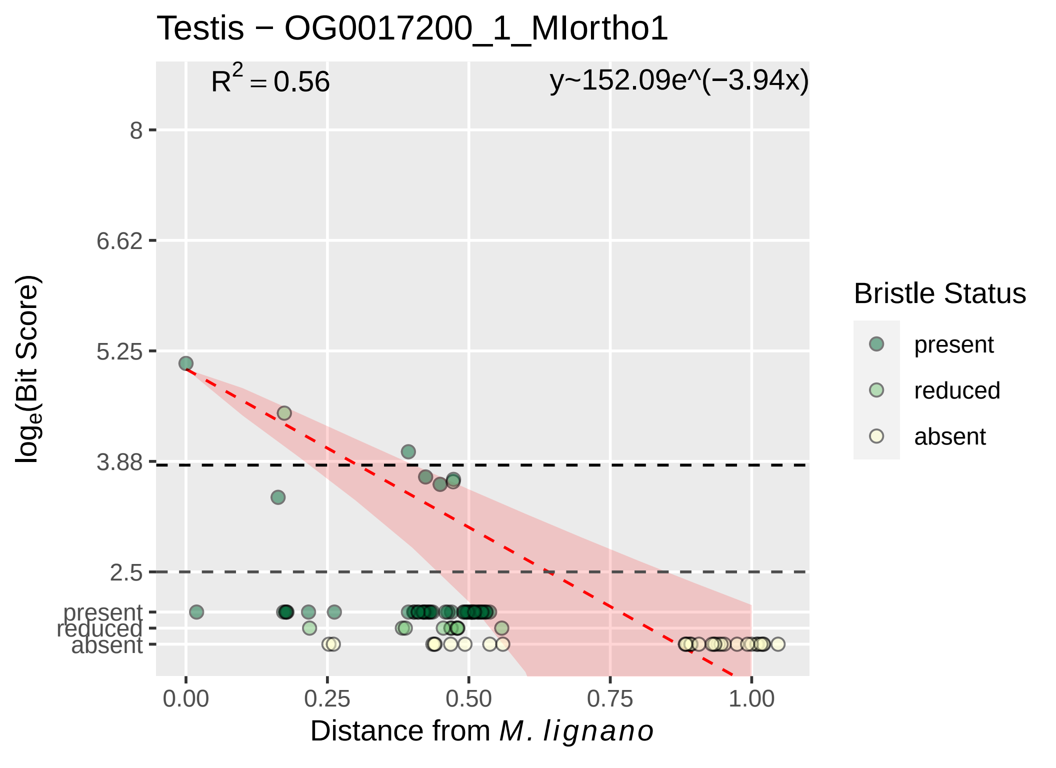

Testis − OG0017200_1_MIortho1
R² = 0.56
y~152.09e^(−3.94x)
8
6.62
5.25
3.88
2.5
present
reduced
absent
0.00
0.25
0.50
0.75
1.00
Distance from M. lignano
loge(Bit Score)
Bristle Status
present
reduced
absent

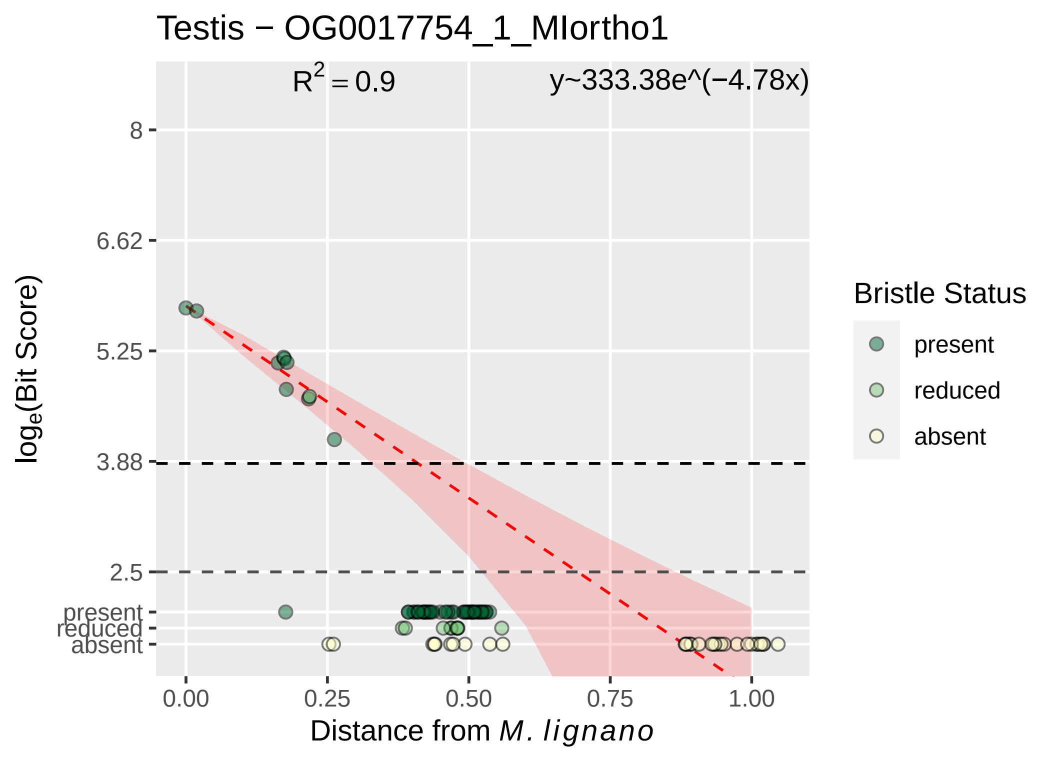

Testis − OG0017754_1_MIortho1
$R^2 = 0.9$
y~333.38e^(−4.78x)
8
6.62
5.25
3.88
2.5
present
reduced
absent
0.00
0.25
0.50
0.75
1.00
Distance from M. lignano
log_e(Bit Score)
Bristle Status
present
reduced
absent

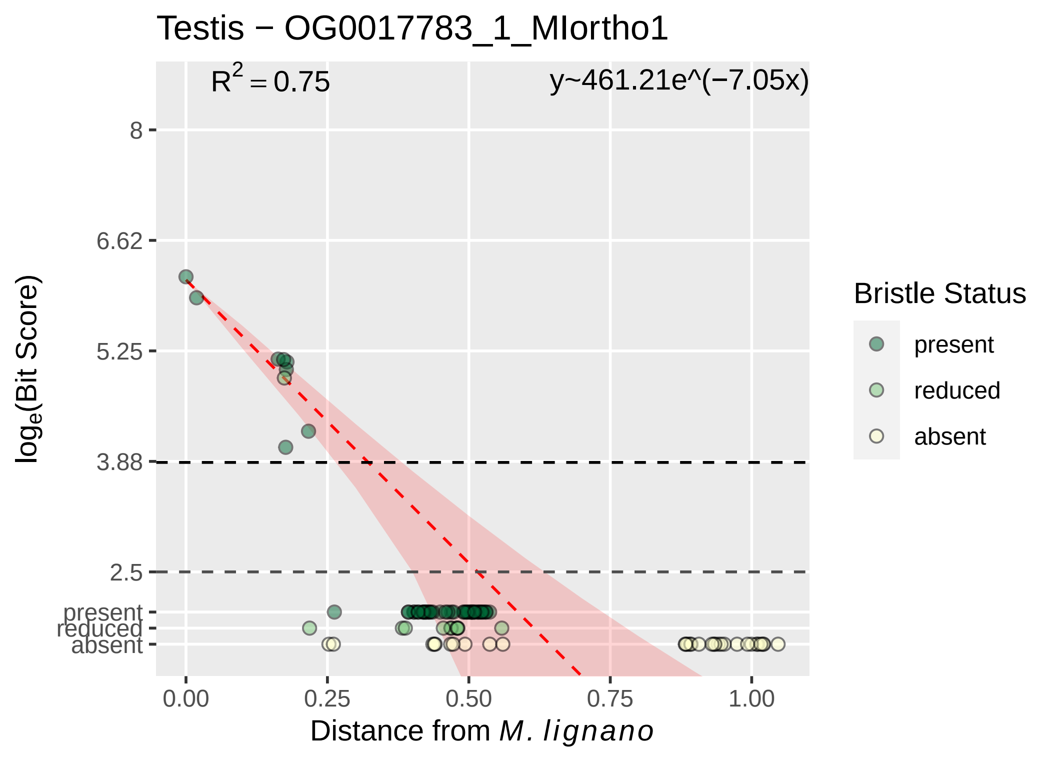

Testis − OG0017783_1_MIortho1
$R^2 = 0.75$
y~461.21e^(−7.05x)
8
6.62
5.25
3.88
2.5
present
reduced
absent
0.00
0.25
0.50
0.75
1.00
Distance from M. lignano
log$_e$(Bit Score)
Bristle Status
present
reduced
absent

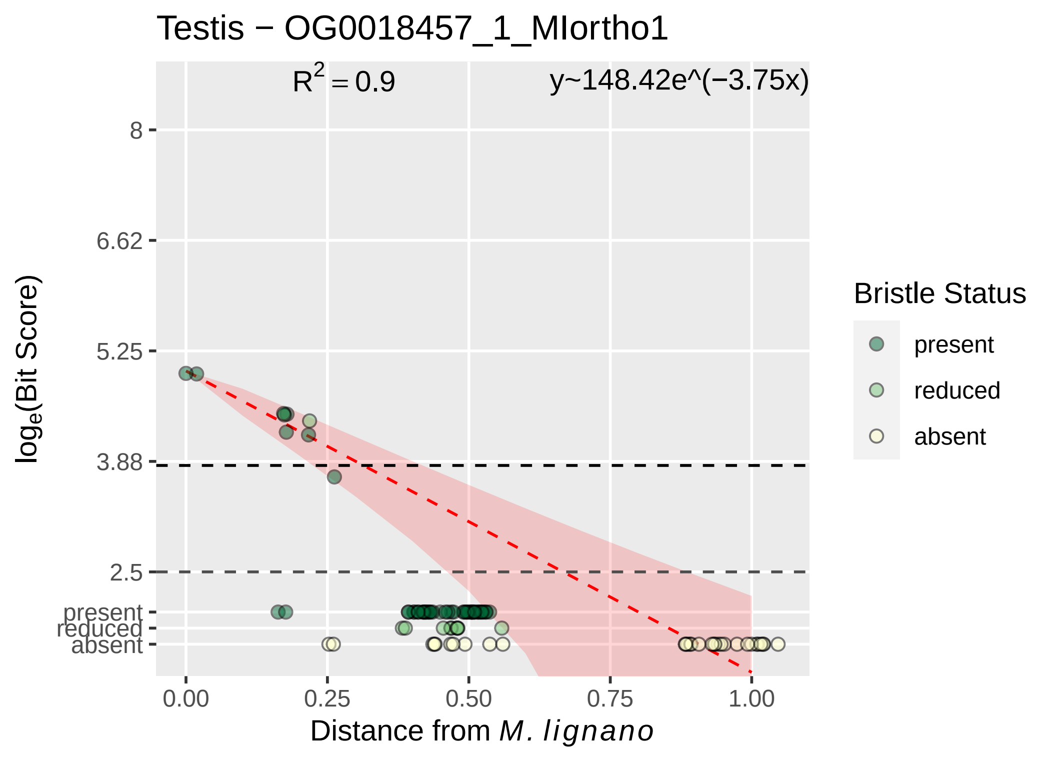

Testis − OG0018457_1_MIortho1
R² = 0.9
y~148.42e^(−3.75x)
8
6.62
5.25
3.88
2.5
present
reduced
absent
0.00
0.25
0.50
0.75
1.00
Distance from M. lignano
loge(Bit Score)
Bristle Status
present
reduced
absent

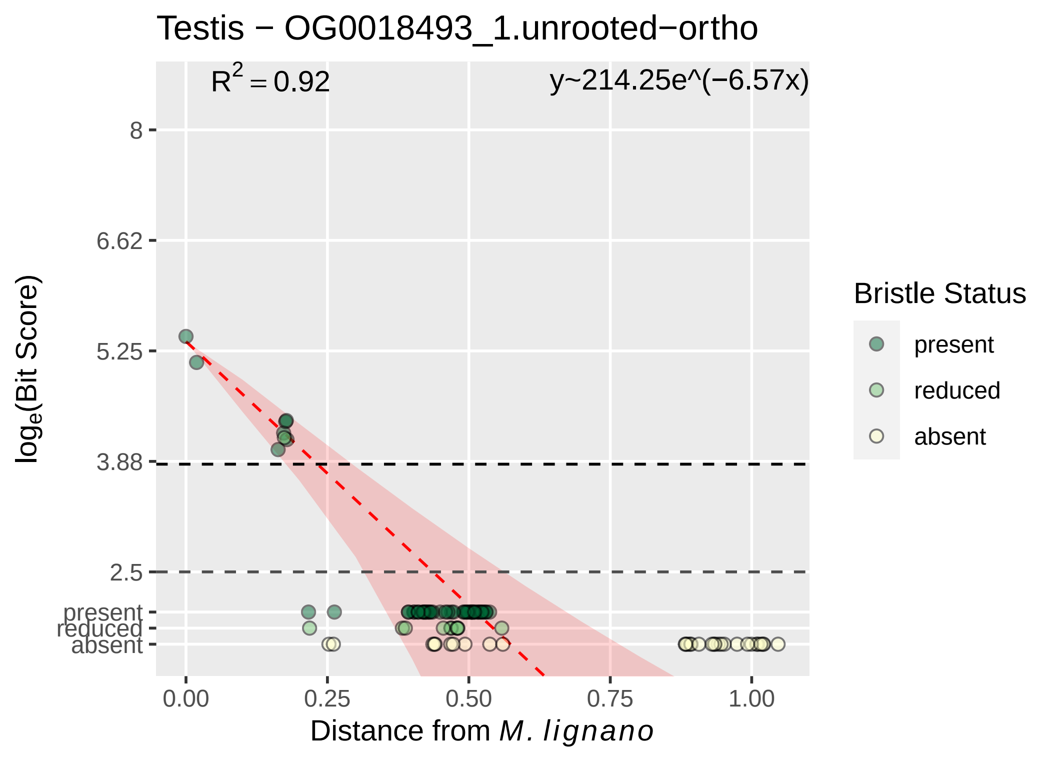

Testis − OG0018493_1.unrooted−ortho
R2 = 0.92
y~214.25e^(−6.57x)
8
6.62
5.25
3.88
2.5
present
reduced
absent
loge(Bit Score)
0.00
0.25
0.50
0.75
1.00
Distance from M. lignano
Bristle Status
present
reduced
absent

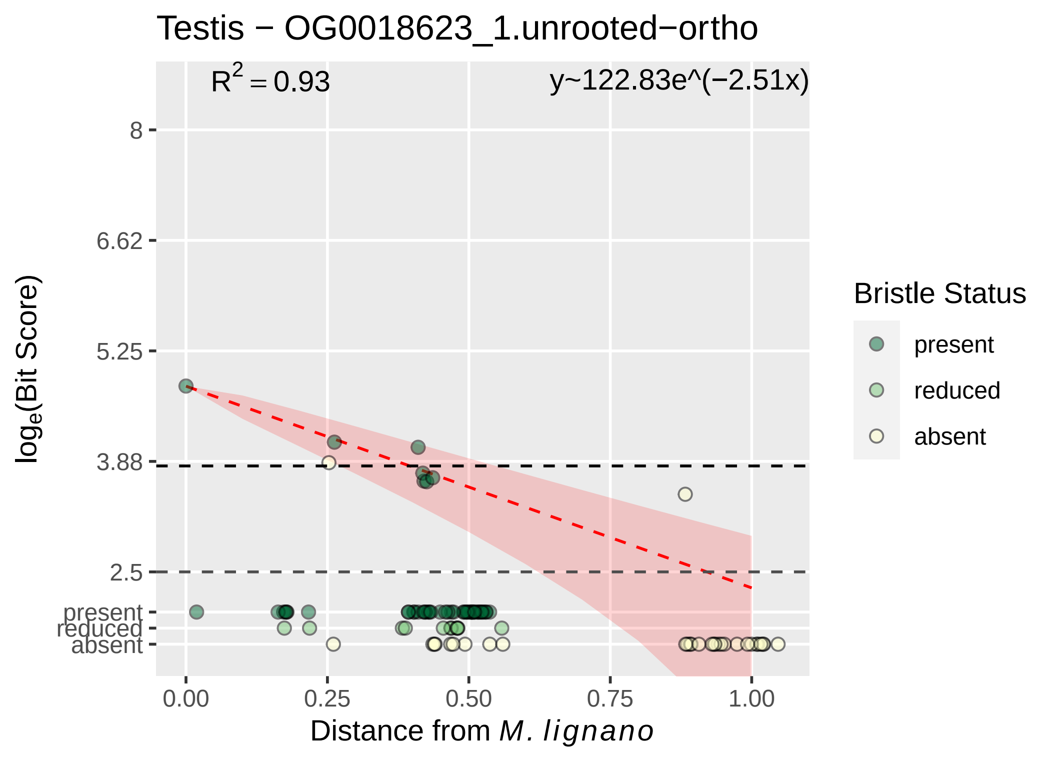

Testis − OG0018623_1.unrooted−ortho
$R^2 = 0.93$
y~122.83e^(−2.51x)
8
6.62
5.25
3.88
2.5
present
reduced
absent
0.00
0.25
0.50
0.75
1.00
$log_e$(Bit Score)
Distance from *M. lignano*
Bristle Status
present
reduced
absent

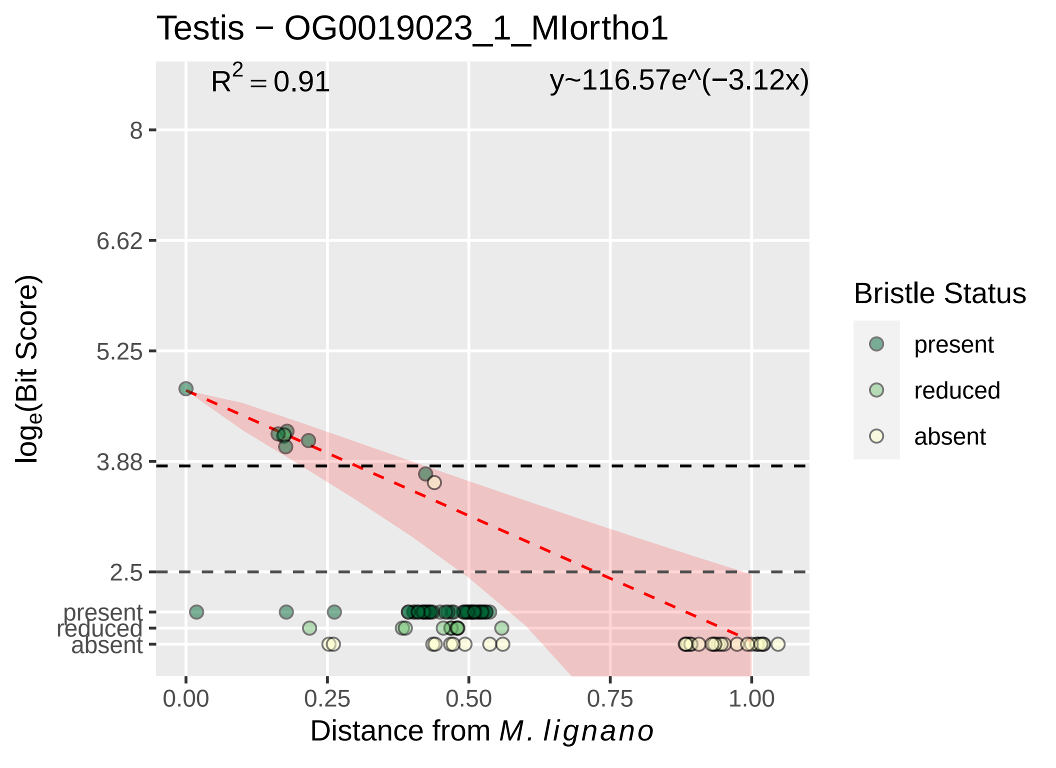

Testis − OG0019023_1_MIortho1
R^2 = 0.91
y~116.57e^(−3.12x)
8
6.62
5.25
3.88
2.5
present
reduced
absent
log_e(Bit Score)
0.00
0.25
0.50
0.75
1.00
Distance from M. lignano
Bristle Status
present
reduced
absent

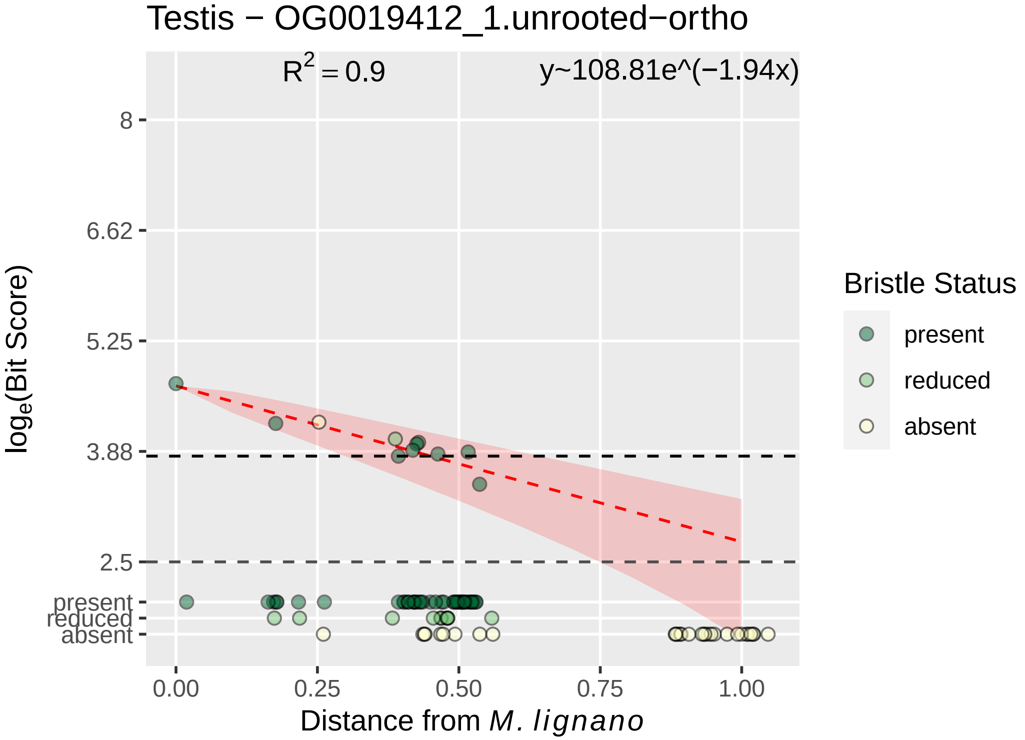

Testis − OG0019412_1.unrooted−ortho
$R^2 = 0.9$
y~108.81e^(−1.94x)
8
6.62
5.25
3.88
2.5
present
reduced
absent
$log_e$(Bit Score)
0.00
0.25
0.50
0.75
1.00
Distance from *M. lignano*
Bristle Status
present
reduced
absent
