## Supplementary material for "Mating strategy predicts gene presence/absence patterns in a genus of simultaneously hermaphroditic flatworms": figure S5B

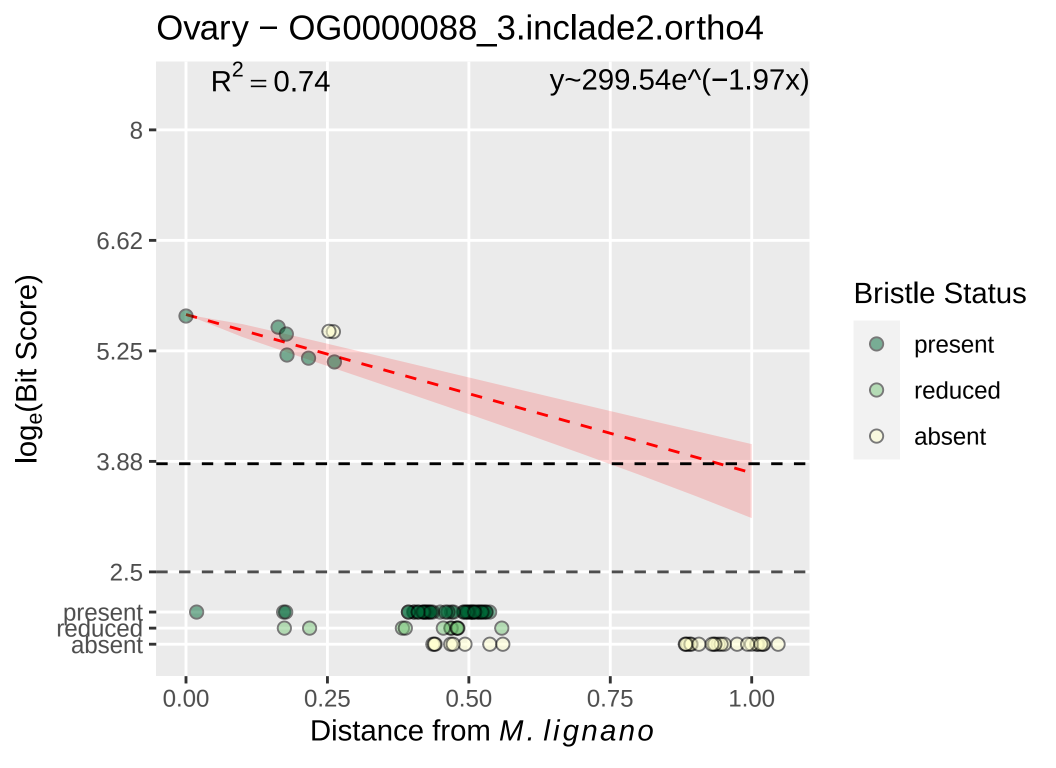

Ovary − OG0000088_3.inclade2.ortho4
R^2 = 0.74
y~299.54e^(−1.97x)
8
6.62
5.25
3.88
2.5
present
reduced
absent
loge(Bit Score)
0.00
0.25
0.50
0.75
1.00
Distance from M. lignano
Bristle Status
present
reduced
absent

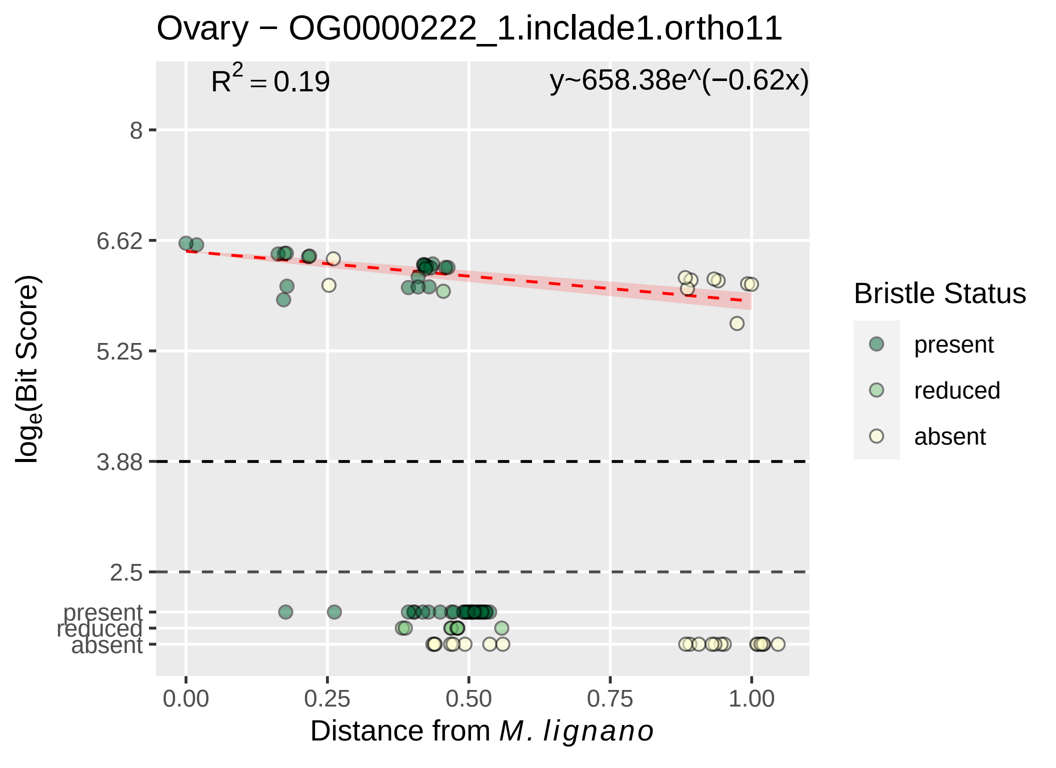

Ovary − OG0000222_1.inclade1.ortho11
R^2 = 0.19
y~658.38e^(−0.62x)
8
6.62
5.25
3.88
2.5
present
reduced
absent
loge(Bit Score)
0.00
0.25
0.50
0.75
1.00
Distance from M. lignano
Bristle Status
present
reduced
absent

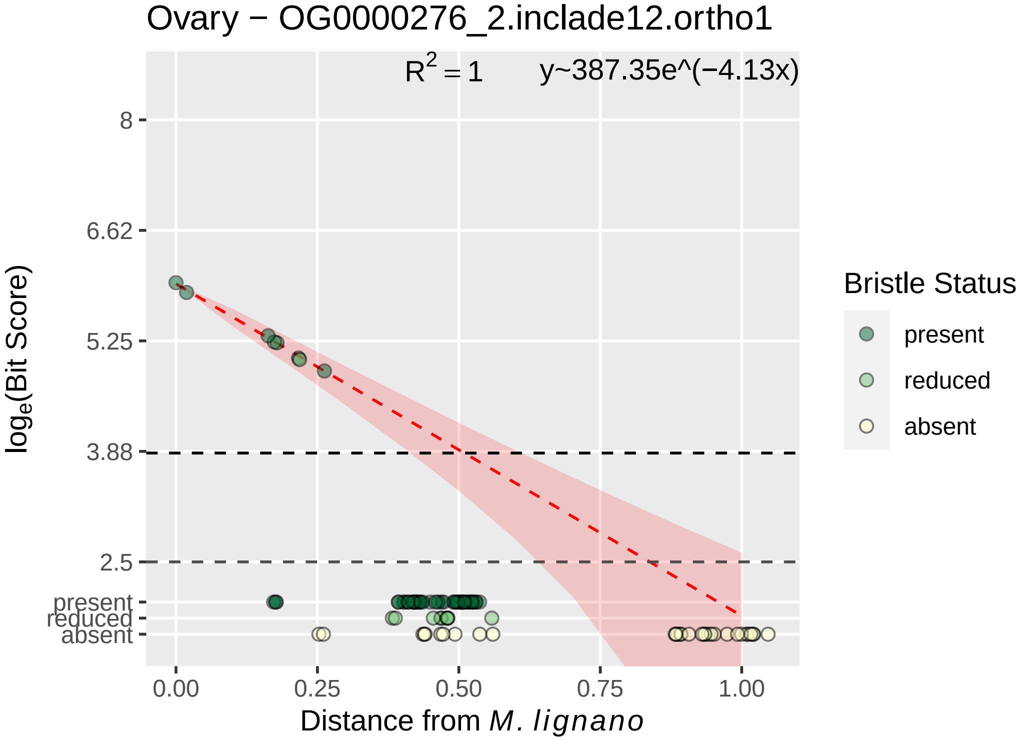

Ovary − OG0000276_2.inclade12.ortho1
$R^2 = 1$
y~387.35e^(−4.13x)
8
6.62
5.25
3.88
2.5
present
reduced
absent
0.00
0.25
0.50
0.75
1.00
$\log_e$(Bit Score)
Distance from *M. lignano*
Bristle Status
present
reduced
absent

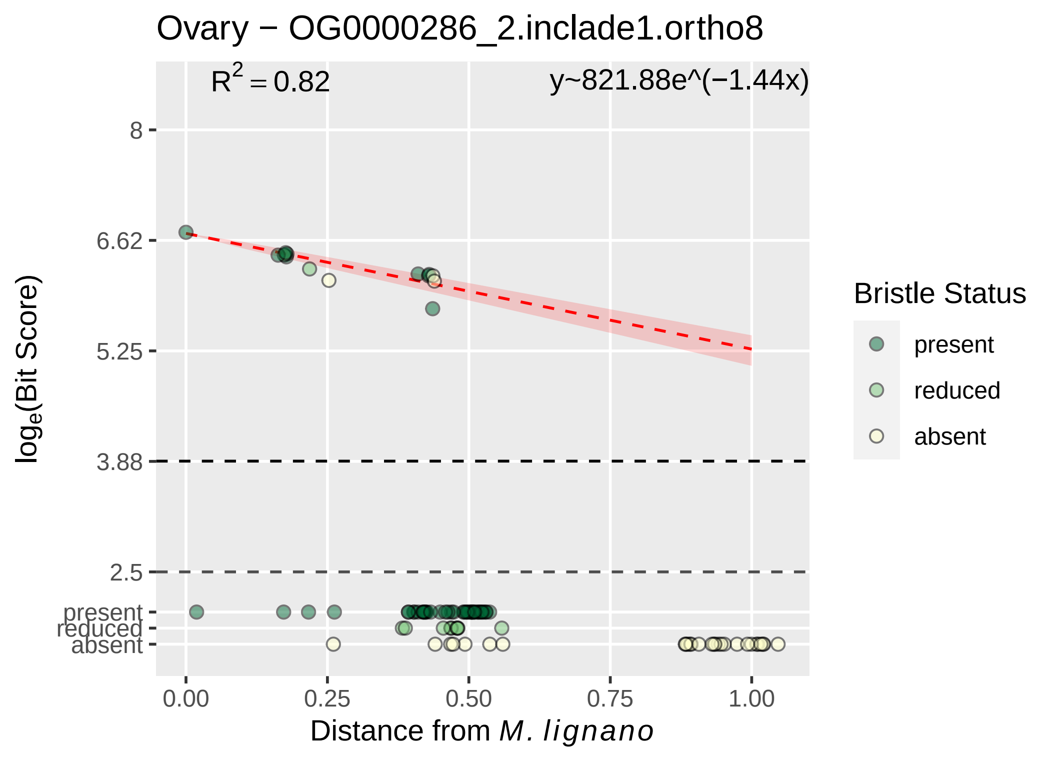

Ovary − OG0000286_2.inclade1.ortho8
R² = 0.82
y~821.88e^(−1.44x)
8
6.62
5.25
3.88
2.5
present
reduced
absent
log_e(Bit Score)
0.00
0.25
0.50
0.75
1.00
Distance from M. lignano
Bristle Status
present
reduced
absent

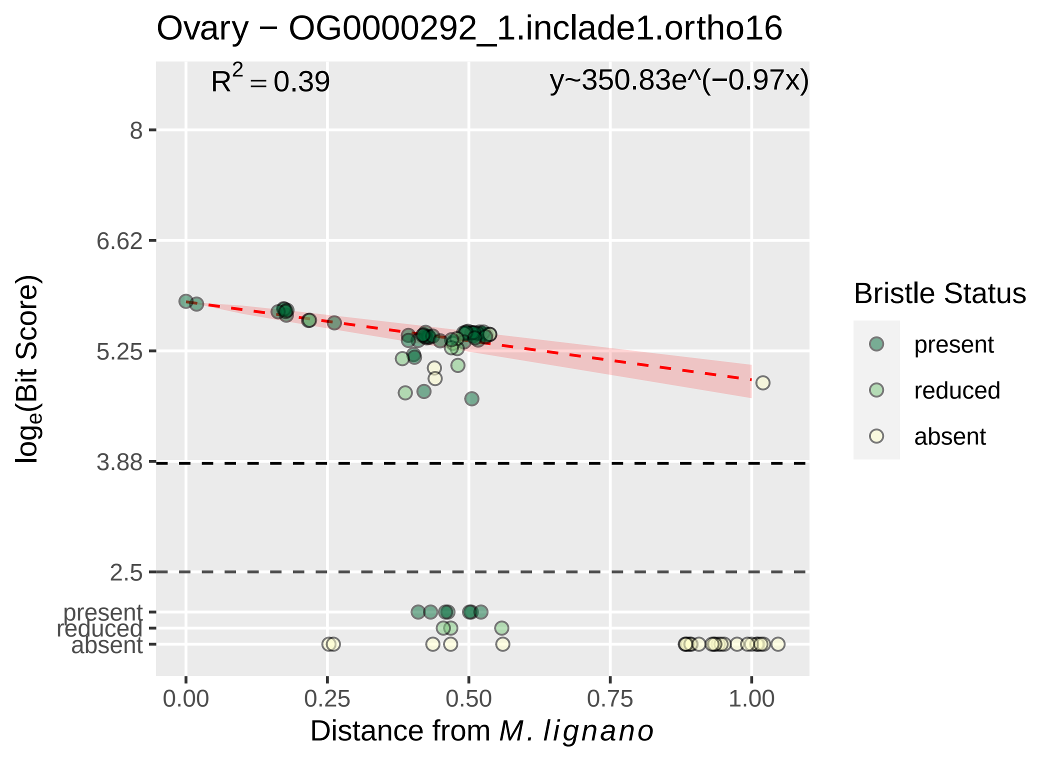

Ovary − OG0000292_1.inclade1.ortho16
R² = 0.39
y~350.83e^(−0.97x)
8
6.62
5.25
3.88
2.5
present
reduced
absent
log_e(Bit Score)
0.00
0.25
0.50
0.75
1.00
Distance from M. lignano
Bristle Status
present
reduced
absent

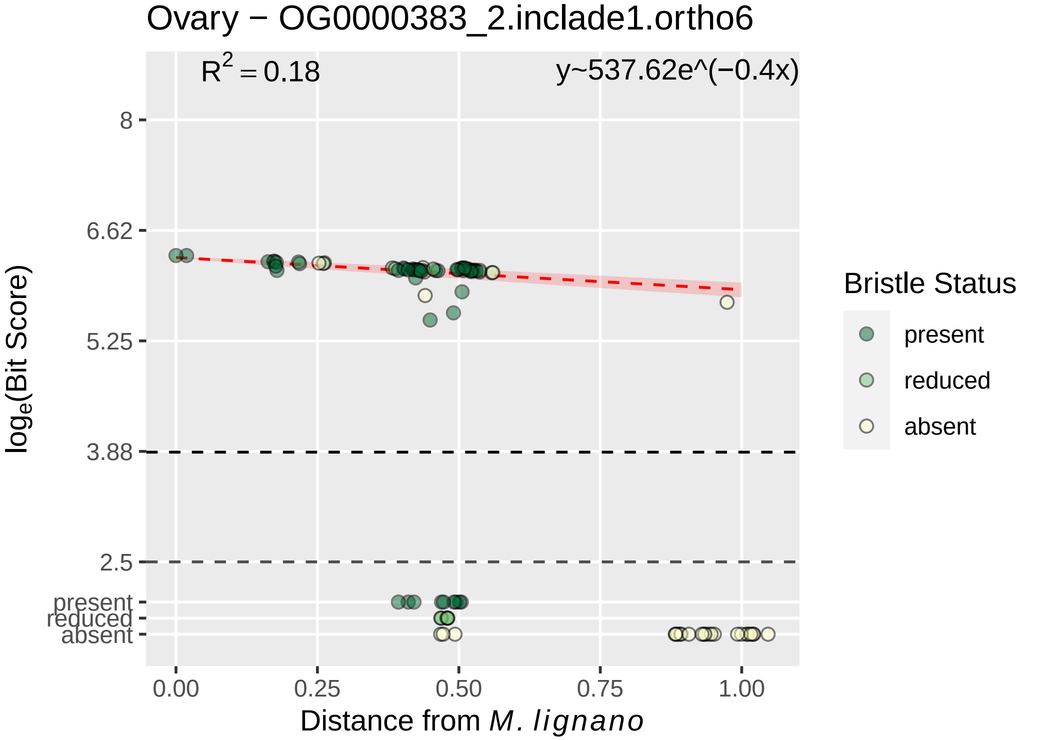
Ovary − OG0000383_2.inclade1.ortho6
$R^2 = 0.18$
y~537.62e^(−0.4x)
8
6.62
5.25
3.88
2.5
present
reduced
absent
log_e(Bit Score)
0.00
0.25
0.50
0.75
1.00
Distance from M. lignano
Bristle Status
present
reduced
absent

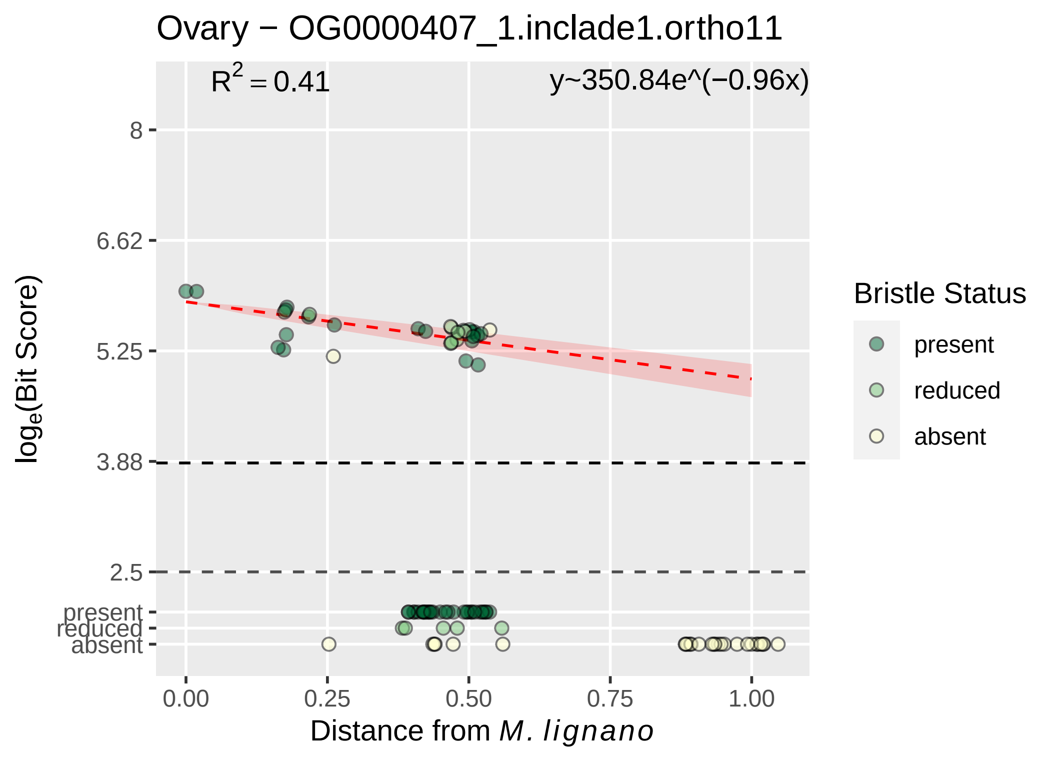

Ovary − OG0000407_1.inclade1.ortho11
$R^2 = 0.41$
y~350.84e^(−0.96x)
8
6.62
5.25
3.88
2.5
present
reduced
absent
log_e(Bit Score)
0.00
0.25
0.50
0.75
1.00
Distance from M. lignano
Bristle Status
present
reduced
absent

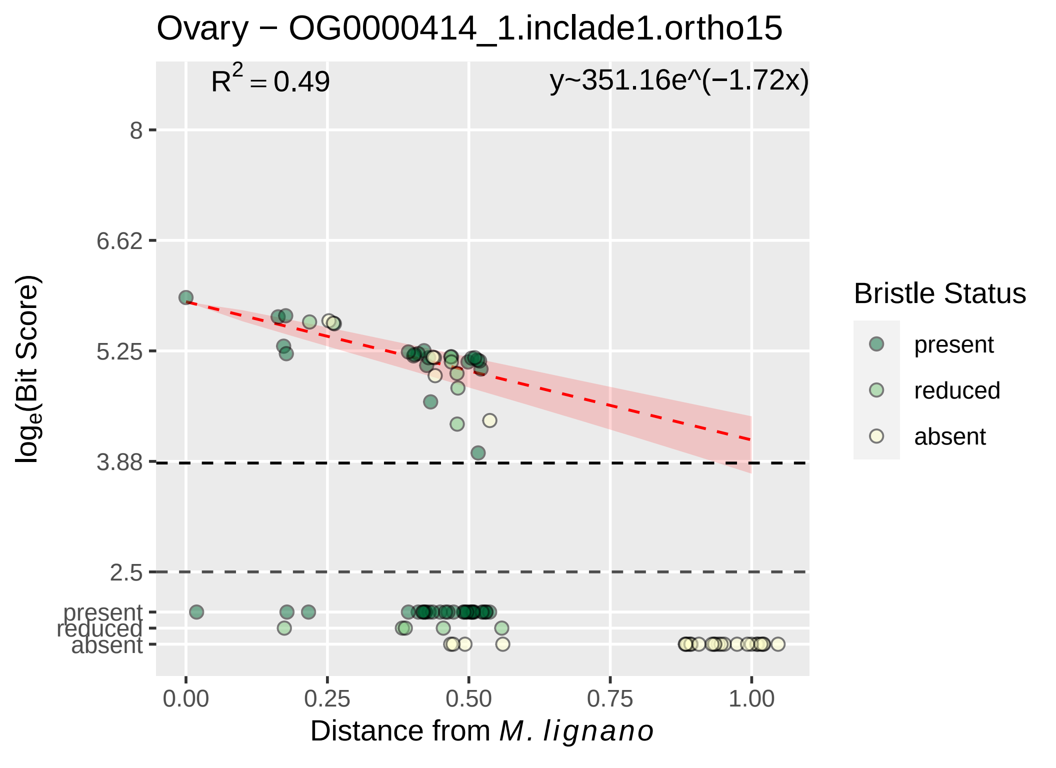

Ovary − OG0000414_1.inclade1.ortho15
R^2 = 0.49
y~351.16e^(−1.72x)
8
6.62
5.25
3.88
2.5
present
reduced
absent
log_e(Bit Score)
0.00
0.25
0.50
0.75
1.00
Distance from M. lignano
Bristle Status
present
reduced
absent

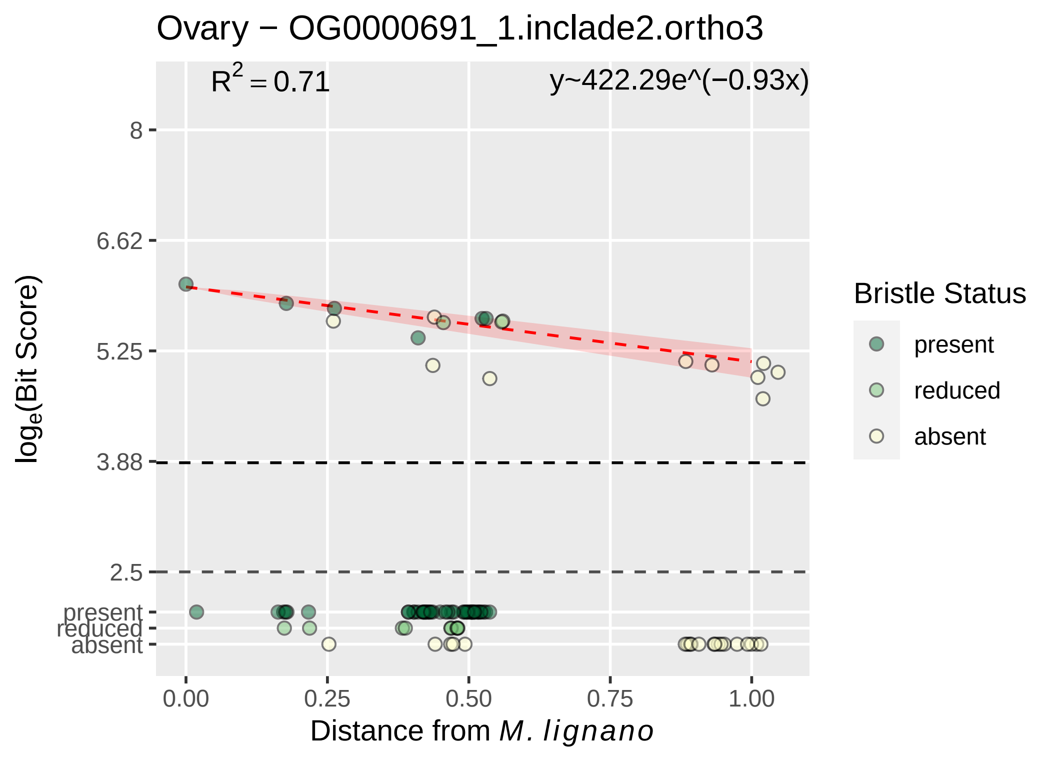

Ovary − OG0000691_1.inclade2.ortho3
R^2 = 0.71
y~422.29e^(−0.93x)
8
6.62
5.25
3.88
2.5
present
reduced
absent
log_e(Bit Score)
0.00
0.25
0.50
0.75
1.00
Distance from M. lignano
Bristle Status
present
reduced
absent

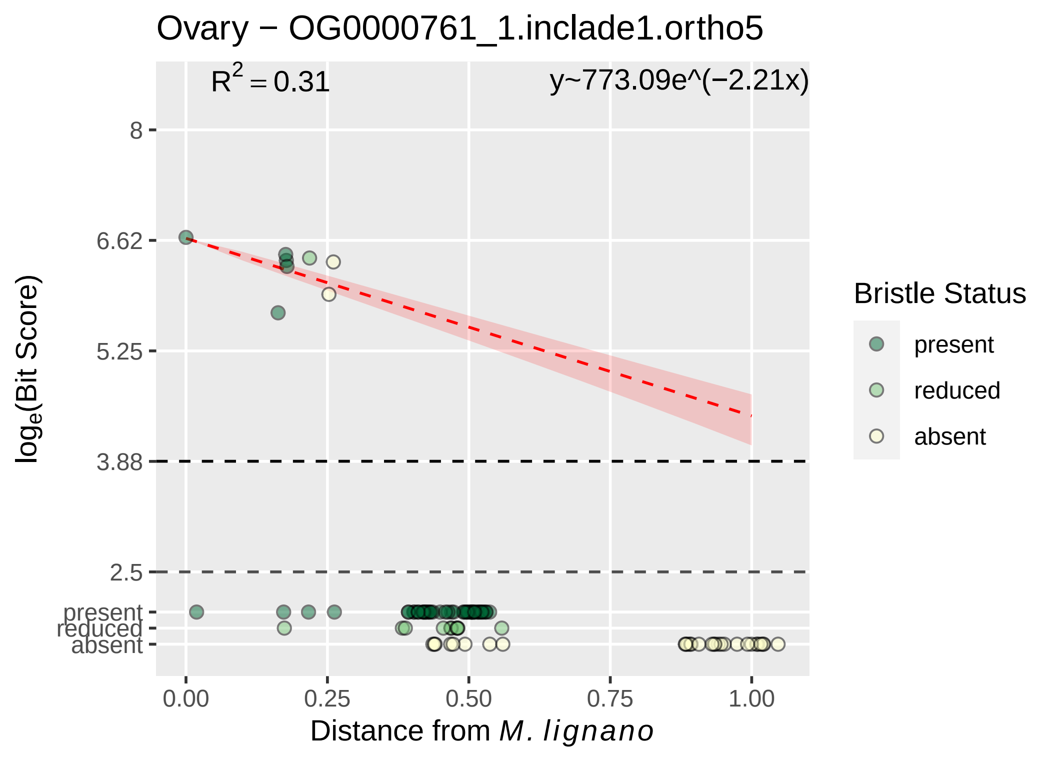

Ovary − OG0000761_1.inclade1.ortho5
R2 = 0.31
y~773.09e^(−2.21x)
8
6.62
5.25
3.88
2.5
present
reduced
absent
loge(Bit Score)
0.00
0.25
0.50
0.75
1.00
Distance from M. lignano
Bristle Status
present
reduced
absent

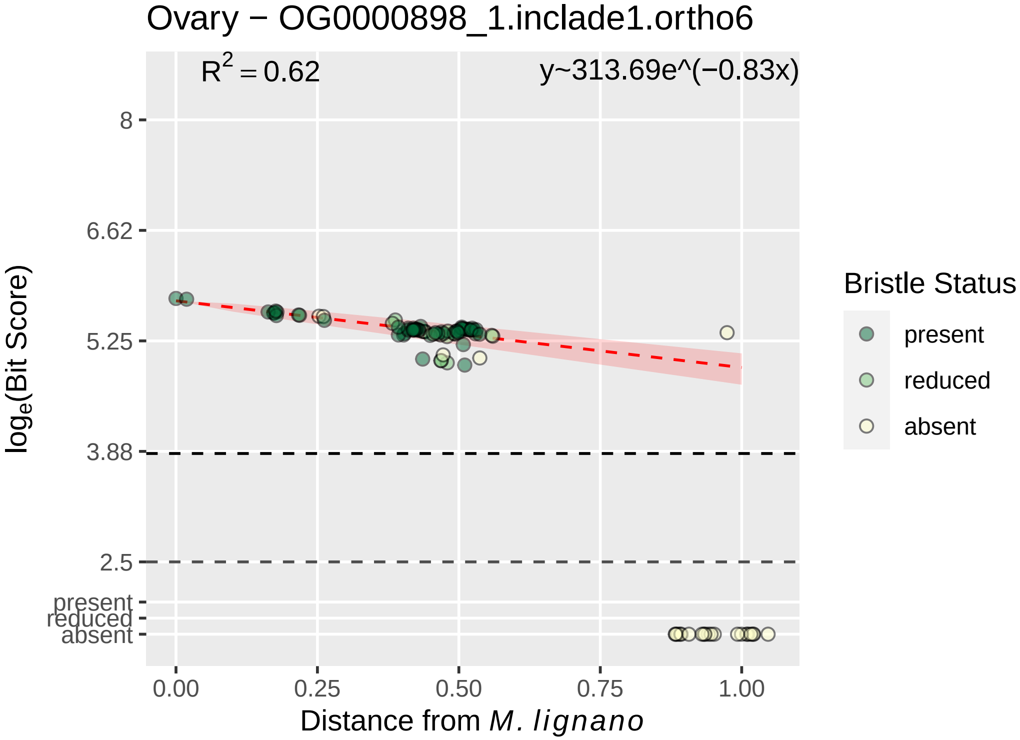

Ovary − OG0000898_1.inclade1.ortho6
R² = 0.62
y~313.69e^(−0.83x)
8
6.62
5.25
3.88
2.5
present
reduced
absent
log_e(Bit Score)
0.00
0.25
0.50
0.75
1.00
Distance from M. lignano
Bristle Status
present
reduced
absent

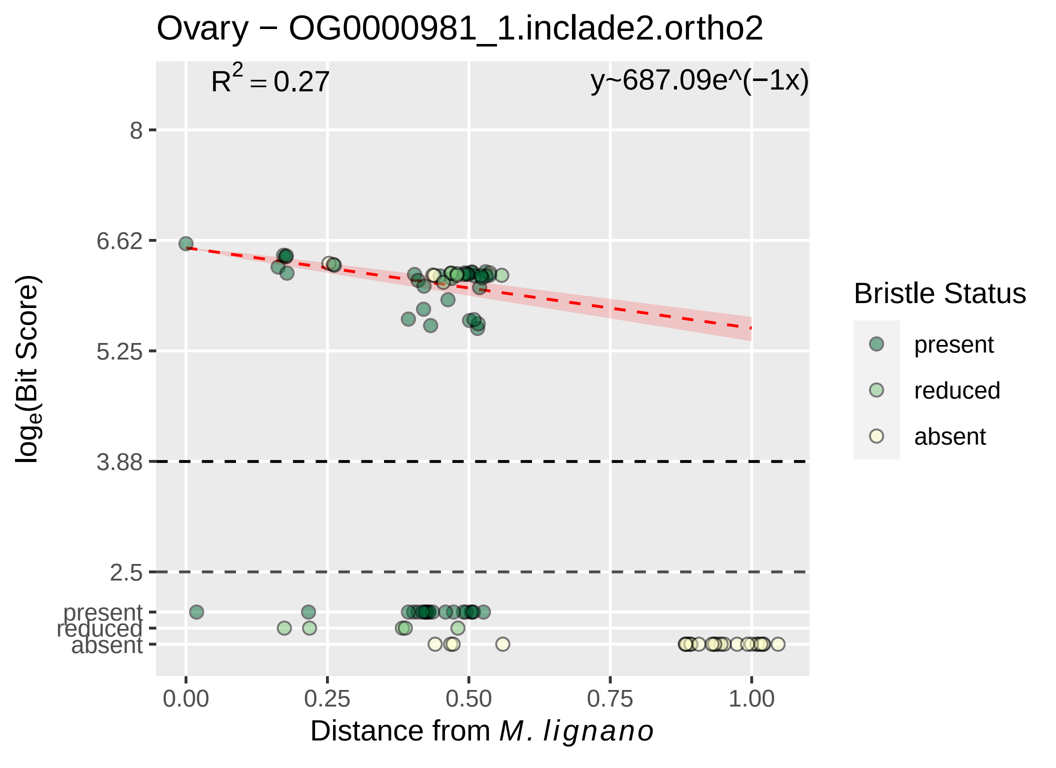
Ovary − OG0000981_1.inclade2.ortho2
$R^2 = 0.27$
y~687.09e^(−1x)
8
6.62
5.25
3.88
2.5
present
reduced
absent
0.00
0.25
0.50
0.75
1.00
Distance from M. lignano
log$_e$(Bit Score)
Bristle Status
present
reduced
absent

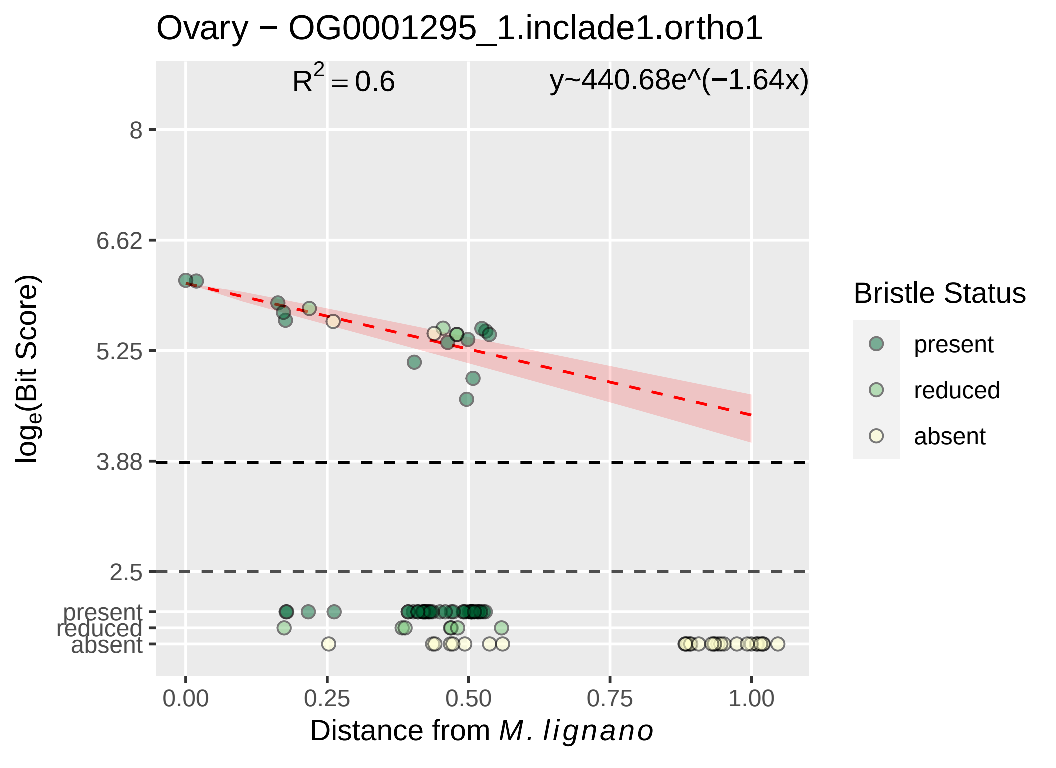

Ovary − OG0001295_1.inclade1.ortho1
R² = 0.6
y~440.68e^(−1.64x)
8
6.62
5.25
3.88
2.5
present
reduced
absent
0.00
0.25
0.50
0.75
1.00
Distance from M. lignano
loge(Bit Score)
Bristle Status
present
reduced
absent

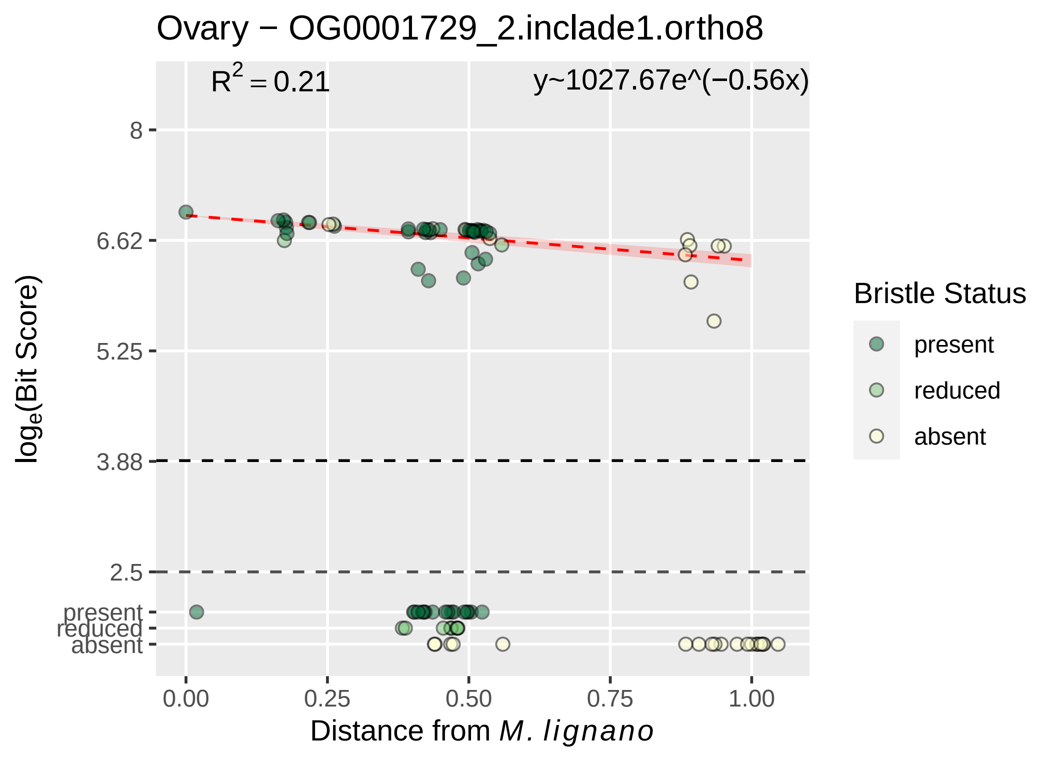

Ovary − OG0001729_2.inclade1.ortho8
R^2 = 0.21
y~1027.67e^(−0.56x)
8
6.62
5.25
3.88
2.5
present
reduced
absent
log_e(Bit Score)
0.00
0.25
0.50
0.75
1.00
Distance from M. lignano
Bristle Status
present
reduced
absent

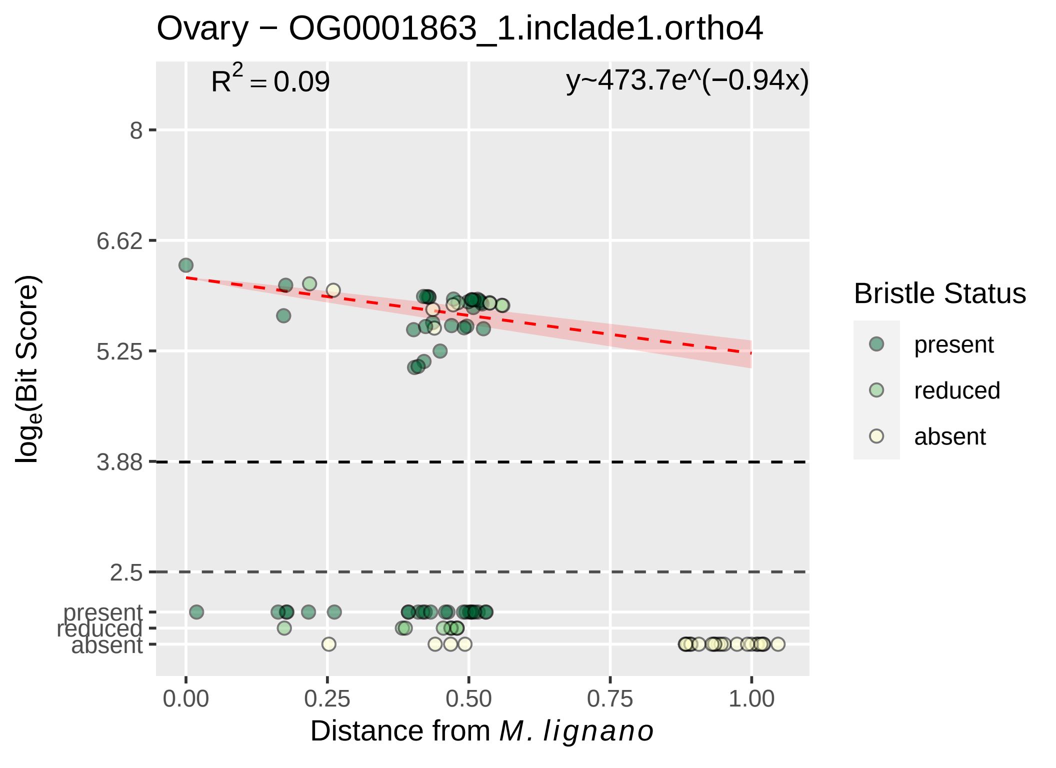

Ovary − OG0001863_1.inclade1.ortho4
$R^2 = 0.09$
y~473.7e^(−0.94x)
8
6.62
5.25
3.88
2.5
present
reduced
absent
log$_e$(Bit Score)
0.00
0.25
0.50
0.75
1.00
Distance from *M. lignano*
Bristle Status
present
reduced
absent

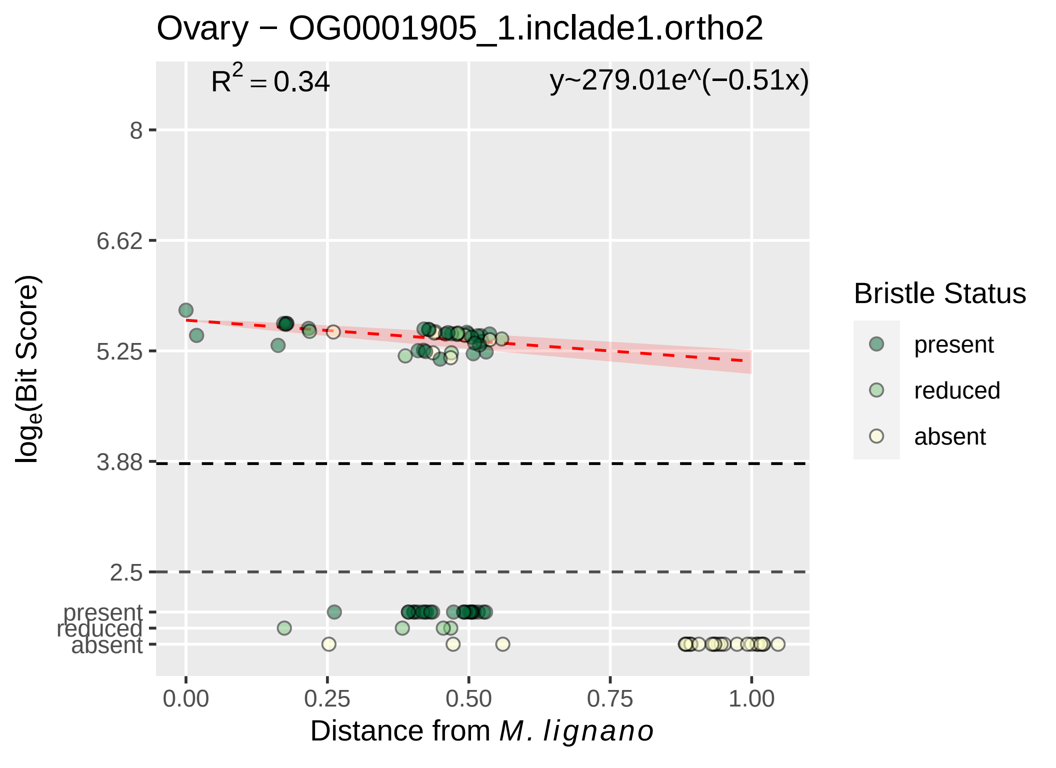

Ovary − OG0001905_1.inclade1.ortho2
R^2 = 0.34
y~279.01e^(−0.51x)
8
6.62
5.25
3.88
2.5
present
reduced
absent
log_e(Bit Score)
0.00
0.25
0.50
0.75
1.00
Distance from M. lignano
Bristle Status
present
reduced
absent

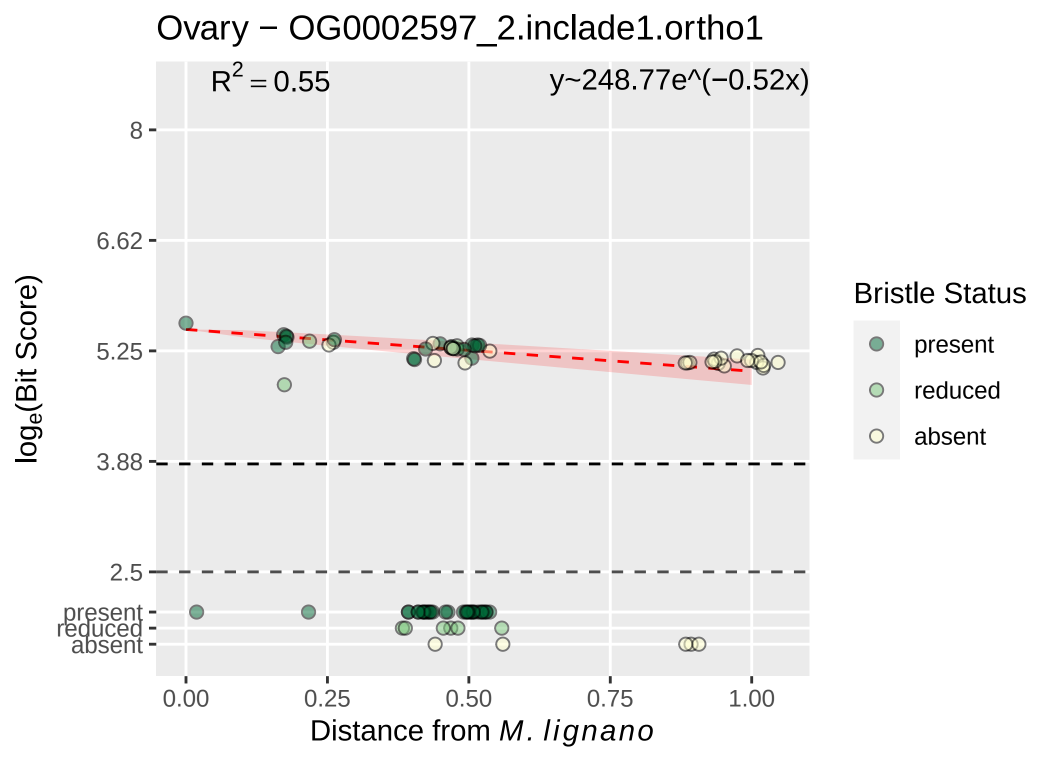

Ovary − OG0002597_2.inclade1.ortho1
R^2 = 0.55
y~248.77e^(−0.52x)
8
6.62
5.25
3.88
2.5
present
reduced
absent
log_e(Bit Score)
0.00
0.25
0.50
0.75
1.00
Distance from M. lignano
Bristle Status
present
reduced
absent

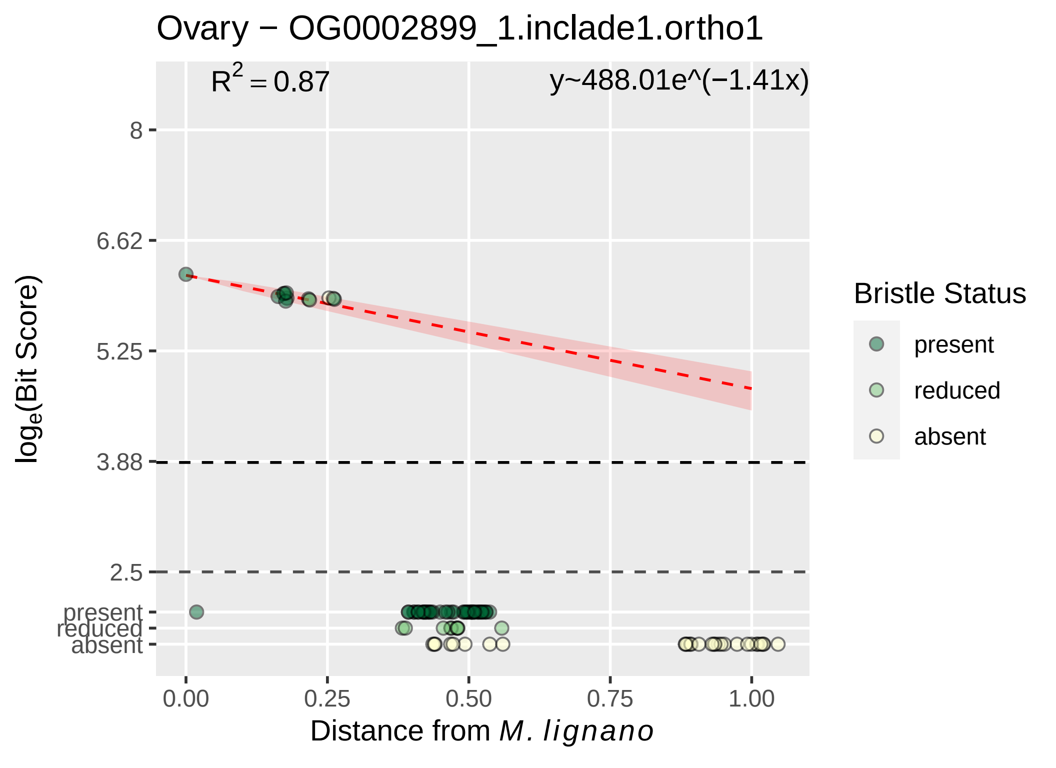

Ovary − OG0002899_1.inclade1.ortho1
R^2 = 0.87
y~488.01e^(−1.41x)
8
6.62
5.25
3.88
2.5
present
reduced
absent
log_e(Bit Score)
0.00
0.25
0.50
0.75
1.00
Distance from M. lignano
Bristle Status
present
reduced
absent

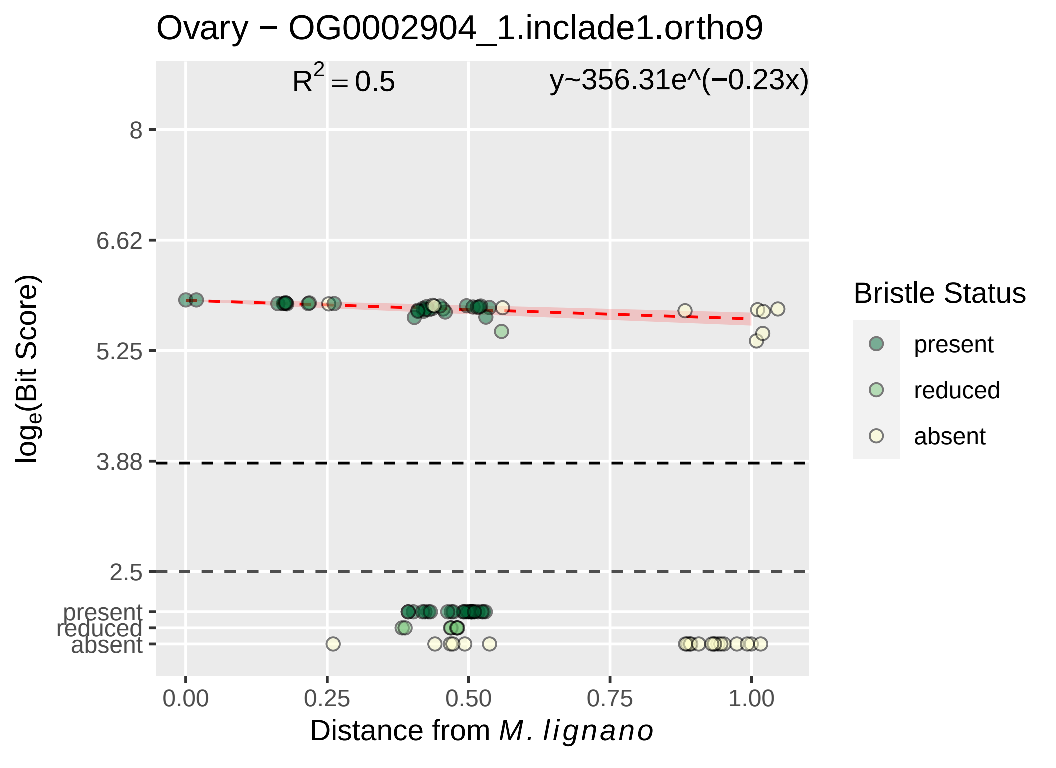

Ovary − OG0002904_1.inclade1.ortho9
$R^2 = 0.5$
y~356.31e^(−0.23x)
8
6.62
5.25
3.88
2.5
present
reduced
absent
0.00
0.25
0.50
0.75
1.00
log$_e$(Bit Score)
Distance from *M. lignano*
Bristle Status
present
reduced
absent

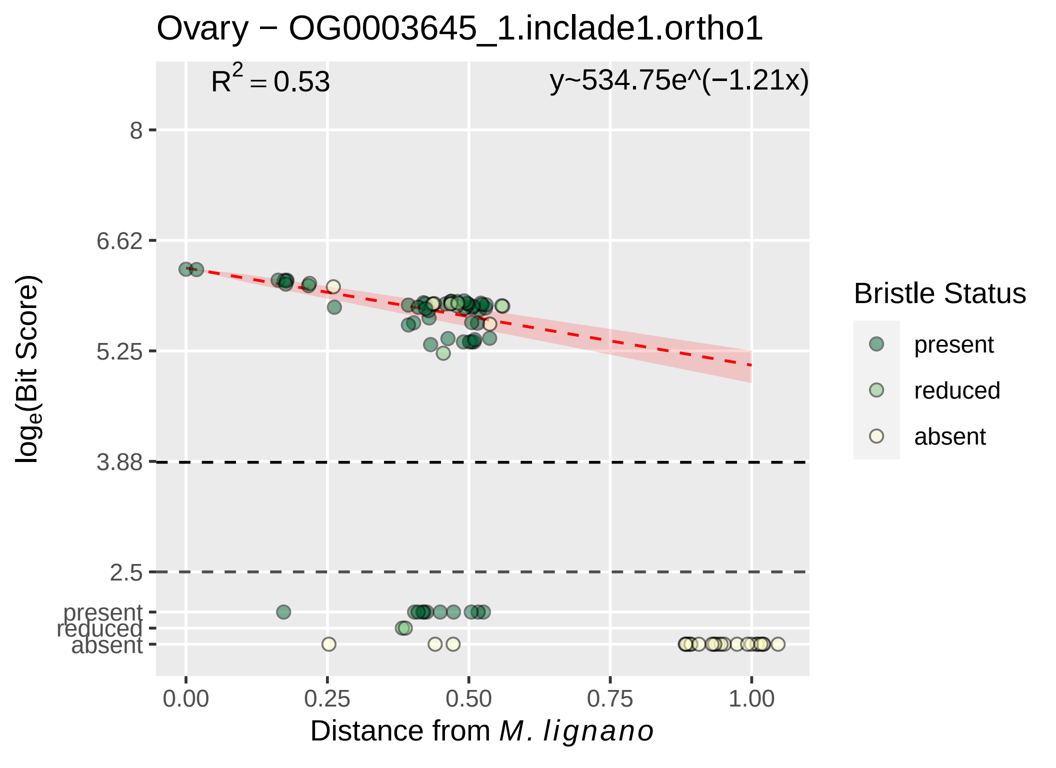

Ovary − OG0003645_1.inclade1.ortho1
R^2 = 0.53
y~534.75e^(−1.21x)
8
6.62
5.25
3.88
2.5
present
reduced
absent
log_e(Bit Score)
0.00
0.25
0.50
0.75
1.00
Distance from M. lignano
Bristle Status
present
reduced
absent

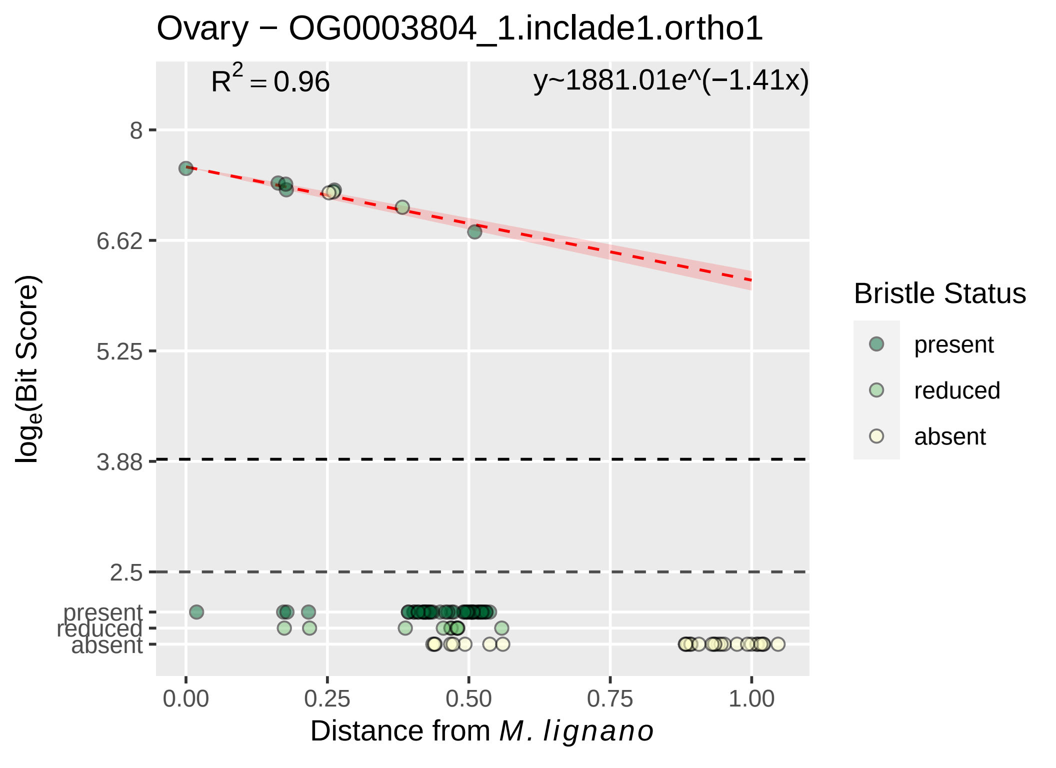
Ovary − OG0003804_1.inclade1.ortho1
R^2 = 0.96
y~1881.01e^(−1.41x)
8
6.62
5.25
3.88
2.5
present
reduced
absent
0.00
0.25
0.50
0.75
1.00
Distance from M. lignano
loge(Bit Score)
Bristle Status
present
reduced
absent

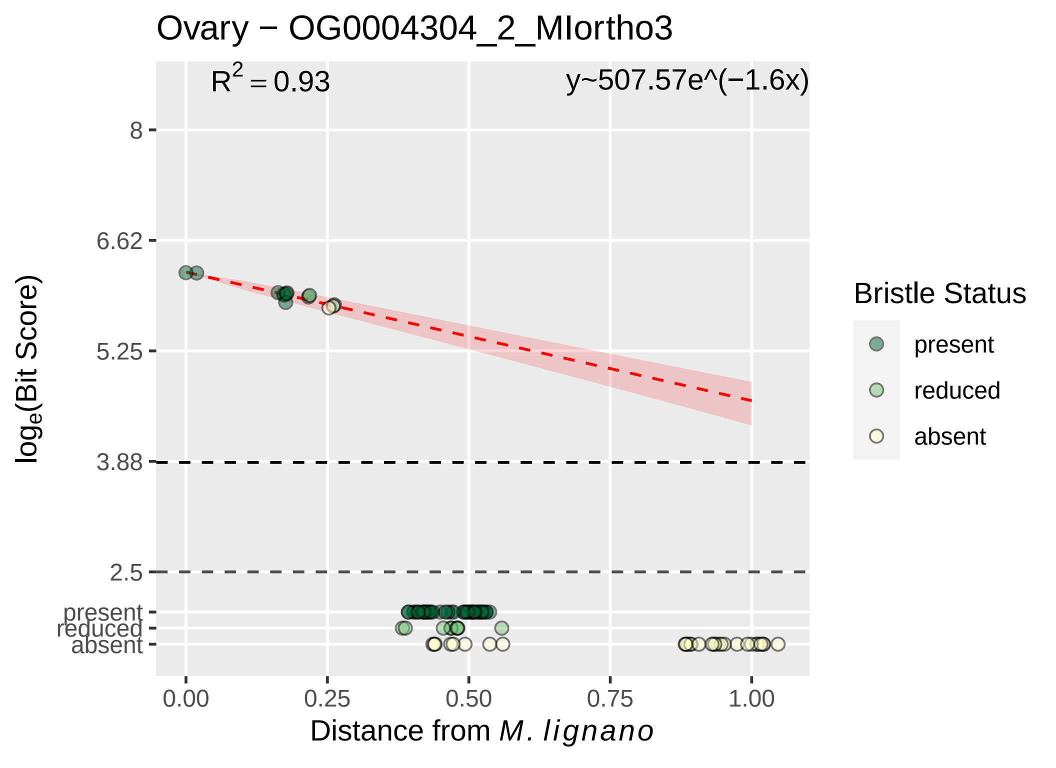

Ovary − OG0004304_2_MIortho3
R^2 = 0.93
y~507.57e^(−1.6x)
8
6.62
5.25
3.88
2.5
present
reduced
absent
log_e(Bit Score)
0.00
0.25
0.50
0.75
1.00
Distance from M. lignano
Bristle Status
present
reduced
absent

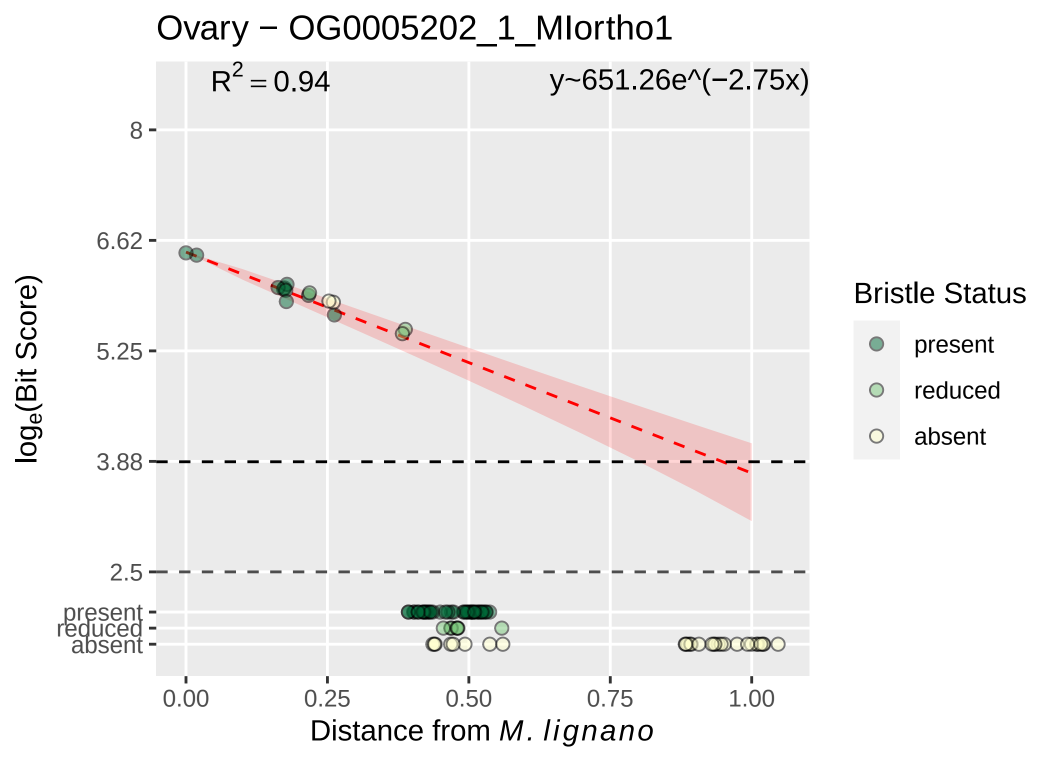

Ovary − OG0005202_1_MIortho1
R^2 = 0.94
y~651.26e^(−2.75x)
8
6.62
5.25
3.88
2.5
present
reduced
absent
log_e(Bit Score)
0.00
0.25
0.50
0.75
1.00
Distance from M. lignano
Bristle Status
present
reduced
absent

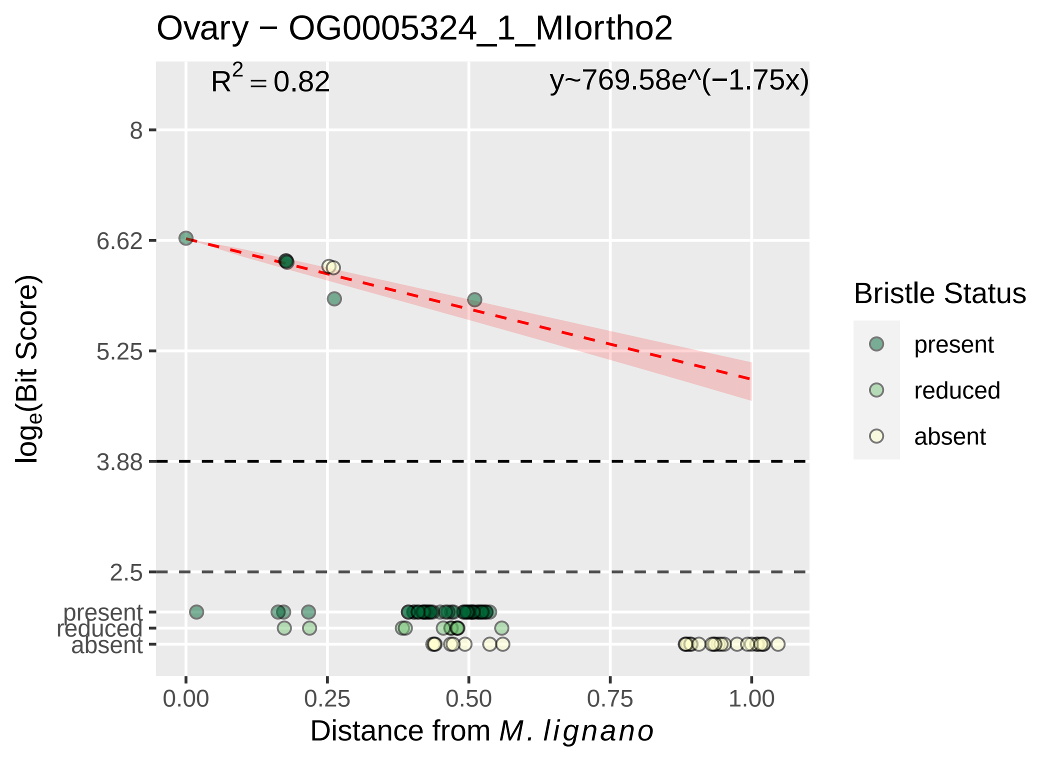

Ovary − OG0005324_1_MIortho2
R2 = 0.82
y~769.58e^(−1.75x)
8
6.62
5.25
3.88
2.5
present
reduced
absent
0.00
0.25
0.50
0.75
1.00
loge(Bit Score)
Distance from M. lignano
Bristle Status
present
reduced
absent

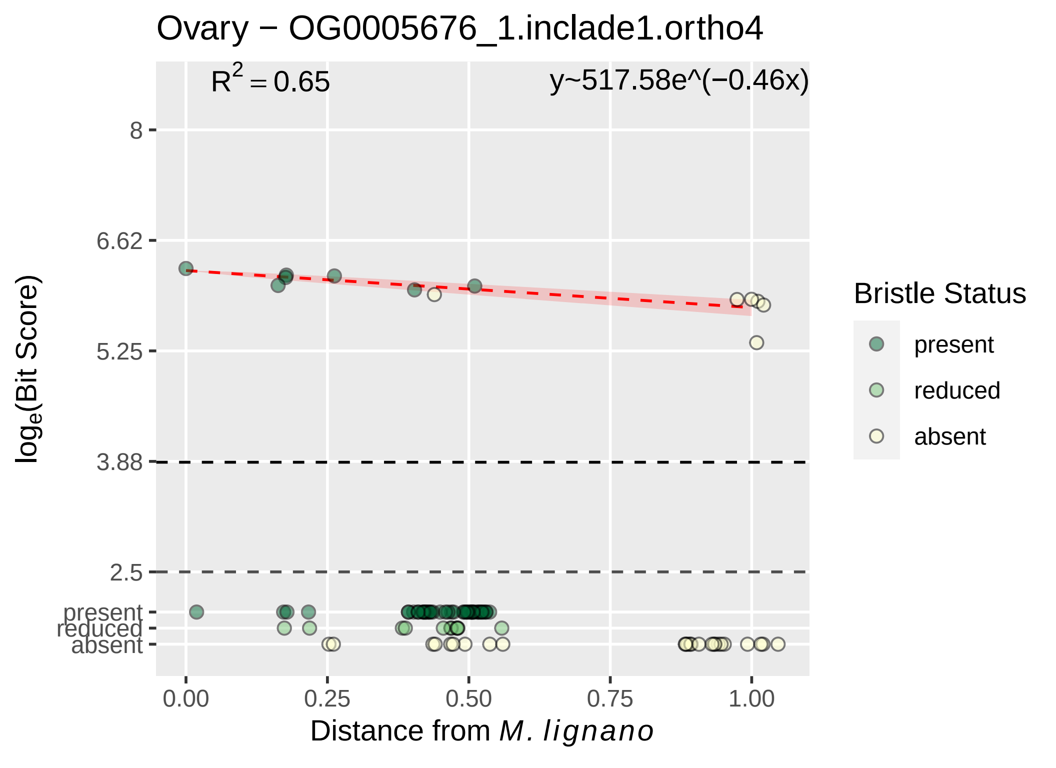

Ovary − OG0005676_1.inclade1.ortho4
$R^2 = 0.65$
y~517.58e^(−0.46x)
8
6.62
5.25
3.88
2.5
present
reduced
absent
0.00
0.25
0.50
0.75
1.00
Distance from M. lignano
log_e(Bit Score)
Bristle Status
present
reduced
absent

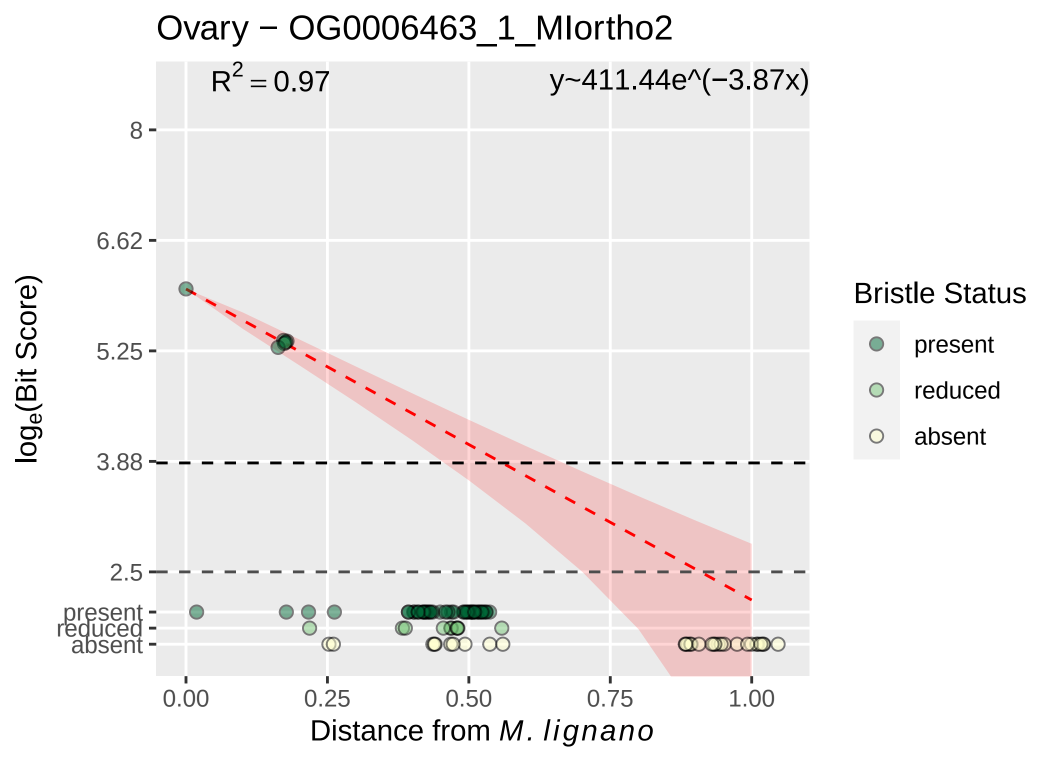

Ovary − OG0006463_1_MIortho2
R2 = 0.97
y~411.44e^(−3.87x)
8
6.62
5.25
3.88
2.5
present
reduced
absent
0.00
0.25
0.50
0.75
1.00
loge(Bit Score)
Distance from M. lignano
Bristle Status
present
reduced
absent

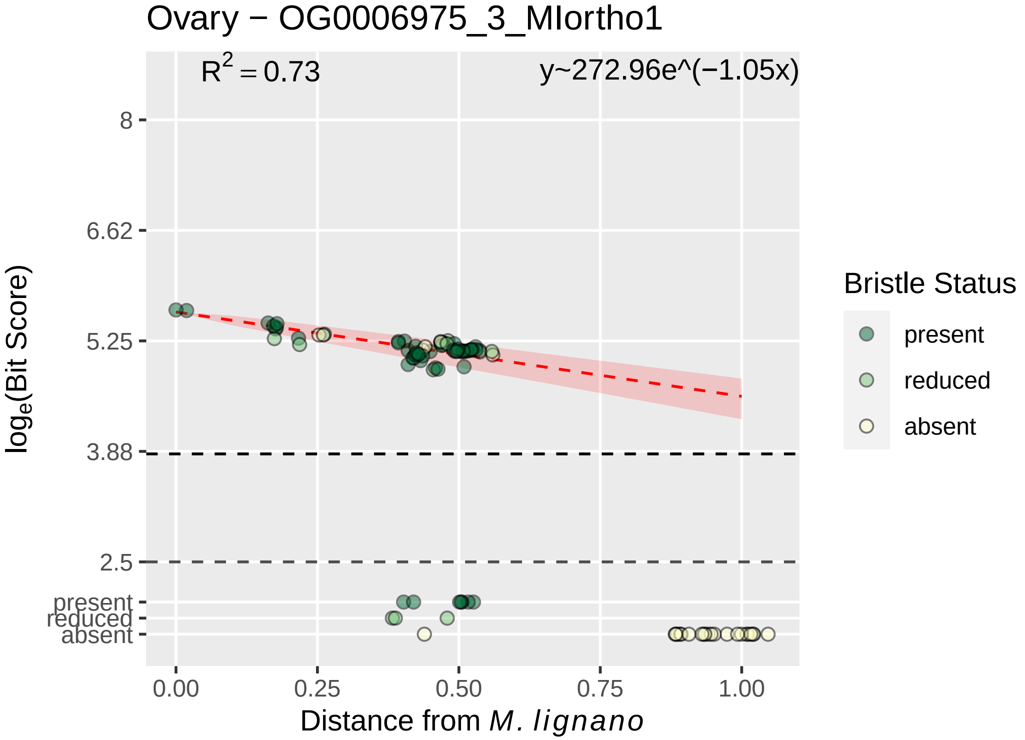
Ovary − OG0006975_3_MIortho1
$R^2 = 0.73$
y~272.96e^(−1.05x)
8
6.62
5.25
3.88
2.5
present
reduced
absent
$\log_e$(Bit Score)
0.00
0.25
0.50
0.75
1.00
Distance from *M. lignano*
Bristle Status
present
reduced
absent

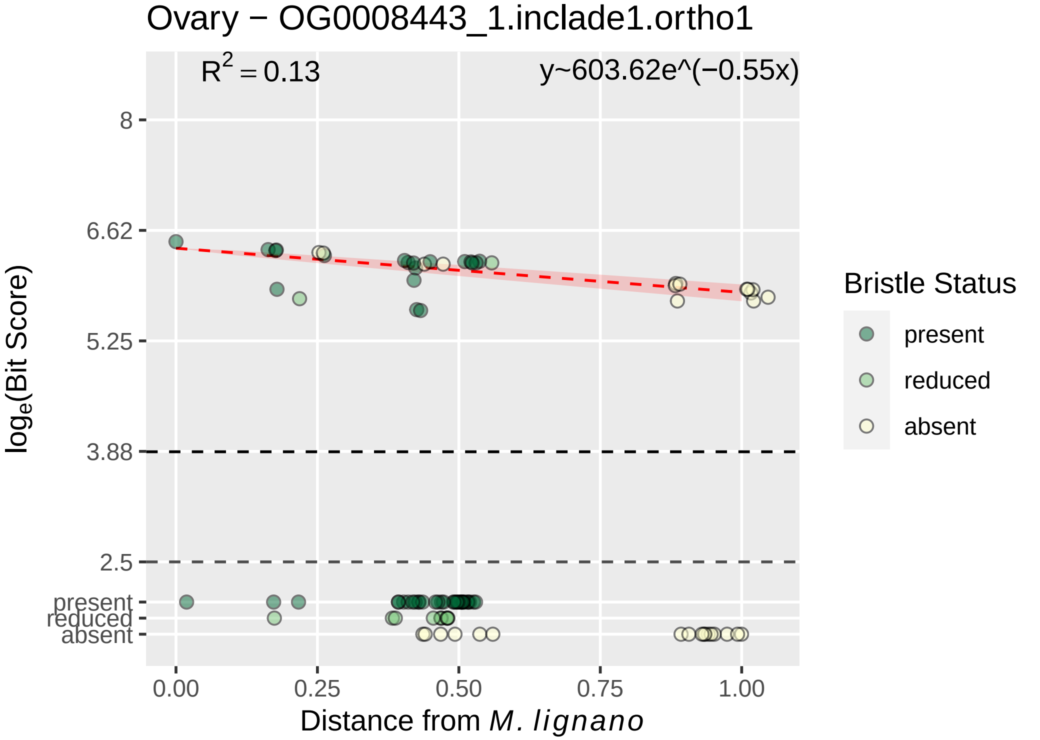
Ovary − OG0008443_1.inclade1.ortho1
$R^2 = 0.13$
y~603.62e^(−0.55x)
8
6.62
5.25
3.88
2.5
present
reduced
absent
$\log_e$(Bit Score)
0.00
0.25
0.50
0.75
1.00
Distance from *M. lignano*
Bristle Status
present
reduced
absent

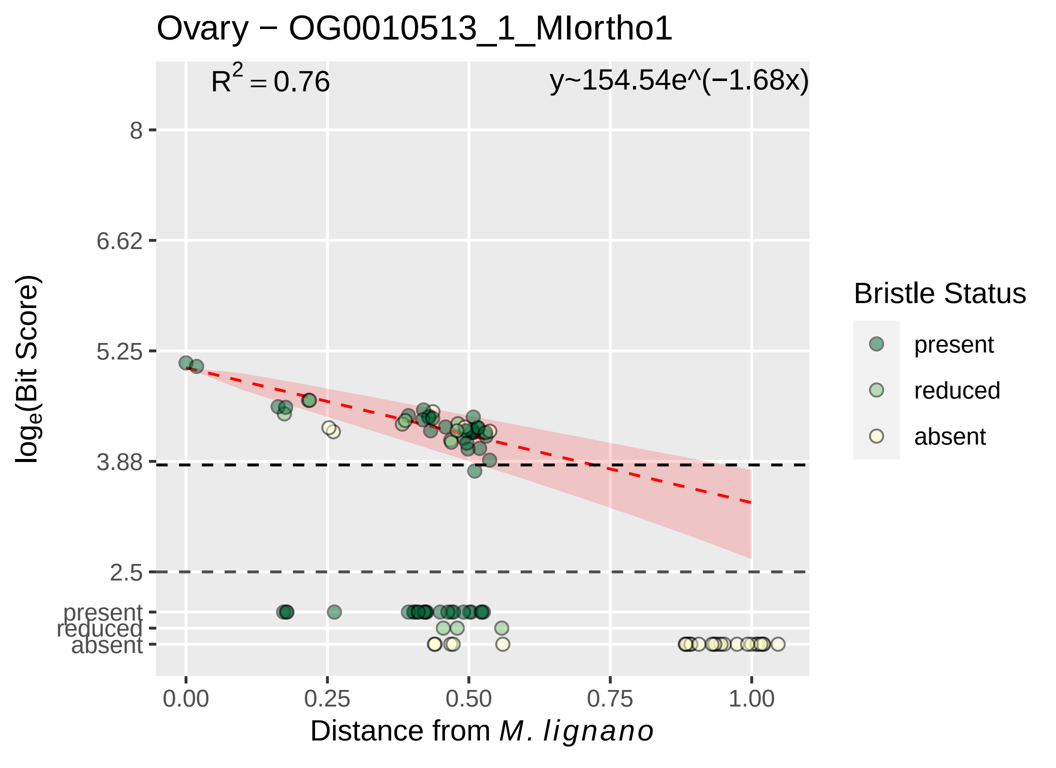
Ovary − OG0010513_1_MIortho1
R2 = 0.76
y~154.54e^(−1.68x)
8
6.62
5.25
3.88
2.5
present
reduced
absent
log_e(Bit Score)
0.00
0.25
0.50
0.75
1.00
Distance from M. lignano
Bristle Status
present
reduced
absent

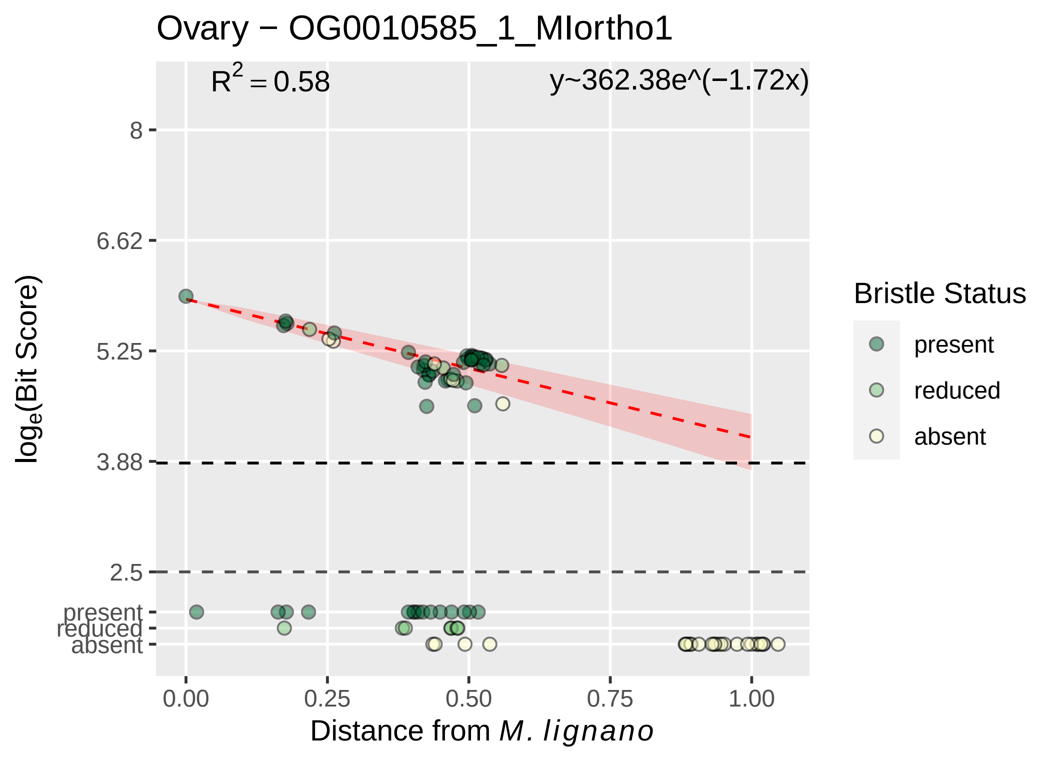

Ovary − OG0010585_1_MIortho1
$R^2 = 0.58$
y~362.38e^(−1.72x)
8
6.62
5.25
3.88
2.5
present
reduced
absent
0.00
0.25
0.50
0.75
1.00
Distance from *M. lignano*
$\log_e$(Bit Score)
Bristle Status
present
reduced
absent

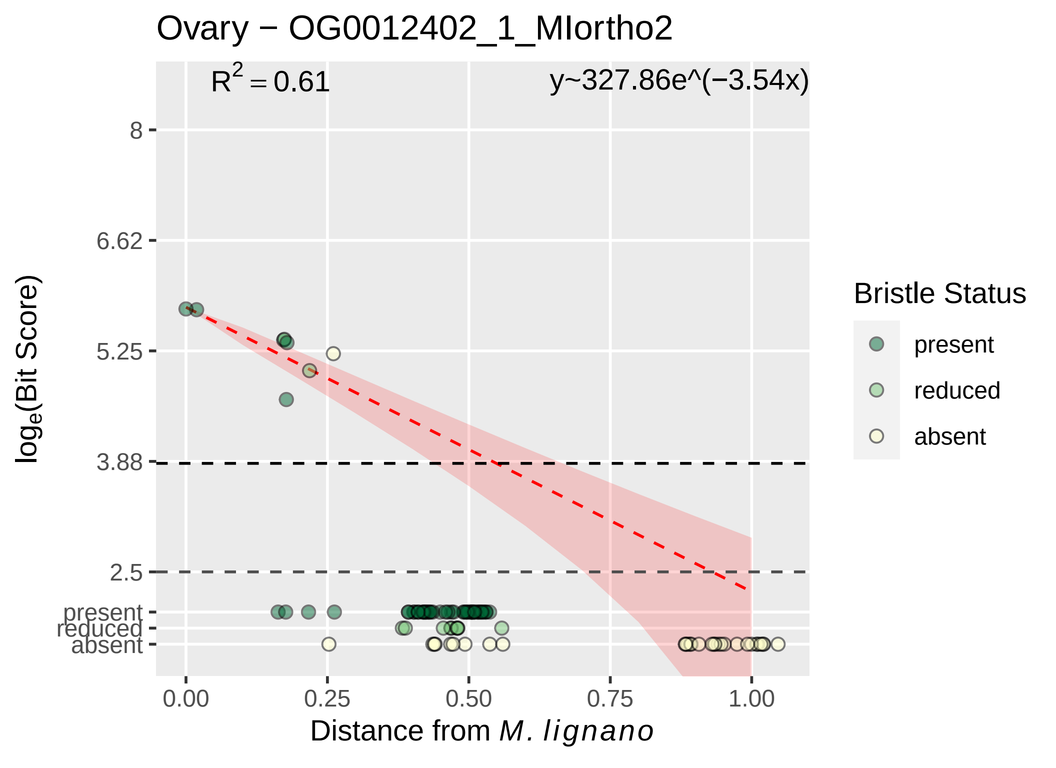

Ovary − OG0012402_1_MIortho2
R2 = 0.61
y~327.86e^(−3.54x)
8
6.62
5.25
3.88
2.5
present
reduced
absent
loge(Bit Score)
0.00
0.25
0.50
0.75
1.00
Distance from M. lignano
Bristle Status
present
reduced
absent

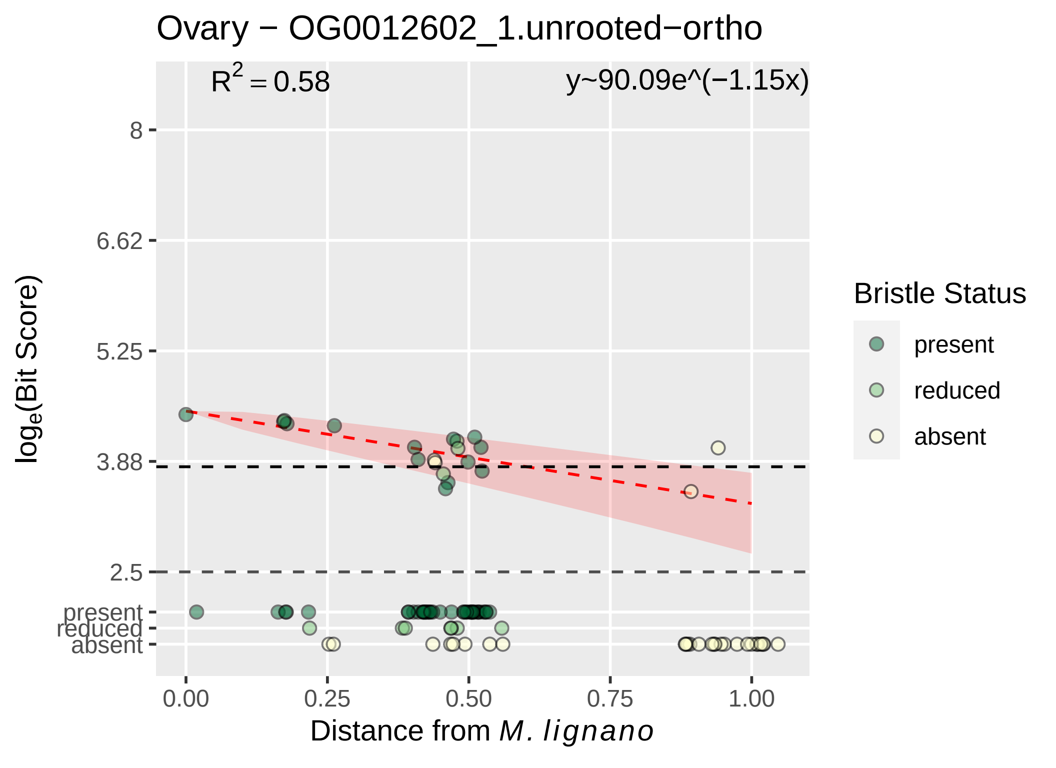

Ovary − OG0012602_1.unrooted−ortho
$R^2 = 0.58$
y~90.09e^(−1.15x)
8
6.62
5.25
3.88
2.5
present
reduced
absent
$\log_e$(Bit Score)
0.00
0.25
0.50
0.75
1.00
Distance from *M. lignano*
Bristle Status
present
reduced
absent

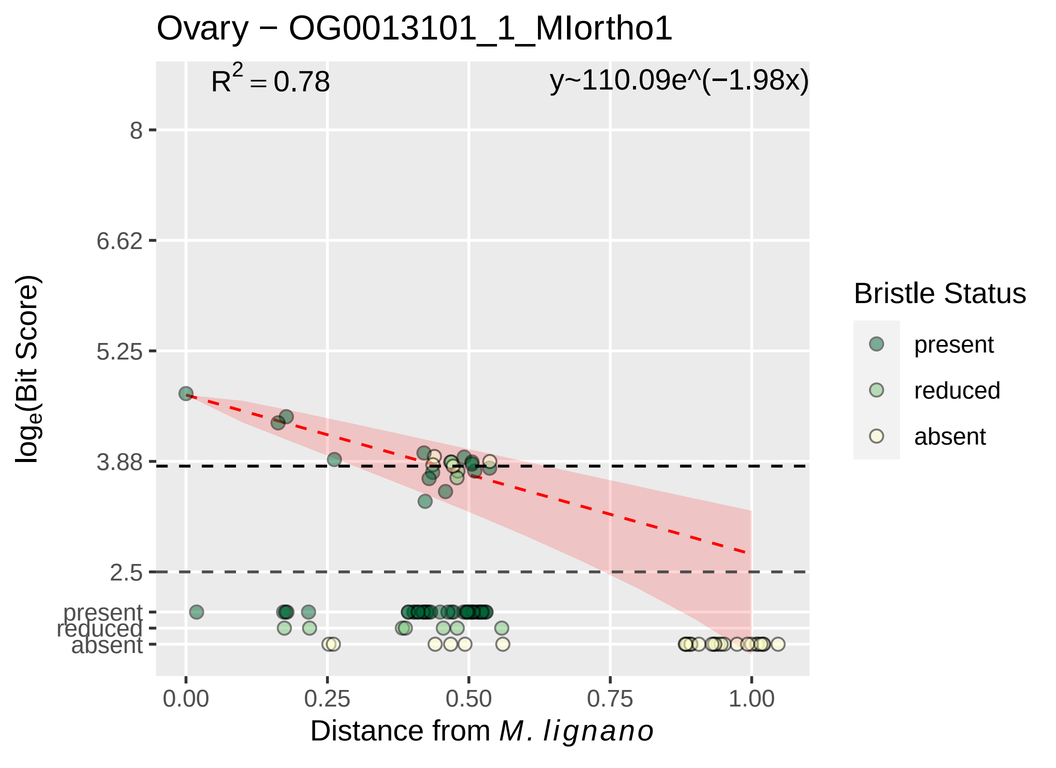

Ovary − OG0013101_1_MIortho1
R^2 = 0.78
y~110.09e^(−1.98x)
8
6.62
5.25
3.88
2.5
present
reduced
absent
log_e(Bit Score)
0.00
0.25
0.50
0.75
1.00
Distance from M. lignano
Bristle Status
present
reduced
absent

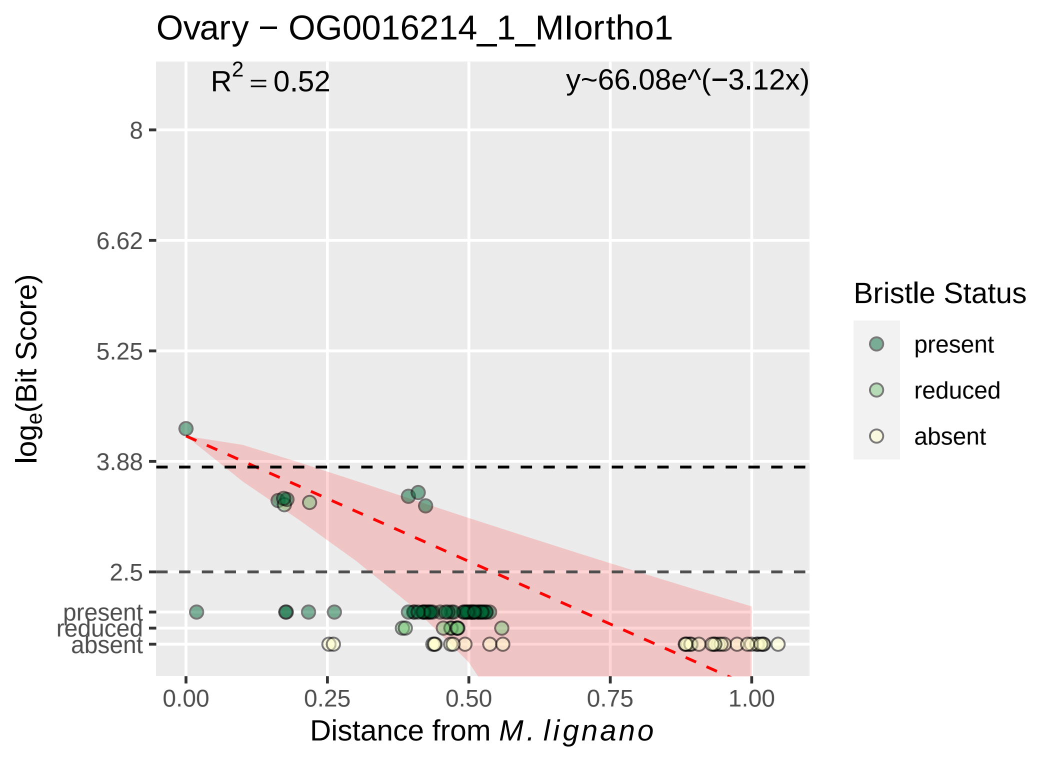

Ovary − OG0016214_1_MIortho1
R^2 = 0.52
y~66.08e^(−3.12x)
8
6.62
5.25
3.88
2.5
present
reduced
absent
0.00
0.25
0.50
0.75
1.00
Distance from M. lignano
loge(Bit Score)
Bristle Status
present
reduced
absent

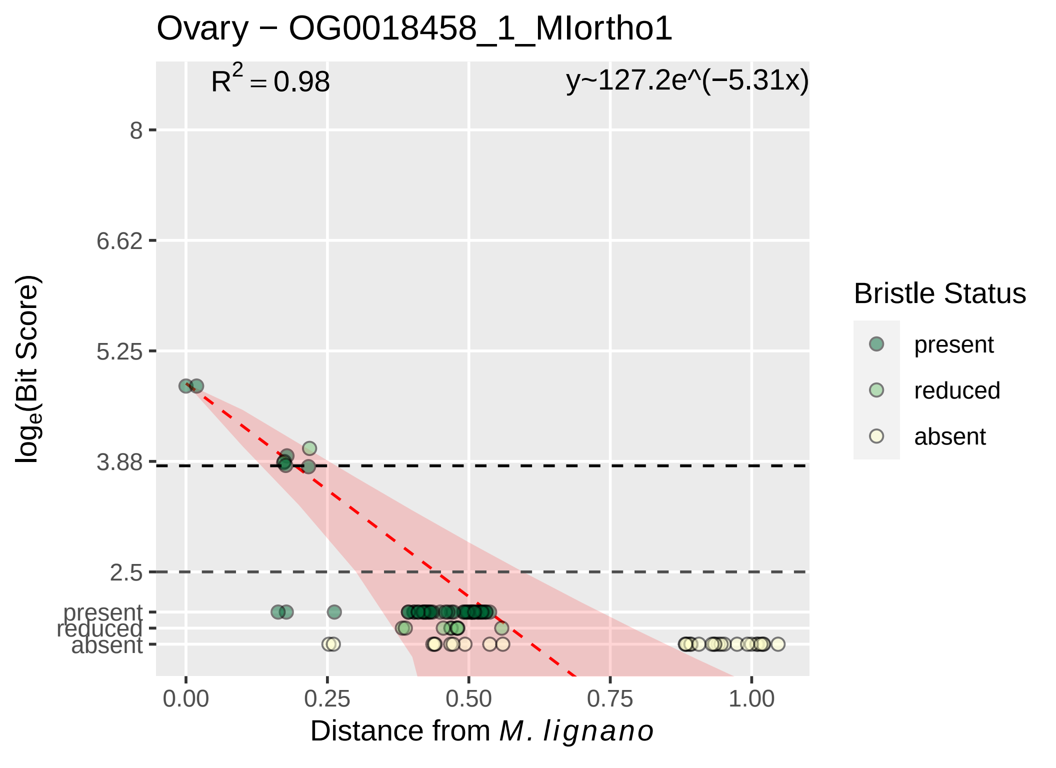

Ovary − OG0018458_1_MIortho1
R^2 = 0.98
y~127.2e^(−5.31x)
8
6.62
5.25
3.88
2.5
present
reduced
absent
0.00
0.25
0.50
0.75
1.00
Distance from M. lignano
loge(Bit Score)
Bristle Status
present
reduced
absent
