## Supplementary material for "Mating strategy predicts gene presence/absence patterns in a genus of simultaneously hermaphroditic flatworms": figure S5C

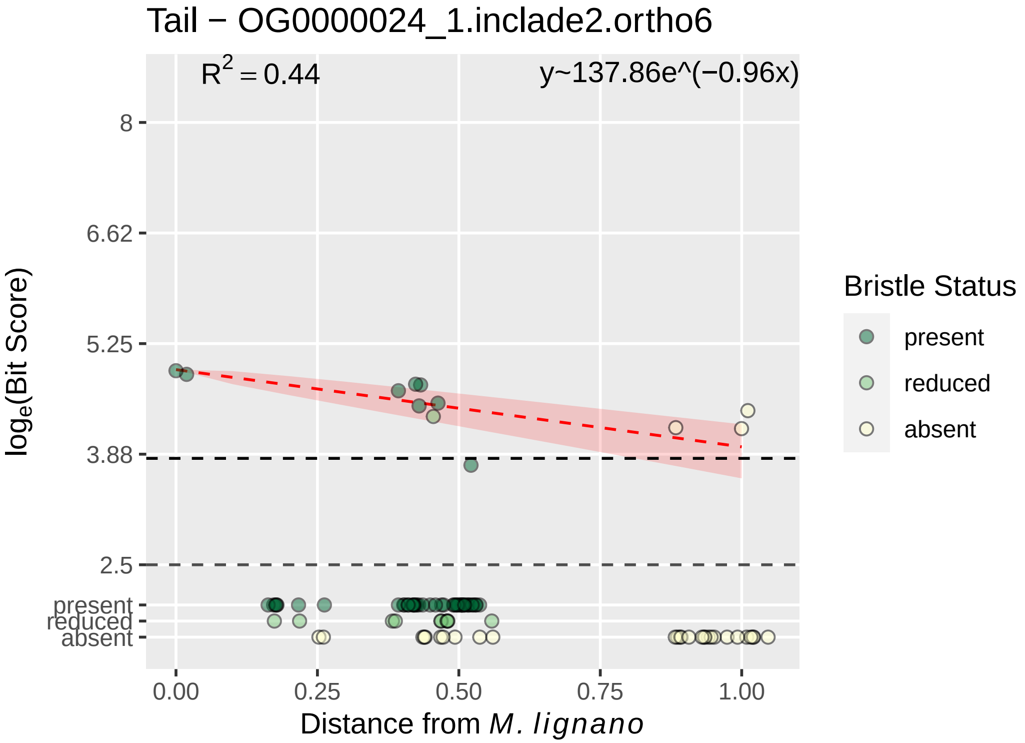

Tail − OG0000024_1.inclade2.ortho6
R2 = 0.44
y~137.86e^(−0.96x)
8
6.62
5.25
3.88
2.5
present
reduced
absent
loge(Bit Score)
0.00
0.25
0.50
0.75
1.00
Distance from M. lignano
Bristle Status
present
reduced
absent

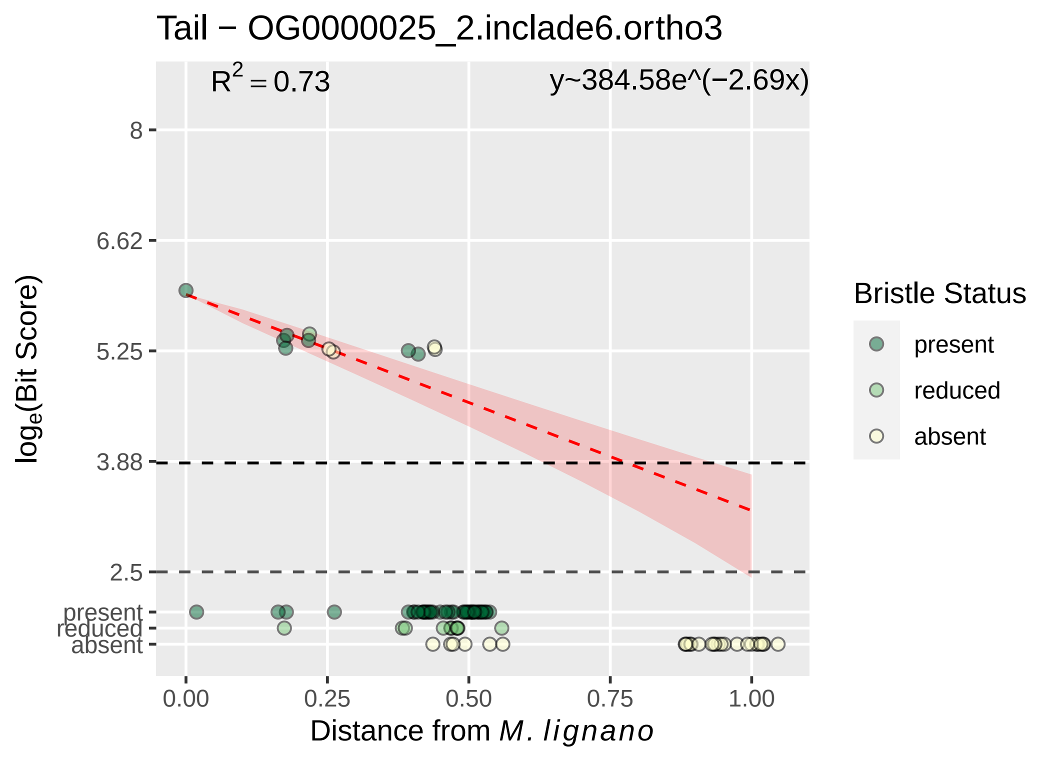

Tail − OG0000025_2.inclade6.ortho3
R^2 = 0.73
y~384.58e^(−2.69x)
8
6.62
5.25
3.88
2.5
present
reduced
absent
0.00
0.25
0.50
0.75
1.00
log_e(Bit Score)
Distance from M. lignano
Bristle Status
present
reduced
absent

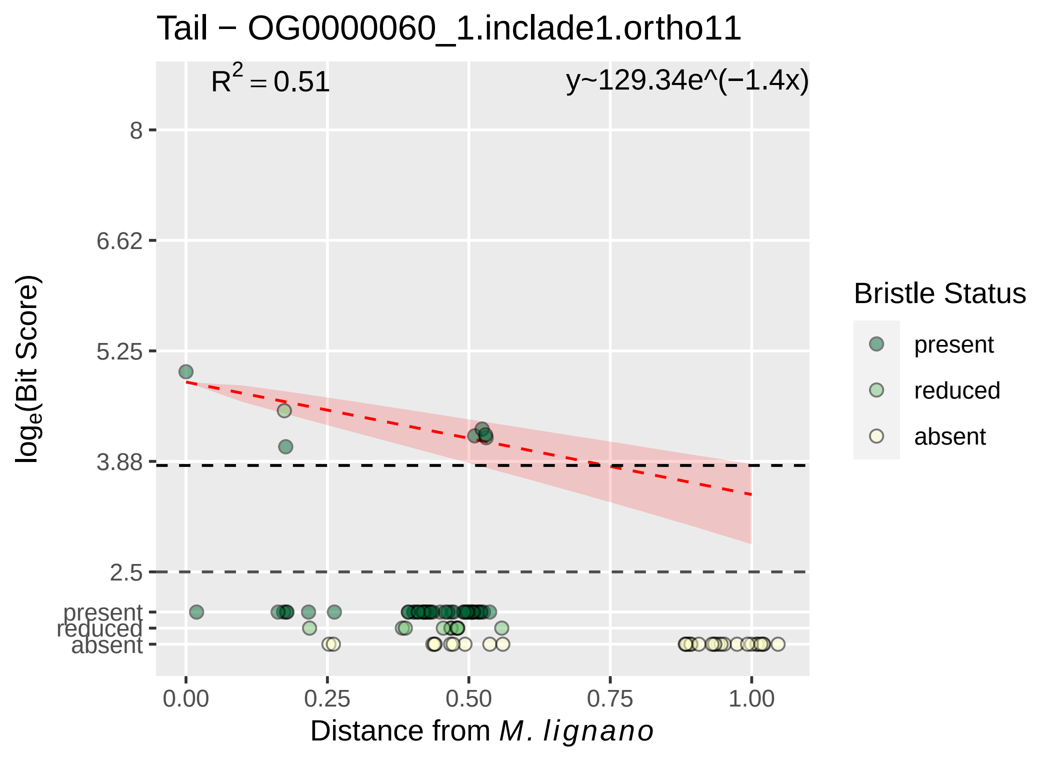

Tail − OG0000060_1.inclade1.ortho11
R^2 = 0.51
y~129.34e^(−1.4x)
8
6.62
5.25
3.88
2.5
present
reduced
absent
0.00
0.25
0.50
0.75
1.00
loge(Bit Score)
Distance from M. lignano
Bristle Status
present
reduced
absent

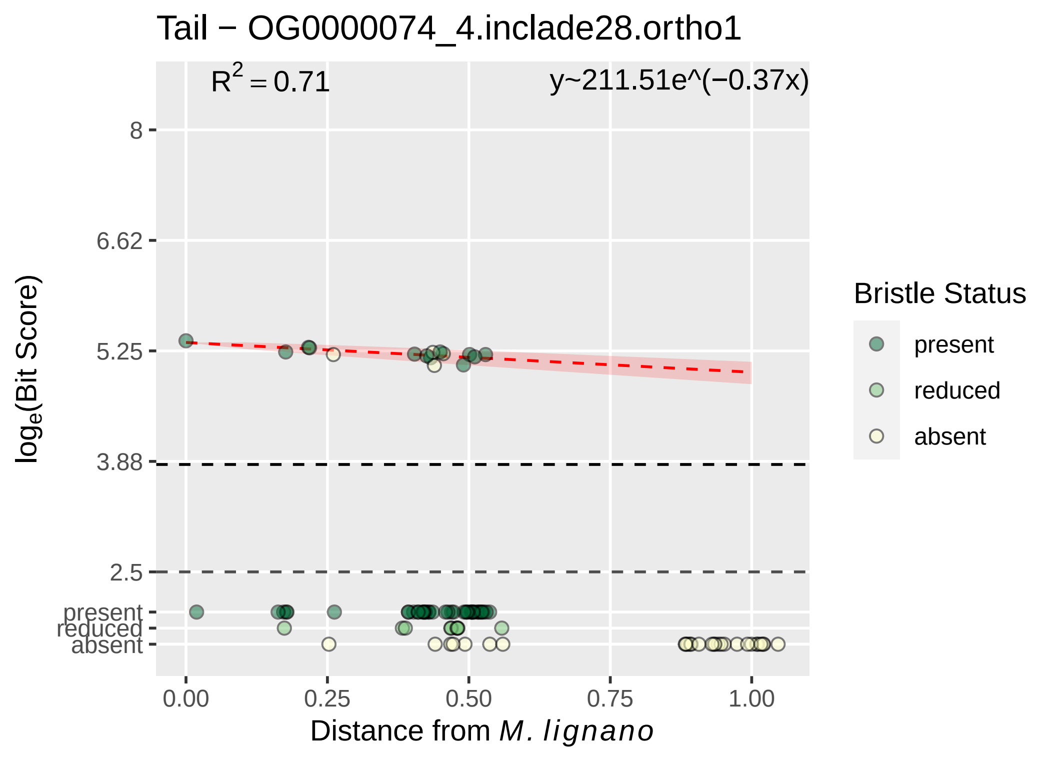

Tail − OG0000074_4.inclade28.ortho1
R^2 = 0.71
y~211.51e^(−0.37x)
8
6.62
5.25
3.88
2.5
present
reduced
absent
0.00
0.25
0.50
0.75
1.00
log_e(Bit Score)
Distance from M. lignano
Bristle Status
present
reduced
absent

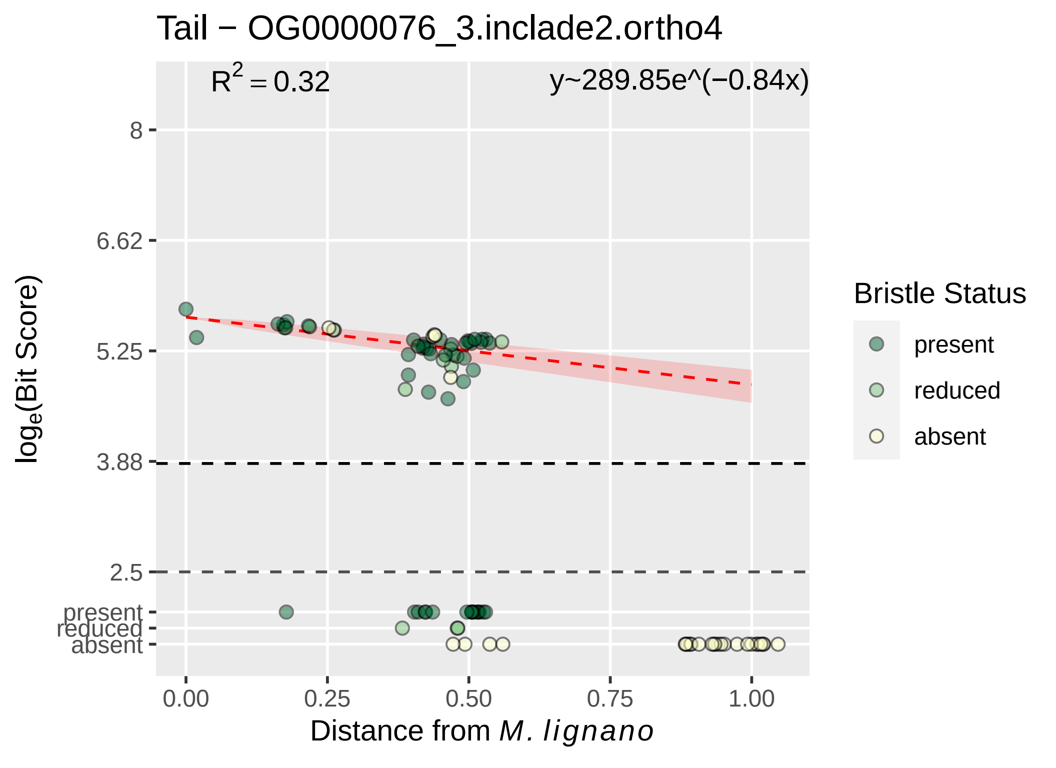

Tail − OG0000076_3.inclade2.ortho4
R^2 = 0.32
y~289.85e^(−0.84x)
8
6.62
5.25
3.88
2.5
present
reduced
absent
0.00
0.25
0.50
0.75
1.00
Distance from M. lignano
loge(Bit Score)
Bristle Status
present
reduced
absent

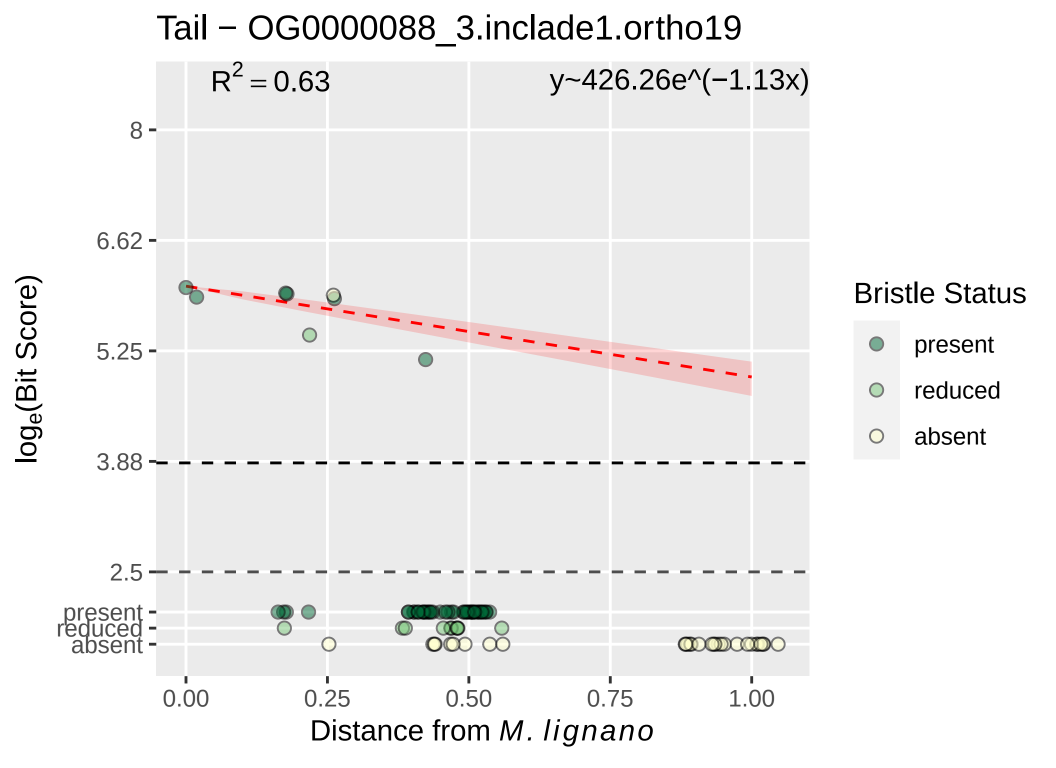
Tail − OG0000088_3.inclade1.ortho19
R2 = 0.63
y~426.26e^(−1.13x)
8
6.62
5.25
3.88
2.5
present
reduced
absent
loge(Bit Score)
0.00
0.25
0.50
0.75
1.00
Distance from M. lignano
Bristle Status
present
reduced
absent

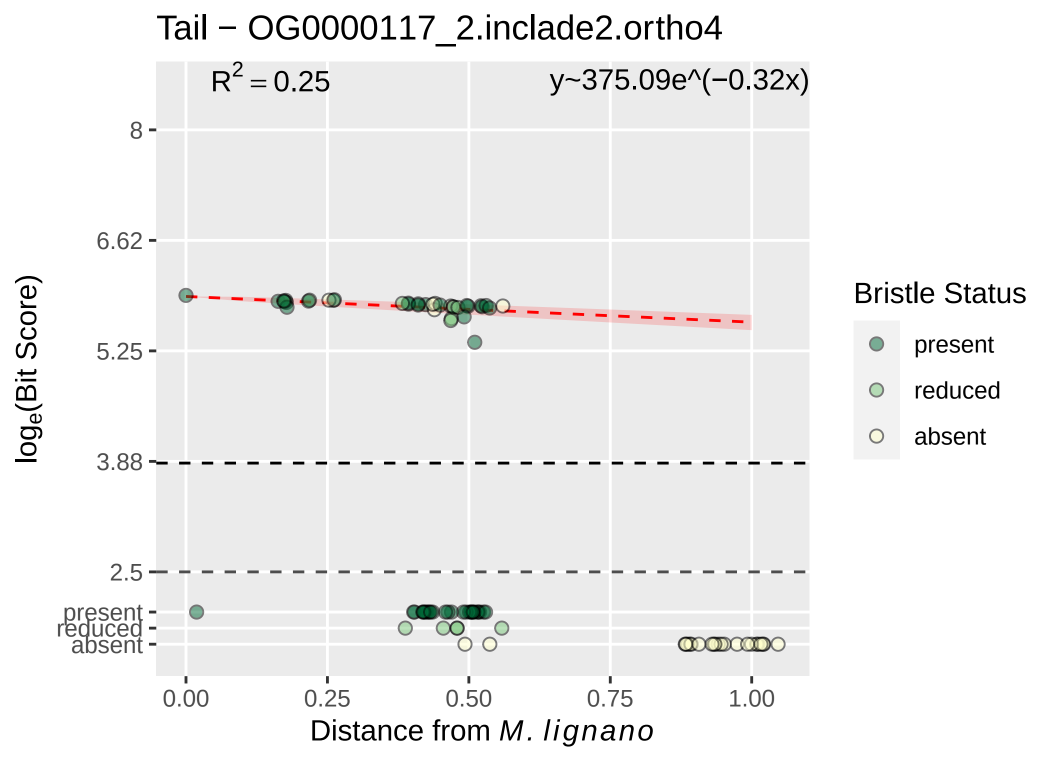
Tail − OG0000117_2.inclade2.ortho4
$R^2 = 0.25$
y~375.09e^(−0.32x)
8
6.62
5.25
3.88
2.5
present
reduced
absent
$\log_e$(Bit Score)
0.00
0.25
0.50
0.75
1.00
Distance from *M. lignano*
Bristle Status
present
reduced
absent

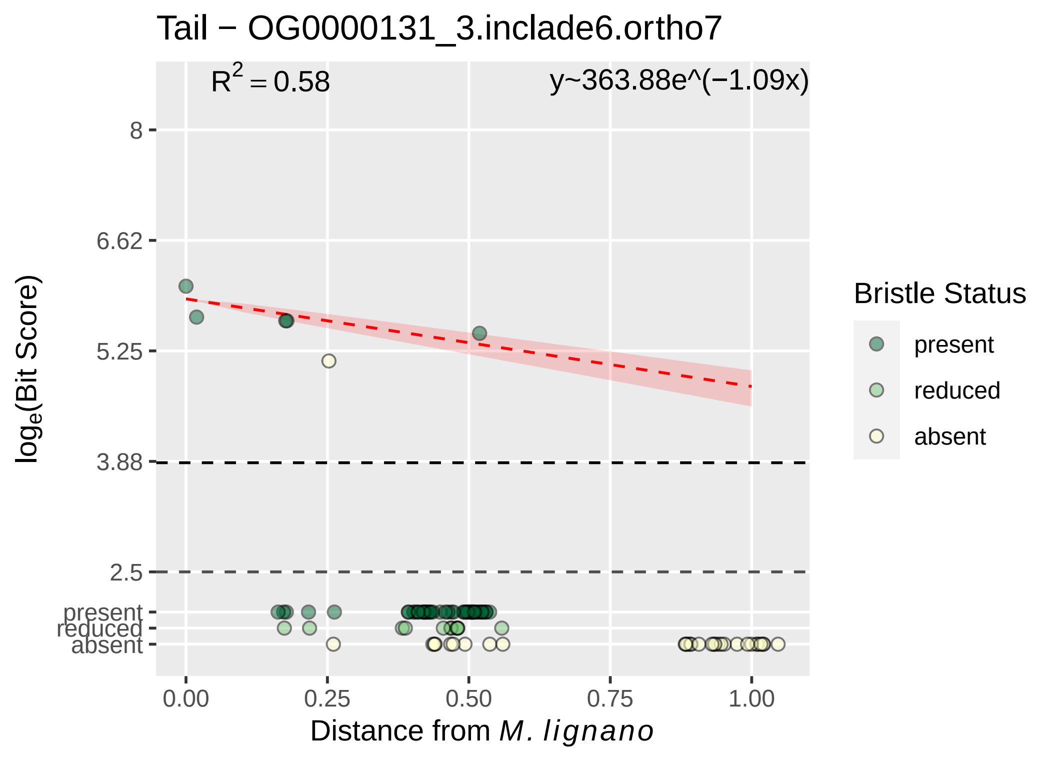

Tail − OG0000131_3.inclade6.ortho7
$R^2 = 0.58$
y~363.88e^(−1.09x)
8
6.62
5.25
3.88
2.5
present
reduced
absent
$\log_e$(Bit Score)
0.00
0.25
0.50
0.75
1.00
Distance from *M. lignano*
Bristle Status
present
reduced
absent

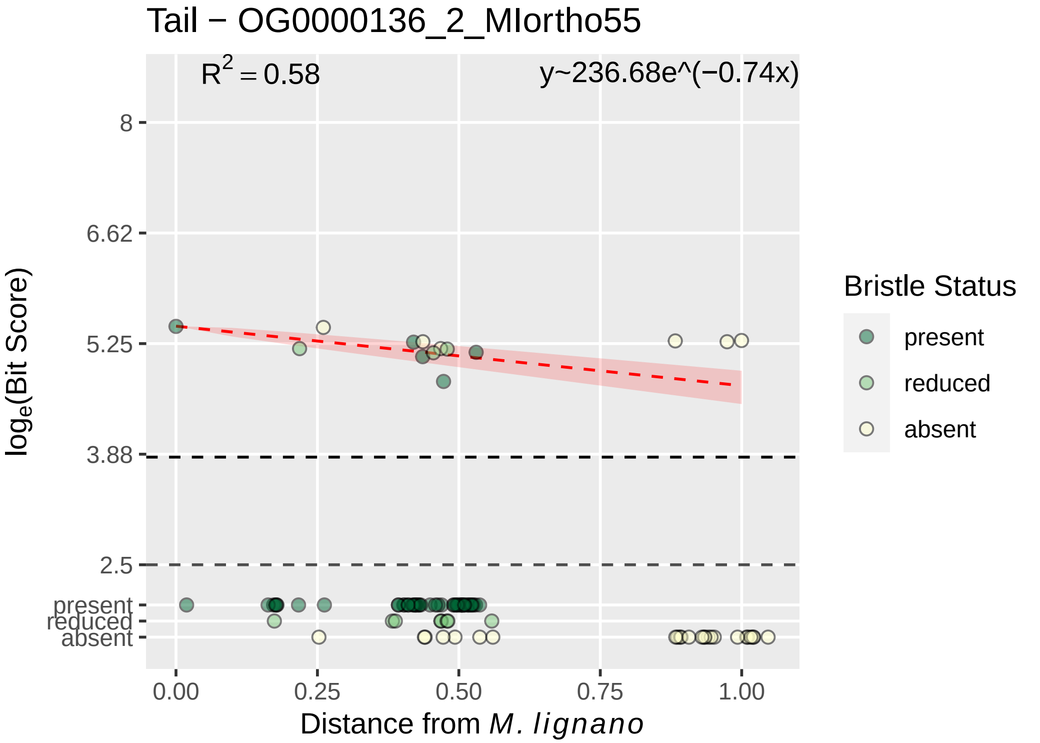
Tail − OG0000136_2_MIortho55
R^2 = 0.58
y~236.68e^(−0.74x)
8
6.62
5.25
3.88
2.5
present
reduced
absent
0.00
0.25
0.50
0.75
1.00
log_e(Bit Score)
Distance from M. lignano
Bristle Status
present
reduced
absent

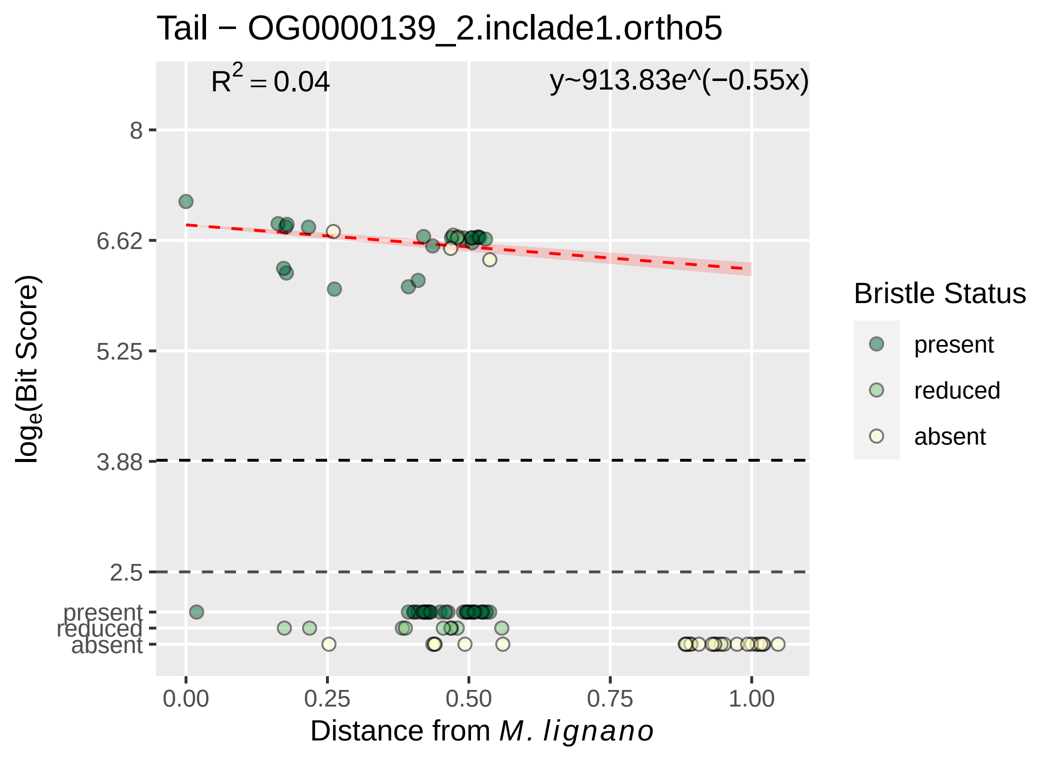

Tail − OG0000139_2.inclade1.ortho5
R^2 = 0.04
y~913.83e^(−0.55x)
8
6.62
5.25
3.88
2.5
present
reduced
absent
log_e(Bit Score)
0.00
0.25
0.50
0.75
1.00
Distance from M. lignano
Bristle Status
present
reduced
absent

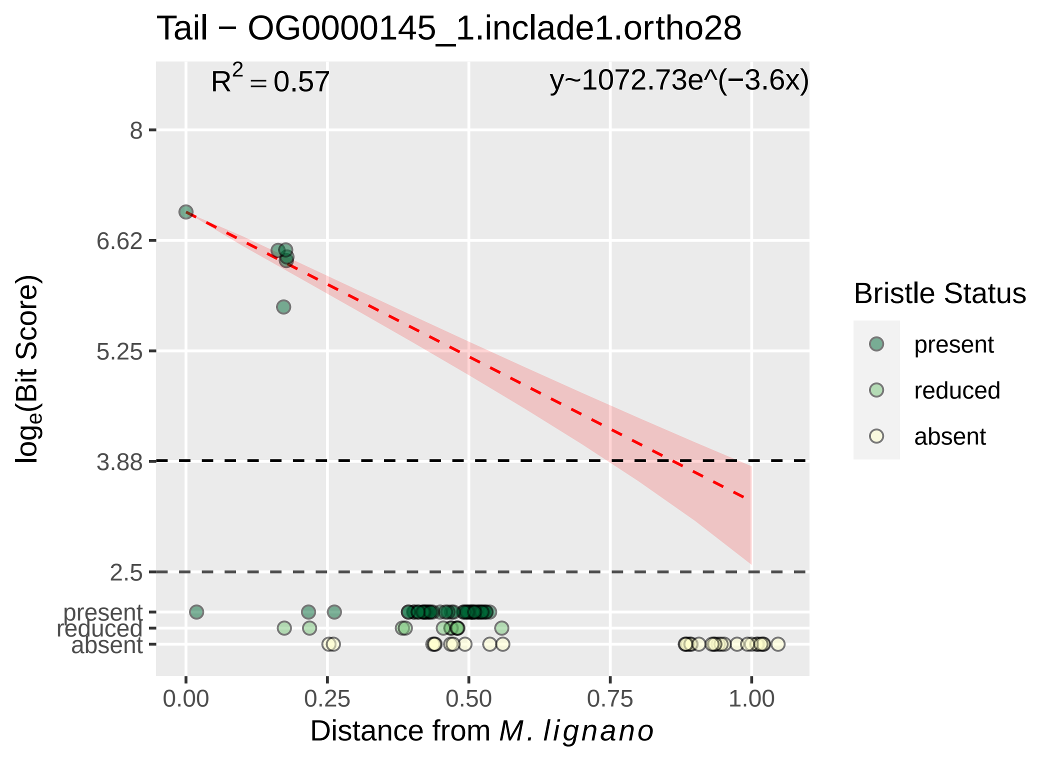

Tail − OG0000145_1.inclade1.ortho28
$R^2 = 0.57$
y~1072.73e^(−3.6x)
8
6.62
5.25
3.88
2.5
present
reduced
absent
0.00
0.25
0.50
0.75
1.00
$\log_e$(Bit Score)
Distance from *M. lignano*
Bristle Status
present
reduced
absent

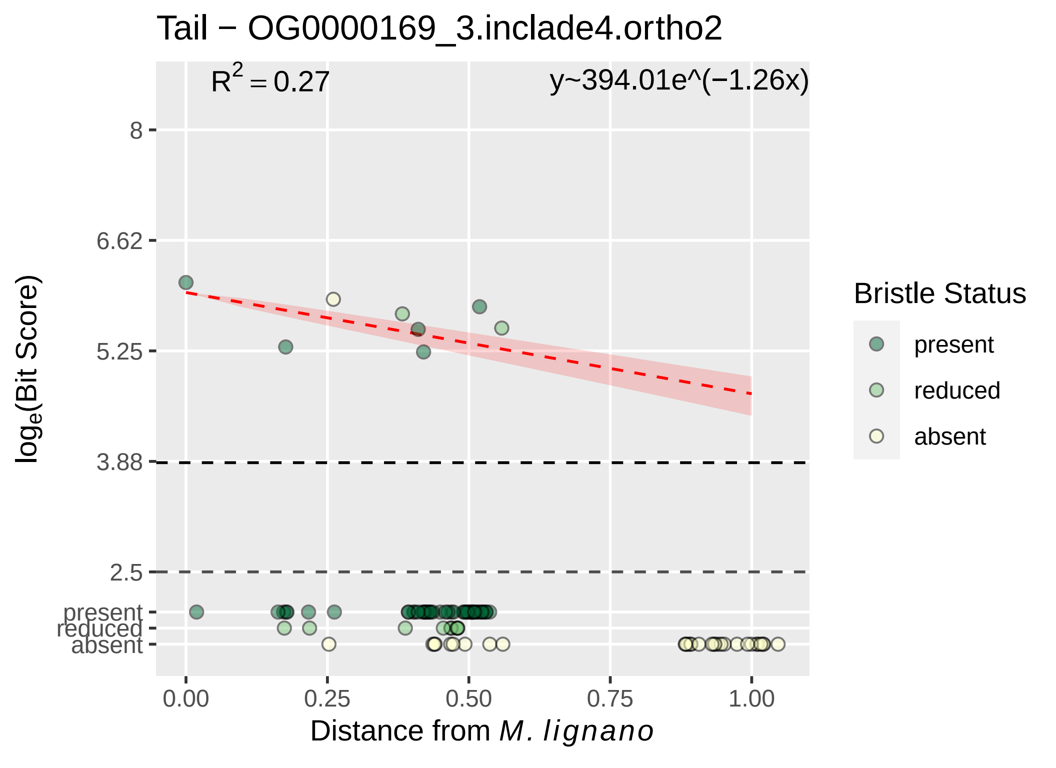

Tail − OG0000169_3.inclade4.ortho2
$R^2 = 0.27$
y~394.01e^(−1.26x)
8
6.62
5.25
3.88
2.5
present
reduced
absent
$log_e$(Bit Score)
0.00
0.25
0.50
0.75
1.00
Distance from *M. lignano*
Bristle Status
present
reduced
absent

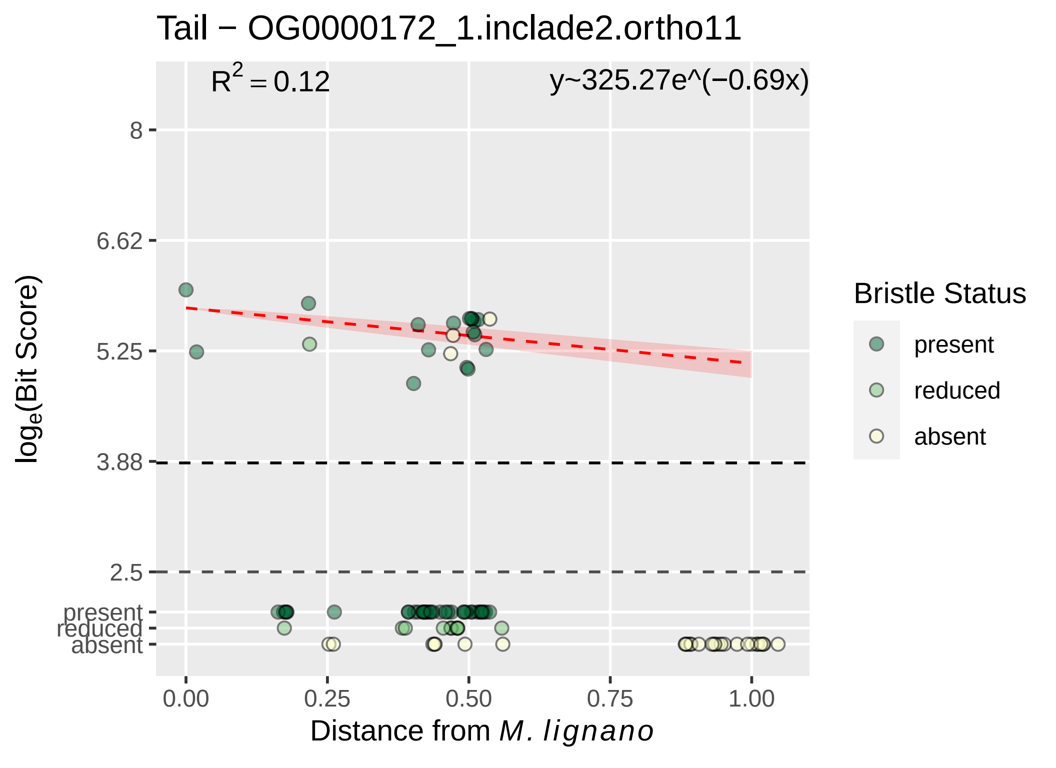

Tail − OG0000172_1.inclade2.ortho11
$R^2 = 0.12$
y~325.27e^(−0.69x)
8
6.62
5.25
3.88
2.5
present
reduced
absent
0.00
0.25
0.50
0.75
1.00
$\log_e$(Bit Score)
Distance from *M. lignano*
Bristle Status
present
reduced
absent

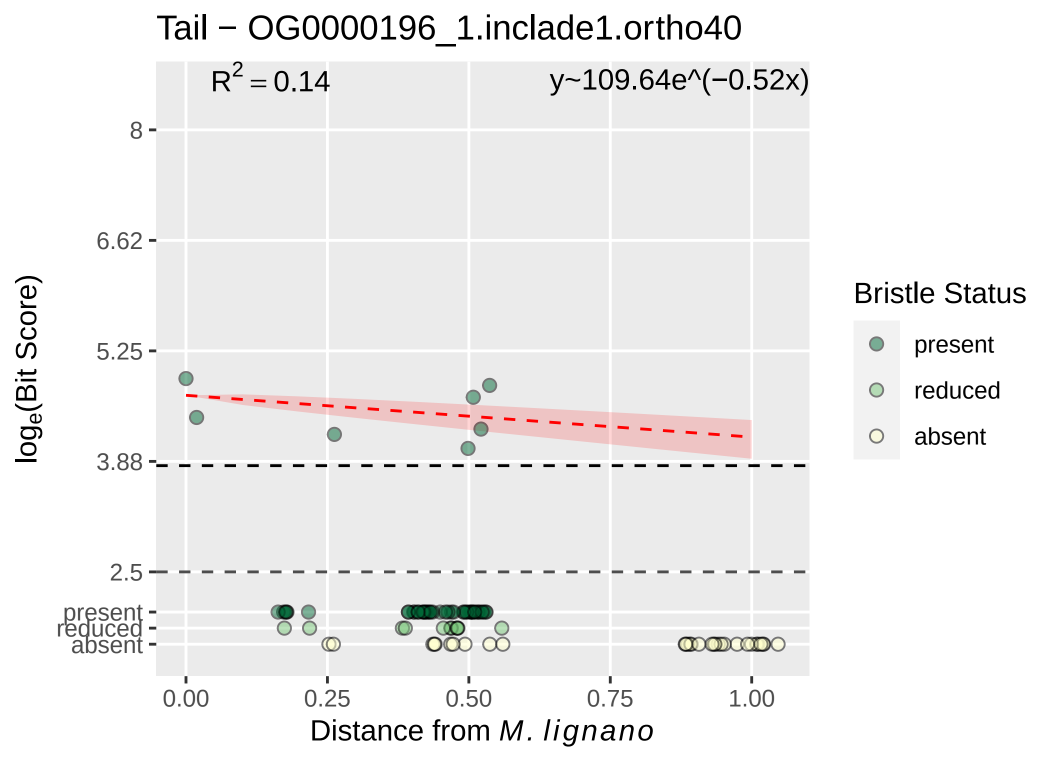

Tail − OG0000196_1.inclade1.ortho40
$R^2 = 0.14$
y~109.64e^(−0.52x)
8
6.62
5.25
3.88
2.5
present
reduced
absent
log_e(Bit Score)
0.00
0.25
0.50
0.75
1.00
Distance from M. lignano
Bristle Status
present
reduced
absent

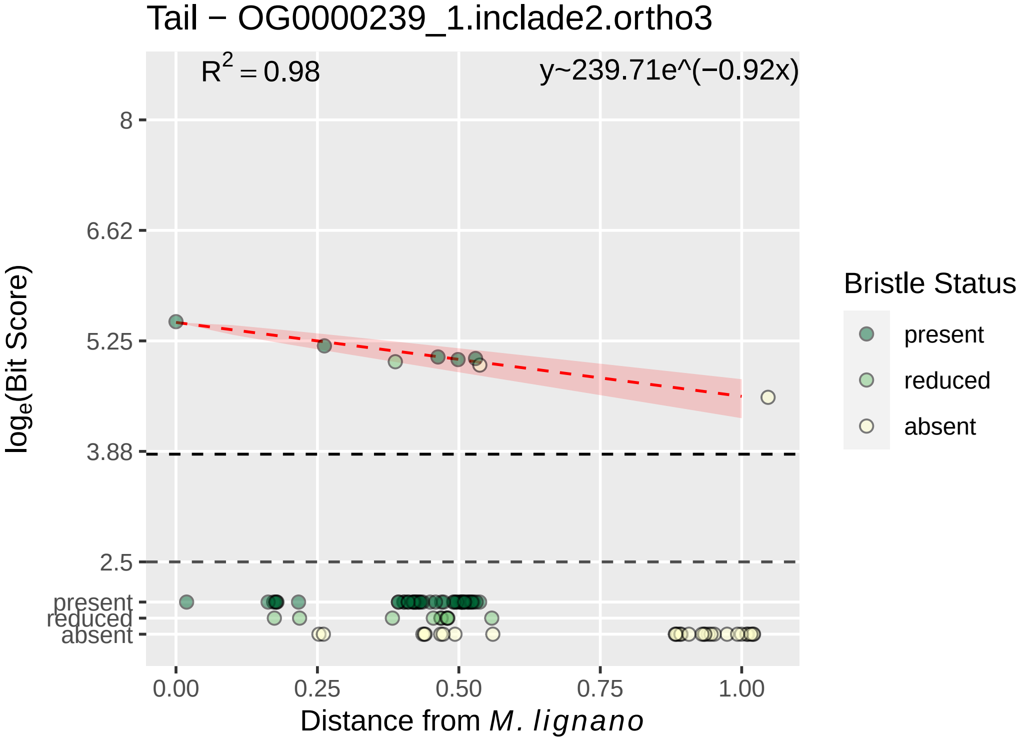

Tail − OG0000239_1.inclade2.ortho3
$R^2 = 0.98$
y~239.71e^(−0.92x)
8
6.62
5.25
3.88
2.5
present
reduced
absent
log_e(Bit Score)
0.00
0.25
0.50
0.75
1.00
Distance from M. lignano
Bristle Status
present
reduced
absent

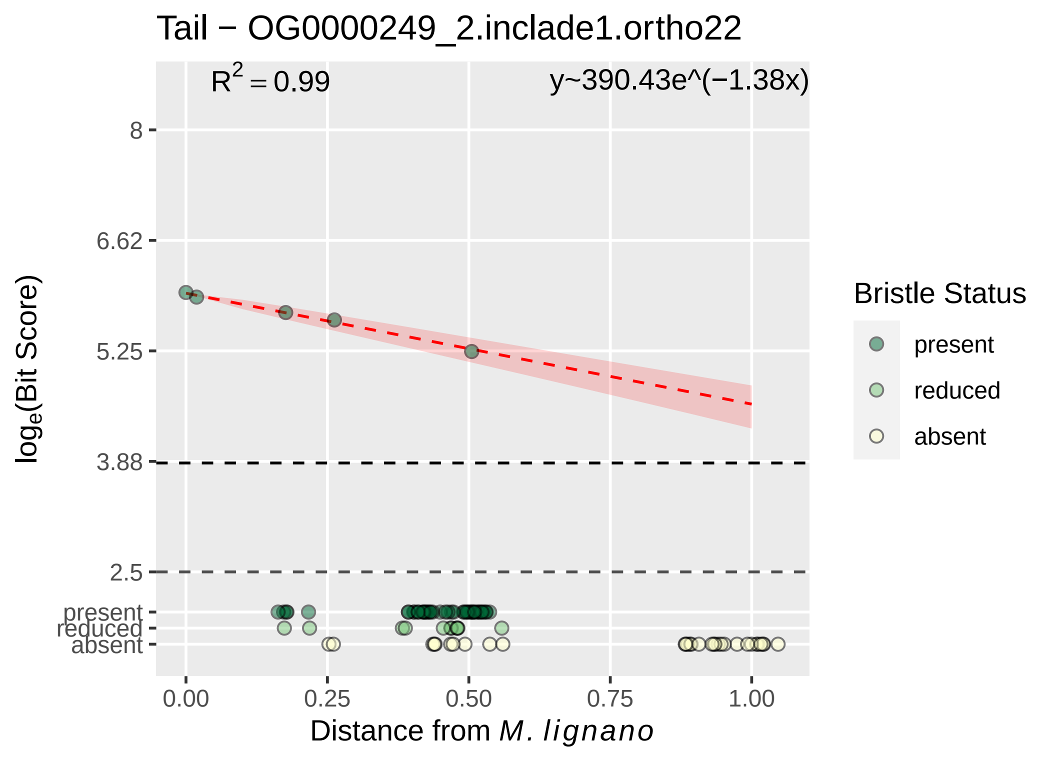

Tail − OG0000249_2.inclade1.ortho22
$R^2 = 0.99$
y~390.43e^(−1.38x)
8
6.62
5.25
3.88
2.5
present
reduced
absent
$\log_e$(Bit Score)
0.00
0.25
0.50
0.75
1.00
Distance from *M. lignano*
Bristle Status
present
reduced
absent

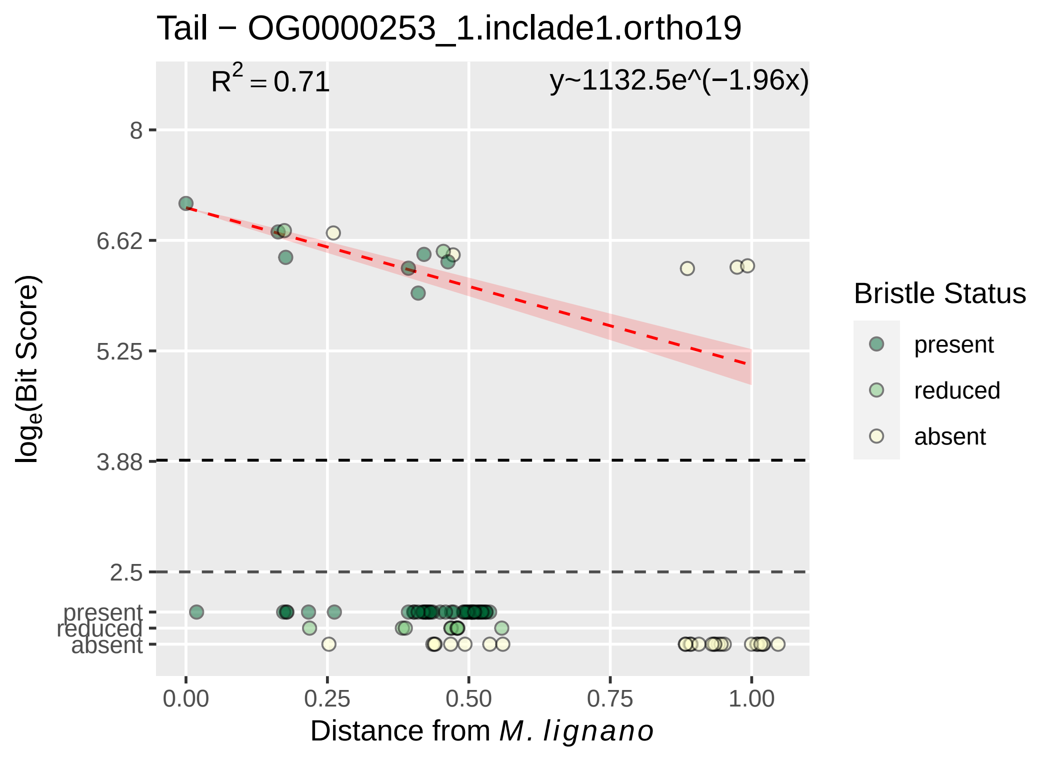

Tail − OG0000253_1.inclade1.ortho19
R2 = 0.71
y~1132.5e^(−1.96x)
8
6.62
5.25
3.88
2.5
present
reduced
absent
0.00
0.25
0.50
0.75
1.00
loge(Bit Score)
Distance from M. lignano
Bristle Status
present
reduced
absent

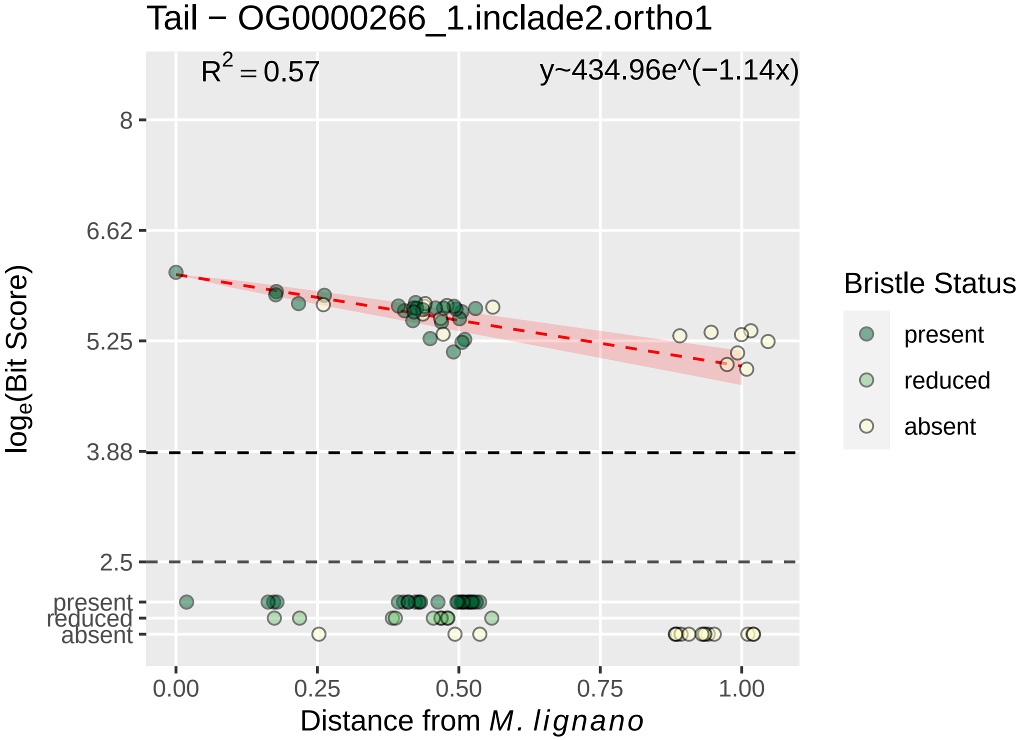

Tail − OG0000266_1.inclade2.ortho1
$R^2 = 0.57$
y~434.96e^(−1.14x)
8
6.62
5.25
3.88
2.5
present
reduced
absent
0.00
0.25
0.50
0.75
1.00
Distance from M. lignano
$\log_e$(Bit Score)
Bristle Status
present
reduced
absent

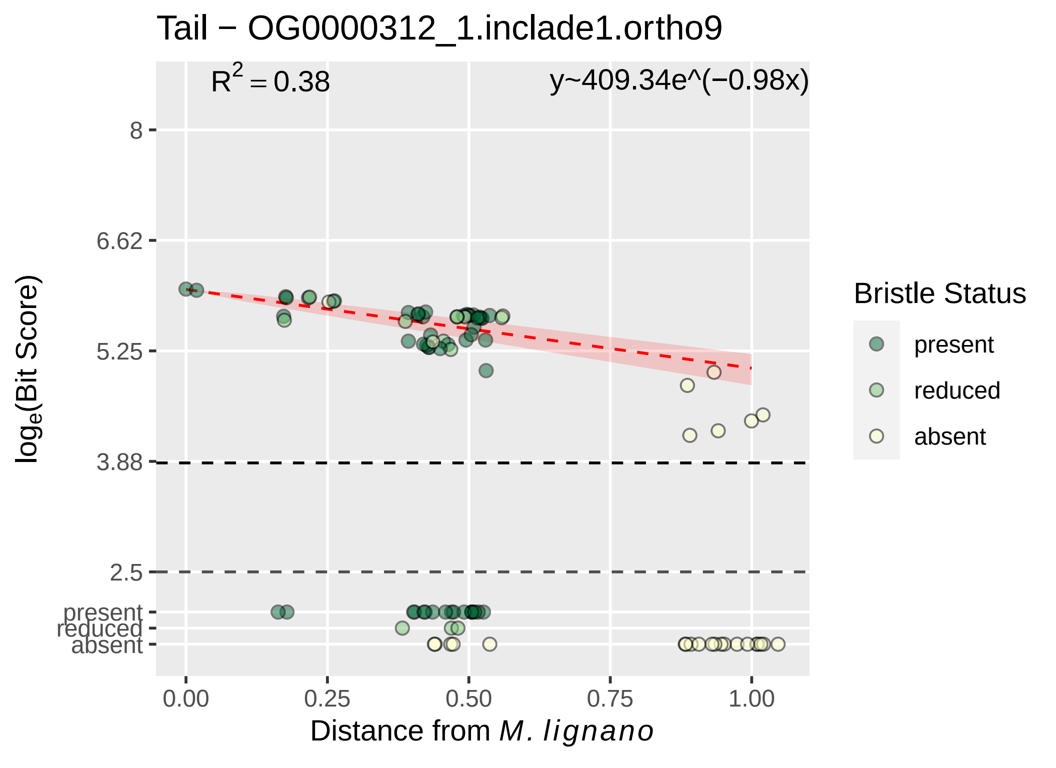

Tail − OG0000312_1.inclade1.ortho9
$R^2 = 0.38$
y~409.34e^(−0.98x)
8
6.62
5.25
3.88
2.5
present
reduced
absent
0.00
0.25
0.50
0.75
1.00
$\log_e$(Bit Score)
Distance from *M. lignano*
Bristle Status
present
reduced
absent

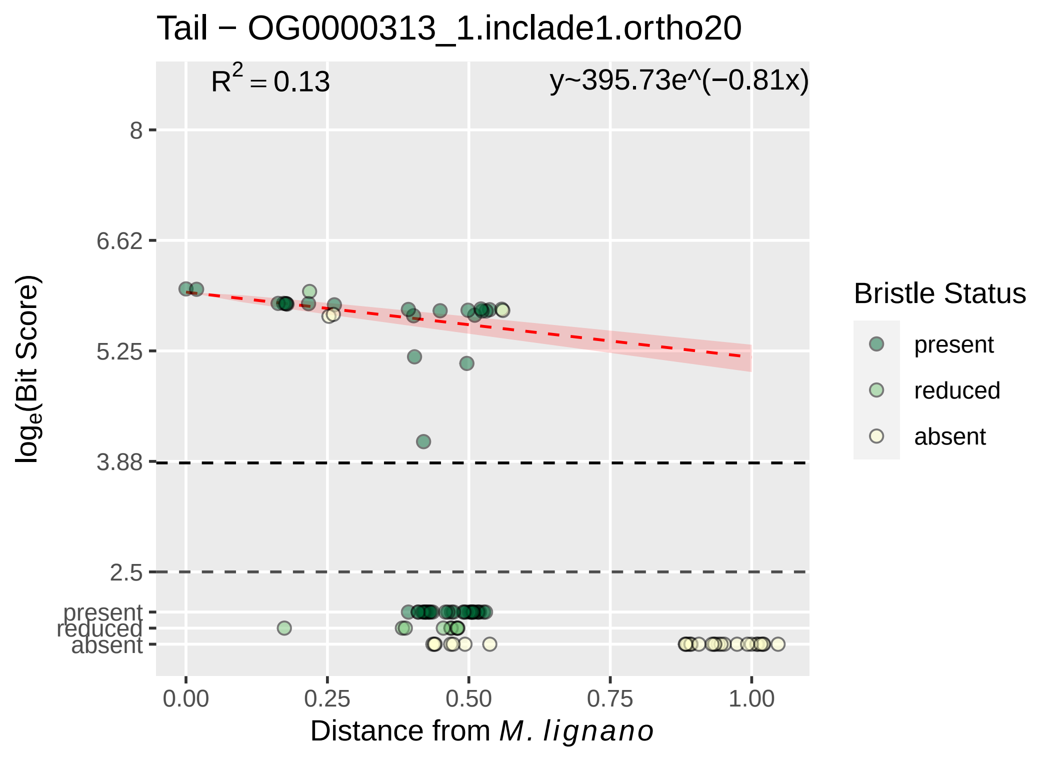
Tail − OG0000313_1.inclade1.ortho20
R² = 0.13
y~395.73e^(−0.81x)
8
6.62
5.25
3.88
2.5
present
reduced
absent
0.00
0.25
0.50
0.75
1.00
Distance from M. lignano
loge(Bit Score)
Bristle Status
present
reduced
absent

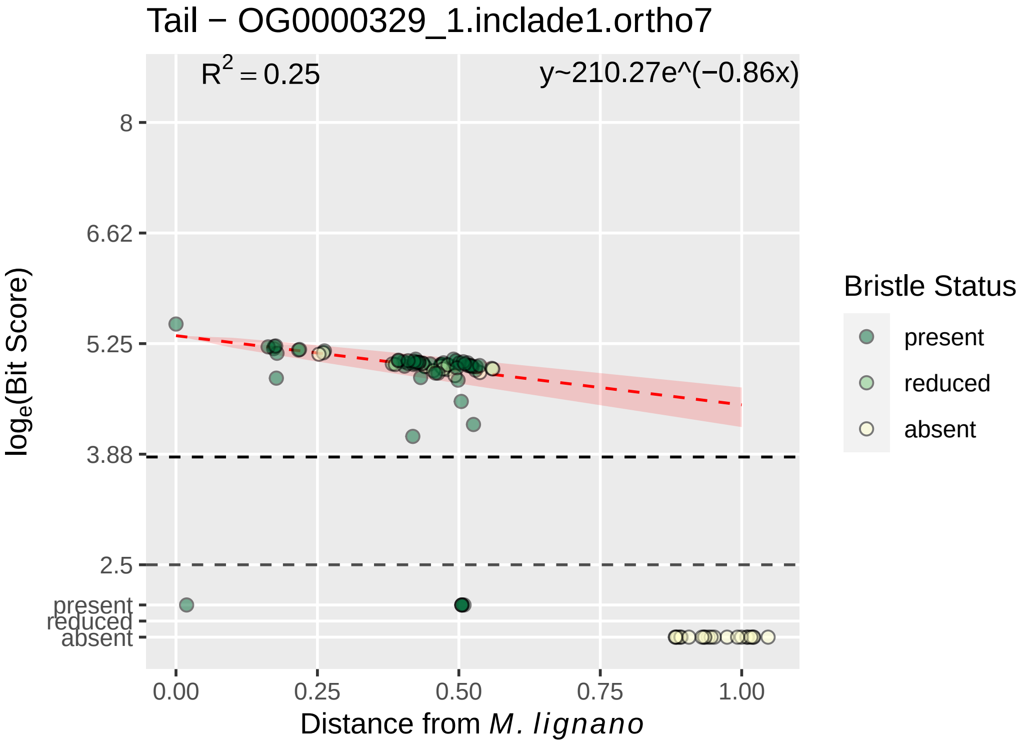

Tail − OG0000329_1.inclade1.ortho7
$R^2 = 0.25$
y~210.27e^(−0.86x)
8
6.62
5.25
3.88
2.5
present
reduced
absent
0.00
0.25
0.50
0.75
1.00
Distance from *M. lignano*
$\log_e$(Bit Score)
Bristle Status
present
reduced
absent

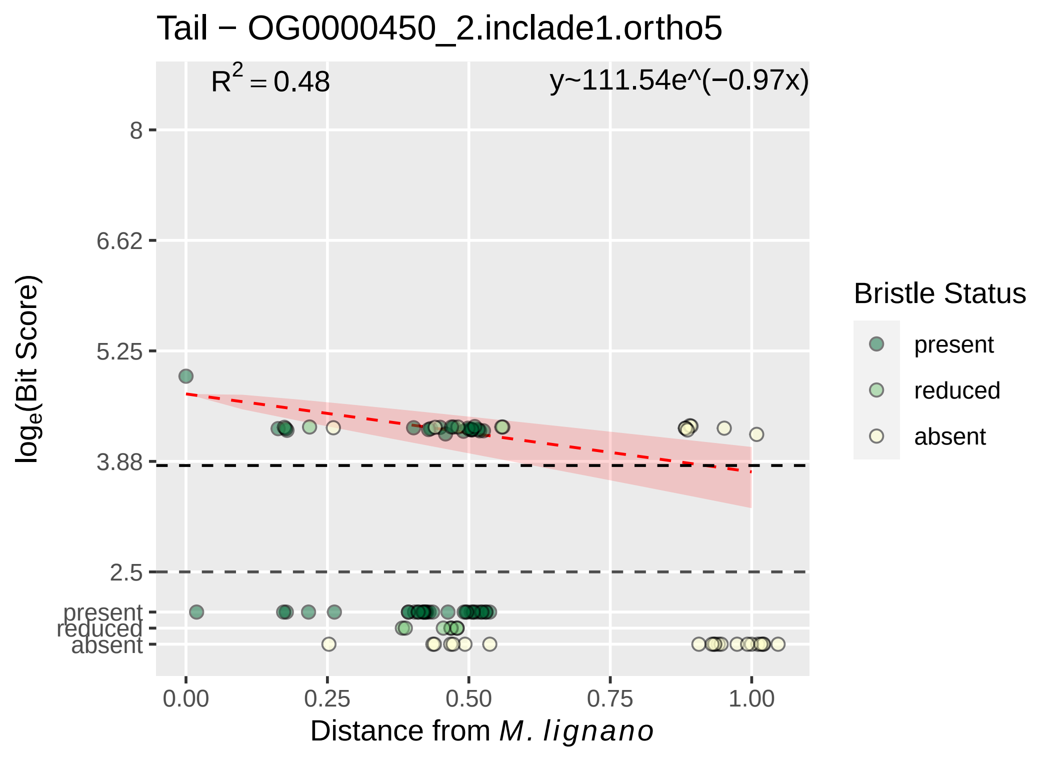

Tail − OG0000450_2.inclade1.ortho5
R² = 0.48
y~111.54e^(−0.97x)
8
6.62
5.25
3.88
2.5
present
reduced
absent
log_e(Bit Score)
0.00
0.25
0.50
0.75
1.00
Distance from M. lignano
Bristle Status
present
reduced
absent

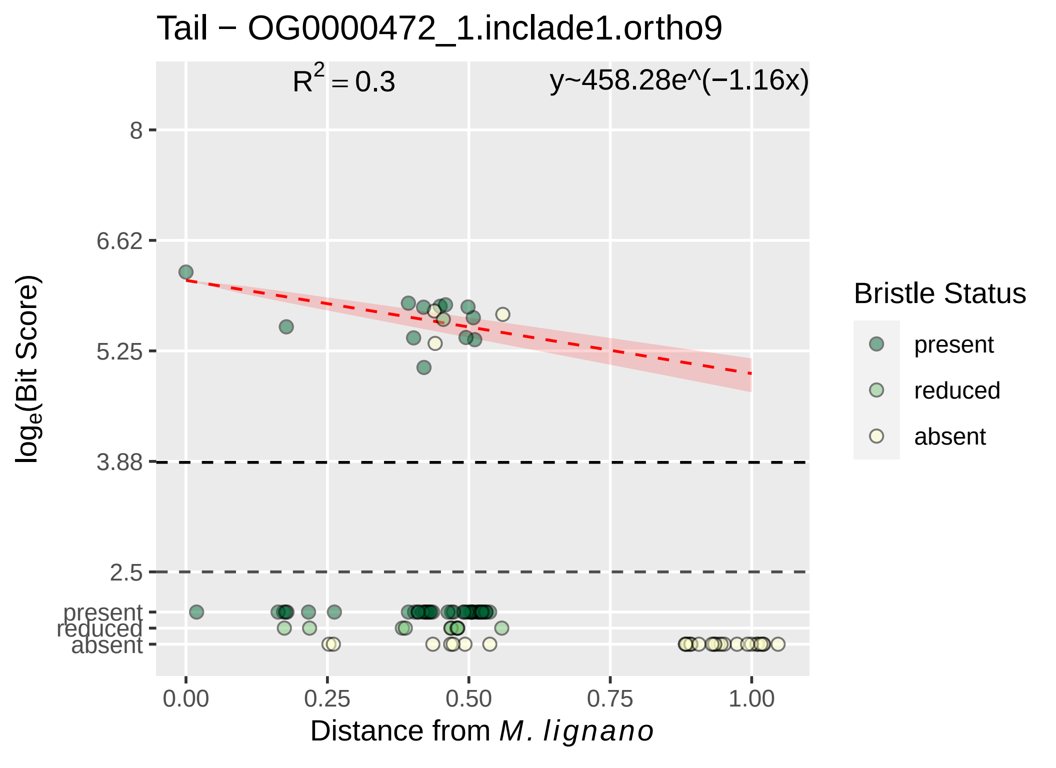

Tail − OG0000472_1.inclade1.ortho9
R^2 = 0.3
y~458.28e^(−1.16x)
8
6.62
5.25
3.88
2.5
present
reduced
absent
log_e(Bit Score)
0.00
0.25
0.50
0.75
1.00
Distance from M. lignano
Bristle Status
present
reduced
absent

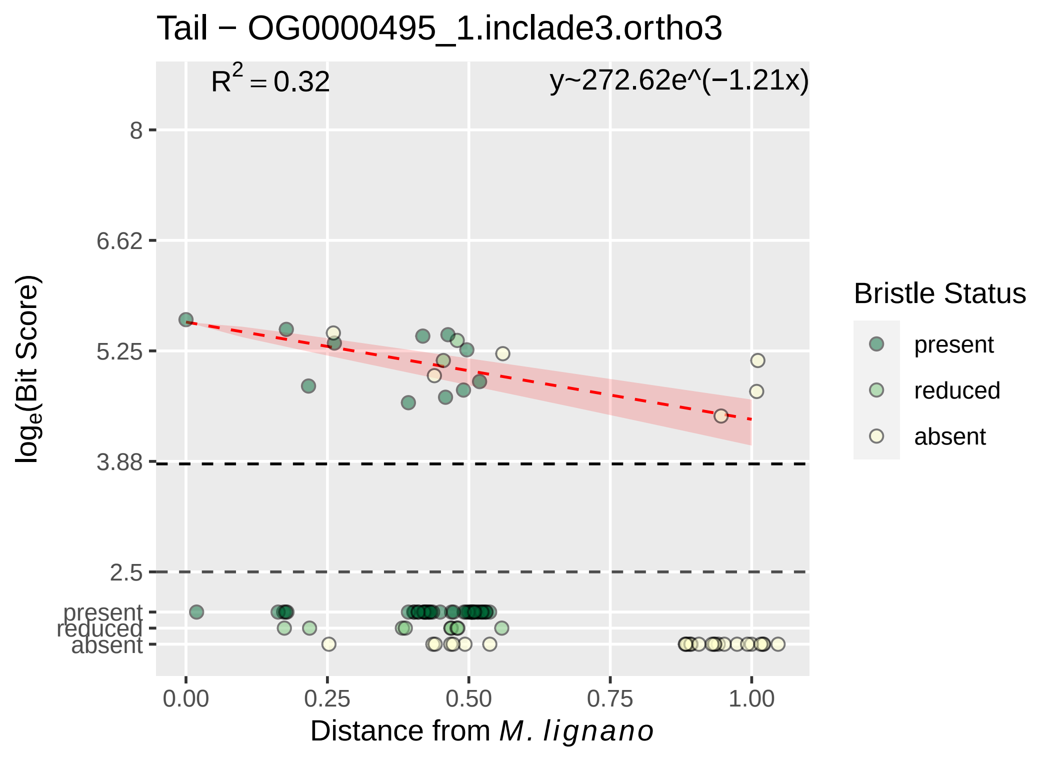

Tail − OG0000495_1.inclade3.ortho3
$R^2 = 0.32$
y~272.62e^(−1.21x)
8
6.62
5.25
3.88
2.5
present
reduced
absent
$\log_e$(Bit Score)
0.00
0.25
0.50
0.75
1.00
Distance from *M. lignano*
Bristle Status
present
reduced
absent

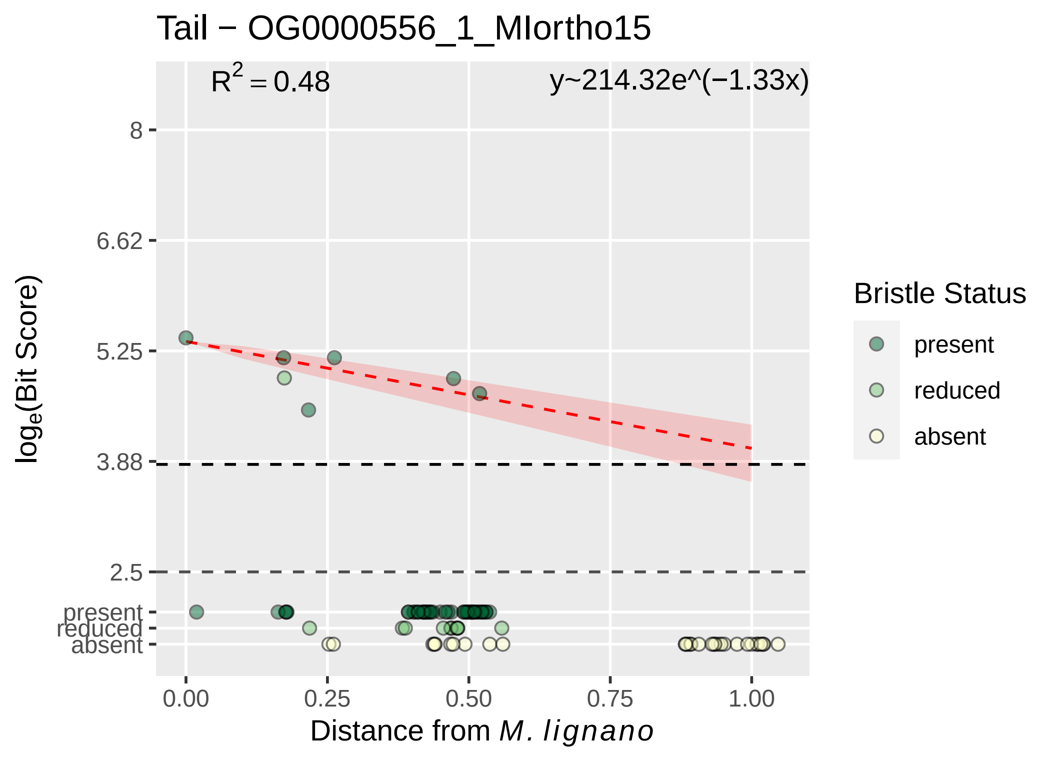

Tail − OG0000556_1_MIortho15
R^2 = 0.48
y~214.32e^(−1.33x)
8
6.62
5.25
3.88
2.5
present
reduced
absent
0.00
0.25
0.50
0.75
1.00
Distance from M. lignano
loge(Bit Score)
Bristle Status
present
reduced
absent

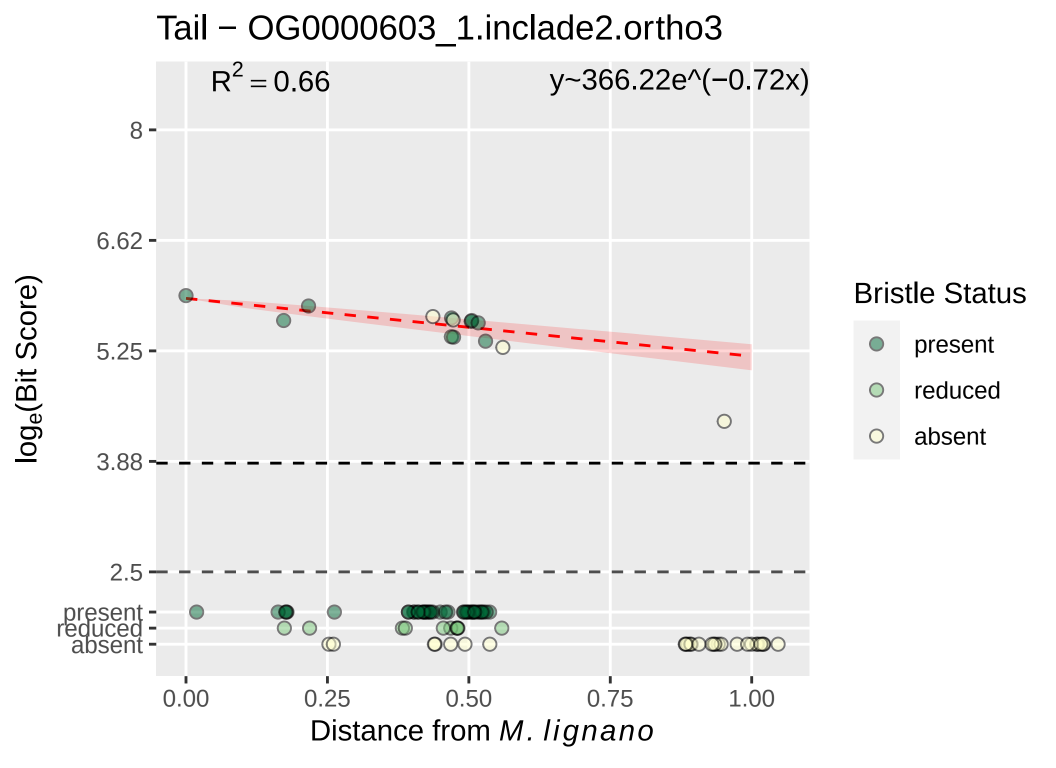

Tail − OG0000603_1.inclade2.ortho3
R2 = 0.66
y~366.22e^(−0.72x)
8
6.62
5.25
3.88
2.5
present
reduced
absent
log_e(Bit Score)
0.00
0.25
0.50
0.75
1.00
Distance from M. lignano
Bristle Status
present
reduced
absent

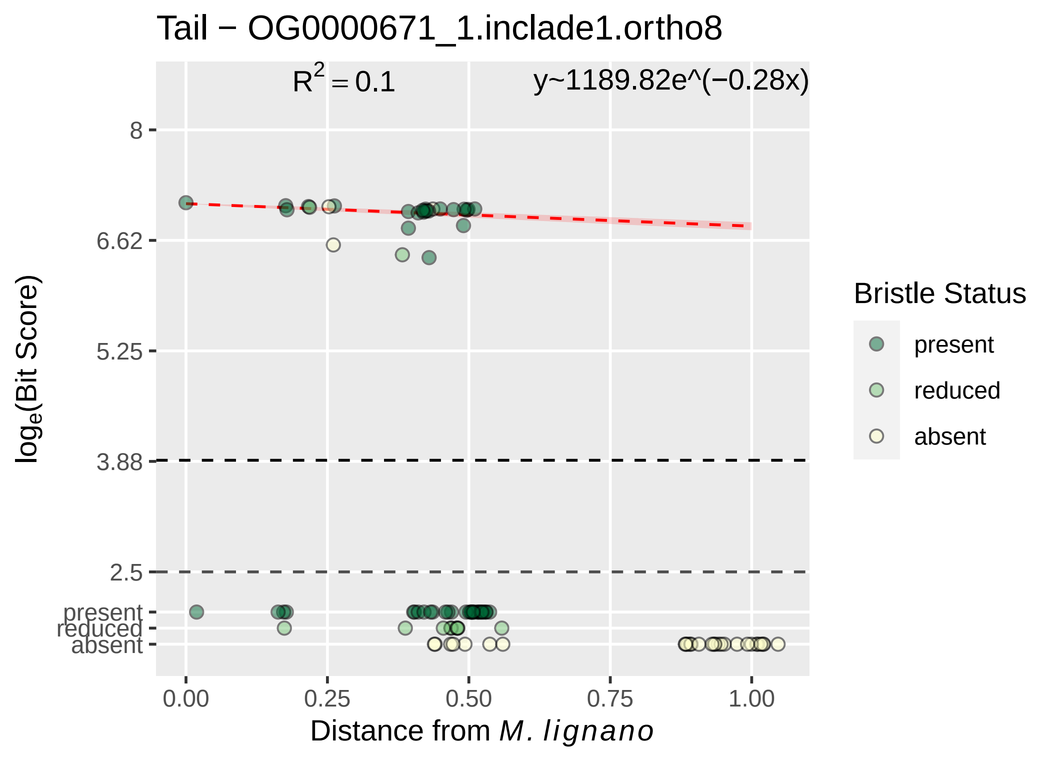

Tail − OG0000671_1.inclade1.ortho8
R^2 = 0.1
y~1189.82e^(−0.28x)
8
6.62
5.25
3.88
2.5
present
reduced
absent
0.00
0.25
0.50
0.75
1.00
Distance from M. lignano
loge(Bit Score)
Bristle Status
present
reduced
absent

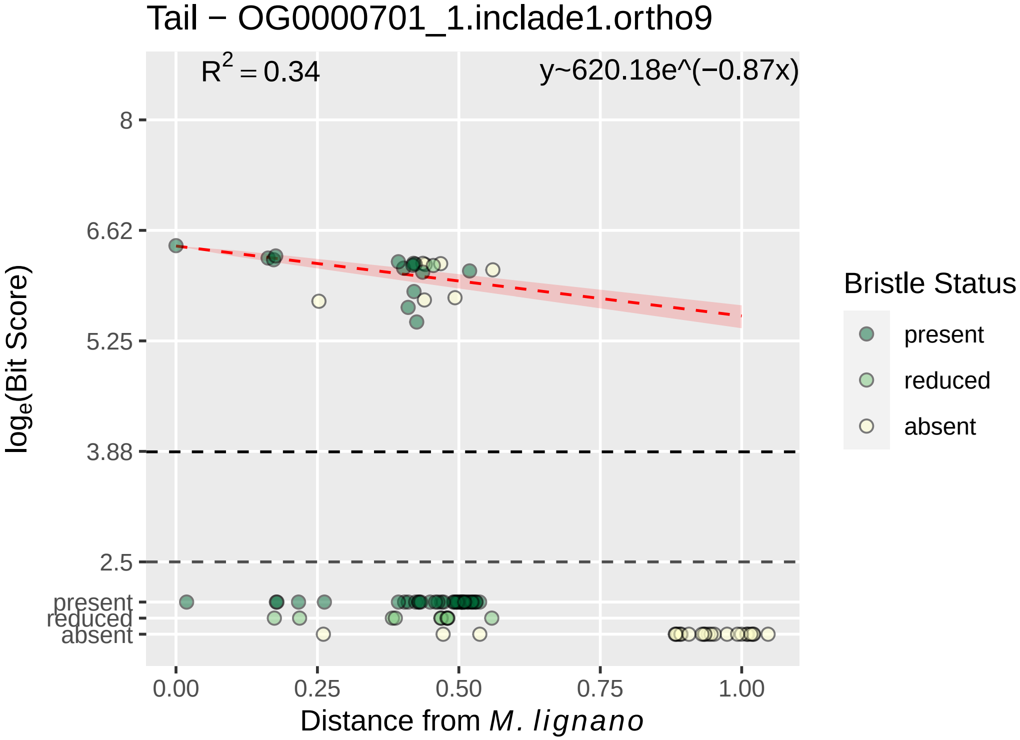

Tail − OG0000701_1.inclade1.ortho9
R^2 = 0.34
y~620.18e^(−0.87x)
8
6.62
5.25
3.88
2.5
present
reduced
absent
0.00
0.25
0.50
0.75
1.00
Distance from M. lignano
log_e(Bit Score)
Bristle Status
present
reduced
absent

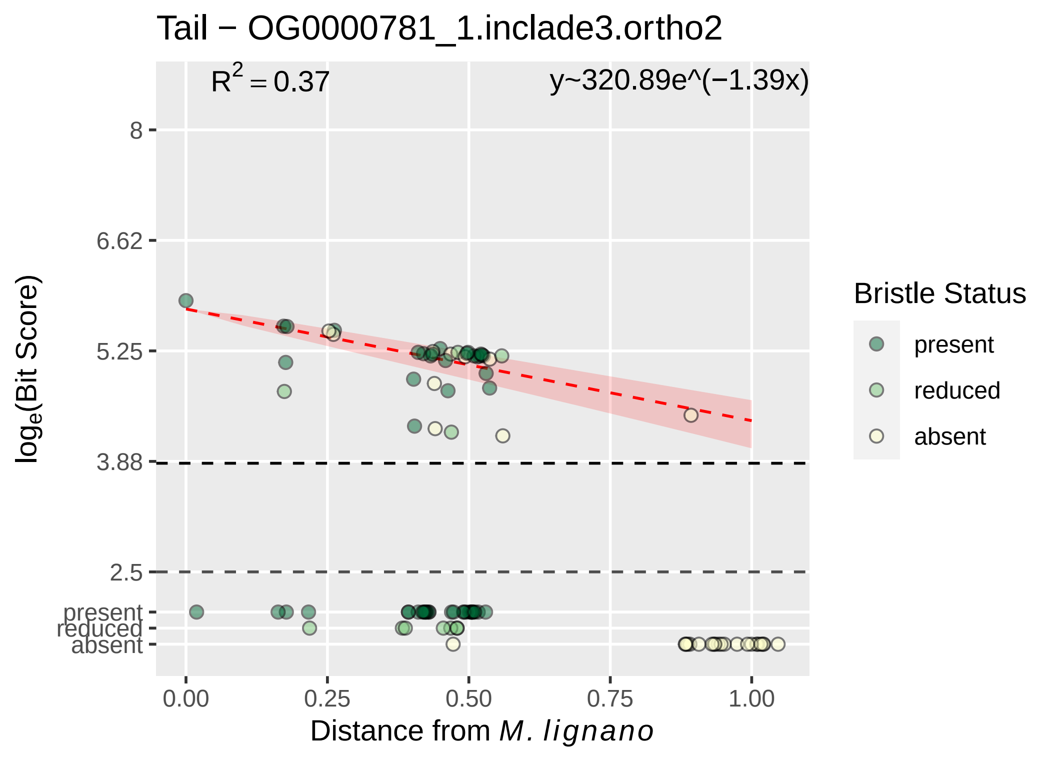

Tail − OG0000781_1.inclade3.ortho2
$R^2 = 0.37$
y~320.89e^(−1.39x)
8
6.62
5.25
3.88
2.5
present
reduced
absent
0.00
0.25
0.50
0.75
1.00
log$_e$(Bit Score)
Distance from *M. lignano*
Bristle Status
present
reduced
absent

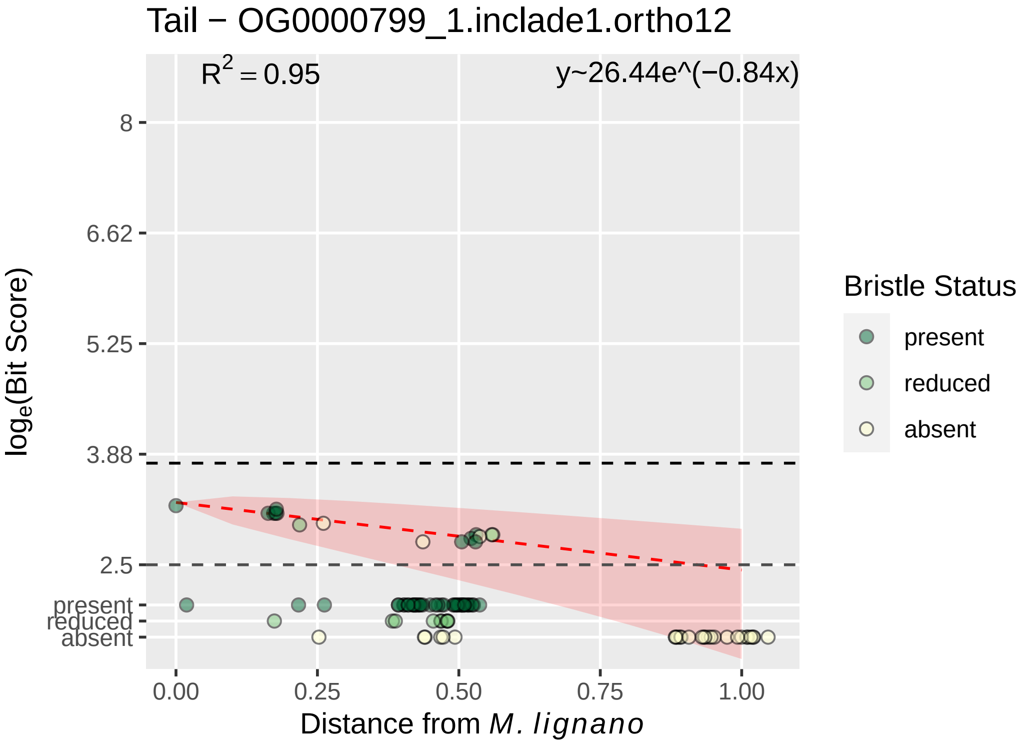
Tail − OG0000799_1.inclade1.ortho12
R2 = 0.95
y~26.44e^(−0.84x)
8
6.62
5.25
3.88
2.5
present
reduced
absent
loge(Bit Score)
0.00
0.25
0.50
0.75
1.00
Distance from M. lignano
Bristle Status
present
reduced
absent

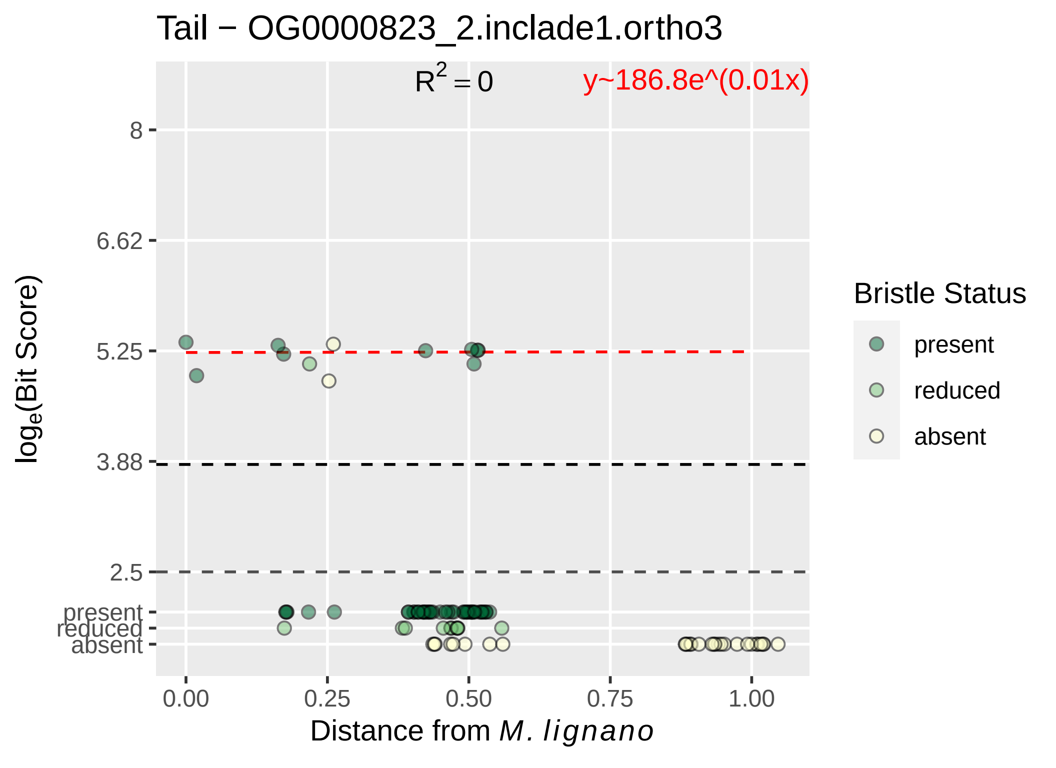

Tail − OG0000823_2.inclade1.ortho3
R^2 = 0
y~186.8e^(0.01x)
log_e(Bit Score)
8
6.62
5.25
3.88
2.5
present
reduced
absent
0.00
0.25
0.50
0.75
1.00
Distance from M. lignano
Bristle Status
present
reduced
absent

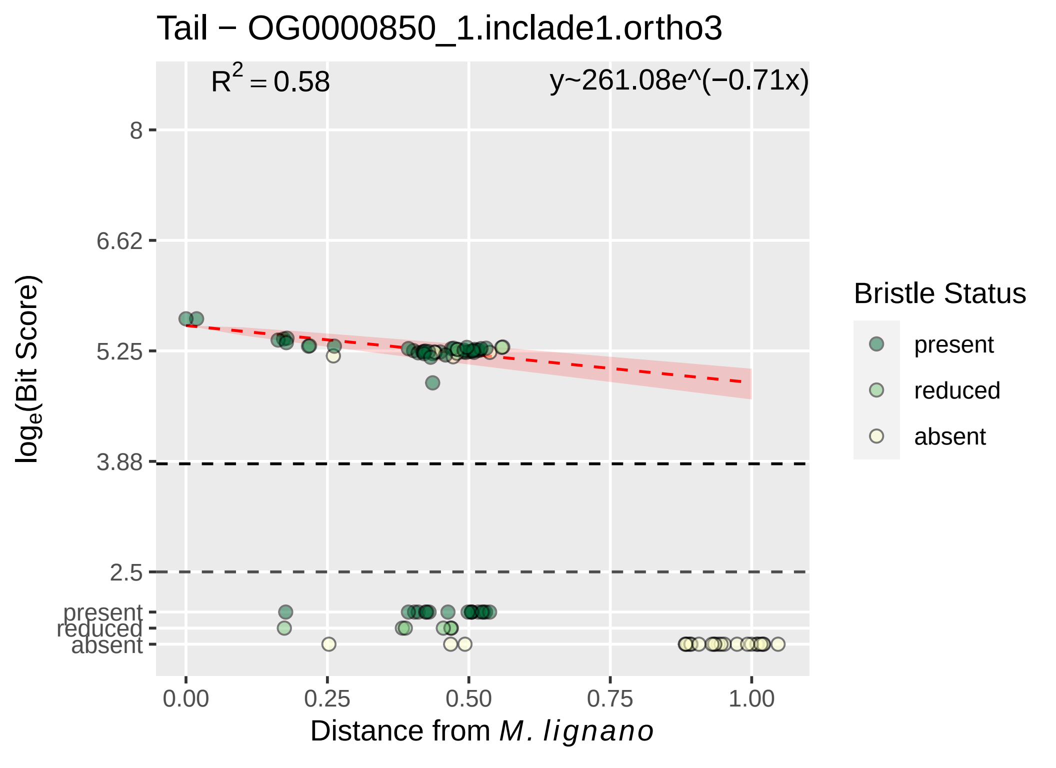
Tail − OG0000850_1.inclade1.ortho3
$R^2 = 0.58$
y~261.08e^(−0.71x)
8
6.62
5.25
3.88
2.5
present
reduced
absent
$\log_e$(Bit Score)
0.00
0.25
0.50
0.75
1.00
Distance from *M. lignano*
Bristle Status
present
reduced
absent

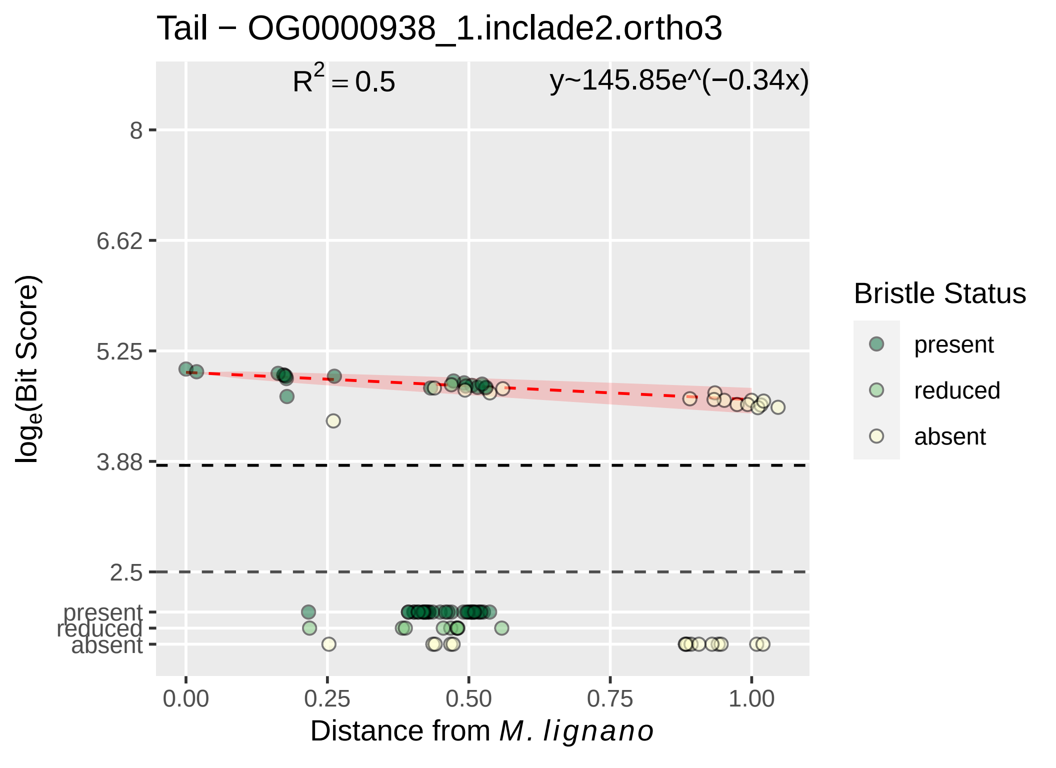

Tail − OG0000938_1.inclade2.ortho3
R² = 0.5
y~145.85e^(−0.34x)
8
6.62
5.25
3.88
2.5
present
reduced
absent
log_e(Bit Score)
0.00
0.25
0.50
0.75
1.00
Distance from M. lignano
Bristle Status
present
reduced
absent

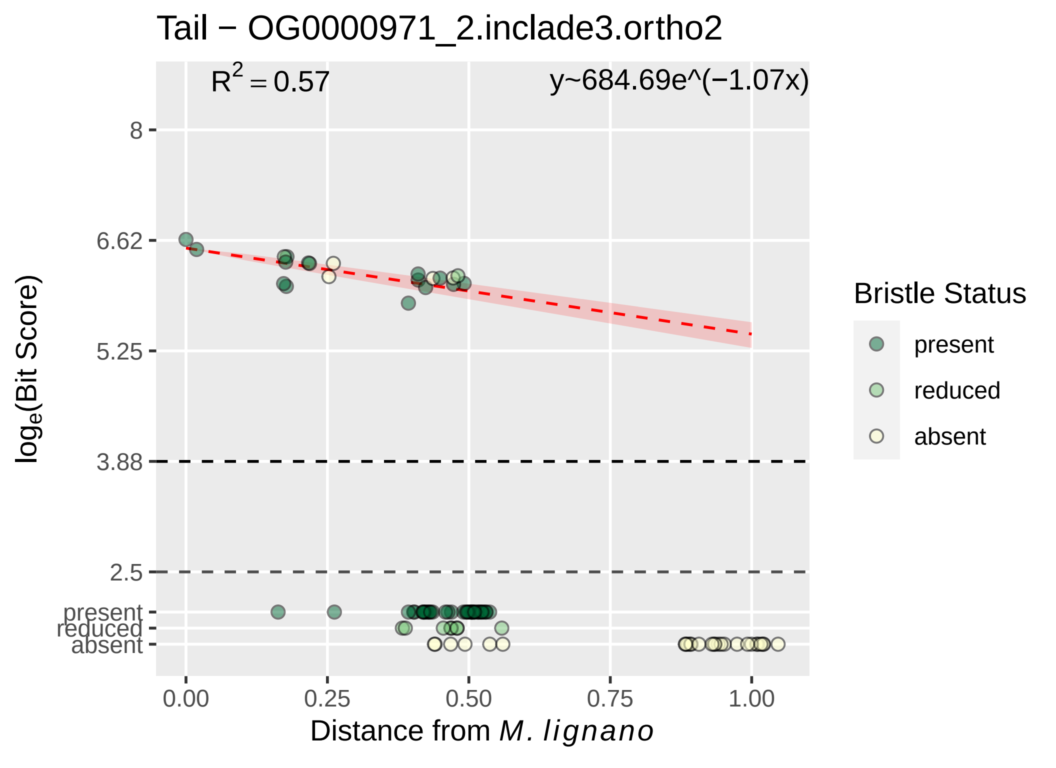

Tail − OG0000971_2.inclade3.ortho2
R^2 = 0.57
y~684.69e^(−1.07x)
8
6.62
5.25
3.88
2.5
present
reduced
absent
log_e(Bit Score)
0.00
0.25
0.50
0.75
1.00
Distance from M. lignano
Bristle Status
present
reduced
absent

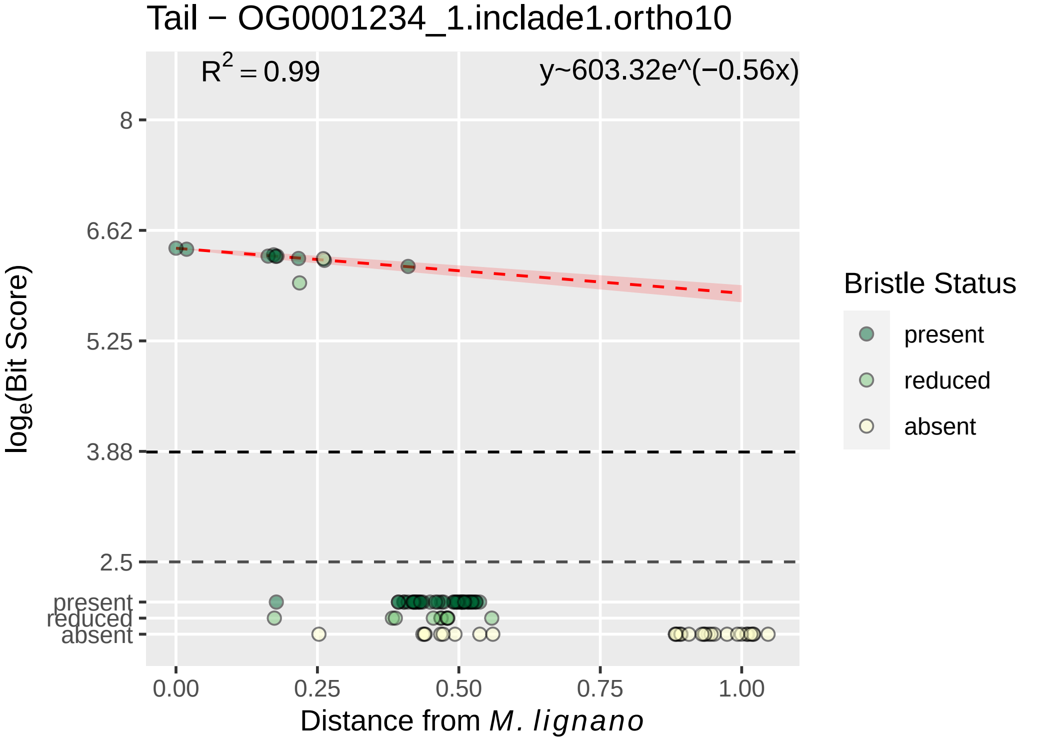

Tail − OG0001234_1.inclade1.ortho10
$R^2 = 0.99$
y~603.32e^(−0.56x)
8
6.62
5.25
3.88
2.5
present
reduced
absent
$\log_e$(Bit Score)
0.00
0.25
0.50
0.75
1.00
Distance from *M. lignano*
Bristle Status
present
reduced
absent

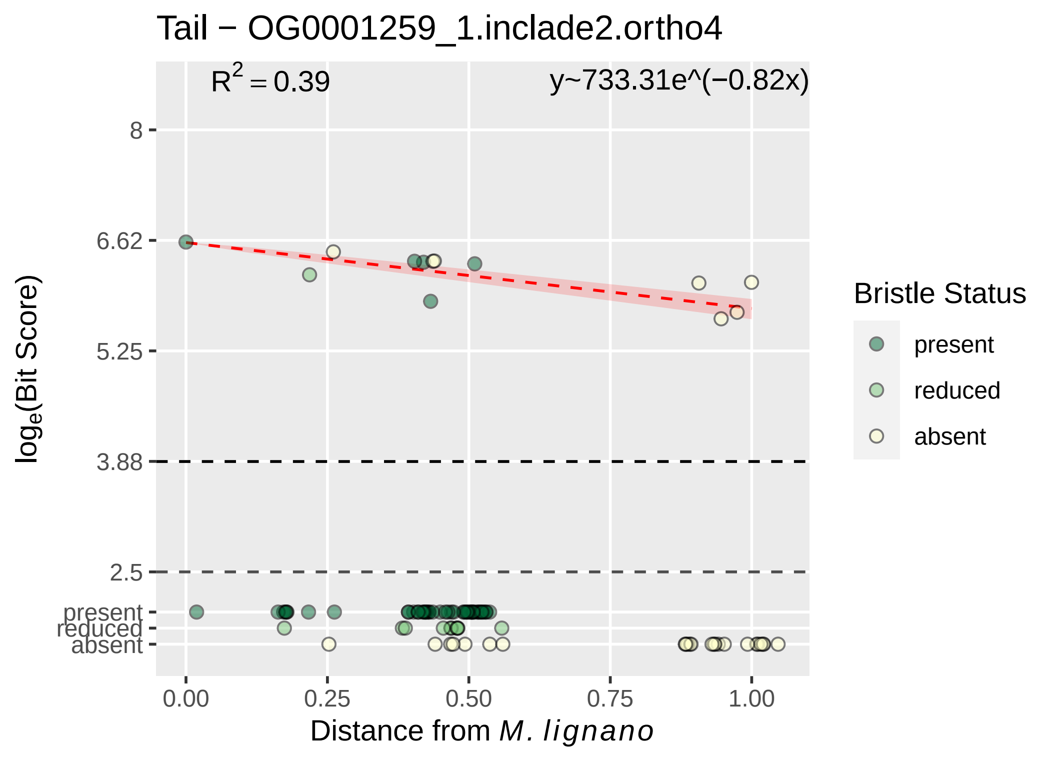

Tail − OG0001259_1.inclade2.ortho4
R2 = 0.39
y~733.31e^(−0.82x)
8
6.62
5.25
3.88
2.5
present
reduced
absent
loge(Bit Score)
0.00
0.25
0.50
0.75
1.00
Distance from M. lignano
Bristle Status
present
reduced
absent

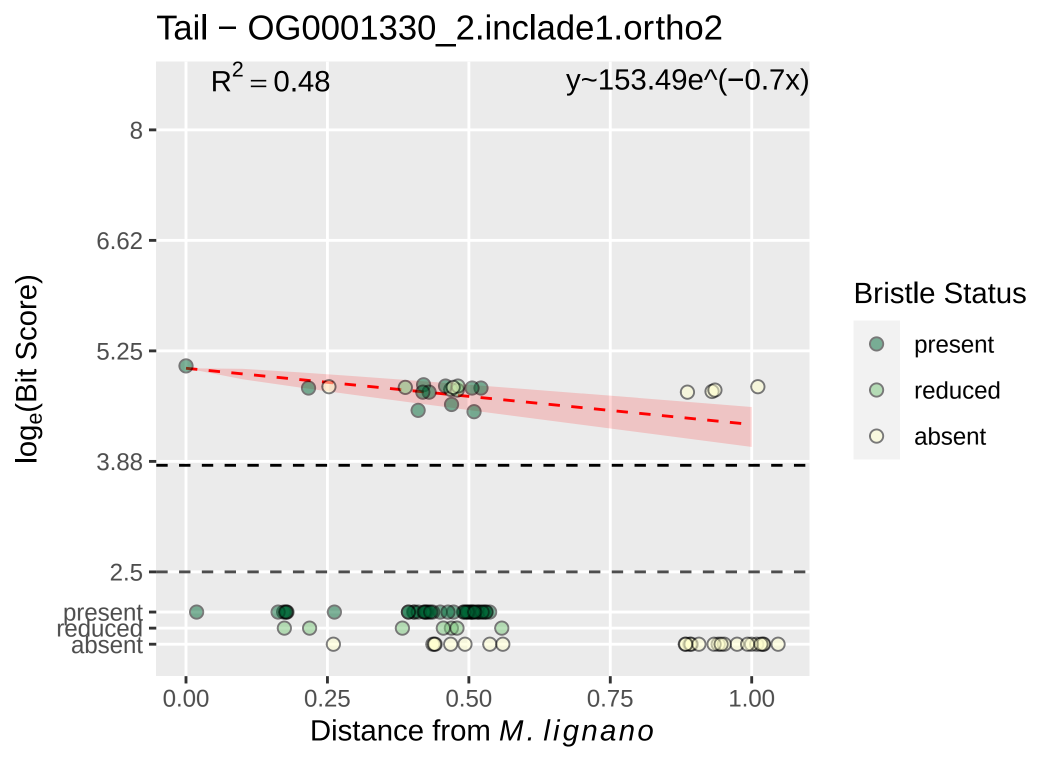

Tail − OG0001330_2.inclade1.ortho2
R^2 = 0.48
y~153.49e^(−0.7x)
8
6.62
5.25
3.88
2.5
present
reduced
absent
log_e(Bit Score)
0.00
0.25
0.50
0.75
1.00
Distance from M. lignano
Bristle Status
present
reduced
absent

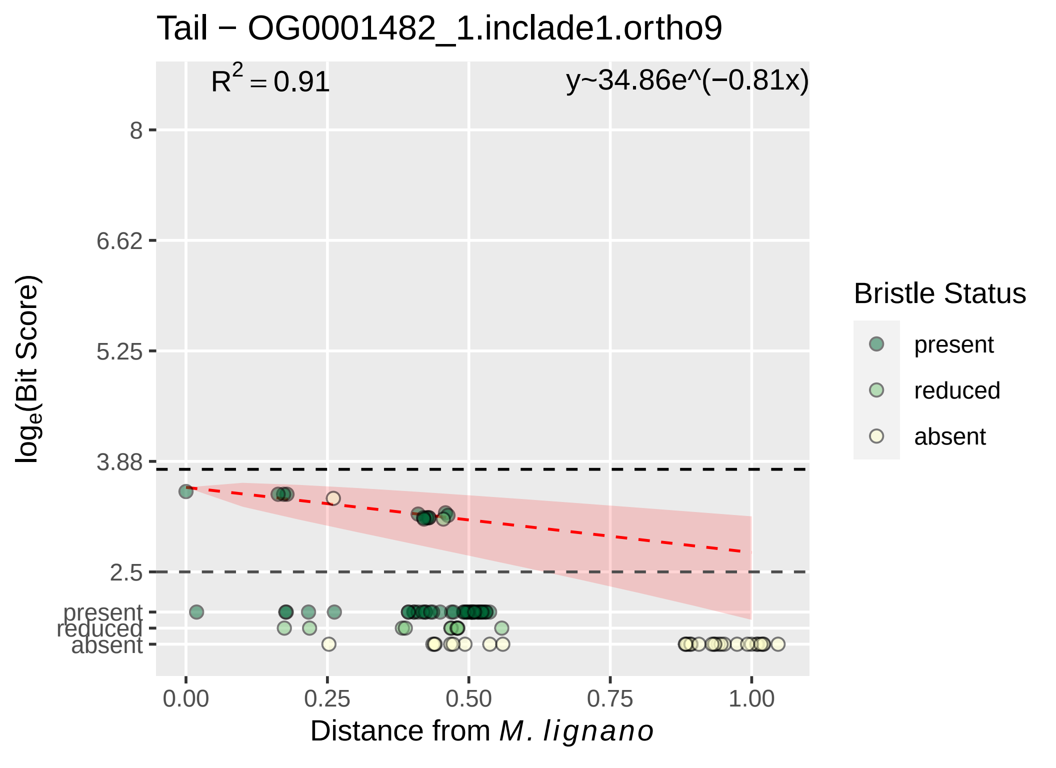
Tail − OG0001482_1.inclade1.ortho9
$R^2 = 0.91$
y~34.86e^(−0.81x)
8
6.62
5.25
3.88
2.5
present
reduced
absent
0.00
0.25
0.50
0.75
1.00
Distance from M. lignano
log_e(Bit Score)
Bristle Status
present
reduced
absent

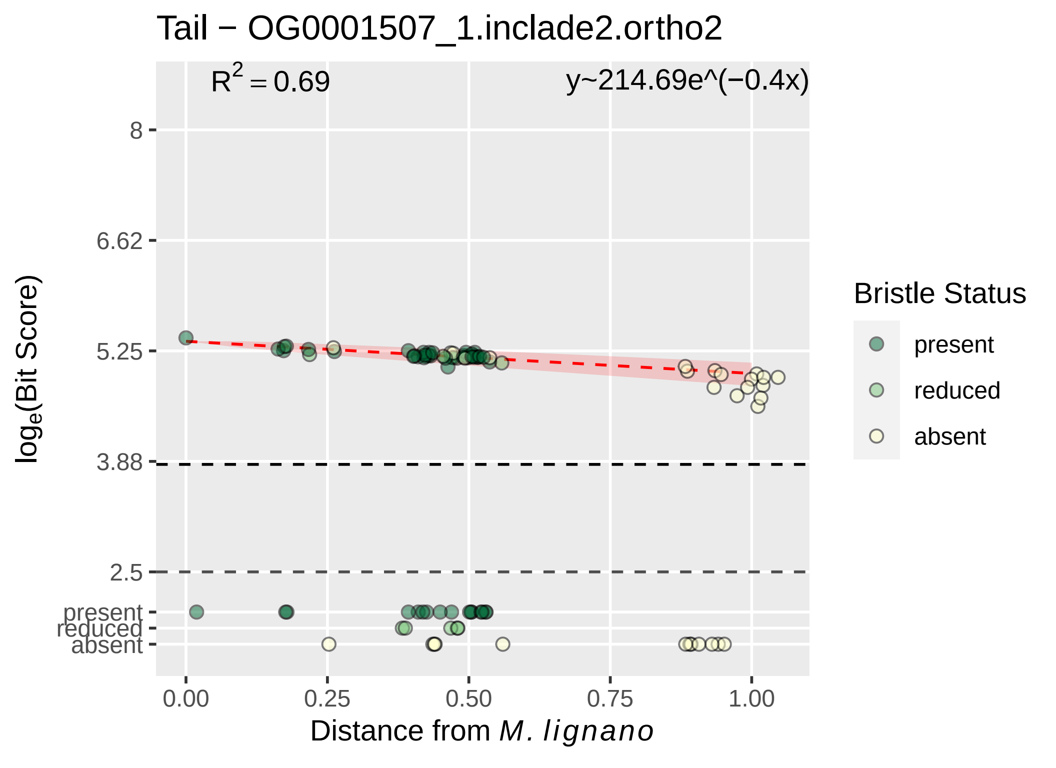

Tail − OG0001507_1.inclade2.ortho2
$R^2 = 0.69$
y~214.69e^(−0.4x)
8
6.62
5.25
3.88
2.5
present
reduced
absent
$\log_e$(Bit Score)
0.00
0.25
0.50
0.75
1.00
Distance from *M. lignano*
Bristle Status
present
reduced
absent

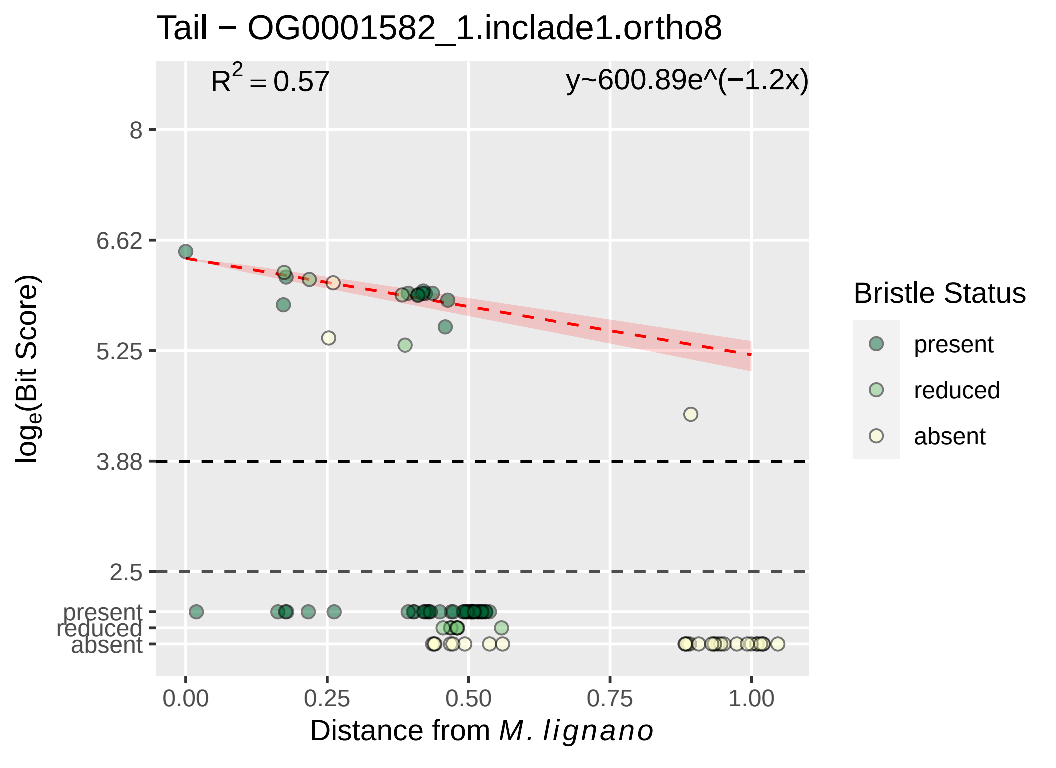

Tail − OG0001582_1.inclade1.ortho8
R^2 = 0.57
y~600.89e^(−1.2x)
8
6.62
5.25
3.88
2.5
present
reduced
absent
log_e(Bit Score)
0.00
0.25
0.50
0.75
1.00
Distance from M. lignano
Bristle Status
present
reduced
absent

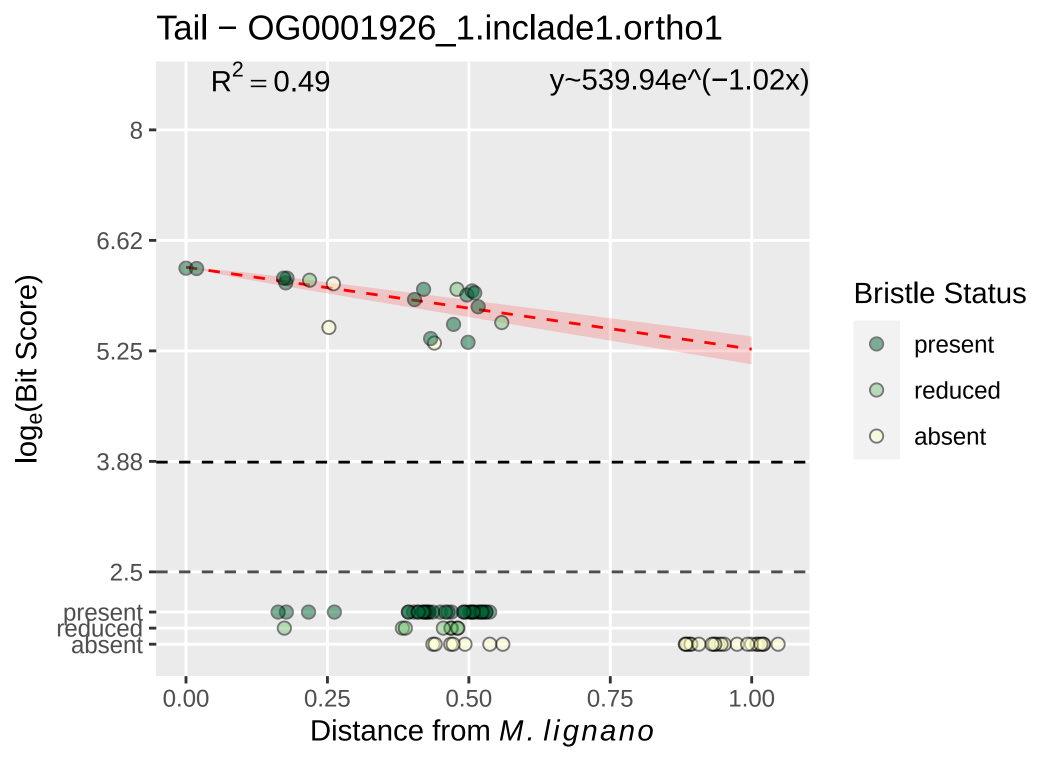
Tail − OG0001926_1.inclade1.ortho1
$R^2 = 0.49$
y~539.94e^(−1.02x)
8
6.62
5.25
3.88
2.5
present
reduced
absent
log$_e$(Bit Score)
0.00
0.25
0.50
0.75
1.00
Distance from M. lignano
Bristle Status
present
reduced
absent

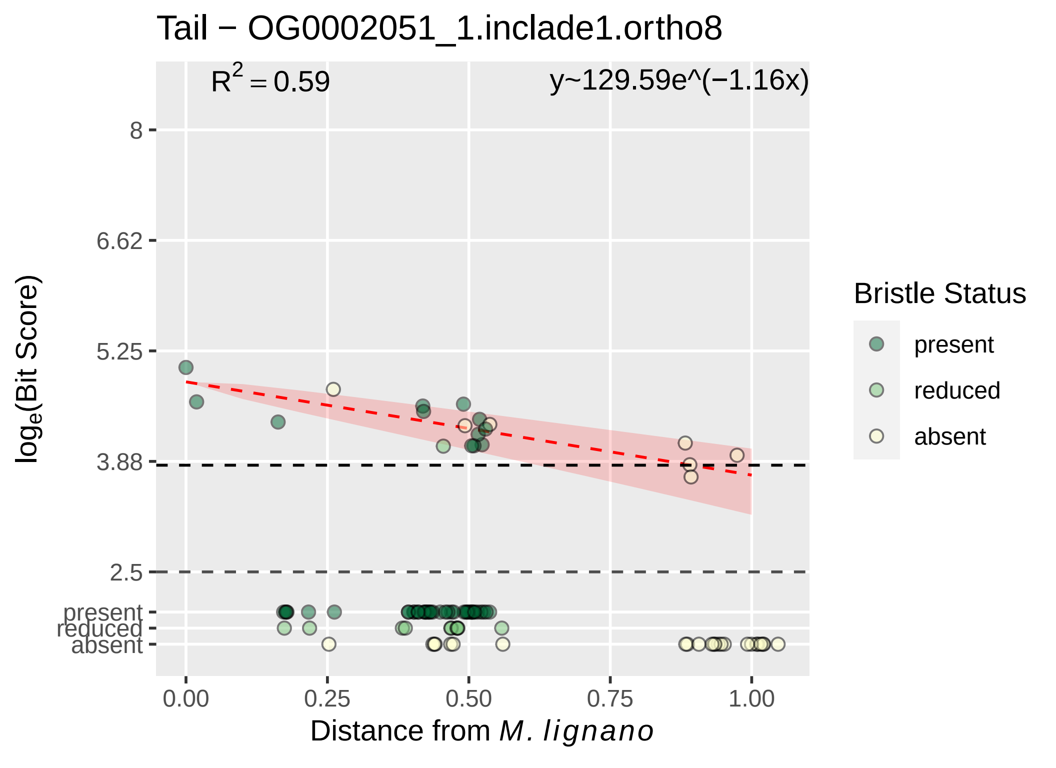
Tail − OG0002051_1.inclade1.ortho8
$R^2 = 0.59$
y~129.59e^(−1.16x)
8
6.62
5.25
3.88
2.5
present
reduced
absent
$\log_e$(Bit Score)
0.00
0.25
0.50
0.75
1.00
Distance from *M. lignano*
Bristle Status
present
reduced
absent

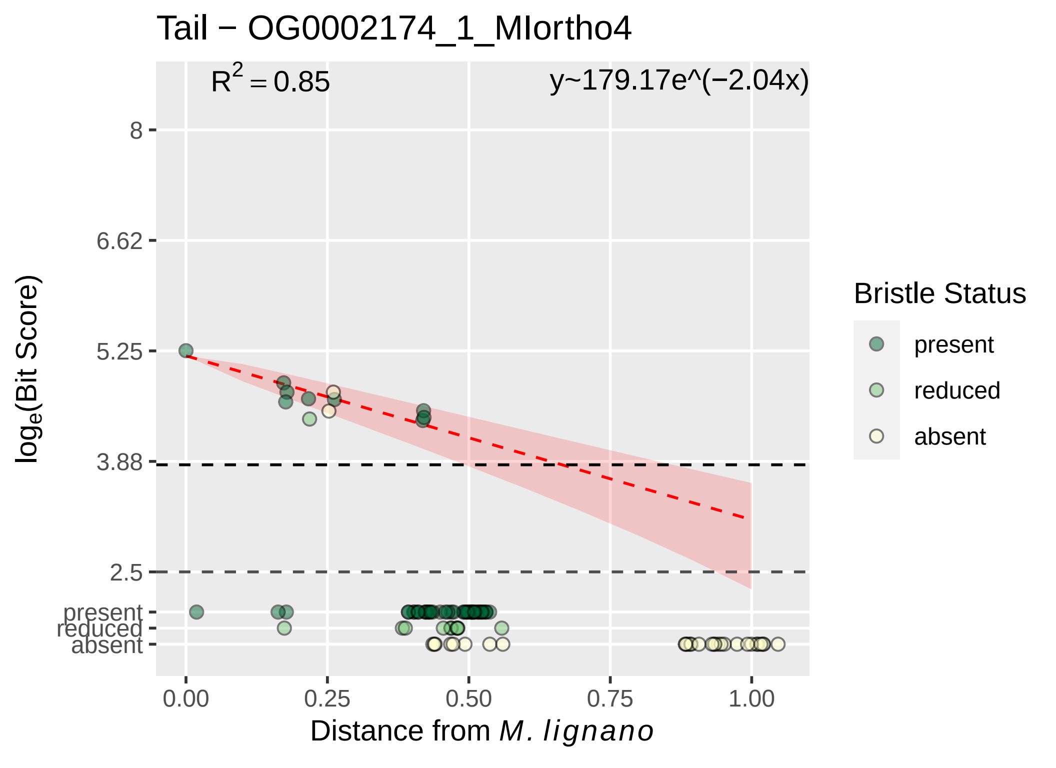
Tail − OG0002174_1_MIortho4
R2 = 0.85
y~179.17e^(−2.04x)
8
6.62
5.25
3.88
2.5
present
reduced
absent
loge(Bit Score)
0.00
0.25
0.50
0.75
1.00
Distance from M. lignano
Bristle Status
present
reduced
absent

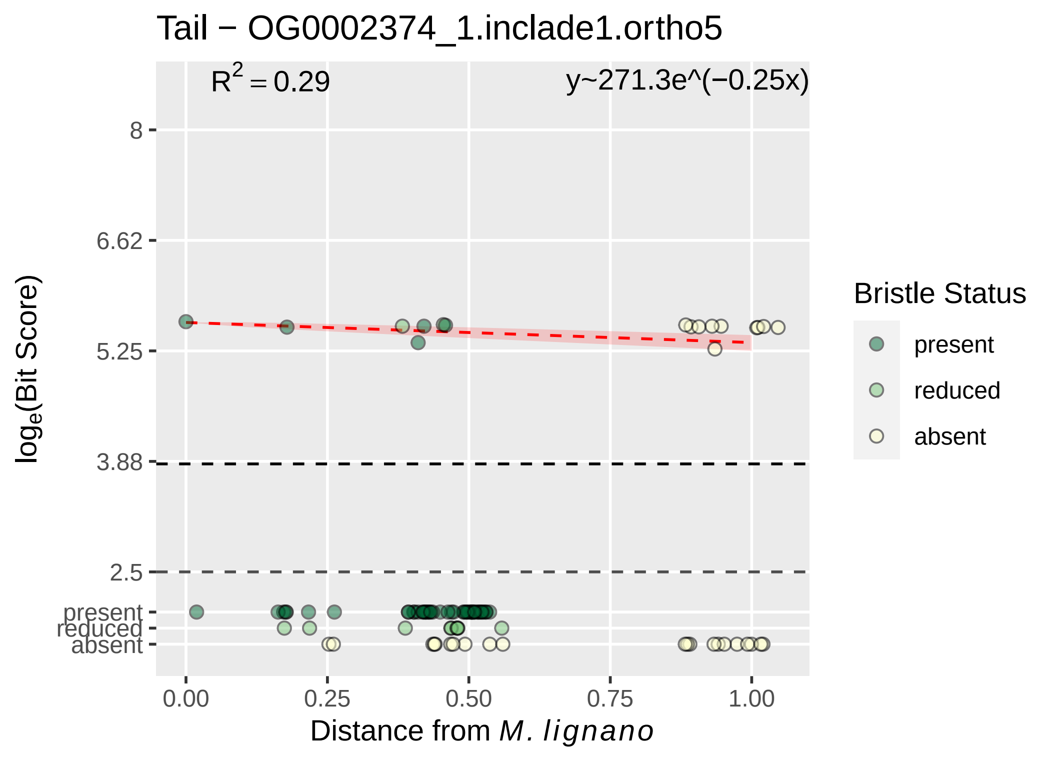

Tail − OG0002374_1.inclade1.ortho5
R2 = 0.29
y~271.3e^(−0.25x)
8
6.62
5.25
3.88
2.5
present
reduced
absent
loge(Bit Score)
0.00
0.25
0.50
0.75
1.00
Distance from M. lignano
Bristle Status
present
reduced
absent

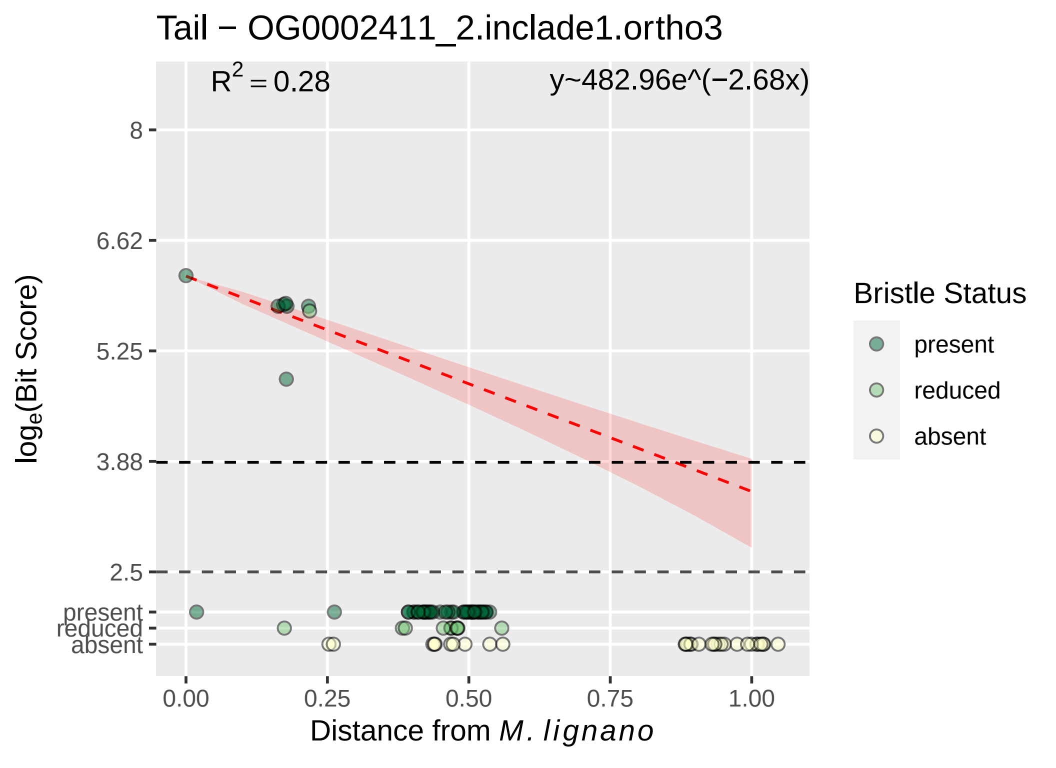

Tail − OG0002411_2.inclade1.ortho3
$R^2 = 0.28$
y~482.96e^(−2.68x)
8
6.62
5.25
3.88
2.5
present
reduced
absent
$\log_e$(Bit Score)
0.00
0.25
0.50
0.75
1.00
Distance from *M. lignano*
Bristle Status
present
reduced
absent

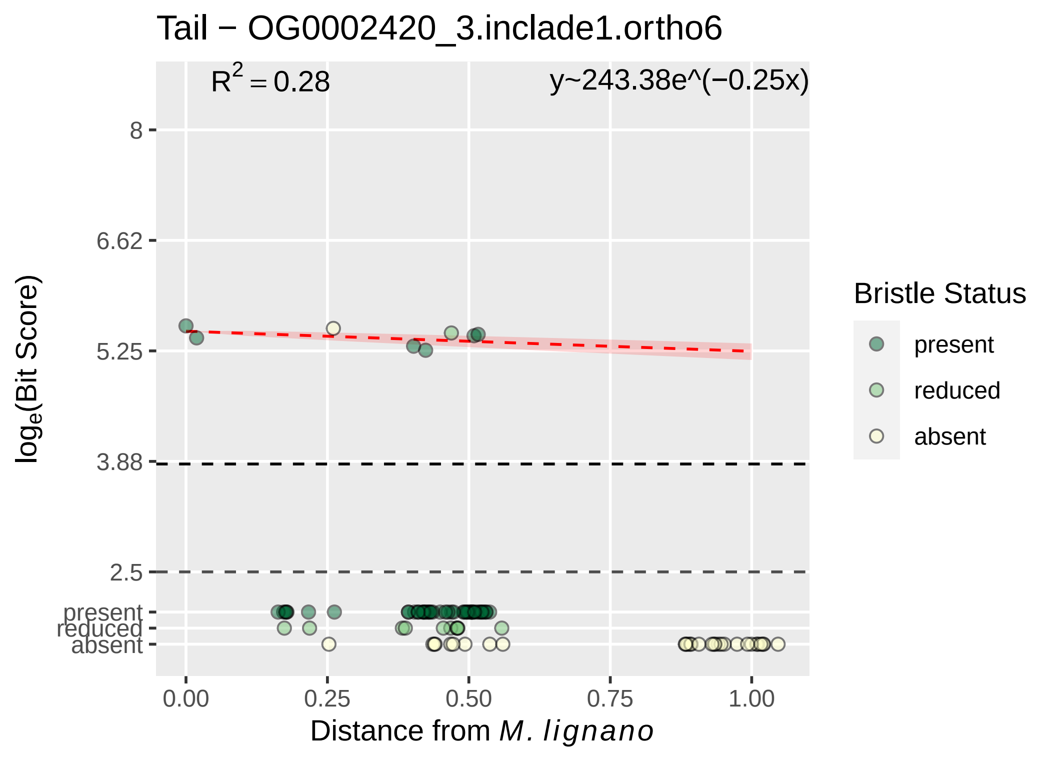

Tail − OG0002420_3.inclade1.ortho6
R^2 = 0.28
y~243.38e^(−0.25x)
8
6.62
5.25
3.88
2.5
present
reduced
absent
log_e(Bit Score)
0.00
0.25
0.50
0.75
1.00
Distance from M. lignano
Bristle Status
present
reduced
absent

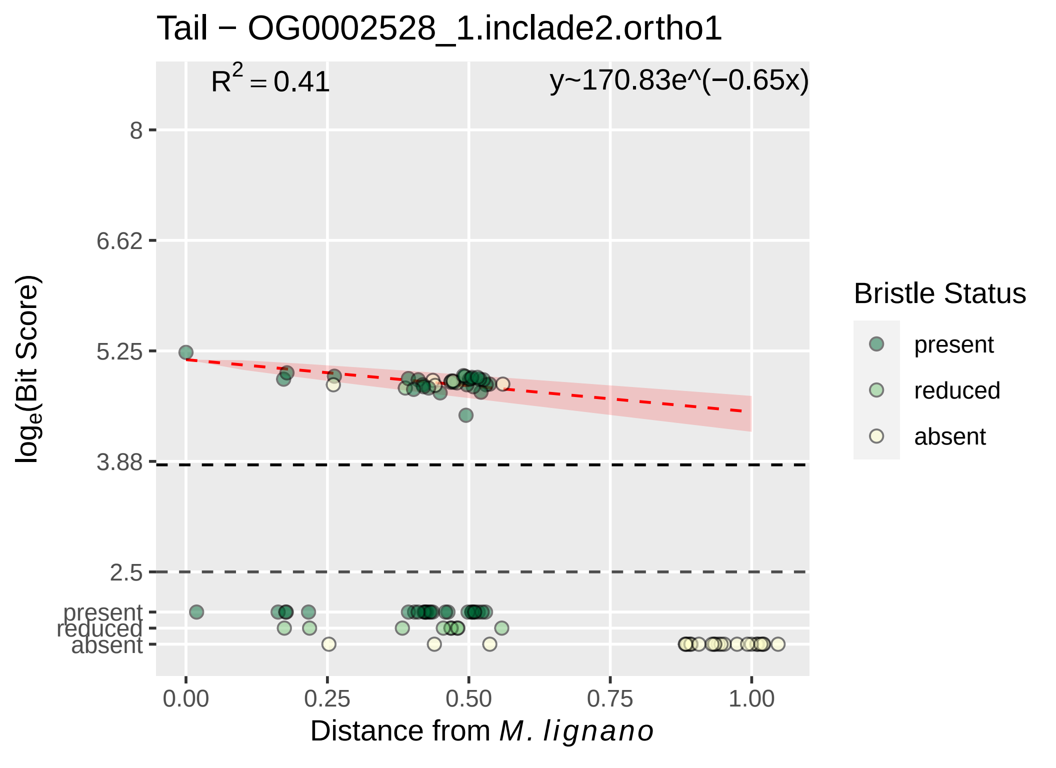

Tail − OG0002528_1.inclade2.ortho1
R^2 = 0.41
y~170.83e^(−0.65x)
log_e(Bit Score)
8
6.62
5.25
3.88
2.5
present
reduced
absent
0.00
0.25
0.50
0.75
1.00
Distance from M. lignano
Bristle Status
present
reduced
absent

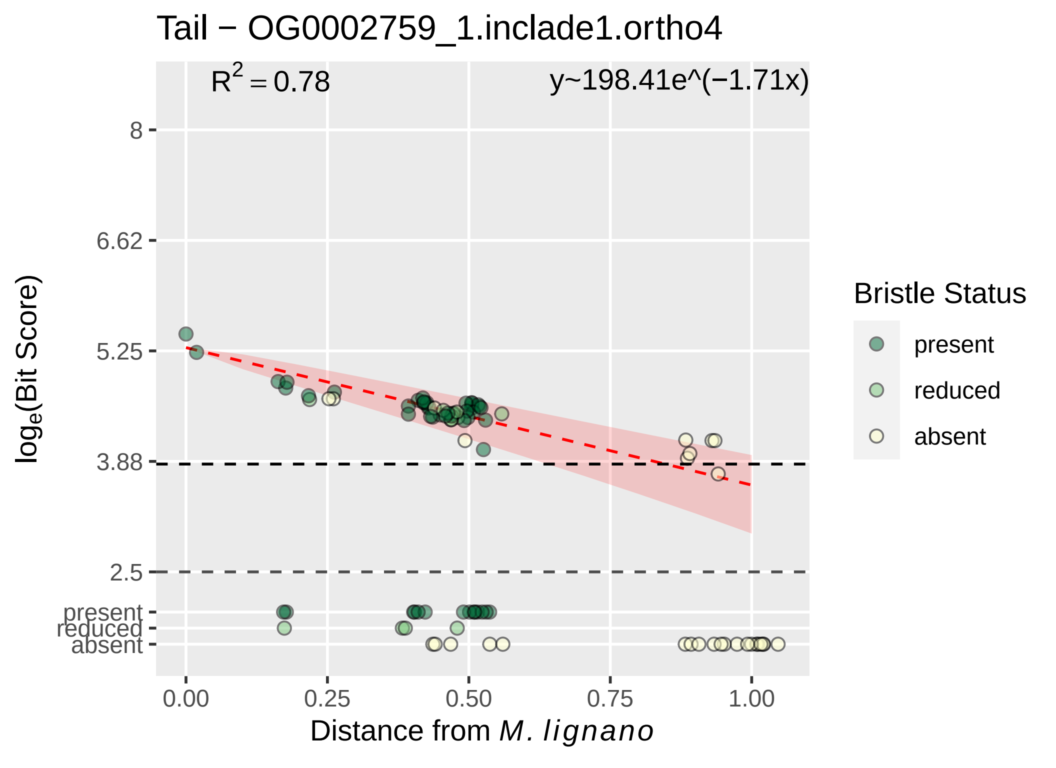

Tail − OG0002759_1.inclade1.ortho4
R^2 = 0.78
y~198.41e^(−1.71x)
8
6.62
5.25
3.88
2.5
present
reduced
absent
0.00
0.25
0.50
0.75
1.00
Distance from M. lignano
log_e(Bit Score)
Bristle Status
present
reduced
absent

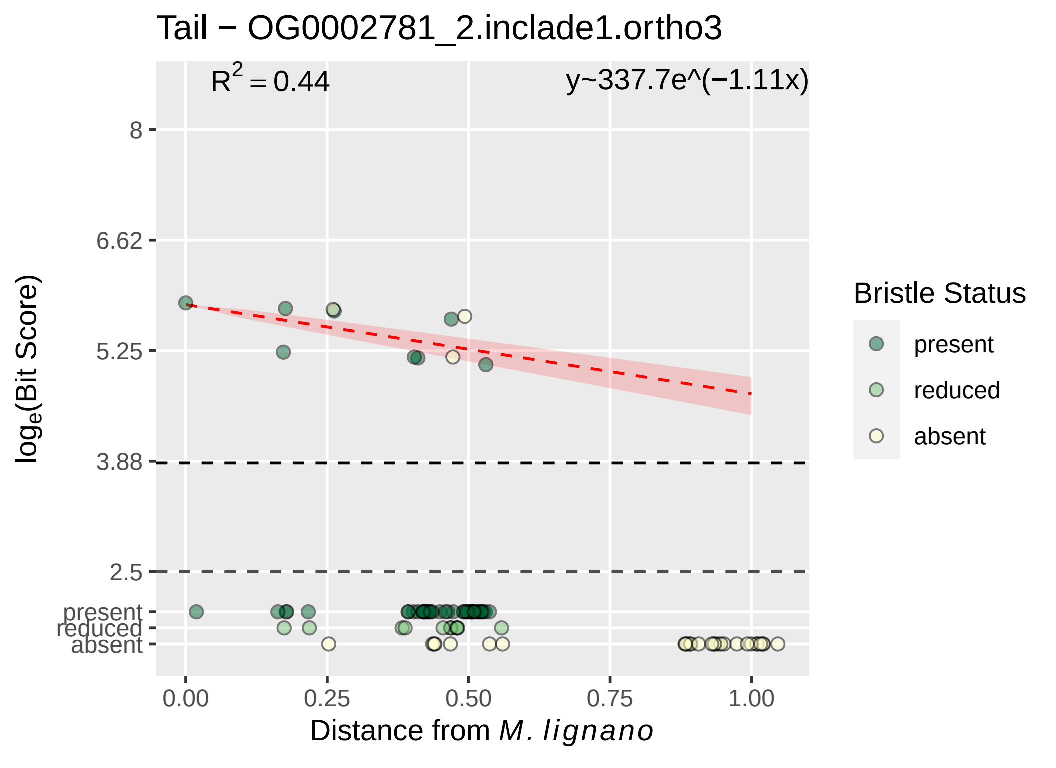

Tail − OG0002781_2.inclade1.ortho3
$R^2 = 0.44$
y~337.7e^(−1.11x)
8
6.62
5.25
3.88
2.5
present
reduced
absent
$log_e$(Bit Score)
0.00
0.25
0.50
0.75
1.00
Distance from *M. lignano*
Bristle Status
present
reduced
absent

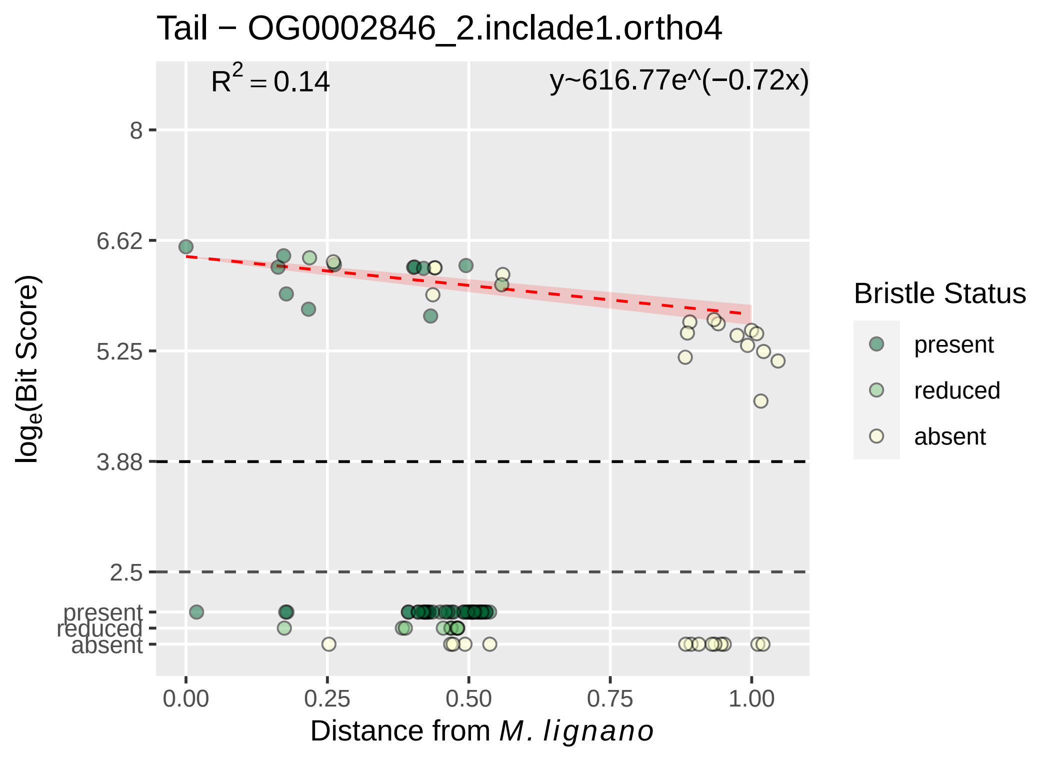

Tail − OG0002846_2.inclade1.ortho4
R2 = 0.14
y~616.77e^(−0.72x)
8
6.62
5.25
3.88
2.5
present
reduced
absent
log_e(Bit Score)
0.00
0.25
0.50
0.75
1.00
Distance from M. lignano
Bristle Status
present
reduced
absent

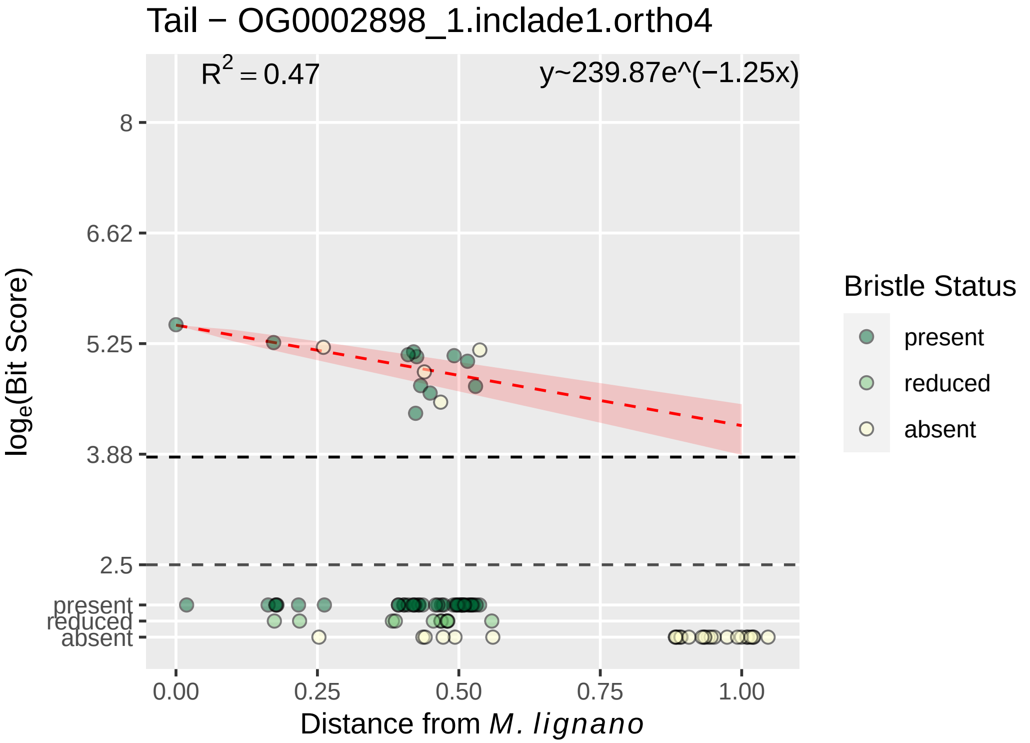

Tail − OG0002898_1.inclade1.ortho4
$R^2 = 0.47$
y~239.87e^(−1.25x)
8
6.62
5.25
3.88
2.5
present
reduced
absent
$\log_e$(Bit Score)
0.00
0.25
0.50
0.75
1.00
Distance from *M. lignano*
Bristle Status
present
reduced
absent

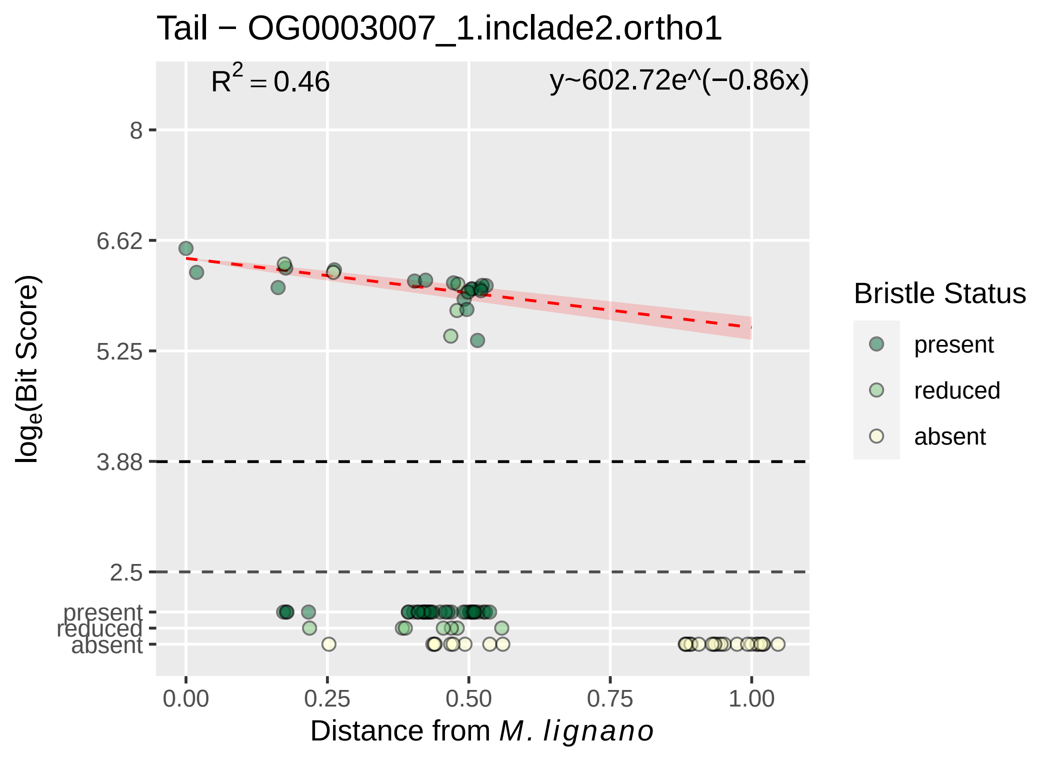

Tail − OG0003007_1.inclade2.ortho1
$R^2 = 0.46$
y~602.72e^(−0.86x)
8
6.62
5.25
3.88
2.5
present
reduced
absent
log$_e$(Bit Score)
0.00
0.25
0.50
0.75
1.00
Distance from *M. lignano*
Bristle Status
present
reduced
absent

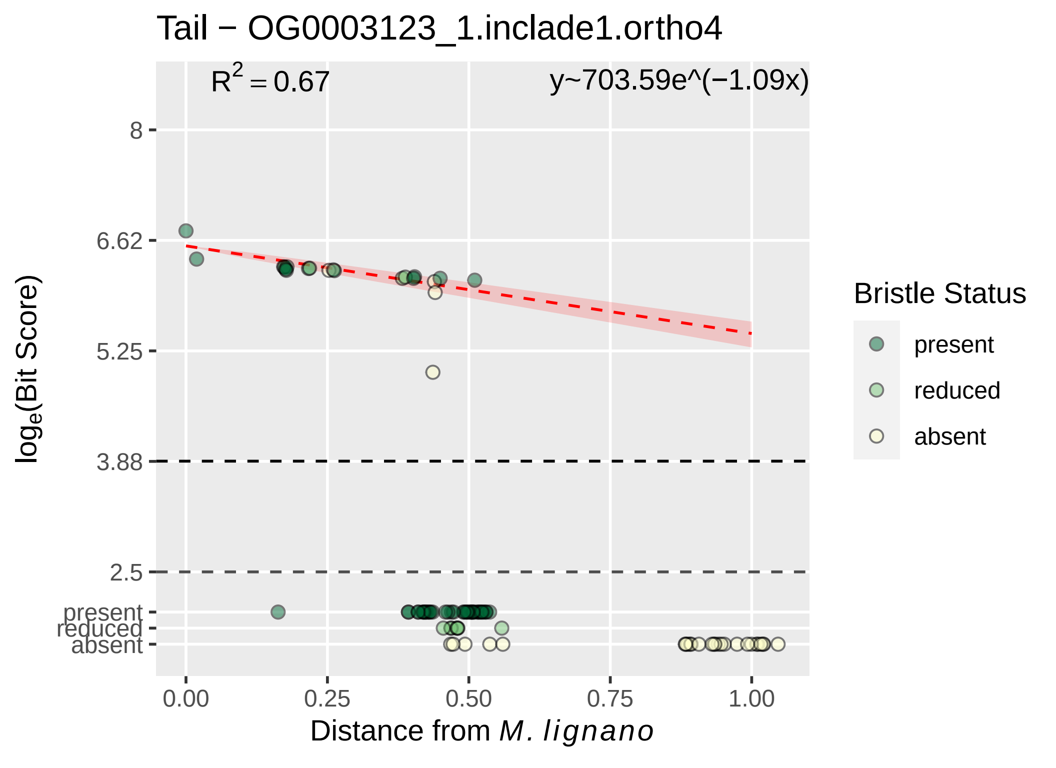

Tail − OG0003123_1.inclade1.ortho4
R^2 = 0.67
y~703.59e^(−1.09x)
8
6.62
5.25
3.88
2.5
present
reduced
absent
loge(Bit Score)
0.00
0.25
0.50
0.75
1.00
Distance from M. lignano
Bristle Status
present
reduced
absent

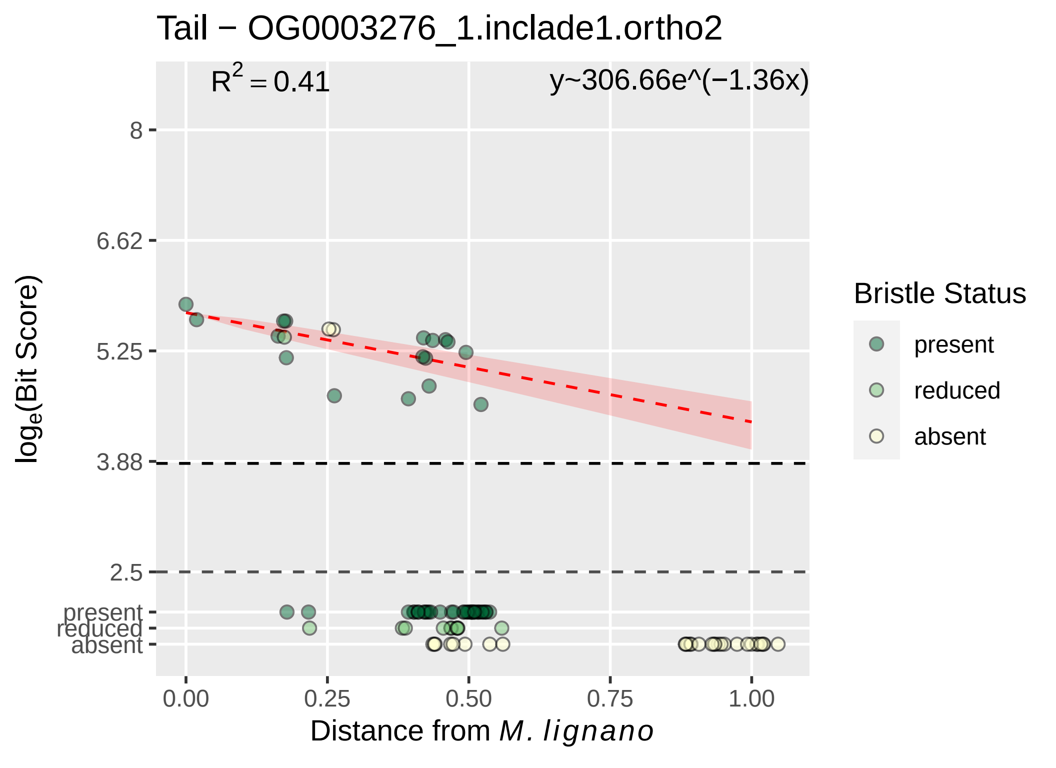

Tail − OG0003276_1.inclade1.ortho2
$R^2 = 0.41$
y~306.66e^(−1.36x)
8
6.62
5.25
3.88
2.5
present
reduced
absent
$\log_e$(Bit Score)
0.00
0.25
0.50
0.75
1.00
Distance from *M. lignano*
Bristle Status
present
reduced
absent

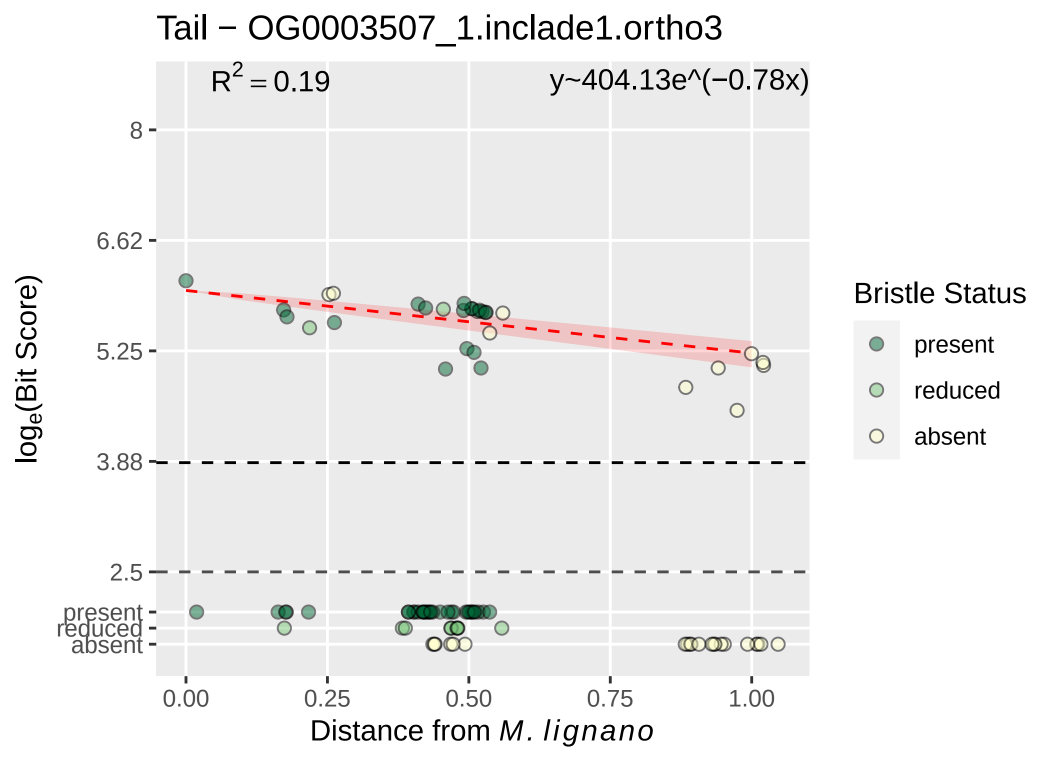

Tail − OG0003507_1.inclade1.ortho3
R2 = 0.19
y~404.13e^(−0.78x)
8
6.62
5.25
3.88
2.5
present
reduced
absent
loge(Bit Score)
0.00
0.25
0.50
0.75
1.00
Distance from M. lignano
Bristle Status
present
reduced
absent

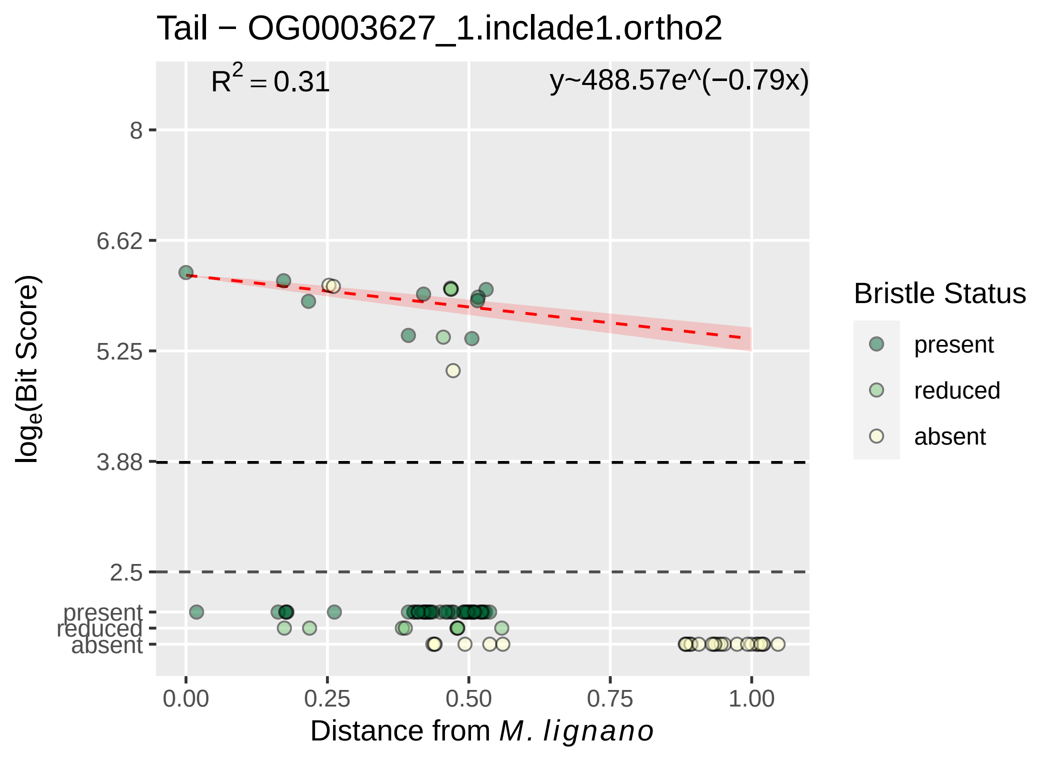

Tail − OG0003627_1.inclade1.ortho2
$R^2 = 0.31$
y~488.57e^(−0.79x)
8
6.62
5.25
3.88
2.5
present
reduced
absent
$log_e$(Bit Score)
0.00
0.25
0.50
0.75
1.00
Distance from *M. lignano*
Bristle Status
present
reduced
absent

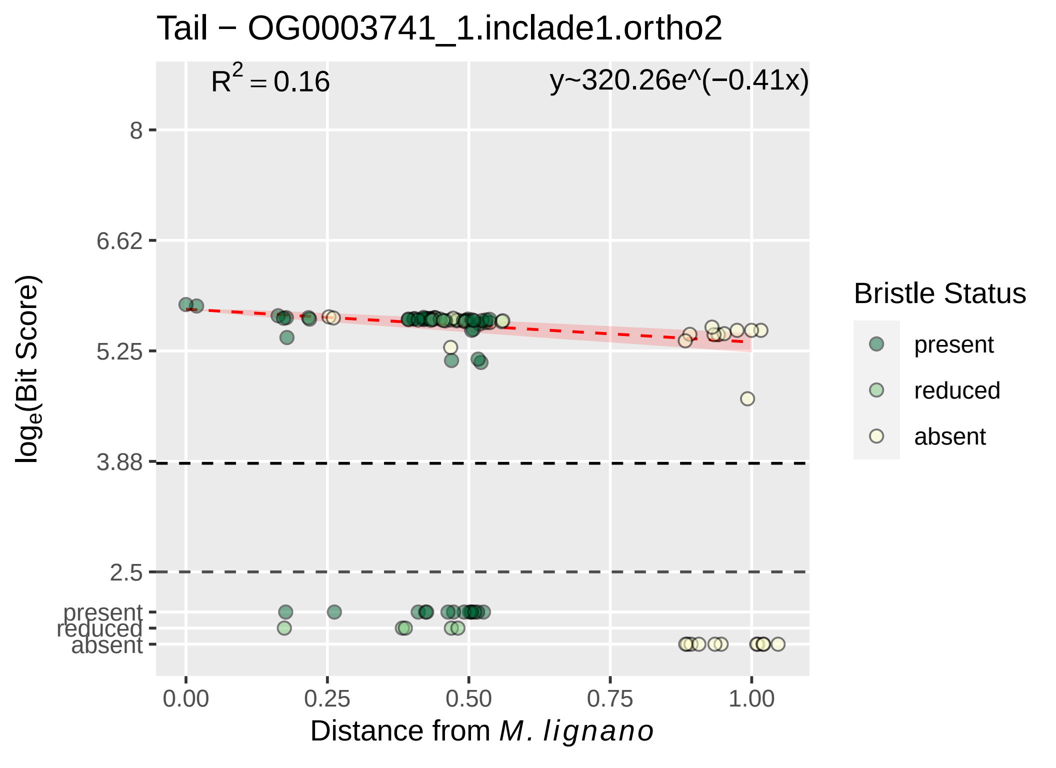

Tail − OG0003741_1.inclade1.ortho2
$R^2 = 0.16$
y~320.26e^(−0.41x)
8
6.62
5.25
3.88
2.5
present
reduced
absent
$log_e$(Bit Score)
0.00
0.25
0.50
0.75
1.00
Distance from *M. lignano*
Bristle Status
present
reduced
absent

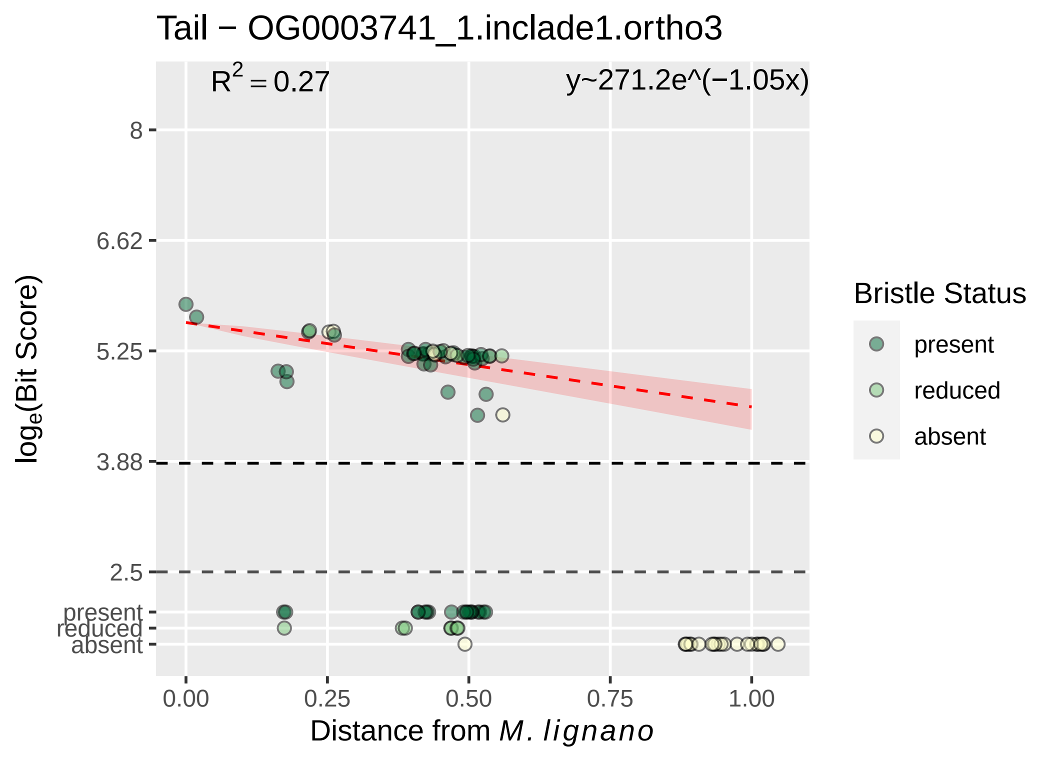

Tail − OG0003741_1.inclade1.ortho3
R² = 0.27
y~271.2e^(−1.05x)
8
6.62
5.25
3.88
2.5
present
reduced
absent
log_e(Bit Score)
0.00
0.25
0.50
0.75
1.00
Distance from M. lignano
Bristle Status
present
reduced
absent

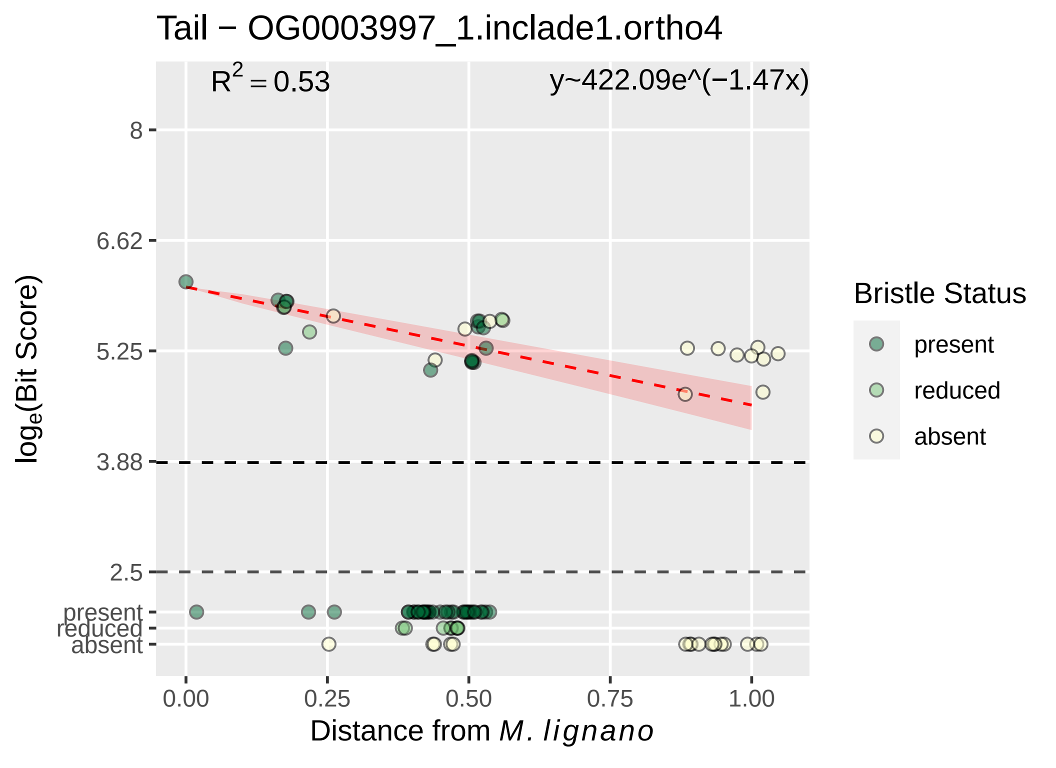

Tail − OG0003997_1.inclade1.ortho4
$R^2 = 0.53$
y~422.09e^(−1.47x)
8
6.62
5.25
3.88
2.5
present
reduced
absent
0.00
0.25
0.50
0.75
1.00
Distance from M. lignano
$\log_e$(Bit Score)
Bristle Status
present
reduced
absent

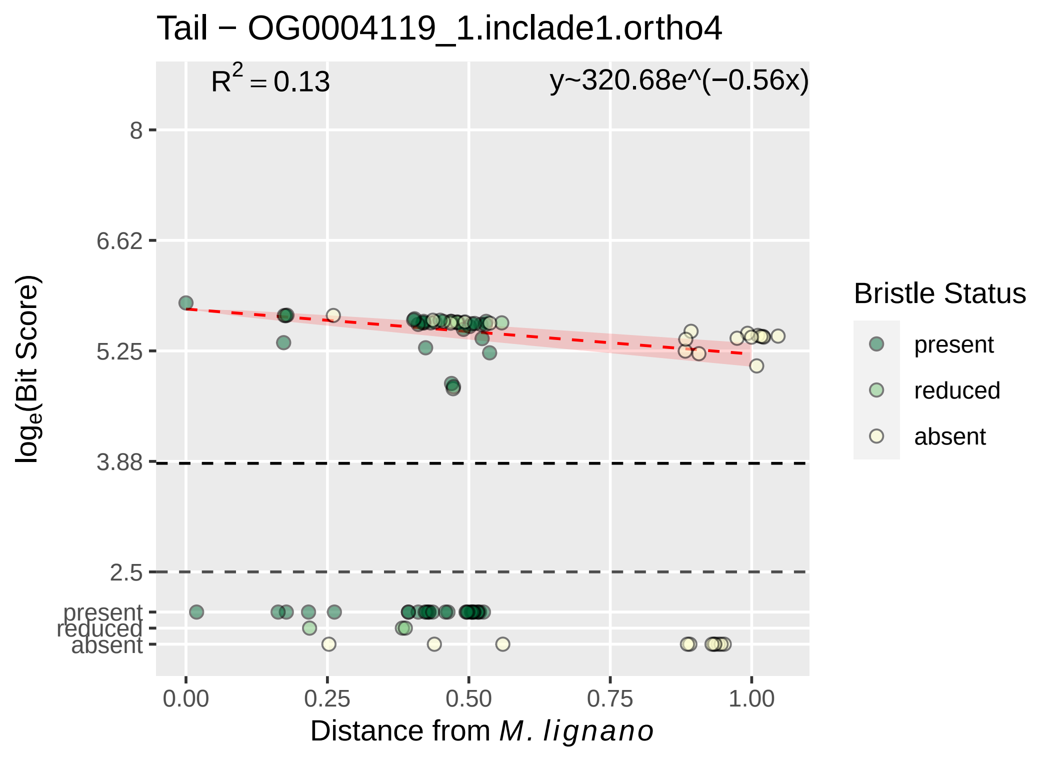
Tail − OG0004119_1.inclade1.ortho4
R^2 = 0.13
y~320.68e^(−0.56x)
8
6.62
5.25
3.88
2.5
present
reduced
absent
log_e(Bit Score)
0.00
0.25
0.50
0.75
1.00
Distance from M. lignano
Bristle Status
present
reduced
absent

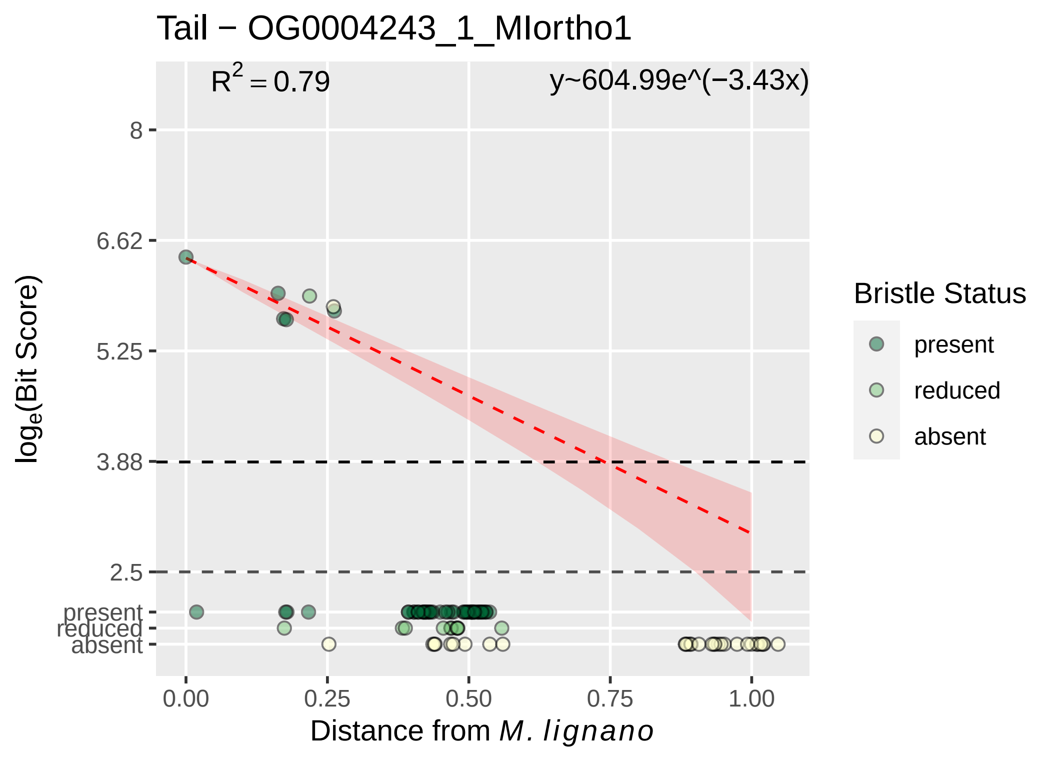

Tail − OG0004243_1_MIortho1
$R^2 = 0.79$
y~604.99e^(−3.43x)
8
6.62
5.25
3.88
2.5
present
reduced
absent
0.00
0.25
0.50
0.75
1.00
$log_e$(Bit Score)
Distance from *M. lignano*
Bristle Status
present
reduced
absent

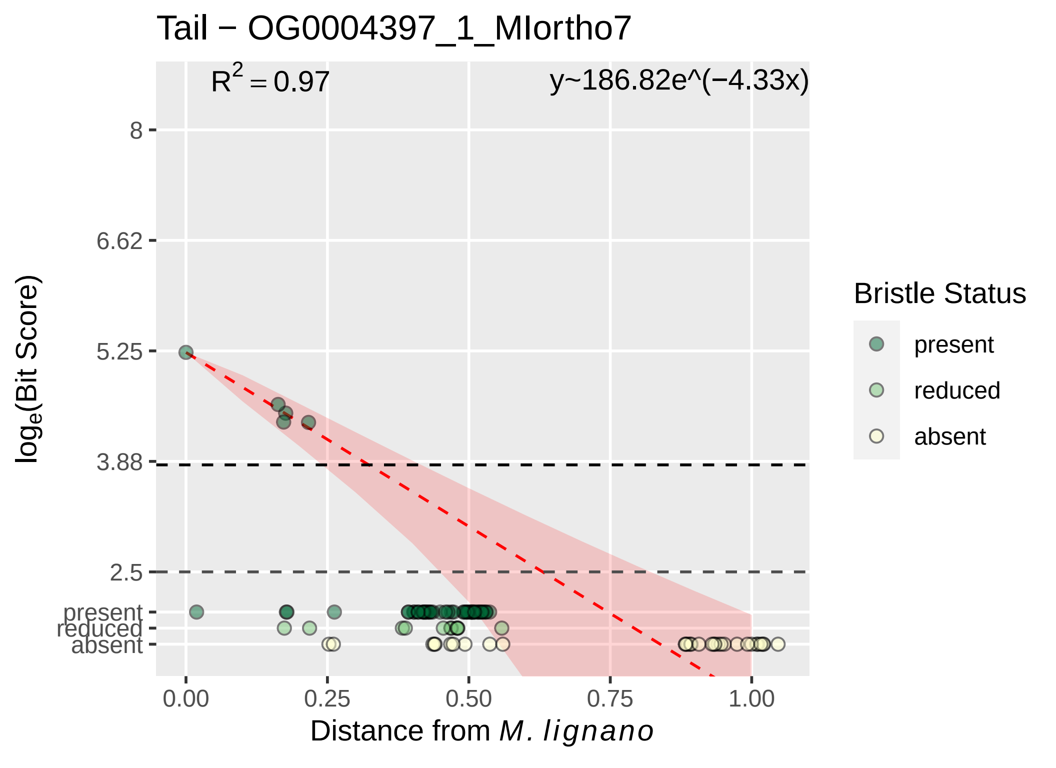
Tail − OG0004397_1_MIortho7
R^2 = 0.97
y~186.82e^(−4.33x)
8
6.62
5.25
3.88
2.5
present
reduced
absent
loge(Bit Score)
0.00
0.25
0.50
0.75
1.00
Distance from M. lignano
Bristle Status
present
reduced
absent

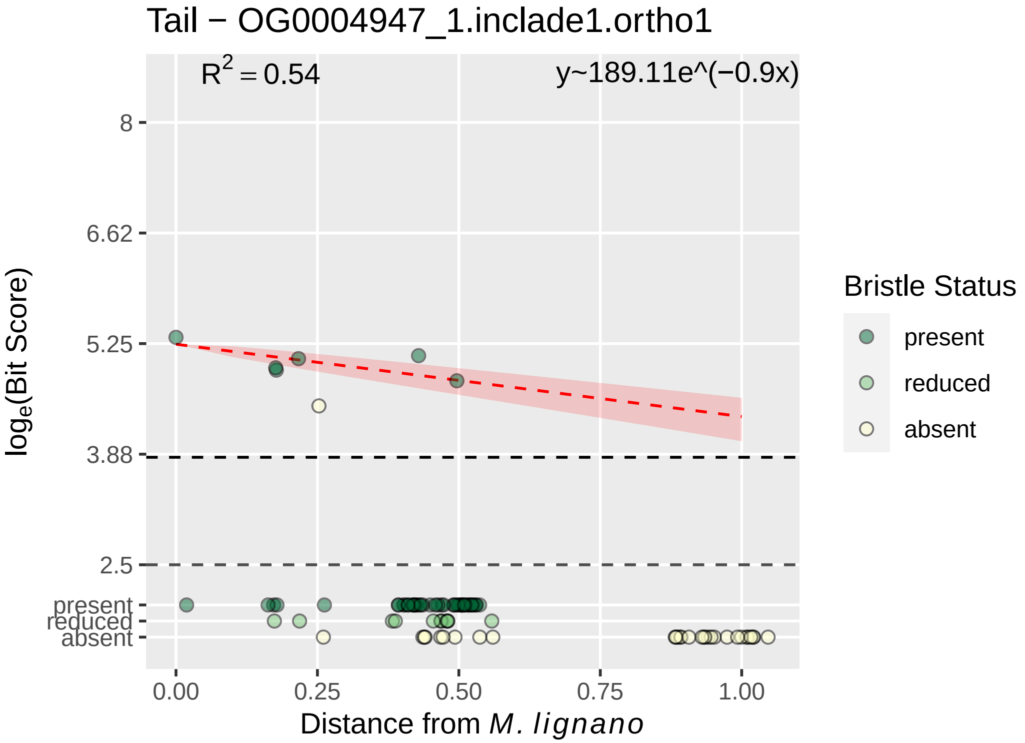

Tail − OG0004947_1.inclade1.ortho1
R^2 = 0.54
y~189.11e^(−0.9x)
8
6.62
5.25
3.88
2.5
present
reduced
absent
log_e(Bit Score)
0.00
0.25
0.50
0.75
1.00
Distance from M. lignano
Bristle Status
present
reduced
absent

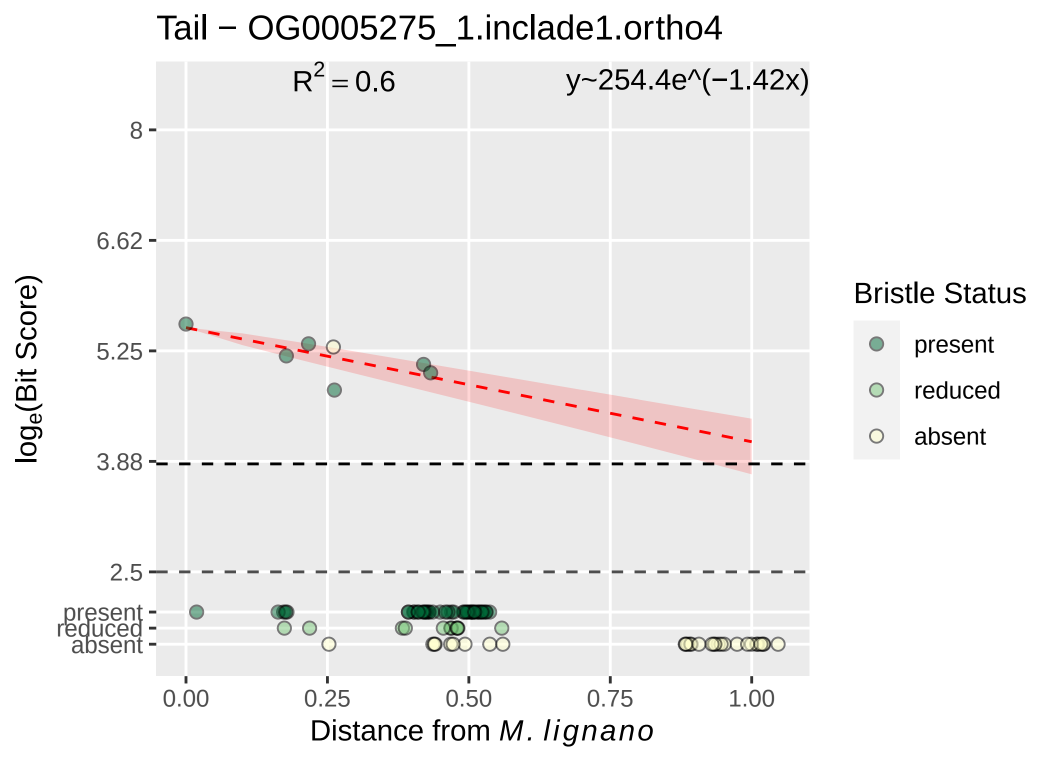

Tail − OG0005275_1.inclade1.ortho4
R^2 = 0.6
y~254.4e^(−1.42x)
8
6.62
5.25
3.88
2.5
present
reduced
absent
0.00
0.25
0.50
0.75
1.00
Distance from M. lignano
loge(Bit Score)
Bristle Status
present
reduced
absent

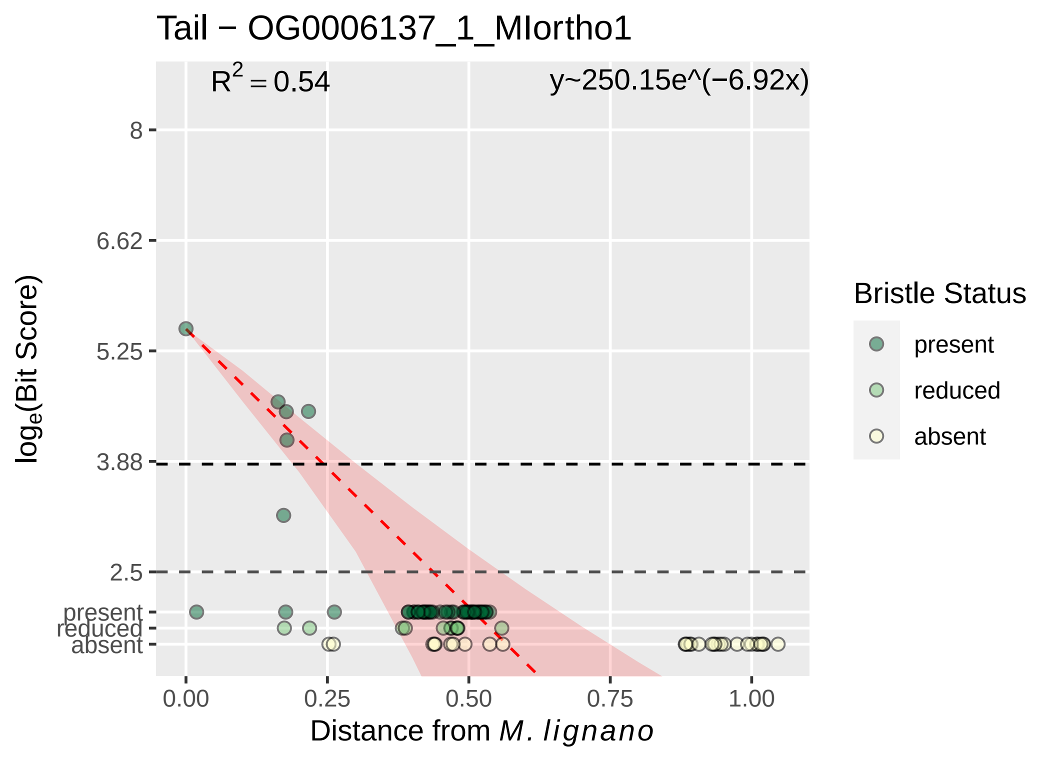

Tail − OG0006137_1_MIortho1
$R^2 = 0.54$
y~250.15e^(−6.92x)
8
6.62
5.25
3.88
2.5
present
reduced
absent
log_e(Bit Score)
0.00
0.25
0.50
0.75
1.00
Distance from M. lignano
Bristle Status
present
reduced
absent

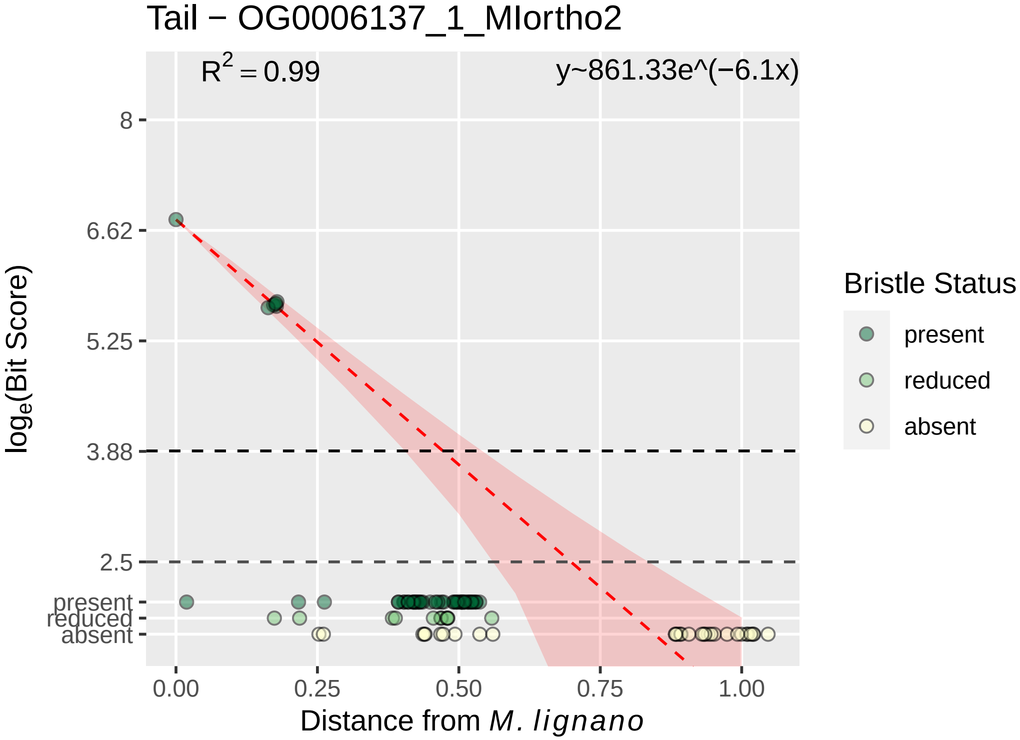

Tail − OG0006137_1_MIortho2
R^2 = 0.99
y~861.33e^(−6.1x)
8
6.62
5.25
3.88
2.5
present
reduced
absent
0.00
0.25
0.50
0.75
1.00
Distance from M. lignano
log_e(Bit Score)
Bristle Status
present
reduced
absent

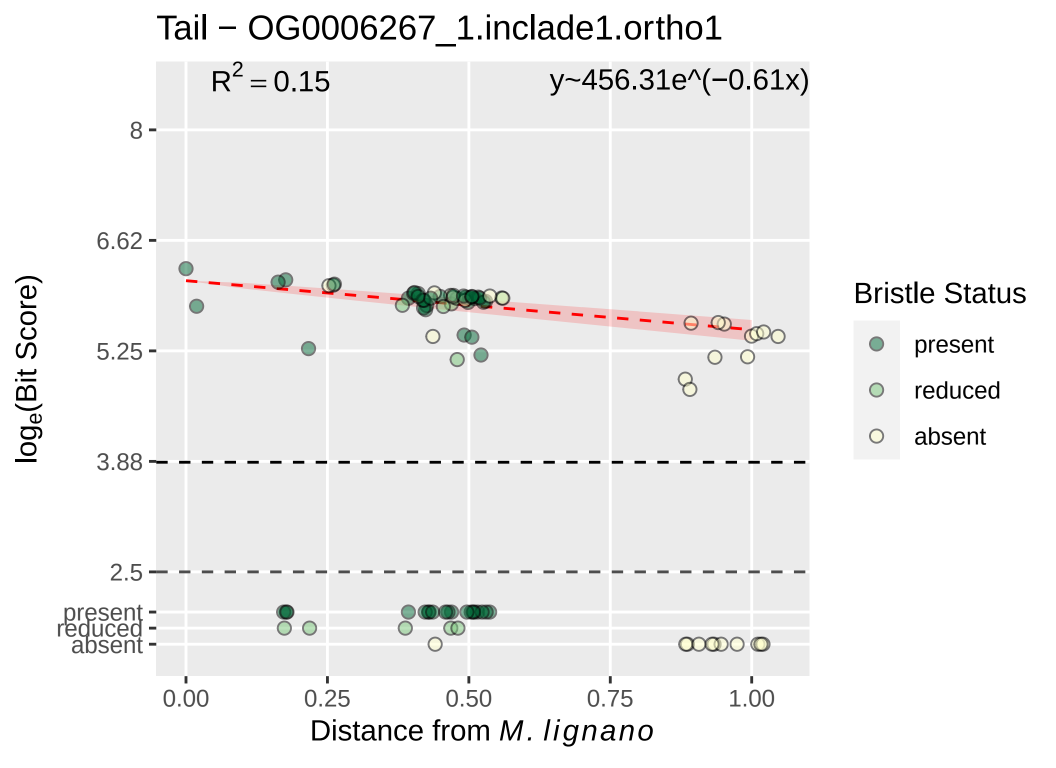

Tail − OG0006267_1.inclade1.ortho1
$R^2 = 0.15$
y~456.31e^(−0.61x)
8
6.62
5.25
3.88
2.5
present
reduced
absent
$\log_e$(Bit Score)
0.00
0.25
0.50
0.75
1.00
Distance from *M. lignano*
Bristle Status
present
reduced
absent

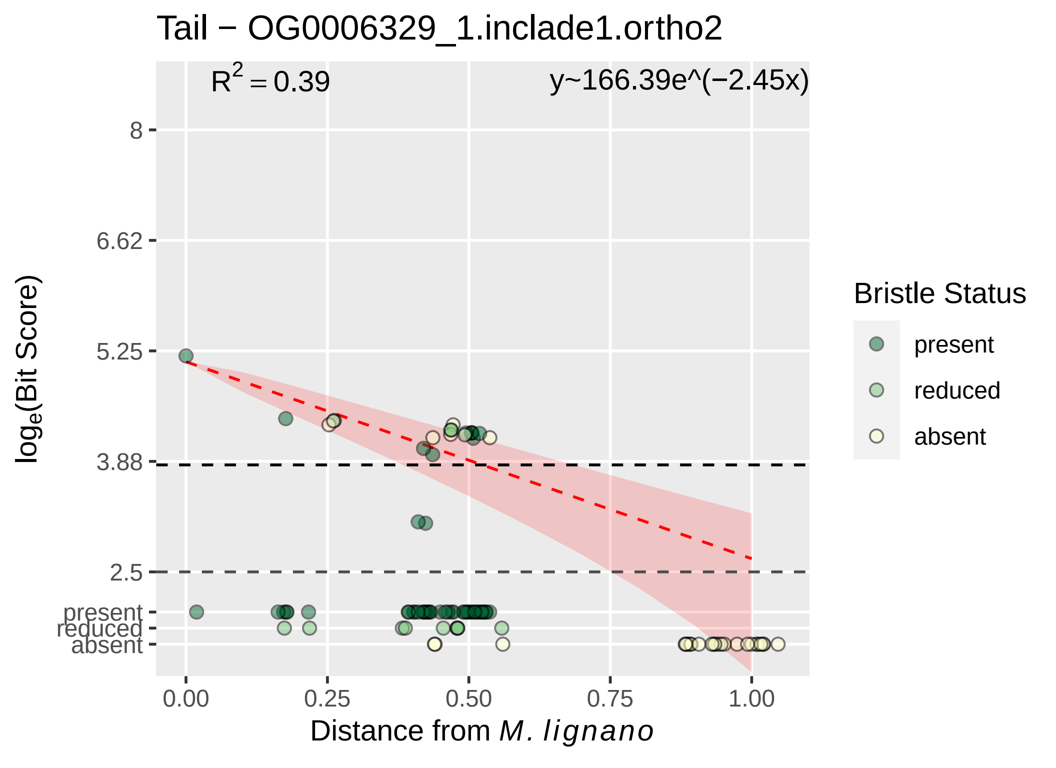
Tail − OG0006329_1.inclade1.ortho2
R^2 = 0.39
y~166.39e^(−2.45x)
8
6.62
5.25
3.88
2.5
present
reduced
absent
log_e(Bit Score)
0.00
0.25
0.50
0.75
1.00
Distance from M. lignano
Bristle Status
present
reduced
absent

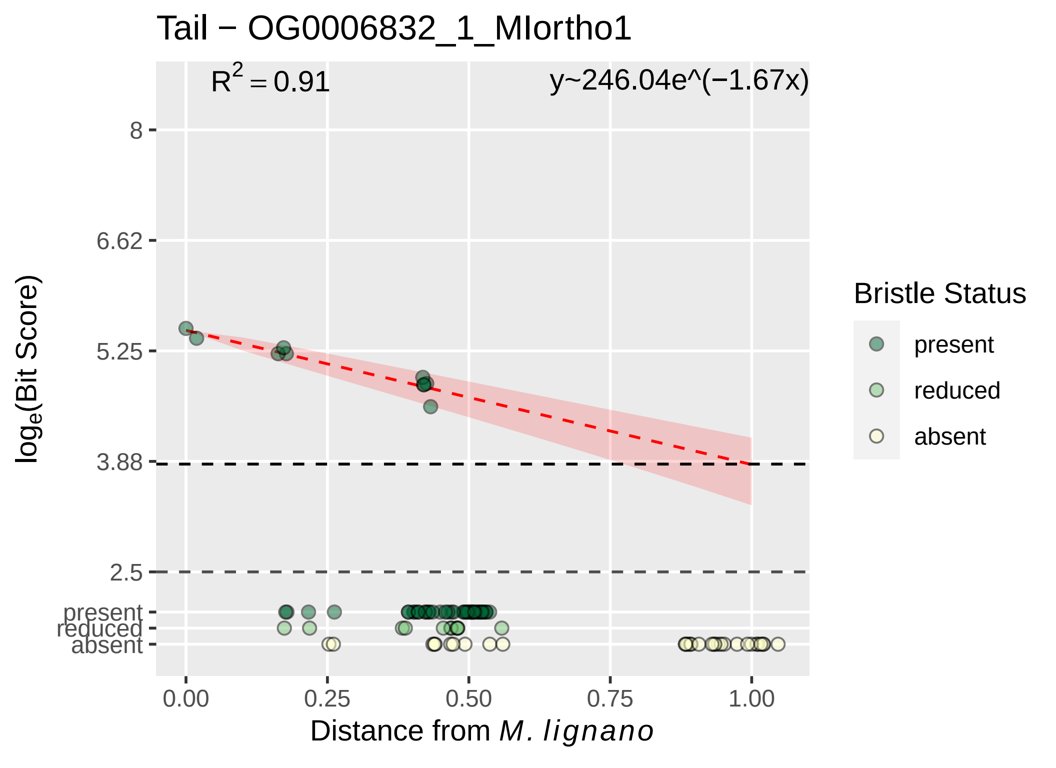

Tail − OG0006832_1_MIortho1
$R^2 = 0.91$
y~246.04e^(−1.67x)
8
6.62
5.25
3.88
2.5
present
reduced
absent
$\log_e$(Bit Score)
0.00
0.25
0.50
0.75
1.00
Distance from *M. lignano*
Bristle Status
present
reduced
absent

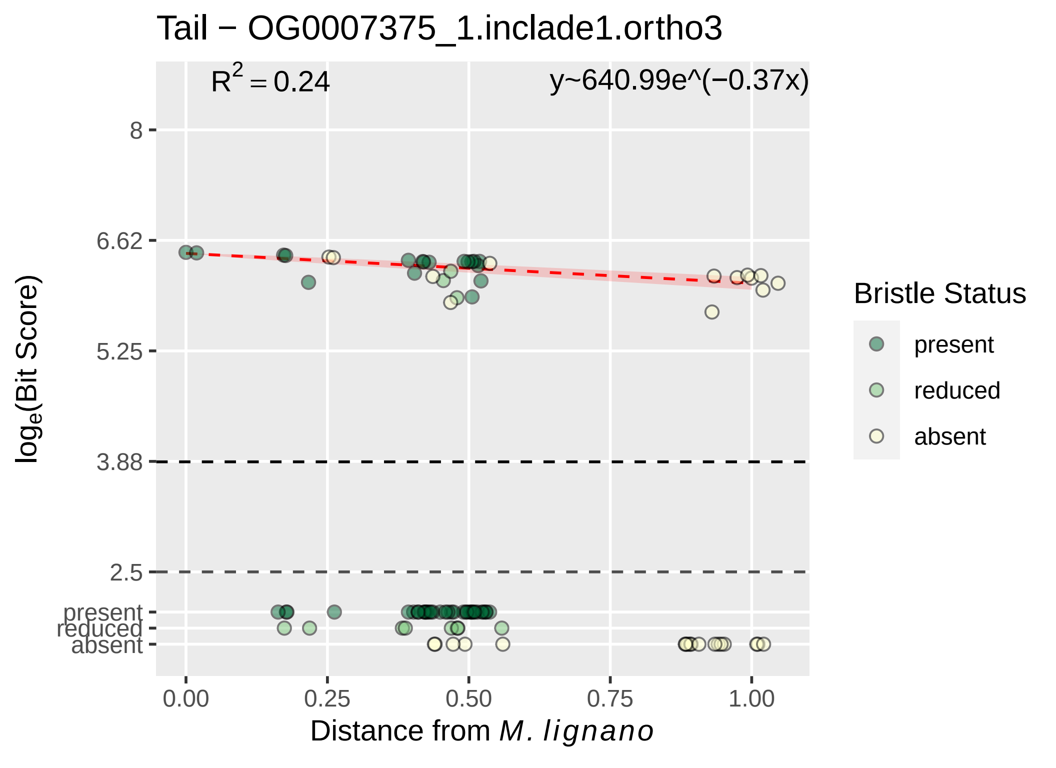

Tail − OG0007375_1.inclade1.ortho3
$R^2 = 0.24$
y~640.99e^(−0.37x)
8
6.62
5.25
3.88
2.5
present
reduced
absent
$\log_e$(Bit Score)
0.00
0.25
0.50
0.75
1.00
Distance from *M. lignano*
Bristle Status
present
reduced
absent

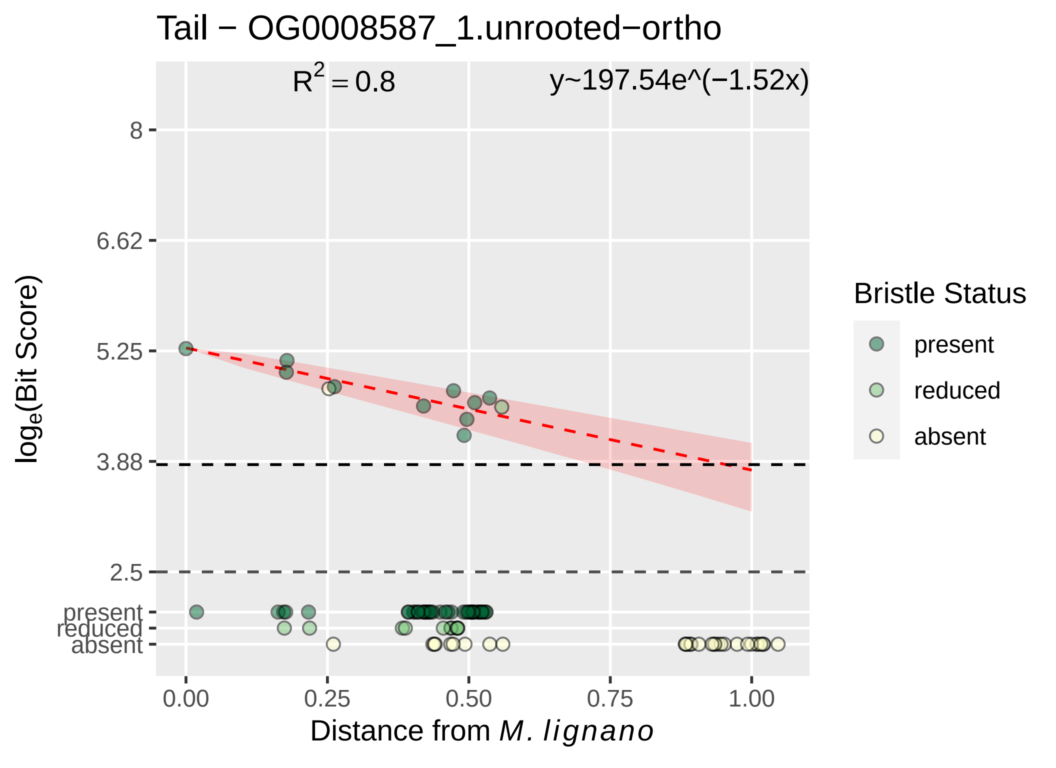

Tail − OG0008587_1.unrooted−ortho
$R^2 = 0.8$
y~197.54e^(−1.52x)
8
6.62
5.25
3.88
2.5
present
reduced
absent
$\log_e$(Bit Score)
0.00
0.25
0.50
0.75
1.00
Distance from *M. lignano*
Bristle Status
present
reduced
absent

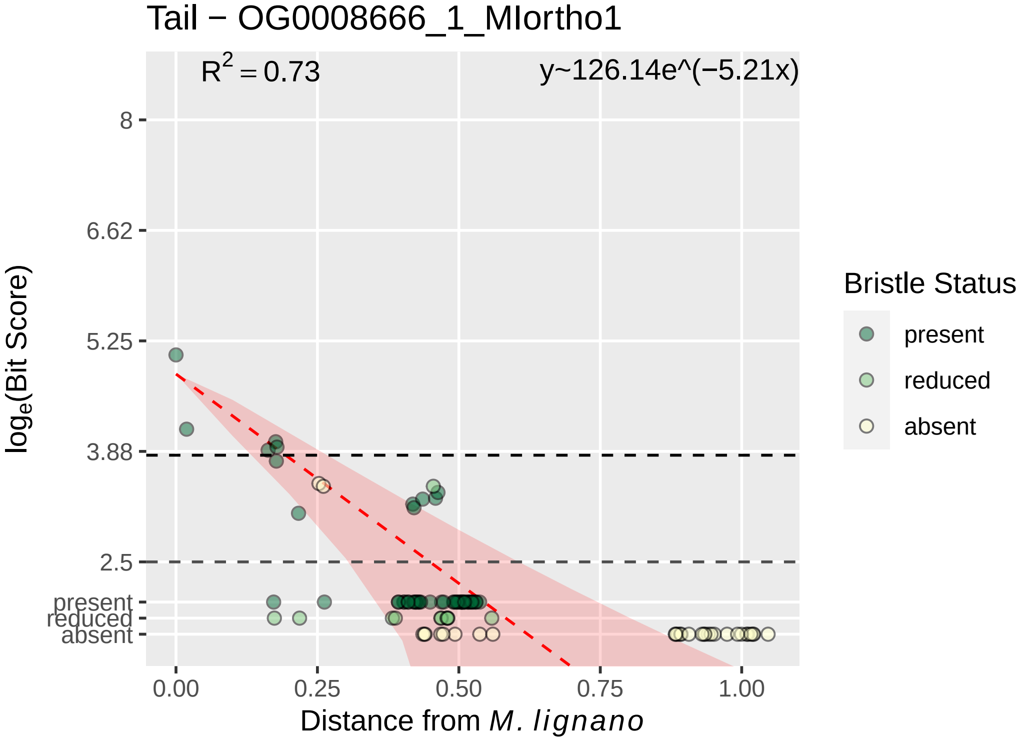

Tail − OG0008666_1_MIortho1
R^2 = 0.73
y~126.14e^(−5.21x)
8
6.62
5.25
3.88
2.5
present
reduced
absent
0.00
0.25
0.50
0.75
1.00
Distance from M. lignano
log_e(Bit Score)
Bristle Status
present
reduced
absent

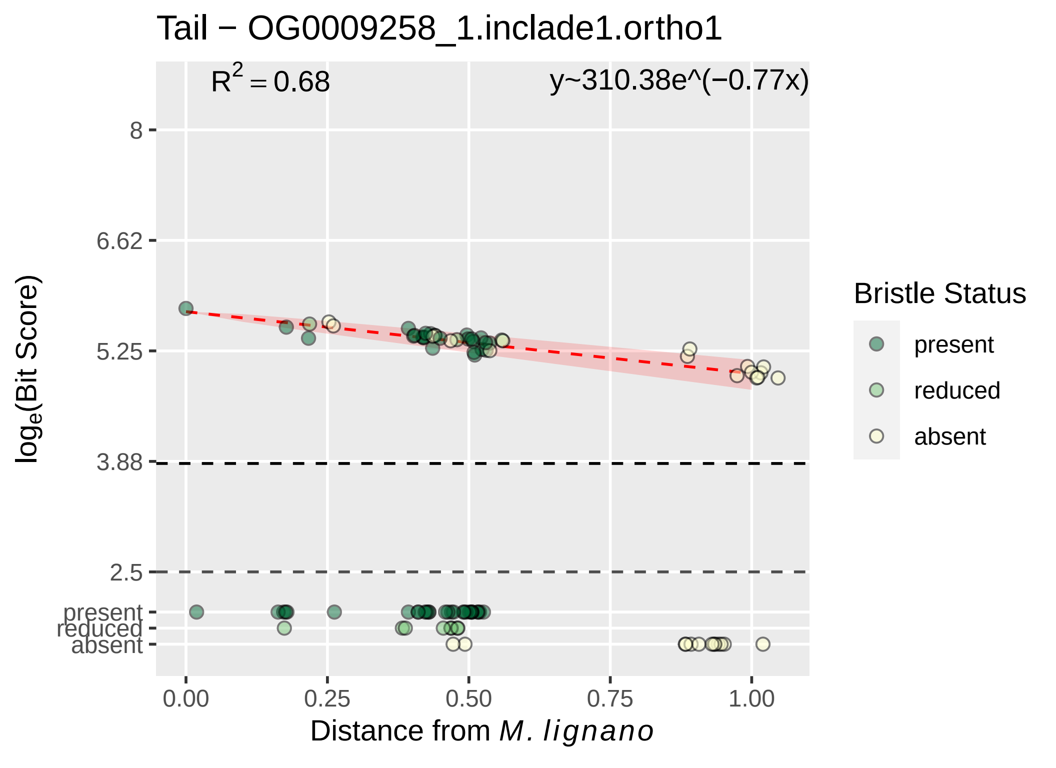

Tail − OG0009258_1.inclade1.ortho1
$R^2 = 0.68$
y~310.38e^(−0.77x)
8
6.62
5.25
3.88
2.5
present
reduced
absent
log$_e$(Bit Score)
0.00
0.25
0.50
0.75
1.00
Distance from *M. lignano*
Bristle Status
present
reduced
absent

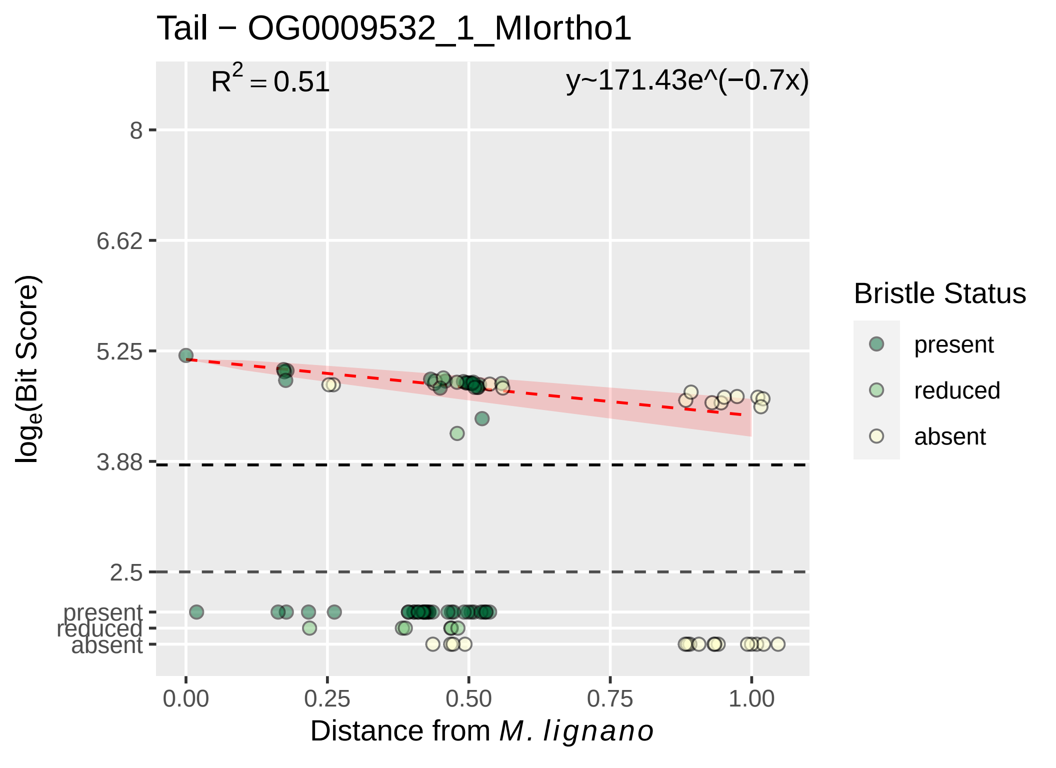

Tail − OG0009532_1_MIortho1
$R^2 = 0.51$
y~171.43e^(−0.7x)
8
6.62
5.25
3.88
2.5
present
reduced
absent
$\log_e$(Bit Score)
0.00
0.25
0.50
0.75
1.00
Distance from *M. lignano*
Bristle Status
present
reduced
absent

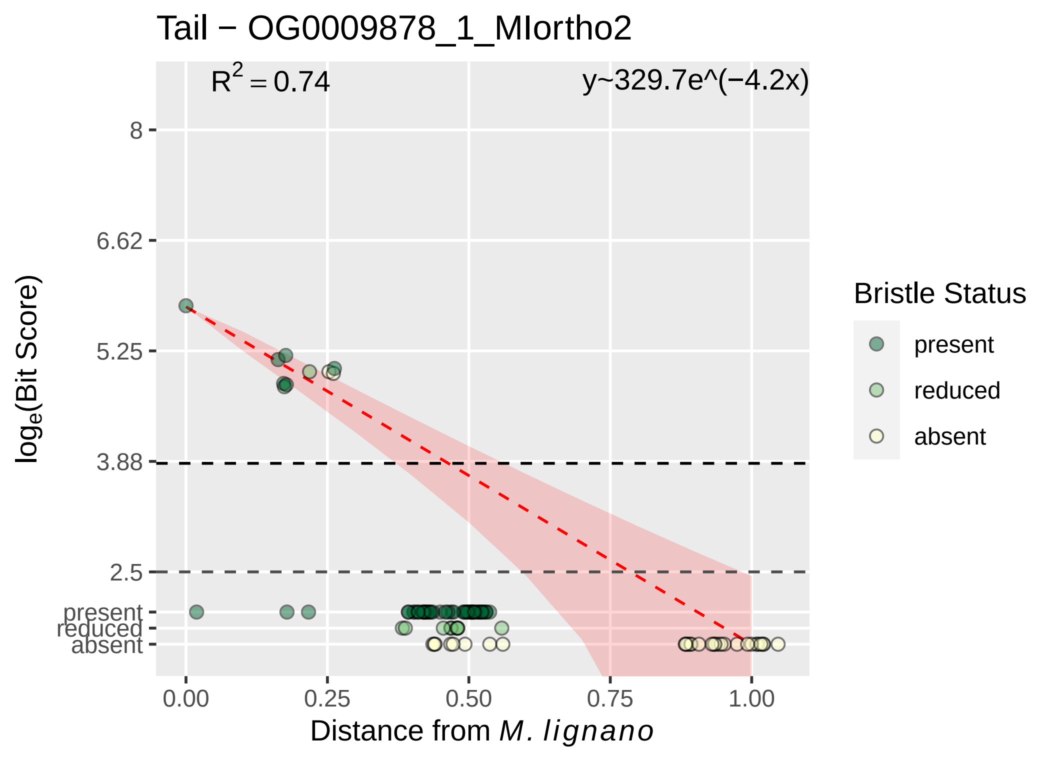

Tail − OG0009878_1_MIortho2
R^2 = 0.74
y~329.7e^(−4.2x)
8
6.62
5.25
3.88
2.5
present
reduced
absent
0.00
0.25
0.50
0.75
1.00
Distance from M. lignano
loge(Bit Score)
Bristle Status
present
reduced
absent

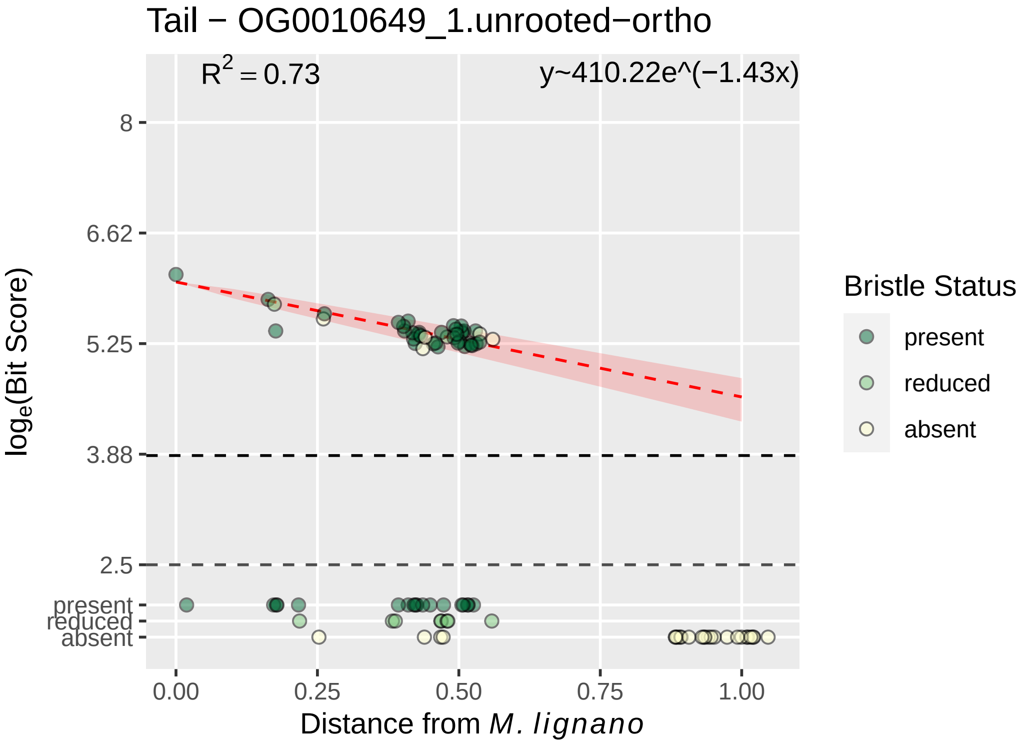

Tail − OG0010649_1.unrooted−ortho
R2 = 0.73
y~410.22e^(−1.43x)
8
6.62
5.25
3.88
2.5
present
reduced
absent
log(Bit Score)
0.00
0.25
0.50
0.75
1.00
Distance from M. lignano
Bristle Status
present
reduced
absent

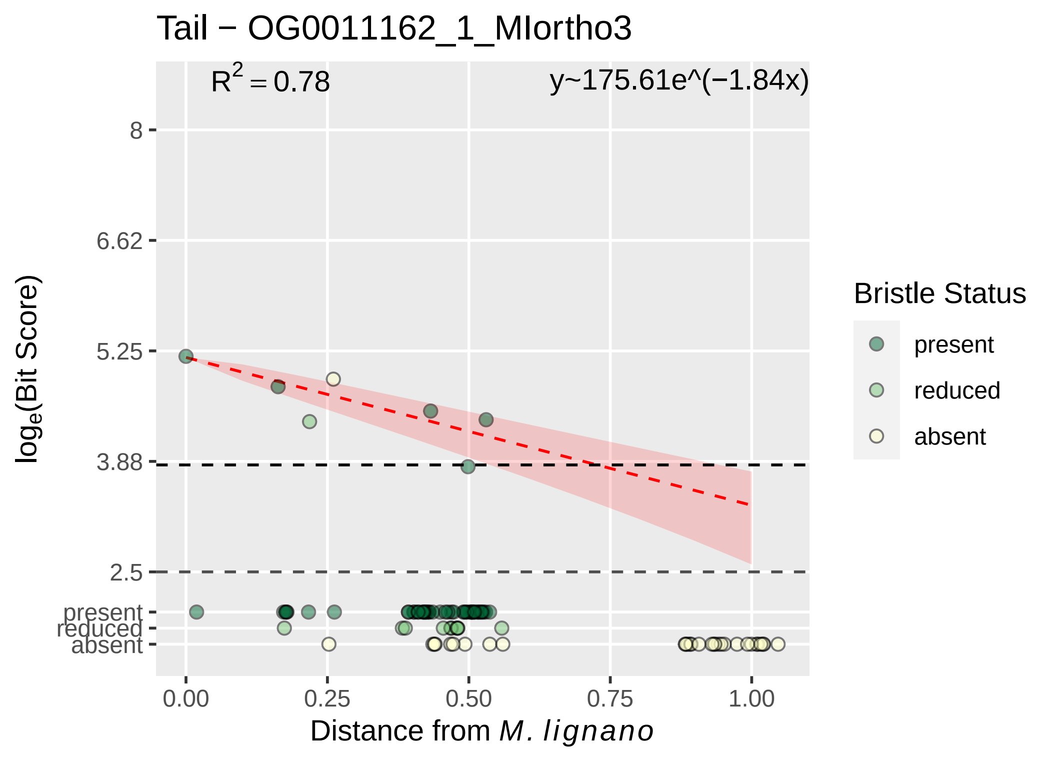

Tail − OG0011162_1_MIortho3
R2 = 0.78
y~175.61e^(−1.84x)
8
6.62
5.25
3.88
2.5
present
reduced
absent
loge(Bit Score)
0.00
0.25
0.50
0.75
1.00
Distance from M. lignano
Bristle Status
present
reduced
absent

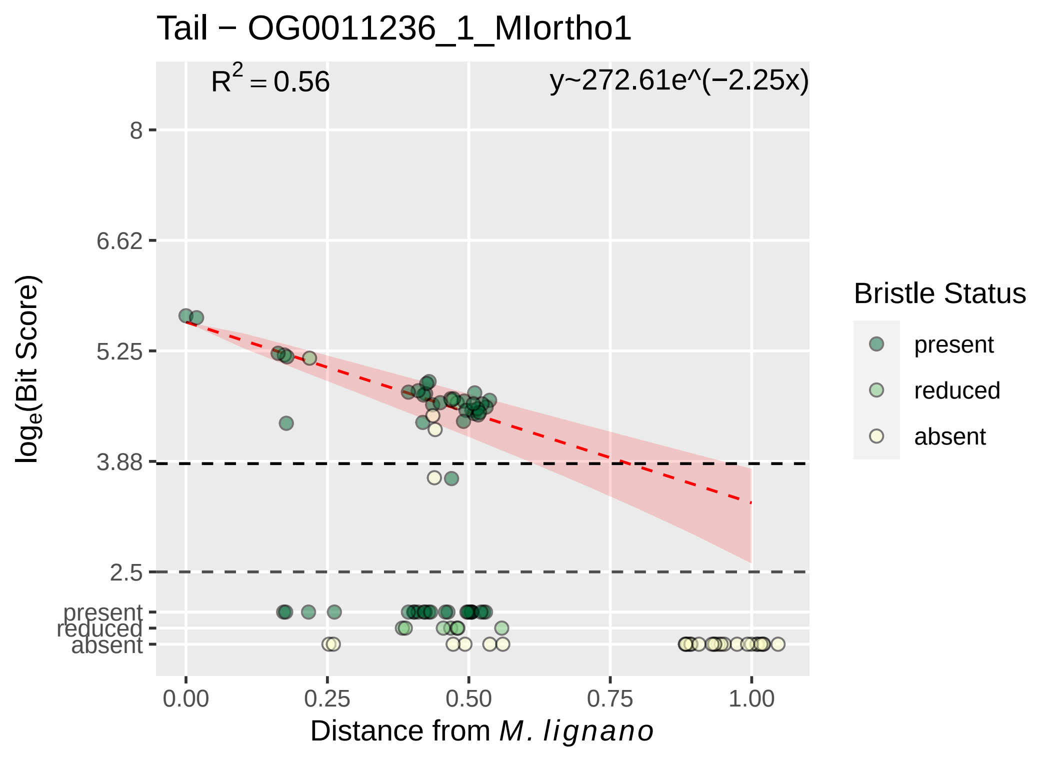

Tail − OG0011236_1_MIortho1
$R^2 = 0.56$
y~272.61e^(−2.25x)
8
6.62
5.25
3.88
2.5
present
reduced
absent
0.00
0.25
0.50
0.75
1.00
Distance from M. lignano
log$_e$(Bit Score)
Bristle Status
present
reduced
absent

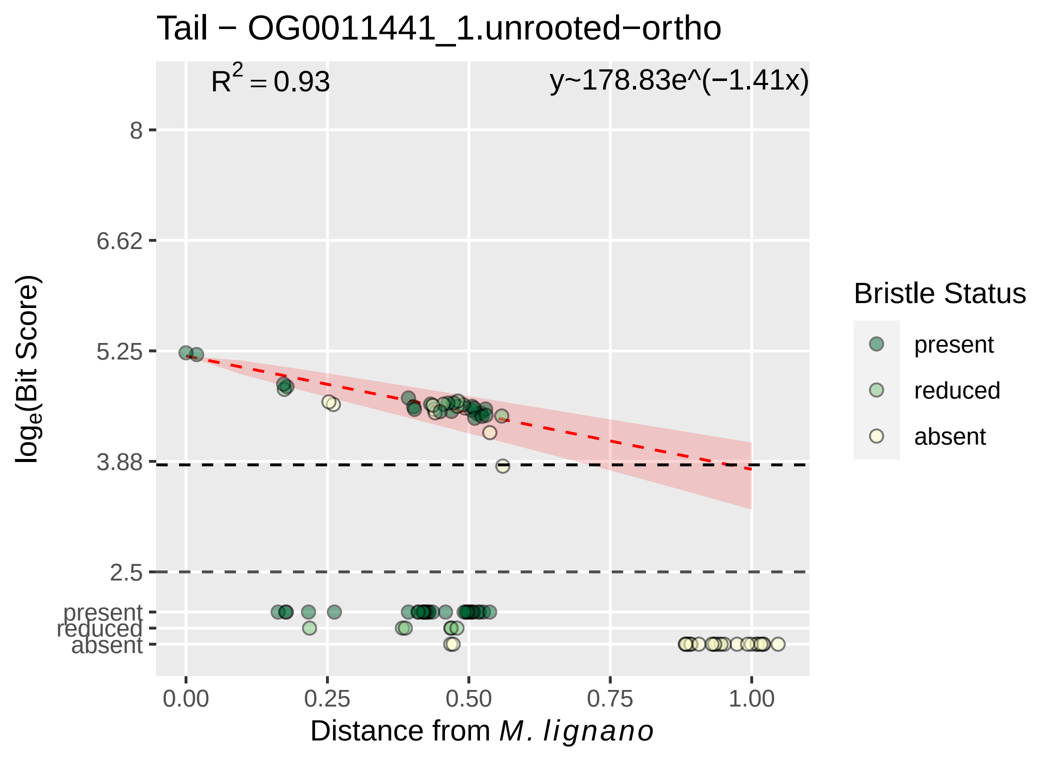
Tail − OG0011441_1.unrooted−ortho
$R^2 = 0.93$
y~178.83e^(−1.41x)
8
6.62
5.25
3.88
2.5
present
reduced
absent
$\log_e$(Bit Score)
0.00
0.25
0.50
0.75
1.00
Distance from *M. lignano*
Bristle Status
present
reduced
absent

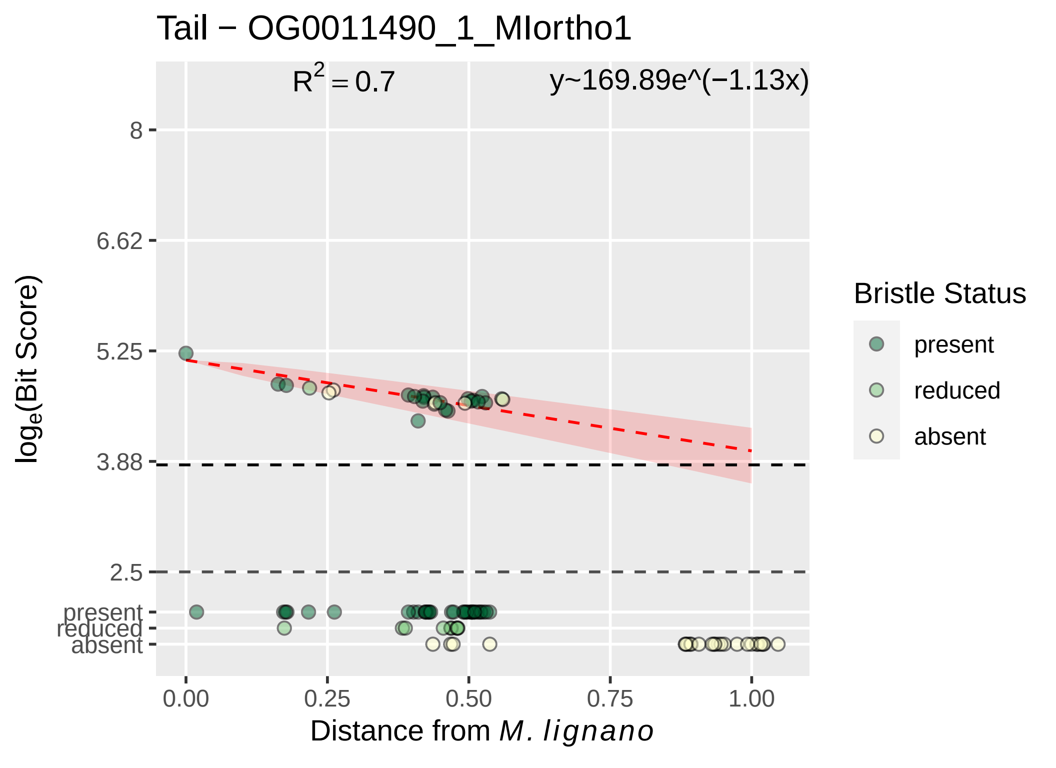

Tail − OG0011490_1_MIortho1
R2 = 0.7
y~169.89e^(−1.13x)
8
6.62
5.25
3.88
2.5
present
reduced
absent
loge(Bit Score)
0.00
0.25
0.50
0.75
1.00
Distance from M. lignano
Bristle Status
present
reduced
absent

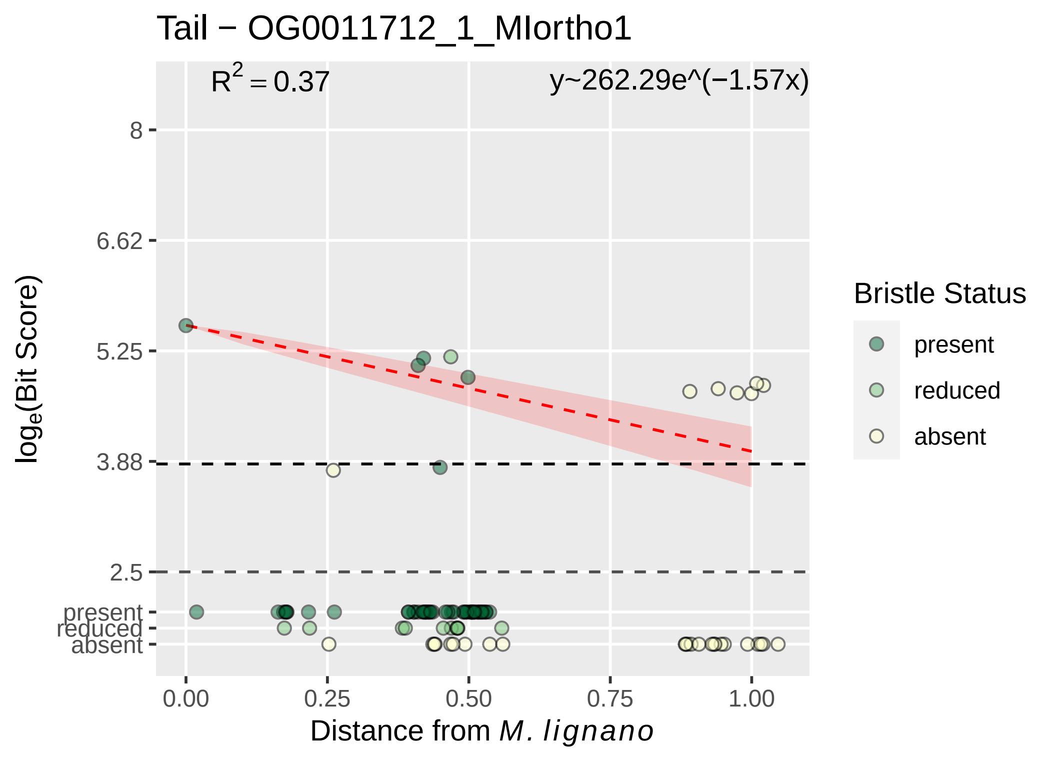

Tail − OG0011712_1_MIortho1
R² = 0.37
y~262.29e^(−1.57x)
8
6.62
5.25
3.88
2.5
present
reduced
absent
log_e(Bit Score)
0.00
0.25
0.50
0.75
1.00
Distance from M. lignano
Bristle Status
present
reduced
absent

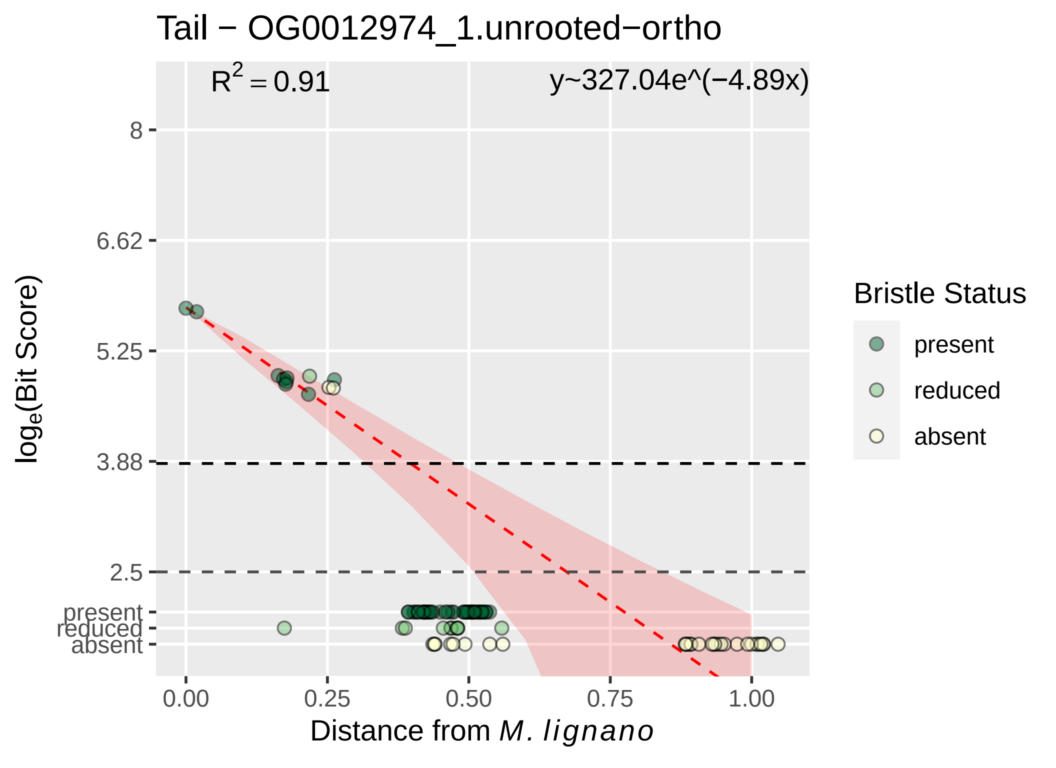

Tail − OG0012974_1.unrooted−ortho
$R^2 = 0.91$
y~327.04e^(−4.89x)
8
6.62
5.25
3.88
2.5
present
reduced
absent
log_e(Bit Score)
0.00
0.25
0.50
0.75
1.00
Distance from M. lignano
Bristle Status
present
reduced
absent

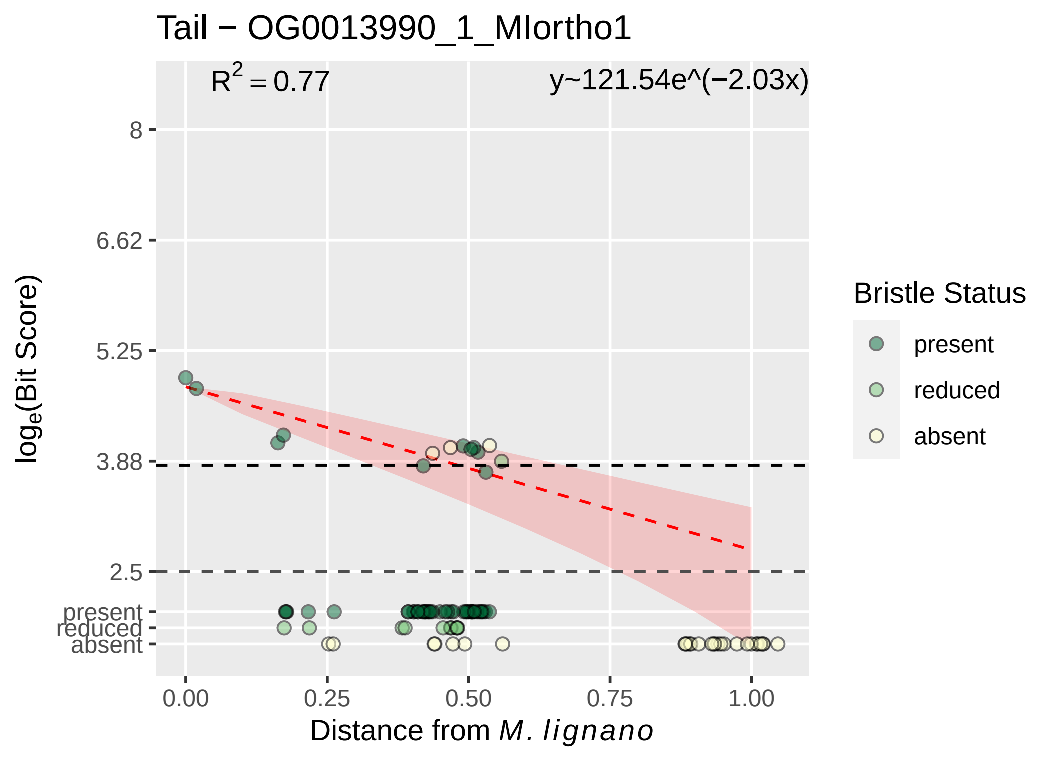

Tail − OG0013990_1_MIortho1
$R^2 = 0.77$
y~121.54e^(−2.03x)
8
6.62
5.25
3.88
2.5
present
reduced
absent
0.00
0.25
0.50
0.75
1.00
Distance from *M. lignano*
$log_e$(Bit Score)
Bristle Status
present
reduced
absent

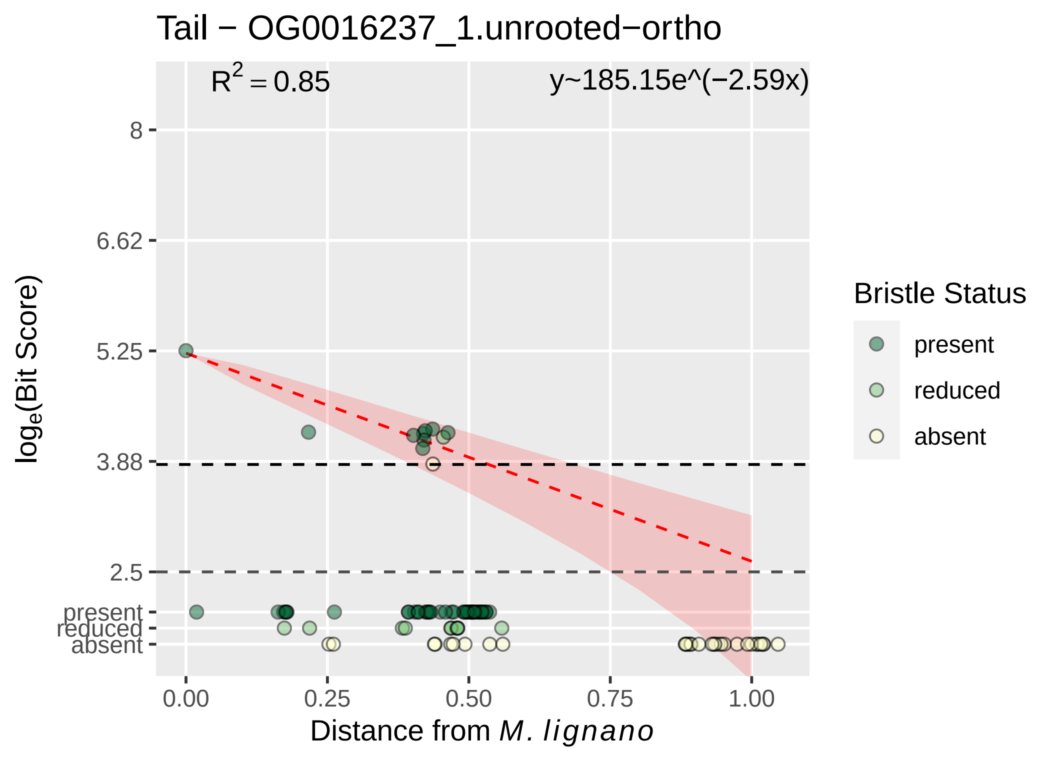

Tail − OG0016237_1.unrooted−ortho
$R^2 = 0.85$
y~185.15e^(−2.59x)
8
6.62
5.25
3.88
2.5
present
reduced
absent
0.00
0.25
0.50
0.75
1.00
Distance from *M. lignano*
$\log_e$(Bit Score)
Bristle Status
present
reduced
absent

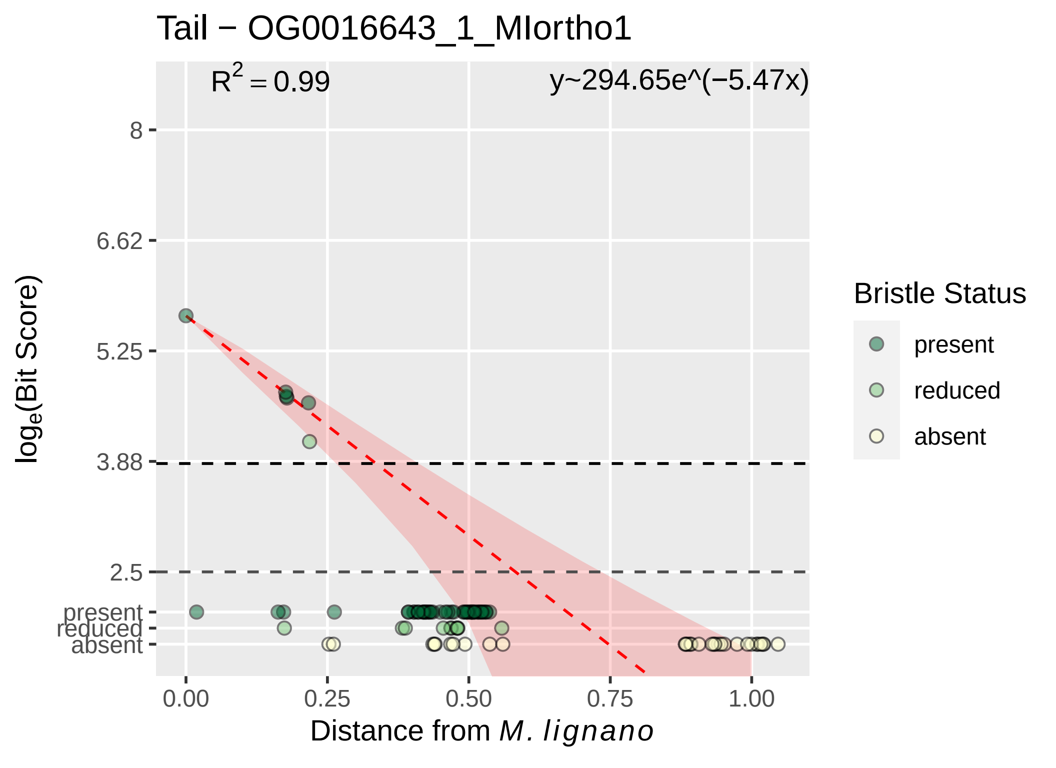

Tail − OG0016643_1_MIortho1
$R^2 = 0.99$
y~294.65e^(−5.47x)
8
6.62
5.25
3.88
2.5
present
reduced
absent
0.00
0.25
0.50
0.75
1.00
Distance from M. lignano
$\log_e$(Bit Score)
Bristle Status
present
reduced
absent

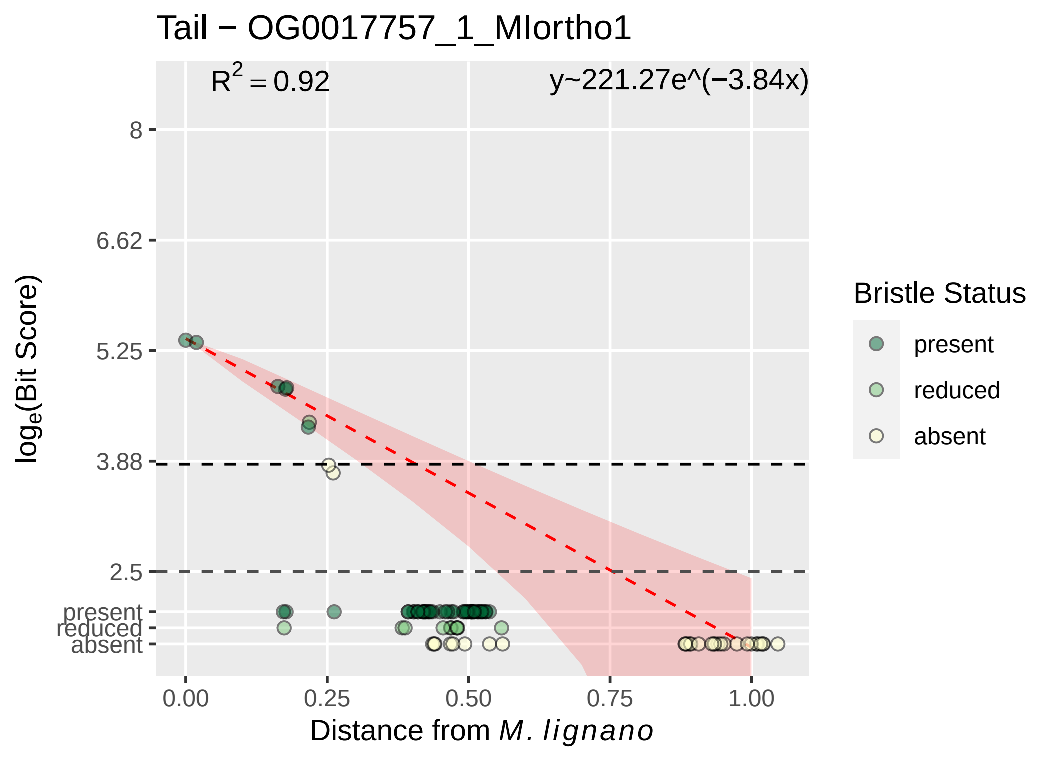

Tail − OG0017757_1_MIortho1
$R^2 = 0.92$
y~221.27e^(−3.84x)
8
6.62
5.25
3.88
2.5
present
reduced
absent
0.00
0.25
0.50
0.75
1.00
log_e(Bit Score)
Distance from M. lignano
Bristle Status
present
reduced
absent

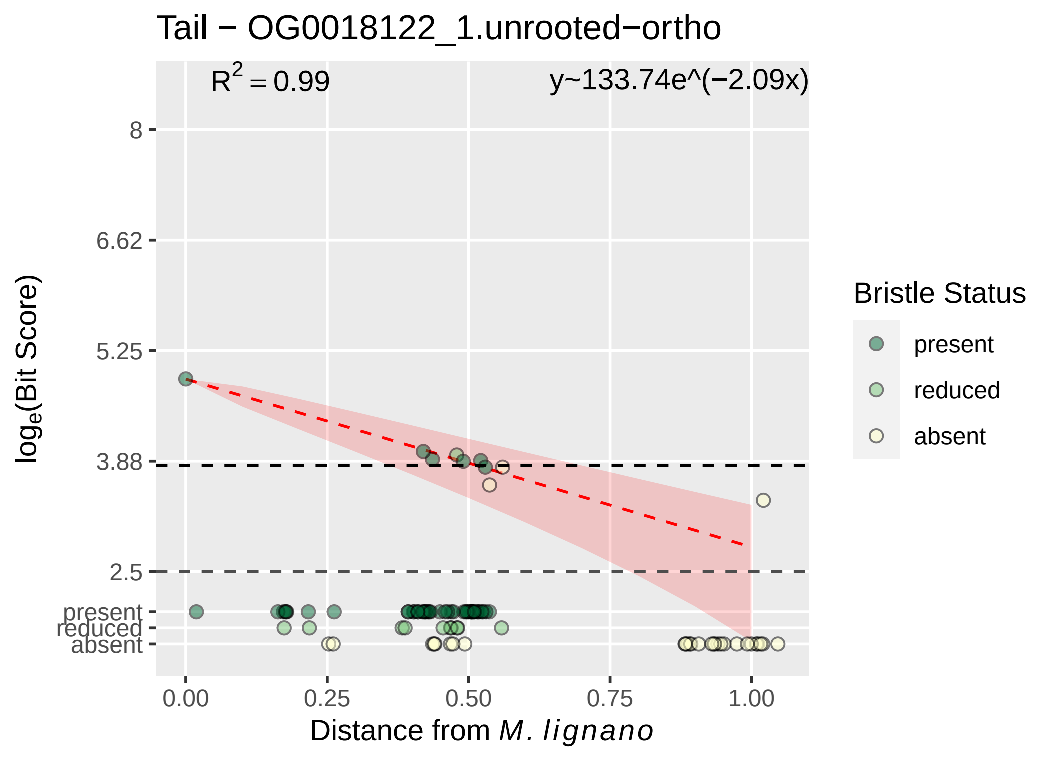

Tail − OG0018122_1.unrooted−ortho
$R^2 = 0.99$
y~133.74e^(−2.09x)
8
6.62
5.25
3.88
2.5
present
reduced
absent
$\log_e$(Bit Score)
0.00
0.25
0.50
0.75
1.00
Distance from M. lignano
Bristle Status
present
reduced
absent

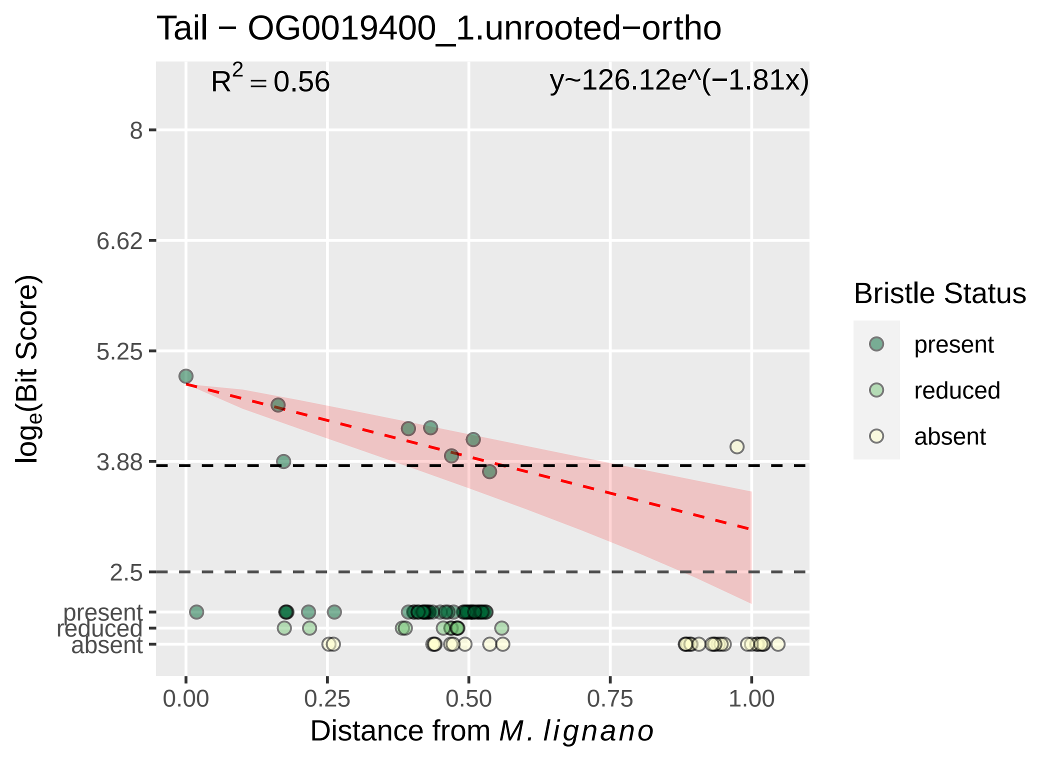

Tail − OG0019400_1.unrooted−ortho
R2 = 0.56
y~126.12e^(−1.81x)
8
6.62
5.25
3.88
2.5
present
reduced
absent
0.00
0.25
0.50
0.75
1.00
loge(Bit Score)
Distance from M. lignano
Bristle Status
present
reduced
absent
